# Permeability transition pore-related changes in the proteome and channel activity of ATP synthase dimers and monomers

**DOI:** 10.1101/2022.09.28.508998

**Authors:** Anna B. Nikiforova, Yulia L. Baburina, Marina P. Borisova, Alexey K. Surin, Ekaterina S. Kharechkina, Olga V. Krestinina, Maria Y. Suvorina, Svetlana A. Kruglova, Alexey G. Kruglov

## Abstract

Monomers, dimers, and individual F_O_F_1_-ATP synthase subunits are, presumably, involved in the formation of the mitochondrial permeability transition pore (PTP), which molecular structure, however, is still unknown. We hypothesized that upon the Ca^2+^-dependent assembly of PTP complex, F-ATP synthase (subunits) recruits mitochondrial proteins that do not interact or weakly interact with F-ATP synthase under normal conditions. Therefore, we examined whether the PTP opening in mitochondria before the separation of supercomplexes by BN-PAGE will increases the channel stability and channel-forming capacity of isolated F-ATP synthase dimers and monomers in planar lipid membranes. Besides, we studied the specific activity and protein composition of F-ATP synthase dimers and monomers from rat liver and heart mitochondria before and after PTP opening. By contrast to our expectations, preliminary PTP opening dramatically suppressed the high-conductance channel activity of F-ATP synthase dimers and monomers and decreased their specific “in gel” activity. The decline in the channel-forming activity correlated with the reduced levels of as few as two proteins in the bands: methylmalonate-semialdehyde dehydrogenase and prohibitin 2. These data indicate that proteins accompanying F-ATP synthase may be important players in the PTP formation and stabilization.

## Introduction

Despite fifty years of extensive research, the molecular structure of the PTP is still the matter of debates [1–13]. PTP models proposed earlier, such as heterooligomeric complexes of adenine nucleotide translocase, voltage-dependent anion channel (VDAC), peripheral benzodiazepine receptor, cyclophilin D (CyPD), creatine kinase, and Bax/Bcl-2 proteins [14–16]; or phosphate transporter and CyPD [17]; or VDAC, CyPD, and spastic paraplegia 7 [18], were rejected on the basis of data on genetic ablation of putative PTP components [19–26].

Recently, novel PTP models have been proposed, which comprise oligomer of the F-ATP synthase subunit c [5, 11] or dimer/oligomer of the F-ATP synthase as a core component [2, 4, 27]. The first model implies the Ca^2+^-dependent separation of the matrix-directed subcomplex F_1_ from the membrane-embedded subcomplex F_O_ comprising a ring of subunits c, which forms a large pore. According to the latter model, PTP is formed as a result of conformational changes in the F- ATP synthase dimer (or oligomer), which are induced by the binding of Ca^2+^ to the subunit β within the F_1_ subcomplex and propagate to the inner membrane through the peripheral stalk, particularly, by the oligomycin-sensitivity-conferring protein (OSCP) [2, 4, 27]. Moreover, the subunits g and f of mammalian F-ATP synthase and g and e of yeast F-ATP synthase are essential for stability and high-conductance activity in planar lipid membranes [8, 9, 28–30].

However, thermodynamically, the pore formed by the oligomer of thе F-ATP synthase subunit c should be extremely unstable [3]. Besides, it was reported that the PTP persisted in mitochondria lacking subunit c [31], though with reduced channel conductance [7].

F-ATP synthase di-(oligo-)mer model was also challenged by the data indicating that the stabilization of the F-ATP synthase dimer inhibits mPTP opening [11] and that the deletion of genes encoding the subunits b, OSCP [12], e and g [8] has a minimal effect on the PTP formation. In addition, the F-ATP synthase monomer was shown to be sufficient for the creation of megachannel-like conductance in membranes [10], while human mitochondria devoid of an assembled F-ATP synthase were able to undergo the PTP opening [8].

Thus, the molecular structure of PTP is still to be understood. Failures in attempts to solve this problem may be due to the following reason. The number of F-ATP synthase complexes (including monomers, dimers, and oligomers) in a mitochondrion is about 15000, while the number of PTP complexes is one or two and does not exceed nine [32]. Hence, PTP opening is a rather rare event, which may require the presence of minor proteins of the inner membrane, the intermembrane space or the matrix. Indeed, besides OXPHOS complexes, the inner membrane contains multiple proteins immersed in, and anchored to the lipids or associated with the matrix and the intermembrane surfaces of the membrane. One can assume that some of these proteins associate with F-ATP synthase complex dimer or monomer in the presence of Ca^2+^ and participate in the formation and stabilization of the PTP complex.

In fact, the PTP in mitochondria does not close spontaneously unless Ca^2+^ is removed and/or strong PTP antagonists are added, indicating a high stability of the PTP structure [33–35]. In addition, mitochondrial megachannels (MMC) in mitoplasts are characterized by stable states of conductance and clear transitions between them [36–38], while isolated and purified dimers and monomers of F-ATP synthase in artificial lipid membranes demonstrate quickly changing conductance with unclear states [2, 4, 10].

We hypothesized that PTP opening in mitochondria before the separation of (super)complexes and other proteins by blue-native gel electrophoresis can increase the level of partner proteins associated with F-ATP synthase dimers and monomers after separation and, thus, increase the stability and activity of channels formed upon the incorporation of dimers/monomers into lipid membranes.

Here we compared the data of mass-spectrometric analysis and conductance of F-ATP synthase (both dimers and monomers) isolated from control mitochondria and mitochondria in which the PTP was opened. The results confirm the role of F-ATP synthase-associated proteins in the PTP formation.

## Results

### Effect of PTP opening on the in gel ATPase activity

Mitochondria from different organs and tissues possess a tissue-specific protein composition, which may predetermine the peculiarities in the PTP regulation (sensitivity to Ca^2+^, ROS, and inhibitors) and dynamics. In addition, the properties of the incubation medium may affect the strength of association of matrix and intermembrane space proteins with lipids and complexes in the inner membrane. Therefore, we first compared the effect of PTP opening on the specific activity of F-ATP synthase dimers (D) and monomers (M) isolated from rat liver and rat heart mitochondria (RLM and RHM) (Fig. 1). Preliminarily, RLM were incubated in the presence of 1 mM EGTA (control samples) or 250 μM CaCl_2_ (PTP samples) either in KCl- or sucrose/mannitol-based media (KCl-BM or SM-BM, respectively). RHM were incubated in SM-BM only. The registration of mitochondrial swelling indicated that in each case, 250 μM Ca^2+^ was sufficient to induce PTP opening during a 15–35-min incubation (Fig. 1A). PTP opening had a minor effect on the specific “in gel” activities in respiratory complexes I (CI), III (CIII), and IV (CIV) and their assemblies (supercomplexes) (Fig. 1B). (Figure shows the activity of RLM complexes incubated in KCl-BM). ATPase activity declined in PTP samples from both RLM and RHM, and in KCl-BM the decrease was more pronounced (Figs. 1B and 1C). Thus, long-term incubation with Ca^2+^ and/or PTP opening destabilizes the F-ATP synthase and reduces its in gel activity after separation.

**Figure 1.**
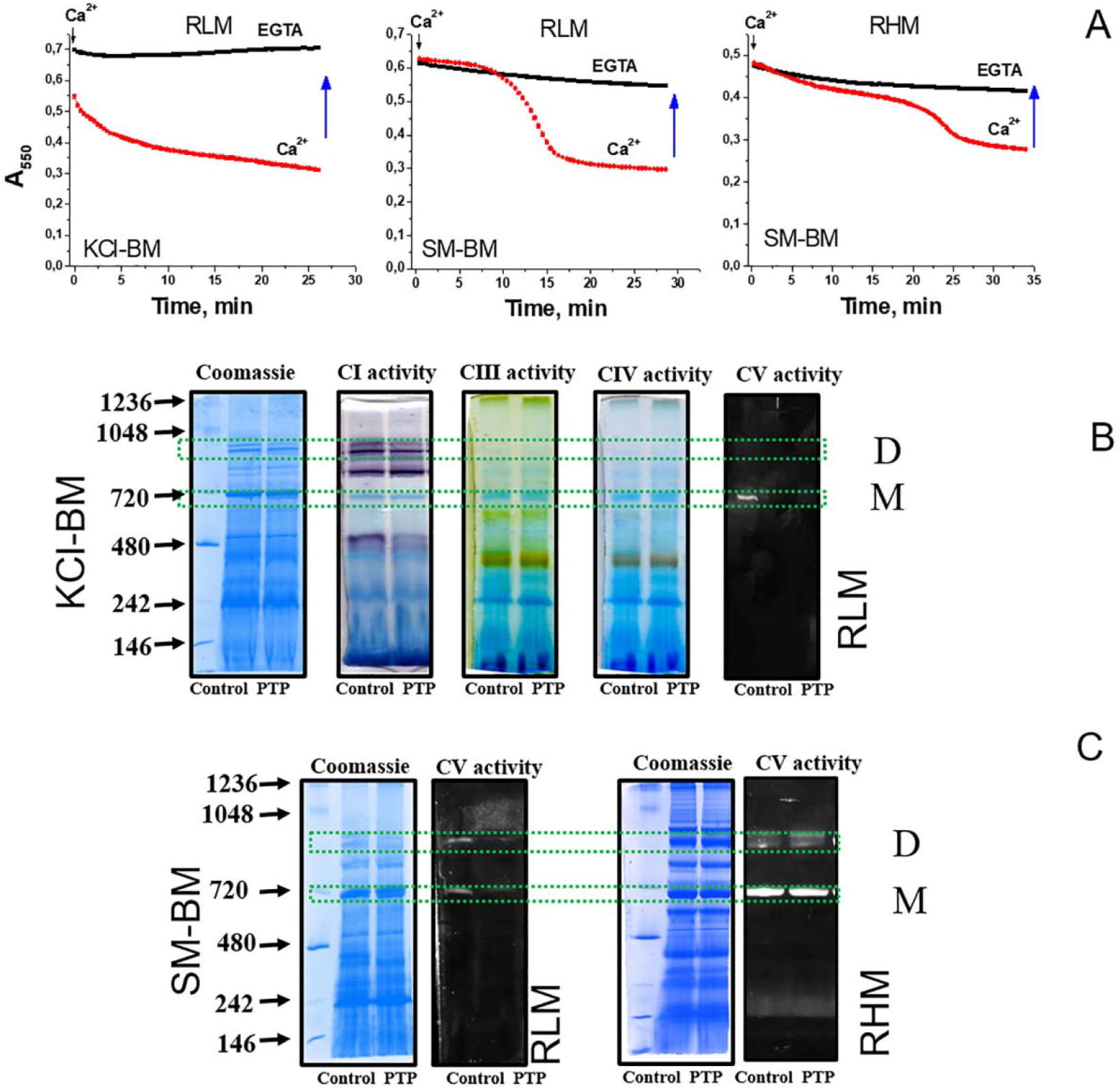
Effect of PTP opening on the in-gel activity of F-ATP synthase monomers and dimers from rat liver and heart mitochondria. A. Swelling of RLM and RHM (0.75 mg protein/ml) in KCl- and SM-BM induced by 250 μM Ca^2+^. Arrows show the time when samples were collected for BN-PAGE. B. In-gel staining of CI, CIII, CIV, and CV specific activity in control and PTP samples from RLM incubated in KCl-BM. Specific activity staining was performed as described in “Materials and Methods”. Monomers and dimers are designated as M and D, respectively. C. In-gel ATPase activity in control and PTP samples from RLM and RHM incubated in SM-BM. All the figures are representative of at least three independent experiments.

### Channel-forming activity of dimers and monomers of F-ATP synthase from control and PTP samples

Then, we examined whether the alterations in the activity of F-ATP synthase monomers and dimers after the PTP opening are associated with changes in their channel-forming capability. Complexes and associated proteins were eluted from D and M bands of RLM control and PTP samples incubated in SM-BM, after which they were incorporated in bilayer soybean lecithin membranes (Fig. 2). The incorporation of F-ATP synthase dimers from control samples into membranes did not induce any channel activity unless Ca^2+^ (300–600 μM) was added at the same (*trans*) side of membranes. Ca^2+^ evoked a delayed by 0.5–2 min but abrupt formation (Fig. 2F) of large and sustainable (for up to several minutes) channels (Fig. 2A, SI Appendix Fig. S1). At *cis* positive voltage (negative from the side of protein addition), channel states and substates were more stable than at *cis* negative voltage. However, the overall conductance was similar. An amplitude histogram revealed that, at *cis* negative voltage, the maximum channel conductance was approximately 1.22 nS with substates of 60, 250, 670, and 915 pS; at positive voltage, the maximum conductance was about 1.53 nS with substates of 100, 650 pS, and 1.09 nS (Fig. 2B). The probability for channels to reside in other conductance states was several orders of magnitude lower. At higher voltage (100 mV) channels were formed more readily, while at lower voltage (50 mV) they were more stable. At middle voltage values, the current–voltage relationship was essentially linear (Fig. 2C).

**Figure 2.**
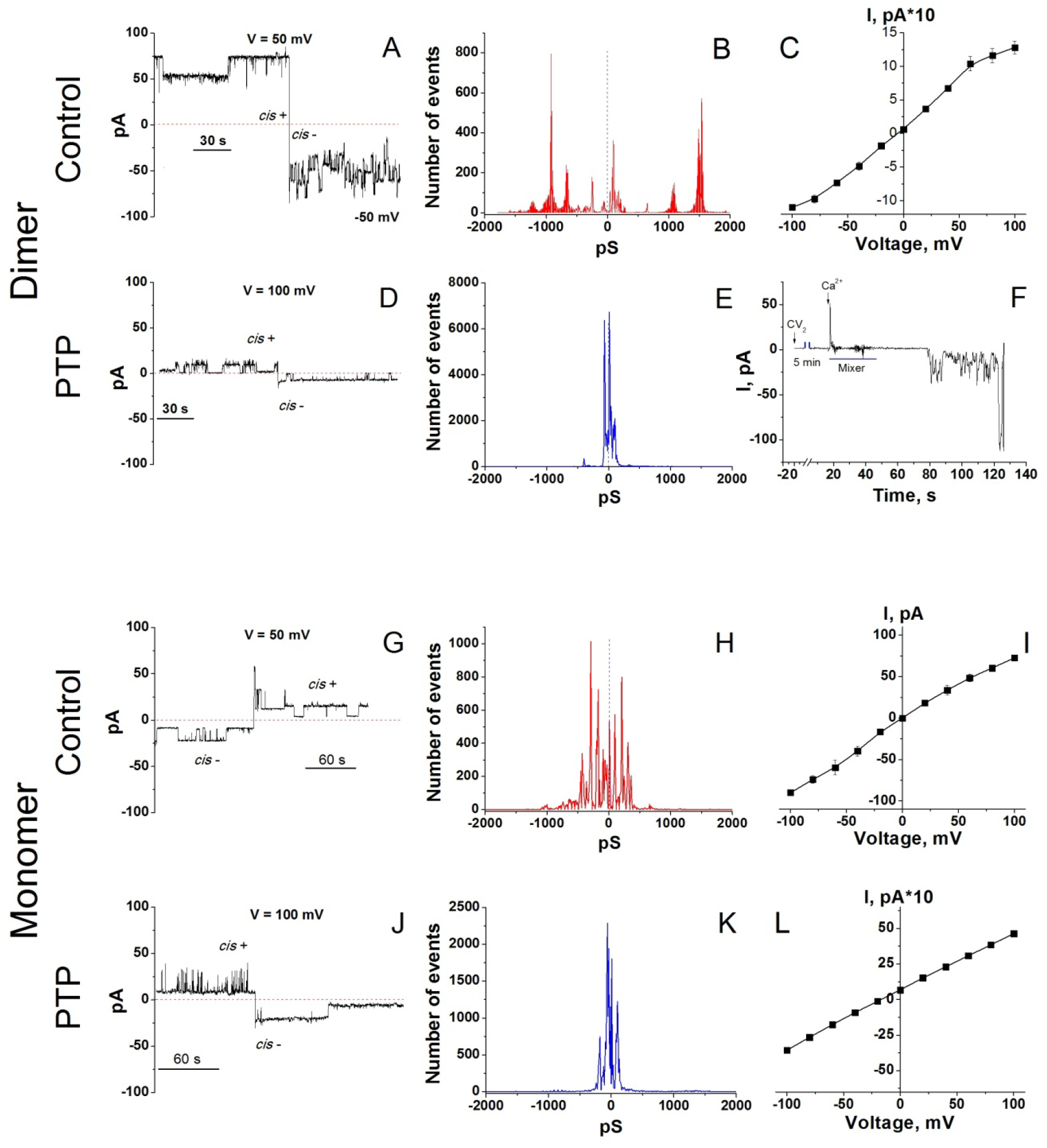
Channels formed in planar lipid membranes by F-ATP synthase dimers and monomers from control and PTP samples. A, D, G, and J. Representative channel activity of dimers from control and PTP samples and monomers from control and PTP samples, respectively. Currents were recorded at indicated (*cis*) positive and negative voltage. Protein eluate (30-90 μl/ml) and calcium (300-900 μM) were added from the *trans* side of the membrane. Dashed lines show the closed state of the channel. B, E, H. and K. Corresponding amplitude histograms of conductance (pS) indicate the probability for channels to reside in different conductance states. Each histogram except the one on panel E summarizes the data of three records from independent experiments. Panel E shows the data of one experiment. Negative and positive values on the abscissa axis correspond to *cis* negative and positive voltage. C, I, and L. Current-voltage relationships obtained for the control dimer, the control monomer, and the PTP monomer, respectively. Values on the curves are means ± S.E.M. (n = 3) for three independent experiments. F. A typical picture of the incorporation of F-ATP synthase dimer (CV_2_, control) and Ca^2+^-dependent channel formation in lipid bilayer. The final Ca^2+^ concentration was 300 μM. The interval between the additions of the protein and Ca^2+^ was about 5 min.

At protein concentrations three times higher, the conductance reached the values of several nS with transitions of ~0.6 and 1 nS, which, presumably, was due to the incorporation of several channel complexes (SI Appendix Fig. S2).

By contrast to our expectations, the incorporation of the dimer from PTP samples followed by Ca^2+^ treatment did not cause any channel activity in three out of four experiments (SI Appendix Fig. S3B). In one experiment, we observed sustainable channels with a conductance of 70 and 100 pS at *cis* negative and positive voltage, respectively (Fig. 2D, SI Appendix Fig. S3A).

The residence of the channels in a state with a conductance of 400 (*cis*-) and 330 pS (*cis*+) was much less probable (Fig. 2E), and individual peaks of 500 and more pS were rare. Opposite to the channel formed by control dimer, this channel was less stable at *cis* positive voltage. Apparently, after the Ca^2+^-mediated assembly of PTP complex, F-ATP synthase dimer loses some components, essential for channel formation or stabilization.

The data on the capability of an F-ATP synthase monomer to form high-conductance channels are controversial [4, 10, 11]. In our experiments, monomers from control RLM samples formed Ca^2+^-activated channels, however, of lower conductance and stability than dimers (Fig. 2G, SI Appendix Fig. S4). Stable states and substates were observed both at *cis* positive and negative voltage. At negative voltage (*cis*), the highest probability of the channel residence was in conductance states of 90, 170, 290 and 430 pS; the conductance of ~650 pS and 1.1 nS was much less probable (Fig. 2H). At positive voltage (*cis*), the channels resided in the states of 100, 210, 310 and 360 pS with rare transitions to 660-pS states. The current–voltage relationship (Fig. 2I) was basically similar to that of a dimer (Fig. 2C).

Monomers from PTP samples demonstrated reduced channel activity in comparison with control samples (Fig. 2J-L, SI Appendix Fig. S5A-F). As channels formed by PTP dimer, these channels were less stable at high-conductance states and *cis* positive voltage. Channels with the highest probability resided in the states of 60 and 180 pS (*cis*-) and 10 and 100 pS (*cis*+), though individual peaks of conductance reached 0.8 and even 1.5 nS (Fig. 2K). The current–voltage relationship was essentially linear (Fig. 2L).

Thus, F-ATP synthase dimer subunits and associated proteins from control samples form large and sustained channels; monomer subunits and associated proteins from control samples compose smaller channels unstable in high-conductance states, while dimers and monomers from PTP samples create the smallest channels with the lowest stability in high-conductance states. Hence, Ca^2+^-dependent PTP opening caused substantial alterations in the bands of F-ATP synthase dimers and monomers (i.e. F-ATP synthase subunits and co-migrating proteins), which affect both specific in-gel activity and channel-forming capacity in planar lipid membranes. One can assume that Ca^2+^-dependent assembly of PTP complex in mitochondria before the separation of supercomplexes diminishes the quantity of available PTP-forming blocks: F-ATP synthase subunits and accompanying proteins. Therefore, eluates from the bands of F-ATP synthase monomers and dimers from PTP samples are devoid of essential components of PTP complex and incapable of forming high-conductance channels.

### Protein composition of the bands of F-ATP synthase monomer and dimer from control and PTP samples

Then, we examined whether the PTP-related changes in the specific and channel-forming activities of F-ATP synthase monomers and dimers are associated with alterations in the protein composition of the bands. Dimer (D) and monomer (M) bands were analyzed by tandem mass spectrometry in control and PTP samples from RLM (KCl- and SM-BM) and RHM (SM-BM) (Dataset_S01-Dataset_S06). Since the PTP is a rare or even a single object in a mitochondrion, for the analysis of MS spectra, we selected all true proteins identified at least by one unique peptide with at least 5% coverage of the protein sequence or by two unique peptides with at least 1% coverage (Dataset_S07). The abundance of a protein in a band was determined as a relative intensity-based absolute quantification (rIBAQ): the sum of peak intensities of all unique peptides of a protein (IBAQ) divided by the sum of peak intensities of all unique peptides of all true proteins in the band (Dataset_S07, sheets designated “All”). The pie diagrams show the total content of subunits of complexes I (CI), III (CIII), IV (CIV), and V (CV) and non-OXPHOS mitochondrial proteins (Other) in the bands (Fig. 3). In addition to CV subunits, D bands contained some amounts of subunits of CI, CIII, and CIV, indicating the incomplete separation from the CI–CIII_2_–CIV_x_ supercomplex, which agrees with the data of WB analysis (SI Appendix, Fig. S6). Besides, D bands comprised from 3 (RHM) to 20% (RLM) of non-OXPHOS (Other) proteins. In all mitochondrial preparations, PTP samples had a reduced percent of “other” proteins in D bands. The protein composition of M bands was much more homogenous than that of D bands: CV subunits composed 85–99% of total protein (Fig. 3, SI Appendix Fig. S6). PTP opening caused bidirectional changes in the protein composition of M bands from different mitochondrial preparations. Thus, mPTP opening caused the elimination of non-OXPHOS proteins from D bands with a minor effect on the protein composition of M bands.

**Figure 3.**
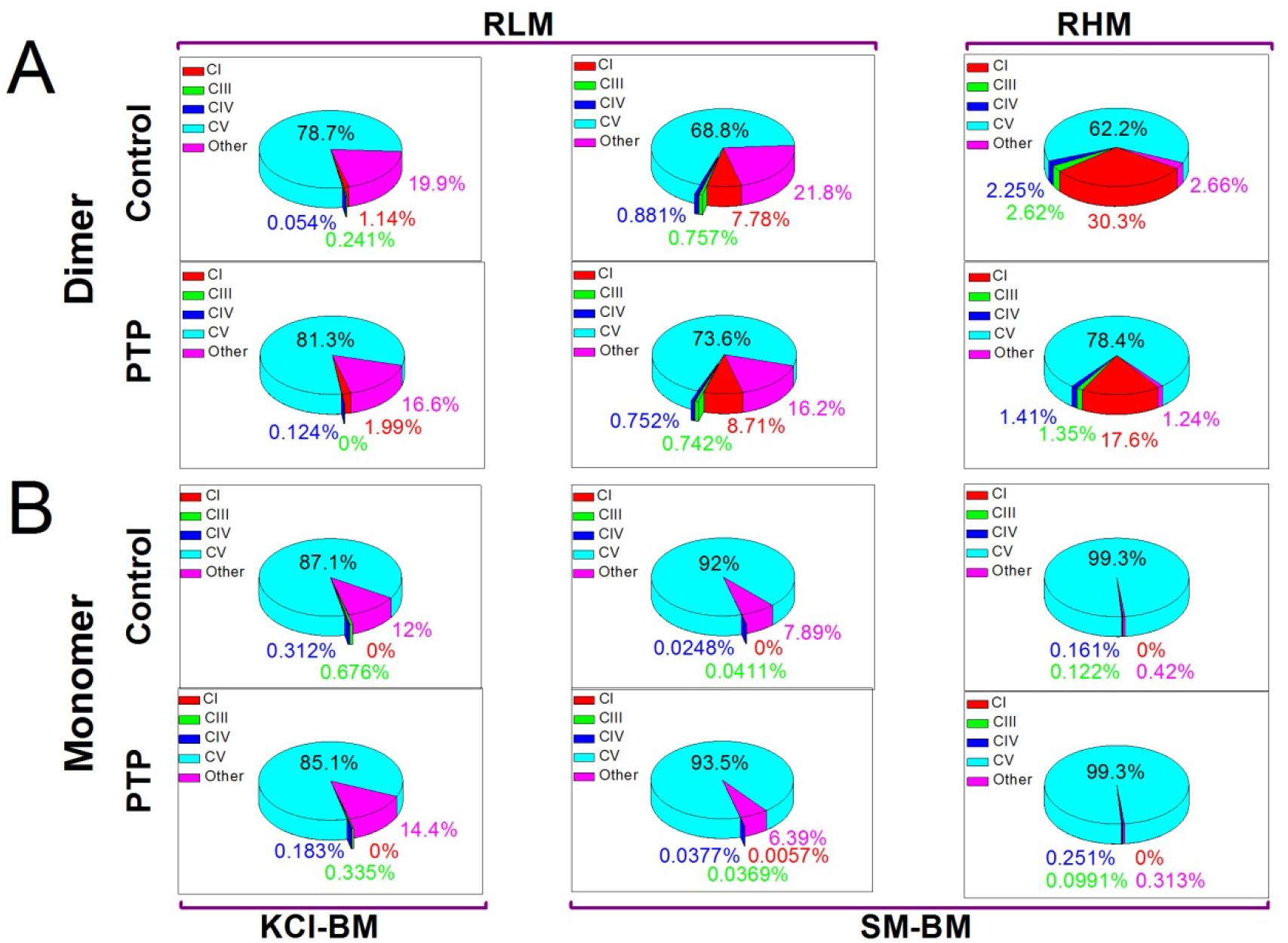
Changes in the protein composition of the bands of F-ATP synthase monomer and dimer associated with PTP opening. For each variant of mitochondria and incubation medium, dimer (A) and monomer (B) bands were collected from four BN-PAGE gels of two independent mitochondrial isolations. Each gel contained equal numbers of control and PTP samples. The total content of all peptides determined in a band by MS-MS analysis (100%) (SI Appendix Table S1) was gathered from IBAQs of peptides of complexes I (CI), III (CIII), IV (CIV), and V (CV) and non-OXPHOS proteins (Other).

It should be stressed that F-ATP synthase dimer bands did not contain mitochondrial Ca^2+^- and voltage-activated potassium and chloride channels and exchangers (VDACs, SK_Ca_, IK_Ca_, BK_Ca_, mitoK_ATP_, TASK3, mitoSLO2, mitoHCNs, mitoKv1.3, mitoKv1.5, mitoKv1.7, mitoKv7.4, CLIC1, CLIC4, CLIC 5, MCLCA1 and Letm1) (Dataset_S07) [39–42]. Other supercomplexes contained some of these proteins (SI Appendix, Table S1). Monomers from PTP samples contained a low quantity of mitoK_ATP_ (KCNJ8) and VDAC1-3 (Dataset_S07). This disproves the role of channels distinct from PTP/MMC in the observed Ca^2+^-induced channel activity.

In accordance with the data of MS-MS analysis, the signals of some proteins in D and M bands from control and PTP samples differed tens of times. To identify proteins that could be essential for channel-forming activity, we selected all proteins whose quantity unidirectionally changed in all pairs of samples: appeared/disappeared and/or increased/decreased by two and more times at least in two pairs (Dataset_S07, sheets designated “Big Difference”). The number of proteins meeting these criteria was found to be few, eight OXPHOS complex subunits and two non-OXPHOS proteins (Table 1). The content of three (NADH:ubiquinone oxidoreductase subunits A7 and A13 and F-ATP synthase subunit e) and two proteins (cytochrome b-c1 complex subunit Rieske and very long-chain specific acyl-CoA dehydrogenase) increased in dimer and monomer bands, respectively. (At least in a part, this can be connected with the partial destruction of CI–CIII_2_–CIV_x_ supercomplex due to PTP-dependent inactivation of complex I [43]. The level of five proteins (cytochrome b-c1 complex subunits 1 and 2, cytochrome c oxidase subunit 5A, methylmalonate-semialdehyde dehydrogenase [acylating] (Aldh6a1), and prohibitin 2 (Phb2)) and one protein (Aldh6a1) decreased in dimer and monomer bands, respectively. (Phb2 was not detected in M bands from both control and PTP samples.) Hence, the PTP-related decrease in the channel-forming activity of dimers and monomers correlates with a loss or absence of two proteins only: Aldh6a1 and Phb2. The level of the Phb2 partner Phb changed not so drastically, thus considerably reducing the Phb2/Phb ratio (Table 1), which indicates the destruction of the Phb-Phb2 complex. These data suggest that non-OXPHOS proteins Aldh6a1 and prohibitin(s) may be essential for PTP/MMC formation and stability.

**Table 1.** The proteins in the bands of monomers and dimers the level of which changed unidirectionally after the PTP opening. **Notes**: ND - not detected; * - absent in all monomer samples. In pairs where a protein was absent (ND) either in a control or a PTP sample, the rIBAQ value is given for the present protein.

| Protein | PTP/Control rIBAQ ratio |  |  |
| --- | --- | --- | --- |
| Dimer | RHM | RLM SM-BM | RLM KCI-BM |
| NADH:ubiquinone oxidoreductase subunit A7 | 7.25 | 1.57 | 0.008/ND |
| NADH:ubiquinone oxidoreductase subunit A13 | 1.26 | 4.51 | 0.038/ND |
| ATP synthase subunit e mitochondrial | 0.322/ND | 1.34 | 0.315/ND |
| Cytochrome b-c1 complex subunit 1 | 0.13 | 0.75 | ND/0.0044 |
| Cytochrome b-c1 complex subunit 2 | 0.49 | 0.75 | ND/0.234 |
| Cytochrome c oxidase subunit 5A | 0.19 | 0.58 | ND/0.003 |
| Methylmalonate-semialdehyde dehydrogenase [acylating] | ND/0.118 | 0.028 | 0.029 |
| Prohibitin-2 | 0.18 | 0.51 | 0.013 |
| Prohibitin | 0.507 | 0.609 | 0.834 |
| Monomer |  |  |  |
| Cytochrome b-c1 complex subunit Rieske | 2.10 | 4.21 | 1.89 |
| Very long-chain specific acyl-CoA dehydrogenase | 0.007/ND | 1.43 | 3.86 |
| Methylmalonate-semialdehyde dehydrogenase [acylating] | ND/ND | 0.40 | 0.348 |
| Prohibitin-2* | ND/ND | ND/ND | ND/ND |
| Prohibitin* | ND/ND | ND/ND | ND/ND |

## Discussion

The data presented disprove our initial hypothesis that preliminary Ca^2+^-dependent assembly of the PTP/MMC complex from F-ATP synthase and other mitochondrial proteins would increase the channel activity of isolated F-ATP synthase dimers and monomers. Though, PTP opening changed the protein composition of F-ATP synthase monomer and dimer bands (Fig. 3, Datasets_S01-S07) and, in accordance with recent observations, reduced the F-ATP synthase stability and/or in gel activity after separation (Fig. 1) [44, 45], it, however, drastically inhibited the channel activity of eluted F-ATP synthase (Fig. 2). Nevertheless, our main idea that proteins not included in the F-ATP synthase complex could participate in the PTP/MMC formation and stabilization is, presumably, correct.

Modern research revealed the role of F-ATP synthase complex and its different subunits in the PTP/MMC formation and stability in a number of protein knockdown- and knockout-based studies [5–8, 12,18, 30, 31, 46–48]. Particularly, subunits c plus δ [8], c [5, 7, 31, 46], DAPIT (k), e, f, 6.8PL (J) [8], g [8; 30], ATP6 (a) [31, 47], ATP8 (A6L) [31, 47], b [12, 30], and OSCP [12, 30] were deleted. The levels of subunits f [18, 48], c, and g [18] were decreased by siRNA or shRNA. In addition, the level of subunits ATP6 (a), ATP8 (A6L), c, d, e, f(1/2), F6, g, 6.8PL (J), DAPIT (k), OSCP, γ, δ, declined to a variable extent due to the deletion of other F-ATP synthase subunits [8, 12, 30, 31]. Although the results and their interpretations are somewhat controversial (apparently, this is due to the parameter used for the registration of PTP opening: calcium retention capacity, quenching of calcein fluorescence, mitochondrial swelling in KCl- and sucrose-based medium, swelling/shrinkage in the presence of PEGs of different size, and channel activity in mitoplasts, bilayer lipid membranes, and vesicles [5–8, 12, 18, 30, 31, 46–48]) several conclusions the on the molecular nature of PTP/MMC can be made. First, subunits g, f, and, perhaps, c are of critical importance for the PTP/MMC formation. In fact, the deletion of subunit g completely suppresses the MMC-like channel activity [30] and inhibits the swelling in KCl-BM [8, 30]. Similarly, the deletion or suppression of subunit f inhibits the swelling in KCl- and sucrose-BM [8, 30]. The deletion of subunit c weakly affects the sum Co^2+^ accumulation upon prolonged incubation [8, 31] but inhibits it in short-term experiments [5, 46]; in addition, it inhibits the swelling in sucrose medium [8] and reduces the size of MMC channels [7]. (The suppression or deletion of all these subunits does not affect the mitochondrial calcium retention capacity [8, 18, 31], presumably, because the release of calcium does not require the formation of high-conductance channels).

Second, intact F-ATP synthase monomer or dimer is, presumably, unnecessary for the assembly of PTP complex [8,10,11]. Third, PTP opening is a relatively rare event and does not require involvement of all critical F-ATP synthase subunits in a mitochondrion [34]. Indeed, even a drastic decrease in the level of some ATP subunits was insufficient for the suppression of channel activity, and their complete removal was required. In the same time, incomplete removal of some subunits decreased the integral index of high-amplitude swelling of mitochondria in sucrose-based medium or in the presence of large PEGs. [8, 12, 30, 31].

Our data agree with the view that F-ATP synthase subunits are the core components of a PTP while entire complexes are not. Indeed, only the eluates from the bands of F-ATP synthase dimers and monomers, but not from bands of CI-CIII_2_-CIV_x_ and CIII_2_-CIV_x_ supercomplexes (SI Appendix, Fig. S7) demonstrated high-conductance channel activity in planar membranes. Further, F-ATP synthase dimers and monomers from PTP samples demonstrated dramatically reduced channel activity (Fig. 2) despite the fact that they possessed the same set of subunits as control dimers and monomers (Dataset_S7). Taking into account that assembled and active F-ATP synthase is avoidable for PTP/MMC formation [8,10,11], partial inactivation of F-ATP synthase during the PTP opening and BN-PAGE (Fig. 1) cannot explain the decrease in channel forming activity in monomers and dimers from PTP samples (Fig. 2) since these bands from both control and PTP samples contain the same set of the complex subunits (Datasets_S01-S07). (Making this conclusion we assume that efficiency of protein elution from both types of samples is similar.)

Nevertheless, the changes in the protein composition of the dimer and monomers bands occurred and covered some OXPHOS subunits and few non-OXPHOS proteins (Table 1). Dimer and monomer bands accumulated complex I and complex III plus IV subunits, respectively, which, presumably connected with partial destruction of CI-CIII_2_-CIV_x_ supercomplex. (The substantial inactivation of complex I due to PTP opening was previously reported [43].) What is more important, the decline in the level of Aldh6a1 (dimer and monomer bands) and Phb2 (dimer bands; monomer bands lacked prohibitins) correlated with the decrease in the channel-forming activity of eluates from PTP samples (Fig. 2, Table 1). These data indicate that minor F-ATP synthase-associated proteins are, presumably, of critical importance for PTP/MMC complex formation, which explains a relative rareness of PTP in a mitochondrion [34]. Remarkable, purified dimers and monomers from bovine and porcine hearts [2, 4, 10] formed comparatively less stable channels than dimers of F-ATP synthase from RLM (Fig. 2), which contained a large portion of attendant proteins (Fig. 3, Dataset_S07). By stability, channels formed by dimers from RLM resemble MMC channels in rat liver mitoplasts [36–38] and mitoplasts from HeLa cells [30]. Thus, presumably, the level of minor F-ATP synthase-associated proteins but not of F-ATP synthase subunits limits the abundance of the PTP complex in a mitochondrion and, therefore, maximal purification of F-ATP synthase dimers and monomers in the studies of channel activity [2, 4, 10] is a dead end in research. On the other hand, Aldh6a1 and Phb2 are obviously insufficient for PTP/MMC formation since eluates from the bands of CI-CIII_2_-CIV_x_ and CIII_2_-CIV_x_ supercomplexes did not demonstrate high-conductance channel activity in planar membranes (SI Appendix, Fig. S7), although the bands contained Aldh6a1 and Phb2 (Datasets_S08-S10,SI Appendix, Table S2).

At the present stage of research, only a speculative model of PTP/MMC channel and mechanism of its formation can be proposed. Along with membrane F-ATP synthase subunits g and f, a stable PTP complex may comprise a matrix Aldh6a1 tetramer. In order to explain the absence of Aldh6a1 and channel activity in monomers and dimers from PTP samples, one may assume that a stable PTP complex (Aldh6a1_4_-g_n_-f_n_-subunit x(?)_n_) loosely associates with an F-ATP synthase complex and migrates separately in BN-PAGE gels. For this to happen, Ca^2+^ should weaken the interaction of F-ATP synthase subunits g and f with their partner subunits a, b, and e [49] and enhance the interaction with proximal available Aldh6a1 (Fig. 4). (Separation of membrane-immersed subcomplex from the matrix-oriented soluble subcomplex during the PTP opening was confirmed in recent works [44, 45].) If this assumption is correct, the molecular mass of the PTP complex should be at least 270–300 kDa and hardly exceed 500 kDa.

**Figure 4.**
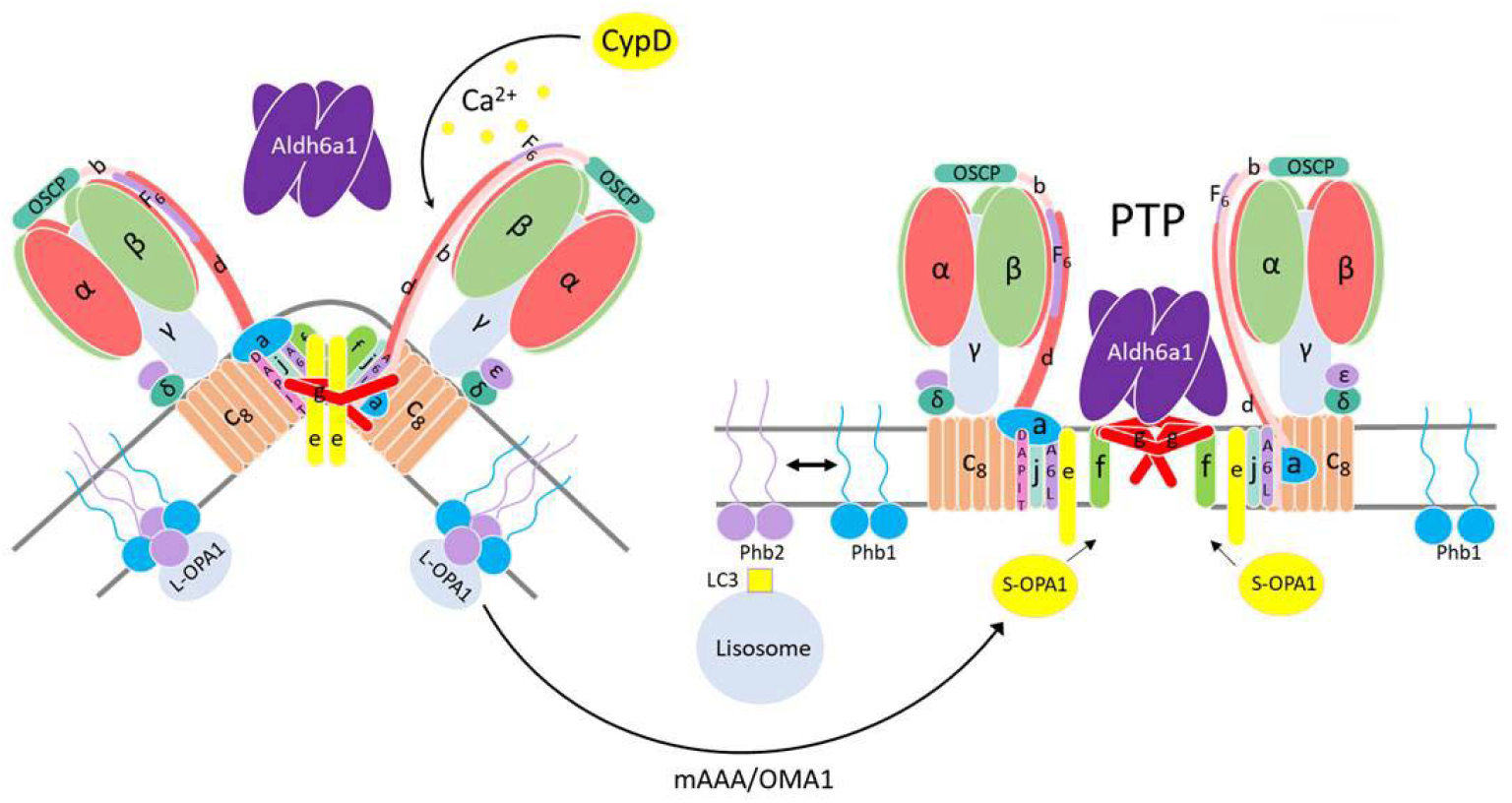
A proposed mechanism for the participation of Aldh6a1 and prohibitins in the PTP formation by subunits of F-ATP synthase dimer. See explanations in the text. mAAA and OMA1, inner mitochondrial membrane ATP-dependent metalloendopeptidases involved in the proteolytic activation of optic atrophy protein 1 (OPA1); L-OPA1 and S-OPA1, full-length and truncated OPA1 forms, respectively.

The data on the involvement of Aldh6a1 in the regulation of PTP opening or cell death are scarсe. It was shown that the expression of Aldh6a1 is reduced ten times in different hepatocellular carcinomas [50], while hepatic carcinoma mitochondria are extremely resistant to calcium [51]. The restoration of Aldh6a1 expression induced the loss of mitochondrial potential and cell death [50]. Matrix NADH inhibited both Aldh6a1 (Ki 3.1 μM) [53] and PTP opening [54]. The level of Aldh6a1 in the liver and kidney was found to be higher than in the heart and the brain [55], which correlates with the ability of mitochondria from these tissues to swell in a PTP-dependent way (Fig. 1) [55, 56]. In addition, tyrosine nitration of Aldh6a1 enhanced in rat kidney mitochondria upon the development of diabetes mellitus Type 1 [57] and acute septic damage [58], and in heart mitochondria in aging [59]. The antagonists of peroxynitrite production suppressed the nitration of Aldh6a1, damage to the kidney, and animal mortality [58]. On the other hand, preconditioning increased the level of Aldh6a1 and F-ATP synthase β subunit in heart mitochondria as well as their resistance to ischemia/reperfusion [60], though the association of Aldh6a1 with F-ATP synthase was not explored.

By contrast, the analysis of publications argues against the direct involvement of Phb2 in the PTP/MMC complex formation. First, the Phb–Phb2 complex is essential for mitochondrial resistance to PTP and cell resistance to apoptotic stimuli [52, 61, 62]. Second, the Phb–Phb2 complex increases the stability of mitochondrial supercomplexes CI–CIII_2_–CIV_x_ and CIII_2_–CIV_n_ [63, 64] and, most probably, F-ATP synthase oligomers through the stabilization of dynamin-related GTPase OPA1 [65–67]. Third, PTP-dependent collapse of the membrane potential triggers mitophagy, which requires the Phb–Phb2 complex destruction and Phb2 binding with LC3 [68, 69]. Therefore, one could assume that the destruction of Phb–Phb2 complex associated with the F-ATP synthase dimer causes the destabilization of the latter and facilitates PTP complex formation [11] (Fig. 4). Concomitantly, liberated Phb2 becomes a receptor that activates the elimination of a damaged mitochondrion.

To conclude, here we describe the effect of the preliminary PTP opening in mitochondria on the protein composition of F-ATP synthase dimer and monomer BN-PAGE bands and high-conductance channel-forming ability of eluates from these bands. The data obtained indicate that proteins accompanying F-ATP synthase, methylmalonate-semialdehyde dehydrogenase (Aldh6a1) and prohibitin 2 (Phb2), may be important players in the PTP/MMC formation and stabilization.

## Materials and Methods

A more detailed description of materials and methods can be found in SI Appendix, Supplementary Materials and Methods.

### Isolation of rat liver and heart mitochondria

All manipulations with animals were performed in accordance with the Helsinki Declaration of 1975 (revised in 1983), national requirements for the care and use of laboratory animals, and protocol 9/2020 of 17.02.2020 approved by the Commission on Biological Safety and Bioethics at the ITEB RAS. Rat liver mitochondria (RLM) were isolated by a standard differential centrifugation procedure [1]. The homogenization medium contained 220 mM mannitol, 70 mM sucrose, 10 mM HEPES (pH adjusted to 7.4 with Trizma Base), 1 mM EGTA, and 0.05% BSA. The mitochondrial pellet was washed three times with a medium devoid of EGTA and BSA. Final pellets were resuspended in this medium to yield ~70 mg protein/ml. Rat heart mitochondria (RHM) were isolated in the same buffer by a similar way except that the mitochondrial pellet was washed twice. The concentration of isolated RHM was approximately 20 mg protein/ml. Mitochondrial protein was assayed by the Biuret method using BSA as a standard [70].

Mitochondria were incubated at 37°C either in a KCl-based medium (KCl-BM) (125 mM KCl, 20 mM sucrose, 10 mM HEPES (pH adjusted to 7.3 with Trizma Base), 2 mM KH_2_PO_4_, 2 mM MgCl_2_, and 10 μM EGTA) or in a sucrose-mannitol-based medium (SM-BM) (220 mM mannitol, 70 mM sucrose, 10 mM HEPES (pH 7.3), 2 mM KH_2_PO_4_, 2 mM MgCl_2_, and 10 μM EGTA) supplemented with 5 mM malate plus 5 mM pyruvate. The quality of isolated mitochondria, indicated by the respiratory control coefficient, was assessed using Oroboros Oxygraph-2k (Austria) by measuring the oxygen consumption rates before, in the course, and after the termination of phosphorylation of known amounts of ADP. In the studies, the respiratory control coefficient was not less than 5.

### Registration of PTP opening

The opening of PTP in isolated mitochondria was registered as EGTA- and CsA-sensitive high-amplitude swelling. Mitochondrial swelling was determined by measuring a decrease in A_550_ in mitochondrial suspension using a plate reader (Infinite 200 Tecan, Austria) and 96-well plates.

### Blue native electrophoresis (BN-PAGE)

BN-PAGE was performed as described [71]. The solubilizing buffer (0.75 M aminocapronic acid, 50 mM Bis-Tris/HCl (pH 7.0), and 10% digitonin (Sigma, Saint-Louis, Missouri, USA)) was added to mitochondria sedimented from control and PTP samples, and the suspension was kept on ice for 20 min. After 10-min centrifugation at 10000 × g, the supernatant was supplemented with 5% Serva Blue G dissolved in 1 M aminocapronic acid. Samples were applied onto 3%–13% gradient gel, 70 μg of the sample per lane. Electrophoresis was performed on ice at 0–4°C. An HMW Calibration Kit for Native Electrophoresis (Sigma-Aldrich, St. Louis, MO, USA) was used as a marker of molecular mass. Two gels were run simultaneously, one of which was used for in-gel activity determination, and the other for SDS-PAGE and immunoblotting.

### Measurement of the in-gel activity of electron transport chain complexes and F-ATP synthase

The in-gel activity of complexes I, IV, and F-ATP synthase (CI, CIV, and CV) was determined as described [71]. To assess CI activity, gels were stained for ~10–30 min with a buffer containing 100 mM Tris-HCl (pH 7.4), 0.14 mM NADH, and nitro blue tetrazolium chloride (1 mg/ml). To determine CIV activity, gels were stained for 1 h with a buffer containing 10 mM KH_2_PO_4_ (рН 7.4), 3,3′-diaminobenzidine (1 mg/ml), and cytochrome с (0.2 mg/ml). The measurement of ATPase activity included the staining of gels for 16 h with a buffer containing 270 mM glycine, 35 mM Tris (HCL) (pH 7.0), 14 mM MgSO_4_, 10 mM ATP, and 0.2% Pb(NO_3_)_2_. After the incubation of gels with appropriate substrates, the reactions were stopped with 10% acetic acid; gels were washed with water and scanned. CIII activity was defined in accordance with [72].

### SDS-PAGE and immunoblotting

The bands of separated complexes were cut out from BN-PAGE gels and applied onto 12.5% SDS-PAGE slabs followed by electrophoresis and immunoblotting. The molecular mass of proteins was calibrated using Precision Plus Pre-stained Standards markers from Bio-Rad Laboratories (Hercules, CA, USA).

The subunits of ETC complexes were detected using a Total Oxphos Rodent WB Antibody Cocktail (Abcam, Cambridge, UK). The cocktail contained the antibodies against F-ATP synthase subunit alpha (CV-ATP5A-55 kDa), cytochrome b-c1 complex subunit 2 (CIII-UQCRC2-48 kDa), mitochondrially encoded cytochrome c oxidase subunit I (CIV-MTCO1-40 kDa), succinate dehydrogenase [ubiquinone] iron-sulfur subunit b (CII-SDHB-30 kDa), and NADH dehydrogenase (ubiquinone) 1 beta subcomplex subunit 8 (CI-NDUFB8-20 kDa). Besides, F-ATP synthase subunit c was detected using the ATP5G antibody from Abcam (Cambridge, UK). Immunoreactivity was studied using appropriate secondary antibodies conjugated to horseradish peroxidase (Jackson Immuno Research, West Grove, PA, USA). The blots were stained with ECL (Bio-Rad, Hercules, CA, USA) and inspected using the ChemiDoc Touch Imaging System (Bio-Rad, Hercules, CA, USA).

### Tandem mass spectrometry (MS-MS) analysis

For each variant of mitochondria and incubation medium, the bands of F-ATP synthase dimer and monomer, complex I, and mitochondrial supercomplexes CI-CIII-CIVn+ CI-CIII-CIVn, and CIII-CIVn were collected from four BN-PAGE gels of two independent mitochondrial isolations. Each gel contained the same number of control and PTP samples. Proteins bands were excised and treated with trypsin (Sigma) at 37°C in a Thermo Mixer thermo shaker (Eppendorf, Germany). The molar enzyme-to-protein ratio was 1/50. The reaction was stopped by the addition of trifluoroacetic acid to the solution. Prior to mass spectrometric analysis, the peptides were separated by reversed-phase HPLC using an Easy-nLC 1000 Nanoflow chromatograph (Thermo Fisher Scientific, USA). The separation was carried out in a –adsorbent with a particle size of 3.6 μm and a pore size of 300 Å. The column was packed under laboratory conditions at a pressure of 350 atm. The peptides were eluted in a gradient of acetonitrile from 5% to 45% for 180 min; the mobile phase flow rate was 0.27 μl/min.

The mass spectra of samples were obtained using an OrbiTrap Elite mass spectrometer (Thermo Scientific, Germany). The method of ionization of peptides was nanoelectrospray. The temperature of the input capillary was 220°C, the voltage between the emitter and the input capillary was 1.9 kV. The panoramic mass spectra were shot with a resolution of 60,000 (for m/z 400). The spectra were recorded in the range of 300–1600 m/z. Ion fragmentation was carried out by the collision-activated dissociation with an inert gas in a high-energy cell (HCD). The resolution of the device when scanning the fragmentation spectra was 15,000.

### MS data analysis and protein quantification

The results of MS-MS analysis were processed using the commercial programs Thermo Xcalibur Qual Browser and PEAKS Studio 7.5/Xpro based on the provided sequences of mitochondrial proteins. The masses of the peptides were determined with an accuracy of at least 5 ppm. The masses of fragments were determined with an accuracy of 0.1 Da. For the analysis of PTP-associated changes in the protein composition of the bands, we selected all true proteins if they were identified at least by one unique peptide with at least 5% coverage of protein sequence or by two unique peptides with at least 1% coverage (Datasets_S01-S07). The abundance of a protein in the band was determined as a relative intensity-based absolute quantification (rIBAQ): the sum of peak intensities of all unique peptides of a protein (IBAQ) divided by the sum of peak intensities of all unique peptides of all true proteins in the band (Dataset_S07).

### Electrophysiology

F-ATP synthase dimers and monomers were eluted from excised protein bands of BN-PAGE gels exactly as described by [4]. The elution buffer contained 25 mM tricine, 10 mM MgSO4, and 7.5 mM Bis-Tris (pH 7.0), with addition of 8 mM ATP-Tris (pH 7.4) and 1% w/v n-heptyl β-D-thioglucopyranoside. After overnight rotation at 4°C, the eluate was centrifuged at 20,000 × *g* for 20 min at 4°C, and the supernatants were collected for reconstitution in electrophysiological analyses. The elution buffer for other mitochondrial complexes and supercomplexes contained 2.5 mM NADH instead of ATP. The electrophysiological properties of mitochondrial complexes and supercomplexes were studied after their insertion into artificial planar lipid membranes. Bilayer lipid membranes were formed according to Muller–Rudin [70] using a solution containing 20 g/l of soybean lecithin (L-α-phosphatidylcholine Type IV–S, Sigma, USA). The surrounding solution contained 0.2 М KCl and 5 mM HEPES-Tris (Sigma), pH 7.3. Voltage clamp conditions were used throughout the experiments. The *cis* compartment was connected to the virtual ground through a Keithle 301 amplifier in the current-to-voltage configuration. The membrane potential was maintained through Ag/AgCl electrodes in 3 M KCl and 2% agar bridges. The data were digitized using a DT2801A board (Data Translation, USA) via AD–DA converters connected to a PC, which collected the data using the software developed in-house by A.Ya. Silberstein. The agent under study was introduced into the chamber part with the measuring electrode at the same side (*trans*) of the membrane. Ca^2+^ was added on the side of mitochondrial complexes. The sign of the potential on the figures refers to the compartment of the cuvette (*cis*). All measurements were performed at room temperature (~22°C).

### Statistics

Data of in-gel activity staining and channel formation are representative of at least three independent experiments. The amplitude histograms of conductance summarize the data of three standard records from independent experiments. Values on the curves of current-voltage relationships are means ± S.E.M. (n = 3) of three experiments. MS data are presented for three Control/PTP pairs of samples; each sample included material from two independent mitochondrial isolations.

## Supporting information

RHM Control Dimer of V compl.

RHM Control Monomer of V compl.

RHM PTP Dimer of V compl.

RHM PTP Monomer of V compl.

RLM KCl-BM control Dimer of V compl.

RLM KCl-BM control Monomer of V compl.

RLM KCl-BM PTP Dimer of V compl.

RLM KCl-BM PTP Monomer of V compl.

RLM SM-BM Control Dimer of V compl.

RLM SM-BM Control Monomer of V compl.

RLM SM-BM PTP Dimer of V compl.

RLM SM-BM PTP Monomer of V compl.

ATP-synthase

ATP-synthase

ATP-synthase

ATP-synthase

ATP-synthase

ATP-synthase

ATP-synthase

ATP-synthase

ATP-synthase

ATP-synthase

ATP-synthase

ATP-synthase

RHM

RHM

RHM

RHM

RHM

RHM

RLM SM-BM

RLM SM-BM

RLM SM-BM

RLM SM-BM

RLM SM-BM

RLM SM-BM

RLM KCl-BM

RLM KCl-BM

RLM KCl-BM

RLM KCl-BM

RLM KCl-BM

RLM KCl-BM

SI Kruglov

## Acknowledgments

This work was supported by a grant to A.K. from the Russian Foundation for Basic Research (RFBR)(19-04-00327a). We sincerely thank Pavel Grigoriev from the Institute of Cell Biophysics for expert advice.

## References

1. Winquist R.J., Gribkoff V.K. Targeting putative components of the mitochondrial permeability transition pore for novel therapeutics. Biochem Pharmacol 177, 113995 (2020).

2. Urbani A., Giorgio V., Carrer A., Franchin C., Arrigoni G., Jiko C., Abe K., Maeda S., Shinzawa-Itoh K., Bogers J.F.M., McMillan D.G.G., Gerle C., Szabò I., Bernardi P. Purified F-ATP synthase forms a Ca^2+^-dependent high-conductance channel matching the mitochondrial permeability transition pore. Nat Commun 10(1), 4341 (2019).

3. Chinopoulos C. Mitochondrial permeability transition pore: Back to the drawing board. Neurochem Int 117, 49–54 (2018).

4. Giorgio V., von Stockum S., Antoniel M., Fabbro A., Fogolari F., Forte M., Glick G.D., Petronilli V., Zoratti M., Szabó I., Lippe G., Bernardi P. Dimers of mitochondrial ATP synthase form the permeability transition pore. Proc Natl Acad Sci U S A 110, 5887–5892 (2013).

5. Alavian K.N., Beutner G., Lazrove E., Sacchetti S., Park H.A., Licznerski P., Li H., Nabili P., Hockensmith K., Graham M., Jr. Porter G.A., Jonas E.A. An uncoupling channel within the c-subunit ring of the F1FO ATP synthase is the mitochondrial permeability transition pore. Proc Natl Acad Sci U S A 111, 10580–10585 (2014).

6. Carraro M., Giorgio V., Šileikyte J., Sartori G., Forte M., Lippe G., Zoratti M., Szabó I., Bernardi P. Channel Formation by Yeast F-ATP Synthase and the Role of Dimerization in the Mitochondrial Permeability Transition. J Biol Chem 289, 15980–15985 (2014).

7. Neginskaya M.A., Solesio M.E., Berezhnaya E.V., Amodeo G.F., Mnatsakanyan N., Jonas E.A., Pavlov E.V. ATP Synthase C-Subunit-Deficient Mitochondria Have a Small Cyclosporine A-Sensitive Channel, but Lack the Permeability Transition Pore. Cell Rep 26(1), 11–17 (2019).

8. Carroll J., He J., Ding S., Fearnley I.M., Walker J.E. Persistence of the permeability transition pore in human mitochondria devoid of an assembled ATP synthase. Proc Natl Acad Sci U S A 116(26), 12816–12821 (2019).

9. Carraro M., Checchetto V., Sartori G., Kucharczyk R., di Rago J.P., Minervini G., Franchin C., Arrigoni G., Giorgio V., Petronilli V., Tosatto S.C.E., Lippe G., Szabó I., Bernardi P. High-Conductance Channel Formation in Yeast Mitochondria is Mediated by F-ATP Synthase e and g Subunits. Cell Physiol Biochem 50(5), 1840–1855 (2018).

10. Mnatsakanyan N., Llaguno M.C., Yang Y., Yan Y., Weber J., Sigworth F.J., Jonas E.A. A mitochondrial megachannel resides in monomeric F_1_F_O_ ATP synthase. Nat Commun 10(1), 5823 (2019).

11. Bonora M., Morganti C., Morciano G., Pedriali G., Lebiedzinska-Arciszewska M., Aquila G., Giorgi C., Rizzo P., Campo G., Ferrari R., Kroemer G., Wieckowski M.R., Galluzzi L., Pinton P. Mitochondrial permeability transition involves dissociation of F_1_F_O_ ATP synthase dimers and C-ring conformation. EMBO Rep 18(7), 1077–1089 (2017).

12. He J., Carroll J., Ding S., Fearnley I.M., Walker J.E. Permeability transition in human mitochondria persists in the absence of peripheral stalk subunits of ATP synthase. Proc Natl Acad Sci U S A 114(34), 9086–9091 (2017).

13. Bernardi P. Mechanisms for Ca^2+^-dependent permeability transition in mitochondria. Proc Natl Acad Sci U S A 117(6), 2743–2744 (2020).

14. Zamzami N., Kroemer G. The mitochondrion in apoptosis: how Pandora’s box opens. Nat Rev Mol Cell Biol. 2(1), 67–71 (2001).

15. Szabo I., Zoratti M. The mitochondrial permeability transition pore may comprise VDAC molecules I. Binary structure and voltage dependence of the pore. FEBS 330(2), 201–205 (1993).

16. Szabo I., De Pinto V., Zoratti M. The mitochondrial permeability transition molecules pore may comprise VDAC II. The electrophysiological properties of VDAC are compatible with those of the mitochondrial megachannel. FEBS 330(2), 206–210 (1993).

17. Leung A.W.C., Varanyuwatana P., Halestrap A.P. The Mitochondrial Phosphate Carrier Interacts with Cyclophilin D and May Play a Key Role in the Permeability Transition. J Biol Chem. 283(39), 26312–26323 (2008).

18. Shanmughapriya S., Rajan S., Hoffman N.E., Higgins A.M., Tomar D., Nemani N., Hines K.J., Smith D.J., Eguchi A., Vallem S., Shaikh F., Cheung M., Leonard N.J., Stolakis R.S., Wolfers M.P., Ibetti J., Chuprun J.K., Jog N.R., Houser S.R., Koch W.J., Elrod J.W., Madesh M. SPG7 Is an Essential and Conserved Component of the Mitochondrial Permeability Transition Pore. Mol Cell. 60(1), 47–62 (2015).

19. Basso E., Fante L., Fowlkes J., Petronilli V., Forte M.A., Bernardi P. Properties of the permeability transition pore in mitochondria devoid of Cyclophilin D. J Biol Chem. 280(19), 18558–18561 (2005).

20. Lohret T.A., Murphy R.C., Drgoñ T., Kinnally K.W. Activity of the mitochondrial multiple conductance channel is independent of the adenine nucleotide translocator. J. Biol. Chem. 271(9), 4846–4849 (1996).

21. Kokoszka J.E., Waymire K.J., Levy S.E., Sligh J.E., Cai J., Jones D.P., MacGregor G.R., Wallace D.C. The ADP/ATP translocator is not essential for the mitochondrial permeability transition pore. Nature 427, 461–465 (2004).

22. Baines C.P., Kaiser R.A., Sheiko T., Craigen W.J., Molkentin J.D. Voltage-dependent anion channels are dispensable for mitochondrial-dependent cell death. Nature Cell Biol. 9 (5), 550–555 (2007).

23. Krauskopf A., Eriksson O., Craigen W.J., Forte M.A., Bernardi P. Properties of the permeability transition in VDAC1(−/−) mitochondria. Biochim. Biophys. Acta 1757 590–595 (2006).

24. Herick K., Krämer R., Lühring H. Patch clamp investigation into the phosphate carrier from Saccharomyces cerevisiae mitochondria. Biochim Biophys Acta. 1321(3), 207–20 (1997).

25. Gutierrez-Aguilar M., Douglas D.L., Gibson A.K., Domeier T.L., Molkentin J.D., Baines S.P. Genetic manipulation of the cardiac mitochondrial phosphate carrier does not affect permeability transition. J. Mol. Cell. Cardiol. 72, 316–325 (2014).

26. König T., Tröder S.E., Bakka K., Korwitz A., Richter-Dennerlein R., Lampe P.A., Patron M., Mühlmeister M., Guerrero-Castillo S., Brandt U., Decker T., Lauria I., Paggio A., Rizzuto R., Rugarli E.I., De Stefani D., Langer T. The m-AAA Protease Associated with Neurodegeneration Limits MCU Activity in Mitochondria. Mol Cell 64, 1–15 (2016).

27. Giorgio V., Burchell V., Schiavone M, Bassot C., Minervini G., Petronilli V., Argenton F., Forte M., Tosatto S., Lippe G., Bernardi P. Ca2+ binding to F-ATP synthase β subunit triggers the mitochondrial permeability transition. EMBO Rep. 18(7), 1065–1076 (2017).

28. Guo L., Carraro M., Carrer A., Minervini G., Urbani A., Masgras I., Tosatto S.C.E., Szabò I., Bernardi P., Lippe G.. Arg-8 of yeast subunit e contributes to the stability of F-ATP synthase dimers and to the generation of the full-conductance mitochondrial megachannel. J Biol Chem.; 294(28), 10987–10997 (2019).

29. Arnold I., Pfeiffer K., Neupert W., Stuart R.A., Schägger H. Yeast mitochondrial F1F0-ATPsynthase exists as a dimer: identification of three dimer-specific subunits. EMBOJ. 17, 7170–7178 (1998).

30. Carrer A., Tommasin L., Šileikytė J., Ciscato F., Filadi R., Urbani A., Forte M.l., Rasola, I. Szabò A., Carraro M., Bernardi P. Defining the molecular mechanisms of the mitochondrial permeability transition through genetic manipulation of F-ATP synthase. Nat. Commun. 12, 4835 (2021).

31. He J., Ford H.C., Carroll J., Ding S, Fearnley I.M., Walker J.E. Persistence of the mitochondrial permeability transition in the absence of subunit c of human ATP synthase. Proc Natl Acad Sci U S A. 114(13), 3409–3414 (2017).

32. Neginskaya M.A., Strubbe J.O., Amodeo G.F., West B.A., Yakar S., Bazil J.N., Pavlov E.V. The very low number of calcium-induced permeability transition pores in the single mitochondrion. J Gen Physiol 152(10), 202012631 (2020).

33. Al-Nasser I., Crompton M. The reversible Ca2+-induced permeabilization of rat liver mitochondria. Biochem. J. 239, 19–29 (1986).

34. Novgorodov S.A., Gudz T.I., Milgrom Y.M., Brierley G.P. The permeability transition in heart mitochondria is regulated synergistically by ADP and cyclosporin A. J Biol Chem., 267(23), 16274–16262 (1992).

35. Broekemeier K.M., Klocek C.K., Pfeiffer D.R. Proton selective substate of the mitochondrial permeability transition pore: regulation by the redox state of the electron transport chain. Biochemistry 37(38), 13059–13065 (1998).

36. Kinnally K.W., Campo M.L., Tedeschi H. Mitochondrial channel activity studied by patch-clamping mitoplasts. Journal of Bioenergetics and Biomembranes 4, 497–506 (1989).

37. Szabó I., Zoratti M. The mitochondrial megachannel is the permeability transition pore. Journal of Bioenergetics and Biomembranes 24(1), 111–117 (1992).

38. Szabó I., Bernardi P., Zoratti M. Modulation of the mitochondrial megachannel by divalent cations and protons. J Biol Chem. 267(5), 2940–2946 (1992).

39. Ponnalagu D., Singh H. Anion Channels of Mitochondria. Handb Exp Pharmacol. 240, 71–101 (2017).

40. Leanza L, Checchetto V., Biasutto L, Rossa A, Costa R, Bachmann M., Zoratti M., Szabo Pharmacological modulation of mitochondrial ion channels. BJP 176(22), 4258–4283 (2019).

41. Wrzosek A, Augustynek B., Żochowska M, Szewczyk A. Mitochondrial Potassium Channels as Druggable Targets. Biomolecules 10(8), 1200 (2020).

42. Kravenska Y, Checchetto V., Szabo I. Routes for Potassium Ions across Mitochondrial Membranes: A Biophysical Point of View with Special Focus on the ATP-Sensitive K ^+^ Channel. Biomolecules 11(8), 1172 (2021).

43. Batandier C., Leverve X., Fontaine E. Opening of the mitochondrial permeability transition pore induces reactive oxygen species production at the level of the respiratory chain complex I. J Biol Chem. 279(17), 17197–204 (2004).

44. Mnatsakanyan N., Park H.A., Wu J., He X., Llaguno M.C., Latta M., Miranda P., Murtishi B., Graham M., Weber J., Levy R.J., Pavlov E.V., Jonas E.A. Mitochondrial ATP synthase c-subunit leak channel triggers cell death upon loss of its F_1_ subcomplex. Cell Death Differ. 29(9), 1874–1887 (2022).

45. Pinke G., Zhou L., Sazanov L.A. Cryo-EM structure of the entire mammalian F-type ATP synthase. Nat Struct Mol Biol. 27(11), 1077–1085 (2020).

46. Bonora M., Bononi A., De Marchi E.,Giorgi C., Lebiedzinska M., Marchi S., Patergnani S., Rimessi A., Suski J.M., Wojtala A., Wieckowski M.R., Kroemer G., Galluzzi L., Pinton P. Role of the c subunit of the F0 ATP synthase in mitochondrial permeability transition. Cell. Cycle. 12(4), 674–683 (2013).

47. Masgras I., Rasola A., Bernardi P. Induction of the permeability transition pore in cells depleted of mitochondrial DNA. Biochim Biophys Acta. 1817(10), 1860–6 (2012).

48. Galber C., Minervini G., Cannino G., Boldrin F., Petronilli V., Tosatto S., Lippe G., Giorgio V. The f subunit of human ATP synthase is essential for normal mitochondrial morphology and permeability transition. Cell Rep. 35(6), 109111 (2021).

49. Spikes T.E., Montgomery M.G., Walker J.E. Structure of the dimeric ATP synthase from bovine mitochondria. Proc Natl Acad Sci U S A. 117(38), 23519–23526 (2020).

50. Shin H., Cha H.J., Lee M.J., Na K., Park D., Kim C.Y., Han D.H., Kim H., Paik Y.K. Identification of ALDH6A1 as a Potential Molecular Signature in Hepatocellular Carcinoma via Quantitative Profiling of the Mitochondrial Proteome. J Proteome Res. 19(4), 1684–1695 (2020).

51. Thorne R.F., Bygrave F.L. The role of mitochondria in modifying the cellular ionic environment. Calcium-induced respiratory activities in mitochondria isolated from various tumour cells. Biochem J. 144(3), 551–8 (1974).

52. Gomez L., Paillard M., Price M., Chen Q., Teixeira G., Spiegel S., Lesnefsky E.J. A novel role for mitochondrial sphingosine-1-phosphate produced by sphingosine kinase-2 in PTP-mediated cell survival during cardioprotection. Basic Res Cardiol. 106(6), 1341–53 (2011).

53. Goodwin G. W., Rougraff P. M., Davis E. J., Harris R. A. Purification and characterization of methylmalonate-semialdehyde dehydrogenase from rat liver. Identity to malonate-semialdehyde dehydrogenase. J Biol Chem. 264(25), 14965–71 (1989).

54. Haworth R.A., Hunter D.R. Allosteric inhibition of the Ca2+-activated hydrophilic channel of the mitochondrial inner membrane by nucleotides. J Membr Biol. 54(3), 231–6 (1980).

55. Kedishvili N. Y., Popov K. M., Rougraff P. M., Zhao Y., Crabb D. W., Harris R. A. CoA-dependent methylmalonate-semialdehyde dehydrogenase, a unique member of the aldehyde dehydrogenase superfamily. cDNA cloning, evolutionary relationships, and tissue distribution. J Biol Chem. 267(27), 19724–9 (1992).

56. Berman S.B., Watkins S.C., Hastings T.G. Quantitative biochemical and ultrastructural comparison of mitochondrial permeability transition in isolated brain and liver mitochondria: evidence for reduced sensitivity of brain mitochondria. Exp Neurol. 164(2), 415–25 (2000).

57. Ishii N., Carmines P.K., Yokoba M, Imaizumi H., Ichikawa T., Ikenagasa H., Kodera Y., Oh-Ishi M., Aoki Y., Maeda T., Takenaka T., Katagiri M. Angiotensin-converting enzyme inhibition curbs tyrosine nitration of mitochondrial proteins in the renal cortex during the early stage of diabetes mellitus in rats. Clin Sci (Lond). 124(8), 543–52 (2013).

58. Seija M., Baccino C., Nin N., Sánchez-Rodríguez C., Granados R., Ferruelo A., Martínez-Caro L., Ruíz-Cabello J., de Paula M., Noboa O., Esteban A., Lorente J.A.. Role of peroxynitrite in sepsis-induced acute kidney injury in an experimental model of sepsis in rats. Shock. 38(4), 403–10 (2012).

59. Kanski J., Behring A., Pelling J., Schöneich C. Proteomic identification of 3-nitrotyrosine-containing rat cardiac proteins: effects of biological aging. Am J Physiol Heart Circ Physiol. 288(1), H371–81 (2005).

60. Zhu W-Z., Wu X-F., Zhang Y., Zhou Z-N. Proteomic analysis of mitochondrial proteins in cardiomyocytes from rats subjected to intermittent hypoxia. Eur J Appl Physiol. 112(3), 1037–46 (2012).

61. Toki S., Yoshimaru T., Matsushita Y., Aihara H., Ono M., Tsuneyama K., Sairyo K., Katagiri T. The survival and proliferation of osteosarcoma cells are dependent on the mitochondrial BIG3-PHB2 complex formation. Cancer Sci. 112(10), 4208–4219 (2021).

62. Xu Y., Wang J., Xu W., Ding F., Ding W. Prohibitin 2-mediated mitophagy attenuates renal tubular epithelial cells injury by regulating mitochondrial dysfunction and NLRP3 inflammasome activation. Am J Physiol Renal Physiol 316(2), F396–F407 (2019).

63. Jian C., Xu F., Hou T., Sun T., Li J., Cheng H., Wang X. Deficiency of PHB complex impairs respiratory supercomplex formation and activates mitochondrial flashes. J. Cell Sci. 130, 2620–2630 (2017).

64. Anderson C.J., Kahl A., Qian L., Stepanova A., Starkov A., Manfredi G., Iadecola C., Zhou P. Prohibitin is a positive modulator of mitochondrial function in PC12 cells under oxidative stress. J. Neurochem. 146, 235–250 (2018).

65. Merkwirth C., Dargazanli S., Tatsuta T., Geimer S., Löwer B., Wunderlich F.T., von Kleist-Retzow J.C., Waisman A., Westermann B., Langer T. Prohibitins control cell proliferation and apoptosis by regulating OPA1-dependent cristae morphogenesis in mitochondria. Genes Dev. 22, 476–488 (2008).

66. Cho S.G., Xiao X., Wang S., Gao H., Rafikov R., Black S., Huang S., Ding H.F., Yoon Y., Kirken R.A., Yin X.M., Wang H.G., Dong Z. Bif-1 Interacts with Prohibitin-2 to Regulate Mitochondrial Inner Membrane during Cell Stress and Apoptosis. J Am Soc Nephrol. 30(7):1174–1191 (2019).

67. Quintana-Cabrera R., Quirin C., Glytsou C., Corrado M., Urbani A., Pellattiero A., Calvo E., Vázquez J., Enríquez J.A., Gerle C. The cristae modulator Optic atrophy 1 requires mitochondrial ATP synthase oligomers to safeguard mitochondrial function. Nat. Commun. 9, 3399 (2018).

68. Wei Y., Chiang W.C., Sumpter R.Jr., Mishra P., Levine B. Prohibitin 2 Is an Inner Mitochondrial Membrane Mitophagy Receptor. Cell 168, 224–238 (2017).

69. Xiao Y., Zhou Y., Lu Y., Zhou K., Cai W. PHB2 interacts with LC3 and SQSTM1 is required for bile acids-induced mitophagy in cholestatic liver. Cell Death Dis. 9(2), 160 (2018).

70. Johnson D., Lardy H. Isolation of liver or kidney mitochondria. Methods in Enzymology, 10, 94–96 (1967).

71. Jha P, Wang X., Auwerx J. Analysis of Mitochondrial Respiratory Chain Supercomplexes Using Blue Native Polyacrylamide Gel Electrophoresis (BN-PAGE) Curr Protoc Mouse Biol. 6(1), 1–14 (2016).

72. Smet J., De Paepe B., Seneca S., Lissens W., Kotarsky H., De Meirleir L., Fellman V., Van Coster R. Complex III staining in blue native polyacrylamide gels. J Inherit Metab Dis. 34(3), 741–7 (2011).

73. Mueller P., Rudin D.O., Tien H.T., Wescott W.C. Reconstitution of cell membrane structure in vitro and its transformation into an excitable system. Nature 194, 979–980 (1962).

