## Supplementary material for "Permeability transition pore-related changes in the proteome and channel activity of ATP synthase dimers and monomers": RHM Control Dimer of V compl.

| Accession | Protein Group | Protein ID | -10lgP | Coverage (%) | Coverage (%) Sample 4 | Area Sample 4 | #Peptides | #Unique | #Spec Sample 4 | PTM | Avg. Mass | Description |
|---|---|---|---|---|---|---|---|---|---|---|---|---|
| P10719\|ATPB_RAT | **2** | 1 | 378.4 | 50 | 50 | 345830000 | 124 | 124 | 582 | Y | 56354 | ATP synthase subunit beta, mitochondrial OS=Rattus norvegicus OX=1011 |
| tr\|G3V6D3\|G3V6D3 | **2** | 2 | 378.4 | 50 | 50 | 345830000 | 124 | 124 | 582 | Y | 56345 | ATP synthase subunit beta OS=Rattus norvegicus OX=10116 GN=Atp5f1b |
| P15999\|ATPA_RAT | **1** | 3 | 312.23 | 48 | 48 | 267690000 | 108 | 108 | 601 | Y | 59754 | ATP synthase subunit alpha, mitochondrial OS=Rattus norvegicus OX=101 |
| tr\|F1LP05\|F1LP05_I | **1** | 4 | 312.23 | 48 | 48 | 267690000 | 108 | 108 | 601 | Y | 59813 | ATP synthase subunit alpha OS=Rattus norvegicus OX=10116 GN=Atp5f1 |
| Q66HF1\|NDUS1_R | **3** | 3421 | 260.31 | 28 | 28 | 48874000 | 56 | 56 | 173 | Y | 79412 | NADH-ubiquinone oxidoreductase 75 kDa subunit, mitochondrial OS=Rattu |
| P31399\|ATP5H_RA | **9** | 7 | 247.72 | 53 | 53 | 20584000 | 21 | 21 | 67 | Y | 18763 | ATP synthase subunit d, mitochondrial OS=Rattus norvegicus OX=10116 ( |
| P35435\|ATPG_RAT | **6** | 5 | 246.27 | 34 | 34 | 22396000 | 26 | 26 | 89 | Y | 30191 | ATP synthase subunit gamma, mitochondrial OS=Rattus norvegicus OX=1 |
| tr\|Q6QI09\|Q6QI09_I | **6** | 6 | 246.27 | 15 | 15 | 22396000 | 26 | 26 | 89 | Y | 67721 | ATP synthase subunit gamma, mitochondrial OS=Rattus norvegicus OX=1 |
| P19234\|NDUV2_RA | **4** | 2870 | 232.09 | 39 | 39 | 66119000 | 31 | 31 | 121 | N | 27378 | NADH dehydrogenase [ubiquinone] flavoprotein 2, mitochondrial OS=Rattu |
| Q5BK63\|NDUA9_R | **7** | 5406 | 228.99 | 38 | 38 | 15159000 | 31 | 30 | 81 | N | 42559 | NADH dehydrogenase [ubiquinone] 1 alpha subcomplex subunit 9, mitoch |
| tr\|B2RYS8\|B2RYS8 | **15** | 2857 | 219.65 | 23 | 23 | 19269000 | 13 | 13 | 38 | Y | 21959 | NADH dehydrogenase [ubiquinone] 1 beta subcomplex subunit 8, mitocho |
| Q641Y2\|NDUS2_R | **5** | 2856 | 210.73 | 30 | 30 | 34707000 | 32 | 32 | 101 | Y | 52562 | NADH dehydrogenase [ubiquinone] iron-sulfur protein 2, mitochondrial OS |
| tr\|D3ZG43\|D3ZG43 | **8** | 2807 | 206.44 | 43 | 43 | 23602000 | 35 | 35 | 79 | N | 30226 | NADH dehydrogenase (Ubiquinone) Fe-S protein 3 (Predicted), isoform CR |
| tr\|D3ZF13\|D3ZF13_ | **13** | 3210 | 194.65 | 28 | 28 | 14275000 | 17 | 17 | 49 | Y | 17514 | Acyl carrier protein OS=Rattus norvegicus OX=10116 GN=Ndufab1 PE=1 |
| Q68FY0\|QCR1_RA | **14** | 118 | 176.06 | 23 | 23 | 10042000 | 23 | 23 | 47 | Y | 52849 | Cytochrome b-c1 complex subunit 1, mitochondrial OS=Rattus norvegicus |
| tr\|D4A565\|D4A565_ | **26** | 5412 | 173.26 | 34 | 34 | 6636500 | 15 | 15 | 24 | N | 21664 | NADH dehydrogenase (Ubiquinone) 1 beta subcomplex, 5 (Predicted), isof |
| tr\|D4A7L4\|D4A7L4_ | **17** | 2872 | 172.5 | 32 | 32 | 14159000 | 13 | 13 | 36 | Y | 17634 | NADH dehydrogenase (Ubiquinone) 1 beta subcomplex, 11 (Predicted) OS |
| tr\|D3ZFQ8\|D3ZFQ8 | **27** | 133 | 167.41 | 12 | 12 | 7580300 | 7 | 7 | 23 | Y | 35435 | Cytochrome c-1 OS=Rattus norvegicus OX=10116 GN=Cyc1 PE=1 SV=3 |
| tr\|B0BNE6\|B0BNE6 | **21** | 5411 | 167.03 | 26 | 26 | 7454200 | 13 | 13 | 32 | Y | 23970 | NADH dehydrogenase (Ubiquinone) Fe-S protein 8 (Predicted), isoform CR |
| P32551\|QCR2_RAT | **10** | 34 | 164.71 | 18 | 18 | 15525000 | 22 | 22 | 55 | N | 48396 | Cytochrome b-c1 complex subunit 2, mitochondrial OS=Rattus norvegicus |
| tr\|Q5XIH3\|Q5XIH3_ | **11** | 5408 | 161.2 | 22 | 22 | 11086000 | 17 | 17 | 52 | N | 50731 | NADH dehydrogenase [ubiquinone] flavoprotein 1, mitochondrial OS=Rattu |
| tr\|Q5PQZ9\|Q5PQZ9 | **16** | 2821 | 154.05 | 27 | 27 | 18181000 | 10 | 10 | 37 | N | 14359 | NADH dehydrogenase [ubiquinone] 1 subunit C2 OS=Rattus norvegicus O |
| P00507\|AATM_RAT | **30** | 19 | 153.42 | 16 | 16 | 2596800 | 10 | 10 | 20 | Y | 47314 | Aspartate aminotransferase, mitochondrial OS=Rattus norvegicus OX=101 |
| Q80W89\|NDUAB_R | **29** | 5440 | 151.01 | 19 | 19 | 6841500 | 7 | 7 | 22 | N | 14854 | NADH dehydrogenase [ubiquinone] 1 alpha subcomplex subunit 11 OS=Ra |
| tr\|A0A0G2JVL6\|A0A | **20** | 3028 | 148.03 | 29 | 29 | 8295600 | 11 | 11 | 32 | N | 19965 | NADH dehydrogenase [ubiquinone] 1 alpha subcomplex subunit 8 OS=Rat |
| tr\|D4A0T0\|D4A0T0_ | **28** | 180 | 142.75 | 21 | 21 | 12726000 | 9 | 9 | 22 | N | 20859 | NADH:ubiquinone oxidoreductase subunit B10 OS=Rattus norvegicus OX= |
| P11240\|COX5A_RA | **25** | 86 | 141.75 | 42 | 42 | 4296900 | 9 | 9 | 25 | N | 16130 | Cytochrome c oxidase subunit 5A, mitochondrial OS=Rattus norvegicus O |
| tr\|Q06QK5\|Q06QK5 | **23** | 5413 | 140.47 | 9 | 9 | 8536000 | 11 | 11 | 27 | Y | 68606 | NADH-ubiquinone oxidoreductase chain 5 OS=Rattus norvegicus OX=1011 |
| P11661\|NU5M_RAT | **23** | 5414 | 140.47 | 9 | 9 | 8536000 | 11 | 11 | 27 | Y | 68618 | NADH-ubiquinone oxidoreductase chain 5 OS=Rattus norvegicus OX=1011 |
| tr\|Q06QG6\|Q06QG6 | **23** | 5415 | 140.47 | 9 | 9 | 8536000 | 11 | 11 | 27 | Y | 68588 | NADH-ubiquinone oxidoreductase chain 5 OS=Rattus norvegicus OX=1011 |
| tr\|Q06QA1\|Q06QA1 | **23** | 5416 | 140.47 | 9 | 9 | 8536000 | 11 | 11 | 27 | Y | 68574 | NADH-ubiquinone oxidoreductase chain 5 OS=Rattus norvegicus OX=1011 |
| tr\|Q8SEZ0\|Q8SEZ0 | **23** | 5417 | 140.47 | 9 | 9 | 8536000 | 11 | 11 | 27 | Y | 68618 | NADH-ubiquinone oxidoreductase chain 5 OS=Rattus norvegicus OX=1011 |
| tr\|A0A096XKT9\|A0A | **23** | 5418 | 140.47 | 9 | 9 | 8536000 | 11 | 11 | 27 | Y | 68584 | NADH-ubiquinone oxidoreductase chain 5 OS=Rattus norvegicus OX=1011 |
| tr\|G3V7Y3\|G3V7Y3_ | **31** | 12 | 135.48 | 33 | 33 | 5030800 | 8 | 8 | 18 | Y | 17563 | ATP synthase subunit delta, mitochondrial OS=Rattus norvegicus OX=101 |
| tr\|Q5UAJ6\|Q5UAJ6 | **24** | 24 | 133.46 | 32 | 32 | 6358700 | 14 | 14 | 26 | Y | 25942 | Cytochrome c oxidase subunit 2 OS=Rattus norvegicus OX=10116 GN=C |
| Q561S0\|NDUAA_R | **18** | 5410 | 130.85 | 33 | 33 | 8793300 | 16 | 16 | 35 | Y | 40493 | NADH dehydrogenase [ubiquinone] 1 alpha subcomplex subunit 10, mitoc |
| tr\|F1LXA0\|F1LXA0_ | **48** | 5458 | 126.04 | 17 | 17 | 5432200 | 3 | 3 | 8 | N | 17178 | NADH dehydrogenase [ubiquinone] 1 alpha subcomplex subunit 12 OS=Ra |
| tr\|A0A411P2I9\|A0A | **33** | 39 | 125.19 | 13 | 13 | 3117800 | 8 | 8 | 17 | N | 40144 | Cytochrome b (Fragment) OS=Rattus norvegicus OX=10116 GN=Cytb PE= |
| tr\|A0A0U2IDR8\|A0A | **33** | 40 | 125.19 | 13 | 13 | 3117800 | 8 | 8 | 17 | N | 40981 | Cytochrome b (Fragment) OS=Rattus norvegicus OX=10116 PE=3 SV=1 |
| tr\|A0A0U2M179\|A0A | **33** | 41 | 125.19 | 13 | 13 | 3117800 | 8 | 8 | 17 | N | 41037 | Cytochrome b (Fragment) OS=Rattus norvegicus OX=10116 PE=3 SV=1 |
| tr\|A0A411P2L0\|A0A | **33** | 42 | 125.19 | 13 | 13 | 3117800 | 8 | 8 | 17 | N | 41017 | Cytochrome b (Fragment) OS=Rattus norvegicus OX=10116 GN=Cytb PE= |
| tr\|F8QU31\|F8QU31_ | **33** | 43 | 125.19 | 13 | 13 | 3117800 | 8 | 8 | 17 | N | 42181 | Cytochrome b (Fragment) OS=Rattus norvegicus OX=10116 GN=cytb PE= |
| tr\|A0A411P2K0\|A0A | **33** | 44 | 125.19 | 13 | 13 | 3117800 | 8 | 8 | 17 | N | 42238 | Cytochrome b (Fragment) OS=Rattus norvegicus OX=10116 GN=Cytb PE= |
| tr\|L0L4L8\|L0L4L8_F | **33** | 45 | 125.19 | 12 | 12 | 3117800 | 8 | 8 | 17 | N | 42470 | Cytochrome b (Fragment) OS=Rattus norvegicus OX=10116 GN=cytb PE= |
| tr\|A0A140GE10\|A0A | **33** | 46 | 125.19 | 12 | 12 | 3117800 | 8 | 8 | 17 | N | 42574 | Cytochrome b (Fragment) OS=Rattus norvegicus OX=10116 PE=3 SV=1 |
| tr\|A0A140GE11\|A0A | **33** | 47 | 125.19 | 12 | 12 | 3117800 | 8 | 8 | 17 | N | 42588 | Cytochrome b (Fragment) OS=Rattus norvegicus OX=10116 PE=3 SV=1 |
| tr\|A0A140GE08\|A0A | **33** | 48 | 125.19 | 12 | 12 | 3117800 | 8 | 8 | 17 | N | 42527 | Cytochrome b (Fragment) OS=Rattus norvegicus OX=10116 PE=3 SV=1 |
| tr\|A0A385HCC9\|A0A | **33** | 49 | 125.19 | 12 | 12 | 3117800 | 8 | 8 | 17 | N | 42766 | Cytochrome b (Fragment) OS=Rattus norvegicus OX=10116 PE=3 SV=1 |
| tr\|A0A3Q8AGE8\|A0 | **33** | 50 | 125.19 | 12 | 12 | 3117800 | 8 | 8 | 17 | N | 42712 | Cytochrome b (Fragment) OS=Rattus norvegicus OX=10116 GN=Cytb PE= |
| tr\|A0A3Q8AC68\|A0A | **33** | 38 | 125.19 | 12 | 12 | 3117800 | 8 | 8 | 17 | N | 42622 | Cytochrome b (Fragment) OS=Rattus norvegicus OX=10116 GN=Cytb PE= |
| tr\|A0A3S6FK13\|A0A | **33** | 51 | 125.19 | 12 | 12 | 3117800 | 8 | 8 | 17 | N | 42702 | Cytochrome b (Fragment) OS=Rattus norvegicus OX=10116 GN=Cytb PE= |
| tr\|A0A3S6FM15\|A0A | **33** | 52 | 125.19 | 12 | 12 | 3117800 | 8 | 8 | 17 | N | 42698 | Cytochrome b (Fragment) OS=Rattus norvegicus OX=10116 GN=Cytb PE= |
| tr\|A0A097PE40\|A0A | **33** | 53 | 125.19 | 12 | 12 | 3117800 | 8 | 8 | 17 | N | 42865 | Cytochrome b OS=Rattus norvegicus OX=10116 GN=CYTB PE=3 SV=1 |
| tr\|A0A220D9Z6\|A0A | **33** | 54 | 125.19 | 12 | 12 | 3117800 | 8 | 8 | 17 | N | 43018 | Cytochrome b OS=Rattus norvegicus OX=10116 PE=3 SV=1 |
| tr\|D6NSP2\|D6NSP2 | **33** | 55 | 125.19 | 12 | 12 | 3117800 | 8 | 8 | 17 | N | 42945 | Cytochrome b (Fragment) OS=Rattus norvegicus OX=10116 GN=cytb PE= |
| tr\|D6NSS6\|D6NSS6 | **33** | 56 | 125.19 | 12 | 12 | 3117800 | 8 | 8 | 17 | N | 42993 | Cytochrome b (Fragment) OS=Rattus norvegicus OX=10116 GN=cytb PE= |

| | | | | | | | | | | | | |
|---|---|---|---|---|---|---|---|---|---|---|---|---|
| tr\|D6NSQ3\|D6NSQ3 | **33** | 57 | 125.19 | 12 | 12 | 3117800 | 8 | 8 | 17 | N | 43012 | Cytochrome b (Fragment) OS=Rattus norvegicus OX=10116 GN=cytb PE= |
| tr\|D6NSR3\|D6NSR3 | **33** | 58 | 125.19 | 12 | 12 | 3117800 | 8 | 8 | 17 | N | 42986 | Cytochrome b (Fragment) OS=Rattus norvegicus OX=10116 GN=cytb PE= |
| tr\|D6NSP4\|D6NSP4 | **33** | 59 | 125.19 | 12 | 12 | 3117800 | 8 | 8 | 17 | N | 43016 | Cytochrome b (Fragment) OS=Rattus norvegicus OX=10116 GN=cytb PE= |
| tr\|D6NSQ8\|D6NSQ8 | **33** | 60 | 125.19 | 12 | 12 | 3117800 | 8 | 8 | 17 | N | 43016 | Cytochrome b (Fragment) OS=Rattus norvegicus OX=10116 GN=cytb PE= |
| tr\|D6NSR8\|D6NSR8 | **33** | 61 | 125.19 | 12 | 12 | 3117800 | 8 | 8 | 17 | N | 42998 | Cytochrome b (Fragment) OS=Rattus norvegicus OX=10116 GN=cytb PE= |
| tr\|F2Q6S6\|F2Q6S6_ | **33** | 63 | 125.19 | 12 | 12 | 3117800 | 8 | 8 | 17 | N | 42938 | Cytochrome b (Fragment) OS=Rattus norvegicus OX=10116 GN=cytb PE= |
| tr\|A0A0A1FZ42\|A0A | **33** | 64 | 125.19 | 12 | 12 | 3117800 | 8 | 8 | 17 | N | 42989 | Cytochrome b OS=Rattus norvegicus OX=10116 GN=CYTB PE=3 SV=1 |
| tr\|D6NSS7\|D6NSS7 | **33** | 65 | 125.19 | 12 | 12 | 3117800 | 8 | 8 | 17 | N | 42948 | Cytochrome b (Fragment) OS=Rattus norvegicus OX=10116 GN=cytb PE= |

| | | | | | | | | | | | | |
|---|---|---|---|---|---|---|---|---|---|---|---|---|
| tr\|Q8SEY9\|Q8SEY9 | **33** | 66 | 125.19 | 12 | 12 | 3117800 | 8 | 8 | 17 | N | 43015 | Cytochrome b OS=Rattus norvegicus OX=10116 GN=cytb PE=3 SV=1 |
| tr\|D6NSR7\|D6NSR7 | **33** | 67 | 125.19 | 12 | 12 | 3117800 | 8 | 8 | 17 | N | 42982 | Cytochrome b (Fragment) OS=Rattus norvegicus OX=10116 GN=cytb PE= |
| tr\|A0A220DA44\|A0A | **33** | 68 | 125.19 | 12 | 12 | 3117800 | 8 | 8 | 17 | N | 42968 | Cytochrome b OS=Rattus norvegicus OX=10116 PE=3 SV=1 |
| tr\|F2Q6S5\|F2Q6S5_ | **33** | 69 | 125.19 | 12 | 12 | 3117800 | 8 | 8 | 17 | N | 42952 | Cytochrome b (Fragment) OS=Rattus norvegicus OX=10116 GN=cytb PE= |
| tr\|D6NSQ6\|D6NSQ6 | **33** | 70 | 125.19 | 12 | 12 | 3117800 | 8 | 8 | 17 | N | 42952 | Cytochrome b (Fragment) OS=Rattus norvegicus OX=10116 GN=cytb PE= |
| tr\|D6NSQ0\|D6NSQ0 | **33** | 71 | 125.19 | 12 | 12 | 3117800 | 8 | 8 | 17 | N | 42968 | Cytochrome b (Fragment) OS=Rattus norvegicus OX=10116 GN=cytb PE= |
| tr\|Q5UAI7\|Q5UAI7_ | **33** | 72 | 125.19 | 12 | 12 | 3117800 | 8 | 8 | 17 | N | 42998 | Cytochrome b OS=Rattus norvegicus OX=10116 GN=CYTB PE=3 SV=1 |
| tr\|D6NSR2\|D6NSR2 | **33** | 73 | 125.19 | 12 | 12 | 3117800 | 8 | 8 | 17 | N | 43073 | Cytochrome b (Fragment) OS=Rattus norvegicus OX=10116 GN=cytb PE= |
| tr\|A0A0S1Z1V9\|A0A | **33** | 74 | 125.19 | 12 | 12 | 3117800 | 8 | 8 | 17 | N | 43002 | Cytochrome b OS=Rattus norvegicus OX=10116 GN=CYTB PE=3 SV=1 |
| tr\|D6NSR1\|D6NSR1 | **33** | 75 | 125.19 | 12 | 12 | 3117800 | 8 | 8 | 17 | N | 43002 | Cytochrome b (Fragment) OS=Rattus norvegicus OX=10116 GN=cytb PE= |
| tr\|Q8HIC4\|Q8HIC4_ | **33** | 76 | 125.19 | 12 | 12 | 3117800 | 8 | 8 | 17 | N | 43012 | Cytochrome b OS=Rattus norvegicus OX=10116 GN=Mt-cyb PE=3 SV=1 |
| tr\|A0A220DA02\|A0A | **33** | 77 | 125.19 | 12 | 12 | 3117800 | 8 | 8 | 17 | N | 42968 | Cytochrome b OS=Rattus norvegicus OX=10116 PE=3 SV=1 |
| tr\|A0A0S1Z1V4\|A0A | **33** | 78 | 125.19 | 12 | 12 | 3117800 | 8 | 8 | 17 | N | 43016 | Cytochrome b OS=Rattus norvegicus OX=10116 GN=CYTB PE=3 SV=1 |
| tr\|R9TKN1\|R9TKN1 | **33** | 119 | 125.19 | 12 | 12 | 3117800 | 8 | 8 | 17 | N | 43012 | Cytochrome b OS=Rattus norvegicus OX=10116 GN=CYTB PE=3 SV=1 |
| P00159\|CYB_RAT | **33** | 79 | 125.19 | 12 | 12 | 3117800 | 8 | 8 | 17 | N | 43012 | Cytochrome b OS=Rattus norvegicus OX=10116 GN=Mt-Cyb PE=3 SV=3 |
| tr\|L0N311\|L0N311_ | **33** | 80 | 125.19 | 12 | 12 | 3117800 | 8 | 8 | 17 | N | 43045 | Cytochrome b (Fragment) OS=Rattus norvegicus OX=10116 GN=cytb PE= |
| tr\|A0A220DA28\|A0A | **33** | 81 | 125.19 | 12 | 12 | 3117800 | 8 | 8 | 17 | N | 42970 | Cytochrome b OS=Rattus norvegicus OX=10116 PE=3 SV=1 |
| tr\|H2KXA0\|H2KXA0 | **33** | 82 | 125.19 | 12 | 12 | 3117800 | 8 | 8 | 17 | N | 42978 | Cytochrome b (Fragment) OS=Rattus norvegicus OX=10116 GN=cytb PE= |
| tr\|A0A096XNM4\|A0A | **33** | 83 | 125.19 | 12 | 12 | 3117800 | 8 | 8 | 17 | N | 43042 | Cytochrome b (Fragment) OS=Rattus norvegicus OX=10116 PE=3 SV=1 |
| tr\|D6NSP5\|D6NSP5 | **33** | 120 | 125.19 | 12 | 12 | 3117800 | 8 | 8 | 17 | N | 43012 | Cytochrome b (Fragment) OS=Rattus norvegicus OX=10116 GN=cytb PE= |
| P67779\|PHB_RAT | **32** | 5407 | 124.95 | 32 | 32 | 2115400 | 11 | 11 | 18 | N | 29820 | Prohibitin OS=Rattus norvegicus OX=10116 GN=Phb PE=1 SV=1 |
| Q06647\|ATPO_RAT | **22** | 8 | 124.51 | 26 | 26 | 7095700 | 10 | 10 | 30 | N | 23398 | ATP synthase subunit O, mitochondrial OS=Rattus norvegicus OX=10116 ( |
| P19511\|AT5F1_RAT | **12** | 10 | 124.23 | 17 | 17 | 27671000 | 10 | 10 | 49 | N | 28869 | ATP synthase F(0) complex subunit B1, mitochondrial OS=Rattus norvegic |
| Q6PDU7\|ATP5L_RA | **19** | 11 | 122.89 | 39 | 39 | 6407800 | 9 | 9 | 33 | Y | 11433 | ATP synthase subunit g, mitochondrial OS=Rattus norvegicus OX=10116 ( |
| P20788\|UCRI_RAT | **44** | 132 | 117.18 | 15 | 15 | 997010 | 5 | 5 | 9 | N | 29446 | Cytochrome b-c1 complex subunit Rieske, mitochondrial OS=Rattus norve |
| P10817\|CX6A2_RA | **54** | 151 | 116.98 | 14 | 14 | 1422000 | 3 | 3 | 6 | N | 10487 | Cytochrome c oxidase subunit 6A2, mitochondrial (Fragment) OS=Rattus r |
| tr\|G3V8M4\|G3V8M4 | **54** | 152 | 116.98 | 13 | 13 | 1422000 | 3 | 3 | 6 | N | 10802 | Cytochrome c oxidase subunit 6A, mitochondrial OS=Rattus norvegicus OX |
| tr\|D3ZE15\|D3ZE15_ | **39** | 5447 | 112.42 | 26 | 26 | 1490000 | 8 | 8 | 11 | N | 16777 | NADH:ubiquinone oxidoreductase subunit A13 OS=Rattus norvegicus OX= |
| tr\|B5DEL8\|B5DEL8_ | **37** | 5442 | 111.74 | 17 | 17 | 3348400 | 3 | 3 | 12 | N | 12700 | NADH dehydrogenase (Ubiquinone) Fe-S protein 5 OS=Rattus norvegicus |
| tr\|A0A0G2JZJ9\|A0A | **37** | 5443 | 111.74 | 17 | 17 | 3348400 | 3 | 3 | 12 | N | 12730 | Uncharacterized protein OS=Rattus norvegicus OX=10116 PE=4 SV=1 |
| Q63362\|NDUA5_RA | **45** | 5439 | 111.71 | 20 | 20 | 1791600 | 4 | 4 | 9 | N | 13412 | NADH dehydrogenase [ubiquinone] 1 alpha subcomplex subunit 5 OS=Rat |
| tr\|A9UMW2\|A9UMW | **43** | 5453 | 104.79 | 34 | 34 | 7056100 | 7 | 7 | 9 | Y | 9255 | Ndufa3 protein (Fragment) OS=Rattus norvegicus OX=10116 GN=Ndufa3 |
| tr\|M0RB63\|M0RB63 | **43** | 5454 | 104.79 | 33 | 33 | 7056100 | 7 | 7 | 9 | Y | 9372 | RCG63041 OS=Rattus norvegicus OX=10116 GN=LOC684509 PE=4 SV=1 |
| tr\|A0A0G2KAA3\|A0A | **43** | 5455 | 104.79 | 31 | 31 | 7056100 | 7 | 7 | 9 | Y | 10170 | NADH:ubiquinone oxidoreductase subunit A3 OS=Rattus norvegicus OX=1 |
| tr\|A0A0G2KB63\|A0A | **34** | 5422 | 102.71 | 13 | 13 | 3542500 | 5 | 5 | 15 | N | 33168 | Prohibitin OS=Rattus norvegicus OX=10116 GN=Phb2 PE=1 SV=1 |
| Q5XIH7\|PHB2_RAT | **34** | 5423 | 102.71 | 13 | 13 | 3542500 | 5 | 5 | 15 | N | 33312 | Prohibitin-2 OS=Rattus norvegicus OX=10116 GN=Phb2 PE=1 SV=1 |
| tr\|F1LPG5\|F1LPG5_ | **63** | 5445 | 101.71 | 19 | 19 | 560640 | 3 | 3 | 4 | N | 15064 | NADH:ubiquinone oxidoreductase subunit B4 OS=Rattus norvegicus OX=1 |
| Q64428\|ECHA_RAT | **42** | 9 | 96.05 | 6 | 6 | 1180300 | 7 | 7 | 10 | N | 82665 | Trifunctional enzyme subunit alpha, mitochondrial OS=Rattus norvegicus ( |
| tr\|D3ZS58\|D3ZS58_ | **41** | 5449 | 91.1 | 23 | 23 | 1593600 | 3 | 3 | 11 | N | 10845 | NADH dehydrogenase [ubiquinone] 1 alpha subcomplex subunit 2 OS=Rat |
| tr\|Q5RJN0\|Q5RJN0 | **35** | 564 | 89.03 | 15 | 15 | 2707400 | 4 | 4 | 13 | Y | 23945 | NADH dehydrogenase (Ubiquinone) Fe-S protein 7 OS=Rattus norvegicus |
| tr\|D3ZZ21\|D3ZZ21_ | **40** | 5448 | 86.16 | 26 | 26 | 1908900 | 4 | 4 | 11 | Y | 15638 | NADH dehydrogenase (Ubiquinone) 1 beta subcomplex, 6 (Predicted) OS= |
| tr\|Q5UAJ5\|Q5UAJ5 | **46** | 32 | 80.38 | 19 | 19 | 7031700 | 3 | 3 | 9 | Y | 7642 | ATP synthase protein 8 OS=Rattus norvegicus OX=10116 GN=ATP8 PE=3 |
| P11608\|ATP8_RAT | **46** | 35 | 80.38 | 19 | 19 | 7031700 | 3 | 3 | 9 | Y | 7630 | ATP synthase protein 8 OS=Rattus norvegicus OX=10116 GN=Mt-atp8 PE |
| tr\|Q8SEZ4\|Q8SEZ4 | **46** | 33 | 80.38 | 19 | 19 | 7031700 | 3 | 3 | 9 | Y | 7632 | ATP synthase protein 8 OS=Rattus norvegicus OX=10116 GN=ATPase8 PE |
| tr\|Q8HIC8\|Q8HIC8_ | **46** | 36 | 80.38 | 19 | 19 | 7031700 | 3 | 3 | 9 | Y | 7630 | ATP synthase protein 8 OS=Rattus norvegicus OX=10116 GN=Mt-atp8 PE |
| Q9ER34\|ACON_RA | **59** | 88 | 78.75 | 4 | 4 | 649530 | 3 | 3 | 5 | N | 85433 | Aconitate hydratase, mitochondrial OS=Rattus norvegicus OX=10116 GN= |
| Q02253\|MMSA_RA | **53** | 2801 | 77.13 | 8 | 8 | 1358900 | 5 | 5 | 6 | Y | 57808 | Methylmalonate-semialdehyde dehydrogenase [acylating], mitochondrial C |
| tr\|G3V7J0\|G3V7J0_ | **53** | 2802 | 77.13 | 8 | 8 | 1358900 | 5 | 5 | 6 | Y | 57748 | Aldehyde dehydrogenase family 6, subfamily A1, isoform CRA_b OS=Rattu |
| tr\|A9UMV9\|A9UMV9 | **55** | 5450 | 76.99 | 21 | 21 | 1509700 | 3 | 3 | 6 | N | 12500 | NADH:ubiquinone oxidoreductase subunit A7 OS=Rattus norvegicus OX=1 |
| Q5XIF3\|NDUS4_RA | **51** | 5457 | 75.88 | 12 | 12 | 973700 | 5 | 5 | 7 | N | 19741 | NADH dehydrogenase [ubiquinone] iron-sulfur protein 4, mitochondrial OS |
| P80432\|COX7C_RA | **65** | 134 | 71.8 | 21 | 21 | 321550 | 2 | 2 | 4 | N | 7375 | Cytochrome c oxidase subunit 7C, mitochondrial OS=Rattus norvegicus OX |
| tr\|Q8SEZ8\|Q8SEZ8 | **38** | 671 | 68.86 | 6 | 6 | 2854700 | 5 | 5 | 11 | N | 36133 | NADH-ubiquinone oxidoreductase chain 1 (Fragment) OS=Rattus norvegic |
| P03889\|NU1M_RAT | **38** | 672 | 68.86 | 6 | 6 | 2854700 | 5 | 5 | 11 | N | 36145 | NADH-ubiquinone oxidoreductase chain 1 OS=Rattus norvegicus OX=1011 |
| tr\|Q8HID1\|Q8HID1_ | **38** | 673 | 68.86 | 6 | 6 | 2854700 | 5 | 5 | 11 | N | 36145 | NADH-ubiquinone oxidoreductase chain 1 OS=Rattus norvegicus OX=1011 |
| tr\|D2E6L7\|D2E6L7_ | **38** | 5456 | 68.86 | 6 | 6 | 2854700 | 5 | 5 | 11 | N | 36062 | NADH-ubiquinone oxidoreductase chain 1 OS=Rattus norvegicus OX=1011 |
| P10888\|COX41_RA | **47** | 290 | 68.37 | 12 | 12 | 2010600 | 3 | 3 | 8 | N | 19515 | Cytochrome c oxidase subunit 4 isoform 1, mitochondrial OS=Rattus norve |
| Q6AXV4\|SAM50_RA | **72** | 2803 | 64.55 | 4 | 4 | 240260 | 2 | 2 | 3 | N | 51960 | Sorting and assembly machinery component 50 homolog OS=Rattus norve |
| P17764\|THIL_RAT | **61** | 2787 | 62.78 | 8 | 8 | 1100400 | 3 | 3 | 4 | N | 44695 | Acetyl-CoA acetyltransferase, mitochondrial OS=Rattus norvegicus OX=10 |
| Q60587\|ECHB_RAT | **58** | 20 | 62.75 | 7 | 7 | 432720 | 4 | 4 | 5 | N | 51414 | Trifunctional enzyme subunit beta, mitochondrial OS=Rattus norvegicus O |
| tr\|D4A3V2\|D4A3V2_ | **88** | 5424 | 61.72 | 8 | 8 | 262320 | 2 | 2 | 2 | N | 15224 | NADH dehydrogenase [ubiquinone] 1 alpha subcomplex subunit 6 OS=Rat |
| P12075\|COX5B_RA | **52** | 139 | 61.53 | 11 | 11 | 3573300 | 3 | 3 | 7 | N | 13915 | Cytochrome c oxidase subunit 5B, mitochondrial OS=Rattus norvegicus OX |
| tr\|A0A0A1G491\|A0A | **50** | 1300 | 57.87 | 10 | 10 | 721790 | 4 | 4 | 7 | N | 38542 | NADH-ubiquinone oxidoreductase chain 2 OS=Rattus norvegicus OX=1011 |

| | | | | | | | | | | | | |
|---|---|---|---|---|---|---|---|---|---|---|---|---|
| tr\|Q5UAJ8\|Q5UAJ8_ | **50** | 1301 | 57.87 | 10 | 10 | 721790 | 4 | 4 | 7 | N | 38455 | NADH-ubiquinone oxidoreductase chain 2 OS=Rattus norvegicus OX=1011 |
| tr\|A0A0G2JVH4\|A0/ | **71** | 147 | 57.78 | 2 | 2 | 73134 | 2 | 2 | 3 | N | 86230 | MICOS complex subunit MIC60 OS=Rattus norvegicus OX=10116 GN=Imr |
| tr\|A0A140TAG5\|A0/ | **71** | 145 | 57.78 | 2 | 2 | 73134 | 2 | 2 | 3 | N | 67049 | MICOS complex subunit MIC60 OS=Rattus norvegicus OX=10116 GN=Imr |
| Q3KR86\|MIC60_RA | **71** | 146 | 57.78 | 2 | 2 | 73134 | 2 | 2 | 3 | N | 67177 | MICOS complex subunit Mic60 (Fragment) OS=Rattus norvegicus OX=101 |

| | | | | | | | | | | | | |
|---|---|---|---|---|---|---|---|---|---|---|---|---|
| tr\|D4A3E8\|D4A3E8_ | **74** | 7776 | 55.17 | 3 | 3 | 238180 | 1 | 1 | 3 | N | 47648 | Mitochondrial ribosomal protein S27 OS=Rattus norvegicus OX=10116 GN |
| tr\|D3ZCZ9\|D3ZCZ9_ | **57** | 5435 | 52.17 | 21 | 21 | 1520800 | 2 | 2 | 5 | N | 13040 | NADH dehydrogenase [ubiquinone] iron-sulfur protein 6, mitochondrial OS |
| P02563\|MYH6_RAT | **60** | 196 | 51.56 | 1 | 1 | 2297500 | 3 | 3 | 4 | Y | 223506 | Myosin-6 OS=Rattus norvegicus OX=10116 GN=Myh6 PE=1 SV=2 |
| tr\|Q7H115\|Q7H115_ | **66** | 2862 | 49.12 | 3 | 3 | 151450 | 1 | 1 | 4 | N | 29871 | Cytochrome c oxidase subunit 3 OS=Rattus norvegicus OX=10116 GN=Mt |
| tr\|I6V4L9\|I6V4L9_R | **66** | 2863 | 49.12 | 3 | 3 | 151450 | 1 | 1 | 4 | N | 29844 | Cytochrome c oxidase subunit 3 OS=Rattus norvegicus OX=10116 GN=CC |
| tr\|Q8M7G4\|Q8M7G | **66** | 2864 | 49.12 | 3 | 3 | 151450 | 1 | 1 | 4 | N | 29861 | Cytochrome c oxidase subunit 3 OS=Rattus norvegicus OX=10116 PE=2 S |
| tr\|A0A096XKT2\|A0A | **66** | 2865 | 49.12 | 3 | 3 | 151450 | 1 | 1 | 4 | N | 29901 | Cytochrome c oxidase subunit 3 OS=Rattus norvegicus OX=10116 GN=CC |
| P05505\|COX3_RAT | **66** | 2866 | 49.12 | 3 | 3 | 151450 | 1 | 1 | 4 | N | 29871 | Cytochrome c oxidase subunit 3 OS=Rattus norvegicus OX=10116 GN=Mt |
| tr\|Q8SEZ2\|Q8SEZ2 | **66** | 2867 | 49.12 | 3 | 3 | 151450 | 1 | 1 | 4 | N | 29870 | Cytochrome c oxidase subunit 3 (Fragment) OS=Rattus norvegicus OX=1( |
| tr\|Q8M7G5\|Q8M7G | **49** | 13 | 47.43 | 8 | 8 | 976850 | 2 | 2 | 7 | Y | 24240 | ATP synthase subunit a (Fragment) OS=Rattus norvegicus OX=10116 PE= |
| P05504\|ATP6_RAT | **49** | 15 | 47.43 | 8 | 8 | 976850 | 2 | 2 | 7 | Y | 25076 | ATP synthase subunit a OS=Rattus norvegicus OX=10116 GN=Mt-atp6 PE |
| tr\|Q8HIC7\|Q8HIC7_ | **49** | 16 | 47.43 | 8 | 8 | 976850 | 2 | 2 | 7 | Y | 25076 | ATP synthase subunit a OS=Rattus norvegicus OX=10116 GN=Mt-atp6 PE |
| tr\|S5S1E9\|S5S1E9_ | **49** | 17 | 47.43 | 8 | 8 | 976850 | 2 | 2 | 7 | Y | 25049 | ATP synthase subunit a OS=Rattus norvegicus OX=10116 GN=ATP6 PE=4 |
| tr\|Q8SEZ3\|Q8SEZ3 | **49** | 23 | 47.43 | 8 | 8 | 976850 | 2 | 2 | 7 | Y | 25030 | ATP synthase subunit a OS=Rattus norvegicus OX=10116 GN=ATPase6 PE |
| tr\|Q549I0\|Q549I0_R | **49** | 14 | 47.43 | 8 | 8 | 976850 | 2 | 2 | 7 | Y | 25050 | ATP synthase subunit a OS=Rattus norvegicus OX=10116 GN=atp6 PE=4 |
| tr\|B2RYW3\|B2RYW | **56** | 5441 | 45.99 | 7 | 7 | 1041600 | 1 | 1 | 5 | Y | 21892 | NADH dehydrogenase (Ubiquinone) 1 beta subcomplex, 9 OS=Rattus norv |
| tr\|Q35733\|Q35733_ | **75** | 5460 | 45.67 | 13 | 13 | 277810 | 1 | 1 | 3 | N | 8389 | NADH-ubiquinone oxidoreductase chain 6 (Fragment) OS=Rattus norvegic |
| tr\|Q06QD9\|Q06QD | **75** | 5461 | 45.67 | 6 | 6 | 277810 | 1 | 1 | 3 | N | 18943 | NADH-ubiquinone oxidoreductase chain 6 OS=Rattus norvegicus OX=1011 |
| tr\|Q7HKW2\|Q7HKW | **75** | 5462 | 45.67 | 6 | 6 | 277810 | 1 | 1 | 3 | N | 18957 | NADH-ubiquinone oxidoreductase chain 6 OS=Rattus norvegicus OX=1011 |
| Q63704\|CPT1B_RA | **122** | 138 | 44.17 | 2 | 2 | 172350 | 1 | 1 | 1 | N | 88217 | Carnitine O-palmitoyltransferase 1, muscle isoform OS=Rattus norvegicus |
| P21571\|ATP5J_RA | **123** | 30 | 39.99 | 10 | 10 | 171430 | 1 | 1 | 1 | N | 12494 | ATP synthase-coupling factor 6, mitochondrial OS=Rattus norvegicus OX= |
| tr\|Q06QA9\|Q06QA9 | **87** | 2790 | 37.15 | 3 | 3 | 105120 | 2 | 2 | 2 | N | 56893 | Cytochrome c oxidase subunit 1 OS=Rattus norvegicus OX=10116 GN=CC |
| tr\|Q06QK0\|Q06QK0 | **87** | 2791 | 37.15 | 3 | 3 | 105120 | 2 | 2 | 2 | N | 56880 | Cytochrome c oxidase subunit 1 OS=Rattus norvegicus OX=10116 GN=CC |
| tr\|Q8SEZ6\|Q8SEZ6 | **87** | 2792 | 37.15 | 3 | 3 | 105120 | 2 | 2 | 2 | N | 56879 | Cytochrome c oxidase subunit 1 OS=Rattus norvegicus OX=10116 GN=CC |
| tr\|A0A0A1FZ34\|A0A | **87** | 2793 | 37.15 | 3 | 3 | 105120 | 2 | 2 | 2 | N | 56937 | Cytochrome c oxidase subunit 1 OS=Rattus norvegicus OX=10116 GN=CC |
| tr\|Q8HIC9\|Q8HIC9_ | **87** | 2794 | 37.15 | 3 | 3 | 105120 | 2 | 2 | 2 | N | 56845 | Cytochrome c oxidase subunit 1 OS=Rattus norvegicus OX=10116 GN=Mt |
| P05503\|COX1_RAT | **87** | 2795 | 37.15 | 3 | 3 | 105120 | 2 | 2 | 2 | N | 56845 | Cytochrome c oxidase subunit 1 OS=Rattus norvegicus OX=10116 GN=Mt |
| tr\|Q95938\|Q95938_ | **87** | 2796 | 37.15 | 3 | 3 | 105120 | 2 | 2 | 2 | N | 56977 | Cytochrome c oxidase subunit 1 OS=Rattus norvegicus OX=10116 GN=Co |
| tr\|B2RZ24\|B2RZ24_ | **124** | 2877 | 36.39 | 4 | 4 | 75060 | 1 | 1 | 1 | N | 47388 | Succinate-CoA ligase subunit beta (Fragment) OS=Rattus norvegicus OX= |
| tr\|F1LM47\|F1LM47_ | **124** | 2878 | 36.39 | 4 | 4 | 75060 | 1 | 1 | 1 | N | 50306 | Succinate--CoA ligase [ADP-forming] subunit beta, mitochondrial OS=Ratt |
| Q5M9I5\|QCR6_RAT | **73** | 2875 | 34.79 | 17 | 17 | 314440 | 3 | 3 | 3 | N | 10424 | Cytochrome b-c1 complex subunit 6, mitochondrial OS=Rattus norvegicus |
| Q9JJW3\|ATPMD_R | **125** | 89 | 33.84 | 21 | 21 | 548650 | 1 | 1 | 1 | N | 6408 | ATP synthase membrane subunit DAPIT, mitochondrial OS=Rattus norvegi |
| tr\|D4A4P3\|D4A4P3_ | **64** | 5451 | 30.36 | 15 | 15 | 726610 | 2 | 2 | 4 | N | 11267 | NADH:ubiquinone oxidoreductase subunit B3 OS=Rattus norvegicus OX=1 |
| Q9R063\|PRDX5_RA | **126** | 10115 | 28.58 | 6 | 6 | 91401 | 1 | 1 | 1 | N | 22179 | Peroxiredoxin-5, mitochondrial OS=Rattus norvegicus OX=10116 GN=Prd |
| tr\|A0A0G2JSS8\|A0A | **126** | 10116 | 28.58 | 6 | 6 | 91401 | 1 | 1 | 1 | N | 22207 | Peroxiredoxin OS=Rattus norvegicus OX=10116 GN=Prdx5 PE=1 SV=1 |
| tr\|B2RZD6\|B2RZD6 | **103** | 568 | 28.49 | 9 | 9 | 46814 | 1 | 1 | 1 | N | 9327 | NDUFA4, mitochondrial complex-associated OS=Rattus norvegicus OX=10 |
| P05508\|NU4M_RAT | **62** | 5425 | 26.36 | 2 | 2 | 362850 | 1 | 1 | 4 | N | 51783 | NADH-ubiquinone oxidoreductase chain 4 OS=Rattus norvegicus OX=1011 |
| tr\|D2E6K0\|D2E6K0_ | **62** | 5426 | 26.36 | 2 | 2 | 362850 | 1 | 1 | 4 | N | 51833 | NADH-ubiquinone oxidoreductase chain 4 OS=Rattus norvegicus OX=1011 |
| tr\|A7XYB9\|A7XYB9_ | **62** | 5427 | 26.36 | 2 | 2 | 362850 | 1 | 1 | 4 | N | 51773 | NADH-ubiquinone oxidoreductase chain 4 OS=Rattus norvegicus OX=1011 |
| tr\|Q06QE1\|Q06QE1 | **62** | 5428 | 26.36 | 2 | 2 | 362850 | 1 | 1 | 4 | N | 51851 | NADH-ubiquinone oxidoreductase chain 4 OS=Rattus norvegicus OX=1011 |
| tr\|Q06QA2\|Q06QA2 | **62** | 5429 | 26.36 | 2 | 2 | 362850 | 1 | 1 | 4 | N | 51787 | NADH-ubiquinone oxidoreductase chain 4 OS=Rattus norvegicus OX=1011 |
| tr\|Q7HKW3\|Q7HKW | **62** | 5430 | 26.36 | 2 | 2 | 362850 | 1 | 1 | 4 | N | 51801 | NADH-ubiquinone oxidoreductase chain 4 (Fragment) OS=Rattus norvegic |
| tr\|Q06QG7\|Q06QG | **62** | 5431 | 26.36 | 2 | 2 | 362850 | 1 | 1 | 4 | N | 51791 | NADH-ubiquinone oxidoreductase chain 4 OS=Rattus norvegicus OX=1011 |
| tr\|Q06Q89\|Q06Q89_ | **62** | 5432 | 26.36 | 2 | 2 | 362850 | 1 | 1 | 4 | N | 51819 | NADH-ubiquinone oxidoreductase chain 4 OS=Rattus norvegicus OX=1011 |
| tr\|Q35737\|Q35737_ | **62** | 5433 | 26.36 | 2 | 2 | 362850 | 1 | 1 | 4 | N | 51765 | NADH-ubiquinone oxidoreductase chain 4 OS=Rattus norvegicus OX=1011 |
| tr\|Q8HIC6\|Q8HIC6_ | **62** | 5434 | 26.36 | 2 | 2 | 362850 | 1 | 1 | 4 | N | 51783 | NADH-ubiquinone oxidoreductase chain 4 OS=Rattus norvegicus OX=1011 |
| Q5M934\|TRUB1_RA | **36** | 284 | 25.68 | 2 | 2 | 2626900 | 1 | 1 | 12 | N | 36404 | Probable tRNA pseudouridine synthase 1 OS=Rattus norvegicus OX=1011 |
| D4A2Z8\|DHX36_RA | **129** | 7802 | 25.53 | 1 | 1 | 1590500 | 1 | 1 | 1 | N | 113843 | ATP-dependent DNA/RNA helicase DHX36 OS=Rattus norvegicus OX=101 |
| tr\|B2RYU0\|B2RYU0 | **108** | 5486 | 24.58 | 9 | 9 | 365080 | 1 | 1 | 1 | N | 11842 | NADH dehydrogenase (Ubiquinone) 1 beta subcomplex, 2 (Predicted), isof |
| P80431\|COX7B_RA | **130** | 2798 | 23.83 | 14 | 14 | 92435 | 1 | 1 | 1 | N | 8995 | Cytochrome c oxidase subunit 7B, mitochondrial OS=Rattus norvegicus O |
| tr\|A0A0G2K6P0\|A0A | **130** | 2860 | 23.83 | 14 | 14 | 92435 | 1 | 1 | 1 | N | 9070 | Cytochrome c oxidase subunit 7B2 OS=Rattus norvegicus OX=10116 GN= |
| P13086\|SUCA_RAT | **109** | 2822 | 44400 | 3 | 3 | 0 | 1 | 1 | 1 | Y | 36148 | Succinate--CoA ligase [ADP/GDP-forming] subunit alpha, mitochondrial OS |
| tr\|A0A0H2UHE1\|A0A | **109** | 2823 | 44400 | 3 | 3 | 0 | 1 | 1 | 1 | Y | 37560 | Succinate--CoA ligase [ADP/GDP-forming] subunit alpha, mitochondrial OS |
| Q8VIJ5\|OGA_RAT | **82** | 207 | 44250 | 1 | 1 | 242720 | 1 | 1 | 2 | N | 102918 | Protein O-GlcNAcase OS=Rattus norvegicus OX=10116 GN=Oga PE=1 SV |
| P52631\|STAT3_RA | **81** | 5489 | 21.55 | 1 | 1 | 126710 | 1 | 1 | 2 | N | 88040 | Signal transducer and activator of transcription 3 OS=Rattus norvegicus O |
| P35171\|CX7A2_RA | **132** | 1062 | 21.25 | 12 | 12 | 44810 | 1 | 1 | 1 | N | 9353 | Cytochrome c oxidase subunit 7A2, mitochondrial OS=Rattus norvegicus C |
| tr\|B2RYS0\|B2RYS0 | **132** | 1063 | 21.25 | 12 | 12 | 44810 | 1 | 1 | 1 | N | 9353 | Cox7a2 protein OS=Rattus norvegicus OX=10116 GN=Cox7a2 PE=2 SV= |
| tr\|A0A0G2JYU2\|A0A | **84** | 2830 | 20.89 | 4 | 4 | 268290 | 1 | 1 | 2 | N | 20750 | Mitochondrial ribosomal protein L11 OS=Rattus norvegicus OX=10116 GN |
| Q5XIE3\|RM11_RAT | **84** | 2831 | 20.89 | 4 | 4 | 268290 | 1 | 1 | 2 | N | 22420 | 39S ribosomal protein L11, mitochondrial OS=Rattus norvegicus OX=1011 |
| tr\|Q6P9Y4\|Q6P9Y4_ | **95** | 1009 | 20.32 | 3 | 3 | 127850 | 1 | 1 | 1 | N | 32904 | ADP/ATP translocase 1 OS=Rattus norvegicus OX=10116 GN=Slc25a4 PE |
| Q05962\|ADT1_RAT | **95** | 1010 | 20.32 | 3 | 3 | 127850 | 1 | 1 | 1 | N | 32989 | ADP/ATP translocase 1 OS=Rattus norvegicus OX=10116 GN=Slc25a4 PE |
| Q09073\|ADT2_RAT | **95** | 1011 | 20.32 | 3 | 3 | 127850 | 1 | 1 | 1 | N | 32901 | ADP/ATP translocase 2 OS=Rattus norvegicus OX=10116 GN=Slc25a5 PE |

| | | | | | | | | | | | | |
|---|---|---|---|---|---|---|---|---|---|---|---|---|
| tr\|D3ZB81\|D3ZB81_ | **95** | 1012 | 20.32 | 2 | 2 | 127850 | 1 | 1 | 1 | N | 35227 | Solute carrier family 25 member 31 OS=Rattus norvegicus OX=10116 GN |
| tr\|A0A1W2Q605\|A0 | **95** | 1008 | 20.32 | 7 | 7 | 127850 | 1 | 1 | 1 | N | 13534 | Solute carrier family 25 member 31 (Fragment) OS=Rattus norvegicus OX |
