## Supplementary material for "Permeability transition pore-related changes in the proteome and channel activity of ATP synthase dimers and monomers": RHM Control Monomer of V compl.

| Accession | Protein Group | Protein ID | -10lgP | Coverage (%) | Coverage (%) | Area Sample 8 | #Peptides | #Unique | #Spec Sample | PTM | Avg. Mass | Description |
|---|---|---|---|---|---|---|---|---|---|---|---|---|
| P10719\|ATPB_RAT | 2 | 1 | 387.46 | 60 | 60 | 1157500000 | 170 | 170 | 874 | Y | 56354 | ATP synthase subunit beta, mitochondrial OS=Rattus norvegicus OX=10116 GN=Atp5f1b PE=1 SV=2 |
| tr\|G3V6D3\|G3V6D3_RAT | 2 | 2 | 387.46 | 60 | 60 | 1157500000 | 170 | 170 | 874 | Y | 56345 | ATP synthase subunit beta OS=Rattus norvegicus OX=10116 GN=Atp5f1b PE=1 SV=1 |
| P15999\|ATPA_RAT | 1 | 3 | 338.57 | 58 | 58 | 716370000 | 168 | 166 | 907 | Y | 59754 | ATP synthase subunit alpha, mitochondrial OS=Rattus norvegicus OX=10116 GN=Atp5f1a PE=1 SV=2 |
| tr\|F1LP05\|F1LP05_RAT | 1 | 4 | 338.57 | 58 | 58 | 716370000 | 168 | 166 | 907 | Y | 59813 | ATP synthase subunit alpha OS=Rattus norvegicus OX=10116 GN=Atp5f1a PE=1 SV=1 |
| P35435\|ATPG_RAT | 3 | 5 | 280.45 | 43 | 43 | 102670000 | 46 | 46 | 221 | Y | 30191 | ATP synthase subunit gamma, mitochondrial OS=Rattus norvegicus OX=10116 GN=Atp5f1c PE=1 SV=2 |
| tr\|Q6QI09\|Q6QI09_RAT | 3 | 6 | 280.45 | 19 | 19 | 102670000 | 46 | 46 | 221 | Y | 67721 | ATP synthase subunit gamma, mitochondrial OS=Rattus norvegicus OX=10116 GN=Taf3 PE=1 SV=1 |
| P31399\|ATP5H_RAT | 4 | 7 | 278.21 | 65 | 65 | 85852000 | 41 | 40 | 154 | Y | 18763 | ATP synthase subunit d, mitochondrial OS=Rattus norvegicus OX=10116 GN=Atp5pd PE=1 SV=3 |
| Q6PDU7\|ATP5L_RAT | 6 | 11 | 171.31 | 45 | 45 | 22118000 | 19 | 19 | 67 | Y | 11433 | ATP synthase subunit g, mitochondrial OS=Rattus norvegicus OX=10116 GN=Atp5mg PE=1 SV=2 |
| P19511\|AT5F1_RAT | 7 | 10 | 167.95 | 23 | 23 | 85675000 | 21 | 21 | 64 | Y | 28869 | ATP synthase F(0) complex subunit B1, mitochondrial OS=Rattus norvegicus OX=10116 GN=Atp5pb PE=1 SV=1 |
| tr\|G3V7Y3\|G3V7Y3_RAT | 8 | 12 | 162.91 | 33 | 33 | 18902000 | 13 | 13 | 47 | Y | 17563 | ATP synthase subunit delta, mitochondrial OS=Rattus norvegicus OX=10116 GN=Atp5f1d PE=1 SV=1 |
| Q06647\|ATPO_RAT | 5 | 8 | 159.85 | 38 | 38 | 30283000 | 21 | 21 | 71 | Y | 23398 | ATP synthase subunit O, mitochondrial OS=Rattus norvegicus OX=10116 GN=Atp5po PE=1 SV=1 |
| P21571\|ATP5J_RAT | 10 | 30 | 159.19 | 39 | 39 | 4939200 | 10 | 10 | 25 | N | 12494 | ATP synthase-coupling factor 6, mitochondrial OS=Rattus norvegicus OX=10116 GN=Atp5pf PE=1 SV=1 |
| tr\|Q5UAJ5\|Q5UAJ5_RAT | 9 | 32 | 134.38 | 48 | 48 | 45606000 | 8 | 8 | 32 | Y | 7642 | ATP synthase protein 8 OS=Rattus norvegicus OX=10116 GN=ATP8 PE=3 SV=1 |
| tr\|Q8SEZ4\|Q8SEZ4_RAT | 9 | 33 | 134.38 | 48 | 48 | 45606000 | 8 | 8 | 32 | Y | 7632 | ATP synthase protein 8 OS=Rattus norvegicus OX=10116 GN=ATPase8 PE=3 SV=1 |
| Q64428\|ECHA_RAT | 12 | 9 | 127.39 | 8 | 8 | 3637100 | 11 | 11 | 21 | N | 82665 | Trifunctional enzyme subunit alpha, mitochondrial OS=Rattus norvegicus OX=10116 GN=Hadha PE=1 SV=2 |
| P00507\|AATM_RAT | 15 | 19 | 126.2 | 12 | 12 | 2748600 | 7 | 7 | 16 | Y | 47314 | Aspartate aminotransferase, mitochondrial OS=Rattus norvegicus OX=10116 GN=Got2 PE=1 SV=2 |
| Q60587\|ECHB_RAT | 13 | 20 | 115.62 | 15 | 15 | 2834100 | 11 | 11 | 19 | N | 51414 | Trifunctional enzyme subunit beta, mitochondrial OS=Rattus norvegicus OX=10116 GN=Hadhb PE=1 SV=1 |
| tr\|A0A0G2K330\|A0A0G2K33 | 13 | 21 | 115.62 | 15 | 15 | 2834100 | 11 | 11 | 19 | N | 52568 | Trifunctional enzyme subunit beta, mitochondrial OS=Rattus norvegicus OX=10116 GN=Hadhb PE=1 SV=1 |
| tr\|D3ZFQ8\|D3ZFQ8_RAT | 21 | 133 | 93.16 | 5 | 5 | 1386800 | 2 | 2 | 6 | N | 35435 | Cytochrome c-1 OS=Rattus norvegicus OX=10116 GN=Cyc1 PE=1 SV=3 |
| P29419\|ATP5I_RAT | 14 | 31 | 91.6 | 49 | 49 | 2316400 | 6 | 6 | 17 | N | 8255 | ATP synthase subunit e, mitochondrial OS=Rattus norvegicus OX=10116 GN=Atp5me PE=1 SV=3 |
| tr\|Q8M7G5\|Q8M7G5_RAT | 11 | 13 | 90.42 | 27 | 27 | 3492600 | 8 | 8 | 22 | Y | 24240 | ATP synthase subunit a (Fragment) OS=Rattus norvegicus OX=10116 PE=2 SV=1 |
| tr\|Q549I0\|Q549I0_RAT | 11 | 14 | 90.42 | 27 | 27 | 3492600 | 8 | 8 | 22 | Y | 25050 | ATP synthase subunit a OS=Rattus norvegicus OX=10116 GN=atp6 PE=4 SV=1 |
| P05504\|ATP6_RAT | 11 | 15 | 90.42 | 27 | 27 | 3492600 | 8 | 8 | 22 | Y | 25076 | ATP synthase subunit a OS=Rattus norvegicus OX=10116 GN=Mt-atp6 PE=1 SV=3 |
| tr\|Q8HIC7\|Q8HIC7_RAT | 11 | 16 | 90.42 | 27 | 27 | 3492600 | 8 | 8 | 22 | Y | 25076 | ATP synthase subunit a OS=Rattus norvegicus OX=10116 GN=Mt-atp6 PE=4 SV=1 |
| tr\|S5S1E9\|S5S1E9_RAT | 11 | 17 | 90.42 | 27 | 27 | 3492600 | 8 | 8 | 22 | Y | 25049 | ATP synthase subunit a OS=Rattus norvegicus OX=10116 GN=ATP6 PE=4 SV=1 |
| P11240\|COX5A_RAT | 19 | 86 | 82.98 | 16 | 16 | 571600 | 3 | 3 | 6 | N | 16130 | Cytochrome c oxidase subunit 5A, mitochondrial OS=Rattus norvegicus OX=10116 GN=Cox5a PE=1 SV=1 |
| P20788\|UCRI_RAT | 75 | 132 | 74.5 | 5 | 5 | 276840 | 1 | 1 | 1 | N | 29446 | Cytochrome b-c1 complex subunit Rieske, mitochondrial OS=Rattus norvegicus OX=10116 GN=Uqcrfs1 PE=1 SV=2 |
| tr\|Q5UAJ6\|Q5UAJ6_RAT | 16 | 24 | 69.65 | 17 | 17 | 843160 | 5 | 5 | 6 | Y | 25942 | Cytochrome c oxidase subunit 2 OS=Rattus norvegicus OX=10116 GN=COX2 PE=3 SV=1 |
| tr\|S5RZM8\|S5RZM8_RAT | 16 | 26 | 69.65 | 17 | 17 | 843160 | 5 | 5 | 6 | Y | 25958 | Cytochrome c oxidase subunit 2 OS=Rattus norvegicus OX=10116 GN=COX2 PE=3 SV=1 |
| P00406\|COX2_RAT | 16 | 27 | 69.65 | 17 | 17 | 843160 | 5 | 5 | 6 | Y | 25928 | Cytochrome c oxidase subunit 2 OS=Rattus norvegicus OX=10116 GN=Mtco2 PE=1 SV=3 |
| tr\|Q8SEZ5\|Q8SEZ5_RAT | 16 | 28 | 69.65 | 17 | 17 | 843160 | 5 | 5 | 6 | Y | 25928 | Cytochrome c oxidase subunit 2 OS=Rattus norvegicus OX=10116 GN=Mt-co2 PE=1 SV=1 |
| tr\|A0A097PE04\|A0A097PE0 | 16 | 29 | 69.65 | 17 | 17 | 843160 | 5 | 5 | 6 | Y | 25894 | Cytochrome c oxidase subunit 2 OS=Rattus norvegicus OX=10116 GN=COX2 PE=3 SV=1 |
| P09605\|KCRS_RAT | 34 | 84 | 63.37 | 6 | 6 | 412600 | 2 | 2 | 2 | Y | 47385 | Creatine kinase S-type, mitochondrial OS=Rattus norvegicus OX=10116 GN=Ckmt2 PE=1 SV=2 |
| P32551\|QCR2_RAT | 17 | 34 | 63.27 | 8 | 8 | 1024500 | 4 | 4 | 6 | N | 48396 | Cytochrome b-c1 complex subunit 2, mitochondrial OS=Rattus norvegicus OX=10116 GN=Uqcrc2 PE=1 SV=2 |

| | | | | | | | | | | | |
|---|---|---|---|---|---|---|---|---|---|---|---|
| Q9ER34\|ACON_RAT | **25** | 88 | 60.11 | 3 | 3 | 552340 | 2 | 2 | 3 | N | 85433 Aconitate hydratase, mitochondrial OS=Rattus norvegicus OX=10116 GN=Aco2 PE=1 SV=2 |
| P80432\|COX7C_RAT | **76** | 134 | 57.04 | 21 | 21 | 152490 | 1 | 1 | 1 | N | 7375 Cytochrome c oxidase subunit 7C, mitochondrial OS=Rattus norvegicus OX=10116 GN=Cox7c PE=1 SV=2 |
| tr\|A0A3Q8AC68\|A0A3Q8AC | **18** | 38 | 51.39 | 8 | 8 | 1491900 | 3 | 3 | 6 | N | 42622 Cytochrome b (Fragment) OS=Rattus norvegicus OX=10116 GN=Cytb PE=3 SV=1 |
| tr\|A0A411P2I9\|A0A411P2I9_ | **18** | 39 | 51.39 | 8 | 8 | 1491900 | 3 | 3 | 6 | N | 40144 Cytochrome b (Fragment) OS=Rattus norvegicus OX=10116 GN=Cytb PE=3 SV=1 |
| tr\|A0A0U2IDR8\|A0A0U2IDR | **18** | 40 | 51.39 | 8 | 8 | 1491900 | 3 | 3 | 6 | N | 40981 Cytochrome b (Fragment) OS=Rattus norvegicus OX=10116 PE=3 SV=1 |
| tr\|A0A0U2M179\|A0A0U2M17 | **18** | 41 | 51.39 | 8 | 8 | 1491900 | 3 | 3 | 6 | N | 41037 Cytochrome b (Fragment) OS=Rattus norvegicus OX=10116 PE=3 SV=1 |
| tr\|A0A411P2L0\|A0A411P2L | **18** | 42 | 51.39 | 8 | 8 | 1491900 | 3 | 3 | 6 | N | 41017 Cytochrome b (Fragment) OS=Rattus norvegicus OX=10116 GN=Cytb PE=3 SV=1 |
| tr\|F8QU31\|F8QU31_RAT | **18** | 43 | 51.39 | 8 | 8 | 1491900 | 3 | 3 | 6 | N | 42181 Cytochrome b (Fragment) OS=Rattus norvegicus OX=10116 GN=cytb PE=3 SV=1 |
| tr\|A0A411P2K0\|A0A411P2K | **18** | 44 | 51.39 | 8 | 8 | 1491900 | 3 | 3 | 6 | N | 42238 Cytochrome b (Fragment) OS=Rattus norvegicus OX=10116 GN=Cytb PE=3 SV=1 |
| tr\|L0L4L8\|L0L4L8_RAT | **18** | 45 | 51.39 | 8 | 8 | 1491900 | 3 | 3 | 6 | N | 42470 Cytochrome b (Fragment) OS=Rattus norvegicus OX=10116 GN=cytb PE=3 SV=1 |
| tr\|A0A140GE10\|A0A140GE1 | **18** | 46 | 51.39 | 8 | 8 | 1491900 | 3 | 3 | 6 | N | 42574 Cytochrome b (Fragment) OS=Rattus norvegicus OX=10116 PE=3 SV=1 |
| tr\|A0A140GE11\|A0A140GE1 | **18** | 47 | 51.39 | 8 | 8 | 1491900 | 3 | 3 | 6 | N | 42588 Cytochrome b (Fragment) OS=Rattus norvegicus OX=10116 PE=3 SV=1 |
| tr\|A0A140GE08\|A0A140GE0 | **18** | 48 | 51.39 | 8 | 8 | 1491900 | 3 | 3 | 6 | N | 42527 Cytochrome b (Fragment) OS=Rattus norvegicus OX=10116 PE=3 SV=1 |
| tr\|A0A385HCC9\|A0A385HC | **18** | 49 | 51.39 | 8 | 8 | 1491900 | 3 | 3 | 6 | N | 42766 Cytochrome b (Fragment) OS=Rattus norvegicus OX=10116 PE=3 SV=1 |
| tr\|A0A3Q8AGE8\|A0A3Q8AG | **18** | 50 | 51.39 | 8 | 8 | 1491900 | 3 | 3 | 6 | N | 42712 Cytochrome b (Fragment) OS=Rattus norvegicus OX=10116 GN=Cytb PE=3 SV=1 |
| tr\|A0A3S6FK13\|A0A3S6FK1 | **18** | 51 | 51.39 | 8 | 8 | 1491900 | 3 | 3 | 6 | N | 42702 Cytochrome b (Fragment) OS=Rattus norvegicus OX=10116 GN=Cytb PE=3 SV=1 |
| tr\|A0A3S6FM15\|A0A3S6FM | **18** | 52 | 51.39 | 8 | 8 | 1491900 | 3 | 3 | 6 | N | 42698 Cytochrome b (Fragment) OS=Rattus norvegicus OX=10116 GN=Cytb PE=3 SV=1 |
| tr\|A0A097PE40\|A0A097PE4 | **18** | 53 | 51.39 | 8 | 8 | 1491900 | 3 | 3 | 6 | N | 42865 Cytochrome b OS=Rattus norvegicus OX=10116 GN=CYTB PE=3 SV=1 |
| tr\|A0A220D9Z6\|A0A220D9Z | **18** | 54 | 51.39 | 8 | 8 | 1491900 | 3 | 3 | 6 | N | 43018 Cytochrome b OS=Rattus norvegicus OX=10116 PE=3 SV=1 |
| tr\|D6NSP2\|D6NSP2_RAT | **18** | 55 | 51.39 | 8 | 8 | 1491900 | 3 | 3 | 6 | N | 42945 Cytochrome b (Fragment) OS=Rattus norvegicus OX=10116 GN=cytb PE=3 SV=1 |
| tr\|D6NSS6\|D6NSS6_RAT | **18** | 56 | 51.39 | 8 | 8 | 1491900 | 3 | 3 | 6 | N | 42993 Cytochrome b (Fragment) OS=Rattus norvegicus OX=10116 GN=cytb PE=3 SV=1 |
| tr\|D6NSQ3\|D6NSQ3_RAT | **18** | 57 | 51.39 | 8 | 8 | 1491900 | 3 | 3 | 6 | N | 43012 Cytochrome b (Fragment) OS=Rattus norvegicus OX=10116 GN=cytb PE=3 SV=1 |
| tr\|D6NSR3\|D6NSR3_RAT | **18** | 58 | 51.39 | 8 | 8 | 1491900 | 3 | 3 | 6 | N | 42986 Cytochrome b (Fragment) OS=Rattus norvegicus OX=10116 GN=cytb PE=3 SV=1 |
| tr\|D6NSP4\|D6NSP4_RAT | **18** | 59 | 51.39 | 8 | 8 | 1491900 | 3 | 3 | 6 | N | 43016 Cytochrome b (Fragment) OS=Rattus norvegicus OX=10116 GN=cytb PE=3 SV=1 |
| tr\|D6NSQ8\|D6NSQ8_RAT | **18** | 60 | 51.39 | 8 | 8 | 1491900 | 3 | 3 | 6 | N | 43016 Cytochrome b (Fragment) OS=Rattus norvegicus OX=10116 GN=cytb PE=3 SV=1 |
| tr\|D6NSR8\|D6NSR8_RAT | **18** | 61 | 51.39 | 8 | 8 | 1491900 | 3 | 3 | 6 | N | 42998 Cytochrome b (Fragment) OS=Rattus norvegicus OX=10116 GN=cytb PE=3 SV=1 |
| tr\|F8QU37\|F8QU37_RAT | **18** | 62 | 51.39 | 8 | 8 | 1491900 | 3 | 3 | 6 | N | 42992 Cytochrome b (Fragment) OS=Rattus norvegicus OX=10116 GN=cytb PE=3 SV=1 |
| tr\|F2Q6S6\|F2Q6S6_RAT | **18** | 63 | 51.39 | 8 | 8 | 1491900 | 3 | 3 | 6 | N | 42938 Cytochrome b (Fragment) OS=Rattus norvegicus OX=10116 GN=cytb PE=3 SV=1 |
| tr\|A0A0A1FZ42\|A0A0A1FZ4 | **18** | 64 | 51.39 | 8 | 8 | 1491900 | 3 | 3 | 6 | N | 42989 Cytochrome b OS=Rattus norvegicus OX=10116 GN=CYTB PE=3 SV=1 |
| tr\|D6NSS7\|D6NSS7_RAT | **18** | 65 | 51.39 | 8 | 8 | 1491900 | 3 | 3 | 6 | N | 42948 Cytochrome b (Fragment) OS=Rattus norvegicus OX=10116 GN=cytb PE=3 SV=1 |
| tr\|Q8SEY9\|Q8SEY9_RAT | **18** | 66 | 51.39 | 8 | 8 | 1491900 | 3 | 3 | 6 | N | 43015 Cytochrome b OS=Rattus norvegicus OX=10116 GN=cytb PE=3 SV=1 |
| tr\|D6NSR7\|D6NSR7_RAT | **18** | 67 | 51.39 | 8 | 8 | 1491900 | 3 | 3 | 6 | N | 42982 Cytochrome b (Fragment) OS=Rattus norvegicus OX=10116 GN=cytb PE=3 SV=1 |
| tr\|A0A220DA44\|A0A220DA4 | **18** | 68 | 51.39 | 8 | 8 | 1491900 | 3 | 3 | 6 | N | 42968 Cytochrome b OS=Rattus norvegicus OX=10116 PE=3 SV=1 |
| tr\|F2Q6S5\|F2Q6S5_RAT | **18** | 69 | 51.39 | 8 | 8 | 1491900 | 3 | 3 | 6 | N | 42952 Cytochrome b (Fragment) OS=Rattus norvegicus OX=10116 GN=cytb PE=3 SV=1 |
| tr\|D6NSQ6\|D6NSQ6_RAT | **18** | 70 | 51.39 | 8 | 8 | 1491900 | 3 | 3 | 6 | N | 42952 Cytochrome b (Fragment) OS=Rattus norvegicus OX=10116 GN=cytb PE=3 SV=1 |
| tr\|D6NSQ0\|D6NSQ0_RAT | **18** | 71 | 51.39 | 8 | 8 | 1491900 | 3 | 3 | 6 | N | 42968 Cytochrome b (Fragment) OS=Rattus norvegicus OX=10116 GN=cytb PE=3 SV=1 |
| tr\|Q5UAI7\|Q5UAI7_RAT | **18** | 72 | 51.39 | 8 | 8 | 1491900 | 3 | 3 | 6 | N | 42998 Cytochrome b OS=Rattus norvegicus OX=10116 GN=CYTB PE=3 SV=1 |
| tr\|D6NSR2\|D6NSR2_RAT | **18** | 73 | 51.39 | 8 | 8 | 1491900 | 3 | 3 | 6 | N | 43073 Cytochrome b (Fragment) OS=Rattus norvegicus OX=10116 GN=cytb PE=3 SV=1 |
| tr\|A0A0S1Z1V9\|A0A0S1Z1V | **18** | 74 | 51.39 | 8 | 8 | 1491900 | 3 | 3 | 6 | N | 43002 Cytochrome b OS=Rattus norvegicus OX=10116 GN=CYTB PE=3 SV=1 |
| tr\|D6NSR1\|D6NSR1_RAT | **18** | 75 | 51.39 | 8 | 8 | 1491900 | 3 | 3 | 6 | N | 43002 Cytochrome b (Fragment) OS=Rattus norvegicus OX=10116 GN=cytb PE=3 SV=1 |
| tr\|Q8HIC4\|Q8HIC4_RAT | **18** | 76 | 51.39 | 8 | 8 | 1491900 | 3 | 3 | 6 | N | 43012 Cytochrome b OS=Rattus norvegicus OX=10116 GN=Mt-cyb PE=3 SV=1 |
| tr\|A0A220DA02\|A0A220DA0 | **18** | 77 | 51.39 | 8 | 8 | 1491900 | 3 | 3 | 6 | N | 42968 Cytochrome b OS=Rattus norvegicus OX=10116 PE=3 SV=1 |

| | | | | | | | | | | | | |
|---|---|---|---|---|---|---|---|---|---|---|---|---|
| tr\|A0A0S1Z1V4\|A0A0S1Z1V4 | **18** | 78 | 51.39 | 8 | 8 | 1491900 | 3 | 3 | 6 | N | 43016 | Cytochrome b OS=Rattus norvegicus OX=10116 GN=CYTB PE=3 SV=1 |
| P00159\|CYB_RAT | **18** | 79 | 51.39 | 8 | 8 | 1491900 | 3 | 3 | 6 | N | 43012 | Cytochrome b OS=Rattus norvegicus OX=10116 GN=Mt-Cyb PE=3 SV=3 |
| tr\|L0N311\|L0N311_RAT | **18** | 80 | 51.39 | 8 | 8 | 1491900 | 3 | 3 | 6 | N | 43045 | Cytochrome b (Fragment) OS=Rattus norvegicus OX=10116 GN=cytb PE=3 SV=1 |
| tr\|A0A220DA28\|A0A220DA2 | **18** | 81 | 51.39 | 8 | 8 | 1491900 | 3 | 3 | 6 | N | 42970 | Cytochrome b OS=Rattus norvegicus OX=10116 PE=3 SV=1 |
| tr\|H2KXA0\|H2KXA0_RAT | **18** | 82 | 51.39 | 8 | 8 | 1491900 | 3 | 3 | 6 | N | 42978 | Cytochrome b (Fragment) OS=Rattus norvegicus OX=10116 GN=cytb PE=3 SV=1 |
| tr\|A0A096XNM4\|A0A096XNM | **18** | 83 | 51.39 | 8 | 8 | 1491900 | 3 | 3 | 6 | N | 43042 | Cytochrome b (Fragment) OS=Rattus norvegicus OX=10116 PE=3 SV=1 |
| Q9JJW3\|ATPMD_RAT | **28** | 89 | 45.84 | 33 | 33 | 927040 | 2 | 2 | 3 | N | 6408 | ATP synthase membrane subunit DAPIT, mitochondrial OS=Rattus norvegicus OX=10116 GN=Atp5md PE=1 SV=1 |
| tr\|Q6QI69\|Q6QI69_RAT | **23** | 87 | 38.2 | 4 | 4 | 424920 | 2 | 1 | 4 | Y | 46014 | LRRGT00139 OS=Rattus norvegicus OX=10116 GN=LOC500350 PE=2 SV=1 |
| Q68FY0\|QCR1_RAT | **35** | 118 | 37.65 | 4 | 4 | 60256 | 2 | 2 | 2 | N | 52849 | Cytochrome b-c1 complex subunit 1, mitochondrial OS=Rattus norvegicus OX=10116 GN=Uqcrc1 PE=1 SV=1 |
| D3ZAF6\|ATPK_RAT | **24** | 85 | 31.38 | 24 | 24 | 380940 | 3 | 3 | 3 | Y | 10452 | ATP synthase subunit f, mitochondrial OS=Rattus norvegicus OX=10116 GN=Atp5mf PE=1 SV=1 |
| Q63704\|CPT1B_RAT | **77** | 138 | 30.76 | 2 | 2 | 143510 | 1 | 1 | 1 | N | 88217 | Carnitine O-palmitoyltransferase 1, muscle isoform OS=Rattus norvegicus OX=10116 GN=Cpt1b PE=1 SV=1 |
| P12075\|COX5B_RAT | **38** | 139 | 28.66 | 9 | 9 | 516610 | 1 | 1 | 2 | N | 13915 | Cytochrome c oxidase subunit 5B, mitochondrial OS=Rattus norvegicus OX=10116 GN=Cox5b PE=1 SV=2 |
| P63039\|CH60_RAT | **78** | 142 | 44558 | 2 | 2 | 219720 | 1 | 1 | 1 | N | 60956 | 60 kDa heat shock protein, mitochondrial OS=Rattus norvegicus OX=10116 GN=Hspd1 PE=1 SV=1 |
| tr\|A0A482IDN3\|A0A482IDN3 | **78** | 143 | 44558 | 2 | 2 | 219720 | 1 | 1 | 1 | N | 60956 | Hsp60 OS=Rattus norvegicus OX=10116 GN=Hspd1 PE=2 SV=1 |
| tr\|A0A0G2K5L6\|A0A0G2K5L | **40** | 127 | 23.19 | 0 | 0 | 0 | 1 | 1 | 1 | N | 275755 | Acetyl-CoA carboxylase beta OS=Rattus norvegicus OX=10116 GN=Acacb PE=1 SV=1 |
| tr\|A0A0G2K1F2\|A0A0G2K1F | **40** | 128 | 23.19 | 0 | 0 | 0 | 1 | 1 | 1 | N | 276393 | Acetyl-CoA carboxylase beta OS=Rattus norvegicus OX=10116 GN=Acacb PE=1 SV=1 |
| tr\|D3ZBE2\|D3ZBE2_RAT | **40** | 129 | 23.19 | 0 | 0 | 0 | 1 | 1 | 1 | N | 275967 | Acetyl-CoA carboxylase beta OS=Rattus norvegicus OX=10116 GN=Acacb PE=1 SV=3 |
| tr\|E9PSQ0\|E9PSQ0_RAT | **40** | 130 | 23.19 | 0 | 0 | 0 | 1 | 1 | 1 | N | 276255 | Acetyl-CoA carboxylase beta OS=Rattus norvegicus OX=10116 GN=Acacb PE=1 SV=2 |
| tr\|O70151\|O70151_RAT | **40** | 140 | 23.19 | 0 | 0 | 0 | 1 | 1 | 1 | N | 276097 | Acetyl-CoA carboxylase OS=Rattus norvegicus OX=10116 GN=Acacb PE=2 SV=1 |
| P11497\|ACACA_RAT | **40** | 165 | 23.19 | 0 | 0 | 0 | 1 | 1 | 1 | N | 265191 | Acetyl-CoA carboxylase 1 OS=Rattus norvegicus OX=10116 GN=Acaca PE=1 SV=1 |
| P10817\|CX6A2_RAT | **66** | 151 | 44370 | 12 | 12 | 214450 | 1 | 1 | 1 | N | 10487 | Cytochrome c oxidase subunit 6A2, mitochondrial (Fragment) OS=Rattus norvegicus OX=10116 GN=Cox6a2 PE=1 SV=3 |
| tr\|G3V8M4\|G3V8M4_RAT | **66** | 152 | 44370 | 11 | 11 | 214450 | 1 | 1 | 1 | N | 10802 | Cytochrome c oxidase subunit 6A, mitochondrial OS=Rattus norvegicus OX=10116 GN=Cox6a2 PE=3 SV=1 |
| tr\|A0A0G2K4C6\|A0A0G2K4C | **67** | 159 | 22.21 | 1 | 1 | 425080 | 1 | 1 | 1 | N | 63436 | Malic enzyme OS=Rattus norvegicus OX=10116 GN=Me3 PE=3 SV=1 |
| P13697\|MAOX_RAT | **67** | 160 | 22.21 | 1 | 1 | 425080 | 1 | 1 | 1 | N | 64003 | NADP-dependent malic enzyme OS=Rattus norvegicus OX=10116 GN=Me1 PE=1 SV=2 |
| tr\|D3ZJH9\|D3ZJH9_RAT | **67** | 161 | 22.21 | 1 | 1 | 425080 | 1 | 1 | 1 | N | 65352 | Malic enzyme OS=Rattus norvegicus OX=10116 GN=Me2 PE=1 SV=1 |
| tr\|A0A0G2K502\|A0A0G2K50 | **67** | 162 | 22.21 | 1 | 1 | 425080 | 1 | 1 | 1 | N | 66310 | Malic enzyme OS=Rattus norvegicus OX=10116 GN=Me2 PE=1 SV=1 |
| tr\|F1M5N4\|F1M5N4_RAT | **67** | 163 | 22.21 | 1 | 1 | 425080 | 1 | 1 | 1 | N | 67209 | Malic enzyme OS=Rattus norvegicus OX=10116 GN=Me3 PE=3 SV=2 |
| Q4V8F9\|HSDL2_RAT | **48** | 121 | 21.81 | 2 | 2 | 179650 | 1 | 1 | 1 | N | 58344 | Hydroxysteroid dehydrogenase-like protein 2 OS=Rattus norvegicus OX=10116 GN=Hsdl2 PE=2 SV=1 |
| tr\|A0A140TAG5\|A0A140TAG | **65** | 145 | 20.88 | 2 | 2 | 45168 | 1 | 1 | 1 | N | 67049 | MICOS complex subunit MIC60 OS=Rattus norvegicus OX=10116 GN=Immt PE=1 SV=1 |
| Q3KR86\|MIC60_RAT | **65** | 146 | 20.88 | 2 | 2 | 45168 | 1 | 1 | 1 | N | 67177 | MICOS complex subunit Mic60 (Fragment) OS=Rattus norvegicus OX=10116 GN=Immt PE=1 SV=1 |
| tr\|A0A0G2JVH4\|A0A0G2JVH | **65** | 147 | 20.88 | 2 | 2 | 45168 | 1 | 1 | 1 | N | 86230 | MICOS complex subunit MIC60 OS=Rattus norvegicus OX=10116 GN=Immt PE=1 SV=1 |
| tr\|D4A4K4\|D4A4K4_RAT | **30** | 173 | 44520 | 0 | 0 | 475940 | 1 | 1 | 2 | N | 418626 | Vacuolar protein sorting 13 homolog C OS=Rattus norvegicus OX=10116 GN=Vps13c PE=1 SV=2 |
