## Supplementary material for "Permeability transition pore-related changes in the proteome and channel activity of ATP synthase dimers and monomers": RHM PTP Dimer of V compl.

| Protein Grc | Protein ID | Accession | -10lgP | Coverage (' | Coverage (' | Area Samp | #Peptides | #Unique | #Spec Sam | PTM | Avg. Mass | Description |
|---|---|---|---|---|---|---|---|---|---|---|---|---|
| 1 | 5 | P10719\|AT | 390.79 | 47 | 47 | 1.33E+09 | 124 | 120 | 681 | Oxidation ( | 56354 | ATP synthase subunit beta mitochondrial OS=Rattus norvegicus OX=10116 GN=Atp5f1b PE=1 SV=2 |
| 1 | 6 | tr\|G3V6D3 | 390.79 | 47 | 47 | 1.33E+09 | 124 | 120 | 681 | Oxidation ( | 56345 | ATP synthase subunit beta OS=Rattus norvegicus OX=10116 GN=Atp5f1b PE=1 SV=1 |
| 2 | 8 | P15999\|AT | 331.26 | 41 | 41 | 7.09E+08 | 109 | 107 | 645 | Carbamidomethylation | | |
| 2 | 9 | tr\|F1LP05\| | 331.26 | 41 | 41 | 7.09E+08 | 109 | 107 | 645 | Carbamidomethylation | | |
| 6 | 65 | P31399\|AT | 277.38 | 61 | 61 | 8.53E+07 | 30 | 30 | 128 | Oxidation ( | 18763 | ATP synthase subunit d mitochondrial OS=Rattus norvegicus OX=10116 GN=Atp5pd PE=1 SV=3 |
| 4 | 63 | P35435\|AT | 262.75 | 37 | 37 | 7.19E+07 | 37 | 37 | 141 | Oxidation ( | 30191 | ATP synthase subunit gamma mitochondrial OS=Rattus norvegicus OX=10116 GN=Atp5f1c PE=1 SV=2 |
| 4 | 64 | tr\|Q6QI09\| | 262.75 | 17 | 17 | 7.19E+07 | 37 | 37 | 141 | Oxidation ( | 67721 | ATP synthase subunit gamma mitochondrial OS=Rattus norvegicus OX=10116 GN=Taf3 PE=1 SV=1 |
| 5 | 20 | Q66HF1\|N | 252.49 | 21 | 21 | 4.49E+07 | 39 | 39 | 140 | Carbamido | 79412 | NADH-ubiquinone oxidoreductase 75 kDa subunit mitochondrial OS=Rattus norvegicus OX=10116 GN=Ndufs1 PE=1 SV=1 |
| 12 | 212 | tr\|B2RYS8\| | 246.45 | 48 | 48 | 2.32E+07 | 23 | 23 | 61 | Oxidation ( | 21959 | NADH dehydrogenase [ubiquinone] 1 beta subcomplex subunit 8 mitochondrial OS=Rattus norvegicus OX=10116 GN=Ndufb8 PE=1 SV=1 |
| 3 | 37 | P19234\|NI | 241.39 | 44 | 44 | 1.07E+08 | 30 | 30 | 145 | | 27378 | NADH dehydrogenase [ubiquinone] flavoprotein 2 mitochondrial OS=Rattus norvegicus OX=10116 GN=Ndufv2 PE=1 SV=2 |
| 15 | 60 | tr\|D4A565 | 232.34 | 34 | 34 | 1.33E+07 | 15 | 15 | 47 | | 21664 | NADH dehydrogenase (Ubiquinone) 1 beta subcomplex 5 (Predicted) isoform CRA_b OS=Rattus norvegicus OX=10116 GN=Ndufb5 PE=1 SV=1 |
| 9 | 26 | Q641Y2\|NI | 204.62 | 26 | 26 | 3.90E+07 | 29 | 29 | 80 | Oxidation ( | 52562 | NADH dehydrogenase [ubiquinone] iron-sulfur protein 2 mitochondrial OS=Rattus norvegicus OX=10116 GN=Ndufs2 PE=1 SV=1 |
| 13 | 77 | tr\|Q5PQZ9 | 201.91 | 37 | 37 | 2.32E+07 | 13 | 13 | 56 | | 14359 | NADH dehydrogenase [ubiquinone] 1 subunit C2 OS=Rattus norvegicus OX=10116 GN=Ndufc2 PE=1 SV=1 |
| 35 | 248 | tr\|A9UMW | 198.26 | 37 | 37 | 1.59E+07 | 9 | 9 | 24 | Oxidation ( | 9255 | Ndufa3 protein (Fragment) OS=Rattus norvegicus OX=10116 GN=Ndufa3 PE=2 SV=1 |
| 35 | 249 | tr\|M0RB63 | 198.26 | 37 | 37 | 1.59E+07 | 9 | 9 | 24 | Oxidation ( | 9372 | RCG63041 OS=Rattus norvegicus OX=10116 GN=LOC684509 PE=4 SV=1 |
| 35 | 250 | tr\|A0A0G2 | 198.26 | 34 | 34 | 1.59E+07 | 9 | 9 | 24 | Oxidation ( | 10170 | NADH:ubiquinone oxidoreductase subunit A3 OS=Rattus norvegicus OX=10116 GN=Ndufa3 PE=1 SV=1 |
| 24 | 59 | tr\|D3ZFQ8 | 196.84 | 13 | 13 | 1.62E+07 | 9 | 9 | 37 | | 35435 | Cytochrome c-1 OS=Rattus norvegicus OX=10116 GN=Cyc1 PE=1 SV=3 |
| 20 | 160 | tr\|D4A7L4\| | 196.05 | 26 | 26 | 2.24E+07 | 9 | 9 | 41 | | 17634 | NADH dehydrogenase (Ubiquinone) 1 beta subcomplex 11 (Predicted) OS=Rattus norvegicus OX=10116 GN=Ndufb11 PE=1 SV=1 |
| 11 | 49 | Q5BK63\|NI | 195.5 | 30 | 30 | 1.85E+07 | 21 | 21 | 61 | | 42559 | NADH dehydrogenase [ubiquinone] 1 alpha subcomplex subunit 9 mitochondrial OS=Rattus norvegicus OX=10116 GN=Ndufa9 PE=1 SV=2 |
| 36 | 58 | P20788\|UC | 186.51 | 17 | 17 | 1.10E+07 | 9 | 9 | 21 | | 29446 | Cytochrome b-c1 complex subunit Rieske mitochondrial OS=Rattus norvegicus OX=10116 GN=Uqcrfs1 PE=1 SV=2 |
| 18 | 252 | tr\|D3ZF13\| | 185.52 | 23 | 23 | 1.58E+07 | 13 | 13 | 44 | Oxidation ( | 17514 | Acyl carrier protein OS=Rattus norvegicus OX=10116 GN=Ndufab1 PE=1 SV=1 |
| 7 | 75 | Q6PDU7\|A | 183.33 | 48 | 48 | 3.41E+07 | 19 | 18 | 89 | Oxidation ( | 11433 | ATP synthase subunit g mitochondrial OS=Rattus norvegicus OX=10116 GN=Atp5mg PE=1 SV=2 |
| 25 | 211 | tr\|F1LXA0\| | 173.13 | 29 | 29 | 1.78E+07 | 8 | 8 | 37 | | 17178 | NADH dehydrogenase [ubiquinone] 1 alpha subcomplex subunit 12 OS=Rattus norvegicus OX=10116 GN=Ndufa12 PE=1 SV=2 |
| 8 | 168 | P19511\|AT | 173.06 | 20 | 20 | 1.15E+08 | 18 | 18 | 81 | | 28869 | ATP synthase F(0) complex subunit B1 mitochondrial OS=Rattus norvegicus OX=10116 GN=Atp5pb PE=1 SV=1 |
| 22 | 199 | tr\|B0BNE6 | 172.57 | 25 | 25 | 1.01E+07 | 13 | 13 | 39 | | 23970 | NADH dehydrogenase (Ubiquinone) Fe-S protein 8 (Predicted) isoform CRA_a OS=Rattus norvegicus OX=10116 GN=Ndufs8 PE=1 SV=1 |
| 42 | 25 | Q68FY0\|Q | 165.24 | 14 | 14 | 3.58E+06 | 9 | 9 | 16 | Carbamido | 52849 | Cytochrome b-c1 complex subunit 1 mitochondrial OS=Rattus norvegicus OX=10116 GN=Uqcrc1 PE=1 SV=1 |
| 31 | 198 | Q80W89\|N | 165.2 | 18 | 18 | 1.66E+07 | 7 | 7 | 26 | | 14854 | NADH dehydrogenase [ubiquinone] 1 alpha subcomplex subunit 11 OS=Rattus norvegicus OX=10116 GN=Ndufa11 PE=2 SV=1 |
| 14 | 74 | tr\|D3ZG43 | 164.22 | 34 | 34 | 1.91E+07 | 22 | 22 | 52 | | 30226 | NADH dehydrogenase (Ubiquinone) Fe-S protein 3 (Predicted) isoform CRA_c OS=Rattus norvegicus OX=10116 GN=Ndufs3 PE=1 SV=1 |
| 10 | 76 | Q06647\|AT | 162.91 | 29 | 29 | 4.16E+07 | 19 | 18 | 72 | | 23398 | ATP synthase subunit O mitochondrial OS=Rattus norvegicus OX=10116 GN=Atp5po PE=1 SV=1 |
| 33 | 45 | P00507\|AA | 162.61 | 17 | 17 | 5.18E+06 | 11 | 11 | 25 | Oxidation ( | 47314 | Aspartate aminotransferase mitochondrial OS=Rattus norvegicus OX=10116 GN=Got2 PE=1 SV=2 |
| 21 | 161 | tr\|A0A0G2 | 161.02 | 44 | 44 | 2.56E+07 | 13 | 13 | 39 | Carbamido | 19965 | NADH dehydrogenase [ubiquinone] 1 alpha subcomplex subunit 8 OS=Rattus norvegicus OX=10116 GN=Ndufa8 PE=1 SV=1 |
| 23 | 172 | tr\|D4A0T0 | 158.95 | 27 | 27 | 1.28E+07 | 10 | 10 | 38 | | 20859 | NADH:ubiquinone oxidoreductase subunit B10 OS=Rattus norvegicus OX=10116 GN=Ndufb10 PE=1 SV=1 |
| 53 | 200 | P11240\|CC | 155.04 | 29 | 29 | 2.22E+06 | 8 | 8 | 12 | | 16130 | Cytochrome c oxidase subunit 5A mitochondrial OS=Rattus norvegicus OX=10116 GN=Cox5a PE=1 SV=1 |
| 26 | 313 | tr\|D3ZZ21\| | 151.15 | 49 | 49 | 8.42E+06 | 10 | 10 | 35 | Oxidation ( | 15638 | NADH dehydrogenase (Ubiquinone) 1 beta subcomplex 6 (Predicted) OS=Rattus norvegicus OX=10116 GN=Ndufb6 PE=1 SV=1 |
| 45 | 251 | P80432\|CC | 143.95 | 21 | 21 | 4.75E+06 | 4 | 4 | 15 | | 7375 | Cytochrome c oxidase subunit 7C mitochondrial OS=Rattus norvegicus OX=10116 GN=Cox7c PE=1 SV=2 |
| 19 | 171 | Q561S0\|NI | 142.62 | 24 | 24 | 7.63E+06 | 14 | 4 | 43 | Oxidation ( | 40493 | NADH dehydrogenase [ubiquinone] 1 alpha subcomplex subunit 10 mitochondrial OS=Rattus norvegicus OX=10116 GN=Ndufa10 PE=1 SV=1 |
| 27 | 137 | tr\|Q06QK5 | 141.53 | 10 | 10 | 9.31E+06 | 12 | 12 | 32 | Oxidation ( | 68606 | NADH-ubiquinone oxidoreductase chain 5 OS=Rattus norvegicus OX=10116 GN=ND5 PE=3 SV=1 |
| 27 | 138 | P11661\|NU | 141.53 | 10 | 10 | 9.31E+06 | 12 | 12 | 32 | Oxidation ( | 68618 | NADH-ubiquinone oxidoreductase chain 5 OS=Rattus norvegicus OX=10116 GN=Mtnd5 PE=3 SV=3 |
| 27 | 139 | tr\|Q06QG6 | 141.53 | 10 | 10 | 9.31E+06 | 12 | 12 | 32 | Oxidation ( | 68588 | NADH-ubiquinone oxidoreductase chain 5 OS=Rattus norvegicus OX=10116 GN=ND5 PE=3 SV=1 |
| 27 | 140 | tr\|Q06QA1 | 141.53 | 10 | 10 | 9.31E+06 | 12 | 12 | 32 | Oxidation ( | 68574 | NADH-ubiquinone oxidoreductase chain 5 OS=Rattus norvegicus OX=10116 GN=ND5 PE=3 SV=1 |
| 27 | 141 | tr\|Q8SEZ0\| | 141.53 | 10 | 10 | 9.31E+06 | 12 | 12 | 32 | Oxidation ( | 68618 | NADH-ubiquinone oxidoreductase chain 5 OS=Rattus norvegicus OX=10116 GN=Mt-nd5 PE=3 SV=1 |
| 27 | 142 | tr\|A0A096 | 141.53 | 10 | 10 | 9.31E+06 | 12 | 12 | 32 | Oxidation ( | 68584 | NADH-ubiquinone oxidoreductase chain 5 OS=Rattus norvegicus OX=10116 GN=ND5 PE=3 SV=1 |
| 43 | 83 | tr\|A0A411 | 140.87 | 7 | 7 | 1.96E+06 | 6 | 6 | 15 | | 40144 | Cytochrome b (Fragment) OS=Rattus norvegicus OX=10116 GN=Cytb PE=3 SV=1 |
| 43 | 84 | tr\|A0A0U2 | 140.87 | 7 | 7 | 1.96E+06 | 6 | 6 | 15 | | 40981 | Cytochrome b (Fragment) OS=Rattus norvegicus OX=10116 PE=3 SV=1 |
| 43 | 85 | tr\|A0A0U2 | 140.87 | 7 | 7 | 1.96E+06 | 6 | 6 | 15 | | 41037 | Cytochrome b (Fragment) OS=Rattus norvegicus OX=10116 PE=3 SV=1 |
| 43 | 86 | tr\|A0A411 | 140.87 | 7 | 7 | 1.96E+06 | 6 | 6 | 15 | | 41017 | Cytochrome b (Fragment) OS=Rattus norvegicus OX=10116 GN=Cytb PE=3 SV=1 |
| 43 | 87 | tr\|F8QU31 | 140.87 | 7 | 7 | 1.96E+06 | 6 | 6 | 15 | | 42181 | Cytochrome b (Fragment) OS=Rattus norvegicus OX=10116 GN=cytb PE=3 SV=1 |
| 43 | 88 | tr\|A0A411 | 140.87 | 7 | 7 | 1.96E+06 | 6 | 6 | 15 | | 42238 | Cytochrome b (Fragment) OS=Rattus norvegicus OX=10116 GN=Cytb PE=3 SV=1 |
| 43 | 89 | tr\|L0L4L8\| | 140.87 | 7 | 7 | 1.96E+06 | 6 | 6 | 15 | | 42470 | Cytochrome b (Fragment) OS=Rattus norvegicus OX=10116 GN=cytb PE=3 SV=1 |
| 43 | 91 | tr\|A0A140 | 140.87 | 7 | 7 | 1.96E+06 | 6 | 6 | 15 | | 42574 | Cytochrome b (Fragment) OS=Rattus norvegicus OX=10116 PE=3 SV=1 |
| 43 | 92 | tr\|A0A140 | 140.87 | 7 | 7 | 1.96E+06 | 6 | 6 | 15 | | 42588 | Cytochrome b (Fragment) OS=Rattus norvegicus OX=10116 PE=3 SV=1 |
| 43 | 93 | tr\|A0A140 | 140.87 | 7 | 7 | 1.96E+06 | 6 | 6 | 15 | | 42527 | Cytochrome b (Fragment) OS=Rattus norvegicus OX=10116 PE=3 SV=1 |
| 43 | 95 | tr\|A0A385 | 140.87 | 7 | 7 | 1.96E+06 | 6 | 6 | 15 | | 42766 | Cytochrome b (Fragment) OS=Rattus norvegicus OX=10116 PE=3 SV=1 |
| 43 | 96 | tr\|A0A3Q8 | 140.87 | 7 | 7 | 1.96E+06 | 6 | 6 | 15 | | 42712 | Cytochrome b (Fragment) OS=Rattus norvegicus OX=10116 GN=Cytb PE=3 SV=1 |
| 43 | 97 | tr\|A0A3Q8 | 140.87 | 7 | 7 | 1.96E+06 | 6 | 6 | 15 | | 42622 | Cytochrome b (Fragment) OS=Rattus norvegicus OX=10116 GN=Cytb PE=3 SV=1 |
| 43 | 98 | tr\|A0A3S6 | 140.87 | 7 | 7 | 1.96E+06 | 6 | 6 | 15 | | 42702 | Cytochrome b (Fragment) OS=Rattus norvegicus OX=10116 GN=Cytb PE=3 SV=1 |
| 43 | 100 | tr\|A0A3S6 | 140.87 | 7 | 7 | 1.96E+06 | 6 | 6 | 15 | | 42698 | Cytochrome b (Fragment) OS=Rattus norvegicus OX=10116 GN=Cytb PE=3 SV=1 |

| | | | | | | | | | | | | |
|---|---|---|---|---|---|---|---|---|---|---|---|---|
| 43 | 101 | tr\|A0A097I | 140.87 | 7 | 7 | 1.96E+06 | 6 | 6 | 15 | | 42865 | Cytochrome b OS=Rattus norvegicus OX=10116 GN=CYTB PE=3 SV=1 |
| 43 | 102 | tr\|A0A220I | 140.87 | 7 | 7 | 1.96E+06 | 6 | 6 | 15 | | 43018 | Cytochrome b OS=Rattus norvegicus OX=10116 PE=3 SV=1 |
| 43 | 103 | tr\|D6NSP2 | 140.87 | 7 | 7 | 1.96E+06 | 6 | 6 | 15 | | 42945 | Cytochrome b (Fragment) OS=Rattus norvegicus OX=10116 GN=cytb PE=3 SV=1 |
| 43 | 104 | tr\|D6NSS6 | 140.87 | 7 | 7 | 1.96E+06 | 6 | 6 | 15 | | 42993 | Cytochrome b (Fragment) OS=Rattus norvegicus OX=10116 GN=cytb PE=3 SV=1 |
| 43 | 106 | tr\|D6NSQ3 | 140.87 | 7 | 7 | 1.96E+06 | 6 | 6 | 15 | | 43012 | Cytochrome b (Fragment) OS=Rattus norvegicus OX=10116 GN=cytb PE=3 SV=1 |
| 43 | 107 | tr\|D6NSR3 | 140.87 | 7 | 7 | 1.96E+06 | 6 | 6 | 15 | | 42986 | Cytochrome b (Fragment) OS=Rattus norvegicus OX=10116 GN=cytb PE=3 SV=1 |
| 43 | 108 | tr\|D6NSP4 | 140.87 | 7 | 7 | 1.96E+06 | 6 | 6 | 15 | | 43016 | Cytochrome b (Fragment) OS=Rattus norvegicus OX=10116 GN=cytb PE=3 SV=1 |
| 43 | 109 | tr\|D6NSQ8 | 140.87 | 7 | 7 | 1.96E+06 | 6 | 6 | 15 | | 43016 | Cytochrome b (Fragment) OS=Rattus norvegicus OX=10116 GN=cytb PE=3 SV=1 |
| 43 | 110 | tr\|D6NSR8 | 140.87 | 7 | 7 | 1.96E+06 | 6 | 6 | 15 | | 42998 | Cytochrome b (Fragment) OS=Rattus norvegicus OX=10116 GN=cytb PE=3 SV=1 |
| 43 | 111 | tr\|F8QU37 | 140.87 | 7 | 7 | 1.96E+06 | 6 | 6 | 15 | | 42992 | Cytochrome b (Fragment) OS=Rattus norvegicus OX=10116 GN=cytb PE=3 SV=1 |
| 43 | 112 | tr\|F2Q6S6\| | 140.87 | 7 | 7 | 1.96E+06 | 6 | 6 | 15 | | 42938 | Cytochrome b (Fragment) OS=Rattus norvegicus OX=10116 GN=cytb PE=3 SV=1 |
| 43 | 113 | tr\|A0A0A1 | 140.87 | 7 | 7 | 1.96E+06 | 6 | 6 | 15 | | 42989 | Cytochrome b OS=Rattus norvegicus OX=10116 GN=CYTB PE=3 SV=1 |
| 43 | 114 | tr\|D6NSS7 | 140.87 | 7 | 7 | 1.96E+06 | 6 | 6 | 15 | | 42948 | Cytochrome b (Fragment) OS=Rattus norvegicus OX=10116 GN=cytb PE=3 SV=1 |
| 43 | 115 | tr\|Q8SEY9\| | 140.87 | 7 | 7 | 1.96E+06 | 6 | 6 | 15 | | 43015 | Cytochrome b OS=Rattus norvegicus OX=10116 GN=cytb PE=3 SV=1 |
| 43 | 116 | tr\|D6NSR7 | 140.87 | 7 | 7 | 1.96E+06 | 6 | 6 | 15 | | 42982 | Cytochrome b (Fragment) OS=Rattus norvegicus OX=10116 GN=cytb PE=3 SV=1 |
| 43 | 117 | tr\|A0A220I | 140.87 | 7 | 7 | 1.96E+06 | 6 | 6 | 15 | | 42968 | Cytochrome b OS=Rattus norvegicus OX=10116 PE=3 SV=1 |
| 43 | 118 | tr\|F2Q6S5\| | 140.87 | 7 | 7 | 1.96E+06 | 6 | 6 | 15 | | 42952 | Cytochrome b (Fragment) OS=Rattus norvegicus OX=10116 GN=cytb PE=3 SV=1 |
| 43 | 119 | tr\|D6NSQ6 | 140.87 | 7 | 7 | 1.96E+06 | 6 | 6 | 15 | | 42952 | Cytochrome b (Fragment) OS=Rattus norvegicus OX=10116 GN=cytb PE=3 SV=1 |
| 43 | 120 | tr\|D6NSQ0 | 140.87 | 7 | 7 | 1.96E+06 | 6 | 6 | 15 | | 42968 | Cytochrome b (Fragment) OS=Rattus norvegicus OX=10116 GN=cytb PE=3 SV=1 |
| 43 | 121 | tr\|Q5UAI7 | 140.87 | 7 | 7 | 1.96E+06 | 6 | 6 | 15 | | 42998 | Cytochrome b OS=Rattus norvegicus OX=10116 GN=CYTB PE=3 SV=1 |
| 43 | 122 | tr\|D6NSR2 | 140.87 | 7 | 7 | 1.96E+06 | 6 | 6 | 15 | | 43073 | Cytochrome b (Fragment) OS=Rattus norvegicus OX=10116 GN=cytb PE=3 SV=1 |
| 43 | 123 | tr\|A0A0S12 | 140.87 | 7 | 7 | 1.96E+06 | 6 | 6 | 15 | | 43002 | Cytochrome b OS=Rattus norvegicus OX=10116 GN=CYTB PE=3 SV=1 |
| 43 | 124 | tr\|D6NSR1 | 140.87 | 7 | 7 | 1.96E+06 | 6 | 6 | 15 | | 43002 | Cytochrome b (Fragment) OS=Rattus norvegicus OX=10116 GN=cytb PE=3 SV=1 |
| 43 | 125 | tr\|Q8HIC4\| | 140.87 | 7 | 7 | 1.96E+06 | 6 | 6 | 15 | | 43012 | Cytochrome b OS=Rattus norvegicus OX=10116 GN=Mt-cyb PE=3 SV=1 |
| 43 | 126 | tr\|A0A220I | 140.87 | 7 | 7 | 1.96E+06 | 6 | 6 | 15 | | 42968 | Cytochrome b OS=Rattus norvegicus OX=10116 PE=3 SV=1 |
| 43 | 128 | tr\|A0A0S12 | 140.87 | 7 | 7 | 1.96E+06 | 6 | 6 | 15 | | 43016 | Cytochrome b OS=Rattus norvegicus OX=10116 GN=CYTB PE=3 SV=1 |
| 43 | 129 | tr\|R9TKN1 | 140.87 | 7 | 7 | 1.96E+06 | 6 | 6 | 15 | | 43012 | Cytochrome b OS=Rattus norvegicus OX=10116 GN=CYTB PE=3 SV=1 |
| 43 | 130 | P00159\|CY | 140.87 | 7 | 7 | 1.96E+06 | 6 | 6 | 15 | | 43012 | Cytochrome b OS=Rattus norvegicus OX=10116 GN=Mt-Cyb PE=3 SV=3 |
| 43 | 131 | tr\|L0N311\| | 140.87 | 7 | 7 | 1.96E+06 | 6 | 6 | 15 | | 43045 | Cytochrome b (Fragment) OS=Rattus norvegicus OX=10116 GN=cytb PE=3 SV=1 |
| 43 | 132 | tr\|A0A220I | 140.87 | 7 | 7 | 1.96E+06 | 6 | 6 | 15 | | 42970 | Cytochrome b OS=Rattus norvegicus OX=10116 PE=3 SV=1 |
| 43 | 133 | tr\|H2KXA0 | 140.87 | 7 | 7 | 1.96E+06 | 6 | 6 | 15 | | 42978 | Cytochrome b (Fragment) OS=Rattus norvegicus OX=10116 GN=cytb PE=3 SV=1 |
| 43 | 134 | tr\|A0A096 | 140.87 | 7 | 7 | 1.96E+06 | 6 | 6 | 15 | | 43042 | Cytochrome b (Fragment) OS=Rattus norvegicus OX=10116 PE=3 SV=1 |
| 43 | 135 | tr\|D6NSP5 | 140.87 | 7 | 7 | 1.96E+06 | 6 | 6 | 15 | | 43012 | Cytochrome b (Fragment) OS=Rattus norvegicus OX=10116 GN=cytb PE=3 SV=1 |
| 16 | 80 | tr\|Q5XIH3\| | 134.88 | 14 | 14 | 1.71E+07 | 12 | 11 | 46 | | 50731 | NADH dehydrogenase [ubiquinone] flavoprotein 1 mitochondrial OS=Rattus norvegicus OX=10116 GN=Ndufv1 PE=1 SV=1 |
| 17 | 50 | P32551\|QC | 134.81 | 11 | 11 | 2.08E+07 | 10 | 10 | 46 | | 48396 | Cytochrome b-c1 complex subunit 2 mitochondrial OS=Rattus norvegicus OX=10116 GN=Uqcrc2 PE=1 SV=2 |
| 39 | 219 | tr\|D3ZS58\| | 133.46 | 12 | 12 | 7.04E+06 | 6 | 6 | 19 | | 10845 | NADH dehydrogenase [ubiquinone] 1 alpha subcomplex subunit 2 OS=Rattus norvegicus OX=10116 GN=Ndufa2 PE=1 SV=1 |
| 30 | 56 | tr\|F1LPG5\| | 128.69 | 35 | 35 | 6.99E+06 | 9 | 9 | 26 | | 15064 | NADH:ubiquinone oxidoreductase subunit B4 OS=Rattus norvegicus OX=10116 GN=Ndufb4 PE=1 SV=1 |
| 30 | 57 | tr\|F1M7T1 | 128.69 | 35 | 35 | 6.99E+06 | 9 | 9 | 26 | | 15160 | NADH dehydrogenase (ubiquinone) 1 beta subcomplex 4 OS=Rattus norvegicus OX=10116 GN=LOC100361934 PE=4 SV=2 |
| 49 | 532 | P67779\|PH | 128.14 | 18 | 18 | 2.91E+06 | 8 | 8 | 12 | | 29820 | Prohibitin OS=Rattus norvegicus OX=10116 GN=Phb PE=1 SV=1 |
| 29 | 319 | P29419\|AT | 125.84 | 51 | 51 | 1.00E+07 | 10 | 10 | 28 | | 8255 | ATP synthase subunit e mitochondrial OS=Rattus norvegicus OX=10116 GN=Atp5me PE=1 SV=3 |
| 34 | 33 | tr\|S5RZM8 | 125.74 | 32 | 32 | 4.45E+06 | 10 | 10 | 24 | Oxidation ( | 25958 | Cytochrome c oxidase subunit 2 OS=Rattus norvegicus OX=10116 GN=COX2 PE=3 SV=1 |
| 34 | 34 | P00406\|CC | 125.74 | 32 | 32 | 4.45E+06 | 10 | 10 | 24 | Oxidation ( | 25928 | Cytochrome c oxidase subunit 2 OS=Rattus norvegicus OX=10116 GN=Mtco2 PE=1 SV=3 |
| 34 | 28 | tr\|Q5UAJ6 | 125.74 | 32 | 32 | 4.45E+06 | 10 | 10 | 24 | Oxidation ( | 25942 | Cytochrome c oxidase subunit 2 OS=Rattus norvegicus OX=10116 GN=COX2 PE=3 SV=1 |
| 34 | 35 | tr\|Q8SEZ5\| | 125.74 | 32 | 32 | 4.45E+06 | 10 | 10 | 24 | Oxidation ( | 25928 | Cytochrome c oxidase subunit 2 OS=Rattus norvegicus OX=10116 GN=Mt-co2 PE=1 SV=1 |
| 34 | 36 | tr\|A0A097I | 125.74 | 32 | 32 | 4.45E+06 | 10 | 10 | 24 | Oxidation ( | 25894 | Cytochrome c oxidase subunit 2 OS=Rattus norvegicus OX=10116 GN=COX2 PE=3 SV=1 |
| 32 | 274 | tr\|A0A1W2 | 125.06 | 23 | 23 | 5.86E+05 | 11 | 1 | 26 | Oxidation ( | 40544 | NADH dehydrogenase [ubiquinone] 1 alpha subcomplex subunit 10 mitochondrial OS=Rattus norvegicus OX=10116 GN=Ndufa10l1 PE=3 SV=1 |
| 41 | 3899 | tr\|B5DEL8\| | 120.07 | 22 | 22 | 4.72E+06 | 6 | 6 | 17 | | 12700 | NADH dehydrogenase (Ubiquinone) Fe-S protein 5 OS=Rattus norvegicus OX=10116 GN=Ndufs5 PE=1 SV=1 |
| 41 | 3900 | tr\|A0A0G2 | 120.07 | 22 | 22 | 4.72E+06 | 6 | 6 | 17 | | 12730 | Uncharacterized protein OS=Rattus norvegicus OX=10116 PE=4 SV=1 |
| 50 | 253 | P10817\|CX | 119.87 | 26 | 26 | 4.72E+06 | 5 | 5 | 12 | | 10487 | Cytochrome c oxidase subunit 6A2 mitochondrial (Fragment) OS=Rattus norvegicus OX=10116 GN=Cox6a2 PE=1 SV=3 |
| 50 | 254 | tr\|G3V8M4 | 119.87 | 25 | 25 | 4.72E+06 | 5 | 5 | 12 | | 10802 | Cytochrome c oxidase subunit 6A mitochondrial OS=Rattus norvegicus OX=10116 GN=Cox6a2 PE=3 SV=1 |
| 40 | 827 | tr\|D4A3V2 | 116.96 | 33 | 33 | 2.86E+06 | 12 | 12 | 19 | | 15224 | NADH dehydrogenase [ubiquinone] 1 alpha subcomplex subunit 6 OS=Rattus norvegicus OX=10116 GN=Ndufa6 PE=1 SV=1 |
| 44 | 169 | tr\|A9UMV9 | 115.75 | 21 | 21 | 2.97E+07 | 4 | 4 | 15 | | 12500 | NADH:ubiquinone oxidoreductase subunit A7 OS=Rattus norvegicus OX=10116 GN=Ndufa7 PE=1 SV=1 |
| 48 | 233 | tr\|Q5UAJ5 | 115.4 | 19 | 19 | 1.01E+07 | 4 | 4 | 13 | Oxidation ( | 7642 | ATP synthase protein 8 OS=Rattus norvegicus OX=10116 GN=ATP8 PE=3 SV=1 |
| 48 | 234 | P11608\|AT | 115.4 | 19 | 19 | 1.01E+07 | 4 | 4 | 13 | Oxidation ( | 7630 | ATP synthase protein 8 OS=Rattus norvegicus OX=10116 GN=Mt-atp8 PE=1 SV=2 |
| 48 | 235 | tr\|Q8SEZ4\| | 115.4 | 19 | 19 | 1.01E+07 | 4 | 4 | 13 | Oxidation ( | 7632 | ATP synthase protein 8 OS=Rattus norvegicus OX=10116 GN=ATPase8 PE=3 SV=1 |
| 48 | 236 | tr\|Q8HIC8\| | 115.4 | 19 | 19 | 1.01E+07 | 4 | 4 | 13 | Oxidation ( | 7630 | ATP synthase protein 8 OS=Rattus norvegicus OX=10116 GN=Mt-atp8 PE=3 SV=1 |
| 28 | 72 | tr\|D3ZE15\| | 114.68 | 28 | 28 | 5.08E+06 | 8 | 7 | 28 | | 16777 | NADH:ubiquinone oxidoreductase subunit A13 OS=Rattus norvegicus OX=10116 GN=Ndufa13 PE=1 SV=1 |
| 54 | 318 | Q5M9I5\|Q | 111.53 | 18 | 18 | 3.13E+06 | 4 | 4 | 12 | | 10424 | Cytochrome b-c1 complex subunit 6 mitochondrial OS=Rattus norvegicus OX=10116 GN=Uqcrh PE=3 SV=1 |

| | | | | | | | | | | | | |
|---|---|---|---|---|---|---|---|---|---|---|---|---|
| 55 | 238 | Q5XIF3\|ND | 107.99 | 12 | 12 | 4.45E+06 | 3 | 3 | 11 | | 19741 | NADH dehydrogenase [ubiquinone] iron-sulfur protein 4 mitochondrial OS=Rattus norvegicus OX=10116 GN=Ndufs4 PE=1 SV=1 |
| 71 | 73 | Q63362\|NI | 97.02 | 17 | 17 | 1.86E+06 | 4 | 4 | 6 | | 13412 | NADH dehydrogenase [ubiquinone] 1 alpha subcomplex subunit 5 OS=Rattus norvegicus OX=10116 GN=Ndufa5 PE=1 SV=3 |
| 75 | 3930 | tr\|Q35733 | 93.65 | 14 | 14 | 1.29E+06 | 3 | 3 | 6 | | 8389 | NADH-ubiquinone oxidoreductase chain 6 (Fragment) OS=Rattus norvegicus OX=10116 PE=3 SV=1 |
| 75 | 3921 | tr\|Q06QD9 | 93.65 | 6 | 6 | 1.29E+06 | 3 | 3 | 6 | | 18943 | NADH-ubiquinone oxidoreductase chain 6 OS=Rattus norvegicus OX=10116 GN=ND6 PE=3 SV=1 |
| 75 | 3922 | tr\|Q7HKW | 93.65 | 6 | 6 | 1.29E+06 | 3 | 3 | 6 | | 18957 | NADH-ubiquinone oxidoreductase chain 6 OS=Rattus norvegicus OX=10116 GN=NADH6 PE=3 SV=1 |
| 47 | 376 | tr\|Q5RJN0 | 91.45 | 21 | 21 | 4.00E+06 | 5 | 5 | 13 | Oxidation ( | 23945 | NADH dehydrogenase (Ubiquinone) Fe-S protein 7 OS=Rattus norvegicus OX=10116 GN=Ndufs7 PE=1 SV=1 |
| 77 | 17 | Q9ER34\|A( | 89.78 | 3 | 3 | 5.92E+05 | 3 | 3 | 5 | | 85433 | Aconitate hydratase mitochondrial OS=Rattus norvegicus OX=10116 GN=Aco2 PE=1 SV=2 |
| 64 | 16 | P02563\|MY | 86.5 | 3 | 3 | 1.49E+06 | 5 | 5 | 7 | Oxidation ( | 223506 | Myosin-6 OS=Rattus norvegicus OX=10116 GN=Myh6 PE=1 SV=2 |
| 61 | 61 | Q63704\|CF | 83.57 | 3 | 3 | 5.01E+05 | 3 | 3 | 7 | | 88217 | Carnitine O-palmitoyltransferase 1 muscle isoform OS=Rattus norvegicus OX=10116 GN=Cpt1b PE=1 SV=1 |
| 58 | 23 | tr\|Q6P9Y4 | 81.73 | 10 | 10 | 1.54E+06 | 5 | 5 | 10 | | 32904 | ADP/ATP translocase 1 OS=Rattus norvegicus OX=10116 GN=Slc25a4 PE=1 SV=1 |
| 58 | 24 | Q05962\|AI | 81.73 | 10 | 10 | 1.54E+06 | 5 | 5 | 10 | | 32989 | ADP/ATP translocase 1 OS=Rattus norvegicus OX=10116 GN=Slc25a4 PE=1 SV=3 |
| 58 | 27 | Q09073\|AI | 81.73 | 10 | 10 | 1.54E+06 | 5 | 5 | 10 | | 32901 | ADP/ATP translocase 2 OS=Rattus norvegicus OX=10116 GN=Slc25a5 PE=1 SV=3 |
| 46 | 3953 | tr\|G3V7Y3 | 81.42 | 11 | 11 | 7.35E+06 | 5 | 5 | 14 | | 17563 | ATP synthase subunit delta mitochondrial OS=Rattus norvegicus OX=10116 GN=Atp5f1d PE=1 SV=1 |
| 46 | 3951 | tr\|B1WBP7 | 81.42 | 15 | 15 | 7.35E+06 | 5 | 5 | 14 | | 12884 | ATP synthase subunit delta mitochondrial OS=Rattus norvegicus OX=10116 GN=Atp5f1d PE=1 SV=1 |
| 85 | 263 | P35171\|CX | 80.03 | 12 | 12 | 1.15E+06 | 2 | 2 | 4 | | 9353 | Cytochrome c oxidase subunit 7A2 mitochondrial OS=Rattus norvegicus OX=10116 GN=Cox7a2 PE=1 SV=1 |
| 85 | 264 | tr\|B2RYS0\| | 80.03 | 12 | 12 | 1.15E+06 | 2 | 2 | 4 | | 9353 | Cox7a2 protein OS=Rattus norvegicus OX=10116 GN=Cox7a2 PE=2 SV=1 |
| 67 | 29 | Q60587\|E( | 79.98 | 8 | 8 | 9.23E+05 | 5 | 5 | 7 | | 51414 | Trifunctional enzyme subunit beta mitochondrial OS=Rattus norvegicus OX=10116 GN=Hadhb PE=1 SV=1 |
| 67 | 40 | tr\|A0A0G2 | 79.98 | 7 | 7 | 9.23E+05 | 5 | 5 | 7 | | 52568 | Trifunctional enzyme subunit beta mitochondrial OS=Rattus norvegicus OX=10116 GN=Hadhb PE=1 SV=1 |
| 56 | 3914 | tr\|B2RYU0 | 78.75 | 26 | 26 | 6.41E+06 | 5 | 4 | 11 | | 11842 | NADH dehydrogenase (Ubiquinone) 1 beta subcomplex 2 (Predicted) isoform CRA_b OS=Rattus norvegicus OX=10116 GN=Ndufb2 PE=1 SV=1 |
| 69 | 14 | Q6AXV4\|SA | 77.24 | 6 | 6 | 3.85E+05 | 3 | 3 | 7 | | 51960 | Sorting and assembly machinery component 50 homolog OS=Rattus norvegicus OX=10116 GN=Samm50 PE=1 SV=1 |
| 70 | 335 | Q5XIH7\|PH | 74.3 | 10 | 10 | 1.76E+06 | 4 | 4 | 6 | | 33312 | Prohibitin-2 OS=Rattus norvegicus OX=10116 GN=Phb2 PE=1 SV=1 |
| 38 | 3918 | tr\|D3ZLT1\| | 74.26 | 16 | 16 | 6.73E+06 | 3 | 3 | 20 | Oxidation ( | 16568 | NADH dehydrogenase (Ubiquinone) 1 beta subcomplex 7 (Predicted) OS=Rattus norvegicus OX=10116 GN=Ndufb7 PE=1 SV=1 |
| 79 | 44 | P17764\|TH | 69.95 | 5 | 5 | 1.29E+06 | 2 | 2 | 5 | | 44695 | Acetyl-CoA acetyltransferase mitochondrial OS=Rattus norvegicus OX=10116 GN=Acat1 PE=1 SV=1 |
| 52 | 3906 | tr\|D3ZCZ9\| | 67.58 | 16 | 16 | 3.71E+06 | 3 | 3 | 12 | | 13040 | NADH dehydrogenase [ubiquinone] iron-sulfur protein 6 mitochondrial OS=Rattus norvegicus OX=10116 GN=LOC100912599 PE=1 SV=1 |
| 110 | 245 | tr\|C8CHS6 | 64.42 | 15 | 15 | 0 | 1 | 1 | 2 | | 7988 | Mitochondrial superoxide dismutase 2 (Fragment) OS=Rattus norvegicus OX=10116 PE=2 SV=1 |
| 110 | 71 | P07895\|SC | 64.42 | 5 | 5 | 0 | 1 | 1 | 2 | | 24674 | Superoxide dismutase [Mn] mitochondrial OS=Rattus norvegicus OX=10116 GN=Sod2 PE=1 SV=2 |
| 60 | 21 | Q64428\|E( | 62.55 | 6 | 6 | 6.21E+05 | 5 | 5 | 7 | | 82665 | Trifunctional enzyme subunit alpha mitochondrial OS=Rattus norvegicus OX=10116 GN=Hadha PE=1 SV=2 |
| 89 | 12037 | P21571\|AT | 60.92 | 24 | 24 | 6.34E+05 | 3 | 3 | 3 | Oxidation ( | 12494 | ATP synthase-coupling factor 6 mitochondrial OS=Rattus norvegicus OX=10116 GN=Atp5pf PE=1 SV=1 |
| 81 | 1021 | tr\|Q06QA9 | 60.72 | 3 | 3 | 6.10E+05 | 2 | 2 | 5 | | 56893 | Cytochrome c oxidase subunit 1 OS=Rattus norvegicus OX=10116 GN=CO1 PE=3 SV=1 |
| 81 | 1022 | tr\|Q06QK0 | 60.72 | 3 | 3 | 6.10E+05 | 2 | 2 | 5 | | 56880 | Cytochrome c oxidase subunit 1 OS=Rattus norvegicus OX=10116 GN=CO1 PE=3 SV=1 |
| 81 | 1023 | tr\|Q8SEZ6\| | 60.72 | 3 | 3 | 6.10E+05 | 2 | 2 | 5 | | 56879 | Cytochrome c oxidase subunit 1 OS=Rattus norvegicus OX=10116 GN=COI PE=3 SV=1 |
| 81 | 1024 | tr\|A0A0A1 | 60.72 | 3 | 3 | 6.10E+05 | 2 | 2 | 5 | | 56937 | Cytochrome c oxidase subunit 1 OS=Rattus norvegicus OX=10116 GN=COX1 PE=3 SV=1 |
| 81 | 1025 | tr\|Q8HIC9\| | 60.72 | 3 | 3 | 6.10E+05 | 2 | 2 | 5 | | 56845 | Cytochrome c oxidase subunit 1 OS=Rattus norvegicus OX=10116 GN=Mt-co1 PE=3 SV=1 |
| 81 | 1026 | P05503\|CC | 60.72 | 3 | 3 | 6.10E+05 | 2 | 2 | 5 | | 56845 | Cytochrome c oxidase subunit 1 OS=Rattus norvegicus OX=10116 GN=Mtco1 PE=2 SV=3 |
| 81 | 1027 | tr\|Q95938 | 60.72 | 3 | 3 | 6.10E+05 | 2 | 2 | 5 | | 56977 | Cytochrome c oxidase subunit 1 OS=Rattus norvegicus OX=10116 GN=Co I PE=3 SV=1 |
| 51 | 384 | tr\|Q06Q97 | 55.93 | 6 | 6 | 1.52E+06 | 3 | 3 | 12 | | 38485 | NADH-ubiquinone oxidoreductase chain 2 OS=Rattus norvegicus OX=10116 GN=ND2 PE=3 SV=1 |
| 51 | 385 | tr\|A0A0A1 | 55.93 | 6 | 6 | 1.52E+06 | 3 | 3 | 12 | | 38542 | NADH-ubiquinone oxidoreductase chain 2 OS=Rattus norvegicus OX=10116 GN=ND2 PE=3 SV=1 |
| 51 | 386 | tr\|Q5UAJ8 | 55.93 | 6 | 6 | 1.52E+06 | 3 | 3 | 12 | | 38455 | NADH-ubiquinone oxidoreductase chain 2 OS=Rattus norvegicus OX=10116 GN=ND2 PE=3 SV=1 |
| 51 | 388 | tr\|Q8HID0 | 55.93 | 6 | 6 | 1.52E+06 | 3 | 3 | 12 | | 38653 | NADH-ubiquinone oxidoreductase chain 2 OS=Rattus norvegicus OX=10116 GN=Mt-nd2 PE=3 SV=1 |
| 51 | 389 | P11662\|NU | 55.93 | 6 | 6 | 1.52E+06 | 3 | 3 | 12 | | 38653 | NADH-ubiquinone oxidoreductase chain 2 OS=Rattus norvegicus OX=10116 GN=Mtnd2 PE=3 SV=3 |
| 51 | 390 | tr\|Q06QH5 | 55.93 | 6 | 6 | 1.52E+06 | 3 | 3 | 12 | | 38626 | NADH-ubiquinone oxidoreductase chain 2 OS=Rattus norvegicus OX=10116 GN=ND2 PE=3 SV=1 |
| 51 | 391 | tr\|Q8SEZ7\| | 55.93 | 6 | 6 | 1.52E+06 | 3 | 3 | 12 | | 38580 | NADH-ubiquinone oxidoreductase chain 2 (Fragment) OS=Rattus norvegicus OX=10116 GN=NADH2 PE=3 SV=1 |
| 51 | 392 | tr\|Q06QB0 | 55.93 | 6 | 6 | 1.52E+06 | 3 | 3 | 12 | | 38534 | NADH-ubiquinone oxidoreductase chain 2 OS=Rattus norvegicus OX=10116 GN=ND2 PE=3 SV=1 |
| 51 | 393 | tr\|D2E6P4 | 55.93 | 6 | 6 | 1.52E+06 | 3 | 3 | 12 | | 38623 | NADH-ubiquinone oxidoreductase chain 2 OS=Rattus norvegicus OX=10116 GN=ND2 PE=3 SV=1 |
| 51 | 394 | tr\|Q06QE9 | 55.93 | 6 | 6 | 1.52E+06 | 3 | 3 | 12 | | 38598 | NADH-ubiquinone oxidoreductase chain 2 OS=Rattus norvegicus OX=10116 GN=ND2 PE=3 SV=1 |
| 90 | 266 | tr\|Q7H115 | 55.61 | 3 | 3 | 0 | 2 | 2 | 3 | | 29871 | Cytochrome c oxidase subunit 3 OS=Rattus norvegicus OX=10116 GN=Mt-co3 PE=3 SV=1 |
| 90 | 267 | tr\|I6V4L9\| | 55.61 | 3 | 3 | 0 | 2 | 2 | 3 | | 29844 | Cytochrome c oxidase subunit 3 OS=Rattus norvegicus OX=10116 GN=COX3 PE=3 SV=1 |
| 90 | 268 | tr\|Q8M7G4 | 55.61 | 3 | 3 | 0 | 2 | 2 | 3 | | 29861 | Cytochrome c oxidase subunit 3 OS=Rattus norvegicus OX=10116 PE=2 SV=1 |
| 90 | 269 | tr\|A0A096 | 55.61 | 3 | 3 | 0 | 2 | 2 | 3 | | 29901 | Cytochrome c oxidase subunit 3 OS=Rattus norvegicus OX=10116 GN=COX3 PE=3 SV=1 |
| 90 | 270 | P05505\|CC | 55.61 | 3 | 3 | 0 | 2 | 2 | 3 | | 29871 | Cytochrome c oxidase subunit 3 OS=Rattus norvegicus OX=10116 GN=Mtco3 PE=1 SV=5 |
| 90 | 271 | tr\|Q8SEZ2\| | 55.61 | 3 | 3 | 0 | 2 | 2 | 3 | | 29870 | Cytochrome c oxidase subunit 3 (Fragment) OS=Rattus norvegicus OX=10116 GN=COIII PE=3 SV=1 |
| 57 | 326 | Q91XJ1\|BE | 54.23 | 3 | 3 | | 3 | 0 | 11 | | 51557 | Beclin-1 OS=Rattus norvegicus OX=10116 GN=Becn1 PE=1 SV=1 |
| 72 | 8248 | B0BN56\|RT | 53.37 | 6 | 6 | 2.44E+05 | 3 | 1 | 6 | | 43962 | 28S ribosomal protein S31 mitochondrial OS=Rattus norvegicus OX=10116 GN=Mrps31 PE=2 SV=1 |
| 86 | 292 | P10888\|CC | 52.97 | 12 | 12 | 8.27E+05 | 3 | 3 | 4 | | 19515 | Cytochrome c oxidase subunit 4 isoform 1 mitochondrial OS=Rattus norvegicus OX=10116 GN=Cox4i1 PE=1 SV=1 |
| 76 | 3911 | tr\|B2RYW3 | 51.96 | 18 | 18 | 1.16E+06 | 4 | 4 | 5 | Oxidation ( | 21892 | NADH dehydrogenase (Ubiquinone) 1 beta subcomplex 9 OS=Rattus norvegicus OX=10116 GN=Ndufb9 PE=1 SV=1 |
| 37 | 256 | tr\|Q8M7G5 | 51.09 | 12 | 12 | 4.26E+06 | 3 | 3 | 21 | Oxidation ( | 24240 | ATP synthase subunit a (Fragment) OS=Rattus norvegicus OX=10116 PE=2 SV=1 |
| 37 | 257 | tr\|Q549I0\| | 51.09 | 12 | 12 | 4.26E+06 | 3 | 3 | 21 | Oxidation ( | 25050 | ATP synthase subunit a OS=Rattus norvegicus OX=10116 GN=atp6 PE=4 SV=1 |
| 37 | 258 | P05504\|AT | 51.09 | 12 | 12 | 4.26E+06 | 3 | 3 | 21 | Oxidation ( | 25076 | ATP synthase subunit a OS=Rattus norvegicus OX=10116 GN=Mt-atp6 PE=1 SV=3 |

| | | | | | | | | | | | | |
|---|---|---|---|---|---|---|---|---|---|---|---|---|
| 37 | 259 | tr\|Q8HIC7\| | 51.09 | 12 | 12 | 4.26E+06 | 3 | 3 | 21 | Oxidation ( | 25076 | ATP synthase subunit a OS=Rattus norvegicus OX=10116 GN=Mt-atp6 PE=4 SV=1 |
| 37 | 260 | tr\|S5S1E9\| | 51.09 | 12 | 12 | 4.26E+06 | 3 | 3 | 21 | Oxidation ( | 25049 | ATP synthase subunit a OS=Rattus norvegicus OX=10116 GN=ATP6 PE=4 SV=1 |
| 37 | 262 | tr\|Q8SEZ3\| | 51.09 | 12 | 12 | 4.26E+06 | 3 | 3 | 21 | Oxidation ( | 25030 | ATP synthase subunit a OS=Rattus norvegicus OX=10116 GN=ATPase6 PE=4 SV=1 |
| 68 | 467 | tr\|D4A4P3 | 49.48 | 30 | 30 | 1.08E+06 | 4 | 4 | 7 | Oxidation ( | 11267 | NADH:ubiquinone oxidoreductase subunit B3 OS=Rattus norvegicus OX=10116 GN=Ndufb3 PE=1 SV=1 |
| 59 | 3901 | tr\|Q8SEZ8\| | 47.26 | 5 | 5 | 2.62E+06 | 3 | 3 | 10 | | 36133 | NADH-ubiquinone oxidoreductase chain 1 (Fragment) OS=Rattus norvegicus OX=10116 GN=NADH1 PE=3 SV=1 |
| 59 | 3902 | P03889\|NU | 47.26 | 5 | 5 | 2.62E+06 | 3 | 3 | 10 | | 36145 | NADH-ubiquinone oxidoreductase chain 1 OS=Rattus norvegicus OX=10116 GN=Mtnd1 PE=1 SV=3 |
| 59 | 3903 | tr\|Q8HID1 | 47.26 | 5 | 5 | 2.62E+06 | 3 | 3 | 10 | | 36145 | NADH-ubiquinone oxidoreductase chain 1 OS=Rattus norvegicus OX=10116 GN=Mt-nd1 PE=3 SV=1 |
| 59 | 3904 | tr\|D2E6L7\| | 47.26 | 5 | 5 | 2.62E+06 | 3 | 3 | 10 | | 36062 | NADH-ubiquinone oxidoreductase chain 1 OS=Rattus norvegicus OX=10116 GN=ND1 PE=3 SV=1 |
| 65 | 420 | tr\|D3ZHK4 | 46.36 | 1 | 1 | | 3 | 0 | 7 | | 182226 | RB1-inducible coiled-coil 1 OS=Rattus norvegicus OX=10116 GN=Rb1cc1 PE=1 SV=1 |
| 66 | 208 | Q7TQ16\|Q | 45.75 | 11 | 11 | 7.27E+05 | 2 | 2 | 7 | | 9849 | Cytochrome b-c1 complex subunit 8 OS=Rattus norvegicus OX=10116 GN=Uqcrq PE=3 SV=1 |
| 73 | 209 | tr\|B2RZD6 | 44.09 | 22 | 22 | 1.09E+06 | 4 | 4 | 6 | | 9327 | NDUFA4 mitochondrial complex-associated OS=Rattus norvegicus OX=10116 GN=Ndufa4 PE=1 SV=1 |
| 80 | 307 | P11497\|AC | 43.76 | 1 | 1 | 0 | 3 | 2 | 5 | | 265191 | Acetyl-CoA carboxylase 1 OS=Rattus norvegicus OX=10116 GN=Acaca PE=1 SV=1 |
| 74 | 300 | tr\|A0A0G2 | 43.67 | 2 | 2 | 2.24E+05 | 2 | 1 | 6 | | 107334 | NLR family member X1 OS=Rattus norvegicus OX=10116 GN=Nlrx1 PE=1 SV=1 |
| 74 | 301 | Q5FVQ8\|N | 43.67 | 2 | 2 | 2.24E+05 | 2 | 1 | 6 | | 107590 | NLR family member X1 OS=Rattus norvegicus OX=10116 GN=Nlrx1 PE=2 SV=1 |
| 105 | 8042 | tr\|Q4G067 | 39.5 | 4 | 4 | 3.61E+05 | 2 | 2 | 2 | | 37440 | Mitochondrial ribosomal protein L44 OS=Rattus norvegicus OX=10116 GN=Mrpl44 PE=1 SV=1 |
| 82 | 151 | P12075\|CC | 38.68 | 11 | 11 | 8.75E+06 | 3 | 3 | 4 | | 13915 | Cytochrome c oxidase subunit 5B mitochondrial OS=Rattus norvegicus OX=10116 GN=Cox5b PE=1 SV=2 |
| 62 | 363 | P05508\|NU | 38.13 | 5 | 5 | 6.73E+05 | 3 | 3 | 7 | | 51783 | NADH-ubiquinone oxidoreductase chain 4 OS=Rattus norvegicus OX=10116 GN=Mtnd4 PE=3 SV=3 |
| 62 | 364 | tr\|D2E6K0 | 38.13 | 5 | 5 | 6.73E+05 | 3 | 3 | 7 | | 51833 | NADH-ubiquinone oxidoreductase chain 4 OS=Rattus norvegicus OX=10116 GN=ND4 PE=3 SV=1 |
| 62 | 365 | tr\|A7XYB9 | 38.13 | 5 | 5 | 6.73E+05 | 3 | 3 | 7 | | 51773 | NADH-ubiquinone oxidoreductase chain 4 OS=Rattus norvegicus OX=10116 GN=Nd4 PE=3 SV=1 |
| 62 | 366 | tr\|Q06QE1 | 38.13 | 5 | 5 | 6.73E+05 | 3 | 3 | 7 | | 51851 | NADH-ubiquinone oxidoreductase chain 4 OS=Rattus norvegicus OX=10116 GN=ND4 PE=3 SV=1 |
| 62 | 367 | tr\|Q06QA2 | 38.13 | 5 | 5 | 6.73E+05 | 3 | 3 | 7 | | 51787 | NADH-ubiquinone oxidoreductase chain 4 OS=Rattus norvegicus OX=10116 GN=ND4 PE=3 SV=1 |
| 62 | 368 | tr\|Q7HKW | 38.13 | 5 | 5 | 6.73E+05 | 3 | 3 | 7 | | 51801 | NADH-ubiquinone oxidoreductase chain 4 (Fragment) OS=Rattus norvegicus OX=10116 GN=NADH4 PE=3 SV=1 |
| 62 | 369 | tr\|Q06QG7 | 38.13 | 5 | 5 | 6.73E+05 | 3 | 3 | 7 | | 51791 | NADH-ubiquinone oxidoreductase chain 4 OS=Rattus norvegicus OX=10116 GN=ND4 PE=3 SV=1 |
| 62 | 370 | tr\|Q06Q89 | 38.13 | 5 | 5 | 6.73E+05 | 3 | 3 | 7 | | 51819 | NADH-ubiquinone oxidoreductase chain 4 OS=Rattus norvegicus OX=10116 GN=ND4 PE=3 SV=1 |
| 62 | 371 | tr\|Q35737 | 38.13 | 5 | 5 | 6.73E+05 | 3 | 3 | 7 | | 51765 | NADH-ubiquinone oxidoreductase chain 4 OS=Rattus norvegicus OX=10116 PE=3 SV=1 |
| 62 | 372 | tr\|Q8HIC6\| | 38.13 | 5 | 5 | 6.73E+05 | 3 | 3 | 7 | | 51783 | NADH-ubiquinone oxidoreductase chain 4 OS=Rattus norvegicus OX=10116 GN=Mt-nd4 PE=3 SV=1 |
| 149 | 31 | P45953\|AC | 35.82 | 2 | 2 | 2.94E+05 | 1 | 1 | 1 | | 70749 | Very long-chain specific acyl-CoA dehydrogenase mitochondrial OS=Rattus norvegicus OX=10116 GN=Acadvl PE=1 SV=1 |
| 149 | 32 | tr\|Q5M9H2 | 35.82 | 2 | 2 | 2.94E+05 | 1 | 1 | 1 | | 70821 | Acyl-Coenzyme A dehydrogenase very long chain OS=Rattus norvegicus OX=10116 GN=Acadvl PE=1 SV=1 |
| 112 | 4487 | tr\|D4A4B1 | 35.63 | 8 | 8 | 1.39E+05 | 2 | 2 | 2 | | 23223 | Mitochondrial ribosomal protein L15 OS=Rattus norvegicus OX=10116 GN=Mrpl15 PE=1 SV=2 |
| 112 | 4488 | tr\|A0A0G2 | 35.63 | 5 | 5 | 1.39E+05 | 2 | 2 | 2 | | 33652 | Mitochondrial ribosomal protein L15 OS=Rattus norvegicus OX=10116 GN=Mrpl15 PE=1 SV=1 |
| 109 | 12038 | Q9JJW3\|AT | 34.48 | 22 | 22 | 1.62E+06 | 2 | 2 | 2 | | 6408 | ATP synthase membrane subunit DAPIT mitochondrial OS=Rattus norvegicus OX=10116 GN=Atp5md PE=1 SV=1 |
| 78 | 12050 | D3ZAF6\|AT | 34.41 | 10 | 10 | 1.88E+06 | 3 | 3 | 5 | | 10452 | ATP synthase subunit f mitochondrial OS=Rattus norvegicus OX=10116 GN=Atp5mf PE=1 SV=1 |
| 101 | 343 | Q924S5\|LC | 34.12 | 1 | 1 | 5.92E+04 | 1 | 1 | 2 | | 105792 | Lon protease homolog mitochondrial OS=Rattus norvegicus OX=10116 GN=Lonp1 PE=2 SV=1 |
| 84 | 572 | F8WLE0\|KI | 30.44 | 2 | 2 | 6.62E+06 | 2 | 1 | 4 | | 115542 | Kinesin-like protein KIF28P OS=Rattus norvegicus OX=10116 GN=Kif28p PE=2 SV=1 |
| 111 | 315 | tr\|G3V7I0\| | 28.27 | 5 | 5 | 3.56E+05 | 1 | 1 | 2 | | 28299 | Peroxiredoxin 3 OS=Rattus norvegicus OX=10116 GN=Prdx3 PE=1 SV=1 |
| 111 | 316 | Q9Z0V6\|PF | 28.27 | 5 | 5 | 3.56E+05 | 1 | 1 | 2 | | 28295 | Thioredoxin-dependent peroxide reductase mitochondrial OS=Rattus norvegicus OX=10116 GN=Prdx3 PE=1 SV=2 |
| 92 | 224 | P11530\|DN | 27.58 | 1 | 1 | 2.25E+06 | 2 | 2 | 2 | Oxidation (M) | | |
| 98 | 3893 | tr\|B2RYS2\| | 26.54 | 16 | 16 | 4.81E+05 | 2 | 2 | 2 | | 13559 | Cytochrome b-c1 complex subunit 7 OS=Rattus norvegicus OX=10116 GN=Uqcrb PE=1 SV=1 |
| 104 | 207 | P11951\|CX | 26.38 | 17 | 17 | 4.28E+05 | 2 | 2 | 2 | | 8455 | Cytochrome c oxidase subunit 6C-2 OS=Rattus norvegicus OX=10116 GN=Cox6c2 PE=1 SV=3 |
| 100 | 344 | tr\|D4A4K4 | 26.29 | 0 | 0 | 3.41E+05 | 2 | 2 | 2 | Formylatio | 418626 | Vacuolar protein sorting 13 homolog C OS=Rattus norvegicus OX=10116 GN=Vps13c PE=1 SV=2 |
| 148 | 4381 | tr\|A0A0G2 | 24.61 | 1 | 1 | 0 | 1 | 1 | 1 | | 63436 | Malic enzyme OS=Rattus norvegicus OX=10116 GN=Me3 PE=3 SV=1 |
| 148 | 4382 | tr\|F1M5N4 | 24.61 | 1 | 1 | 0 | 1 | 1 | 1 | | 67209 | Malic enzyme OS=Rattus norvegicus OX=10116 GN=Me3 PE=3 SV=2 |
| 148 | 1308 | P13697\|MA | 24.61 | 1 | 1 | 0 | 1 | 1 | 1 | | 64003 | NADP-dependent malic enzyme OS=Rattus norvegicus OX=10116 GN=Me1 PE=1 SV=2 |
| 148 | 4463 | tr\|D3ZJH9\| | 24.61 | 1 | 1 | 0 | 1 | 1 | 1 | | 65352 | Malic enzyme OS=Rattus norvegicus OX=10116 GN=Me2 PE=1 SV=1 |
| 148 | 4464 | tr\|A0A0G2 | 24.61 | 1 | 1 | 0 | 1 | 1 | 1 | | 66310 | Malic enzyme OS=Rattus norvegicus OX=10116 GN=Me2 PE=1 SV=1 |
| 113 | 12041 | tr\|D3Z8Q5 | 24.29 | 6 | 6 | 2.94E+05 | 1 | 1 | 2 | | 12124 | RCG25747 OS=Rattus norvegicus OX=10116 GN=Tusc2 PE=4 SV=1 |
| 166 | 8178 | tr\|D4A9Z6 | 23.58 | 2 | 2 | 5.25E+05 | 1 | 1 | 1 | | 36207 | Mitochondrial ribosomal protein S35 OS=Rattus norvegicus OX=10116 GN=Mrps35 PE=1 SV=1 |
| 63 | 1269 | Q5M934\|T | 23.08 | 2 | 2 | 1.71E+07 | 1 | 1 | 7 | | 36404 | Probable tRNA pseudouridine synthase 1 OS=Rattus norvegicus OX=10116 GN=Trub1 PE=2 SV=1 |
| 115 | 12048 | tr\|B2RYK8\| | 22.85 | 1 | 1 | 5.06E+06 | 1 | 1 | 2 | | 58738 | RCG28722 isoform CRA_a OS=Rattus norvegicus OX=10116 GN=Slc9b2 PE=2 SV=1 |
| 151 | 4466 | Q58FK9\|KA | 21.61 | 2 | 2 | 1.54E+06 | 1 | 1 | 1 | | 51044 | Kynurenine--oxoglutarate transaminase 3 OS=Rattus norvegicus OX=10116 GN=Kyat3 PE=2 SV=1 |
| 167 | 66 | P56574\|ID | 21.49 | 2 | 2 | 8.91E+04 | 1 | 1 | 1 | | 50967 | Isocitrate dehydrogenase [NADP] mitochondrial OS=Rattus norvegicus OX=10116 GN=Idh2 PE=1 SV=2 |
| 142 | 3917 | tr\|D3ZJG4\| | 21.33 | 1 | 1 | 1.74E+06 | 1 | 1 | 1 | Oxidation ( | 95972 | Phosphofurin acidic cluster sorting protein 2 OS=Rattus norvegicus OX=10116 GN=Pacs2 PE=1 SV=2 |
| 168 | 3923 | P80431\|CC | 21.01 | 14 | 14 | 1.17E+05 | 1 | 1 | 1 | | 8995 | Cytochrome c oxidase subunit 7B mitochondrial OS=Rattus norvegicus OX=10116 GN=Cox7b PE=1 SV=3 |
| 168 | 4304 | tr\|A0A0G2 | 21.01 | 14 | 14 | 1.17E+05 | 1 | 1 | 1 | | 9070 | Cytochrome c oxidase subunit 7B2 OS=Rattus norvegicus OX=10116 GN=Cox7b2 PE=4 SV=1 |
| 141 | 11 | tr\|A0A140 | 20.78 | 1 | 1 | 1.05E+05 | 1 | 1 | 1 | | 67049 | MICOS complex subunit MIC60 OS=Rattus norvegicus OX=10116 GN=Immt PE=1 SV=1 |
| 141 | 12 | Q3KR86\|M | 20.78 | 1 | 1 | 1.05E+05 | 1 | 1 | 1 | | 67177 | MICOS complex subunit Mic60 (Fragment) OS=Rattus norvegicus OX=10116 GN=Immt PE=1 SV=1 |
| 141 | 10 | tr\|A0A0G2 | 20.78 | 1 | 1 | 1.05E+05 | 1 | 1 | 1 | | 86230 | MICOS complex subunit MIC60 OS=Rattus norvegicus OX=10116 GN=Immt PE=1 SV=1 |
| 133 | 310 | Q75Q40\|T | 20.09 | 4 | 4 | 1.55E+05 | 1 | 1 | 1 | | 37919 | Mitochondrial import receptor subunit TOM40 homolog OS=Rattus norvegicus OX=10116 GN=Tomm40 PE=1 SV=1 |

| | | | | | | | | | | | | |
|---|---|---|---|---|---|---|---|---|---|---|---|---|
| 133 | 311 | tr\|G3V8F5 | 20.09 | 4 | 4 | 1.55E+05 | 1 | 1 | 1 | | 37920 | Mitochondrial import receptor subunit TOM40 homolog OS=Rattus norvegicus OX=10116 GN=Tomm40 PE=1 SV=1 |
| 128 | 289 | Q4KM98\|N | 20.05 | 3 | 3 | 0 | 1 | 1 | 1 | Formylatio | 24971 | Mitochondrial fission factor OS=Rattus norvegicus OX=10116 GN=Mff PE=1 SV=1 |
| 128 | 239 | tr\|A0A0H2 | 20.05 | 2 | 2 | 0 | 1 | 1 | 1 | Formylatio | 27523 | Mitochondrial fission factor-like OS=Rattus norvegicus OX=10116 GN=LOC102556337 PE=1 SV=1 |
| 128 | 290 | tr\|A0A0G2 | 20.05 | 2 | 2 | 0 | 1 | 1 | 1 | Formylatio | 32801 | Mitochondrial fission factor OS=Rattus norvegicus OX=10116 GN=Mff PE=1 SV=1 |
| 128 | 240 | tr\|A0A0G2 | 20.05 | 2 | 2 | 0 | 1 | 1 | 1 | Formylatio | 39707 | Mitochondrial fission factor-like OS=Rattus norvegicus OX=10116 GN=LOC102556337 PE=1 SV=1 |
