## Supplementary material for "Permeability transition pore-related changes in the proteome and channel activity of ATP synthase dimers and monomers": RHM PTP Monomer of V compl.

| Protein Grc | Protein ID | Accession | -10lgP | Coverage (' | Coverage (' | Area Samp | #Peptides | #Unique | #Spec Sam | PTM | Avg. Mass | Description |
|---|---|---|---|---|---|---|---|---|---|---|---|---|
| 1 | 5 | P10719\|AT | 412.63 | 52 | 52 | 2.70E+09 | 151 | 148 | 1008 | Oxidation ( | 56354 | ATP synthase subunit beta mitochondrial OS=Rattus norvegicus OX=10116 GN=Atp5f1b PE=1 SV=2 |
| 1 | 6 | tr\|G3V6D3 | 412.63 | 52 | 52 | 2.70E+09 | 151 | 148 | 1008 | Oxidation ( | 56345 | ATP synthase subunit beta OS=Rattus norvegicus OX=10116 GN=Atp5f1b PE=1 SV=1 |
| 2 | 8 | P15999\|AT | 351.84 | 44 | 44 | 1.48E+09 | 147 | 143 | 897 | Carbamidomethylation | | |
| 2 | 9 | tr\|F1LP05\| | 351.84 | 44 | 44 | 1.48E+09 | 147 | 143 | 897 | Carbamidomethylation | | |
| 3 | 65 | P31399\|AT | 317.64 | 66 | 66 | 2.03E+08 | 47 | 46 | 200 | Oxidation (M) | | |
| 4 | 63 | P35435\|AT | 297.5 | 48 | 48 | 1.50E+08 | 47 | 47 | 172 | Carbamidomethylation | | |
| 4 | 64 | tr\|Q6QI09\| | 297.5 | 22 | 22 | 1.50E+08 | 47 | 47 | 172 | Carbamidomethylation | | |
| 5 | 168 | P19511\|AT | 208.61 | 32 | 32 | 2.53E+08 | 32 | 31 | 128 | Formylatio | 28869 | ATP synthase F(0) complex subunit B1 mitochondrial OS=Rattus norvegicus OX=10116 GN=Atp5pb PE=1 SV=1 |
| 6 | 76 | Q06647\|AT | 199.86 | 42 | 42 | 1.11E+08 | 34 | 33 | 116 | Carbamidomethylation | | |
| 7 | 75 | Q6PDU7\|A | 186.13 | 50 | 50 | 7.71E+07 | 22 | 22 | 103 | Oxidation ( | 11433 | ATP synthase subunit g mitochondrial OS=Rattus norvegicus OX=10116 GN=Atp5mg PE=1 SV=2 |
| 9 | 45 | P00507\|AA | 175.16 | 17 | 17 | 7.50E+06 | 13 | 13 | 32 | Oxidation ( | 47314 | Aspartate aminotransferase mitochondrial OS=Rattus norvegicus OX=10116 GN=Got2 PE=1 SV=2 |
| 11 | 12037 | P21571\|AT | 170.28 | 49 | 49 | 7.65E+06 | 13 | 12 | 28 | Oxidation ( | 12494 | ATP synthase-coupling factor 6 mitochondrial OS=Rattus norvegicus OX=10116 GN=Atp5pf PE=1 SV=1 |
| 8 | 319 | P29419\|AT | 157.03 | 65 | 65 | 3.22E+07 | 17 | 17 | 54 | Oxidation ( | 8255 | ATP synthase subunit e mitochondrial OS=Rattus norvegicus OX=10116 GN=Atp5me PE=1 SV=3 |
| 17 | 59 | tr\|D3ZFQ8 | 142.67 | 13 | 13 | 2.83E+06 | 5 | 5 | 12 | | 35435 | Cytochrome c-1 OS=Rattus norvegicus OX=10116 GN=Cyc1 PE=1 SV=3 |
| 13 | 29 | Q60587\|EC | 131.94 | 14 | 14 | 2.83E+06 | 12 | 12 | 20 | | 51414 | Trifunctional enzyme subunit beta mitochondrial OS=Rattus norvegicus OX=10116 GN=Hadhb PE=1 SV=1 |
| 29 | 58 | P20788\|UC | 126.55 | 5 | 5 | 1.30E+06 | 2 | 2 | 4 | | 29446 | Cytochrome b-c1 complex subunit Rieske mitochondrial OS=Rattus norvegicus OX=10116 GN=Uqcrfs1 PE=1 SV=2 |
| 21 | 200 | P11240\|CC | 118.98 | 23 | 23 | 9.24E+05 | 6 | 6 | 10 | | 16130 | Cytochrome c oxidase subunit 5A mitochondrial OS=Rattus norvegicus OX=10116 GN=Cox5a PE=1 SV=1 |
| 16 | 233 | tr\|Q5UAJ5 | 111.96 | 28 | 28 | 1.70E+07 | 5 | 5 | 14 | Oxidation ( | 7642 | ATP synthase protein 8 OS=Rattus norvegicus OX=10116 GN=ATP8 PE=3 SV=1 |
| 16 | 235 | tr\|Q8SEZ4\| | 111.96 | 28 | 28 | 1.70E+07 | 5 | 5 | 14 | Oxidation ( | 7632 | ATP synthase protein 8 OS=Rattus norvegicus OX=10116 GN=ATPase8 PE=3 SV=1 |
| 12 | 3953 | tr\|G3V7Y3 | 105.59 | 19 | 19 | 1.91E+07 | 9 | 9 | 25 | | 17563 | ATP synthase subunit delta mitochondrial OS=Rattus norvegicus OX=10116 GN=Atp5f1d PE=1 SV=1 |
| 18 | 21 | Q64428\|EC | 102.43 | 8 | 8 | 1.63E+06 | 7 | 7 | 11 | | 82665 | Trifunctional enzyme subunit alpha mitochondrial OS=Rattus norvegicus OX=10116 GN=Hadha PE=1 SV=2 |
| 14 | 33 | tr\|S5RZM8 | 99.64 | 27 | 27 | 3.16E+06 | 8 | 8 | 17 | Oxidation ( | 25958 | Cytochrome c oxidase subunit 2 OS=Rattus norvegicus OX=10116 GN=COX2 PE=3 SV=1 |
| 14 | 34 | P00406\|CC | 99.64 | 27 | 27 | 3.16E+06 | 8 | 8 | 17 | Oxidation ( | 25928 | Cytochrome c oxidase subunit 2 OS=Rattus norvegicus OX=10116 GN=Mtco2 PE=1 SV=3 |
| 14 | 28 | tr\|Q5UAJ6 | 99.64 | 27 | 27 | 3.16E+06 | 8 | 8 | 17 | Oxidation ( | 25942 | Cytochrome c oxidase subunit 2 OS=Rattus norvegicus OX=10116 GN=COX2 PE=3 SV=1 |
| 14 | 35 | tr\|Q8SEZ5\| | 99.64 | 27 | 27 | 3.16E+06 | 8 | 8 | 17 | Oxidation ( | 25928 | Cytochrome c oxidase subunit 2 OS=Rattus norvegicus OX=10116 GN=Mt-co2 PE=1 SV=1 |
| 14 | 36 | tr\|A0A097I | 99.64 | 27 | 27 | 3.16E+06 | 8 | 8 | 17 | Oxidation ( | 25894 | Cytochrome c oxidase subunit 2 OS=Rattus norvegicus OX=10116 GN=COX2 PE=3 SV=1 |
| 43 | 263 | P35171\|CX | 89.33 | 12 | 12 | 7.48E+05 | 2 | 2 | 3 | | 9353 | Cytochrome c oxidase subunit 7A2 mitochondrial OS=Rattus norvegicus OX=10116 GN=Cox7a2 PE=1 SV=1 |
| 43 | 264 | tr\|B2RYS0\| | 89.33 | 12 | 12 | 7.48E+05 | 2 | 2 | 3 | | 9353 | Cox7a2 protein OS=Rattus norvegicus OX=10116 GN=Cox7a2 PE=2 SV=1 |
| 26 | 17 | Q9ER34\|AC | 79.49 | 3 | 3 | 4.41E+05 | 3 | 3 | 5 | | 85433 | Aconitate hydratase mitochondrial OS=Rattus norvegicus OX=10116 GN=Aco2 PE=1 SV=2 |
| 52 | 251 | P80432\|CC | 78.41 | 21 | 21 | 8.95E+05 | 1 | 1 | 3 | | 7375 | Cytochrome c oxidase subunit 7C mitochondrial OS=Rattus norvegicus OX=10116 GN=Cox7c PE=1 SV=2 |
| 51 | 315 | tr\|G3V7I0\| | 77.98 | 8 | 8 | 9.74E+05 | 2 | 2 | 3 | | 28299 | Peroxiredoxin 3 OS=Rattus norvegicus OX=10116 GN=Prdx3 PE=1 SV=1 |
| 51 | 316 | Q9Z0V6\|PF | 77.98 | 8 | 8 | 9.74E+05 | 2 | 2 | 3 | | 28295 | Thioredoxin-dependent peroxide reductase mitochondrial OS=Rattus norvegicus OX=10116 GN=Prdx3 PE=1 SV=2 |
| 42 | 61 | Q63704\|CF | 77.4 | 2 | 2 | 5.81E+05 | 2 | 2 | 3 | | 88217 | Carnitine O-palmitoyltransferase 1 muscle isoform OS=Rattus norvegicus OX=10116 GN=Cpt1b PE=1 SV=1 |
| 20 | 50 | P32551\|QC | 77.02 | 5 | 5 | 2.76E+06 | 4 | 4 | 10 | | 48396 | Cytochrome b-c1 complex subunit 2 mitochondrial OS=Rattus norvegicus OX=10116 GN=Uqcrc2 PE=1 SV=2 |
| 10 | 256 | tr\|Q8M7G5 | 74.9 | 18 | 18 | 1.07E+07 | 7 | 7 | 28 | Oxidation ( | 24240 | ATP synthase subunit a (Fragment) OS=Rattus norvegicus OX=10116 PE=2 SV=1 |
| 10 | 257 | tr\|Q549I0\| | 74.9 | 18 | 18 | 1.07E+07 | 7 | 7 | 28 | Oxidation ( | 25050 | ATP synthase subunit a OS=Rattus norvegicus OX=10116 GN=atp6 PE=4 SV=1 |
| 10 | 258 | P05504\|AT | 74.9 | 18 | 18 | 1.07E+07 | 7 | 7 | 28 | Oxidation ( | 25076 | ATP synthase subunit a OS=Rattus norvegicus OX=10116 GN=Mt-atp6 PE=1 SV=3 |
| 10 | 259 | tr\|Q8HIC7\| | 74.9 | 18 | 18 | 1.07E+07 | 7 | 7 | 28 | Oxidation ( | 25076 | ATP synthase subunit a OS=Rattus norvegicus OX=10116 GN=Mt-atp6 PE=4 SV=1 |
| 10 | 260 | tr\|S5S1E9\| | 74.9 | 18 | 18 | 1.07E+07 | 7 | 7 | 28 | Oxidation ( | 25049 | ATP synthase subunit a OS=Rattus norvegicus OX=10116 GN=ATP6 PE=4 SV=1 |
| 10 | 262 | tr\|Q8SEZ3\| | 74.9 | 18 | 18 | 1.07E+07 | 7 | 7 | 28 | Oxidation ( | 25030 | ATP synthase subunit a OS=Rattus norvegicus OX=10116 GN=ATPase6 PE=4 SV=1 |
| 77 | 266 | tr\|Q7H115 | 67.76 | 3 | 3 | 4.14E+05 | 2 | 2 | 2 | | 29871 | Cytochrome c oxidase subunit 3 OS=Rattus norvegicus OX=10116 GN=Mt-co3 PE=3 SV=1 |
| 77 | 267 | tr\|I6V4L9\| | 67.76 | 3 | 3 | 4.14E+05 | 2 | 2 | 2 | | 29844 | Cytochrome c oxidase subunit 3 OS=Rattus norvegicus OX=10116 GN=COX3 PE=3 SV=1 |
| 77 | 268 | tr\|Q8M7G4 | 67.76 | 3 | 3 | 4.14E+05 | 2 | 2 | 2 | | 29861 | Cytochrome c oxidase subunit 3 OS=Rattus norvegicus OX=10116 PE=2 SV=1 |
| 77 | 269 | tr\|A0A096X | 67.76 | 3 | 3 | 4.14E+05 | 2 | 2 | 2 | | 29901 | Cytochrome c oxidase subunit 3 OS=Rattus norvegicus OX=10116 GN=COX3 PE=3 SV=1 |
| 77 | 270 | P05505\|CC | 67.76 | 3 | 3 | 4.14E+05 | 2 | 2 | 2 | | 29871 | Cytochrome c oxidase subunit 3 OS=Rattus norvegicus OX=10116 GN=Mtco3 PE=1 SV=5 |
| 77 | 271 | tr\|Q8SEZ2\| | 67.76 | 3 | 3 | 4.14E+05 | 2 | 2 | 2 | | 29870 | Cytochrome c oxidase subunit 3 (Fragment) OS=Rattus norvegicus OX=10116 GN=COIII PE=3 SV=1 |
| 22 | 12038 | Q9JJW3\|AT | 61.59 | 40 | 40 | 5.67E+06 | 4 | 4 | 7 | | 6408 | ATP synthase membrane subunit DAPIT mitochondrial OS=Rattus norvegicus OX=10116 GN=Atp5md PE=1 SV=1 |
| 15 | 326 | Q91XJ1\|BE | 57.26 | 3 | 3 | | 3 | 0 | 17 | | 51557 | Beclin-1 OS=Rattus norvegicus OX=10116 GN=Becn1 PE=1 SV=1 |
| 70 | 25 | Q68FY0\|QC | 53.21 | 4 | 4 | 8.95E+05 | 2 | 2 | 2 | | 52849 | Cytochrome b-c1 complex subunit 1 mitochondrial OS=Rattus norvegicus OX=10116 GN=Uqcrc1 PE=1 SV=1 |
| 72 | 31 | P45953\|AC | 51.63 | 3 | 3 | 3.55E+05 | 2 | 2 | 2 | | 70749 | Very long-chain specific acyl-CoA dehydrogenase mitochondrial OS=Rattus norvegicus OX=10116 GN=Acadvl PE=1 SV=1 |
| 72 | 32 | tr\|Q5M9H2 | 51.63 | 3 | 3 | 3.55E+05 | 2 | 2 | 2 | | 70821 | Acyl-Coenzyme A dehydrogenase very long chain OS=Rattus norvegicus OX=10116 GN=Acadvl PE=1 SV=1 |
| 19 | 12050 | D3ZAF6\|AT | 50.42 | 20 | 20 | 3.96E+06 | 6 | 6 | 10 | Oxidation ( | 10452 | ATP synthase subunit f mitochondrial OS=Rattus norvegicus OX=10116 GN=Atp5mf PE=1 SV=1 |
| 24 | 253 | P10817\|CX | 48.06 | 12 | 12 | 2.19E+06 | 2 | 2 | 6 | | 10487 | Cytochrome c oxidase subunit 6A2 mitochondrial (Fragment) OS=Rattus norvegicus OX=10116 GN=Cox6a2 PE=1 SV=3 |
| 24 | 254 | tr\|G3V8M4 | 48.06 | 11 | 11 | 2.19E+06 | 2 | 2 | 6 | | 10802 | Cytochrome c oxidase subunit 6A mitochondrial OS=Rattus norvegicus OX=10116 GN=Cox6a2 PE=3 SV=1 |
| 31 | 46 | P09605\|KC | 46.99 | 7 | 7 | 4.04E+05 | 3 | 3 | 4 | | 47385 | Creatine kinase S-type mitochondrial OS=Rattus norvegicus OX=10116 GN=Ckmt2 PE=1 SV=2 |
| 23 | 420 | tr\|D3ZHK4 | 43.42 | 1 | 1 | | 3 | 0 | 6 | | 182226 | RB1-inducible coiled-coil 1 OS=Rattus norvegicus OX=10116 GN=Rb1cc1 PE=1 SV=1 |
| 47 | 209 | tr\|B2RZD6 | 37.86 | 10 | 10 | 1.19E+05 | 2 | 2 | 3 | | 9327 | NDUFA4 mitochondrial complex-associated OS=Rattus norvegicus OX=10116 GN=Ndufa4 PE=1 SV=1 |

| | | | | | | | | | | | | |
|---|---|---|---|---|---|---|---|---|---|---|---|---|
| 27 | 12057 | tr\|Q64592 | 37.12 | 6 | 6 | 0 | 3 | 1 | 5 | | 32894 | Delta3 delta2-enoyl-CoA isomerase OS=Rattus norvegicus OX=10116 PE=2 SV=1 |
| 64 | 44 | P17764\|TH | 35.59 | 3 | 3 | 4.27E+05 | 1 | 1 | 2 | | 44695 | Acetyl-CoA acetyltransferase mitochondrial OS=Rattus norvegicus OX=10116 GN=Acat1 PE=1 SV=1 |
| 73 | 66 | P56574\|ID | 34.89 | 3 | 3 | 1.82E+05 | 1 | 1 | 2 | | 50967 | Isocitrate dehydrogenase [NADP] mitochondrial OS=Rattus norvegicus OX=10116 GN=Idh2 PE=1 SV=2 |
| 32 | 318 | Q5M9I5\|Q | 34.77 | 17 | 17 | 8.64E+05 | 1 | 1 | 4 | | 10424 | Cytochrome b-c1 complex subunit 6 mitochondrial OS=Rattus norvegicus OX=10116 GN=Uqcrh PE=3 SV=1 |
| 40 | 410 | tr\|F1LNF0\| | 33.06 | 0 | 0 | | 2 | 0 | 3 | | 228912 | Myosin heavy chain 14 OS=Rattus norvegicus OX=10116 GN=Myh14 PE=1 SV=1 |
| 71 | 144 | P13086\|SU | 32.88 | 6 | 6 | 1.24E+05 | 2 | 2 | 2 | | 36148 | Succinate--CoA ligase [ADP/GDP-forming] subunit alpha mitochondrial OS=Rattus norvegicus OX=10116 GN=Suclg1 PE=2 SV=2 |
| 71 | 145 | tr\|A0A0H2 | 32.88 | 6 | 6 | 1.24E+05 | 2 | 2 | 2 | | 37560 | Succinate--CoA ligase [ADP/GDP-forming] subunit alpha mitochondrial OS=Rattus norvegicus OX=10116 GN=Suclg1 PE=1 SV=1 |
| 33 | 8248 | B0BN56\|RT | 32.7 | 3 | 3 | | 2 | 0 | 4 | | 43962 | 28S ribosomal protein S31 mitochondrial OS=Rattus norvegicus OX=10116 GN=Mrps31 PE=2 SV=1 |
| 48 | 4381 | tr\|A0A0G2 | 31.76 | 1 | 1 | 5.56E+05 | 1 | 1 | 3 | | 63436 | Malic enzyme OS=Rattus norvegicus OX=10116 GN=Me3 PE=3 SV=1 |
| 48 | 4382 | tr\|F1M5N4 | 31.76 | 1 | 1 | 5.56E+05 | 1 | 1 | 3 | | 67209 | Malic enzyme OS=Rattus norvegicus OX=10116 GN=Me3 PE=3 SV=2 |
| 48 | 1308 | P13697\|MA | 31.76 | 1 | 1 | 5.56E+05 | 1 | 1 | 3 | | 64003 | NADP-dependent malic enzyme OS=Rattus norvegicus OX=10116 GN=Me1 PE=1 SV=2 |
| 48 | 4463 | tr\|D3ZJH9\| | 31.76 | 1 | 1 | 5.56E+05 | 1 | 1 | 3 | | 65352 | Malic enzyme OS=Rattus norvegicus OX=10116 GN=Me2 PE=1 SV=1 |
| 48 | 4464 | tr\|A0A0G2 | 31.76 | 1 | 1 | 5.56E+05 | 1 | 1 | 3 | | 66310 | Malic enzyme OS=Rattus norvegicus OX=10116 GN=Me2 PE=1 SV=1 |
| 146 | 292 | P10888\|CC | 31.61 | 4 | 4 | 1.61E+05 | 1 | 1 | 1 | | 19515 | Cytochrome c oxidase subunit 4 isoform 1 mitochondrial OS=Rattus norvegicus OX=10116 GN=Cox4i1 PE=1 SV=1 |
| 41 | 158 | Q9R1Z0\|VD | 31.48 | 8 | 8 | 1.64E+06 | 2 | 2 | 3 | Formylatio | 30798 | Voltage-dependent anion-selective channel protein 3 OS=Rattus norvegicus OX=10116 GN=Vdac3 PE=1 SV=2 |
| 118 | 403 | tr\|A0A0G2 | 30.67 | 1 | 1 | 1.49E+05 | 1 | 1 | 1 | | 79812 | Propionyl-CoA carboxylase alpha chain mitochondrial OS=Rattus norvegicus OX=10116 GN=Pcca PE=1 SV=1 |
| 118 | 405 | P14882\|PC | 30.67 | 1 | 1 | 1.49E+05 | 1 | 1 | 1 | | 81623 | Propionyl-CoA carboxylase alpha chain mitochondrial OS=Rattus norvegicus OX=10116 GN=Pcca PE=1 SV=3 |
| 118 | 404 | tr\|A0A0G2 | 30.67 | 1 | 1 | 1.49E+05 | 1 | 1 | 1 | | 81413 | Propionyl-CoA carboxylase alpha chain mitochondrial OS=Rattus norvegicus OX=10116 GN=Pcca PE=1 SV=1 |
| 38 | 151 | P12075\|CC | 30.38 | 11 | 11 | 1.47E+06 | 2 | 2 | 3 | | 13915 | Cytochrome c oxidase subunit 5B mitochondrial OS=Rattus norvegicus OX=10116 GN=Cox5b PE=1 SV=2 |
| 45 | 1231 | tr\|D4A1D3 | 30.28 | 0 | 0 | | 2 | 0 | 3 | | 521487 | Sacsin molecular chaperone OS=Rattus norvegicus OX=10116 GN=Sacs PE=1 SV=2 |
| 56 | 737 | tr\|A0A0G2 | 29.89 | 1 | 1 | 1.32E+05 | 2 | 2 | 2 | Carbamidomethylation | | |
| 56 | 738 | tr\|A0A0G2 | 29.89 | 1 | 1 | 1.32E+05 | 2 | 2 | 2 | Carbamidomethylation | | |
| 56 | 739 | tr\|D3ZBE2\| | 29.89 | 1 | 1 | 1.32E+05 | 2 | 2 | 2 | Carbamidomethylation | | |
| 56 | 740 | tr\|E9PSQ0 | 29.89 | 1 | 1 | 1.32E+05 | 2 | 2 | 2 | Carbamidomethylation | | |
| 56 | 426 | tr\|O70151 | 29.89 | 1 | 1 | 1.32E+05 | 2 | 2 | 2 | Carbamidomethylation | | |
| 65 | 8226 | tr\|D3Z899\| | 29.6 | 3 | 3 | 1.72E+05 | 2 | 1 | 2 | Formylatio | 61393 | Mitoguardin 2 OS=Rattus norvegicus OX=10116 GN=Miga2 PE=1 SV=1 |
| 65 | 8227 | tr\|A0A0G2 | 29.6 | 2 | 2 | 1.72E+05 | 2 | 1 | 2 | Formylatio | 65660 | Mitoguardin 2 OS=Rattus norvegicus OX=10116 GN=Miga2 PE=1 SV=1 |
| 55 | 647 | O55207\|SY | 29.35 | 1 | 1 | 1.43E+05 | 2 | 2 | 3 | | 165263 | Synaptojanin-2 OS=Rattus norvegicus OX=10116 GN=Synj2 PE=1 SV=2 |
| 62 | 310 | Q75Q40\|T | 27.14 | 4 | 4 | 2.54E+05 | 1 | 1 | 2 | | 37919 | Mitochondrial import receptor subunit TOM40 homolog OS=Rattus norvegicus OX=10116 GN=Tomm40 PE=1 SV=1 |
| 62 | 311 | tr\|G3V8F5 | 27.14 | 4 | 4 | 2.54E+05 | 1 | 1 | 2 | | 37920 | Mitochondrial import receptor subunit TOM40 homolog OS=Rattus norvegicus OX=10116 GN=Tomm40 PE=1 SV=1 |
| 148 | 321 | tr\|D4AE90 | 25.24 | 2 | 2 | 0 | 1 | 1 | 1 | Formylatio | 50113 | RCC1-like OS=Rattus norvegicus OX=10116 GN=Rcc1l PE=1 SV=1 |
| 79 | 12048 | tr\|B2RYK8\| | 24.95 | 1 | 1 | 4.55E+06 | 1 | 1 | 2 | | 58738 | RCG28722 isoform CRA_a OS=Rattus norvegicus OX=10116 GN=Slc9b2 PE=2 SV=1 |
| 96 | 8293 | P07633\|PC | 24.66 | 1 | 1 | 6.22E+04 | 1 | 1 | 1 | | 58626 | Propionyl-CoA carboxylase beta chain mitochondrial OS=Rattus norvegicus OX=10116 GN=Pccb PE=2 SV=1 |
| 96 | 8294 | tr\|Q68FZ8\| | 24.66 | 1 | 1 | 6.22E+04 | 1 | 1 | 1 | | 58678 | Propionyl coenzyme A carboxylase beta polypeptide OS=Rattus norvegicus OX=10116 GN=Pccb PE=1 SV=1 |
| 25 | 1269 | Q5M934\|T | 23.57 | 2 | 2 | 4.52E+07 | 1 | 1 | 5 | | 36404 | Probable tRNA pseudouridine synthase 1 OS=Rattus norvegicus OX=10116 GN=Trub1 PE=2 SV=1 |
| 122 | 350 | tr\|G3V761 | 23.42 | 4 | 4 | 0 | 1 | 1 | 1 | | 16714 | Interferon alpha-inducible protein 27 OS=Rattus norvegicus OX=10116 GN=Ifi27 PE=4 SV=1 |
| 122 | 351 | tr\|Q6P9X5 | 23.42 | 4 | 4 | 0 | 1 | 1 | 1 | | 16682 | Interferon alpha-inducible protein 27-like OS=Rattus norvegicus OX=10116 GN=Ifi27 PE=2 SV=1 |
| 120 | 298 | tr\|Q5RKL4 | 23.11 | 1 | 1 | 0 | 1 | 1 | 1 | | 95977 | Dimethylglycine dehydrogenase OS=Rattus norvegicus OX=10116 GN=Dmgdh PE=1 SV=1 |
| 120 | 299 | Q63342\|M | 23.11 | 1 | 1 | 0 | 1 | 1 | 1 | | 96047 | Dimethylglycine dehydrogenase mitochondrial OS=Rattus norvegicus OX=10116 GN=Dmgdh PE=1 SV=1 |
| 120 | 303 | tr\|A0A0G2 | 23.11 | 1 | 1 | 0 | 1 | 1 | 1 | | 98864 | Dimethylglycine dehydrogenase mitochondrial OS=Rattus norvegicus OX=10116 GN=Dmgdh PE=1 SV=1 |
| 54 | 12095 | D3Z9R8\|AT | 22.95 | 8 | 8 | 2.30E+06 | 1 | 1 | 3 | | 6914 | ATP synthase subunit ATP5MPL mitochondrial OS=Rattus norvegicus OX=10116 GN=Atp5mpl PE=1 SV=1 |
| 54 | 12112 | Q9WVK3\|P | 22.95 | 2 | 2 | 2.30E+06 | 1 | 1 | 3 | | 32433 | Peroxisomal trans-2-enoyl-CoA reductase OS=Rattus norvegicus OX=10116 GN=Pecr PE=2 SV=1 |
| 54 | 12113 | tr\|A0A0G2 | 22.95 | 2 | 2 | 2.30E+06 | 1 | 1 | 3 | | 32737 | Peroxisomal trans-2-enoyl-CoA reductase OS=Rattus norvegicus OX=10116 GN=Pecr PE=1 SV=1 |
| 106 | 152 | Q8VHF5\|CI | 22.82 | 2 | 2 | 9.65E+04 | 1 | 1 | 1 | | 51867 | Citrate synthase mitochondrial OS=Rattus norvegicus OX=10116 GN=Cs PE=1 SV=1 |
| 106 | 153 | tr\|G3V936 | 22.82 | 2 | 2 | 9.65E+04 | 1 | 1 | 1 | | 51831 | Citrate synthase OS=Rattus norvegicus OX=10116 GN=Cs PE=1 SV=1 |
| 78 | 18 | P26284\|OD | 22.75 | 3 | 3 | 1.47E+05 | 1 | 1 | 2 | | 43227 | Pyruvate dehydrogenase E1 component subunit alpha somatic form mitochondrial OS=Rattus norvegicus OX=10116 GN=Pdha1 PE=1 SV=2 |
| 97 | 83 | tr\|A0A411I | 21.45 | 3 | 3 | 1.40E+05 | 1 | 1 | 1 | | 40144 | Cytochrome b (Fragment) OS=Rattus norvegicus OX=10116 GN=Cytb PE=3 SV=1 |
| 97 | 84 | tr\|A0A0U2 | 21.45 | 3 | 3 | 1.40E+05 | 1 | 1 | 1 | | 40981 | Cytochrome b (Fragment) OS=Rattus norvegicus OX=10116 PE=3 SV=1 |
| 97 | 85 | tr\|A0A0U2 | 21.45 | 3 | 3 | 1.40E+05 | 1 | 1 | 1 | | 41037 | Cytochrome b (Fragment) OS=Rattus norvegicus OX=10116 PE=3 SV=1 |
| 97 | 86 | tr\|A0A411I | 21.45 | 3 | 3 | 1.40E+05 | 1 | 1 | 1 | | 41017 | Cytochrome b (Fragment) OS=Rattus norvegicus OX=10116 GN=Cytb PE=3 SV=1 |
| 97 | 87 | tr\|F8QU31 | 21.45 | 3 | 3 | 1.40E+05 | 1 | 1 | 1 | | 42181 | Cytochrome b (Fragment) OS=Rattus norvegicus OX=10116 GN=cytb PE=3 SV=1 |
| 97 | 88 | tr\|A0A411I | 21.45 | 3 | 3 | 1.40E+05 | 1 | 1 | 1 | | 42238 | Cytochrome b (Fragment) OS=Rattus norvegicus OX=10116 GN=Cytb PE=3 SV=1 |
| 97 | 89 | tr\|L0L4L8\| | 21.45 | 3 | 3 | 1.40E+05 | 1 | 1 | 1 | | 42470 | Cytochrome b (Fragment) OS=Rattus norvegicus OX=10116 GN=cytb PE=3 SV=1 |
| 97 | 90 | tr\|A0A140 | 21.45 | 3 | 3 | 1.40E+05 | 1 | 1 | 1 | | 42569 | Cytochrome b (Fragment) OS=Rattus norvegicus OX=10116 PE=3 SV=1 |
| 97 | 91 | tr\|A0A140 | 21.45 | 3 | 3 | 1.40E+05 | 1 | 1 | 1 | | 42574 | Cytochrome b (Fragment) OS=Rattus norvegicus OX=10116 PE=3 SV=1 |
| 97 | 92 | tr\|A0A140 | 21.45 | 3 | 3 | 1.40E+05 | 1 | 1 | 1 | | 42588 | Cytochrome b (Fragment) OS=Rattus norvegicus OX=10116 PE=3 SV=1 |
| 97 | 93 | tr\|A0A140 | 21.45 | 3 | 3 | 1.40E+05 | 1 | 1 | 1 | | 42527 | Cytochrome b (Fragment) OS=Rattus norvegicus OX=10116 PE=3 SV=1 |
| 97 | 95 | tr\|A0A385I | 21.45 | 3 | 3 | 1.40E+05 | 1 | 1 | 1 | | 42766 | Cytochrome b (Fragment) OS=Rattus norvegicus OX=10116 PE=3 SV=1 |

| | | | | | | | | | | | | |
|---|---|---|---|---|---|---|---|---|---|---|---|---|
| 97 | 96 | tr\|A0A3Q8 | 21.45 | 3 | 3 | 1.40E+05 | 1 | 1 | 1 | | 42712 | Cytochrome b (Fragment) OS=Rattus norvegicus OX=10116 GN=Cytb PE=3 SV=1 |
| 97 | 97 | tr\|A0A3Q8 | 21.45 | 3 | 3 | 1.40E+05 | 1 | 1 | 1 | | 42622 | Cytochrome b (Fragment) OS=Rattus norvegicus OX=10116 GN=Cytb PE=3 SV=1 |
| 97 | 98 | tr\|A0A3S6I | 21.45 | 3 | 3 | 1.40E+05 | 1 | 1 | 1 | | 42702 | Cytochrome b (Fragment) OS=Rattus norvegicus OX=10116 GN=Cytb PE=3 SV=1 |
| 97 | 100 | tr\|A0A3S6I | 21.45 | 3 | 3 | 1.40E+05 | 1 | 1 | 1 | | 42698 | Cytochrome b (Fragment) OS=Rattus norvegicus OX=10116 GN=Cytb PE=3 SV=1 |
| 97 | 101 | tr\|A0A097I | 21.45 | 3 | 3 | 1.40E+05 | 1 | 1 | 1 | | 42865 | Cytochrome b OS=Rattus norvegicus OX=10116 GN=CYTB PE=3 SV=1 |
| 97 | 102 | tr\|A0A220I | 21.45 | 3 | 3 | 1.40E+05 | 1 | 1 | 1 | | 43018 | Cytochrome b OS=Rattus norvegicus OX=10116 PE=3 SV=1 |
| 97 | 103 | tr\|D6NSP2 | 21.45 | 3 | 3 | 1.40E+05 | 1 | 1 | 1 | | 42945 | Cytochrome b (Fragment) OS=Rattus norvegicus OX=10116 GN=cytb PE=3 SV=1 |
| 97 | 104 | tr\|D6NSS6 | 21.45 | 3 | 3 | 1.40E+05 | 1 | 1 | 1 | | 42993 | Cytochrome b (Fragment) OS=Rattus norvegicus OX=10116 GN=cytb PE=3 SV=1 |
| 97 | 105 | tr\|Q06Q99 | 21.45 | 3 | 3 | 1.40E+05 | 1 | 1 | 1 | | 42997 | Cytochrome b OS=Rattus norvegicus OX=10116 GN=CYB PE=3 SV=1 |
| 97 | 106 | tr\|D6NSQ3 | 21.45 | 3 | 3 | 1.40E+05 | 1 | 1 | 1 | | 43012 | Cytochrome b (Fragment) OS=Rattus norvegicus OX=10116 GN=cytb PE=3 SV=1 |
| 97 | 107 | tr\|D6NSR3 | 21.45 | 3 | 3 | 1.40E+05 | 1 | 1 | 1 | | 42986 | Cytochrome b (Fragment) OS=Rattus norvegicus OX=10116 GN=cytb PE=3 SV=1 |
| 97 | 108 | tr\|D6NSP4 | 21.45 | 3 | 3 | 1.40E+05 | 1 | 1 | 1 | | 43016 | Cytochrome b (Fragment) OS=Rattus norvegicus OX=10116 GN=cytb PE=3 SV=1 |
| 97 | 109 | tr\|D6NSQ8 | 21.45 | 3 | 3 | 1.40E+05 | 1 | 1 | 1 | | 43016 | Cytochrome b (Fragment) OS=Rattus norvegicus OX=10116 GN=cytb PE=3 SV=1 |
| 97 | 110 | tr\|D6NSR8 | 21.45 | 3 | 3 | 1.40E+05 | 1 | 1 | 1 | | 42998 | Cytochrome b (Fragment) OS=Rattus norvegicus OX=10116 GN=cytb PE=3 SV=1 |
| 97 | 111 | tr\|F8QU37 | 21.45 | 3 | 3 | 1.40E+05 | 1 | 1 | 1 | | 42992 | Cytochrome b (Fragment) OS=Rattus norvegicus OX=10116 GN=cytb PE=3 SV=1 |
| 97 | 112 | tr\|F2Q6S6\| | 21.45 | 3 | 3 | 1.40E+05 | 1 | 1 | 1 | | 42938 | Cytochrome b (Fragment) OS=Rattus norvegicus OX=10116 GN=cytb PE=3 SV=1 |
| 97 | 113 | tr\|A0A0A1 | 21.45 | 3 | 3 | 1.40E+05 | 1 | 1 | 1 | | 42989 | Cytochrome b OS=Rattus norvegicus OX=10116 GN=CYTB PE=3 SV=1 |
| 97 | 114 | tr\|D6NSS7 | 21.45 | 3 | 3 | 1.40E+05 | 1 | 1 | 1 | | 42948 | Cytochrome b (Fragment) OS=Rattus norvegicus OX=10116 GN=cytb PE=3 SV=1 |
| 97 | 115 | tr\|Q8SEY9\| | 21.45 | 3 | 3 | 1.40E+05 | 1 | 1 | 1 | | 43015 | Cytochrome b OS=Rattus norvegicus OX=10116 GN=cytb PE=3 SV=1 |
| 97 | 116 | tr\|D6NSR7 | 21.45 | 3 | 3 | 1.40E+05 | 1 | 1 | 1 | | 42982 | Cytochrome b (Fragment) OS=Rattus norvegicus OX=10116 GN=cytb PE=3 SV=1 |
| 97 | 117 | tr\|A0A220I | 21.45 | 3 | 3 | 1.40E+05 | 1 | 1 | 1 | | 42968 | Cytochrome b OS=Rattus norvegicus OX=10116 PE=3 SV=1 |
| 97 | 118 | tr\|F2Q6S5\| | 21.45 | 3 | 3 | 1.40E+05 | 1 | 1 | 1 | | 42952 | Cytochrome b (Fragment) OS=Rattus norvegicus OX=10116 GN=cytb PE=3 SV=1 |
| 97 | 119 | tr\|D6NSQ6 | 21.45 | 3 | 3 | 1.40E+05 | 1 | 1 | 1 | | 42952 | Cytochrome b (Fragment) OS=Rattus norvegicus OX=10116 GN=cytb PE=3 SV=1 |
| 97 | 120 | tr\|D6NSQ0 | 21.45 | 3 | 3 | 1.40E+05 | 1 | 1 | 1 | | 42968 | Cytochrome b (Fragment) OS=Rattus norvegicus OX=10116 GN=cytb PE=3 SV=1 |
| 97 | 121 | tr\|Q5UAI7 | 21.45 | 3 | 3 | 1.40E+05 | 1 | 1 | 1 | | 42998 | Cytochrome b OS=Rattus norvegicus OX=10116 GN=CYTB PE=3 SV=1 |
| 97 | 122 | tr\|D6NSR2 | 21.45 | 3 | 3 | 1.40E+05 | 1 | 1 | 1 | | 43073 | Cytochrome b (Fragment) OS=Rattus norvegicus OX=10116 GN=cytb PE=3 SV=1 |
| 97 | 123 | tr\|A0A0S12 | 21.45 | 3 | 3 | 1.40E+05 | 1 | 1 | 1 | | 43002 | Cytochrome b OS=Rattus norvegicus OX=10116 GN=CYTB PE=3 SV=1 |
| 97 | 124 | tr\|D6NSR1 | 21.45 | 3 | 3 | 1.40E+05 | 1 | 1 | 1 | | 43002 | Cytochrome b (Fragment) OS=Rattus norvegicus OX=10116 GN=cytb PE=3 SV=1 |
| 97 | 125 | tr\|Q8HIC4\| | 21.45 | 3 | 3 | 1.40E+05 | 1 | 1 | 1 | | 43012 | Cytochrome b OS=Rattus norvegicus OX=10116 GN=Mt-cyb PE=3 SV=1 |
| 97 | 126 | tr\|A0A220I | 21.45 | 3 | 3 | 1.40E+05 | 1 | 1 | 1 | | 42968 | Cytochrome b OS=Rattus norvegicus OX=10116 PE=3 SV=1 |
| 97 | 127 | tr\|S5RKC8\| | 21.45 | 3 | 3 | 1.40E+05 | 1 | 1 | 1 | | 42997 | Cytochrome b OS=Rattus norvegicus OX=10116 GN=CYTB PE=3 SV=1 |
| 97 | 128 | tr\|A0A0S12 | 21.45 | 3 | 3 | 1.40E+05 | 1 | 1 | 1 | | 43016 | Cytochrome b OS=Rattus norvegicus OX=10116 GN=CYTB PE=3 SV=1 |
| 97 | 130 | P00159\|CY | 21.45 | 3 | 3 | 1.40E+05 | 1 | 1 | 1 | | 43012 | Cytochrome b OS=Rattus norvegicus OX=10116 GN=Mt-Cyb PE=3 SV=3 |
| 97 | 132 | tr\|A0A220I | 21.45 | 3 | 3 | 1.40E+05 | 1 | 1 | 1 | | 42970 | Cytochrome b OS=Rattus norvegicus OX=10116 PE=3 SV=1 |
| 97 | 133 | tr\|H2KXA0 | 21.45 | 3 | 3 | 1.40E+05 | 1 | 1 | 1 | | 42978 | Cytochrome b (Fragment) OS=Rattus norvegicus OX=10116 GN=cytb PE=3 SV=1 |
| 97 | 134 | tr\|A0A096X | 21.45 | 3 | 3 | 1.40E+05 | 1 | 1 | 1 | | 43042 | Cytochrome b (Fragment) OS=Rattus norvegicus OX=10116 PE=3 SV=1 |
| 97 | 136 | tr\|Q9G898 | 21.45 | 3 | 3 | 1.40E+05 | 1 | 1 | 1 | | 42830 | Cytochrome b OS=Rattus norvegicus OX=10116 PE=2 SV=1 |
| 97 | 94 | tr\|A0A385I | 21.45 | 3 | 3 | 1.40E+05 | 1 | 1 | 1 | | 42780 | Cytochrome b (Fragment) OS=Rattus norvegicus OX=10116 PE=3 SV=1 |
| 97 | 99 | tr\|A0A385I | 21.45 | 3 | 3 | 1.40E+05 | 1 | 1 | 1 | | 42751 | Cytochrome b (Fragment) OS=Rattus norvegicus OX=10116 PE=3 SV=1 |
| 97 | 129 | tr\|R9TKN1 | 21.45 | 3 | 3 | 1.40E+05 | 1 | 1 | 1 | | 43012 | Cytochrome b OS=Rattus norvegicus OX=10116 GN=CYTB PE=3 SV=1 |
| 97 | 131 | tr\|L0N311\| | 21.45 | 3 | 3 | 1.40E+05 | 1 | 1 | 1 | | 43045 | Cytochrome b (Fragment) OS=Rattus norvegicus OX=10116 GN=cytb PE=3 SV=1 |
| 97 | 135 | tr\|D6NSP5 | 21.45 | 3 | 3 | 1.40E+05 | 1 | 1 | 1 | | 43012 | Cytochrome b (Fragment) OS=Rattus norvegicus OX=10116 GN=cytb PE=3 SV=1 |
| 99 | 953 | tr\|D4ACE9 | 21.16 | 1 | 1 | | 1 | 0 | 1 | | 103115 | Alpha-aminoadipic semialdehyde synthase mitochondrial OS=Rattus norvegicus OX=10116 GN=Aass PE=1 SV=3 |
| 99 | 954 | A2VCW9\|A | 21.16 | 1 | 1 | | 1 | 0 | 1 | | 102908 | Alpha-aminoadipic semialdehyde synthase mitochondrial OS=Rattus norvegicus OX=10116 GN=Aass PE=2 SV=1 |
| 98 | 560 | P28494\|M | 20.51 | 1 | 1 | 1.79E+06 | 1 | 1 | 1 | | 131242 | Alpha-mannosidase 2 OS=Rattus norvegicus OX=10116 GN=Man2a1 PE=1 SV=2 |
| 74 | 12052 | P32198\|CP | 20.51 | 1 | 1 | 3.46E+05 | 1 | 1 | 2 | Carbamido | 88126 | Carnitine O-palmitoyltransferase 1 liver isoform OS=Rattus norvegicus OX=10116 GN=Cpt1a PE=1 SV=2 |
| 151 | 23092 | Q63664\|K | 20.47 | 2 | 2 | 3.78E+05 | 1 | 1 | 1 | | 47963 | ATP-sensitive inward rectifier potassium channel 8 OS=Rattus norvegicus OX=10116 GN=Kcnj8 PE=1 SV=1 |
