## Supplementary material for "Permeability transition pore-related changes in the proteome and channel activity of ATP synthase dimers and monomers": RLM KCl-BM control Dimer of V compl.

<table>
<tr><th>Protein Group</th><th>Protein ID</th><th>Accession</th><th>-10lgP</th><th>Coverage (%)</th><th>Coverage (%) Sample 3</th><th>Area Sample 3</th><th>#Peptides</th><th>#Unique</th><th>#Spec Sample 3</th><th>PTM</th><th>Avg. Mass</th><th>Description</th></tr>
<tr><td>2</td><td>14</td><td>P15999|ATPA_RAT</td><td>382.48</td><td>66</td><td>66</td><td>277360000</td><td>51</td><td>51</td><td>109</td><td>Y</td><td>59754</td><td>ATP synthase subunit alpha, mitochondrial OS=Rattus norvegicus OX=10116 GN=Atp5f1a PE=1 SV=2</td></tr>
<tr><td>1</td><td>7</td><td>P10719|ATPB_RAT</td><td>366.27</td><td>86</td><td>86</td><td>369900000</td><td>50</td><td>50</td><td>149</td><td>Y</td><td>56354</td><td>ATP synthase subunit beta, mitochondrial OS=Rattus norvegicus OX=10116 GN=Atp5f1b PE=1 SV=2</td></tr>
<tr><td>1</td><td>8</td><td>tr|G3V6D3|G3V6D3_RAT</td><td>366.27</td><td>86</td><td>86</td><td>369900000</td><td>50</td><td>50</td><td>149</td><td>Y</td><td>56345</td><td>ATP synthase subunit beta OS=Rattus norvegicus OX=10116 GN=Atp5f1b PE=1 SV=1</td></tr>
<tr><td>4</td><td>4</td><td>P67779|PHB_RAT</td><td>275.36</td><td>86</td><td>86</td><td>71777000</td><td>23</td><td>23</td><td>38</td><td>N</td><td>29820</td><td>Prohibitin OS=Rattus norvegicus OX=10116 GN=Phb PE=1 SV=1</td></tr>
<tr><td>3</td><td>11</td><td>Q5XIH7|PHB2_RAT</td><td>267.91</td><td>73</td><td>73</td><td>66230000</td><td>28</td><td>28</td><td>48</td><td>Y</td><td>33312</td><td>Prohibitin-2 OS=Rattus norvegicus OX=10116 GN=Phb2 PE=1 SV=1</td></tr>
<tr><td>7</td><td>2</td><td>Q02253|MMSA_RAT</td><td>242.74</td><td>48</td><td>48</td><td>23705000</td><td>21</td><td>21</td><td>30</td><td>Y</td><td>57808</td><td>Methylmalonate-semialdehyde dehydrogenase [acylating], mitochondrial OS=Rattus norvegicus OX=10116 GN=Aldh6a1 PE=1 SV=1</td></tr>
<tr><td>7</td><td>3</td><td>tr|G3V7J0|G3V7J0_RAT</td><td>242.74</td><td>48</td><td>48</td><td>23705000</td><td>21</td><td>21</td><td>30</td><td>Y</td><td>57748</td><td>Aldehyde dehydrogenase family 6, subfamily A1, isoform CRA_b OS=Rattus norvegicus OX=10116 GN=Aldh6a1 PE=1 SV=1</td></tr>
<tr><td>6</td><td>116</td><td>P35435|ATPG_RAT</td><td>229.61</td><td>57</td><td>57</td><td>513120</td><td>16</td><td>1</td><td>35</td><td>Y</td><td>30191</td><td>ATP synthase subunit gamma, mitochondrial OS=Rattus norvegicus OX=10116 GN=Atp5f1c PE=1 SV=2</td></tr>
<tr><td>5</td><td>117</td><td>tr|Q6QI09|Q6QI09_RAT</td><td>221.49</td><td>25</td><td>25</td><td>2453300</td><td>16</td><td>1</td><td>35</td><td>Y</td><td>67721</td><td>ATP synthase subunit gamma, mitochondrial OS=Rattus norvegicus OX=10116 GN=Taf3 PE=1 SV=1</td></tr>
<tr><td>10</td><td>4260</td><td>P31399|ATP5H_RAT</td><td>218.85</td><td>74</td><td>74</td><td>14806000</td><td>12</td><td>12</td><td>17</td><td>Y</td><td>18763</td><td>ATP synthase subunit d, mitochondrial OS=Rattus norvegicus OX=10116 GN=Atp5pd PE=1 SV=3</td></tr>
<tr><td>8</td><td>41</td><td>P19511|AT5F1_RAT</td><td>208.5</td><td>46</td><td>46</td><td>44029000</td><td>13</td><td>12</td><td>26</td><td>N</td><td>28869</td><td>ATP synthase F(0) complex subunit B1, mitochondrial OS=Rattus norvegicus OX=10116 GN=Atp5pb PE=1 SV=1</td></tr>
<tr><td>9</td><td>9</td><td>P00507|AATM_RAT</td><td>202.5</td><td>40</td><td>40</td><td>23694000</td><td>16</td><td>16</td><td>22</td><td>Y</td><td>47314</td><td>Aspartate aminotransferase, mitochondrial OS=Rattus norvegicus OX=10116 GN=Got2 PE=1 SV=2</td></tr>
<tr><td>12</td><td>13</td><td>P07756|CPSM_RAT</td><td>170.01</td><td>6</td><td>6</td><td>1524800</td><td>9</td><td>9</td><td>12</td><td>Y</td><td>164579</td><td>Carbamoyl-phosphate synthase [ammonia], mitochondrial OS=Rattus norvegicus OX=10116 GN=Cps1 PE=1 SV=1</td></tr>
<tr><td>16</td><td>21</td><td>Q64428|ECHA_RAT</td><td>164.25</td><td>10</td><td>10</td><td>4189000</td><td>6</td><td>6</td><td>7</td><td>N</td><td>82665</td><td>Trifunctional enzyme subunit alpha, mitochondrial OS=Rattus norvegicus OX=10116 GN=Hadha PE=1 SV=2</td></tr>
<tr><td>15</td><td>25</td><td>tr|D3ZG43|D3ZG43_RAT</td><td>158.7</td><td>22</td><td>22</td><td>2680900</td><td>5</td><td>5</td><td>8</td><td>Y</td><td>30226</td><td>NADH dehydrogenase (Ubiquinone) Fe-S protein 3 (Predicted), isoform CRA_c OS=Rattus norvegicus OX=10116 GN=Ndufs3 PE=1 SV=1</td></tr>
<tr><td>11</td><td>2163</td><td>Q06647|ATPO_RAT</td><td>158.27</td><td>49</td><td>49</td><td>17943000</td><td>10</td><td>10</td><td>14</td><td>Y</td><td>23398</td><td>ATP synthase subunit O, mitochondrial OS=Rattus norvegicus OX=10116 GN=Atp5po PE=1 SV=1</td></tr>
<tr><td>14</td><td>6</td><td>Q66HF1|NDUS1_RAT</td><td>155.91</td><td>13</td><td>13</td><td>4688500</td><td>7</td><td>7</td><td>10</td><td>N</td><td>79412</td><td>NADH-ubiquinone oxidoreductase 75 kDa subunit, mitochondrial OS=Rattus norvegicus OX=10116 GN=Ndufs1 PE=1 SV=1</td></tr>
<tr><td>13</td><td>4261</td><td>tr|G3V7Y3|G3V7Y3_RAT</td><td>146.01</td><td>61</td><td>61</td><td>24198000</td><td>7</td><td>7</td><td>11</td><td>Y</td><td>17563</td><td>ATP synthase subunit delta, mitochondrial OS=Rattus norvegicus OX=10116 GN=Atp5f1d PE=1 SV=1</td></tr>
</table>

| | | | | | | | | | | | | |
|---|---|---|---|---|---|---|---|---|---|---|---|---|
| **22** | 4263 | Q6PDU7\|ATP5L_RAT | 127.48 | 46 | 46 | 1928000 | 3 | 3 | 4 | Y | 11433 | ATP synthase subunit g, mitochondrial OS=Rattus norvegicus OX=10116 GN=Atp5mg PE=1 SV=2 |
| **29** | 4270 | Q9JJW3\|ATPMD_RAT | 126.91 | 38 | 38 | 2369300 | 3 | 3 | 3 | N | 6408 | ATP synthase membrane subunit DAPIT, mitochondrial OS=Rattus norvegicus OX=10116 GN=Atp5md PE=1 SV=1 |
| **19** | 1 | P32551\|QCR2_RAT | 125.75 | 17 | 17 | 2291000 | 5 | 5 | 6 | N | 48396 | Cytochrome b-c1 complex subunit 2, mitochondrial OS=Rattus norvegicus OX=10116 GN=Uqcrc2 PE=1 SV=2 |
| **17** | 15 | Q5BK63\|NDUA9_RAT | 121.86 | 16 | 16 | 701120 | 4 | 4 | 7 | Y | 42559 | NADH dehydrogenase [ubiquinone] 1 alpha subcomplex subunit 9, mitochondrial OS=Rattus norvegicus OX=10116 GN=Ndufa9 PE=1 SV=2 |
| **20** | 33 | Q561S0\|NDUAA_RAT | 102.64 | 17 | 17 | 1898400 | 4 | 4 | 5 | Y | 40493 | NADH dehydrogenase [ubiquinone] 1 alpha subcomplex subunit 10, mitochondrial OS=Rattus norvegicus OX=10116 GN=Ndufa10 PE=1 SV=1 |
| **21** | 4264 | tr\|Q5UAJ5\|Q5UAJ5_RAT | 99.48 | 46 | 46 | 5060900 | 3 | 3 | 4 | N | 7642 | ATP synthase protein 8 OS=Rattus norvegicus OX=10116 GN=ATP8 PE=3 SV=1 |
| **35** | 12 | Q68FY0\|QCR1_RAT | 85.46 | 5 | 5 | 43626 | 2 | 2 | 2 | N | 52849 | Cytochrome b-c1 complex subunit 1, mitochondrial OS=Rattus norvegicus OX=10116 GN=Uqcrc1 PE=1 SV=1 |
| **26** | 23 | tr\|Q5XIH3\|Q5XIH3_RAT | 73.84 | 10 | 10 | 146380 | 3 | 3 | 3 | N | 50731 | NADH dehydrogenase [ubiquinone] flavoprotein 1, mitochondrial OS=Rattus norvegicus OX=10116 GN=Ndufv1 PE=1 SV=1 |
| **95** | 95 | Q63362\|NDUA5_RAT | 67.98 | 22 | 22 | 0 | 1 | 1 | 1 | N | 13412 | NADH dehydrogenase [ubiquinone] 1 alpha subcomplex subunit 5 OS=Rattus norvegicus OX=10116 GN=Ndufa5 PE=1 SV=3 |
| **24** | 19 | P22791\|HMCS2_RAT | 63.14 | 4 | 4 | 1662400 | 2 | 2 | 3 | N | 56912 | Hydroxymethylglutaryl-CoA synthase, mitochondrial OS=Rattus norvegicus OX=10116 GN=Hmgcs2 PE=1 SV=1 |
| **24** | 20 | tr\|Q68G44\|Q68G44_RAT | 63.14 | 4 | 4 | 1662400 | 2 | 2 | 3 | N | 56886 | 3-hydroxy-3-methylglutaryl coenzyme A synthase OS=Rattus norvegicus OX=10116 GN=Hmgcs2 PE=1 SV=1 |
| **28** | 4266 | D3ZAF6\|ATPK_RAT | 52.4 | 30 | 30 | 3083900 | 3 | 3 | 3 | Y | 10452 | ATP synthase subunit f, mitochondrial OS=Rattus norvegicus OX=10116 GN=Atp5mf PE=1 SV=1 |
| **96** | 42 | tr\|D4A565\|D4A565_RAT | 50.27 | 12 | 12 | 69212 | 1 | 1 | 1 | N | 21664 | NADH dehydrogenase (Ubiquinone) 1 beta subcomplex, 5 (Predicted), isoform CRA_b OS=Rattus norvegicus OX=10116 GN=Ndufb5 PE=1 SV=1 |
| **36** | 109 | P11240\|COX5A_RAT | 49.51 | 21 | 21 | 29933 | 1 | 1 | 2 | N | 16130 | Cytochrome c oxidase subunit 5A, mitochondrial OS=Rattus norvegicus OX=10116 GN=Cox5a PE=1 SV=1 |
| **97** | 29 | tr\|D4A0T0\|D4A0T0_RAT | 41.71 | 8 | 8 | 282740 | 1 | 1 | 1 | N | 20859 | NADH:ubiquinone oxidoreductase subunit B10 OS=Rattus norvegicus OX=10116 GN=Ndufb10 PE=1 SV=1 |
| **98** | 4272 | tr\|M0R601\|M0R601_RAT | 39.16 | 16 | 16 | 437370 | 1 | 1 | 1 | N | 5582 | Uncharacterized protein OS=Rattus norvegicus OX=10116 PE=4 SV=2 |
| **98** | 4273 | P29418\|ATP5E_RAT | 39.16 | 16 | 16 | 437370 | 1 | 1 | 1 | N | 5767 | ATP synthase subunit epsilon, mitochondrial OS=Rattus norvegicus OX=10116 GN=Atp5f1e PE=1 SV=2 |
| **37** | 77 | tr\|Q06QK5\|Q06QK5_RAT | 38.16 | 2 | 2 | 636560 | 1 | 1 | 2 | N | 68606 | NADH-ubiquinone oxidoreductase chain 5 OS=Rattus norvegicus OX=10116 GN=ND5 PE=3 SV=1 |
| **37** | 81 | P11661\|NU5M_RAT | 38.16 | 2 | 2 | 636560 | 1 | 1 | 2 | N | 68618 | NADH-ubiquinone oxidoreductase chain 5 OS=Rattus norvegicus OX=10116 GN=Mtnd5 PE=3 SV=3 |

| | | | | | | | | | | | | |
|---|---|---|---|---|---|---|---|---|---|---|---|---|
| **37** | 82 | tr\|Q06QG6\|Q06QG6_RAT | 38.16 | 2 | 2 | 636560 | 1 | 1 | 2 | N | 68588 | NADH-ubiquinone oxidoreductase chain 5 OS=Rattus norvegicus OX=10116 GN=ND5 PE=3 SV=1 |
| **37** | 83 | tr\|Q06QA1\|Q06QA1_RAT | 38.16 | 2 | 2 | 636560 | 1 | 1 | 2 | N | 68574 | NADH-ubiquinone oxidoreductase chain 5 OS=Rattus norvegicus OX=10116 GN=ND5 PE=3 SV=1 |
| **37** | 76 | tr\|A7XYD5\|A7XYD5_RAT | 38.16 | 2 | 2 | 636560 | 1 | 1 | 2 | N | 68856 | NADH-ubiquinone oxidoreductase chain 5 OS=Rattus norvegicus OX=10116 GN=Nd5 PE=3 SV=1 |
| **37** | 78 | tr\|Q8SEZ0\|Q8SEZ0_RAT | 38.16 | 2 | 2 | 636560 | 1 | 1 | 2 | N | 68618 | NADH-ubiquinone oxidoreductase chain 5 OS=Rattus norvegicus OX=10116 GN=Mt-nd5 PE=3 SV=1 |
| **37** | 79 | tr\|A0A096XKT9\|A0A096XKT9_RAT | 38.16 | 2 | 2 | 636560 | 1 | 1 | 2 | N | 68584 | NADH-ubiquinone oxidoreductase chain 5 OS=Rattus norvegicus OX=10116 GN=ND5 PE=3 SV=1 |
| **37** | 80 | tr\|A7XYC1\|A7XYC1_RAT | 38.16 | 2 | 2 | 636560 | 1 | 1 | 2 | N | 68841 | NADH-ubiquinone oxidoreductase chain 5 OS=Rattus norvegicus OX=10116 GN=Nd5 PE=3 SV=1 |
| **37** | 84 | tr\|A0A0A1FZN8\|A0A0A1FZN8_RAT | 38.16 | 2 | 2 | 636560 | 1 | 1 | 2 | N | 68970 | NADH-ubiquinone oxidoreductase chain 5 OS=Rattus norvegicus OX=10116 GN=ND5 PE=3 SV=1 |
| **18** | 179 | F1M775\|DIAP1_RAT | 36.06 | 2 | 2 | 2036300 | 3 | 2 | 6 | Y | 140418 | Protein diaphanous homolog 1 OS=Rattus norvegicus OX=10116 GN=Diaph1 PE=1 SV=3 |
| **38** | 4275 | tr\|D3ZA45\|D3ZA45_RAT | 35.36 | 2 | 2 | 780470 | 1 | 1 | 2 | N | 52491 | Autophagy-related protein 13 OS=Rattus norvegicus OX=10116 GN=Atg13 PE=1 SV=2 |
| **99** | 34 | tr\|Q5RJN0\|Q5RJN0_RAT | 34.81 | 6 | 6 | 196860 | 1 | 1 | 1 | N | 23945 | NADH dehydrogenase (Ubiquinone) Fe-S protein 7 OS=Rattus norvegicus OX=10116 GN=Ndufs7 PE=1 SV=1 |
| **100** | 22 | P19234\|NDUV2_RAT | 33.1 | 5 | 5 | 2019900 | 1 | 1 | 1 | N | 27378 | NADH dehydrogenase [ubiquinone] flavoprotein 2, mitochondrial OS=Rattus norvegicus OX=10116 GN=Ndufv2 PE=1 SV=2 |
| **47** | 53 | P52873\|PYC_RAT | 32.92 | 2 | 2 | 312410 | 1 | 1 | 1 | N | 129777 | Pyruvate carboxylase, mitochondrial OS=Rattus norvegicus OX=10116 GN=Pc PE=1 SV=2 |
| **47** | 54 | tr\|A0A0G2JTL5\|A0A0G2JTL5_RAT | 32.92 | 2 | 2 | 312410 | 1 | 1 | 1 | N | 140005 | Pyruvate carboxylase, mitochondrial OS=Rattus norvegicus OX=10116 GN=Pc PE=1 SV=1 |
| **101** | 157 | tr\|F1LTG5\|F1LTG5_RAT | 32.32 | 14 | 14 | 493200 | 1 | 1 | 1 | N | 13829 | Uncharacterized protein OS=Rattus norvegicus OX=10116 PE=4 SV=2 |
| **101** | 158 | P12075\|COX5B_RAT | 32.32 | 14 | 14 | 493200 | 1 | 1 | 1 | N | 13915 | Cytochrome c oxidase subunit 5B, mitochondrial OS=Rattus norvegicus OX=10116 GN=Cox5b PE=1 SV=2 |
| **25** | 369 | P11530\|DMD_RAT | 30.6 | 0 | 0 | 1032100 | 2 | 2 | 3 | Y | 425830 | Dystrophin OS=Rattus norvegicus OX=10116 GN=Dmd PE=1 SV=2 |
| **39** | 4278 | tr\|A0A0U1RRQ6\|A0A0U1RRQ6_RAT | 27.67 | 3 | 3 | 1497800 | 1 | 1 | 2 | Y | 35344 | Solute carrier family 25, member 44 OS=Rattus norvegicus OX=10116 GN=Slc25a44 PE=1 SV=1 |
| **39** | 4279 | tr\|D3ZSN7\|D3ZSN7_RAT | 27.67 | 2 | 2 | 1497800 | 1 | 1 | 2 | Y | 37879 | Similar to CG5805-PA (Predicted), isoform CRA_a OS=Rattus norvegicus OX=10116 GN=Slc25a44 PE=1 SV=1 |
| **30** | 390 | tr\|O70151\|O70151_RAT | 44161 | 1 | 1 | 0 | 2 | 2 | 2 | Y | 276097 | Acetyl-CoA carboxylase OS=Rattus norvegicus OX=10116 GN=Acacb PE=2 SV=1 |

| | | | | | | | | | | | | |
|---|---|---|---|---|---|---|---|---|---|---|---|---|
| **105** | 2187 | Q06645\|AT5G1_RAT | 25.94 | 5 | 5 | 3196100 | 1 | 1 | 1 | N | 14244 | ATP synthase F(0) complex subunit C1, mitochondrial OS=Rattus norvegicus OX=10116 GN=Atp5mc1 PE=1 SV=1 |
| **105** | 2189 | tr\|Q499S2\|Q499S2_RAT | 25.94 | 5 | 5 | 3196100 | 1 | 1 | 1 | N | 14745 | ATP synthase F(0) complex subunit C3, mitochondrial OS=Rattus norvegicus OX=10116 GN=Atp5mc3 PE=2 SV=1 |
| **105** | 2188 | Q06646\|AT5G2_RAT | 25.94 | 5 | 5 | 3196100 | 1 | 1 | 1 | N | 14918 | ATP synthase F(0) complex subunit C2, mitochondrial OS=Rattus norvegicus OX=10116 GN=Atp5mc2 PE=1 SV=1 |
| **105** | 2190 | Q71S46\|AT5G3_RAT | 25.94 | 5 | 5 | 3196100 | 1 | 1 | 1 | N | 14693 | ATP synthase F(0) complex subunit C3, mitochondrial OS=Rattus norvegicus OX=10116 GN=Atp5mc3 PE=1 SV=1 |
| **105** | 2191 | tr\|A0A0G2JTN8\|A0A0G2JTN8_RAT | 25.94 | 5 | 5 | 3196100 | 1 | 1 | 1 | N | 15465 | ATP synthase F(0) complex subunit C2, mitochondrial OS=Rattus norvegicus OX=10116 GN=Atp5mc2 PE=3 SV=1 |
| **108** | 2238 | tr\|B5DFK5\|B5DFK5_RAT | 24.34 | 1 | 1 | 0 | 1 | 1 | 1 | Y | 119424 | Hip1r protein OS=Rattus norvegicus OX=10116 GN=Hip1r PE=2 SV=1 |
| **108** | 2240 | tr\|F1LML7\|F1LML7_RAT | 24.34 | 1 | 1 | 0 | 1 | 1 | 1 | Y | 119553 | Huntingtin-interacting protein 1-related OS=Rattus norvegicus OX=10116 GN=Hip1r PE=1 SV=1 |
| **108** | 2241 | tr\|Q99PW9\|Q99PW9_RAT | 24.34 | 1 | 1 | 0 | 1 | 1 | 1 | Y | 120572 | Huntingtin interacting protein 1 related (Fragment) OS=Rattus norvegicus OX=10116 GN=Hip1r PE=2 SV=1 |
| **45** | 140 | tr\|F1LPI6\|F1LPI6_RAT | 44067 | 0 | 0 | 0 | 1 | 1 | 2 | N | 200075 | Peripheral-type benzodiazepine receptor-associated protein 1 OS=Rattus norvegicus OX=10116 GN=Tspoap1 PE=4 SV=1 |
| **45** | 141 | Q9JIR0\|RIMB1_RAT | 44067 | 0 | 0 | 0 | 1 | 1 | 2 | N | 200202 | Peripheral-type benzodiazepine receptor-associated protein 1 OS=Rattus norvegicus OX=10116 GN=Tspoap1 PE=1 SV=2 |
| **111** | 51 | P10888\|COX41_RAT | 43914 | 4 | 4 | 0 | 1 | 1 | 1 | N | 19515 | Cytochrome c oxidase subunit 4 isoform 1, mitochondrial OS=Rattus norvegicus OX=10116 GN=Cox4i1 PE=1 SV=1 |
| **59** | 283 | tr\|F1MAR6\|F1MAR6_RAT | 23.86 | 1 | 1 | 0 | 1 | 1 | 1 | N | 68095 | Proline dehydrogenase OS=Rattus norvegicus OX=10116 GN=Prodh1 PE=1 SV=2 |
| **110** | 38 | tr\|D4A3V2\|D4A3V2_RAT | 23.69 | 8 | 8 | 134410 | 1 | 1 | 1 | N | 15224 | NADH dehydrogenase [ubiquinone] 1 alpha subcomplex subunit 6 OS=Rattus norvegicus OX=10116 GN=Ndufa6 PE=1 SV=1 |
| **112** | 164 | Q5M9G9\|FAKD4_RAT | 23.5 | 1 | 1 | 0 | 1 | 1 | 1 | N | 71181 | FAST kinase domain-containing protein 4 OS=Rattus norvegicus OX=10116 GN=Tbrg4 PE=2 SV=1 |
| **116** | 2199 | Q32Q86\|CRY1_RAT | 23.13 | 1 | 1 | 1294500 | 1 | 1 | 1 | N | 66231 | Cryptochrome-1 OS=Rattus norvegicus OX=10116 GN=Cry1 PE=1 SV=1 |
| **48** | 2554 | tr\|F1M4Y5\|F1M4Y5_RAT | 22.83 | 1 | 1 | 7464100 | 1 | 1 | 1 | Y | 108218 | Storkhead box 1 OS=Rattus norvegicus OX=10116 GN=Stox1 PE=4 SV=2 |
| **48** | 2544 | tr\|A0A0G2K418\|A0A0G2K418_RAT | 22.83 | 1 | 1 | 7464100 | 1 | 1 | 1 | Y | 96381 | Storkhead box 1 OS=Rattus norvegicus OX=10116 GN=Stox1 PE=4 SV=1 |
| **115** | 4293 | Q63449\|PRGR_RAT | 22.77 | 1 | 1 | 0 | 1 | 1 | 1 | N | 99408 | Progesterone receptor OS=Rattus norvegicus OX=10116 GN=Pgr PE=2 SV=1 |
| **113** | 4292 | B0K035\|MTFR2_RAT | 22.71 | 7 | 7 | 2025700 | 1 | 1 | 1 | Y | 40772 | Mitochondrial fission regulator 2 OS=Rattus norvegicus OX=10116 GN=Mtfr2 PE=2 SV=1 |

| | | | | | | | | | | | | |
|---|---|---|---|---|---|---|---|---|---|---|---|---|
| **114** | 435 | Q5XIJ9\|SCO2A_RAT | 22.7 | 4 | 4 | 0 | 1 | 1 | 1 | Y | 56900 | Succinyl-CoA:3-ketoacid coenzyme A transferase 2A, mitochondrial OS=Rattus norvegicus OX=10116 GN=Oxct2a PE=1 SV=1 |
| **117** | 2443 | tr\|A0A0G2JWL4\|A0A0G2JWL4_RAT | 22.25 | 4 | 4 | 30417 | 1 | 1 | 1 | Y | 68423 | FA complementation group G OS=Rattus norvegicus OX=10116 GN=Fancg PE=4 SV=1 |
| **119** | 52 | tr\|B0BNE6\|B0BNE6_RAT | 21.84 | 4 | 4 | 0 | 1 | 1 | 1 | N | 23970 | NADH dehydrogenase (Ubiquinone) Fe-S protein 8 (Predicted), isoform CRA_a OS=Rattus norvegicus OX=10116 GN=Ndufs8 PE=1 SV=1 |
| **93** | 4365 | tr\|A0A0G2K1N9\|A0A0G2K1N9_RAT | 20.81 | 1 | 1 | 1525500 | 1 | 1 | 1 | N | 74235 | Selenoprotein O OS=Rattus norvegicus OX=10116 GN=Selenoo PE=1 SV=1 |
| **93** | 4366 | tr\|B2GVA1\|B2GVA1_RAT | 20.81 | 1 | 1 | 1525500 | 1 | 1 | 1 | N | 74385 | Selenoprotein O OS=Rattus norvegicus OX=10116 GN=Selenoo PE=2 SV=1 |
| **126** | 4307 | Q1HCL7\|NAKD2_RAT | 20.57 | 1 | 1 | 0 | 1 | 1 | 1 | N | 48116 | NAD kinase 2, mitochondrial OS=Rattus norvegicus OX=10116 GN=Nadk2 PE=1 SV=1 |
| **123** | 4302 | tr\|D4A676\|D4A676_RAT | 20.48 | 4 | 4 | 646040 | 1 | 1 | 1 | N | 24898 | GrpE protein homolog OS=Rattus norvegicus OX=10116 GN=Grpel2 PE=3 SV=1 |
| **124** | 2566 | tr\|F1LSY7\|F1LSY7_RAT | 20.24 | 1 | 1 | 11066000 | 1 | 1 | 1 | Y | 109031 | Endoplasmic reticulum to nucleus-signaling 1 OS=Rattus norvegicus OX=10116 GN=Ern1 PE=4 SV=2 |
| **124** | 2567 | tr\|A0A0G2K2H4\|A0A0G2K2H4_RAT | 20.24 | 1 | 1 | 11066000 | 1 | 1 | 1 | Y | 110150 | Endoplasmic reticulum to nucleus-signaling 1 OS=Rattus norvegicus OX=10116 GN=Ern1 PE=2 SV=1 |
| **44** | 4305 | tr\|G3V8G1\|G3V8G1_RAT | 20.2 | 1 | 1 | 2305300 | 1 | 1 | 2 | Y | 137113 | ATP/GTP binding protein 1 (Predicted), isoform CRA_a OS=Rattus norvegicus OX=10116 GN=Agtpbp1 PE=4 SV=1 |
| **total 85 proteins** | | | | | | | | | | | | |
