## Supplementary material for "Permeability transition pore-related changes in the proteome and channel activity of ATP synthase dimers and monomers": RLM KCl-BM control Monomer of V compl.

| Protein Group | Protein ID | Accession | -10lgP | Coverage (%) | Coverage (%) Sample 7 | Area Sample 7 | #Peptides | #Unique | #Spec Sample 7 | PTM | Avg. Mass | Description |
|---|---|---|---|---|---|---|---|---|---|---|---|---|
| **2** | 14 | P15999\|ATPA_RAT | 432.09 | 77 | 77 | 2723700000 | 104 | 104 | 270 | Y | 59754 | ATP synthase subunit alpha, mitochondrial OS=Rattus norvegicus OX=10116 GN=Atp5f1a PE=1 SV=2 |
| **3** | 53 | P52873\|PYC_RAT | 423.33 | 69 | 69 | 485200000 | 89 | 87 | 137 | Y | 129777 | Pyruvate carboxylase, mitochondrial OS=Rattus norvegicus OX=10116 GN=Pc PE=1 SV=2 |
| **3** | 54 | tr\|A0A0G2JTL5\|A0A0G2JTL5_RA | 423.33 | 64 | 64 | 485200000 | 89 | 87 | 137 | Y | 140005 | Pyruvate carboxylase, mitochondrial OS=Rattus norvegicus OX=10116 GN=Pc PE=1 SV=1 |
| **1** | 7 | P10719\|ATPB_RAT | 408.87 | 82 | 82 | 3460600000 | 84 | 83 | 301 | Y | 56354 | ATP synthase subunit beta, mitochondrial OS=Rattus norvegicus OX=10116 GN=Atp5f1b PE=1 SV=2 |
| **1** | 8 | tr\|G3V6D3\|G3V6D3_RAT | 408.87 | 82 | 82 | 3460600000 | 84 | 83 | 301 | Y | 56345 | ATP synthase subunit beta OS=Rattus norvegicus OX=10116 GN=Atp5f1b PE=1 SV=1 |
| **5** | 4260 | P31399\|ATP5H_RAT | 358.66 | 93 | 93 | 461400000 | 40 | 40 | 67 | Y | 18763 | ATP synthase subunit d, mitochondrial OS=Rattus norvegicus OX=10116 GN=Atp5pd PE=1 SV=3 |
| **10** | 21 | Q64428\|ECHA_RAT | 348.58 | 45 | 45 | 94713000 | 36 | 36 | 48 | Y | 82665 | Trifunctional enzyme subunit alpha, mitochondrial OS=Rattus norvegicus OX=10116 GN=Hadha PE=1 SV=2 |
| **6** | 13 | P07756\|CPSM_RAT | 335.39 | 32 | 32 | 99521000 | 38 | 37 | 53 | Y | 164579 | Carbamoyl-phosphate synthase [ammonia], mitochondrial OS=Rattus norvegicus OX=10116 GN=Cps1 PE=1 SV=1 |
| **9** | 6796 | P04762\|CATA_RAT | 325.85 | 71 | 71 | 159850000 | 34 | 33 | 48 | Y | 59757 | Catalase OS=Rattus norvegicus OX=10116 GN=Cat PE=1 SV=3 |
| **8** | 116 | P35435\|ATPG_RAT | 283.13 | 68 | 68 | 222780000 | 29 | 29 | 50 | Y | 30191 | ATP synthase subunit gamma, mitochondrial OS=Rattus norvegicus OX=10116 GN=Atp5f1c PE=1 SV=2 |
| **12** | 9 | P00507\|AATM_RAT | 281.21 | 64 | 64 | 95293000 | 25 | 25 | 35 | Y | 47314 | Aspartate aminotransferase, mitochondrial OS=Rattus norvegicus OX=10116 GN=Got2 PE=1 SV=2 |
| **4** | 41 | P19511\|AT5F1_RAT | 277.66 | 71 | 71 | 460220000 | 31 | 30 | 71 | Y | 28869 | ATP synthase F(0) complex subunit B1, mitochondrial OS=Rattus norvegicus OX=10116 GN=Atp5pb PE=1 SV=1 |
| **21** | 19 | P22791\|HMCS2_RAT | 262.65 | 32 | 32 | 21445000 | 16 | 16 | 16 | Y | 56912 | Hydroxymethylglutaryl-CoA synthase, mitochondrial OS=Rattus norvegicus OX=10116 GN=Hmgcs2 PE=1 SV=1 |
| **21** | 20 | tr\|Q68G44\|Q68G44_RAT | 262.65 | 32 | 32 | 21445000 | 16 | 16 | 16 | Y | 56886 | 3-hydroxy-3-methylglutaryl coenzyme A synthase OS=Rattus norvegicus OX=10116 GN=Hmgcs2 PE=1 SV=1 |
| **7** | 4261 | tr\|G3V7Y3\|G3V7Y3_RAT | 256.07 | 82 | 82 | 658280000 | 20 | 20 | 52 | Y | 17563 | ATP synthase subunit delta, mitochondrial OS=Rattus norvegicus OX=10116 GN=Atp5f1d PE=1 SV=1 |
| **11** | 2163 | Q06647\|ATPO_RAT | 255.64 | 76 | 76 | 507150000 | 27 | 27 | 45 | Y | 23398 | ATP synthase subunit O, mitochondrial OS=Rattus norvegicus OX=10116 GN=Atp5po PE=1 SV=1 |
| **15** | 85 | tr\|A0A0G2JVH4\|A0A0G2JVH4_R | 248.58 | 25 | 25 | 10599000 | 20 | 20 | 20 | Y | 86230 | MICOS complex subunit MIC60 OS=Rattus norvegicus OX=10116 GN=Immt PE=1 SV=1 |
| **13** | 40 | Q60587\|ECHB_RAT | 243.61 | 37 | 37 | 25243000 | 20 | 20 | 24 | Y | 51414 | Trifunctional enzyme subunit beta, mitochondrial OS=Rattus norvegicus OX=10116 GN=Hadhb PE=1 SV=1 |
| **16** | 2302 | P16970\|ABCD3_RAT | 241.35 | 27 | 27 | 18504000 | 15 | 15 | 20 | Y | 75316 | ATP-binding cassette sub-family D member 3 OS=Rattus norvegicus OX=10116 GN=Abcd3 PE=1 SV=3 |
| **14** | 2 | Q02253\|MMSA_RAT | 240.93 | 47 | 47 | 38664000 | 18 | 18 | 24 | Y | 57808 | Methylmalonate-semialdehyde dehydrogenase [acylating], mitochondrial OS=Rattus norvegicus OX=10116 GN=Aldh6a1 PE=1 SV=1 |
| **14** | 3 | tr\|G3V7J0\|G3V7J0_RAT | 240.93 | 47 | 47 | 38664000 | 18 | 18 | 24 | Y | 57748 | Aldehyde dehydrogenase family 6, subfamily A1, isoform CRA_b OS=Rattus norvegicus OX=10116 GN=Aldh6a1 PE=1 SV=1 |
| **17** | 12 | Q68FY0\|QCR1_RAT | 239.36 | 43 | 43 | 22165000 | 14 | 14 | 20 | Y | 52849 | Cytochrome b-c1 complex subunit 1, mitochondrial OS=Rattus norvegicus OX=10116 GN=Uqcrc1 PE=1 SV=1 |
| **19** | 1 | P32551\|QCR2_RAT | 237.64 | 45 | 45 | 35141000 | 15 | 15 | 17 | N | 48396 | Cytochrome b-c1 complex subunit 2, mitochondrial OS=Rattus norvegicus OX=10116 GN=Uqcrc2 PE=1 SV=2 |
| **20** | 4373 | P0C2X9\|AL4A1_RAT | 233.98 | 34 | 34 | 47000000 | 13 | 13 | 16 | Y | 61869 | Delta-1-pyrroline-5-carboxylate dehydrogenase, mitochondrial OS=Rattus norvegicus OX=10116 GN=Aldh4a1 PE=1 SV=1 |
| **29** | 542 | P10860\|DHE3_RAT | 224.76 | 20 | 20 | 6627100 | 8 | 8 | 9 | N | 61416 | Glutamate dehydrogenase 1, mitochondrial OS=Rattus norvegicus OX=10116 GN=Glud1 PE=1 SV=2 |
| **18** | 9289 | P24329\|THTR_RAT | 224.6 | 40 | 40 | 56148000 | 14 | 14 | 19 | Y | 33407 | Thiosulfate sulfurtransferase OS=Rattus norvegicus OX=10116 GN=Tst PE=1 SV=3 |
| **24** | 229 | A2VCW9\|AASS_RAT | 217.43 | 15 | 15 | 780490 | 13 | 1 | 15 | N | 102908 | Alpha-aminoadipic semialdehyde synthase, mitochondrial OS=Rattus norvegicus OX=10116 GN=Aass PE=2 SV=1 |

| | | | | | | | | | | | | |
|---|---|---|---|---|---|---|---|---|---|---|---|---|
| **27** | 35 | P13086\|SUCA_RAT | 213.6 | 29 | 29 | 8325500 | 11 | 11 | 12 | Y | 36148 | Succinate--CoA ligase [ADP/GDP-forming] subunit alpha, mitochondrial OS=Rattus norvegicus OX=10116 GN=Suclg1 PE=2 SV=2 |
| **27** | 36 | tr\|A0A0H2UHE1\|A0A0H2UHE1_ | 213.6 | 28 | 28 | 8325500 | 11 | 11 | 12 | Y | 37560 | Succinate--CoA ligase [ADP/GDP-forming] subunit alpha, mitochondrial OS=Rattus norvegicus OX=10116 GN=Suclg1 PE=1 SV=1 |
| **23** | 228 | tr\|D4ACE9\|D4ACE9_RAT | 212.29 | 15 | 15 | 1908500 | 13 | 1 | 15 | Y | 103115 | Alpha-aminoadipic semialdehyde synthase, mitochondrial OS=Rattus norvegicus OX=10116 GN=Aass PE=1 SV=3 |
| **28** | 39 | P17764\|THIL_RAT | 208.41 | 24 | 24 | 13235000 | 8 | 8 | 11 | N | 44695 | Acetyl-CoA acetyltransferase, mitochondrial OS=Rattus norvegicus OX=10116 GN=Acat1 PE=1 SV=1 |
| **26** | 4263 | Q6PDU7\|ATP5L_RAT | 197.76 | 89 | 89 | 83153000 | 7 | 7 | 13 | Y | 11433 | ATP synthase subunit g, mitochondrial OS=Rattus norvegicus OX=10116 GN=Atp5mg PE=1 SV=2 |
| **32** | 289 | P07872\|ACOX1_RAT | 188.9 | 11 | 11 | 3079100 | 7 | 7 | 7 | Y | 74679 | Peroxisomal acyl-coenzyme A oxidase 1 OS=Rattus norvegicus OX=10116 GN=Acox1 PE=1 SV=1 |
| **30** | 4602 | B3DMA2\|ACD11_RAT | 182.87 | 11 | 11 | 7627400 | 8 | 8 | 8 | Y | 87371 | Acyl-CoA dehydrogenase family member 11 OS=Rattus norvegicus OX=10116 GN=Acad11 PE=1 SV=1 |
| **33** | 4338 | P63039\|CH60_RAT | 170.4 | 16 | 16 | 2991300 | 7 | 6 | 7 | N | 60956 | 60 kDa heat shock protein, mitochondrial OS=Rattus norvegicus OX=10116 GN=Hspd1 PE=1 SV=1 |
| **33** | 4339 | tr\|A0A482IDN3\|A0A482IDN3_R | 170.4 | 16 | 16 | 2991300 | 7 | 6 | 7 | N | 60956 | Hsp60 OS=Rattus norvegicus OX=10116 GN=Hspd1 PE=2 SV=1 |
| **36** | 13889 | P21571\|ATP5J_RAT | 169.58 | 37 | 37 | 11661000 | 5 | 5 | 6 | N | 12494 | ATP synthase-coupling factor 6, mitochondrial OS=Rattus norvegicus OX=10116 GN=Atp5pf PE=1 SV=1 |
| **31** | 13896 | D3Z9R8\|ATP68_RAT | 159.56 | 37 | 37 | 12692000 | 5 | 5 | 7 | Y | 6914 | ATP synthase subunit ATP5MPL, mitochondrial OS=Rattus norvegicus OX=10116 GN=Atp5mpl PE=1 SV=1 |
| **34** | 28 | tr\|D3ZFQ8\|D3ZFQ8_RAT | 142.12 | 14 | 14 | 6386600 | 5 | 5 | 7 | N | 35435 | Cytochrome c-1 OS=Rattus norvegicus OX=10116 GN=Cyc1 PE=1 SV=3 |
| **37** | 51 | P10888\|COX41_RAT | 141.37 | 30 | 30 | 6270700 | 5 | 5 | 6 | N | 19515 | Cytochrome c oxidase subunit 4 isoform 1, mitochondrial OS=Rattus norvegicus OX=10116 GN=Cox4i1 PE=1 SV=1 |
| **42** | 4264 | tr\|Q5UAJ5\|Q5UAJ5_RAT | 140.51 | 55 | 55 | 96459000 | 4 | 4 | 5 | Y | 7642 | ATP synthase protein 8 OS=Rattus norvegicus OX=10116 GN=ATP8 PE=3 SV=1 |
| **39** | 9290 | P04636\|MDHM_RAT | 137.22 | 19 | 19 | 3989000 | 5 | 5 | 6 | Y | 35684 | Malate dehydrogenase, mitochondrial OS=Rattus norvegicus OX=10116 GN=Mdh2 PE=1 SV=2 |
| **22** | 6960 | P29419\|ATP5I_RAT | 136 | 69 | 69 | 80884000 | 12 | 11 | 16 | N | 8255 | ATP synthase subunit e, mitochondrial OS=Rattus norvegicus OX=10116 GN=Atp5me PE=1 SV=3 |
| **40** | 4270 | Q9JJW3\|ATPMD_RAT | 134.56 | 45 | 45 | 56515000 | 3 | 3 | 6 | N | 6408 | ATP synthase membrane subunit DAPIT, mitochondrial OS=Rattus norvegicus OX=10116 GN=Atp5md PE=1 SV=1 |
| **25** | 4266 | D3ZAF6\|ATPK_RAT | 128.8 | 56 | 56 | 78115000 | 10 | 9 | 13 | Y | 10452 | ATP synthase subunit f, mitochondrial OS=Rattus norvegicus OX=10116 GN=Atp5mf PE=1 SV=1 |
| **35** | 4857 | P32198\|CPT1A_RAT | 127.43 | 10 | 10 | 3293100 | 5 | 5 | 6 | Y | 88126 | Carnitine O-palmitoyltransferase 1, liver isoform OS=Rattus norvegicus OX=10116 GN=Cpt1a PE=1 SV=2 |
| **41** | 278 | tr\|D3ZFJ6\|D3ZFJ6_RAT | 126.49 | 15 | 15 | 7551900 | 4 | 4 | 5 | N | 60420 | Lactamase, beta OS=Rattus norvegicus OX=10116 GN=Lactb PE=1 SV=1 |
| **47** | 106 | tr\|G3V6I4\|G3V6I4_RAT | 124.05 | 12 | 12 | 4104600 | 2 | 2 | 4 | N | 37791 | Mitochondrial amidoxime reducing component 1 OS=Rattus norvegicus OX=10116 GN=Marc1 PE=1 SV=1 |
| **57** | 4472 | P45953\|ACADV_RAT | 123.85 | 6 | 6 | 1443600 | 3 | 3 | 3 | N | 70749 | Very long-chain specific acyl-CoA dehydrogenase, mitochondrial OS=Rattus norvegicus OX=10116 GN=Acadvl PE=1 SV=1 |
| **57** | 4473 | tr\|Q5M9H2\|Q5M9H2_RAT | 123.85 | 6 | 6 | 1443600 | 3 | 3 | 3 | N | 70821 | Acyl-Coenzyme A dehydrogenase, very long chain OS=Rattus norvegicus OX=10116 GN=Acadvl PE=1 SV=1 |
| **45** | 98 | P29147\|BDH_RAT | 122.39 | 13 | 13 | 4000100 | 4 | 4 | 4 | N | 38202 | D-beta-hydroxybutyrate dehydrogenase, mitochondrial OS=Rattus norvegicus OX=10116 GN=Bdh1 PE=1 SV=2 |
| **45** | 99 | tr\|A0A0G2JSH2\|A0A0G2JSH2_R | 122.39 | 13 | 13 | 4000100 | 4 | 4 | 4 | N | 38333 | 3-hydroxybutyrate dehydrogenase, type 1, isoform CRA_a OS=Rattus norvegicus OX=10116 GN=Bdh1 PE=1 SV=1 |
| **66** | 13899 | Q5XIC2\|ECSIT_RAT | 118.41 | 8 | 8 | 991930 | 2 | 2 | 2 | Y | 49620 | Evolutionarily conserved signaling intermediate in Toll pathway, mitochondrial OS=Rattus norvegicus OX=10116 GN=Ecsit PE=1 SV=1 |
| **59** | 9321 | tr\|Q5EBA4\|Q5EBA4_RAT | 115.77 | 7 | 7 | 1666300 | 3 | 3 | 3 | N | 33215 | Nipsnap1 protein (Fragment) OS=Rattus norvegicus OX=10116 GN=Nipsnap1 PE=2 SV=1 |
| **59** | 9322 | tr\|G3V728\|G3V728_RAT | 115.77 | 7 | 7 | 1666300 | 3 | 3 | 3 | N | 33346 | 4-nitrophenylphosphatase domain and non-neuronal SNAP25-like protein homolog 1 (C. elegans), isoform CRA_b OS=Rattus norvegicus OX=10116 GN=Nipsnap1 PE=1 SV=1 |

| | | | | | | | | | | | | |
|---|---|---|---|---|---|---|---|---|---|---|---|---|
| **71** | 37 | P20788\|UCRI_RAT | 115.38 | 18 | 18 | 1933300 | 2 | 2 | 2 | N | 29446 | Cytochrome b-c1 complex subunit Rieske, mitochondrial OS=Rattus norvegicus OX=10116 GN=Uqcrfs1 PE=1 SV=2 |
| **43** | 110 | Q9WVK7\|HCDH_RAT | 113.73 | 16 | 16 | 1933000 | 3 | 3 | 4 | N | 34448 | Hydroxyacyl-coenzyme A dehydrogenase, mitochondrial OS=Rattus norvegicus OX=10116 GN=Hadh PE=2 SV=1 |
| **73** | 93 | tr\|A0A0G2K8Q8\|A0A0G2K8Q8_ | 110.17 | 39 | 39 | 1652700 | 2 | 2 | 2 | N | 7099 | Ubiquinol-cytochrome c reductase, complex III subunit X OS=Rattus norvegicus OX=10116 GN=Uqcr10 PE=1 SV=1 |
| **73** | 94 | tr\|B2RYX1\|B2RYX1_RAT | 110.17 | 38 | 38 | 1652700 | 2 | 2 | 2 | N | 7462 | LOC685322 protein OS=Rattus norvegicus OX=10116 GN=Uqcr10 PE=2 SV=1 |
| **54** | 2174 | O70351\|HCD2_RAT | 109.85 | 22 | 22 | 4123800 | 3 | 3 | 3 | N | 27246 | 3-hydroxyacyl-CoA dehydrogenase type-2 OS=Rattus norvegicus OX=10116 GN=Hsd17b10 PE=1 SV=3 |
| **54** | 2175 | tr\|B0BMW2\|B0BMW2_RAT | 109.85 | 22 | 22 | 4123800 | 3 | 3 | 3 | N | 27250 | 3-hydroxyacyl-CoA dehydrogenase type-2 OS=Rattus norvegicus OX=10116 GN=Hsd17b10 PE=1 SV=1 |
| **63** | 2188 | Q06646\|AT5G2_RAT | 109.43 | 27 | 27 | 10200000 | 2 | 2 | 2 | N | 14918 | ATP synthase F(0) complex subunit C2, mitochondrial OS=Rattus norvegicus OX=10116 GN=Atp5mc2 PE=1 SV=1 |
| **63** | 2187 | Q06645\|AT5G1_RAT | 109.43 | 28 | 28 | 10200000 | 2 | 2 | 2 | N | 14244 | ATP synthase F(0) complex subunit C1, mitochondrial OS=Rattus norvegicus OX=10116 GN=Atp5mc1 PE=1 SV=1 |
| **63** | 2189 | tr\|Q499S2\|Q499S2_RAT | 109.43 | 27 | 27 | 10200000 | 2 | 2 | 2 | N | 14745 | ATP synthase F(0) complex subunit C3, mitochondrial OS=Rattus norvegicus OX=10116 GN=Atp5mc3 PE=2 SV=1 |
| **63** | 2190 | Q71S46\|AT5G3_RAT | 109.43 | 27 | 27 | 10200000 | 2 | 2 | 2 | N | 14693 | ATP synthase F(0) complex subunit C3, mitochondrial OS=Rattus norvegicus OX=10116 GN=Atp5mc3 PE=1 SV=1 |
| **60** | 109 | P11240\|COX5A_RAT | 107.55 | 30 | 30 | 8579600 | 3 | 2 | 3 | N | 16130 | Cytochrome c oxidase subunit 5A, mitochondrial OS=Rattus norvegicus OX=10116 GN=Cox5a PE=1 SV=1 |
| **49** | 13895 | B1WC61\|ACAD9_RAT | 102.09 | 6 | 6 | 867700 | 2 | 2 | 3 | N | 68843 | Complex I assembly factor ACAD9, mitochondrial OS=Rattus norvegicus OX=10116 GN=Acad9 PE=1 SV=1 |
| **38** | 4273 | P29418\|ATP5E_RAT | 101.81 | 69 | 69 | 38088000 | 5 | 5 | 6 | Y | 5767 | ATP synthase subunit epsilon, mitochondrial OS=Rattus norvegicus OX=10116 GN=Atp5f1e PE=1 SV=2 |
| **56** | 13890 | P05504\|ATP6_RAT | 99.6 | 18 | 18 | 4736200 | 3 | 3 | 3 | Y | 25076 | ATP synthase subunit a OS=Rattus norvegicus OX=10116 GN=Mt-atp6 PE=1 SV=3 |
| **56** | 13891 | tr\|Q8HIC7\|Q8HIC7_RAT | 99.6 | 18 | 18 | 4736200 | 3 | 3 | 3 | Y | 25076 | ATP synthase subunit a OS=Rattus norvegicus OX=10116 GN=Mt-atp6 PE=4 SV=1 |
| **56** | 13892 | tr\|S5S1E9\|S5S1E9_RAT | 99.6 | 18 | 18 | 4736200 | 3 | 3 | 3 | Y | 25049 | ATP synthase subunit a OS=Rattus norvegicus OX=10116 GN=ATP6 PE=4 SV=1 |
| **56** | 13893 | tr\|Q06QE5\|Q06QE5_RAT | 99.6 | 18 | 18 | 4736200 | 3 | 3 | 3 | Y | 25075 | ATP synthase subunit a OS=Rattus norvegicus OX=10116 GN=ATP6 PE=4 SV=1 |
| **56** | 13894 | tr\|Q8SEZ3\|Q8SEZ3_RAT | 99.6 | 18 | 18 | 4736200 | 3 | 3 | 3 | Y | 25030 | ATP synthase subunit a OS=Rattus norvegicus OX=10116 GN=ATPase6 PE=4 SV=1 |
| **50** | 603 | A0A0G2K047\|ACSS3_RAT | 98.87 | 3 | 3 | 1012600 | 2 | 2 | 3 | N | 74675 | Acyl-CoA synthetase short-chain family member 3, mitochondrial OS=Rattus norvegicus OX=10116 GN=Acss3 PE=1 SV=1 |
| **102** | 178 | P62909\|RS3_RAT | 96.06 | 11 | 11 | 1123600 | 1 | 1 | 1 | N | 26674 | 40S ribosomal protein S3 OS=Rattus norvegicus OX=10116 GN=Rps3 PE=1 SV=1 |
| **51** | 11723 | tr\|G3V734\|G3V734_RAT | 95.53 | 8 | 8 | 1212000 | 2 | 1 | 3 | N | 36133 | 2,4-dienoyl CoA reductase 1, mitochondrial, isoform CRA_a OS=Rattus norvegicus OX=10116 GN=Decr1 PE=1 SV=1 |
| **51** | 11724 | Q64591\|DECR_RAT | 95.53 | 8 | 8 | 1212000 | 2 | 1 | 3 | N | 36133 | 2,4-dienoyl-CoA reductase, mitochondrial OS=Rattus norvegicus OX=10116 GN=Decr1 PE=1 SV=2 |
| **64** | 11616 | P10818\|CX6A1_RAT | 93.76 | 27 | 27 | 1546100 | 2 | 2 | 2 | N | 12301 | Cytochrome c oxidase subunit 6A1, mitochondrial OS=Rattus norvegicus OX=10116 GN=Cox6a1 PE=1 SV=2 |
| **77** | 2167 | Q5M9I5\|QCR6_RAT | 90.49 | 21 | 21 | 952880 | 1 | 1 | 2 | N | 10424 | Cytochrome b-c1 complex subunit 6, mitochondrial OS=Rattus norvegicus OX=10116 GN=Uqcrh PE=3 SV=1 |
| **58** | 158 | P12075\|COX5B_RAT | 88.52 | 29 | 29 | 6944000 | 3 | 3 | 3 | N | 13915 | Cytochrome c oxidase subunit 5B, mitochondrial OS=Rattus norvegicus OX=10116 GN=Cox5b PE=1 SV=2 |
| **46** | 47 | Q9WVK3\|PECR_RAT | 88.14 | 9 | 9 | 1768600 | 3 | 3 | 4 | N | 32433 | Peroxisomal trans-2-enoyl-CoA reductase OS=Rattus norvegicus OX=10116 GN=Pecr PE=2 SV=1 |
| **46** | 48 | tr\|A0A0G2JVG4\|A0A0G2JVG4_R | 88.14 | 9 | 9 | 1768600 | 3 | 3 | 4 | N | 32737 | Peroxisomal trans-2-enoyl-CoA reductase OS=Rattus norvegicus OX=10116 GN=Pecr PE=1 SV=1 |
| **125** | 13900 | P19804\|NDKB_RAT | 84.02 | 9 | 9 | 933500 | 1 | 1 | 1 | N | 17283 | Nucleoside diphosphate kinase B OS=Rattus norvegicus OX=10116 GN=Nme2 PE=1 SV=1 |

| | | | | | | | | | | | | |
|---|---|---|---|---|---|---|---|---|---|---|---|---|
| **125** | 13901 | Q05982\|NDKA_RAT | 84.02 | 9 | 9 | 933500 | 1 | 1 | 1 | N | 17193 | Nucleoside diphosphate kinase A OS=Rattus norvegicus OX=10116 GN=Nme1 PE=1 SV=1 |
| **126** | 11610 | tr\|Q7H115\|Q7H115_RAT | 82.94 | 5 | 5 | 911840 | 1 | 1 | 1 | N | 29871 | Cytochrome c oxidase subunit 3 OS=Rattus norvegicus OX=10116 GN=Mt-co3 PE=3 SV=1 |
| **126** | 11611 | tr\|I6V4L9\|I6V4L9_RAT | 82.94 | 5 | 5 | 911840 | 1 | 1 | 1 | N | 29844 | Cytochrome c oxidase subunit 3 OS=Rattus norvegicus OX=10116 GN=COX3 PE=3 SV=1 |
| **126** | 11612 | tr\|Q8M7G4\|Q8M7G4_RAT | 82.94 | 5 | 5 | 911840 | 1 | 1 | 1 | N | 29861 | Cytochrome c oxidase subunit 3 OS=Rattus norvegicus OX=10116 PE=2 SV=1 |
| **126** | 11613 | tr\|A0A096XKT2\|A0A096XKT2_R/ | 82.94 | 5 | 5 | 911840 | 1 | 1 | 1 | N | 29901 | Cytochrome c oxidase subunit 3 OS=Rattus norvegicus OX=10116 GN=COX3 PE=3 SV=1 |
| **126** | 11614 | P05505\|COX3_RAT | 82.94 | 5 | 5 | 911840 | 1 | 1 | 1 | N | 29871 | Cytochrome c oxidase subunit 3 OS=Rattus norvegicus OX=10116 GN=Mtco3 PE=1 SV=5 |
| **126** | 11615 | tr\|Q8SEZ2\|Q8SEZ2_RAT | 82.94 | 5 | 5 | 911840 | 1 | 1 | 1 | N | 29870 | Cytochrome c oxidase subunit 3 (Fragment) OS=Rattus norvegicus OX=10116 GN=COIII PE=3 SV=1 |
| **44** | 56 | tr\|S5RZM8\|S5RZM8_RAT | 76.74 | 11 | 11 | 4079900 | 3 | 3 | 4 | Y | 25958 | Cytochrome c oxidase subunit 2 OS=Rattus norvegicus OX=10116 GN=COX2 PE=3 SV=1 |
| **44** | 57 | P00406\|COX2_RAT | 76.74 | 11 | 11 | 4079900 | 3 | 3 | 4 | Y | 25928 | Cytochrome c oxidase subunit 2 OS=Rattus norvegicus OX=10116 GN=Mtco2 PE=1 SV=3 |
| **44** | 59 | tr\|Q5UAJ6\|Q5UAJ6_RAT | 76.74 | 11 | 11 | 4079900 | 3 | 3 | 4 | Y | 25942 | Cytochrome c oxidase subunit 2 OS=Rattus norvegicus OX=10116 GN=COX2 PE=3 SV=1 |
| **44** | 58 | tr\|Q37652\|Q37652_RAT | 76.74 | 11 | 11 | 4079900 | 3 | 3 | 4 | Y | 26007 | Cytochrome c oxidase subunit 2 OS=Rattus norvegicus OX=10116 GN=COXII PE=3 SV=1 |
| **44** | 60 | tr\|Q8SEZ5\|Q8SEZ5_RAT | 76.74 | 11 | 11 | 4079900 | 3 | 3 | 4 | Y | 25928 | Cytochrome c oxidase subunit 2 OS=Rattus norvegicus OX=10116 GN=Mt-co2 PE=1 SV=1 |
| **44** | 61 | tr\|A0A097PE04\|A0A097PE04_R/ | 76.74 | 11 | 11 | 4079900 | 3 | 3 | 4 | Y | 25894 | Cytochrome c oxidase subunit 2 OS=Rattus norvegicus OX=10116 GN=COX2 PE=3 SV=1 |
| **67** | 6851 | Q5PQT3\|GLYAT_RAT | 75.82 | 5 | 5 | 516050 | 1 | 1 | 2 | N | 33899 | Glycine N-acyltransferase OS=Rattus norvegicus OX=10116 GN=Glyat PE=2 SV=1 |
| **67** | 6852 | tr\|A4PB92\|A4PB92_RAT | 75.82 | 5 | 5 | 516050 | 1 | 1 | 2 | N | 33899 | Glycine N-acyltransferase OS=Rattus norvegicus OX=10116 GN=Glyat PE=2 SV=1 |
| **74** | 11618 | P11951\|CX6C2_RAT | 75.46 | 21 | 21 | 1308400 | 2 | 2 | 2 | N | 8455 | Cytochrome c oxidase subunit 6C-2 OS=Rattus norvegicus OX=10116 GN=Cox6c2 PE=1 SV=3 |
| **72** | 30 | Q7TQ16\|QCR8_RAT | 75.25 | 23 | 23 | 1354300 | 2 | 2 | 2 | N | 9849 | Cytochrome b-c1 complex subunit 8 OS=Rattus norvegicus OX=10116 GN=Uqcrq PE=3 SV=1 |
| **127** | 6797 | P08011\|MGST1_RAT | 73.01 | 10 | 10 | 408280 | 1 | 1 | 1 | N | 17472 | Microsomal glutathione S-transferase 1 OS=Rattus norvegicus OX=10116 GN=Mgst1 PE=1 SV=3 |
| **127** | 6798 | tr\|B6DYQ4\|B6DYQ4_RAT | 73.01 | 10 | 10 | 408280 | 1 | 1 | 1 | N | 17472 | Microsomal glutathione S-transferase OS=Rattus norvegicus OX=10116 GN=Mgst1 PE=2 SV=1 |
| **128** | 13902 | P70619\|GSHR_RAT | 73.01 | 5 | 5 | 302750 | 1 | 1 | 1 | N | 46301 | Glutathione reductase (Fragment) OS=Rattus norvegicus OX=10116 GN=Gsr PE=2 SV=2 |
| **130** | 9340 | D4A7N1\|MIC25_RAT | 72.62 | 5 | 5 | 295080 | 1 | 1 | 1 | N | 29211 | MICOS complex subunit Mic25 OS=Rattus norvegicus OX=10116 GN=Chchd6 PE=1 SV=1 |
| **129** | 376 | tr\|D3ZXF9\|D3ZXF9_RAT | 69.73 | 9 | 9 | 290590 | 1 | 1 | 1 | N | 29441 | Mitochondrial ribosomal protein L12 OS=Rattus norvegicus OX=10116 GN=Mrpl12 PE=1 SV=1 |
| **131** | 2210 | Q8VID1\|DHRS4_RAT | 66.41 | 5 | 5 | 831730 | 1 | 1 | 1 | N | 29822 | Dehydrogenase/reductase SDR family member 4 OS=Rattus norvegicus OX=10116 GN=Dhrs4 PE=2 SV=2 |
| **65** | 760 | tr\|F1M953\|F1M953_RAT | 66.21 | 4 | 4 | 160410 | 2 | 2 | 2 | N | 73745 | Stress-70 protein, mitochondrial OS=Rattus norvegicus OX=10116 GN=Hspa9 PE=1 SV=1 |
| **65** | 761 | P48721\|GRP75_RAT | 66.21 | 4 | 4 | 160410 | 2 | 2 | 2 | N | 73858 | Stress-70 protein, mitochondrial OS=Rattus norvegicus OX=10116 GN=Hspa9 PE=1 SV=3 |
| **132** | 11617 | P80432\|COX7C_RAT | 65.38 | 14 | 14 | 252730 | 1 | 1 | 1 | N | 7375 | Cytochrome c oxidase subunit 7C, mitochondrial OS=Rattus norvegicus OX=10116 GN=Cox7c PE=1 SV=2 |
| **55** | 101 | tr\|B2RYT5\|B2RYT5_RAT | 64.98 | 24 | 24 | 2231100 | 3 | 3 | 3 | N | 12651 | Cox7a2l protein OS=Rattus norvegicus OX=10116 GN=Cox7a2l PE=2 SV=1 |
| **75** | 205 | tr\|D3Z900\|D3Z900_RAT | 64.17 | 5 | 5 | 1193300 | 2 | 2 | 2 | N | 38239 | Mitochondrial amidoxime reducing component 2 OS=Rattus norvegicus OX=10116 GN=Marc2 PE=1 SV=2 |

| | | | | | | | | | | | | |
|---|---|---|---|---|---|---|---|---|---|---|---|---|
| **75** | 206 | O88994\|MARC2_RAT | 64.17 | 5 | 5 | 1193300 | 2 | 2 | 2 | N | 38249 | Mitochondrial amidoxime reducing component 2 OS=Rattus norvegicus OX=10116 GN=Marc2 PE=2 SV=1 |
| **75** | 207 | tr\|M0R6N2\|M0R6N2_RAT | 64.17 | 5 | 5 | 1193300 | 2 | 2 | 2 | N | 38175 | MOSC domain-containing protein 2, mitochondrial-like OS=Rattus norvegicus OX=10116 GN=LOC100910481 PE=1 SV=1 |
| **133** | 4512 | Q5RJR8\|LRC59_RAT | 58.38 | 3 | 3 | 131730 | 1 | 1 | 1 | N | 34869 | Leucine-rich repeat-containing protein 59 OS=Rattus norvegicus OX=10116 GN=Lrrc59 PE=1 SV=1 |
| **101** | 7185 | Q924S5\|LONM_RAT | 58.08 | 4 | 4 | 3606500 | 1 | 1 | 1 | N | 105792 | Lon protease homolog, mitochondrial OS=Rattus norvegicus OX=10116 GN=Lonp1 PE=2 SV=1 |
| **134** | 13903 | P35171\|CX7A2_RAT | 53.43 | 10 | 10 | 364200 | 1 | 1 | 1 | N | 9353 | Cytochrome c oxidase subunit 7A2, mitochondrial OS=Rattus norvegicus OX=10116 GN=Cox7a2 PE=1 SV=1 |
| **134** | 13904 | tr\|B2RYS0\|B2RYS0_RAT | 53.43 | 10 | 10 | 364200 | 1 | 1 | 1 | N | 9353 | Cox7a2 protein OS=Rattus norvegicus OX=10116 GN=Cox7a2 PE=2 SV=1 |
| **62** | 504 | tr\|A0A0G2K5L6\|A0A0G2K5L6_R/ | 53.22 | 1 | 1 | 10507000 | 2 | 1 | 2 | Y | 275755 | Acetyl-CoA carboxylase beta OS=Rattus norvegicus OX=10116 GN=Acacb PE=1 SV=1 |
| **62** | 505 | tr\|A0A0G2K1F2\|A0A0G2K1F2_R/ | 53.22 | 1 | 1 | 10507000 | 2 | 1 | 2 | Y | 276393 | Acetyl-CoA carboxylase beta OS=Rattus norvegicus OX=10116 GN=Acacb PE=1 SV=1 |
| **62** | 390 | tr\|O70151\|O70151_RAT | 53.22 | 1 | 1 | 10507000 | 2 | 1 | 2 | Y | 276097 | Acetyl-CoA carboxylase OS=Rattus norvegicus OX=10116 GN=Acacb PE=2 SV=1 |
| **62** | 506 | tr\|D3ZBE2\|D3ZBE2_RAT | 53.22 | 1 | 1 | 10507000 | 2 | 1 | 2 | Y | 275967 | Acetyl-CoA carboxylase beta OS=Rattus norvegicus OX=10116 GN=Acacb PE=1 SV=3 |
| **62** | 507 | tr\|E9PSQ0\|E9PSQ0_RAT | 53.22 | 1 | 1 | 10507000 | 2 | 1 | 2 | Y | 276255 | Acetyl-CoA carboxylase beta OS=Rattus norvegicus OX=10116 GN=Acacb PE=1 SV=2 |
| **48** | 179 | F1M775\|DIAP1_RAT | 49.15 | 2 | 2 | 1275600 | 3 | 1 | 4 | Y | 140418 | Protein diaphanous homolog 1 OS=Rattus norvegicus OX=10116 GN=Diaph1 PE=1 SV=3 |
| **135** | 11619 | P80431\|COX7B_RAT | 47.35 | 9 | 9 | 447640 | 1 | 1 | 1 | N | 8995 | Cytochrome c oxidase subunit 7B, mitochondrial OS=Rattus norvegicus OX=10116 GN=Cox7b PE=1 SV=3 |
| **76** | 9291 | tr\|D3ZUX5\|D3ZUX5_RAT | 47.25 | 7 | 7 | 154490 | 2 | 2 | 2 | N | 26435 | MICOS complex subunit OS=Rattus norvegicus OX=10116 GN=Chchd3 PE=1 SV=1 |
| **52** | 4633 | P00481\|OTC_RAT | 40.49 | 10 | 10 | 1044200 | 1 | 1 | 3 | N | 39886 | Ornithine carbamoyltransferase, mitochondrial OS=Rattus norvegicus OX=10116 GN=Otc PE=1 SV=1 |
| **136** | 11623 | tr\|B2RZD6\|B2RZD6_RAT | 39.08 | 10 | 10 | 543290 | 1 | 1 | 1 | N | 9327 | NDUFA4, mitochondrial complex-associated OS=Rattus norvegicus OX=10116 GN=Ndufa4 PE=1 SV=1 |
| **139** | 13905 | tr\|C7EMI4\|C7EMI4_RAT | 37.25 | 4 | 4 | 741150 | 1 | 1 | 1 | N | 20146 | Cytochrome b (Fragment) OS=Rattus norvegicus OX=10116 PE=4 SV=1 |
| **139** | 13906 | tr\|U3M4V1\|U3M4V1_RAT | 37.25 | 4 | 4 | 741150 | 1 | 1 | 1 | N | 20795 | Cytochrome b (Fragment) OS=Rattus norvegicus OX=10116 GN=cyt b PE=4 SV=1 |
| **139** | 13907 | tr\|A4UHZ3\|A4UHZ3_RAT | 37.25 | 3 | 3 | 741150 | 1 | 1 | 1 | N | 25311 | Cytochrome b (Fragment) OS=Rattus norvegicus OX=10116 GN=cytb PE=4 SV=1 |
| **139** | 13908 | tr\|A4UHZ2\|A4UHZ2_RAT | 37.25 | 3 | 3 | 741150 | 1 | 1 | 1 | N | 25312 | Cytochrome b (Fragment) OS=Rattus norvegicus OX=10116 GN=cytb PE=4 SV=1 |
| **139** | 13909 | tr\|B3Y999\|B3Y999_RAT | 37.25 | 3 | 3 | 741150 | 1 | 1 | 1 | N | 28378 | Cytochrome b (Fragment) OS=Rattus norvegicus OX=10116 GN=cytb PE=4 SV=1 |
| **139** | 13910 | tr\|M9TEK6\|M9TEK6_RAT | 37.25 | 2 | 2 | 741150 | 1 | 1 | 1 | N | 35982 | Cytochrome b (Fragment) OS=Rattus norvegicus OX=10116 GN=cytb PE=3 SV=1 |
| **139** | 13911 | tr\|E0A0C7\|E0A0C7_RAT | 37.25 | 2 | 2 | 741150 | 1 | 1 | 1 | N | 41009 | Cytochrome b (Fragment) OS=Rattus norvegicus OX=10116 GN=cytb PE=3 SV=1 |
| **139** | 13912 | tr\|E0A016\|E0A016_RAT | 37.25 | 2 | 2 | 741150 | 1 | 1 | 1 | N | 41634 | Cytochrome b (Fragment) OS=Rattus norvegicus OX=10116 GN=cytb PE=3 SV=1 |
| **139** | 13913 | tr\|A0A411P2K0\|A0A411P2K0_R/ | 37.25 | 2 | 2 | 741150 | 1 | 1 | 1 | N | 42238 | Cytochrome b (Fragment) OS=Rattus norvegicus OX=10116 GN=Cytb PE=3 SV=1 |
| **139** | 13914 | tr\|E0A075\|E0A075_RAT | 37.25 | 2 | 2 | 741150 | 1 | 1 | 1 | N | 42410 | Cytochrome b (Fragment) OS=Rattus norvegicus OX=10116 GN=cytb PE=3 SV=1 |
| **139** | 13915 | tr\|L0L4L8\|L0L4L8_RAT | 37.25 | 2 | 2 | 741150 | 1 | 1 | 1 | N | 42470 | Cytochrome b (Fragment) OS=Rattus norvegicus OX=10116 GN=cytb PE=3 SV=1 |
| **139** | 13916 | tr\|A0A140GE09\|A0A140GE09_R/ | 37.25 | 2 | 2 | 741150 | 1 | 1 | 1 | N | 42569 | Cytochrome b (Fragment) OS=Rattus norvegicus OX=10116 PE=3 SV=1 |

| | | | | | | | | | | | | |
|---|---|---|---|---|---|---|---|---|---|---|---|---|
| **139** | 13917 | tr\|A0A140GE10\|A0A140GE10_R. | 37.25 | 2 | 2 | 741150 | 1 | 1 | 1 | N | 42574 | Cytochrome b (Fragment) OS=Rattus norvegicus OX=10116 PE=3 SV=1 |
| **139** | 13918 | tr\|A0A140GE11\|A0A140GE11_R. | 37.25 | 2 | 2 | 741150 | 1 | 1 | 1 | N | 42588 | Cytochrome b (Fragment) OS=Rattus norvegicus OX=10116 PE=3 SV=1 |
| **139** | 13919 | tr\|A0A385HC91\|A0A385HC91_R | 37.25 | 2 | 2 | 741150 | 1 | 1 | 1 | N | 42780 | Cytochrome b (Fragment) OS=Rattus norvegicus OX=10116 PE=3 SV=1 |
| **139** | 13920 | tr\|A0A385HCC9\|A0A385HCC9_R | 37.25 | 2 | 2 | 741150 | 1 | 1 | 1 | N | 42766 | Cytochrome b (Fragment) OS=Rattus norvegicus OX=10116 PE=3 SV=1 |
| **139** | 13921 | tr\|A0A3Q8AGE8\|A0A3Q8AGE8_I | 37.25 | 2 | 2 | 741150 | 1 | 1 | 1 | N | 42712 | Cytochrome b (Fragment) OS=Rattus norvegicus OX=10116 GN=Cytb PE=3 SV=1 |
| **139** | 13922 | tr\|A0A3Q8AC68\|A0A3Q8AC68_F | 37.25 | 2 | 2 | 741150 | 1 | 1 | 1 | N | 42622 | Cytochrome b (Fragment) OS=Rattus norvegicus OX=10116 GN=Cytb PE=3 SV=1 |
| **139** | 13923 | tr\|A0A3S6FK13\|A0A3S6FK13_RA | 37.25 | 2 | 2 | 741150 | 1 | 1 | 1 | N | 42702 | Cytochrome b (Fragment) OS=Rattus norvegicus OX=10116 GN=Cytb PE=3 SV=1 |
| **139** | 13924 | tr\|A0A385HCK1\|A0A385HCK1_R | 37.25 | 2 | 2 | 741150 | 1 | 1 | 1 | N | 42824 | Cytochrome b (Fragment) OS=Rattus norvegicus OX=10116 PE=3 SV=1 |
| **139** | 13925 | tr\|A0A3Q8AZT4\|A0A3Q8AZT4_R | 37.25 | 2 | 2 | 741150 | 1 | 1 | 1 | N | 42723 | Cytochrome b (Fragment) OS=Rattus norvegicus OX=10116 GN=Cytb PE=3 SV=1 |
| **139** | 13926 | tr\|A0A3S6FK10\|A0A3S6FK10_RA | 37.25 | 2 | 2 | 741150 | 1 | 1 | 1 | N | 42726 | Cytochrome b (Fragment) OS=Rattus norvegicus OX=10116 GN=Cytb PE=3 SV=1 |
| **139** | 13927 | tr\|A0A385HC93\|A0A385HC93_R | 37.25 | 2 | 2 | 741150 | 1 | 1 | 1 | N | 42751 | Cytochrome b (Fragment) OS=Rattus norvegicus OX=10116 PE=3 SV=1 |
| **139** | 13928 | tr\|A0A3S6FM15\|A0A3S6FM15_F | 37.25 | 2 | 2 | 741150 | 1 | 1 | 1 | N | 42698 | Cytochrome b (Fragment) OS=Rattus norvegicus OX=10116 GN=Cytb PE=3 SV=1 |
| **139** | 13929 | tr\|A0A097PE40\|A0A097PE40_R/ | 37.25 | 2 | 2 | 741150 | 1 | 1 | 1 | N | 42865 | Cytochrome b OS=Rattus norvegicus OX=10116 GN=CYTB PE=3 SV=1 |
| **139** | 13930 | tr\|A0A220D9Z6\|A0A220D9Z6_R/ | 37.25 | 2 | 2 | 741150 | 1 | 1 | 1 | N | 43018 | Cytochrome b OS=Rattus norvegicus OX=10116 PE=3 SV=1 |
| **139** | 13931 | tr\|D6NSP2\|D6NSP2_RAT | 37.25 | 2 | 2 | 741150 | 1 | 1 | 1 | N | 42945 | Cytochrome b (Fragment) OS=Rattus norvegicus OX=10116 GN=cytb PE=3 SV=1 |
| **139** | 13932 | tr\|D6NSS6\|D6NSS6_RAT | 37.25 | 2 | 2 | 741150 | 1 | 1 | 1 | N | 42993 | Cytochrome b (Fragment) OS=Rattus norvegicus OX=10116 GN=cytb PE=3 SV=1 |
| **139** | 13933 | tr\|Q06Q99\|Q06Q99_RAT | 37.25 | 2 | 2 | 741150 | 1 | 1 | 1 | N | 42997 | Cytochrome b OS=Rattus norvegicus OX=10116 GN=CYB PE=3 SV=1 |
| **139** | 13934 | tr\|L0N490\|L0N490_RAT | 37.25 | 2 | 2 | 741150 | 1 | 1 | 1 | N | 43056 | Cytochrome b (Fragment) OS=Rattus norvegicus OX=10116 GN=cytb PE=3 SV=1 |
| **139** | 13935 | tr\|D6NSQ3\|D6NSQ3_RAT | 37.25 | 2 | 2 | 741150 | 1 | 1 | 1 | N | 43012 | Cytochrome b (Fragment) OS=Rattus norvegicus OX=10116 GN=cytb PE=3 SV=1 |
| **139** | 13936 | tr\|D6NSR3\|D6NSR3_RAT | 37.25 | 2 | 2 | 741150 | 1 | 1 | 1 | N | 42986 | Cytochrome b (Fragment) OS=Rattus norvegicus OX=10116 GN=cytb PE=3 SV=1 |
| **139** | 13937 | tr\|D6NSP4\|D6NSP4_RAT | 37.25 | 2 | 2 | 741150 | 1 | 1 | 1 | N | 43016 | Cytochrome b (Fragment) OS=Rattus norvegicus OX=10116 GN=cytb PE=3 SV=1 |
| **139** | 13938 | tr\|D6NSQ8\|D6NSQ8_RAT | 37.25 | 2 | 2 | 741150 | 1 | 1 | 1 | N | 43016 | Cytochrome b (Fragment) OS=Rattus norvegicus OX=10116 GN=cytb PE=3 SV=1 |
| **139** | 13939 | tr\|D6NSR8\|D6NSR8_RAT | 37.25 | 2 | 2 | 741150 | 1 | 1 | 1 | N | 42998 | Cytochrome b (Fragment) OS=Rattus norvegicus OX=10116 GN=cytb PE=3 SV=1 |
| **139** | 13940 | tr\|F8QU37\|F8QU37_RAT | 37.25 | 2 | 2 | 741150 | 1 | 1 | 1 | N | 42992 | Cytochrome b (Fragment) OS=Rattus norvegicus OX=10116 GN=cytb PE=3 SV=1 |
| **139** | 13941 | tr\|F2Q6S6\|F2Q6S6_RAT | 37.25 | 2 | 2 | 741150 | 1 | 1 | 1 | N | 42938 | Cytochrome b (Fragment) OS=Rattus norvegicus OX=10116 GN=cytb PE=3 SV=1 |
| **139** | 13942 | tr\|A0A0A1FZ42\|A0A0A1FZ42_R/ | 37.25 | 2 | 2 | 741150 | 1 | 1 | 1 | N | 42989 | Cytochrome b OS=Rattus norvegicus OX=10116 GN=CYTB PE=3 SV=1 |
| **139** | 13943 | tr\|D6NSS7\|D6NSS7_RAT | 37.25 | 2 | 2 | 741150 | 1 | 1 | 1 | N | 42948 | Cytochrome b (Fragment) OS=Rattus norvegicus OX=10116 GN=cytb PE=3 SV=1 |
| **139** | 13944 | tr\|Q8SEY9\|Q8SEY9_RAT | 37.25 | 2 | 2 | 741150 | 1 | 1 | 1 | N | 43015 | Cytochrome b OS=Rattus norvegicus OX=10116 GN=cytb PE=3 SV=1 |

| | | | | | | | | | | | | |
|---|---|---|---|---|---|---|---|---|---|---|---|---|
| **139** | 13945 | tr\|D6NSR7\|D6NSR7_RAT | 37.25 | 2 | 2 | 741150 | 1 | 1 | 1 | N | 42982 | Cytochrome b (Fragment) OS=Rattus norvegicus OX=10116 GN=cytb PE=3 SV=1 |
| **139** | 13946 | tr\|A0A220DA44\|A0A220DA44_R | 37.25 | 2 | 2 | 741150 | 1 | 1 | 1 | N | 42968 | Cytochrome b OS=Rattus norvegicus OX=10116 PE=3 SV=1 |
| **139** | 13947 | tr\|D6NSP0\|D6NSP0_RAT | 37.25 | 2 | 2 | 741150 | 1 | 1 | 1 | N | 43023 | Cytochrome b (Fragment) OS=Rattus norvegicus OX=10116 GN=cytb PE=3 SV=1 |
| **139** | 13948 | tr\|D6NSQ6\|D6NSQ6_RAT | 37.25 | 2 | 2 | 741150 | 1 | 1 | 1 | N | 42952 | Cytochrome b (Fragment) OS=Rattus norvegicus OX=10116 GN=cytb PE=3 SV=1 |
| **139** | 13949 | tr\|L0N495\|L0N495_RAT | 37.25 | 2 | 2 | 741150 | 1 | 1 | 1 | N | 42992 | Cytochrome b (Fragment) OS=Rattus norvegicus OX=10116 GN=cytb PE=3 SV=1 |
| **139** | 13950 | tr\|D6NSQ0\|D6NSQ0_RAT | 37.25 | 2 | 2 | 741150 | 1 | 1 | 1 | N | 42968 | Cytochrome b (Fragment) OS=Rattus norvegicus OX=10116 GN=cytb PE=3 SV=1 |
| **139** | 13951 | tr\|Q5UAI7\|Q5UAI7_RAT | 37.25 | 2 | 2 | 741150 | 1 | 1 | 1 | N | 42998 | Cytochrome b OS=Rattus norvegicus OX=10116 GN=CYTB PE=3 SV=1 |
| **139** | 13952 | tr\|D6NSR2\|D6NSR2_RAT | 37.25 | 2 | 2 | 741150 | 1 | 1 | 1 | N | 43073 | Cytochrome b (Fragment) OS=Rattus norvegicus OX=10116 GN=cytb PE=3 SV=1 |
| **139** | 13953 | tr\|A0A0S1Z1V9\|A0A0S1Z1V9_RA | 37.25 | 2 | 2 | 741150 | 1 | 1 | 1 | N | 43002 | Cytochrome b OS=Rattus norvegicus OX=10116 GN=CYTB PE=3 SV=1 |
| **139** | 13954 | tr\|B0M1Q8\|B0M1Q8_RAT | 37.25 | 2 | 2 | 741150 | 1 | 1 | 1 | N | 43026 | Cytochrome b (Fragment) OS=Rattus norvegicus OX=10116 GN=cytb PE=3 SV=1 |
| **139** | 13955 | tr\|D6NSR9\|D6NSR9_RAT | 37.25 | 2 | 2 | 741150 | 1 | 1 | 1 | N | 42970 | Cytochrome b (Fragment) OS=Rattus norvegicus OX=10116 GN=cytb PE=3 SV=1 |
| **139** | 13956 | tr\|D6NSR1\|D6NSR1_RAT | 37.25 | 2 | 2 | 741150 | 1 | 1 | 1 | N | 43002 | Cytochrome b (Fragment) OS=Rattus norvegicus OX=10116 GN=cytb PE=3 SV=1 |
| **139** | 13957 | tr\|Q8HIC4\|Q8HIC4_RAT | 37.25 | 2 | 2 | 741150 | 1 | 1 | 1 | N | 43012 | Cytochrome b OS=Rattus norvegicus OX=10116 GN=Mt-cyb PE=3 SV=1 |
| **139** | 13958 | tr\|A0A220DA02\|A0A220DA02_R | 37.25 | 2 | 2 | 741150 | 1 | 1 | 1 | N | 42968 | Cytochrome b OS=Rattus norvegicus OX=10116 PE=3 SV=1 |
| **139** | 13959 | tr\|S5RKC8\|S5RKC8_RAT | 37.25 | 2 | 2 | 741150 | 1 | 1 | 1 | N | 42997 | Cytochrome b OS=Rattus norvegicus OX=10116 GN=CYTB PE=3 SV=1 |
| **139** | 13960 | tr\|A0A0S1Z1V4\|A0A0S1Z1V4_RA | 37.25 | 2 | 2 | 741150 | 1 | 1 | 1 | N | 43016 | Cytochrome b OS=Rattus norvegicus OX=10116 GN=CYTB PE=3 SV=1 |
| **139** | 13961 | tr\|R9TKN1\|R9TKN1_RAT | 37.25 | 2 | 2 | 741150 | 1 | 1 | 1 | N | 43012 | Cytochrome b OS=Rattus norvegicus OX=10116 GN=CYTB PE=3 SV=1 |
| **139** | 13962 | P00159\|CYB_RAT | 37.25 | 2 | 2 | 741150 | 1 | 1 | 1 | N | 43012 | Cytochrome b OS=Rattus norvegicus OX=10116 GN=Mt-Cyb PE=3 SV=3 |
| **139** | 13963 | tr\|D6NSP8\|D6NSP8_RAT | 37.25 | 2 | 2 | 741150 | 1 | 1 | 1 | N | 43100 | Cytochrome b (Fragment) OS=Rattus norvegicus OX=10116 GN=cytb PE=3 SV=1 |
| **139** | 13964 | tr\|L0N311\|L0N311_RAT | 37.25 | 2 | 2 | 741150 | 1 | 1 | 1 | N | 43045 | Cytochrome b (Fragment) OS=Rattus norvegicus OX=10116 GN=cytb PE=3 SV=1 |
| **139** | 13965 | tr\|A0A220DA20\|A0A220DA20_R | 37.25 | 2 | 2 | 741150 | 1 | 1 | 1 | N | 43010 | Cytochrome b OS=Rattus norvegicus OX=10116 PE=3 SV=1 |
| **139** | 13966 | tr\|A0A220D9W5\|A0A220D9W5_ | 37.25 | 2 | 2 | 741150 | 1 | 1 | 1 | N | 43028 | Cytochrome b OS=Rattus norvegicus OX=10116 GN=cyt-b PE=3 SV=1 |
| **139** | 13967 | tr\|A0A220DA28\|A0A220DA28_R | 37.25 | 2 | 2 | 741150 | 1 | 1 | 1 | N | 42970 | Cytochrome b OS=Rattus norvegicus OX=10116 PE=3 SV=1 |
| **139** | 13968 | tr\|H6RXT5\|H6RXT5_RAT | 37.25 | 2 | 2 | 741150 | 1 | 1 | 1 | N | 43044 | Cytochrome b OS=Rattus norvegicus OX=10116 GN=cytb PE=3 SV=1 |
| **139** | 13969 | tr\|H2KXA0\|H2KXA0_RAT | 37.25 | 2 | 2 | 741150 | 1 | 1 | 1 | N | 42978 | Cytochrome b (Fragment) OS=Rattus norvegicus OX=10116 GN=cytb PE=3 SV=1 |
| **139** | 13970 | tr\|L0N1V8\|L0N1V8_RAT | 37.25 | 2 | 2 | 741150 | 1 | 1 | 1 | N | 43058 | Cytochrome b (Fragment) OS=Rattus norvegicus OX=10116 GN=cytb PE=3 SV=1 |
| **139** | 13971 | tr\|A0A096XNM4\|A0A096XNM4_ | 37.25 | 2 | 2 | 741150 | 1 | 1 | 1 | N | 43042 | Cytochrome b (Fragment) OS=Rattus norvegicus OX=10116 PE=3 SV=1 |
| **139** | 13972 | tr\|D6NSP7\|D6NSP7_RAT | 37.25 | 2 | 2 | 741150 | 1 | 1 | 1 | N | 43046 | Cytochrome b (Fragment) OS=Rattus norvegicus OX=10116 GN=cytb PE=3 SV=1 |

| | | | | | | | | | | | | |
|---|---|---|---|---|---|---|---|---|---|---|---|---|
| **139** | 13973 | tr\|A0A220D9Z3\|A0A220D9Z3_RA | 37.25 | 2 | 2 | 741150 | 1 | 1 | 1 | N | 43024 | Cytochrome b OS=Rattus norvegicus OX=10116 PE=3 SV=1 |
| **139** | 13974 | tr\|D6NSP5\|D6NSP5_RAT | 37.25 | 2 | 2 | 741150 | 1 | 1 | 1 | N | 43012 | Cytochrome b (Fragment) OS=Rattus norvegicus OX=10116 GN=cytb PE=3 SV=1 |
| **139** | 13975 | tr\|D6NSP6\|D6NSP6_RAT | 37.25 | 2 | 2 | 741150 | 1 | 1 | 1 | N | 43050 | Cytochrome b (Fragment) OS=Rattus norvegicus OX=10116 GN=cytb PE=3 SV=1 |
| **139** | 13976 | tr\|Q9G898\|Q9G898_RAT | 37.25 | 2 | 2 | 741150 | 1 | 1 | 1 | N | 42830 | Cytochrome b OS=Rattus norvegicus OX=10116 PE=2 SV=1 |
| **137** | 622 | P18163\|ACSL1_RAT | 34.62 | 2 | 2 | 1753800 | 1 | 1 | 1 | N | 78179 | Long-chain-fatty-acid--CoA ligase 1 OS=Rattus norvegicus OX=10116 GN=Acsl1 PE=1 SV=1 |
| **78** | 181 | P61980\|HNRPK_RAT | 33.22 | 3 | 3 | 1064400 | 2 | 1 | 2 | N | 50976 | Heterogeneous nuclear ribonucleoprotein K OS=Rattus norvegicus OX=10116 GN=Hnrnpk PE=1 SV=1 |
| **79** | 2538 | D3ZG52\|DNA2_RAT | 31.94 | 1 | 1 | | 2 | 0 | 2 | N | 119588 | DNA replication ATP-dependent helicase/nuclease DNA2 OS=Rattus norvegicus OX=10116 GN=Dna2 PE=3 SV=1 |
| **79** | 2550 | tr\|A0A0H2UH92\|A0A0H2UH92_ | 31.94 | 1 | 1 | | 2 | 0 | 2 | N | 136526 | Graves disease carrier protein OS=Rattus norvegicus OX=10116 GN=Slc25a16 PE=1 SV=1 |
| **103** | 6940 | P06761\|BIP_RAT | 30.93 | 1 | 1 | 0 | 1 | 1 | 1 | N | 72347 | Endoplasmic reticulum chaperone BiP OS=Rattus norvegicus OX=10116 GN=Hspa5 PE=1 SV=1 |
| **140** | 4278 | tr\|A0A0U1RRQ6\|A0A0U1RRQ6_ | 29.99 | 3 | 3 | 1315400 | 1 | 1 | 1 | Y | 35344 | Solute carrier family 25, member 44 OS=Rattus norvegicus OX=10116 GN=Slc25a44 PE=1 SV=1 |
| **140** | 4279 | tr\|D3ZSN7\|D3ZSN7_RAT | 29.99 | 2 | 2 | 1315400 | 1 | 1 | 1 | Y | 37879 | Similar to CG5805-PA (Predicted), isoform CRA_a OS=Rattus norvegicus OX=10116 GN=Slc25a44 PE=1 SV=1 |
| **153** | 140 | tr\|F1LPI6\|F1LPI6_RAT | 26.62 | 0 | 0 | 555250 | 1 | 1 | 1 | N | 200075 | Peripheral-type benzodiazepine receptor-associated protein 1 OS=Rattus norvegicus OX=10116 GN=Tspoap1 PE=4 SV=1 |
| **153** | 141 | Q9JIR0\|RIMB1_RAT | 26.62 | 0 | 0 | 555250 | 1 | 1 | 1 | N | 200202 | Peripheral-type benzodiazepine receptor-associated protein 1 OS=Rattus norvegicus OX=10116 GN=Tspoap1 PE=1 SV=2 |
| **69** | 2194 | tr\|Q5U2T0\|Q5U2T0_RAT | 23.77 | 2 | 2 | 0 | 1 | 1 | 2 | N | 44494 | Death associated protein 3 OS=Rattus norvegicus OX=10116 GN=Dap3 PE=2 SV=1 |
| **69** | 2195 | tr\|F7EZZ0\|F7EZZ0_RAT | 23.77 | 2 | 2 | 0 | 1 | 1 | 2 | N | 45112 | Death-associated protein 3 OS=Rattus norvegicus OX=10116 GN=Dap3 PE=1 SV=1 |
| **69** | 256 | tr\|A0A0G2K264\|A0A0G2K264_R | 23.77 | 2 | 2 | 0 | 1 | 1 | 2 | N | 46666 | Death-associated protein 3 OS=Rattus norvegicus OX=10116 GN=Dap3 PE=1 SV=1 |
| **81** | 4406 | tr\|A0A0G2JYD4\|A0A0G2JYD4_RA | 23.58 | 0 | 0 | 2622700 | 1 | 1 | 1 | N | 487857 | Vacuolar protein sorting 13 homolog D OS=Rattus norvegicus OX=10116 GN=Vps13d PE=1 SV=1 |
| **81** | 4407 | tr\|D3ZKC6\|D3ZKC6_RAT | 23.58 | 0 | 0 | 2622700 | 1 | 1 | 1 | N | 488962 | Vacuolar protein sorting 13 homolog D OS=Rattus norvegicus OX=10116 GN=Vps13d PE=1 SV=1 |
| **105** | 9327 | O54902\|NRAM2_RAT | 22.9 | 2 | 2 | 1733600 | 1 | 1 | 1 | N | 62277 | Natural resistance-associated macrophage protein 2 OS=Rattus norvegicus OX=10116 GN=Slc11a2 PE=1 SV=1 |
| **148** | 6827 | Q9Z0U4\|GABR1_RAT | 22.26 | 1 | 1 | 451400 | 1 | 1 | 1 | N | 111534 | Gamma-aminobutyric acid type B receptor subunit 1 OS=Rattus norvegicus OX=10116 GN=Gabbr1 PE=1 SV=1 |
| **93** | 4527 | Q5XHZ0\|TRAP1_RAT | 43973 | 4 | 4 | 430990 | 1 | 1 | 1 | N | 80461 | Heat shock protein 75 kDa, mitochondrial OS=Rattus norvegicus OX=10116 GN=Trap1 PE=1 SV=1 |
| **96** | 4666 | tr\|D3ZDM5\|D3ZDM5_RAT | 21.54 | 1 | 1 | | 1 | 0 | 1 | N | 99970 | NACHT, leucine rich repeat and PYD containing 5 (Predicted) OS=Rattus norvegicus OX=10116 GN=Nlrp5 PE=4 SV=1 |
| **150** | 6949 | Q66HA8\|HS105_RAT | 20.97 | 1 | 1 | 467670 | 1 | 1 | 1 | Y | 96419 | Heat shock protein 105 kDa OS=Rattus norvegicus OX=10116 GN=Hsph1 PE=1 SV=1 |
| **109** | 14004 | P07308\|ACOD1_RAT | 20.9 | 1 | 1 | 299660 | 1 | 1 | 1 | N | 41467 | Acyl-CoA desaturase 1 OS=Rattus norvegicus OX=10116 GN=Scd1 PE=1 SV=2 |
| **109** | 96 | P08461\|ODP2_RAT | 20.9 | 1 | 1 | 299660 | 1 | 1 | 1 | N | 67166 | Dihydrolipoyllysine-residue acetyltransferase component of pyruvate dehydrogenase complex, mitochondrial OS=Rattus norvegicus OX=10116 GN=Dlat PE=1 SV=3 |
| **109** | 7046 | O35303\|DNM1L_RAT | 20.9 | 1 | 1 | 299660 | 1 | 1 | 1 | N | 83908 | Dynamin-1-like protein OS=Rattus norvegicus OX=10116 GN=Dnm1l PE=1 SV=1 |
| **152** | 2204 | Q63484\|AKT3_RAT | 20.54 | 2 | 2 | 264270 | 1 | 1 | 1 | Y | 55797 | RAC-gamma serine/threonine-protein kinase OS=Rattus norvegicus OX=10116 GN=Akt3 PE=2 SV=2 |
| **total 222 proteins** | | | | | | | | | | | | |
