## Supplementary material for "Permeability transition pore-related changes in the proteome and channel activity of ATP synthase dimers and monomers": RLM KCl-BM PTP Dimer of V compl.

| Protein Group | Protein ID | Accession | -10lgP | Coverage (%) | Coverage (%) Sample 2 | Area Sample 2 | #Peptides | #Unique | #Spec Sample 2 | PTM | Avg. Mass | Description |
|---|---|---|---|---|---|---|---|---|---|---|---|---|
| **2** | 13 | P15999\|ATPA_RAT | 460.42 | 70 | 70 | 1415100000 | 75 | 75 | 245 | Y | 59754 | ATP synthase subunit alpha, mitochondrial OS=Rattus norvegicus OX=10116 GN=Atp5f1a PE=1 SV=2 |
| **1** | 8 | P10719\|ATPB_RAT | 449.64 | 81 | 81 | 1687800000 | 64 | 64 | 308 | Y | 56354 | ATP synthase subunit beta, mitochondrial OS=Rattus norvegicus OX=10116 GN=Atp5f1b PE=1 SV=2 |
| **1** | 9 | tr\|G3V6D3\|G3V6D3_RAT | 449.64 | 81 | 81 | 1687800000 | 64 | 64 | 308 | Y | 56345 | ATP synthase subunit beta OS=Rattus norvegicus OX=10116 GN=Atp5f1b PE=1 SV=1 |
| **6** | 3 | P07756\|CPSM_RAT | 355.54 | 32 | 32 | 106500000 | 37 | 37 | 47 | Y | 164579 | Carbamoyl-phosphate synthase [ammonia], mitochondrial OS=Rattus norvegicus OX=10116 GN=Cps1 PE=1 SV=1 |
| **8** | 10 | P22791\|HMCS2_RAT | 330.14 | 44 | 44 | 87135000 | 27 | 27 | 32 | Y | 56912 | Hydroxymethylglutaryl-CoA synthase, mitochondrial OS=Rattus norvegicus OX=10116 GN=Hmgcs2 PE=1 SV=1 |
| **8** | 11 | tr\|Q68G44\|Q68G44_RAT | 330.14 | 44 | 44 | 87135000 | 27 | 27 | 32 | Y | 56886 | 3-hydroxy-3-methylglutaryl coenzyme A synthase OS=Rattus norvegicus OX=10116 GN=Hmgcs2 PE=1 SV=1 |
| **16** | 2 | P10860\|DHE3_RAT | 319.64 | 36 | 36 | 61338000 | 19 | 19 | 24 | Y | 61416 | Glutamate dehydrogenase 1, mitochondrial OS=Rattus norvegicus OX=10116 GN=Glud1 PE=1 SV=2 |
| **10** | 5 | Q64428\|ECHA_RAT | 315.18 | 36 | 36 | 36389000 | 26 | 25 | 30 | Y | 82665 | Trifunctional enzyme subunit alpha, mitochondrial OS=Rattus norvegicus OX=10116 GN=Hadha PE=1 SV=2 |
| **5** | 16 | P67779\|PHB_RAT | 293.28 | 85 | 85 | 298710000 | 30 | 29 | 57 | Y | 29820 | Prohibitin OS=Rattus norvegicus OX=10116 GN=Phb PE=1 SV=1 |
| **7** | 97 | P35435\|ATPG_RAT | 283.88 | 65 | 65 | 16360000 | 19 | 1 | 37 | Y | 30191 | ATP synthase subunit gamma, mitochondrial OS=Rattus norvegicus OX=10116 GN=Atp5f1c PE=1 SV=2 |
| **11** | 95 | tr\|Q6QI09\|Q6QI09_RAT | 277.73 | 30 | 30 | 14501000 | 20 | 2 | 29 | Y | 67721 | ATP synthase subunit gamma, mitochondrial OS=Rattus norvegicus OX=10116 GN=Taf3 PE=1 SV=1 |
| **3** | 18 | Q5XIH7\|PHB2_RAT | 275.42 | 75 | 75 | 4403900 | 30 | 1 | 62 | Y | 33312 | Prohibitin-2 OS=Rattus norvegicus OX=10116 GN=Phb2 PE=1 SV=1 |
| **9** | 114 | P19511\|AT5F1_RAT | 275.26 | 55 | 55 | 250850000 | 19 | 19 | 31 | Y | 28869 | ATP synthase F(0) complex subunit B1, mitochondrial OS=Rattus norvegicus OX=10116 GN=Atp5pb PE=1 SV=1 |
| **4** | 19 | tr\|A0A0G2KB63\|A0A0G2KB6 | 274.47 | 76 | 76 | 2528600 | 30 | 1 | 61 | Y | 33168 | Prohibitin OS=Rattus norvegicus OX=10116 GN=Phb2 PE=1 SV=1 |
| **13** | 204 | P31399\|ATP5H_RAT | 266.61 | 80 | 80 | 116440000 | 16 | 16 | 26 | Y | 18763 | ATP synthase subunit d, mitochondrial OS=Rattus norvegicus OX=10116 GN=Atp5pd PE=1 SV=3 |
| **17** | 168 | tr\|G3V7Y3\|G3V7Y3_RAT | 258.55 | 61 | 61 | 76560000 | 11 | 11 | 15 | Y | 17563 | ATP synthase subunit delta, mitochondrial OS=Rattus norvegicus OX=10116 GN=Atp5f1d PE=1 SV=1 |
| **14** | 6 | P52873\|PYC_RAT | 258 | 20 | 20 | 34013000 | 20 | 20 | 25 | Y | 129777 | Pyruvate carboxylase, mitochondrial OS=Rattus norvegicus OX=10116 GN=Pc PE=1 SV=2 |
| **14** | 7 | tr\|A0A0G2JTL5\|A0A0G2JTL5 | 258 | 19 | 19 | 34013000 | 20 | 20 | 25 | Y | 140005 | Pyruvate carboxylase, mitochondrial OS=Rattus norvegicus OX=10116 GN=Pc PE=1 SV=1 |
| **12** | 188 | Q06647\|ATPO_RAT | 245.52 | 65 | 65 | 217880000 | 22 | 22 | 28 | Y | 23398 | ATP synthase subunit O, mitochondrial OS=Rattus norvegicus OX=10116 GN=Atp5po PE=1 SV=1 |
| **19** | 12 | Q66HF1\|NDUS1_RAT | 242.11 | 22 | 22 | 32472000 | 13 | 13 | 14 | N | 79412 | NADH-ubiquinone oxidoreductase 75 kDa subunit, mitochondrial OS=Rattus norvegicus OX=10116 GN=Ndufs1 PE=1 SV=1 |
| **15** | 4 | P00507\|AATM_RAT | 238.5 | 39 | 39 | 83207000 | 18 | 18 | 25 | Y | 47314 | Aspartate aminotransferase, mitochondrial OS=Rattus norvegicus OX=10116 GN=Got2 PE=1 SV=2 |
| **20** | 28 | P18163\|ACSL1_RAT | 213.5 | 15 | 15 | 10176000 | 11 | 8 | 12 | N | 78179 | Long-chain-fatty-acid--CoA ligase 1 OS=Rattus norvegicus OX=10116 GN=Acsl1 PE=1 SV=1 |
| **23** | 48 | Q9WVK3\|PECR_RAT | 204.95 | 37 | 37 | 11917000 | 8 | 8 | 10 | Y | 32433 | Peroxisomal trans-2-enoyl-CoA reductase OS=Rattus norvegicus OX=10116 GN=Pecr PE=2 SV=1 |
| **23** | 49 | tr\|A0A0G2JVG4\|A0A0G2JVG | 204.95 | 36 | 36 | 11917000 | 8 | 8 | 10 | Y | 32737 | Peroxisomal trans-2-enoyl-CoA reductase OS=Rattus norvegicus OX=10116 GN=Pecr PE=1 SV=1 |
| **34** | 54 | tr\|D4A0T0\|D4A0T0_RAT | 200.87 | 28 | 28 | 6537200 | 5 | 5 | 5 | Y | 20859 | NADH:ubiquinone oxidoreductase subunit B10 OS=Rattus norvegicus OX=10116 GN=Ndufb10 PE=1 SV=1 |
| **22** | 14 | Q5BK63\|NDUA9_RAT | 196.06 | 27 | 27 | 8445600 | 9 | 9 | 10 | Y | 42559 | NADH dehydrogenase [ubiquinone] 1 alpha subcomplex subunit 9, mitochondrial OS=Rattus norvegicus OX=10116 GN=Ndufa9 PE=1 SV=2 |
| **24** | 32 | P13086\|SUCA_RAT | 193.77 | 17 | 17 | 8044300 | 5 | 5 | 8 | Y | 36148 | Succinate--CoA ligase [ADP/GDP-forming] subunit alpha, mitochondrial OS=Rattus norvegicus OX=10116 GN=Suclg1 PE=2 SV=2 |
| **24** | 33 | tr\|A0A0H2UHE1\|A0A0H2UH | 193.77 | 16 | 16 | 8044300 | 5 | 5 | 8 | Y | 37560 | Succinate--CoA ligase [ADP/GDP-forming] subunit alpha, mitochondrial OS=Rattus norvegicus OX=10116 GN=Suclg1 PE=1 SV=1 |
| **40** | 36 | tr\|D3ZG43\|D3ZG43_RAT | 185.13 | 21 | 21 | 14115000 | 4 | 4 | 4 | N | 30226 | NADH dehydrogenase (Ubiquinone) Fe-S protein 3 (Predicted), isoform CRA_c OS=Rattus norvegicus OX=10116 GN=Ndufs3 PE=1 SV=1 |
| **21** | 31 | Q60587\|ECHB_RAT | 176.17 | 22 | 22 | 13498000 | 9 | 9 | 11 | Y | 51414 | Trifunctional enzyme subunit beta, mitochondrial OS=Rattus norvegicus OX=10116 GN=Hadhb PE=1 SV=1 |
| **37** | 47 | P0C2X9\|AL4A1_RAT | 171.46 | 9 | 9 | 3731900 | 4 | 4 | 4 | N | 61869 | Delta-1-pyrroline-5-carboxylate dehydrogenase, mitochondrial OS=Rattus norvegicus OX=10116 GN=Aldh4a1 PE=1 SV=1 |
| **27** | 55 | P24329\|THTR_RAT | 169.31 | 23 | 23 | 7435600 | 5 | 5 | 6 | N | 33407 | Thiosulfate sulfurtransferase OS=Rattus norvegicus OX=10116 GN=Tst PE=1 SV=3 |
| **44** | 93 | tr\|D4A565\|D4A565_RAT | 167.21 | 28 | 28 | 4243200 | 4 | 4 | 4 | N | 21664 | NADH dehydrogenase (Ubiquinone) 1 beta subcomplex, 5 (Predicted), isoform CRA_b OS=Rattus norvegicus OX=10116 GN=Ndufb5 PE=1 SV=1 |
| **41** | 46 | P17764\|THIL_RAT | 162.52 | 12 | 12 | 2189300 | 4 | 4 | 4 | N | 44695 | Acetyl-CoA acetyltransferase, mitochondrial OS=Rattus norvegicus OX=10116 GN=Acat1 PE=1 SV=1 |
| **32** | 39 | P29147\|BDH_RAT | 155.31 | 15 | 15 | 6003700 | 5 | 5 | 5 | N | 38202 | D-beta-hydroxybutyrate dehydrogenase, mitochondrial OS=Rattus norvegicus OX=10116 GN=Bdh1 PE=1 SV=2 |
| **32** | 40 | tr\|A0A0G2JSH2\|A0A0G2JSH | 155.31 | 15 | 15 | 6003700 | 5 | 5 | 5 | N | 38333 | 3-hydroxybutyrate dehydrogenase, type 1, isoform CRA_a OS=Rattus norvegicus OX=10116 GN=Bdh1 PE=1 SV=1 |
| **55** | 64 | Q02253\|MMSA_RAT | 153.04 | 10 | 10 | 3458800 | 3 | 3 | 3 | N | 57808 | Methylmalonate-semialdehyde dehydrogenase [acylating], mitochondrial OS=Rattus norvegicus OX=10116 GN=Aldh6a1 PE=1 SV=1 |
| **55** | 65 | tr\|G3V7J0\|G3V7J0_RAT | 153.04 | 10 | 10 | 3458800 | 3 | 3 | 3 | N | 57748 | Aldehyde dehydrogenase family 6, subfamily A1, isoform CRA_b OS=Rattus norvegicus OX=10116 GN=Aldh6a1 PE=1 SV=1 |
| **26** | 21 | tr\|Q5XIH3\|Q5XIH3_RAT | 151.14 | 18 | 18 | 4726100 | 7 | 7 | 7 | N | 50731 | NADH dehydrogenase [ubiquinone] flavoprotein 1, mitochondrial OS=Rattus norvegicus OX=10116 GN=Ndufv1 PE=1 SV=1 |
| **30** | 27 | P85834\|EFTU_RAT | 145 | 13 | 13 | 4037800 | 5 | 5 | 5 | N | 49522 | Elongation factor Tu, mitochondrial OS=Rattus norvegicus OX=10116 GN=Tufm PE=1 SV=1 |
| **29** | 61 | P63039\|CH60_RAT | 139.8 | 7 | 7 | 18976000 | 4 | 4 | 5 | N | 60956 | 60 kDa heat shock protein, mitochondrial OS=Rattus norvegicus OX=10116 GN=Hspd1 PE=1 SV=1 |
| **29** | 62 | tr\|A0A482IDN3\|A0A482IDN | 139.8 | 7 | 7 | 18976000 | 4 | 4 | 5 | N | 60956 | Hsp60 OS=Rattus norvegicus OX=10116 GN=Hspd1 PE=2 SV=1 |
| **90** | 4499 | Q9JJW3\|ATPMD_RAT | 139.63 | 28 | 28 | 10314000 | 2 | 2 | 2 | N | 6408 | ATP synthase membrane subunit DAPIT, mitochondrial OS=Rattus norvegicus OX=10116 GN=Atp5md PE=1 SV=1 |
| **35** | 4490 | Q6PDU7\|ATP5L_RAT | 133.19 | 34 | 34 | 11005000 | 2 | 2 | 5 | Y | 11433 | ATP synthase subunit g, mitochondrial OS=Rattus norvegicus OX=10116 GN=Atp5mg PE=1 SV=2 |
| **58** | 35 | tr\|Q6P6W6\|Q6P6W6_RAT | 128.01 | 20 | 20 | 1170700 | 3 | 3 | 3 | N | 31701 | NADH dehydrogenase [ubiquinone] 1 alpha subcomplex subunit 10, mitochondrial OS=Rattus norvegicus OX=10116 GN=Ndufa10 PE=2 SV=1 |
| **58** | 23 | Q561S0\|NDUAA_RAT | 128.01 | 15 | 15 | 1170700 | 3 | 3 | 3 | N | 40493 | NADH dehydrogenase [ubiquinone] 1 alpha subcomplex subunit 10, mitochondrial OS=Rattus norvegicus OX=10116 GN=Ndufa10 PE=1 SV=1 |
| **58** | 26 | tr\|A0A1W2Q6F8\|A0A1W2Q | 128.01 | 15 | 15 | 1170700 | 3 | 3 | 3 | N | 40544 | NADH dehydrogenase [ubiquinone] 1 alpha subcomplex subunit 10, mitochondrial OS=Rattus norvegicus OX=10116 GN=Ndufa10l1 PE=3 SV=1 |
| **33** | 58 | P19234\|NDUV2_RAT | 126.7 | 24 | 24 | 14809000 | 4 | 4 | 5 | N | 27378 | NADH dehydrogenase [ubiquinone] flavoprotein 2, mitochondrial OS=Rattus norvegicus OX=10116 GN=Ndufv2 PE=1 SV=2 |
| **42** | 50 | P10888\|COX41_RAT | 126.18 | 24 | 24 | 4134200 | 4 | 4 | 4 | N | 19515 | Cytochrome c oxidase subunit 4 isoform 1, mitochondrial OS=Rattus norvegicus OX=10116 GN=Cox4i1 PE=1 SV=1 |

| | | | | | | | | | | | | |
|---|---|---|---|---|---|---|---|---|---|---|---|---|
| **61** | 4494 | D3Z9R8\|ATP68_RAT | 123.36 | 32 | 32 | 4396500 | 3 | 3 | 3 | Y | 6914 | ATP synthase subunit ATP5MPL, mitochondrial OS=Rattus norvegicus OX=10116 GN=Atp5mpl PE=1 SV=1 |
| **49** | 44 | P16970\|ABCD3_RAT | 122.68 | 5 | 5 | 1969000 | 2 | 2 | 3 | N | 75316 | ATP-binding cassette sub-family D member 3 OS=Rattus norvegicus OX=10116 GN=Abcd3 PE=1 SV=3 |
| **48** | 76 | O70351\|HCD2_RAT | 121.91 | 16 | 16 | 2753200 | 3 | 3 | 3 | N | 27246 | 3-hydroxyacyl-CoA dehydrogenase type-2 OS=Rattus norvegicus OX=10116 GN=Hsd17b10 PE=1 SV=3 |
| **48** | 77 | tr\|B0BMW2\|B0BMW2_RAT | 121.91 | 16 | 16 | 2753200 | 3 | 3 | 3 | N | 27250 | 3-hydroxyacyl-CoA dehydrogenase type-2 OS=Rattus norvegicus OX=10116 GN=Hsd17b10 PE=1 SV=1 |
| **56** | 105 | tr\|D3ZF13\|D3ZF13_RAT | 118.09 | 15 | 15 | 3263000 | 3 | 3 | 3 | N | 17514 | Acyl carrier protein OS=Rattus norvegicus OX=10116 GN=Ndufab1 PE=1 SV=1 |
| **71** | 24 | Q641Y2\|NDUS2_RAT | 108.84 | 5 | 5 | 4160900 | 2 | 2 | 2 | N | 52562 | NADH dehydrogenase [ubiquinone] iron-sulfur protein 2, mitochondrial OS=Rattus norvegicus OX=10116 GN=Ndufs2 PE=1 SV=1 |
| **85** | 73 | tr\|B2RYS8\|B2RYS8_RAT | 106.22 | 15 | 15 | 1986100 | 2 | 2 | 2 | N | 21959 | NADH dehydrogenase [ubiquinone] 1 beta subcomplex subunit 8, mitochondrial OS=Rattus norvegicus OX=10116 GN=Ndufb8 PE=1 SV=1 |
| **45** | 4487 | tr\|Q5UAJ5\|Q5UAJ5_RAT | 105.48 | 46 | 46 | 26910000 | 3 | 3 | 4 | N | 7642 | ATP synthase protein 8 OS=Rattus norvegicus OX=10116 GN=ATP8 PE=3 SV=1 |
| **83** | 1 | P32551\|QCR2_RAT | 104.52 | 6 | 6 | 0 | 2 | 2 | 2 | N | 48396 | Cytochrome b-c1 complex subunit 2, mitochondrial OS=Rattus norvegicus OX=10116 GN=Uqcrc2 PE=1 SV=2 |
| **43** | 4486 | P29419\|ATP5I_RAT | 101.05 | 52 | 52 | 15392000 | 4 | 4 | 4 | N | 8255 | ATP synthase subunit e, mitochondrial OS=Rattus norvegicus OX=10116 GN=Atp5me PE=1 SV=3 |
| **63** | 4496 | P21571\|ATP5J_RAT | 98.83 | 28 | 28 | 830950 | 2 | 2 | 3 | N | 12494 | ATP synthase-coupling factor 6, mitochondrial OS=Rattus norvegicus OX=10116 GN=Atp5pf PE=1 SV=1 |
| **106** | 248 | tr\|G3V6I4\|G3V6I4_RAT | 98.24 | 9 | 9 | 1562300 | 1 | 1 | 1 | N | 37791 | Mitochondrial amidoxime reducing component 1 OS=Rattus norvegicus OX=10116 GN=Marc1 PE=1 SV=1 |
| **196** | 170 | Q63362\|NDUA5_RAT | 97.71 | 22 | 22 | 0 | 1 | 1 | 1 | N | 13412 | NADH dehydrogenase [ubiquinone] 1 alpha subcomplex subunit 5 OS=Rattus norvegicus OX=10116 GN=Ndufa5 PE=1 SV=3 |
| **64** | 112 | tr\|Q5PQZ9\|Q5PQZ9_RAT | 97.41 | 21 | 21 | 638550 | 3 | 3 | 3 | N | 14359 | NADH dehydrogenase [ubiquinone] 1 subunit C2 OS=Rattus norvegicus OX=10116 GN=Ndufc2 PE=1 SV=1 |
| **25** | 4488 | D3ZAF6\|ATPK_RAT | 95.42 | 40 | 40 | 25060000 | 7 | 7 | 8 | Y | 10452 | ATP synthase subunit f, mitochondrial OS=Rattus norvegicus OX=10116 GN=Atp5mf PE=1 SV=1 |
| **38** | 102 | tr\|F1LPB3\|F1LPB3_RAT | 94.17 | 4 | 4 | 288720 | 3 | 1 | 4 | N | 76559 | Long-chain-fatty-acid--CoA ligase 5 OS=Rattus norvegicus OX=10116 GN=Acsl5 PE=1 SV=3 |
| **38** | 101 | O88813\|ACSL5_RAT | 94.17 | 4 | 4 | 288720 | 3 | 1 | 4 | N | 76405 | Long-chain-fatty-acid--CoA ligase 5 OS=Rattus norvegicus OX=10116 GN=Acsl5 PE=1 SV=1 |
| **39** | 53 | B3DMA2\|ACD11_RAT | 89.68 | 4 | 4 | 3247200 | 4 | 4 | 4 | N | 87371 | Acyl-CoA dehydrogenase family member 11 OS=Rattus norvegicus OX=10116 GN=Acad11 PE=1 SV=1 |
| **77** | 51 | tr\|Q5EBA4\|Q5EBA4_RAT | 88.7 | 8 | 8 | 2658700 | 2 | 2 | 2 | N | 33215 | Nipsnap1 protein (Fragment) OS=Rattus norvegicus OX=10116 GN=Nipsnap1 PE=2 SV=1 |
| **77** | 52 | tr\|G3V728\|G3V728_RAT | 88.7 | 8 | 8 | 2658700 | 2 | 2 | 2 | N | 33346 | 4-nitrophenylphosphatase domain and non-neuronal SNAP25-like protein homolog 1 (C. elegans), isoform CRA_b OS=Rattus norvegicus OX=10116 GN=Nipsnap1 PE=1 SV=1 |
| **84** | 71 | tr\|F1LPG5\|F1LPG5_RAT | 82.67 | 29 | 29 | 1846400 | 2 | 2 | 2 | N | 15064 | NADH:ubiquinone oxidoreductase subunit B4 OS=Rattus norvegicus OX=10116 GN=Ndufb4 PE=1 SV=1 |
| **36** | 29 | tr\|A0A140TAG5\|A0A140TAG | 81.55 | 6 | 6 | 5965000 | 3 | 3 | 4 | N | 67049 | MICOS complex subunit MIC60 OS=Rattus norvegicus OX=10116 GN=Immt PE=1 SV=1 |
| **36** | 22 | tr\|A0A0G2JVH4\|A0A0G2JVH | 81.55 | 5 | 5 | 5965000 | 3 | 3 | 4 | N | 86230 | MICOS complex subunit MIC60 OS=Rattus norvegicus OX=10116 GN=Immt PE=1 SV=1 |
| **36** | 30 | Q3KR86\|MIC60_RAT | 81.55 | 6 | 6 | 5965000 | 3 | 3 | 4 | N | 67177 | MICOS complex subunit Mic60 (Fragment) OS=Rattus norvegicus OX=10116 GN=Immt PE=1 SV=1 |
| **88** | 106 | tr\|D3ZXF9\|D3ZXF9_RAT | 80.57 | 9 | 9 | 952090 | 2 | 2 | 2 | Y | 29441 | Mitochondrial ribosomal protein L12 OS=Rattus norvegicus OX=10116 GN=Mrpl12 PE=1 SV=1 |
| **59** | 4489 | P29418\|ATP5E_RAT | 79.32 | 41 | 41 | 5361400 | 3 | 3 | 3 | N | 5767 | ATP synthase subunit epsilon, mitochondrial OS=Rattus norvegicus OX=10116 GN=Atp5f1e PE=1 SV=2 |
| **62** | 67 | tr\|Q5RJN0\|Q5RJN0_RAT | 78.15 | 11 | 11 | 2352400 | 2 | 2 | 3 | N | 23945 | NADH dehydrogenase (Ubiquinone) Fe-S protein 7 OS=Rattus norvegicus OX=10116 GN=Ndufs7 PE=1 SV=1 |
| **60** | 103 | P04762\|CATA_RAT | 75.08 | 8 | 8 | 1307100 | 3 | 3 | 3 | N | 59757 | Catalase OS=Rattus norvegicus OX=10116 GN=Cat PE=1 SV=3 |
| **51** | 63 | Q09073\|ADT2_RAT | 69.12 | 10 | 10 | 2645600 | 3 | 3 | 3 | N | 32901 | ADP/ATP translocase 2 OS=Rattus norvegicus OX=10116 GN=Slc25a5 PE=1 SV=3 |
| **153** | 66 | Q8VID1\|DHRS4_RAT | 67.48 | 5 | 5 | 319920 | 1 | 1 | 1 | N | 29822 | Dehydrogenase/reductase SDR family member 4 OS=Rattus norvegicus OX=10116 GN=Dhrs4 PE=2 SV=2 |
| **197** | 158 | P11240\|COX5A_RAT | 62.99 | 17 | 17 | 0 | 1 | 1 | 1 | N | 16130 | Cytochrome c oxidase subunit 5A, mitochondrial OS=Rattus norvegicus OX=10116 GN=Cox5a PE=1 SV=1 |
| **152** | 104 | tr\|A9UMV9\|A9UMV9_RAT | 61.39 | 11 | 11 | 407550 | 1 | 1 | 1 | N | 12500 | NADH:ubiquinone oxidoreductase subunit A7 OS=Rattus norvegicus OX=10116 GN=Ndufa7 PE=1 SV=1 |
| **151** | 110 | tr\|D3ZCZ9\|D3ZCZ9_RAT | 58.66 | 9 | 9 | 865090 | 1 | 1 | 1 | N | 13040 | NADH dehydrogenase [ubiquinone] iron-sulfur protein 6, mitochondrial OS=Rattus norvegicus OX=10116 GN=LOC100912599 PE=1 SV=1 |
| **151** | 178 | P52504\|NDUS6_RAT | 58.66 | 9 | 9 | 865090 | 1 | 1 | 1 | N | 12783 | NADH dehydrogenase [ubiquinone] iron-sulfur protein 6, mitochondrial OS=Rattus norvegicus OX=10116 GN=Ndufs6 PE=3 SV=1 |
| **91** | 100 | tr\|D3ZE15\|D3ZE15_RAT | 54.88 | 7 | 7 | 1861700 | 2 | 2 | 2 | N | 16777 | NADH:ubiquinone oxidoreductase subunit A13 OS=Rattus norvegicus OX=10116 GN=Ndufa13 PE=1 SV=1 |
| **124** | 111 | Q5RJR8\|LRC59_RAT | 54.45 | 4 | 4 | 723070 | 1 | 1 | 1 | N | 34869 | Leucine-rich repeat-containing protein 59 OS=Rattus norvegicus OX=10116 GN=Lrrc59 PE=1 SV=1 |
| **150** | 160 | Q8VHF5\|CISY_RAT | 50.03 | 2 | 2 | 0 | 1 | 1 | 1 | N | 51867 | Citrate synthase, mitochondrial OS=Rattus norvegicus OX=10116 GN=Cs PE=1 SV=1 |
| **150** | 161 | tr\|G3V936\|G3V936_RAT | 50.03 | 2 | 2 | 0 | 1 | 1 | 1 | N | 51831 | Citrate synthase OS=Rattus norvegicus OX=10116 GN=Cs PE=1 SV=1 |
| **198** | 234 | tr\|Q06QK5\|Q06QK5_RAT | 46.65 | 2 | 2 | 2808500 | 1 | 1 | 1 | N | 68606 | NADH-ubiquinone oxidoreductase chain 5 OS=Rattus norvegicus OX=10116 GN=ND5 PE=3 SV=1 |
| **198** | 235 | P11661\|NU5M_RAT | 46.65 | 2 | 2 | 2808500 | 1 | 1 | 1 | N | 68618 | NADH-ubiquinone oxidoreductase chain 5 OS=Rattus norvegicus OX=10116 GN=Mtnd5 PE=3 SV=3 |
| **198** | 239 | tr\|Q06QG6\|Q06QG6_RAT | 46.65 | 2 | 2 | 2808500 | 1 | 1 | 1 | N | 68588 | NADH-ubiquinone oxidoreductase chain 5 OS=Rattus norvegicus OX=10116 GN=ND5 PE=3 SV=1 |
| **198** | 236 | tr\|Q06QA1\|Q06QA1_RAT | 46.65 | 2 | 2 | 2808500 | 1 | 1 | 1 | N | 68574 | NADH-ubiquinone oxidoreductase chain 5 OS=Rattus norvegicus OX=10116 GN=ND5 PE=3 SV=1 |
| **198** | 240 | tr\|A7XYD5\|A7XYD5_RAT | 46.65 | 2 | 2 | 2808500 | 1 | 1 | 1 | N | 68856 | NADH-ubiquinone oxidoreductase chain 5 OS=Rattus norvegicus OX=10116 GN=Nd5 PE=3 SV=1 |
| **198** | 237 | tr\|Q8SEZ0\|Q8SEZ0_RAT | 46.65 | 2 | 2 | 2808500 | 1 | 1 | 1 | N | 68618 | NADH-ubiquinone oxidoreductase chain 5 OS=Rattus norvegicus OX=10116 GN=Mt-nd5 PE=3 SV=1 |
| **198** | 238 | tr\|A0A096XKT9\|A0A096XKT | 46.65 | 2 | 2 | 2808500 | 1 | 1 | 1 | N | 68584 | NADH-ubiquinone oxidoreductase chain 5 OS=Rattus norvegicus OX=10116 GN=ND5 PE=3 SV=1 |
| **198** | 241 | tr\|A7XYC1\|A7XYC1_RAT | 46.65 | 2 | 2 | 2808500 | 1 | 1 | 1 | N | 68841 | NADH-ubiquinone oxidoreductase chain 5 OS=Rattus norvegicus OX=10116 GN=Nd5 PE=3 SV=1 |
| **198** | 242 | tr\|A0A0A1FZN8\|A0A0A1FZN | 46.65 | 2 | 2 | 2808500 | 1 | 1 | 1 | N | 68970 | NADH-ubiquinone oxidoreductase chain 5 OS=Rattus norvegicus OX=10116 GN=ND5 PE=3 SV=1 |
| **89** | 113 | tr\|B2RYW3\|B2RYW3_RAT | 44.46 | 11 | 11 | 1696700 | 2 | 2 | 2 | N | 21892 | NADH dehydrogenase (Ubiquinone) 1 beta subcomplex, 9 OS=Rattus norvegicus OX=10116 GN=Ndufb9 PE=1 SV=1 |
| **86** | 80 | tr\|S5RZM8\|S5RZM8_RAT | 44.27 | 7 | 7 | 2403600 | 2 | 2 | 2 | Y | 25958 | Cytochrome c oxidase subunit 2 OS=Rattus norvegicus OX=10116 GN=COX2 PE=3 SV=1 |
| **86** | 81 | P00406\|COX2_RAT | 44.27 | 7 | 7 | 2403600 | 2 | 2 | 2 | Y | 25928 | Cytochrome c oxidase subunit 2 OS=Rattus norvegicus OX=10116 GN=Mtco2 PE=1 SV=3 |
| **86** | 68 | tr\|Q5UAJ6\|Q5UAJ6_RAT | 44.27 | 7 | 7 | 2403600 | 2 | 2 | 2 | Y | 25942 | Cytochrome c oxidase subunit 2 OS=Rattus norvegicus OX=10116 GN=COX2 PE=3 SV=1 |

<table>
<tr><th></th><th></th><th></th><th></th><th></th><th></th><th></th><th></th><th></th><th></th><th></th><th></th><th></th></tr>
<tr><td>86</td><td>109</td><td>tr|Q37652|Q37652_RAT</td><td>44.27</td><td>7</td><td>7</td><td>2403600</td><td>2</td><td>2</td><td>2</td><td>Y</td><td>26007</td><td>Cytochrome c oxidase subunit 2 OS=Rattus norvegicus OX=10116 GN=COXII PE=3 SV=1</td></tr>
<tr><td>86</td><td>82</td><td>tr|Q8SEZ5|Q8SEZ5_RAT</td><td>44.27</td><td>7</td><td>7</td><td>2403600</td><td>2</td><td>2</td><td>2</td><td>Y</td><td>25928</td><td>Cytochrome c oxidase subunit 2 OS=Rattus norvegicus OX=10116 GN=Mt-co2 PE=1 SV=1</td></tr>
<tr><td>86</td><td>83</td><td>tr|A0A097PE04|A0A097PE0</td><td>44.27</td><td>7</td><td>7</td><td>2403600</td><td>2</td><td>2</td><td>2</td><td>Y</td><td>25894</td><td>Cytochrome c oxidase subunit 2 OS=Rattus norvegicus OX=10116 GN=COX2 PE=3 SV=1</td></tr>
<tr><td>87</td><td>60</td><td>tr|B0BNE6|B0BNE6_RAT</td><td>43.56</td><td>8</td><td>8</td><td>129890</td><td>2</td><td>2</td><td>2</td><td>Y</td><td>23970</td><td>NADH dehydrogenase (Ubiquinone) Fe-S protein 8 (Predicted), isoform CRA_a OS=Rattus norvegicus OX=10116 GN=Ndufs8 PE=1 SV=1</td></tr>
<tr><td>154</td><td>15</td><td>Q68FY0|QCR1_RAT</td><td>43.38</td><td>5</td><td>5</td><td>0</td><td>1</td><td>1</td><td>1</td><td>N</td><td>52849</td><td>Cytochrome b-c1 complex subunit 1, mitochondrial OS=Rattus norvegicus OX=10116 GN=Uqcrc1 PE=1 SV=1</td></tr>
<tr><td>50</td><td>1061</td><td>tr|D3ZKG9|D3ZKG9_RAT</td><td>41.98</td><td>2</td><td>2</td><td>6711400</td><td>3</td><td>2</td><td>3</td><td>Y</td><td>145348</td><td>Clustered mitochondria protein homolog OS=Rattus norvegicus OX=10116 GN=Cluh PE=1 SV=3</td></tr>
<tr><td>50</td><td>1062</td><td>tr|A0A0U1RRV5|A0A0U1RR</td><td>41.98</td><td>2</td><td>2</td><td>6711400</td><td>3</td><td>2</td><td>3</td><td>Y</td><td>151165</td><td>Clustered mitochondria protein homolog OS=Rattus norvegicus OX=10116 GN=Cluh PE=1 SV=1</td></tr>
<tr><td>199</td><td>88</td><td>tr|D4A3V2|D4A3V2_RAT</td><td>41.62</td><td>8</td><td>8</td><td>621300</td><td>1</td><td>1</td><td>1</td><td>N</td><td>15224</td><td>NADH dehydrogenase [ubiquinone] 1 alpha subcomplex subunit 6 OS=Rattus norvegicus OX=10116 GN=Ndufa6 PE=1 SV=1</td></tr>
<tr><td>149</td><td>1110</td><td>Q06437|ODPAT_RAT</td><td>39.87</td><td>3</td><td>3</td><td>1103200</td><td>1</td><td>1</td><td>1</td><td>N</td><td>43393</td><td>Pyruvate dehydrogenase E1 component subunit alpha, testis-specific form, mitochondrial OS=Rattus norvegicus OX=10116 GN=Pdha2 PE=1 SV=1</td></tr>
<tr><td>200</td><td>228</td><td>tr|Q7TP12|Q7TP12_RAT</td><td>39.81</td><td>10</td><td>10</td><td>1252600</td><td>1</td><td>1</td><td>1</td><td>N</td><td>14969</td><td>Cc2-17 OS=Rattus norvegicus OX=10116 GN=Sqor PE=1 SV=1</td></tr>
<tr><td>200</td><td>38</td><td>tr|B0BMT9|B0BMT9_RAT</td><td>39.81</td><td>3</td><td>3</td><td>1252600</td><td>1</td><td>1</td><td>1</td><td>N</td><td>50202</td><td>Sqrdl protein OS=Rattus norvegicus OX=10116 GN=Sqor PE=1 SV=1</td></tr>
<tr><td>125</td><td>96</td><td>tr|D3ZFJ6|D3ZFJ6_RAT</td><td>37.93</td><td>7</td><td>7</td><td>889180</td><td>1</td><td>1</td><td>1</td><td>N</td><td>60420</td><td>Lactamase, beta OS=Rattus norvegicus OX=10116 GN=Lactb PE=1 SV=1</td></tr>
<tr><td>201</td><td>276</td><td>tr|F1LTG5|F1LTG5_RAT</td><td>37.15</td><td>14</td><td>14</td><td>1812200</td><td>1</td><td>1</td><td>1</td><td>N</td><td>13829</td><td>Uncharacterized protein OS=Rattus norvegicus OX=10116 PE=4 SV=2</td></tr>
<tr><td>201</td><td>133</td><td>P12075|COX5B_RAT</td><td>37.15</td><td>14</td><td>14</td><td>1812200</td><td>1</td><td>1</td><td>1</td><td>N</td><td>13915</td><td>Cytochrome c oxidase subunit 5B, mitochondrial OS=Rattus norvegicus OX=10116 GN=Cox5b PE=1 SV=2</td></tr>
<tr><td>31</td><td>539</td><td>F1M775|DIAP1_RAT</td><td>36.97</td><td>2</td><td>2</td><td>21662000</td><td>3</td><td>3</td><td>5</td><td>Y</td><td>140418</td><td>Protein diaphanous homolog 1 OS=Rattus norvegicus OX=10116 GN=Diaph1 PE=1 SV=3</td></tr>
<tr><td>95</td><td>365</td><td>tr|G3V8G1|G3V8G1_RAT</td><td>33.48</td><td>1</td><td>1</td><td>12791000</td><td>2</td><td>1</td><td>2</td><td>Y</td><td>137113</td><td>ATP/GTP binding protein 1 (Predicted), isoform CRA_a OS=Rattus norvegicus OX=10116 GN=Agtpbp1 PE=4 SV=1</td></tr>
<tr><td>93</td><td>484</td><td>Q80W57|ABCG2_RAT</td><td>33.48</td><td>3</td><td>3</td><td>3650700</td><td>2</td><td>1</td><td>2</td><td>N</td><td>72961</td><td>Broad substrate specificity ATP-binding cassette transporter ABCG2 OS=Rattus norvegicus OX=10116 GN=Abcg2 PE=1 SV=1</td></tr>
<tr><td>202</td><td>373</td><td>Q641Y8|DDX1_RAT</td><td>31.84</td><td>2</td><td>2</td><td>1115600</td><td>1</td><td>1</td><td>1</td><td>Y</td><td>82497</td><td>ATP-dependent RNA helicase DDX1 OS=Rattus norvegicus OX=10116 GN=Ddx1 PE=1 SV=1</td></tr>
<tr><td>74</td><td>482</td><td>tr|D3ZXA6|D3ZXA6_RAT</td><td>29.61</td><td>3</td><td>3</td><td>2048500</td><td>2</td><td>2</td><td>2</td><td>N</td><td>98819</td><td>Pyruvate dehydrogenase phosphatase regulatory subunit OS=Rattus norvegicus OX=10116 GN=Pdpr PE=1 SV=1</td></tr>
<tr><td>69</td><td>321</td><td>tr|F1LNF0|F1LNF0_RAT</td><td>29.17</td><td>1</td><td>1</td><td>4369000</td><td>2</td><td>2</td><td>2</td><td>N</td><td>228912</td><td>Myosin heavy chain 14 OS=Rattus norvegicus OX=10116 GN=Myh14 PE=1 SV=1</td></tr>
<tr><td>47</td><td>421</td><td>P11497|ACACA_RAT</td><td>28.67</td><td>1</td><td>1</td><td>1605200</td><td>3</td><td>2</td><td>3</td><td>Y</td><td>265191</td><td>Acetyl-CoA carboxylase 1 OS=Rattus norvegicus OX=10116 GN=Acaca PE=1 SV=1</td></tr>
<tr><td>203</td><td>941</td><td>tr|D4A6J4|D4A6J4_RAT</td><td>28.сен</td><td>1</td><td>1</td><td>1023600</td><td>1</td><td>1</td><td>1</td><td>N</td><td>111741</td><td>DNA topoisomerase OS=Rattus norvegicus OX=10116 GN=Top3a PE=3 SV=3</td></tr>
<tr><td>75</td><td>515</td><td>tr|A0A0G2K5L6|A0A0G2K5L</td><td>27.72</td><td>1</td><td>1</td><td>0</td><td>2</td><td>1</td><td>2</td><td>Y</td><td>275755</td><td>Acetyl-CoA carboxylase beta OS=Rattus norvegicus OX=10116 GN=Acacb PE=1 SV=1</td></tr>
<tr><td>75</td><td>516</td><td>tr|A0A0G2K1F2|A0A0G2K1F</td><td>27.72</td><td>1</td><td>1</td><td>0</td><td>2</td><td>1</td><td>2</td><td>Y</td><td>276393</td><td>Acetyl-CoA carboxylase beta OS=Rattus norvegicus OX=10116 GN=Acacb PE=1 SV=1</td></tr>
<tr><td>75</td><td>518</td><td>tr|D3ZBE2|D3ZBE2_RAT</td><td>27.72</td><td>1</td><td>1</td><td>0</td><td>2</td><td>1</td><td>2</td><td>Y</td><td>275967</td><td>Acetyl-CoA carboxylase beta OS=Rattus norvegicus OX=10116 GN=Acacb PE=1 SV=3</td></tr>
<tr><td>75</td><td>519</td><td>tr|E9PSQ0|E9PSQ0_RAT</td><td>27.72</td><td>1</td><td>1</td><td>0</td><td>2</td><td>1</td><td>2</td><td>Y</td><td>276255</td><td>Acetyl-CoA carboxylase beta OS=Rattus norvegicus OX=10116 GN=Acacb PE=1 SV=2</td></tr>
<tr><td>75</td><td>517</td><td>tr|O70151|O70151_RAT</td><td>27.72</td><td>1</td><td>1</td><td>0</td><td>2</td><td>1</td><td>2</td><td>Y</td><td>276097</td><td>Acetyl-CoA carboxylase OS=Rattus norvegicus OX=10116 GN=Acacb PE=2 SV=1</td></tr>
<tr><td>112</td><td>1624</td><td>tr|D3ZUJ5|D3ZUJ5_RAT</td><td>25.99</td><td>8</td><td>8</td><td>2021600</td><td>1</td><td>1</td><td>1</td><td>Y</td><td>23973</td><td>Deoxythymidylate kinase OS=Rattus norvegicus OX=10116 GN=Dtymk PE=1 SV=1</td></tr>
<tr><td>204</td><td>119</td><td>P32198|CPT1A_RAT</td><td>25.76</td><td>2</td><td>2</td><td>292360</td><td>1</td><td>1</td><td>1</td><td>N</td><td>88126</td><td>Carnitine O-palmitoyltransferase 1, liver isoform OS=Rattus norvegicus OX=10116 GN=Cpt1a PE=1 SV=2</td></tr>
<tr><td>163</td><td>4506</td><td>Q5BJQ0|COQ8A_RAT</td><td>25.69</td><td>1</td><td>1</td><td>0</td><td>1</td><td>1</td><td>1</td><td>N</td><td>72226</td><td>Atypical kinase COQ8A, mitochondrial OS=Rattus norvegicus OX=10116 GN=Coq8a PE=2 SV=1</td></tr>
<tr><td>159</td><td>784</td><td>tr|A0A0G2JX15|A0A0G2JX1</td><td>25.окт</td><td>1</td><td>1</td><td>715910</td><td>1</td><td>1</td><td>1</td><td>N</td><td>121101</td><td>Cation-transporting ATPase OS=Rattus norvegicus OX=10116 GN=Atp13a2 PE=3 SV=1</td></tr>
<tr><td>92</td><td>4509</td><td>Q06645|AT5G1_RAT</td><td>24.35</td><td>5</td><td>5</td><td>11261000</td><td>1</td><td>1</td><td>2</td><td>N</td><td>14244</td><td>ATP synthase F(0) complex subunit C1, mitochondrial OS=Rattus norvegicus OX=10116 GN=Atp5mc1 PE=1 SV=1</td></tr>
<tr><td>92</td><td>4510</td><td>tr|Q499S2|Q499S2_RAT</td><td>24.35</td><td>5</td><td>5</td><td>11261000</td><td>1</td><td>1</td><td>2</td><td>N</td><td>14745</td><td>ATP synthase F(0) complex subunit C3, mitochondrial OS=Rattus norvegicus OX=10116 GN=Atp5mc3 PE=2 SV=1</td></tr>
<tr><td>92</td><td>4511</td><td>Q06646|AT5G2_RAT</td><td>24.35</td><td>5</td><td>5</td><td>11261000</td><td>1</td><td>1</td><td>2</td><td>N</td><td>14918</td><td>ATP synthase F(0) complex subunit C2, mitochondrial OS=Rattus norvegicus OX=10116 GN=Atp5mc2 PE=1 SV=1</td></tr>
<tr><td>92</td><td>4512</td><td>Q71S46|AT5G3_RAT</td><td>24.35</td><td>5</td><td>5</td><td>11261000</td><td>1</td><td>1</td><td>2</td><td>N</td><td>14693</td><td>ATP synthase F(0) complex subunit C3, mitochondrial OS=Rattus norvegicus OX=10116 GN=Atp5mc3 PE=1 SV=1</td></tr>
<tr><td>92</td><td>4513</td><td>tr|A0A0G2JTN8|A0A0G2JTN</td><td>24.35</td><td>5</td><td>5</td><td>11261000</td><td>1</td><td>1</td><td>2</td><td>N</td><td>15465</td><td>ATP synthase F(0) complex subunit C2, mitochondrial OS=Rattus norvegicus OX=10116 GN=Atp5mc2 PE=3 SV=1</td></tr>
<tr><td>81</td><td>701</td><td>tr|F1LMJ8|F1LMJ8_RAT</td><td>24.29</td><td>5</td><td>5</td><td>6004200</td><td>2</td><td>2</td><td>2</td><td>N</td><td>49215</td><td>Calcium uptake protein 2, mitochondrial OS=Rattus norvegicus OX=10116 GN=Micu2 PE=1 SV=1</td></tr>
<tr><td>160</td><td>1534</td><td>tr|D3ZSM9|D3ZSM9_RAT</td><td>24.20</td><td>1</td><td>1</td><td>934160</td><td>1</td><td>1</td><td>1</td><td>Y</td><td>88476</td><td>BRCA1-associated ATM activator 1 OS=Rattus norvegicus OX=10116 GN=Brat1 PE=1 SV=3</td></tr>
<tr><td>120</td><td>549</td><td>P05708|HXK1_RAT</td><td>24.14</td><td>1</td><td>1</td><td>540180</td><td>1</td><td>1</td><td>1</td><td>N</td><td>102408</td><td>Hexokinase-1 OS=Rattus norvegicus OX=10116 GN=Hk1 PE=1 SV=4</td></tr>
<tr><td>120</td><td>533</td><td>tr|M0RAQ6|M0RAQ6_RAT</td><td>24.14</td><td>1</td><td>1</td><td>540180</td><td>1</td><td>1</td><td>1</td><td>N</td><td>99557</td><td>Hexokinase-1 OS=Rattus norvegicus OX=10116 GN=Hk1 PE=1 SV=2</td></tr>
<tr><td>120</td><td>468</td><td>Q63704|CPT1B_RAT</td><td>24.14</td><td>1</td><td>1</td><td>540180</td><td>1</td><td>1</td><td>1</td><td>N</td><td>88217</td><td>Carnitine O-palmitoyltransferase 1, muscle isoform OS=Rattus norvegicus OX=10116 GN=Cpt1b PE=1 SV=1</td></tr>
<tr><td>120</td><td>489</td><td>F1LN46|CPT1C_RAT</td><td>24.14</td><td>1</td><td>1</td><td>540180</td><td>1</td><td>1</td><td>1</td><td>N</td><td>90170</td><td>Carnitine O-palmitoyltransferase 1, brain isoform OS=Rattus norvegicus OX=10116 GN=Cpt1c PE=1 SV=1</td></tr>
<tr><td>208</td><td>137</td><td>Q9WVK7|HCDH_RAT</td><td>23.96</td><td>3</td><td>3</td><td>0</td><td>1</td><td>1</td><td>1</td><td>N</td><td>34448</td><td>Hydroxyacyl-coenzyme A dehydrogenase, mitochondrial OS=Rattus norvegicus OX=10116 GN=Hadh PE=2 SV=1</td></tr>
<tr><td>94</td><td>4538</td><td>tr|B1WBM1|B1WBM1_RAT</td><td>23.77</td><td>3</td><td>3</td><td>949520</td><td>2</td><td>1</td><td>2</td><td>Y</td><td>58354</td><td>FAST kinase domain-containing protein 3, mitochondrial OS=Rattus norvegicus OX=10116 GN=Fastkd3 PE=2 SV=1</td></tr>
<tr><td>94</td><td>4541</td><td>Q68FN9|FAKD3_RAT</td><td>23.77</td><td>2</td><td>2</td><td>949520</td><td>2</td><td>1</td><td>2</td><td>Y</td><td>75248</td><td>FAST kinase domain-containing protein 3, mitochondrial OS=Rattus norvegicus OX=10116 GN=Fastkd3 PE=2 SV=2</td></tr>
<tr><td>209</td><td>4519</td><td>tr|D3ZA45|D3ZA45_RAT</td><td>23.63</td><td>2</td><td>2</td><td>8430000</td><td>1</td><td>1</td><td>1</td><td>N</td><td>52491</td><td>Autophagy-related protein 13 OS=Rattus norvegicus OX=10116 GN=Atg13 PE=1 SV=2</td></tr>
<tr><td>107</td><td>162</td><td>tr|D3Z900|D3Z900_RAT</td><td>23.39</td><td>3</td><td>3</td><td>0</td><td>1</td><td>1</td><td>1</td><td>N</td><td>38239</td><td>Mitochondrial amidoxime reducing component 2 OS=Rattus norvegicus OX=10116 GN=Marc2 PE=1 SV=2</td></tr>
<tr><td>107</td><td>163</td><td>O88994|MARC2_RAT</td><td>23.39</td><td>3</td><td>3</td><td>0</td><td>1</td><td>1</td><td>1</td><td>N</td><td>38249</td><td>Mitochondrial amidoxime reducing component 2 OS=Rattus norvegicus OX=10116 GN=Marc2 PE=2 SV=1</td></tr>
<tr><td>107</td><td>164</td><td>tr|M0R6N2|M0R6N2_RAT</td><td>23.39</td><td>3</td><td>3</td><td>0</td><td>1</td><td>1</td><td>1</td><td>N</td><td>38175</td><td>MOSC domain-containing protein 2, mitochondrial-like OS=Rattus norvegicus OX=10116 GN=LOC100910481 PE=1 SV=1</td></tr>
<tr><td>127</td><td>4501</td><td>tr|F1LUC0|F1LUC0_RAT</td><td>23.мар</td><td>1</td><td>1</td><td>0</td><td>1</td><td>1</td><td>1</td><td>Y</td><td>72937</td><td>Similar to RIKEN cDNA 5730410E15 gene (Predicted), isoform CRA_a OS=Rattus norvegicus OX=10116 GN=Sybu PE=1 SV=2</td></tr>
<tr><td>28</td><td>4514</td><td>tr|F1LSY7|F1LSY7_RAT</td><td>22.67</td><td>1</td><td>1</td><td>76808000</td><td>1</td><td>1</td><td>5</td><td>Y</td><td>109031</td><td>Endoplasmic reticulum to nucleus-signaling 1 OS=Rattus norvegicus OX=10116 GN=Ern1 PE=4 SV=2</td></tr>
<tr><td>28</td><td>4515</td><td>tr|A0A0G2K2H4|A0A0G2K2H</td><td>22.67</td><td>1</td><td>1</td><td>76808000</td><td>1</td><td>1</td><td>5</td><td>Y</td><td>110150</td><td>Endoplasmic reticulum to nucleus-signaling 1 OS=Rattus norvegicus OX=10116 GN=Ern1 PE=2 SV=1</td></tr>
<tr><td>168</td><td>1019</td><td>Q2M2R8|PEX5_RAT</td><td>22.32</td><td>1</td><td>1</td><td>1913400</td><td>1</td><td>1</td><td>1</td><td>N</td><td>71022</td><td>Peroxisomal targeting signal 1 receptor OS=Rattus norvegicus OX=10116 GN=Pex5 PE=1 SV=2</td></tr>
<tr><td>170</td><td>1370</td><td>Q6IMY1|MTUS1_RAT</td><td>21.96</td><td>2</td><td>2</td><td>0</td><td>1</td><td>1</td><td>1</td><td>N</td><td>50745</td><td>Microtubule-associated tumor suppressor 1 homolog OS=Rattus norvegicus OX=10116 GN=Mtus1 PE=1 SV=1</td></tr>
</table>

<table>
<tr><td>170</td><td>1371</td><td>tr|B2GVA4|B2GVA4_RAT</td><td>21.96</td><td>2</td><td>2</td><td>0</td><td>1</td><td>1</td><td>1</td><td>N</td><td>50728</td><td>Mitochondrial tumor suppressor 1 OS=Rattus norvegicus OX=10116 GN=Mtus1 PE=2 SV=1</td></tr>
<tr><td>169</td><td>4525</td><td>tr|R9PXR4|R9PXR4_RAT</td><td>21.95</td><td>1</td><td>1</td><td>9344600</td><td>1</td><td>1</td><td>1</td><td>N</td><td>61939</td><td>Mitochondrial import receptor subunit TOM70 OS=Rattus norvegicus OX=10116 GN=Tomm70 PE=1 SV=1</td></tr>
<tr><td>169</td><td>4526</td><td>Q75Q39|TOM70_RAT</td><td>21.95</td><td>1</td><td>1</td><td>9344600</td><td>1</td><td>1</td><td>1</td><td>N</td><td>67445</td><td>Mitochondrial import receptor subunit TOM70 OS=Rattus norvegicus OX=10116 GN=Tomm70 PE=1 SV=1</td></tr>
<tr><td>100</td><td>396</td><td>P11530|DMD_RAT</td><td>21.17</td><td>0</td><td>0</td><td>1781200</td><td>1</td><td>1</td><td>1</td><td>N</td><td>425830</td><td>Dystrophin OS=Rattus norvegicus OX=10116 GN=Dmd PE=1 SV=2</td></tr>
<tr><td>98</td><td>834</td><td>tr|A0A0G2JYD4|A0A0G2JYD</td><td>20.72</td><td>0</td><td>0</td><td>4631000</td><td>1</td><td>1</td><td>1</td><td>N</td><td>487857</td><td>Vacuolar protein sorting 13 homolog D OS=Rattus norvegicus OX=10116 GN=Vps13d PE=1 SV=1</td></tr>
<tr><td>98</td><td>835</td><td>tr|D3ZKC6|D3ZKC6_RAT</td><td>20.72</td><td>0</td><td>0</td><td>4631000</td><td>1</td><td>1</td><td>1</td><td>N</td><td>488962</td><td>Vacuolar protein sorting 13 homolog D OS=Rattus norvegicus OX=10116 GN=Vps13d PE=1 SV=1</td></tr>
<tr><td>213</td><td>4555</td><td>Q5EBA0|ERAL1_RAT</td><td>20.48</td><td>2</td><td>2</td><td>0</td><td>1</td><td>1</td><td>1</td><td>N</td><td>48379</td><td>GTPase Era, mitochondrial OS=Rattus norvegicus OX=10116 GN=Eral1 PE=2 SV=2</td></tr>
<tr><td>214</td><td>4556</td><td>Q8VIJ5|OGA_RAT</td><td>20.14</td><td>1</td><td>1</td><td>1932800</td><td>1</td><td>1</td><td>1</td><td>N</td><td>102918</td><td>Protein O-GlcNAcase OS=Rattus norvegicus OX=10116 GN=Oga PE=1 SV=1</td></tr>
<tr><td>total 162 proteins</td><td></td><td></td><td></td><td></td><td></td><td></td><td></td><td></td><td></td><td></td><td></td><td></td></tr>
</table>

Prepared with
PEAKS ™
(bioinfor.com)
