## Supplementary material for "Permeability transition pore-related changes in the proteome and channel activity of ATP synthase dimers and monomers": RLM KCl-BM PTP Monomer of V compl.

| Protein Group | Protein ID | Accession | -10lgP | Coverage (%) | Coverage (%) Sample 7 | Area Sample 7 | #Peptides | #Unique | #Spec Sample 7 | PTM | Avg. Mass | Description |
|---|---|---|---|---|---|---|---|---|---|---|---|---|
| **2** | 13 | P15999\|ATPA_RAT | 488.87 | 85 | 85 | 5592100000 | 104 | 103 | 244 | Y | 59754 | ATP synthase subunit alpha, mitochondrial OS=Rattus norvegicus OX=10116 GN=Atp5f1a PE=1 SV=2 |
| **1** | 9 | tr\|G3V6D3\|G3V6D3_RAT | 479.96 | 84 | 84 | 4460400000 | 75 | 74 | 248 | Y | 56345 | ATP synthase subunit beta OS=Rattus norvegicus OX=10116 GN=Atp5f1b PE=1 SV=1 |
| **1** | 8 | P10719\|ATPB_RAT | 479.96 | 84 | 84 | 4460400000 | 75 | 74 | 248 | Y | 56354 | ATP synthase subunit beta, mitochondrial OS=Rattus norvegicus OX=10116 GN=Atp5f1b PE=1 SV=2 |
| **4** | 3 | P07756\|CPSM_RAT | 398.05 | 45 | 45 | 315720000 | 63 | 62 | 76 | Y | 164579 | Carbamoyl-phosphate synthase [ammonia], mitochondrial OS=Rattus norvegicus OX=10116 GN=Cps1 PE=1 SV=1 |
| **3** | 6 | P52873\|PYC_RAT | 378.67 | 50 | 50 | 693540000 | 65 | 61 | 85 | Y | 129777 | Pyruvate carboxylase, mitochondrial OS=Rattus norvegicus OX=10116 GN=Pc PE=1 SV=2 |
| **3** | 7 | tr\|A0A0G2JTL5\|A0A0G2JTL5_RAT | 378.67 | 46 | 46 | 693540000 | 65 | 61 | 85 | Y | 140005 | Pyruvate carboxylase, mitochondrial OS=Rattus norvegicus OX=10116 GN=Pc PE=1 SV=1 |
| **7** | 204 | P31399\|ATP5H_RAT | 372.39 | 76 | 76 | 434080000 | 26 | 26 | 39 | Y | 18763 | ATP synthase subunit d, mitochondrial OS=Rattus norvegicus OX=10116 GN=Atp5pd PE=1 SV=3 |
| **6** | 114 | P19511\|AT5F1_RAT | 337.53 | 63 | 63 | 863160000 | 28 | 28 | 49 | Y | 28869 | ATP synthase F(0) complex subunit B1, mitochondrial OS=Rattus norvegicus OX=10116 GN=Atp5pb PE=1 SV=1 |
| **11** | 5 | Q64428\|ECHA_RAT | 332.89 | 38 | 38 | 144360000 | 27 | 27 | 30 | Y | 82665 | Trifunctional enzyme subunit alpha, mitochondrial OS=Rattus norvegicus OX=10116 GN=Hadha PE=1 SV=2 |
| **8** | 10 | P22791\|HMCS2_RAT | 331.56 | 45 | 45 | 251280000 | 29 | 29 | 38 | Y | 56912 | Hydroxymethylglutaryl-CoA synthase, mitochondrial OS=Rattus norvegicus OX=10116 GN=Hmgcs2 PE=1 SV=1 |
| **9** | 103 | P04762\|CATA_RAT | 324.96 | 47 | 47 | 226900000 | 28 | 27 | 37 | Y | 59757 | Catalase OS=Rattus norvegicus OX=10116 GN=Cat PE=1 SV=3 |
| **5** | 97 | P35435\|ATPG_RAT | 297.27 | 71 | 71 | 616000000 | 24 | 24 | 49 | Y | 30191 | ATP synthase subunit gamma, mitochondrial OS=Rattus norvegicus OX=10116 GN=Atp5f1c PE=1 SV=2 |
| **13** | 44 | P16970\|ABCD3_RAT | 268.43 | 29 | 29 | 35871000 | 17 | 16 | 24 | N | 75316 | ATP-binding cassette sub-family D member 3 OS=Rattus norvegicus OX=10116 GN=Abcd3 PE=1 SV=3 |
| **16** | 28 | P18163\|ACSL1_RAT | 264.34 | 26 | 26 | 43871000 | 17 | 16 | 18 | Y | 78179 | Long-chain-fatty-acid--CoA ligase 1 OS=Rattus norvegicus OX=10116 GN=Acsl1 PE=1 SV=1 |
| **14** | 1580 | A2VCW9\|AASS_RAT | 261.93 | 19 | 19 | 41627000 | 17 | 17 | 20 | Y | 102908 | Alpha-aminoadipic semialdehyde synthase, mitochondrial OS=Rattus norvegicus OX=10116 GN=Aass PE=2 SV=1 |
| **15** | 61 | P63039\|CH60_RAT | 260.80 | 28 | 28 | 52406000 | 17 | 16 | 20 | Y | 60956 | 60 kDa heat shock protein, mitochondrial OS=Rattus norvegicus OX=10116 GN=Hspd1 PE=1 SV=1 |
| **15** | 62 | tr\|A0A482IDN3\|A0A482IDN3_RAT | 260.80 | 28 | 28 | 52406000 | 17 | 16 | 20 | Y | 60956 | Hsp60 OS=Rattus norvegicus OX=10116 GN=Hspd1 PE=2 SV=1 |
| **10** | 188 | Q06647\|ATPO_RAT | 255.92 | 68 | 68 | 846650000 | 24 | 24 | 33 | Y | 23398 | ATP synthase subunit O, mitochondrial OS=Rattus norvegicus OX=10116 GN=Atp5po PE=1 SV=1 |
| **12** | 31 | Q60587\|ECHB_RAT | 255.39 | 41 | 41 | 97359000 | 22 | 21 | 27 | Y | 51414 | Trifunctional enzyme subunit beta, mitochondrial OS=Rattus norvegicus OX=10116 GN=Hadhb PE=1 SV=1 |
| **18** | 2 | P10860\|DHE3_RAT | 245.57 | 26 | 26 | 59549000 | 13 | 13 | 17 | Y | 61416 | Glutamate dehydrogenase 1, mitochondrial OS=Rattus norvegicus OX=10116 GN=Glud1 PE=1 SV=2 |
| **20** | 168 | tr\|G3V7Y3\|G3V7Y3_RAT | 243.79 | 61 | 61 | 183980000 | 10 | 10 | 14 | Y | 17563 | ATP synthase subunit delta, mitochondrial OS=Rattus norvegicus OX=10116 GN=Atp5f1d PE=1 SV=1 |
| **21** | 46 | P17764\|THIL_RAT | 236.91 | 27 | 27 | 17844000 | 11 | 11 | 13 | N | 44695 | Acetyl-CoA acetyltransferase, mitochondrial OS=Rattus norvegicus OX=10116 GN=Acat1 PE=1 SV=1 |

| | | | | | | | | | | | | |
|---|---|---|---|---|---|---|---|---|---|---|---|---|
| **31** | 15 | Q68FY0\|QCR1_RAT | 232.07 | 23 | 23 | 15442000 | 8 | 8 | 8 | Y | 52849 | Cytochrome b-c1 complex subunit 1, mitochondrial OS=Rattus norvegicus OX=10116 GN=Uqcrc1 PE=1 SV=1 |
| **23** | 32 | P13086\|SUCA_RAT | 228.38 | 27 | 27 | 16128000 | 8 | 8 | 11 | Y | 36148 | Succinate--CoA ligase [ADP/GDP-forming] subunit alpha, mitochondrial OS=Rattus norvegicus OX=10116 GN=Suclg1 PE=2 SV=2 |
| **23** | 33 | tr\|A0A0H2UHE1\|A0A0H2UHE1_RAT | 228.38 | 26 | 26 | 16128000 | 8 | 8 | 11 | Y | 37560 | Succinate--CoA ligase [ADP/GDP-forming] subunit alpha, mitochondrial OS=Rattus norvegicus OX=10116 GN=Suclg1 PE=1 SV=1 |
| **26** | 1 | P32551\|QCR2_RAT | 224.98 | 25 | 25 | 23364000 | 9 | 9 | 10 | N | 48396 | Cytochrome b-c1 complex subunit 2, mitochondrial OS=Rattus norvegicus OX=10116 GN=Uqcrc2 PE=1 SV=2 |
| **17** | 55 | P24329\|THTR_RAT | 219.50 | 39 | 39 | 56702000 | 15 | 15 | 18 | Y | 33407 | Thiosulfate sulfurtransferase OS=Rattus norvegicus OX=10116 GN=Tst PE=1 SV=3 |
| **22** | 64 | Q02253\|MMSA_RAT | 215.93 | 22 | 22 | 20733000 | 11 | 11 | 11 | Y | 57808 | Methylmalonate-semialdehyde dehydrogenase [acylating], mitochondrial OS=Rattus norvegicus OX=10116 GN=Aldh6a1 PE=1 SV=1 |
| **22** | 65 | tr\|G3V7J0\|G3V7J0_RAT | 215.93 | 22 | 22 | 20733000 | 11 | 11 | 11 | Y | 57748 | Aldehyde dehydrogenase family 6, subfamily A1, isoform CRA_b OS=Rattus norvegicus OX=10116 GN=Aldh6a1 PE=1 SV=1 |
| **19** | 4 | P00507\|AATM_RAT | 213.50 | 29 | 29 | 50146000 | 14 | 14 | 17 | Y | 47314 | Aspartate aminotransferase, mitochondrial OS=Rattus norvegicus OX=10116 GN=Got2 PE=1 SV=2 |
| **32** | 53 | B3DMA2\|ACD11_RAT | 206.36 | 13 | 13 | 10317000 | 8 | 8 | 8 | N | 87371 | Acyl-CoA dehydrogenase family member 11 OS=Rattus norvegicus OX=10116 GN=Acad11 PE=1 SV=1 |
| **24** | 47 | P0C2X9\|AL4A1_RAT | 205.82 | 19 | 19 | 25744000 | 9 | 9 | 11 | N | 61869 | Delta-1-pyrroline-5-carboxylate dehydrogenase, mitochondrial OS=Rattus norvegicus OX=10116 GN=Aldh4a1 PE=1 SV=1 |
| **44** | 12439 | Q5XIT9\|MCCB_RAT | 204.16 | 13 | 13 | 5586100 | 5 | 5 | 5 | N | 61517 | Methylcrotonoyl-CoA carboxylase beta chain, mitochondrial OS=Rattus norvegicus OX=10116 GN=Mccc2 PE=2 SV=1 |
| **34** | 4490 | Q6PDU7\|ATP5L_RAT | 193.65 | 66 | 66 | 30454000 | 7 | 7 | 8 | Y | 11433 | ATP synthase subunit g, mitochondrial OS=Rattus norvegicus OX=10116 GN=Atp5mg PE=1 SV=2 |
| **27** | 4499 | Q9JJW3\|ATPMD_RAT | 185.83 | 45 | 45 | 79179000 | 6 | 6 | 10 | N | 6408 | ATP synthase membrane subunit DAPIT, mitochondrial OS=Rattus norvegicus OX=10116 GN=Atp5md PE=1 SV=1 |
| **36** | 56 | P04636\|MDHM_RAT | 185.21 | 24 | 24 | 8121200 | 7 | 7 | 7 | N | 35684 | Malate dehydrogenase, mitochondrial OS=Rattus norvegicus OX=10116 GN=Mdh2 PE=1 SV=2 |
| **38** | 4496 | P21571\|ATP5J_RAT | 174.24 | 44 | 44 | 17048000 | 6 | 6 | 6 | N | 12494 | ATP synthase-coupling factor 6, mitochondrial OS=Rattus norvegicus OX=10116 GN=Atp5pf PE=1 SV=1 |
| **30** | 4487 | tr\|Q5UAJ5\|Q5UAJ5_RAT | 168.07 | 55 | 55 | 137000000 | 4 | 4 | 9 | Y | 7642 | ATP synthase protein 8 OS=Rattus norvegicus OX=10116 GN=ATP8 PE=3 SV=1 |
| **29** | 119 | P32198\|CPT1A_RAT | 165.28 | 13 | 13 | 6892600 | 8 | 8 | 9 | Y | 88126 | Carnitine O-palmitoyltransferase 1, liver isoform OS=Rattus norvegicus OX=10116 GN=Cpt1a PE=1 SV=2 |
| **48** | 92 | tr\|Q5M9H2\|Q5M9H2_RAT | 160.33 | 9 | 9 | 8584400 | 4 | 4 | 4 | N | 70821 | Acyl-Coenzyme A dehydrogenase, very long chain OS=Rattus norvegicus OX=10116 GN=Acadvl PE=1 SV=1 |
| **48** | 91 | P45953\|ACADV_RAT | 160.33 | 9 | 9 | 8584400 | 4 | 4 | 4 | N | 70749 | Very long-chain specific acyl-CoA dehydrogenase, mitochondrial OS=Rattus norvegicus OX=10116 GN=Acadvl PE=1 SV=1 |
| **50** | 5017 | Q5PQT3\|GLYAT_RAT | 158.82 | 14 | 14 | 2057700 | 3 | 3 | 4 | N | 33899 | Glycine N-acyltransferase OS=Rattus norvegicus OX=10116 GN=Glyat PE=2 SV=1 |
| **50** | 5018 | tr\|A4PB92\|A4PB92_RAT | 158.82 | 14 | 14 | 2057700 | 3 | 3 | 4 | N | 33899 | Glycine N-acyltransferase OS=Rattus norvegicus OX=10116 GN=Glyat PE=2 SV=1 |
| **55** | 1310 | P07872\|ACOX1_RAT | 154.71 | 7 | 7 | 2511300 | 3 | 3 | 3 | N | 74679 | Peroxisomal acyl-coenzyme A oxidase 1 OS=Rattus norvegicus OX=10116 GN=Acox1 PE=1 SV=1 |
| **45** | 42 | tr\|D3ZFQ8\|D3ZFQ8_RAT | 145.33 | 14 | 14 | 8479100 | 4 | 4 | 5 | N | 35435 | Cytochrome c-1 OS=Rattus norvegicus OX=10116 GN=Cyc1 PE=1 SV=3 |

| | | | | | | | | | | | | |
|---|---|---|---|---|---|---|---|---|---|---|---|---|
| **40** | 22 | tr\|A0A0G2JVH4\|A0A0G2JVH4_RAT | 138.58 | 9 | 9 | 7827100 | 6 | 6 | 6 | Y | 86230 | MICOS complex subunit MIC60 OS=Rattus norvegicus OX=10116 GN=Immt PE=1 SV=1 |
| **39** | 51 | tr\|Q5EBA4\|Q5EBA4_RAT | 130.16 | 10 | 10 | 10793000 | 4 | 4 | 6 | N | 33215 | Nipsnap1 protein (Fragment) OS=Rattus norvegicus OX=10116 GN=Nipsnap1 PE=2 SV=1 |
| **39** | 52 | tr\|G3V728\|G3V728_RAT | 130.16 | 10 | 10 | 10793000 | 4 | 4 | 6 | N | 33346 | 4-nitrophenylphosphatase domain and non-neuronal SNAP25-like protein homolog 1 (C. elegans), isoform CRA_b OS=Rattus norvegicus OX=10116 GN=Nipsnap1 PE=1 SV=1 |
| **28** | 96 | tr\|D3ZFJ6\|D3ZFJ6_RAT | 129.48 | 18 | 18 | 21094000 | 8 | 8 | 9 | N | 60420 | Lactamase, beta OS=Rattus norvegicus OX=10116 GN=Lactb PE=1 SV=1 |
| **60** | 66 | Q8VID1\|DHRS4_RAT | 118.76 | 12 | 12 | 13709000 | 3 | 3 | 3 | N | 29822 | Dehydrogenase/reductase SDR family member 4 OS=Rattus norvegicus OX=10116 GN=Dhrs4 PE=2 SV=2 |
| **33** | 39 | P29147\|BDH_RAT | 117.79 | 10 | 10 | 15385000 | 8 | 8 | 8 | Y | 38202 | D-beta-hydroxybutyrate dehydrogenase, mitochondrial OS=Rattus norvegicus OX=10116 GN=Bdh1 PE=1 SV=2 |
| **33** | 40 | tr\|A0A0G2JSH2\|A0A0G2JSH2_RAT | 117.79 | 10 | 10 | 15385000 | 8 | 8 | 8 | Y | 38333 | 3-hydroxybutyrate dehydrogenase, type 1, isoform CRA_a OS=Rattus norvegicus OX=10116 GN=Bdh1 PE=1 SV=1 |
| **25** | 4486 | P29419\|ATP5I_RAT | 115.83 | 55 | 55 | 92867000 | 7 | 7 | 11 | N | 8255 | ATP synthase subunit e, mitochondrial OS=Rattus norvegicus OX=10116 GN=Atp5me PE=1 SV=3 |
| **66** | 133 | P12075\|COX5B_RAT | 112.28 | 30 | 30 | 6379300 | 3 | 3 | 3 | N | 13915 | Cytochrome c oxidase subunit 5B, mitochondrial OS=Rattus norvegicus OX=10116 GN=Cox5b PE=1 SV=2 |
| **65** | 111 | Q5RJR8\|LRC59_RAT | 112.19 | 10 | 10 | 2438700 | 3 | 3 | 3 | Y | 34869 | Leucine-rich repeat-containing protein 59 OS=Rattus norvegicus OX=10116 GN=Lrrc59 PE=1 SV=1 |
| **81** | 117 | tr\|G3V734\|G3V734_RAT | 111.64 | 10 | 10 | 3269600 | 2 | 2 | 2 | N | 36133 | 2,4-dienoyl CoA reductase 1, mitochondrial, isoform CRA_a OS=Rattus norvegicus OX=10116 GN=Decr1 PE=1 SV=1 |
| **81** | 118 | Q64591\|DECR_RAT | 111.64 | 10 | 10 | 3269600 | 2 | 2 | 2 | N | 36133 | 2,4-dienoyl-CoA reductase, mitochondrial OS=Rattus norvegicus OX=10116 GN=Decr1 PE=1 SV=2 |
| **89** | 176 | P08011\|MGST1_RAT | 111.12 | 19 | 19 | 1746700 | 2 | 2 | 2 | Y | 17472 | Microsomal glutathione S-transferase 1 OS=Rattus norvegicus OX=10116 GN=Mgst1 PE=1 SV=3 |
| **89** | 177 | tr\|B6DYQ4\|B6DYQ4_RAT | 111.12 | 19 | 19 | 1746700 | 2 | 2 | 2 | Y | 17472 | Microsomal glutathione S-transferase OS=Rattus norvegicus OX=10116 GN=Mgst1 PE=2 SV=1 |
| **52** | 45 | P20788\|UCRI_RAT | 108.70 | 17 | 17 | 5644100 | 4 | 4 | 4 | N | 29446 | Cytochrome b-c1 complex subunit Rieske, mitochondrial OS=Rattus norvegicus OX=10116 GN=Uqcrfs1 PE=1 SV=2 |
| **35** | 4488 | D3ZAF6\|ATPK_RAT | 108.39 | 47 | 47 | 33896000 | 7 | 6 | 8 | Y | 10452 | ATP synthase subunit f, mitochondrial OS=Rattus norvegicus OX=10116 GN=Atp5mf PE=1 SV=1 |
| **56** | 4532 | Q5XIN6\|LETM1_RAT | 106.67 | 7 | 7 | 2969100 | 3 | 3 | 3 | N | 83060 | Mitochondrial proton/calcium exchanger protein OS=Rattus norvegicus OX=10116 GN=Letm1 PE=1 SV=1 |
| **53** | 8241 | tr\|F1LM33\|F1LM33_RAT | 105.60 | 2 | 2 | 2652900 | 3 | 3 | 3 | Y | 156679 | Leucine-rich PPR motif-containing protein, mitochondrial OS=Rattus norvegicus OX=10116 GN=Lrpprc PE=1 SV=2 |
| **53** | 8242 | Q5SGE0\|LPPRC_RAT | 105.60 | 2 | 2 | 2652900 | 3 | 3 | 3 | Y | 156652 | Leucine-rich PPR motif-containing protein, mitochondrial OS=Rattus norvegicus OX=10116 GN=Lrpprc PE=1 SV=1 |
| **49** | 416 | tr\|F1M953\|F1M953_RAT | 102.45 | 7 | 7 | 2133600 | 4 | 4 | 4 | N | 73745 | Stress-70 protein, mitochondrial OS=Rattus norvegicus OX=10116 GN=Hspa9 PE=1 SV=1 |
| **49** | 417 | P48721\|GRP75_RAT | 102.45 | 7 | 7 | 2133600 | 4 | 4 | 4 | N | 73858 | Stress-70 protein, mitochondrial OS=Rattus norvegicus OX=10116 GN=Hspa9 PE=1 SV=3 |
| **155** | 244 | Q66H15\|RMD3_RAT | 101.79 | 5 | 5 | 335860 | 1 | 1 | 1 | N | 52312 | Regulator of microtubule dynamics protein 3 OS=Rattus norvegicus OX=10116 GN=Rmdn3 PE=1 SV=1 |
| **37** | 63 | Q09073\|ADT2_RAT | 101.27 | 16 | 16 | 9141500 | 6 | 6 | 6 | Y | 32901 | ADP/ATP translocase 2 OS=Rattus norvegicus OX=10116 GN=Slc25a5 PE=1 SV=3 |

| | | | | | | | | | | | | |
|---|---|---|---|---|---|---|---|---|---|---|---|---|
| **51** | 76 | O70351\|HCD2_RAT | 100.59 | 18 | 18 | 6197400 | 4 | 4 | 4 | N | 27246 | 3-hydroxyacyl-CoA dehydrogenase type-2 OS=Rattus norvegicus OX=10116 GN=Hsd17b10 PE=1 SV=3 |
| **51** | 77 | tr\|B0BMW2\|B0BMW2_RAT | 100.59 | 18 | 18 | 6197400 | 4 | 4 | 4 | N | 27250 | 3-hydroxyacyl-CoA dehydrogenase type-2 OS=Rattus norvegicus OX=10116 GN=Hsd17b10 PE=1 SV=1 |
| **72** | 79 | tr\|A0A0H2UHN7\|A0A0H2UHN7_RAT | 99.93 | 7 | 7 | 1143900 | 2 | 2 | 2 | N | 62492 | RCG24013, isoform CRA_a OS=Rattus norvegicus OX=10116 GN=Cyp27a1 PE=1 SV=1 |
| **72** | 78 | P17178\|CP27A_RAT | 99.93 | 7 | 7 | 1143900 | 2 | 2 | 2 | N | 60733 | Sterol 26-hydroxylase, mitochondrial OS=Rattus norvegicus OX=10116 GN=Cyp27a1 PE=1 SV=1 |
| **96** | 158 | P11240\|COX5A_RAT | 96.56 | 10 | 10 | 1361900 | 1 | 1 | 2 | N | 16130 | Cytochrome c oxidase subunit 5A, mitochondrial OS=Rattus norvegicus OX=10116 GN=Cox5a PE=1 SV=1 |
| **63** | 282 | P19804\|NDKB_RAT | 95.49 | 20 | 20 | 6514400 | 3 | 3 | 3 | N | 17283 | Nucleoside diphosphate kinase B OS=Rattus norvegicus OX=10116 GN=Nme2 PE=1 SV=1 |
| **64** | 48 | Q9WVK3\|PECR_RAT | 91.06 | 9 | 9 | 7976400 | 3 | 3 | 3 | N | 32433 | Peroxisomal trans-2-enoyl-CoA reductase OS=Rattus norvegicus OX=10116 GN=Pecr PE=2 SV=1 |
| **64** | 49 | tr\|A0A0G2JVG4\|A0A0G2JVG4_RAT | 91.06 | 9 | 9 | 7976400 | 3 | 3 | 3 | N | 32737 | Peroxisomal trans-2-enoyl-CoA reductase OS=Rattus norvegicus OX=10116 GN=Pecr PE=1 SV=1 |
| **42** | 1035 | Q9Z2L0\|VDAC1_RAT | 90.61 | 18 | 18 | 3497000 | 5 | 4 | 5 | Y | 30756 | Voltage-dependent anion-selective channel protein 1 OS=Rattus norvegicus OX=10116 GN=Vdac1 PE=1 SV=4 |
| **46** | 4489 | P29418\|ATP5E_RAT | 88.50 | 55 | 55 | 71086000 | 4 | 4 | 5 | Y | 5767 | ATP synthase subunit epsilon, mitochondrial OS=Rattus norvegicus OX=10116 GN=Atp5f1e PE=1 SV=2 |
| **203** | 254 | tr\|Q7H115\|Q7H115_RAT | 85.02 | 5 | 5 | 410410 | 1 | 1 | 1 | N | 29871 | Cytochrome c oxidase subunit 3 OS=Rattus norvegicus OX=10116 GN=Mt-co3 PE=3 SV=1 |
| **203** | 255 | tr\|I6V4L9\|I6V4L9_RAT | 85.02 | 5 | 5 | 410410 | 1 | 1 | 1 | N | 29844 | Cytochrome c oxidase subunit 3 OS=Rattus norvegicus OX=10116 GN=COX3 PE=3 SV=1 |
| **203** | 256 | tr\|Q8M7G4\|Q8M7G4_RAT | 85.02 | 5 | 5 | 410410 | 1 | 1 | 1 | N | 29861 | Cytochrome c oxidase subunit 3 OS=Rattus norvegicus OX=10116 PE=2 SV=1 |
| **203** | 257 | tr\|A0A096XKT2\|A0A096XKT2_RAT | 85.02 | 5 | 5 | 410410 | 1 | 1 | 1 | N | 29901 | Cytochrome c oxidase subunit 3 OS=Rattus norvegicus OX=10116 GN=COX3 PE=3 SV=1 |
| **203** | 258 | P05505\|COX3_RAT | 85.02 | 5 | 5 | 410410 | 1 | 1 | 1 | N | 29871 | Cytochrome c oxidase subunit 3 OS=Rattus norvegicus OX=10116 GN=Mtco3 PE=1 SV=5 |
| **203** | 259 | tr\|Q8SEZ2\|Q8SEZ2_RAT | 85.02 | 5 | 5 | 410410 | 1 | 1 | 1 | N | 29870 | Cytochrome c oxidase subunit 3 (Fragment) OS=Rattus norvegicus OX=10116 GN=COIII PE=3 SV=1 |
| **204** | 12427 | tr\|F1LP30\|F1LP30_RAT | 77.89 | 3 | 3 | 845590 | 1 | 1 | 1 | N | 79296 | Methylcrotonoyl-CoA carboxylase subunit alpha, mitochondrial OS=Rattus norvegicus OX=10116 GN=Mccc1 PE=1 SV=1 |
| **204** | 12428 | Q5I0C3\|MCCA_RAT | 77.89 | 3 | 3 | 845590 | 1 | 1 | 1 | N | 79330 | Methylcrotonoyl-CoA carboxylase subunit alpha, mitochondrial OS=Rattus norvegicus OX=10116 GN=Mccc1 PE=1 SV=1 |
| **57** | 80 | tr\|S5RZM8\|S5RZM8_RAT | 74.72 | 11 | 11 | 5183300 | 3 | 3 | 3 | Y | 25958 | Cytochrome c oxidase subunit 2 OS=Rattus norvegicus OX=10116 GN=COX2 PE=3 SV=1 |
| **57** | 81 | P00406\|COX2_RAT | 74.72 | 11 | 11 | 5183300 | 3 | 3 | 3 | Y | 25928 | Cytochrome c oxidase subunit 2 OS=Rattus norvegicus OX=10116 GN=Mtco2 PE=1 SV=3 |
| **57** | 68 | tr\|Q5UAJ6\|Q5UAJ6_RAT | 74.72 | 11 | 11 | 5183300 | 3 | 3 | 3 | Y | 25942 | Cytochrome c oxidase subunit 2 OS=Rattus norvegicus OX=10116 GN=COX2 PE=3 SV=1 |
| **57** | 109 | tr\|Q37652\|Q37652_RAT | 74.72 | 11 | 11 | 5183300 | 3 | 3 | 3 | Y | 26007 | Cytochrome c oxidase subunit 2 OS=Rattus norvegicus OX=10116 GN=COXII PE=3 SV=1 |
| **57** | 82 | tr\|Q8SEZ5\|Q8SEZ5_RAT | 74.72 | 11 | 11 | 5183300 | 3 | 3 | 3 | Y | 25928 | Cytochrome c oxidase subunit 2 OS=Rattus norvegicus OX=10116 GN=Mt-co2 PE=1 SV=1 |

| | | | | | | | | | | | | |
|---|---|---|---|---|---|---|---|---|---|---|---|---|
| **57** | 83 | tr\|A0A097PE04\|A0A097PE04_RAT | 74.72 | 11 | 11 | 5183300 | 3 | 3 | 3 | Y | 25894 | Cytochrome c oxidase subunit 2 OS=Rattus norvegicus OX=10116 GN=COX2 PE=3 SV=1 |
| **59** | 50 | P10888\|COX41_RAT | 74.54 | 17 | 17 | 7908300 | 3 | 3 | 3 | N | 19515 | Cytochrome c oxidase subunit 4 isoform 1, mitochondrial OS=Rattus norvegicus OX=10116 GN=Cox4i1 PE=1 SV=1 |
| **205** | 145 | P62909\|RS3_RAT | 74.18 | 7 | 7 | 1028500 | 1 | 1 | 1 | Y | 26674 | 40S ribosomal protein S3 OS=Rattus norvegicus OX=10116 GN=Rps3 PE=1 SV=1 |
| **207** | 12424 | tr\|G3V7I0\|G3V7I0_RAT | 73.93 | 4 | 4 | 998460 | 1 | 1 | 1 | N | 28299 | Peroxiredoxin 3 OS=Rattus norvegicus OX=10116 GN=Prdx3 PE=1 SV=1 |
| **207** | 12425 | Q9Z0V6\|PRDX3_RAT | 73.93 | 4 | 4 | 998460 | 1 | 1 | 1 | N | 28295 | Thioredoxin-dependent peroxide reductase, mitochondrial OS=Rattus norvegicus OX=10116 GN=Prdx3 PE=1 SV=2 |
| **206** | 12874 | Q6MGB5\|DHB8_RAT | 72.97 | 5 | 5 | 504990 | 1 | 1 | 1 | N | 26791 | Estradiol 17-beta-dehydrogenase 8 OS=Rattus norvegicus OX=10116 GN=Hsd17b8 PE=1 SV=1 |
| **67** | 1593 | tr\|Q8M7G5\|Q8M7G5_RAT | 70.24 | 7 | 7 | 1908800 | 2 | 1 | 3 | N | 24240 | ATP synthase subunit a (Fragment) OS=Rattus norvegicus OX=10116 PE=2 SV=1 |
| **67** | 1594 | tr\|Q549I0\|Q549I0_RAT | 70.24 | 7 | 7 | 1908800 | 2 | 1 | 3 | N | 25050 | ATP synthase subunit a OS=Rattus norvegicus OX=10116 GN=atp6 PE=4 SV=1 |
| **67** | 1595 | P05504\|ATP6_RAT | 70.24 | 7 | 7 | 1908800 | 2 | 1 | 3 | N | 25076 | ATP synthase subunit a OS=Rattus norvegicus OX=10116 GN=Mt-atp6 PE=1 SV=3 |
| **67** | 1596 | tr\|Q8HIC7\|Q8HIC7_RAT | 70.24 | 7 | 7 | 1908800 | 2 | 1 | 3 | N | 25076 | ATP synthase subunit a OS=Rattus norvegicus OX=10116 GN=Mt-atp6 PE=4 SV=1 |
| **67** | 1597 | tr\|S5S1E9\|S5S1E9_RAT | 70.24 | 7 | 7 | 1908800 | 2 | 1 | 3 | N | 25049 | ATP synthase subunit a OS=Rattus norvegicus OX=10116 GN=ATP6 PE=4 SV=1 |
| **67** | 1598 | tr\|Q06QE5\|Q06QE5_RAT | 70.24 | 7 | 7 | 1908800 | 2 | 1 | 3 | N | 25075 | ATP synthase subunit a OS=Rattus norvegicus OX=10116 GN=ATP6 PE=4 SV=1 |
| **67** | 1599 | tr\|Q8SEZ3\|Q8SEZ3_RAT | 70.24 | 7 | 7 | 1908800 | 2 | 1 | 3 | N | 25030 | ATP synthase subunit a OS=Rattus norvegicus OX=10116 GN=ATPase6 PE=4 SV=1 |
| **73** | 38 | tr\|B0BMT9\|B0BMT9_RAT | 67.51 | 3 | 3 | 2672100 | 1 | 1 | 2 | N | 50202 | Sqrdl protein OS=Rattus norvegicus OX=10116 GN=Sqor PE=1 SV=1 |
| **58** | 162 | tr\|D3Z900\|D3Z900_RAT | 67.29 | 7 | 7 | 4790300 | 3 | 3 | 3 | N | 38239 | Mitochondrial amidoxime reducing component 2 OS=Rattus norvegicus OX=10116 GN=Marc2 PE=1 SV=2 |
| **58** | 163 | O88994\|MARC2_RAT | 67.29 | 7 | 7 | 4790300 | 3 | 3 | 3 | N | 38249 | Mitochondrial amidoxime reducing component 2 OS=Rattus norvegicus OX=10116 GN=Marc2 PE=2 SV=1 |
| **58** | 164 | tr\|M0R6N2\|M0R6N2_RAT | 67.29 | 7 | 7 | 4790300 | 3 | 3 | 3 | N | 38175 | MOSC domain-containing protein 2, mitochondrial-like OS=Rattus norvegicus OX=10116 GN=LOC100910481 PE=1 SV=1 |
| **82** | 248 | tr\|G3V6I4\|G3V6I4_RAT | 64.95 | 5 | 5 | 1660600 | 2 | 2 | 2 | N | 37791 | Mitochondrial amidoxime reducing component 1 OS=Rattus norvegicus OX=10116 GN=Marc1 PE=1 SV=1 |
| **90** | 137 | Q9WVK7\|HCDH_RAT | 61.57 | 4 | 4 | 2042300 | 2 | 2 | 2 | N | 34448 | Hydroxyacyl-coenzyme A dehydrogenase, mitochondrial OS=Rattus norvegicus OX=10116 GN=Hadh PE=2 SV=1 |
| **208** | 409 | P56574\|IDHP_RAT | 59.41 | 2 | 2 | 621240 | 1 | 1 | 1 | N | 50967 | Isocitrate dehydrogenase [NADP], mitochondrial OS=Rattus norvegicus OX=10116 GN=Idh2 PE=1 SV=2 |
| **84** | 141 | Q924S5\|LONM_RAT | 54.31 | 2 | 2 | 1413600 | 2 | 2 | 2 | Y | 105792 | Lon protease homolog, mitochondrial OS=Rattus norvegicus OX=10116 GN=Lonp1 PE=2 SV=1 |
| **54** | 459 | tr\|F1LPI6\|F1LPI6_RAT | 53.88 | 1 | 1 | 3637800 | 3 | 2 | 3 | N | 200075 | Peripheral-type benzodiazepine receptor-associated protein 1 OS=Rattus norvegicus OX=10116 GN=Tspoap1 PE=4 SV=1 |
| **54** | 460 | Q9JIR0\|RIMB1_RAT | 53.88 | 1 | 1 | 3637800 | 3 | 2 | 3 | N | 200202 | Peripheral-type benzodiazepine receptor-associated protein 1 OS=Rattus norvegicus OX=10116 GN=Tspoap1 PE=1 SV=2 |

| | | | | | | | | | | | | |
|---|---|---|---|---|---|---|---|---|---|---|---|---|
| **209** | 1706 | Q63716\|PRDX1_RAT | 53.40 | 6 | 6 | 2002900 | 1 | 1 | 1 | N | 22109 | Peroxiredoxin-1 OS=Rattus norvegicus OX=10116 GN=Prdx1 PE=1 SV=1 |
| **61** | 515 | tr\|A0A0G2K5L6\|A0A0G2K5L6_RAT | 53.01 | 1 | 1 | 1590300 | 3 | 1 | 3 | Y | 275755 | Acetyl-CoA carboxylase beta OS=Rattus norvegicus OX=10116 GN=Acacb PE=1 SV=1 |
| **61** | 516 | tr\|A0A0G2K1F2\|A0A0G2K1F2_RAT | 53.01 | 1 | 1 | 1590300 | 3 | 1 | 3 | Y | 276393 | Acetyl-CoA carboxylase beta OS=Rattus norvegicus OX=10116 GN=Acacb PE=1 SV=1 |
| **61** | 518 | tr\|D3ZBE2\|D3ZBE2_RAT | 53.01 | 1 | 1 | 1590300 | 3 | 1 | 3 | Y | 275967 | Acetyl-CoA carboxylase beta OS=Rattus norvegicus OX=10116 GN=Acacb PE=1 SV=3 |
| **61** | 519 | tr\|E9PSQ0\|E9PSQ0_RAT | 53.01 | 1 | 1 | 1590300 | 3 | 1 | 3 | Y | 276255 | Acetyl-CoA carboxylase beta OS=Rattus norvegicus OX=10116 GN=Acacb PE=1 SV=2 |
| **210** | 290 | tr\|M0RAM5\|M0RAM5_RAT | 52.02 | 6 | 6 | 856390 | 1 | 1 | 1 | N | 22155 | Glutathione peroxidase OS=Rattus norvegicus OX=10116 GN=Gpx1 PE=1 SV=1 |
| **210** | 291 | P04041\|GPX1_RAT | 52.02 | 6 | 6 | 856390 | 1 | 1 | 1 | N | 22305 | Glutathione peroxidase 1 OS=Rattus norvegicus OX=10116 GN=Gpx1 PE=1 SV=4 |
| **211** | 301 | tr\|B2RZ24\|B2RZ24_RAT | 49.31 | 2 | 2 | 922450 | 1 | 1 | 1 | N | 47388 | Succinate-CoA ligase subunit beta (Fragment) OS=Rattus norvegicus OX=10116 GN=Sucla2 PE=2 SV=1 |
| **211** | 303 | tr\|F1LM47\|F1LM47_RAT | 49.31 | 2 | 2 | 922450 | 1 | 1 | 1 | N | 50306 | Succinate--CoA ligase [ADP-forming] subunit beta, mitochondrial OS=Rattus norvegicus OX=10116 GN=Sucla2 PE=1 SV=1 |
| **91** | 12430 | Q9R1Z0\|VDAC3_RAT | 48.96 | 6 | 6 | 786220 | 2 | 1 | 2 | N | 30798 | Voltage-dependent anion-selective channel protein 3 OS=Rattus norvegicus OX=10116 GN=Vdac3 PE=1 SV=2 |
| **91** | 12431 | tr\|A0A0G2JSR0\|A0A0G2JSR0_RAT | 48.96 | 6 | 6 | 786220 | 2 | 1 | 2 | N | 30784 | Voltage-dependent anion-selective channel protein 3 OS=Rattus norvegicus OX=10116 GN=Vdac3 PE=1 SV=1 |
| **91** | 1402 | P81155\|VDAC2_RAT | 48.96 | 5 | 5 | 786220 | 2 | 1 | 2 | N | 31746 | Voltage-dependent anion-selective channel protein 2 OS=Rattus norvegicus OX=10116 GN=Vdac2 PE=1 SV=2 |
| **138** | 160 | Q8VHF5\|CISY_RAT | 48.65 | 2 | 2 | 1553600 | 1 | 1 | 1 | N | 51867 | Citrate synthase, mitochondrial OS=Rattus norvegicus OX=10116 GN=Cs PE=1 SV=1 |
| **138** | 161 | tr\|G3V936\|G3V936_RAT | 48.65 | 2 | 2 | 1553600 | 1 | 1 | 1 | N | 51831 | Citrate synthase OS=Rattus norvegicus OX=10116 GN=Cs PE=1 SV=1 |
| **68** | 1710 | D3ZHR2\|ABCD1_RAT | 46.99 | 2 | 2 | 755240 | 3 | 1 | 3 | N | 81938 | ATP-binding cassette sub-family D member 1 OS=Rattus norvegicus OX=10116 GN=Abcd1 PE=1 SV=1 |
| **213** | 191 | tr\|B2RZD6\|B2RZD6_RAT | 45.95 | 10 | 10 | 1625700 | 1 | 1 | 1 | N | 9327 | NDUFA4, mitochondrial complex-associated OS=Rattus norvegicus OX=10116 GN=Ndufa4 PE=1 SV=1 |
| **212** | 277 | P0CG51\|UBB_RAT | 45.20 | 3 | 3 | 951350 | 1 | 1 | 1 | N | 34369 | Polyubiquitin-B OS=Rattus norvegicus OX=10116 GN=Ubb PE=1 SV=1 |
| **214** | 269 | tr\|A0A0G2K642\|A0A0G2K642_RAT | 43.47 | 4 | 4 | 2627400 | 1 | 1 | 1 | N | 41754 | 3-ketoacyl-CoA thiolase, mitochondrial OS=Rattus norvegicus OX=10116 GN=Acaa2 PE=1 SV=1 |
| **214** | 270 | P13437\|THIM_RAT | 43.47 | 4 | 4 | 2627400 | 1 | 1 | 1 | N | 41871 | 3-ketoacyl-CoA thiolase, mitochondrial OS=Rattus norvegicus OX=10116 GN=Acaa2 PE=1 SV=1 |
| **214** | 271 | tr\|G3V9U2\|G3V9U2_RAT | 43.47 | 4 | 4 | 2627400 | 1 | 1 | 1 | N | 41885 | 3-ketoacyl-CoA thiolase, mitochondrial OS=Rattus norvegicus OX=10116 GN=Acaa2 PE=1 SV=3 |
| **215** | 4531 | P06761\|BIP_RAT | 42.77 | 2 | 2 | 431510 | 1 | 1 | 1 | N | 72347 | Endoplasmic reticulum chaperone BiP OS=Rattus norvegicus OX=10116 GN=Hspa5 PE=1 SV=1 |
| **137** | 4731 | tr\|A0A0G2K3W1\|A0A0G2K3W1_RAT | 41.44 | 2 | 2 | 347510 | 1 | 1 | 1 | N | 53760 | von Willebrand factor A domain-containing 8 OS=Rattus norvegicus OX=10116 GN=Vwa8 PE=1 SV=1 |
| **216** | 31543 | tr\|B2RYM8\|B2RYM8_RAT | 39.01 | 7 | 7 | 68923 | 1 | 1 | 1 | N | 20157 | Family with sequence similarity 210, member B OS=Rattus norvegicus OX=10116 GN=Fam210b PE=1 SV=1 |

| | | | | | | | | | | | | |
|---|---|---|---|---|---|---|---|---|---|---|---|---|
| **93** | 4920 | tr\|F1LPV8\|F1LPV8_RAT | 38.94 | 3 | 3 | 1047100 | 2 | 2 | 2 | N | 46639 | Succinate--CoA ligase [GDP-forming] subunit beta, mitochondrial OS=Rattus norvegicus OX=10116 GN=Suclg2 PE=1 SV=2 |
| **93** | 1660 | tr\|B1H270\|B1H270_RAT | 38.94 | 3 | 3 | 1047100 | 2 | 2 | 2 | N | 46988 | Succinate--CoA ligase [GDP-forming] subunit beta, mitochondrial OS=Rattus norvegicus OX=10116 GN=Suclg2 PE=2 SV=1 |
| **69** | 1344 | D3ZG52\|DNA2_RAT | 37.97 | 1 | 1 | 31878000 | 3 | 2 | 3 | N | 119588 | DNA replication ATP-dependent helicase/nuclease DNA2 OS=Rattus norvegicus OX=10116 GN=Dna2 PE=3 SV=1 |
| **69** | 1345 | tr\|A0A0H2UH92\|A0A0H2UH92_RAT | 37.97 | 1 | 1 | 31878000 | 3 | 2 | 3 | N | 136526 | Graves disease carrier protein OS=Rattus norvegicus OX=10116 GN=Slc25a16 PE=1 SV=1 |
| **47** | 834 | tr\|A0A0G2JYD4\|A0A0G2JYD4_RAT | 37.97 | 0 | 0 | 24109000 | 3 | 3 | 4 | N | 487857 | Vacuolar protein sorting 13 homolog D OS=Rattus norvegicus OX=10116 GN=Vps13d PE=1 SV=1 |
| **47** | 835 | tr\|D3ZKC6\|D3ZKC6_RAT | 37.97 | 0 | 0 | 24109000 | 3 | 3 | 4 | N | 488962 | Vacuolar protein sorting 13 homolog D OS=Rattus norvegicus OX=10116 GN=Vps13d PE=1 SV=1 |
| **217** | 680 | tr\|A0A0G2K7K2\|A0A0G2K7K2_RAT | 37.91 | 2 | 2 | 239570 | 1 | 1 | 1 | N | 66134 | Apoptosis-inducing factor 1, mitochondrial OS=Rattus norvegicus OX=10116 GN=Aifm1 PE=1 SV=1 |
| **217** | 681 | Q9JM53\|AIFM1_RAT | 37.91 | 2 | 2 | 239570 | 1 | 1 | 1 | N | 66723 | Apoptosis-inducing factor 1, mitochondrial OS=Rattus norvegicus OX=10116 GN=Aifm1 PE=1 SV=1 |
| **218** | 4992 | P80431\|COX7B_RAT | 37.86 | 9 | 9 | 290430 | 1 | 1 | 1 | N | 8995 | Cytochrome c oxidase subunit 7B, mitochondrial OS=Rattus norvegicus OX=10116 GN=Cox7b PE=1 SV=3 |
| **85** | 12478 | Q5BK22\|GTPB8_RAT | 35.74 | 7 | 7 | 2574300 | 2 | 2 | 2 | Y | 32232 | GTP-binding protein 8 OS=Rattus norvegicus OX=10116 GN=Gtpbp8 PE=2 SV=1 |
| **105** | 8253 | Q5BJS0\|DHX30_RAT | 34.16 | 1 | 1 | 4048900 | 2 | 1 | 2 | N | 133997 | ATP-dependent RNA helicase DHX30 OS=Rattus norvegicus OX=10116 GN=Dhx30 PE=1 SV=1 |
| **154** | 8376 | Q2V057\|HYPDH_RAT | 32.42 | 2 | 2 | 449640 | 1 | 1 | 1 | N | 51002 | Hydroxyproline dehydrogenase OS=Rattus norvegicus OX=10116 GN=Prodh2 PE=2 SV=1 |
| **156** | 478 | Q64578\|AT2A1_RAT | 31.21 | 1 | 1 | 2598600 | 1 | 1 | 1 | Y | 109409 | Sarcoplasmic/endoplasmic reticulum calcium ATPase 1 OS=Rattus norvegicus OX=10116 GN=Atp2a1 PE=1 SV=1 |
| **77** | 321 | tr\|F1LNF0\|F1LNF0_RAT | 30.51 | 1 | 1 | 2943900 | 2 | 2 | 2 | N | 228912 | Myosin heavy chain 14 OS=Rattus norvegicus OX=10116 GN=Myh14 PE=1 SV=1 |
| **100** | 721 | Q5QJC9\|BAG5_RAT | 29.ноя | 3 | 3 | 2669000 | 2 | 1 | 2 | Y | 51031 | BAG family molecular chaperone regulator 5 OS=Rattus norvegicus OX=10116 GN=Bag5 PE=1 SV=1 |
| **71** | 421 | P11497\|ACACA_RAT | 28.янв | 1 | 1 | 1950300 | 2 | 1 | 2 | N | 265191 | Acetyl-CoA carboxylase 1 OS=Rattus norvegicus OX=10116 GN=Acaca PE=1 SV=1 |
| **101** | 1110 | Q06437\|ODPAT_RAT | 26.70 | 5 | 5 | 11204000 | 2 | 2 | 2 | N | 43393 | Pyruvate dehydrogenase E1 component subunit alpha, testis-specific form, mitochondrial OS=Rattus norvegicus OX=10116 GN=Pdha2 PE=1 SV=1 |
| **104** | 106 | tr\|D3ZXF9\|D3ZXF9_RAT | 25.84 | 6 | 6 | 2346300 | 2 | 2 | 2 | Y | 29441 | Mitochondrial ribosomal protein L12 OS=Rattus norvegicus OX=10116 GN=Mrpl12 PE=1 SV=1 |
| **221** | 4556 | Q8VIJ5\|OGA_RAT | 25.май | 1 | 1 | 969110 | 1 | 1 | 1 | N | 102918 | Protein O-GlcNAcase OS=Rattus norvegicus OX=10116 GN=Oga PE=1 SV=1 |
| **98** | 4494 | D3Z9R8\|ATP68_RAT | 24.24 | 12 | 12 | 8282000 | 1 | 1 | 2 | Y | 6914 | ATP synthase subunit ATP5MPL, mitochondrial OS=Rattus norvegicus OX=10116 GN=Atp5mpl PE=1 SV=1 |
| **182** | 468 | Q63704\|CPT1B_RAT | 23.85 | 1 | 1 | 15421000 | 1 | 1 | 1 | N | 88217 | Carnitine O-palmitoyltransferase 1, muscle isoform OS=Rattus norvegicus OX=10116 GN=Cpt1b PE=1 SV=1 |
| **159** | 498 | tr\|Q5U2X8\|Q5U2X8_RAT | 23.71 | 2 | 2 | 11530000 | 1 | 1 | 1 | N | 50444 | Acyl-CoA thioesterase 9 OS=Rattus norvegicus OX=10116 GN=Acot9 PE=1 SV=1 |
| **158** | 4725 | Q63484\|AKT3_RAT | 23.15 | 2 | 2 | 260280 | 1 | 1 | 1 | Y | 55797 | RAC-gamma serine/threonine-protein kinase OS=Rattus norvegicus OX=10116 GN=Akt3 PE=2 SV=2 |

| | | | | | | | | | | | | |
|---|---|---|---|---|---|---|---|---|---|---|---|---|
| **149** | 396 | P11530\|DMD_RAT | 23.14 | 0 | 0 | 2238800 | 1 | 1 | 1 | N | 425830 | Dystrophin OS=Rattus norvegicus OX=10116 GN=Dmd PE=1 SV=2 |
| **160** | 4526 | Q75Q39\|TOM70_RAT | 22.66 | 1 | 1 | 5719200 | 1 | 1 | 1 | N | 67445 | Mitochondrial import receptor subunit TOM70 OS=Rattus norvegicus OX=10116 GN=Tomm70 PE=1 SV=1 |
| **160** | 4525 | tr\|R9PXR4\|R9PXR4_RAT | 22.66 | 1 | 1 | 5719200 | 1 | 1 | 1 | N | 61939 | Mitochondrial import receptor subunit TOM70 OS=Rattus norvegicus OX=10116 GN=Tomm70 PE=1 SV=1 |
| **224** | 1037 | tr\|G3V6E8\|G3V6E8_RAT | 22.35 | 2 | 2 | 1139600 | 1 | 1 | 1 | N | 56439 | Myocilin OS=Rattus norvegicus OX=10116 GN=Myoc PE=4 SV=1 |
| **224** | 1038 | Q9R1J4\|MYOC_RAT | 22.35 | 2 | 2 | 1139600 | 1 | 1 | 1 | N | 56442 | Myocilin OS=Rattus norvegicus OX=10116 GN=Myoc PE=2 SV=1 |
| **142** | 31545 | Q5FVR2\|TYPH_RAT | 22.34 | 2 | 2 | 0 | 1 | 1 | 1 | N | 49901 | Thymidine phosphorylase OS=Rattus norvegicus OX=10116 GN=Tymp PE=1 SV=1 |
| **124** | 4522 | P02563\|MYH6_RAT | 21.88 | 1 | 1 | 0 | 1 | 1 | 1 | N | 223506 | Myosin-6 OS=Rattus norvegicus OX=10116 GN=Myh6 PE=1 SV=2 |
| **226** | 8311 | tr\|Q6IEA8\|Q6IEA8_RAT | 21.87 | 8 | 8 | 1211800 | 1 | 1 | 1 | Y | 8528 | Interferon, alpha-inducible protein 27-like 2B OS=Rattus norvegicus OX=10116 GN=Ifi27l2b PE=2 SV=1 |
| **228** | 438 | tr\|D4A7V6\|D4A7V6_RAT | 21.фев | 1 | 1 | 4314800 | 1 | 1 | 1 | N | 219689 | Caspase 8-associated protein 2 OS=Rattus norvegicus OX=10116 GN=Casp8ap2 PE=1 SV=1 |
| **129** | 447 | Q2PQA9\|KINH_RAT | 20.82 | 1 | 1 | 187680 | 1 | 1 | 1 | N | 109531 | Kinesin-1 heavy chain OS=Rattus norvegicus OX=10116 GN=Kif5b PE=1 SV=1 |
| **229** | 23466 | Q5M7W1\|TXNIP_RAT | 20.75 | 2 | 2 | 867100 | 1 | 1 | 1 | N | 44018 | Thioredoxin-interacting protein OS=Rattus norvegicus OX=10116 GN=Txnip PE=2 SV=1 |
| **126** | 1697 | tr\|A0A0G2QC41\|A0A0G2QC41_RAT | 20.53 | 1 | 1 | 27177000 | 1 | 1 | 1 | N | 125495 | Histone deacetylase 6 OS=Rattus norvegicus OX=10116 GN=Hdac6 PE=1 SV=1 |
| **123** | 273 | Q7TT47\|SPG7_RAT | 20.21 | 2 | 2 | 342020 | 1 | 1 | 1 | N | 86103 | Paraplegin OS=Rattus norvegicus OX=10116 GN=Spg7 PE=2 SV=2 |
| **127** | 1506 | tr\|D4A1D3\|D4A1D3_RAT | 20.июл | 0 | 0 | 3320600 | 1 | 1 | 1 | Y | 521487 | Sacsin molecular chaperone OS=Rattus norvegicus OX=10116 GN=Sacs PE=1 SV=2 |
| **143** | 1291 | tr\|B1WC65\|B1WC65_RAT | 20.апр | 1 | 1 | 672150 | 1 | 1 | 1 | N | 69836 | HAUS augmin-like complex, subunit 3 OS=Rattus norvegicus OX=10116 GN=Haus3 PE=2 SV=1 |
