## Supplementary material for "Permeability transition pore-related changes in the proteome and channel activity of ATP synthase dimers and monomers": RLM SM-BM Control Dimer of V compl.

| Protein Grc | Protein ID | Accession | -10lgP | Coverage (' | Coverage (' | Area Sample 3 | #Peptides | #Unique | #Spec Sam | PTM | Avg. Mass | Description |
|---|---|---|---|---|---|---|---|---|---|---|---|---|
| 2 | 17 | P10719\|ATPB_RAT | 398.18 | 64 | 64 | 657620000 | 183 | 180 | 407 | Oxidation (M) | | ATP synthase subunit beta, mitochondrial |
| 2 | 18 | tr\|G3V6D3\|G3V6D3_RAT | 398.18 | 64 | 64 | 657620000 | 183 | 180 | 407 | Oxidation (M) | | ATP synthase subunit beta |
| 1 | 20 | P15999\|ATPA_RAT | 358.35 | 57 | 57 | 613900000 | 160 | 156 | 494 | Carbamidomethylatior | | ATP synthase subunit alpha, mitochondrial |
| 1 | 21 | tr\|F1LP05\|F1LP05_RAT | 358.35 | 57 | 57 | 613900000 | 160 | 156 | 494 | Carbamidomethylatior | | ATP synthase subunit alpha |
| 3 | 1 | Q02253\|MMSA_RAT | 325.94 | 51 | 51 | 81132000 | 90 | 90 | 162 | Carbamidomethylatior | | Methylmalonate-semialdehyde dehydrogenase [acylating], mitochondrial |
| 3 | 2 | tr\|G3V7J0\|G3V7J0_RAT | 325.94 | 51 | 51 | 81132000 | 90 | 90 | 162 | Carbamidomethylatior | | Aldehyde dehydrogenase family 6, subfamily A1, isoform CRA_b |
| 4 | 8 | P07756\|CPSM_RAT | 320.50 | 34 | 34 | 73977000 | 96 | 96 | 149 | Oxidation ( | 164579 | Carbamoyl-phosphate synthase [ammonia] mitochondrial OS=Rattus norvegicus OX=10116 GN=Cps1 PE=1 SV=1 |
| 10 | 280 | P31399\|ATP5H_RAT | 309.21 | 71 | 71 | 68444000 | 49 | 49 | 82 | Oxidation ( | 18763 | ATP synthase subunit d mitochondrial OS=Rattus norvegicus OX=10116 GN=Atp5pd PE=1 SV=3 |
| 5 | 11 | P67779\|PHB_RAT | 283.84 | 62 | 62 | 104200000 | 68 | 66 | 131 | | 29820 | Prohibitin OS=Rattus norvegicus OX=10116 GN=Phb PE=1 SV=1 |
| 7 | 35 | P52873\|PYC_RAT | 282.20 | 29 | 29 | 34267000 | 66 | 65 | 110 | Oxidation ( | 129777 | Pyruvate carboxylase mitochondrial OS=Rattus norvegicus OX=10116 GN=Pc PE=1 SV=2 |
| 7 | 34 | tr\|A0A0G2JTL5\|A0A0G2JTL5_RAT | 282.20 | 27 | 27 | 34267000 | 66 | 65 | 110 | Oxidation ( | 140005 | Pyruvate carboxylase mitochondrial OS=Rattus norvegicus OX=10116 GN=Pc PE=1 SV=1 |
| 6 | 19 | Q5XIH7\|PHB2_RAT | 272.00 | 61 | 61 | 79864000 | 67 | 64 | 129 | Oxidation ( | 33312 | Prohibitin-2 OS=Rattus norvegicus OX=10116 GN=Phb2 PE=1 SV=1 |
| 13 | 14 | P10860\|DHE3_RAT | 263.23 | 37 | 37 | 21074000 | 44 | 44 | 61 | Oxidation ( | 61416 | Glutamate dehydrogenase 1 mitochondrial OS=Rattus norvegicus OX=10116 GN=Glud1 PE=1 SV=2 |
| 9 | 301 | P35435\|ATPG_RAT | 259.98 | 40 | 40 | 74174000 | 35 | 35 | 84 | Oxidation ( | 30191 | ATP synthase subunit gamma mitochondrial OS=Rattus norvegicus OX=10116 GN=Atp5f1c PE=1 SV=2 |
| 9 | 297 | tr\|Q6QI09\|Q6QI09_RAT | 259.98 | 18 | 18 | 74174000 | 35 | 35 | 84 | Oxidation ( | 67721 | ATP synthase subunit gamma mitochondrial OS=Rattus norvegicus OX=10116 GN=Taf3 PE=1 SV=1 |
| 11 | 10 | Q66HF1\|NDUS1_RAT | 253.84 | 21 | 21 | 32389000 | 42 | 42 | 67 | | 79412 | NADH-ubiquinone oxidoreductase 75 kDa subunit mitochondrial OS=Rattus norvegicus OX=10116 GN=Ndufs1 PE=1 SV=1 |
| 12 | 15 | P00507\|AATM_RAT | 248.51 | 36 | 36 | 30717000 | 36 | 35 | 63 | Oxidation ( | 47314 | Aspartate aminotransferase mitochondrial OS=Rattus norvegicus OX=10116 GN=Got2 PE=1 SV=2 |
| 15 | 30 | P19234\|NDUV2_RAT | 231.93 | 36 | 36 | 26646000 | 24 | 24 | 47 | | 27378 | NADH dehydrogenase [ubiquinone] flavoprotein 2 mitochondrial OS=Rattus norvegicus OX=10116 GN=Ndufv2 PE=1 SV=2 |
| 19 | 289 | tr\|D3ZFJ6\|D3ZFJ6_RAT | 231.44 | 25 | 25 | 10034000 | 21 | 21 | 31 | | 60420 | Lactamase beta OS=Rattus norvegicus OX=10116 GN=Lactb PE=1 SV=1 |
| 8 | 268 | P19511\|AT5F1_RAT | 225.77 | 30 | 30 | 81333000 | 32 | 32 | 87 | | 28869 | ATP synthase F(0) complex subunit B1 mitochondrial OS=Rattus norvegicus OX=10116 GN=Atp5pb PE=1 SV=1 |
| 17 | 9 | Q5BK63\|NDUA9_RAT | 223.35 | 36 | 36 | 8186200 | 28 | 25 | 41 | | 42559 | NADH dehydrogenase [ubiquinone] 1 alpha subcomplex subunit 9 mitochondrial OS=Rattus norvegicus OX=10116 GN=Ndufa9 PE=1 SV=2 |
| 16 | 26 | Q641Y2\|NDUS2_RAT | 214.17 | 28 | 28 | 13080000 | 23 | 23 | 45 | Oxidation ( | 52562 | NADH dehydrogenase [ubiquinone] iron-sulfur protein 2 mitochondrial OS=Rattus norvegicus OX=10116 GN=Ndufs2 PE=1 SV=1 |
| 25 | 76 | tr\|Q68G44\|Q68G44_RAT | 201.57 | 25 | 25 | 7912400 | 20 | 20 | 24 | | 56886 | 3-hydroxy-3-methylglutaryl coenzyme A synthase OS=Rattus norvegicus OX=10116 GN=Hmgcs2 PE=1 SV=1 |
| 37 | 29 | P07895\|SODM_RAT | 196.80 | 27 | 27 | 7993800 | 9 | 9 | 16 | | 24674 | Superoxide dismutase [Mn] mitochondrial OS=Rattus norvegicus OX=10116 GN=Sod2 PE=1 SV=2 |
| 21 | 387 | tr\|G3V7Y3\|G3V7Y3_RAT | 194.19 | 36 | 36 | 2.1029E7 | 16 | 16 | 30 | Oxidation ( | 17563 | ATP synthase subunit delta mitochondrial OS=Rattus norvegicus OX=10116 GN=Atp5f1d PE=1 SV=1 |
| 23 | 33 | Q64428\|ECHA_RAT | 192.46 | 15 | 15 | 7.333E6 | 21 | 21 | 26 | Formylatio | 82665 | Trifunctional enzyme subunit alpha mitochondrial OS=Rattus norvegicus OX=10116 GN=Hadha PE=1 SV=2 |
| 18 | 5937 | Q6PDU7\|ATP5L_RAT | 190.11 | 39 | 39 | 2.7382E7 | 13 | 13 | 32 | Oxidation ( | 11433 | ATP synthase subunit g mitochondrial OS=Rattus norvegicus OX=10116 GN=Atp5mg PE=1 SV=2 |
| 14 | 5938 | Q06647\|ATPO_RAT | 188.95 | 56 | 56 | 2.5753E7 | 30 | 30 | 51 | Carbamidomethylatior | | ATP synthase subunit O, mitochondrial |
| 45 | 45 | tr\|B2RYS8\|B2RYS8_RAT | 188.09 | 25 | 25 | 3.7714E6 | 8 | 8 | 12 | | 21959 | NADH dehydrogenase [ubiquinone] 1 beta subcomplex subunit 8 mitochondrial OS=Rattus norvegicus OX=10116 GN=Ndufb8 PE=1 SV=1 |
| 24 | 164 | tr\|D3ZF13\|D3ZF13_RAT | 184.14 | 23 | 23 | 1.0259E7 | 13 | 13 | 25 | Oxidation ( | 17514 | Acyl carrier protein OS=Rattus norvegicus OX=10116 GN=Ndufab1 PE=1 SV=1 |
| 35 | 228 | P16970\|ABCD3_RAT | 175.41 | 13 | 13 | 1.2428E7 | 14 | 14 | 17 | | 75316 | ATP-binding cassette sub-family D member 3 OS=Rattus norvegicus OX=10116 GN=Abcd3 PE=1 SV=3 |
| 39 | 183 | tr\|D4A7L4\|D4A7L4_RAT | 174.56 | 32 | 32 | 6.2574E6 | 9 | 9 | 14 | | 17634 | NADH dehydrogenase (Ubiquinone) 1 beta subcomplex 11 (Predicted) OS=Rattus norvegicus OX=10116 GN=Ndufb11 PE=1 SV=1 |
| 28 | 36 | tr\|Q06QK5\|Q06QK5_RAT | 173.28 | 14 | 14 | 6.3817E6 | 15 | 15 | 21 | | 68606 | NADH-ubiquinone oxidoreductase chain 5 OS=Rattus norvegicus OX=10116 GN=ND5 PE=3 SV=1 |
| 28 | 37 | P11661\|NU5M_RAT | 173.28 | 14 | 14 | 6.3817E6 | 15 | 15 | 21 | | 68618 | NADH-ubiquinone oxidoreductase chain 5 OS=Rattus norvegicus OX=10116 GN=Mtnd5 PE=3 SV=3 |
| 28 | 38 | tr\|Q06QA1\|Q06QA1_RAT | 173.28 | 14 | 14 | 6.3817E6 | 15 | 15 | 21 | | 68574 | NADH-ubiquinone oxidoreductase chain 5 OS=Rattus norvegicus OX=10116 GN=ND5 PE=3 SV=1 |
| 28 | 39 | tr\|Q8SEZ0\|Q8SEZ0_RAT | 173.28 | 14 | 14 | 6.3817E6 | 15 | 15 | 21 | | 68618 | NADH-ubiquinone oxidoreductase chain 5 OS=Rattus norvegicus OX=10116 GN=Mt-nd5 PE=3 SV=1 |
| 28 | 40 | tr\|A0A096XKT9\|A0A096XKT9_RAT | 173.28 | 14 | 14 | 6.3817E6 | 15 | 15 | 21 | | 68584 | NADH-ubiquinone oxidoreductase chain 5 OS=Rattus norvegicus OX=10116 GN=ND5 PE=3 SV=1 |
| 26 | 12 | Q68FY0\|QCR1_RAT | 173.08 | 15 | 15 | 6.3117E6 | 12 | 12 | 22 | | 52849 | Cytochrome b-c1 complex subunit 1 mitochondrial OS=Rattus norvegicus OX=10116 GN=Uqcrc1 PE=1 SV=1 |
| 31 | 169 | Q60587\|ECHB_RAT | 171.47 | 18 | 18 | 4.9076E6 | 15 | 15 | 19 | | 51414 | Trifunctional enzyme subunit beta mitochondrial OS=Rattus norvegicus OX=10116 GN=Hadhb PE=1 SV=1 |
| 30 | 13 | P32551\|QCR2_RAT | 168.74 | 15 | 15 | 8.2061E6 | 13 | 13 | 20 | | 48396 | Cytochrome b-c1 complex subunit 2 mitochondrial OS=Rattus norvegicus OX=10116 GN=Uqcrc2 PE=1 SV=2 |
| 47 | 246 | tr\|Q5PQZ9\|Q5PQZ9_RAT | 167.44 | 27 | 27 | 1.0909E7 | 7 | 7 | 12 | | 14359 | NADH dehydrogenase [ubiquinone] 1 subunit C2 OS=Rattus norvegicus OX=10116 GN=Ndufc2 PE=1 SV=1 |

| | | | | | | | | | | | | |
|---|---|---|---|---|---|---|---|---|---|---|---|---|
| 20 | 24 | tr\|D3ZG43\|D3ZG43_RAT | 165.05 | 34 | 34 | 9.9423E6 | 20 | 20 | 30 | | 30226 | NADH dehydrogenase (Ubiquinone) Fe-S protein 3 (Predicted) isoform CRA_c OS=Rattus norvegicus OX=10116 GN=Ndufs3 PE=1 SV=1 |
| 59 | 22 | tr\|D3ZFQ8\|D3ZFQ8_RAT | 159.23 | 9 | 9 | 4.8892E6 | 5 | 5 | 8 | | 35435 | Cytochrome c-1 OS=Rattus norvegicus OX=10116 GN=Cyc1 PE=1 SV=3 |
| 43 | 416 | tr\|Q5UAJ5\|Q5UAJ5_RAT | 156.33 | 48 | 48 | 1.482E7 | 7 | 7 | 13 | Oxidation ( | 7642 | ATP synthase protein 8 OS=Rattus norvegicus OX=10116 GN=ATP8 PE=3 SV=1 |
| 43 | 418 | tr\|Q8SEZ4\|Q8SEZ4_RAT | 156.33 | 48 | 48 | 1.482E7 | 7 | 7 | 13 | Oxidation ( | 7632 | ATP synthase protein 8 OS=Rattus norvegicus OX=10116 GN=ATPase8 PE=3 SV=1 |
| 33 | 42 | tr\|D4A0T0\|D4A0T0_RAT | 155.37 | 35 | 35 | 5.5919E6 | 12 | 12 | 19 | | 20859 | NADH:ubiquinone oxidoreductase subunit B10 OS=Rattus norvegicus OX=10116 GN=Ndufb10 PE=1 SV=1 |
| 60 | 261 | P24329\|THTR_RAT | 154.41 | 10 | 10 | 1.6588E6 | 5 | 5 | 8 | | 33407 | Thiosulfate sulfurtransferase OS=Rattus norvegicus OX=10116 GN=Tst PE=1 SV=3 |
| 51 | 165 | tr\|D4A565\|D4A565_RAT | 153.90 | 24 | 24 | 1.7247E6 | 6 | 6 | 10 | | 21664 | NADH dehydrogenase (Ubiquinone) 1 beta subcomplex 5 (Predicted) isoform CRA_b OS=Rattus norvegicus OX=10116 GN=Ndufb5 PE=1 SV=1 |
| 38 | 266 | P04762\|CATA_RAT | 152.68 | 13 | 13 | 3.0877E6 | 11 | 11 | 15 | | 59757 | Catalase OS=Rattus norvegicus OX=10116 GN=Cat PE=1 SV=3 |
| 42 | 135 | tr\|B0BNE6\|B0BNE6_RAT | 152.14 | 20 | 20 | 3.3957E6 | 13 | 11 | 13 | | 23970 | NADH dehydrogenase (Ubiquinone) Fe-S protein 8 (Predicted) isoform CRA_a OS=Rattus norvegicus OX=10116 GN=Ndufs8 PE=1 SV=1 |
| 40 | 242 | Q80W89\|NDUAB_RAT | 150.70 | 21 | 21 | 6.2084E6 | 9 | 9 | 14 | | 14854 | NADH dehydrogenase [ubiquinone] 1 alpha subcomplex subunit 11 OS=Rattus norvegicus OX=10116 GN=Ndufa11 PE=2 SV=1 |
| 62 | 1939 | tr\|D4ACE9\|D4ACE9_RAT | 146.87 | 7 | 7 | 2.0224E6 | 6 | 6 | 7 | | 103115 | Alpha-aminoadipic semialdehyde synthase mitochondrial OS=Rattus norvegicus OX=10116 GN=Aass PE=1 SV=3 |
| 62 | 1940 | A2VCW9\|AASS_RAT | 146.87 | 7 | 7 | 2.0224E6 | 6 | 6 | 7 | | 102908 | Alpha-aminoadipic semialdehyde synthase mitochondrial OS=Rattus norvegicus OX=10116 GN=Aass PE=2 SV=1 |
| 34 | 100 | P17764\|THIL_RAT | 144.23 | 21 | 21 | 6.2144E6 | 13 | 13 | 18 | | 44695 | Acetyl-CoA acetyltransferase mitochondrial OS=Rattus norvegicus OX=10116 GN=Acat1 PE=1 SV=1 |
| 29 | 23 | tr\|Q5XIH3\|Q5XIH3_RAT | 140.68 | 14 | 14 | 1.2894E7 | 10 | 10 | 21 | | 50731 | NADH dehydrogenase [ubiquinone] flavoprotein 1 mitochondrial OS=Rattus norvegicus OX=10116 GN=Ndufv1 PE=1 SV=1 |
| 27 | 138 | P05508\|NU4M_RAT | 139.23 | 15 | 15 | 3.1924E6 | 13 | 12 | 21 | Oxidation (M) | | NADH-ubiquinone oxidoreductase chain 4 |
| 27 | 141 | tr\|Q06QA2\|Q06QA2_RAT | 139.23 | 15 | 15 | 3.1924E6 | 13 | 12 | 21 | Oxidation (M) | | |
| 27 | 142 | tr\|Q7HKW3\|Q7HKW3_RAT | 139.23 | 15 | 15 | 3.1924E6 | 13 | 12 | 21 | Oxidation (M) | | NADH-ubiquinone oxidoreductase chain 4 |
| 27 | 143 | tr\|Q06Q89\|Q06Q89_RAT | 139.23 | 15 | 15 | 3.1924E6 | 13 | 12 | 21 | Oxidation (M) | | |
| 27 | 145 | tr\|Q8HIC6\|Q8HIC6_RAT | 139.23 | 15 | 15 | 3.1924E6 | 13 | 12 | 21 | Oxidation (M) | | NADH-ubiquinone oxidoreductase chain 4 |
| 27 | 151 | tr\|A7XYB9\|A7XYB9_RAT | 139.23 | 15 | 15 | 3.1924E6 | 13 | 12 | 21 | Oxidation (M) | | |
| 27 | 152 | tr\|Q06QG7\|Q06QG7_RAT | 139.23 | 15 | 15 | 3.1924E6 | 13 | 12 | 21 | Oxidation (M) | | |
| 27 | 144 | tr\|Q35737\|Q35737_RAT | 139.23 | 15 | 15 | 3.1924E6 | 13 | 12 | 21 | Oxidation (M) | | NADH-ubiquinone oxidoreductase chain 4 |
| 27 | 139 | tr\|D2E6K0\|D2E6K0_RAT | 139.23 | 15 | 15 | 3.1924E6 | 13 | 12 | 21 | Oxidation (M) | | |
| 27 | 140 | tr\|Q06QE1\|Q06QE1_RAT | 139.23 | 15 | 15 | 3.1924E6 | 13 | 12 | 21 | Oxidation (M) | | |
| 52 | 179 | P11240\|COX5A_RAT | 136.75 | 33 | 33 | 5.2151E6 | 6 | 6 | 10 | | 16130 | Cytochrome c oxidase subunit 5A mitochondrial OS=Rattus norvegicus OX=10116 GN=Cox5a PE=1 SV=1 |
| 41 | 81 | tr\|S5RZM8\|S5RZM8_RAT | 135.94 | 26 | 26 | 3.3919E6 | 9 | 9 | 13 | Oxidation ( | 25958 | Cytochrome c oxidase subunit 2 OS=Rattus norvegicus OX=10116 GN=COX2 PE=3 SV=1 |
| 41 | 82 | P00406\|COX2_RAT | 135.94 | 26 | 26 | 3.3919E6 | 9 | 9 | 13 | Oxidation ( | 25928 | Cytochrome c oxidase subunit 2 OS=Rattus norvegicus OX=10116 GN=Mtco2 PE=1 SV=3 |
| 41 | 41 | tr\|Q5UAJ6\|Q5UAJ6_RAT | 135.94 | 26 | 26 | 3.3919E6 | 9 | 9 | 13 | Oxidation ( | 25942 | Cytochrome c oxidase subunit 2 OS=Rattus norvegicus OX=10116 GN=COX2 PE=3 SV=1 |
| 41 | 83 | tr\|Q8SEZ5\|Q8SEZ5_RAT | 135.94 | 26 | 26 | 3.3919E6 | 9 | 9 | 13 | Oxidation ( | 25928 | Cytochrome c oxidase subunit 2 OS=Rattus norvegicus OX=10116 GN=Mt-co2 PE=1 SV=1 |
| 41 | 84 | tr\|A0A097PE04\|A0A097PE04_RAT | 135.94 | 26 | 26 | 3.3919E6 | 9 | 9 | 13 | Oxidation ( | 25894 | Cytochrome c oxidase subunit 2 OS=Rattus norvegicus OX=10116 GN=COX2 PE=3 SV=1 |
| 46 | 176 | P13086\|SUCA_RAT | 134.04 | 14 | 14 | 3.3719E6 | 8 | 8 | 12 | | 36148 | Succinate--CoA ligase [ADP/GDP-forming] subunit alpha mitochondrial OS=Rattus norvegicus OX=10116 GN=Suclg1 PE=2 SV=2 |
| 46 | 177 | tr\|A0A0H2UHE1\|A0A0H2UHE1_RA | 134.04 | 14 | 14 | 3.3719E6 | 8 | 8 | 12 | | 37560 | Succinate--CoA ligase [ADP/GDP-forming] subunit alpha mitochondrial OS=Rattus norvegicus OX=10116 GN=Suclg1 PE=1 SV=1 |
| 64 | 133 | tr\|F1LPG5\|F1LPG5_RAT | 132.26 | 19 | 19 | 2.1435E6 | 4 | 4 | 7 | | 15064 | NADH:ubiquinone oxidoreductase subunit B4 OS=Rattus norvegicus OX=10116 GN=Ndufb4 PE=1 SV=1 |
| 53 | 154 | tr\|Q06QK0\|Q06QK0_RAT | 126.29 | 11 | 11 | 1.7259E6 | 6 | 6 | 9 | Oxidation (M) | | |
| 53 | 155 | tr\|Q8SEZ6\|Q8SEZ6_RAT | 126.29 | 11 | 11 | 1.7259E6 | 6 | 6 | 9 | Oxidation (M) | | |
| 53 | 156 | tr\|A0A0A1FZ34\|A0A0A1FZ34_RAT | 126.29 | 11 | 11 | 1.7259E6 | 6 | 6 | 9 | Oxidation (M) | | |
| 53 | 157 | tr\|Q8HIC9\|Q8HIC9_RAT | 126.29 | 11 | 11 | 1.7259E6 | 6 | 6 | 9 | Oxidation (M) | | Cytochrome c oxidase subunit 1 |
| 53 | 158 | P05503\|COX1_RAT | 126.29 | 11 | 11 | 1.7259E6 | 6 | 6 | 9 | Oxidation (M) | | Cytochrome c oxidase subunit 1 |
| 53 | 159 | tr\|Q95938\|Q95938_RAT | 126.29 | 11 | 11 | 1.7259E6 | 6 | 6 | 9 | Oxidation (M) | | Cytochrome c oxidase subunit 1 |
| 44 | 110 | tr\|A0A0G2JVL6\|A0A0G2JVL6_RAT | 125.01 | 19 | 19 | 3.8662E6 | 6 | 6 | 13 | | 19965 | NADH dehydrogenase [ubiquinone] 1 alpha subcomplex subunit 8 OS=Rattus norvegicus OX=10116 GN=Ndufa8 PE=1 SV=1 |
| 57 | 175 | tr\|D3ZCZ9\|D3ZCZ9_RAT | 124.09 | 21 | 21 | 2.1504E6 | 5 | 5 | 8 | | 13040 | NADH dehydrogenase [ubiquinone] iron-sulfur protein 6 mitochondrial OS=Rattus norvegicus OX=10116 GN=LOC100912599 PE=1 SV=1 |
| 22 | 5942 | P05504\|ATP6_RAT | 122.12 | 32 | 32 | 9.3815E6 | 13 | 13 | 29 | Oxidation (M) | | ATP synthase subunit a |

| | | | | | | | | | | | | |
|---|---|---|---|---|---|---|---|---|---|---|---|---|
| 22 | 5943 | tr\|Q8HIC7\|Q8HIC7_RAT | 122.12 | 32 | 32 | 9.3815E6 | 13 | 13 | 29 | Oxidation (M) | | ATP synthase subunit a |
| 22 | 5944 | tr\|S5S1E9\|S5S1E9_RAT | 122.12 | 32 | 32 | 9.3815E6 | 13 | 13 | 29 | Oxidation (M) | | |
| 80 | 814 | P45953\|ACADV_RAT | 116.80 | 3 | 3 | 8.2097E5 | 4 | 4 | 4 | | 70749 | Very long-chain specific acyl-CoA dehydrogenase mitochondrial OS=Rattus norvegicus OX=10116 GN=Acadvl PE=1 SV=1 |
| 80 | 815 | tr\|Q5M9H2\|Q5M9H2_RAT | 116.80 | 3 | 3 | 8.2097E5 | 4 | 4 | 4 | | 70821 | Acyl-Coenzyme A dehydrogenase very long chain OS=Rattus norvegicus OX=10116 GN=Acadvl PE=1 SV=1 |
| 36 | 28 | Q561S0\|NDUAA_RAT | 115.14 | 17 | 17 | 1.8575E6 | 11 | 3 | 17 | Oxidation ( | 40493 | NADH dehydrogenase [ubiquinone] 1 alpha subcomplex subunit 10 mitochondrial OS=Rattus norvegicus OX=10116 GN=Ndufa10 PE=1 SV=1 |
| 66 | 291 | tr\|Q5XFW4\|Q5XFW4_RAT | 114.39 | 10 | 10 | 3.9767E5 | 5 | 5 | 7 | | 20541 | Mitochondrial ribosomal protein L13 OS=Rattus norvegicus OX=10116 GN=Mrpl13 PE=1 SV=1 |
| 70 | 168 | tr\|F1LXA0\|F1LXA0_RAT | 113.54 | 21 | 21 | 1.6995E6 | 4 | 4 | 6 | | 17178 | NADH dehydrogenase [ubiquinone] 1 alpha subcomplex subunit 12 OS=Rattus norvegicus OX=10116 GN=Ndufa12 PE=1 SV=2 |
| 65 | 198 | P85834\|EFTU_RAT | 113.46 | 10 | 10 | 9.1198E5 | 5 | 5 | 7 | | 49522 | Elongation factor Tu mitochondrial OS=Rattus norvegicus OX=10116 GN=Tufm PE=1 SV=1 |
| 55 | 111 | Q5XIF3\|NDUS4_RAT | 112.51 | 12 | 12 | 3.5964E6 | 5 | 5 | 8 | | 19741 | NADH dehydrogenase [ubiquinone] iron-sulfur protein 4 mitochondrial OS=Rattus norvegicus OX=10116 GN=Ndufs4 PE=1 SV=1 |
| 67 | 218 | tr\|M0RAM5\|M0RAM5_RAT | 112.27 | 18 | 18 | 1.2065E6 | 5 | 5 | 6 | | 22155 | Glutathione peroxidase OS=Rattus norvegicus OX=10116 GN=Gpx1 PE=1 SV=1 |
| 67 | 219 | P04041\|GPX1_RAT | 112.27 | 18 | 18 | 1.2065E6 | 5 | 5 | 6 | | 22305 | Glutathione peroxidase 1 OS=Rattus norvegicus OX=10116 GN=Gpx1 PE=1 SV=4 |
| 68 | 44 | tr\|D6NSR7\|D6NSR7_RAT | 107.30 | 14 | 14 | 1.2547E6 | 5 | 5 | 6 | | 42982 | Cytochrome b (Fragment) OS=Rattus norvegicus OX=10116 GN=cytb PE=3 SV=1 |
| 68 | 113 | tr\|A0A411P2I9\|A0A411P2I9_RAT | 107.30 | 15 | 15 | 1.2547E6 | 5 | 5 | 6 | | 40144 | Cytochrome b (Fragment) OS=Rattus norvegicus OX=10116 GN=Cytb PE=3 SV=1 |
| 68 | 114 | tr\|A0A0U2IDR8\|A0A0U2IDR8_RAT | 107.30 | 15 | 15 | 1.2547E6 | 5 | 5 | 6 | | 40981 | Cytochrome b (Fragment) OS=Rattus norvegicus OX=10116 PE=3 SV=1 |
| 68 | 115 | tr\|A0A0U2M179\|A0A0U2M179_RA | 107.30 | 15 | 15 | 1.2547E6 | 5 | 5 | 6 | | 41037 | Cytochrome b (Fragment) OS=Rattus norvegicus OX=10116 PE=3 SV=1 |
| 68 | 116 | tr\|A0A411P2L0\|A0A411P2L0_RAT | 107.30 | 15 | 15 | 1.2547E6 | 5 | 5 | 6 | | 41017 | Cytochrome b (Fragment) OS=Rattus norvegicus OX=10116 GN=Cytb PE=3 SV=1 |
| 68 | 90 | tr\|F8QU31\|F8QU31_RAT | 107.30 | 14 | 14 | 1.2547E6 | 5 | 5 | 6 | | 42181 | Cytochrome b (Fragment) OS=Rattus norvegicus OX=10116 GN=cytb PE=3 SV=1 |
| 68 | 91 | tr\|A0A411P2K0\|A0A411P2K0_RAT | 107.30 | 14 | 14 | 1.2547E6 | 5 | 5 | 6 | | 42238 | Cytochrome b (Fragment) OS=Rattus norvegicus OX=10116 GN=Cytb PE=3 SV=1 |
| 68 | 46 | tr\|L0L4L8\|L0L4L8_RAT | 107.30 | 14 | 14 | 1.2547E6 | 5 | 5 | 6 | | 42470 | Cytochrome b (Fragment) OS=Rattus norvegicus OX=10116 GN=cytb PE=3 SV=1 |
| 68 | 47 | tr\|A0A140GE10\|A0A140GE10_RAT | 107.30 | 14 | 14 | 1.2547E6 | 5 | 5 | 6 | | 42574 | Cytochrome b (Fragment) OS=Rattus norvegicus OX=10116 PE=3 SV=1 |
| 68 | 48 | tr\|A0A140GE11\|A0A140GE11_RAT | 107.30 | 14 | 14 | 1.2547E6 | 5 | 5 | 6 | | 42588 | Cytochrome b (Fragment) OS=Rattus norvegicus OX=10116 PE=3 SV=1 |
| 68 | 78 | tr\|A0A140GE08\|A0A140GE08_RAT | 107.30 | 14 | 14 | 1.2547E6 | 5 | 5 | 6 | | 42527 | Cytochrome b (Fragment) OS=Rattus norvegicus OX=10116 PE=3 SV=1 |
| 68 | 92 | tr\|A0A385HCC9\|A0A385HCC9_RAT | 107.30 | 14 | 14 | 1.2547E6 | 5 | 5 | 6 | | 42766 | Cytochrome b (Fragment) OS=Rattus norvegicus OX=10116 PE=3 SV=1 |
| 68 | 49 | tr\|A0A3Q8AGE8\|A0A3Q8AGE8_RA | 107.30 | 14 | 14 | 1.2547E6 | 5 | 5 | 6 | | 42712 | Cytochrome b (Fragment) OS=Rattus norvegicus OX=10116 GN=Cytb PE=3 SV=1 |
| 68 | 50 | tr\|A0A3Q8AC68\|A0A3Q8AC68_RA | 107.30 | 14 | 14 | 1.2547E6 | 5 | 5 | 6 | | 42622 | Cytochrome b (Fragment) OS=Rattus norvegicus OX=10116 GN=Cytb PE=3 SV=1 |
| 68 | 51 | tr\|A0A3S6FM15\|A0A3S6FM15_RA | 107.30 | 14 | 14 | 1.2547E6 | 5 | 5 | 6 | | 42698 | Cytochrome b (Fragment) OS=Rattus norvegicus OX=10116 GN=Cytb PE=3 SV=1 |
| 68 | 85 | tr\|A0A097PE40\|A0A097PE40_RAT | 107.30 | 14 | 14 | 1.2547E6 | 5 | 5 | 6 | | 42865 | Cytochrome b OS=Rattus norvegicus OX=10116 GN=CYTB PE=3 SV=1 |
| 68 | 52 | tr\|A0A220D9Z6\|A0A220D9Z6_RAT | 107.30 | 14 | 14 | 1.2547E6 | 5 | 5 | 6 | | 43018 | Cytochrome b OS=Rattus norvegicus OX=10116 PE=3 SV=1 |
| 68 | 99 | tr\|D6NSP2\|D6NSP2_RAT | 107.30 | 14 | 14 | 1.2547E6 | 5 | 5 | 6 | | 42945 | Cytochrome b (Fragment) OS=Rattus norvegicus OX=10116 GN=cytb PE=3 SV=1 |
| 68 | 53 | tr\|D6NSS6\|D6NSS6_RAT | 107.30 | 14 | 14 | 1.2547E6 | 5 | 5 | 6 | | 42993 | Cytochrome b (Fragment) OS=Rattus norvegicus OX=10116 GN=cytb PE=3 SV=1 |
| 68 | 86 | tr\|D6NSQ3\|D6NSQ3_RAT | 107.30 | 14 | 14 | 1.2547E6 | 5 | 5 | 6 | | 43012 | Cytochrome b (Fragment) OS=Rattus norvegicus OX=10116 GN=cytb PE=3 SV=1 |
| 68 | 54 | tr\|D6NSR3\|D6NSR3_RAT | 107.30 | 14 | 14 | 1.2547E6 | 5 | 5 | 6 | | 42986 | Cytochrome b (Fragment) OS=Rattus norvegicus OX=10116 GN=cytb PE=3 SV=1 |
| 68 | 93 | tr\|D6NSP4\|D6NSP4_RAT | 107.30 | 14 | 14 | 1.2547E6 | 5 | 5 | 6 | | 43016 | Cytochrome b (Fragment) OS=Rattus norvegicus OX=10116 GN=cytb PE=3 SV=1 |
| 68 | 55 | tr\|D6NSR8\|D6NSR8_RAT | 107.30 | 14 | 14 | 1.2547E6 | 5 | 5 | 6 | | 42998 | Cytochrome b (Fragment) OS=Rattus norvegicus OX=10116 GN=cytb PE=3 SV=1 |
| 68 | 56 | tr\|A0A0A1FZ42\|A0A0A1FZ42_RAT | 107.30 | 14 | 14 | 1.2547E6 | 5 | 5 | 6 | | 42989 | Cytochrome b OS=Rattus norvegicus OX=10116 GN=CYTB PE=3 SV=1 |
| 68 | 57 | tr\|D6NSS7\|D6NSS7_RAT | 107.30 | 14 | 14 | 1.2547E6 | 5 | 5 | 6 | | 42948 | Cytochrome b (Fragment) OS=Rattus norvegicus OX=10116 GN=cytb PE=3 SV=1 |
| 68 | 58 | tr\|Q8SEY9\|Q8SEY9_RAT | 107.30 | 14 | 14 | 1.2547E6 | 5 | 5 | 6 | | 43015 | Cytochrome b OS=Rattus norvegicus OX=10116 GN=cytb PE=3 SV=1 |
| 68 | 59 | tr\|A0A220DA44\|A0A220DA44_RAT | 107.30 | 14 | 14 | 1.2547E6 | 5 | 5 | 6 | | 42968 | Cytochrome b OS=Rattus norvegicus OX=10116 PE=3 SV=1 |
| 68 | 79 | tr\|F2Q6S5\|F2Q6S5_RAT | 107.30 | 14 | 14 | 1.2547E6 | 5 | 5 | 6 | | 42952 | Cytochrome b (Fragment) OS=Rattus norvegicus OX=10116 GN=cytb PE=3 SV=1 |
| 68 | 60 | tr\|D6NSQ0\|D6NSQ0_RAT | 107.30 | 14 | 14 | 1.2547E6 | 5 | 5 | 6 | | 42968 | Cytochrome b (Fragment) OS=Rattus norvegicus OX=10116 GN=cytb PE=3 SV=1 |
| 68 | 61 | tr\|Q5UAI7\|Q5UAI7_RAT | 107.30 | 14 | 14 | 1.2547E6 | 5 | 5 | 6 | | 42998 | Cytochrome b OS=Rattus norvegicus OX=10116 GN=CYTB PE=3 SV=1 |
| 68 | 94 | tr\|D6NSR2\|D6NSR2_RAT | 107.30 | 14 | 14 | 1.2547E6 | 5 | 5 | 6 | | 43073 | Cytochrome b (Fragment) OS=Rattus norvegicus OX=10116 GN=cytb PE=3 SV=1 |

| | | | | | | | | | | | | |
|---|---|---|---|---|---|---|---|---|---|---|---|---|
| 68 | 87 | tr\|A0A0S1Z1V9\|A0A0S1Z1V9_RAT | 107.30 | 14 | 14 | 1.2547E6 | 5 | 5 | 6 | | 43002 | Cytochrome b OS=Rattus norvegicus OX=10116 GN=CYTB PE=3 SV=1 |
| 68 | 95 | tr\|D6NSR1\|D6NSR1_RAT | 107.30 | 14 | 14 | 1.2547E6 | 5 | 5 | 6 | | 43002 | Cytochrome b (Fragment) OS=Rattus norvegicus OX=10116 GN=cytb PE=3 SV=1 |
| 68 | 62 | tr\|Q8HIC4\|Q8HIC4_RAT | 107.30 | 14 | 14 | 1.2547E6 | 5 | 5 | 6 | | 43012 | Cytochrome b OS=Rattus norvegicus OX=10116 GN=Mt-cyb PE=3 SV=1 |
| 68 | 63 | tr\|A0A220DA02\|A0A220DA02_RAT | 107.30 | 14 | 14 | 1.2547E6 | 5 | 5 | 6 | | 42968 | Cytochrome b OS=Rattus norvegicus OX=10116 PE=3 SV=1 |
| 68 | 64 | tr\|A0A0S1Z1V4\|A0A0S1Z1V4_RAT | 107.30 | 14 | 14 | 1.2547E6 | 5 | 5 | 6 | | 43016 | Cytochrome b OS=Rattus norvegicus OX=10116 GN=CYTB PE=3 SV=1 |
| 68 | 88 | tr\|R9TKN1\|R9TKN1_RAT | 107.30 | 14 | 14 | 1.2547E6 | 5 | 5 | 6 | | 43012 | Cytochrome b OS=Rattus norvegicus OX=10116 GN=CYTB PE=3 SV=1 |
| 68 | 65 | P00159\|CYB_RAT | 107.30 | 14 | 14 | 1.2547E6 | 5 | 5 | 6 | | 43012 | Cytochrome b OS=Rattus norvegicus OX=10116 GN=Mt-Cyb PE=3 SV=3 |
| 68 | 66 | tr\|L0N311\|L0N311_RAT | 107.30 | 14 | 14 | 1.2547E6 | 5 | 5 | 6 | | 43045 | Cytochrome b (Fragment) OS=Rattus norvegicus OX=10116 GN=cytb PE=3 SV=1 |
| 68 | 67 | tr\|A0A220DA28\|A0A220DA28_RAT | 107.30 | 14 | 14 | 1.2547E6 | 5 | 5 | 6 | | 42970 | Cytochrome b OS=Rattus norvegicus OX=10116 PE=3 SV=1 |
| 68 | 68 | tr\|H2KXA0\|H2KXA0_RAT | 107.30 | 14 | 14 | 1.2547E6 | 5 | 5 | 6 | | 42978 | Cytochrome b (Fragment) OS=Rattus norvegicus OX=10116 GN=cytb PE=3 SV=1 |
| 68 | 101 | tr\|A0A096XNM4\|A0A096XNM4_RA | 107.30 | 14 | 14 | 1.2547E6 | 5 | 5 | 6 | | 43042 | Cytochrome b (Fragment) OS=Rattus norvegicus OX=10116 PE=3 SV=1 |
| 68 | 89 | tr\|D6NSP5\|D6NSP5_RAT | 107.30 | 14 | 14 | 1.2547E6 | 5 | 5 | 6 | | 43012 | Cytochrome b (Fragment) OS=Rattus norvegicus OX=10116 GN=cytb PE=3 SV=1 |
| 125 | 667 | tr\|B2RZ24\|B2RZ24_RAT | 106.95 | 5 | 5 | 3.3814E5 | 2 | 2 | 2 | | 47388 | Succinate-CoA ligase subunit beta (Fragment) OS=Rattus norvegicus OX=10116 GN=Sucla2 PE=2 SV=1 |
| 125 | 668 | tr\|F1LM47\|F1LM47_RAT | 106.95 | 5 | 5 | 3.3814E5 | 2 | 2 | 2 | | 50306 | Succinate--CoA ligase [ADP-forming] subunit beta mitochondrial OS=Rattus norvegicus OX=10116 GN=Sucla2 PE=1 SV=1 |
| 71 | 1219 | tr\|D3ZGM1\|D3ZGM1_RAT | 106.77 | 6 | 6 | 7.3668E5 | 5 | 5 | 5 | | 77891 | Pentatricopeptide repeat domain 3 OS=Rattus norvegicus OX=10116 GN=Ptcd3 PE=1 SV=1 |
| 71 | 1220 | tr\|A0A0G2JU15\|A0A0G2JU15_RAT | 106.77 | 6 | 6 | 7.3668E5 | 5 | 5 | 5 | | 84557 | Pentatricopeptide repeat domain 3 OS=Rattus norvegicus OX=10116 GN=Ptcd3 PE=1 SV=1 |
| 72 | 80 | P20788\|UCRI_RAT | 104.55 | 11 | 11 | 1.8463E6 | 3 | 3 | 5 | | 29446 | Cytochrome b-c1 complex subunit Rieske mitochondrial OS=Rattus norvegicus OX=10116 GN=Uqcrfs1 PE=1 SV=2 |
| 56 | 244 | P07824\|ARGI1_RAT | 104.16 | 13 | 13 | 2.3213E6 | 5 | 5 | 8 | | 34973 | Arginase-1 OS=Rattus norvegicus OX=10116 GN=Arg1 PE=1 SV=2 |
| 89 | 196 | Q63362\|NDUA5_RAT | 102.41 | 18 | 18 | 2.1437E6 | 4 | 4 | 4 | | 13412 | NADH dehydrogenase [ubiquinone] 1 alpha subcomplex subunit 5 OS=Rattus norvegicus OX=10116 GN=Ndufa5 PE=1 SV=3 |
| 49 | 31 | tr\|A0A1W2Q6F8\|A0A1W2Q6F8_RA | 102.30 | 16 | 16 | 4.4632E5 | 9 | 1 | 11 | Oxidation ( | 40544 | NADH dehydrogenase [ubiquinone] 1 alpha subcomplex subunit 10 mitochondrial OS=Rattus norvegicus OX=10116 GN=Ndufa10l1 PE=3 SV=1 |
| 81 | 112 | tr\|D4A3V2\|D4A3V2_RAT | 101.94 | 28 | 28 | 7.894E5 | 4 | 4 | 4 | | 15224 | NADH dehydrogenase [ubiquinone] 1 alpha subcomplex subunit 6 OS=Rattus norvegicus OX=10116 GN=Ndufa6 PE=1 SV=1 |
| 102 | 332 | Q9ER34\|ACON_RAT | 100.95 | 6 | 6 | 5.3443E5 | 3 | 3 | 3 | | 85433 | Aconitate hydratase mitochondrial OS=Rattus norvegicus OX=10116 GN=Aco2 PE=1 SV=2 |
| 54 | 245 | P18163\|ACSL1_RAT | 96.91 | 8 | 8 | 1.3122E6 | 7 | 7 | 9 | | 78179 | Long-chain-fatty-acid--CoA ligase 1 OS=Rattus norvegicus OX=10116 GN=Acsl1 PE=1 SV=1 |
| 63 | 5947 | Q924S5\|LONM_RAT | 96.84 | 3 | 3 | 1.1877E6 | 4 | 4 | 7 | | 105792 | Lon protease homolog mitochondrial OS=Rattus norvegicus OX=10116 GN=Lonp1 PE=2 SV=1 |
| 100 | 348 | P09139\|SPYA_RAT | 83.80 | 9 | 9 | 3.0747E5 | 3 | 3 | 3 | | 45834 | Serine--pyruvate aminotransferase mitochondrial OS=Rattus norvegicus OX=10116 GN=Agxt PE=1 SV=1 |
| 48 | 250 | tr\|Q06Q97\|Q06Q97_RAT | 83.44 | 12 | 12 | 1.3901E6 | 6 | 6 | 11 | | 38485 | NADH-ubiquinone oxidoreductase chain 2 OS=Rattus norvegicus OX=10116 GN=ND2 PE=3 SV=1 |
| 48 | 251 | tr\|A0A0A1G491\|A0A0A1G491_RAT | 83.44 | 12 | 12 | 1.3901E6 | 6 | 6 | 11 | | 38542 | NADH-ubiquinone oxidoreductase chain 2 OS=Rattus norvegicus OX=10116 GN=ND2 PE=3 SV=1 |
| 48 | 252 | tr\|Q5UAJ8\|Q5UAJ8_RAT | 83.44 | 12 | 12 | 1.3901E6 | 6 | 6 | 11 | | 38455 | NADH-ubiquinone oxidoreductase chain 2 OS=Rattus norvegicus OX=10116 GN=ND2 PE=3 SV=1 |
| 48 | 255 | tr\|Q06QH5\|Q06QH5_RAT | 83.44 | 12 | 12 | 1.3901E6 | 6 | 6 | 11 | | 38626 | NADH-ubiquinone oxidoreductase chain 2 OS=Rattus norvegicus OX=10116 GN=ND2 PE=3 SV=1 |
| 48 | 256 | tr\|Q06QB0\|Q06QB0_RAT | 83.44 | 12 | 12 | 1.3901E6 | 6 | 6 | 11 | | 38534 | NADH-ubiquinone oxidoreductase chain 2 OS=Rattus norvegicus OX=10116 GN=ND2 PE=3 SV=1 |
| 48 | 257 | tr\|D2E6P4\|D2E6P4_RAT | 83.44 | 12 | 12 | 1.3901E6 | 6 | 6 | 11 | | 38623 | NADH-ubiquinone oxidoreductase chain 2 OS=Rattus norvegicus OX=10116 GN=ND2 PE=3 SV=1 |
| 48 | 249 | tr\|Q06QE9\|Q06QE9_RAT | 83.44 | 12 | 12 | 1.3901E6 | 6 | 6 | 11 | | 38598 | NADH-ubiquinone oxidoreductase chain 2 OS=Rattus norvegicus OX=10116 GN=ND2 PE=3 SV=1 |
| 48 | 253 | tr\|Q8HID0\|Q8HID0_RAT | 83.44 | 12 | 12 | 1.3901E6 | 6 | 6 | 11 | | 38653 | NADH-ubiquinone oxidoreductase chain 2 OS=Rattus norvegicus OX=10116 GN=Mt-nd2 PE=3 SV=1 |
| 48 | 254 | P11662\|NU2M_RAT | 83.44 | 12 | 12 | 1.3901E6 | 6 | 6 | 11 | | 38653 | NADH-ubiquinone oxidoreductase chain 2 OS=Rattus norvegicus OX=10116 GN=Mtnd2 PE=3 SV=3 |
| 48 | 258 | tr\|Q8SEZ7\|Q8SEZ7_RAT | 83.44 | 12 | 12 | 1.3901E6 | 6 | 6 | 11 | | 38580 | NADH-ubiquinone oxidoreductase chain 2 (Fragment) OS=Rattus norvegicus OX=10116 GN=NADH2 PE=3 SV=1 |
| 151 | 283 | P80431\|COX7B_RAT | 82.69 | 29 | 29 | 6.8926E5 | 2 | 2 | 2 | | 8995 | Cytochrome c oxidase subunit 7B mitochondrial OS=Rattus norvegicus OX=10116 GN=Cox7b PE=1 SV=3 |
| 58 | 231 | tr\|Q5M949\|Q5M949_RAT | 80.79 | 14 | 14 | 2.2508E6 | 6 | 6 | 8 | | 28340 | Nipsnap homolog 3A (C. elegans) OS=Rattus norvegicus OX=10116 GN=Nipsnap3b PE=1 SV=1 |
| 61 | 11914 | P29419\|ATP5I_RAT | 79.60 | 27 | 27 | 3.0402E6 | 3 | 3 | 8 | | 8255 | ATP synthase subunit e mitochondrial OS=Rattus norvegicus OX=10116 GN=Atp5me PE=1 SV=3 |
| 77 | 346 | Q63750\|RM23_RAT | 79.17 | 19 | 19 | 5.4653E5 | 3 | 3 | 5 | | 17050 | 39S ribosomal protein L23 mitochondrial OS=Rattus norvegicus OX=10116 GN=Mrpl23 PE=2 SV=1 |
| 78 | 178 | tr\|D3ZZ21\|D3ZZ21_RAT | 77.40 | 12 | 12 | 7.0255E5 | 3 | 3 | 5 | | 15638 | NADH dehydrogenase (Ubiquinone) 1 beta subcomplex 6 (Predicted) OS=Rattus norvegicus OX=10116 GN=Ndufb6 PE=1 SV=1 |
| 50 | 171 | tr\|Q8SEZ8\|Q8SEZ8_RAT | 77.13 | 18 | 18 | 1.7499E6 | 8 | 8 | 11 | Oxidation ( | 36133 | NADH-ubiquinone oxidoreductase chain 1 (Fragment) OS=Rattus norvegicus OX=10116 GN=NADH1 PE=3 SV=1 |
| 50 | 172 | P03889\|NU1M_RAT | 77.13 | 18 | 18 | 1.7499E6 | 8 | 8 | 11 | Oxidation ( | 36145 | NADH-ubiquinone oxidoreductase chain 1 OS=Rattus norvegicus OX=10116 GN=Mtnd1 PE=1 SV=3 |

| | | | | | | | | | | | | |
|---|---|---|---|---|---|---|---|---|---|---|---|---|
| 50 | 173 | tr\|Q8HID1\|Q8HID1_RAT | 77.13 | 18 | 18 | 1.7499E6 | 8 | 8 | 11 | Oxidation ( | 36145 | NADH-ubiquinone oxidoreductase chain 1 OS=Rattus norvegicus OX=10116 GN=Mt-nd1 PE=3 SV=1 |
| 50 | 174 | tr\|D2E6L7\|D2E6L7_RAT | 77.13 | 18 | 18 | 1.7499E6 | 8 | 8 | 11 | Oxidation ( | 36062 | NADH-ubiquinone oxidoreductase chain 1 OS=Rattus norvegicus OX=10116 GN=ND1 PE=3 SV=1 |
| 74 | 182 | P10888\|COX41_RAT | 74.44 | 17 | 17 | 1.1557E6 | 4 | 4 | 5 | | 19515 | Cytochrome c oxidase subunit 4 isoform 1 mitochondrial OS=Rattus norvegicus OX=10116 GN=Cox4i1 PE=1 SV=1 |
| 75 | 194 | Q9R063\|PRDX5_RAT | 73.85 | 17 | 17 | 1.5202E6 | 4 | 4 | 5 | | 22179 | Peroxiredoxin-5 mitochondrial OS=Rattus norvegicus OX=10116 GN=Prdx5 PE=1 SV=1 |
| 75 | 195 | tr\|A0A0G2JSS8\|A0A0G2JSS8_RAT | 73.85 | 17 | 17 | 1.5202E6 | 4 | 4 | 5 | | 22207 | Peroxiredoxin OS=Rattus norvegicus OX=10116 GN=Prdx5 PE=1 SV=1 |
| 105 | 264 | P02770\|ALBU_RAT | 72.98 | 3 | 3 | 6.0016E5 | 3 | 3 | 3 | Formylatio | 68731 | Serum albumin OS=Rattus norvegicus OX=10116 GN=Alb PE=1 SV=2 |
| 105 | 263 | tr\|A0A0G2JSH5\|A0A0G2JSH5_RAT | 72.98 | 3 | 3 | 6.0016E5 | 3 | 3 | 3 | Formylatio | 68759 | Serum albumin OS=Rattus norvegicus OX=10116 GN=Alb PE=1 SV=1 |
| 69 | 148 | tr\|Q5RJN0\|Q5RJN0_RAT | 72.84 | 14 | 14 | 1.5232E6 | 4 | 4 | 6 | Oxidation ( | 23945 | NADH dehydrogenase (Ubiquinone) Fe-S protein 7 OS=Rattus norvegicus OX=10116 GN=Ndufs7 PE=1 SV=1 |
| 91 | 290 | tr\|Q4G067\|Q4G067_RAT | 68.98 | 8 | 8 | 1.0023E6 | 3 | 3 | 4 | | 37440 | Mitochondrial ribosomal protein L44 OS=Rattus norvegicus OX=10116 GN=Mrpl44 PE=1 SV=1 |
| 108 | 1292 | tr\|G3V6I4\|G3V6I4_RAT | 68.42 | 5 | 5 | 3.3735E5 | 2 | 2 | 3 | | 37791 | Mitochondrial amidoxime reducing component 1 OS=Rattus norvegicus OX=10116 GN=Marc1 PE=1 SV=1 |
| 82 | 3 | tr\|B2GV15\|B2GV15_RAT | 67.54 | 4 | 4 | 5.5589E5 | 2 | 2 | 4 | | 53274 | Dihydrolipoamide acetyltransferase component of pyruvate dehydrogenase complex OS=Rattus norvegicus OX=10116 GN=Dbt PE=1 SV=1 |
| 106 | 11911 | P21571\|ATP5J_RAT | 63.45 | 19 | 19 | 8.4415E5 | 2 | 2 | 3 | | 12494 | ATP synthase-coupling factor 6 mitochondrial OS=Rattus norvegicus OX=10116 GN=Atp5pf PE=1 SV=1 |
| 88 | 232 | tr\|M0R6J0\|M0R6J0_RAT | 61.62 | 6 | 6 | 4.5281E5 | 2 | 2 | 4 | | 38374 | Mitochondrial ribosomal protein L39 OS=Rattus norvegicus OX=10116 GN=Mrpl39 PE=1 SV=2 |
| 90 | 816 | tr\|Q5U2T0\|Q5U2T0_RAT | 60.31 | 5 | 5 | 1.4915E5 | 3 | 2 | 4 | | 44494 | Death associated protein 3 OS=Rattus norvegicus OX=10116 GN=Dap3 PE=2 SV=1 |
| 90 | 817 | tr\|F7EZZ0\|F7EZZ0_RAT | 60.31 | 5 | 5 | 1.4915E5 | 3 | 2 | 4 | | 45112 | Death-associated protein 3 OS=Rattus norvegicus OX=10116 GN=Dap3 PE=1 SV=1 |
| 90 | 1036 | tr\|A0A0G2K264\|A0A0G2K264_RAT | 60.31 | 5 | 5 | 1.4915E5 | 3 | 2 | 4 | | 46666 | Death-associated protein 3 OS=Rattus norvegicus OX=10116 GN=Dap3 PE=1 SV=1 |
| 87 | 233 | P29147\|BDH_RAT | 60.12 | 10 | 10 | 7.5888E5 | 4 | 3 | 4 | | 38202 | D-beta-hydroxybutyrate dehydrogenase mitochondrial OS=Rattus norvegicus OX=10116 GN=Bdh1 PE=1 SV=2 |
| 87 | 234 | tr\|A0A0G2JSH2\|A0A0G2JSH2_RAT | 60.12 | 10 | 10 | 7.5888E5 | 4 | 3 | 4 | | 38333 | 3-hydroxybutyrate dehydrogenase type 1 isoform CRA_a OS=Rattus norvegicus OX=10116 GN=Bdh1 PE=1 SV=1 |
| 120 | 6630 | tr\|F1LZW6\|F1LZW6_RAT | 58.01 | 2 | 2 | 2.8788E4 | 2 | 2 | 2 | | 54099 | Solute carrier family 25 member 13 OS=Rattus norvegicus OX=10116 GN=Slc25a13 PE=1 SV=2 |
| 120 | 11912 | tr\|A0A0G2K2J7\|A0A0G2K2J7_RAT | 58.01 | 3 | 3 | 2.8788E4 | 2 | 2 | 2 | | 41504 | Solute carrier family 25 member 12 OS=Rattus norvegicus OX=10116 GN=Slc25a12 PE=1 SV=1 |
| 120 | 11913 | tr\|F1LX07\|F1LX07_RAT | 58.01 | 2 | 2 | 2.8788E4 | 2 | 2 | 2 | | 71917 | Solute carrier family 25 member 12 OS=Rattus norvegicus OX=10116 GN=Slc25a12 PE=1 SV=3 |
| 103 | 322 | Q09073\|ADT2_RAT | 57.96 | 8 | 8 | 1.6523E6 | 3 | 3 | 3 | | 32901 | ADP/ATP translocase 2 OS=Rattus norvegicus OX=10116 GN=Slc25a5 PE=1 SV=3 |
| 76 | 399 | Q91XJ1\|BECN1_RAT | 57.72 | 3 | 3 | | 3 | 0 | 5 | | 51557 | Beclin-1 OS=Rattus norvegicus OX=10116 GN=Becn1 PE=1 SV=1 |
| 104 | 295 | Q8VID1\|DHRS4_RAT | 57.55 | 9 | 9 | 3.8865E5 | 3 | 3 | 3 | | 29822 | Dehydrogenase/reductase SDR family member 4 OS=Rattus norvegicus OX=10116 GN=Dhrs4 PE=2 SV=2 |
| 258 | 11920 | tr\|D4A3E8\|D4A3E8_RAT | 57.36 | 3 | 3 | 3.6939E5 | 1 | 1 | 1 | | 47648 | Mitochondrial ribosomal protein S27 OS=Rattus norvegicus OX=10116 GN=Mrps27 PE=1 SV=1 |
| 156 | 705 | Q64565\|AGT2_RAT | 57.04 | 3 | 3 | 3.5709E4 | 2 | 2 | 2 | | 57201 | Alanine--glyoxylate aminotransferase 2 mitochondrial OS=Rattus norvegicus OX=10116 GN=Agxt2 PE=1 SV=2 |
| 157 | 119 | tr\|D3ZE15\|D3ZE15_RAT | 56.60 | 7 | 7 | 1.7342E5 | 2 | 2 | 2 | | 16777 | NADH:ubiquinone oxidoreductase subunit A13 OS=Rattus norvegicus OX=10116 GN=Ndufa13 PE=1 SV=1 |
| 153 | 343 | tr\|D3ZX69\|D3ZX69_RAT | 55.13 | 5 | 5 | 4.1453E5 | 1 | 1 | 2 | | 29466 | 39S ribosomal protein L10 mitochondrial OS=Rattus norvegicus OX=10116 GN=Mrpl10 PE=1 SV=1 |
| 153 | 344 | P0C2C4\|RM10_RAT | 55.13 | 5 | 5 | 4.1453E5 | 1 | 1 | 2 | | 29940 | 39S ribosomal protein L10 mitochondrial OS=Rattus norvegicus OX=10116 GN=Mrpl10 PE=1 SV=1 |
| 142 | 267 | tr\|I6V4L9\|I6V4L9_RAT | 54.89 | 5 | 5 | 2.0885E5 | 1 | 1 | 2 | | 29844 | Cytochrome c oxidase subunit 3 OS=Rattus norvegicus OX=10116 GN=COX3 PE=3 SV=1 |
| 85 | 753 | P08011\|MGST1_RAT | 52.39 | 8 | 8 | 4.5451E5 | 3 | 3 | 4 | | 17472 | Microsomal glutathione S-transferase 1 OS=Rattus norvegicus OX=10116 GN=Mgst1 PE=1 SV=3 |
| 85 | 754 | tr\|B6DYQ4\|B6DYQ4_RAT | 52.39 | 8 | 8 | 4.5451E5 | 3 | 3 | 4 | | 17472 | Microsomal glutathione S-transferase OS=Rattus norvegicus OX=10116 GN=Mgst1 PE=2 SV=1 |
| 152 | 281 | tr\|A9UMW2\|A9UMW2_RAT | 52.35 | 31 | 31 | 1.4716E6 | 2 | 2 | 2 | Oxidation ( | 9255 | Ndufa3 protein (Fragment) OS=Rattus norvegicus OX=10116 GN=Ndufa3 PE=2 SV=1 |
| 152 | 272 | tr\|M0RB63\|M0RB63_RAT | 52.35 | 31 | 31 | 1.4716E6 | 2 | 2 | 2 | Oxidation ( | 9372 | RCG63041 OS=Rattus norvegicus OX=10116 GN=LOC684509 PE=4 SV=1 |
| 152 | 282 | tr\|A0A0G2KAA3\|A0A0G2KAA3_RA | 52.35 | 29 | 29 | 1.4716E6 | 2 | 2 | 2 | Oxidation ( | 10170 | NADH:ubiquinone oxidoreductase subunit A3 OS=Rattus norvegicus OX=10116 GN=Ndufa3 PE=1 SV=1 |
| 107 | 276 | tr\|A0A140TAG5\|A0A140TAG5_RAT | 51.63 | 3 | 3 | 6.2663E5 | 2 | 2 | 3 | | 67049 | MICOS complex subunit MIC60 OS=Rattus norvegicus OX=10116 GN=Immt PE=1 SV=1 |
| 107 | 277 | Q3KR86\|MIC60_RAT | 51.63 | 3 | 3 | 6.2663E5 | 2 | 2 | 3 | | 67177 | MICOS complex subunit Mic60 (Fragment) OS=Rattus norvegicus OX=10116 GN=Immt PE=1 SV=1 |
| 107 | 273 | tr\|A0A0G2JVH4\|A0A0G2JVH4_RAT | 51.63 | 3 | 3 | 6.2663E5 | 2 | 2 | 3 | | 86230 | MICOS complex subunit MIC60 OS=Rattus norvegicus OX=10116 GN=Immt PE=1 SV=1 |
| 259 | 11921 | tr\|D4ACA5\|D4ACA5_RAT | 50.85 | 9 | 9 | 2.5943E5 | 1 | 1 | 1 | | 18377 | Mitochondrial import inner membrane translocase subunit TIM17 OS=Rattus norvegicus OX=10116 GN=LOC100911130 PE=1 SV=1 |
| 95 | 380 | O70351\|HCD2_RAT | 49.73 | 9 | 9 | 1.8286E5 | 2 | 2 | 3 | | 27246 | 3-hydroxyacyl-CoA dehydrogenase type-2 OS=Rattus norvegicus OX=10116 GN=Hsd17b10 PE=1 SV=3 |
| 95 | 381 | tr\|B0BMW2\|B0BMW2_RAT | 49.73 | 9 | 9 | 1.8286E5 | 2 | 2 | 3 | | 27250 | 3-hydroxyacyl-CoA dehydrogenase type-2 OS=Rattus norvegicus OX=10116 GN=Hsd17b10 PE=1 SV=1 |
| 260 | 243 | tr\|D3ZS58\|D3ZS58_RAT | 49.55 | 11 | 11 | 2.7487E5 | 1 | 1 | 1 | | 10845 | NADH dehydrogenase [ubiquinone] 1 alpha subcomplex subunit 2 OS=Rattus norvegicus OX=10116 GN=Ndufa2 PE=1 SV=1 |

| | | | | | | | | | | | | | |
|---|---|---|---|---|---|---|---|---|---|---|---|---|---|
| 154 | 5939 | P56571\|ES1_RAT | 48.89 | 4 | 4 | | 0 | 1 | 1 | 2 | | 28173 | ES1 protein homolog mitochondrial OS=Rattus norvegicus OX=10116 PE=1 SV=2 |
| 158 | 396 | tr\|D3ZH23\|D3ZH23_RAT | 47.52 | 11 | 11 | 1.5168E4 | | 2 | 2 | 2 | Oxidation ( | 15195 | Similar to mitochondrial ribosomal protein L41 OS=Rattus norvegicus OX=10116 GN=RGD1560917 PE=4 SV=1 |
| 158 | 397 | Q5BJX1\|RM41_RAT | 47.52 | 11 | 11 | 1.5168E4 | | 2 | 2 | 2 | Oxidation ( | 15193 | 39S ribosomal protein L41 mitochondrial OS=Rattus norvegicus OX=10116 GN=Mrpl41 PE=1 SV=1 |
| 261 | 6074 | B1WC61\|ACAD9_RAT | 46.22 | 2 | 2 | 2.0868E5 | | 1 | 1 | 1 | | 68843 | Complex I assembly factor ACAD9 mitochondrial OS=Rattus norvegicus OX=10116 GN=Acad9 PE=1 SV=1 |
| 155 | 11923 | P32089\|TXTP_RAT | 45.33 | 3 | 3 | 1.2085E5 | | 1 | 1 | 2 | | 33835 | Tricarboxylate transport protein mitochondrial OS=Rattus norvegicus OX=10116 GN=Slc25a1 PE=1 SV=1 |
| 109 | 11922 | Q9JJW3\|ATPMD_RAT | 44.46 | 21 | 21 | 1.8356E6 | | 1 | 1 | 3 | | 6408 | ATP synthase membrane subunit DAPIT mitochondrial OS=Rattus norvegicus OX=10116 GN=Atp5md PE=1 SV=1 |
| 129 | 1045 | Q2V057\|HYPDH_RAT | 43.98 | 4 | 4 | 5.658E6 | | 2 | 2 | 2 | | 51002 | Hydroxyproline dehydrogenase OS=Rattus norvegicus OX=10116 GN=Prodh2 PE=2 SV=1 |
| 73 | 220 | P12075\|COX5B_RAT | 43.58 | 22 | 22 | 3.1884E6 | | 4 | 4 | 5 | | 13915 | Cytochrome c oxidase subunit 5B mitochondrial OS=Rattus norvegicus OX=10116 GN=Cox5b PE=1 SV=2 |
| 164 | 427 | tr\|G3V9J8\|G3V9J8_RAT | 42.96 | 2 | 2 | 2.1743E5 | | 1 | 1 | 1 | | 87521 | Glycerol-3-phosphate acyltransferase 1 mitochondrial OS=Rattus norvegicus OX=10116 GN=Gpam PE=1 SV=2 |
| 164 | 413 | tr\|A0A0G2K2U7\|A0A0G2K2U7_RA | 42.96 | 2 | 2 | 2.1743E5 | | 1 | 1 | 1 | | 93728 | Glycerol-3-phosphate acyltransferase 1 mitochondrial OS=Rattus norvegicus OX=10116 GN=Gpam PE=1 SV=1 |
| 164 | 414 | P97564\|GPAT1_RAT | 42.96 | 2 | 2 | 2.1743E5 | | 1 | 1 | 1 | | 93715 | Glycerol-3-phosphate acyltransferase 1 mitochondrial OS=Rattus norvegicus OX=10116 GN=Gpam PE=1 SV=3 |
| 201 | 354 | tr\|Q3B8N9\|Q3B8N9_RAT | 42.89 | 3 | 3 | 2.4401E5 | | 1 | 1 | 1 | | 32823 | Biphenyl hydrolase-like OS=Rattus norvegicus OX=10116 GN=Bphl PE=1 SV=1 |
| 32 | 368 | tr\|A0A0G2JWK2\|A0A0G2JWK2_RA | 42.20 | 1 | 1 | 3.0277E5 | | 1 | 1 | 19 | | 53049 | Methyl-CpG-binding protein 2 OS=Rattus norvegicus OX=10116 GN=Mecp2 PE=1 SV=1 |
| 32 | 369 | Q00566\|MECP2_RAT | 42.20 | 1 | 1 | 3.0277E5 | | 1 | 1 | 19 | | 53048 | Methyl-CpG-binding protein 2 OS=Rattus norvegicus OX=10116 GN=Mecp2 PE=1 SV=1 |
| 184 | 1422 | tr\|F1LPV8\|F1LPV8_RAT | 41.09 | 2 | 2 | 1.4723E5 | | 1 | 1 | 1 | | 46639 | Succinate--CoA ligase [GDP-forming] subunit beta mitochondrial OS=Rattus norvegicus OX=10116 GN=Suclg2 PE=1 SV=2 |
| 184 | 1423 | tr\|B1H270\|B1H270_RAT | 41.09 | 2 | 2 | 1.4723E5 | | 1 | 1 | 1 | | 46988 | Succinate--CoA ligase [GDP-forming] subunit beta mitochondrial OS=Rattus norvegicus OX=10116 GN=Suclg2 PE=2 SV=1 |
| 226 | 11915 | Q63159\|COQ3_RAT | 41.03 | 3 | 3 | 1.824E5 | | 1 | 1 | 1 | | 38708 | Ubiquinone biosynthesis O-methyltransferase mitochondrial OS=Rattus norvegicus OX=10116 GN=Coq3 PE=2 SV=2 |
| 229 | 309 | P63039\|CH60_RAT | 40.87 | 2 | 2 | 4.4997E5 | | 1 | 1 | 1 | | 60956 | 60 kDa heat shock protein mitochondrial OS=Rattus norvegicus OX=10116 GN=Hspd1 PE=1 SV=1 |
| 229 | 310 | tr\|A0A482IDN3\|A0A482IDN3_RAT | 40.87 | 2 | 2 | 4.4997E5 | | 1 | 1 | 1 | | 60956 | Hsp60 OS=Rattus norvegicus OX=10116 GN=Hspd1 PE=2 SV=1 |
| 128 | 293 | Q9WVK3\|PECR_RAT | 40.85 | 7 | 7 | 6.4124E5 | | 2 | 2 | 2 | Oxidation ( | 32433 | Peroxisomal trans-2-enoyl-CoA reductase OS=Rattus norvegicus OX=10116 GN=Pecr PE=2 SV=1 |
| 128 | 294 | tr\|A0A0G2JVG4\|A0A0G2JVG4_RA | 40.85 | 7 | 7 | 6.4124E5 | | 2 | 2 | 2 | Oxidation ( | 32737 | Peroxisomal trans-2-enoyl-CoA reductase OS=Rattus norvegicus OX=10116 GN=Pecr PE=1 SV=1 |
| 263 | 5949 | tr\|B2RYU0\|B2RYU0_RAT | 39.86 | 9 | 9 | 4.5063E5 | | 1 | 1 | 1 | | 11842 | NADH dehydrogenase (Ubiquinone) 1 beta subcomplex 2 (Predicted) isoform CRA_b OS=Rattus norvegicus OX=10116 GN=Ndufb2 PE=1 SV=1 |
| 262 | 350 | tr\|Q35733\|Q35733_RAT | 39.48 | 13 | 13 | 1.5553E5 | | 1 | 1 | 1 | | 8389 | NADH-ubiquinone oxidoreductase chain 6 (Fragment) OS=Rattus norvegicus OX=10116 PE=3 SV=1 |
| 262 | 351 | tr\|Q06QD9\|Q06QD9_RAT | 39.48 | 6 | 6 | 1.5553E5 | | 1 | 1 | 1 | | 18943 | NADH-ubiquinone oxidoreductase chain 6 OS=Rattus norvegicus OX=10116 GN=ND6 PE=3 SV=1 |
| 262 | 352 | tr\|Q7HKW2\|Q7HKW2_RAT | 39.48 | 6 | 6 | 1.5553E5 | | 1 | 1 | 1 | | 18957 | NADH-ubiquinone oxidoreductase chain 6 OS=Rattus norvegicus OX=10116 GN=NADH6 PE=3 SV=1 |
| 135 | 511 | P32198\|CPT1A_RAT | 38.74 | 2 | 2 | 3.6389E5 | | 2 | 1 | 2 | | 88126 | Carnitine O-palmitoyltransferase 1 liver isoform OS=Rattus norvegicus OX=10116 GN=Cpt1a PE=1 SV=2 |
| 83 | 601 | Q5BJP6\|RRF2M_RAT | 38.72 | 2 | 2 | 1.6099E6 | | 2 | 2 | 4 | Formylatio | 85914 | Ribosome-releasing factor 2 mitochondrial OS=Rattus norvegicus OX=10116 GN=Gfm2 PE=2 SV=2 |
| 83 | 602 | tr\|F1LMZ4\|F1LMZ4_RAT | 38.72 | 2 | 2 | 1.6099E6 | | 2 | 2 | 4 | Formylatio | 85914 | Ribosome-releasing factor 2 mitochondrial OS=Rattus norvegicus OX=10116 GN=Gfm2 PE=1 SV=2 |
| 84 | 286 | tr\|A9UMV9\|A9UMV9_RAT | 38.55 | 11 | 11 | 1.8232E6 | | 1 | 1 | 4 | | 12500 | NADH:ubiquinone oxidoreductase subunit A7 OS=Rattus norvegicus OX=10116 GN=Ndufa7 PE=1 SV=1 |
| 264 | 1169 | Q9Z2Y0\|KEG1_RAT | 37.65 | 3 | 3 | 1.1821E5 | | 1 | 1 | 1 | | 34016 | Glycine N-acyltransferase-like protein Keg1 OS=Rattus norvegicus OX=10116 GN=Keg1 PE=1 SV=2 |
| 264 | 1170 | tr\|B1H250\|B1H250_RAT | 37.65 | 3 | 3 | 1.1821E5 | | 1 | 1 | 1 | | 33986 | Glycine-N-acyltransferase-like 1 OS=Rattus norvegicus OX=10116 GN=Glyatl1 PE=1 SV=1 |
| 264 | 1171 | Q5PQT3\|GLYAT_RAT | 37.65 | 3 | 3 | 1.1821E5 | | 1 | 1 | 1 | | 33899 | Glycine N-acyltransferase OS=Rattus norvegicus OX=10116 GN=Glyat PE=2 SV=1 |
| 264 | 1172 | tr\|A4PB92\|A4PB92_RAT | 37.65 | 3 | 3 | 1.1821E5 | | 1 | 1 | 1 | | 33899 | Glycine N-acyltransferase OS=Rattus norvegicus OX=10116 GN=Glyat PE=2 SV=1 |
| 264 | 1173 | tr\|A0A0G2JWI5\|A0A0G2JWI5_RAT | 37.65 | 3 | 3 | 1.1821E5 | | 1 | 1 | 1 | | 39453 | Glycine N-acyltransferase OS=Rattus norvegicus OX=10116 GN=Glyat PE=1 SV=1 |
| 204 | 623 | tr\|G3V6I5\|G3V6I5_RAT | 36.15 | 3 | 3 | 2.9761E5 | | 1 | 1 | 1 | | 46777 | DnaJ heat shock protein family (Hsp40) member A3 OS=Rattus norvegicus OX=10116 GN=Dnaja3 PE=1 SV=2 |
| 204 | 624 | tr\|Q2TVU3\|Q2TVU3_RAT | 36.15 | 3 | 3 | 2.9761E5 | | 1 | 1 | 1 | | 49412 | TID1 OS=Rattus norvegicus OX=10116 GN=Dnaja3 PE=2 SV=1 |
| 204 | 625 | tr\|A0A0G2K4Y1\|A0A0G2K4Y1_RA | 36.15 | 3 | 3 | 2.9761E5 | | 1 | 1 | 1 | | 49416 | DnaJ heat shock protein family (Hsp40) member A3 OS=Rattus norvegicus OX=10116 GN=Dnaja3 PE=1 SV=1 |
| 204 | 626 | tr\|Q2UZS7\|Q2UZS7_RAT | 36.15 | 2 | 2 | 2.9761E5 | | 1 | 1 | 1 | | 52399 | Tid-1 long isoform OS=Rattus norvegicus OX=10116 GN=Dnaja3 PE=2 SV=1 |
| 204 | 627 | tr\|A0A0G2K5E4\|A0A0G2K5E4_RA | 36.15 | 2 | 2 | 2.9761E5 | | 1 | 1 | 1 | | 52403 | DnaJ heat shock protein family (Hsp40) member A3 OS=Rattus norvegicus OX=10116 GN=Dnaja3 PE=1 SV=1 |
| 228 | 162 | tr\|Q5EBA4\|Q5EBA4_RAT | 34.65 | 4 | 4 | 1.1944E5 | | 1 | 1 | 1 | | 33215 | Nipsnap1 protein (Fragment) OS=Rattus norvegicus OX=10116 GN=Nipsnap1 PE=2 SV=1 |
| 228 | 163 | tr\|G3V728\|G3V728_RAT | 34.65 | 4 | 4 | 1.1944E5 | | 1 | 1 | 1 | | 33346 | 4-nitrophenylphosphatase domain and non-neuronal SNAP25-like protein homolog 1 (C. elegans) isoform CRA_b OS=Rattus norvegicus OX=10116 GN=Nipsnap1 PE=1 SV=1 |
| 205 | 762 | B0BN56\|RT31_RAT | 33.49 | 4 | 4 | 2.4645E5 | | 1 | 1 | 1 | | 43962 | 28S ribosomal protein S31 mitochondrial OS=Rattus norvegicus OX=10116 GN=Mrps31 PE=2 SV=1 |

| | | | | | | | | | | | | | |
|---|---|---|---|---|---|---|---|---|---|---|---|---|---|
| 232 | 11918 | tr\|D3ZXF8\|D3ZXF8_RAT | 33.45 | 5 | 5 | 2.937E5 | | 1 | 1 | 1 | | 17575 | Mitochondrial ribosomal protein L43 OS=Rattus norvegicus OX=10116 GN=Mrpl43 PE=1 SV=3 |
| 265 | 6121 | tr\|A0A0G2K401\|A0A0G2K401_RAT | 33.08 | 1 | 1 | 2.8489E5 | | 1 | 1 | 1 | | 79812 | Propionyl-CoA carboxylase alpha chain mitochondrial OS=Rattus norvegicus OX=10116 GN=Pcca PE=1 SV=1 |
| 265 | 6122 | tr\|A0A0G2K5P9\|A0A0G2K5P9_RAT | 33.08 | 1 | 1 | 2.8489E5 | | 1 | 1 | 1 | | 81413 | Propionyl-CoA carboxylase alpha chain mitochondrial OS=Rattus norvegicus OX=10116 GN=Pcca PE=1 SV=1 |
| 265 | 1119 | P14882\|PCCA_RAT | 33.08 | 1 | 1 | 2.8489E5 | | 1 | 1 | 1 | | 81623 | Propionyl-CoA carboxylase alpha chain mitochondrial OS=Rattus norvegicus OX=10116 GN=Pcca PE=1 SV=3 |
| 227 | 11916 | Q5I0C5\|FMT_RAT | 32.46 | 2 | 2 | 7.0512E5 | | 1 | 1 | 1 | Formylatio | 42997 | Methionyl-tRNA formyltransferase mitochondrial OS=Rattus norvegicus OX=10116 GN=Mtfmt PE=2 SV=1 |
| 139 | 1477 | tr\|D4AE90\|D4AE90_RAT | 31.93 | 3 | 3 | | 0 | 2 | 1 | 2 | Formylatio | 50113 | RCC1-like OS=Rattus norvegicus OX=10116 GN=Rcc1l PE=1 SV=1 |
| 233 | 5951 | tr\|B2RZ57\|B2RZ57_RAT | 31.74 | 7 | 7 | 1.8033E5 | | 1 | 1 | 1 | | 20651 | Mitochondrial ribosomal protein L18 OS=Rattus norvegicus OX=10116 GN=Mrpl18 PE=1 SV=1 |
| 136 | 1546 | Q9EPJ3\|RT26_RAT | 31.65 | 12 | 12 | 6.1742E4 | | 2 | 2 | 2 | Oxidation ( | 23350 | 28S ribosomal protein S26 mitochondrial OS=Rattus norvegicus OX=10116 GN=Mrps26 PE=1 SV=1 |
| 101 | 6035 | Q68FU7\|COQ6_RAT | 31.59 | 3 | 3 | 3.8314E5 | | 2 | 2 | 3 | | 51496 | Ubiquinone biosynthesis monooxygenase COQ6 mitochondrial OS=Rattus norvegicus OX=10116 GN=Coq6 PE=2 SV=1 |
| 119 | 1285 | Q80ZG6\|BBC3_RAT | 31.46 | 6 | 6 | 7.01E6 | | 2 | 2 | 2 | | 20718 | Bcl-2-binding component 3 OS=Rattus norvegicus OX=10116 GN=Bbc3 PE=1 SV=1 |
| 110 | 374 | P48037\|ANXA6_RAT | 31.24 | 1 | 1 | 1.6184E6 | | 1 | 1 | 3 | | 75754 | Annexin A6 OS=Rattus norvegicus OX=10116 GN=Anxa6 PE=1 SV=2 |
| 110 | 375 | tr\|Q6IMZ3\|Q6IMZ3_RAT | 31.24 | 1 | 1 | 1.6184E6 | | 1 | 1 | 3 | | 75756 | Annexin OS=Rattus norvegicus OX=10116 GN=Anxa6 PE=1 SV=1 |
| 266 | 11924 | tr\|A9UMV2\|A9UMV2_RAT | 31.09 | 5 | 5 | 1.4656E5 | | 1 | 1 | 1 | | 18742 | Mitochondrial ribosomal protein S24 OS=Rattus norvegicus OX=10116 GN=Mrps24 PE=1 SV=1 |
| 266 | 11925 | tr\|A0A0G2K9G3\|A0A0G2K9G3_RAT | 31.09 | 4 | 4 | 1.4656E5 | | 1 | 1 | 1 | | 19827 | Mitochondrial ribosomal protein S24 OS=Rattus norvegicus OX=10116 GN=Mrps24 PE=1 SV=1 |
| 267 | 1612 | tr\|D3ZY44\|D3ZY44_RAT | 30.43 | 4 | 4 | 3.7195E4 | | 1 | 1 | 1 | | 32176 | Mitochondrial ribosomal protein S2 OS=Rattus norvegicus OX=10116 GN=Mrps2 PE=1 SV=1 |
| 202 | 1984 | tr\|B0BMT9\|B0BMT9_RAT | 30.авг | 2 | 2 | 1.0593E5 | | 1 | 1 | 1 | | 50202 | Sqrdl protein OS=Rattus norvegicus OX=10116 GN=Sqor PE=1 SV=1 |
| 111 | 11926 | D3ZAF6\|ATPK_RAT | 29.98 | 10 | 10 | 1.2867E6 | | 1 | 1 | 3 | | 10452 | ATP synthase subunit f mitochondrial OS=Rattus norvegicus OX=10116 GN=Atp5mf PE=1 SV=1 |
| 141 | 16 | tr\|F1LN92\|F1LN92_RAT | 29.83 | 2 | 2 | 2.4553E6 | | 2 | 2 | 2 | | 89352 | AFG3-like matrix AAA peptidase subunit 2 OS=Rattus norvegicus OX=10116 GN=Afg3l2 PE=1 SV=1 |
| 143 | 347 | P80433\|COX8A_RAT | 29.79 | 14 | 14 | 2.4312E5 | | 1 | 1 | 2 | | 7672 | Cytochrome c oxidase subunit 8A mitochondrial OS=Rattus norvegicus OX=10116 GN=Cox8a PE=1 SV=3 |
| 79 | 428 | tr\|D4A1D3\|D4A1D3_RAT | 28.64 | 0 | 0 | 3.0836E6 | | 2 | 1 | 4 | | 521487 | Sacsin molecular chaperone OS=Rattus norvegicus OX=10116 GN=Sacs PE=1 SV=2 |
| 137 | 338 | tr\|A0A0G2JYU2\|A0A0G2JYU2_RAT | 28.62 | 4 | 4 | 8.1859E5 | | 1 | 1 | 2 | | 20750 | Mitochondrial ribosomal protein L11 OS=Rattus norvegicus OX=10116 GN=mrpl11 PE=1 SV=1 |
| 137 | 339 | Q5XIE3\|RM11_RAT | 28.62 | 4 | 4 | 8.1859E5 | | 1 | 1 | 2 | | 22420 | 39S ribosomal protein L11 mitochondrial OS=Rattus norvegicus OX=10116 GN=Mrpl11 PE=2 SV=1 |
| 140 | 406 | tr\|D3ZD23\|D3ZD23_RAT | 26.82 | 1 | 1 | 3.7192E5 | | 1 | 1 | 2 | | 67300 | ATP-binding cassette subfamily E member 1 OS=Rattus norvegicus OX=10116 GN=Abce1 PE=1 SV=1 |
| 270 | 11928 | tr\|B2GV62\|B2GV62_RAT | 26.49 | 7 | 7 | 2.3788E5 | | 1 | 1 | 1 | | 17526 | Mitochondrial ribosomal protein L20 OS=Rattus norvegicus OX=10116 GN=Mrpl20 PE=1 SV=1 |
| 231 | 6178 | P06761\|BIP_RAT | 25.61 | 1 | 1 | 1.196E5 | | 1 | 1 | 1 | | 72347 | Endoplasmic reticulum chaperone BiP OS=Rattus norvegicus OX=10116 GN=Hspa5 PE=1 SV=1 |
| 231 | 11940 | P55063\|HS71L_RAT | 25.61 | 1 | 1 | 1.196E5 | | 1 | 1 | 1 | | 70549 | Heat shock 70 kDa protein 1-like OS=Rattus norvegicus OX=10116 GN=Hspa1l PE=2 SV=2 |
| 231 | 6652 | P0DMW0\|HS71A_RAT | 25.61 | 1 | 1 | 1.196E5 | | 1 | 1 | 1 | | 70185 | Heat shock 70 kDa protein 1A OS=Rattus norvegicus OX=10116 GN=Hspa1a PE=2 SV=1 |
| 231 | 6653 | P0DMW1\|HS71B_RAT | 25.61 | 1 | 1 | 1.196E5 | | 1 | 1 | 1 | | 70185 | Heat shock 70 kDa protein 1B OS=Rattus norvegicus OX=10116 GN=Hspa1b PE=2 SV=1 |
| 131 | 495 | tr\|D4ACN9\|D4ACN9_RAT | 25.39 | 4 | 4 | 6.4002E5 | | 2 | 1 | 2 | | 34267 | Solute carrier family 25 member 36 OS=Rattus norvegicus OX=10116 GN=Slc25a36 PE=3 SV=1 |
| 271 | 6717 | tr\|A0A0G2K2R2\|A0A0G2K2R2_RAT | 24.89 | 4 | 4 | 9.1766E4 | | 1 | 1 | 1 | | 30788 | Enoyl-CoA delta isomerase 1 mitochondrial OS=Rattus norvegicus OX=10116 GN=Eci1 PE=1 SV=1 |
| 271 | 6718 | P23965\|ECI1_RAT | 24.89 | 4 | 4 | 9.1766E4 | | 1 | 1 | 1 | | 32254 | Enoyl-CoA delta isomerase 1 mitochondrial OS=Rattus norvegicus OX=10116 GN=Eci1 PE=1 SV=1 |
| 271 | 6719 | tr\|Q68G41\|Q68G41_RAT | 24.89 | 4 | 4 | 9.1766E4 | | 1 | 1 | 1 | | 32264 | Dodecenoyl-Coenzyme A delta isomerase (3 2 trans-enoyl-Coenzyme A isomerase) OS=Rattus norvegicus OX=10116 GN=Eci1 PE=1 SV=1 |
| 271 | 6720 | tr\|Q64592\|Q64592_RAT | 24.89 | 4 | 4 | 9.1766E4 | | 1 | 1 | 1 | | 32894 | Delta3 delta2-enoyl-CoA isomerase OS=Rattus norvegicus OX=10116 PE=2 SV=1 |
| 174 | 382 | Q01062\|PDE2A_RAT | 24.86 | 2 | 2 | 5.9259E5 | | 1 | 1 | 1 | | 104664 | cGMP-dependent 3' 5'-cyclic phosphodiesterase OS=Rattus norvegicus OX=10116 GN=Pde2a PE=1 SV=2 |
| 174 | 383 | tr\|F8WFW5\|F8WFW5_RAT | 24.86 | 1 | 1 | 5.9259E5 | | 1 | 1 | 1 | | 105234 | Phosphodiesterase OS=Rattus norvegicus OX=10116 GN=Pde2a PE=1 SV=1 |
| 203 | 1261 | tr\|Q5RKL4\|Q5RKL4_RAT | 23.97 | 1 | 1 | | 0 | 1 | 1 | 1 | | 95977 | Dimethylglycine dehydrogenase OS=Rattus norvegicus OX=10116 GN=Dmgdh PE=1 SV=1 |
| 203 | 1262 | Q63342\|M2GD_RAT | 23.97 | 1 | 1 | | 0 | 1 | 1 | 1 | | 96047 | Dimethylglycine dehydrogenase mitochondrial OS=Rattus norvegicus OX=10116 GN=Dmgdh PE=1 SV=1 |
| 203 | 1268 | tr\|A0A0G2K9Y2\|A0A0G2K9Y2_RAT | 23.97 | 1 | 1 | | 0 | 1 | 1 | 1 | | 98864 | Dimethylglycine dehydrogenase mitochondrial OS=Rattus norvegicus OX=10116 GN=Dmgdh PE=1 SV=1 |
| 272 | 11942 | tr\|A0A0U1RRX6\|A0A0U1RRX6_RAT | 23.63 | 9 | 9 | 6.3576E4 | | 1 | 1 | 1 | | 10726 | 5-demethoxyubiquinone hydroxylase mitochondrial (Fragment) OS=Rattus norvegicus OX=10116 GN=Coq7 PE=1 SV=1 |
| 272 | 11943 | tr\|G3V879\|G3V879_RAT | 23.63 | 5 | 5 | 6.3576E4 | | 1 | 1 | 1 | | 20126 | 5-demethoxyubiquinone hydroxylase mitochondrial OS=Rattus norvegicus OX=10116 GN=Coq7 PE=1 SV=4 |
| 273 | 192 | tr\|B5DEL8\|B5DEL8_RAT | 23.49 | 14 | 14 | 2.8915E5 | | 1 | 1 | 1 | | 12700 | NADH dehydrogenase (Ubiquinone) Fe-S protein 5 OS=Rattus norvegicus OX=10116 GN=Ndufs5 PE=1 SV=1 |
| 273 | 193 | tr\|A0A0G2JZJ9\|A0A0G2JZJ9_RAT | 23.49 | 14 | 14 | 2.8915E5 | | 1 | 1 | 1 | | 12730 | Uncharacterized protein OS=Rattus norvegicus OX=10116 PE=4 SV=1 |

| | | | | | | | | | | | | | |
|---|---|---|---|---|---|---|---|---|---|---|---|---|---|
| 277 | 400 | tr\|D4A7Q5\|D4A7Q5_RAT | 22.34 | 1 | 1 | 2.7474E5 | | 1 | 1 | 1 | | 59940 | DEAD (Asp-Glu-Ala-Asp) box polypeptide 28 (Predicted) OS=Rattus norvegicus OX=10116 GN=Ddx28 PE=4 SV=1 |
| 114 | 501 | tr\|D4A4K4\|D4A4K4_RAT | 21.96 | 0 | 0 | 4.2781E6 | | 1 | 1 | 2 | | 418626 | Vacuolar protein sorting 13 homolog C OS=Rattus norvegicus OX=10116 GN=Vps13c PE=1 SV=2 |
| 276 | 6337 | P17178\|CP27A_RAT | 21.85 | 2 | 2 | 1.4126E5 | | 1 | 1 | 1 | | 60733 | Sterol 26-hydroxylase mitochondrial OS=Rattus norvegicus OX=10116 GN=Cyp27a1 PE=1 SV=1 |
| 276 | 6338 | tr\|A0A0H2UHN7\|A0A0H2UHN7_RA | 21.85 | 2 | 2 | 1.4126E5 | | 1 | 1 | 1 | | 62492 | RCG24013 isoform CRA_a OS=Rattus norvegicus OX=10116 GN=Cyp27a1 PE=1 SV=1 |
| 86 | 403 | Q01728\|NAC1_RAT | 21.61 | 1 | 1 | | 7.88E+06 | 1 | 1 | 4 | Formylatio | 108184 | Sodium/calcium exchanger 1 OS=Rattus norvegicus OX=10116 GN=Slc8a1 PE=1 SV=3 |
| 86 | 404 | P70549\|NAC3_RAT | 21.61 | 1 | 1 | 7.8776E6 | | 1 | 1 | 4 | Formylatio | 103163 | Sodium/calcium exchanger 3 OS=Rattus norvegicus OX=10116 GN=Slc8a3 PE=1 SV=1 |
| 219 | 1309 | tr\|B2GUW4\|B2GUW4_RAT | 21.49 | 1 | 1 | | | 1 | 0 | 1 | | 74060 | Exdl2 protein OS=Rattus norvegicus OX=10116 GN=Exd2 PE=2 SV=1 |
| 279 | 6661 | M0R7Z9\|PLIN5_RAT | 21.40 | 1 | 1 | | 0 | 1 | 1 | 1 | | 51805 | Perilipin-5 OS=Rattus norvegicus OX=10116 GN=Plin5 PE=1 SV=1 |
| 132 | 471 | tr\|A0A0G2JYY2\|A0A0G2JYY2_RAT | 21.30 | 2 | 2 | 3.8014E6 | | 1 | 1 | 2 | | 62571 | 5-aminolevulinate synthase OS=Rattus norvegicus OX=10116 GN=Alas1 PE=1 SV=1 |
| 132 | 472 | tr\|A0A0G2K962\|A0A0G2K962_RA | 21.30 | 2 | 2 | 3.8014E6 | | 1 | 1 | 2 | | 71039 | 5-aminolevulinate synthase OS=Rattus norvegicus OX=10116 GN=Alas1 PE=1 SV=1 |
| 132 | 473 | P13195\|HEM1_RAT | 21.30 | 2 | 2 | 3.8014E6 | | 1 | 1 | 2 | | 71021 | 5-aminolevulinate synthase nonspecific mitochondrial OS=Rattus norvegicus OX=10116 GN=Alas1 PE=2 SV=2 |
| 236 | 1390 | Q811R2\|PRGC2_RAT | 20.74 | 1 | 1 | 2.2862E5 | | 1 | 1 | 1 | | 111589 | Peroxisome proliferator-activated receptor gamma coactivator 1-beta OS=Rattus norvegicus OX=10116 GN=Ppargc1b PE=2 SV=2 |
| 196 | 733 | tr\|F1MA54\|F1MA54_RAT | 20.72 | 1 | 1 | 1.6936E5 | | 1 | 1 | 1 | | 49099 | Protein-serine/threonine kinase OS=Rattus norvegicus OX=10116 GN=Pdk1 PE=1 SV=2 |
| 196 | 734 | Q63065\|PDK1_RAT | 20.72 | 1 | 1 | 1.6936E5 | | 1 | 1 | 1 | | 49081 | [Pyruvate dehydrogenase (acetyl-transferring)] kinase isozyme 1 mitochondrial OS=Rattus norvegicus OX=10116 GN=Pdk1 PE=1 SV=1 |
| 196 | 735 | tr\|Q5FVT5\|Q5FVT5_RAT | 20.72 | 1 | 1 | 1.6936E5 | | 1 | 1 | 1 | | 49095 | Protein-serine/threonine kinase OS=Rattus norvegicus OX=10116 GN=Pdk1 PE=2 SV=1 |
| 196 | 1109 | tr\|B5DFI9\|B5DFI9_RAT | 20.72 | 1 | 1 | 1.6936E5 | | 1 | 1 | 1 | | 47944 | Protein-serine/threonine kinase OS=Rattus norvegicus OX=10116 GN=Pdk3 PE=2 SV=1 |
| 280 | 300 | tr\|A0A0G2K9B4\|A0A0G2K9B4_RA | 20.68 | 4 | 4 | 5.2364E5 | | 1 | 1 | 1 | | 33652 | Mitochondrial ribosomal protein L15 OS=Rattus norvegicus OX=10116 GN=Mrpl15 PE=1 SV=1 |
| 281 | 226 | tr\|D3ZXF9\|D3ZXF9_RAT | 20.30 | 4 | 4 | 5.2364E5 | | 1 | 1 | 1 | | 29441 | Mitochondrial ribosomal protein L12 OS=Rattus norvegicus OX=10116 GN=Mrpl12 PE=1 SV=1 |
| 97 | 531 | P02563\|MYH6_RAT | 20.28 | 1 | 1 | 7.7659E6 | | 1 | 1 | 3 | Formylatio | 223506 | Myosin-6 OS=Rattus norvegicus OX=10116 GN=Myh6 PE=1 SV=2 |
