## Supplementary material for "Permeability transition pore-related changes in the proteome and channel activity of ATP synthase dimers and monomers": RLM SM-BM Control Monomer of V compl.

| Protein Gro | Protein ID | Accession | -10lgP | Coverage (% | Coverage (% | Area Sample 7 | #Peptides | #Unique | #Spec Sam | PTM | Avg. Mass | Description |
|---|---|---|---|---|---|---|---|---|---|---|---|---|
| 2 | 17 | P10719\|ATPB_RAT | 404.32 | 61 | 61 | 3757700000 | 202 | 197 | 966 | Oxidation (M) | | |
| 2 | 18 | tr\|G3V6D3\|G3V6D3_RAT | 404.32 | 61 | 61 | 3757700000 | 202 | 197 | 966 | Oxidation (M) | | |
| 1 | 20 | P15999\|ATPA_RAT | 363.73 | 59 | 59 | 3050800000 | 199 | 194 | 1112 | Carbamidomethylation | | |
| 1 | 21 | tr\|F1LP05\|F1LP05_RAT | 363.73 | 59 | 59 | 3050800000 | 199 | 194 | 1112 | Carbamidomethylation | | |
| 3 | 35 | P52873\|PYC_RAT | 360.14 | 39 | 39 | 425370000 | 153 | 150 | 375 | Oxidation ( | 129777 | Pyruvate carboxylase mitochondrial OS=Rattus norvegicus OX=10116 GN=Pc PE=1 SV=2 |
| 3 | 34 | tr\|A0A0G2JTL5\|A0A0G2J | 360.14 | 36 | 36 | 425370000 | 153 | 150 | 375 | Oxidation ( | 140005 | Pyruvate carboxylase mitochondrial OS=Rattus norvegicus OX=10116 GN=Pc PE=1 SV=1 |
| 5 | 280 | P31399\|ATP5H_RAT | 326.71 | 73 | 73 | 306560000 | 58 | 56 | 168 | Carbamidomethylation | | |
| 4 | 301 | P35435\|ATPG_RAT | 297.02 | 56 | 56 | 433430000 | 67 | 66 | 204 | Oxidation ( | 30191 | ATP synthase subunit gamma mitochondrial OS=Rattus norvegicus OX=10116 GN=Atp5f1c PE=1 SV=2 |
| 4 | 297 | tr\|Q6QI09\|Q6QI09_RAT | 297.02 | 25 | 25 | 433430000 | 67 | 66 | 204 | Oxidation ( | 67721 | ATP synthase subunit gamma mitochondrial OS=Rattus norvegicus OX=10116 GN=Taf3 PE=1 SV=1 |
| 7 | 8 | P07756\|CPSM_RAT | 252 | 25 | 25 | 93640000 | 70 | 68 | 131 | Carbamidomethylation | | |
| 6 | 268 | P19511\|AT5F1_RAT | 226.48 | 29 | 29 | 376000000 | 36 | 36 | 134 | Formylatio | 28869 | ATP synthase F(0) complex subunit B1 mitochondrial OS=Rattus norvegicus OX=10116 GN=Atp5pb PE=1 SV=1 |
| 18 | 289 | tr\|D3ZFJ6\|D3ZFJ6_RAT | 222.73 | 17 | 17 | 15061000 | 17 | 17 | 27 | | 60420 | Lactamase beta OS=Rattus norvegicus OX=10116 GN=Lactb PE=1 SV=1 |
| 8 | 5938 | Q06647\|ATPO_RAT | 218.5 | 48 | 48 | 123830000 | 45 | 43 | 115 | Carbamidomethylation | | |
| 9 | 5937 | Q6PDU7\|ATP5L_RAT | 215.17 | 39 | 39 | 109760000 | 25 | 25 | 66 | Oxidation ( | 11433 | ATP synthase subunit g mitochondrial OS=Rattus norvegicus OX=10116 GN=Atp5mg PE=1 SV=2 |
| 15 | 15 | P00507\|AATM_RAT | 208.27 | 24 | 24 | 35689000 | 22 | 21 | 36 | Oxidation ( | 47314 | Aspartate aminotransferase mitochondrial OS=Rattus norvegicus OX=10116 GN=Got2 PE=1 SV=2 |
| 10 | 387 | tr\|G3V7Y3\|G3V7Y3_RAT | 203.07 | 36 | 36 | 122500000 | 19 | 19 | 55 | Oxidation ( | 17563 | ATP synthase subunit delta mitochondrial OS=Rattus norvegicus OX=10116 GN=Atp5f1d PE=1 SV=1 |
| 11 | 1 | Q02253\|MMSA_RAT | 196.28 | 19 | 19 | 15340000 | 26 | 25 | 51 | Oxidation ( | 57808 | Methylmalonate-semialdehyde dehydrogenase [acylating] mitochondrial OS=Rattus norvegicus OX=10116 GN=Aldh6a1 PE=1 SV=1 |
| 11 | 2 | tr\|G3V7J0\|G3V7J0_RAT | 196.28 | 19 | 19 | 15340000 | 26 | 25 | 51 | Oxidation ( | 57748 | Aldehyde dehydrogenase family 6 subfamily A1 isoform CRA_b OS=Rattus norvegicus OX=10116 GN=Aldh6a1 PE=1 SV=1 |
| 14 | 266 | P04762\|CATA_RAT | 195.48 | 27 | 27 | 19116000 | 21 | 21 | 36 | Oxidation ( | 59757 | Catalase OS=Rattus norvegicus OX=10116 GN=Cat PE=1 SV=3 |
| 21 | 14 | P10860\|DHE3_RAT | 193.93 | 20 | 20 | 8.45E+06 | 17 | 17 | 21 | Oxidation (M) | | |
| 12 | 367 | P0C2X9\|AL4A1_RAT | 168.52 | 28 | 28 | 2.39E+07 | 25 | 24 | 48 | Oxidation ( | 61869 | Delta-1-pyrroline-5-carboxylate dehydrogenase mitochondrial OS=Rattus norvegicus OX=10116 GN=Aldh4a1 PE=1 SV=1 |
| 19 | 309 | P63039\|CH60_RAT | 165.54 | 16 | 16 | 1.68E+07 | 14 | 14 | 25 | Oxidation ( | 60956 | 60 kDa heat shock protein mitochondrial OS=Rattus norvegicus OX=10116 GN=Hspd1 PE=1 SV=1 |
| 19 | 310 | tr\|A0A482IDN3\|A0A482I | 165.54 | 16 | 16 | 1.68E+07 | 14 | 14 | 25 | Oxidation ( | 60956 | Hsp60 OS=Rattus norvegicus OX=10116 GN=Hspd1 PE=2 SV=1 |
| 17 | 33 | Q64428\|ECHA_RAT | 163.49 | 12 | 12 | 1.63E+07 | 16 | 15 | 31 | Carbamidomethylation | | |
| 16 | 169 | Q60587\|ECHB_RAT | 155.82 | 16 | 16 | 1.27E+07 | 17 | 17 | 32 | Oxidation ( | 51414 | Trifunctional enzyme subunit beta mitochondrial OS=Rattus norvegicus OX=10116 GN=Hadhb PE=1 SV=1 |
| 16 | 170 | tr\|A0A0G2K330\|A0A0G2 | 155.82 | 16 | 16 | 1.27E+07 | 17 | 17 | 32 | Oxidation ( | 52568 | Trifunctional enzyme subunit beta mitochondrial OS=Rattus norvegicus OX=10116 GN=Hadhb PE=1 SV=1 |
| 57 | 667 | tr\|B2RZ24\|B2RZ24_RAT | 144.95 | 5 | 5 | 3.46E+05 | 4 | 4 | 4 | | 47388 | Succinate-CoA ligase subunit beta (Fragment) OS=Rattus norvegicus OX=10116 GN=Sucla2 PE=2 SV=1 |
| 57 | 668 | tr\|F1LM47\|F1LM47_RAT | 144.95 | 5 | 5 | 3.46E+05 | 4 | 4 | 4 | | 50306 | Succinate--CoA ligase [ADP-forming] subunit beta mitochondrial OS=Rattus norvegicus OX=10116 GN=Sucla2 PE=1 SV=1 |
| 22 | 245 | P18163\|ACSL1_RAT | 141.49 | 9 | 9 | 8.03E+06 | 10 | 10 | 20 | Formylatio | 78179 | Long-chain-fatty-acid--CoA ligase 1 OS=Rattus norvegicus OX=10116 GN=Acsl1 PE=1 SV=1 |
| 33 | 11911 | P21571\|ATP5J_RAT | 137.22 | 31 | 31 | 4.15E+06 | 6 | 6 | 8 | | 12494 | ATP synthase-coupling factor 6 mitochondrial OS=Rattus norvegicus OX=10116 GN=Atp5pf PE=1 SV=1 |
| 23 | 77 | P22791\|HMCS2_RAT | 132.53 | 13 | 13 | 5.84E+06 | 12 | 12 | 17 | | 56912 | Hydroxymethylglutaryl-CoA synthase mitochondrial OS=Rattus norvegicus OX=10116 GN=Hmgcs2 PE=1 SV=1 |
| 23 | 76 | tr\|Q68G44\|Q68G44_RAT | 132.53 | 13 | 13 | 5.84E+06 | 12 | 12 | 17 | | 56886 | 3-hydroxy-3-methylglutaryl coenzyme A synthase OS=Rattus norvegicus OX=10116 GN=Hmgcs2 PE=1 SV=1 |
| 24 | 100 | P17764\|THIL_RAT | 127.55 | 13 | 13 | 5.61E+06 | 8 | 8 | 13 | | 44695 | Acetyl-CoA acetyltransferase mitochondrial OS=Rattus norvegicus OX=10116 GN=Acat1 PE=1 SV=1 |
| 20 | 416 | tr\|Q5UAJ5\|Q5UAJ5_RAT | 124.72 | 42 | 42 | 5.01E+07 | 7 | 6 | 23 | Oxidation ( | 7642 | ATP synthase protein 8 OS=Rattus norvegicus OX=10116 GN=ATP8 PE=3 SV=1 |
| 20 | 418 | tr\|Q8SEZ4\|Q8SEZ4_RAT | 124.72 | 42 | 42 | 5.01E+07 | 7 | 6 | 23 | Oxidation ( | 7632 | ATP synthase protein 8 OS=Rattus norvegicus OX=10116 GN=ATPase8 PE=3 SV=1 |
| 13 | 5942 | P05504\|ATP6_RAT | 121.92 | 35 | 35 | 3.25E+07 | 18 | 17 | 45 | Oxidation (M) | | |
| 13 | 5943 | tr\|Q8HIC7\|Q8HIC7_RAT | 121.92 | 35 | 35 | 3.25E+07 | 18 | 17 | 45 | Oxidation (M) | | |
| 13 | 5944 | tr\|S5S1E9\|S5S1E9_RAT | 121.92 | 35 | 35 | 3.25E+07 | 18 | 17 | 45 | Oxidation (M) | | |
| 49 | 814 | P45953\|ACADV_RAT | 120.51 | 5 | 5 | 6.07E+05 | 4 | 4 | 4 | | 70749 | Very long-chain specific acyl-CoA dehydrogenase mitochondrial OS=Rattus norvegicus OX=10116 GN=Acadvl PE=1 SV=1 |
| 49 | 815 | tr\|Q5M9H2\|Q5M9H2_R | 120.51 | 5 | 5 | 6.07E+05 | 4 | 4 | 4 | | 70821 | Acyl-Coenzyme A dehydrogenase very long chain OS=Rattus norvegicus OX=10116 GN=Acadvl PE=1 SV=1 |
| 27 | 1292 | tr\|G3V6I4\|G3V6I4_RAT | 111.07 | 12 | 12 | 1.38E+06 | 6 | 6 | 9 | Formylatio | 37791 | Mitochondrial amidoxime reducing component 1 OS=Rattus norvegicus OX=10116 GN=Marc1 PE=1 SV=1 |
| 44 | 22 | tr\|D3ZFQ8\|D3ZFQ8_RAT | 102.41 | 6 | 6 | 1.85E+06 | 3 | 2 | 5 | | 35435 | Cytochrome c-1 OS=Rattus norvegicus OX=10116 GN=Cyc1 PE=1 SV=3 |
| 32 | 6503 | B3DMA2\|ACD11_RAT | 96.59 | 6 | 6 | 1.93E+06 | 6 | 4 | 8 | | 87371 | Acyl-CoA dehydrogenase family member 11 OS=Rattus norvegicus OX=10116 GN=Acad11 PE=1 SV=1 |
| 35 | 176 | P13086\|SUCA_RAT | 94.66 | 9 | 9 | 1.56E+06 | 5 | 5 | 6 | | 36148 | Succinate--CoA ligase [ADP/GDP-forming] subunit alpha mitochondrial OS=Rattus norvegicus OX=10116 GN=Suclg1 PE=2 SV=2 |
| 35 | 177 | tr\|A0A0H2UHE1\|A0A0H2 | 94.66 | 9 | 9 | 1.56E+06 | 5 | 5 | 6 | | 37560 | Succinate--CoA ligase [ADP/GDP-forming] subunit alpha mitochondrial OS=Rattus norvegicus OX=10116 GN=Suclg1 PE=1 SV=1 |
| 42 | 261 | P24329\|THTR_RAT | 92.35 | 6 | 6 | 1.24E+06 | 2 | 2 | 5 | | 33407 | Thiosulfate sulfurtransferase OS=Rattus norvegicus OX=10116 GN=Tst PE=1 SV=3 |
| 28 | 13 | P32551\|QCR2_RAT | 88.74 | 5 | 5 | 2.49E+06 | 6 | 6 | 9 | | 48396 | Cytochrome b-c1 complex subunit 2 mitochondrial OS=Rattus norvegicus OX=10116 GN=Uqcrc2 PE=1 SV=2 |
| 40 | 12 | Q68FY0\|QCR1_RAT | 78.39 | 6 | 6 | 2.66E+06 | 3 | 3 | 5 | | 52849 | Cytochrome b-c1 complex subunit 1 mitochondrial OS=Rattus norvegicus OX=10116 GN=Uqcrc1 PE=1 SV=1 |
| 151 | 80 | P20788\|UCRI_RAT | 77.02 | 5 | 5 | 3.70E+05 | 1 | 1 | 1 | | 29446 | Cytochrome b-c1 complex subunit Rieske mitochondrial OS=Rattus norvegicus OX=10116 GN=Uqcrfs1 PE=1 SV=2 |
| 52 | 1610 | A0A0G2K047\|ACSS3_RAT | 74.15 | 3 | 3 | 5.10E+05 | 2 | 2 | 4 | | 74675 | Acyl-CoA synthetase short-chain family member 3 mitochondrial OS=Rattus norvegicus OX=10116 GN=Acss3 PE=1 SV=1 |
| 85 | 11963 | tr\|Q5BJZ3\|Q5BJZ3_RAT | 66.58 | 2 | 2 | 2.75E+06 | 2 | 2 | 2 | | 113869 | Nicotinamide nucleotide transhydrogenase OS=Rattus norvegicus OX=10116 GN=Nnt PE=1 SV=1 |
| 50 | 5947 | Q924S5\|LONM_RAT | 63.57 | 2 | 2 | 4.06E+05 | 2 | 2 | 4 | | 105792 | Lon protease homolog mitochondrial OS=Rattus norvegicus OX=10116 GN=Lonp1 PE=2 SV=1 |
| 115 | 233 | P29147\|BDH_RAT | 63.43 | 4 | 4 | 5.82E+05 | 2 | 2 | 2 | | 38202 | D-beta-hydroxybutyrate dehydrogenase mitochondrial OS=Rattus norvegicus OX=10116 GN=Bdh1 PE=1 SV=2 |
| 115 | 234 | tr\|A0A0G2JSH2\|A0A0G2 | 63.43 | 4 | 4 | 5.82E+05 | 2 | 2 | 2 | | 38333 | 3-hydroxybutyrate dehydrogenase type 1 isoform CRA_a OS=Rattus norvegicus OX=10116 GN=Bdh1 PE=1 SV=1 |
| 29 | 11914 | P29419\|ATP5I_RAT | 62.94 | 37 | 37 | 5.32E+06 | 4 | 4 | 9 | | 8255 | ATP synthase subunit e mitochondrial OS=Rattus norvegicus OX=10116 GN=Atp5me PE=1 SV=3 |
| 191 | 1171 | Q5PQT3\|GLYAT_RAT | 58.49 | 4 | 4 | 9.74E+04 | 1 | 1 | 1 | | 33899 | Glycine N-acyltransferase OS=Rattus norvegicus OX=10116 GN=Glyat PE=2 SV=1 |
| 191 | 1172 | tr\|A4PB92\|A4PB92_RAT | 58.49 | 4 | 4 | 9.74E+04 | 1 | 1 | 1 | | 33899 | Glycine N-acyltransferase OS=Rattus norvegicus OX=10116 GN=Glyat PE=2 SV=1 |
| 191 | 1173 | tr\|A0A0G2JWI5\|A0A0G2 | 58.49 | 3 | 3 | 9.74E+04 | 1 | 1 | 1 | | 39453 | Glycine N-acyltransferase OS=Rattus norvegicus OX=10116 GN=Glyat PE=1 SV=1 |
| 191 | 1170 | tr\|B1H250\|B1H250_RAT | 58.49 | 4 | 4 | 9.74E+04 | 1 | 1 | 1 | | 33986 | Glycine-N-acyltransferase-like 1 OS=Rattus norvegicus OX=10116 GN=Glyatl1 PE=1 SV=1 |
| 67 | 1939 | tr\|D4ACE9\|D4ACE9_RAT | 58.4 | 2 | 2 | 4.18E+05 | 2 | 2 | 3 | | 103115 | Alpha-aminoadipic semialdehyde synthase mitochondrial OS=Rattus norvegicus OX=10116 GN=Aass PE=1 SV=3 |
| 67 | 1940 | A2VCW9\|AASS_RAT | 58.4 | 2 | 2 | 4.18E+05 | 2 | 2 | 3 | | 102908 | Alpha-aminoadipic semialdehyde synthase mitochondrial OS=Rattus norvegicus OX=10116 GN=Aass PE=2 SV=1 |
| 43 | 11922 | Q9JJW3\|ATPMD_RAT | 55.76 | 29 | 29 | 4.97E+06 | 3 | 3 | 5 | | 6408 | ATP synthase membrane subunit DAPIT mitochondrial OS=Rattus norvegicus OX=10116 GN=Atp5md PE=1 SV=1 |
| 26 | 399 | Q91XJ1\|BECN1_RAT | 52.24 | 3 | 3 | | 3 | 0 | 9 | | 51557 | Beclin-1 OS=Rattus norvegicus OX=10116 GN=Becn1 PE=1 SV=1 |
| 69 | 293 | Q9WVK3\|PECR_RAT | 51.98 | 12 | 12 | 7.68E+05 | 3 | 3 | 3 | Oxidation ( | 32433 | Peroxisomal trans-2-enoyl-CoA reductase OS=Rattus norvegicus OX=10116 GN=Pecr PE=2 SV=1 |
| 69 | 294 | tr\|A0A0G2JVG4\|A0A0G2 | 51.98 | 11 | 11 | 7.68E+05 | 3 | 3 | 3 | Oxidation ( | 32737 | Peroxisomal trans-2-enoyl-CoA reductase OS=Rattus norvegicus OX=10116 GN=Pecr PE=1 SV=1 |
| 112 | 218 | tr\|M0RAM5\|M0RAM5_R | 51.73 | 6 | 6 | 5.07E+05 | 2 | 2 | 2 | | 22155 | Glutathione peroxidase OS=Rattus norvegicus OX=10116 GN=Gpx1 PE=1 SV=1 |
| 112 | 219 | P04041\|GPX1_RAT | 51.73 | 6 | 6 | 5.07E+05 | 2 | 2 | 2 | | 22305 | Glutathione peroxidase 1 OS=Rattus norvegicus OX=10116 GN=Gpx1 PE=1 SV=4 |
| 30 | 11926 | D3ZAF6\|ATPK_RAT | 48.39 | 17 | 17 | 1.03E+07 | 4 | 4 | 8 | | 10452 | ATP synthase subunit f mitochondrial OS=Rattus norvegicus OX=10116 GN=Atp5mf PE=1 SV=1 |
| 65 | 427 | tr\|G3V9J8\|G3V9J8_RAT | 46.83 | 3 | 3 | 1.44E+07 | 2 | 2 | 3 | Formylatio | 87521 | Glycerol-3-phosphate acyltransferase 1 mitochondrial OS=Rattus norvegicus OX=10116 GN=Gpam PE=1 SV=2 |
| 65 | 413 | tr\|A0A0G2K2U7\|A0A0G2 | 46.83 | 3 | 3 | 1.44E+07 | 2 | 2 | 3 | Formylatio | 93728 | Glycerol-3-phosphate acyltransferase 1 mitochondrial OS=Rattus norvegicus OX=10116 GN=Gpam PE=1 SV=1 |

| | | | | | | | | | | | | | |
|---|---|---|---|---|---|---|---|---|---|---|---|---|---|
| 65 | 414 | P97564\|GPAT1_RAT | 46.83 | 3 | 3 | | 1.44E+07 | 2 | 2 | 3 | Formylatio | 93715 | Glycerol-3-phosphate acyltransferase 1 mitochondrial OS=Rattus norvegicus OX=10116 GN=Gpam PE=1 SV=3 |
| 215 | 11952 | P13803\|ETFA_RAT | 46.58 | 3 | 3 | | 2.56E+05 | 1 | 1 | 1 | | 34951 | Electron transfer flavoprotein subunit alpha mitochondrial OS=Rattus norvegicus OX=10116 GN=Etfa PE=1 SV=4 |
| 110 | 2022 | Q9Z0V6\|PRDX3_RAT | 46.07 | 5 | 5 | | 8.45E+05 | 1 | 1 | 2 | | 28295 | Thioredoxin-dependent peroxide reductase mitochondrial OS=Rattus norvegicus OX=10116 GN=Prdx3 PE=1 SV=2 |
| 110 | 2021 | tr\|G3V7I0\|G3V7I0_RAT | 46.07 | 5 | 5 | | 8.45E+05 | 1 | 1 | 2 | | 28299 | Peroxiredoxin 3 OS=Rattus norvegicus OX=10116 GN=Prdx3 PE=1 SV=1 |
| 102 | 394 | P46462\|TERA_RAT | 45.01 | 3 | 3 | | 4.61E+05 | 2 | 2 | 2 | | 89349 | Transitional endoplasmic reticulum ATPase OS=Rattus norvegicus OX=10116 GN=Vcp PE=1 SV=3 |
| 77 | 244 | P07824\|ARGI1_RAT | 43.78 | 6 | 6 | | 2.14E+05 | 2 | 1 | 3 | | 34973 | Arginase-1 OS=Rattus norvegicus OX=10116 GN=Arg1 PE=1 SV=2 |
| 71 | 41 | tr\|Q5UAJ6\|Q5UAJ6_RAT | 43.53 | 7 | 7 | | 4.15E+05 | 2 | 2 | 3 | Oxidation ( | 25942 | Cytochrome c oxidase subunit 2 OS=Rattus norvegicus OX=10116 GN=COX2 PE=3 SV=1 |
| 71 | 81 | tr\|S5RZM8\|S5RZM8_RA | 43.53 | 7 | 7 | | 4.15E+05 | 2 | 2 | 3 | Oxidation ( | 25958 | Cytochrome c oxidase subunit 2 OS=Rattus norvegicus OX=10116 GN=COX2 PE=3 SV=1 |
| 71 | 82 | P00406\|COX2_RAT | 43.53 | 7 | 7 | | 4.15E+05 | 2 | 2 | 3 | Oxidation ( | 25928 | Cytochrome c oxidase subunit 2 OS=Rattus norvegicus OX=10116 GN=Mtco2 PE=1 SV=3 |
| 71 | 83 | tr\|Q8SEZ5\|Q8SEZ5_RAT | 43.53 | 7 | 7 | | 4.15E+05 | 2 | 2 | 3 | Oxidation ( | 25928 | Cytochrome c oxidase subunit 2 OS=Rattus norvegicus OX=10116 GN=Mt-co2 PE=1 SV=1 |
| 71 | 84 | tr\|A0A097PE04\|A0A097 | 43.53 | 7 | 7 | | 4.15E+05 | 2 | 2 | 3 | Oxidation ( | 25894 | Cytochrome c oxidase subunit 2 OS=Rattus norvegicus OX=10116 GN=COX2 PE=3 SV=1 |
| 216 | 198 | P85834\|EFTU_RAT | 42.96 | 3 | 3 | | 7.43E+04 | 1 | 1 | 1 | | 49522 | Elongation factor Tu mitochondrial OS=Rattus norvegicus OX=10116 GN=Tufm PE=1 SV=1 |
| 116 | 380 | O70351\|HCD2_RAT | 42.91 | 7 | 7 | | 5.65E+05 | 2 | 2 | 2 | | 27246 | 3-hydroxyacyl-CoA dehydrogenase type-2 OS=Rattus norvegicus OX=10116 GN=Hsd17b10 PE=1 SV=3 |
| 116 | 381 | tr\|B0BMW2\|B0BMW2_R | 42.91 | 7 | 7 | | 5.65E+05 | 2 | 2 | 2 | | 27250 | 3-hydroxyacyl-CoA dehydrogenase type-2 OS=Rattus norvegicus OX=10116 GN=Hsd17b10 PE=1 SV=1 |
| 169 | 162 | tr\|Q5EBA4\|Q5EBA4_RA | 42.44 | 4 | 4 | | 5.00E+05 | 1 | 1 | 1 | Oxidation ( | 33215 | Nipsnap1 protein (Fragment) OS=Rattus norvegicus OX=10116 GN=Nipsnap1 PE=2 SV=1 |
| 169 | 163 | tr\|G3V728\|G3V728_RAT | 42.44 | 4 | 4 | | 5.00E+05 | 1 | 1 | 1 | Oxidation ( | 33346 | 4-nitrophenylphosphatase domain and non-neuronal SNAP25-like protein homolog 1 (C. elegans) isoform CRA_b OS=Rattus norvegicus OX=10116 GN=Nipsnap1 PE=1 SV=1 |
| 25 | 317 | tr\|A0A0G2JYD4\|A0A0G2 | 41.32 | 1 | 1 | | 6.14E+06 | 4 | 4 | 10 | Formylatio | 487857 | Vacuolar protein sorting 13 homolog D OS=Rattus norvegicus OX=10116 GN=Vps13d PE=1 SV=1 |
| 25 | 318 | tr\|D3ZKC6\|D3ZKC6_RAT | 41.32 | 1 | 1 | | 6.14E+06 | 4 | 4 | 10 | Formylatio | 488962 | Vacuolar protein sorting 13 homolog D OS=Rattus norvegicus OX=10116 GN=Vps13d PE=1 SV=1 |
| 66 | 1985 | tr\|Q6QI69\|Q6QI69_RAT | 41.18 | 5 | 5 | | 0 | 2 | 1 | 3 | | 46014 | LRRGT00139 OS=Rattus norvegicus OX=10116 GN=LOC500350 PE=2 SV=1 |
| 72 | 12296 | tr\|G3V734\|G3V734_RAT | 41.16 | 4 | 4 | | 0 | 2 | 2 | 3 | | 36133 | 2 4-dienoyl CoA reductase 1 mitochondrial isoform CRA_a OS=Rattus norvegicus OX=10116 GN=Decr1 PE=1 SV=1 |
| 72 | 12297 | Q64591\|DECR_RAT | 41.16 | 4 | 4 | | 0 | 2 | 2 | 3 | | 36133 | 2 4-dienoyl-CoA reductase mitochondrial OS=Rattus norvegicus OX=10116 GN=Decr1 PE=1 SV=2 |
| 111 | 87 | tr\|A0A0S1Z1V9\|A0A0S1 | 40.87 | 3 | 3 | | 3.23E+05 | 1 | 1 | 2 | | 43002 | Cytochrome b OS=Rattus norvegicus OX=10116 GN=CYTB PE=3 SV=1 |
| 111 | 225 | tr\|A0A411AID0\|A0A411 | 40.87 | 6 | 6 | | 3.23E+05 | 1 | 1 | 2 | | 23039 | Cytochrome b (Fragment) OS=Rattus norvegicus OX=10116 GN=Cytb PE=3 SV=1 |
| 111 | 224 | tr\|H8Y1J4\|H8Y1J4_RAT | 40.87 | 6 | 6 | | 3.23E+05 | 1 | 1 | 2 | | 23531 | Cytochrome b (Fragment) OS=Rattus norvegicus OX=10116 GN=cytb PE=4 SV=1 |
| 111 | 189 | tr\|A4UHZ2\|A4UHZ2_RA | 40.87 | 5 | 5 | | 3.23E+05 | 1 | 1 | 2 | | 25312 | Cytochrome b (Fragment) OS=Rattus norvegicus OX=10116 GN=cytb PE=4 SV=1 |
| 111 | 190 | tr\|B3Y999\|B3Y999_RAT | 40.87 | 5 | 5 | | 3.23E+05 | 1 | 1 | 2 | | 28378 | Cytochrome b (Fragment) OS=Rattus norvegicus OX=10116 GN=cytb PE=4 SV=1 |
| 111 | 185 | tr\|K7T8L3\|K7T8L3_RAT | 40.87 | 5 | 5 | | 3.23E+05 | 1 | 1 | 2 | | 29017 | Cytochrome b (Fragment) OS=Rattus norvegicus OX=10116 GN=cytb PE=4 SV=1 |
| 111 | 186 | tr\|A0A0U3BXX1\|A0A0U | 40.87 | 4 | 4 | | 3.23E+05 | 1 | 1 | 2 | | 33684 | Cytochrome b (Fragment) OS=Rattus norvegicus OX=10116 GN=Cytb PE=3 SV=1 |
| 111 | 187 | tr\|G8HZ47\|G8HZ47_RAT | 40.87 | 4 | 4 | | 3.23E+05 | 1 | 1 | 2 | | 35455 | Cytochrome b (Fragment) OS=Rattus norvegicus OX=10116 GN=cytb PE=3 SV=1 |
| 111 | 167 | tr\|M9TEK6\|M9TEK6_RA | 40.87 | 4 | 4 | | 3.23E+05 | 1 | 1 | 2 | | 35982 | Cytochrome b (Fragment) OS=Rattus norvegicus OX=10116 GN=cytb PE=3 SV=1 |
| 111 | 113 | tr\|A0A411P2I9\|A0A411 | 40.87 | 3 | 3 | | 3.23E+05 | 1 | 1 | 2 | | 40144 | Cytochrome b (Fragment) OS=Rattus norvegicus OX=10116 GN=Cytb PE=3 SV=1 |
| 111 | 146 | tr\|E0A0C7\|E0A0C7_RAT | 40.87 | 3 | 3 | | 3.23E+05 | 1 | 1 | 2 | | 41009 | Cytochrome b (Fragment) OS=Rattus norvegicus OX=10116 GN=cytb PE=3 SV=1 |
| 111 | 114 | tr\|A0A0U2IDR8\|A0A0U2 | 40.87 | 3 | 3 | | 3.23E+05 | 1 | 1 | 2 | | 40981 | Cytochrome b (Fragment) OS=Rattus norvegicus OX=10116 PE=3 SV=1 |
| 111 | 166 | tr\|A0A0U2UDS4\|A0A0U | 40.87 | 3 | 3 | | 3.23E+05 | 1 | 1 | 2 | | 41049 | Cytochrome b (Fragment) OS=Rattus norvegicus OX=10116 PE=3 SV=1 |
| 111 | 115 | tr\|A0A0U2M179\|A0A0U | 40.87 | 3 | 3 | | 3.23E+05 | 1 | 1 | 2 | | 41037 | Cytochrome b (Fragment) OS=Rattus norvegicus OX=10116 PE=3 SV=1 |
| 111 | 116 | tr\|A0A411P2L0\|A0A411 | 40.87 | 3 | 3 | | 3.23E+05 | 1 | 1 | 2 | | 41017 | Cytochrome b (Fragment) OS=Rattus norvegicus OX=10116 GN=Cytb PE=3 SV=1 |
| 111 | 147 | tr\|E0A016\|E0A016_RAT | 40.87 | 3 | 3 | | 3.23E+05 | 1 | 1 | 2 | | 41634 | Cytochrome b (Fragment) OS=Rattus norvegicus OX=10116 GN=cytb PE=3 SV=1 |
| 111 | 90 | tr\|F8QU31\|F8QU31_RA | 40.87 | 3 | 3 | | 3.23E+05 | 1 | 1 | 2 | | 42181 | Cytochrome b (Fragment) OS=Rattus norvegicus OX=10116 GN=cytb PE=3 SV=1 |
| 111 | 91 | tr\|A0A411P2K0\|A0A411 | 40.87 | 3 | 3 | | 3.23E+05 | 1 | 1 | 2 | | 42238 | Cytochrome b (Fragment) OS=Rattus norvegicus OX=10116 GN=Cytb PE=3 SV=1 |
| 111 | 126 | tr\|E0A075\|E0A075_RAT | 40.87 | 3 | 3 | | 3.23E+05 | 1 | 1 | 2 | | 42410 | Cytochrome b (Fragment) OS=Rattus norvegicus OX=10116 GN=cytb PE=3 SV=1 |
| 111 | 46 | tr\|L0L4L8\|L0L4L8_RAT | 40.87 | 3 | 3 | | 3.23E+05 | 1 | 1 | 2 | | 42470 | Cytochrome b (Fragment) OS=Rattus norvegicus OX=10116 GN=cytb PE=3 SV=1 |
| 111 | 47 | tr\|A0A140GE10\|A0A140 | 40.87 | 3 | 3 | | 3.23E+05 | 1 | 1 | 2 | | 42574 | Cytochrome b (Fragment) OS=Rattus norvegicus OX=10116 PE=3 SV=1 |
| 111 | 48 | tr\|A0A140GE11\|A0A140 | 40.87 | 3 | 3 | | 3.23E+05 | 1 | 1 | 2 | | 42588 | Cytochrome b (Fragment) OS=Rattus norvegicus OX=10116 PE=3 SV=1 |
| 111 | 78 | tr\|A0A140GE08\|A0A140 | 40.87 | 3 | 3 | | 3.23E+05 | 1 | 1 | 2 | | 42527 | Cytochrome b (Fragment) OS=Rattus norvegicus OX=10116 PE=3 SV=1 |
| 111 | 92 | tr\|A0A385HCC9\|A0A385 | 40.87 | 3 | 3 | | 3.23E+05 | 1 | 1 | 2 | | 42766 | Cytochrome b (Fragment) OS=Rattus norvegicus OX=10116 PE=3 SV=1 |
| 111 | 49 | tr\|A0A3Q8AGE8\|A0A3Q | 40.87 | 3 | 3 | | 3.23E+05 | 1 | 1 | 2 | | 42712 | Cytochrome b (Fragment) OS=Rattus norvegicus OX=10116 GN=Cytb PE=3 SV=1 |
| 111 | 50 | tr\|A0A3Q8AC68\|A0A3Q | 40.87 | 3 | 3 | | 3.23E+05 | 1 | 1 | 2 | | 42622 | Cytochrome b (Fragment) OS=Rattus norvegicus OX=10116 GN=Cytb PE=3 SV=1 |
| 111 | 98 | tr\|A0A3S6FK13\|A0A3S6 | 40.87 | 3 | 3 | | 3.23E+05 | 1 | 1 | 2 | | 42702 | Cytochrome b (Fragment) OS=Rattus norvegicus OX=10116 GN=Cytb PE=3 SV=1 |
| 111 | 127 | tr\|A0A385HCK1\|A0A385 | 40.87 | 3 | 3 | | 3.23E+05 | 1 | 1 | 2 | | 42824 | Cytochrome b (Fragment) OS=Rattus norvegicus OX=10116 PE=3 SV=1 |
| 111 | 102 | tr\|A0A3S6FK10\|A0A3S6 | 40.87 | 3 | 3 | | 3.23E+05 | 1 | 1 | 2 | | 42726 | Cytochrome b (Fragment) OS=Rattus norvegicus OX=10116 GN=Cytb PE=3 SV=1 |
| 111 | 51 | tr\|A0A3S6FM15\|A0A3S6 | 40.87 | 3 | 3 | | 3.23E+05 | 1 | 1 | 2 | | 42698 | Cytochrome b (Fragment) OS=Rattus norvegicus OX=10116 GN=Cytb PE=3 SV=1 |
| 111 | 188 | tr\|E0A019\|E0A019_RAT | 40.87 | 3 | 3 | | 3.23E+05 | 1 | 1 | 2 | | 43024 | Cytochrome b (Fragment) OS=Rattus norvegicus OX=10116 GN=cytb PE=3 SV=1 |
| 111 | 85 | tr\|A0A097PE40\|A0A097 | 40.87 | 3 | 3 | | 3.23E+05 | 1 | 1 | 2 | | 42865 | Cytochrome b OS=Rattus norvegicus OX=10116 GN=CYTB PE=3 SV=1 |
| 111 | 52 | tr\|A0A220D9Z6\|A0A220 | 40.87 | 3 | 3 | | 3.23E+05 | 1 | 1 | 2 | | 43018 | Cytochrome b OS=Rattus norvegicus OX=10116 PE=3 SV=1 |
| 111 | 99 | tr\|D6NSP2\|D6NSP2_RAT | 40.87 | 3 | 3 | | 3.23E+05 | 1 | 1 | 2 | | 42945 | Cytochrome b (Fragment) OS=Rattus norvegicus OX=10116 GN=cytb PE=3 SV=1 |
| 111 | 53 | tr\|D6NSS6\|D6NSS6_RAT | 40.87 | 3 | 3 | | 3.23E+05 | 1 | 1 | 2 | | 42993 | Cytochrome b (Fragment) OS=Rattus norvegicus OX=10116 GN=cytb PE=3 SV=1 |
| 111 | 103 | tr\|L0N490\|L0N490_RAT | 40.87 | 3 | 3 | | 3.23E+05 | 1 | 1 | 2 | | 43056 | Cytochrome b (Fragment) OS=Rattus norvegicus OX=10116 GN=cytb PE=3 SV=1 |
| 111 | 86 | tr\|D6NSQ3\|D6NSQ3_RA | 40.87 | 3 | 3 | | 3.23E+05 | 1 | 1 | 2 | | 43012 | Cytochrome b (Fragment) OS=Rattus norvegicus OX=10116 GN=cytb PE=3 SV=1 |
| 111 | 54 | tr\|D6NSR3\|D6NSR3_RAT | 40.87 | 3 | 3 | | 3.23E+05 | 1 | 1 | 2 | | 42986 | Cytochrome b (Fragment) OS=Rattus norvegicus OX=10116 GN=cytb PE=3 SV=1 |
| 111 | 93 | tr\|D6NSP4\|D6NSP4_RAT | 40.87 | 3 | 3 | | 3.23E+05 | 1 | 1 | 2 | | 43016 | Cytochrome b (Fragment) OS=Rattus norvegicus OX=10116 GN=cytb PE=3 SV=1 |
| 111 | 69 | tr\|D6NSQ8\|D6NSQ8_RA | 40.87 | 3 | 3 | | 3.23E+05 | 1 | 1 | 2 | | 43016 | Cytochrome b (Fragment) OS=Rattus norvegicus OX=10116 GN=cytb PE=3 SV=1 |
| 111 | 55 | tr\|D6NSR8\|D6NSR8_RAT | 40.87 | 3 | 3 | | 3.23E+05 | 1 | 1 | 2 | | 42998 | Cytochrome b (Fragment) OS=Rattus norvegicus OX=10116 GN=cytb PE=3 SV=1 |
| 111 | 120 | tr\|F8QU37\|F8QU37_RAT | 40.87 | 3 | 3 | | 3.23E+05 | 1 | 1 | 2 | | 42992 | Cytochrome b (Fragment) OS=Rattus norvegicus OX=10116 GN=cytb PE=3 SV=1 |
| 111 | 96 | tr\|F2Q6S6\|F2Q6S6_RAT | 40.87 | 3 | 3 | | 3.23E+05 | 1 | 1 | 2 | | 42938 | Cytochrome b (Fragment) OS=Rattus norvegicus OX=10116 GN=cytb PE=3 SV=1 |
| 111 | 56 | tr\|A0A0A1FZ42\|A0A0A1 | 40.87 | 3 | 3 | | 3.23E+05 | 1 | 1 | 2 | | 42989 | Cytochrome b OS=Rattus norvegicus OX=10116 GN=CYTB PE=3 SV=1 |
| 111 | 57 | tr\|D6NSS7\|D6NSS7_RAT | 40.87 | 3 | 3 | | 3.23E+05 | 1 | 1 | 2 | | 42948 | Cytochrome b (Fragment) OS=Rattus norvegicus OX=10116 GN=cytb PE=3 SV=1 |
| 111 | 58 | tr\|Q8SEY9\|Q8SEY9_RAT | 40.87 | 3 | 3 | | 3.23E+05 | 1 | 1 | 2 | | 43015 | Cytochrome b OS=Rattus norvegicus OX=10116 GN=cytb PE=3 SV=1 |
| 111 | 44 | tr\|D6NSR7\|D6NSR7_RAT | 40.87 | 3 | 3 | | 3.23E+05 | 1 | 1 | 2 | | 42982 | Cytochrome b (Fragment) OS=Rattus norvegicus OX=10116 GN=cytb PE=3 SV=1 |
| 111 | 59 | tr\|A0A220DA44\|A0A220 | 40.87 | 3 | 3 | | 3.23E+05 | 1 | 1 | 2 | | 42968 | Cytochrome b OS=Rattus norvegicus OX=10116 PE=3 SV=1 |
| 111 | 79 | tr\|F2Q6S5\|F2Q6S5_RAT | 40.87 | 3 | 3 | | 3.23E+05 | 1 | 1 | 2 | | 42952 | Cytochrome b (Fragment) OS=Rattus norvegicus OX=10116 GN=cytb PE=3 SV=1 |
| 111 | 70 | tr\|D6NSQ6\|D6NSQ6_RA | 40.87 | 3 | 3 | | 3.23E+05 | 1 | 1 | 2 | | 42952 | Cytochrome b (Fragment) OS=Rattus norvegicus OX=10116 GN=cytb PE=3 SV=1 |

| | | | | | | | | | | | | |
|---|---|---|---|---|---|---|---|---|---|---|---|---|
| 111 | 104 | tr\|L0N495\|L0N495_RAT | 40.87 | 3 | 3 | 3.23E+05 | 1 | 1 | 2 | | 42992 | Cytochrome b (Fragment) OS=Rattus norvegicus OX=10116 GN=cytb PE=3 SV=1 |
| 111 | 60 | tr\|D6NSQ0\|D6NSQ0_RA | 40.87 | 3 | 3 | 3.23E+05 | 1 | 1 | 2 | | 42968 | Cytochrome b (Fragment) OS=Rattus norvegicus OX=10116 GN=cytb PE=3 SV=1 |
| 111 | 61 | tr\|Q5UAI7\|Q5UAI7_RAT | 40.87 | 3 | 3 | 3.23E+05 | 1 | 1 | 2 | | 42998 | Cytochrome b OS=Rattus norvegicus OX=10116 GN=CYTB PE=3 SV=1 |
| 111 | 94 | tr\|D6NSR2\|D6NSR2_RAT | 40.87 | 3 | 3 | 3.23E+05 | 1 | 1 | 2 | | 43073 | Cytochrome b (Fragment) OS=Rattus norvegicus OX=10116 GN=cytb PE=3 SV=1 |
| 111 | 105 | tr\|B0M1Q8\|B0M1Q8_RA | 40.87 | 3 | 3 | 3.23E+05 | 1 | 1 | 2 | | 43026 | Cytochrome b (Fragment) OS=Rattus norvegicus OX=10116 GN=cytb PE=3 SV=1 |
| 111 | 109 | tr\|D6NSR9\|D6NSR9_RAT | 40.87 | 3 | 3 | 3.23E+05 | 1 | 1 | 2 | | 42970 | Cytochrome b (Fragment) OS=Rattus norvegicus OX=10116 GN=cytb PE=3 SV=1 |
| 111 | 95 | tr\|D6NSR1\|D6NSR1_RAT | 40.87 | 3 | 3 | 3.23E+05 | 1 | 1 | 2 | | 43002 | Cytochrome b (Fragment) OS=Rattus norvegicus OX=10116 GN=cytb PE=3 SV=1 |
| 111 | 62 | tr\|Q8HIC4\|Q8HIC4_RAT | 40.87 | 3 | 3 | 3.23E+05 | 1 | 1 | 2 | | 43012 | Cytochrome b OS=Rattus norvegicus OX=10116 GN=Mt-cyb PE=3 SV=1 |
| 111 | 63 | tr\|A0A220DA02\|A0A220 | 40.87 | 3 | 3 | 3.23E+05 | 1 | 1 | 2 | | 42968 | Cytochrome b OS=Rattus norvegicus OX=10116 PE=3 SV=1 |
| 111 | 64 | tr\|A0A0S1Z1V4\|A0A0S1 | 40.87 | 3 | 3 | 3.23E+05 | 1 | 1 | 2 | | 43016 | Cytochrome b OS=Rattus norvegicus OX=10116 GN=CYTB PE=3 SV=1 |
| 111 | 88 | tr\|R9TKN1\|R9TKN1_RAT | 40.87 | 3 | 3 | 3.23E+05 | 1 | 1 | 2 | | 43012 | Cytochrome b OS=Rattus norvegicus OX=10116 GN=CYTB PE=3 SV=1 |
| 111 | 65 | P00159\|CYB_RAT | 40.87 | 3 | 3 | 3.23E+05 | 1 | 1 | 2 | | 43012 | Cytochrome b OS=Rattus norvegicus OX=10116 GN=Mt-Cyb PE=3 SV=3 |
| 111 | 128 | tr\|D6NSP8\|D6NSP8_RAT | 40.87 | 3 | 3 | 3.23E+05 | 1 | 1 | 2 | | 43100 | Cytochrome b (Fragment) OS=Rattus norvegicus OX=10116 GN=cytb PE=3 SV=1 |
| 111 | 66 | tr\|L0N311\|L0N311_RAT | 40.87 | 3 | 3 | 3.23E+05 | 1 | 1 | 2 | | 43045 | Cytochrome b (Fragment) OS=Rattus norvegicus OX=10116 GN=cytb PE=3 SV=1 |
| 111 | 121 | tr\|A0A220DA20\|A0A220 | 40.87 | 3 | 3 | 3.23E+05 | 1 | 1 | 2 | | 43010 | Cytochrome b OS=Rattus norvegicus OX=10116 PE=3 SV=1 |
| 111 | 106 | tr\|A0A220D9W5\|A0A22 | 40.87 | 3 | 3 | 3.23E+05 | 1 | 1 | 2 | | 43028 | Cytochrome b OS=Rattus norvegicus OX=10116 GN=cyt-b PE=3 SV=1 |
| 111 | 67 | tr\|A0A220DA28\|A0A220 | 40.87 | 3 | 3 | 3.23E+05 | 1 | 1 | 2 | | 42970 | Cytochrome b OS=Rattus norvegicus OX=10116 PE=3 SV=1 |
| 111 | 130 | tr\|R9TL61\|R9TL61_RAT | 40.87 | 3 | 3 | 3.23E+05 | 1 | 1 | 2 | | 42980 | Cytochrome b OS=Rattus norvegicus OX=10116 GN=CYTB PE=3 SV=1 |
| 111 | 107 | tr\|H6RXT5\|H6RXT5_RAT | 40.87 | 3 | 3 | 3.23E+05 | 1 | 1 | 2 | | 43044 | Cytochrome b OS=Rattus norvegicus OX=10116 GN=cytb PE=3 SV=1 |
| 111 | 68 | tr\|H2KXA0\|H2KXA0_RAT | 40.87 | 3 | 3 | 3.23E+05 | 1 | 1 | 2 | | 42978 | Cytochrome b (Fragment) OS=Rattus norvegicus OX=10116 GN=cytb PE=3 SV=1 |
| 111 | 108 | tr\|L0N1V8\|L0N1V8_RAT | 40.87 | 3 | 3 | 3.23E+05 | 1 | 1 | 2 | | 43058 | Cytochrome b (Fragment) OS=Rattus norvegicus OX=10116 GN=cytb PE=3 SV=1 |
| 111 | 101 | tr\|A0A096XNM4\|A0A09 | 40.87 | 3 | 3 | 3.23E+05 | 1 | 1 | 2 | | 43042 | Cytochrome b (Fragment) OS=Rattus norvegicus OX=10116 PE=3 SV=1 |
| 111 | 129 | tr\|D6NSP7\|D6NSP7_RAT | 40.87 | 3 | 3 | 3.23E+05 | 1 | 1 | 2 | | 43046 | Cytochrome b (Fragment) OS=Rattus norvegicus OX=10116 GN=cytb PE=3 SV=1 |
| 111 | 122 | tr\|A0A220D9Z3\|A0A220 | 40.87 | 3 | 3 | 3.23E+05 | 1 | 1 | 2 | | 43024 | Cytochrome b OS=Rattus norvegicus OX=10116 PE=3 SV=1 |
| 111 | 89 | tr\|D6NSP5\|D6NSP5_RAT | 40.87 | 3 | 3 | 3.23E+05 | 1 | 1 | 2 | | 43012 | Cytochrome b (Fragment) OS=Rattus norvegicus OX=10116 GN=cytb PE=3 SV=1 |
| 111 | 97 | tr\|D6NSP6\|D6NSP6_RAT | 40.87 | 3 | 3 | 3.23E+05 | 1 | 1 | 2 | | 43050 | Cytochrome b (Fragment) OS=Rattus norvegicus OX=10116 GN=cytb PE=3 SV=1 |
| 39 | 428 | tr\|D4A1D3\|D4A1D3_RAT | 40.86 | 1 | 1 | 2.47E+06 | 3 | 2 | 5 | Carbamidomethylation | | |
| 37 | 12306 | Q8CGU6\|NICA_RAT | 39.55 | 3 | 3 | 1.88E+06 | 3 | 2 | 6 | | 78400 | Nicastrin OS=Rattus norvegicus OX=10116 GN=Ncstn PE=1 SV=1 |
| 34 | 314 | tr\|D3ZHK4\|D3ZHK4_RAT | 38.21 | 1 | 1 | | 3 | 0 | 7 | | 182226 | RB1-inducible coiled-coil 1 OS=Rattus norvegicus OX=10116 GN=Rb1cc1 PE=1 SV=1 |
| 107 | 623 | tr\|G3V6I5\|G3V6I5_RAT | 38.05 | 5 | 5 | 8.52E+05 | 2 | 2 | 2 | | 46777 | DnaJ heat shock protein family (Hsp40) member A3 OS=Rattus norvegicus OX=10116 GN=Dnaja3 PE=1 SV=2 |
| 107 | 624 | tr\|Q2TVU3\|Q2TVU3_RA | 38.05 | 5 | 5 | 8.52E+05 | 2 | 2 | 2 | | 49412 | TID1 OS=Rattus norvegicus OX=10116 GN=Dnaja3 PE=2 SV=1 |
| 107 | 625 | tr\|A0A0G2K4Y1\|A0A0G2 | 38.05 | 5 | 5 | 8.52E+05 | 2 | 2 | 2 | | 49416 | DnaJ heat shock protein family (Hsp40) member A3 OS=Rattus norvegicus OX=10116 GN=Dnaja3 PE=1 SV=1 |
| 107 | 626 | tr\|Q2UZS7\|Q2UZS7_RAT | 38.05 | 5 | 5 | 8.52E+05 | 2 | 2 | 2 | | 52399 | Tid-1 long isoform OS=Rattus norvegicus OX=10116 GN=Dnaja3 PE=2 SV=1 |
| 107 | 627 | tr\|A0A0G2K5E4\|A0A0G2 | 38.05 | 5 | 5 | 8.52E+05 | 2 | 2 | 2 | | 52403 | DnaJ heat shock protein family (Hsp40) member A3 OS=Rattus norvegicus OX=10116 GN=Dnaja3 PE=1 SV=1 |
| 45 | 6541 | tr\|D4A175\|D4A175_RAT | 36.95 | 2 | 2 | | 2 | 0 | 5 | | 53667 | tRNA dimethylallyltransferase OS=Rattus norvegicus OX=10116 GN=Trit1 PE=3 SV=1 |
| 164 | 11913 | tr\|F1LX07\|F1LX07_RAT | 35.84 | 1 | 1 | 3.93E+05 | 1 | 1 | 1 | | 71917 | Solute carrier family 25 member 12 OS=Rattus norvegicus OX=10116 GN=Slc25a12 PE=1 SV=3 |
| 164 | 6630 | tr\|F1LZW6\|F1LZW6_RAT | 35.84 | 2 | 2 | 3.93E+05 | 1 | 1 | 1 | | 54099 | Solute carrier family 25 member 13 OS=Rattus norvegicus OX=10116 GN=Slc25a13 PE=1 SV=2 |
| 164 | 11912 | tr\|A0A0G2K2J7\|A0A0G2 | 35.84 | 2 | 2 | 3.93E+05 | 1 | 1 | 1 | | 41504 | Solute carrier family 25 member 12 OS=Rattus norvegicus OX=10116 GN=Slc25a12 PE=1 SV=1 |
| 70 | 589 | tr\|F1LM33\|F1LM33_RAT | 35.33 | 1 | 1 | 0 | 2 | 1 | 3 | | 156679 | Leucine-rich PPR motif-containing protein mitochondrial OS=Rattus norvegicus OX=10116 GN=Lrpprc PE=1 SV=2 |
| 70 | 590 | Q5SGE0\|LPPRC_RAT | 35.33 | 1 | 1 | 0 | 2 | 1 | 3 | | 156652 | Leucine-rich PPR motif-containing protein mitochondrial OS=Rattus norvegicus OX=10116 GN=Lrpprc PE=1 SV=1 |
| 31 | 888 | O70600\|RSAD2_RAT | 35.28 | 6 | 6 | 1.01E+07 | 2 | 2 | 8 | Formylatio | 41255 | Radical S-adenosyl methionine domain-containing protein 2 OS=Rattus norvegicus OX=10116 GN=Rsad2 PE=1 SV=1 |
| 31 | 889 | tr\|A0A0H2UHF4\|A0A0H | 35.28 | 6 | 6 | 1.01E+07 | 2 | 2 | 8 | Formylatio | 41444 | RCG62278 OS=Rattus norvegicus OX=10116 GN=Rsad2 PE=4 SV=1 |
| 190 | 279 | Q5M9I5\|QCR6_RAT | 35.26 | 17 | 17 | 3.96E+05 | 1 | 1 | 1 | | 10424 | Cytochrome b-c1 complex subunit 6 mitochondrial OS=Rattus norvegicus OX=10116 GN=Uqcrh PE=3 SV=1 |
| 38 | 333 | tr\|A0A0G2K1Q1\|A0A0G | 34.66 | 1 | 1 | | 2 | 0 | 6 | | 107334 | NLR family member X1 OS=Rattus norvegicus OX=10116 GN=Nlrx1 PE=1 SV=1 |
| 38 | 334 | Q5FVQ8\|NLRX1_RAT | 34.66 | 1 | 1 | | 2 | 0 | 6 | | 107590 | NLR family member X1 OS=Rattus norvegicus OX=10116 GN=Nlrx1 PE=2 SV=1 |
| 192 | 6579 | tr\|A0A0G2JYE8\|A0A0G2 | 34.09 | 6 | 6 | 1.34E+05 | 1 | 1 | 1 | | 20804 | Serine--pyruvate aminotransferase mitochondrial OS=Rattus norvegicus OX=10116 GN=Agxt PE=1 SV=1 |
| 192 | 348 | P09139\|SPYA_RAT | 34.09 | 3 | 3 | 1.34E+05 | 1 | 1 | 1 | | 45834 | Serine--pyruvate aminotransferase mitochondrial OS=Rattus norvegicus OX=10116 GN=Agxt PE=1 SV=1 |
| 61 | 5986 | tr\|D3Z899\|D3Z899_RAT | 32.73 | 3 | 3 | 7.43E+05 | 2 | 1 | 4 | Formylatio | 61393 | Mitoguardin 2 OS=Rattus norvegicus OX=10116 GN=Miga2 PE=1 SV=1 |
| 61 | 5993 | tr\|A0A0G2KAE6\|A0A0G2 | 32.73 | 2 | 2 | 7.43E+05 | 2 | 1 | 4 | Formylatio | 65660 | Mitoguardin 2 OS=Rattus norvegicus OX=10116 GN=Miga2 PE=1 SV=1 |
| 189 | 1273 | tr\|A0A0G2K3W1\|A0A0G | 31.46 | 3 | 3 | 1.76E+05 | 1 | 1 | 1 | | 53760 | von Willebrand factor A domain-containing 8 OS=Rattus norvegicus OX=10116 GN=Vwa8 PE=1 SV=1 |
| 95 | 6227 | tr\|A0A0H2UH92\|A0A0H | 30.79 | 1 | 1 | 5.43E+06 | 2 | 2 | 2 | Formylatio | 136526 | Graves disease carrier protein OS=Rattus norvegicus OX=10116 GN=Slc25a16 PE=1 SV=1 |
| 95 | 6075 | D3ZG52\|DNA2_RAT | 30.79 | 1 | 1 | 5.43E+06 | 2 | 2 | 2 | Formylatio | 119588 | DNA replication ATP-dependent helicase/nuclease DNA2 OS=Rattus norvegicus OX=10116 GN=Dna2 PE=3 SV=1 |
| 119 | 1436 | P30277\|CCNB1_RAT | 30.04 | 4 | 4 | 2.71E+05 | 2 | 1 | 2 | | 47391 | G2/mitotic-specific cyclin-B1 OS=Rattus norvegicus OX=10116 GN=Ccnb1 PE=2 SV=1 |
| 81 | 6369 | tr\|A0A0G2JVK4\|A0A0G2 | 29.4 | 1 | 1 | | 2 | 0 | 3 | | 77802 | Neurolysin mitochondrial OS=Rattus norvegicus OX=10116 GN=Nln PE=1 SV=1 |
| 81 | 6128 | tr\|A0A0G2JSY3\|A0A0G2 | 29.4 | 1 | 1 | | 2 | 0 | 3 | | 80282 | Neurolysin (Metallopeptidase M3 family) isoform CRA_a OS=Rattus norvegicus OX=10116 GN=Nln PE=1 SV=1 |
| 81 | 6129 | P42676\|NEUL_RAT | 29.4 | 1 | 1 | | 2 | 0 | 3 | | 80254 | Neurolysin mitochondrial OS=Rattus norvegicus OX=10116 GN=Nln PE=1 SV=1 |
| 68 | 595 | D4A929\|WDR81_RAT | 28.88 | 1 | 1 | 0 | 2 | 1 | 3 | | 212258 | WD repeat-containing protein 81 OS=Rattus norvegicus OX=10116 GN=Wdr81 PE=3 SV=1 |
| 47 | 762 | B0BN56\|RT31_RAT | 28.2 | 3 | 3 | | 2 | 0 | 5 | | 43962 | 28S ribosomal protein S31 mitochondrial OS=Rattus norvegicus OX=10116 GN=Mrps31 PE=2 SV=1 |
| 97 | 6664 | tr\|D3ZUM2\|D3ZUM2_RA | 27.67 | 2 | 2 | 1.17E+05 | 2 | 1 | 2 | Formylatio | 79688 | Sterile alpha and TIR motif containing 1 (Predicted) OS=Rattus norvegicus OX=10116 GN=Sarm1 PE=1 SV=1 |
| 59 | 6015 | tr\|D4A5P3\|D4A5P3_RAT | 27.11 | 2 | 2 | | 2 | 0 | 4 | | 59813 | Mitoguardin 1 OS=Rattus norvegicus OX=10116 GN=Miga1 PE=4 SV=1 |
| 59 | 5978 | tr\|A0A0G2K2T2\|A0A0G2 | 27.11 | 2 | 2 | | 2 | 0 | 4 | | 67482 | Mitoguardin 1 OS=Rattus norvegicus OX=10116 GN=Miga1 PE=4 SV=1 |
| 108 | 228 | P16970\|ABCD3_RAT | 26.63 | 3 | 3 | 3.01E+05 | 2 | 2 | 2 | | 75316 | ATP-binding cassette sub-family D member 3 OS=Rattus norvegicus OX=10116 GN=Abcd3 PE=1 SV=3 |
| 79 | 957 | Q5M821\|PPM1H_RAT | 25.62 | 2 | 2 | 5.67E+05 | 2 | 1 | 3 | | 56380 | Protein phosphatase 1H OS=Rattus norvegicus OX=10116 GN=Ppm1h PE=1 SV=2 |
| 218 | 23250 | tr\|Q9ESE3\|Q9ESE3_RAT | 25.32 | 5 | 5 | 2.29E+05 | 1 | 1 | 1 | | 18186 | 5-aminolevulinate synthase (Fragment) OS=Rattus norvegicus OX=10116 GN=ALS2 PE=2 SV=1 |
| 218 | 677 | Q63147\|HEM0_RAT | 25.32 | 2 | 2 | 2.29E+05 | 1 | 1 | 1 | | 64842 | 5-aminolevulinate synthase erythroid-specific mitochondrial OS=Rattus norvegicus OX=10116 GN=Alas2 PE=1 SV=1 |
| 219 | 1261 | tr\|Q5RKL4\|Q5RKL4_RAT | 23.94 | 1 | 1 | 0 | 1 | 1 | 1 | | 95977 | Dimethylglycine dehydrogenase OS=Rattus norvegicus OX=10116 GN=Dmgdh PE=1 SV=1 |
| 219 | 1262 | Q63342\|M2GD_RAT | 23.94 | 1 | 1 | 0 | 1 | 1 | 1 | | 96047 | Dimethylglycine dehydrogenase mitochondrial OS=Rattus norvegicus OX=10116 GN=Dmgdh PE=1 SV=1 |
| 219 | 1268 | tr\|A0A0G2K9Y2\|A0A0G2 | 23.94 | 1 | 1 | 0 | 1 | 1 | 1 | | 98864 | Dimethylglycine dehydrogenase mitochondrial OS=Rattus norvegicus OX=10116 GN=Dmgdh PE=1 SV=1 |
| 152 | 545 | P11497\|ACACA_RAT | 23.78 | 0 | 0 | 5.86E+05 | 1 | 1 | 1 | | 265191 | Acetyl-CoA carboxylase 1 OS=Rattus norvegicus OX=10116 GN=Acaca PE=1 SV=1 |
| 63 | 11978 | tr\|A0A0A0MXU8\|A0A0A | 23.7 | 4 | 4 | 1.80E+07 | 1 | 1 | 3 | | 18236 | Caveolin OS=Rattus norvegicus OX=10116 GN=Cav2 PE=1 SV=1 |
| 63 | 11979 | Q2IBC5\|CAV2_RAT | 23.7 | 4 | 4 | 1.80E+07 | 1 | 1 | 3 | | 18266 | Caveolin-2 OS=Rattus norvegicus OX=10116 GN=Cav2 PE=1 SV=2 |

<table>
<tr><td>195</td><td>6337</td><td>P17178|CP27A_RAT</td><td>23.69</td><td>2</td><td>2</td><td>2.45E+05</td><td>1</td><td>1</td><td>1</td><td></td><td>60733</td><td>Sterol 26-hydroxylase  mitochondrial OS=Rattus norvegicus OX=10116 GN=Cyp27a1 PE=1 SV=1</td></tr>
<tr><td>195</td><td>6338</td><td>tr|A0A0H2UHN7|A0A0H</td><td>23.69</td><td>2</td><td>2</td><td>2.45E+05</td><td>1</td><td>1</td><td>1</td><td></td><td>62492</td><td>RCG24013  isoform CRA_a OS=Rattus norvegicus OX=10116 GN=Cyp27a1 PE=1 SV=1</td></tr>
<tr><td>220</td><td>17931</td><td>O88767|PARK7_RAT</td><td>22.97</td><td>5</td><td>5</td><td>0</td><td>1</td><td>1</td><td>1</td><td></td><td>19974</td><td>Protein/nucleic acid deglycase DJ-1 OS=Rattus norvegicus OX=10116 GN=Park7 PE=1 SV=1</td></tr>
<tr><td>220</td><td>23225</td><td>tr|Q5BKC3|Q5BKC3_RAT</td><td>22.97</td><td>5</td><td>5</td><td>0</td><td>1</td><td>1</td><td>1</td><td></td><td>22503</td><td>Park7 protein OS=Rattus norvegicus OX=10116 GN=Park7 PE=1 SV=1</td></tr>
<tr><td>118</td><td>33514</td><td>Q68FX0|IDH3B_RAT</td><td>22.7</td><td>2</td><td>2</td><td>7.44E+05</td><td>1</td><td>1</td><td>2</td><td></td><td>42354</td><td>Isocitrate dehydrogenase [NAD] subunit beta  mitochondrial OS=Rattus norvegicus OX=10116 GN=Idh3B PE=2 SV=1</td></tr>
<tr><td>193</td><td>1503</td><td>tr|F1LSY7|F1LSY7_RAT</td><td>22.27</td><td>1</td><td>1</td><td>4.46E+05</td><td>1</td><td>1</td><td>1</td><td></td><td>109031</td><td>Endoplasmic reticulum to nucleus-signaling 1 OS=Rattus norvegicus OX=10116 GN=Ern1 PE=4 SV=2</td></tr>
<tr><td>193</td><td>1504</td><td>tr|A0A0G2K2H4|A0A0G2</td><td>22.27</td><td>1</td><td>1</td><td>4.46E+05</td><td>1</td><td>1</td><td>1</td><td></td><td>110150</td><td>Endoplasmic reticulum to nucleus-signaling 1 OS=Rattus norvegicus OX=10116 GN=Ern1 PE=2 SV=1</td></tr>
<tr><td>91</td><td>537</td><td>tr|D3ZKV7|D3ZKV7_RAT</td><td>21.73</td><td>0</td><td>0</td><td>3.05E+05</td><td>1</td><td>1</td><td>2</td><td>Formylatio</td><td>309807</td><td>Trinucleotide repeat-containing 18 OS=Rattus norvegicus OX=10116 GN=Tnrc18 PE=1 SV=3</td></tr>
<tr><td>223</td><td>17980</td><td>Q5M9G9|FAKD4_RAT</td><td>21.39</td><td>1</td><td>1</td><td>1.56E+06</td><td>1</td><td>1</td><td>1</td><td></td><td>71181</td><td>FAST kinase domain-containing protein 4 OS=Rattus norvegicus OX=10116 GN=Tbrg4 PE=2 SV=1</td></tr>
<tr><td>106</td><td>28163</td><td>Q9R1Z0|VDAC3_RAT</td><td>21.38</td><td>5</td><td>5</td><td>4.46E+06</td><td>1</td><td>1</td><td>2</td><td>Formylatio</td><td>30798</td><td>Voltage-dependent anion-selective channel protein 3 OS=Rattus norvegicus OX=10116 GN=Vdac3 PE=1 SV=2</td></tr>
<tr><td>147</td><td>1776</td><td>Q9Z286|ADCYA_RAT</td><td>21.37</td><td>0</td><td>0</td><td></td><td>1</td><td>0</td><td>1</td><td></td><td>185852</td><td>Adenylate cyclase type 10 OS=Rattus norvegicus OX=10116 GN=Adcy10 PE=1 SV=1</td></tr>
<tr><td>201</td><td>6675</td><td>Q5RKH7|S35F6_RAT</td><td>21.12</td><td>3</td><td>3</td><td>1.05E+06</td><td>1</td><td>1</td><td>1</td><td>Carbamido</td><td>41092</td><td>Solute carrier family 35 member F6 OS=Rattus norvegicus OX=10116 GN=Slc35f6 PE=2 SV=1</td></tr>
<tr><td>94</td><td>1723</td><td>O88658|KIF1B_RAT</td><td>20.58</td><td>0</td><td>0</td><td>2.00E+06</td><td>1</td><td>1</td><td>2</td><td></td><td>204169</td><td>Kinesin-like protein KIF1B OS=Rattus norvegicus OX=10116 GN=Kif1b PE=1 SV=2</td></tr>
<tr><td>196</td><td>465</td><td>P07633|PCCB_RAT</td><td>20.22</td><td>2</td><td>2</td><td>1.05E+05</td><td>1</td><td>1</td><td>1</td><td>Oxidation (</td><td>58626</td><td>Propionyl-CoA carboxylase beta chain  mitochondrial OS=Rattus norvegicus OX=10116 GN=Pccb PE=2 SV=1</td></tr>
<tr><td>196</td><td>466</td><td>tr|Q68FZ8|Q68FZ8_RAT</td><td>20.22</td><td>2</td><td>2</td><td>1.05E+05</td><td>1</td><td>1</td><td>1</td><td>Oxidation (</td><td>58678</td><td>Propionyl coenzyme A carboxylase  beta polypeptide OS=Rattus norvegicus OX=10116 GN=Pccb PE=1 SV=1</td></tr>
<tr><td>41</td><td>733</td><td>tr|F1MA54|F1MA54_RA</td><td>20.1</td><td>2</td><td>2</td><td>4.58E+07</td><td>1</td><td>1</td><td>5</td><td>Formylatio</td><td>49099</td><td>Protein-serine/threonine kinase OS=Rattus norvegicus OX=10116 GN=Pdk1 PE=1 SV=2</td></tr>
<tr><td>41</td><td>734</td><td>Q63065|PDK1_RAT</td><td>20.1</td><td>2</td><td>2</td><td>4.58E+07</td><td>1</td><td>1</td><td>5</td><td>Formylatio</td><td>49081</td><td>[Pyruvate dehydrogenase (acetyl-transferring)] kinase isozyme 1  mitochondrial OS=Rattus norvegicus OX=10116 GN=Pdk1 PE=1 SV=1</td></tr>
<tr><td>41</td><td>735</td><td>tr|Q5FVT5|Q5FVT5_RAT</td><td>20.1</td><td>2</td><td>2</td><td>4.58E+07</td><td>1</td><td>1</td><td>5</td><td>Formylatio</td><td>49095</td><td>Protein-serine/threonine kinase OS=Rattus norvegicus OX=10116 GN=Pdk1 PE=2 SV=1</td></tr>
<tr><td>80</td><td>377</td><td>A0A0G2JZ79|SIR1_RAT</td><td>20.05</td><td>1</td><td>1</td><td>5.41E+06</td><td>1</td><td>1</td><td>3</td><td></td><td>62059</td><td>NAD-dependent protein deacetylase sirtuin-1 OS=Rattus norvegicus OX=10116 GN=Sirt1 PE=3 SV=2</td></tr>
<tr><td>80</td><td>378</td><td>tr|A0A182DWI7|A0A182</td><td>20.05</td><td>1</td><td>1</td><td>5.41E+06</td><td>1</td><td>1</td><td>3</td><td></td><td>81778</td><td>NAD-dependent protein deacetylase sirtuin-1 OS=Rattus norvegicus OX=10116 GN=Sirt1 PE=4 SV=1</td></tr>
</table>
