## Supplementary material for "Permeability transition pore-related changes in the proteome and channel activity of ATP synthase dimers and monomers": RLM SM-BM PTP Dimer of V compl.

| Protein Gr | Protein ID | Accession | -10lgP | Coverage ( | Coverage ( | Area Samp | #Peptides | #Unique | #Spec Sam | PTM | Avg. Mass | Description |
|---|---|---|---|---|---|---|---|---|---|---|---|---|
| 1 | 1 | P10860\|DH | 384.07 | 68 | 68 | 4.92E+08 | 228 | 225 | 442 | Carbamidomethylation | | |
| 3 | 8 | P10719\|AT | 359.3 | 65 | 65 | 4.21E+08 | 177 | 174 | 322 | Oxidation ( | 56354 | ATP synthase subunit beta mitochondrial OS=Rattus norvegicus OX=10116 GN=Atp5f1b PE=1 SV=2 |
| 3 | 9 | tr\|G3V6D3 | 359.3 | 65 | 65 | 4.21E+08 | 177 | 174 | 322 | Oxidation ( | 56345 | ATP synthase subunit beta OS=Rattus norvegicus OX=10116 GN=Atp5f1b PE=1 SV=1 |
| 2 | 10 | P15999\|AT | 326.78 | 60 | 60 | 4.56E+08 | 159 | 157 | 432 | Carbamidomethylation | | |
| 2 | 11 | tr\|F1LP05\| | 326.78 | 60 | 60 | 4.56E+08 | 159 | 157 | 432 | Carbamidomethylation | | |
| 8 | 119 | P31399\|AT | 264.38 | 61 | 61 | 3.42E+07 | 36 | 35 | 59 | Carbamidomethylation | | |
| 4 | 4 | P07756\|CP | 257.16 | 30 | 30 | 4.89E+07 | 79 | 79 | 115 | Oxidation ( | 164579 | Carbamoyl-phosphate synthase [ammonia] mitochondrial OS=Rattus norvegicus OX=10116 GN=Cps1 PE=1 SV=1 |
| 5 | 15 | P67779\|PH | 248.83 | 57 | 57 | 5.23E+07 | 55 | 55 | 83 | | 29820 | Prohibitin OS=Rattus norvegicus OX=10116 GN=Phb PE=1 SV=1 |
| 6 | 64 | P35435\|AT | 245.02 | 37 | 37 | 4.80E+07 | 36 | 36 | 82 | Oxidation ( | 30191 | ATP synthase subunit gamma mitochondrial OS=Rattus norvegicus OX=10116 GN=Atp5f1c PE=1 SV=2 |
| 6 | 63 | tr\|Q6QI09 | 245.02 | 16 | 16 | 4.80E+07 | 36 | 36 | 82 | Oxidation ( | 67721 | ATP synthase subunit gamma mitochondrial OS=Rattus norvegicus OX=10116 GN=Taf3 PE=1 SV=1 |
| 7 | 12 | Q5XIH7\|PH | 225.53 | 48 | 48 | 3.37E+07 | 46 | 45 | 71 | Oxidation ( | 33312 | Prohibitin-2 OS=Rattus norvegicus OX=10116 GN=Phb2 PE=1 SV=1 |
| 9 | 30 | P52873\|PY | 222.84 | 24 | 24 | 1.68E+07 | 41 | 41 | 55 | Oxidation ( | 129777 | Pyruvate carboxylase mitochondrial OS=Rattus norvegicus OX=10116 GN=Pc PE=1 SV=2 |
| 9 | 31 | tr\|A0A0G2 | 222.84 | 22 | 22 | 1.68E+07 | 41 | 41 | 55 | Oxidation ( | 140005 | Pyruvate carboxylase mitochondrial OS=Rattus norvegicus OX=10116 GN=Pc PE=1 SV=1 |
| 11 | 20 | P19234\|ND | 218.11 | 39 | 39 | 2.35E+07 | 29 | 29 | 49 | Formylatio | 27378 | NADH dehydrogenase [ubiquinone] flavoprotein 2 mitochondrial OS=Rattus norvegicus OX=10116 GN=Ndufv2 PE=1 SV=2 |
| 13 | 3 | P85834\|EF | 214.69 | 27 | 27 | 1.20E+07 | 30 | 30 | 40 | Oxidation ( | 49522 | Elongation factor Tu mitochondrial OS=Rattus norvegicus OX=10116 GN=Tufm PE=1 SV=1 |
| 12 | 13 | Q66HF1\|N | 210.51 | 20 | 20 | 1.40E+07 | 31 | 31 | 42 | | 79412 | NADH-ubiquinone oxidoreductase 75 kDa subunit mitochondrial OS=Rattus norvegicus OX=10116 GN=Ndufs1 PE=1 SV=1 |
| 10 | 246 | P19511\|AT | 199.97 | 30 | 30 | 4.79E+07 | 27 | 27 | 54 | | 28869 | ATP synthase F(0) complex subunit B1 mitochondrial OS=Rattus norvegicus OX=10116 GN=Atp5pb PE=1 SV=1 |
| 15 | 26 | P00507\|AA | 196.49 | 27 | 27 | 1.48E+07 | 23 | 23 | 33 | Oxidation ( | 47314 | Aspartate aminotransferase mitochondrial OS=Rattus norvegicus OX=10116 GN=Got2 PE=1 SV=2 |
| 16 | 22 | tr\|Q68G44 | 193.93 | 29 | 29 | 6.08E+06 | 25 | 25 | 31 | | 56886 | 3-hydroxy-3-methylglutaryl coenzyme A synthase OS=Rattus norvegicus OX=10116 GN=Hmgcs2 PE=1 SV=1 |
| 32 | 49 | tr\|B2RYS8 | 187.03 | 21 | 21 | 4.63E+06 | 11 | 11 | 15 | Oxidation ( | 21959 | NADH dehydrogenase [ubiquinone] 1 beta subcomplex subunit 8 mitochondrial OS=Rattus norvegicus OX=10116 GN=Ndufb8 PE=1 SV=1 |
| 17 | 19 | Q641Y2\|N | 183.77 | 28 | 28 | 1.10E+07 | 23 | 23 | 31 | Oxidation ( | 52562 | NADH dehydrogenase [ubiquinone] iron-sulfur protein 2 mitochondrial OS=Rattus norvegicus OX=10116 GN=Ndufs2 PE=1 SV=1 |
| 18 | 7 | Q5BK63\|N | 182.64 | 32 | 32 | 5.96E+06 | 22 | 21 | 29 | | 42559 | NADH dehydrogenase [ubiquinone] 1 alpha subcomplex subunit 9 mitochondrial OS=Rattus norvegicus OX=10116 GN=Ndufa9 PE=1 SV=2 |
| 26 | 171 | tr\|D3ZFJ6\| | 177.72 | 22 | 22 | 4.51E+06 | 15 | 15 | 17 | Carbamido | 60420 | Lactamase beta OS=Rattus norvegicus OX=10116 GN=Lactb PE=1 SV=1 |
| 45 | 136 | P24329\|TH | 168.08 | 13 | 13 | 1.75E+06 | 8 | 8 | 9 | | 33407 | Thiosulfate sulfurtransferase OS=Rattus norvegicus OX=10116 GN=Tst PE=1 SV=3 |
| 21 | 350 | tr\|G3V7Y3 | 167.78 | 30 | 30 | 1.57E+07 | 14 | 14 | 22 | Oxidation ( | 17563 | ATP synthase subunit delta mitochondrial OS=Rattus norvegicus OX=10116 GN=Atp5f1d PE=1 SV=1 |
| 31 | 32 | P07895\|SO | 164.03 | 28 | 28 | 1.51E+06 | 10 | 10 | 15 | | 24674 | Superoxide dismutase [Mn] mitochondrial OS=Rattus norvegicus OX=10116 GN=Sod2 PE=1 SV=2 |
| 22 | 48 | tr\|D3ZF13 | 163.06 | 29 | 29 | 7.94E+06 | 13 | 13 | 21 | Oxidation ( | 17514 | Acyl carrier protein OS=Rattus norvegicus OX=10116 GN=Ndufab1 PE=1 SV=1 |
| 14 | 5802 | Q06647\|AT | 162.52 | 45 | 45 | 1.47E+07 | 21 | 20 | 36 | Carbamidomethylation | | |
| 34 | 204 | tr\|Q5PQZ9 | 160.2 | 30 | 30 | 6.18E+06 | 10 | 10 | 14 | | 14359 | NADH dehydrogenase [ubiquinone] 1 subunit C2 OS=Rattus norvegicus OX=10116 GN=Ndufc2 PE=1 SV=1 |
| 20 | 28 | tr\|D3ZG43 | 157.32 | 33 | 33 | 7.06E+06 | 18 | 18 | 24 | | 30226 | NADH dehydrogenase (Ubiquinone) Fe-S protein 3 (Predicted) isoform CRA_c OS=Rattus norvegicus OX=10116 GN=Ndufs3 PE=1 SV=1 |
| 25 | 16 | Q68FY0\|Q | 154.18 | 16 | 16 | 3.90E+06 | 15 | 15 | 18 | Carbamido | 52849 | Cytochrome b-c1 complex subunit 1 mitochondrial OS=Rattus norvegicus OX=10116 GN=Uqcrc1 PE=1 SV=1 |
| 41 | 58 | tr\|D4A565 | 147.34 | 21 | 21 | 3.33E+06 | 7 | 7 | 9 | | 21664 | NADH dehydrogenase (Ubiquinone) 1 beta subcomplex 5 (Predicted) isoform CRA_b OS=Rattus norvegicus OX=10116 GN=Ndufb5 PE=1 SV=1 |
| 29 | 27 | Q60587\|EC | 145.31 | 21 | 21 | 3.51E+06 | 15 | 15 | 16 | | 51414 | Trifunctional enzyme subunit beta mitochondrial OS=Rattus norvegicus OX=10116 GN=Hadhb PE=1 SV=1 |
| 23 | 24 | Q64428\|EC | 144.98 | 12 | 12 | 6.44E+06 | 15 | 15 | 20 | Formylatio | 82665 | Trifunctional enzyme subunit alpha mitochondrial OS=Rattus norvegicus OX=10116 GN=Hadha PE=1 SV=2 |
| 30 | 50 | Q02253\|M | 140.27 | 17 | 17 | 1.87E+06 | 15 | 15 | 16 | Carbamidomethylation | | |
| 30 | 51 | tr\|G3V7J0 | 140.27 | 17 | 17 | 1.87E+06 | 15 | 15 | 16 | Carbamidomethylation | | |
| 24 | 35 | P17764\|TH | 139.39 | 22 | 22 | 5.64E+06 | 13 | 13 | 19 | Formylatio | 44695 | Acetyl-CoA acetyltransferase mitochondrial OS=Rattus norvegicus OX=10116 GN=Acat1 PE=1 SV=1 |
| 39 | 137 | tr\|D4A7L4 | 138.67 | 26 | 26 | 6.93E+06 | 7 | 7 | 10 | | 17634 | NADH dehydrogenase (Ubiquinone) 1 beta subcomplex 11 (Predicted) OS=Rattus norvegicus OX=10116 GN=Ndufb11 PE=1 SV=1 |
| 19 | 1646 | Q6PDU7\|A | 137.93 | 39 | 39 | 1.24E+07 | 13 | 13 | 25 | Oxidation ( | 11433 | ATP synthase subunit g mitochondrial OS=Rattus norvegicus OX=10116 GN=Atp5mg PE=1 SV=2 |
| 48 | 38 | tr\|D3ZFQ8 | 132.42 | 9 | 9 | 4.20E+06 | 4 | 4 | 8 | | 35435 | Cytochrome c-1 OS=Rattus norvegicus OX=10116 GN=Cyc1 PE=1 SV=3 |
| 51 | 1044 | tr\|D4ACE9 | 130.76 | 9 | 9 | 2.75E+06 | 7 | 7 | 7 | Formylatio | 103115 | Alpha-aminoadipic semialdehyde synthase mitochondrial OS=Rattus norvegicus OX=10116 GN=Aass PE=1 SV=3 |
| 51 | 742 | A2VCW9\|A | 130.76 | 9 | 9 | 2.75E+06 | 7 | 7 | 7 | Formylatio | 102908 | Alpha-aminoadipic semialdehyde synthase mitochondrial OS=Rattus norvegicus OX=10116 GN=Aass PE=2 SV=1 |
| 27 | 46 | P13086\|SU | 129.53 | 13 | 13 | 2.65E+06 | 9 | 9 | 17 | | 36148 | Succinate--CoA ligase [ADP/GDP-forming] subunit alpha mitochondrial OS=Rattus norvegicus OX=10116 GN=Suclg1 PE=2 SV=2 |
| 27 | 47 | tr\|A0A0H2 | 129.53 | 13 | 13 | 2.65E+06 | 9 | 9 | 17 | | 37560 | Succinate--CoA ligase [ADP/GDP-forming] subunit alpha mitochondrial OS=Rattus norvegicus OX=10116 GN=Suclg1 PE=1 SV=1 |
| 35 | 66 | tr\|D4A0T0 | 129.03 | 33 | 33 | 4.56E+06 | 8 | 8 | 11 | Oxidation ( | 20859 | NADH:ubiquinone oxidoreductase subunit B10 OS=Rattus norvegicus OX=10116 GN=Ndufb10 PE=1 SV=1 |
| 36 | 45 | P04762\|CA | 127.85 | 9 | 9 | 1.34E+06 | 9 | 9 | 11 | | 59757 | Catalase OS=Rattus norvegicus OX=10116 GN=Cat PE=1 SV=3 |
| 54 | 44 | P11240\|CO | 126.62 | 31 | 31 | 2.49E+06 | 5 | 5 | 7 | | 16130 | Cytochrome c oxidase subunit 5A mitochondrial OS=Rattus norvegicus OX=10116 GN=Cox5a PE=1 SV=1 |
| 50 | 55 | tr\|B0BNE6 | 125.8 | 16 | 16 | 1.49E+06 | 7 | 6 | 8 | | 23970 | NADH dehydrogenase (Ubiquinone) Fe-S protein 8 (Predicted) isoform CRA_a OS=Rattus norvegicus OX=10116 GN=Ndufs8 PE=1 SV=1 |
| 49 | 181 | Q80W89\|N | 125.66 | 18 | 18 | 4.91E+06 | 6 | 6 | 8 | | 14854 | NADH dehydrogenase [ubiquinone] 1 alpha subcomplex subunit 11 OS=Rattus norvegicus OX=10116 GN=Ndufa11 PE=2 SV=1 |
| 57 | 142 | tr\|D3ZCZ9 | 118.49 | 21 | 21 | 1.72E+06 | 5 | 5 | 7 | | 13040 | NADH dehydrogenase [ubiquinone] iron-sulfur protein 6 mitochondrial OS=Rattus norvegicus OX=10116 GN=LOC100912599 PE=1 SV=1 |
| 28 | 43 | tr\|Q5XIH3 | 115.48 | 11 | 11 | 8.39E+06 | 8 | 8 | 16 | Formylatio | 50731 | NADH dehydrogenase [ubiquinone] flavoprotein 1 mitochondrial OS=Rattus norvegicus OX=10116 GN=Ndufv1 PE=1 SV=1 |
| 38 | 138 | Q5XIF3\|ND | 114.98 | 22 | 22 | 2.52E+06 | 8 | 8 | 10 | | 19741 | NADH dehydrogenase [ubiquinone] iron-sulfur protein 4 mitochondrial OS=Rattus norvegicus OX=10116 GN=Ndufs4 PE=1 SV=1 |
| 44 | 57 | Q561S0\|N | 113.27 | 15 | 15 | 3.53E+06 | 7 | 7 | 9 | Oxidation ( | 40493 | NADH dehydrogenase [ubiquinone] 1 alpha subcomplex subunit 10 mitochondrial OS=Rattus norvegicus OX=10116 GN=Ndufa10 PE=1 SV=1 |
| 53 | 41 | tr\|D4A3V2 | 111.96 | 29 | 29 | 1.52E+06 | 7 | 7 | 7 | Oxidation ( | 15224 | NADH dehydrogenase [ubiquinone] 1 alpha subcomplex subunit 6 OS=Rattus norvegicus OX=10116 GN=Ndufa6 PE=1 SV=1 |
| 37 | 34 | tr\|A0A0G2 | 110.04 | 25 | 25 | 1.55E+06 | 6 | 6 | 11 | | 19965 | NADH dehydrogenase [ubiquinone] 1 alpha subcomplex subunit 8 OS=Rattus norvegicus OX=10116 GN=Ndufa8 PE=1 SV=1 |
| 33 | 21 | P32551\|QC | 108.59 | 11 | 11 | 5.07E+06 | 10 | 10 | 14 | | 48396 | Cytochrome b-c1 complex subunit 2 mitochondrial OS=Rattus norvegicus OX=10116 GN=Uqcrc2 PE=1 SV=2 |
| 62 | 249 | tr\|F1LXA0\| | 101.36 | 21 | 21 | 1.57E+06 | 4 | 4 | 5 | | 17178 | NADH dehydrogenase [ubiquinone] 1 alpha subcomplex subunit 12 OS=Rattus norvegicus OX=10116 GN=Ndufa12 PE=1 SV=2 |
| 42 | 182 | tr\|Q06QG6 | 101.34 | 8 | 8 | 1.35E+06 | 7 | 7 | 9 | | 68588 | NADH-ubiquinone oxidoreductase chain 5 OS=Rattus norvegicus OX=10116 GN=ND5 PE=3 SV=1 |
| 67 | 42 | P20788\|UC | 97.39 | 8 | 8 | 7.71E+05 | 3 | 3 | 4 | | 29446 | Cytochrome b-c1 complex subunit Rieske mitochondrial OS=Rattus norvegicus OX=10116 GN=Uqcrfs1 PE=1 SV=2 |
| 59 | 194 | tr\|Q5RJN0 | 95.2 | 15 | 15 | 8.14E+05 | 5 | 5 | 5 | Oxidation ( | 23945 | NADH dehydrogenase (Ubiquinone) Fe-S protein 7 OS=Rattus norvegicus OX=10116 GN=Ndufs7 PE=1 SV=1 |
| 52 | 111 | tr\|S5RZM8 | 91.71 | 29 | 29 | 2.11E+06 | 7 | 7 | 7 | Oxidation (M) | | |
| 52 | 112 | P00406\|CO | 91.71 | 29 | 29 | 2.11E+06 | 7 | 7 | 7 | Oxidation (M) | | |

| | | | | | | | | | | | |
|---|---|---|---|---|---|---|---|---|---|---|---|
| 52 | 104 | tr\|Q5UAJ6 | 91.71 | 29 | 29 | 2.11E+06 | 7 | 7 | 7 | Oxidation (M) | | |
| 52 | 113 | tr\|Q8SEZ5 | 91.71 | 29 | 29 | 2.11E+06 | 7 | 7 | 7 | Oxidation (M) | | |
| 52 | 114 | tr\|A0A097 | 91.71 | 29 | 29 | 2.11E+06 | 7 | 7 | 7 | Oxidation (M) | | |
| 55 | 210 | tr\|Q5M949 | 90.29 | 13 | 13 | 1.62E+06 | 5 | 5 | 7 | | 28340 | Nipsnap homolog 3A (C. elegans) OS=Rattus norvegicus OX=10116 GN=Nipsnap3b PE=1 SV=1 |
| 46 | 1095 | tr\|Q5UAJ5 | 89.84 | 30 | 30 | 7.24E+06 | 5 | 4 | 9 | Oxidation ( | 7642 | ATP synthase protein 8 OS=Rattus norvegicus OX=10116 GN=ATP8 PE=3 SV=1 |
| 46 | 1097 | tr\|Q8SEZ4 | 89.84 | 30 | 30 | 7.24E+06 | 5 | 4 | 9 | Oxidation ( | 7632 | ATP synthase protein 8 OS=Rattus norvegicus OX=10116 GN=ATPase8 PE=3 SV=1 |
| 47 | 11740 | P29419\|AT | 85.39 | 34 | 34 | 3.35E+06 | 4 | 4 | 9 | | 8255 | ATP synthase subunit e mitochondrial OS=Rattus norvegicus OX=10116 GN=Atp5me PE=1 SV=3 |
| 60 | 135 | tr\|D3ZZ21 | 83.7 | 27 | 27 | 1.74E+06 | 4 | 4 | 5 | | 15638 | NADH dehydrogenase (Ubiquinone) 1 beta subcomplex 6 (Predicted) OS=Rattus norvegicus OX=10116 GN=Ndufb6 PE=1 SV=1 |
| 72 | 61 | P29147\|BD | 83.4 | 8 | 8 | 3.35E+05 | 4 | 3 | 4 | | 38202 | D-beta-hydroxybutyrate dehydrogenase mitochondrial OS=Rattus norvegicus OX=10116 GN=Bdh1 PE=1 SV=2 |
| 72 | 62 | tr\|A0A0G2 | 83.4 | 8 | 8 | 3.35E+05 | 4 | 3 | 4 | | 38333 | 3-hydroxybutyrate dehydrogenase type 1 isoform CRA_a OS=Rattus norvegicus OX=10116 GN=Bdh1 PE=1 SV=1 |
| 40 | 147 | P05508\|NU | 82.09 | 9 | 9 | 6.83E+05 | 6 | 6 | 9 | | 51783 | NADH-ubiquinone oxidoreductase chain 4 OS=Rattus norvegicus OX=10116 GN=Mtnd4 PE=3 SV=3 |
| 40 | 148 | tr\|D2E6K0 | 82.09 | 9 | 9 | 6.83E+05 | 6 | 6 | 9 | | 51833 | NADH-ubiquinone oxidoreductase chain 4 OS=Rattus norvegicus OX=10116 GN=ND4 PE=3 SV=1 |
| 40 | 149 | tr\|A7XYB9 | 82.09 | 9 | 9 | 6.83E+05 | 6 | 6 | 9 | | 51773 | NADH-ubiquinone oxidoreductase chain 4 OS=Rattus norvegicus OX=10116 GN=Nd4 PE=3 SV=1 |
| 40 | 150 | tr\|Q06QE1 | 82.09 | 9 | 9 | 6.83E+05 | 6 | 6 | 9 | | 51851 | NADH-ubiquinone oxidoreductase chain 4 OS=Rattus norvegicus OX=10116 GN=ND4 PE=3 SV=1 |
| 40 | 151 | tr\|Q06QA2 | 82.09 | 9 | 9 | 6.83E+05 | 6 | 6 | 9 | | 51787 | NADH-ubiquinone oxidoreductase chain 4 OS=Rattus norvegicus OX=10116 GN=ND4 PE=3 SV=1 |
| 40 | 152 | tr\|Q7HKW | 82.09 | 9 | 9 | 6.83E+05 | 6 | 6 | 9 | | 51801 | NADH-ubiquinone oxidoreductase chain 4 (Fragment) OS=Rattus norvegicus OX=10116 GN=NADH4 PE=3 SV=1 |
| 40 | 153 | tr\|Q06QG7 | 82.09 | 9 | 9 | 6.83E+05 | 6 | 6 | 9 | | 51791 | NADH-ubiquinone oxidoreductase chain 4 OS=Rattus norvegicus OX=10116 GN=ND4 PE=3 SV=1 |
| 40 | 154 | tr\|Q06Q89 | 82.09 | 9 | 9 | 6.83E+05 | 6 | 6 | 9 | | 51819 | NADH-ubiquinone oxidoreductase chain 4 OS=Rattus norvegicus OX=10116 GN=ND4 PE=3 SV=1 |
| 40 | 155 | tr\|Q35737 | 82.09 | 9 | 9 | 6.83E+05 | 6 | 6 | 9 | | 51765 | NADH-ubiquinone oxidoreductase chain 4 OS=Rattus norvegicus OX=10116 PE=3 SV=1 |
| 40 | 156 | tr\|Q8HIC6 | 82.09 | 9 | 9 | 6.83E+05 | 6 | 6 | 9 | | 51783 | NADH-ubiquinone oxidoreductase chain 4 OS=Rattus norvegicus OX=10116 GN=Mt-nd4 PE=3 SV=1 |
| 73 | 120 | tr\|A0A411 | 81.26 | 10 | 10 | 8.50E+05 | 4 | 4 | 4 | | 40144 | Cytochrome b (Fragment) OS=Rattus norvegicus OX=10116 GN=Cytb PE=3 SV=1 |
| 73 | 132 | tr\|A0A0U2 | 81.26 | 9 | 9 | 8.50E+05 | 4 | 4 | 4 | | 40981 | Cytochrome b (Fragment) OS=Rattus norvegicus OX=10116 PE=3 SV=1 |
| 73 | 121 | tr\|A0A0U2 | 81.26 | 9 | 9 | 8.50E+05 | 4 | 4 | 4 | | 41037 | Cytochrome b (Fragment) OS=Rattus norvegicus OX=10116 PE=3 SV=1 |
| 73 | 122 | tr\|A0A411 | 81.26 | 9 | 9 | 8.50E+05 | 4 | 4 | 4 | | 41017 | Cytochrome b (Fragment) OS=Rattus norvegicus OX=10116 GN=Cytb PE=3 SV=1 |
| 73 | 123 | tr\|F8QU31 | 81.26 | 9 | 9 | 8.50E+05 | 4 | 4 | 4 | | 42181 | Cytochrome b (Fragment) OS=Rattus norvegicus OX=10116 GN=cytb PE=3 SV=1 |
| 73 | 124 | tr\|A0A411 | 81.26 | 9 | 9 | 8.50E+05 | 4 | 4 | 4 | | 42238 | Cytochrome b (Fragment) OS=Rattus norvegicus OX=10116 GN=Cytb PE=3 SV=1 |
| 73 | 67 | tr\|L0L4L8\| | 81.26 | 9 | 9 | 8.50E+05 | 4 | 4 | 4 | | 42470 | Cytochrome b (Fragment) OS=Rattus norvegicus OX=10116 GN=cytb PE=3 SV=1 |
| 73 | 68 | tr\|A0A140 | 81.26 | 9 | 9 | 8.50E+05 | 4 | 4 | 4 | | 42574 | Cytochrome b (Fragment) OS=Rattus norvegicus OX=10116 PE=3 SV=1 |
| 73 | 69 | tr\|A0A140 | 81.26 | 9 | 9 | 8.50E+05 | 4 | 4 | 4 | | 42588 | Cytochrome b (Fragment) OS=Rattus norvegicus OX=10116 PE=3 SV=1 |
| 73 | 70 | tr\|A0A140 | 81.26 | 9 | 9 | 8.50E+05 | 4 | 4 | 4 | | 42527 | Cytochrome b (Fragment) OS=Rattus norvegicus OX=10116 PE=3 SV=1 |
| 73 | 125 | tr\|A0A385 | 81.26 | 9 | 9 | 8.50E+05 | 4 | 4 | 4 | | 42766 | Cytochrome b (Fragment) OS=Rattus norvegicus OX=10116 PE=3 SV=1 |
| 73 | 71 | tr\|A0A3Q8 | 81.26 | 9 | 9 | 8.50E+05 | 4 | 4 | 4 | | 42712 | Cytochrome b (Fragment) OS=Rattus norvegicus OX=10116 GN=Cytb PE=3 SV=1 |
| 73 | 72 | tr\|A0A3Q8 | 81.26 | 9 | 9 | 8.50E+05 | 4 | 4 | 4 | | 42622 | Cytochrome b (Fragment) OS=Rattus norvegicus OX=10116 GN=Cytb PE=3 SV=1 |
| 73 | 73 | tr\|A0A3S6 | 81.26 | 9 | 9 | 8.50E+05 | 4 | 4 | 4 | | 42702 | Cytochrome b (Fragment) OS=Rattus norvegicus OX=10116 GN=Cytb PE=3 SV=1 |
| 73 | 126 | tr\|A0A385 | 81.26 | 9 | 9 | 8.50E+05 | 4 | 4 | 4 | | 42751 | Cytochrome b (Fragment) OS=Rattus norvegicus OX=10116 PE=3 SV=1 |
| 73 | 74 | tr\|A0A3S6 | 81.26 | 9 | 9 | 8.50E+05 | 4 | 4 | 4 | | 42698 | Cytochrome b (Fragment) OS=Rattus norvegicus OX=10116 GN=Cytb PE=3 SV=1 |
| 73 | 76 | tr\|A0A220 | 81.26 | 9 | 9 | 8.50E+05 | 4 | 4 | 4 | | 43018 | Cytochrome b OS=Rattus norvegicus OX=10116 PE=3 SV=1 |
| 73 | 77 | tr\|D6NSP2 | 81.26 | 9 | 9 | 8.50E+05 | 4 | 4 | 4 | | 42945 | Cytochrome b (Fragment) OS=Rattus norvegicus OX=10116 GN=cytb PE=3 SV=1 |
| 73 | 78 | tr\|D6NSS6 | 81.26 | 9 | 9 | 8.50E+05 | 4 | 4 | 4 | | 42993 | Cytochrome b (Fragment) OS=Rattus norvegicus OX=10116 GN=cytb PE=3 SV=1 |
| 73 | 79 | tr\|D6NSQ3 | 81.26 | 9 | 9 | 8.50E+05 | 4 | 4 | 4 | | 43012 | Cytochrome b (Fragment) OS=Rattus norvegicus OX=10116 GN=cytb PE=3 SV=1 |
| 73 | 80 | tr\|D6NSR3 | 81.26 | 9 | 9 | 8.50E+05 | 4 | 4 | 4 | | 42986 | Cytochrome b (Fragment) OS=Rattus norvegicus OX=10116 GN=cytb PE=3 SV=1 |
| 73 | 127 | tr\|D6NSP4 | 81.26 | 9 | 9 | 8.50E+05 | 4 | 4 | 4 | | 43016 | Cytochrome b (Fragment) OS=Rattus norvegicus OX=10116 GN=cytb PE=3 SV=1 |
| 73 | 81 | tr\|D6NSQ8 | 81.26 | 9 | 9 | 8.50E+05 | 4 | 4 | 4 | | 43016 | Cytochrome b (Fragment) OS=Rattus norvegicus OX=10116 GN=cytb PE=3 SV=1 |
| 73 | 82 | tr\|D6NSR8 | 81.26 | 9 | 9 | 8.50E+05 | 4 | 4 | 4 | | 42998 | Cytochrome b (Fragment) OS=Rattus norvegicus OX=10116 GN=cytb PE=3 SV=1 |
| 73 | 103 | tr\|F8QU37 | 81.26 | 9 | 9 | 8.50E+05 | 4 | 4 | 4 | | 42992 | Cytochrome b (Fragment) OS=Rattus norvegicus OX=10116 GN=cytb PE=3 SV=1 |
| 73 | 83 | tr\|F2Q6S6 | 81.26 | 9 | 9 | 8.50E+05 | 4 | 4 | 4 | | 42938 | Cytochrome b (Fragment) OS=Rattus norvegicus OX=10116 GN=cytb PE=3 SV=1 |
| 73 | 84 | tr\|A0A0A1 | 81.26 | 9 | 9 | 8.50E+05 | 4 | 4 | 4 | | 42989 | Cytochrome b OS=Rattus norvegicus OX=10116 GN=CYTB PE=3 SV=1 |
| 73 | 85 | tr\|D6NSS7 | 81.26 | 9 | 9 | 8.50E+05 | 4 | 4 | 4 | | 42948 | Cytochrome b (Fragment) OS=Rattus norvegicus OX=10116 GN=cytb PE=3 SV=1 |
| 73 | 86 | tr\|Q8SEY9 | 81.26 | 9 | 9 | 8.50E+05 | 4 | 4 | 4 | | 43015 | Cytochrome b OS=Rattus norvegicus OX=10116 GN=cytb PE=3 SV=1 |
| 73 | 87 | tr\|D6NSR7 | 81.26 | 9 | 9 | 8.50E+05 | 4 | 4 | 4 | | 42982 | Cytochrome b (Fragment) OS=Rattus norvegicus OX=10116 GN=cytb PE=3 SV=1 |
| 73 | 88 | tr\|A0A220 | 81.26 | 9 | 9 | 8.50E+05 | 4 | 4 | 4 | | 42968 | Cytochrome b OS=Rattus norvegicus OX=10116 PE=3 SV=1 |
| 73 | 89 | tr\|F2Q6S5 | 81.26 | 9 | 9 | 8.50E+05 | 4 | 4 | 4 | | 42952 | Cytochrome b (Fragment) OS=Rattus norvegicus OX=10116 GN=cytb PE=3 SV=1 |
| 73 | 90 | tr\|D6NSQ6 | 81.26 | 9 | 9 | 8.50E+05 | 4 | 4 | 4 | | 42952 | Cytochrome b (Fragment) OS=Rattus norvegicus OX=10116 GN=cytb PE=3 SV=1 |
| 73 | 116 | tr\|D6NSQ0 | 81.26 | 9 | 9 | 8.50E+05 | 4 | 4 | 4 | | 42968 | Cytochrome b (Fragment) OS=Rattus norvegicus OX=10116 GN=cytb PE=3 SV=1 |
| 73 | 91 | tr\|Q5UAI7 | 81.26 | 9 | 9 | 8.50E+05 | 4 | 4 | 4 | | 42998 | Cytochrome b OS=Rattus norvegicus OX=10116 GN=CYTB PE=3 SV=1 |
| 73 | 128 | tr\|D6NSR2 | 81.26 | 9 | 9 | 8.50E+05 | 4 | 4 | 4 | | 43073 | Cytochrome b (Fragment) OS=Rattus norvegicus OX=10116 GN=cytb PE=3 SV=1 |
| 73 | 92 | tr\|A0A0S1 | 81.26 | 9 | 9 | 8.50E+05 | 4 | 4 | 4 | | 43002 | Cytochrome b OS=Rattus norvegicus OX=10116 GN=CYTB PE=3 SV=1 |
| 73 | 129 | tr\|D6NSR1 | 81.26 | 9 | 9 | 8.50E+05 | 4 | 4 | 4 | | 43002 | Cytochrome b (Fragment) OS=Rattus norvegicus OX=10116 GN=cytb PE=3 SV=1 |
| 73 | 93 | tr\|Q8HIC4 | 81.26 | 9 | 9 | 8.50E+05 | 4 | 4 | 4 | | 43012 | Cytochrome b OS=Rattus norvegicus OX=10116 GN=Mt-cyb PE=3 SV=1 |
| 73 | 94 | tr\|A0A220 | 81.26 | 9 | 9 | 8.50E+05 | 4 | 4 | 4 | | 42968 | Cytochrome b OS=Rattus norvegicus OX=10116 PE=3 SV=1 |
| 73 | 95 | tr\|A0A0S1 | 81.26 | 9 | 9 | 8.50E+05 | 4 | 4 | 4 | | 43016 | Cytochrome b OS=Rattus norvegicus OX=10116 GN=CYTB PE=3 SV=1 |
| 73 | 96 | tr\|R9TKN1 | 81.26 | 9 | 9 | 8.50E+05 | 4 | 4 | 4 | | 43012 | Cytochrome b OS=Rattus norvegicus OX=10116 GN=CYTB PE=3 SV=1 |
| 73 | 97 | P00159\|CY | 81.26 | 9 | 9 | 8.50E+05 | 4 | 4 | 4 | | 43012 | Cytochrome b OS=Rattus norvegicus OX=10116 GN=Mt-Cyb PE=3 SV=3 |

| | | | | | | | | | | | | |
|---|---|---|---|---|---|---|---|---|---|---|---|---|
| 73 | 98 | tr\|L0N311 | 81.26 | 9 | 9 | 8.50E+05 | 4 | 4 | 4 | | 43045 | Cytochrome b (Fragment) OS=Rattus norvegicus OX=10116 GN=cytb PE=3 SV=1 |
| 73 | 99 | tr\|A0A220 | 81.26 | 9 | 9 | 8.50E+05 | 4 | 4 | 4 | | 42970 | Cytochrome b OS=Rattus norvegicus OX=10116 PE=3 SV=1 |
| 73 | 100 | tr\|H2KXA0 | 81.26 | 9 | 9 | 8.50E+05 | 4 | 4 | 4 | | 42978 | Cytochrome b (Fragment) OS=Rattus norvegicus OX=10116 GN=cytb PE=3 SV=1 |
| 73 | 101 | tr\|A0A096 | 81.26 | 9 | 9 | 8.50E+05 | 4 | 4 | 4 | | 43042 | Cytochrome b (Fragment) OS=Rattus norvegicus OX=10116 PE=3 SV=1 |
| 73 | 102 | tr\|D6NSP5 | 81.26 | 9 | 9 | 8.50E+05 | 4 | 4 | 4 | | 43012 | Cytochrome b (Fragment) OS=Rattus norvegicus OX=10116 GN=cytb PE=3 SV=1 |
| 126 | 145 | tr\|F1LPG5 | 80.25 | 11 | 11 | 8.74E+05 | 2 | 2 | 2 | | 15064 | NADH:ubiquinone oxidoreductase subunit B4 OS=Rattus norvegicus OX=10116 GN=Ndufb4 PE=1 SV=1 |
| 106 | 258 | tr\|M0RB63 | 76.64 | 31 | 31 | 0 | 2 | 2 | 2 | Oxidation ( | 9372 | RCG63041 OS=Rattus norvegicus OX=10116 GN=LOC684509 PE=4 SV=1 |
| 106 | 257 | tr\|A9UMW | 76.64 | 31 | 31 | 0 | 2 | 2 | 2 | Oxidation ( | 9255 | Ndufa3 protein (Fragment) OS=Rattus norvegicus OX=10116 GN=Ndufa3 PE=2 SV=1 |
| 106 | 259 | tr\|A0A0G2 | 76.64 | 29 | 29 | 0 | 2 | 2 | 2 | Oxidation ( | 10170 | NADH:ubiquinone oxidoreductase subunit A3 OS=Rattus norvegicus OX=10116 GN=Ndufa3 PE=1 SV=1 |
| 98 | 202 | tr\|Q5M9H | 74.08 | 3 | 3 | 3.56E+05 | 2 | 2 | 2 | | 70821 | Acyl-Coenzyme A dehydrogenase very long chain OS=Rattus norvegicus OX=10116 GN=Acadvl PE=1 SV=1 |
| 98 | 201 | P45953\|AC | 74.08 | 3 | 3 | 3.56E+05 | 2 | 2 | 2 | | 70749 | Very long-chain specific acyl-CoA dehydrogenase mitochondrial OS=Rattus norvegicus OX=10116 GN=Acadvl PE=1 SV=1 |
| 100 | 109 | tr\|A0A140 | 72.48 | 4 | 4 | 3.83E+05 | 2 | 2 | 2 | | 67049 | MICOS complex subunit MIC60 OS=Rattus norvegicus OX=10116 GN=Immt PE=1 SV=1 |
| 100 | 110 | Q3KR86\|M | 72.48 | 4 | 4 | 3.83E+05 | 2 | 2 | 2 | | 67177 | MICOS complex subunit Mic60 (Fragment) OS=Rattus norvegicus OX=10116 GN=Immt PE=1 SV=1 |
| 100 | 115 | tr\|A0A0G2 | 72.48 | 3 | 3 | 3.83E+05 | 2 | 2 | 2 | | 86230 | MICOS complex subunit MIC60 OS=Rattus norvegicus OX=10116 GN=Immt PE=1 SV=1 |
| 79 | 139 | Q6AXV4\|SA | 72.26 | 6 | 6 | 1.30E+05 | 3 | 3 | 3 | | 51960 | Sorting and assembly machinery component 50 homolog OS=Rattus norvegicus OX=10116 GN=Samm50 PE=1 SV=1 |
| 89 | 140 | tr\|F1LN88 | 71.9 | 6 | 6 | 2.54E+05 | 3 | 3 | 3 | | 56517 | Aldehyde dehydrogenase mitochondrial OS=Rattus norvegicus OX=10116 GN=Aldh2 PE=1 SV=2 |
| 89 | 141 | P11884\|AL | 71.9 | 6 | 6 | 2.54E+05 | 3 | 3 | 3 | | 56488 | Aldehyde dehydrogenase mitochondrial OS=Rattus norvegicus OX=10116 GN=Aldh2 PE=1 SV=1 |
| 85 | 346 | tr\|D3ZS58 | 71.84 | 23 | 23 | 6.43E+05 | 3 | 3 | 3 | | 10845 | NADH dehydrogenase [ubiquinone] 1 alpha subcomplex subunit 2 OS=Rattus norvegicus OX=10116 GN=Ndufa2 PE=1 SV=1 |
| 61 | 192 | tr\|A0A0G2 | 71.21 | 5 | 5 | 2.56E+06 | 4 | 4 | 5 | Formylatio | 68759 | Serum albumin OS=Rattus norvegicus OX=10116 GN=Alb PE=1 SV=1 |
| 63 | 245 | tr\|A9UMV | 71.05 | 19 | 19 | 2.36E+06 | 2 | 2 | 5 | | 12500 | NADH:ubiquinone oxidoreductase subunit A7 OS=Rattus norvegicus OX=10116 GN=Ndufa7 PE=1 SV=1 |
| 88 | 65 | tr\|D3ZE15 | 69.36 | 19 | 19 | 6.45E+05 | 3 | 3 | 3 | Oxidation ( | 16777 | NADH:ubiquinone oxidoreductase subunit A13 OS=Rattus norvegicus OX=10116 GN=Ndufa13 PE=1 SV=1 |
| 71 | 297 | Q924S5\|LC | 68.69 | 3 | 3 | 3.43E+05 | 3 | 3 | 4 | | 105792 | Lon protease homolog mitochondrial OS=Rattus norvegicus OX=10116 GN=Lonp1 PE=2 SV=1 |
| 261 | 277 | tr\|B2RZ24 | 67.94 | 4 | 4 | 0 | 1 | 1 | 1 | | 47388 | Succinate-CoA ligase subunit beta (Fragment) OS=Rattus norvegicus OX=10116 GN=Sucla2 PE=2 SV=1 |
| 261 | 278 | tr\|F1LM47 | 67.94 | 4 | 4 | 0 | 1 | 1 | 1 | | 50306 | Succinate--CoA ligase [ADP-forming] subunit beta mitochondrial OS=Rattus norvegicus OX=10116 GN=Sucla2 PE=1 SV=1 |
| 56 | 54 | P18163\|AC | 67.92 | 5 | 5 | 8.09E+05 | 5 | 3 | 7 | | 78179 | Long-chain-fatty-acid--CoA ligase 1 OS=Rattus norvegicus OX=10116 GN=Acsl1 PE=1 SV=1 |
| 65 | 190 | P07824\|AR | 67.48 | 12 | 12 | 7.37E+05 | 3 | 3 | 4 | | 34973 | Arginase-1 OS=Rattus norvegicus OX=10116 GN=Arg1 PE=1 SV=2 |
| 87 | 143 | tr\|Q5EBA4 | 66.89 | 11 | 11 | 4.99E+05 | 3 | 3 | 3 | Oxidation ( | 33215 | Nipsnap1 protein (Fragment) OS=Rattus norvegicus OX=10116 GN=Nipsnap1 PE=2 SV=1 |
| 87 | 144 | tr\|G3V728 | 66.89 | 11 | 11 | 4.99E+05 | 3 | 3 | 3 | Oxidation ( | 33346 | 4-nitrophenylphosphatase domain and non-neuronal SNAP25-like protein homolog 1 (C. elegans) isoform CRA_b OS=Rattus norvegicus OX=10116 GN=Nipsnap1 PE=1 SV=1 |
| 43 | 689 | tr\|Q549I0\| | 64.28 | 18 | 18 | 7.12E+05 | 5 | 5 | 9 | Oxidation (M) | | |
| 43 | 690 | P05504\|AT | 64.28 | 18 | 18 | 7.12E+05 | 5 | 5 | 9 | Oxidation (M) | | |
| 43 | 691 | tr\|Q8HIC7 | 64.28 | 18 | 18 | 7.12E+05 | 5 | 5 | 9 | Oxidation (M) | | |
| 43 | 692 | tr\|S5S1E9\| | 64.28 | 18 | 18 | 7.12E+05 | 5 | 5 | 9 | Oxidation (M) | | |
| 43 | 693 | tr\|Q06QE5 | 64.28 | 18 | 18 | 7.12E+05 | 5 | 5 | 9 | Oxidation (M) | | |
| 43 | 459 | tr\|Q8SEZ3 | 64.28 | 18 | 18 | 7.12E+05 | 5 | 5 | 9 | Oxidation (M) | | |
| 80 | 207 | tr\|G3V6I4\| | 59.34 | 7 | 7 | 2.30E+05 | 2 | 2 | 3 | | 37791 | Mitochondrial amidoxime reducing component 1 OS=Rattus norvegicus OX=10116 GN=Marc1 PE=1 SV=1 |
| 125 | 222 | tr\|Q4G067 | 58.68 | 4 | 4 | 3.51E+05 | 2 | 2 | 2 | | 37440 | Mitochondrial ribosomal protein L44 OS=Rattus norvegicus OX=10116 GN=Mrpl44 PE=1 SV=1 |
| 69 | 366 | P33124\|AC | 57.69 | 3 | 3 | 1.05E+05 | 3 | 1 | 4 | Formylatio | 78180 | Long-chain-fatty-acid--CoA ligase 6 OS=Rattus norvegicus OX=10116 GN=Acsl6 PE=1 SV=1 |
| 58 | 446 | Q91XJ1\|BE | 56.12 | 3 | 3 | | 3 | 0 | 6 | | 51557 | Beclin-1 OS=Rattus norvegicus OX=10116 GN=Becn1 PE=1 SV=1 |
| 187 | 248 | tr\|D3ZUX5 | 55.73 | 5 | 5 | 1.87E+05 | 1 | 1 | 1 | | 26435 | MICOS complex subunit OS=Rattus norvegicus OX=10116 GN=Chchd3 PE=1 SV=1 |
| 91 | 5817 | P21571\|AT | 54.59 | 19 | 19 | 5.43E+05 | 2 | 2 | 3 | | 12494 | ATP synthase-coupling factor 6 mitochondrial OS=Rattus norvegicus OX=10116 GN=Atp5pf PE=1 SV=1 |
| 77 | 225 | tr\|Q8SEZ8 | 53.86 | 7 | 7 | 8.04E+05 | 3 | 3 | 3 | | 36133 | NADH-ubiquinone oxidoreductase chain 1 (Fragment) OS=Rattus norvegicus OX=10116 GN=NADH1 PE=3 SV=1 |
| 77 | 226 | P03889\|NU | 53.86 | 7 | 7 | 8.04E+05 | 3 | 3 | 3 | | 36145 | NADH-ubiquinone oxidoreductase chain 1 OS=Rattus norvegicus OX=10116 GN=Mtnd1 PE=1 SV=3 |
| 77 | 227 | tr\|Q8HID1 | 53.86 | 7 | 7 | 8.04E+05 | 3 | 3 | 3 | | 36145 | NADH-ubiquinone oxidoreductase chain 1 OS=Rattus norvegicus OX=10116 GN=Mt-nd1 PE=3 SV=1 |
| 77 | 228 | tr\|D2E6L7 | 53.86 | 7 | 7 | 8.04E+05 | 3 | 3 | 3 | | 36062 | NADH-ubiquinone oxidoreductase chain 1 OS=Rattus norvegicus OX=10116 GN=ND1 PE=3 SV=1 |
| 93 | 188 | O70351\|H | 52.99 | 7 | 7 | 2.70E+05 | 2 | 2 | 3 | | 27246 | 3-hydroxyacyl-CoA dehydrogenase type-2 OS=Rattus norvegicus OX=10116 GN=Hsd17b10 PE=1 SV=3 |
| 93 | 189 | tr\|B0BMW | 52.99 | 7 | 7 | 2.70E+05 | 2 | 2 | 3 | | 27250 | 3-hydroxyacyl-CoA dehydrogenase type-2 OS=Rattus norvegicus OX=10116 GN=Hsd17b10 PE=1 SV=1 |
| 81 | 250 | P08011\|M | 52.63 | 15 | 15 | 6.43E+05 | 3 | 3 | 3 | | 17472 | Microsomal glutathione S-transferase 1 OS=Rattus norvegicus OX=10116 GN=Mgst1 PE=1 SV=3 |
| 81 | 251 | tr\|B6DYQ4 | 52.63 | 15 | 15 | 6.43E+05 | 3 | 3 | 3 | | 17472 | Microsomal glutathione S-transferase OS=Rattus norvegicus OX=10116 GN=Mgst1 PE=2 SV=1 |
| 117 | 306 | Q2V057\|H | 50.95 | 4 | 4 | 1.31E+06 | 2 | 2 | 2 | | 51002 | Hydroxyproline dehydrogenase OS=Rattus norvegicus OX=10116 GN=Prodh2 PE=2 SV=1 |
| 92 | 247 | P10888\|CC | 50.93 | 12 | 12 | 3.89E+05 | 2 | 2 | 3 | | 19515 | Cytochrome c oxidase subunit 4 isoform 1 mitochondrial OS=Rattus norvegicus OX=10116 GN=Cox4i1 PE=1 SV=1 |
| 133 | 195 | P49432\|OI | 50.13 | 3 | 3 | 5.85E+04 | 2 | 2 | 2 | | 38982 | Pyruvate dehydrogenase E1 component subunit beta mitochondrial OS=Rattus norvegicus OX=10116 GN=Pdhb PE=1 SV=2 |
| 133 | 196 | tr\|A0A0G2 | 50.13 | 2 | 2 | 5.85E+04 | 2 | 2 | 2 | | 46193 | Pyruvate dehydrogenase E1 component subunit beta OS=Rattus norvegicus OX=10116 GN=Pdhb PE=1 SV=1 |
| 101 | 229 | tr\|Q06QK0 | 49.91 | 4 | 4 | 3.48E+05 | 2 | 2 | 2 | | 56880 | Cytochrome c oxidase subunit 1 OS=Rattus norvegicus OX=10116 GN=CO1 PE=3 SV=1 |
| 101 | 230 | tr\|Q8SEZ6 | 49.91 | 4 | 4 | 3.48E+05 | 2 | 2 | 2 | | 56879 | Cytochrome c oxidase subunit 1 OS=Rattus norvegicus OX=10116 GN=COI PE=3 SV=1 |
| 101 | 232 | tr\|Q8HIC9 | 49.91 | 4 | 4 | 3.48E+05 | 2 | 2 | 2 | | 56845 | Cytochrome c oxidase subunit 1 OS=Rattus norvegicus OX=10116 GN=Mt-co1 PE=3 SV=1 |
| 101 | 233 | P05503\|CC | 49.91 | 4 | 4 | 3.48E+05 | 2 | 2 | 2 | | 56845 | Cytochrome c oxidase subunit 1 OS=Rattus norvegicus OX=10116 GN=Mtco1 PE=2 SV=3 |
| 101 | 241 | tr\|Q06QA9 | 49.91 | 4 | 4 | 3.48E+05 | 2 | 2 | 2 | | 56893 | Cytochrome c oxidase subunit 1 OS=Rattus norvegicus OX=10116 GN=CO1 PE=3 SV=1 |
| 101 | 231 | tr\|A0A0A1 | 49.91 | 4 | 4 | 3.48E+05 | 2 | 2 | 2 | | 56937 | Cytochrome c oxidase subunit 1 OS=Rattus norvegicus OX=10116 GN=COX1 PE=3 SV=1 |
| 101 | 234 | tr\|Q95938 | 49.91 | 4 | 4 | 3.48E+05 | 2 | 2 | 2 | | 56977 | Cytochrome c oxidase subunit 1 OS=Rattus norvegicus OX=10116 GN=Co I PE=3 SV=1 |
| 112 | 218 | P63039\|CH | 49.63 | 4 | 4 | 1.98E+05 | 2 | 2 | 2 | Oxidation ( | 60956 | 60 kDa heat shock protein mitochondrial OS=Rattus norvegicus OX=10116 GN=Hspd1 PE=1 SV=1 |
| 112 | 219 | tr\|A0A482 | 49.63 | 4 | 4 | 1.98E+05 | 2 | 2 | 2 | Oxidation ( | 60956 | Hsp60 OS=Rattus norvegicus OX=10116 GN=Hspd1 PE=2 SV=1 |
| 164 | 208 | P13803\|ET | 47.83 | 3 | 3 | 3.79E+05 | 1 | 1 | 1 | | 34951 | Electron transfer flavoprotein subunit alpha mitochondrial OS=Rattus norvegicus OX=10116 GN=Etfa PE=1 SV=4 |
| 114 | 235 | tr\|B2RYW3 | 46.63 | 13 | 13 | 1.76E+05 | 2 | 2 | 2 | Oxidation ( | 21892 | NADH dehydrogenase (Ubiquinone) 1 beta subcomplex 9 OS=Rattus norvegicus OX=10116 GN=Ndufb9 PE=1 SV=1 |

| | | | | | | | | | | | | |
|---|---|---|---|---|---|---|---|---|---|---|---|---|
| 90 | 307 | P16970\|AE | 45.09 | 5 | 5 | 8.50E+05 | 3 | 3 | 3 | | 75316 | ATP-binding cassette sub-family D member 3 OS=Rattus norvegicus OX=10116 GN=Abcd3 PE=1 SV=3 |
| 68 | 322 | P12075\|CC | 44.85 | 28 | 28 | 1.11E+06 | 4 | 4 | 4 | | 13915 | Cytochrome c oxidase subunit 5B mitochondrial OS=Rattus norvegicus OX=10116 GN=Cox5b PE=1 SV=2 |
| 83 | 133 | Q63362\|N | 44.2 | 22 | 22 | 5.09E+05 | 3 | 2 | 3 | | 13412 | NADH dehydrogenase [ubiquinone] 1 alpha subcomplex subunit 5 OS=Rattus norvegicus OX=10116 GN=Ndufa5 PE=1 SV=3 |
| 107 | 401 | tr\|A0A0G2 | 42.53 | 1 | 1 | 0 | 2 | 2 | 2 | | 275755 | Acetyl-CoA carboxylase beta OS=Rattus norvegicus OX=10116 GN=Acacb PE=1 SV=1 |
| 107 | 327 | tr\|A0A0G2 | 42.53 | 1 | 1 | 0 | 2 | 2 | 2 | | 276393 | Acetyl-CoA carboxylase beta OS=Rattus norvegicus OX=10116 GN=Acacb PE=1 SV=1 |
| 107 | 402 | tr\|O70151 | 42.53 | 1 | 1 | 0 | 2 | 2 | 2 | | 276097 | Acetyl-CoA carboxylase OS=Rattus norvegicus OX=10116 GN=Acacb PE=2 SV=1 |
| 107 | 328 | tr\|D3ZBE2 | 42.53 | 1 | 1 | 0 | 2 | 2 | 2 | | 275967 | Acetyl-CoA carboxylase beta OS=Rattus norvegicus OX=10116 GN=Acacb PE=1 SV=3 |
| 107 | 329 | tr\|E9PSQ0 | 42.53 | 1 | 1 | 0 | 2 | 2 | 2 | | 276255 | Acetyl-CoA carboxylase beta OS=Rattus norvegicus OX=10116 GN=Acacb PE=1 SV=2 |
| 121 | 146 | tr\|B0BMT9 | 41.79 | 3 | 3 | 3.56E+05 | 2 | 1 | 2 | | 50202 | Sqrdl protein OS=Rattus norvegicus OX=10116 GN=Sqor PE=1 SV=1 |
| 111 | 436 | D4A2Z8\|DI | 41.13 | 2 | 2 | 7.38E+05 | 2 | 1 | 2 | | 113843 | ATP-dependent DNA/RNA helicase DHX36 OS=Rattus norvegicus OX=10116 GN=Dhx36 PE=1 SV=1 |
| 74 | 482 | tr\|D3ZHF8 | 39.93 | 1 | 1 | 1.79E+06 | 3 | 2 | 4 | Formylatio | 73207 | Translation factor GUF1 mitochondrial OS=Rattus norvegicus OX=10116 GN=Guf1 PE=1 SV=1 |
| 84 | 216 | Q9R063\|PI | 38.9 | 9 | 9 | 1.10E+06 | 2 | 2 | 3 | | 22179 | Peroxiredoxin-5 mitochondrial OS=Rattus norvegicus OX=10116 GN=Prdx5 PE=1 SV=1 |
| 84 | 217 | tr\|A0A0G2 | 38.9 | 9 | 9 | 1.10E+06 | 2 | 2 | 3 | | 22207 | Peroxiredoxin OS=Rattus norvegicus OX=10116 GN=Prdx5 PE=1 SV=1 |
| 155 | 281 | tr\|F1LPV8\| | 38.46 | 2 | 2 | 1.64E+05 | 1 | 1 | 1 | | 46639 | Succinate--CoA ligase [GDP-forming] subunit beta mitochondrial OS=Rattus norvegicus OX=10116 GN=Suclg2 PE=1 SV=2 |
| 155 | 282 | tr\|B1H270 | 38.46 | 2 | 2 | 1.64E+05 | 1 | 1 | 1 | | 46988 | Succinate--CoA ligase [GDP-forming] subunit beta mitochondrial OS=Rattus norvegicus OX=10116 GN=Suclg2 PE=2 SV=1 |
| 217 | 341 | tr\|G3V6I5\| | 37.59 | 3 | 3 | 1.85E+05 | 1 | 1 | 1 | | 46777 | DnaJ heat shock protein family (Hsp40) member A3 OS=Rattus norvegicus OX=10116 GN=Dnaja3 PE=1 SV=2 |
| 217 | 342 | tr\|Q2TVU3 | 37.59 | 3 | 3 | 1.85E+05 | 1 | 1 | 1 | | 49412 | TID1 OS=Rattus norvegicus OX=10116 GN=Dnaja3 PE=2 SV=1 |
| 217 | 343 | tr\|A0A0G2 | 37.59 | 3 | 3 | 1.85E+05 | 1 | 1 | 1 | | 49416 | DnaJ heat shock protein family (Hsp40) member A3 OS=Rattus norvegicus OX=10116 GN=Dnaja3 PE=1 SV=1 |
| 217 | 344 | tr\|Q2UZS7 | 37.59 | 2 | 2 | 1.85E+05 | 1 | 1 | 1 | | 52399 | Tid-1 long isoform OS=Rattus norvegicus OX=10116 GN=Dnaja3 PE=2 SV=1 |
| 217 | 345 | tr\|A0A0G2 | 37.59 | 2 | 2 | 1.85E+05 | 1 | 1 | 1 | | 52403 | DnaJ heat shock protein family (Hsp40) member A3 OS=Rattus norvegicus OX=10116 GN=Dnaja3 PE=1 SV=1 |
| 189 | 264 | tr\|D3ZTW8 | 37.32 | 8 | 8 | 1.28E+05 | 1 | 1 | 1 | | 15823 | Mitochondrial ribosomal protein L27 OS=Rattus norvegicus OX=10116 GN=Mrpl27 PE=1 SV=2 |
| 95 | 11746 | Q9JJW3\|AT | 36.62 | 21 | 21 | 8.21E+05 | 2 | 2 | 3 | | 6408 | ATP synthase membrane subunit DAPIT mitochondrial OS=Rattus norvegicus OX=10116 GN=Atp5md PE=1 SV=1 |
| 123 | 283 | Q9WVK3\|P | 36.01 | 7 | 7 | 1.78E+05 | 2 | 2 | 2 | Oxidation ( | 32433 | Peroxisomal trans-2-enoyl-CoA reductase OS=Rattus norvegicus OX=10116 GN=Pecr PE=2 SV=1 |
| 123 | 284 | tr\|A0A0G2 | 36.01 | 7 | 7 | 1.78E+05 | 2 | 2 | 2 | Oxidation ( | 32737 | Peroxisomal trans-2-enoyl-CoA reductase OS=Rattus norvegicus OX=10116 GN=Pecr PE=1 SV=1 |
| 75 | 555 | tr\|D4A1D3 | 35.67 | 0 | 0 | 3.07E+06 | 3 | 1 | 3 | | 521487 | Sacsin molecular chaperone OS=Rattus norvegicus OX=10116 GN=Sacs PE=1 SV=2 |
| 263 | 11742 | P56571\|ES | 35.33 | 4 | 4 | 1.35E+05 | 1 | 1 | 1 | | 28173 | ES1 protein homolog mitochondrial OS=Rattus norvegicus OX=10116 PE=1 SV=2 |
| 82 | 531 | tr\|D3ZHK4 | 34.97 | 1 | 1 | | 2 | 0 | 3 | | 182226 | RB1-inducible coiled-coil 1 OS=Rattus norvegicus OX=10116 GN=Rb1cc1 PE=1 SV=1 |
| 118 | 318 | tr\|F1LM33 | 34.76 | 1 | 1 | | 2 | 0 | 2 | | 156679 | Leucine-rich PPR motif-containing protein mitochondrial OS=Rattus norvegicus OX=10116 GN=Lrpprc PE=1 SV=2 |
| 118 | 319 | Q5SGE0\|LI | 34.76 | 1 | 1 | | 2 | 0 | 2 | | 156652 | Leucine-rich PPR motif-containing protein mitochondrial OS=Rattus norvegicus OX=10116 GN=Lrpprc PE=1 SV=1 |
| 66 | 236 | tr\|Q5U2T0 | 34.01 | 4 | 4 | 0 | 2 | 1 | 4 | | 44494 | Death associated protein 3 OS=Rattus norvegicus OX=10116 GN=Dap3 PE=2 SV=1 |
| 66 | 237 | tr\|F7EZZ0\| | 34.01 | 4 | 4 | 0 | 2 | 1 | 4 | | 45112 | Death-associated protein 3 OS=Rattus norvegicus OX=10116 GN=Dap3 PE=1 SV=1 |
| 66 | 242 | tr\|A0A0G2 | 34.01 | 4 | 4 | 0 | 2 | 1 | 4 | | 46666 | Death-associated protein 3 OS=Rattus norvegicus OX=10116 GN=Dap3 PE=1 SV=1 |
| 264 | 244 | P09139\|SP | 33.97 | 3 | 3 | 0 | 1 | 1 | 1 | | 45834 | Serine--pyruvate aminotransferase mitochondrial OS=Rattus norvegicus OX=10116 GN=Agxt PE=1 SV=1 |
| 265 | 5796 | B1WC61\|A | 33.34 | 2 | 2 | 8.85E+04 | 1 | 1 | 1 | | 68843 | Complex I assembly factor ACAD9 mitochondrial OS=Rattus norvegicus OX=10116 GN=Acad9 PE=1 SV=1 |
| 266 | 5819 | tr\|B2RYU0 | 32.86 | 9 | 9 | 2.43E+05 | 1 | 1 | 1 | | 11842 | NADH dehydrogenase (Ubiquinone) 1 beta subcomplex 2 (Predicted) isoform CRA_b OS=Rattus norvegicus OX=10116 GN=Ndufb2 PE=1 SV=1 |
| 113 | 517 | tr\|F1M4Y5 | 32.7 | 1 | 1 | 3.05E+05 | 2 | 2 | 2 | Formylatio | 108218 | Storkhead box 1 OS=Rattus norvegicus OX=10116 GN=Stox1 PE=4 SV=2 |
| 267 | 312 | tr\|D3ZGM1 | 32.68 | 1 | 1 | 1.68E+05 | 1 | 1 | 1 | | 77891 | Pentatricopeptide repeat domain 3 OS=Rattus norvegicus OX=10116 GN=Ptcd3 PE=1 SV=1 |
| 267 | 313 | tr\|A0A0G2 | 32.68 | 1 | 1 | 1.68E+05 | 1 | 1 | 1 | | 84557 | Pentatricopeptide repeat domain 3 OS=Rattus norvegicus OX=10116 GN=Ptcd3 PE=1 SV=1 |
| 103 | 474 | tr\|A0A0G2 | 32.67 | 2 | 2 | 1.04E+06 | 2 | 2 | 2 | Formylatio | 133221 | Adhesion G protein-coupled receptor G6 OS=Rattus norvegicus OX=10116 GN=Adgrg6 PE=1 SV=1 |
| 173 | 52 | tr\|M0RAM | 32.37 | 6 | 6 | 1.78E+05 | 1 | 1 | 1 | | 22155 | Glutathione peroxidase OS=Rattus norvegicus OX=10116 GN=Gpx1 PE=1 SV=1 |
| 173 | 53 | P04041\|GI | 32.37 | 5 | 5 | 1.78E+05 | 1 | 1 | 1 | | 22305 | Glutathione peroxidase 1 OS=Rattus norvegicus OX=10116 GN=Gpx1 PE=1 SV=4 |
| 94 | 1525 | Q62925\|M | 32.14 | 2 | 2 | 2.45E+05 | 3 | 3 | 3 | Oxidation (M) | | |
| 268 | 5783 | tr\|Q35733 | 32.1 | 13 | 13 | 9.21E+04 | 1 | 1 | 1 | | 8389 | NADH-ubiquinone oxidoreductase chain 6 (Fragment) OS=Rattus norvegicus OX=10116 PE=3 SV=1 |
| 268 | 5784 | tr\|Q06QD9 | 32.1 | 6 | 6 | 9.21E+04 | 1 | 1 | 1 | | 18943 | NADH-ubiquinone oxidoreductase chain 6 OS=Rattus norvegicus OX=10116 GN=ND6 PE=3 SV=1 |
| 268 | 5785 | tr\|Q7HKW | 32.1 | 6 | 6 | 9.21E+04 | 1 | 1 | 1 | | 18957 | NADH-ubiquinone oxidoreductase chain 6 OS=Rattus norvegicus OX=10116 GN=NADH6 PE=3 SV=1 |
| 269 | 11743 | P35171\|CX | 32.05 | 11 | 11 | 0 | 1 | 1 | 1 | | 9353 | Cytochrome c oxidase subunit 7A2 mitochondrial OS=Rattus norvegicus OX=10116 GN=Cox7a2 PE=1 SV=1 |
| 269 | 11744 | tr\|B2RYS0 | 32.05 | 11 | 11 | 0 | 1 | 1 | 1 | | 9353 | Cox7a2 protein OS=Rattus norvegicus OX=10116 GN=Cox7a2 PE=2 SV=1 |
| 270 | 361 | P80433\|CC | 31.69 | 14 | 14 | 2.07E+05 | 1 | 1 | 1 | | 7672 | Cytochrome c oxidase subunit 8A mitochondrial OS=Rattus norvegicus OX=10116 GN=Cox8a PE=1 SV=3 |
| 271 | 197 | tr\|M0R6J0 | 31.13 | 3 | 3 | 1.45E+05 | 1 | 1 | 1 | | 38374 | Mitochondrial ribosomal protein L39 OS=Rattus norvegicus OX=10116 GN=Mrpl39 PE=1 SV=2 |
| 129 | 656 | tr\|D3ZD23 | 31.1 | 1 | 1 | 9.92E+05 | 2 | 2 | 2 | | 67300 | ATP-binding cassette subfamily E member 1 OS=Rattus norvegicus OX=10116 GN=Abce1 PE=1 SV=1 |
| 216 | 290 | tr\|A0A0G2 | 30.61 | 4 | 4 | 2.94E+05 | 1 | 1 | 1 | | 20750 | Mitochondrial ribosomal protein L11 OS=Rattus norvegicus OX=10116 GN=mrpl11 PE=1 SV=1 |
| 216 | 291 | Q5XIE3\|RM | 30.61 | 4 | 4 | 2.94E+05 | 1 | 1 | 1 | | 22420 | 39S ribosomal protein L11 mitochondrial OS=Rattus norvegicus OX=10116 GN=Mrpl11 PE=2 SV=1 |
| 134 | 414 | tr\|D4A4P3 | 30.54 | 15 | 15 | 5.86E+05 | 2 | 2 | 2 | | 11267 | NADH:ubiquinone oxidoreductase subunit B3 OS=Rattus norvegicus OX=10116 GN=Ndufb3 PE=1 SV=1 |
| 218 | 321 | tr\|Q5BJZ3\| | 30.53 | 1 | 1 | 4.58E+05 | 1 | 1 | 1 | | 113869 | Nicotinamide nucleotide transhydrogenase OS=Rattus norvegicus OX=10116 GN=Nnt PE=1 SV=1 |
| 190 | 289 | tr\|Q5XFW4 | 29.56 | 7 | 7 | 1.42E+05 | 1 | 1 | 1 | | 20541 | Mitochondrial ribosomal protein L13 OS=Rattus norvegicus OX=10116 GN=Mrpl13 PE=1 SV=1 |
| 145 | 6514 | tr\|D4A3D8 | 28.61 | 1 | 1 | 5.51E+04 | 2 | 1 | 2 | | 104860 | XPC complex subunit DNA damage recognition and repair factor OS=Rattus norvegicus OX=10116 GN=Xpc PE=1 SV=1 |
| 272 | 11745 | P32089\|TX | 28.46 | 4 | 4 | 0 | 1 | 1 | 1 | | 33835 | Tricarboxylate transport protein mitochondrial OS=Rattus norvegicus OX=10116 GN=Slc25a1 PE=1 SV=1 |
| 127 | 1412 | Q8CGU6\|N | 27.86 | 3 | 3 | 1.42E+05 | 2 | 1 | 2 | | 78400 | Nicastrin OS=Rattus norvegicus OX=10116 GN=Ncstn PE=1 SV=1 |
| 105 | 380 | P11530\|DI | 27.4 | 0 | 0 | 3.63E+05 | 2 | 2 | 2 | | 425830 | Dystrophin OS=Rattus norvegicus OX=10116 GN=Dmd PE=1 SV=2 |
| 119 | 1248 | Q80ZG6\|B | 27.12 | 10 | 10 | 5.05E+05 | 2 | 2 | 2 | | 20718 | Bcl-2-binding component 3 OS=Rattus norvegicus OX=10116 GN=Bbc3 PE=1 SV=1 |
| 64 | 461 | tr\|A0A0G2 | 26.98 | 0 | 0 | 9.47E+05 | 2 | 1 | 4 | | 487857 | Vacuolar protein sorting 13 homolog D OS=Rattus norvegicus OX=10116 GN=Vps13d PE=1 SV=1 |
| 64 | 462 | tr\|D3ZKC6 | 26.98 | 0 | 0 | 9.47E+05 | 2 | 1 | 4 | | 488962 | Vacuolar protein sorting 13 homolog D OS=Rattus norvegicus OX=10116 GN=Vps13d PE=1 SV=1 |
| 274 | 379 | tr\|D3ZDP2 | 26.3 | 4 | 4 | 1.59E+05 | 1 | 1 | 1 | | 23470 | Mitochondrial ribosomal protein L58 OS=Rattus norvegicus OX=10116 GN=Mrpl58 PE=1 SV=1 |

<table>
<tr><td>137</td><td>562</td><td>P21575|DY</td><td>26.07</td><td>1</td><td>1</td><td>4.60E+05</td><td>1</td><td>1</td><td>2</td><td></td><td>97295</td><td>Dynamin-1 OS=Rattus norvegicus OX=10116 GN=Dnm1 PE=1 SV=2</td></tr>
<tr><td>275</td><td>349</td><td>Q5I0C5|FM</td><td>25.81</td><td>2</td><td>2</td><td>0</td><td>1</td><td>1</td><td>1</td><td>Formylatio</td><td>42997</td><td>Methionyl-tRNA formyltransferase  mitochondrial OS=Rattus norvegicus OX=10116 GN=Mtfmt PE=2 SV=1</td></tr>
<tr><td>116</td><td>398</td><td>P48037|AN</td><td>25.78</td><td>1</td><td>1</td><td>9.49E+05</td><td>1</td><td>1</td><td>2</td><td></td><td>75754</td><td>Annexin A6 OS=Rattus norvegicus OX=10116 GN=Anxa6 PE=1 SV=2</td></tr>
<tr><td>116</td><td>399</td><td>tr|Q6IMZ3</td><td>25.78</td><td>1</td><td>1</td><td>9.49E+05</td><td>1</td><td>1</td><td>2</td><td></td><td>75756</td><td>Annexin OS=Rattus norvegicus OX=10116 GN=Anxa6 PE=1 SV=1</td></tr>
<tr><td>96</td><td>514</td><td>tr|Q5RKL4</td><td>25.52</td><td>1</td><td>1</td><td>9.39E+04</td><td>1</td><td>1</td><td>3</td><td></td><td>95977</td><td>Dimethylglycine dehydrogenase OS=Rattus norvegicus OX=10116 GN=Dmgdh PE=1 SV=1</td></tr>
<tr><td>96</td><td>515</td><td>Q63342|M</td><td>25.52</td><td>1</td><td>1</td><td>9.39E+04</td><td>1</td><td>1</td><td>3</td><td></td><td>96047</td><td>Dimethylglycine dehydrogenase  mitochondrial OS=Rattus norvegicus OX=10116 GN=Dmgdh PE=1 SV=1</td></tr>
<tr><td>96</td><td>516</td><td>tr|A0A0G2</td><td>25.52</td><td>1</td><td>1</td><td>9.39E+04</td><td>1</td><td>1</td><td>3</td><td></td><td>98864</td><td>Dimethylglycine dehydrogenase  mitochondrial OS=Rattus norvegicus OX=10116 GN=Dmgdh PE=1 SV=1</td></tr>
<tr><td>277</td><td>1802</td><td>tr|D3ZY44</td><td>25.4</td><td>3</td><td>3</td><td>0</td><td>1</td><td>1</td><td>1</td><td></td><td>32176</td><td>Mitochondrial ribosomal protein S2 OS=Rattus norvegicus OX=10116 GN=Mrps2 PE=1 SV=1</td></tr>
<tr><td>177</td><td>520</td><td>tr|A0A0G2</td><td>25.21</td><td>1</td><td>1</td><td>1.85E+05</td><td>1</td><td>1</td><td>1</td><td></td><td>74235</td><td>Selenoprotein O OS=Rattus norvegicus OX=10116 GN=Selenoo PE=1 SV=1</td></tr>
<tr><td>177</td><td>521</td><td>tr|B2GVA1</td><td>25.21</td><td>1</td><td>1</td><td>1.85E+05</td><td>1</td><td>1</td><td>1</td><td></td><td>74385</td><td>Selenoprotein O OS=Rattus norvegicus OX=10116 GN=Selenoo PE=2 SV=1</td></tr>
<tr><td>131</td><td>11756</td><td>D3ZAF6|AT</td><td>25.19</td><td>14</td><td>14</td><td>3.59E+05</td><td>2</td><td>1</td><td>2</td><td></td><td>10452</td><td>ATP synthase subunit f  mitochondrial OS=Rattus norvegicus OX=10116 GN=Atp5mf PE=1 SV=1</td></tr>
<tr><td>140</td><td>432</td><td>tr|F1LSP2|</td><td>25.09</td><td>1</td><td>1</td><td>4.26E+04</td><td>2</td><td>1</td><td>2</td><td></td><td>119190</td><td>Acyl-CoA dehydrogenase family  member 10 OS=Rattus norvegicus OX=10116 GN=Acad10 PE=1 SV=2</td></tr>
<tr><td>279</td><td>338</td><td>Q5M9I5|Q</td><td>24.93</td><td>17</td><td>17</td><td>1.26E+05</td><td>1</td><td>1</td><td>1</td><td></td><td>10424</td><td>Cytochrome b-c1 complex subunit 6  mitochondrial OS=Rattus norvegicus OX=10116 GN=Uqcrh PE=3 SV=1</td></tr>
<tr><td>144</td><td>2046</td><td>tr|A0A0H2</td><td>24.86</td><td>1</td><td>1</td><td>0</td><td>2</td><td>1</td><td>2</td><td></td><td>118367</td><td>Valine--tRNA ligase  mitochondrial OS=Rattus norvegicus OX=10116 GN=Vars2 PE=1 SV=1</td></tr>
<tr><td>144</td><td>2047</td><td>Q6MG21|S</td><td>24.86</td><td>1</td><td>1</td><td>0</td><td>2</td><td>1</td><td>2</td><td></td><td>118894</td><td>Valine--tRNA ligase  mitochondrial OS=Rattus norvegicus OX=10116 GN=Vars2 PE=3 SV=1</td></tr>
<tr><td>280</td><td>1055</td><td>tr|D3ZL85|</td><td>24.56</td><td>4</td><td>4</td><td>3.31E+05</td><td>1</td><td>1</td><td>1</td><td></td><td>31130</td><td>Cytochrome c heme lyase OS=Rattus norvegicus OX=10116 GN=Hccs PE=1 SV=1</td></tr>
<tr><td>128</td><td>1719</td><td>Q673L6|TE</td><td>24.25</td><td>2</td><td>2</td><td>7.50E+05</td><td>2</td><td>2</td><td>2</td><td></td><td>126934</td><td>Telomerase reverse transcriptase OS=Rattus norvegicus OX=10116 GN=Tert PE=2 SV=1</td></tr>
<tr><td>136</td><td>419</td><td>Q01062|PI</td><td>23.71</td><td>2</td><td>2</td><td>9.59E+05</td><td>1</td><td>1</td><td>2</td><td></td><td>104664</td><td>cGMP-dependent 3' 5'-cyclic phosphodiesterase OS=Rattus norvegicus OX=10116 GN=Pde2a PE=1 SV=2</td></tr>
<tr><td>136</td><td>420</td><td>tr|F8WFW</td><td>23.71</td><td>1</td><td>1</td><td>9.59E+05</td><td>1</td><td>1</td><td>2</td><td></td><td>105234</td><td>Phosphodiesterase OS=Rattus norvegicus OX=10116 GN=Pde2a PE=1 SV=1</td></tr>
<tr><td>281</td><td>11749</td><td>tr|A0A0U1</td><td>23.67</td><td>9</td><td>9</td><td>1.15E+05</td><td>1</td><td>1</td><td>1</td><td></td><td>10726</td><td>5-demethoxyubiquinone hydroxylase  mitochondrial (Fragment) OS=Rattus norvegicus OX=10116 GN=Coq7 PE=1 SV=1</td></tr>
<tr><td>281</td><td>11750</td><td>tr|G3V879</td><td>23.67</td><td>5</td><td>5</td><td>1.15E+05</td><td>1</td><td>1</td><td>1</td><td></td><td>20126</td><td>5-demethoxyubiquinone hydroxylase  mitochondrial OS=Rattus norvegicus OX=10116 GN=Coq7 PE=1 SV=4</td></tr>
<tr><td>174</td><td>760</td><td>D3ZG52|D</td><td>23.6</td><td>1</td><td>1</td><td>3.08E+05</td><td>1</td><td>1</td><td>1</td><td></td><td>119588</td><td>DNA replication ATP-dependent helicase/nuclease DNA2 OS=Rattus norvegicus OX=10116 GN=Dna2 PE=3 SV=1</td></tr>
<tr><td>174</td><td>761</td><td>tr|A0A0H2</td><td>23.6</td><td>1</td><td>1</td><td>3.08E+05</td><td>1</td><td>1</td><td>1</td><td></td><td>136526</td><td>Graves disease carrier protein OS=Rattus norvegicus OX=10116 GN=Slc25a16 PE=1 SV=1</td></tr>
<tr><td>174</td><td>11751</td><td>P61980|HN</td><td>23.6</td><td>2</td><td>2</td><td>3.08E+05</td><td>1</td><td>1</td><td>1</td><td></td><td>50976</td><td>Heterogeneous nuclear ribonucleoprotein K OS=Rattus norvegicus OX=10116 GN=Hnrnpk PE=1 SV=1</td></tr>
<tr><td>97</td><td>481</td><td>tr|D4A4K4</td><td>23.29</td><td>0</td><td>0</td><td>2.96E+04</td><td>2</td><td>2</td><td>2</td><td></td><td>418626</td><td>Vacuolar protein sorting 13 homolog C OS=Rattus norvegicus OX=10116 GN=Vps13c PE=1 SV=2</td></tr>
<tr><td>109</td><td>294</td><td>Q01728|N</td><td>22.99</td><td>1</td><td>1</td><td>5.57E+06</td><td>1</td><td>1</td><td>2</td><td>Formylatio</td><td>108184</td><td>Sodium/calcium exchanger 1 OS=Rattus norvegicus OX=10116 GN=Slc8a1 PE=1 SV=3</td></tr>
<tr><td>109</td><td>359</td><td>P70549|NA</td><td>22.99</td><td>1</td><td>1</td><td>5.57E+06</td><td>1</td><td>1</td><td>2</td><td>Formylatio</td><td>103163</td><td>Sodium/calcium exchanger 3 OS=Rattus norvegicus OX=10116 GN=Slc8a3 PE=1 SV=1</td></tr>
<tr><td>193</td><td>725</td><td>tr|F1LP30|</td><td>22.53</td><td>1</td><td>1</td><td></td><td>1</td><td>0</td><td>1</td><td></td><td>79296</td><td>Methylcrotonoyl-CoA carboxylase subunit alpha  mitochondrial OS=Rattus norvegicus OX=10116 GN=Mccc1 PE=1 SV=1</td></tr>
<tr><td>193</td><td>726</td><td>Q5I0C3|M</td><td>22.53</td><td>1</td><td>1</td><td></td><td>1</td><td>0</td><td>1</td><td></td><td>79330</td><td>Methylcrotonoyl-CoA carboxylase subunit alpha  mitochondrial OS=Rattus norvegicus OX=10116 GN=Mccc1 PE=1 SV=1</td></tr>
<tr><td>167</td><td>1089</td><td>tr|A0A0G2</td><td>22.27</td><td>1</td><td>1</td><td>0</td><td>1</td><td>1</td><td>1</td><td></td><td>77504</td><td>MMR_HSR1 domain containing protein RGD1359460  isoform CRA_a OS=Rattus norvegicus OX=10116 GN=Noa1 PE=4 SV=1</td></tr>
<tr><td>167</td><td>1081</td><td>tr|Q5XHZ7</td><td>22.27</td><td>1</td><td>1</td><td>0</td><td>1</td><td>1</td><td>1</td><td></td><td>66660</td><td>MMR_HSR1 domain containing protein RGD1359460 OS=Rattus norvegicus OX=10116 GN=Noa1 PE=2 SV=1</td></tr>
<tr><td>167</td><td>1082</td><td>tr|A0A0G2</td><td>22.27</td><td>1</td><td>1</td><td>0</td><td>1</td><td>1</td><td>1</td><td></td><td>66708</td><td>Nitric oxide-associated 1 OS=Rattus norvegicus OX=10116 GN=Noa1 PE=4 SV=1</td></tr>
<tr><td>157</td><td>1405</td><td>tr|A0A0H2</td><td>22.02</td><td>1</td><td>1</td><td>0</td><td>1</td><td>1</td><td>1</td><td></td><td>99265</td><td>Alanine--tRNA ligase  mitochondrial OS=Rattus norvegicus OX=10116 GN=Aars2 PE=1 SV=1</td></tr>
<tr><td>157</td><td>1414</td><td>D3ZX08|SY</td><td>22.02</td><td>1</td><td>1</td><td>0</td><td>1</td><td>1</td><td>1</td><td></td><td>107806</td><td>Alanine--tRNA ligase  mitochondrial OS=Rattus norvegicus OX=10116 GN=Aars2 PE=3 SV=1</td></tr>
<tr><td>284</td><td>872</td><td>Q5FWU3|A</td><td>21.26</td><td>1</td><td>1</td><td>3.98E+04</td><td>1</td><td>1</td><td>1</td><td>Oxidation (</td><td>94488</td><td>Autophagy-related protein 9A OS=Rattus norvegicus OX=10116 GN=Atg9a PE=1 SV=1</td></tr>
<tr><td>70</td><td>599</td><td>Q00918|LT</td><td>21.03</td><td>1</td><td>1</td><td>2.15E+06</td><td>1</td><td>1</td><td>4</td><td>Carbamido</td><td>186598</td><td>Latent-transforming growth factor beta-binding protein 1 OS=Rattus norvegicus OX=10116 GN=Ltbp1 PE=1 SV=1</td></tr>
<tr><td>148</td><td>593</td><td>P15205|M</td><td>20.91</td><td>0</td><td>0</td><td></td><td>1</td><td>0</td><td>1</td><td></td><td>269640</td><td>Microtubule-associated protein 1B OS=Rattus norvegicus OX=10116 GN=Map1b PE=1 SV=3</td></tr>
<tr><td>182</td><td>6206</td><td>tr|B5DFK5</td><td>20.91</td><td>0</td><td>0</td><td></td><td>1</td><td>0</td><td>1</td><td></td><td>119424</td><td>Hip1r protein OS=Rattus norvegicus OX=10116 GN=Hip1r PE=2 SV=1</td></tr>
<tr><td>182</td><td>6207</td><td>tr|F1LML7</td><td>20.91</td><td>0</td><td>0</td><td></td><td>1</td><td>0</td><td>1</td><td></td><td>119553</td><td>Huntingtin-interacting protein 1-related OS=Rattus norvegicus OX=10116 GN=Hip1r PE=1 SV=1</td></tr>
<tr><td>182</td><td>6221</td><td>tr|Q99PW</td><td>20.91</td><td>0</td><td>0</td><td></td><td>1</td><td>0</td><td>1</td><td></td><td>120572</td><td>Huntingtin interacting protein 1 related (Fragment) OS=Rattus norvegicus OX=10116 GN=Hip1r PE=2 SV=1</td></tr>
<tr><td>286</td><td>5893</td><td>tr|F1LP46|</td><td>20.68</td><td>1</td><td>1</td><td>1.87E+05</td><td>1</td><td>1</td><td>1</td><td></td><td>78923</td><td>ATP-dependent RNA helicase SUPV3L1  mitochondrial OS=Rattus norvegicus OX=10116 GN=Supv3l1 PE=1 SV=2</td></tr>
<tr><td>286</td><td>5894</td><td>tr|A0A0G2</td><td>20.68</td><td>1</td><td>1</td><td>1.87E+05</td><td>1</td><td>1</td><td>1</td><td></td><td>84180</td><td>ATP-dependent RNA helicase SUPV3L1  mitochondrial OS=Rattus norvegicus OX=10116 GN=Supv3l1 PE=1 SV=1</td></tr>
<tr><td>286</td><td>5895</td><td>Q5EBA1|SU</td><td>20.68</td><td>1</td><td>1</td><td>1.87E+05</td><td>1</td><td>1</td><td>1</td><td></td><td>86706</td><td>ATP-dependent RNA helicase SUPV3L1  mitochondrial OS=Rattus norvegicus OX=10116 GN=Supv3l1 PE=2 SV=1</td></tr>
<tr><td>191</td><td>683</td><td>O88658|KI</td><td>20.68</td><td>0</td><td>0</td><td>6.77E+05</td><td>1</td><td>1</td><td>1</td><td></td><td>204169</td><td>Kinesin-like protein KIF1B OS=Rattus norvegicus OX=10116 GN=Kif1b PE=1 SV=2</td></tr>
</table>
