## Supplementary material for "Permeability transition pore-related changes in the proteome and channel activity of ATP synthase dimers and monomers": RLM SM-BM PTP Monomer of V compl.

| Protein Gr | Protein ID | Accession | -10lgP | Coverage ( | Coverage ( | Area Samp | #Peptides | #Unique | #Spec Sam | PTM | Avg. Mass | Description |
|---|---|---|---|---|---|---|---|---|---|---|---|---|
| 2 | 8 | P10719\|AT | 405.85 | 71 | 71 | 4.98E+09 | 241 | 233 | 875 | Oxidation (M) | | |
| 2 | 9 | tr\|G3V6D3 | 405.85 | 71 | 71 | 4.98E+09 | 241 | 233 | 875 | Oxidation (M) | | |
| 1 | 10 | P15999\|AT | 367.07 | 63 | 63 | 3.88E+09 | 248 | 237 | 1140 | Carbamidomethylation | | |
| 1 | 11 | tr\|F1LP05\| | 367.07 | 63 | 63 | 3.88E+09 | 248 | 237 | 1140 | Carbamidomethylation | | |
| 4 | 119 | P31399\|AT | 349.97 | 82 | 82 | 6.50E+08 | 87 | 87 | 194 | Carbamidomethylation | | |
| 3 | 30 | P52873\|PY | 337.26 | 43 | 43 | 4.12E+08 | 164 | 159 | 292 | Oxidation (M) | | |
| 3 | 31 | tr\|A0A0G2 | 337.26 | 39 | 39 | 4.12E+08 | 164 | 159 | 292 | Oxidation (M) | | |
| 6 | 64 | P35435\|AT | 305.24 | 53 | 53 | 5.30E+08 | 80 | 80 | 182 | Carbamidomethylation | | |
| 6 | 63 | tr\|Q6QI09 | 305.24 | 24 | 24 | 5.30E+08 | 80 | 80 | 182 | Carbamidomethylation | | |
| 8 | 1 | P10860\|DH | 293.87 | 41 | 41 | 1.18E+08 | 66 | 66 | 107 | Oxidation ( | 61416 | Glutamate dehydrogenase 1 mitochondrial OS=Rattus norvegicus OX=10116 GN=Glud1 PE=1 SV=2 |
| 19 | 171 | tr\|D3ZFJ6\| | 266.34 | 25 | 25 | 1.96E+07 | 26 | 26 | 32 | | 60420 | Lactamase beta OS=Rattus norvegicus OX=10116 GN=Lactb PE=1 SV=1 |
| 9 | 4 | P07756\|CP | 256.7 | 26 | 26 | 7.50E+07 | 74 | 71 | 98 | Carbamidomethylation | | |
| 5 | 246 | P19511\|AT | 238.91 | 48 | 48 | 6.10E+08 | 59 | 58 | 189 | Formylatio | 28869 | ATP synthase F(0) complex subunit B1 mitochondrial OS=Rattus norvegicus OX=10116 GN=Atp5pb PE=1 SV=1 |
| 12 | 26 | P00507\|AA | 237.65 | 36 | 36 | 7.64E+07 | 40 | 39 | 58 | Oxidation ( | 47314 | Aspartate aminotransferase mitochondrial OS=Rattus norvegicus OX=10116 GN=Got2 PE=1 SV=2 |
| 7 | 5802 | Q06647\|AT | 236.85 | 54 | 54 | 2.64E+08 | 63 | 61 | 124 | Carbamidomethylation | | |
| 15 | 5817 | P21571\|AT | 234.2 | 58 | 58 | 4.55E+07 | 29 | 28 | 45 | Oxidation (M) | | |
| 10 | 1646 | Q6PDU7\|A | 228.57 | 50 | 50 | 2.59E+08 | 38 | 38 | 94 | Oxidation ( | 11433 | ATP synthase subunit g mitochondrial OS=Rattus norvegicus OX=10116 GN=Atp5mg PE=1 SV=2 |
| 11 | 350 | tr\|G3V7Y3 | 213.55 | 48 | 48 | 1.91E+08 | 26 | 26 | 63 | Oxidation ( | 17563 | ATP synthase subunit delta mitochondrial OS=Rattus norvegicus OX=10116 GN=Atp5f1d PE=1 SV=1 |
| 14 | 45 | P04762\|CA | 211.13 | 27 | 27 | 4.90E+07 | 30 | 30 | 46 | | 59757 | Catalase OS=Rattus norvegicus OX=10116 GN=Cat PE=1 SV=3 |
| 20 | 218 | P63039\|CH | 193.69 | 24 | 24 | 1.88E+07 | 25 | 25 | 29 | Oxidation ( | 60956 | 60 kDa heat shock protein mitochondrial OS=Rattus norvegicus OX=10116 GN=Hspd1 PE=1 SV=1 |
| 20 | 219 | tr\|A0A482 | 193.69 | 24 | 24 | 1.88E+07 | 25 | 25 | 29 | Oxidation ( | 60956 | Hsp60 OS=Rattus norvegicus OX=10116 GN=Hspd1 PE=2 SV=1 |
| 18 | 27 | Q60587\|EC | 190.48 | 27 | 27 | 2.30E+07 | 25 | 25 | 33 | | 51414 | Trifunctional enzyme subunit beta mitochondrial OS=Rattus norvegicus OX=10116 GN=Hadhb PE=1 SV=1 |
| 16 | 24 | Q64428\|EC | 185.48 | 16 | 16 | 2.21E+07 | 20 | 20 | 35 | Carbamido | 82665 | Trifunctional enzyme subunit alpha mitochondrial OS=Rattus norvegicus OX=10116 GN=Hadha PE=1 SV=2 |
| 22 | 23 | P22791\|HM | 176.62 | 23 | 23 | 1.23E+07 | 21 | 21 | 27 | | 56912 | Hydroxymethylglutaryl-CoA synthase mitochondrial OS=Rattus norvegicus OX=10116 GN=Hmgcs2 PE=1 SV=1 |
| 22 | 22 | tr\|Q68G44 | 176.62 | 23 | 23 | 1.23E+07 | 21 | 21 | 27 | | 56886 | 3-hydroxy-3-methylglutaryl coenzyme A synthase OS=Rattus norvegicus OX=10116 GN=Hmgcs2 PE=1 SV=1 |
| 30 | 136 | P24329\|TH | 171.16 | 18 | 18 | 4.65E+06 | 7 | 7 | 9 | | 33407 | Thiosulfate sulfurtransferase OS=Rattus norvegicus OX=10116 GN=Tst PE=1 SV=3 |
| 17 | 352 | P0C2X9\|AL | 165.56 | 23 | 23 | 1.59E+07 | 26 | 26 | 34 | Oxidation ( | 61869 | Delta-1-pyrroline-5-carboxylate dehydrogenase mitochondrial OS=Rattus norvegicus OX=10116 GN=Aldh4a1 PE=1 SV=1 |
| 24 | 54 | P18163\|AC | 158.35 | 15 | 15 | 1.53E+07 | 17 | 17 | 21 | Formylatio | 78179 | Long-chain-fatty-acid--CoA ligase 1 OS=Rattus norvegicus OX=10116 GN=Acsl1 PE=1 SV=1 |
| 25 | 50 | Q02253\|M | 156.3 | 12 | 12 | 8.45E+06 | 13 | 13 | 16 | Oxidation ( | 57808 | Methylmalonate-semialdehyde dehydrogenase [acylating] mitochondrial OS=Rattus norvegicus OX=10116 GN=Aldh6a1 PE=1 SV=1 |
| 25 | 51 | tr\|G3V7J0 | 156.3 | 12 | 12 | 8.45E+06 | 13 | 13 | 16 | Oxidation ( | 57748 | Aldehyde dehydrogenase family 6 subfamily A1 isoform CRA_b OS=Rattus norvegicus OX=10116 GN=Aldh6a1 PE=1 SV=1 |
| 13 | 11740 | P29419\|AT | 155.85 | 73 | 73 | 6.61E+07 | 28 | 26 | 56 | | 8255 | ATP synthase subunit e mitochondrial OS=Rattus norvegicus OX=10116 GN=Atp5me PE=1 SV=3 |
| 36 | 201 | P45953\|AC | 151.85 | 9 | 9 | 1.19E+06 | 6 | 6 | 6 | | 70749 | Very long-chain specific acyl-CoA dehydrogenase mitochondrial OS=Rattus norvegicus OX=10116 GN=Acadvl PE=1 SV=1 |
| 36 | 202 | tr\|Q5M9H | 151.85 | 9 | 9 | 1.19E+06 | 6 | 6 | 6 | | 70821 | Acyl-Coenzyme A dehydrogenase very long chain OS=Rattus norvegicus OX=10116 GN=Acadvl PE=1 SV=1 |
| 26 | 35 | P17764\|TH | 142.6 | 15 | 15 | 9.93E+06 | 10 | 10 | 16 | | 44695 | Acetyl-CoA acetyltransferase mitochondrial OS=Rattus norvegicus OX=10116 GN=Acat1 PE=1 SV=1 |
| 27 | 46 | P13086\|SU | 141.11 | 17 | 17 | 4.12E+06 | 12 | 12 | 16 | Formylatio | 36148 | Succinate--CoA ligase [ADP/GDP-forming] subunit alpha mitochondrial OS=Rattus norvegicus OX=10116 GN=Suclg1 PE=2 SV=2 |
| 27 | 47 | tr\|A0A0H2 | 141.11 | 16 | 16 | 4.12E+06 | 12 | 12 | 16 | Formylatio | 37560 | Succinate--CoA ligase [ADP/GDP-forming] subunit alpha mitochondrial OS=Rattus norvegicus OX=10116 GN=Suclg1 PE=1 SV=1 |
| 34 | 742 | A2VCW9\|A | 139.69 | 8 | 8 | 1.97E+06 | 7 | 7 | 7 | Carbamido | 102908 | Alpha-aminoadipic semialdehyde synthase mitochondrial OS=Rattus norvegicus OX=10116 GN=Aass PE=2 SV=1 |
| 23 | 1095 | tr\|Q5UAJ5 | 135.59 | 69 | 69 | 9.31E+07 | 10 | 10 | 22 | Oxidation ( | 7642 | ATP synthase protein 8 OS=Rattus norvegicus OX=10116 GN=ATP8 PE=3 SV=1 |
| 23 | 1097 | tr\|Q8SEZ4 | 135.59 | 69 | 69 | 9.31E+07 | 10 | 10 | 22 | Oxidation ( | 7632 | ATP synthase protein 8 OS=Rattus norvegicus OX=10116 GN=ATPase8 PE=3 SV=1 |
| 72 | 277 | tr\|B2RZ24 | 128.18 | 5 | 5 | 5.06E+05 | 3 | 3 | 3 | | 47388 | Succinate-CoA ligase subunit beta (Fragment) OS=Rattus norvegicus OX=10116 GN=Sucla2 PE=2 SV=1 |
| 72 | 278 | tr\|F1LM47 | 128.18 | 5 | 5 | 5.06E+05 | 3 | 3 | 3 | | 50306 | Succinate--CoA ligase [ADP-forming] subunit beta mitochondrial OS=Rattus norvegicus OX=10116 GN=Sucla2 PE=1 SV=1 |
| 48 | 42 | P20788\|UC | 125.14 | 10 | 10 | 2.14E+06 | 4 | 4 | 5 | Formylatio | 29446 | Cytochrome b-c1 complex subunit Rieske mitochondrial OS=Rattus norvegicus OX=10116 GN=Uqcrfs1 PE=1 SV=2 |
| 39 | 28533 | P26772\|CH | 122.56 | 38 | 38 | 1.08E+06 | 6 | 6 | 6 | Oxidation (M) | | |
| 33 | 11785 | B3DMA2\|A | 113.7 | 8 | 8 | 3.55E+06 | 7 | 7 | 8 | | 87371 | Acyl-CoA dehydrogenase family member 11 OS=Rattus norvegicus OX=10116 GN=Acad11 PE=1 SV=1 |
| 35 | 207 | tr\|G3V6I4\| | 112.27 | 8 | 8 | 1.57E+06 | 4 | 4 | 7 | | 37791 | Mitochondrial amidoxime reducing component 1 OS=Rattus norvegicus OX=10116 GN=Marc1 PE=1 SV=1 |
| 21 | 690 | P05504\|AT | 108.74 | 38 | 38 | 2.32E+07 | 18 | 16 | 27 | Oxidation (M) | | |
| 21 | 691 | tr\|Q8HIC7 | 108.74 | 38 | 38 | 2.32E+07 | 18 | 16 | 27 | Oxidation (M) | | |
| 21 | 692 | tr\|S5S1E9\| | 108.74 | 38 | 38 | 2.32E+07 | 18 | 16 | 27 | Oxidation (M) | | |
| 45 | 38 | tr\|D3ZFQ8 | 105.46 | 6 | 6 | 3.11E+06 | 3 | 3 | 5 | | 35435 | Cytochrome c-1 OS=Rattus norvegicus OX=10116 GN=Cyc1 PE=1 SV=3 |
| 29 | 21 | P32551\|QC | 104.27 | 5 | 5 | 4.67E+06 | 6 | 6 | 11 | | 48396 | Cytochrome b-c1 complex subunit 2 mitochondrial OS=Rattus norvegicus OX=10116 GN=Uqcrc2 PE=1 SV=2 |
| 31 | 11746 | Q9JJW3\|AT | 103.21 | 43 | 43 | 1.22E+07 | 6 | 6 | 9 | | 6408 | ATP synthase membrane subunit DAPIT mitochondrial OS=Rattus norvegicus OX=10116 GN=Atp5md PE=1 SV=1 |
| 49 | 68 | tr\|A0A140 | 102.62 | 10 | 10 | 9.56E+05 | 4 | 4 | 5 | | 42574 | Cytochrome b (Fragment) OS=Rattus norvegicus OX=10116 PE=3 SV=1 |
| 49 | 69 | tr\|A0A140 | 102.62 | 10 | 10 | 9.56E+05 | 4 | 4 | 5 | | 42588 | Cytochrome b (Fragment) OS=Rattus norvegicus OX=10116 PE=3 SV=1 |
| 49 | 70 | tr\|A0A140 | 102.62 | 10 | 10 | 9.56E+05 | 4 | 4 | 5 | | 42527 | Cytochrome b (Fragment) OS=Rattus norvegicus OX=10116 PE=3 SV=1 |
| 49 | 92 | tr\|A0A0S1 | 102.62 | 9 | 9 | 9.56E+05 | 4 | 4 | 5 | | 43002 | Cytochrome b OS=Rattus norvegicus OX=10116 GN=CYTB PE=3 SV=1 |
| 49 | 95 | tr\|A0A0S1 | 102.62 | 9 | 9 | 9.56E+05 | 4 | 4 | 5 | | 43016 | Cytochrome b OS=Rattus norvegicus OX=10116 GN=CYTB PE=3 SV=1 |
| 49 | 72 | tr\|A0A3Q8 | 102.62 | 10 | 10 | 9.56E+05 | 4 | 4 | 5 | | 42622 | Cytochrome b (Fragment) OS=Rattus norvegicus OX=10116 GN=Cytb PE=3 SV=1 |
| 49 | 120 | tr\|A0A411 | 102.62 | 10 | 10 | 9.56E+05 | 4 | 4 | 5 | | 40144 | Cytochrome b (Fragment) OS=Rattus norvegicus OX=10116 GN=Cytb PE=3 SV=1 |
| 49 | 132 | tr\|A0A0U2 | 102.62 | 10 | 10 | 9.56E+05 | 4 | 4 | 5 | | 40981 | Cytochrome b (Fragment) OS=Rattus norvegicus OX=10116 PE=3 SV=1 |
| 49 | 122 | tr\|A0A411 | 102.62 | 10 | 10 | 9.56E+05 | 4 | 4 | 5 | | 41017 | Cytochrome b (Fragment) OS=Rattus norvegicus OX=10116 GN=Cytb PE=3 SV=1 |
| 49 | 121 | tr\|A0A0U2 | 102.62 | 10 | 10 | 9.56E+05 | 4 | 4 | 5 | | 41037 | Cytochrome b (Fragment) OS=Rattus norvegicus OX=10116 PE=3 SV=1 |
| 49 | 123 | tr\|F8QU31 | 102.62 | 10 | 10 | 9.56E+05 | 4 | 4 | 5 | | 42181 | Cytochrome b (Fragment) OS=Rattus norvegicus OX=10116 GN=cytb PE=3 SV=1 |

| | | | | | | | | | | | | |
|---|---|---|---|---|---|---|---|---|---|---|---|---|
| 49 | 124 | tr\|A0A411 | 102.62 | 10 | 10 | 9.56E+05 | 4 | 4 | 5 | | 42238 | Cytochrome b (Fragment) OS=Rattus norvegicus OX=10116 GN=Cytb PE=3 SV=1 |
| 49 | 67 | tr\|L0L4L8\| | 102.62 | 10 | 10 | 9.56E+05 | 4 | 4 | 5 | | 42470 | Cytochrome b (Fragment) OS=Rattus norvegicus OX=10116 GN=cytb PE=3 SV=1 |
| 49 | 125 | tr\|A0A385 | 102.62 | 10 | 10 | 9.56E+05 | 4 | 4 | 5 | | 42766 | Cytochrome b (Fragment) OS=Rattus norvegicus OX=10116 PE=3 SV=1 |
| 49 | 71 | tr\|A0A3Q8 | 102.62 | 10 | 10 | 9.56E+05 | 4 | 4 | 5 | | 42712 | Cytochrome b (Fragment) OS=Rattus norvegicus OX=10116 GN=Cytb PE=3 SV=1 |
| 49 | 73 | tr\|A0A3S6 | 102.62 | 10 | 10 | 9.56E+05 | 4 | 4 | 5 | | 42702 | Cytochrome b (Fragment) OS=Rattus norvegicus OX=10116 GN=Cytb PE=3 SV=1 |
| 49 | 74 | tr\|A0A3S6 | 102.62 | 10 | 10 | 9.56E+05 | 4 | 4 | 5 | | 42698 | Cytochrome b (Fragment) OS=Rattus norvegicus OX=10116 GN=Cytb PE=3 SV=1 |
| 49 | 80 | tr\|D6NSR3 | 102.62 | 9 | 9 | 9.56E+05 | 4 | 4 | 5 | | 42986 | Cytochrome b (Fragment) OS=Rattus norvegicus OX=10116 GN=cytb PE=3 SV=1 |
| 49 | 81 | tr\|D6NSQ8 | 102.62 | 9 | 9 | 9.56E+05 | 4 | 4 | 5 | | 43016 | Cytochrome b (Fragment) OS=Rattus norvegicus OX=10116 GN=cytb PE=3 SV=1 |
| 49 | 82 | tr\|D6NSR8 | 102.62 | 9 | 9 | 9.56E+05 | 4 | 4 | 5 | | 42998 | Cytochrome b (Fragment) OS=Rattus norvegicus OX=10116 GN=cytb PE=3 SV=1 |
| 49 | 103 | tr\|F8QU37 | 102.62 | 9 | 9 | 9.56E+05 | 4 | 4 | 5 | | 42992 | Cytochrome b (Fragment) OS=Rattus norvegicus OX=10116 GN=cytb PE=3 SV=1 |
| 49 | 86 | tr\|Q8SEY9 | 102.62 | 9 | 9 | 9.56E+05 | 4 | 4 | 5 | | 43015 | Cytochrome b OS=Rattus norvegicus OX=10116 GN=cytb PE=3 SV=1 |
| 49 | 87 | tr\|D6NSR7 | 102.62 | 9 | 9 | 9.56E+05 | 4 | 4 | 5 | | 42982 | Cytochrome b (Fragment) OS=Rattus norvegicus OX=10116 GN=cytb PE=3 SV=1 |
| 49 | 89 | tr\|F2Q6S5 | 102.62 | 9 | 9 | 9.56E+05 | 4 | 4 | 5 | | 42952 | Cytochrome b (Fragment) OS=Rattus norvegicus OX=10116 GN=cytb PE=3 SV=1 |
| 49 | 90 | tr\|D6NSQ6 | 102.62 | 9 | 9 | 9.56E+05 | 4 | 4 | 5 | | 42952 | Cytochrome b (Fragment) OS=Rattus norvegicus OX=10116 GN=cytb PE=3 SV=1 |
| 49 | 116 | tr\|D6NSQ0 | 102.62 | 9 | 9 | 9.56E+05 | 4 | 4 | 5 | | 42968 | Cytochrome b (Fragment) OS=Rattus norvegicus OX=10116 GN=cytb PE=3 SV=1 |
| 49 | 91 | tr\|Q5UAI7 | 102.62 | 9 | 9 | 9.56E+05 | 4 | 4 | 5 | | 42998 | Cytochrome b OS=Rattus norvegicus OX=10116 GN=CYTB PE=3 SV=1 |
| 49 | 129 | tr\|D6NSR1 | 102.62 | 9 | 9 | 9.56E+05 | 4 | 4 | 5 | | 43002 | Cytochrome b (Fragment) OS=Rattus norvegicus OX=10116 GN=cytb PE=3 SV=1 |
| 49 | 93 | tr\|Q8HIC4 | 102.62 | 9 | 9 | 9.56E+05 | 4 | 4 | 5 | | 43012 | Cytochrome b OS=Rattus norvegicus OX=10116 GN=Mt-cyb PE=3 SV=1 |
| 49 | 97 | P00159\|CY | 102.62 | 9 | 9 | 9.56E+05 | 4 | 4 | 5 | | 43012 | Cytochrome b OS=Rattus norvegicus OX=10116 GN=Mt-Cyb PE=3 SV=3 |
| 49 | 99 | tr\|A0A220 | 102.62 | 9 | 9 | 9.56E+05 | 4 | 4 | 5 | | 42970 | Cytochrome b OS=Rattus norvegicus OX=10116 PE=3 SV=1 |
| 49 | 100 | tr\|H2KXA0 | 102.62 | 9 | 9 | 9.56E+05 | 4 | 4 | 5 | | 42978 | Cytochrome b (Fragment) OS=Rattus norvegicus OX=10116 GN=cytb PE=3 SV=1 |
| 49 | 101 | tr\|A0A096 | 102.62 | 9 | 9 | 9.56E+05 | 4 | 4 | 5 | | 43042 | Cytochrome b (Fragment) OS=Rattus norvegicus OX=10116 PE=3 SV=1 |
| 49 | 76 | tr\|A0A220 | 102.62 | 9 | 9 | 9.56E+05 | 4 | 4 | 5 | | 43018 | Cytochrome b OS=Rattus norvegicus OX=10116 PE=3 SV=1 |
| 49 | 77 | tr\|D6NSP2 | 102.62 | 9 | 9 | 9.56E+05 | 4 | 4 | 5 | | 42945 | Cytochrome b (Fragment) OS=Rattus norvegicus OX=10116 GN=cytb PE=3 SV=1 |
| 49 | 78 | tr\|D6NSS6 | 102.62 | 9 | 9 | 9.56E+05 | 4 | 4 | 5 | | 42993 | Cytochrome b (Fragment) OS=Rattus norvegicus OX=10116 GN=cytb PE=3 SV=1 |
| 49 | 79 | tr\|D6NSQ3 | 102.62 | 9 | 9 | 9.56E+05 | 4 | 4 | 5 | | 43012 | Cytochrome b (Fragment) OS=Rattus norvegicus OX=10116 GN=cytb PE=3 SV=1 |
| 49 | 127 | tr\|D6NSP4 | 102.62 | 9 | 9 | 9.56E+05 | 4 | 4 | 5 | | 43016 | Cytochrome b (Fragment) OS=Rattus norvegicus OX=10116 GN=cytb PE=3 SV=1 |
| 49 | 83 | tr\|F2Q6S6 | 102.62 | 9 | 9 | 9.56E+05 | 4 | 4 | 5 | | 42938 | Cytochrome b (Fragment) OS=Rattus norvegicus OX=10116 GN=cytb PE=3 SV=1 |
| 49 | 84 | tr\|A0A0A1 | 102.62 | 9 | 9 | 9.56E+05 | 4 | 4 | 5 | | 42989 | Cytochrome b OS=Rattus norvegicus OX=10116 GN=CYTB PE=3 SV=1 |
| 49 | 85 | tr\|D6NSS7 | 102.62 | 9 | 9 | 9.56E+05 | 4 | 4 | 5 | | 42948 | Cytochrome b (Fragment) OS=Rattus norvegicus OX=10116 GN=cytb PE=3 SV=1 |
| 49 | 88 | tr\|A0A220 | 102.62 | 9 | 9 | 9.56E+05 | 4 | 4 | 5 | | 42968 | Cytochrome b OS=Rattus norvegicus OX=10116 PE=3 SV=1 |
| 49 | 128 | tr\|D6NSR2 | 102.62 | 9 | 9 | 9.56E+05 | 4 | 4 | 5 | | 43073 | Cytochrome b (Fragment) OS=Rattus norvegicus OX=10116 GN=cytb PE=3 SV=1 |
| 49 | 94 | tr\|A0A220 | 102.62 | 9 | 9 | 9.56E+05 | 4 | 4 | 5 | | 42968 | Cytochrome b OS=Rattus norvegicus OX=10116 PE=3 SV=1 |
| 49 | 96 | tr\|R9TKN1 | 102.62 | 9 | 9 | 9.56E+05 | 4 | 4 | 5 | | 43012 | Cytochrome b OS=Rattus norvegicus OX=10116 GN=CYTB PE=3 SV=1 |
| 49 | 98 | tr\|L0N311 | 102.62 | 9 | 9 | 9.56E+05 | 4 | 4 | 5 | | 43045 | Cytochrome b (Fragment) OS=Rattus norvegicus OX=10116 GN=cytb PE=3 SV=1 |
| 49 | 102 | tr\|D6NSP5 | 102.62 | 9 | 9 | 9.56E+05 | 4 | 4 | 5 | | 43012 | Cytochrome b (Fragment) OS=Rattus norvegicus OX=10116 GN=cytb PE=3 SV=1 |
| 44 | 189 | tr\|B0BMW | 98.87 | 13 | 13 | 1.97E+06 | 5 | 5 | 5 | | 27250 | 3-hydroxyacyl-CoA dehydrogenase type-2 OS=Rattus norvegicus OX=10116 GN=Hsd17b10 PE=1 SV=1 |
| 44 | 188 | O70351\|H | 98.87 | 13 | 13 | 1.97E+06 | 5 | 5 | 5 | | 27246 | 3-hydroxyacyl-CoA dehydrogenase type-2 OS=Rattus norvegicus OX=10116 GN=Hsd17b10 PE=1 SV=3 |
| 50 | 3 | P85834\|EF | 94.86 | 7 | 7 | 1.44E+06 | 4 | 4 | 5 | | 49522 | Elongation factor Tu mitochondrial OS=Rattus norvegicus OX=10116 GN=Tufm PE=1 SV=1 |
| 107 | 44 | P11240\|CC | 90.83 | 17 | 17 | 5.29E+05 | 2 | 2 | 2 | | 16130 | Cytochrome c oxidase subunit 5A mitochondrial OS=Rattus norvegicus OX=10116 GN=Cox5a PE=1 SV=1 |
| 61 | 5862 | tr\|G3V7I0\| | 84.56 | 9 | 9 | 2.07E+06 | 3 | 3 | 4 | | 28299 | Peroxiredoxin 3 OS=Rattus norvegicus OX=10116 GN=Prdx3 PE=1 SV=1 |
| 61 | 5863 | Q9Z0V6\|PI | 84.56 | 9 | 9 | 2.07E+06 | 3 | 3 | 4 | | 28295 | Thioredoxin-dependent peroxide reductase mitochondrial OS=Rattus norvegicus OX=10116 GN=Prdx3 PE=1 SV=2 |
| 57 | 16 | Q68FY0\|Q | 82.22 | 7 | 7 | 1.50E+06 | 4 | 4 | 4 | | 52849 | Cytochrome b-c1 complex subunit 1 mitochondrial OS=Rattus norvegicus OX=10116 GN=Uqcrc1 PE=1 SV=1 |
| 40 | 297 | Q924S5\|LC | 81.14 | 3 | 3 | 1.17E+06 | 4 | 4 | 6 | | 105792 | Lon protease homolog mitochondrial OS=Rattus norvegicus OX=10116 GN=Lonp1 PE=2 SV=1 |
| 28 | 11756 | D3ZAF6\|AT | 77.21 | 30 | 30 | 2.00E+07 | 8 | 7 | 15 | Oxidation ( | 10452 | ATP synthase subunit f mitochondrial OS=Rattus norvegicus OX=10116 GN=Atp5mf PE=1 SV=1 |
| 32 | 6375 | tr\|D3ZTP0 | 76.77 | 6 | 6 | 2.78E+06 | 6 | 6 | 8 | | 101806 | 10-formyltetrahydrofolate dehydrogenase OS=Rattus norvegicus OX=10116 GN=Aldh1l2 PE=1 SV=3 |
| 78 | 34377 | P19804\|NI | 74.8 | 15 | 15 | 7.93E+05 | 2 | 2 | 3 | Formylatio | 17283 | Nucleoside diphosphate kinase B OS=Rattus norvegicus OX=10116 GN=Nme2 PE=1 SV=1 |
| 90 | 115 | tr\|A0A0G2 | 74.63 | 2 | 2 | 1.52E+06 | 2 | 2 | 2 | | 86230 | MICOS complex subunit MIC60 OS=Rattus norvegicus OX=10116 GN=Immt PE=1 SV=1 |
| 90 | 109 | tr\|A0A140 | 74.63 | 3 | 3 | 1.52E+06 | 2 | 2 | 2 | | 67049 | MICOS complex subunit MIC60 OS=Rattus norvegicus OX=10116 GN=Immt PE=1 SV=1 |
| 90 | 110 | Q3KR86\|M | 74.63 | 3 | 3 | 1.52E+06 | 2 | 2 | 2 | | 67177 | MICOS complex subunit Mic60 (Fragment) OS=Rattus norvegicus OX=10116 GN=Immt PE=1 SV=1 |
| 47 | 5804 | tr\|A0A0G2 | 72.95 | 2 | 2 | 1.36E+05 | 4 | 1 | 5 | | 79812 | Propionyl-CoA carboxylase alpha chain mitochondrial OS=Rattus norvegicus OX=10116 GN=Pcca PE=1 SV=1 |
| 47 | 5805 | tr\|A0A0G2 | 72.95 | 2 | 2 | 1.36E+05 | 4 | 1 | 5 | | 81413 | Propionyl-CoA carboxylase alpha chain mitochondrial OS=Rattus norvegicus OX=10116 GN=Pcca PE=1 SV=1 |
| 47 | 5806 | P14882\|PC | 72.95 | 2 | 2 | 1.36E+05 | 4 | 1 | 5 | | 81623 | Propionyl-CoA carboxylase alpha chain mitochondrial OS=Rattus norvegicus OX=10116 GN=Pcca PE=1 SV=3 |
| 67 | 240 | A0A0G2K0 | 72.24 | 3 | 3 | 1.52E+05 | 2 | 2 | 3 |  | 74675 | Acyl-CoA synthetase short-chain family member 3 mitochondrial OS=Rattus norvegicus OX=10116 GN=Acss3 PE=1 SV=1 |
| 38 | 251 | tr\|B6DYQ4 | 71.07 | 15 | 15 | 1.10E+06 | 5 | 5 | 6 | | 17472 | Microsomal glutathione S-transferase OS=Rattus norvegicus OX=10116 GN=Mgst1 PE=2 SV=1 |
| 38 | 250 | P08011\|M | 71.07 | 15 | 15 | 1.10E+06 | 5 | 5 | 6 | | 17472 | Microsomal glutathione S-transferase 1 OS=Rattus norvegicus OX=10116 GN=Mgst1 PE=1 SV=3 |
| 83 | 143 | tr\|Q5EBA4 | 70.7 | 8 | 8 | 4.40E+05 | 2 | 2 | 3 | Oxidation ( | 33215 | Nipsnap1 protein (Fragment) OS=Rattus norvegicus OX=10116 GN=Nipsnap1 PE=2 SV=1 |
| 83 | 144 | tr\|G3V728 | 70.7 | 8 | 8 | 4.40E+05 | 2 | 2 | 3 | Oxidation ( | 33346 | 4-nitrophenylphosphatase domain and non-neuronal SNAP25-like protein homolog 1 (C. elegans) isoform CRA_b OS=Rattus norvegicus OX=10116 GN=Nipsnap1 PE=1 SV=1 |
| 236 | 11743 | P35171\|CX | 65.29 | 14 | 14 | 3.12E+05 | 1 | 1 | 1 | | 9353 | Cytochrome c oxidase subunit 7A2 mitochondrial OS=Rattus norvegicus OX=10116 GN=Cox7a2 PE=1 SV=1 |
| 236 | 11744 | tr\|B2RYS0 | 65.29 | 14 | 14 | 3.12E+05 | 1 | 1 | 1 | | 9353 | Cox7a2 protein OS=Rattus norvegicus OX=10116 GN=Cox7a2 PE=2 SV=1 |
| 76 | 1322 | P32198\|CF | 62.37 | 3 | 3 | 1.70E+05 | 2 | 2 | 3 | | 88126 | Carnitine O-palmitoyltransferase 1 liver isoform OS=Rattus norvegicus OX=10116 GN=Cpt1a PE=1 SV=2 |
| 41 | 446 | Q91XJ1\|BE | 58.57 | 3 | 3 | | 3 | 0 | 6 | | 51557 | Beclin-1 OS=Rattus norvegicus OX=10116 GN=Becn1 PE=1 SV=1 |
| 70 | 52 | tr\|M0RAM | 56.38 | 13 | 13 | 4.42E+05 | 3 | 3 | 3 | | 22155 | Glutathione peroxidase OS=Rattus norvegicus OX=10116 GN=Gpx1 PE=1 SV=1 |

| | | | | | | | | | | | |
|---|---|---|---|---|---|---|---|---|---|---|---|
| 70 | 53 | P04041\|GI | 56.38 | 13 | 13 | 4.42E+05 | 3 | 3 | 3 | | 22305 Glutathione peroxidase 1 OS=Rattus norvegicus OX=10116 GN=Gpx1 PE=1 SV=4 |
| 52 | 283 | Q9WVK3\|I | 55.65 | 13 | 13 | 3.89E+05 | 4 | 3 | 5 | Oxidation ( | 32433 Peroxisomal trans-2-enoyl-CoA reductase OS=Rattus norvegicus OX=10116 GN=Pecr PE=2 SV=1 |
| 52 | 284 | tr\|A0A0G2 | 55.65 | 13 | 13 | 3.89E+05 | 4 | 3 | 5 | Oxidation ( | 32737 Peroxisomal trans-2-enoyl-CoA reductase OS=Rattus norvegicus OX=10116 GN=Pecr PE=1 SV=1 |
| 95 | 321 | tr\|Q5BJZ3 | 55.11 | 2 | 2 | 4.81E+05 | 2 | 2 | 2 | | 113869 Nicotinamide nucleotide transhydrogenase OS=Rattus norvegicus OX=10116 GN=Nnt PE=1 SV=1 |
| 60 | 34376 | D3Z9R8\|AT | 51.97 | 32 | 32 | 3.06E+05 | 3 | 2 | 4 | Oxidation ( | 6914 ATP synthase subunit ATP5MPL mitochondrial OS=Rattus norvegicus OX=10116 GN=Atp5mpl PE=1 SV=1 |
| 79 | 294 | Q01728\|N | 51.52 | 2 | 2 | 6.24E+06 | 3 | 1 | 3 | Formylatio | 108184 Sodium/calcium exchanger 1 OS=Rattus norvegicus OX=10116 GN=Slc8a1 PE=1 SV=3 |
| 202 | 725 | tr\|F1LP30\| | 51.4 | 2 | 2 | 1.87E+05 | 1 | 1 | 1 | | 79296 Methylcrotonoyl-CoA carboxylase subunit alpha mitochondrial OS=Rattus norvegicus OX=10116 GN=Mccc1 PE=1 SV=1 |
| 202 | 726 | Q5I0C3\|M | 51.4 | 2 | 2 | 1.87E+05 | 1 | 1 | 1 | | 79330 Methylcrotonoyl-CoA carboxylase subunit alpha mitochondrial OS=Rattus norvegicus OX=10116 GN=Mccc1 PE=1 SV=1 |
| 159 | 208 | P13803\|ET | 49.5 | 3 | 3 | 2.19E+05 | 1 | 1 | 1 | | 34951 Electron transfer flavoprotein subunit alpha mitochondrial OS=Rattus norvegicus OX=10116 GN=Etfa PE=1 SV=4 |
| 200 | 287 | tr\|D3ZX69 | 47.83 | 5 | 5 | 0 | 1 | 1 | 1 | | 29466 39S ribosomal protein L10 mitochondrial OS=Rattus norvegicus OX=10116 GN=Mrpl10 PE=1 SV=1 |
| 200 | 223 | P0C2C4\|RI | 47.83 | 5 | 5 | 0 | 1 | 1 | 1 | | 29940 39S ribosomal protein L10 mitochondrial OS=Rattus norvegicus OX=10116 GN=Mrpl10 PE=1 SV=1 |
| 65 | 1178 | tr\|G3V9J8 | 47.36 | 3 | 3 | 1.16E+06 | 2 | 2 | 3 | Formylatio | 87521 Glycerol-3-phosphate acyltransferase 1 mitochondrial OS=Rattus norvegicus OX=10116 GN=Gpam PE=1 SV=2 |
| 65 | 1180 | P97564\|GI | 47.36 | 3 | 3 | 1.16E+06 | 2 | 2 | 3 | Formylatio | 93715 Glycerol-3-phosphate acyltransferase 1 mitochondrial OS=Rattus norvegicus OX=10116 GN=Gpam PE=1 SV=3 |
| 65 | 1179 | tr\|A0A0G2 | 47.36 | 3 | 3 | 1.16E+06 | 2 | 2 | 3 | Formylatio | 93728 Glycerol-3-phosphate acyltransferase 1 mitochondrial OS=Rattus norvegicus OX=10116 GN=Gpam PE=1 SV=1 |
| 237 | 17425 | tr\|A0A0G2 | 47.29 | 6 | 6 | 3.85E+05 | 1 | 1 | 1 | | 20804 Serine--pyruvate aminotransferase mitochondrial OS=Rattus norvegicus OX=10116 GN=Agxt PE=1 SV=1 |
| 237 | 244 | P09139\|SP | 47.29 | 3 | 3 | 3.85E+05 | 1 | 1 | 1 | | 45834 Serine--pyruvate aminotransferase mitochondrial OS=Rattus norvegicus OX=10116 GN=Agxt PE=1 SV=1 |
| 123 | 369 | tr\|G3V734 | 46.27 | 3 | 3 | 2.25E+05 | 2 | 1 | 2 | | 36133 2 4-dienoyl CoA reductase 1 mitochondrial isoform CRA_a OS=Rattus norvegicus OX=10116 GN=Decr1 PE=1 SV=1 |
| 123 | 370 | Q64591\|D | 46.27 | 3 | 3 | 2.25E+05 | 2 | 1 | 2 | | 36133 2 4-dienoyl-CoA reductase mitochondrial OS=Rattus norvegicus OX=10116 GN=Decr1 PE=1 SV=2 |
| 37 | 1405 | tr\|A0A0H2 | 46.08 | 2 | 2 | 3.05E+05 | 3 | 2 | 6 | | 99265 Alanine--tRNA ligase mitochondrial OS=Rattus norvegicus OX=10116 GN=Aars2 PE=1 SV=1 |
| 37 | 1414 | D3ZX08\|SY | 46.08 | 2 | 2 | 3.05E+05 | 3 | 2 | 6 | | 107806 Alanine--tRNA ligase mitochondrial OS=Rattus norvegicus OX=10116 GN=Aars2 PE=3 SV=1 |
| 43 | 531 | tr\|D3ZHK4 | 46.03 | 1 | 1 | 1.30E+06 | 4 | 1 | 5 | | 182226 RB1-inducible coiled-coil 1 OS=Rattus norvegicus OX=10116 GN=Rb1cc1 PE=1 SV=1 |
| 197 | 300 | Q5PQT3\|G | 45.52 | 3 | 3 | 0 | 1 | 1 | 1 | | 33899 Glycine N-acyltransferase OS=Rattus norvegicus OX=10116 GN=Glyat PE=2 SV=1 |
| 197 | 301 | tr\|A4PB92 | 45.52 | 3 | 3 | 0 | 1 | 1 | 1 | | 33899 Glycine N-acyltransferase OS=Rattus norvegicus OX=10116 GN=Glyat PE=2 SV=1 |
| 197 | 302 | tr\|A0A0G2 | 45.52 | 3 | 3 | 0 | 1 | 1 | 1 | | 39453 Glycine N-acyltransferase OS=Rattus norvegicus OX=10116 GN=Glyat PE=1 SV=1 |
| 197 | 1988 | Q9Z2Y0\|KI | 45.52 | 3 | 3 | 0 | 1 | 1 | 1 | | 34016 Glycine N-acyltransferase-like protein Keg1 OS=Rattus norvegicus OX=10116 GN=Keg1 PE=1 SV=2 |
| 197 | 303 | tr\|B1H250 | 45.52 | 3 | 3 | 0 | 1 | 1 | 1 | | 33986 Glycine-N-acyltransferase-like 1 OS=Rattus norvegicus OX=10116 GN=Glyatl1 PE=1 SV=1 |
| 238 | 137 | tr\|D4A7L4 | 44.08 | 9 | 9 | 1.36E+05 | 1 | 1 | 1 | | 17634 NADH dehydrogenase (Ubiquinone) 1 beta subcomplex 11 (Predicted) OS=Rattus norvegicus OX=10116 GN=Ndufb11 PE=1 SV=1 |
| 108 | 61 | P29147\|BI | 42.54 | 5 | 5 | 1.47E+06 | 2 | 2 | 2 | | 38202 D-beta-hydroxybutyrate dehydrogenase mitochondrial OS=Rattus norvegicus OX=10116 GN=Bdh1 PE=1 SV=2 |
| 108 | 62 | tr\|A0A0G2 | 42.54 | 5 | 5 | 1.47E+06 | 2 | 2 | 2 | | 38333 3-hydroxybutyrate dehydrogenase type 1 isoform CRA_a OS=Rattus norvegicus OX=10116 GN=Bdh1 PE=1 SV=1 |
| 239 | 1075 | Q9WVK7\|I | 42.53 | 5 | 5 | 1.76E+05 | 1 | 1 | 1 | Oxidation ( | 34448 Hydroxyacyl-coenzyme A dehydrogenase mitochondrial OS=Rattus norvegicus OX=10116 GN=Hadh PE=2 SV=1 |
| 59 | 619 | P35559\|ID | 41.38 | 1 | 1 | | 3 | 0 | 4 | | 117710 Insulin-degrading enzyme OS=Rattus norvegicus OX=10116 GN=Ide PE=1 SV=1 |
| 109 | 111 | tr\|S5RZM8 | 40.45 | 7 | 7 | 4.96E+05 | 2 | 2 | 2 | | 25958 Cytochrome c oxidase subunit 2 OS=Rattus norvegicus OX=10116 GN=COX2 PE=3 SV=1 |
| 109 | 112 | P00406\|CC | 40.45 | 7 | 7 | 4.96E+05 | 2 | 2 | 2 | | 25928 Cytochrome c oxidase subunit 2 OS=Rattus norvegicus OX=10116 GN=Mtco2 PE=1 SV=3 |
| 109 | 114 | tr\|A0A097 | 40.45 | 7 | 7 | 4.96E+05 | 2 | 2 | 2 | | 25894 Cytochrome c oxidase subunit 2 OS=Rattus norvegicus OX=10116 GN=COX2 PE=3 SV=1 |
| 109 | 104 | tr\|Q5UAJ6 | 40.45 | 7 | 7 | 4.96E+05 | 2 | 2 | 2 | | 25942 Cytochrome c oxidase subunit 2 OS=Rattus norvegicus OX=10116 GN=COX2 PE=3 SV=1 |
| 109 | 113 | tr\|Q8SEZ5 | 40.45 | 7 | 7 | 4.96E+05 | 2 | 2 | 2 | | 25928 Cytochrome c oxidase subunit 2 OS=Rattus norvegicus OX=10116 GN=Mt-co2 PE=1 SV=1 |
| 198 | 190 | P07824\|AI | 38.36 | 4 | 4 | 0 | 1 | 1 | 1 | | 34973 Arginase-1 OS=Rattus norvegicus OX=10116 GN=Arg1 PE=1 SV=2 |
| 180 | 306 | Q2V057\|H | 38.16 | 2 | 2 | 1.71E+05 | 1 | 1 | 1 | | 51002 Hydroxyproline dehydrogenase OS=Rattus norvegicus OX=10116 GN=Prodh2 PE=2 SV=1 |
| 62 | 11752 | tr\|D4A175 | 38.03 | 2 | 2 | | 2 | 0 | 4 | | 53667 tRNA dimethylallyltransferase OS=Rattus norvegicus OX=10116 GN=Trit1 PE=3 SV=1 |
| 240 | 247 | P10888\|CC | 37.76 | 8 | 8 | 2.38E+05 | 1 | 1 | 1 | | 19515 Cytochrome c oxidase subunit 4 isoform 1 mitochondrial OS=Rattus norvegicus OX=10116 GN=Cox4i1 PE=1 SV=1 |
| 199 | 34 | tr\|A0A0G2 | 37.02 | 7 | 7 | 1.39E+05 | 1 | 1 | 1 | | 19965 NADH dehydrogenase [ubiquinone] 1 alpha subcomplex subunit 8 OS=Rattus norvegicus OX=10116 GN=Ndufa8 PE=1 SV=1 |
| 120 | 5882 | Q5XII9\|MI | 35.65 | 5 | 5 | 3.83E+05 | 2 | 1 | 2 | | 31730 Mitochondrial fission regulator 1-like OS=Rattus norvegicus OX=10116 GN=Mtfr1l PE=1 SV=1 |
| 51 | 872 | Q5FWU3\|/ | 35.58 | 1 | 1 | 2.68E+06 | 2 | 2 | 5 | Oxidation ( | 94488 Autophagy-related protein 9A OS=Rattus norvegicus OX=10116 GN=Atg9a PE=1 SV=1 |
| 55 | 761 | tr\|A0A0H2 | 35.17 | 2 | 2 | 1.29E+07 | 4 | 3 | 4 | Formylatio | 136526 Graves disease carrier protein OS=Rattus norvegicus OX=10116 GN=Slc25a16 PE=1 SV=1 |
| 55 | 760 | D3ZG52\|D | 35.17 | 3 | 3 | 1.29E+07 | 4 | 3 | 4 | Formylatio | 119588 DNA replication ATP-dependent helicase/nuclease DNA2 OS=Rattus norvegicus OX=10116 GN=Dna2 PE=3 SV=1 |
| 103 | 5935 | tr\|B2GUW | 34.74 | 2 | 2 | 6.15E+05 | 2 | 1 | 2 | | 74060 Exdl2 protein OS=Rattus norvegicus OX=10116 GN=Exd2 PE=2 SV=1 |
| 42 | 462 | tr\|D3ZKC6 | 34.69 | 0 | 0 | 5.09E+06 | 2 | 2 | 5 | | 488962 Vacuolar protein sorting 13 homolog D OS=Rattus norvegicus OX=10116 GN=Vps13d PE=1 SV=1 |
| 42 | 461 | tr\|A0A0G2 | 34.69 | 0 | 0 | 5.09E+06 | 2 | 2 | 5 | | 487857 Vacuolar protein sorting 13 homolog D OS=Rattus norvegicus OX=10116 GN=Vps13d PE=1 SV=1 |
| 241 | 338 | Q5M9I5\|Q | 34.58 | 17 | 17 | 0 | 1 | 1 | 1 | | 10424 Cytochrome b-c1 complex subunit 6 mitochondrial OS=Rattus norvegicus OX=10116 GN=Uqcrh PE=3 SV=1 |
| 68 | 5813 | tr\|F1LNF0 | 33.81 | 1 | 1 | 3.74E+05 | 3 | 3 | 3 | | 228912 Myosin heavy chain 14 OS=Rattus norvegicus OX=10116 GN=Myh14 PE=1 SV=1 |
| 82 | 702 | tr\|D3ZT64 | 33.62 | 1 | 1 | 3.34E+06 | 2 | 1 | 3 | | 211362 Autophagy-related 2A OS=Rattus norvegicus OX=10116 GN=Atg2a PE=1 SV=1 |
| 73 | 1013 | tr\|D3ZJH9 | 33.27 | 1 | 1 | 0 | 1 | 1 | 3 | | 65352 Malic enzyme OS=Rattus norvegicus OX=10116 GN=Me2 PE=1 SV=1 |
| 73 | 1014 | tr\|A0A0G2 | 33.27 | 1 | 1 | 0 | 1 | 1 | 3 | | 66310 Malic enzyme OS=Rattus norvegicus OX=10116 GN=Me2 PE=1 SV=1 |
| 73 | 11929 | tr\|A0A0G2 | 33.27 | 1 | 1 | 0 | 1 | 1 | 3 | | 63436 Malic enzyme OS=Rattus norvegicus OX=10116 GN=Me3 PE=3 SV=1 |
| 73 | 6137 | P13697\|M | 33.27 | 1 | 1 | 0 | 1 | 1 | 3 | | 64003 NADP-dependent malic enzyme OS=Rattus norvegicus OX=10116 GN=Me1 PE=1 SV=2 |
| 73 | 11934 | tr\|F1M5N4 | 33.27 | 1 | 1 | 0 | 1 | 1 | 3 | | 67209 Malic enzyme OS=Rattus norvegicus OX=10116 GN=Me3 PE=3 SV=2 |
| 56 | 1230 | tr\|D3ZIE9\| | 32.86 | 1 | 1 | | 2 | 0 | 4 | | 87329 Delta-1-pyrroline-5-carboxylate synthase OS=Rattus norvegicus OX=10116 GN=Aldh18a1 PE=3 SV=1 |
| 85 | 12007 | B0BN56\|R | 32.47 | 3 | 3 | | 2 | 0 | 3 | | 43962 28S ribosomal protein S31 mitochondrial OS=Rattus norvegicus OX=10116 GN=Mrps31 PE=2 SV=1 |
| 110 | 59 | P17178\|CI | 32.33 | 5 | 5 | 2.76E+05 | 2 | 2 | 2 | | 60733 Sterol 26-hydroxylase mitochondrial OS=Rattus norvegicus OX=10116 GN=Cyp27a1 PE=1 SV=1 |
| 110 | 60 | tr\|A0A0H2 | 32.33 | 5 | 5 | 2.76E+05 | 2 | 2 | 2 | | 62492 RCG24013 isoform CRA_a OS=Rattus norvegicus OX=10116 GN=Cyp27a1 PE=1 SV=1 |
| 119 | 28662 | tr\|M0R601 | 31.66 | 26 | 26 | 7.28E+05 | 2 | 2 | 2 | | 5582 Uncharacterized protein OS=Rattus norvegicus OX=10116 PE=4 SV=2 |
| 119 | 34379 | P29418\|AT | 31.66 | 25 | 25 | 7.28E+05 | 2 | 2 | 2 | | 5767 ATP synthase subunit epsilon mitochondrial OS=Rattus norvegicus OX=10116 GN=Atp5f1e PE=1 SV=2 |
| 127 | 656 | tr\|D3ZD23 | 31.47 | 1 | 1 | 7.74E+05 | 2 | 2 | 2 | | 67300 ATP-binding cassette subfamily E member 1 OS=Rattus norvegicus OX=10116 GN=Abce1 PE=1 SV=1 |

| | | | | | | | | | | | |
|---|---|---|---|---|---|---|---|---|---|---|---|
| 204 | 146 | tr\|B0BMT9 | 31.03 | 2 | 2 | 0 | 1 | 1 | 1 | | 50202 Sqrdl protein OS=Rattus norvegicus OX=10116 GN=Sqor PE=1 SV=1 |
| 91 | 626 | tr\|D4A054 | 30.39 | 1 | 1 | 2.47E+06 | 2 | 2 | 2 | Formylatio | 341403 RAN-binding protein 2 OS=Rattus norvegicus OX=10116 GN=Ranbp2 PE=1 SV=2 |
| 91 | 627 | tr\|M0R3M | 30.39 | 1 | 1 | 2.47E+06 | 2 | 2 | 2 | Formylatio | 344396 RAN-binding protein 2 OS=Rattus norvegicus OX=10116 GN=Ranbp2 PE=1 SV=1 |
| 92 | 650 | D4A929\|W | 28.68 | 1 | 1 | 6.52E+05 | 2 | 1 | 2 | | 212258 WD repeat-containing protein 81 OS=Rattus norvegicus OX=10116 GN=Wdr81 PE=3 SV=1 |
| 160 | 341 | tr\|G3V6I5\| | 28.64 | 3 | 3 | 2.11E+05 | 1 | 1 | 1 | | 46777 DnaJ heat shock protein family (Hsp40) member A3 OS=Rattus norvegicus OX=10116 GN=Dnaja3 PE=1 SV=2 |
| 160 | 342 | tr\|Q2TVU3 | 28.64 | 3 | 3 | 2.11E+05 | 1 | 1 | 1 | | 49412 TID1 OS=Rattus norvegicus OX=10116 GN=Dnaja3 PE=2 SV=1 |
| 160 | 343 | tr\|A0A0G2 | 28.64 | 3 | 3 | 2.11E+05 | 1 | 1 | 1 | | 49416 DnaJ heat shock protein family (Hsp40) member A3 OS=Rattus norvegicus OX=10116 GN=Dnaja3 PE=1 SV=1 |
| 160 | 344 | tr\|Q2UZS7 | 28.64 | 2 | 2 | 2.11E+05 | 1 | 1 | 1 | | 52399 Tid-1 long isoform OS=Rattus norvegicus OX=10116 GN=Dnaja3 PE=2 SV=1 |
| 160 | 345 | tr\|A0A0G2 | 28.64 | 2 | 2 | 2.11E+05 | 1 | 1 | 1 | | 52403 DnaJ heat shock protein family (Hsp40) member A3 OS=Rattus norvegicus OX=10116 GN=Dnaja3 PE=1 SV=1 |
| 80 | 5883 | Q8VIJ5\|OG | 28.21 | 1 | 1 | 7.52E+05 | 1 | 1 | 3 | | 102918 Protein O-GlcNAcase OS=Rattus norvegicus OX=10116 GN=Oga PE=1 SV=1 |
| 118 | 1487 | P48768\|NA | 28.2 | 1 | 1 | 2.03E+06 | 2 | 1 | 2 | Formylatio | 100523 Sodium/calcium exchanger 2 OS=Rattus norvegicus OX=10116 GN=Slc8a2 PE=1 SV=1 |
| 176 | 281 | tr\|F1LPV8\| | 27.77 | 3 | 3 | 2.85E+05 | 1 | 1 | 1 | | 46639 Succinate--CoA ligase [GDP-forming] subunit beta mitochondrial OS=Rattus norvegicus OX=10116 GN=Suclg2 PE=1 SV=2 |
| 176 | 282 | tr\|B1H270 | 27.77 | 3 | 3 | 2.85E+05 | 1 | 1 | 1 | | 46988 Succinate--CoA ligase [GDP-forming] subunit beta mitochondrial OS=Rattus norvegicus OX=10116 GN=Suclg2 PE=2 SV=1 |
| 242 | 255 | tr\|I6V4L9\| | 27.5 | 3 | 3 | 5.54E+04 | 1 | 1 | 1 | | 29844 Cytochrome c oxidase subunit 3 OS=Rattus norvegicus OX=10116 GN=COX3 PE=3 SV=1 |
| 201 | 678 | Q63147\|H | 27.39 | 2 | 2 | 2.56E+05 | 1 | 1 | 1 | | 64842 5-aminolevulinate synthase erythroid-specific mitochondrial OS=Rattus norvegicus OX=10116 GN=Alas2 PE=1 SV=1 |
| 201 | 34380 | tr\|Q9ESE3 | 27.39 | 5 | 5 | 2.56E+05 | 1 | 1 | 1 | | 18186 5-aminolevulinate synthase (Fragment) OS=Rattus norvegicus OX=10116 GN=ALS2 PE=2 SV=1 |
| 122 | 1374 | tr\|A0A0G2 | 26.95 | 3 | 3 | 2.71E+05 | 1 | 1 | 2 | | 53760 von Willebrand factor A domain-containing 8 OS=Rattus norvegicus OX=10116 GN=Vwa8 PE=1 SV=1 |
| 87 | 1081 | tr\|Q5XHZ7 | 26.86 | 2 | 2 | 0 | 2 | 1 | 3 | | 66660 MMR_HSR1 domain containing protein RGD1359460 OS=Rattus norvegicus OX=10116 GN=Noa1 PE=2 SV=1 |
| 87 | 1082 | tr\|A0A0G2 | 26.86 | 2 | 2 | 0 | 2 | 1 | 3 | | 66708 Nitric oxide-associated 1 OS=Rattus norvegicus OX=10116 GN=Noa1 PE=4 SV=1 |
| 87 | 1089 | tr\|A0A0G2 | 26.86 | 2 | 2 | 0 | 2 | 1 | 3 | | 77504 MMR_HSR1 domain containing protein RGD1359460 isoform CRA_a OS=Rattus norvegicus OX=10116 GN=Noa1 PE=4 SV=1 |
| 177 | 5924 | P07150\|AN | 26.42 | 3 | 3 | 1.19E+05 | 1 | 1 | 1 | | 38829 Annexin A1 OS=Rattus norvegicus OX=10116 GN=Anxa1 PE=1 SV=2 |
| 77 | 5791 | tr\|F1LSY7\| | 25.86 | 1 | 1 | 1.76E+07 | 2 | 2 | 3 | | 109031 Endoplasmic reticulum to nucleus-signaling 1 OS=Rattus norvegicus OX=10116 GN=Ern1 PE=4 SV=2 |
| 77 | 5792 | tr\|A0A0G2 | 25.86 | 1 | 1 | 1.76E+07 | 2 | 2 | 3 | | 110150 Endoplasmic reticulum to nucleus-signaling 1 OS=Rattus norvegicus OX=10116 GN=Ern1 PE=2 SV=1 |
| 178 | 5893 | tr\|F1LP46\| | 25.79 | 1 | 1 | 2.96E+05 | 1 | 1 | 1 | | 78923 ATP-dependent RNA helicase SUPV3L1 mitochondrial OS=Rattus norvegicus OX=10116 GN=Supv3l1 PE=1 SV=2 |
| 178 | 5894 | tr\|A0A0G2 | 25.79 | 1 | 1 | 2.96E+05 | 1 | 1 | 1 | | 84180 ATP-dependent RNA helicase SUPV3L1 mitochondrial OS=Rattus norvegicus OX=10116 GN=Supv3l1 PE=1 SV=1 |
| 178 | 5895 | Q5EBA1\|SU | 25.79 | 1 | 1 | 2.96E+05 | 1 | 1 | 1 | | 86706 ATP-dependent RNA helicase SUPV3L1 mitochondrial OS=Rattus norvegicus OX=10116 GN=Supv3l1 PE=2 SV=1 |
| 124 | 5796 | B1WC61\|A | 25.51 | 2 | 2 | 1.46E+05 | 1 | 1 | 2 | | 68843 Complex I assembly factor ACAD9 mitochondrial OS=Rattus norvegicus OX=10116 GN=Acad9 PE=1 SV=1 |
| 69 | 555 | tr\|D4A1D3 | 24.54 | 0 | 0 | 0 | 2 | 1 | 3 | Carbamido | 521487 Sacsin molecular chaperone OS=Rattus norvegicus OX=10116 GN=Sacs PE=1 SV=2 |
| 74 | 485 | Q2PQA9\|K | 24.48 | 2 | 2 | 0 | 2 | 1 | 3 | Formylatio | 109531 Kinesin-1 heavy chain OS=Rattus norvegicus OX=10116 GN=Kif5b PE=1 SV=1 |
| 158 | 253 | Q9ER34\|AC | 24.46 | 1 | 1 | 2.68E+05 | 1 | 1 | 1 | | 85433 Aconitate hydratase mitochondrial OS=Rattus norvegicus OX=10116 GN=Aco2 PE=1 SV=2 |
| 126 | 34383 | Q32Q86\|C | 23.56 | 1 | 1 | 3.28E+05 | 1 | 1 | 2 | | 66231 Cryptochrome-1 OS=Rattus norvegicus OX=10116 GN=Cry1 PE=1 SV=1 |
| 81 | 356 | O70600\|RS | 23.54 | 5 | 5 | 7.27E+06 | 2 | 2 | 3 | | 41255 Radical S-adenosyl methionine domain-containing protein 2 OS=Rattus norvegicus OX=10116 GN=Rsad2 PE=1 SV=1 |
| 81 | 357 | tr\|A0A0H2 | 23.54 | 5 | 5 | 7.27E+06 | 2 | 2 | 3 | | 41444 RCG62278 OS=Rattus norvegicus OX=10116 GN=Rsad2 PE=4 SV=1 |
| 100 | 683 | O88658\|KI | 23.34 | 0 | 0 | 9.88E+05 | 1 | 1 | 2 | | 204169 Kinesin-like protein KIF1B OS=Rattus norvegicus OX=10116 GN=Kif1b PE=1 SV=2 |
| 64 | 489 | tr\|D3ZL86\| | 23.09 | 1 | 1 | 1.84E+06 | 1 | 1 | 3 | | 241219 HEAT repeat-containing protein 1 OS=Rattus norvegicus OX=10116 GN=Heatr1 PE=1 SV=2 |
| 205 | 34382 | Q66HG6\|C | 22.69 | 2 | 2 | 1.93E+05 | 1 | 1 | 1 | | 36598 Carbonic anhydrase 5B mitochondrial OS=Rattus norvegicus OX=10116 GN=Ca5b PE=2 SV=1 |
| 179 | 622 | tr\|A0A0G2 | 22.46 | 1 | 1 | 2.38E+05 | 1 | 1 | 1 | | 90015 Centrosomal protein 89 OS=Rattus norvegicus OX=10116 GN=Cep89 PE=4 SV=1 |
| 179 | 621 | tr\|B1WBZ3 | 22.46 | 1 | 1 | 2.38E+05 | 1 | 1 | 1 | | 72885 Ccdc123 protein OS=Rattus norvegicus OX=10116 GN=Cep89 PE=2 SV=1 |
| 247 | 267 | tr\|Q06Q97 | 22.43 | 2 | 2 | 0 | 1 | 1 | 1 | | 38485 NADH-ubiquinone oxidoreductase chain 2 OS=Rattus norvegicus OX=10116 GN=ND2 PE=3 SV=1 |
| 247 | 268 | tr\|A0A0A1 | 22.43 | 2 | 2 | 0 | 1 | 1 | 1 | | 38542 NADH-ubiquinone oxidoreductase chain 2 OS=Rattus norvegicus OX=10116 GN=ND2 PE=3 SV=1 |
| 247 | 269 | tr\|Q5UAJ8 | 22.43 | 2 | 2 | 0 | 1 | 1 | 1 | | 38455 NADH-ubiquinone oxidoreductase chain 2 OS=Rattus norvegicus OX=10116 GN=ND2 PE=3 SV=1 |
| 247 | 270 | tr\|Q8HID0 | 22.43 | 2 | 2 | 0 | 1 | 1 | 1 | | 38653 NADH-ubiquinone oxidoreductase chain 2 OS=Rattus norvegicus OX=10116 GN=Mt-nd2 PE=3 SV=1 |
| 247 | 271 | P11662\|NU | 22.43 | 2 | 2 | 0 | 1 | 1 | 1 | | 38653 NADH-ubiquinone oxidoreductase chain 2 OS=Rattus norvegicus OX=10116 GN=Mtnd2 PE=3 SV=3 |
| 247 | 274 | tr\|Q06QB0 | 22.43 | 2 | 2 | 0 | 1 | 1 | 1 | | 38534 NADH-ubiquinone oxidoreductase chain 2 OS=Rattus norvegicus OX=10116 GN=ND2 PE=3 SV=1 |
| 247 | 416 | tr\|S5RF74\| | 22.43 | 2 | 2 | 0 | 1 | 1 | 1 | | 38641 NADH-ubiquinone oxidoreductase chain 2 OS=Rattus norvegicus OX=10116 GN=ND2 PE=3 SV=1 |
| 247 | 275 | tr\|D2E6P4 | 22.43 | 2 | 2 | 0 | 1 | 1 | 1 | | 38623 NADH-ubiquinone oxidoreductase chain 2 OS=Rattus norvegicus OX=10116 GN=ND2 PE=3 SV=1 |
| 145 | 6379 | O08561\|PI | 22.15 | 1 | 1 | 0 | 1 | 1 | 1 | Formylatio | 91655 Phosphatidylinositol 4-kinase beta OS=Rattus norvegicus OX=10116 GN=Pi4kb PE=1 SV=1 |
| 249 | 355 | tr\|D4AE90 | 21.9 | 2 | 2 | 0 | 1 | 1 | 1 | Formylatio | 50113 RCC1-like OS=Rattus norvegicus OX=10116 GN=Rcc1l PE=1 SV=1 |
| 257 | 1475 | P04177\|TY | 21.53 | 1 | 1 | 1.56E+06 | 1 | 1 | 1 | | 55966 Tyrosine 3-monooxygenase OS=Rattus norvegicus OX=10116 GN=Th PE=1 SV=3 |
| 253 | 382 | tr\|D3ZU73 | 21.16 | 3 | 3 | 9.96E+05 | 1 | 1 | 1 | | 28584 Nipsnap homolog 3A (C. elegans) OS=Rattus norvegicus OX=10116 GN=Nipsnap3a PE=4 SV=2 |
| 255 | 6385 | O88767\|PA | 20.89 | 5 | 5 | 0 | 1 | 1 | 1 | | 19974 Protein/nucleic acid deglycase DJ-1 OS=Rattus norvegicus OX=10116 GN=Park7 PE=1 SV=1 |
| 255 | 6386 | tr\|Q5BKC3 | 20.89 | 5 | 5 | 0 | 1 | 1 | 1 | | 22503 Park7 protein OS=Rattus norvegicus OX=10116 GN=Park7 PE=1 SV=1 |
| 256 | 66 | tr\|D4A0T0 | 20.77 | 10 | 10 | 4.34E+05 | 1 | 1 | 1 | | 20859 NADH:ubiquinone oxidoreductase subunit B10 OS=Rattus norvegicus OX=10116 GN=Ndufb10 PE=1 SV=1 |
| 261 | 1953 | D4A5C3\|H | 20.62 | 2 | 2 | 4.16E+05 | 1 | 1 | 1 | | 36549 3-hydroxy-3-methylglutaryl-CoA lyase cytoplasmic OS=Rattus norvegicus OX=10116 GN=Hmgcll1 PE=1 SV=1 |
| 262 | 1059 | tr\|A0A0G2 | 20.27 | 2 | 2 | 3.68E+05 | 1 | 1 | 1 | | 46686 39S ribosomal protein L37 mitochondrial OS=Rattus norvegicus OX=10116 GN=Mrpl37 PE=1 SV=1 |
| 262 | 1060 | Q6AXT0\|R | 20.27 | 2 | 2 | 3.68E+05 | 1 | 1 | 1 | | 48366 39S ribosomal protein L37 mitochondrial OS=Rattus norvegicus OX=10116 GN=Mrpl37 PE=2 SV=1 |
