## Supplementary material for "Permeability transition pore-related changes in the proteome and channel activity of ATP synthase dimers and monomers": ATP-synthase

| Pro tein Gro up | Protei n ID | Score (%) | -10lgP | Cover age (%) | #Pe ptid es | #Uni que | Area (IBAQ) | riBAQ | PTM | Avg. Mass | Accession | riBAQ PTP/Control | Pro tein Gro up | Protein ID | Score (%) | -10lgP | Coverage (%) | #Pepti des | #Uniq ue | Area (IBAQ) | riBAQ | PTM | Avg. Mass | Descriptio n |
|---|---|---|---|---|---|---|---|---|---|---|---|---|---|---|---|---|---|---|---|---|---|---|---|---|
| | | | | | | | CONTROL | % | | | | | | | | | | | | PTP | % | | | |
| **Complex III** | | | | | | | | | | | | | | | | | | | | | | | | |
| 17 | 12 | 99.2 | 239.36 | 43 | 14 | 14 | 22165000 | 0.21475999 | Y | 52849 | Q68FY0\|QCR1_RAT | 0.451805576 | 31 | 15 | 99.1 | 232.07 | 23 | 8 | 8 | 15442000.00 | 0.09702976 | Y | 52849 | Cytochrome b-c1 complex subunit 1, mitochondrial OS=Rattus norvegicus OX=10116 GN=Uqcrc1 PE=1 SV=1 |
| 19 | 1 | 99.2 | 237.64 | 45 | 15 | 15 | 35141000 | 0.34048639 | N | 48396 | P32551\|QCR2_RAT | 0.431170317 | 26 | 1 | 99.2 | 224.98 | 25 | 9 | 9 | 23364000.00 | 0.14680762 | N | 48396 | Cytochrome b-c1 complex subunit 2, mitochondrial OS=Rattus norvegicus OX=10116 GN=Uqcrc2 PE=1 SV=2 |
| 34 | 28 | 98.9 | 142.12 | 14 | 5 | 5 | 6386600 | 0.06188072 | N | 35435 | tr\|D3ZFQ8\|D3ZFQ8_RA | 0.860985454 | 45 | 42 | 98.8 | 145.33 | 14 | 4 | 4 | 8479100.00 | 0.0532784 | N | 35435 | Cytochrome c-1 OS=Rattus norvegicus OX=10116 GN=Cyc1 PE=1 SV=3 |
| 71 | 37 | 84.4 | 115.38 | 18 | 2 | 2 | 1933300 | 0.01873203 | N | 29446 | P20788\|UCRI_RAT | 1.893264199 | 52 | 45 | 98.4 | 108.7 | 17 | 4 | 4 | 5644100.00 | 0.03546469 | N | 29446 | Cytochrome b-c1 complex subunit Rieske, mitochondrial OS=Rattus norvegicus OX=10116 GN=Uqcrfs1 PE=1 SV=2 |
| 73 | 93 | 84.4 | 110.17 | 39 | 2 | 2 | 1652700 | 0.01601326 | N | 7099 | tr\|A0A0G2K8Q8\|A0A0( | - | | | | | | | | | 0 | | | Ubiquinol-cytochrome c reductase, complex III subunit X OS=Rattus norvegicus OX=10116 GN=Uqcr10 PE=1 SV=1 |
| 77 | 2167 | 61.7 | 90.49 | 21 | 1 | 1 | 952880 | 0.0092326 | N | 10424 | Q5M9I5\|QCR6_RAT | - | | | | | | | | | 0 | | | Cytochrome b-c1 complex subunit 6, mitochondrial OS=Rattus norvegicus OX=10116 GN=Uqcrh PE=3 SV=1 |
| 72 | 30 | 84.4 | 75.25 | 23 | 2 | 2 | 1354300 | 0.01312201 | N | 9849 | Q7TQ16\|QCR8_RAT | - | | | | | | | | | 0 | | | Cytochrome b-c1 complex subunit 8 OS=Rattus norvegicus OX=10116 GN=Uqcrq PE=3 SV=1 |
| **Complex IV** | | | | | | | | | | | | | | | | | | | | | | | | |
| 37 | 51 | 99 | 141.37 | 30 | 5 | 5 | 6270700 | 0.06075775 | N | 19515 | P10888\|COX41_RAT | 0.817867377 | 59 | 50 | 98 | 74.54 | 17 | 3 | 3 | 7908300.00 | 0.04969178 | N | 19515 | Cytochrome c oxidase subunit 4 isoform 1, mitochondrial OS=Rattus norvegicus OX=10116 GN=Cox4i1 PE=1 SV=1 |
| 60 | 109 | 84.3 | 107.55 | 30 | 3 | 2 | 8579600 | 0.08312902 | N | 16130 | P11240\|COX5A_RAT | 0.10294232 | 96 | 158 | 61.7 | 96.56 | 10 | 1 | 1 | 1361900.00 | 0.00855749 | N | 16130 | Cytochrome c oxidase subunit 5A, mitochondrial OS=Rattus norvegicus OX=10116 GN=Cox5a PE=1 SV=1 |
| 64 | 11616 | 84.4 | 93.76 | 27 | 2 | 2 | 1546100 | 0.01498039 | N | 12301 | P10818\|CX6A1_RAT | - | | | | | | | | | 0 | | | Cytochrome c oxidase subunit 6A1, mitochondrial OS=Rattus norvegicus OX=10116 GN=Cox6a1 PE=1 SV=2 |
| 58 | 158 | 84.4 | 88.52 | 29 | 3 | 3 | 6944000 | 0.06728145 | N | 13915 | P12075\|COX5B_RAT | 0.595770627 | 66 | 133 | 91.6 | 112.28 | 30 | 3 | 3 | 6379300.00 | 0.04008431 | N | 13915 | Cytochrome c oxidase subunit 5B, mitochondrial OS=Rattus norvegicus OX=10116 GN=Cox5b PE=1 SV=2 |
| 44 | 57 | 98.4 | 76.74 | 11 | 3 | 3 | 4079900 | 0.03953076 | Y | 25928 | P00406\|COX2_RAT | 0.823896392 | 57 | 81 | 98.4 | 74.72 | 11 | 3 | 3 | 5183300.00 | 0.03256925 | Y | 25928 | Cytochrome c oxidase subunit 2 OS=Rattus norvegicus OX=10116 GN=Mtco2 PE=1 SV=3 |
| 74 | 11618 | 84.2 | 75.46 | 21 | 2 | 2 | 1308400 | 0.01267728 | N | 8455 | P11951\|CX6C2_RAT | - | | | | | | | | | 0 | | | Cytochrome c oxidase subunit 6C-2 OS=Rattus norvegicus OX=10116 GN=Cox6c2 PE=1 SV=3 |
| 132 | 11617 | 61.7 | 65.38 | 14 | 1 | 1 | 252730 | 0.00244874 | N | 7375 | P80432\|COX7C_RAT | - | | | | | | | | | 0 | | | Cytochrome c oxidase subunit 7C, mitochondrial OS=Rattus norvegicus OX=10116 GN=Cox7c PE=1 SV=2 |
| 134 | 13903 | 61.6 | 53.43 | 10 | 1 | 1 | 2595300 | 0.02514625 | N | 9353 | P35171\|CX7A2_RAT | - | | | | | | | | | 0 | | | Cytochrome c oxidase subunit 7A2, mitochondrial OS=Rattus norvegicus OX=10116 GN=Cox7a2 PE=1 SV=1 |
| 136 | 11623 | 61.2 | 39.08 | 10 | 1 | 1 | 543290 | 0.00526402 | N | 9327 | tr\|B2RZD6\|B2RZD6_RA | 1.94054835 | 213 | 191 | 61.5 | 45.95 | 10 | 1 | 1 | 1625700.00 | 0.01021508 | N | 9327 | NDUFA4, mitochondrial complex-associated OS=Rattus norvegicus OX=10116 GN=Ndufa4 PE=1 SV=1 |
| 49 | 13895 | 92.5 | 102.09 | 6 | 2 | 2 | 867700 | 0.00840727 | N | 68843 | B1WC61\|ACAD9_RAT | - | | | | | | | | | 0 | | | Complex I assembly factor ACAD9, mitochondrial OS=Rattus norvegicus OX=10116 GN=Acad9 PE=1 SV=1 |
| 126 | 11614 | 61.7 | 82.94 | 5 | 1 | 1 | 911840 | 0.00883495 | N | 29871 | P05505\|COX3_RAT | 0.291887212 | 203 | 258 | 61.7 | 85.02 | 5 | 1 | 1 | 410410.00 | 0.00257881 | N | 29871 | Cytochrome c oxidase subunit 3 OS=Rattus norvegicus OX=10116 GN=Mtco3 PE=1 SV=5 |
| 135 | 11619 | 61.4 | 47.35 | 9 | 1 | 1 | 447640 | 0.00433725 | N | 8995 | P80431\|COX7B_RAT | 0.420754105 | 218 | 4992 | 60.6 | 37.86 | 9 | 1 | 1 | 290430.00 | 0.00182492 | N | 8995 | Cytochrome c oxidase subunit 7B, mitochondrial OS=Rattus norvegicus OX=10116 GN=Cox7b PE=1 SV=3 |
| **125** | 13900 | | 84.02 | 9 | 1 | 1 | 933500 | 0.00904482 | N | 17283 | P19804\|NDKB_RAT | 4.525596951 | 63 | 282 | 98.4 | 95.49 | 20 | 3 | 3 | 6514400.00 | 0.04093321 | N | 17283 | Nucleoside diphosphate kinase B OS=Rattus norvegicus OX=10116 GN=Nme2 PE=1 SV=1 |
| **Complex V** | | | | | | | | | | | | | | | | | | | | | | | | |
| 2 | 14 | 99.2 | 432.09 | 77 | 104 | 104 | 2723700000 | 26.3903354 | Y | 59754 | P15999\|ATPA_RAT | 1.331470142 | 2 | 13 | 99.2 | 488.87 | 85 | 104 | 103 | 5592100000.00 | 35.1379436 | Y | 59754 | ATP synthase subunit alpha, mitochondrial OS=Rattus norvegicus OX=10116 GN=Atp5f1a PE=1 SV=2 |
| 1 | 7 | 99.2 | 408.87 | 82 | 84 | 83 | 3460600000 | 33.5302693 | Y | 56354 | P10719\|ATPB_RAT | 0.83586892 | 1 | 8 | 99.2 | 479.96 | 84 | 75 | 74 | 4460400000.00 | 28.02691 | Y | 56354 | ATP synthase subunit beta, mitochondrial OS=Rattus norvegicus OX=10116 GN=Atp5f1b PE=1 SV=2 |
| 5 | 4260 | 99.2 | 358.66 | 93 | 40 | 40 | 461400000 | 4.47057339 | Y | 18763 | P31399\|ATP5H_RAT | 0.610109742 | 7 | 204 | 99.1 | 372.39 | 76 | 26 | 26 | 434080000.00 | 2.72754038 | Y | 18763 | ATP synthase subunit d, mitochondrial OS=Rattus norvegicus OX=10116 GN=Atp5pd PE=1 SV=3 |
| 8 | 116 | 99.2 | 283.13 | 68 | 29 | 29 | 222780000 | 2.15854863 | Y | 30191 | P35435\|ATPG_RAT | 1.793165135 | 5 | 97 | 99.2 | 297.27 | 71 | 24 | 24 | 616000000.00 | 3.87063415 | Y | 30191 | ATP synthase subunit gamma, mitochondrial OS=Rattus norvegicus OX=10116 GN=Atp5f1c PE=1 SV=2 |
| 4 | 41 | 99.2 | 277.66 | 71 | 31 | 30 | 460220000 | 4.45914019 | Y | 28869 | P19511\|AT5F1_RAT | 1.216302479 | 6 | 114 | 99.2 | 337.53 | 63 | 28 | 28 | 863160000.00 | 5.42366327 | Y | 28869 | ATP synthase F(0) complex subunit B1, mitochondrial OS=Rattus norvegicus OX=10116 GN=Atp5pb PE=1 SV=1 |
| 7 | 4261 | 99.1 | 256.07 | 82 | 20 | 20 | 658280000 | 6.37817306 | Y | 17563 | tr\|G3V7Y3\|G3V7Y3_RA | 0.181249045 | 20 | 168 | 99.1 | 243.79 | 61 | 10 | 10 | 183980000.00 | 1.15603778 | Y | 17563 | ATP synthase subunit delta, mitochondrial OS=Rattus norvegicus OX=10116 GN=Atp5f1d PE=1 SV=1 |
| 11 | 2163 | 99.2 | 255.64 | 76 | 27 | 27 | 507150000 | 4.91385196 | Y | 23398 | Q06647\|ATPO_RAT | 1.082637975 | 10 | 188 | 99.2 | 255.92 | 68 | 24 | 24 | 846650000.00 | 5.31992273 | Y | 23398 | ATP synthase subunit O, mitochondrial OS=Rattus norvegicus OX=10116 GN=Atp5po PE=1 SV=1 |
| 26 | 4263 | 99.1 | 197.76 | 89 | 7 | 7 | 83153000 | 0.80568181 | Y | 11433 | Q6PDU7\|ATP5L_RAT | 0.237510161 | 34 | 4490 | 99.1 | 193.65 | 66 | 7 | 7 | 30454000.00 | 0.19135762 | Y | 11433 | ATP synthase subunit g, mitochondrial OS=Rattus norvegicus OX=10116 GN=Atp5mg PE=1 SV=2 |
| 36 | 13889 | 98.3 | 169.58 | 37 | 5 | 5 | 11661000 | 0.11298517 | N | 12494 | P21571\|ATP5J_RAT | 0.948098403 | 38 | 4496 | 98.8 | 174.24 | 44 | 6 | 6 | 17048000.00 | 0.10712106 | N | 12494 | ATP synthase-coupling factor 6, mitochondrial OS=Rattus norvegicus OX=10116 GN=Atp5pf PE=1 SV=1 |
| 31 | 13896 | 84.3 | 159.56 | 37 | 5 | 5 | 12692000 | 0.12297468 | Y | 6914 | D3Z9R8\|ATP68_RAT | 0.42317591 | 98 | 4494 | 35.5 | 24.24 | 12 | 1 | 1 | 8282000.00 | 0.05203992 | Y | 6914 | ATP synthase subunit ATP5MPL, mitochondrial OS=Rattus norvegicus OX=10116 GN=Atp5mpl PE=1 SV=1 |
| 42 | 4264 | 98.4 | 140.51 | 55 | 4 | 4 | 96459000 | 0.93460563 | Y | 7642 | tr\|Q5UAJ5\|Q5UAJ5_RA | 0.921072011 | 30 | 4487 | 98.4 | 168.07 | 55 | 4 | 4 | 137000000.00 | 0.86083909 | Y | 7642 | ATP synthase protein 8 OS=Rattus norvegicus OX=10116 GN=ATP8 PE=3 SV=1 |
| 22 | 6960 | 99.1 | 136 | 69 | 12 | 11 | 80884000 | 0.78369713 | N | 8255 | P29419\|ATP5I_RAT | 0.744585491 | 25 | 4486 | 99 | 115.83 | 55 | 7 | 7 | 92867000.00 | 0.58352952 | N | 8255 | ATP synthase subunit e, mitochondrial OS=Rattus norvegicus OX=10116 GN=Atp5me PE=1 SV=3 |
| 40 | 4270 | 84.4 | 134.56 | 45 | 3 | 3 | 56515000 | 0.54758226 | N | 6408 | Q9JJW3\|ATPMD_RAT | 0.90857766 | 27 | 4499 | 98.7 | 185.83 | 45 | 6 | 6 | 79179000.00 | 0.49752101 | N | 6408 | ATP synthase membrane subunit DAPIT, mitochondrial OS=Rattus norvegicus OX=10116 GN=Atp5md PE=1 SV=1 |
| 25 | 4266 | 99 | 128.8 | 56 | 10 | 9 | 78115000 | 0.75686788 | Y | 10452 | D3ZAF6\|ATPK_RAT | 0.28140369 | 35 | 4488 | 98.8 | 108.39 | 47 | 7 | 6 | 33896000.00 | 0.21298541 | Y | 10452 | ATP synthase subunit f, mitochondrial OS=Rattus norvegicus OX=10116 GN=Atp5mf PE=1 SV=1 |
| 63 | 2187 | 84.3 | 109.43 | 28 | 2 | 2 | 10200000 | 0.09882932 | N | 14244 | Q06645\|AT5G1_RAT | - | | | | | | | | | 0 | | | ATP synthase F(0) complex subunit C1, mitochondrial OS=Rattus norvegicus OX=10116 GN=Atp5mc1 PE=1 SV=1 |
| 38 | 4273 | 98.9 | 101.81 | 69 | 5 | 5 | 38088000 | 0.36904031 | Y | 5767 | P29418\|ATP5E_RAT | 1.210351973 | 46 | 4489 | 98.6 | 88.5 | 55 | 4 | 4 | 71086000.00 | 0.44666867 | Y | 5767 | ATP synthase subunit epsilon, mitochondrial OS=Rattus norvegicus OX=10116 GN=Atp5f1e PE=1 SV=2 |
| 56 | 13890 | 96.3 | 99.6 | 18 | 3 | 3 | 4736200 | 0.04588975 | Y | 25076 | P05504\|ATP6_RAT | 0.26136424 | 67 | 1595 | 74.6 | 70.24 | 7 | 2 | 1 | 1908800.00 | 0.01199394 | N | 25076 | ATP synthase subunit a OS=Rattus norvegicus OX=10116 GN=Mt-atp6 PE=1 SV=3 |
| **Other** | | | | | | | | | | | | | | | | | | | | | | | | |
| 3 | 53 | 99.2 | 423.33 | 69 | 89 | 87 | 485200000 | 4.70117514 | Y | 129777 | P52873\|PYC_RAT | 0.926971743 | 3 | 6 | 99.2 | 378.67 | 50 | 65 | 61 | 693540000.00 | 4.35785651 | Y | 129777 | Pyruvate carboxylase, mitochondrial OS=Rattus norvegicus OX=10116 GN=Pc PE=1 SV=2 |
| 10 | 21 | 99.2 | 348.58 | 45 | 36 | 36 | 94713000 | 0.91768838 | Y | 82665 | Q64428\|ECHA_RAT | 0.988446241 | 11 | 5 | 99.2 | 332.89 | 38 | 27 | 27 | 144360000.00 | 0.90708563 | Y | 82665 | Trifunctional enzyme subunit alpha, mitochondrial OS=Rattus norvegicus OX=10116 GN=Hadha PE=1 SV=2 |
| 6 | 13 | 99.2 | 335.39 | 32 | 38 | 37 | 99521000 | 0.96427381 | Y | 164579 | P07756\|CPSM_RAT | 2.057326104 | 4 | 3 | 99.2 | 398.05 | 45 | 63 | 62 | 315720000.00 | 1.98382567 | Y | 164579 | Carbamoyl-phosphate synthase [ammonia], mitochondrial OS=Rattus norvegicus OX=10116 GN=Cps1 PE=1 SV=1 |
| 9 | 6796 | 99.2 | 325.85 | 71 | 34 | 33 | 159850000 | 1.54881048 | Y | 59757 | P04762\|CATA_RAT | 0.920529326 | 9 | 103 | 99.2 | 324.96 | 47 | 28 | 27 | 226900000.00 | 1.42572547 | Y | 59757 | Catalase OS=Rattus norvegicus OX=10116 GN=Cat PE=1 SV=3 |
| 12 | 9 | 99.2 | 281.21 | 64 | 25 | 25 | 95293000 | 0.92330808 | Y | 47314 | P00507\|AATM_RAT | 0.341264466 | 19 | 4 | 99.2 | 213.5 | 29 | 14 | 14 | 50146000.00 | 0.31509224 | Y | 47314 | Aspartate aminotransferase, mitochondrial OS=Rattus norvegicus OX=10116 GN=Got2 PE=1 SV=2 |
| 21 | 19 | 99.2 | 262.65 | 32 | 16 | 16 | 21445000 | 0.2077838 | Y | 56912 | P22791\|HMCS2_RAT | 7.598846059 | 8 | 10 | 99.2 | 331.56 | 45 | 29 | 29 | 251280000.00 | 1.57891713 | Y | 56912 | Hydroxymethylglutaryl-CoA synthase, mitochondrial OS=Rattus norvegicus OX=10116 GN=Hmgcs2 PE=1 SV=1 |
| 15 | 85 | 99.2 | 248.58 | 25 | 20 | 20 | 10599000 | 0.10269529 | Y | 86230 | tr\|A0A0G2JVH4\|A0A0( | 0.478907638 | 40 | 22 | 99 | 138.58 | 9 | 6 | 6 | 7827100.00 | 0.04918156 | Y | 86230 | MICOS complex subunit MIC60 OS=Rattus norvegicus OX=10116 GN=Immt PE=1 SV=1 |
| 13 | 40 | 99.2 | 243.61 | 37 | 20 | 20 | 25243000 | 0.24458319 | Y | 51414 | Q60587\|ECHB_RAT | 2.501214345 | 12 | 31 | 99.2 | 255.39 | 41 | 22 | 21 | 97359000.00 | 0.61175498 | Y | 51414 | Trifunctional enzyme subunit beta, mitochondrial OS=Rattus norvegicus OX=10116 GN=Hadhb PE=1 SV=1 |
| 16 | 2302 | 99.2 | 241.35 | 27 | 15 | 15 | 18504000 | 0.17928801 | Y | 75316 | P16970\|ABCD3_RAT | 1.257168926 | 13 | 44 | 99.2 | 268.43 | 29 | 17 | 16 | 35871000.00 | 0.22539532 | N | 75316 | ATP-binding cassette sub-family D member 3 OS=Rattus norvegicus OX=10116 GN=Abcd3 PE=1 SV=3 |
| 14 | 2 | 99.2 | 240.93 | 47 | 18 | 18 | 38664000 | 0.37462126 | Y | 57808 | Q02253\|MMSA_RAT | 0.347753203 | 22 | 64 | 99.2 | 215.93 | 22 | 11 | 11 | 20733000.00 | 0.13027574 | Y | 57808 | Methylmalonate-semialdehyde dehydrogenase [acylating], mitochondrial OS=Rattus norvegicus OX=10116 GN=Aldh6a1 PE=1 SV=1 |
| 20 | 4373 | 99.2 | 233.98 | 34 | 13 | 13 | 47000000 | 0.45539001 | Y | 61869 | P0C2X9\|AL4A1_RAT | 0.355217164 | 24 | 47 | 99.2 | 205.82 | 19 | 9 | 9 | 25744000.00 | 0.16176235 | N | 61869 | Delta-1-pyrroline-5-carboxylate dehydrogenase, mitochondrial OS=Rattus norvegicus OX=10116 GN=Aldh4a1 PE=1 SV=1 |
| 29 | 542 | 99.1 | 224.76 | 20 | 8 | 8 | 6627100 | 0.06421096 | N | 61416 | P10860\|DHE3_RAT | 5.827291219 | 18 | 2 | 99.2 | 245.57 | 26 | 13 | 13 | 59549000.00 | 0.37417596 | Y | 61416 | Glutamate dehydrogenase 1, mitochondrial OS=Rattus norvegicus OX=10116 GN=Glud1 PE=1 SV=2 |
| 18 | 9289 | 99.2 | 224.6 | 40 | 14 | 14 | 56148000 | 0.54402634 | Y | 33407 | P24329\|THTR_RAT | 0.654907343 | 17 | 55 | 99.2 | 219.5 | 39 | 15 | 15 | 56702000.00 | 0.35628685 | Y | 33407 | Thiosulfate sulfurtransferase OS=Rattus norvegicus OX=10116 GN=Tst PE=1 SV=3 |
| 24 | 229 | 99.2 | 217.43 | 15 | 13 | 1 | 780490 | 0.00756228 | N | 102908 | A2VCW9\|AASS_RAT | 34.58784815 | 14 | 1580 | 99.2 | 261.93 | 19 | 17 | 17 | 41627000.00 | 0.26156313 | Y | 102908 | Alpha-aminoadipic semialdehyde synthase, mitochondrial OS=Rattus norvegicus OX=10116 GN=Aass PE=2 SV=1 |
| 27 | 35 | 99.2 | 213.6 | 29 | 11 | 11 | 8325500 | 0.08066701 | Y | 36148 | P13086\|SUCA_RAT | 1.256278604 | 23 | 32 | 99.1 | 228.38 | 27 | 8 | 8 | 16128000.00 | 0.10134024 | Y | 36148 | Succinate--CoA ligase [ADP/GDP-forming] subunit alpha, mitochondrial OS=Rattus norvegicus OX=10116 GN=Suclg1 PE=2 SV=2 |
| 28 | 39 | 99.1 | 208.41 | 24 | 8 | 8 | 13235000 | 0.12823589 | N | 44695 | P17764\|THIL_RAT | 0.87434744 | 21 | 46 | 99.2 | 236.91 | 27 | 11 | 11 | 17844000.00 | 0.11212272 | N | 44695 | Acetyl-CoA acetyltransferase, mitochondrial OS=Rattus norvegicus OX=10116 GN=Acat1 PE=1 SV=1 |
| 32 | 289 | 98.9 | 188.9 | 11 | 7 | 7 | 3079100 | 0.02983386 | Y | 74679 | P07872\|ACOX1_RAT | 0.528920715 | 55 | 1310 | 98.4 | 154.71 | 7 | 3 | 3 | 2511300.00 | 0.01577975 | N | 74679 | Peroxisomal acyl-coenzyme A oxidase 1 OS=Rattus norvegicus OX=10116 GN=Acox1 PE=1 SV=1 |
| 30 | 4602 | 99.1 | 182.87 | 11 | 8 | 8 | 7627400 | 0.07390302 | Y | 87371 | B3DMA2\|ACD11_RAT | 0.877188 | 32 | 53 | 99.1 | 206.36 | 13 | 8 | 8 | 10317000.00 | 0.06482684 | N | 87371 | Acyl-CoA dehydrogenase family member 11 OS=Rattus norvegicus OX=10116 GN=Acad11 PE=1 SV=1 |
| 33 | 4338 | 99.1 | 170.4 | 16 | 7 | 6 | 2991300 | 0.02898315 | N | 60956 | P63039\|CH60_RAT | 11.36152989 | 15 | 61 | 99.2 | 260.8 | 28 | 17 | 16 | 52406000.00 | 0.32929294 | Y | 60956 | 60 kDa heat shock protein, mitochondrial OS=Rattus norvegicus OX=10116 GN=Hspd1 PE=1 SV=1 |
| 39 | 9290 | 98.9 | 137.22 | 19 | 5 | 5 | 3989000 | 0.03865002 | Y | 35684 | P04636\|MDHM_RAT | 1.320297933 | 36 | 56 | 99.1 | 185.21 | 24 | 7 | 7 | 8121200.00 | 0.05102954 | N | 35684 | Malate dehydrogenase, mitochondrial OS=Rattus norvegicus OX=10116 GN=Mdh2 PE=1 SV=2 |
| 35 | 4857 | 99 | 127.43 | 10 | 5 | 5 | 3293100 | 0.03190734 | Y | 88126 | P32198\|CPT1A_RAT | 1.357356511 | 29 | 119 | 99.1 | 165.28 | 13 | 8 | 8 | 6892600.00 | 0.04330963 | Y | 88126 | Carnitine O-palmitoyltransferase 1, liver isoform OS=Rattus norvegicus OX=10116 GN=Cpt1a PE=1 SV=2 |
| 41 | 278 | 98.4 | 126.49 | 15 | 4 | 4 | 7551900 | 0.07317148 | N | 60420 | tr\|D3ZFJ6\|D3ZFJ6_RAT | 1.81141719 | 28 | 96 | 99.1 | 129.48 | 18 | 8 | 8 | 21094000.00 | 0.13254409 | N | 60420 | Lactamase, beta OS=Rattus norvegicus OX=10116 GN=Lactb PE=1 SV=1 |
| 47 | 106 | 84.4 | 124.05 | 12 | 2 | 2 | 4104600 | 0.03977008 | N | 37791 | tr\|G3V6I4\|G3V6I4_RAT | 0.262367457 | 82 | 248 | 88.2 | 64.95 | 5 | 2 | 2 | 1660600.00 | 0.01043438 | N | 37791 | Mitochondrial amidoxime reducing component 1 OS=Rattus norvegicus OX=10116 GN=Marc1 PE=1 SV=1 |
| 45 | 98 | 98.7 | 122.39 | 13 | 4 | 4 | 4000100 | 0.03875757 | N | 38202 | P29147\|BDH_RAT | 2.494264041 | 33 | 39 | 98.9 | 117.79 | 10 | 8 | 8 | 15385000.00 | 0.0966716 | Y | 38202 | D-beta-hydroxybutyrate dehydrogenase, mitochondrial OS=Rattus norvegicus OX=10116 GN=Bdh1 PE=1 SV=2 |
| 66 | 13899 | 70.9 | 118.41 | 8 | 2 | 2 | 991930 | 0.00961096 | Y | 49620 | Q5XIC2\|ECSIT_RAT | - | | | | | | | | | 0 | | | Evolutionarily conserved signaling intermediate in Toll pathway, mitochondrial OS=Rattus norvegicus OX=10116 GN=Ecsit PE=1 SV=1 |
| 59 | 9321 | 84.4 | 115.77 | 7 | 3 | 3 | 1666300 | 0.01614503 | N | 33215 | tr\|Q5EBA4\|Q5EBA4_R/ | 4.200536438 | 39 | 51 | 98.4 | 130.16 | 10 | 4 | 4 | 10793000.00 | 0.06781778 | N | 33215 | Nipsnap1 protein (Fragment) OS=Rattus norvegicus OX=10116 GN=Nipsnap1 PE=2 SV=1 |
| 43 | 110 | 98.4 | 113.73 | 16 | 3 | 3 | 1933000 | 0.01872913 | N | 34448 | Q9WVK7\|HCDH_RAT | 0.685178074 | 90 | 137 | 80.3 | 61.57 | 4 | 2 | 2 | 2042300.00 | 0.01283279 | N | 34448 | Hydroxyacyl-coenzyme A dehydrogenase, mitochondrial OS=Rattus norvegicus OX=10116 GN=Hadh PE=2 SV=1 |
| 54 | 2174 | 98.4 | 109.85 | 22 | 3 | 3 | 4123800 | 0.03995611 | N | 27246 | O70351\|HCD2_RAT | 0.974602918 | 51 | 76 | 98.6 | 100.59 | 18 | 4 | 4 | 6197400.00 | 0.03894134 | N | 27246 | 3-hydroxyacyl-CoA dehydrogenase type-2 OS=Rattus norvegicus OX=10116 GN=Hsd17b10 PE=1 SV=3 |
| 102 | 178 | 68.7 | 96.06 | 11 | 1 | 1 | 1123600 | 0.01088673 | N | 26674 | P62909\|RS3_RAT | 0.593619747 | 205 | 145 | 61.7 | 74.18 | 7 | 1 | 1 | 1028500.00 | 0.00646258 | Y | 26674 | 40S ribosomal protein S3 OS=Rattus norvegicus OX=10116 GN=Rps3 PE=1 SV=1 |

| | | | | | | | | | | | | | | | | | | | | | | | | |
|---|---|---|---|---|---|---|---|---|---|---|---|---|---|---|---|---|---|---|---|---|---|---|---|---|
| 46 | 47 | 98.3 | 88.14 | 9 | 3 | 3 | 1768600 | **0.01713623** | N | 32433 | Q9WVK3\|PECR_RAT | **2.924779148** | 64 | 48 | 98.3 | 91.06 | 9 | 3 | 3 | 7976400.00 | **0.05011969** | N | 32433 | Peroxisomal trans-2-enoyl-CoA reductase OS=Rattus norvegicus OX=10116 GN=Pecr PE=2 SV=1 |
| 127 | 6797 | 61.7 | 73.01 | 10 | 1 | 1 | 408280 | **0.00395589** | N | 17472 | P08011\|MGST1_RAT | **2.774444156** | 89 | 176 | 84.4 | 111.12 | 19 | 2 | 2 | 1746700.00 | **0.01097538** | Y | 17472 | Microsomal glutathione S-transferase 1 OS=Rattus norvegicus OX=10116 GN=Mgst1 PE=1 SV=3 |
| 76 | 9291 | 79.6 | 47.25 | 7 | 2 | 2 | 154490 | **0.00149688** | N | 26435 | tr\|D3ZUX5\|D3ZUX5_RA | - | | | | | | | | | **0** | | | MICOS complex subunit OS=Rattus norvegicus OX=10116 GN=Chchd3 PE=1 SV=1 |
| 52 | 4633 | 61.5 | 40.49 | 10 | 1 | 1 | 1044200 | **0.01011741** | N | 39886 | P00481\|OTC_RAT | - | | | | | | | | | **0** | | | Ornithine carbamoyltransferase, mitochondrial OS=Rattus norvegicus OX=10116 GN=Otc PE=1 SV=1 |
| 48 | 179 | 76.3 | 49.15 | 2 | 3 | 1 | 1275600 | **0.01235948** | Y | 140418 | F1M775\|DIAP1_RAT | - | | | | | | | | | **0** | | | Protein diaphanous homolog 1 OS=Rattus norvegicus OX=10116 GN=Diaph1 PE=1 SV=3 |
| 50 | 603 | 90.6 | 98.87 | 3 | 2 | 2 | 1012600 | **0.00981123** | N | 74675 | A0A0G2K047\|ACSS3_R | - | | | | | | | | | **0** | | | Acyl-CoA synthetase short-chain family member 3, mitochondrial OS=Rattus norvegicus OX=10116 GN=Acss3 PE=1 SV=1 |
| 51 | 11724 | 89.7 | 95.53 | 8 | 2 | 1 | 1212000 | **0.01174325** | N | 36133 | Q64591\|DECR_RAT | **1.749475152** | 81 | 118 | 90.1 | 111.64 | 10 | 2 | 2 | 3269600.00 | **0.02054452** | N | 36133 | 2,4-dienoyl-CoA reductase, mitochondrial OS=Rattus norvegicus OX=10116 GN=Decr1 PE=1 SV=2 |
| 57 | 4472 | 93.7 | 123.85 | 6 | 3 | 3 | 1443600 | **0.01398726** | N | 70749 | P45953\|ACADV_RAT | **3.856371336** | 48 | 91 | 98.6 | 160.33 | 9 | 4 | 4 | 8584400.00 | **0.05394005** | N | 70749 | Very long-chain specific acyl-CoA dehydrogenase, mitochondrial OS=Rattus norvegicus OX=10116 GN=Acadvl PE=1 SV=1 |
| 65 | 761 | 84 | 66.21 | 4 | 2 | 2 | 160410 | **0.00155424** | N | 73858 | P48721\|GRP75_RAT | **8.625759348** | 49 | 417 | 97.8 | 102.45 | 7 | 4 | 4 | 2133600.00 | **0.01340647** | N | 73858 | Stress-70 protein, mitochondrial OS=Rattus norvegicus OX=10116 GN=Hspa9 PE=1 SV=3 |
| 67 | 6851 | 68.3 | 75.82 | 5 | 1 | 1 | 516050 | **0.00500009** | N | 33899 | Q5PQT3\|GLYAT_RAT | **2.585866196** | 50 | 5017 | 98.5 | 158.82 | 14 | 3 | 3 | 2057700.00 | **0.01292955** | N | 33899 | Glycine N-acyltransferase OS=Rattus norvegicus OX=10116 GN=Glyat PE=2 SV=1 |
| 75 | 206 | 83.7 | 64.17 | 5 | 2 | 2 | 1193300 | **0.01156206** | N | 38249 | O88994\|MARC2_RAT | **2.603327737** | 58 | 163 | 98.3 | 67.29 | 7 | 3 | 3 | 4790300.00 | **0.03009984** | N | 38249 | Mitochondrial amidoxime reducing component 2 OS=Rattus norvegicus OX=10116 GN=Marc2 PE=2 SV=1 |
| 78 | 181 | 58 | 33.22 | 3 | 2 | 1 | 1064400 | **0.01031313** | N | 50976 | P61980\|HNRPK_RAT | - | | | | | | | | | **0** | | | Heterogeneous nuclear ribonucleoprotein K OS=Rattus norvegicus OX=10116 GN=Hnrnpk PE=1 SV=1 |
| 93 | 4527 | 41.8 | 43973 | 4 | 1 | 1 | 430990 | **0.00417593** | N | 80461 | Q5XHZ0\|TRAP1_RAT | - | | | | | | | | | **0** | | | Heat shock protein 75 kDa, mitochondrial OS=Rattus norvegicus OX=10116 GN=Trap1 PE=1 SV=1 |
| 101 | 7185 | 70.5 | 58.08 | 4 | 1 | 1 | 3606500 | **0.03494392** | N | 105792 | Q924S5\|LONM_RAT | **0.254188778** | 84 | 141 | 73.5 | 54.31 | 2 | 2 | 2 | 1413600.00 | **0.00888235** | Y | 105792 | Lon protease homolog, mitochondrial OS=Rattus norvegicus OX=10116 GN=Lonp1 PE=2 SV=1 |
| 125 | 13901 | 61.7 | 84.02 | 9 | 1 | 1 | 933500 | **0.00904482** | N | 17193 | Q05982\|NDKA_RAT | - | | | | | | | | | **0** | | | Nucleoside diphosphate kinase A OS=Rattus norvegicus OX=10116 GN=Nme1 PE=1 SV=1 |
| 128 | 13902 | 61.7 | 73.01 | 5 | 1 | 1 | 302750 | **0.00293339** | N | 46301 | P70619\|GSHR_RAT | - | | | | | | | | | **0** | | | Glutathione reductase (Fragment) OS=Rattus norvegicus OX=10116 GN=Gsr PE=2 SV=2 |
| 129 | 376 | 61.7 | 69.73 | 9 | 1 | 1 | 290590 | **0.00281557** | N | 29441 | tr\|D3ZXF9\|D3ZXF9_RAT | **5.236229213** | 104 | 106 | 43882 | 25.84 | 6 | 2 | 2 | 2346300.00 | **0.01474297** | Y | 29441 | Mitochondrial ribosomal protein L12 OS=Rattus norvegicus OX=10116 GN=Mrpl12 PE=1 SV=1 |
| 130 | 9340 | 61.7 | 72.62 | 5 | 1 | 1 | 295080 | **0.00285907** | N | 29211 | D4A7N1\|MIC25_RAT | - | | | | | | | | | **0** | | | MICOS complex subunit Mic25 OS=Rattus norvegicus OX=10116 GN=Chchd6 PE=1 SV=1 |
| 131 | 2210 | 61.7 | 66.41 | 5 | 1 | 1 | 831730 | **0.00805876** | N | 29822 | Q8VID1\|DHRS4_RAT | **10.68905185** | 60 | 66 | 94.9 | 118.76 | 12 | 3 | 3 | 13709000.00 | **0.08614046** | N | 29822 | Dehydrogenase/reductase SDR family member 4 OS=Rattus norvegicus OX=10116 GN=Dhrs4 PE=2 SV=2 |
| 133 | 4512 | 61.7 | 58.38 | 3 | 1 | 1 | 131730 | **0.00127635** | N | 34869 | Q5RJR8\|LRC59_RAT | **12.00575455** | 65 | 111 | 93.4 | 112.19 | 10 | 3 | 3 | 2438700.00 | **0.01532356** | Y | 34869 | Leucine-rich repeat-containing protein 59 OS=Rattus norvegicus OX=10116 GN=Lrrc59 PE=1 SV=1 |
| 140 | 4278 | 58.2 | 29.99 | 3 | 1 | 1 | 1315400 | **0.01274511** | Y | 35344 | tr\|A0A0U1RRQ6\|A0A0 | - | | | | | | | | | **0** | | | Solute carrier family 25, member 44 OS=Rattus norvegicus OX=10116 GN=Slc25a44 PE=1 SV=1 |
| **137** | 622 | | 34.62 | 2 | 1 | 1 | 1753800 | **0.01699283** | N | 78179 | P18163\|ACSL1_RAT | **16.22233038** | 16 | 28 | 99.2 | 264.34 | 26 | 17 | 16 | 43871000.00 | **0.2756633** | Y | 78179 | Long-chain-fatty-acid--CoA ligase 1 OS=Rattus norvegicus OX=10116 GN=Acsl1 PE=1 SV=1 |
| **153** | 141 | | 26.62 | 0 | 1 | 1 | 555250 | **0.0053799** | N | 200202 | Q9JIR0\|RIMB1_RAT | **4.248797423** | 54 | 460 | 83.1 | 53.88 | 1 | 3 | 2 | 3637800.00 | **0.02285811** | N | 200202 | Peripheral-type benzodiazepine receptor-associated protein 1 OS=Rattus norvegicus OX=10116 GN=Tspoap1 PE=1 SV=2 |
| **79** | 2538 | | 31.94 | 1 | 2 | 0 | | **0** | N | 119588 | D3ZG52\|DNA2_RAT | **+** | 69 | 1344 | 44004 | 37.97 | 1 | 3 | 2 | 31878000.00 | **0.20030532** | N | 119588 | DNA replication ATP-dependent helicase/nuclease DNA2 OS=Rattus norvegicus OX=10116 GN=Dna2 PE=3 SV=1 |
| | | | | | | | | **0** | | | Q5XIT9\|MCCB_RAT | **+** | 44 | 12439 | 99 | 204.16 | 13 | 5 | 5 | 5586100.00 | **0.03510024** | N | 61517 | Methylcrotonoyl-CoA carboxylase beta chain, mitochondrial OS=Rattus norvegicus OX=10116 GN=Mccc2 PE=2 SV=1 |
| | | | | | | | | **0** | | | Q5SGE0\|LPPRC_RAT | **+** | 53 | 8242 | 93.6 | 105.6 | 2 | 3 | 3 | 2652900.00 | **0.01666949** | Y | 156652 | Leucine-rich PPR motif-containing protein, mitochondrial OS=Rattus norvegicus OX=10116 GN=Lrpprc PE=1 SV=1 |
| | | | | | | | | **0** | | | Q5XIN6\|LETM1_RAT | **+** | 56 | 4532 | 98.4 | 106.67 | 7 | 3 | 3 | 2969100.00 | **0.01865633** | N | 83060 | Mitochondrial proton/calcium exchanger protein OS=Rattus norvegicus OX=10116 GN=Letm1 PE=1 SV=1 |
| | | | | | | | | **0** | | | P17178\|CP27A_RAT | **+** | 72 | 78 | 91.6 | 99.93 | 7 | 2 | 2 | 1143900.00 | **0.00718769** | N | 60733 | Sterol 26-hydroxylase, mitochondrial OS=Rattus norvegicus OX=10116 GN=Cyp27a1 PE=1 SV=1 |
| | | | | | | | | **0** | | | Q5BK22\|GTPB8_RAT | **+** | 85 | 12478 | 60.6 | 35.74 | 7 | 2 | 2 | 2574300.00 | **0.01617561** | Y | 32232 | GTP-binding protein 8 OS=Rattus norvegicus OX=10116 GN=Gtpbp8 PE=2 SV=1 |
| | | | | | | | | **0** | | | tr\|F1LPV8\|F1LPV8_RAT | **+** | 93 | 4920 | 68.8 | 38.94 | 3 | 2 | 2 | 1047100.00 | **0.00657945** | N | 46639 | Succinate--CoA ligase [GDP-forming] subunit beta, mitochondrial OS=Rattus norvegicus OX=10116 GN=Suclg2 PE=1 SV=2 |
| | | | | | | | | **0** | | | Q66H15\|RMD3_RAT | **+** | 155 | 244 | 65.4 | 101.79 | 5 | 1 | 1 | 335860.00 | **0.00211038** | N | 52312 | Regulator of microtubule dynamics protein 3 OS=Rattus norvegicus OX=10116 GN=Rmdn3 PE=1 SV=1 |
| | | | | | | | | **0** | | | Q6MGB5\|DHB8_RAT | **+** | 206 | 12874 | 61.7 | 72.97 | 5 | 1 | 1 | 504990.00 | **0.0031731** | N | 26791 | Estradiol 17-beta-dehydrogenase 8 OS=Rattus norvegicus OX=10116 GN=Hsd17b8 PE=1 SV=1 |
| | | | | | | | | **0** | | | Q63716\|PRDX1_RAT | **+** | 209 | 1706 | 61.7 | 53.4 | 6 | 1 | 1 | 2002900.00 | **0.01258522** | N | 22109 | Peroxiredoxin-1 OS=Rattus norvegicus OX=10116 GN=Prdx1 PE=1 SV=1 |
| | | | | | | | | **0** | | | P04041\|GPX1_RAT | **+** | 210 | 291 | 61.7 | 52.02 | 6 | 1 | 1 | 856390.00 | **0.00538112** | N | 22305 | Glutathione peroxidase 1 OS=Rattus norvegicus OX=10116 GN=Gpx1 PE=1 SV=4 |
| | | | | | | | | **0** | | | tr\|B2RYM8\|B2RYM8_R | **+** | 216 | 31543 | 61.1 | 39.01 | 7 | 1 | 1 | 68923.00 | **0.00043308** | N | 20157 | Family with sequence similarity 210, member B OS=Rattus norvegicus OX=10116 GN=Fam210b PE=1 SV=1 |
| | | | | | | | | **0** | | | tr\|Q6IEA8\|Q6IEA8_RAT | **+** | 226 | 8311 | 43855 | 21.87 | 8 | 1 | 1 | 1211800.00 | **0.00761434** | Y | 8528 | Interferon, alpha-inducible protein 27-like 2B OS=Rattus norvegicus OX=10116 GN=Ifi27l2b PE=2 SV=1 |
| | | | | | | | | **0** | | | Q09073\|ADT2_RAT | **+** | 37 | 63 | 99 | 101.27 | 16 | 6 | 6 | 9141500.00 | **0.05744059** | Y | 32901 | ADP/ATP translocase 2 OS=Rattus norvegicus OX=10116 GN=Slc25a5 PE=1 SV=3 |
| | | | | | | | | **0** | | | Q9Z2L0\|VDAC1_RAT | **+** | 42 | 1035 | 98.5 | 90.61 | 18 | 5 | 4 | 3497000.00 | **0.02197339** | Y | 30756 | Voltage-dependent anion-selective channel protein 1 OS=Rattus norvegicus OX=10116 GN=Vdac1 PE=1 SV=4 |
| | | | | | | | 10320823750 | | | | P81155\|VDAC2_RAT | **+** | 91 | 1402 | 77.1 | 48.96 | 5 | 2 | 1 | 786220.00 | **0.00494021** | N | 31746 | Voltage-dependent anion-selective channel protein 2 OS=Rattus norvegicus OX=10116 GN=Vdac2 PE=1 SV=2 |
| | | | | | | | | **100** | | | | | | | | | | | | | **100** | | | |
| | | | | | | | 103208237.5 | | | | | | | | | | | | | | | | | |
| | | | | | | | 103208237.5 | | | | | | | | | | | | | | | | | |
