## Supplementary material for "Permeability transition pore-related changes in the proteome and channel activity of ATP synthase dimers and monomers": ATP-synthase

| Protein Group | Protein ID | -10lgP | Coverage (%) | Coverage (%) Sample 1 | Area (IBAQ) | riBAQ | #Peptides | #Unique | #Spec Sample 1 | PTM | Avg. Mass | Accession | riBAQ PTP/Control | Protein Group | Protein ID | -10lgP | Coverage (%) | Coverage (%) Sample 1 | Area (IBAQ) | riBAQ | #Peptides | #Unique | #Spec Sample 1 | PTM | Avg. Mass | Description |
|---|---|---|---|---|---|---|---|---|---|---|---|---|---|---|---|---|---|---|---|---|---|---|---|---|---|---|
| | | | | | CONTROL | % | | | | | | | | | | | | | PTP | % | | | | | | |
| Complex I | | | | | | | | | | | | | | | | | | | | | | | | | | |
| 3 | 3421 | 260.31 | 28 | 28 | 48874000 | 4.260063092 | 56 | 56 | 173 | Y | 79412 | Q66HF1\|NDUS1_RAT | 0.339180642 | 5 | 20 | 252.49 | 21 | 21 | 44940000 | 1.444930933 | 39 | 39 | 140 | Carbam | 79412 | NADH-ubiquinone oxidoreductase 75 kDa subunit mitochondrial OS=Rattus norvegicus OX=10116 GN=Ndufs1 PE=1 SV=1 |
| 4 | 2870 | 232.09 | 39 | 39 | 66119000 | 5.763209714 | 31 | 31 | 121 | N | 27378 | P19234\|NDUV2_RAT | 0.597557383 | 3 | 37 | 241.39 | 44 | 44 | 107110000 | 3.443848515 | 30 | 30 | 145 | | 27378 | NADH dehydrogenase [ubiquinone] flavoprotein 2 mitochondrial OS=Rattus norvegicus OX=10116 GN=Ndufv2 PE=1 SV=2 |
| 5 | 2856 | 210.73 | 30 | 30 | 34707000 | 3.025207876 | 32 | 32 | 101 | Y | 52562 | Q641Y2\|NDUS2_RAT | 0.41439264 | 9 | 26 | 204.62 | 26 | 26 | 38990000 | 1.253623878 | 29 | 29 | 80 | Oxidati | 52562 | NADH dehydrogenase [ubiquinone] iron-sulfur protein 2 mitochondrial OS=Rattus norvegicus OX=10116 GN=Ndufs2 PE=1 SV=1 |
| 7 | 5406 | 228.99 | 38 | 38 | 15159000 | 1.321322102 | 31 | 30 | 81 | N | 42559 | Q5BK63\|NDUA9_RAT | 0.450073184 | 11 | 49 | 195.5 | 30 | 30 | 18496000 | 0.594691645 | 21 | 21 | 61 | | 42559 | NADH dehydrogenase [ubiquinone] 1 alpha subcomplex subunit 9 mitochondrial OS=Rattus norvegicus OX=10116 GN=Ndufa9 PE=1 SV=2 |
| 15 | 2857 | 219.65 | 23 | 23 | 19269000 | 1.679566962 | 13 | 13 | 38 | Y | 21959 | tr\|B2RYS8\|B2RYS8_RAT | 0.444832751 | 12 | 212 | 246.45 | 48 | 48 | 23237000 | 0.747126393 | 23 | 23 | 61 | Oxidati | 21959 | NADH dehydrogenase [ubiquinone] 1 beta subcomplex subunit 8 mitochondrial OS=Rattus norvegicus OX=10116 GN=Ndufb8 PE=1 SV=1 |
| 16 | 2821 | 154.05 | 27 | 27 | 18181000 | 1.584732313 | 10 | 10 | 37 | N | 14359 | tr\|Q5PQZ9\|Q5PQZ9_RAT | 0.470499164 | 13 | 77 | 201.91 | 37 | 37 | 23190000 | 0.745615228 | 13 | 13 | 56 | | 14359 | NADH dehydrogenase [ubiquinone] 1 subunit C2 OS=Rattus norvegicus OX=10116 GN=Ndufc2 PE=1 SV=1 |
| 8 | 2807 | 206.44 | 43 | 43 | 23602000 | 2.057249439 | 35 | 35 | 79 | N | 30226 | tr\|D3ZG43\|D3ZG43_RAT | 0.29913622 | 14 | 74 | 164.22 | 34 | 34 | 19140000 | 0.615397821 | 22 | 22 | 52 | | 30226 | NADH dehydrogenase (Ubiquinone) Fe-S protein 3 (Predicted) isoform CRA_c OS=Rattus norvegicus OX=10116 GN=Ndufs3 PE=1 SV=1 |
| 26 | 5412 | 173.26 | 34 | 34 | 6636500 | 0.578465211 | 15 | 15 | 24 | N | 21664 | tr\|D4A565\|D4A565_RAT | 0.739800858 | 15 | 60 | 232.34 | 34 | 34 | 13310000 | 0.427949059 | 15 | 15 | 47 | | 21664 | NADH dehydrogenase (Ubiquinone) 1 beta subcomplex 5 (Predicted) isoform CRA_b OS=Rattus norvegicus OX=10116 GN=Ndufb5 PE=1 SV=1 |
| 11 | 5408 | 161.2 | 22 | 22 | 11086000 | 0.966302317 | 17 | 17 | 52 | N | 50731 | tr\|Q5XIH3\|Q5XIH3_RAT | 0.569579069 | 16 | 80 | 134.88 | 14 | 14 | 17118000 | 0.550385574 | 12 | 11 | 46 | | 50731 | NADH dehydrogenase [ubiquinone] flavoprotein 1 mitochondrial OS=Rattus norvegicus OX=10116 GN=Ndufv1 PE=1 SV=1 |
| 18 | 5410 | 130.85 | 33 | 33 | 8793300 | 0.766460956 | 16 | 16 | 35 | Y | 40493 | Q561S0\|NDUAA_RAT | 0.344564675 | 19 | 171 | 142.62 | 24 | 24 | 8213850 | 0.26409537 | 14 | 4 | 43 | Oxidati | 40493 | NADH dehydrogenase [ubiquinone] 1 alpha subcomplex subunit 10 mitochondrial OS=Rattus norvegicus OX=10116 GN=Ndufa10 PE=1 SV=1 |
| 17 | 2872 | 172.5 | 32 | 32 | 14159000 | 1.234157902 | 13 | 13 | 36 | Y | 17634 | tr\|D4A7L4\|D4A7L4_RAT | 0.584609872 | 20 | 160 | 196.05 | 26 | 26 | 22440000 | 0.721500893 | 9 | 9 | 41 | | 17634 | NADH dehydrogenase (Ubiquinone) 1 beta subcomplex 11 (Predicted) OS=Rattus norvegicus OX=10116 GN=Ndufb11 PE=1 SV=1 |
| 20 | 3028 | 148.03 | 29 | 29 | 8295600 | 0.723079334 | 11 | 11 | 32 | N | 19965 | tr\|A0A0G2JVL6\|A0A0G2JVL6_ | 1.139752531 | 21 | 161 | 161.02 | 44 | 44 | 25632000 | 0.824131501 | 13 | 13 | 39 | Carbam | 19965 | NADH dehydrogenase [ubiquinone] 1 alpha subcomplex subunit 8 OS=Rattus norvegicus OX=10116 GN=Ndufa8 PE=1 SV=1 |
| 21 | 5411 | 167.03 | 26 | 26 | 7454200 | 0.649739377 | 13 | 13 | 32 | Y | 23970 | tr\|B0BNE6\|B0BNE6_RAT | 0.500987592 | 22 | 199 | 172.57 | 25 | 25 | 10124000 | 0.325511366 | 13 | 13 | 39 | | 23970 | NADH dehydrogenase (Ubiquinone) Fe-S protein 8 (Predicted) isoform CRA_a OS=Rattus norvegicus OX=10116 GN=Ndufs8 PE=1 SV=1 |
| 28 | 180 | 142.75 | 21 | 21 | 12726000 | 1.109251604 | 9 | 9 | 22 | N | 20859 | tr\|D4A0T0\|D4A0T0_RAT | 0.371944641 | 23 | 172 | 158.95 | 27 | 27 | 12832000 | 0.41258019 | 10 | 10 | 38 | | 20859 | NADH:ubiquinone oxidoreductase subunit B10 OS=Rattus norvegicus OX=10116 GN=Ndufb10 PE=1 SV=1 |
| 48 | 5458 | 126.04 | 17 | 17 | 5432200 | 0.473493365 | 3 | 3 | 8 | N | 17178 | tr\|F1LXA0\|F1LXA0_RAT | 1.205309224 | 25 | 211 | 173.13 | 29 | 29 | 17750000 | 0.57070592 | 8 | 8 | 37 | | 17178 | NADH dehydrogenase [ubiquinone] 1 alpha subcomplex subunit 12 OS=Rattus norvegicus OX=10116 GN=Ndufa12 PE=1 SV=2 |
| 40 | 5448 | 86.16 | 26 | 26 | 1908900 | 0.166387741 | 4 | 4 | 11 | Y | 15638 | tr\|D3ZZ21\|D3ZZ21_RAT | 1.626156348 | 26 | 313 | 151.15 | 49 | 49 | 8415300 | 0.270572481 | 10 | 10 | 35 | Oxidati | 15638 | NADH dehydrogenase (Ubiquinone) 1 beta subcomplex 6 (Predicted) OS=Rattus norvegicus OX=10116 GN=Ndufb6 PE=1 SV=1 |
| 23 | 5414 | 140.47 | 9 | 9 | 8536000 | 0.744033608 | 11 | 11 | 27 | Y | 68618 | P11661\|NU5M_RAT | 0.402501058 | 27 | 138 | 141.53 | 10 | 10 | 9314200 | 0.299474315 | 12 | 12 | 32 | Oxidati | 68618 | NADH-ubiquinone oxidoreductase chain 5 OS=Rattus norvegicus OX=10116 GN=Mtnd5 PE=3 SV=3 |
| 39 | 5447 | 112.42 | 26 | 26 | 1490000 | 0.129874657 | 8 | 8 | 11 | N | 16777 | tr\|D3ZE15\|D3ZE15_RAT | 1.257111356 | 28 | 72 | 114.68 | 28 | 28 | 5077900 | 0.163266907 | 8 | 7 | 28 | | 16777 | NADH:ubiquinone oxidoreductase subunit A13 OS=Rattus norvegicus OX=10116 GN=Ndufa13 PE=1 SV=1 |
| 63 | 5445 | 101.71 | 19 | 19 | 560640 | 0.048867737 | 3 | 3 | 4 | N | 15064 | tr\|F1LPG5\|F1LPG5_RAT | 4.59813775 | 30 | 56 | 128.69 | 35 | 35 | 6988600 | 0.224700586 | 9 | 9 | 26 | | 15064 | NADH:ubiquinone oxidoreductase subunit B4 OS=Rattus norvegicus OX=10116 GN=Ndufb4 PE=1 SV=1 |
| 29 | 5440 | 151.01 | 19 | 19 | 6841500 | 0.596333872 | 7 | 7 | 22 | N | 14854 | Q80W89\|NDUAB_RAT | 0.8976617 | 31 | 198 | 165.2 | 18 | 18 | 16649000 | 0.535306077 | 7 | 7 | 26 | | 14854 | NADH dehydrogenase [ubiquinone] 1 alpha subcomplex subunit 11 OS=Rattus norvegicus OX=10116 GN=Ndufa11 PE=2 SV=1 |
| 43 | 5455 | 104.79 | 31 | 31 | 7056100 | 0.615039309 | 7 | 7 | 9 | Y | 10170 | tr\|A0A0G2KAA3\|A0A0G2KAA | 0.829846034 | 35 | 250 | 198.26 | 34 | 34 | 15874000 | 0.510387931 | 9 | 9 | 24 | Oxidati | 10170 | NADH:ubiquinone oxidoreductase subunit A3 OS=Rattus norvegicus OX=10116 GN=Ndufa3 PE=1 SV=1 |
| | | | | | | 0 | | | | | | tr\|D3ZLT1\|D3ZLT1_RAT | + | 38 | 3918 | 74.26 | 16 | 16 | 6726500 | 0.216273429 | 3 | 3 | 20 | Oxidati | 16568 | NADH dehydrogenase (Ubiquinone) 1 beta subcomplex 7 (Predicted) OS=Rattus norvegicus OX=10116 GN=Ndufb7 PE=1 SV=1 |
| 41 | 5449 | 91.1 | 23 | 23 | 1593600 | 0.138904869 | 3 | 3 | 11 | N | 10845 | tr\|D3ZS58\|D3ZS58_RAT | 1.628815001 | 39 | 219 | 133.46 | 12 | 12 | 7036800 | 0.226250334 | 6 | 6 | 19 | | 10845 | NADH dehydrogenase [ubiquinone] 1 alpha subcomplex subunit 2 OS=Rattus norvegicus OX=10116 GN=Ndufa2 PE=1 SV=1 |
| 88 | 5424 | 61.72 | 8 | 8 | 262320 | 0.022864913 | 2 | 2 | 2 | N | 15224 | tr\|D4A3V2\|D4A3V2_RAT | 4.020020277 | 40 | 827 | 116.96 | 33 | 33 | 2858800 | 0.091917413 | 12 | 12 | 19 | | 15224 | NADH dehydrogenase [ubiquinone] 1 alpha subcomplex subunit 6 OS=Rattus norvegicus OX=10116 GN=Ndufa6 PE=1 SV=1 |
| 37 | 5442 | 111.74 | 17 | 17 | 3348400 | 0.291860606 | 3 | 3 | 12 | N | 12700 | tr\|B5DEL8\|B5DEL8_RAT | 0.519752369 | 41 | 3899 | 120.07 | 22 | 22 | 4718000 | 0.151695241 | 6 | 6 | 17 | | 12700 | NADH dehydrogenase (Ubiquinone) Fe-S protein 5 OS=Rattus norvegicus OX=10116 GN=Ndufs5 PE=1 SV=1 |
| 55 | 5450 | 76.99 | 21 | 21 | 1509700 | 0.131591792 | 3 | 3 | 6 | N | 12500 | tr\|A9UMV9\|A9UMV9_RAT | 7.246235361 | 44 | 169 | 115.75 | 21 | 21 | 29657000 | 0.953545098 | 4 | 4 | 15 | | 12500 | NADH:ubiquinone oxidoreductase subunit A7 OS=Rattus norvegicus OX=10116 GN=Ndufa7 PE=1 SV=1 |
| 35 | 564 | 89.03 | 15 | 15 | 2707400 | 0.235988354 | 4 | 4 | 13 | Y | 23945 | tr\|Q5RJN0\|Q5RJN0_RAT | 0.545092606 | 47 | 376 | 91.45 | 21 | 21 | 4000800 | 0.128635507 | 5 | 5 | 13 | Oxidati | 23945 | NADH dehydrogenase (Ubiquinone) Fe-S protein 7 OS=Rattus norvegicus OX=10116 GN=Ndufs7 PE=1 SV=1 |
| 50 | 1301 | 57.87 | 10 | 10 | 721790 | 0.062914248 | 4 | 4 | 7 | N | 38455 | P11662\|NU2M_RAT | 0.778536475 | 51 | 389 | 55.93 | 6 | 6 | 1523400 | 0.048981037 | 3 | 3 | 12 | | 38653 | NADH-ubiquinone oxidoreductase chain 2 OS=Rattus norvegicus OX=10116 GN=Mtnd2 PE=3 SV=3 |
| 57 | 5435 | 52.17 | 21 | 21 | 1520800 | 0.132559315 | 2 | 2 | 5 | N | 13040 | tr\|D3ZCZ9\|D3ZCZ9_RAT | 0.898919711 | 52 | 3906 | 67.58 | 16 | 16 | 3706100 | 0.119160181 | 3 | 3 | 12 | | 13040 | NADH dehydrogenase [ubiquinone] iron-sulfur protein 6 mitochondrial OS=Rattus norvegicus OX=10116 GN=LOC100912599 PE=1 SV=1 |
| 51 | 5457 | 75.88 | 12 | 12 | 973700 | 0.084871781 | 5 | 5 | 7 | N | 19741 | Q5XIF3\|NDUS4_RAT | 1.685363507 | 55 | 238 | 107.99 | 12 | 12 | 4448800 | 0.143039803 | 3 | 3 | 11 | | 19741 | NADH dehydrogenase [ubiquinone] iron-sulfur protein 4 mitochondrial OS=Rattus norvegicus OX=10116 GN=Ndufs4 PE=1 SV=1 |
| 108 | 5486 | 24.58 | 9 | 9 | 365080 | 0.031821906 | 1 | 1 | 1 | N | 11842 | tr\|B2RYU0\|B2RYU0_RAT | 6.476682935 | 56 | 3914 | 78.75 | 26 | 26 | 6410100 | 0.206100396 | 5 | 4 | 11 | | 11842 | NADH dehydrogenase (Ubiquinone) 1 beta subcomplex 2 (Predicted) isoform CRA_b OS=Rattus norvegicus OX=10116 GN=Ndufb2 PE=1 SV=1 |
| 38 | 672 | 68.86 | 6 | 6 | 2854700 | 0.248827641 | 5 | 5 | 11 | N | 36145 | P03889\|NU1M_RAT | 0.337924987 | 59 | 3902 | 47.26 | 5 | 5 | 2615200 | 0.084085077 | 3 | 3 | 10 | | 36145 | NADH-ubiquinone oxidoreductase chain 1 OS=Rattus norvegicus OX=10116 GN=Mtnd1 PE=1 SV=3 |
| 62 | 5425 | 26.36 | 2 | 2 | 362850 | 0.03162753 | 1 | 1 | 4 | N | 51783 | P05508\|NU4M_RAT | 0.684403476 | 62 | 363 | 38.13 | 5 | 5 | 673230 | 0.021645991 | 3 | 3 | 7 | | 51783 | NADH-ubiquinone oxidoreductase chain 4 OS=Rattus norvegicus OX=10116 GN=Mtnd4 PE=3 SV=3 |
| 64 | 5451 | 30.36 | 15 | 15 | 726610 | 0.063334379 | 2 | 2 | 4 | N | 11267 | tr\|D4A4P3\|D4A4P3_RAT | 0.547107145 | 68 | 467 | 49.48 | 30 | 30 | 1077700 | 0.034650691 | 4 | 4 | 7 | Oxidati | 11267 | NADH:ubiquinone oxidoreductase subunit B3 OS=Rattus norvegicus OX=10116 GN=Ndufb3 PE=1 SV=1 |
| 45 | 5439 | 111.71 | 20 | 20 | 1791600 | 0.15616338 | 4 | 4 | 9 | N | 13412 | Q63362\|NDUA5_RAT | 0.38229617 | 71 | 73 | 97.02 | 17 | 17 | 1856800 | 0.059700662 | 4 | 4 | 6 | | 13412 | NADH dehydrogenase [ubiquinone] 1 alpha subcomplex subunit 5 OS=Rattus norvegicus OX=10116 GN=Ndufa5 PE=1 SV=3 |
| 75 | 5460 | 45.67 | 13 | 13 | 277810 | 0.024215086 | 1 | 1 | 3 | N | 8389 | tr\|Q35733\|Q35733_RAT | 1.71669418 | 75 | 3930 | 93.65 | 14 | 14 | 1292900 | 0.041569898 | 3 | 3 | 6 | | 8389 | NADH-ubiquinone oxidoreductase chain 6 (Fragment) OS=Rattus norvegicus OX=10116 PE=3 SV=1 |
| 56 | 5441 | 45.99 | 7 | 7 | 1041600 | 0.09079023 | 1 | 1 | 5 | Y | 21892 | tr\|B2RYW3\|B2RYW3_RAT | 0.411970986 | 76 | 3911 | 51.96 | 18 | 18 | 1163300 | 0.037402941 | 4 | 4 | 5 | Oxidati | 21892 | NADH dehydrogenase (Ubiquinone) 1 beta subcomplex 9 OS=Rattus norvegicus OX=10116 GN=Ndufb9 PE=1 SV=1 |
| 13 | 3210 | 194.65 | 28 | 28 | 14275000 | 1.24426895 | 17 | 17 | 49 | Y | 17514 | tr\|D3ZF13\|D3ZF13_RAT | 0.407529441 | 18 | 252 | 185.52 | 23 | 23 | 15771000 | 0.507076229 | 13 | 13 | 44 | Oxidati | 17514 | Acyl carrier protein OS=Rattus norvegicus OX=10116 GN=Ndufab1 PE=1 SV=1 |
| Complex III | | | | | | | | | | | | | | | | | | | | | | | | | | |
| 10 | 34 | 164.71 | 18 | 18 | 15525000 | 1.353224199 | 22 | 22 | 55 | N | 48396 | P32551\|QCR2_RAT | 0.494490648 | 17 | 50 | 134.81 | 11 | 11 | 20812000 | 0.669156711 | 10 | 10 | 46 | | 48396 | Cytochrome b-c1 complex subunit 2 mitochondrial OS=Rattus norvegicus OX=10116 GN=Uqcrc2 PE=1 SV=2 |
| 44 | 132 | 117.18 | 15 | 15 | 997010 | 0.086903579 | 5 | 5 | 9 | N | 29446 | P20788\|UCRI_RAT | 4.076421914 | 36 | 58 | 186.51 | 17 | 17 | 11018000 | 0.354255652 | 9 | 9 | 21 | | 29446 | Cytochrome b-c1 complex subunit Rieske mitochondrial OS=Rattus norvegicus OX=10116 GN=Uqcrfs1 PE=1 SV=2 |
| 14 | 118 | 176.06 | 23 | 23 | 10042000 | 0.875302893 | 23 | 23 | 47 | Y | 52849 | Q68FY0\|QCR1_RAT | 0.131577381 | 42 | 25 | 165.24 | 14 | 14 | 3582000 | 0.115170062 | 9 | 9 | 16 | Carbam | 52849 | Cytochrome b-c1 complex subunit 1 mitochondrial OS=Rattus norvegicus OX=10116 GN=Uqcrc1 PE=1 SV=1 |
| 33 | 79 | 125.19 | 12 | 12 | 3117800 | 0.271760542 | 8 | 8 | 17 | N | 43012 | P00159\|CYB_RAT | 0.231571451 | 43 | 130 | 140.87 | 7 | 7 | 1957300 | 0.062931983 | 6 | 6 | 15 | | 43012 | Cytochrome b OS=Rattus norvegicus OX=10116 GN=Mt-Cyb PE=3 SV=3 |
| 73 | 2875 | 34.79 | 17 | 17 | 314440 | 0.027407911 | 3 | 3 | 3 | N | 10424 | Q5M9I5\|QCR6_RAT | 3.676873165 | 54 | 318 | 111.53 | 18 | 18 | 3134300 | 0.100775412 | 4 | 4 | 12 | | 10424 | Cytochrome b-c1 complex subunit 6 mitochondrial OS=Rattus norvegicus OX=10116 GN=Uqcrh PE=3 SV=1 |
| | | | | | | 0 | | | | | | Q7TQ16\|QCR8_RAT | + | 66 | 208 | 45.75 | 11 | 11 | 727260 | 0.023383188 | 2 | 2 | 7 | | 9849 | Cytochrome b-c1 complex subunit 8 OS=Rattus norvegicus OX=10116 GN=Uqcrq PE=3 SV=1 |
| | | | | | | 0 | | | | | | tr\|B2RYS2\|B2RYS2_RAT | + | 98 | 3893 | 26.54 | 16 | 16 | 480600 | 0.015452466 | 2 | 2 | 2 | | 13559 | Cytochrome b-c1 complex subunit 7 OS=Rattus norvegicus OX=10116 GN=Uqcrb PE=1 SV=1 |
| Complex IV | | | | | | | | | | | | | | | | | | | | | | | | | | |
| 27 | 133 | 167.41 | 12 | 12 | 7580300 | 0.660730782 | 7 | 7 | 23 | Y | 35435 | tr\|D3ZFQ8\|D3ZFQ8_RAT | 0.788566849 | 24 | 59 | 196.84 | 13 | 13 | 16205000 | 0.521030391 | 9 | 9 | 37 | | 35435 | Cytochrome c-1 OS=Rattus norvegicus OX=10116 GN=Cyc1 PE=1 SV=3 |
| 24 | 24 | 133.46 | 32 | 32 | 6358700 | 0.554250996 | 14 | 14 | 26 | Y | 25942 | P00406\|COX2_RAT | 0.25843153 | 34 | 34 | 125.74 | 32 | 32 | 4454900 | 0.143235933 | 10 | 10 | 24 | Oxidati | 25928 | Cytochrome c oxidase subunit 2 OS=Rattus norvegicus OX=10116 GN=Mtco2 PE=1 SV=3 |
| 65 | 134 | 71.8 | 21 | 21 | 321550 | 0.028027648 | 2 | 2 | 4 | N | 7375 | P80432\|COX7C_RAT | 5.454214476 | 45 | 251 | 143.95 | 21 | 21 | 4754500 | 0.152868806 | 4 | 4 | 15 | | 7375 | Cytochrome c oxidase subunit 7C mitochondrial OS=Rattus norvegicus OX=10116 GN=Cox7c PE=1 SV=2 |
| 54 | 151 | 116.98 | 14 | 14 | 1422000 | 0.123947492 | 3 | 3 | 6 | N | 10487 | P10817\|CX6A2_RAT | 1.224904585 | 50 | 253 | 119.87 | 26 | 26 | 4722000 | 0.151823851 | 5 | 5 | 12 | | 10487 | Cytochrome c oxidase subunit 6A2 mitochondrial (Fragment) OS=Rattus norvegicus OX=10116 GN=Cox6a2 PE=1 SV=3 |
| 25 | 86 | 141.75 | 42 | 42 | 4296900 | 0.374535849 | 9 | 9 | 25 | N | 16130 | P11240\|COX5A_RAT | 0.190544029 | 53 | 200 | 155.04 | 29 | 29 | 2219600 | 0.07136557 | 8 | 8 | 12 | | 16130 | Cytochrome c oxidase subunit 5A mitochondrial OS=Rattus norvegicus OX=10116 GN=Cox5a PE=1 SV=1 |
| 87 | 2795 | 37.15 | 3 | 3 | 105120 | 0.009162701 | 2 | 2 | 2 | N | 56845 | P05503\|COX1_RAT | 2.139367268 | 81 | 1026 | 60.72 | 3 | 3 | 609670 | 0.019602382 | 2 | 2 | 5 | | 56845 | Cytochrome c oxidase subunit 1 OS=Rattus norvegicus OX=10116 GN=Mtco1 PE=2 SV=3 |
| 52 | 139 | 61.53 | 11 | 11 | 3573300 | 0.311463834 | 3 | 3 | 7 | N | 13915 | P12075\|COX5B_RAT | 0.903149918 | 82 | 151 | 38.68 | 11 | 11 | 8748900 | 0.281298537 | 3 | 3 | 4 | | 13915 | Cytochrome c oxidase subunit 5B mitochondrial OS=Rattus norvegicus OX=10116 GN=Cox5b PE=1 SV=2 |
| 132 | 1062 | 21.25 | 12 | 12 | 44810 | 0.003905828 | 1 | 1 | 1 | N | 9353 | P35171\|CX7A2_RAT | 9.506216592 | 85 | 263 | 80.03 | 12 | 12 | 1154800 | 0.037129645 | 2 | 2 | 4 | | 9353 | Cytochrome c oxidase subunit 7A2 mitochondrial OS=Rattus norvegicus OX=10116 GN=Cox7a2 PE=1 SV=1 |
| 47 | 290 | 68.37 | 12 | 12 | 2010600 | 0.17525234 | 3 | 3 | 8 | N | 19515 | P10888\|COX41_RAT | 0.15178137 | 86 | 292 | 52.97 | 12 | 12 | 827310 | 0.02660004 | 3 | 3 | 4 | | 19515 | Cytochrome c oxidase subunit 4 isoform 1 mitochondrial OS=Rattus norvegicus OX=10116 GN=Cox4i1 PE=1 SV=1 |
| 66 | 2866 | 49.12 | 3 | 3 | 151450 | 0.013201018 | 1 | 1 | 4 | N | 29871 | P05505\|COX3_RAT | - | 90 | 270 | 55.61 | 3 | 3 | 0 | 0 | 2 | 2 | 3 | | 29871 | Cytochrome c oxidase subunit 3 OS=Rattus norvegicus OX=10116 GN=Mtco3 PE=1 SV=5 |
| | | | | | | 0 | | | | | | P11951\|CX6C2_RAT | + | 104 | 207 | 26.38 | 17 | 17 | 427700 | 0.013751601 | 2 | 2 | 2 | | 8455 | Cytochrome c oxidase subunit 6C-2 OS=Rattus norvegicus OX=10116 GN=Cox6c2 PE=1 SV=3 |
| 130 | 2798 | 23.83 | 14 | 14 | 92435 | 0.008057023 | 1 | 1 | 1 | N | 8995 | P80431\|COX7B_RAT | 0.466382743 | 168 | 3923 | 21.01 | 14 | 14 | 116870 | 0.003757656 | 1 | 1 | 1 | | 8995 | Cytochrome c oxidase subunit 7B mitochondrial OS=Rattus norvegicus OX=10116 GN=Cox7b PE=1 SV=3 |
| 103 | 568 | 28.49 | 9 | 9 | 46814 | 0.004080505 | 1 | 1 | 1 | N | 9327 | tr\|B2RZD6\|B2RZD6_RAT | 8.562681499 | 73 | 209 | 44.09 | 22 | 22 | 1086700 | 0.034940063 | 4 | 4 | 6 | | 9327 | NDUFA4 mitochondrial complex-associated OS=Rattus norvegicus OX=10116 GN=Ndufa4 PE=1 SV=1 |
| | | | | | | 0 | | | | | | | | | | | | | | 0 | | | | | | |
| Complex V | | | | | | 0 | | | | | | | | | | | | | | 0 | | | | | | |
| 2 | 1 | 378.4 | 50 | 50 | 345830000 | 30.14399515 | 124 | 124 | 582 | Y | 56354 | P10719\|ATPB_RAT | 1.420749245 | 1 | 5 | 390.79 | 47 | 47 | 1332000000 | 42.82705836 | 124 | 120 | 681 | Oxidati | 56354 | ATP synthase subunit beta mitochondrial OS=Rattus norvegicus OX=10116 GN=Atp5f1b PE=1 SV=2 |
| 1 | 3 | 312.23 | 48 | 48 | 267690000 | 23.3329846 | 108 | 108 | 601 | Y | 59754 | P15999\|ATPA_RAT | 0.976300655 | 2 | 8 | 331.26 | 41 | 41 | 708500000 | 22.78000815 | 109 | 107 | 645 | Carbamidomethyla | | ATP synthase subunit alpha, mitochondrial |
| 6 | 5 | 246.27 | 34 | 34 | 22396000 | 1.952129415 | 26 | 26 | 89 | Y | 30191 | P35435\|ATPG_RAT | 1.184571098 | 4 | 63 | 262.75 | 37 | 37 | 71921000 | 2.312436085 | 37 | 37 | 141 | Oxidati | 30191 | ATP synthase subunit gamma mitochondrial OS=Rattus norvegicus OX=10116 GN=Atp5f1c PE=1 SV=2 |
| 9 | 7 | 247.72 | 53 | 53 | 20584000 | 1.794187885 | 21 | 21 | 67 | Y | 18763 | P31399\|ATP5H_RAT | 1.529392988 | 6 | 65 | 277.38 | 61 | 61 | 85344000 | 2.74401837 | 30 | 30 | 128 | Oxidati | 18763 | ATP synthase subunit d mitochondrial OS=Rattus norvegicus OX=10116 GN=Atp5pd PE=1 SV=3 |
| 19 | 11 | 122.89 | 39 | 39 | 6407800 | 0.558530758 | 9 | 9 | 33 | Y | 11433 | Q6PDU7\|ATP5L_RAT | 1.96438613 | 7 | 75 | 183.33 | 48 | 48 | 34124000 | 1.097170075 | 19 | 18 | 89 | Oxidati | 11433 | ATP synthase subunit g mitochondrial OS=Rattus norvegicus OX=10116 GN=Atp5mg PE=1 SV=2 |
| 12 | 10 | 124.23 | 17 | 17 | 27671000 | 2.411920568 | 10 | 10 | 49 | N | 28869 | P19511\|AT5F1_RAT | 1.538089311 | 8 | 168 | 173.06 | 20 | 20 | 115380000 | 3.709749245 | 18 | 18 | 81 | | 28869 | ATP synthase F(0) complex subunit B1 mitochondrial OS=Rattus norvegicus OX=10116 GN=Atp5pb PE=1 SV=1 |
| 22 | 8 | 124.51 | 26 | 26 | 7095700 | 0.618491011 | 10 | 10 | 30 | N | 23398 | Q06647\|ATPO_RAT | 2.162536862 | 10 | 76 | 162.91 | 29 | 29 | 41599000 | 1.33750961 | 19 | 18 | 72 | | 23398 | ATP synthase subunit O mitochondrial OS=Rattus norvegicus OX=10116 GN=Atp5po PE=1 SV=1 |
| | | | | | | 0 | | | | | | P29419\|ATP5I_RAT | + | 29 | 319 | 125.84 | 51 | 51 | 10020000 | 0.322167511 | 10 | 10 | 28 | | 8255 | ATP synthase subunit e mitochondrial OS=Rattus norvegicus OX=10116 GN=Atp5me PE=1 SV=3 |
| 49 | 15 | 47.43 | 8 | 8 | 976850 | 0.085146348 | 2 | 2 | 7 | Y | 25076 | P05504\|ATP6_RAT | 1.606860505 | 37 | 258 | 51.09 | 12 | 12 | 4255300 | 0.136818304 | 3 | 3 | 21 | Oxidati | 25076 | ATP synthase subunit a OS=Rattus norvegicus OX=10116 GN=Mt-atp6 PE=1 SV=3 |
| 31 | 12 | 135.48 | 33 | 33 | 5030800 | 0.438505655 | 8 | 8 | 18 | Y | 17563 | tr\|G3V7Y3\|G3V7Y3_RAT | 0.538585023 | 46 | 3953 | 81.42 | 11 | 11 | 7345400 | 0.236172578 | 5 | 5 | 14 | | 17563 | ATP synthase subunit delta mitochondrial OS=Rattus norvegicus OX=10116 GN=Atp5f1d PE=1 SV=1 |
| 46 | 35 | 80.38 | 19 | 19 | 7031700 | 0.612912502 | 3 | 3 | 9 | Y | 7630 | P11608\|ATP8_RAT | 0.531928788 | 48 | 234 | 115.4 | 19 | 19 | 10140000 | 0.326025805 | 4 | 4 | 13 | Oxidati | 7630 | ATP synthase protein 8 OS=Rattus norvegicus OX=10116 GN=Mt-atp8 PE=1 SV=2 |
| | | | | | | 0 | | | | | | D3ZAF6\|ATPK_RAT | + | 78 | #### | 34.41 | 10 | 10 | 1883500 | 0.060559132 | 3 | 3 | 5 | | 10452 | ATP synthase subunit f mitochondrial OS=Rattus norvegicus OX=10116 GN=Atp5mf PE=1 SV=1 |

| | | | | | | | | | | | | | | | | | | | | | | | | | | |
|---|---|---|---|---|---|---|---|---|---|---|---|---|---|---|---|---|---|---|---|---|---|---|---|---|---|---|
| 125 | 89 | 33.84 | 21 | 21 | 548650 | **0.047822638** | 1 | 1 | 1 | N | 6408 | Q9JJW3\|ATPMD_RAT | **1.091388263** | 109 | #### | 34.48 | 22 | 22 | 1623300 | **0.052193066** | 2 | 2 | 2 | | 6408 | ATP synthase membrane subunit DAPIT mitochondrial OS=Rattus norvegicus OX=10116 GN=Atp5md PE=1 SV=1 |
| 123 | 30 | 39.99 | 10 | 10 | 171430 | **0.014942559** | 1 | 1 | 1 | N | 12494 | P21571\|ATP5J_RAT | **1.363878061** | 89 | #### | 60.92 | 24 | 24 | 633850 | **0.020379828** | 3 | 3 | 3 | Oxidati | 12494 | ATP synthase-coupling factor 6 mitochondrial OS=Rattus norvegicus OX=10116 GN=Atp5pf PE=1 SV=1 |
| Others | | | | | | | | | | | | | | | | | | | | | | | | | | |
| 30 | 19 | 153.42 | 16 | 16 | 2596800 | **0.226347994** | 10 | 10 | 20 | Y | 47314 | P00507\|AATM_RAT | **0.735201639** | 33 | 45 | 162.61 | 17 | 17 | 5175700 | **0.166411416** | 11 | 11 | 25 | Oxidati | 47314 | Aspartate aminotransferase mitochondrial OS=Rattus norvegicus OX=10116 GN=Got2 PE=1 SV=2 |
| 32 | 5407 | 124.95 | 32 | 32 | 2115400 | **0.184387148** | 11 | 11 | 18 | N | 29820 | P67779\|PHB_RAT | **0.506889699** | 49 | 532 | 128.14 | 18 | 18 | 2906900 | **0.093463946** | 8 | 8 | 12 | | 29820 | Prohibitin OS=Rattus norvegicus OX=10116 GN=Phb PE=1 SV=1 |
| | | | | | | **0** | | | | | | Q91XJ1\|BECN1_RAT | **+** | 57 | 326 | 54.23 | 3 | 3 | | **0** | 3 | 0 | 11 | | 51557 | Beclin-1 OS=Rattus norvegicus OX=10116 GN=Becn1 PE=1 SV=1 |
| 95 | 1010 | 20.32 | 3 | 3 | 127850 | **0.011143943** | 1 | 1 | 1 | N | 32989 | Q05962\|ADT1_RAT | **4.450701506** | 58 | 24 | 81.73 | 10 | 10 | 1542600 | **0.049598364** | 5 | 5 | 10 | | 32989 | ADP/ATP translocase 1 OS=Rattus norvegicus OX=10116 GN=Slc25a4 PE=1 SV=3 |
| 42 | 9 | 96.05 | 6 | 6 | 1180300 | **0.102879905** | 7 | 7 | 10 | N | 82665 | Q64428\|ECHA_RAT | **0.194005565** | 60 | 21 | 62.55 | 6 | 6 | 620770 | **0.019959274** | 5 | 5 | 7 | | 82665 | Trifunctional enzyme subunit alpha mitochondrial OS=Rattus norvegicus OX=10116 GN=Hadha PE=1 SV=2 |
| 122 | 138 | 44.17 | 2 | 2 | 172350 | **0.01502275** | 1 | 1 | 1 | N | 88217 | Q63704\|CPT1B_RAT | **1.073121549** | 61 | 61 | 83.57 | 3 | 3 | 501400 | **0.016121237** | 3 | 3 | 7 | | 88217 | Carnitine O-palmitoyltransferase 1 muscle isoform OS=Rattus norvegicus OX=10116 GN=Cpt1b PE=1 SV=1 |
| 36 | 284 | 25.68 | 2 | 2 | 2626900 | **0.228971636** | 1 | 1 | 12 | N | 36404 | Q5M934\|TRUB1_RAT | **2.394881654** | 63 | 1269 | 23.08 | 2 | 2 | 17055000 | **0.54835997** | 1 | 1 | 7 | | 36404 | Probable tRNA pseudouridine synthase 1 OS=Rattus norvegicus OX=10116 GN=Trub1 PE=2 SV=1 |
| 60 | 196 | 51.56 | 1 | 1 | 2297500 | **0.200259749** | 3 | 3 | 4 | Y | 223506 | P02563\|MYH6_RAT | **0.239000257** | 64 | 16 | 86.5 | 3 | 3 | 1488600 | **0.047862131** | 5 | 5 | 7 | Oxidati | 223506 | Myosin-6 OS=Rattus norvegicus OX=10116 GN=Myh6 PE=1 SV=2 |
| 58 | 20 | 62.75 | 7 | 7 | 432720 | **0.037717692** | 4 | 4 | 5 | N | 51414 | Q60587\|ECHB_RAT | **0.786657891** | 67 | 29 | 79.98 | 8 | 8 | 922820 | **0.02967092** | 5 | 5 | 7 | | 51414 | Trifunctional enzyme subunit beta mitochondrial OS=Rattus norvegicus OX=10116 GN=Hadhb PE=1 SV=1 |
| 72 | 2803 | 64.55 | 4 | 4 | 240260 | **0.020942071** | 2 | 2 | 3 | N | 51960 | Q6AXV4\|SAM50_RAT | **0.590539357** | 69 | 14 | 77.24 | 6 | 6 | 384640 | **0.012367117** | 3 | 3 | 7 | | 51960 | Sorting and assembly machinery component 50 homolog OS=Rattus norvegicus OX=10116 GN=Samm50 PE=1 SV=1 |
| 34 | 5423 | 102.71 | 13 | 13 | 3542500 | **0.308779177** | 5 | 5 | 15 | N | 33312 | Q5XIH7\|PHB2_RAT | **0.183347944** | 70 | 335 | 74.3 | 10 | 10 | 1760800 | **0.056614027** | 4 | 4 | 6 | | 33312 | Prohibitin-2 OS=Rattus norvegicus OX=10116 GN=Phb2 PE=1 SV=1 |
| | | | | | | **0** | | | | | | B0BN56\|RT31_RAT | **+** | 72 | 8248 | 53.37 | 6 | 6 | 244140 | **0.007849698** | 3 | 1 | 6 | | 43962 | 28S ribosomal protein S31 mitochondrial OS=Rattus norvegicus OX=10116 GN=Mrps31 PE=2 SV=1 |
| 59 | 88 | 78.75 | 4 | 4 | 649530 | **0.056615763** | 3 | 3 | 5 | N | 85433 | Q9ER34\|ACON_RAT | **0.336456066** | 77 | 17 | 89.78 | 3 | 3 | 592450 | **0.019048717** | 3 | 3 | 5 | | 85433 | Aconitate hydratase mitochondrial OS=Rattus norvegicus OX=10116 GN=Aco2 PE=1 SV=2 |
| 61 | 2787 | 62.78 | 8 | 8 | 1100400 | **0.095915485** | 3 | 3 | 4 | N | 44695 | P17764\|THIL_RAT | **0.432764407** | 79 | 44 | 69.95 | 5 | 5 | 1291000 | **0.041508808** | 2 | 2 | 5 | | 44695 | Acetyl-CoA acetyltransferase mitochondrial OS=Rattus norvegicus OX=10116 GN=Acat1 PE=1 SV=1 |
| | | | | | | **0** | | | | | | P11530\|DMD_RAT | **+** | 92 | 224 | 27.58 | 1 | 1 | 2253100 | **0.072442677** | 2 | 2 | 2 | Oxidation (M) | | Dystrophin |
| | | | | | | **0** | | | | | | tr\|Q4G067\|Q4G067_RAT | **+** | 105 | 8042 | 39.5 | 4 | 4 | 361430 | **0.011620859** | 2 | 2 | 2 | | 37440 | Mitochondrial ribosomal protein L44 OS=Rattus norvegicus OX=10116 GN=Mrpl44 PE=1 SV=1 |
| | | | | | | **0** | | | | | | Q9Z0V6\|PRDX3_RAT | **+** | 111 | 316 | 28.27 | 5 | 5 | 356470 | **0.011461383** | 1 | 1 | 2 | | 28295 | Thioredoxin-dependent peroxide reductase mitochondrial OS=Rattus norvegicus OX=10116 GN=Prdx3 PE=1 SV=2 |
| | | | | | | **0** | | | | | | tr\|D4A4B1\|D4A4B1_RAT | **+** | 112 | 4487 | 35.63 | 8 | 8 | 139410 | **0.004482373** | 2 | 2 | 2 | | 23223 | Mitochondrial ribosomal protein L15 OS=Rattus norvegicus OX=10116 GN=Mrpl15 PE=1 SV=2 |
| | | | | | | **0** | | | | | | tr\|D3Z8Q5\|D3Z8Q5_RAT | **+** | 113 | #### | 24.29 | 6 | 6 | 293930 | **0.009450569** | 1 | 1 | 2 | | 12124 | RCG25747 OS=Rattus norvegicus OX=10116 GN=Tusc2 PE=4 SV=1 |
| 71 | 146 | 57.78 | 2 | 2 | 73134 | **0.006374667** | 2 | 2 | 3 | N | 67177 | Q3KR86\|MIC60_RAT | **0.529900029** | 141 | 12 | 20.78 | 1 | 1 | 105060 | **0.003377936** | 1 | 1 | 1 | | 67177 | MICOS complex subunit Mic60 (Fragment) OS=Rattus norvegicus OX=10116 GN=Immt PE=1 SV=1 |
| 53 | 2801 | 77.13 | 8 | 8 | 1358900 | **0.118447431** | 5 | 5 | 6 | Y | 57808 | Q02253\|MMSA_RAT | - | | | | | | | | | | | | | Methylmalonate-semialdehyde dehydrogenase [acylating], mitochondrial OS=Rattus norvegicus OX=10116 GN=Aldh6a1 PE=1 SV=1 |
| 126 | #### | 28.58 | 6 | 6 | 91401 | **0.007966895** | 1 | 1 | 1 | N | 22179 | Q9R063\|PRDX5_RAT | - | | | | | | | | | | | | | Peroxiredoxin-5, mitochondrial OS=Rattus norvegicus OX=10116 GN=Prdx5 PE=1 SV=1 |
| | | | | | 1147260004 | **100** | | | | | | | | | | | | | 3110183260 | **100** | | | | | | |
| | | | | | | | | | | | | | | | | | | | 31101832.6 | | | | | | | |
