## Supplementary material for "Permeability transition pore-related changes in the proteome and channel activity of ATP synthase dimers and monomers": ATP-synthase

| Area (IBAQ) | riBAQ | Accession | rIBAQ PTP/Control | Area (IBAQ) | riBAQ | Description |
|---|---|---|---|---|---|---|
| CONTROL | % | | | PTP | % | |
| **Complex I** | | | | | | |
| 48874000 | **4.26006309** | Q66HF1\|NDUS1_RAT | **0.33918064** | 44940000 | **1.444930933** | NADH-ubiquinone oxidoreductase 75 kDa subunit mitochondrial OS=Rattus norvegicus OX=10116 GN=Ndufs1 PE=1 SV=1 |
| 34707000 | **3.02520788** | Q641Y2\|NDUS2_RAT | **0.41439264** | 38990000 | **1.253623878** | NADH dehydrogenase [ubiquinone] iron-sulfur protein 2 mitochondrial OS=Rattus norvegicus OX=10116 GN=Ndufs2 PE=1 SV=1 |
| 15159000 | **1.3213221** | Q5BK63\|NDUA9_RAT | **0.45007318** | 18496000 | **0.594691645** | NADH dehydrogenase [ubiquinone] 1 alpha subcomplex subunit 9 mitochondrial OS=Rattus norvegicus OX=10116 GN=Ndufa9 PE=1 SV=2 |
| 19269000 | **1.67956696** | tr\|B2RYS8\|B2RYS8_RAT | **0.44483275** | 23237000 | **0.747126393** | NADH dehydrogenase [ubiquinone] 1 beta subcomplex subunit 8 mitochondrial OS=Rattus norvegicus OX=10116 GN=Ndufb8 PE=1 SV=1 |
| 18181000 | **1.58473231** | tr\|Q5PQZ9\|Q5PQZ9_RAT | **0.47049916** | 23190000 | **0.745615228** | NADH dehydrogenase [ubiquinone] 1 subunit C2 OS=Rattus norvegicus OX=10116 GN=Ndufc2 PE=1 SV=1 |
| 23602000 | **2.05724944** | tr\|D3ZG43\|D3ZG43_RAT | **0.29913622** | 19140000 | **0.615397821** | NADH dehydrogenase (Ubiquinone) Fe-S protein 3 (Predicted) isoform CRA_c OS=Rattus norvegicus OX=10116 GN=Ndufs3 PE=1 SV=1 |
| 8793300 | **0.76646096** | Q561S0\|NDUAA_RAT | **0.34456468** | 8213850 | **0.26409537** | NADH dehydrogenase [ubiquinone] 1 alpha subcomplex subunit 10 mitochondrial OS=Rattus norvegicus OX=10116 GN=Ndufa10 PE=1 SV=1 |
| 12726000 | **1.1092516** | tr\|D4A0T0\|D4A0T0_RAT | **0.37194464** | 12832000 | **0.41258019** | NADH:ubiquinone oxidoreductase subunit B10 OS=Rattus norvegicus OX=10116 GN=Ndufb10 PE=1 SV=1 |
| 8536000 | **0.74403361** | P11661\|NU5M_RAT | **0.40250106** | 9314200 | **0.299474315** | NADH-ubiquinone oxidoreductase chain 5 OS=Rattus norvegicus OX=10116 GN=Mtnd5 PE=3 SV=3 |
| 560640 | **0.04886774** | tr\|F1LPG5\|F1LPG5_RAT | **4.59813775** | 6988600 | **0.224700586** | NADH:ubiquinone oxidoreductase subunit B4 OS=Rattus norvegicus OX=10116 GN=Ndufb4 PE=1 SV=1 |
| | **0** | tr\|D3ZLT1\|D3ZLT1_RAT | **+** | 6726500 | **0.216273429** | NADH dehydrogenase (Ubiquinone) 1 beta subcomplex 7 (Predicted) OS=Rattus norvegicus OX=10116 GN=Ndufb7 PE=1 SV=1 |
| 262320 | **0.02286491** | tr\|D4A3V2\|D4A3V2_RAT | **4.02002028** | 2858800 | **0.091917413** | NADH dehydrogenase [ubiquinone] 1 alpha subcomplex subunit 6 OS=Rattus norvegicus OX=10116 GN=Ndufa6 PE=1 SV=1 |
| 1509700 | **0.13159179** | tr\|A9UMV9\|A9UMV9_RAT | **7.24623536** | 29657000 | **0.953545098** | NADH:ubiquinone oxidoreductase subunit A7 OS=Rattus norvegicus OX=10116 GN=Ndufa7 PE=1 SV=1 |
| 365080 | **0.03182191** | tr\|B2RYU0\|B2RYU0_RAT | **6.47668293** | 6410100 | **0.206100396** | NADH dehydrogenase (Ubiquinone) 1 beta subcomplex 2 (Predicted) isoform CRA_b OS=Rattus norvegicus OX=10116 GN=Ndufb2 PE=1 SV=1 |
| 2854700 | **0.24882764** | P03889\|NU1M_RAT | **0.33792499** | 2615200 | **0.084085077** | NADH-ubiquinone oxidoreductase chain 1 OS=Rattus norvegicus OX=10116 GN=Mtnd1 PE=1 SV=3 |
| 1791600 | **0.15616338** | Q63362\|NDUA5_RAT | **0.38229617** | 1856800 | **0.059700662** | NADH dehydrogenase [ubiquinone] 1 alpha subcomplex subunit 5 OS=Rattus norvegicus OX=10116 GN=Ndufa5 PE=1 SV=3 |
| 1041600 | **0.09079023** | tr\|B2RYW3\|B2RYW3_RAT | **0.41197099** | 1163300 | **0.037402941** | NADH dehydrogenase (Ubiquinone) 1 beta subcomplex 9 OS=Rattus norvegicus OX=10116 GN=Ndufb9 PE=1 SV=1 |
| 14275000 | **1.24426895** | tr\|D3ZF13\|D3ZF13_RAT | **0.40752944** | 15771000 | **0.507076229** | Acyl carrier protein OS=Rattus norvegicus OX=10116 GN=Ndufab1 PE=1 SV=1 |
| **Complex III** | | | | | | |
| 15525000 | **1.3532242** | P32551\|QCR2_RAT | **0.49449065** | 20812000 | **0.669156711** | Cytochrome b-c1 complex subunit 2 mitochondrial OS=Rattus norvegicus OX=10116 GN=Uqcrc2 PE=1 SV=2 |
| 997010 | **0.08690358** | P20788\|UCRI_RAT | **4.07642191** | 11018000 | **0.354255652** | Cytochrome b-c1 complex subunit Rieske mitochondrial OS=Rattus norvegicus OX=10116 GN=Uqcrfs1 PE=1 SV=2 |
| 10042000 | **0.87530289** | Q68FY0\|QCR1_RAT | **0.13157738** | 3582000 | **0.115170062** | Cytochrome b-c1 complex subunit 1 mitochondrial OS=Rattus norvegicus OX=10116 GN=Uqcrc1 PE=1 SV=1 |
| 3117800 | **0.27176054** | P00159\|CYB_RAT | **0.23157145** | 1957300 | **0.062931983** | Cytochrome b OS=Rattus norvegicus OX=10116 GN=Mt-Cyb PE=3 SV=3 |
| 314440 | **0.02740791** | Q5M9I5\|QCR6_RAT | **3.67687316** | 3134300 | **0.100775412** | Cytochrome b-c1 complex subunit 6 mitochondrial OS=Rattus norvegicus OX=10116 GN=Uqcrh PE=3 SV=1 |
| | **0** | Q7TQ16\|QCR8_RAT | **+** | 727260 | **0.023383188** | Cytochrome b-c1 complex subunit 8 OS=Rattus norvegicus OX=10116 GN=Uqcrq PE=3 SV=1 |
| | **0** | tr\|B2RYS2\|B2RYS2_RAT | **+** | 480600 | **0.015452466** | Cytochrome b-c1 complex subunit 7 OS=Rattus norvegicus OX=10116 GN=Uqcrb PE=1 SV=1 |
| **Complex IV** | | | | | | |
| 6358700 | **0.554251** | P00406\|COX2_RAT | **0.25843153** | 4454900 | **0.143235933** | Cytochrome c oxidase subunit 2 OS=Rattus norvegicus OX=10116 GN=Mtco2 PE=1 SV=3 |
| 321550 | **0.02802765** | P80432\|COX7C_RAT | **5.45421448** | 4754500 | **0.152868806** | Cytochrome c oxidase subunit 7C mitochondrial OS=Rattus norvegicus OX=10116 GN=Cox7c PE=1 SV=2 |
| 4296900 | **0.37453585** | P11240\|COX5A_RAT | **0.19054403** | 2219600 | **0.07136557** | Cytochrome c oxidase subunit 5A mitochondrial OS=Rattus norvegicus OX=10116 GN=Cox5a PE=1 SV=1 |
| 105120 | **0.0091627** | P05503\|COX1_RAT | **2.13936727** | 609670 | **0.019602382** | Cytochrome c oxidase subunit 1 OS=Rattus norvegicus OX=10116 GN=Mtco1 PE=2 SV=3 |
| 44810 | **0.00390583** | P35171\|CX7A2_RAT | **9.50621659** | 1154800 | **0.037129645** | Cytochrome c oxidase subunit 7A2 mitochondrial OS=Rattus norvegicus OX=10116 GN=Cox7a2 PE=1 SV=1 |
| 2010600 | **0.17525234** | P10888\|COX41_RAT | **0.15178137** | 827310 | **0.02660004** | Cytochrome c oxidase subunit 4 isoform 1 mitochondrial OS=Rattus norvegicus OX=10116 GN=Cox4i1 PE=1 SV=1 |
| 151450 | **0.01320102** | P05505\|COX3_RAT | **-** | 0 | **0** | Cytochrome c oxidase subunit 3 OS=Rattus norvegicus OX=10116 GN=Mtco3 PE=1 SV=5 |
| | **0** | P11951\|CX6C2_RAT | **+** | 427700 | **0.013751601** | Cytochrome c oxidase subunit 6C-2 OS=Rattus norvegicus OX=10116 GN=Cox6c2 PE=1 SV=3 |
| 92435 | **0.00805702** | P80431\|COX7B_RAT | **0.46638274** | 116870 | **0.003757656** | Cytochrome c oxidase subunit 7B mitochondrial OS=Rattus norvegicus OX=10116 GN=Cox7b PE=1 SV=3 |
| 46814 | **0.0040805** | tr\|B2RZD6\|B2RZD6_RAT | **8.5626815** | 1086700 | **0.034940063** | NDUFA4 mitochondrial complex-associated OS=Rattus norvegicus OX=10116 GN=Ndufa4 PE=1 SV=1 |
| **Complex V** | | | | | **0** | |
| 7095700 | **0.61849101** | Q06647\|ATPO_RAT | **2.16253686** | 41599000 | **1.33750961** | ATP synthase subunit O mitochondrial OS=Rattus norvegicus OX=10116 GN=Atp5po PE=1 SV=1 |
| | **0** | P29419\|ATP5I_RAT | **+** | 10020000 | **0.322167511** | ATP synthase subunit e mitochondrial OS=Rattus norvegicus OX=10116 GN=Atp5me PE=1 SV=3 |
| | **0** | D3ZAF6\|ATPK_RAT | **+** | 1883500 | **0.060559132** | ATP synthase subunit f mitochondrial OS=Rattus norvegicus OX=10116 GN=Atp5mf PE=1 SV=1 |

| | | | | | | |
|---|---|---|---|---|---|---|
| **Others** | | | | | | |
| 127850 | **0.01114394** | Q05962\|ADT1_RAT | **4.45070151** | 1542600 | **0.049598364** | ADP/ATP translocase 1 OS=Rattus norvegicus OX=10116 GN=Slc25a4 PE=1 SV=3 |
| 1180300 | **0.1028799** | Q64428\|ECHA_RAT | **0.19400556** | 620770 | **0.019959274** | Trifunctional enzyme subunit alpha mitochondrial OS=Rattus norvegicus OX=10116 GN=Hadha PE=1 SV=2 |
| 2626900 | **0.22897164** | Q5M934\|TRUB1_RAT | **2.39488165** | 17055000 | **0.54835997** | Probable tRNA pseudouridine synthase 1 OS=Rattus norvegicus OX=10116 GN=Trub1 PE=2 SV=1 |
| 2297500 | **0.20025975** | P02563\|MYH6_RAT | **0.23900026** | 1488600 | **0.047862131** | Myosin-6 OS=Rattus norvegicus OX=10116 GN=Myh6 PE=1 SV=2 |
| 3542500 | **0.30877918** | Q5XIH7\|PHB2_RAT | **0.18334794** | 1760800 | **0.056614027** | Prohibitin-2 OS=Rattus norvegicus OX=10116 GN=Phb2 PE=1 SV=1 |
| | **0** | B0BN56\|RT31_RAT | **+** | 244140 | **0.007849698** | 28S ribosomal protein S31 mitochondrial OS=Rattus norvegicus OX=10116 GN=Mrps31 PE=2 SV=1 |
| 649530 | **0.05661576** | Q9ER34\|ACON_RAT | **0.33645607** | 592450 | **0.019048717** | Aconitate hydratase mitochondrial OS=Rattus norvegicus OX=10116 GN=Aco2 PE=1 SV=2 |
| 1100400 | **0.09591549** | P17764\|THIL_RAT | **0.43276441** | 1291000 | **0.041508808** | Acetyl-CoA acetyltransferase mitochondrial OS=Rattus norvegicus OX=10116 GN=Acat1 PE=1 SV=1 |
| | **0** | P11530\|DMD_RAT | **+** | 2253100 | **0.072442677** | Dystrophin |
| | **0** | tr\|Q4G067\|Q4G067_RAT | **+** | 361430 | **0.011620859** | Mitochondrial ribosomal protein L44 OS=Rattus norvegicus OX=10116 GN=Mrpl44 PE=1 SV=1 |
| | **0** | Q9Z0V6\|PRDX3_RAT | **+** | 356470 | **0.011461383** | Thioredoxin-dependent peroxide reductase mitochondrial OS=Rattus norvegicus OX=10116 GN=Prdx3 PE=1 SV=2 |
| | **0** | tr\|D4A4B1\|D4A4B1_RAT | **+** | 139410 | **0.004482373** | Mitochondrial ribosomal protein L15 OS=Rattus norvegicus OX=10116 GN=Mrpl15 PE=1 SV=2 |
| | **0** | tr\|D3Z8Q5\|D3Z8Q5_RAT | **+** | 293930 | **0.009450569** | RCG25747 OS=Rattus norvegicus OX=10116 GN=Tusc2 PE=4 SV=1 |
| 1358900 | **0.11844743** | Q02253\|MMSA_RAT | **-** | | | Methylmalonate-semialdehyde dehydrogenase [acylating], mitochondrial OS=Rattus norvegicus OX=10116 GN=Aldh6a1 PE=1 SV=1 |
| 91401 | **0.0079669** | Q9R063\|PRDX5_RAT | **-** | | | Peroxiredoxin-5, mitochondrial OS=Rattus norvegicus OX=10116 GN=Prdx5 PE=1 SV=1 |
