## Supplementary material for "Permeability transition pore-related changes in the proteome and channel activity of ATP synthase dimers and monomers": ATP-synthase

| Protein Group | Protein ID | -10lgP | Coverage (%) | Coverage (%) Sample 1 | Area (IBAQ) | riBAQ | #Peptides | #Unique | #Spec Sample 1 | PTM | Avg. Mass | Accession | rIBAQ PTP/Control | Protein Group | Protein ID | -10lgP | Coverage (%) | Coverage (%) Sample 1 | Area (IBAQ) | riBAQ | #Peptides | #Unique | #Spec Sample 1 | PTM | Avg. Mass | |
|---|---|---|---|---|---|---|---|---|---|---|---|---|---|---|---|---|---|---|---|---|---|---|---|---|---|---|
| | | | | | CONTROL | % | | | | | | | | | | | | | PTP | % | | | | | | |
| **Complex III** | | | | | | | | | | | | | | | | | | | | | | | | | | |
| 17 | 34 | 63.27 | 8 | 8 | 1024500 | **0.044661948** | 4 | 4 | 6 | N | 48396 | P32551\|QCR2_RAT | **1.209733733** | 20 | 50 | 77.02 | 5 | 5 | 2756400 | **0.054029065** | 4 | 4 | 10 | | 29446 | Cytochrome b-c1 complex subunit 2 mitochondrial OS=Rattus norvegicus OX=10116 GN=Uqcrc2 PE=1 SV=2 |
| 75 | 132 | 74.5 | 5 | 5 | 276840 | **0.012068535** | 1 | 1 | 1 | N | 29446 | P20788\|UCRI_RAT | **2.104595548** | 29 | 58 | 126.55 | 5 | 5 | 1295800 | **0.025399384** | 2 | 2 | 4 | | 48396 | Cytochrome b-c1 complex subunit 1 mitochondrial OS=Rattus norvegicus OX=10116 GN=Uqcrc1 PE=1 SV=1 |
| | | | | | | **0** | | | | | | Q5M9I5\|QCR6_RAT | **+** | 32 | 318 | 34.77 | 17 | 17 | 864070 | **0.016936908** | 1 | 1 | 4 | | 52849 | Cytochrome b-c1 complex subunit 6 mitochondrial OS=Rattus norvegicus OX=10116 GN=Uqcrh PE=3 SV=1 |
| 35 | 118 | 37.65 | 4 | 4 | 60256 | **0.002626794** | 2 | 2 | 2 | N | 52849 | Q68FY0\|QCR1_RAT | **6.682207453** | 70 | 25 | 53.21 | 4 | 4 | 895490 | **0.017552782** | 2 | 2 | 2 | | 10424 | Cytochrome b (Fragment) OS=Rattus norvegicus OX=10116 GN=Cytb PE=3 SV=1 |
| 18 | 79 | 51.39 | 8 | 8 | 1491900 | **0.065037736** | 3 | 3 | 6 | N | 43012 | P00159\|CYB_RAT | **0.042247971** | 97 | 130 | 21.45 | 3 | 3 | 140180 | **0.002747712** | 1 | 1 | 1 | | 43012 | Cytochrome b OS=Rattus norvegicus OX=10116 PE=3 SV=1 |
| | | | | | | | | | | | | | | | | | | | | **0** | | | | | | |
| **Complex IV** | | | | | | | | | | | | | | | | | | | | **0** | | | | | | |
| 16 | 27 | 69.65 | 17 | 17 | 843160 | **0.036756631** | 5 | 5 | 6 | Y | 25928 | P00406\|COX2_RAT | **1.682795975** | 14 | 34 | 99.64 | 27 | 27 | 3155600 | **0.06185391** | 8 | 8 | 17 | Oxid | 25928 | Cytochrome c oxidase subunit 2 OS=Rattus norvegicus OX=10116 GN=Mtco2 PE=1 SV=3 |
| 21 | 133 | 93.16 | 5 | 5 | 1386800 | **0.060456017** | 2 | 2 | 6 | N | 35435 | tr\|D3ZFQ8\|D3ZFQ8_RAT | **0.91784682** | 17 | 59 | 142.67 | 13 | 13 | 2830900 | **0.055489363** | 5 | 5 | 12 | | 35435 | Cytochrome c-1 OS=Rattus norvegicus OX=10116 GN=Cyc1 PE=1 SV=3 |
| 19 | 86 | 82.98 | 16 | 16 | 571600 | **0.024918272** | 3 | 3 | 6 | N | 16130 | P11240\|COX5A_RAT | **0.726990201** | 21 | 200 | 118.98 | 23 | 23 | 924190 | **0.01811534** | 6 | 6 | 10 | | 16130 | Cytochrome c oxidase subunit 5A mitochondrial OS=Rattus norvegicus OX=10116 GN=Cox5a PE=1 SV=1 |
| 66 | 151 | 44370 | 12 | 12 | 214450 | **0.009348711** | 1 | 1 | 1 | N | 10487 | P10817\|CX6A2_RAT | **4.581469279** | 24 | 253 | 48.06 | 12 | 12 | 2185100 | **0.042830834** | 2 | 2 | 6 | | 10487 | Cytochrome c oxidase subunit 6A2 mitochondrial (Fragment) OS=Rattus norvegicus OX=10116 GN=Cox6a2 PE=1 SV=3 |
| 38 | 139 | 28.66 | 9 | 9 | 516610 | **0.022521044** | 1 | 1 | 2 | N | 13915 | P12075\|COX5B_RAT | **1.280989301** | 38 | 151 | 30.38 | 11 | 11 | 1471800 | **0.028849216** | 2 | 2 | 3 | | 13915 | Cytochrome c oxidase subunit 5B mitochondrial OS=Rattus norvegicus OX=10116 GN=Cox5b PE=1 SV=2 |
| | | | | | | **0** | | | | | | P35171\|CX7A2_RAT | **+** | 43 | 263 | 89.33 | 12 | 12 | 748420 | **0.014670016** | 2 | 2 | 3 | | 9353 | Cytochrome c oxidase subunit 7A2 mitochondrial OS=Rattus norvegicus OX=10116 GN=Cox7a2 PE=1 SV=1 |
| 76 | 134 | 57.04 | 21 | 21 | 152490 | **0.006647633** | 1 | 1 | 1 | N | 7375 | P80432\|COX7C_RAT | **2.639777535** | 52 | 251 | 78.41 | 21 | 21 | 895260 | **0.017548273** | 1 | 1 | 3 | | 7375 | Cytochrome c oxidase subunit 7C mitochondrial OS=Rattus norvegicus OX=10116 GN=Cox7c PE=1 SV=2 |
| | | | | | | | | | | | | P05505\|COX3_RAT | **+** | 77 | 270 | 67.76 | 3 | 3 | 413690 | **0.008108868** | 2 | 2 | 2 | | 29871 | Cytochrome c oxidase subunit 3 OS=Rattus norvegicus OX=10116 GN=Mtco3 PE=1 SV=5 |
| | | | | | | **0** | | | | | | tr\|B2RZD6\|B2RZD6_RAT | **+** | 47 | 209 | 37.86 | 10 | 10 | 119080 | **0.002334125** | 2 | 2 | 3 | | 9327 | NDUFA4 mitochondrial complex-associated OS=Rattus norvegicus OX=10116 GN=Ndufa4 PE=1 SV=1 |
| **Complex V** | | | | | | | | | | | | | | | | | | | | **0** | | | | | | |
| 2 | 1 | 387.5 | 60 | 60 | 1157500000 | **50.45993663** | 170 | 170 | 874 | Y | 56354 | P10719\|ATPB_RAT | **1.047113986** | 1 | 5 | 412.63 | 52 | 52 | 2695600000 | **52.83730538** | 151 | 148 | 1008 | Oxid | 56354 | ATP synthase subunit beta mitochondrial OS=Rattus norvegicus OX=10116 GN=Atp5f1b PE=1 SV=2 |
| 1 | 3 | 338.6 | 58 | 58 | 716370000 | **31.22936052** | 168 | 166 | 907 | Y | 59754 | P15999\|ATPA_RAT | **0.928241393** | 2 | 8 | 351.84 | 44 | 44 | 1478900000 | **28.98838512** | 147 | 143 | 897 | Carbamidomet | | ATP synthase subunit alpha, mitochondrial |
| 4 | 7 | 278.2 | 65 | 65 | 85852000 | **3.742623309** | 41 | 40 | 154 | Y | 18763 | P31399\|ATP5H_RAT | **1.061918967** | 3 | 65 | 317.64 | 66 | 66 | 202760000 | **3.974362679** | 47 | 46 | 200 | Oxidation (M) | | ATP synthase subunit d, mitochondrial |
| 3 | 5 | 280.5 | 43 | 43 | 102670000 | **4.475785481** | 46 | 46 | 221 | Y | 30191 | P35435\|ATPG_RAT | **0.657306168** | 4 | 63 | 297.5 | 48 | 48 | 150090000 | **2.941961405** | 47 | 47 | 172 | Carbamidomet | | ATP synthase subunit gamma, mitochondrial |
| 7 | 10 | 168 | 23 | 23 | 85675000 | **3.734907189** | 21 | 21 | 64 | Y | 28869 | P19511\|AT5F1_RAT | **1.325732607** | 5 | 168 | 208.61 | 32 | 32 | 252610000 | **4.951488244** | 32 | 31 | 128 | Form | 28869 | ATP synthase F(0) complex subunit B1 mitochondrial OS=Rattus norvegicus OX=10116 GN=Atp5pb PE=1 SV=1 |
| 5 | 8 | 159.9 | 38 | 38 | 30283000 | **1.320154005** | 21 | 21 | 71 | Y | 23398 | Q06647\|ATPO_RAT | **1.645279061** | 6 | 76 | 199.86 | 42 | 42 | 110810000 | **2.172021742** | 34 | 33 | 116 | Carbamidomet | | ATP synthase subunit O, mitochondrial |
| 6 | 11 | 171.3 | 45 | 45 | 22118000 | **0.96420983** | 19 | 19 | 67 | Y | 11433 | Q6PDU7\|ATP5L_RAT | **1.567052459** | 7 | 75 | 186.13 | 50 | 50 | 77085000 | **1.510967386** | 22 | 22 | 103 | Oxid | 11433 | ATP synthase subunit g mitochondrial OS=Rattus norvegicus OX=10116 GN=Atp5mg PE=1 SV=2 |
| 14 | 31 | 91.6 | 49 | 49 | 2316400 | **0.100980905** | 6 | 6 | 17 | N | 8255 | P29419\|ATP5I_RAT | **6.247984576** | 8 | 319 | 157.03 | 65 | 65 | 32188000 | **0.630927135** | 17 | 17 | 54 | Oxid | 8255 | ATP synthase subunit e mitochondrial OS=Rattus norvegicus OX=10116 GN=Atp5me PE=1 SV=3 |
| 11 | 15 | 90.42 | 27 | 27 | 3492600 | **0.152256047** | 8 | 8 | 22 | Y | 25076 | P05504\|ATP6_RAT | **1.37519168** | 10 | 258 | 74.9 | 18 | 18 | 10682000 | **0.209381249** | 7 | 7 | 28 | Oxid | 25076 | ATP synthase subunit a OS=Rattus norvegicus OX=10116 GN=Mt-atp6 PE=1 SV=3 |
| 10 | 30 | 159.2 | 39 | 39 | 4939200 | **0.21531898** | 10 | 10 | 25 | N | 12494 | P21571\|ATP5J_RAT | **0.696063008** | 11 | 12037 | 170.28 | 49 | 49 | 7646200 | **0.149875577** | 13 | 12 | 28 | Oxid | 12494 | ATP synthase-coupling factor 6 mitochondrial OS=Rattus norvegicus OX=10116 GN=Atp5pf PE=1 SV=1 |
| 8 | 12 | 162.9 | 33 | 33 | 18902000 | **0.824011855** | 13 | 13 | 47 | Y | 17563 | tr\|G3V7Y3\|G3V7Y3_RAT | **0.454915242** | 12 | 3953 | 105.59 | 19 | 19 | 19124000 | **0.374855553** | 9 | 9 | 25 | | 17563 | ATP synthase subunit delta mitochondrial OS=Rattus norvegicus OX=10116 GN=Atp5f1d PE=1 SV=1 |
| 9 | 32 | 134.4 | 48 | 48 | 45606000 | **1.9881433** | 8 | 8 | 32 | Y | 7642 | tr\|Q5UAJ5\|Q5UAJ5_RAT | **0.167240013** | 16 | 233 | 111.96 | 28 | 28 | 16963000 | **0.332497111** | 5 | 5 | 14 | Oxid | 7642 | ATP synthase protein 8 OS=Rattus norvegicus OX=10116 GN=ATP8 PE=3 SV=1 |
| 24 | 85 | 31.38 | 24 | 24 | 380940 | **0.016606659** | 3 | 3 | 3 | Y | 10452 | D3ZAF6\|ATPK_RAT | **4.671504581** | 19 | 12050 | 50.42 | 20 | 20 | 3957800 | **0.077578085** | 6 | 6 | 10 | Oxid | 10452 | ATP synthase subunit f mitochondrial OS=Rattus norvegicus OX=10116 GN=Atp5mf PE=1 SV=1 |
| 28 | 89 | 45.84 | 33 | 33 | 927040 | **0.040413287** | 2 | 2 | 3 | N | 6408 | Q9JJW3\|ATPMD_RAT | **2.749053666** | 22 | 12038 | 61.59 | 40 | 40 | 5667900 | **0.111098295** | 4 | 4 | 7 | | 6408 | ATP synthase membrane subunit DAPIT mitochondrial OS=Rattus norvegicus OX=10116 GN=Atp5md PE=1 SV=1 |
| | | | | | | | | | | | | D3Z9R8\|ATP68_RAT | **+** | 54 | 12095 | 22.95 | 8 | 8 | 2296500 | **0.04501442** | 1 | 1 | 3 | | 6914 | ATP synthase subunit ATP5MPL mitochondrial OS=Rattus norvegicus OX=10116 GN=Atp5mpl PE=1 SV=1 |
| | | | | | | | | | | | | | | | | | | | | **0** | | | | | | |
| **Others** | | | | | | | | | | | | | | | | | | | | **0** | | | | | | |
| 15 | 19 | 126.2 | 12 | 12 | 2748600 | **0.119822187** | 7 | 7 | 16 | Y | 47314 | P00507\|AATM_RAT | **1.226638451** | 9 | 45 | 175.16 | 17 | 17 | 7498400 | **0.146978502** | 13 | 13 | 32 | Oxid | 47314 | Aspartate aminotransferase mitochondrial OS=Rattus norvegicus OX=10116 GN=Got2 PE=1 SV=2 |
| 13 | 20 | 115.6 | 15 | 15 | 2834100 | **0.123549466** | 11 | 11 | 19 | N | 51414 | Q60587\|ECHB_RAT | **0.449285348** | 13 | 29 | 131.94 | 14 | 14 | 2831900 | **0.055508965** | 12 | 12 | 20 | | 51414 | Trifunctional enzyme subunit beta mitochondrial OS=Rattus norvegicus OX=10116 GN=Hadhb PE=1 SV=1 |
| 12 | 9 | 127.4 | 8 | 8 | 3637100 | **0.158555365** | 11 | 11 | 21 | N | 82665 | Q64428\|ECHA_RAT | **0.201557256** | 18 | 21 | 102.43 | 8 | 8 | 1630400 | **0.031957984** | 7 | 7 | 11 | | 82665 | Trifunctional enzyme subunit alpha mitochondrial OS=Rattus norvegicus OX=10116 GN=Hadha PE=1 SV=2 |
| 25 | 88 | 60.11 | 3 | 3 | 552340 | **0.024078653** | 2 | 2 | 3 | N | 85433 | Q9ER34\|ACON_RAT | **0.359388396** | 26 | 17 | 79.49 | 3 | 3 | 441480 | **0.008653589** | 3 | 3 | 5 | | 85433 | Aconitate hydratase mitochondrial OS=Rattus norvegicus OX=10116 GN=Aco2 PE=1 SV=2 |
| 34 | 84 | 63.37 | 6 | 6 | 412600 | **0.017986842** | 2 | 2 | 2 | Y | 47385 | P09605\|KCRS_RAT | **0.439739374** | 31 | 46 | 46.99 | 7 | 7 | 403520 | **0.007909523** | 3 | 3 | 4 | | 47385 | Creatine kinase S-type mitochondrial OS=Rattus norvegicus OX=10116 GN=Ckmt2 PE=1 SV=2 |
| 77 | 138 | 30.76 | 2 | 2 | 143510 | **0.00625616** | 1 | 1 | 1 | N | 88217 | Q63704\|CPT1B_RAT | **1.820845357** | 42 | 61 | 77.4 | 2 | 2 | 581160 | **0.0113915** | 2 | 2 | 3 | | 88217 | Carnitine O-palmitoyltransferase 1 muscle isoform OS=Rattus norvegicus OX=10116 GN=Cpt1b PE=1 SV=1 |
| | | | | | | **0** | | | | | | Q9Z0V6\|PRDX3_RAT | **+** | 51 | 316 | 77.98 | 8 | 8 | 974260 | **0.019096777** | 2 | 2 | 3 | | 28295 | Thioredoxin-dependent peroxide reductase mitochondrial OS=Rattus norvegicus OX=10116 GN=Prdx3 PE=1 SV=2 |
| | | | | | | **0** | | | | | | O55207\|SYNJ2_RAT | **+** | 55 | 647 | 29.35 | 1 | 1 | 142720 | **0.0027975** | 2 | 2 | 3 | | 165263 | Synaptojanin-2 OS=Rattus norvegicus OX=10116 GN=Synj2 PE=1 SV=2 |
| | | | | | | **0** | | | | | | P13086\|SUCA_RAT | **+** | 71 | 144 | 32.88 | 6 | 6 | 124000 | **0.002430563** | 2 | 2 | 2 | | 36148 | Succinate--CoA ligase [ADP/GDP-forming] subunit alpha mitochondrial OS=Rattus norvegicus OX=10116 GN=Suclg1 PE=2 SV=2 |
| | | | | | | **0** | | | | | | P45953\|ACADV_RAT | **+** | 72 | 31 | 51.63 | 3 | 3 | 354850 | **0.006955527** | 2 | 2 | 2 | | 70749 | Very long-chain specific acyl-CoA dehydrogenase mitochondrial OS=Rattus norvegicus OX=10116 GN=Acadvl PE=1 SV=1 |
| | | | | | | | | | | | | Q9R1Z0\|VDAC3_RAT | **+** | 41 | 158 | 31.48 | 8 | 8 | 1639400 | **0.032134396** | 2 | 2 | 3 | Form | 30798 | Voltage-dependent anion-selective channel protein 3 OS=Rattus norvegicus OX=10116 GN=Vdac3 PE=1 SV=2 |
| | | | | | | **100** | | | | | | | | | | | | | 5101698470 | **100** | | | | | | |
| | | | | | | | | | | | | | | | | | | | 51016984.7 | | | | | | | |
