## Supplementary material for "Permeability transition pore-related changes in the proteome and channel activity of ATP synthase dimers and monomers": ATP-synthase

| Area (IBAQ) | riBAQ | Accession | rIBAQ PTP/Control | Area (IBAQ) | riBAQ | |
|---|---|---|---|---|---|---|
| CONTROL | % | | | PTP | % | |
| **Complex III** | | | | | | |
| 276840 | **0.012068535** | P20788\|UCRI_RAT | **2.104595548** | 1295800 | **0.025399384** | Cytochrome b-c1 complex subunit 1 mitochondrial OS=Rattus norvegicus OX=10116 GN=Uqcrc1 PE=1 SV=1 |
| | **0** | Q5M9I5\|QCR6_RAT | **+** | 864070 | **0.016936908** | Cytochrome b-c1 complex subunit 6 mitochondrial OS=Rattus norvegicus OX=10116 GN=Uqcrh PE=3 SV=1 |
| 60256 | **0.002626794** | Q68FY0\|QCR1_RAT | **6.682207453** | 895490 | **0.017552782** | Cytochrome b (Fragment) OS=Rattus norvegicus OX=10116 GN=Cytb PE=3 SV=1 |
| 1491900 | **0.065037736** | P00159\|CYB_RAT | **0.042247971** | 140180 | **0.002747712** | Cytochrome b OS=Rattus norvegicus OX=10116 PE=3 SV=1 |
| | | | | | **0** | |
| **Complex IV** | | | | | **0** | |
| 214450 | **0.009348711** | P10817\|CX6A2_RAT | **4.581469279** | 2185100 | **0.042830834** | Cytochrome c oxidase subunit 6A2 mitochondrial (Fragment) OS=Rattus norvegicus OX=10116 GN=Cox6a2 PE=1 SV=3 |
| | **0** | P35171\|CX7A2_RAT | **+** | 748420 | **0.014670016** | Cytochrome c oxidase subunit 7A2 mitochondrial OS=Rattus norvegicus OX=10116 GN=Cox7a2 PE=1 SV=1 |
| 152490 | **0.006647633** | P80432\|COX7C_RAT | **2.639777535** | 895260 | **0.017548273** | Cytochrome c oxidase subunit 7C mitochondrial OS=Rattus norvegicus OX=10116 GN=Cox7c PE=1 SV=2 |
| | | P05505\|COX3_RAT | **+** | 413690 | **0.008108868** | Cytochrome c oxidase subunit 3 OS=Rattus norvegicus OX=10116 GN=Mtco3 PE=1 SV=5 |
| | **0** | tr\|B2RZD6\|B2RZD6_RAT | **+** | 119080 | **0.002334125** | NDUFA4 mitochondrial complex-associated OS=Rattus norvegicus OX=10116 GN=Ndufa4 PE=1 SV=1 |
| **Complex V** | | | | | **0** | |
| 2316400 | **0.100980905** | P29419\|ATP5I_RAT | **6.247984576** | 32188000 | **0.630927135** | ATP synthase subunit e mitochondrial OS=Rattus norvegicus OX=10116 GN=Atp5me PE=1 SV=3 |
| 18902000 | **0.824011855** | tr\|G3V7Y3\|G3V7Y3_RAT | **0.454915242** | 19124000 | **0.374855553** | ATP synthase subunit delta mitochondrial OS=Rattus norvegicus OX=10116 GN=Atp5f1d PE=1 SV=1 |
| 45606000 | **1.9881433** | tr\|Q5UAJ5\|Q5UAJ5_RAT | **0.167240013** | 16963000 | **0.332497111** | ATP synthase protein 8 OS=Rattus norvegicus OX=10116 GN=ATP8 PE=3 SV=1 |
| 380940 | **0.016606659** | D3ZAF6\|ATPK_RAT | **4.671504581** | 3957800 | **0.077578085** | ATP synthase subunit f mitochondrial OS=Rattus norvegicus OX=10116 GN=Atp5mf PE=1 SV=1 |
| 927040 | **0.040413287** | Q9JJW3\|ATPMD_RAT | **2.749053666** | 5667900 | **0.111098295** | ATP synthase membrane subunit DAPIT mitochondrial OS=Rattus norvegicus OX=10116 GN=Atp5md PE=1 SV=1 |
| | | D3Z9R8\|ATP68_RAT | **+** | 2296500 | **0.04501442** | ATP synthase subunit ATP5MPL mitochondrial OS=Rattus norvegicus OX=10116 GN=Atp5mpl PE=1 SV=1 |
| | | | | | **0** | |
| **Others** | | | | | **0** | |
| 2834100 | **0.123549466** | Q60587\|ECHB_RAT | **0.449285348** | 2831900 | **0.055508965** | Trifunctional enzyme subunit beta mitochondrial OS=Rattus norvegicus OX=10116 GN=Hadhb PE=1 SV=1 |
| 3637100 | **0.158555365** | Q64428\|ECHA_RAT | **0.201557256** | 1630400 | **0.031957984** | Trifunctional enzyme subunit alpha mitochondrial OS=Rattus norvegicus OX=10116 GN=Hadha PE=1 SV=2 |
| 552340 | **0.024078653** | Q9ER34\|ACON_RAT | **0.359388396** | 441480 | **0.008653589** | Aconitate hydratase mitochondrial OS=Rattus norvegicus OX=10116 GN=Aco2 PE=1 SV=2 |
| 412600 | **0.017986842** | P09605\|KCRS_RAT | **0.439739374** | 403520 | **0.007909523** | Creatine kinase S-type mitochondrial OS=Rattus norvegicus OX=10116 GN=Ckmt2 PE=1 SV=2 |
| | **0** | Q9Z0V6\|PRDX3_RAT | **+** | 974260 | **0.019096777** | Thioredoxin-dependent peroxide reductase mitochondrial OS=Rattus norvegicus OX=10116 GN=Prdx3 PE=1 SV=2 |
| | **0** | O55207\|SYNJ2_RAT | **+** | 142720 | **0.0027975** | Synaptojanin-2 OS=Rattus norvegicus OX=10116 GN=Synj2 PE=1 SV=2 |
| | **0** | P13086\|SUCA_RAT | **+** | 124000 | **0.002430563** | Succinate--CoA ligase [ADP/GDP-forming] subunit alpha mitochondrial OS=Rattus norvegicus OX=10116 GN=Suclg1 PE=2 SV=2 |
| | **0** | P45953\|ACADV_RAT | **+** | 354850 | **0.006955527** | Very long-chain specific acyl-CoA dehydrogenase mitochondrial OS=Rattus norvegicus OX=10116 GN=Acadvl PE=1 SV=1 |
| | | Q9R1Z0\|VDAC3_RAT | **+** | 1639400 | **0.032134396** | Voltage-dependent anion-selective channel protein 3 OS=Rattus norvegicus OX=10116 GN=Vdac3 PE=1 SV=2 |
