## Supplementary material for "Permeability transition pore-related changes in the proteome and channel activity of ATP synthase dimers and monomers": ATP-synthase

| Protein Group | Protein ID | Score (%) | -10lgP | Coverage (%) | #Peptides | #Unique | Area (IBAQ) | riBAQ | PTM | Avg. Mass | Accession | rIBAQ PTP/Control | Protein Group | Protein ID | Score (%) | -10lgP | Coverage (%) | #Peptides | #Unique | Area (IBAQ) | riBAQ | PTM | Avg. Mass | Description |
|---|---|---|---|---|---|---|---|---|---|---|---|---|---|---|---|---|---|---|---|---|---|---|---|---|
| | | | | | | | CONTROL | % | | | | | | | | | | | | PTP | % | | | |
| **Complex I** | | | | | | | | | | | | | | | | | | | | | | | | |
| 15 | 25 | 98.9 | 158.7 | 22 | 5 | 5 | 2680900 | 0.27353 | Y | 30226 | tr\|D3ZG43\|D3ZG43_RAT | 1.055695468 | 40 | 36 | 98.9 | 185 | 21 | 4 | 4 | 14115000 | 0.288764702 | N | 30226 | NADH dehydrogenase (Ubiquinone) Fe-S protein 3 (Predicted), isoform CRA_c OS=Rattus norvegicus OX=10116 GN=Ndufs3 PE=1 SV=1 |
| 14 | 6 | 99.1 | 155.91 | 13 | 7 | 7 | 4688500 | 0.47836 | N | 79412 | Q66HF1\|NDUS1_RAT | 1.388716218 | 19 | 12 | 99.2 | 242 | 22 | 13 | 13 | 32472000 | 0.664312251 | N | 79412 | NADH-ubiquinone oxidoreductase 75 kDa subunit, mitochondrial OS=Rattus norvegicus OX=10116 GN=Ndufs1 PE=1 SV=1 |
| 17 | 15 | 98.7 | 121.86 | 16 | 4 | 4 | 701120 | 0.07153 | Y | 42559 | Q5BK63\|NDUA9_RAT | 2.41533037 | 22 | 14 | 99.1 | 196 | 27 | 9 | 9 | 8445600 | 0.172780104 | Y | 42559 | NADH dehydrogenase [ubiquinone] 1 alpha subcomplex subunit 9, mitochondrial OS=Rattus norvegicus OX=10116 GN=Ndufa9 PE=1 SV=2 |
| 20 | 33 | 97.9 | 102.64 | 17 | 4 | 4 | 1898400 | 0.19369 | Y | 40493 | Q561S0\|NDUAA_RAT | 0.123650616 | 58 | 23 | 98.3 | 128 | 15 | 3 | 3 | 1170700 | 0.023950183 | N | 40493 | NADH dehydrogenase [ubiquinone] 1 alpha subcomplex subunit 10, mitochondrial OS=Rattus norvegicus OX=10116 GN=Ndufa10 PE=1 SV=1 |
| 26 | 23 | 97.3 | 73.84 | 10 | 3 | 3 | 146380 | 0.01494 | N | 50731 | tr\|Q5XIH3\|Q5XIH3_RAT | 6.473804099 | 26 | 21 | 99.1 | 151 | 18 | 7 | 7 | 4726100 | 0.096686565 | N | 50731 | NADH dehydrogenase [ubiquinone] flavoprotein 1, mitochondrial OS=Rattus norvegicus OX=10116 GN=Ndufv1 PE=1 SV=1 |
| 95 | 95 | 61.7 | 67.98 | 22 | 1 | 1 | 0 | 0.00000 | N | 13412 | Q63362\|NDUA5_RAT | + | 196 | 170 | 61.7 | 98 | 22 | 1 | 1 | 0 | 0 | N | 13412 | NADH dehydrogenase [ubiquinone] 1 alpha subcomplex subunit 5 OS=Rattus norvegicus OX=10116 GN=Ndufa5 PE=1 SV=3 |
| 96 | 42 | 61.6 | 50.27 | 12 | 1 | 1 | 69212 | 0.00706 | N | 21664 | tr\|D4A565\|D4A565_RAT | 12.29279085 | 44 | 93 | 98.6 | 167 | 28 | 4 | 4 | 4243200 | 0.086807395 | N | 21664 | NADH dehydrogenase (Ubiquinone) 1 beta subcomplex, 5 (Predicted), isoform CRA_b OS=Rattus norvegicus OX=10116 GN=Ndufb5 PE=1 SV=1 |
| 97 | 29 | 61.3 | 41.71 | 8 | 1 | 1 | 282740 | 0.02885 | N | 20859 | tr\|D4A0T0\|D4A0T0_RAT | 4.635994455 | 34 | 54 | 98.6 | 201 | 28 | 5 | 5 | 6537200 | 0.133738053 | Y | 20859 | NADH:ubiquinone oxidoreductase subunit B10 OS=Rattus norvegicus OX=10116 GN=Ndufb10 PE=1 SV=1 |
| 99 | 34 | 60 | 34.81 | 6 | 1 | 1 | 196860 | 0.02009 | N | 23945 | tr\|Q5RJN0\|Q5RJN0_RAT | 2.396029051 | 62 | 67 | 83.1 | 78 | 11 | 2 | 2 | 2352400 | 0.048125405 | N | 23945 | NADH dehydrogenase (Ubiquinone) Fe-S protein 7 OS=Rattus norvegicus OX=10116 GN=Ndufs7 PE=1 SV=1 |
| 100 | 22 | 58.1 | 33.1 | 5 | 1 | 1 | 2019900 | 0.20609 | N | 27378 | P19234\|NDUV2_RAT | 1.470057267 | 33 | 58 | 98.8 | 127 | 24 | 4 | 4 | 14809000 | 0.302962556 | N | 27378 | NADH dehydrogenase [ubiquinone] flavoprotein 2, mitochondrial OS=Rattus norvegicus OX=10116 GN=Ndufv2 PE=1 SV=2 |
| 119 | 52 | 33.3 | 21.84 | 4 | 1 | 1 | 0 | 0.00000 | N | 23970 | tr\|B0BNE6\|B0BNE6_RAT | + | 87 | 60 | 77.1 | 44 | 8 | 2 | 2 | 129890 | 0.00265729 | Y | 23970 | NADH dehydrogenase (Ubiquinone) Fe-S protein 8 (Predicted), isoform CRA_a OS=Rattus norvegicus OX=10116 GN=Ndufs8 PE=1 SV=1 |
| 110 | 38 | 40.7 | 23.69 | 8 | 1 | 1 | 134410 | 0.01371 | N | 15224 | tr\|D4A3V2\|D4A3V2_RAT | 0.926847242 | 199 | 88 | 61.3 | 42 | 8 | 1 | 1 | 621300 | 0.012710557 | N | 15224 | NADH dehydrogenase [ubiquinone] 1 alpha subcomplex subunit 6 OS=Rattus norvegicus OX=10116 GN=Ndufa6 PE=1 SV=1 |
| | | | | | | | | 0.00000 | | | tr\|B2RYS8\|B2RYS8_RAT | + | 85 | 73 | 84.3 | 106 | 15 | 2 | 2 | 1986100 | 0.040631638 | N | 21959 | NADH dehydrogenase [ubiquinone] 1 beta subcomplex subunit 8, mitochondrial OS=Rattus norvegicus OX=10116 GN=Ndufb8 PE=1 SV=1 |
| | | | | | | | | 0.00000 | | | tr\|Q5PQZ9\|Q5PQZ9_RAT | + | 64 | 112 | 76.1 | 97 | 21 | 3 | 3 | 638550 | 0.013063457 | N | 14359 | NADH dehydrogenase [ubiquinone] 1 subunit C2 OS=Rattus norvegicus OX=10116 GN=Ndufc2 PE=1 SV=1 |
| | | | | | | | | 0.00000 | | | tr\|F1LPG5\|F1LPG5_RAT | + | 84 | 71 | 84.4 | 83 | 29 | 2 | 2 | 1846400 | 0.037773655 | N | 15064 | NADH:ubiquinone oxidoreductase subunit B4 OS=Rattus norvegicus OX=10116 GN=Ndufb4 PE=1 SV=1 |
| | | | | | | | | 0.00000 | | | tr\|A9UMV9\|A9UMV9_RA | + | 152 | 104 | 68.9 | 61 | 11 | 1 | 1 | 407550 | 0.008337659 | N | 12500 | NADH:ubiquinone oxidoreductase subunit A7 OS=Rattus norvegicus OX=10116 GN=Ndufa7 PE=1 SV=1 |
| | | | | | | | | 0.00000 | | | tr\|B2RYW3\|B2RYW3_RAT | + | 89 | 113 | 73.5 | 44 | 11 | 2 | 2 | 1696700 | 0.034711093 | N | 21892 | NADH dehydrogenase (Ubiquinone) 1 beta subcomplex, 9 OS=Rattus norvegicus OX=10116 GN=Ndufb9 PE=1 SV=1 |
| | | | | | | | | 0.00000 | | | Q641Y2\|NDUS2_RAT | + | 71 | 24 | 94.1 | 109 | 5 | 2 | 2 | 4160900 | 0.085123702 | N | 52562 | NADH dehydrogenase [ubiquinone] iron-sulfur protein 2, mitochondrial OS=Rattus norvegicus OX=10116 GN=Ndufs2 PE=1 SV=1 |
| | | | | | | | | 0.00000 | | | tr\|D3ZE15\|D3ZE15_RAT | + | 91 | 100 | 61.3 | 55 | 7 | 2 | 2 | 1861700 | 0.038086663 | N | 16777 | NADH:ubiquinone oxidoreductase subunit A13 OS=Rattus norvegicus OX=10116 GN=Ndufa13 PE=1 SV=1 |
| | | | | | | | | 0.00000 | | | P52504\|NDUS6_RAT | + | 151 | 178 | 61.7 | 59 | 9 | 1 | 1 | 865090 | 0.017698013 | N | 12783 | NADH dehydrogenase [ubiquinone] iron-sulfur protein 6, mitochondrial OS=Rattus norvegicus OX=10116 GN=Ndufs6 PE=3 SV=1 |
| | | | | | | | | 0.00000 | | | tr\|D3ZF13\|D3ZF13_RAT | + | 56 | 105 | 92.8 | 118 | 15 | 3 | 3 | 3263000 | 0.066754462 | N | 17514 | Acyl carrier protein OS=Rattus norvegicus OX=10116 GN=Ndufab1 PE=1 SV=1 |
| **Complex III** | | | | | | | | | | | | | | | | | | | | | | | | |
| 19 | 1 | 98.6 | 125.75 | 17 | 5 | 5 | 2291000 | 0.23375 | N | 48396 | P32551\|QCR2_RAT | - | 83 | 1 | 84.4 | 105 | 6 | 2 | 2 | 0 | 0 | N | 48396 | Cytochrome b-c1 complex subunit 2, mitochondrial OS=Rattus norvegicus OX=10116 GN=Uqcrc2 PE=1 SV=2 |
| 35 | 12 | 61.7 | 85.46 | 5 | 2 | 2 | 43626 | 0.00445 | N | 52849 | Q68FY0\|QCR1_RAT | - | 154 | 15 | 64.2 | 43 | 5 | 1 | 1 | 0 | 0 | N | 52849 | Cytochrome b-c1 complex subunit 1, mitochondrial OS=Rattus norvegicus OX=10116 GN=Uqcrc1 PE=1 SV=1 |
| **Complex IV** | | | | | | | | | | | | | | | | | | | | | | | | |
| 36 | 109 | 61.6 | 49.51 | 21 | 1 | 1 | 29933 | 0.00305 | N | 16130 | P11240\|COX5A_RAT | - | 197 | 158 | 61.7 | 63 | 17 | 1 | 1 | 0 | 0 | N | 16130 | Cytochrome c oxidase subunit 5A, mitochondrial OS=Rattus norvegicus OX=10116 GN=Cox5a PE=1 SV=1 |
| 101 | 158 | 57.7 | 32.32 | 14 | 1 | 1 | 493200 | 0.05032 | N | 13915 | P12075\|COX5B_RAT | 0.736752211 | 201 | 133 | 60.2 | 37 | 14 | 1 | 1 | 1812200 | 0.037073992 | N | 13915 | Cytochrome c oxidase subunit 5B, mitochondrial OS=Rattus norvegicus OX=10116 GN=Cox5b PE=1 SV=2 |
| 111 | 51 | 40.5 | 43914 | 4 | 1 | 1 | 0 | 0.00000 | N | 19515 | P10888\|COX41_RAT | + | 42 | 50 | 98.8 | 126 | 24 | 4 | 4 | 4134200 | 0.084577473 | N | 19515 | Cytochrome c oxidase subunit 4 isoform 1, mitochondrial OS=Rattus norvegicus OX=10116 GN=Cox4i1 PE=1 SV=1 |
| | | | | | | | | 0.00000 | | | P00406\|COX2_RAT | + | 86 | 81 | 77.3 | 44 | 7 | 2 | 2 | 2403600 | 0.049172854 | Y | 25928 | Cytochrome c oxidase subunit 2 OS=Rattus norvegicus OX=10116 GN=Mtco2 PE=1 SV=3 |
| **Complex V** | | | | | | | | | | | | | | | | | | | | | | | | |
| 2 | 14 | 99.2 | 382.48 | 66 | 51 | 51 | 277360000 | 28.29884 | Y | 59754 | P15999\|ATPA_RAT | 1.02301427 | 2 | 13 | 99.2 | 460 | 70 | 75 | 75 | 1415100000 | 28.95011905 | Y | 59754 | ATP synthase subunit alpha, mitochondrial OS=Rattus norvegicus OX=10116 GN=Atp5f1a PE=1 SV=2 |
| 1 | 7 | 99.2 | 366.27 | 86 | 50 | 50 | 369900000 | 37.74063 | Y | 56354 | P10719\|ATPB_RAT | 0.914902977 | 1 | 8 | 99.2 | 450 | 81 | 64 | 64 | 1687800000 | 34.52901628 | Y | 56354 | ATP synthase subunit beta, mitochondrial OS=Rattus norvegicus OX=10116 GN=Atp5f1b PE=1 SV=2 |
| 6 | 116 | 99.2 | 229.61 | 57 | 16 | 1 | 513120 | 0.05235 | Y | 30191 | P35435\|ATPG_RAT | 6.392971174 | 7 | 97 | 99.2 | 284 | 65 | 19 | 1 | 16360000 | 0.334692918 | Y | 30191 | ATP synthase subunit gamma, mitochondrial OS=Rattus norvegicus OX=10116 GN=Atp5f1c PE=1 SV=2 |
| 10 | 4260 | 99.2 | 218.85 | 74 | 12 | 12 | 14806000 | 1.51065 | Y | 18763 | P31399\|ATP5H_RAT | 1.576895236 | 13 | 204 | 99.2 | 267 | 80 | 16 | 16 | 116440000 | 2.382129788 | Y | 18763 | ATP synthase subunit d, mitochondrial OS=Rattus norvegicus OX=10116 GN=Atp5pd PE=1 SV=3 |
| 8 | 41 | 99.1 | 208.5 | 46 | 13 | 12 | 44029000 | 4.49225 | N | 28869 | P19511\|AT5F1_RAT | 1.142388107 | 9 | 114 | 99.2 | 275 | 55 | 19 | 19 | 250850000 | 5.131889877 | Y | 28869 | ATP synthase F(0) complex subunit B1, mitochondrial OS=Rattus norvegicus OX=10116 GN=Atp5pb PE=1 SV=1 |
| 11 | 2163 | 99.1 | 158.27 | 49 | 10 | 10 | 17943000 | 1.83071 | Y | 23398 | Q06647\|ATPO_RAT | 2.434785456 | 12 | 188 | 99.2 | 246 | 65 | 22 | 22 | 217880000 | 4.457389541 | Y | 23398 | ATP synthase subunit O, mitochondrial OS=Rattus norvegicus OX=10116 GN=Atp5po PE=1 SV=1 |
| 13 | 4261 | 99.1 | 146.01 | 61 | 7 | 7 | 24198000 | 2.46890 | Y | 17563 | tr\|G3V7Y3\|G3V7Y3_RAT | 0.634396593 | 17 | 168 | 99.1 | 259 | 61 | 11 | 11 | 76560000 | 1.566264656 | Y | 17563 | ATP synthase subunit delta, mitochondrial OS=Rattus norvegicus OX=10116 GN=Atp5f1d PE=1 SV=1 |
| 22 | 4263 | 98.4 | 127.48 | 46 | 3 | 3 | 1928000 | 0.19671 | Y | 11433 | Q6PDU7\|ATP5L_RAT | 1.144514784 | 35 | 4490 | 84.4 | 133 | 34 | 2 | 2 | 11005000 | 0.225140315 | Y | 11433 | ATP synthase subunit g, mitochondrial OS=Rattus norvegicus OX=10116 GN=Atp5mg PE=1 SV=2 |
| 29 | 4270 | 84.2 | 126.91 | 38 | 3 | 3 | 2369300 | 0.24174 | N | 6408 | Q9JJW3\|ATPMD_RAT | 0.872861749 | 90 | 4499 | 61.7 | 140 | 28 | 2 | 2 | 10314000 | 0.211003836 | N | 6408 | ATP synthase membrane subunit DAPIT, mitochondrial OS=Rattus norvegicus OX=10116 GN=Atp5md PE=1 SV=1 |
| 21 | 4264 | 98.4 | 99.48 | 46 | 3 | 3 | 5060900 | 0.51636 | N | 7642 | tr\|Q5UAJ5\|Q5UAJ5_RAT | 1.066164778 | 45 | 4487 | 98.4 | 105 | 46 | 3 | 3 | 26910000 | 0.550524842 | N | 7642 | ATP synthase protein 8 OS=Rattus norvegicus OX=10116 GN=ATP8 PE=3 SV=1 |
| 28 | 4266 | 96.2 | 52.4 | 30 | 3 | 3 | 3083900 | 0.31465 | Y | 10452 | D3ZAF6\|ATPK_RAT | 1.629367949 | 25 | 4488 | 97.5 | 95 | 40 | 7 | 7 | 25060000 | 0.512677538 | Y | 10452 | ATP synthase subunit f, mitochondrial OS=Rattus norvegicus OX=10116 GN=Atp5mf PE=1 SV=1 |
| 98 | 4273 | 60.9 | 39.16 | 16 | 1 | 1 | 437370 | 0.04462 | N | 5767 | P29418\|ATP5E_RAT | 2.457919238 | 59 | 4489 | 95.4 | 79 | 41 | 3 | 3 | 5361400 | 0.109683534 | N | 5767 | ATP synthase subunit epsilon, mitochondrial OS=Rattus norvegicus OX=10116 GN=Atp5f1e PE=1 SV=2 |
| 105 | 2187 | 41.7 | 25.94 | 5 | 1 | 1 | 3196100 | 0.32610 | N | 14244 | Q06645\|AT5G1_RAT | 0.706472057 | 92 | 4509 | 38.3 | 24 | 5 | 1 | 1 | 11261000 | 0.230377564 | N | 14244 | ATP synthase F(0) complex subunit C1, mitochondrial OS=Rattus norvegicus OX=10116 GN=Atp5mc1 PE=1 SV=1 |
| **5** | 117 | | 221.49 | 25 | 16 | 1 | 2453300 | 0.25031 | Y | 67721 | tr\|Q6QI09\|Q6QI09_RAT | 1.185183736 | 11 | 95 | 99.2 | 278 | 30 | 20 | 2 | 14501000 | 0.296661491 | Y | 67721 | ATP synthase subunit gamma, mitochondrial OS=Rattus norvegicus OX=10116 GN=Taf3 PE=1 SV=1 |
| | | | | | | | | 0.00000 | | | D3Z9R8\|ATP68_RAT | + | 61 | 4494 | 84.3 | 123 | 32 | 3 | 3 | 4396500 | 0.089943607 | Y | 6914 | ATP synthase subunit ATP5MPL, mitochondrial OS=Rattus norvegicus OX=10116 GN=Atp5mpl PE=1 SV=1 |
| | | | | | | | | 0.00000 | | | P29419\|ATP5I_RAT | + | 43 | 4486 | 98.6 | 101 | 52 | 4 | 4 | 15392000 | 0.314889571 | N | 8255 | ATP synthase subunit e, mitochondrial OS=Rattus norvegicus OX=10116 GN=Atp5me PE=1 SV=3 |
| | | | | | | | | 0.00000 | | | P21571\|ATP5J_RAT | + | 63 | 4496 | 81.8 | 99 | 28 | 2 | 2 | 830950 | 0.016999577 | N | 12494 | ATP synthase-coupling factor 6, mitochondrial OS=Rattus norvegicus OX=10116 GN=Atp5pf PE=1 SV=1 |
| **Other** | | | | | | | | | | | | | | | | | | | | | | | | |
| R | 4 | 99.2 | 275.36 | 86 | 23 | 23 | 71777000 | 7.32336 | N | 29820 | P67779\|PHB_RAT | 0.834454861 | 5 | 16 | 99.2 | 293 | 85 | 30 | 29 | 298710000 | 6.111009867 | Y | 29820 | Prohibitin OS=Rattus norvegicus OX=10116 GN=Phb PE=1 SV=1 |
| 3 | 11 | 99.2 | 267.91 | 73 | 28 | 28 | 66230000 | 6.75740 | Y | 33312 | Q5XIH7\|PHB2_RAT | 0.013332791 | 3 | 18 | 99.2 | 275 | 75 | 30 | 1 | 4403900 | 0.090094996 | Y | 33312 | Prohibitin-2 OS=Rattus norvegicus OX=10116 GN=Phb2 PE=1 SV=1 |
| 7 | 2 | 99.2 | 242.74 | 48 | 21 | 21 | 23705000 | 2.41860 | Y | 57808 | Q02253\|MMSA_RAT | 0.029256602 | 55 | 64 | 98.4 | 153 | 10 | 3 | 3 | 3458800 | 0.070760138 | N | 57808 | Methylmalonate-semialdehyde dehydrogenase [acylating], mitochondrial OS=Rattus norvegicus OX=10116 GN=Aldh6a1 PE=1 SV=1 |
| 9 | 9 | 99.2 | 202.5 | 40 | 16 | 16 | 23694000 | 2.41748 | Y | 47314 | P00507\|AATM_RAT | 0.704141384 | 15 | 4 | 99.2 | 239 | 39 | 18 | 18 | 83207000 | 1.702248997 | Y | 47314 | Aspartate aminotransferase, mitochondrial OS=Rattus norvegicus OX=10116 GN=Got2 PE=1 SV=2 |
| 12 | 13 | 99.1 | 170.01 | 6 | 9 | 9 | 1524800 | 0.15557 | Y | 164579 | P07756\|CPSM_RAT | 14.00474205 | 6 | 3 | 99.2 | 356 | 32 | 37 | 37 | 106500000 | 2.178777245 | Y | 2E+05 | Carbamoyl-phosphate synthase [ammonia], mitochondrial OS=Rattus norvegicus OX=10116 GN=Cps1 PE=1 SV=1 |
| 16 | 21 | 98.6 | 164.25 | 10 | 6 | 6 | 4189000 | 0.42740 | N | 82665 | Q64428\|ECHA_RAT | 1.741799454 | 10 | 5 | 99.2 | 315 | 36 | 26 | 25 | 36389000 | 0.744446246 | Y | 82665 | Trifunctional enzyme subunit alpha, mitochondrial OS=Rattus norvegicus OX=10116 GN=Hadha PE=1 SV=2 |
| 24 | 19 | 93.3 | 63.14 | 4 | 2 | 2 | 1662400 | 0.16961 | N | 56912 | P22791\|HMCS2_RAT | 10.50982525 | 8 | 10 | 99.2 | 330 | 44 | 27 | 27 | 87135000 | 1.78260803 | Y | 56912 | Hydroxymethylglutaryl-CoA synthase, mitochondrial OS=Rattus norvegicus OX=10116 GN=Hmgcs2 PE=1 SV=1 |
| 18 | 179 | 80.3 | 36.06 | 2 | 3 | 2 | 2036300 | 0.20776 | Y | 140418 | F1M775\|DIAP1_RAT | 2.133021238 | 31 | 539 | 61.2 | 37 | 2 | 3 | 3 | 21662000 | 0.443161246 | Y | 1E+05 | Protein diaphanous homolog 1 OS=Rattus norvegicus OX=10116 GN=Diaph1 PE=1 SV=3 |
| 113 | 4292 | 34.3 | 22.71 | 7 | 1 | 1 | 2025700 | 0.20668 | Y | 40772 | B0K035\|MTFR2_RAT | - | | | | | | | | | 0 | | | Mitochondrial fission regulator 2 OS=Rattus norvegicus OX=10116 GN=Mtfr2 PE=2 SV=1 |
| **47** | 53 | | 32.92 | 2 | 1 | 1 | 312410 | 0.03187 | N | 129777 | P52873\|PYC_RAT | 21.83023454 | 14 | 6 | 99.2 | 258 | 20 | 20 | 20 | 34013000 | 0.695838032 | Y | 1E+05 | Pyruvate carboxylase, mitochondrial OS=Rattus norvegicus OX=10116 GN=Pc PE=1 SV=2 |
| | | | | | | | | 0.00000 | | | P10860\|DHE3_RAT | + | 16 | 2 | 99.2 | 320 | 36 | 19 | 19 | 61338000 | 1.254852945 | Y | 61416 | Glutamate dehydrogenase 1, mitochondrial OS=Rattus norvegicus OX=10116 GN=Glud1 PE=1 SV=2 |
| | | | | | | | | 0.00000 | | | tr\|A0A0G2KB63\|A0A0G2K | + | 4 | 19 | 99.2 | 274 | 76 | 30 | 1 | 2528600 | 0.051730105 | Y | 33168 | Prohibitin OS=Rattus norvegicus OX=10116 GN=Phb2 PE=1 SV=1 |
| | | | | | | | | 0.00000 | | | P18163\|ACSL1_RAT | + | 20 | 28 | 99.2 | 214 | 15 | 11 | 8 | 10176000 | 0.208180631 | N | 78179 | Long-chain-fatty-acid--CoA ligase 1 OS=Rattus norvegicus OX=10116 GN=Acsl1 PE=1 SV=1 |
| | | | | | | | | 0.00000 | | | Q9WVK3\|PECR_RAT | + | 23 | 48 | 99.1 | 205 | 37 | 8 | 8 | 11917000 | 0.243798013 | Y | 32433 | Peroxisomal trans-2-enoyl-CoA reductase OS=Rattus norvegicus OX=10116 GN=Pecr PE=2 SV=1 |
| | | | | | | | | 0.00000 | | | P13086\|SUCA_RAT | + | 24 | 32 | 99 | 194 | 17 | 5 | 5 | 8044300 | 0.164570308 | Y | 36148 | Succinate--CoA ligase [ADP/GDP-forming] subunit alpha, mitochondrial OS=Rattus norvegicus OX=10116 GN=Suclg1 PE=2 SV=2 |

| | | | | | | | | | | | | | | | |
|---|---|---|---|---|---|---|---|---|---|---|---|---|---|---|---|
| | **0.00000** | Q60587\|ECHB_RAT | + | 21 | 31 | 99.2 | 176 | 22 | 9 | 9 | 13498000 | **0.276142115** | Y | 51414 | Trifunctional enzyme subunit beta, mitochondrial OS=Rattus norvegicus OX=10116 GN=Hadhb PE=1 SV=1 |
| | **0.00000** | P24329\|THTR_RAT | + | 27 | 55 | 99 | 169 | 23 | 5 | 5 | 7435600 | **0.152117522** | N | 33407 | Thiosulfate sulfurtransferase OS=Rattus norvegicus OX=10116 GN=Tst PE=1 SV=3 |
| | **0.00000** | P17764\|THIL_RAT | + | 41 | 46 | 98.9 | 163 | 12 | 4 | 4 | 2189300 | **0.044788704** | N | 44695 | Acetyl-CoA acetyltransferase, mitochondrial OS=Rattus norvegicus OX=10116 GN=Acat1 PE=1 SV=1 |
| | **0.00000** | P29147\|BDH_RAT | + | 32 | 39 | 98.9 | 155 | 15 | 5 | 5 | 6003700 | **0.122823708** | N | 38202 | D-beta-hydroxybutyrate dehydrogenase, mitochondrial OS=Rattus norvegicus OX=10116 GN=Bdh1 PE=1 SV=2 |
| | **0.00000** | P85834\|EFTU_RAT | + | 30 | 27 | 99 | 145 | 13 | 5 | 5 | 4037800 | **0.082605322** | N | 49522 | Elongation factor Tu, mitochondrial OS=Rattus norvegicus OX=10116 GN=Tufm PE=1 SV=1 |
| | **0.00000** | O70351\|HCD2_RAT | + | 48 | 76 | 95.6 | 122 | 16 | 3 | 3 | 2753200 | **0.056324972** | N | 27246 | 3-hydroxyacyl-CoA dehydrogenase type-2 OS=Rattus norvegicus OX=10116 GN=Hsd17b10 PE=1 SV=3 |
| | **0.00000** | tr\|Q7TP12\|Q7TP12_RAT | + | 200 | 228 | 61.2 | 40 | 10 | 1 | 1 | 1252600 | **0.025625694** | N | 14969 | Cc2-17 OS=Rattus norvegicus OX=10116 GN=Sqor PE=1 SV=1 |
| | **0.00000** | P63039\|CH60_RAT | + | 29 | 61 | 94.2 | 140 | 7 | 4 | 4 | 18976000 | **0.388211052** | N | 60956 | 60 kDa heat shock protein, mitochondrial OS=Rattus norvegicus OX=10116 GN=Hspd1 PE=1 SV=1 |
| | **0.00000** | Q3KR86\|MIC60_RAT | + | 36 | 30 | 94 | 82 | 6 | 3 | 3 | 5965000 | **0.122031984** | N | 67177 | MICOS complex subunit Mic60 (Fragment) OS=Rattus norvegicus OX=10116 GN=Immt PE=1 SV=1 |
| | **0.00000** | P0C2X9\|AL4A1_RAT | + | 37 | 47 | 98.8 | 171 | 9 | 4 | 4 | 3731900 | **0.076347219** | N | 61869 | Delta-1-pyrroline-5-carboxylate dehydrogenase, mitochondrial OS=Rattus norvegicus OX=10116 GN=Aldh4a1 PE=1 SV=1 |
| | **0.00000** | B3DMA2\|ACD11_RAT | + | 39 | 53 | 95.4 | 90 | 4 | 4 | 4 | 3247200 | **0.066431225** | N | 87371 | Acyl-CoA dehydrogenase family member 11 OS=Rattus norvegicus OX=10116 GN=Acad11 PE=1 SV=1 |
| | **0.00000** | P16970\|ABCD3_RAT | + | 49 | 44 | 94.2 | 123 | 5 | 2 | 2 | 1969000 | **0.040281807** | N | 75316 | ATP-binding cassette sub-family D member 3 OS=Rattus norvegicus OX=10116 GN=Abcd3 PE=1 SV=3 |
| | **0.00000** | tr\|A0A0U1RRV5\|A0A0U1I | + | 50 | 1062 | 77.7 | 42 | 2 | 3 | 2 | 6711400 | **0.137301837** | Y | 2E+05 | Clustered mitochondria protein homolog OS=Rattus norvegicus OX=10116 GN=Cluh PE=1 SV=1 |
| | **0.00000** | P04762\|CATA_RAT | + | 60 | 103 | 94.7 | 75 | 8 | 3 | 3 | 1307100 | **0.026740655** | N | 59757 | Catalase OS=Rattus norvegicus OX=10116 GN=Cat PE=1 SV=3 |
| | **0.00000** | tr\|D3ZXA6\|D3ZXA6_RAT | + | 74 | 482 | 63.2 | 30 | 3 | 2 | 2 | 2048500 | **0.041908218** | N | 98819 | Pyruvate dehydrogenase phosphatase regulatory subunit OS=Rattus norvegicus OX=10116 GN=Pdpr PE=1 SV=1 |
| | **0.00000** | tr\|G3V728\|G3V728_RAT | + | 77 | 52 | 89.8 | 89 | 8 | 2 | 2 | 2658700 | **0.054391691** | N | 33346 | 4-nitrophenylphosphatase domain and non-neuronal SNAP25-like protein homolog 1 (C. elegans), isoform CRA_b OS=Rattus norvegicus OX=10116 GN=Nipsnap1 PE=1 SV=1 |
| | **0.00000** | tr\|F1LMJ8\|F1LMJ8_RAT | + | 81 | 701 | 47 | 24 | 5 | 2 | 2 | 6004200 | **0.122833937** | N | 49215 | Calcium uptake protein 2, mitochondrial OS=Rattus norvegicus OX=10116 GN=Micu2 PE=1 SV=1 |
| | **0.00000** | tr\|D3ZXF9\|D3ZXF9_RAT | + | 88 | 106 | 74.2 | 81 | 9 | 2 | 2 | 952090 | **0.019477859** | Y | 29441 | Mitochondrial ribosomal protein L12 OS=Rattus norvegicus OX=10116 GN=Mrpl12 PE=1 SV=1 |
| | **0.00000** | tr\|G3V6I4\|G3V6I4_RAT | + | 106 | 248 | 76 | 98 | 9 | 1 | 1 | 1562300 | **0.031961537** | N | 37791 | Mitochondrial amidoxime reducing component 1 OS=Rattus norvegicus OX=10116 GN=Marc1 PE=1 SV=1 |
| | **0.00000** | tr\|D3ZUJ5\|D3ZUJ5_RAT | + | 112 | 1624 | 55.1 | 26 | 8 | 1 | 1 | 2021600 | **0.041357897** | Y | 23973 | Deoxythymidylate kinase OS=Rattus norvegicus OX=10116 GN=Dtymk PE=1 SV=1 |
| | **0.00000** | tr\|D3ZFJ6\|D3ZFJ6_RAT | + | 125 | 96 | 74 | 38 | 7 | 1 | 1 | 889180 | **0.018190846** | N | 60420 | Lactamase, beta OS=Rattus norvegicus OX=10116 GN=Lactb PE=1 SV=1 |
| | **0.00000** | Q09073\|ADT2_RAT | + | 51 | 63 | 98 | 69 | 10 | 3 | 3 | 2645600 | **0.054123691** | N | 32901 | ADP/ATP translocase 2 OS=Rattus norvegicus OX=10116 GN=Slc25a5 PE=1 SV=3 |
| 980110781.00 | **100.00000** | | | | | | | | | | 4888062800.00 | **100** | | | |
| 9801107.81 | | | | | | | | | | | | | | | |
| | | | | | | | | | | | 48880628 | | | | |
| 9801107.81 | | | | | | | | | | | | | | | |
| | | | | | | | | | | | 48880628 | | | | |
