## Supplementary material for "Permeability transition pore-related changes in the proteome and channel activity of ATP synthase dimers and monomers": ATP-synthase

| Area (IBAQ) | riBAQ | Accession | rIBAQ PTP/Control | Area (IBAQ) | riBAQ | Description |
|---|---|---|---|---|---|---|
| CONTROL | % | | | PTP | % | |
| **Complex I** | | | | | | |
| 1898400 | **0.19369** | Q561S0\|NDUAA_RAT | **0.123650616** | 1170700 | **0.023950183** | NADH dehydrogenase [ubiquinone] 1 alpha subcomplex subunit 10, mitochondrial OS=Rattus norvegicus OX=10116 GN=Ndufa10 PE=1 SV=1 |
| 146380 | **0.01494** | tr\|Q5XIH3\|Q5XIH3_RAT | **6.473804099** | 4726100 | **0.096686565** | NADH dehydrogenase [ubiquinone] flavoprotein 1, mitochondrial OS=Rattus norvegicus OX=10116 GN=Ndufv1 PE=1 SV=1 |
| 69212 | **0.00706** | tr\|D4A565\|D4A565_RAT | **12.29279085** | 4243200 | **0.086807395** | NADH dehydrogenase (Ubiquinone) 1 beta subcomplex, 5 (Predicted), isoform CRA_b OS=Rattus norvegicus OX=10116 GN=Ndufb5 PE=1 SV=1 |
| 282740 | **0.02885** | tr\|D4A0T0\|D4A0T0_RAT | **4.635994455** | 6537200 | **0.133738053** | NADH:ubiquinone oxidoreductase subunit B10 OS=Rattus norvegicus OX=10116 GN=Ndufb10 PE=1 SV=1 |
| 196860 | **0.02009** | tr\|Q5RJN0\|Q5RJN0_RAT | **2.396029051** | 2352400 | **0.048125405** | NADH dehydrogenase (Ubiquinone) Fe-S protein 7 OS=Rattus norvegicus OX=10116 GN=Ndufs7 PE=1 SV=1 |
| 0 | **0.00000** | tr\|B0BNE6\|B0BNE6_RAT | **+** | 129890 | **0.00265729** | NADH dehydrogenase (Ubiquinone) Fe-S protein 8 (Predicted), isoform CRA_a OS=Rattus norvegicus OX=10116 GN=Ndufs8 PE=1 SV=1 |
| | **0.00000** | tr\|B2RYS8\|B2RYS8_RAT | **+** | 1986100 | **0.040631638** | NADH dehydrogenase [ubiquinone] 1 beta subcomplex subunit 8, mitochondrial OS=Rattus norvegicus OX=10116 GN=Ndufb8 PE=1 SV=1 |
| | **0.00000** | tr\|Q5PQZ9\|Q5PQZ9_RAT | **+** | 638550 | **0.013063457** | NADH dehydrogenase [ubiquinone] 1 subunit C2 OS=Rattus norvegicus OX=10116 GN=Ndufc2 PE=1 SV=1 |
| | **0.00000** | tr\|F1LPG5\|F1LPG5_RAT | **+** | 1846400 | **0.037773655** | NADH:ubiquinone oxidoreductase subunit B4 OS=Rattus norvegicus OX=10116 GN=Ndufb4 PE=1 SV=1 |
| | **0.00000** | tr\|A9UMV9\|A9UMV9_RA | **+** | 407550 | **0.008337659** | NADH:ubiquinone oxidoreductase subunit A7 OS=Rattus norvegicus OX=10116 GN=Ndufa7 PE=1 SV=1 |
| | **0.00000** | tr\|B2RYW3\|B2RYW3_RAT | **+** | 1696700 | **0.034711093** | NADH dehydrogenase (Ubiquinone) 1 beta subcomplex, 9 OS=Rattus norvegicus OX=10116 GN=Ndufb9 PE=1 SV=1 |
| | **0.00000** | Q641Y2\|NDUS2_RAT | **+** | 4160900 | **0.085123702** | NADH dehydrogenase [ubiquinone] iron-sulfur protein 2, mitochondrial OS=Rattus norvegicus OX=10116 GN=Ndufs2 PE=1 SV=1 |
| | **0.00000** | tr\|D3ZE15\|D3ZE15_RAT | **+** | 1861700 | **0.038086663** | NADH:ubiquinone oxidoreductase subunit A13 OS=Rattus norvegicus OX=10116 GN=Ndufa13 PE=1 SV=1 |
| | **0.00000** | P52504\|NDUS6_RAT | **+** | 865090 | **0.017698013** | NADH dehydrogenase [ubiquinone] iron-sulfur protein 6, mitochondrial OS=Rattus norvegicus OX=10116 GN=Ndufs6 PE=3 SV=1 |
| | **0.00000** | tr\|D3ZF13\|D3ZF13_RAT | **+** | 3263000 | **0.066754462** | Acyl carrier protein OS=Rattus norvegicus OX=10116 GN=Ndufab1 PE=1 SV=1 |
| **Complex III** | | | | | | |
| 2291000 | **0.23375** | P32551\|QCR2_RAT | **-** | 0 | **0** | Cytochrome b-c1 complex subunit 2, mitochondrial OS=Rattus norvegicus OX=10116 GN=Uqcrc2 PE=1 SV=2 |
| 43626 | **0.00445** | Q68FY0\|QCR1_RAT | **-** | 0 | **0** | Cytochrome b-c1 complex subunit 1, mitochondrial OS=Rattus norvegicus OX=10116 GN=Uqcrc1 PE=1 SV=1 |
| **Complex IV** | | | | | | |
| 29933 | **0.00305** | P11240\|COX5A_RAT | **-** | 0 | **0** | Cytochrome c oxidase subunit 5A, mitochondrial OS=Rattus norvegicus OX=10116 GN=Cox5a PE=1 SV=1 |
| 0 | **0.00000** | P10888\|COX41_RAT | **+** | 4134200 | **0.084577473** | Cytochrome c oxidase subunit 4 isoform 1, mitochondrial OS=Rattus norvegicus OX=10116 GN=Cox4i1 PE=1 SV=1 |
| | **0.00000** | P00406\|COX2_RAT | **+** | 2403600 | **0.049172854** | Cytochrome c oxidase subunit 2 OS=Rattus norvegicus OX=10116 GN=Mtco2 PE=1 SV=3 |
| **Complex V** | | | | | | |
| 513120 | **0.05235** | P35435\|ATPG_RAT | **6.392971174** | 16360000 | **0.334692918** | ATP synthase subunit gamma, mitochondrial OS=Rattus norvegicus OX=10116 GN=Atp5f1c PE=1 SV=2 |
| 17943000 | **1.83071** | Q06647\|ATPO_RAT | **2.434785456** | 217880000 | **4.457389541** | ATP synthase subunit O, mitochondrial OS=Rattus norvegicus OX=10116 GN=Atp5po PE=1 SV=1 |
| 437370 | **0.04462** | P29418\|ATP5E_RAT | **2.457919238** | 5361400 | **0.109683534** | ATP synthase subunit epsilon, mitochondrial OS=Rattus norvegicus OX=10116 GN=Atp5f1e PE=1 SV=2 |
| | **0.00000** | D3Z9R8\|ATP68_RAT | **+** | 4396500 | **0.089943607** | ATP synthase subunit ATP5MPL, mitochondrial OS=Rattus norvegicus OX=10116 GN=Atp5mpl PE=1 SV=1 |
| | **0.00000** | P29419\|ATP5I_RAT | **+** | 15392000 | **0.314889571** | ATP synthase subunit e, mitochondrial OS=Rattus norvegicus OX=10116 GN=Atp5me PE=1 SV=3 |
| | **0.00000** | P21571\|ATP5J_RAT | **+** | 830950 | **0.016999577** | ATP synthase-coupling factor 6, mitochondrial OS=Rattus norvegicus OX=10116 GN=Atp5pf PE=1 SV=1 |
| **Other** | | | | | | |
| 66230000 | **6.75740** | Q5XIH7\|PHB2_RAT | **0.013332791** | 4403900 | **0.090094996** | Prohibitin-2 OS=Rattus norvegicus OX=10116 GN=Phb2 PE=1 SV=1 |
| 23705000 | **2.41860** | Q02253\|MMSA_RAT | **0.029256602** | 3458800 | **0.070760138** | Methylmalonate-semialdehyde dehydrogenase [acylating], mitochondrial OS=Rattus norvegicus OX=10116 GN=Aldh6a1 PE=1 SV=1 |
| 1524800 | **0.15557** | P07756\|CPSM_RAT | **14.00474205** | 106500000 | **2.178777245** | Carbamoyl-phosphate synthase [ammonia], mitochondrial OS=Rattus norvegicus OX=10116 GN=Cps1 PE=1 SV=1 |
| 1662400 | **0.16961** | P22791\|HMCS2_RAT | **10.50982525** | 87135000 | **1.78260803** | Hydroxymethylglutaryl-CoA synthase, mitochondrial OS=Rattus norvegicus OX=10116 GN=Hmgcs2 PE=1 SV=1 |
| 2036300 | **0.20776** | F1M775\|DIAP1_RAT | **2.133021238** | 21662000 | **0.443161246** | Protein diaphanous homolog 1 OS=Rattus norvegicus OX=10116 GN=Diaph1 PE=1 SV=3 |

| | | | | | | |
|---|---|---|---|---|---|---|
| 2025700 | **0.20668** | B0K035\|MTFR2_RAT | - | | **0** | Mitochondrial fission regulator 2 OS=Rattus norvegicus OX=10116 GN=Mtfr2 PE=2 SV=1 |
| 312410 | **0.03187** | P52873\|PYC_RAT | **21.83023454** | 34013000 | **0.695838032** | Pyruvate carboxylase, mitochondrial OS=Rattus norvegicus OX=10116 GN=Pc PE=1 SV=2 |
| | **0.00000** | P10860\|DHE3_RAT | **+** | 61338000 | **1.254852945** | Glutamate dehydrogenase 1, mitochondrial OS=Rattus norvegicus OX=10116 GN=Glud1 PE=1 SV=2 |
| | **0.00000** | tr\|A0A0G2KB63\|A0A0G2K | **+** | 2528600 | **0.051730105** | Prohibitin OS=Rattus norvegicus OX=10116 GN=Phb2 PE=1 SV=1 |
| | **0.00000** | P18163\|ACSL1_RAT | **+** | 10176000 | **0.208180631** | Long-chain-fatty-acid--CoA ligase 1 OS=Rattus norvegicus OX=10116 GN=Acsl1 PE=1 SV=1 |
| | **0.00000** | Q9WVK3\|PECR_RAT | **+** | 11917000 | **0.243798013** | Peroxisomal trans-2-enoyl-CoA reductase OS=Rattus norvegicus OX=10116 GN=Pecr PE=2 SV=1 |
| | **0.00000** | P13086\|SUCA_RAT | **+** | 8044300 | **0.164570308** | Succinate--CoA ligase [ADP/GDP-forming] subunit alpha, mitochondrial OS=Rattus norvegicus OX=10116 GN=Suclg1 PE=2 SV=2 |
| | **0.00000** | Q60587\|ECHB_RAT | **+** | 13498000 | **0.276142115** | Trifunctional enzyme subunit beta, mitochondrial OS=Rattus norvegicus OX=10116 GN=Hadhb PE=1 SV=1 |
| | **0.00000** | P24329\|THTR_RAT | **+** | 7435600 | **0.152117522** | Thiosulfate sulfurtransferase OS=Rattus norvegicus OX=10116 GN=Tst PE=1 SV=3 |
| | **0.00000** | P17764\|THIL_RAT | **+** | 2189300 | **0.044788704** | Acetyl-CoA acetyltransferase, mitochondrial OS=Rattus norvegicus OX=10116 GN=Acat1 PE=1 SV=1 |
| | **0.00000** | P29147\|BDH_RAT | **+** | 6003700 | **0.122823708** | D-beta-hydroxybutyrate dehydrogenase, mitochondrial OS=Rattus norvegicus OX=10116 GN=Bdh1 PE=1 SV=2 |
| | **0.00000** | P85834\|EFTU_RAT | **+** | 4037800 | **0.082605322** | Elongation factor Tu, mitochondrial OS=Rattus norvegicus OX=10116 GN=Tufm PE=1 SV=1 |
| | **0.00000** | O70351\|HCD2_RAT | **+** | 2753200 | **0.056324972** | 3-hydroxyacyl-CoA dehydrogenase type-2 OS=Rattus norvegicus OX=10116 GN=Hsd17b10 PE=1 SV=3 |
| | **0.00000** | tr\|Q7TP12\|Q7TP12_RAT | **+** | 1252600 | **0.025625694** | Cc2-17 OS=Rattus norvegicus OX=10116 GN=Sqor PE=1 SV=1 |
| | **0.00000** | P63039\|CH60_RAT | **+** | 18976000 | **0.388211052** | 60 kDa heat shock protein, mitochondrial OS=Rattus norvegicus OX=10116 GN=Hspd1 PE=1 SV=1 |
| | **0.00000** | Q3KR86\|MIC60_RAT | **+** | 5965000 | **0.122031984** | MICOS complex subunit Mic60 (Fragment) OS=Rattus norvegicus OX=10116 GN=Immt PE=1 SV=1 |
| | **0.00000** | P0C2X9\|AL4A1_RAT | **+** | 3731900 | **0.076347219** | Delta-1-pyrroline-5-carboxylate dehydrogenase, mitochondrial OS=Rattus norvegicus OX=10116 GN=Aldh4a1 PE=1 SV=1 |
| | **0.00000** | B3DMA2\|ACD11_RAT | **+** | 3247200 | **0.066431225** | Acyl-CoA dehydrogenase family member 11 OS=Rattus norvegicus OX=10116 GN=Acad11 PE=1 SV=1 |
| | **0.00000** | P16970\|ABCD3_RAT | **+** | 1969000 | **0.040281807** | ATP-binding cassette sub-family D member 3 OS=Rattus norvegicus OX=10116 GN=Abcd3 PE=1 SV=3 |
| | **0.00000** | tr\|A0A0U1RRV5\|A0A0U1F | **+** | 6711400 | **0.137301837** | Clustered mitochondria protein homolog OS=Rattus norvegicus OX=10116 GN=Cluh PE=1 SV=1 |
| | **0.00000** | P04762\|CATA_RAT | **+** | 1307100 | **0.026740655** | Catalase OS=Rattus norvegicus OX=10116 GN=Cat PE=1 SV=3 |
| | **0.00000** | tr\|D3ZXA6\|D3ZXA6_RAT | **+** | 2048500 | **0.041908218** | Pyruvate dehydrogenase phosphatase regulatory subunit OS=Rattus norvegicus OX=10116 GN=Pdpr PE=1 SV=1 |
| | **0.00000** | tr\|G3V728\|G3V728_RAT | **+** | 2658700 | **0.054391691** | 4-nitrophenylphosphatase domain and non-neuronal SNAP25-like protein homolog 1 (C. elegans), isoform CRA_b OS=Rattus norvegicus OX=10116 GN=Nipsnap1 PE=1 SV=1 |
| | **0.00000** | tr\|F1LMJ8\|F1LMJ8_RAT | **+** | 6004200 | **0.122833937** | Calcium uptake protein 2, mitochondrial OS=Rattus norvegicus OX=10116 GN=Micu2 PE=1 SV=1 |
| | **0.00000** | tr\|D3ZXF9\|D3ZXF9_RAT | **+** | 952090 | **0.019477859** | Mitochondrial ribosomal protein L12 OS=Rattus norvegicus OX=10116 GN=Mrpl12 PE=1 SV=1 |
| | **0.00000** | tr\|G3V6I4\|G3V6I4_RAT | **+** | 1562300 | **0.031961537** | Mitochondrial amidoxime reducing component 1 OS=Rattus norvegicus OX=10116 GN=Marc1 PE=1 SV=1 |
| | **0.00000** | tr\|D3ZUJ5\|D3ZUJ5_RAT | **+** | 2021600 | **0.041357897** | Deoxythymidylate kinase OS=Rattus norvegicus OX=10116 GN=Dtymk PE=1 SV=1 |
| | **0.00000** | tr\|D3ZFJ6\|D3ZFJ6_RAT | **+** | 889180 | **0.018190846** | Lactamase, beta OS=Rattus norvegicus OX=10116 GN=Lactb PE=1 SV=1 |
| | **0.00000** | Q09073\|ADT2_RAT | **+** | 2645600 | **0.054123691** | ADP/ATP translocase 2 OS=Rattus norvegicus OX=10116 GN=Slc25a5 PE=1 SV=3 |
