## Supplementary material for "Permeability transition pore-related changes in the proteome and channel activity of ATP synthase dimers and monomers": ATP-synthase

| Area (IBAQ) CONTROL | riBAQ % | Accession | rIBAQ PTP/Control | Area (IBAQ) PTP | riBAQ % | Description |
|---|---|---|---|---|---|---|
| **Complex III** | | | | | | |
| 22165000 | **0.21475999** | Q68FY0\|QCR1_RAT | **0.451805576** | 15442000.00 | **0.097029761** | Cytochrome b-c1 complex subunit 1, mitochondrial OS=Rattus norvegicus OX=10116 GN=Uqcrc1 PE=1 SV=1 |
| 35141000 | **0.34048639** | P32551\|QCR2_RAT | **0.431170317** | 23364000.00 | **0.146807624** | Cytochrome b-c1 complex subunit 2, mitochondrial OS=Rattus norvegicus OX=10116 GN=Uqcrc2 PE=1 SV=2 |
| 1652700 | **0.01601326** | tr\|A0A0G2K8Q8\|A0A0G | - | | **0** | Ubiquinol-cytochrome c reductase, complex III subunit X OS=Rattus norvegicus OX=10116 GN=Uqcr10 PE=1 SV=1 |
| 952880 | **0.0092326** | Q5M9I5\|QCR6_RAT | - | | **0** | Cytochrome b-c1 complex subunit 6, mitochondrial OS=Rattus norvegicus OX=10116 GN=Uqcrh PE=3 SV=1 |
| 1354300 | **0.01312201** | Q7TQ16\|QCR8_RAT | - | | **0** | Cytochrome b-c1 complex subunit 8 OS=Rattus norvegicus OX=10116 GN=Uqcrq PE=3 SV=1 |
| **Complex IV** | | | | | | |
| 8579600 | **0.08312902** | P11240\|COX5A_RAT | **0.10294232** | 1361900.00 | **0.008557495** | Cytochrome c oxidase subunit 5A, mitochondrial OS=Rattus norvegicus OX=10116 GN=Cox5a PE=1 SV=1 |
| 1546100 | **0.01498039** | P10818\|CX6A1_RAT | - | | **0** | Cytochrome c oxidase subunit 6A1, mitochondrial OS=Rattus norvegicus OX=10116 GN=Cox6a1 PE=1 SV=2 |
| 1308400 | **0.01267728** | P11951\|CX6C2_RAT | - | | **0** | Cytochrome c oxidase subunit 6C-2 OS=Rattus norvegicus OX=10116 GN=Cox6c2 PE=1 SV=3 |
| 252730 | **0.00244874** | P80432\|COX7C_RAT | - | | **0** | Cytochrome c oxidase subunit 7C, mitochondrial OS=Rattus norvegicus OX=10116 GN=Cox7c PE=1 SV=2 |
| 2595300 | **0.02514625** | P35171\|CX7A2_RAT | - | | **0** | Cytochrome c oxidase subunit 7A2, mitochondrial OS=Rattus norvegicus OX=10116 GN=Cox7a2 PE=1 SV=1 |
| 867700 | **0.00840727** | B1WC61\|ACAD9_RAT | - | | **0** | Complex I assembly factor ACAD9, mitochondrial OS=Rattus norvegicus OX=10116 GN=Acad9 PE=1 SV=1 |
| 911840 | **0.00883495** | P05505\|COX3_RAT | **0.291887212** | 410410.00 | **0.00257881** | Cytochrome c oxidase subunit 3 OS=Rattus norvegicus OX=10116 GN=Mtco3 PE=1 SV=5 |
| 447640 | **0.00433725** | P80431\|COX7B_RAT | **0.420754105** | 290430.00 | **0.001824916** | Cytochrome c oxidase subunit 7B, mitochondrial OS=Rattus norvegicus OX=10116 GN=Cox7b PE=1 SV=3 |
| 933500 | **0.00904482** | P19804\|NDKB_RAT | **4.525596951** | 6514400.00 | **0.040933213** | Nucleoside diphosphate kinase B OS=Rattus norvegicus OX=10116 GN=Nme2 PE=1 SV=1 |
| **Complex V** | | | | | | |
| 658280000 | **6.37817306** | tr\|G3V7Y3\|G3V7Y3_RA | **0.181249045** | 183980000.00 | **1.156037778** | ATP synthase subunit delta, mitochondrial OS=Rattus norvegicus OX=10116 GN=Atp5f1d PE=1 SV=1 |
| 83153000 | **0.80568181** | Q6PDU7\|ATP5L_RAT | **0.237510161** | 30454000.00 | **0.191357618** | ATP synthase subunit g, mitochondrial OS=Rattus norvegicus OX=10116 GN=Atp5mg PE=1 SV=2 |
| 12692000 | **0.12297468** | D3Z9R8\|ATP68_RAT | **0.42317591** | 8282000.00 | **0.052039922** | ATP synthase subunit ATP5MPL, mitochondrial OS=Rattus norvegicus OX=10116 GN=Atp5mpl PE=1 SV=1 |
| 78115000 | **0.75686788** | D3ZAF6\|ATPK_RAT | **0.28140369** | 33896000.00 | **0.212985414** | ATP synthase subunit f, mitochondrial OS=Rattus norvegicus OX=10116 GN=Atp5mf PE=1 SV=1 |
| 10200000 | **0.09882932** | Q06645\|AT5G1_RAT | - | | **0** | ATP synthase F(0) complex subunit C1, mitochondrial OS=Rattus norvegicus OX=10116 GN=Atp5mc1 PE=1 SV=1 |
| 4736200 | **0.04588975** | P05504\|ATP6_RAT | **0.26136424** | 1908800.00 | **0.011993939** | ATP synthase subunit a OS=Rattus norvegicus OX=10116 GN=Mt-atp6 PE=1 SV=3 |
| **Other** | | | | | | |
| 99521000 | **0.96427381** | P07756\|CPSM_RAT | **2.057326104** | 315720000.00 | **1.983825673** | Carbamoyl-phosphate synthase [ammonia], mitochondrial OS=Rattus norvegicus OX=10116 GN=Cps1 PE=1 SV=1 |
| 95293000 | **0.92330808** | P00507\|AATM_RAT | **0.341264466** | 50146000.00 | **0.315092241** | Aspartate aminotransferase, mitochondrial OS=Rattus norvegicus OX=10116 GN=Got2 PE=1 SV=2 |
| 21445000 | **0.2077838** | P22791\|HMCS2_RAT | **7.598846059** | 251280000.00 | **1.578917126** | Hydroxymethylglutaryl-CoA synthase, mitochondrial OS=Rattus norvegicus OX=10116 GN=Hmgcs2 PE=1 SV=1 |
| 10599000 | **0.10269529** | tr\|A0A0G2JVH4\|A0A0G | **0.478907638** | 7827100.00 | **0.049181559** | MICOS complex subunit MIC60 OS=Rattus norvegicus OX=10116 GN=Immt PE=1 SV=1 |
| 25243000 | **0.24458319** | Q60587\|ECHB_RAT | **2.501214345** | 97359000.00 | **0.611754984** | Trifunctional enzyme subunit beta, mitochondrial OS=Rattus norvegicus OX=10116 GN=Hadhb PE=1 SV=1 |
| 38664000 | **0.37462126** | Q02253\|MMSA_RAT | **0.347753203** | 20733000.00 | **0.130275743** | Methylmalonate-semialdehyde dehydrogenase [acylating], mitochondrial OS=Rattus norvegicus OX=10116 GN=Aldh6a1 PE=1 SV=1 |
| 47000000 | **0.45539001** | P0C2X9\|AL4A1_RAT | **0.355217164** | 25744000.00 | **0.161762347** | Delta-1-pyrroline-5-carboxylate dehydrogenase, mitochondrial OS=Rattus norvegicus OX=10116 GN=Aldh4a1 PE=1 SV=1 |
| 6627100 | **0.06421096** | P10860\|DHE3_RAT | **5.827291219** | 59549000.00 | **0.374175963** | Glutamate dehydrogenase 1, mitochondrial OS=Rattus norvegicus OX=10116 GN=Glud1 PE=1 SV=2 |
| 780490 | **0.00756228** | A2VCW9\|AASS_RAT | **34.58784815** | 41627000.00 | **0.26156313** | Alpha-aminoadipic semialdehyde synthase, mitochondrial OS=Rattus norvegicus OX=10116 GN=Aass PE=2 SV=1 |
| 2991300 | **0.02898315** | P63039\|CH60_RAT | **11.36152989** | 52406000.00 | **0.329292944** | 60 kDa heat shock protein, mitochondrial OS=Rattus norvegicus OX=10116 GN=Hspd1 PE=1 SV=1 |
| 4104600 | **0.03977008** | tr\|G3V6I4\|G3V6I4_RAT | **0.262367457** | 1660600.00 | **0.010434375** | Mitochondrial amidoxime reducing component 1 OS=Rattus norvegicus OX=10116 GN=Marc1 PE=1 SV=1 |
| 4000100 | **0.03875757** | P29147\|BDH_RAT | **2.494264041** | 15385000.00 | **0.096671601** | D-beta-hydroxybutyrate dehydrogenase, mitochondrial OS=Rattus norvegicus OX=10116 GN=Bdh1 PE=1 SV=2 |
| 991930 | **0.00961096** | Q5XIC2\|ECSIT_RAT | - | | **0** | Evolutionarily conserved signaling intermediate in Toll pathway, mitochondrial OS=Rattus norvegicus OX=10116 GN=Ecsit PE=1 SV=1 |
| 1666300 | **0.01614503** | tr\|Q5EBA4\|Q5EBA4_RA | **4.200536438** | 10793000.00 | **0.067817783** | Nipsnap1 protein (Fragment) OS=Rattus norvegicus OX=10116 GN=Nipsnap1 PE=2 SV=1 |
| 1768600 | **0.01713623** | Q9WVK3\|PECR_RAT | **2.924779148** | 7976400.00 | **0.050119685** | Peroxisomal trans-2-enoyl-CoA reductase OS=Rattus norvegicus OX=10116 GN=Pecr PE=2 SV=1 |

| | | | | | | |
|---|---|---|---|---|---|---|
| 408280 | **0.00395589** | P08011\|MGST1_RAT | **2.774444156** | 1746700.00 | **0.010975384** | Microsomal glutathione S-transferase 1 OS=Rattus norvegicus OX=10116 GN=Mgst1 PE=1 SV=3 |
| 154490 | **0.00149688** | tr\|D3ZUX5\|D3ZUX5_RA | **-** | | **0** | MICOS complex subunit OS=Rattus norvegicus OX=10116 GN=Chchd3 PE=1 SV=1 |
| 1044200 | **0.01011741** | P00481\|OTC_RAT | **-** | | **0** | Ornithine carbamoyltransferase, mitochondrial OS=Rattus norvegicus OX=10116 GN=Otc PE=1 SV=1 |
| 1275600 | **0.01235948** | F1M775\|DIAP1_RAT | **-** | | **0** | Protein diaphanous homolog 1 OS=Rattus norvegicus OX=10116 GN=Diaph1 PE=1 SV=3 |
| 1012600 | **0.00981123** | A0A0G2K047\|ACSS3_RA | **-** | | **0** | Acyl-CoA synthetase short-chain family member 3, mitochondrial OS=Rattus norvegicus OX=10116 GN=Acss3 PE=1 SV=1 |
| 1443600 | **0.01398726** | P45953\|ACADV_RAT | **3.856371336** | 8584400.00 | **0.053940052** | Very long-chain specific acyl-CoA dehydrogenase, mitochondrial OS=Rattus norvegicus OX=10116 GN=Acadvl PE=1 SV=1 |
| 160410 | **0.00155424** | P48721\|GRP75_RAT | **8.625759348** | 2133600.00 | **0.013406469** | Stress-70 protein, mitochondrial OS=Rattus norvegicus OX=10116 GN=Hspa9 PE=1 SV=3 |
| 516050 | **0.00500009** | Q5PQT3\|GLYAT_RAT | **2.585866196** | 2057700.00 | **0.012929552** | Glycine N-acyltransferase OS=Rattus norvegicus OX=10116 GN=Glyat PE=2 SV=1 |
| 1193300 | **0.01156206** | O88994\|MARC2_RAT | **2.603327737** | 4790300.00 | **0.030099836** | Mitochondrial amidoxime reducing component 2 OS=Rattus norvegicus OX=10116 GN=Marc2 PE=2 SV=1 |
| 1064400 | **0.01031313** | P61980\|HNRPK_RAT | **-** | | **0** | Heterogeneous nuclear ribonucleoprotein K OS=Rattus norvegicus OX=10116 GN=Hnrnpk PE=1 SV=1 |
| 430990 | **0.00417593** | Q5XHZ0\|TRAP1_RAT | **-** | | **0** | Heat shock protein 75 kDa, mitochondrial OS=Rattus norvegicus OX=10116 GN=Trap1 PE=1 SV=1 |
| 3606500 | **0.03494392** | Q924S5\|LONM_RAT | **0.254188778** | 1413600.00 | **0.008882351** | Lon protease homolog, mitochondrial OS=Rattus norvegicus OX=10116 GN=Lonp1 PE=2 SV=1 |
| 933500 | **0.00904482** | Q05982\|NDKA_RAT | **-** | | **0** | Nucleoside diphosphate kinase A OS=Rattus norvegicus OX=10116 GN=Nme1 PE=1 SV=1 |
| 302750 | **0.00293339** | P70619\|GSHR_RAT | **-** | | **0** | Glutathione reductase (Fragment) OS=Rattus norvegicus OX=10116 GN=Gsr PE=2 SV=2 |
| 290590 | **0.00281557** | tr\|D3ZXF9\|D3ZXF9_RAT | **5.236229213** | 2346300.00 | **0.014742969** | Mitochondrial ribosomal protein L12 OS=Rattus norvegicus OX=10116 GN=Mrpl12 PE=1 SV=1 |
| 295080 | **0.00285907** | D4A7N1\|MIC25_RAT | **-** | | **0** | MICOS complex subunit Mic25 OS=Rattus norvegicus OX=10116 GN=Chchd6 PE=1 SV=1 |
| 831730 | **0.00805876** | Q8VID1\|DHRS4_RAT | **10.68905185** | 13709000.00 | **0.08614046** | Dehydrogenase/reductase SDR family member 4 OS=Rattus norvegicus OX=10116 GN=Dhrs4 PE=2 SV=2 |
| 131730 | **0.00127635** | Q5RJR8\|LRC59_RAT | **12.00575455** | 2438700.00 | **0.015323564** | Leucine-rich repeat-containing protein 59 OS=Rattus norvegicus OX=10116 GN=Lrrc59 PE=1 SV=1 |
| 1315400 | **0.01274511** | tr\|A0A0U1RRQ6\|A0A0U | **-** | | **0** | Solute carrier family 25, member 44 OS=Rattus norvegicus OX=10116 GN=Slc25a44 PE=1 SV=1 |
| 1753800 | **0.01699283** | P18163\|ACSL1_RAT | **16.22233038** | 43871000.00 | **0.275663297** | Long-chain-fatty-acid--CoA ligase 1 OS=Rattus norvegicus OX=10116 GN=Acsl1 PE=1 SV=1 |
| 555250 | **0.0053799** | Q9JIR0\|RIMB1_RAT | **4.248797423** | 3637800.00 | **0.022858105** | Peripheral-type benzodiazepine receptor-associated protein 1 OS=Rattus norvegicus OX=10116 GN=Tspoap1 PE=1 SV=2 |
| | **0** | D3ZG52\|DNA2_RAT | **+** | 31878000.00 | **0.200305317** | DNA replication ATP-dependent helicase/nuclease DNA2 OS=Rattus norvegicus OX=10116 GN=Dna2 PE=3 SV=1 |
| | **0** | Q5XIT9\|MCCB_RAT | **+** | 5586100.00 | **0.035100243** | Methylcrotonoyl-CoA carboxylase beta chain, mitochondrial OS=Rattus norvegicus OX=10116 GN=Mccc2 PE=2 SV=1 |
| | **0** | Q5SGE0\|LPPRC_RAT | **+** | 2652900.00 | **0.016669489** | Leucine-rich PPR motif-containing protein, mitochondrial OS=Rattus norvegicus OX=10116 GN=Lrpprc PE=1 SV=1 |
| | **0** | Q5XIN6\|LETM1_RAT | **+** | 2969100.00 | **0.018656331** | Mitochondrial proton/calcium exchanger protein OS=Rattus norvegicus OX=10116 GN=Letm1 PE=1 SV=1 |
| | **0** | P17178\|CP27A_RAT | **+** | 1143900.00 | **0.007187692** | Sterol 26-hydroxylase, mitochondrial OS=Rattus norvegicus OX=10116 GN=Cyp27a1 PE=1 SV=1 |
| | **0** | Q5BK22\|GTPB8_RAT | **+** | 2574300.00 | **0.016175606** | GTP-binding protein 8 OS=Rattus norvegicus OX=10116 GN=Gtpbp8 PE=2 SV=1 |
| | **0** | tr\|F1LPV8\|F1LPV8_RAT | **+** | 1047100.00 | **0.00657945** | Succinate--CoA ligase [GDP-forming] subunit beta, mitochondrial OS=Rattus norvegicus OX=10116 GN=Suclg2 PE=1 SV=2 |
| | **0** | Q66H15\|RMD3_RAT | **+** | 335860.00 | **0.002110375** | Regulator of microtubule dynamics protein 3 OS=Rattus norvegicus OX=10116 GN=Rmdn3 PE=1 SV=1 |
| | **0** | Q6MGB5\|DHB8_RAT | **+** | 504990.00 | **0.003173103** | Estradiol 17-beta-dehydrogenase 8 OS=Rattus norvegicus OX=10116 GN=Hsd17b8 PE=1 SV=1 |
| | **0** | Q63716\|PRDX1_RAT | **+** | 2002900.00 | **0.012585216** | Peroxiredoxin-1 OS=Rattus norvegicus OX=10116 GN=Prdx1 PE=1 SV=1 |
| | **0** | P04041\|GPX1_RAT | **+** | 856390.00 | **0.005381124** | Glutathione peroxidase 1 OS=Rattus norvegicus OX=10116 GN=Gpx1 PE=1 SV=4 |
| | **0** | tr\|B2RYM8\|B2RYM8_RA | **+** | 68923.00 | **0.000433077** | Family with sequence similarity 210, member B OS=Rattus norvegicus OX=10116 GN=Fam210b PE=1 SV=1 |
| | **0** | tr\|Q6IEA8\|Q6IEA8_RAT | **+** | 1211800.00 | **0.007614342** | Interferon, alpha-inducible protein 27-like 2B OS=Rattus norvegicus OX=10116 GN=Ifi27l2b PE=2 SV=1 |
| | **0** | Q09073\|ADT2_RAT | **+** | 9141500.00 | **0.057440588** | ADP/ATP translocase 2 OS=Rattus norvegicus OX=10116 GN=Slc25a5 PE=1 SV=3 |
| | **0** | Q9Z2L0\|VDAC1_RAT | **+** | 3497000.00 | **0.021973389** | Voltage-dependent anion-selective channel protein 1 OS=Rattus norvegicus OX=10116 GN=Vdac1 PE=1 SV=4 |
| | | P81155\|VDAC2_RAT | **+** | 786220.00 | **0.004940211** | Voltage-dependent anion-selective channel protein 2 OS=Rattus norvegicus OX=10116 GN=Vdac2 PE=1 SV=2 |
