## Supplementary material for "Permeability transition pore-related changes in the proteome and channel activity of ATP synthase dimers and monomers": ATP-synthase

| Protein Group | Protein ID | -10lgP | Coverage (%) | Coverage (%) Sample 1 | Area (IBAQ) | riBAQ | #Peptides | #Unique | #Spec Sample 1 | PTM | Avg. Mass | Accession | riBAQ PTP/Control | Protein Group | Protein ID | -10lgP | Coverage (%) | Coverage (%) Sample 1 | Area (IBAQ) | riBAQ | #Peptides | #Unique | #Spec Sample 1 | PTM | Avg. Mass | Description |
|---|---|---|---|---|---|---|---|---|---|---|---|---|---|---|---|---|---|---|---|---|---|---|---|---|---|---|
| | | | | | CONTROL | % | | | | | | | | | | | | | PTP | % | | | | | | |
| Complex I | | | | | | | | | | | | | | | | | | | | | | | | | | |
| 11 | 10 | 254 | 21 | 21 | 32389000 | 1.37376131 | 42 | 42 | 67 | | 79412 | Q66HF1\|NDUS1_RAT | 0.523734108 | 12 | 13 | 210.5 | 20 | 20 | 13976000 | 0.719485655 | 31 | 31 | 42 | | 79412 | NADH-ubiquinone oxidoreductase 75 kDa subunit mitochondrial OS=Rattus norvegicus OX=10116 GN=Ndufs1 PE=1 SV=1 |
| 15 | 30 | 232 | 36 | 36 | 26646000 | 1.130175179 | 24 | 24 | 47 | | 27378 | P19234\|NDUV2_RAT | 1.070938629 | 11 | 20 | 218.1 | 39 | 39 | 23511000 | 1.210348256 | 29 | 29 | 49 | Formyl | 27378 | NADH dehydrogenase [ubiquinone] flavoprotein 2 mitochondrial OS=Rattus norvegicus OX=10116 GN=Ndufv2 PE=1 SV=2 |
| 16 | 26 | 214 | 28 | 28 | 13080000 | 0.55478088 | 23 | 23 | 45 | Oxidatic | 52562 | Q641Y2\|NDUS2_RAT | 1.02295603 | 17 | 19 | 183.8 | 28 | 28 | 11024000 | 0.567516447 | 23 | 23 | 31 | Oxidati | 52562 | NADH dehydrogenase [ubiquinone] iron-sulfur protein 2 mitochondrial OS=Rattus norvegicus OX=10116 GN=Ndufs2 PE=1 SV=1 |
| 17 | 9 | 223 | 36 | 36 | 8186200 | 0.347213092 | 28 | 25 | 41 | | 42559 | Q5BK63\|NDUA9_RAT | 0.88350549 | 18 | 7 | 182.6 | 32 | 32 | 5958900 | 0.306764673 | 22 | 21 | 29 | | 42559 | NADH dehydrogenase [ubiquinone] 1 alpha subcomplex subunit 9 mitochondrial OS=Rattus norvegicus OX=10116 GN=Ndufa9 PE=1 SV=2 |
| 20 | 24 | 165 | 34 | 34 | 9942300 | 0.421697091 | 20 | 20 | 30 | | 30226 | tr\|D3ZG43\|D3ZG43_RAT | 0.861397029 | 20 | 28 | 157.3 | 33 | 33 | 7056100 | 0.363248621 | 18 | 18 | 24 | | 30226 | NADH dehydrogenase (Ubiquinone) Fe-S protein 3 (Predicted) isoform CRA_c OS=Rattus norvegicus OX=10116 GN=Ndufs3 PE=1 SV=1 |
| 27 | 138 | 139 | 15 | 15 | 3192400 | 0.13540386 | 13 | 12 | 21 | Oxidation (M) | | P05508\|NU4M_RAT | 0.259484165 | 40 | 147 | 82.09 | 9 | 9 | 682500 | 0.035135157 | 6 | 6 | 9 | | 51783 | NADH-ubiquinone oxidoreductase chain 4 |
| 28 | 37 | 173 | 14 | 14 | 6381700 | 0.270676234 | 15 | 15 | 21 | | 68618 | P11661\|NU5M_RAT | - | | | | | | | 0 | | | | | | NADH-ubiquinone oxidoreductase chain 5 OS=Rattus norvegicus OX=10116 GN=Mtnd5 PE=3 SV=3 |
| 29 | 23 | 141 | 14 | 14 | 12894000 | 0.546891794 | 10 | 10 | 21 | | 50731 | tr\|Q5XIH3\|Q5XIH3_RAT | 0.789514331 | 28 | 43 | 115.5 | 11 | 11 | 8387300 | 0.431778909 | 8 | 8 | 16 | Formyl | 50731 | NADH dehydrogenase [ubiquinone] flavoprotein 1 mitochondrial OS=Rattus norvegicus OX=10116 GN=Ndufv1 PE=1 SV=1 |
| 33 | 42 | 155 | 35 | 35 | 5591900 | 0.237177309 | 12 | 12 | 19 | | 20859 | tr\|D4A0T0\|D4A0T0_RAT | 0.98963217 | 35 | 66 | 129 | 33 | 33 | 4559400 | 0.234718295 | 8 | 8 | 11 | Oxidati | 20859 | NADH:ubiquinone oxidoreductase subunit B10 OS=Rattus norvegicus OX=10116 GN=Ndufb10 PE=1 SV=1 |
| 36 | 28 | 115 | 17 | 17 | 2303820 | 0.097715236 | 11 | 3 | 17 | Oxidatic | 40493 | Q561S0\|NDUAA_RAT | 1.860264418 | 44 | 57 | 113.3 | 15 | 15 | 3531000 | 0.181776177 | 7 | 7 | 9 | Oxidati | 40493 | NADH dehydrogenase [ubiquinone] 1 alpha subcomplex subunit 10 mitochondrial OS=Rattus norvegicus OX=10116 GN=Ndufa10 PE=1 SV=1 |
| 39 | 183 | 175 | 32 | 32 | 6257400 | 0.265404119 | 9 | 9 | 14 | | 17634 | tr\|D4A7L4\|D4A7L4_RAT | 1.343523993 | 39 | 137 | 138.7 | 26 | 26 | 6926500 | 0.356576802 | 7 | 7 | 10 | | 17634 | NADH dehydrogenase (Ubiquinone) 1 beta subcomplex 11 (Predicted) OS=Rattus norvegicus OX=10116 GN=Ndufb11 PE=1 SV=1 |
| 40 | 242 | 151 | 21 | 21 | 6208400 | 0.263325812 | 9 | 9 | 14 | | 14854 | Q80W89\|NDUAB_RAT | 0.95959011 | 49 | 181 | 125.7 | 18 | 18 | 4908400 | 0.252684845 | 6 | 6 | 8 | | 14854 | NADH dehydrogenase [ubiquinone] 1 alpha subcomplex subunit 11 OS=Rattus norvegicus OX=10116 GN=Ndufa11 PE=2 SV=1 |
| 42 | 135 | 152 | 20 | 20 | 3395700 | 0.144026715 | 13 | 11 | 13 | | 23970 | tr\|B0BNE6\|B0BNE6_RAT | 0.534257006 | 50 | 55 | 125.8 | 16 | 16 | 1494700 | 0.076947282 | 7 | 6 | 8 | | 23970 | NADH dehydrogenase (Ubiquinone) Fe-S protein 8 (Predicted) isoform CRA_a OS=Rattus norvegicus OX=10116 GN=Ndufs8 PE=1 SV=1 |
| 44 | 110 | 125 | 19 | 19 | 3866200 | 0.163982709 | 6 | 6 | 13 | | 19965 | tr\|A0A0G2JVL6\|A0A0G2JVL6_ | 0.485533495 | 37 | 34 | 110 | 25 | 25 | 1546600 | 0.079619098 | 6 | 6 | 11 | | 19965 | NADH dehydrogenase [ubiquinone] 1 alpha subcomplex subunit 8 OS=Rattus norvegicus OX=10116 GN=Ndufa8 PE=1 SV=1 |
| 45 | 45 | 188 | 25 | 25 | 3771400 | 0.15996182 | 8 | 8 | 12 | | 21959 | tr\|B2RYS8\|B2RYS8_RAT | 1.488805367 | 32 | 49 | 187 | 21 | 21 | 4626100 | 0.238152017 | 11 | 11 | 15 | Oxidati | 21959 | NADH dehydrogenase [ubiquinone] 1 beta subcomplex subunit 8 mitochondrial OS=Rattus norvegicus OX=10116 GN=Ndufb8 PE=1 SV=1 |
| 47 | 246 | 167 | 27 | 27 | 10909000 | 0.46269913 | 7 | 7 | 12 | | 14359 | tr\|Q5PQZ9\|Q5PQZ9_RAT | 0.687299911 | 34 | 204 | 160.2 | 30 | 30 | 6177400 | 0.318013071 | 10 | 10 | 14 | | 14359 | NADH dehydrogenase [ubiquinone] 1 subunit C2 OS=Rattus norvegicus OX=10116 GN=Ndufc2 PE=1 SV=1 |
| 48 | 254 | 83.4 | 12 | 12 | 1390100 | 0.058960314 | 6 | 6 | 11 | | 38653 | P11662\|NU2M_RAT | - | | | | | | | 0 | | | | | | NADH-ubiquinone oxidoreductase chain 2 OS=Rattus norvegicus OX=10116 GN=Mtnd2 PE=3 SV=3 |
| 50 | 172 | 77.1 | 18 | 18 | 1749900 | 0.074221029 | 8 | 8 | 11 | Oxidatic | 36145 | P03889\|NU1M_RAT | 0.557492031 | 77 | 226 | 53.86 | 7 | 7 | 803760 | 0.041377632 | 3 | 3 | 3 | | 36145 | NADH-ubiquinone oxidoreductase chain 1 OS=Rattus norvegicus OX=10116 GN=Mtnd1 PE=1 SV=3 |
| 51 | 165 | 154 | 24 | 24 | 1724700 | 0.073152185 | 6 | 6 | 10 | | 21664 | tr\|D4A565\|D4A565_RAT | 2.343522987 | 41 | 58 | 147.3 | 21 | 21 | 3330100 | 0.171433828 | 7 | 7 | 9 | | 21664 | NADH dehydrogenase (Ubiquinone) 1 beta subcomplex 5 (Predicted) isoform CRA_b OS=Rattus norvegicus OX=10116 GN=Ndufb5 PE=1 SV=1 |
| 55 | 111 | 113 | 12 | 12 | 3596400 | 0.152539293 | 5 | 5 | 8 | | 19741 | Q5XIF3\|NDUS4_RAT | 0.850333162 | 38 | 138 | 115 | 22 | 22 | 2519600 | 0.12970922 | 8 | 8 | 10 | | 19741 | NADH dehydrogenase [ubiquinone] iron-sulfur protein 4 mitochondrial OS=Rattus norvegicus OX=10116 GN=Ndufs4 PE=1 SV=1 |
| 57 | 175 | 124 | 21 | 21 | 2150400 | 0.091208013 | 5 | 5 | 8 | | 13040 | tr\|D3ZCZ9\|D3ZCZ9_RAT | 0.968779099 | 57 | 142 | 118.5 | 21 | 21 | 1716400 | 0.088360416 | 5 | 5 | 7 | | 13040 | NADH dehydrogenase [ubiquinone] iron-sulfur protein 6 mitochondrial OS=Rattus norvegicus OX=10116 GN=LOC100912599 PE=1 SV=1 |
| 64 | 133 | 132 | 19 | 19 | 2143500 | 0.090915353 | 4 | 4 | 7 | | 15064 | tr\|F1LPG5\|F1LPG5_RAT | 0.495048324 | 126 | 145 | 80.25 | 11 | 11 | 874270 | 0.045007493 | 2 | 2 | 2 | | 15064 | NADH:ubiquinone oxidoreductase subunit B4 OS=Rattus norvegicus OX=10116 GN=Ndufb4 PE=1 SV=1 |
| 69 | 148 | 72.8 | 14 | 14 | 1523200 | 0.064605676 | 4 | 4 | 6 | Oxidatic | 23945 | tr\|Q5RJN0\|Q5RJN0_RAT | 0.648416772 | 59 | 194 | 95.2 | 15 | 15 | 813740 | 0.041891404 | 5 | 5 | 5 | Oxidati | 23945 | NADH dehydrogenase (Ubiquinone) Fe-S protein 7 OS=Rattus norvegicus OX=10116 GN=Ndufs7 PE=1 SV=1 |
| 70 | 168 | 114 | 21 | 21 | 1699500 | 0.072083341 | 4 | 4 | 6 | | 17178 | tr\|F1LXA0\|F1LXA0_RAT | 1.120754037 | 62 | 249 | 101.4 | 21 | 21 | 1569300 | 0.080787696 | 4 | 4 | 5 | | 17178 | NADH dehydrogenase [ubiquinone] 1 alpha subcomplex subunit 12 OS=Rattus norvegicus OX=10116 GN=Ndufa12 PE=1 SV=2 |
| 78 | 178 | 77.4 | 12 | 12 | 702550 | 0.029798265 | 3 | 3 | 5 | | 15638 | tr\|D3ZZ21\|D3ZZ21_RAT | 3.003640442 | 60 | 135 | 83.7 | 27 | 27 | 1738600 | 0.089503274 | 4 | 4 | 5 | | 15638 | NADH dehydrogenase (Ubiquinone) 1 beta subcomplex 6 (Predicted) OS=Rattus norvegicus OX=10116 GN=Ndufb6 PE=1 SV=1 |
| 81 | 112 | 102 | 28 | 28 | 789400 | 0.033481959 | 4 | 4 | 4 | | 15224 | tr\|D4A3V2\|D4A3V2_RAT | 2.329998636 | 53 | 41 | 112 | 29 | 29 | 1515400 | 0.078012919 | 7 | 7 | 7 | Oxidati | 15224 | NADH dehydrogenase [ubiquinone] 1 alpha subcomplex subunit 6 OS=Rattus norvegicus OX=10116 GN=Ndufa6 PE=1 SV=1 |
| 84 | 286 | 38.6 | 11 | 11 | 1823200 | 0.077330008 | 1 | 1 | 4 | | 12500 | tr\|A9UMV9\|A9UMV9_RAT | 1.5730283 | 63 | 245 | 71.05 | 19 | 19 | 2362900 | 0.121642291 | 2 | 2 | 5 | | 12500 | NADH:ubiquinone oxidoreductase subunit A7 OS=Rattus norvegicus OX=10116 GN=Ndufa7 PE=1 SV=1 |
| 89 | 196 | 102 | 18 | 18 | 2143700 | 0.090923836 | 4 | 4 | 4 | | 13412 | Q63362\|NDUA5_RAT | 0.288026053 | 83 | 133 | 44.2 | 22 | 22 | 508710 | 0.026188434 | 3 | 2 | 3 | | 13412 | NADH dehydrogenase [ubiquinone] 1 alpha subcomplex subunit 5 OS=Rattus norvegicus OX=10116 GN=Ndufa5 PE=1 SV=3 |
| 152 | 282 | 52.4 | 29 | 29 | 1471600 | 0.06241709 | 2 | 2 | 2 | Oxidatic | 10170 | tr\|A0A0G2KAA3\|A0A0G2KAA | - | 106 | 259 | 76.64 | 29 | 29 | 0 | 0 | 2 | 2 | 2 | Oxidati | 10170 | NADH:ubiquinone oxidoreductase subunit A3 OS=Rattus norvegicus OX=10116 GN=Ndufa3 PE=1 SV=1 |
| 157 | 119 | 56.6 | 7 | 7 | 173420 | 0.007355512 | 2 | 2 | 2 | | 16777 | tr\|D3ZE15\|D3ZE15_RAT | 4.511664918 | 88 | 65 | 69.36 | 19 | 19 | 644630 | 0.033185607 | 3 | 3 | 3 | Oxidati | 16777 | NADH:ubiquinone oxidoreductase subunit A13 OS=Rattus norvegicus OX=10116 GN=Ndufa13 PE=1 SV=1 |
| 260 | 243 | 49.6 | 11 | 11 | 274870 | 0.011658457 | 1 | 1 | 1 | | 10845 | tr\|D3ZS58\|D3ZS58_RAT | 2.837210598 | 85 | 346 | 71.84 | 23 | 23 | 642530 | 0.033077498 | 3 | 3 | 3 | | 10845 | NADH dehydrogenase [ubiquinone] 1 alpha subcomplex subunit 2 OS=Rattus norvegicus OX=10116 GN=Ndufa2 PE=1 SV=1 |
| 262 | 350 | 39.5 | 13 | 13 | 155530 | 0.006596718 | 1 | 1 | 1 | | 8389 | tr\|Q35733\|Q35733_RAT | 0.718379612 | 268 | 5783 | 32.1 | 13 | 13 | 92054 | 0.004738948 | 1 | 1 | 1 | | 8389 | NADH-ubiquinone oxidoreductase chain 6 (Fragment) OS=Rattus norvegicus OX=10116 PE=3 SV=1 |
| 263 | 5949 | 39.9 | 9 | 9 | 450630 | 0.019113219 | 1 | 1 | 1 | | 11842 | tr\|B2RYU0\|B2RYU0_RAT | 0.653371816 | 266 | 5819 | 32.86 | 9 | 9 | 242580 | 0.012488039 | 1 | 1 | 1 | | 11842 | NADH dehydrogenase (Ubiquinone) 1 beta subcomplex 2 (Predicted) isoform CRA_b OS=Rattus norvegicus OX=10116 GN=Ndufb2 PE=1 SV=1 |
| 273 | 192 | 23.5 | 14 | 14 | 289150 | 0.012264135 | 1 | 1 | 1 | | 12700 | tr\|B5DEL8\|B5DEL8_RAT | - | | | | | | | 0 | | | | | | NADH dehydrogenase (Ubiquinone) Fe-S protein 5 OS=Rattus norvegicus OX=10116 GN=Ndufs5 PE=1 SV=1 |
| | | | | | | 0 | | | | | | tr\|Q06QG6\|Q06QG6_RAT | + | 42 | 182 | 101.3 | 8 | 8 | 1353800 | 0.069693738 | 7 | 7 | 9 | | 68588 | NADH-ubiquinone oxidoreductase chain 5 OS=Rattus norvegicus OX=10116 GN=ND5 PE=3 SV=1 |
| | | | | | | 0 | | | | | | tr\|B2RYW3\|B2RYW3_RAT | + | 114 | 235 | 46.63 | 13 | 13 | 175610 | 0.009040418 | 2 | 2 | 2 | Oxidati | 21892 | NADH dehydrogenase (Ubiquinone) 1 beta subcomplex 9 OS=Rattus norvegicus OX=10116 GN=Ndufb9 PE=1 SV=1 |
| | | | | | | 0 | | | | | | tr\|D4A4P3\|D4A4P3_RAT | + | 134 | 414 | 30.54 | 15 | 15 | 586050 | 0.030169903 | 2 | 2 | 2 | | 11267 | NADH:ubiquinone oxidoreductase subunit B3 OS=Rattus norvegicus OX=10116 GN=Ndufb3 PE=1 SV=1 |
| 24 | 164 | 184 | 23 | 23 | 10259000 | 0.435129744 | 13 | 13 | 25 | Oxidatic | 17514 | tr\|D3ZF13\|D3ZF13_RAT | 0.939805199 | 22 | 48 | 163.1 | 29 | 29 | 7943600 | 0.408937196 | 13 | 13 | 21 | Oxidati | 17514 | Acyl carrier protein OS=Rattus norvegicus OX=10116 GN=Ndufab1 PE=1 SV=1 |
| Complex III | | | | | | | | | | | | | | | | | | | | | | | | | | |
| 26 | 12 | 173 | 15 | 15 | 6311700 | 0.267707223 | 12 | 12 | 22 | | 52849 | Q68FY0\|QCR1_RAT | 0.750296691 | 25 | 16 | 154.2 | 16 | 16 | 3901700 | 0.200859844 | 15 | 15 | 18 | Carbam | 52849 | Cytochrome b-c1 complex subunit 1 mitochondrial OS=Rattus norvegicus OX=10116 GN=Uqcrc1 PE=1 SV=1 |
| 30 | 13 | 169 | 15 | 15 | 8206100 | 0.348057139 | 13 | 13 | 20 | | 48396 | P32551\|QCR2_RAT | 0.749355977 | 33 | 21 | 108.6 | 11 | 11 | 5066400 | 0.260818698 | 10 | 10 | 14 | | 48396 | Cytochrome b-c1 complex subunit 2 mitochondrial OS=Rattus norvegicus OX=10116 GN=Uqcrc2 PE=1 SV=2 |
| 68 | 65 | 107 | 14 | 14 | 1254700 | 0.053217398 | 5 | 5 | 6 | | 43012 | P00159\|CYB_RAT | 0.822551113 | 73 | 97 | 81.26 | 9 | 9 | 850310 | 0.04377403 | 4 | 4 | 4 | | 43012 | Cytochrome b OS=Rattus norvegicus OX=10116 GN=Mt-Cyb PE=3 SV=3 |
| 72 | 80 | 105 | 11 | 11 | 1846300 | 0.078309781 | 3 | 3 | 5 | | 29446 | P20788\|UCRI_RAT | 0.506979335 | 67 | 42 | 97.39 | 8 | 8 | 771200 | 0.039701441 | 3 | 3 | 4 | | 29446 | Cytochrome b-c1 complex subunit Rieske mitochondrial OS=Rattus norvegicus OX=10116 GN=Uqcrfs1 PE=1 SV=2 |
| | | | | | | 0 | | | | | | Q5M9I5\|QCR6_RAT | + | 279 | 338 | 24.93 | 17 | 17 | 126470 | 0.006510686 | 1 | 1 | 1 | | 10424 | Cytochrome b-c1 complex subunit 6 mitochondrial OS=Rattus norvegicus OX=10116 GN=Uqcrh PE=3 SV=1 |
| Complex IV | | | | | | | | | | | | | | | | | | | | | | | | | | |
| 41 | 82 | 136 | 26 | 26 | 3391900 | 0.14386554 | 9 | 9 | 13 | Oxidatic | 25928 | P00406\|COX2_RAT | 0.756319445 | 52 | 112 | 91.71 | 29 | 29 | 2113600 | 0.108808306 | 7 | 7 | 7 | Oxidation (M) | | Cytochrome c oxidase subunit 2 OS=Rattus norvegicus OX=10116 GN=Mtco2 PE=1 SV=3 |
| 52 | 179 | 137 | 33 | 33 | 5215100 | 0.221195548 | 6 | 6 | 10 | | 16130 | P11240\|COX5A_RAT | 0.578650584 | 54 | 44 | 126.6 | 31 | 31 | 2486300 | 0.127994933 | 5 | 5 | 7 | | 16130 | Cytochrome c oxidase subunit 5A mitochondrial OS=Rattus norvegicus OX=10116 GN=Cox5a PE=1 SV=1 |
| 53 | 158 | 126 | 11 | 11 | 1725900 | 0.073203083 | 6 | 6 | 9 | Oxidation (M) | | P05503\|COX1_RAT | 0.245040507 | 101 | 233 | 49.91 | 4 | 4 | 348440 | 0.01793772 | 2 | 2 | 2 | | 56845 | Cytochrome c oxidase subunit 1 |
| 59 | 22 | 159 | 9 | 9 | 4889200 | 0.207372682 | 5 | 5 | 8 | | 35435 | tr\|D3ZFQ8\|D3ZFQ8_RAT | 1.042944167 | 48 | 38 | 132.4 | 9 | 9 | 4201200 | 0.216278129 | 4 | 4 | 8 | | 35435 | Cytochrome c-1 OS=Rattus norvegicus OX=10116 GN=Cyc1 PE=1 SV=3 |
| 73 | 220 | 43.6 | 22 | 22 | 3188400 | 0.135234202 | 4 | 4 | 5 | | 13915 | P12075\|COX5B_RAT | 0.424374877 | 68 | 322 | 44.85 | 28 | 28 | 1114800 | 0.057389998 | 4 | 4 | 4 | | 13915 | Cytochrome c oxidase subunit 5B mitochondrial OS=Rattus norvegicus OX=10116 GN=Cox5b PE=1 SV=2 |
| 74 | 182 | 74.4 | 17 | 17 | 1155700 | 0.049018369 | 4 | 4 | 5 | | 19515 | P10888\|COX41_RAT | 0.408094589 | 92 | 247 | 50.93 | 12 | 12 | 388580 | 0.020004131 | 2 | 2 | 3 | | 19515 | Cytochrome c oxidase subunit 4 isoform 1 mitochondrial OS=Rattus norvegicus OX=10116 GN=Cox4i1 PE=1 SV=1 |
| 143 | 347 | 29.8 | 14 | 14 | 243120 | 0.010311799 | 1 | 1 | 2 | | 7672 | P80433\|COX8A_RAT | 1.035812287 | 270 | 361 | 31.69 | 14 | 14 | 207480 | 0.010681088 | 1 | 1 | 1 | | 7672 | Cytochrome c oxidase subunit 8A mitochondrial OS=Rattus norvegicus OX=10116 GN=Cox8a PE=1 SV=3 |
| 151 | 283 | 82.7 | 29 | 29 | 689260 | 0.029234577 | 2 | 2 | 2 | | 8995 | P80431\|COX7B_RAT | - | | | | | | | 0 | | | | | | Cytochrome c oxidase subunit 7B mitochondrial OS=Rattus norvegicus OX=10116 GN=Cox7b PE=1 SV=3 |
| Complex V | | | | | | | | | | | | | | | | | | | | | | | | | | |
| 1 | 20 | 358 | 57 | 57 | 613900000 | 26.03822496 | 160 | 156 | 494 | Carbamidomet | | P15999\|ATPA_RAT | 0.900784947 | 2 | 10 | 326.8 | 60 | 60 | 455610000 | 23.45484109 | 159 | 157 | 432 | Carbamidometl | | ATP synthase subunit alpha, mitochondrial |
| 2 | 17 | 398 | 64 | 64 | 657620000 | 27.89258429 | 183 | 180 | 407 | Oxidation (M) | | P10719\|ATPB_RAT | 0.777888158 | 3 | 8 | 359.3 | 65 | 65 | 421470000 | 21.69731102 | 177 | 174 | 322 | Oxidati | 56354 | ATP synthase subunit beta, mitochondrial |
| 8 | 268 | 226 | 30 | 30 | 81333000 | 3.44969368 | 32 | 32 | 87 | | 28869 | P19511\|AT5F1_RAT | 0.715069631 | 10 | 246 | 200 | 30 | 30 | 47917000 | 2.466771187 | 27 | 27 | 54 | | 28869 | ATP synthase F(0) complex subunit B1 mitochondrial OS=Rattus norvegicus OX=10116 GN=Atp5pb PE=1 SV=1 |
| 9 | 301 | 260 | 40 | 40 | 74174000 | 3.146048702 | 35 | 35 | 84 | Oxidatic | 30191 | P35435\|ATPG_RAT | 0.786196386 | 6 | 64 | 245 | 37 | 37 | 48046000 | 2.473412118 | 36 | 36 | 82 | Oxidati | 30191 | ATP synthase subunit gamma mitochondrial OS=Rattus norvegicus OX=10116 GN=Atp5f1c PE=1 SV=2 |
| 10 | 280 | 309 | 71 | 71 | 68444000 | 2.903013958 | 49 | 49 | 82 | Oxidatic | 18763 | P31399\|ATP5H_RAT | 0.605681645 | 8 | 119 | 264.4 | 61 | 61 | 34155000 | 1.758302271 | 36 | 35 | 59 | Carbamidometl | | ATP synthase subunit d mitochondrial OS=Rattus norvegicus OX=10116 GN=Atp5pd PE=1 SV=3 |
| 14 | 5938 | 189 | 56 | 56 | 25753000 | 1.092299083 | 30 | 30 | 51 | Carbamidomet | | Q06647\|ATPO_RAT | 0.690926179 | 14 | 5802 | 162.5 | 45 | 45 | 14660000 | 0.754698032 | 21 | 20 | 36 | Carbamidometl | | ATP synthase subunit O, mitochondrial |
| 18 | 5937 | 190 | 39 | 39 | 27382000 | 1.161392207 | 13 | 13 | 32 | Oxidatic | 11433 | Q6PDU7\|ATP5L_RAT | 0.551018787 | 19 | 1646 | 137.9 | 39 | 39 | 12431000 | 0.639948925 | 13 | 13 | 25 | Oxidati | 11433 | ATP synthase subunit g mitochondrial OS=Rattus norvegicus OX=10116 GN=Atp5mg PE=1 SV=2 |
| 21 | 387 | 194 | 36 | 36 | 21029000 | 0.891933267 | 16 | 16 | 30 | Oxidatic | 17563 | tr\|G3V7Y3\|G3V7Y3_RAT | 0.904258768 | 21 | 350 | 167.8 | 30 | 30 | 15667000 | 0.806538477 | 14 | 14 | 22 | Oxidati | 17563 | ATP synthase subunit delta mitochondrial OS=Rattus norvegicus OX=10116 GN=Atp5f1d PE=1 SV=1 |
| 22 | 5942 | 122 | 32 | 32 | 9381500 | 0.397911073 | 13 | 13 | 29 | Oxidation (M) | | P05504\|ATP6_RAT | 0.092111725 | 43 | 690 | 64.28 | 18 | 18 | 711970 | 0.036652275 | 5 | 5 | 9 | Oxidation (M) | | ATP synthase subunit a |
| 43 | 416 | 156 | 48 | 48 | 14820000 | 0.628582007 | 7 | 7 | 13 | Oxidatic | 7642 | tr\|Q5UAJ5\|Q5UAJ5_RAT | 0.5929388 | 46 | 1095 | 89.84 | 30 | 30 | 7239900 | 0.372710661 | 5 | 4 | 9 | Oxidati | 7642 | ATP synthase protein 8 OS=Rattus norvegicus OX=10116 GN=ATP8 PE=3 SV=1 |
| 61 | #### | 79.6 | 27 | 27 | 3040200 | 0.128948382 | 3 | 3 | 8 | | 8255 | P29419\|ATP5I_RAT | 1.336063679 | 47 | 11740 | 85.39 | 34 | 34 | 3346600 | 0.172283249 | 4 | 4 | 9 | | 8255 | ATP synthase subunit e mitochondrial OS=Rattus norvegicus OX=10116 GN=Atp5me PE=1 SV=3 |
| 106 | #### | 63.5 | 19 | 19 | 844150 | 0.03580415 | 2 | 2 | 3 | | 12494 | P21571\|ATP5J_RAT | 0.780451141 | 91 | 5817 | 54.59 | 19 | 19 | 542800 | 0.02794339 | 2 | 2 | 3 | | 12494 | ATP synthase-coupling factor 6 mitochondrial OS=Rattus norvegicus OX=10116 GN=Atp5pf PE=1 SV=1 |
| 109 | #### | 44.5 | 21 | 21 | 1835600 | 0.077855947 | 1 | 1 | 3 | | 6408 | Q9JJW3\|ATPMD_RAT | 0.543008933 | 95 | 11746 | 36.62 | 21 | 21 | 821220 | 0.042276475 | 2 | 2 | 3 | | 6408 | ATP synthase membrane subunit DAPIT mitochondrial OS=Rattus norvegicus OX=10116 GN=Atp5md PE=1 SV=1 |
| 111 | #### | 30 | 10 | 10 | 1286700 | 0.05457466 | 1 | 1 | 3 | | 10452 | D3ZAF6\|ATPK_RAT | 0.338973583 | 131 | 11756 | 25.19 | 14 | 14 | 359350 | 0.018499368 | 2 | 1 | 2 | | 10452 | ATP synthase subunit f mitochondrial OS=Rattus norvegicus OX=10116 GN=Atp5mf PE=1 SV=1 |
| Others | | | | | | | | | | | | | | | | | | | | | | | | | | |
| 3 | 1 | 326 | 51 | 51 | 81132000 | 3.441168378 | 90 | 90 | 162 | Carbamidomet | | Q02253\|MMSA_RAT | 0.027939407 | 30 | 50 | 140.3 | 17 | 17 | 1867600 | 0.096144205 | 15 | 15 | 16 | Carbamidometl | | Methylmalonate-semialdehyde dehydrogenase [acylating], mitochondrial |
| 4 | 8 | 321 | 34 | 34 | 73977000 | 3.137693057 | 96 | 96 | 149 | Oxidatic | 2E+05 | P07756\|CPSM_RAT | 0.802186735 | 4 | 4 | 257.2 | 30 | 30 | 48893000 | 2.51701575 | 79 | 79 | 115 | Oxidati | 2E+05 | Carbamoyl-phosphate synthase [ammonia] mitochondrial OS=Rattus norvegicus OX=10116 GN=Cps1 PE=1 SV=1 |

| | | | | | | | | | | | | | | | | | | | | | | | | | | |
|---|---|---|---|---|---|---|---|---|---|---|---|---|---|---|---|---|---|---|---|---|---|---|---|---|---|---|
| **5** | 11 | 284 | 62 | 62 | 104200000 | **4.419584689** | 68 | 66 | 131 | | 29820 | P67779\|PHB_RAT | **0.609141172** | 5 | 15 | 248.8 | 57 | 57 | 52295000 | **2.692150995** | 55 | 55 | 83 | | 29820 | Prohibitin OS=Rattus norvegicus OX=10116 GN=Phb PE=1 SV=1 |
| **6** | 19 | 272 | 61 | 61 | 79864000 | **3.387386867** | 67 | 64 | 129 | Oxidatic | 33312 | Q5XIH7\|PHB2_RAT | **0.511824111** | 7 | 12 | 225.5 | 48 | 48 | 33678000 | **1.733746271** | 46 | 45 | 71 | Oxidati | 33312 | Prohibitin-2 OS=Rattus norvegicus OX=10116 GN=Phb2 PE=1 SV=1 |
| **7** | 35 | 282 | 29 | 29 | 34267000 | **1.453415629** | 66 | 65 | 110 | Oxidatic | 1E+05 | P52873\|PYC_RAT | **0.596119787** | 9 | 30 | 222.8 | 24 | 24 | 16830000 | **0.866409815** | 41 | 41 | 55 | Oxidati | 1E+05 | Pyruvate carboxylase mitochondrial OS=Rattus norvegicus OX=10116 GN=Pc PE=1 SV=2 |
| **12** | 15 | 249 | 36 | 36 | 30717000 | **1.302844365** | 36 | 35 | 63 | Oxidatic | 47314 | P00507\|AATM_RAT | **0.582825746** | 15 | 26 | 196.5 | 27 | 27 | 14750000 | **0.75933124** | 23 | 23 | 33 | Oxidati | 47314 | Aspartate aminotransferase mitochondrial OS=Rattus norvegicus OX=10116 GN=Got2 PE=1 SV=2 |
| **13** | 14 | 263 | 37 | 37 | 21074000 | **0.893841917** | 44 | 44 | 61 | Oxidatic | 61416 | P10860\|DHE3_RAT | **28.32769292** | 1 | 1 | 384.1 | 68 | 68 | 491850000 | **25.32047934** | 228 | 225 | 442 | Carbamidometl | 61416 | Glutamate dehydrogenase 1 mitochondrial OS=Rattus norvegicus OX=10116 GN=Glud1 PE=1 SV=2 |
| **19** | 289 | 231 | 25 | 25 | 10034000 | **0.425586495** | 21 | 21 | 31 | | 60420 | tr\|D3ZFJ6\|D3ZFJ6_RAT | **0.545118329** | 26 | 171 | 177.7 | 22 | 22 | 4506500 | **0.231994999** | 15 | 15 | 17 | Carbam | 60420 | Lactamase beta OS=Rattus norvegicus OX=10116 GN=Lactb PE=1 SV=1 |
| **23** | 33 | 192 | 15 | 15 | 7333000 | **0.311025091** | 21 | 21 | 26 | Formyla | 82665 | Q64428\|ECHA_RAT | **1.065783496** | 23 | 24 | 145 | 12 | 12 | 6439100 | **0.331485409** | 15 | 15 | 20 | Formyl | 82665 | Trifunctional enzyme subunit alpha mitochondrial OS=Rattus norvegicus OX=10116 GN=Hadha PE=1 SV=2 |
| **25** | 76 | 202 | 25 | 25 | 7912400 | **0.335600018** | 20 | 20 | 24 | | 56886 | tr\|Q68G44\|Q68G44_RAT | **0.932209779** | 16 | 22 | 193.9 | 29 | 29 | 6077100 | **0.312849619** | 25 | 25 | 31 | | 56886 | 3-hydroxy-3-methylglutaryl coenzyme A synthase OS=Rattus norvegicus OX=10116 GN=Hmgcs2 PE=1 SV=1 |
| **31** | 169 | 171 | 18 | 18 | 4907600 | **0.208153108** | 15 | 15 | 19 | | 51414 | Q60587\|ECHB_RAT | **0.868606793** | 29 | 27 | 145.3 | 21 | 21 | 3512100 | **0.180803203** | 15 | 15 | 16 | | 51414 | Trifunctional enzyme subunit beta mitochondrial OS=Rattus norvegicus OX=10116 GN=Hadhb PE=1 SV=1 |
| **34** | 100 | 144 | 21 | 21 | 6214400 | **0.263580298** | 13 | 13 | 18 | | 44695 | P17764\|THIL_RAT | **1.102099892** | 24 | 35 | 139.4 | 22 | 22 | 5642800 | **0.290491818** | 13 | 13 | 19 | Formyl | 44695 | Acetyl-CoA acetyltransferase mitochondrial OS=Rattus norvegicus OX=10116 GN=Acat1 PE=1 SV=1 |
| **35** | 228 | 175 | 13 | 13 | 12428000 | **0.527126665** | 14 | 14 | 17 | | 75316 | P16970\|ABCD3_RAT | **0.082971424** | 90 | 307 | 45.09 | 5 | 5 | 849580 | **0.04373645** | 3 | 3 | 3 | | 75316 | ATP-binding cassette sub-family D member 3 OS=Rattus norvegicus OX=10116 GN=Abcd3 PE=1 SV=3 |
| **37** | 29 | 197 | 27 | 27 | 7993800 | **0.339052554** | 9 | 9 | 16 | | 24674 | P07895\|SODM_RAT | **0.230015018** | 31 | 32 | 164 | 28 | 28 | 1514900 | **0.077987179** | 10 | 10 | 15 | | 24674 | Superoxide dismutase [Mn] mitochondrial OS=Rattus norvegicus OX=10116 GN=Sod2 PE=1 SV=2 |
| **38** | 266 | 153 | 13 | 13 | 3087700 | **0.130963068** | 11 | 11 | 15 | | 59757 | P04762\|CATA_RAT | **0.5286255** | 36 | 45 | 127.9 | 9 | 9 | 1344800 | **0.069230417** | 9 | 9 | 11 | | 59757 | Catalase OS=Rattus norvegicus OX=10116 GN=Cat PE=1 SV=3 |
| **46** | 176 | 134 | 14 | 14 | 3371900 | **0.143017252** | 8 | 8 | 12 | | 36148 | P13086\|SUCA_RAT | **0.954282433** | 27 | 46 | 129.5 | 13 | 13 | 2651100 | **0.136478851** | 9 | 9 | 17 | | 36148 | Succinate--CoA ligase [ADP/GDP-forming] subunit alpha mitochondrial OS=Rattus norvegicus OX=10116 GN=Suclg1 PE=2 SV=2 |
| **54** | 245 | 96.9 | 8 | 8 | 1312200 | **0.055656229** | 7 | 7 | 9 | | 78179 | P18163\|ACSL1_RAT | **0.748518975** | 56 | 54 | 67.92 | 5 | 5 | 809240 | **0.041659743** | 5 | 3 | 7 | | 78179 | Long-chain-fatty-acid--CoA ligase 1 OS=Rattus norvegicus OX=10116 GN=Acsl1 PE=1 SV=1 |
| **56** | 244 | 104 | 13 | 13 | 2321300 | **0.09845664** | 5 | 5 | 8 | | 34973 | P07824\|ARGI1_RAT | **0.385460217** | 65 | 190 | 67.48 | 12 | 12 | 737200 | **0.037951118** | 3 | 3 | 4 | | 34973 | Arginase-1 OS=Rattus norvegicus OX=10116 GN=Arg1 PE=1 SV=2 |
| **58** | 231 | 80.8 | 14 | 14 | 2250800 | **0.095466422** | 6 | 6 | 8 | | 28340 | tr\|Q5M949\|Q5M949_RAT | **0.873581873** | 55 | 210 | 90.29 | 13 | 13 | 1620000 | **0.083397736** | 5 | 5 | 7 | | 28340 | Nipsnap homolog 3A (C. elegans) OS=Rattus norvegicus OX=10116 GN=Nipsnap3b PE=1 SV=1 |
| **60** | 261 | 154 | 10 | 10 | 1658800 | **0.070357074** | 5 | 5 | 8 | | 33407 | P24329\|THTR_RAT | **1.278348432** | 45 | 136 | 168.1 | 13 | 13 | 1747100 | **0.089940855** | 8 | 8 | 9 | | 33407 | Thiosulfate sulfurtransferase OS=Rattus norvegicus OX=10116 GN=Tst PE=1 SV=3 |
| **62** | 1940 | 147 | 7 | 7 | 2022400 | **0.085778964** | 6 | 6 | 7 | | 1E+05 | A2VCW9\|AASS_RAT | **1.651007476** | 51 | 742 | 130.8 | 9 | 9 | 2751000 | **0.141621711** | 7 | 7 | 7 | Formyl | 1E+05 | Alpha-aminoadipic semialdehyde synthase mitochondrial OS=Rattus norvegicus OX=10116 GN=Aass PE=2 SV=1 |
| **63** | 5947 | 96.8 | 3 | 3 | 1187700 | **0.050375631** | 4 | 4 | 7 | | 1E+05 | Q924S5\|LONM_RAT | **0.350009091** | 71 | 297 | 68.69 | 3 | 3 | 342500 | **0.017631929** | 3 | 3 | 4 | | 1E+05 | Lon protease homolog mitochondrial OS=Rattus norvegicus OX=10116 GN=Lonp1 PE=2 SV=1 |
| **65** | 198 | 113 | 10 | 10 | 911980 | **0.038681121** | 5 | 5 | 7 | | 49522 | P85834\|EFTU_RAT | **15.91737219** | 13 | 3 | 214.7 | 27 | 27 | 11960000 | **0.615701805** | 30 | 30 | 40 | Oxidati | 49522 | Elongation factor Tu mitochondrial OS=Rattus norvegicus OX=10116 GN=Tufm PE=1 SV=1 |
| **66** | 291 | 114 | 10 | 10 | 397670 | **0.016866951** | 5 | 5 | 7 | | 20541 | tr\|Q5XFW4\|Q5XFW4_RAT | **0.434439833** | 190 | 289 | 29.56 | 7 | 7 | 142340 | **0.007327675** | 1 | 1 | 1 | | 20541 | Mitochondrial ribosomal protein L13 OS=Rattus norvegicus OX=10116 GN=Mrpl13 PE=1 SV=1 |
| **67** | 219 | 112 | 18 | 18 | 1206500 | **0.051173022** | 5 | 5 | 6 | | 22305 | P04041\|GPX1_RAT | **0.179530842** | 173 | 53 | 32.37 | 5 | 5 | 178460 | **0.009187136** | 1 | 1 | 1 | | 22305 | Glutathione peroxidase 1 OS=Rattus norvegicus OX=10116 GN=Gpx1 PE=1 SV=4 |
| **71** | 1219 | 107 | 6 | 6 | 736680 | **0.03124587** | 5 | 5 | 5 | | 77891 | tr\|D3ZGM1\|D3ZGM1_RAT | **0.276908844** | 267 | 312 | 32.68 | 1 | 1 | 168070 | **0.008652258** | 1 | 1 | 1 | | 77891 | Pentatricopeptide repeat domain 3 OS=Rattus norvegicus OX=10116 GN=Ptcd3 PE=1 SV=1 |
| **75** | 194 | 73.9 | 17 | 17 | 1520200 | **0.064478432** | 4 | 4 | 5 | | 22179 | Q9R063\|PRDX5_RAT | **0.880324454** | 84 | 216 | 38.9 | 9 | 9 | 1102600 | **0.056761941** | 2 | 2 | 3 | | 22179 | Peroxiredoxin-5 mitochondrial OS=Rattus norvegicus OX=10116 GN=Prdx5 PE=1 SV=1 |
| **77** | 346 | 79.2 | 19 | 19 | 546530 | **0.023180764** | 3 | 3 | 5 | | 17050 | Q63750\|RM23_RAT | - | | | | | | | **0** | | | | | | 39S ribosomal protein L23 mitochondrial OS=Rattus norvegicus OX=10116 GN=Mrpl23 PE=2 SV=1 |
| **80** | 814 | 117 | 3 | 3 | 820970 | **0.034820983** | 4 | 4 | 4 | | 70749 | P45953\|ACADV_RAT | **0.526569324** | 98 | 201 | 74.08 | 3 | 3 | 356170 | **0.018335662** | 2 | 2 | 2 | | 70749 | Very long-chain specific acyl-CoA dehydrogenase mitochondrial OS=Rattus norvegicus OX=10116 GN=Acadvl PE=1 SV=1 |
| **82** | 3 | 67.5 | 4 | 4 | 555890 | **0.023577763** | 2 | 2 | 4 | | 53274 | tr\|B2GV15\|B2GV15_RAT | - | | | | | | | **0** | | | | | | Dihydrolipoamide acetyltransferase component of pyruvate dehydrogenase complex OS=Rattus norvegicus OX=10116 GN=Dbt PE=1 SV=1 |
| **85** | 753 | 52.4 | 8 | 8 | 454510 | **0.019277787** | 3 | 3 | 4 | | 17472 | P08011\|MGST1_RAT | **1.716689344** | 81 | 250 | 52.63 | 15 | 15 | 642850 | **0.033093972** | 3 | 3 | 3 | | 17472 | Microsomal glutathione S-transferase 1 OS=Rattus norvegicus OX=10116 GN=Mgst1 PE=1 SV=3 |
| **87** | 233 | 60.1 | 10 | 10 | 758880 | **0.032187471** | 4 | 3 | 4 | | 38202 | P29147\|BDH_RAT | **0.535409358** | 72 | 61 | 83.4 | 8 | 8 | 334760 | **0.017233473** | 4 | 3 | 4 | | 38202 | D-beta-hydroxybutyrate dehydrogenase mitochondrial OS=Rattus norvegicus OX=10116 GN=Bdh1 PE=1 SV=2 |
| **88** | 232 | 61.6 | 6 | 6 | 452810 | **0.019205683** | 2 | 2 | 4 | | 38374 | tr\|M0R6J0\|M0R6J0_RAT | **0.388506021** | 271 | 197 | 31.13 | 3 | 3 | 144940 | **0.007461523** | 1 | 1 | 1 | | 38374 | Mitochondrial ribosomal protein L39 OS=Rattus norvegicus OX=10116 GN=Mrpl39 PE=1 SV=2 |
| **90** | 816 | 60.3 | 5 | 5 | 149150 | **0.006326114** | 3 | 2 | 4 | | 44494 | tr\|Q5U2T0\|Q5U2T0_RAT | - | 66 | 236 | 34.01 | 4 | 4 | 0 | **0** | 2 | 1 | 4 | | 44494 | Death associated protein 3 OS=Rattus norvegicus OX=10116 GN=Dap3 PE=2 SV=1 |
| **91** | 290 | 69 | 8 | 8 | 1002300 | **0.042511994** | 3 | 3 | 4 | | 37440 | tr\|Q4G067\|Q4G067_RAT | **0.425178186** | 125 | 222 | 58.68 | 4 | 4 | 351110 | **0.018075172** | 2 | 2 | 2 | | 37440 | Mitochondrial ribosomal protein L44 OS=Rattus norvegicus OX=10116 GN=Mrpl44 PE=1 SV=1 |
| **95** | 380 | 49.7 | 9 | 9 | 182860 | **0.007755905** | 2 | 2 | 3 | | 27246 | O70351\|HCD2_RAT | **1.792731689** | 93 | 188 | 52.99 | 7 | 7 | 270090 | **0.013904256** | 2 | 2 | 3 | | 27246 | 3-hydroxyacyl-CoA dehydrogenase type-2 OS=Rattus norvegicus OX=10116 GN=Hsd17b10 PE=1 SV=3 |
| **100** | 348 | 83.8 | 9 | 9 | 307470 | **0.013041168** | 3 | 3 | 3 | | 45834 | P09139\|SPYA_RAT | - | 264 | 244 | 33.97 | 3 | 3 | 0 | **0** | 1 | 1 | 1 | | 45834 | Serine--pyruvate aminotransferase mitochondrial OS=Rattus norvegicus OX=10116 GN=Agxt PE=1 SV=1 |
| **101** | 6035 | 31.6 | 3 | 3 | 383140 | **0.016250669** | 2 | 2 | 3 | | 51496 | Q68FU7\|COQ6_RAT | - | | | | | | | **0** | | | | | | Ubiquinone biosynthesis monooxygenase COQ6 mitochondrial OS=Rattus norvegicus OX=10116 GN=Coq6 PE=2 SV=1 |
| **102** | 332 | 101 | 6 | 6 | 534430 | **0.022667549** | 3 | 3 | 3 | | 85433 | Q9ER34\|ACON_RAT | - | | | | | | | **0** | | | | | | Aconitate hydratase mitochondrial OS=Rattus norvegicus OX=10116 GN=Aco2 PE=1 SV=2 |
| **103** | 322 | 58 | 8 | 8 | 1652300 | **0.07008138** | 3 | 3 | 3 | | 32901 | Q09073\|ADT2_RAT | - | | | | | | | **0** | | | | | | ADP/ATP translocase 2 OS=Rattus norvegicus OX=10116 GN=Slc25a5 PE=1 SV=3 |
| **104** | 295 | 57.6 | 9 | 9 | 388650 | **0.016484372** | 3 | 3 | 3 | | 29822 | Q8VID1\|DHRS4_RAT | - | | | | | | | **0** | | | | | | Dehydrogenase/reductase SDR family member 4 OS=Rattus norvegicus OX=10116 GN=Dhrs4 PE=2 SV=2 |
| **105** | 264 | 73 | 3 | 3 | 600160 | **0.025455451** | 3 | 3 | 3 | Formyla | 68731 | P02770\|ALBU_RAT | - | | | | | | | **0** | | | | | | Serum albumin OS=Rattus norvegicus OX=10116 GN=Alb PE=1 SV=2 |
| **107** | 277 | 51.6 | 3 | 3 | 626630 | **0.026578161** | 2 | 2 | 3 | | 67177 | Q3KR86\|MIC60_RAT | **0.741922346** | 100 | 110 | 72.48 | 4 | 4 | 383040 | **0.019718931** | 2 | 2 | 2 | | 67177 | MICOS complex subunit Mic60 (Fragment) OS=Rattus norvegicus OX=10116 GN=Immt PE=1 SV=1 |
| **108** | 1292 | 68.4 | 5 | 5 | 337350 | **0.014308511** | 2 | 2 | 3 | | 37791 | tr\|G3V6I4\|G3V6I4_RAT | **0.828120449** | 80 | 207 | 59.34 | 7 | 7 | 230170 | **0.011849171** | 2 | 2 | 3 | | 37791 | Mitochondrial amidoxime reducing component 1 OS=Rattus norvegicus OX=10116 GN=Marc1 PE=1 SV=1 |
| **119** | 1285 | 31.5 | 6 | 6 | 7010000 | **0.297325227** | 2 | 2 | 2 | | 20718 | Q80ZG6\|BBC3_RAT | **0.087455043** | 119 | 1248 | 27.12 | 10 | 10 | 505100 | **0.02600259** | 2 | 2 | 2 | | 20718 | Bcl-2-binding component 3 OS=Rattus norvegicus OX=10116 GN=Bbc3 PE=1 SV=1 |
| **125** | 667 | 107 | 5 | 5 | 338140 | **0.014342019** | 2 | 2 | 2 | | 47388 | tr\|B2RZ24\|B2RZ24_RAT | - | 261 | 277 | 67.94 | 4 | 4 | 0 | **0** | 1 | 1 | 1 | | 47388 | Succinate-CoA ligase subunit beta (Fragment) OS=Rattus norvegicus OX=10116 GN=Sucla2 PE=2 SV=1 |
| **128** | 293 | 40.9 | 7 | 7 | 641240 | **0.027197836** | 2 | 2 | 2 | Oxidatic | 32433 | Q9WVK3\|PECR_RAT | **0.337581015** | 123 | 283 | 36.01 | 7 | 7 | 178350 | **0.009181473** | 2 | 2 | 2 | Oxidati | 32433 | Peroxisomal trans-2-enoyl-CoA reductase OS=Rattus norvegicus OX=10116 GN=Pecr PE=2 SV=1 |
| **129** | 1045 | 44 | 4 | 4 | 5658000 | **0.239980904** | 2 | 2 | 2 | | 51002 | Q2V057\|HYPDH_RAT | **0.28005249** | 117 | 306 | 50.95 | 4 | 4 | 1305500 | **0.06720725** | 2 | 2 | 2 | | 51002 | Hydroxyproline dehydrogenase OS=Rattus norvegicus OX=10116 GN=Prodh2 PE=2 SV=1 |
| **136** | 1546 | 31.7 | 12 | 12 | 61742 | **0.002618752** | 2 | 2 | 2 | Oxidatic | 23350 | Q9EPJ3\|RT26_RAT | - | | | | | | | **0** | | | | | | 28S ribosomal protein S26 mitochondrial OS=Rattus norvegicus OX=10116 GN=Mrps26 PE=1 SV=1 |
| **140** | 406 | 26.8 | 1 | 1 | 371920 | **0.015774779** | 1 | 1 | 2 | | 67300 | tr\|D3ZD23\|D3ZD23_RAT | **3.238542491** | 129 | 656 | 31.1 | 1 | 1 | 992370 | **0.051087291** | 2 | 2 | 2 | | 67300 | ATP-binding cassette subfamily E member 1 OS=Rattus norvegicus OX=10116 GN=Abce1 PE=1 SV=1 |
| **158** | 397 | 47.5 | 11 | 11 | 15168 | **0.000643342** | 2 | 2 | 2 | Oxidatic | 15193 | Q5BJX1\|RM41_RAT | - | | | | | | | **0** | | | | | | 39S ribosomal protein L41 mitochondrial OS=Rattus norvegicus OX=10116 GN=Mrpl41 PE=1 SV=1 |
| **228** | 162 | 34.7 | 4 | 4 | 119440 | **0.005065981** | 1 | 1 | 1 | | 33215 | tr\|Q5EBA4\|Q5EBA4_RAT | **5.073236236** | 87 | 143 | 66.89 | 11 | 11 | 499240 | **0.025700917** | 3 | 3 | 3 | Oxidati | 33215 | Nipsnap1 protein (Fragment) OS=Rattus norvegicus OX=10116 GN=Nipsnap1 PE=2 SV=1 |
| **229** | 309 | 40.9 | 2 | 2 | 449970 | **0.019085226** | 1 | 1 | 1 | | 60956 | P63039\|CH60_RAT | **0.534485617** | 112 | 218 | 49.63 | 4 | 4 | 198150 | **0.010200779** | 2 | 2 | 2 | Oxidati | 60956 | 60 kDa heat shock protein mitochondrial OS=Rattus norvegicus OX=10116 GN=Hspd1 PE=1 SV=1 |
| **233** | 5951 | 31.7 | 7 | 7 | 180330 | **0.007648596** | 1 | 1 | 1 | | 20651 | tr\|B2RZ57\|B2RZ57_RAT | - | | | | | | | **0** | | | | | | Mitochondrial ribosomal protein L18 OS=Rattus norvegicus OX=10116 GN=Mrpl18 PE=1 SV=1 |
| **259** | #### | 50.9 | 9 | 9 | 259430 | **0.011003578** | 1 | 1 | 1 | | 18377 | tr\|D4ACA5\|D4ACA5_RAT | - | | | | | | | **0** | | | | | | Mitochondrial import inner membrane translocase subunit TIM17 OS=Rattus norvegicus OX=10116 GN=LOC100911130 PE=1 SV=1 |
| **270** | #### | 26.5 | 7 | 7 | 237880 | **0.010089547** | 1 | 1 | 1 | | 17526 | tr\|B2GV62\|B2GV62_RAT | - | | | | | | | **0** | | | | | | Mitochondrial ribosomal protein L20 OS=Rattus norvegicus OX=10116 GN=Mrpl20 PE=1 SV=1 |
| **272** | #### | 23.6 | 9 | 9 | 63576 | **0.00269654** | 1 | 1 | 1 | | 10726 | tr\|A0A0U1RRX6\|A0A0U1RRX( | **2.187465055** | 281 | 11749 | 23.67 | 9 | 9 | 114580 | **0.005898588** | 1 | 1 | 1 | | 10726 | 5-demethoxyubiquinone hydroxylase mitochondrial (Fragment) OS=Rattus norvegicus OX=10116 GN=Coq7 PE=1 SV=1 |
| | | | | | | **0** | | | | | | tr\|A0A0G2JSH5\|A0A0G2JSH5 | **+** | 61 | 192 | 71.21 | 5 | 5 | 2562900 | **0.131938307** | 4 | 4 | 5 | Formyl | 68759 | Serum albumin OS=Rattus norvegicus OX=10116 GN=Alb PE=1 SV=1 |
| | | | | | | **0** | | | | | | Q6AXV4\|SAM50_RAT | **+** | 79 | 139 | 72.26 | 6 | 6 | 129970 | **0.006690867** | 3 | 3 | 3 | | 51960 | Sorting and assembly machinery component 50 homolog OS=Rattus norvegicus OX=10116 GN=Samm50 PE=1 SV=1 |
| | | | | | | **0** | | | | | | P11884\|ALDH2_RAT | **+** | 89 | 141 | 71.9 | 6 | 6 | 254000 | **0.013075941** | 3 | 3 | 3 | | 56488 | Aldehyde dehydrogenase mitochondrial OS=Rattus norvegicus OX=10116 GN=Aldh2 PE=1 SV=1 |
| | | | | | | **0** | | | | | | Q62925\|M3K1_RAT | **+** | 94 | 1525 | 32.14 | 2 | 2 | 244650 | **0.012594603** | 3 | 3 | 3 | Oxidation (M) | | Mitogen-activated protein kinase kinase kinase 1 |
| | | | | | | **0** | | | | | | P49432\|ODPB_RAT | **+** | 133 | 195 | 50.13 | 3 | 3 | 58467 | **0.003009886** | 2 | 2 | 2 | | 38982 | Pyruvate dehydrogenase E1 component subunit beta mitochondrial OS=Rattus norvegicus OX=10116 GN=Pdhb PE=1 SV=2 |
| | | | | | 2357687596 | | | | | | | tr\|D3ZTW8\|D3ZTW8_RAT | **+** | 189 | 264 | 37.32 | 8 | 8 | 128420 | **0.006611072** | 1 | 1 | 1 | | 15823 | Mitochondrial ribosomal protein L27 OS=Rattus norvegicus OX=10116 GN=Mrpl27 PE=1 SV=2 |
| | | | | | | **100** | | | | | | | | | | | | | 1942498771 | **100** | | | | | | |
| | | | | | 23576875.96 | | | | | | | | | | | | | | | | | | | | | |
| | | | | | | | | | | | | | | | | | | | 19424987.71 | | | | | | | |
