## Supplementary material for "Permeability transition pore-related changes in the proteome and channel activity of ATP synthase dimers and monomers": ATP-synthase

| Area (IBAQ) | riBAQ | Accession | rIBAQ PTP/Control | Area (IBAQ) | riBAQ | Description |
|---|---|---|---|---|---|---|
| CONTROL | % | | | PTP | % | |
| **Complex I** | | | | | | |
| 3192400 | **0.13540386** | P05508\|NU4M_RAT | **0.259484165** | 682500 | **0.035135157** | NADH-ubiquinone oxidoreductase chain 4 |
| 6381700 | **0.270676234** | P11661\|NU5M_RAT | **0.257480078** | 1353800 | **0.069693738** | NADH-ubiquinone oxidoreductase chain 5 OS=Rattus norvegicus OX=10116 GN=Mtnd5 PE=3 SV=3 |
| 3866200 | **0.163982709** | tr\|A0A0G2JVL6\|A0A0G2JVL6_ | **0.485533495** | 1546600 | **0.079619098** | NADH dehydrogenase [ubiquinone] 1 alpha subcomplex subunit 8 OS=Rattus norvegicus OX=10116 GN=Ndufa8 PE=1 SV=1 |
| 1390100 | **0.058960314** | P11662\|NU2M_RAT | - | | **0** | NADH-ubiquinone oxidoreductase chain 2 OS=Rattus norvegicus OX=10116 GN=Mtnd2 PE=3 SV=3 |
| 1724700 | **0.073152185** | tr\|D4A565\|D4A565_RAT | **2.343522987** | 3330100 | **0.171433828** | NADH dehydrogenase (Ubiquinone) 1 beta subcomplex 5 (Predicted) isoform CRA_b OS=Rattus norvegicus OX=10116 GN=Ndufb5 PE=1 SV=1 |
| 2143500 | **0.090915353** | tr\|F1LPG5\|F1LPG5_RAT | **0.495048324** | 874270 | **0.045007493** | NADH:ubiquinone oxidoreductase subunit B4 OS=Rattus norvegicus OX=10116 GN=Ndufb4 PE=1 SV=1 |
| 702550 | **0.029798265** | tr\|D3ZZ21\|D3ZZ21_RAT | **3.003640442** | 1738600 | **0.089503274** | NADH dehydrogenase (Ubiquinone) 1 beta subcomplex 6 (Predicted) OS=Rattus norvegicus OX=10116 GN=Ndufb6 PE=1 SV=1 |
| 789400 | **0.033481959** | tr\|D4A3V2\|D4A3V2_RAT | **2.329998636** | 1515400 | **0.078012919** | NADH dehydrogenase [ubiquinone] 1 alpha subcomplex subunit 6 OS=Rattus norvegicus OX=10116 GN=Ndufa6 PE=1 SV=1 |
| 2143700 | **0.090923836** | Q63362\|NDUA5_RAT | **0.288026053** | 508710 | **0.026188434** | NADH dehydrogenase [ubiquinone] 1 alpha subcomplex subunit 5 OS=Rattus norvegicus OX=10116 GN=Ndufa5 PE=1 SV=3 |
| 1471600 | **0.06241709** | tr\|A0A0G2KAA3\|A0A0G2KAA3 | - | 0 | **0** | NADH:ubiquinone oxidoreductase subunit A3 OS=Rattus norvegicus OX=10116 GN=Ndufa3 PE=1 SV=1 |
| 173420 | **0.007355512** | tr\|D3ZE15\|D3ZE15_RAT | **4.511664918** | 644630 | **0.033185607** | NADH:ubiquinone oxidoreductase subunit A13 OS=Rattus norvegicus OX=10116 GN=Ndufa13 PE=1 SV=1 |
| 274870 | **0.011658457** | tr\|D3ZS58\|D3ZS58_RAT | **2.837210598** | 642530 | **0.033077498** | NADH dehydrogenase [ubiquinone] 1 alpha subcomplex subunit 2 OS=Rattus norvegicus OX=10116 GN=Ndufa2 PE=1 SV=1 |
| 289150 | **0.012264135** | tr\|B5DEL8\|B5DEL8_RAT | - | | **0** | NADH dehydrogenase (Ubiquinone) Fe-S protein 5 OS=Rattus norvegicus OX=10116 GN=Ndufs5 PE=1 SV=1 |
| | **0** | tr\|B2RYW3\|B2RYW3_RAT | **+** | 175610 | **0.009040418** | NADH dehydrogenase (Ubiquinone) 1 beta subcomplex 9 OS=Rattus norvegicus OX=10116 GN=Ndufb9 PE=1 SV=1 |
| | **0** | tr\|D4A4P3\|D4A4P3_RAT | **+** | 586050 | **0.030169903** | NADH:ubiquinone oxidoreductase subunit B3 OS=Rattus norvegicus OX=10116 GN=Ndufb3 PE=1 SV=1 |
| **Complex III** | | | | | | |
| | **0** | Q5M9I5\|QCR6_RAT | **+** | 126470 | **0.006510686** | Cytochrome b-c1 complex subunit 6 mitochondrial OS=Rattus norvegicus OX=10116 GN=Uqcrh PE=3 SV=1 |
| **Complex IV** | | | | | | |
| 1725900 | **0.073203083** | P05503\|COX1_RAT | **0.245040507** | 348440 | **0.01793772** | Cytochrome c oxidase subunit 1 |
| 3188400 | **0.135234202** | P12075\|COX5B_RAT | **0.424374877** | 1114800 | **0.057389998** | Cytochrome c oxidase subunit 5B mitochondrial OS=Rattus norvegicus OX=10116 GN=Cox5b PE=1 SV=2 |
| 1155700 | **0.049018369** | P10888\|COX41_RAT | **0.408094589** | 388580 | **0.020004131** | Cytochrome c oxidase subunit 4 isoform 1 mitochondrial OS=Rattus norvegicus OX=10116 GN=Cox4i1 PE=1 SV=1 |
| 689260 | **0.029234577** | P80431\|COX7B_RAT | - | | **0** | Cytochrome c oxidase subunit 7B mitochondrial OS=Rattus norvegicus OX=10116 GN=Cox7b PE=1 SV=3 |
| **Complex V** | | | | | | |
| 9381500 | **0.397911073** | P05504\|ATP6_RAT | **0.092111725** | 711970 | **0.036652275** | ATP synthase subunit a |
| 1286700 | **0.05457466** | D3ZAF6\|ATPK_RAT | **0.338973583** | 359350 | **0.018499368** | ATP synthase subunit f mitochondrial OS=Rattus norvegicus OX=10116 GN=Atp5mf PE=1 SV=1 |
| **Other** | | | | | | |
| 81132000 | **3.441168378** | Q02253\|MMSA_RAT | **0.027939407** | 1867600 | **0.096144205** | Methylmalonate-semialdehyde dehydrogenase [acylating], mitochondrial |
| 21074000 | **0.893841917** | P10860\|DHE3_RAT | **28.32769292** | 491850000 | **25.32047934** | Glutamate dehydrogenase 1 mitochondrial OS=Rattus norvegicus OX=10116 GN=Glud1 PE=1 SV=2 |
| 12428000 | **0.527126665** | P16970\|ABCD3_RAT | **0.082971424** | 849580 | **0.04373645** | ATP-binding cassette sub-family D member 3 OS=Rattus norvegicus OX=10116 GN=Abcd3 PE=1 SV=3 |
| 7993800 | **0.339052554** | P07895\|SODM_RAT | **0.230015018** | 1514900 | **0.077987179** | Superoxide dismutase [Mn] mitochondrial OS=Rattus norvegicus OX=10116 GN=Sod2 PE=1 SV=2 |
| 2321300 | **0.09845664** | P07824\|ARGI1_RAT | **0.385460217** | 737200 | **0.037951118** | Arginase-1 OS=Rattus norvegicus OX=10116 GN=Arg1 PE=1 SV=2 |
| 1187700 | **0.050375631** | Q924S5\|LONM_RAT | **0.350009091** | 342500 | **0.017631929** | Lon protease homolog mitochondrial OS=Rattus norvegicus OX=10116 GN=Lonp1 PE=2 SV=1 |
| 911980 | **0.038681121** | P85834\|EFTU_RAT | **15.91737219** | 11960000 | **0.615701805** | Elongation factor Tu mitochondrial OS=Rattus norvegicus OX=10116 GN=Tufm PE=1 SV=1 |
| 397670 | **0.016866951** | tr\|Q5XFW4\|Q5XFW4_RAT | **0.434439833** | 142340 | **0.007327675** | Mitochondrial ribosomal protein L13 OS=Rattus norvegicus OX=10116 GN=Mrpl13 PE=1 SV=1 |
| 1206500 | **0.051173022** | P04041\|GPX1_RAT | **0.179530842** | 178460 | **0.009187136** | Glutathione peroxidase 1 OS=Rattus norvegicus OX=10116 GN=Gpx1 PE=1 SV=4 |
| 736680 | **0.03124587** | tr\|D3ZGM1\|D3ZGM1_RAT | **0.276908844** | 168070 | **0.008652258** | Pentatricopeptide repeat domain 3 OS=Rattus norvegicus OX=10116 GN=Ptcd3 PE=1 SV=1 |
| 546530 | **0.023180764** | Q63750\|RM23_RAT | - | | **0** | 39S ribosomal protein L23 mitochondrial OS=Rattus norvegicus OX=10116 GN=Mrpl23 PE=2 SV=1 |
| 555890 | **0.023577763** | tr\|B2GV15\|B2GV15_RAT | - | | **0** | Dihydrolipoamide acetyltransferase component of pyruvate dehydrogenase complex OS=Rattus norvegicus OX=10116 GN=Dbt PE=1 SV=1 |
| 452810 | **0.019205683** | tr\|M0R6J0\|M0R6J0_RAT | **0.388506021** | 144940 | **0.007461523** | Mitochondrial ribosomal protein L39 OS=Rattus norvegicus OX=10116 GN=Mrpl39 PE=1 SV=2 |
| 149150 | **0.006326114** | tr\|Q5U2T0\|Q5U2T0_RAT | - | 0 | **0** | Death associated protein 3 OS=Rattus norvegicus OX=10116 GN=Dap3 PE=2 SV=1 |
| 1002300 | **0.042511994** | tr\|Q4G067\|Q4G067_RAT | **0.425178186** | 351110 | **0.018075172** | Mitochondrial ribosomal protein L44 OS=Rattus norvegicus OX=10116 GN=Mrpl44 PE=1 SV=1 |
| 307470 | **0.013041168** | P09139\|SPYA_RAT | **0** | 0 | **0** | Serine--pyruvate aminotransferase mitochondrial OS=Rattus norvegicus OX=10116 GN=Agxt PE=1 SV=1 |
| 383140 | **0.016250669** | Q68FU7\|COQ6_RAT | **0** | | **0** | Ubiquinone biosynthesis monooxygenase COQ6 mitochondrial OS=Rattus norvegicus OX=10116 GN=Coq6 PE=2 SV=1 |
| 534430 | **0.022667549** | Q9ER34\|ACON_RAT | **0** | | **0** | Aconitate hydratase mitochondrial OS=Rattus norvegicus OX=10116 GN=Aco2 PE=1 SV=2 |

<table>
<tr><td>1652300</td><td>0.07008138</td><td>Q09073|ADT2_RAT</td><td>0</td><td></td><td>0</td><td>ADP/ATP translocase 2 OS=Rattus norvegicus OX=10116 GN=Slc25a5 PE=1 SV=3</td></tr>
<tr><td>388650</td><td>0.016484372</td><td>Q8VID1|DHRS4_RAT</td><td>0</td><td></td><td>0</td><td>Dehydrogenase/reductase SDR family member 4 OS=Rattus norvegicus OX=10116 GN=Dhrs4 PE=2 SV=2</td></tr>
<tr><td>600160</td><td>0.025455451</td><td>P02770|ALBU_RAT</td><td>5.183106352</td><td>2562900</td><td>0.131938307</td><td>Serum albumin OS=Rattus norvegicus OX=10116 GN=Alb PE=1 SV=2</td></tr>
<tr><td>7010000</td><td>0.297325227</td><td>Q80ZG6|BBC3_RAT</td><td>0.087455043</td><td>505100</td><td>0.02600259</td><td>Bcl-2-binding component 3 OS=Rattus norvegicus OX=10116 GN=Bbc3 PE=1 SV=1</td></tr>
<tr><td>338140</td><td>0.014342019</td><td>tr|B2RZ24|B2RZ24_RAT</td><td>-</td><td>0</td><td>0</td><td>Succinate-CoA ligase subunit beta (Fragment) OS=Rattus norvegicus OX=10116 GN=Sucla2 PE=2 SV=1</td></tr>
<tr><td>641240</td><td>0.027197836</td><td>Q9WVK3|PECR_RAT</td><td>0.337581015</td><td>178350</td><td>0.009181473</td><td>Peroxisomal trans-2-enoyl-CoA reductase OS=Rattus norvegicus OX=10116 GN=Pecr PE=2 SV=1</td></tr>
<tr><td>5658000</td><td>0.239980904</td><td>Q2V057|HYPDH_RAT</td><td>0.28005249</td><td>1305500</td><td>0.06720725</td><td>Hydroxyproline dehydrogenase OS=Rattus norvegicus OX=10116 GN=Prodh2 PE=2 SV=1</td></tr>
<tr><td>61742</td><td>0.002618752</td><td>Q9EPJ3|RT26_RAT</td><td>-</td><td></td><td>0</td><td>28S ribosomal protein S26  mitochondrial OS=Rattus norvegicus OX=10116 GN=Mrps26 PE=1 SV=1</td></tr>
<tr><td>371920</td><td>0.015774779</td><td>tr|D3ZD23|D3ZD23_RAT</td><td>3.238542491</td><td>992370</td><td>0.051087291</td><td>ATP-binding cassette subfamily E member 1 OS=Rattus norvegicus OX=10116 GN=Abce1 PE=1 SV=1</td></tr>
<tr><td>15168</td><td>0.000643342</td><td>Q5BJX1|RM41_RAT</td><td>-</td><td></td><td>0</td><td>39S ribosomal protein L41  mitochondrial OS=Rattus norvegicus OX=10116 GN=Mrpl41 PE=1 SV=1</td></tr>
<tr><td>119440</td><td>0.005065981</td><td>tr|Q5EBA4|Q5EBA4_RAT</td><td>5.073236236</td><td>499240</td><td>0.025700917</td><td>Nipsnap1 protein (Fragment) OS=Rattus norvegicus OX=10116 GN=Nipsnap1 PE=2 SV=1</td></tr>
<tr><td>180330</td><td>0.007648596</td><td>tr|B2RZ57|B2RZ57_RAT</td><td>-</td><td></td><td>0</td><td>Mitochondrial ribosomal protein L18 OS=Rattus norvegicus OX=10116 GN=Mrpl18 PE=1 SV=1</td></tr>
<tr><td>259430</td><td>0.011003578</td><td>tr|D4ACA5|D4ACA5_RAT</td><td>-</td><td></td><td>0</td><td>Mitochondrial import inner membrane translocase subunit TIM17 OS=Rattus norvegicus OX=10116 GN=LOC100911130 PE=1 SV=1</td></tr>
<tr><td>237880</td><td>0.010089547</td><td>tr|B2GV62|B2GV62_RAT</td><td>-</td><td></td><td>0</td><td>Mitochondrial ribosomal protein L20 OS=Rattus norvegicus OX=10116 GN=Mrpl20 PE=1 SV=1</td></tr>
<tr><td>63576</td><td>0.00269654</td><td>tr|A0A0U1RRX6|A0A0U1RRX6</td><td>2.187465055</td><td>114580</td><td>0.005898588</td><td>5-demethoxyubiquinone hydroxylase  mitochondrial (Fragment) OS=Rattus norvegicus OX=10116 GN=Coq7 PE=1 SV=1</td></tr>
<tr><td></td><td>0</td><td>Q6AXV4|SAM50_RAT</td><td>+</td><td>129970</td><td>0.006690867</td><td>Sorting and assembly machinery component 50 homolog OS=Rattus norvegicus OX=10116 GN=Samm50 PE=1 SV=1</td></tr>
<tr><td></td><td>0</td><td>P11884|ALDH2_RAT</td><td>+</td><td>254000</td><td>0.013075941</td><td>Aldehyde dehydrogenase  mitochondrial OS=Rattus norvegicus OX=10116 GN=Aldh2 PE=1 SV=1</td></tr>
<tr><td></td><td>0</td><td>Q62925|M3K1_RAT</td><td>+</td><td>244650</td><td>0.012594603</td><td>Mitogen-activated protein kinase kinase kinase 1</td></tr>
<tr><td></td><td>0</td><td>P49432|ODPB_RAT</td><td>+</td><td>58467</td><td>0.003009886</td><td>Pyruvate dehydrogenase E1 component subunit beta  mitochondrial OS=Rattus norvegicus OX=10116 GN=Pdhb PE=1 SV=2</td></tr>
<tr><td></td><td></td><td>tr|D3ZTW8|D3ZTW8_RAT</td><td>+</td><td>128420</td><td>0.006611072</td><td>Mitochondrial ribosomal protein L27 OS=Rattus norvegicus OX=10116 GN=Mrpl27 PE=1 SV=2</td></tr>
</table>
