## Supplementary material for "Permeability transition pore-related changes in the proteome and channel activity of ATP synthase dimers and monomers": ATP-synthase

| Protein Group | Protein ID | -10lgP | Coverage (%) | Coverage (%) Sample 1 | Area (IBAQ) | riBAQ | #Peptides | #Unique | #Spec Sample 1 | PTM | Avg. Mass | Accession | riBAQ PTP/Control | Protein Group | Protein ID | -10lgP | Coverage (%) | Coverage (%) Sample 1 | Area (IBAQ) | riBAQ | #Peptides | #Unique | #Spec Sample 1 | PTM | Avg. Mass | Description |
|---|---|---|---|---|---|---|---|---|---|---|---|---|---|---|---|---|---|---|---|---|---|---|---|---|---|---|
| | | | | | CONTROL | % | | | | | | | | | | | | | PTP | % | | | | | | |
| **Complex I** | | | | | | | | | | | | | | | | | | | | | | | | | | |
| | | | | | | | | | | | | tr\|A0A0G2JVL6\|A0 | + | 199 | 34 | 37.02 | 7 | 7 | 139230 | 0.001106384 | 1 | 1 | 1 | | 17634 | NADH dehydrogenase (Ubiquinone) 1 beta subcomplex 11 (Predicted) OS=Rattus norvegicus OX=10116 GN=Ndufb11 PE=1 SV=1 |
| | | | | | | | | | | | | tr\|D4A7L4\|D4A7L4 | + | 238 | 137 | 44.08 | 9 | 9 | 135780 | 0.001078969 | 1 | 1 | 1 | | 19965 | NADH dehydrogenase [ubiquinone] 1 alpha subcomplex subunit 8 OS=Rattus norvegicus OX=10116 GN=Ndufa8 PE=1 SV=1 |
| | | | | | | | | | | | | tr\|D4A0T0\|D4A0T0 | + | 256 | 66 | 20.77 | 10 | 10 | 434220 | 0.003450507 | 1 | 1 | 1 | | 20859 | NADH:ubiquinone oxidoreductase subunit B10 OS=Rattus norvegicus OX=10116 GN=Ndufb10 PE=1 SV=1 |
| | | | | | | | | | | | | | | | | | | | | 0 | | | | | | |
| **Complex III** | | | | | | | | | | | | | | | | | | | | 0 | | | | | | |
| 28 | 13 | 88.7 | 5 | 5 | 2492000 | 0.027213397 | 6 | 6 | 9 | | 48396 | P32551\|QCR2_RAT | 1.363634483 | 29 | 21 | 104.27 | 5 | 5 | 4669900 | 0.037109126 | 6 | 6 | 11 | | 48396 | Cytochrome b-c1 complex subunit 2 mitochondrial OS=Rattus norvegicus OX=10116 GN=Uqcrc2 PE=1 SV=2 |
| 40 | 12 | 78.4 | 6 | 6 | 2660100 | 0.0290491 | 3 | 3 | 5 | | 52849 | Q68FY0\|QCR1_RAT | 0.410219122 | 57 | 16 | 82.22 | 7 | 7 | 1499600 | 0.011916496 | 4 | 4 | 4 | | 52849 | Cytochrome b-c1 complex subunit 1 mitochondrial OS=Rattus norvegicus OX=10116 GN=Uqcrc1 PE=1 SV=1 |
| 111 | 65 | 40.9 | 3 | 3 | 323100 | 0.00352835 | 1 | 1 | 2 | | 43012 | P00159\|CYB_RAT | 2.152085076 | 49 | 97 | 102.62 | 9 | 9 | 955560 | 0.00759331 | 4 | 4 | 5 | | 43012 | Cytochrome b OS=Rattus norvegicus OX=10116 GN=CYTB PE=3 SV=1 |
| 151 | 80 | 77 | 5 | 5 | 369530 | 0.00403538 | 1 | 1 | 1 | | 43050 | P20788\|UCRI_RAT | 4.210138965 | 48 | 42 | 125.14 | 10 | 10 | 2138000 | 0.01698951 | 4 | 4 | 5 | Formyla | 29446 | Cytochrome b (Fragment) OS=Rattus norvegicus OX=10116 GN=cytb PE=3 SV=1 |
| 190 | 279 | 35.3 | 17 | 17 | 396150 | 0.004326078 | 1 | 1 | 1 | | 10424 | Q5M9I5\|QCR6_RAT | - | 241 | 338 | 34.58 | 17 | 17 | 0 | 0 | 1 | 1 | 1 | | 10424 | Cytochrome b-c1 complex subunit 6 mitochondrial OS=Rattus norvegicus OX=10116 GN=Uqcrh PE=3 SV=1 |
| | | | | | | 0 | | | | | | | | | | | | | | 0 | | | | | | |
| **Complex IV** | | | | | | 0 | | | | | | | | | | | | | | 0 | | | | | | |
| 44 | 22 | 102 | 6 | 6 | 1848400 | 0.020185089 | 3 | 2 | 5 | | 35435 | tr\|D3ZFQ8\|D3ZFQ8 | 1.223633697 | 45 | 38 | 105.46 | 6 | 6 | 3108200 | 0.024699155 | 3 | 3 | 5 | | 35435 | Cytochrome c-1 OS=Rattus norvegicus OX=10116 GN=Cyc1 PE=1 SV=3 |
| 71 | 82 | 43.5 | 7 | 7 | 414580 | 0.004527339 | 2 | 2 | 3 | Oxi | 25928 | P00406\|COX2_RAT | 0.871218042 | 109 | 112 | 40.45 | 7 | 7 | 496360 | 0.0039443 | 2 | 2 | 2 | | 25958 | Cytochrome c oxidase subunit 2 OS=Rattus norvegicus OX=10116 GN=Mtco2 PE=1 SV=3 |
| | | | | | | 0 | | | | | | P11240\|COX5A_RA | + | 107 | 44 | 90.83 | 17 | 17 | 529460 | 0.004207327 | 2 | 2 | 2 | | 16130 | Cytochrome c oxidase subunit 5A mitochondrial OS=Rattus norvegicus OX=10116 GN=Cox5a PE=1 SV=1 |
| | | | | | | 0 | | | | | | P35171\|CX7A2_RA | + | 236 | 11743 | 65.29 | 14 | 14 | 311870 | 0.002478259 | 1 | 1 | 1 | | 25928 | Cytochrome c oxidase subunit 2 OS=Rattus norvegicus OX=10116 GN=Mt-co2 PE=1 SV=1 |
| | | | | | | 0 | | | | | | P10888\|COX41_RA | + | 240 | 247 | 37.76 | 8 | 8 | 237750 | 0.001889268 | 1 | 1 | 1 | | 19515 | Cytochrome c oxidase subunit 4 isoform 1 mitochondrial OS=Rattus norvegicus OX=10116 GN=Cox4i1 PE=1 SV=1 |
| | | | | | | 0 | | | | | | | | | | | | | | 0 | | | | | | |
| **Complex V** | | | | | | 0 | | | | | | | | | | | | | | 0 | | | | | | |
| 1 | 20 | 364 | 59 | 59 | 3050800000 | 33.31566247 | 199 | 194 | 1112 | Carbamido | | P15999\|ATPA_RAT | 0.925862857 | 1 | 10 | 367.07 | 63 | 63 | 3881700000 | 30.84573445 | 248 | 237 | 1140 | Carbamidomethy | | ATP synthase subunit alpha, mitochondrial |
| 2 | 17 | 404 | 61 | 61 | 3757700000 | 41.03522514 | 202 | 197 | 966 | Oxidation ( | | P10719\|ATPB_RAT | 0.965187721 | 2 | 8 | 405.85 | 71 | 71 | 4984200000 | 39.60669542 | 241 | 233 | 875 | Oxidation (M) | | ATP synthase subunit beta, mitochondrial |
| 4 | 301 | 297 | 56 | 56 | 433430000 | 4.733187224 | 67 | 66 | 204 | Oxi | 30191 | P35435\|ATPG_RAT | 0.889335922 | 6 | 64 | 305.24 | 53 | 53 | 529720000 | 4.209393423 | 80 | 80 | 182 | Carbamidomethy | | ATP synthase subunit gamma mitochondrial OS=Rattus norvegicus OX=10116 GN=Atp5f1c PE=1 SV=2 |
| 5 | 280 | 327 | 73 | 73 | 306560000 | 3.347728296 | 58 | 56 | 168 | Carbamido | | P31399\|ATP5H_RA | 1.542111434 | 4 | 119 | 349.97 | 82 | 82 | 649670000 | 5.162570084 | 87 | 87 | 194 | Carbamidomethy | | ATP synthase subunit d, mitochondrial |
| 6 | 268 | 226 | 29 | 29 | 376000000 | 4.106034184 | 36 | 36 | 134 | For | 28869 | P19511\|AT5F1_RAT | 1.180384401 | 5 | 246 | 238.91 | 48 | 48 | 609920000 | 4.846698702 | 59 | 58 | 189 | Formyla | 28869 | ATP synthase F(0) complex subunit B1 mitochondrial OS=Rattus norvegicus OX=10116 GN=Atp5pb PE=1 SV=1 |
| 8 | 5938 | 219 | 48 | 48 | 123830000 | 1.352261205 | 45 | 43 | 115 | Carbamido | | Q06647\|ATPO_RAT | 1.553019315 | 7 | 5802 | 236.85 | 54 | 54 | 264280000 | 2.100087771 | 63 | 61 | 124 | Carbamidomethy | | ATP synthase subunit O, mitochondrial |
| 9 | 5937 | 215 | 39 | 39 | 109760000 | 1.198612532 | 25 | 25 | 66 | Oxi | 11433 | Q6PDU7\|ATP5L_RA | 1.715304086 | 10 | 1646 | 228.57 | 50 | 50 | 258730000 | 2.055984974 | 38 | 38 | 94 | Oxidati | 11433 | ATP synthase subunit g mitochondrial OS=Rattus norvegicus OX=10116 GN=Atp5mg PE=1 SV=2 |
| 10 | 387 | 203 | 36 | 36 | 122500000 | 1.337737201 | 19 | 19 | 55 | Oxi | 17563 | tr\|G3V7Y3\|G3V7Y3 | 1.13749224 | 11 | 350 | 213.55 | 48 | 48 | 191490000 | 1.521665685 | 26 | 26 | 63 | Oxidati | 17563 | ATP synthase subunit delta mitochondrial OS=Rattus norvegicus OX=10116 GN=Atp5f1d PE=1 SV=1 |
| 13 | 5942 | 122 | 35 | 35 | 32507000 | 0.354986312 | 18 | 17 | 45 | Oxidation ( | | P05504\|ATP6_RAT | 0.51967309 | 21 | 690 | 108.74 | 38 | 38 | 23215000 | 0.184476834 | 18 | 16 | 27 | Oxidation (M) | | ATP synthase subunit a |
| 20 | 416 | 125 | 42 | 42 | 50060000 | 0.546670402 | 7 | 6 | 23 | Oxi | 7642 | tr\|Q5UAJ5\|Q5UAJ5 | 1.353019206 | 23 | 1095 | 135.59 | 69 | 69 | 93080000 | 0.739655554 | 10 | 10 | 22 | Oxidati | 7642 | ATP synthase protein 8 OS=Rattus norvegicus OX=10116 GN=ATP8 PE=3 SV=1 |
| 29 | #### | 62.9 | 37 | 37 | 5323500 | 0.058134237 | 4 | 4 | 9 | | 8255 | P29419\|ATP5I_RAT | 9.036804771 | 13 | 11740 | 155.85 | 73 | 73 | 66111000 | 0.525347747 | 28 | 26 | 56 | | 8255 | ATP synthase subunit e mitochondrial OS=Rattus norvegicus OX=10116 GN=Atp5me PE=1 SV=3 |
| 30 | #### | 48.4 | 17 | 17 | 10349000 | 0.113014223 | 4 | 4 | 8 | | 10452 | D3ZAF6\|ATPK_RAT | 1.409719674 | 28 | 11756 | 77.21 | 30 | 30 | 20049000 | 0.159318373 | 8 | 7 | 15 | Oxidati | 10452 | ATP synthase subunit f mitochondrial OS=Rattus norvegicus OX=10116 GN=Atp5mf PE=1 SV=1 |
| 33 | #### | 137 | 31 | 31 | 4150100 | 0.045320352 | 6 | 6 | 8 | | 12494 | P21571\|ATP5J_RAT | 7.97829989 | 15 | 5817 | 234.2 | 58 | 58 | 45502000 | 0.361579362 | 29 | 28 | 45 | Oxidation (M) | | ATP synthase-coupling factor 6 mitochondrial OS=Rattus norvegicus OX=10116 GN=Atp5pf PE=1 SV=1 |
| 43 | #### | 55.8 | 29 | 29 | 4965000 | 0.054219308 | 3 | 3 | 5 | | 6408 | Q9JJW3\|ATPMD_RA | 1.783064245 | 31 | 11746 | 103.21 | 43 | 43 | 12166000 | 0.096676509 | 6 | 6 | 9 | | 6408 | ATP synthase membrane subunit DAPIT mitochondrial OS=Rattus norvegicus OX=10116 GN=Atp5md PE=1 SV=1 |
| | | | | | | 0 | | | | | | D3Z9R8\|ATP68_RA | + | 60 | 34376 | 51.97 | 32 | 32 | 306390 | 0.002434713 | 3 | 2 | 4 | Oxidati | 6914 | ATP synthase subunit ATP5MPL mitochondrial OS=Rattus norvegicus OX=10116 GN=Atp5mpl PE=1 SV=1 |
| | | | | | | 0 | | | | | | P29418\|ATP5E_RA | + | 119 | 34379 | 31.66 | 25 | 25 | 727560 | 0.005781519 | 2 | 2 | 2 | | 5767 | ATP synthase subunit epsilon mitochondrial OS=Rattus norvegicus OX=10116 GN=Atp5f1e PE=1 SV=2 |
| | | | | | | 0 | | | | | | | | | | | | | | 0 | | | | | | |
| **Other** | | | | | | 0 | | | | | | | | | | | | | | 0 | | | | | | |
| 3 | 35 | 360 | 39 | 39 | 425370000 | 4.645169577 | 153 | 150 | 375 | Oxi | 1E+05 | P52873\|PYC_RAT | 0.705198158 | 3 | 30 | 337.26 | 43 | 43 | 412230000 | 3.275765028 | 164 | 159 | 292 | Oxidation (M) | | Pyruvate carboxylase mitochondrial OS=Rattus norvegicus OX=10116 GN=Pc PE=1 SV=2 |
| 7 | 8 | 252 | 25 | 25 | 93640000 | 1.022577237 | 70 | 68 | 131 | Carbamido | | P07756\|CPSM_RAT | 0.583058288 | 9 | 4 | 256.7 | 26 | 26 | 75030000 | 0.596222133 | 74 | 71 | 98 | Carbamidomethy | | Carbamoyl-phosphate synthase [ammonia], mitochondrial |
| 11 | 1 | 196 | 19 | 19 | 15340000 | 0.167517458 | 26 | 25 | 51 | Oxi | 57808 | Q02253\|MMSA_RA | 0.40094793 | 25 | 50 | 156.3 | 12 | 12 | 8452300 | 0.067165778 | 13 | 13 | 16 | Oxidati | 57808 | Methylmalonate-semialdehyde dehydrogenase [acylating] mitochondrial OS=Rattus norvegicus OX=10116 GN=Aldh6a1 PE=1 SV=1 |
| 12 | 367 | 169 | 28 | 28 | 23897000 | 0.260962497 | 25 | 24 | 48 | Oxi | 61869 | P0C2X9\|AL4A1_RA | 0.483036971 | 17 | 352 | 165.56 | 23 | 23 | 15863000 | 0.126054534 | 26 | 26 | 34 | Oxidati | 61869 | Delta-1-pyrroline-5-carboxylate dehydrogenase mitochondrial OS=Rattus norvegicus OX=10116 GN=Aldh4a1 PE=1 SV=1 |
| 14 | 266 | 195 | 27 | 27 | 19116000 | 0.208752525 | 21 | 21 | 36 | Oxi | 59757 | P04762\|CATA_RAT | 1.867117148 | 14 | 45 | 211.13 | 27 | 27 | 49049000 | 0.389765419 | 30 | 30 | 46 | | 59757 | Catalase OS=Rattus norvegicus OX=10116 GN=Cat PE=1 SV=3 |
| 15 | 15 | 208 | 24 | 24 | 35689000 | 0.389734718 | 22 | 21 | 36 | Oxi | 47314 | P00507\|AATM_RAT | 1.558258484 | 12 | 26 | 237.65 | 36 | 36 | 76425000 | 0.607307431 | 40 | 39 | 58 | Oxidati | 47314 | Aspartate aminotransferase mitochondrial OS=Rattus norvegicus OX=10116 GN=Got2 PE=1 SV=2 |
| 16 | 169 | 156 | 16 | 16 | 12681000 | 0.138480371 | 17 | 17 | 32 | Oxi | 51414 | Q60587\|ECHB_RAT | 1.320043638 | 18 | 27 | 190.48 | 27 | 27 | 23004000 | 0.182800133 | 25 | 25 | 33 | | 51414 | Trifunctional enzyme subunit beta mitochondrial OS=Rattus norvegicus OX=10116 GN=Hadhb PE=1 SV=1 |
| 17 | 33 | 163 | 12 | 12 | 16344000 | 0.178481443 | 16 | 15 | 31 | Carbamido | | Q64428\|ECHA_RAT | 0.98457178 | 16 | 24 | 185.48 | 16 | 16 | 22114000 | 0.175727792 | 20 | 20 | 35 | Carbam | 82665 | Trifunctional enzyme subunit alpha, mitochondrial |
| 18 | 289 | 223 | 17 | 17 | 15061000 | 0.164470694 | 17 | 17 | 27 | | 60420 | tr\|D3ZFJ6\|D3ZFJ6_ | 0.948767422 | 19 | 171 | 266.34 | 25 | 25 | 19637000 | 0.156044436 | 26 | 26 | 32 | | 60420 | Lactamase beta OS=Rattus norvegicus OX=10116 GN=Lactb PE=1 SV=1 |
| 19 | 309 | 166 | 16 | 16 | 16778000 | 0.183220855 | 14 | 14 | 25 | Oxi | 60956 | P63039\|CH60_RAT | 0.814852108 | 20 | 218 | 193.69 | 24 | 24 | 18788000 | 0.1492979 | 25 | 25 | 29 | Oxidati | 60956 | 60 kDa heat shock protein mitochondrial OS=Rattus norvegicus OX=10116 GN=Hspd1 PE=1 SV=1 |
| 21 | 14 | 194 | 20 | 20 | 8447200 | 0.092245989 | 17 | 17 | 21 | Oxidation ( | | P10860\|DHE3_RAT | 10.15983789 | 8 | 1 | 293.87 | 41 | 41 | 117940000 | 0.937204297 | 66 | 66 | 107 | Oxidati | 61416 | Glutamate dehydrogenase 1, mitochondrial |
| 22 | 245 | 141 | 9 | 9 | 8032700 | 0.087719523 | 10 | 10 | 20 | For | 78179 | P18163\|ACSL1_RAT | 1.38329855 | 24 | 54 | 158.35 | 15 | 15 | 15270000 | 0.121342289 | 17 | 17 | 21 | Formyla | 78179 | Long-chain-fatty-acid--CoA ligase 1 OS=Rattus norvegicus OX=10116 GN=Acsl1 PE=1 SV=1 |
| 23 | 77 | 133 | 13 | 13 | 5839200 | 0.063765837 | 12 | 12 | 17 | | 56912 | P22791\|HMCS2_RA | 1.537053646 | 22 | 23 | 176.62 | 23 | 23 | 12334000 | 0.098011513 | 21 | 21 | 27 | | 56912 | Hydroxymethylglutaryl-CoA synthase mitochondrial OS=Rattus norvegicus OX=10116 GN=Hmgcs2 PE=1 SV=1 |
| 24 | 100 | 128 | 13 | 13 | 5613000 | 0.061295665 | 8 | 8 | 13 | | 44695 | P17764\|THIL_RAT | 1.287247397 | 26 | 35 | 142.6 | 15 | 15 | 9929300 | 0.078902685 | 10 | 10 | 16 | | 44695 | Acetyl-CoA acetyltransferase mitochondrial OS=Rattus norvegicus OX=10116 GN=Acat1 PE=1 SV=1 |
| 25 | 317 | 41.3 | 1 | 1 | 6135900 | 0.067005891 | 4 | 4 | 10 | For | 5E+05 | tr\|A0A0G2JYD4\|A0 | 0.603485733 | 42 | 461 | 34.69 | 0 | 0 | 5088700 | 0.040437099 | 2 | 2 | 5 | | 487857 | Vacuolar protein sorting 13 homolog D OS=Rattus norvegicus OX=10116 GN=Vps13d PE=1 SV=1 |
| 27 | 1292 | 111 | 12 | 12 | 1379200 | 0.015061283 | 6 | 6 | 9 | For | 37791 | tr\|G3V6I4\|G3V6I4_ | 0.827552789 | 35 | 207 | 112.27 | 8 | 8 | 1568500 | 0.012464007 | 4 | 4 | 7 | | 37791 | Mitochondrial amidoxime reducing component 1 OS=Rattus norvegicus OX=10116 GN=Marc1 PE=1 SV=1 |
| 31 | 888 | 35.3 | 6 | 6 | 10104000 | 0.110338748 | 2 | 2 | 8 | For | 41255 | O70600\|RSAD2_RA | 0.52392862 | 81 | 356 | 23.54 | 5 | 5 | 7274900 | 0.057809628 | 2 | 2 | 3 | | 41255 | Radical S-adenosyl methionine domain-containing protein 2 OS=Rattus norvegicus OX=10116 GN=Rsad2 PE=1 SV=1 |
| 32 | 6503 | 96.6 | 6 | 6 | 1930500 | 0.021081646 | 6 | 4 | 8 | | 87371 | B3DMA2\|ACD11_RA | 1.338201295 | 33 | 11785 | 113.7 | 8 | 8 | 3550200 | 0.028211486 | 7 | 7 | 8 | | 87371 | Acyl-CoA dehydrogenase family member 11 OS=Rattus norvegicus OX=10116 GN=Acad11 PE=1 SV=1 |
| 35 | 176 | 94.7 | 9 | 9 | 1558900 | 0.017023661 | 5 | 5 | 6 | | 36148 | P13086\|SUCA_RAT | 1.922188483 | 27 | 46 | 141.11 | 17 | 17 | 4117900 | 0.032722686 | 12 | 12 | 16 | Formyla | 36148 | Succinate--CoA ligase [ADP/GDP-forming] subunit alpha mitochondrial OS=Rattus norvegicus OX=10116 GN=Suclg1 PE=2 SV=2 |
| 37 | #### | 39.6 | 3 | 3 | 1883600 | 0.020569484 | 3 | 2 | 6 | | 78400 | Q8CGU6\|NICA_RAT | - | | | | | | | 0 | | | | | | Nicastrin OS=Rattus norvegicus OX=10116 GN=Ncstn PE=1 SV=1 |
| 42 | 261 | 92.4 | 6 | 6 | 1237900 | 0.013518244 | 2 | 2 | 5 | | 33407 | P24329\|THTR_RAT | 2.732534818 | 30 | 136 | 171.16 | 18 | 18 | 4648500 | 0.036939072 | 7 | 7 | 9 | | 33407 | Thiosulfate sulfurtransferase OS=Rattus norvegicus OX=10116 GN=Tst PE=1 SV=3 |
| 49 | 814 | 121 | 5 | 5 | 607180 | 0.00663059 | 4 | 4 | 4 | | 70749 | P45953\|ACADV_RA | 1.427716951 | 36 | 201 | 151.85 | 9 | 9 | 1191300 | 0.009466606 | 6 | 6 | 6 | | 70749 | Very long-chain specific acyl-CoA dehydrogenase mitochondrial OS=Rattus norvegicus OX=10116 GN=Acadvl PE=1 SV=1 |
| 50 | 5947 | 63.6 | 2 | 2 | 406090 | 0.004434626 | 2 | 2 | 4 | | 1E+05 | Q924S5\|LONM_RA | 2.102089349 | 40 | 297 | 81.14 | 3 | 3 | 1173100 | 0.00932198 | 4 | 4 | 6 | | 105792 | Lon protease homolog mitochondrial OS=Rattus norvegicus OX=10116 GN=Lonp1 PE=2 SV=1 |
| 52 | 1610 | 74.2 | 3 | 3 | 509790 | 0.005567062 | 2 | 2 | 4 | | 74675 | A0A0G2K047\|ACSS | 0.217122528 | 67 | 240 | 72.24 | 3 | 3 | 152110 | 0.001208734 | 2 | 2 | 3 | | 74675 | Acyl-CoA synthetase short-chain family member 3 mitochondrial OS=Rattus norvegicus OX=10116 GN=Acss3 PE=1 SV=1 |
| 57 | 667 | 145 | 5 | 5 | 345780 | 0.003776023 | 4 | 4 | 4 | | 47388 | tr\|B2RZ24\|B2RZ24_ | 1.064199199 | 72 | 277 | 128.18 | 5 | 5 | 505690 | 0.00401844 | 3 | 3 | 3 | | 47388 | Succinate-CoA ligase subunit beta (Fragment) OS=Rattus norvegicus OX=10116 GN=Sucla2 PE=2 SV=1 |
| 65 | 414 | 46.8 | 3 | 3 | 14366000 | 0.156881083 | 2 | 2 | 3 | For | 93715 | P97564\|GPAT1_RA | 0.058878694 | 65 | 1180 | 47.36 | 3 | 3 | 1162400 | 0.009236953 | 2 | 2 | 3 | Formyla | 93715 | Glycerol-3-phosphate acyltransferase 1 mitochondrial OS=Rattus norvegicus OX=10116 GN=Gpam PE=1 SV=3 |
| 67 | 1940 | 58.4 | 2 | 2 | 418350 | 0.004568509 | 2 | 2 | 3 | | 1E+05 | A2VCW9\|AASS_RA | 3.429394667 | 34 | 742 | 139.69 | 8 | 8 | 1971600 | 0.015667221 | 7 | 7 | 7 | Carbam | 102908 | Alpha-aminoadipic semialdehyde synthase mitochondrial OS=Rattus norvegicus OX=10116 GN=Aass PE=2 SV=1 |
| 69 | 293 | 52 | 12 | 12 | 767510 | 0.008381442 | 3 | 3 | 3 | Oxi | 32433 | Q9WVK3\|PECR_RA | 0.369114498 | 52 | 283 | 55.65 | 13 | 13 | 389320 | 0.003093712 | 4 | 3 | 5 | Oxidati | 32433 | Peroxisomal trans-2-enoyl-CoA reductase OS=Rattus norvegicus OX=10116 GN=Pecr PE=2 SV=1 |
| 77 | 244 | 43.8 | 6 | 6 | 214350 | 0.002340767 | 2 | 1 | 3 | | 34973 | P07824\|ARGI1_RAT | - | 198 | 190 | 38.36 | 4 | 4 | 0 | 0 | 1 | 1 | 1 | | 34973 | Arginase-1 OS=Rattus norvegicus OX=10116 GN=Arg1 PE=1 SV=2 |
| 85 | #### | 66.6 | 2 | 2 | 2745300 | 0.02997951 | 2 | 2 | 2 | | 1E+05 | tr\|Q5BJZ3\|Q5BJZ3_ | 0.127383834 | 95 | 321 | 55.11 | 2 | 2 | 480580 | 0.003818905 | 2 | 2 | 2 | | 113869 | Nicotinamide nucleotide transhydrogenase OS=Rattus norvegicus OX=10116 GN=Nnt PE=1 SV=1 |
| 95 | 6075 | 30.8 | 1 | 1 | 5427400 | 0.059268856 | 2 | 2 | 2 | For | 1E+05 | D3ZG52\|DNA2_RAT | 1.730501228 | 55 | 760 | 35.17 | 3 | 3 | 12907000 | 0.102564828 | 4 | 3 | 4 | Formyla | 119588 | DNA replication ATP-dependent helicase/nuclease DNA2 OS=Rattus norvegicus OX=10116 GN=Dna2 PE=3 SV=1 |
| 107 | 623 | 38.1 | 5 | 5 | 851720 | 0.009301041 | 2 | 2 | 2 | | 46777 | tr\|G3V6I5\|G3V6I5_ | 0.180466745 | 160 | 341 | 28.64 | 3 | 3 | 211230 | 0.001678529 | 1 | 1 | 1 | | 46777 | DnaJ heat shock protein family (Hsp40) member A3 OS=Rattus norvegicus OX=10116 GN=Dnaja3 PE=1 SV=2 |
| 108 | 228 | 26.6 | 3 | 3 | 300990 | 0.003286902 | 2 | 2 | 2 | | 75316 | P16970\|ABCD3_RA | - | | | | | | | 0 | | | | | | ATP-binding cassette sub-family D member 3 OS=Rattus norvegicus OX=10116 GN=Abcd3 PE=1 SV=3 |

| | | | | | | | | | | | | | | | | | | | | | | | | | | |
|---|---|---|---|---|---|---|---|---|---|---|---|---|---|---|---|---|---|---|---|---|---|---|---|---|---|---|
| 110 | 2022 | 46.1 | 5 | 5 | 844660 | **0.009223944** | 1 | 1 | 2 | | 28295 | Q9Z0V6\|PRDX3_RA | **1.786325277** | 61 | 5863 | 84.56 | 9 | 9 | 2073500 | **0.016476964** | 3 | 3 | 4 | | 28295 | Thioredoxin-dependent peroxide reductase mitochondrial OS=Rattus norvegicus OX=10116 GN=Prdx3 PE=1 SV=2 |
| 112 | 219 | 51.7 | 6 | 6 | 506980 | **0.005536376** | 2 | 2 | 2 | | 22305 | P04041\|GPX1_RAT | **0.634725557** | 70 | 53 | 56.38 | 13 | 13 | 442220 | **0.003514079** | 3 | 3 | 3 | | 22305 | Glutathione peroxidase 1 OS=Rattus norvegicus OX=10116 GN=Gpx1 PE=1 SV=4 |
| 115 | 233 | 63.4 | 4 | 4 | 581710 | **0.00635245** | 2 | 2 | 2 | | 38202 | P29147\|BDH_RAT | **1.843490848** | 108 | 61 | 42.54 | 5 | 5 | 1473700 | **0.011710683** | 2 | 2 | 2 | | 38202 | D-beta-hydroxybutyrate dehydrogenase mitochondrial OS=Rattus norvegicus OX=10116 GN=Bdh1 PE=1 SV=2 |
| 116 | 380 | 42.9 | 7 | 7 | 565230 | **0.006172483** | 2 | 2 | 2 | | 27246 | O70351\|HCD2_RAT | **2.542484638** | 44 | 188 | 98.87 | 13 | 13 | 1974900 | **0.015693444** | 5 | 5 | 5 | | 27246 | 3-hydroxyacyl-CoA dehydrogenase type-2 OS=Rattus norvegicus OX=10116 GN=Hsd17b10 PE=1 SV=3 |
| 169 | 162 | 42.4 | 4 | 4 | 499600 | **0.005455784** | 1 | 1 | 1 | Oxi | 33215 | tr\|Q5EBA4\|Q5EBA4 | **0.641013788** | 83 | 143 | 70.7 | 8 | 8 | 440100 | **0.003497233** | 2 | 2 | 3 | Oxidati | 33215 | Nipsnap1 protein (Fragment) OS=Rattus norvegicus OX=10116 GN=Nipsnap1 PE=2 SV=1 |
| 195 | 6337 | 23.7 | 2 | 2 | 244940 | **0.002674819** | 1 | 1 | 1 | | 60733 | P17178\|CP27A_RAT | **0.81891102** | 110 | 59 | 32.33 | 5 | 5 | 275650 | **0.002190439** | 2 | 2 | 2 | | 60733 | Sterol 26-hydroxylase mitochondrial OS=Rattus norvegicus OX=10116 GN=Cyp27a1 PE=1 SV=1 |
| 216 | 198 | 43 | 3 | 3 | 74258 | **0.00081092** | 1 | 1 | 1 | | 49522 | P85834\|EFTU_RAT | **14.142354** | 50 | 3 | 94.86 | 7 | 7 | 1443200 | **0.011468316** | 4 | 4 | 5 | | 49522 | Elongation factor Tu mitochondrial OS=Rattus norvegicus OX=10116 GN=Tufm PE=1 SV=1 |
| | | | | | | **0** | | | | | | tr\|D3ZTP0\|D3ZTP0_ | **+** | 32 | 6375 | 76.77 | 6 | 6 | 2779100 | **0.022083979** | 6 | 6 | 8 | | 101806 | 10-formyltetrahydrofolate dehydrogenase OS=Rattus norvegicus OX=10116 GN=Aldh1l2 PE=1 SV=3 |
| | | | | | | **0** | | | | | | D3ZX08\|SYAM_RAT | **+** | 37 | 1414 | 46.08 | 2 | 2 | 305160 | **0.002424939** | 3 | 2 | 6 | | 107806 | Alanine--tRNA ligase mitochondrial OS=Rattus norvegicus OX=10116 GN=Aars2 PE=3 SV=1 |
| | | | | | | **0** | | | | | | P08011\|MGST1_RA | **+** | 38 | 250 | 71.07 | 15 | 15 | 1101000 | **0.008749041** | 5 | 5 | 6 | | 17472 | Microsomal glutathione S-transferase 1 OS=Rattus norvegicus OX=10116 GN=Mgst1 PE=1 SV=3 |
| | | | | | | **0** | | | | | | P26772\|CH10_RAT | **+** | 39 | 28533 | 122.56 | 38 | 38 | 1075300 | **0.008544818** | 6 | 6 | 6 | Oxidation (M) | | 10 kDa heat shock protein, mitochondrial |
| | | | | | | **0** | | | | | | P32198\|CPT1A_RAT | **+** | 76 | 1322 | 62.37 | 3 | 3 | 169930 | **0.00135034** | 2 | 2 | 3 | | 88126 | Carnitine O-palmitoyltransferase 1 liver isoform OS=Rattus norvegicus OX=10116 GN=Cpt1a PE=1 SV=2 |
| | | | | | | **0** | | | | | | P19804\|NDKB_RAT | **+** | 78 | 34377 | 74.8 | 15 | 15 | 793250 | **0.006303521** | 2 | 2 | 3 | Formyla | 17283 | Nucleoside diphosphate kinase B OS=Rattus norvegicus OX=10116 GN=Nme2 PE=1 SV=1 |
| | | | | | | **0** | | | | | | Q3KR86\|MIC60_RA | **+** | 90 | 110 | 74.63 | 3 | 3 | 1524400 | **0.012113568** | 2 | 2 | 2 | | 67177 | MICOS complex subunit Mic60 (Fragment) OS=Rattus norvegicus OX=10116 GN=Immt PE=1 SV=1 |
| | | | | | | **0** | | | | | | Q5XII9\|MFR1L_RAT | **+** | 120 | 5882 | 35.65 | 5 | 5 | 383020 | **0.003043649** | 2 | 1 | 2 | | 31730 | Mitochondrial fission regulator 1-like OS=Rattus norvegicus OX=10116 GN=Mtfr1l PE=1 SV=1 |
| 106 | #### | 21.4 | 5 | 5 | 4460100 | **0.048705646** | 1 | 1 | 2 | For | 30798 | Q9R1Z0\|VDAC3_RA | **-** | | | | | | | **0** | | | | | | Voltage-dependent anion-selective channel protein 3 OS=Rattus norvegicus OX=10116 GN=Vdac3 PE=1 SV=2 |
| | | | | | 9157254498 | **100** | | | | | | | | | | | | | 12584235940 | **100** | | | | | | |
| | | | | | | | | | | | | | | | | | | | 125842359.4 | | | | | | | |
| | | | | | 91572544.98 | | | | | | | | | | | | | | | | | | | | | |
| | | | | | | | | | | | | | | | | | | | 125842359.4 | | | | | | | |
