## Supplementary material for "Permeability transition pore-related changes in the proteome and channel activity of ATP synthase dimers and monomers": ATP-synthase

| Area (IBAQ) CONTROL | riBAQ % | Accession | rIBAQ PTP/Control | Area (IBAQ) PTP | riBAQ % | Description |
|---|---|---|---|---|---|---|
| Complex I | | | | | | |
| | | tr\|A0A0G2JVL6\|A0A0G2J | + | 139230 | **0.001106384** | NADH dehydrogenase (Ubiquinone) 1 beta subcomplex 11 (Predicted) OS=Rattus norvegicus OX=10116 GN=Ndufb11 PE=1 SV=1 |
| | | tr\|D4A7L4\|D4A7L4_RAT | + | 135780 | **0.001078969** | NADH dehydrogenase [ubiquinone] 1 alpha subcomplex subunit 8 OS=Rattus norvegicus OX=10116 GN=Ndufa8 PE=1 SV=1 |
| | | tr\|D4A0T0\|D4A0T0_RAT | + | 434220 | **0.003450507** | NADH:ubiquinone oxidoreductase subunit B10 OS=Rattus norvegicus OX=10116 GN=Ndufb10 PE=1 SV=1 |
| | | | | | **0** | |
| Complex III | | | | | **0** | |
| 2660100 | **0.0290491** | Q68FY0\|QCR1_RAT | **0.410219122** | 1499600 | **0.011916496** | Cytochrome b-c1 complex subunit 1 mitochondrial OS=Rattus norvegicus OX=10116 GN=Uqcrc1 PE=1 SV=1 |
| 323100 | **0.00352835** | P00159\|CYB_RAT | **2.152085076** | 955560 | **0.00759331** | Cytochrome b OS=Rattus norvegicus OX=10116 GN=CYTB PE=3 SV=1 |
| 369530 | **0.00403538** | P20788\|UCRI_RAT | **4.210138965** | 2138000 | **0.01698951** | Cytochrome b (Fragment) OS=Rattus norvegicus OX=10116 GN=cytb PE=3 SV=1 |
| 396150 | **0.004326078** | Q5M9I5\|QCR6_RAT | - | 0 | **0** | Cytochrome b-c1 complex subunit 6 mitochondrial OS=Rattus norvegicus OX=10116 GN=Uqcrh PE=3 SV=1 |
| | **0** | | | | **0** | |
| Complex IV | | | | | **0** | |
| | **0** | P11240\|COX5A_RAT | + | 529460 | **0.004207327** | Cytochrome c oxidase subunit 5A mitochondrial OS=Rattus norvegicus OX=10116 GN=Cox5a PE=1 SV=1 |
| | **0** | P35171\|CX7A2_RAT | + | 311870 | **0.002478259** | Cytochrome c oxidase subunit 2 OS=Rattus norvegicus OX=10116 GN=Mt-co2 PE=1 SV=1 |
| | **0** | P10888\|COX41_RAT | + | 237750 | **0.001889268** | Cytochrome c oxidase subunit 4 isoform 1 mitochondrial OS=Rattus norvegicus OX=10116 GN=Cox4i1 PE=1 SV=1 |
| | **0** | | | | **0** | |
| Complex V | | | | | **0** | |
| 5323500 | **0.058134237** | P29419\|ATP5I_RAT | **9.036804771** | 66111000 | **0.525347747** | ATP synthase subunit e mitochondrial OS=Rattus norvegicus OX=10116 GN=Atp5me PE=1 SV=3 |
| 4150100 | **0.045320352** | P21571\|ATP5J_RAT | **7.97829989** | 45502000 | **0.361579362** | ATP synthase-coupling factor 6 mitochondrial OS=Rattus norvegicus OX=10116 GN=Atp5pf PE=1 SV=1 |
| | **0** | D3Z9R8\|ATP68_RAT | + | 306390 | **0.002434713** | ATP synthase subunit ATP5MPL mitochondrial OS=Rattus norvegicus OX=10116 GN=Atp5mpl PE=1 SV=1 |
| | **0** | P29418\|ATP5E_RAT | + | 727560 | **0.005781519** | ATP synthase subunit epsilon mitochondrial OS=Rattus norvegicus OX=10116 GN=Atp5f1e PE=1 SV=2 |
| | **0** | | | | **0** | |
| Other | | | | | **0** | |
| 15340000 | **0.167517458** | Q02253\|MMSA_RAT | **0.40094793** | 8452300 | **0.067165778** | Methylmalonate-semialdehyde dehydrogenase [acylating] mitochondrial OS=Rattus norvegicus OX=10116 GN=Aldh6a1 PE=1 SV=1 |
| 23897000 | **0.260962497** | P0C2X9\|AL4A1_RAT | **0.483036971** | 15863000 | **0.126054534** | Delta-1-pyrroline-5-carboxylate dehydrogenase mitochondrial OS=Rattus norvegicus OX=10116 GN=Aldh4a1 PE=1 SV=1 |
| 8447200 | **0.092245989** | P10860\|DHE3_RAT | **10.15983789** | 117940000 | **0.937204297** | Glutamate dehydrogenase 1, mitochondrial |
| 1883600 | **0.020569484** | Q8CGU6\|NICA_RAT | - | | **0** | Nicastrin OS=Rattus norvegicus OX=10116 GN=Ncstn PE=1 SV=1 |
| 1237900 | **0.013518244** | P24329\|THTR_RAT | **2.732534818** | 4648500 | **0.036939072** | Thiosulfate sulfurtransferase OS=Rattus norvegicus OX=10116 GN=Tst PE=1 SV=3 |
| 406090 | **0.004434626** | Q924S5\|LONM_RAT | **2.102089349** | 1173100 | **0.00932198** | Lon protease homolog mitochondrial OS=Rattus norvegicus OX=10116 GN=Lonp1 PE=2 SV=1 |
| 509790 | **0.005567062** | A0A0G2K047\|ACSS3_RAT | **0.217122528** | 152110 | **0.001208734** | Acyl-CoA synthetase short-chain family member 3 mitochondrial OS=Rattus norvegicus OX=10116 GN=Acss3 PE=1 SV=1 |
| 14366000 | **0.156881083** | P97564\|GPAT1_RAT | **0.058878694** | 1162400 | **0.009236953** | Glycerol-3-phosphate acyltransferase 1 mitochondrial OS=Rattus norvegicus OX=10116 GN=Gpam PE=1 SV=3 |
| 418350 | **0.004568509** | A2VCW9\|AASS_RAT | **3.429394667** | 1971600 | **0.015667221** | Alpha-aminoadipic semialdehyde synthase mitochondrial OS=Rattus norvegicus OX=10116 GN=Aass PE=2 SV=1 |
| 767510 | **0.008381442** | Q9WVK3\|PECR_RAT | **0.369114498** | 389320 | **0.003093712** | Peroxisomal trans-2-enoyl-CoA reductase OS=Rattus norvegicus OX=10116 GN=Pecr PE=2 SV=1 |
| 214350 | **0.002340767** | P07824\|ARGI1_RAT | - | 0 | **0** | Arginase-1 OS=Rattus norvegicus OX=10116 GN=Arg1 PE=1 SV=2 |
| 2745300 | **0.02997951** | tr\|Q5BJZ3\|Q5BJZ3_RAT | **0.127383834** | 480580 | **0.003818905** | Nicotinamide nucleotide transhydrogenase OS=Rattus norvegicus OX=10116 GN=Nnt PE=1 SV=1 |
| 851720 | **0.009301041** | tr\|G3V6I5\|G3V6I5_RAT | **0.180466745** | 211230 | **0.001678529** | DnaJ heat shock protein family (Hsp40) member A3 OS=Rattus norvegicus OX=10116 GN=Dnaja3 PE=1 SV=2 |
| 300990 | **0.003286902** | P16970\|ABCD3_RAT | - | | **0** | ATP-binding cassette sub-family D member 3 OS=Rattus norvegicus OX=10116 GN=Abcd3 PE=1 SV=3 |
| 565230 | **0.006172483** | O70351\|HCD2_RAT | **2.542484638** | 1974900 | **0.015693444** | 3-hydroxyacyl-CoA dehydrogenase type-2 OS=Rattus norvegicus OX=10116 GN=Hsd17b10 PE=1 SV=3 |
| 74258 | **0.00081092** | P85834\|EFTU_RAT | **14.142354** | 1443200 | **0.011468316** | Elongation factor Tu mitochondrial OS=Rattus norvegicus OX=10116 GN=Tufm PE=1 SV=1 |
| | **0** | tr\|D3ZTP0\|D3ZTP0_RAT | + | 2779100 | **0.022083979** | 10-formyltetrahydrofolate dehydrogenase OS=Rattus norvegicus OX=10116 GN=Aldh1l2 PE=1 SV=3 |
| | **0** | D3ZX08\|SYAM_RAT | + | 305160 | **0.002424939** | Alanine--tRNA ligase mitochondrial OS=Rattus norvegicus OX=10116 GN=Aars2 PE=3 SV=1 |
| | **0** | P08011\|MGST1_RAT | + | 1101000 | **0.008749041** | Microsomal glutathione S-transferase 1 OS=Rattus norvegicus OX=10116 GN=Mgst1 PE=1 SV=3 |

| | | | | | | |
|---|---|---|---|---|---|---|
| | **0** | P26772\|CH10_RAT | **+** | 1075300 | **0.008544818** | 10 kDa heat shock protein, mitochondrial |
| | **0** | P32198\|CPT1A_RAT | **+** | 169930 | **0.00135034** | Carnitine O-palmitoyltransferase 1 liver isoform OS=Rattus norvegicus OX=10116 GN=Cpt1a PE=1 SV=2 |
| | **0** | P19804\|NDKB_RAT | **+** | 793250 | **0.006303521** | Nucleoside diphosphate kinase B OS=Rattus norvegicus OX=10116 GN=Nme2 PE=1 SV=1 |
| | **0** | Q3KR86\|MIC60_RAT | **+** | 1524400 | **0.012113568** | MICOS complex subunit Mic60 (Fragment) OS=Rattus norvegicus OX=10116 GN=Immt PE=1 SV=1 |
| | **0** | Q5XII9\|MFR1L_RAT | **+** | 383020 | **0.003043649** | Mitochondrial fission regulator 1-like OS=Rattus norvegicus OX=10116 GN=Mtfr1l PE=1 SV=1 |
| 4460100 | **0.048705646** | Q9R1Z0\|VDAC3_RAT | **-** | | **0** | Voltage-dependent anion-selective channel protein 3 OS=Rattus norvegicus OX=10116 GN=Vdac3 PE=1 SV=2 |
