## Supplementary material for "Permeability transition pore-related changes in the proteome and channel activity of ATP synthase dimers and monomers": RHM

| Accession | Protein Group | Protein ID | -10lgP | Coverage (%) | Coverage (%) Sample 7 | IBAQ | #Peptides | #Unique | #Spec Sample 7 | PTM | Avg. Mass | Description |
|---|---|---|---|---|---|---|---|---|---|---|---|---|
| P10719\|AT | 2 | 1 | 327.61 | 37 | 37 | 55442000 | 81 | 81 | 184 | Y | 56354 | ATP synthase subunit beta, mitochondrial OS=Rattus norvegicus OX=10116 GN=Atp5f1b PE=1 SV=2 |
| tr\|G3V6D3 | 2 | 2 | 327.61 | 37 | 37 | 55442000 | 81 | 81 | 184 | Y | 56345 | ATP synthase subunit beta OS=Rattus norvegicus OX=10116 GN=Atp5f1b PE=1 SV=1 |
| P15999\|AT | 1 | 3 | 278.06 | 44 | 44 | 50080000 | 81 | 79 | 222 | Y | 59754 | ATP synthase subunit alpha, mitochondrial OS=Rattus norvegicus OX=10116 GN=Atp5f1a PE=1 SV=2 |
| tr\|F1LP05\| | 1 | 4 | 278.06 | 44 | 44 | 50080000 | 81 | 79 | 222 | Y | 59813 | ATP synthase subunit alpha OS=Rattus norvegicus OX=10116 GN=Atp5f1a PE=1 SV=1 |
| P35435\|AT | 5 | 5 | 236.59 | 32 | 32 | 10349000 | 26 | 26 | 65 | Y | 30191 | ATP synthase subunit gamma, mitochondrial OS=Rattus norvegicus OX=10116 GN=Atp5f1c PE=1 SV=2 |
| tr\|Q6QI09\| | 5 | 6 | 236.59 | 14 | 14 | 10349000 | 26 | 26 | 65 | Y | 67721 | ATP synthase subunit gamma, mitochondrial OS=Rattus norvegicus OX=10116 GN=Taf3 PE=1 SV=1 |
| tr\|D3ZFQ8 | 9 | 133 | 235.99 | 29 | 29 | 9715200 | 22 | 22 | 52 | Y | 35435 | Cytochrome c-1 OS=Rattus norvegicus OX=10116 GN=Cyc1 PE=1 SV=3 |
| Q68FY0\|Q | 3 | 118 | 221.36 | 31 | 31 | 23228000 | 39 | 39 | 99 | Y | 52849 | Cytochrome b-c1 complex subunit 1, mitochondrial OS=Rattus norvegicus OX=10116 GN=Uqcrc1 PE=1 SV=1 |
| Q64428\|E | 7 | 9 | 216.18 | 23 | 23 | 8785200 | 34 | 34 | 59 | Y | 82665 | Trifunctional enzyme subunit alpha, mitochondrial OS=Rattus norvegicus OX=10116 GN=Hadha PE=1 SV=2 |
| Q60587\|E | 10 | 20 | 214.31 | 23 | 23 | 11020000 | 23 | 23 | 39 | Y | 51414 | Trifunctional enzyme subunit beta, mitochondrial OS=Rattus norvegicus OX=10116 GN=Hadhb PE=1 SV=1 |
| P00507\|A | 8 | 19 | 209.44 | 29 | 29 | 8210600 | 27 | 27 | 53 | Y | 47314 | Aspartate aminotransferase, mitochondrial OS=Rattus norvegicus OX=10116 GN=Got2 PE=1 SV=2 |
| P20788\|U | 12 | 132 | 201.11 | 37 | 37 | 4900700 | 22 | 22 | 36 | N | 29446 | Cytochrome b-c1 complex subunit Rieske, mitochondrial OS=Rattus norvegicus OX=10116 GN=Uqcrfs1 PE=1 SV=2 |
| P32551\|Q | 4 | 34 | 198.35 | 30 | 30 | 19133000 | 38 | 38 | 87 | Y | 48396 | Cytochrome b-c1 complex subunit 2, mitochondrial OS=Rattus norvegicus OX=10116 GN=Uqcrc2 PE=1 SV=2 |
| P11240\|CC | 11 | 86 | 183.95 | 52 | 52 | 7033400 | 18 | 18 | 38 | N | 16130 | Cytochrome c oxidase subunit 5A, mitochondrial OS=Rattus norvegicus OX=10116 GN=Cox5a PE=1 SV=1 |
| P31399\|AT | 16 | 7 | 183.4 | 39 | 39 | 4120100 | 16 | 16 | 28 | N | 18763 | ATP synthase subunit d, mitochondrial OS=Rattus norvegicus OX=10116 GN=Atp5pd PE=1 SV=3 |
| tr\|A0A097 | 13 | 53 | 175.7 | 16 | 16 | 572310 | 15 | 1 | 35 | N | 42865 | Cytochrome b OS=Rattus norvegicus OX=10116 GN=CYTB PE=3 SV=1 |
| tr\|A0A411 | 13 | 44 | 175.7 | 16 | 16 | 572310 | 15 | 1 | 35 | N | 42238 | Cytochrome b (Fragment) OS=Rattus norvegicus OX=10116 GN=Cytb PE=3 SV=1 |
| tr\|L0L4L8\| | 13 | 45 | 175.7 | 16 | 16 | 572310 | 15 | 1 | 35 | N | 42470 | Cytochrome b (Fragment) OS=Rattus norvegicus OX=10116 GN=cytb PE=3 SV=1 |
| tr\|A0A140 | 13 | 46 | 175.7 | 16 | 16 | 572310 | 15 | 1 | 35 | N | 42574 | Cytochrome b (Fragment) OS=Rattus norvegicus OX=10116 PE=3 SV=1 |
| tr\|A0A140 | 13 | 47 | 175.7 | 16 | 16 | 572310 | 15 | 1 | 35 | N | 42588 | Cytochrome b (Fragment) OS=Rattus norvegicus OX=10116 PE=3 SV=1 |
| tr\|A0A385 | 13 | 49 | 175.7 | 16 | 16 | 572310 | 15 | 1 | 35 | N | 42766 | Cytochrome b (Fragment) OS=Rattus norvegicus OX=10116 PE=3 SV=1 |
| tr\|A0A3Q8 | 13 | 50 | 175.7 | 16 | 16 | 572310 | 15 | 1 | 35 | N | 42712 | Cytochrome b (Fragment) OS=Rattus norvegicus OX=10116 GN=Cytb PE=3 SV=1 |
| tr\|A0A3Q8 | 13 | 38 | 175.7 | 16 | 16 | 572310 | 15 | 1 | 35 | N | 42622 | Cytochrome b (Fragment) OS=Rattus norvegicus OX=10116 GN=Cytb PE=3 SV=1 |
| tr\|A0A3S6 | 13 | 52 | 175.7 | 16 | 16 | 572310 | 15 | 1 | 35 | N | 42698 | Cytochrome b (Fragment) OS=Rattus norvegicus OX=10116 GN=Cytb PE=3 SV=1 |
| tr\|A0A220 | 13 | 54 | 175.7 | 16 | 16 | 572310 | 15 | 1 | 35 | N | 43018 | Cytochrome b OS=Rattus norvegicus OX=10116 PE=3 SV=1 |
| tr\|D6NSP2 | 13 | 55 | 175.7 | 16 | 16 | 572310 | 15 | 1 | 35 | N | 42945 | Cytochrome b (Fragment) OS=Rattus norvegicus OX=10116 GN=cytb PE=3 SV=1 |
| tr\|D6NSS6 | 13 | 56 | 175.7 | 16 | 16 | 572310 | 15 | 1 | 35 | N | 42993 | Cytochrome b (Fragment) OS=Rattus norvegicus OX=10116 GN=cytb PE=3 SV=1 |
| tr\|D6NSQ3 | 13 | 57 | 175.7 | 16 | 16 | 572310 | 15 | 1 | 35 | N | 43012 | Cytochrome b (Fragment) OS=Rattus norvegicus OX=10116 GN=cytb PE=3 SV=1 |
| tr\|D6NSR3 | 13 | 58 | 175.7 | 16 | 16 | 572310 | 15 | 1 | 35 | N | 42986 | Cytochrome b (Fragment) OS=Rattus norvegicus OX=10116 GN=cytb PE=3 SV=1 |
| tr\|D6NSP4 | 13 | 59 | 175.7 | 16 | 16 | 572310 | 15 | 1 | 35 | N | 43016 | Cytochrome b (Fragment) OS=Rattus norvegicus OX=10116 GN=cytb PE=3 SV=1 |
| tr\|D6NSR8 | 13 | 61 | 175.7 | 16 | 16 | 572310 | 15 | 1 | 35 | N | 42998 | Cytochrome b (Fragment) OS=Rattus norvegicus OX=10116 GN=cytb PE=3 SV=1 |
| tr\|F8QU37 | 13 | 62 | 175.7 | 16 | 16 | 572310 | 15 | 1 | 35 | N | 42992 | Cytochrome b (Fragment) OS=Rattus norvegicus OX=10116 GN=cytb PE=3 SV=1 |
| tr\|F2Q6S6\| | 13 | 63 | 175.7 | 16 | 16 | 572310 | 15 | 1 | 35 | N | 42938 | Cytochrome b (Fragment) OS=Rattus norvegicus OX=10116 GN=cytb PE=3 SV=1 |
| tr\|A0A0A1 | 13 | 64 | 175.7 | 16 | 16 | 572310 | 15 | 1 | 35 | N | 42989 | Cytochrome b OS=Rattus norvegicus OX=10116 GN=CYTB PE=3 SV=1 |
| tr\|D6NSS7 | 13 | 65 | 175.7 | 16 | 16 | 572310 | 15 | 1 | 35 | N | 42948 | Cytochrome b (Fragment) OS=Rattus norvegicus OX=10116 GN=cytb PE=3 SV=1 |
| tr\|Q8SEY9\| | 13 | 66 | 175.7 | 16 | 16 | 572310 | 15 | 1 | 35 | N | 43015 | Cytochrome b OS=Rattus norvegicus OX=10116 GN=cytb PE=3 SV=1 |
| tr\|D6NSR7 | 13 | 67 | 175.7 | 16 | 16 | 572310 | 15 | 1 | 35 | N | 42982 | Cytochrome b (Fragment) OS=Rattus norvegicus OX=10116 GN=cytb PE=3 SV=1 |
| tr\|A0A220 | 13 | 68 | 175.7 | 16 | 16 | 572310 | 15 | 1 | 35 | N | 42968 | Cytochrome b OS=Rattus norvegicus OX=10116 PE=3 SV=1 |
| tr\|D6NSQ0 | 13 | 71 | 175.7 | 16 | 16 | 572310 | 15 | 1 | 35 | N | 42968 | Cytochrome b (Fragment) OS=Rattus norvegicus OX=10116 GN=cytb PE=3 SV=1 |
| tr\|Q5UAI7 | 13 | 72 | 175.7 | 16 | 16 | 572310 | 15 | 1 | 35 | N | 42998 | Cytochrome b OS=Rattus norvegicus OX=10116 GN=CYTB PE=3 SV=1 |
| tr\|D6NSR2 | 13 | 73 | 175.7 | 16 | 16 | 572310 | 15 | 1 | 35 | N | 43073 | Cytochrome b (Fragment) OS=Rattus norvegicus OX=10116 GN=cytb PE=3 SV=1 |
| tr\|A0A0S1 | 13 | 74 | 175.7 | 16 | 16 | 572310 | 15 | 1 | 35 | N | 43002 | Cytochrome b OS=Rattus norvegicus OX=10116 GN=CYTB PE=3 SV=1 |
| tr\|D6NSR1 | 13 | 75 | 175.7 | 16 | 16 | 572310 | 15 | 1 | 35 | N | 43002 | Cytochrome b (Fragment) OS=Rattus norvegicus OX=10116 GN=cytb PE=3 SV=1 |
| tr\|Q8HIC4\| | 13 | 76 | 175.7 | 16 | 16 | 572310 | 15 | 1 | 35 | N | 43012 | Cytochrome b OS=Rattus norvegicus OX=10116 GN=Mt-cyb PE=3 SV=1 |
| tr\|A0A220 | 13 | 77 | 175.7 | 16 | 16 | 572310 | 15 | 1 | 35 | N | 42968 | Cytochrome b OS=Rattus norvegicus OX=10116 PE=3 SV=1 |
| tr\|A0A0S1 | 13 | 78 | 175.7 | 16 | 16 | 572310 | 15 | 1 | 35 | N | 43016 | Cytochrome b OS=Rattus norvegicus OX=10116 GN=CYTB PE=3 SV=1 |
| tr\|R9TKN1 | 13 | 119 | 175.7 | 16 | 16 | 572310 | 15 | 1 | 35 | N | 43012 | Cytochrome b OS=Rattus norvegicus OX=10116 GN=CYTB PE=3 SV=1 |
| P00159\|CY | 13 | 79 | 175.7 | 16 | 16 | 572310 | 15 | 1 | 35 | N | 43012 | Cytochrome b OS=Rattus norvegicus OX=10116 GN=Mt-Cyb PE=3 SV=3 |
| tr\|L0N311\| | 13 | 80 | 175.7 | 16 | 16 | 572310 | 15 | 1 | 35 | N | 43045 | Cytochrome b (Fragment) OS=Rattus norvegicus OX=10116 GN=cytb PE=3 SV=1 |
| tr\|A0A220 | 13 | 81 | 175.7 | 16 | 16 | 572310 | 15 | 1 | 35 | N | 42970 | Cytochrome b OS=Rattus norvegicus OX=10116 PE=3 SV=1 |
| tr\|H2KXA0 | 13 | 82 | 175.7 | 16 | 16 | 572310 | 15 | 1 | 35 | N | 42978 | Cytochrome b (Fragment) OS=Rattus norvegicus OX=10116 GN=cytb PE=3 SV=1 |
| tr\|A0A096 | 13 | 83 | 175.7 | 16 | 16 | 572310 | 15 | 1 | 35 | N | 43042 | Cytochrome b (Fragment) OS=Rattus norvegicus OX=10116 PE=3 SV=1 |
| tr\|D6NSP5 | 13 | 120 | 175.7 | 16 | 16 | 572310 | 15 | 1 | 35 | N | 43012 | Cytochrome b (Fragment) OS=Rattus norvegicus OX=10116 GN=cytb PE=3 SV=1 |
| Q9ER34\|A | 17 | 88 | 175.14 | 13 | 13 | 3358900 | 17 | 17 | 27 | N | 85433 | Aconitate hydratase, mitochondrial OS=Rattus norvegicus OX=10116 GN=Aco2 PE=1 SV=2 |
| tr\|Q5UAJ6 | 6 | 24 | 173.47 | 49 | 49 | 10697000 | 33 | 33 | 60 | Y | 25942 | Cytochrome c oxidase subunit 2 OS=Rattus norvegicus OX=10116 GN=COX2 PE=3 SV=1 |
| P14882\|PC | 14 | 2783 | 161.44 | 16 | 16 | 2493200 | 19 | 17 | 32 | Y | 81623 | Propionyl-CoA carboxylase alpha chain, mitochondrial OS=Rattus norvegicus OX=10116 GN=Pcca PE=1 SV=3 |
| tr\|A0A385 | 15 | 212 | 161.08 | 16 | 16 | 46246 | 15 | 1 | 32 | N | 42751 | Cytochrome b (Fragment) OS=Rattus norvegicus OX=10116 PE=3 SV=1 |

| | | | | | | | | | | | | |
|---|---|---|---|---|---|---|---|---|---|---|---|---|
| tr\|A0A0G2 | **18** | 147 | 134.38 | 14 | 14 | 3172100 | 15 | 15 | 23 | Y | 86230 | MICOS complex subunit MIC60 OS=Rattus norvegicus OX=10116 GN=Immt PE=1 SV=1 |
| P10817\|CX | **27** | 151 | 129.89 | 15 | 15 | 2204700 | 5 | 5 | 10 | N | 10487 | Cytochrome c oxidase subunit 6A2, mitochondrial (Fragment) OS=Rattus norvegicus OX=10116 GN=Cox6a2 PE=1 SV=3 |
| tr\|G3V8M4 | **27** | 152 | 129.89 | 14 | 14 | 2204700 | 5 | 5 | 10 | N | 10802 | Cytochrome c oxidase subunit 6A, mitochondrial OS=Rattus norvegicus OX=10116 GN=Cox6a2 PE=3 SV=1 |
| P80431\|CC | **33** | 2798 | 121.84 | 29 | 29 | 1469200 | 6 | 6 | 7 | N | 8995 | Cytochrome c oxidase subunit 7B, mitochondrial OS=Rattus norvegicus OX=10116 GN=Cox7b PE=1 SV=3 |
| tr\|B2RYT5\| | **20** | 2788 | 117.53 | 43 | 43 | 1649100 | 12 | 12 | 19 | Y | 12651 | Cox7a2l protein OS=Rattus norvegicus OX=10116 GN=Cox7a2l PE=2 SV=1 |
| Q6AXV4\|SA | **31** | 2803 | 108.68 | 9 | 9 | 330220 | 6 | 6 | 8 | N | 51960 | Sorting and assembly machinery component 50 homolog OS=Rattus norvegicus OX=10116 GN=Samm50 PE=1 SV=1 |
| P10888\|CC | **19** | 290 | 106.92 | 25 | 25 | 1754300 | 10 | 10 | 20 | Y | 19515 | Cytochrome c oxidase subunit 4 isoform 1, mitochondrial OS=Rattus norvegicus OX=10116 GN=Cox4i1 PE=1 SV=1 |
| P56574\|ID | **25** | 2786 | 101.89 | 13 | 13 | 843110 | 9 | 9 | 10 | N | 50967 | Isocitrate dehydrogenase [NADP], mitochondrial OS=Rattus norvegicus OX=10116 GN=Idh2 PE=1 SV=2 |
| P04636\|MI | **21** | 561 | 97.44 | 18 | 18 | 943230 | 10 | 9 | 15 | Y | 35684 | Malate dehydrogenase, mitochondrial OS=Rattus norvegicus OX=10116 GN=Mdh2 PE=1 SV=2 |
| Q06647\|AT | **23** | 8 | 92.62 | 13 | 13 | 1125300 | 6 | 6 | 12 | N | 23398 | ATP synthase subunit O, mitochondrial OS=Rattus norvegicus OX=10116 GN=Atp5po PE=1 SV=1 |
| tr\|Q5BJZ3\| | **35** | 1175 | 91.49 | 4 | 4 | 391640 | 4 | 4 | 6 | N | 113869 | Nicotinamide nucleotide transhydrogenase OS=Rattus norvegicus OX=10116 GN=Nnt PE=1 SV=1 |
| P17764\|TH | **28** | 2787 | 88.78 | 15 | 15 | 766390 | 7 | 7 | 9 | N | 44695 | Acetyl-CoA acetyltransferase, mitochondrial OS=Rattus norvegicus OX=10116 GN=Acat1 PE=1 SV=1 |
| P45953\|AC | **26** | 503 | 88.01 | 8 | 8 | 1324300 | 6 | 6 | 10 | N | 70749 | Very long-chain specific acyl-CoA dehydrogenase, mitochondrial OS=Rattus norvegicus OX=10116 GN=Acadvl PE=1 SV=1 |
| tr\|Q5M9H2 | **26** | 504 | 88.01 | 8 | 8 | 1324300 | 6 | 6 | 10 | N | 70821 | Acyl-Coenzyme A dehydrogenase, very long chain OS=Rattus norvegicus OX=10116 GN=Acadvl PE=1 SV=1 |
| P12075\|CC | **22** | 139 | 86.99 | 22 | 22 | 6260800 | 6 | 6 | 15 | Y | 13915 | Cytochrome c oxidase subunit 5B, mitochondrial OS=Rattus norvegicus OX=10116 GN=Cox5b PE=1 SV=2 |
| Q6PDU7\|A | **34** | 11 | 85.86 | 26 | 26 | 666270 | 3 | 3 | 6 | Y | 11433 | ATP synthase subunit g, mitochondrial OS=Rattus norvegicus OX=10116 GN=Atp5mg PE=1 SV=2 |
| tr\|Q06QA9 | **29** | 2790 | 82.74 | 7 | 7 | 602690 | 5 | 5 | 9 | Y | 56893 | Cytochrome c oxidase subunit 1 OS=Rattus norvegicus OX=10116 GN=CO1 PE=3 SV=1 |
| tr\|Q06QK0 | **29** | 2791 | 82.74 | 7 | 7 | 602690 | 5 | 5 | 9 | Y | 56880 | Cytochrome c oxidase subunit 1 OS=Rattus norvegicus OX=10116 GN=CO1 PE=3 SV=1 |
| tr\|Q8SEZ6\| | **29** | 2792 | 82.74 | 7 | 7 | 602690 | 5 | 5 | 9 | Y | 56879 | Cytochrome c oxidase subunit 1 OS=Rattus norvegicus OX=10116 GN=COI PE=3 SV=1 |
| tr\|A0A0A1 | **29** | 2793 | 82.74 | 7 | 7 | 602690 | 5 | 5 | 9 | Y | 56937 | Cytochrome c oxidase subunit 1 OS=Rattus norvegicus OX=10116 GN=COX1 PE=3 SV=1 |
| tr\|Q8HIC9\| | **29** | 2794 | 82.74 | 7 | 7 | 602690 | 5 | 5 | 9 | Y | 56845 | Cytochrome c oxidase subunit 1 OS=Rattus norvegicus OX=10116 GN=Mt-co1 PE=3 SV=1 |
| P05503\|CC | **29** | 2795 | 82.74 | 7 | 7 | 602690 | 5 | 5 | 9 | Y | 56845 | Cytochrome c oxidase subunit 1 OS=Rattus norvegicus OX=10116 GN=Mtco1 PE=2 SV=3 |
| tr\|Q95938 | **29** | 2796 | 82.74 | 7 | 7 | 602690 | 5 | 5 | 9 | Y | 56977 | Cytochrome c oxidase subunit 1 OS=Rattus norvegicus OX=10116 GN=Co I PE=3 SV=1 |
| tr\|D3ZUX5 | **53** | 2815 | 80.37 | 9 | 9 | 426040 | 2 | 2 | 3 | N | 26435 | MICOS complex subunit OS=Rattus norvegicus OX=10116 GN=Chchd3 PE=1 SV=1 |
| P11507\|AT | **49** | 2797 | 78.39 | 3 | 3 | 308840 | 3 | 3 | 3 | N | 114768 | Sarcoplasmic/endoplasmic reticulum calcium ATPase 2 OS=Rattus norvegicus OX=10116 GN=Atp2a2 PE=1 SV=1 |
| tr\|G3V7I0\| | **55** | 2858 | 77.47 | 5 | 5 | 292970 | 2 | 2 | 3 | N | 28299 | Peroxiredoxin 3 OS=Rattus norvegicus OX=10116 GN=Prdx3 PE=1 SV=1 |
| Q9Z0V6\|PF | **55** | 2859 | 77.47 | 5 | 5 | 292970 | 2 | 2 | 3 | N | 28295 | Thioredoxin-dependent peroxide reductase, mitochondrial OS=Rattus norvegicus OX=10116 GN=Prdx3 PE=1 SV=2 |
| P19511\|AT | **32** | 10 | 76.7 | 9 | 9 | 2794800 | 3 | 3 | 7 | N | 28869 | ATP synthase F(0) complex subunit B1, mitochondrial OS=Rattus norvegicus OX=10116 GN=Atp5pb PE=1 SV=1 |
| P11951\|CX | **39** | 2816 | 75.09 | 22 | 22 | 875870 | 3 | 3 | 5 | N | 8455 | Cytochrome c oxidase subunit 6C-2 OS=Rattus norvegicus OX=10116 GN=Cox6c2 PE=1 SV=3 |
| tr\|Q5UAJ5 | **40** | 32 | 74.57 | 19 | 19 | 2120600 | 3 | 3 | 5 | Y | 7642 | ATP synthase protein 8 OS=Rattus norvegicus OX=10116 GN=ATP8 PE=3 SV=1 |
| P11608\|AT | **40** | 35 | 74.57 | 19 | 19 | 2120600 | 3 | 3 | 5 | Y | 7630 | ATP synthase protein 8 OS=Rattus norvegicus OX=10116 GN=Mt-atp8 PE=1 SV=2 |
| tr\|Q8SEZ4\| | **40** | 33 | 74.57 | 19 | 19 | 2120600 | 3 | 3 | 5 | Y | 7632 | ATP synthase protein 8 OS=Rattus norvegicus OX=10116 GN=ATPase8 PE=3 SV=1 |
| tr\|Q8HIC8\| | **40** | 36 | 74.57 | 19 | 19 | 2120600 | 3 | 3 | 5 | Y | 7630 | ATP synthase protein 8 OS=Rattus norvegicus OX=10116 GN=Mt-atp8 PE=3 SV=1 |
| P07633\|PC | **24** | 2799 | 74.26 | 7 | 7 | 988350 | 5 | 5 | 10 | N | 58626 | Propionyl-CoA carboxylase beta chain, mitochondrial OS=Rattus norvegicus OX=10116 GN=Pccb PE=2 SV=1 |
| tr\|Q68FZ8\| | **24** | 2800 | 74.26 | 7 | 7 | 988350 | 5 | 5 | 10 | N | 58678 | Propionyl coenzyme A carboxylase, beta polypeptide OS=Rattus norvegicus OX=10116 GN=Pccb PE=1 SV=1 |
| tr\|F1LP30\| | **54** | 2818 | 72.91 | 3 | 3 | 36580 | 2 | 1 | 3 | N | 79296 | Methylcrotonoyl-CoA carboxylase subunit alpha, mitochondrial OS=Rattus norvegicus OX=10116 GN=Mccc1 PE=1 SV=1 |
| Q5I0C3\|M | **54** | 2819 | 72.91 | 3 | 3 | 36580 | 2 | 1 | 3 | N | 79330 | Methylcrotonoyl-CoA carboxylase subunit alpha, mitochondrial OS=Rattus norvegicus OX=10116 GN=Mccc1 PE=1 SV=1 |
| P52873\|PY | **45** | 2805 | 70.08 | 3 | 3 | 79061 | 3 | 2 | 4 | N | 129777 | Pyruvate carboxylase, mitochondrial OS=Rattus norvegicus OX=10116 GN=Pc PE=1 SV=2 |
| tr\|A0A0G2 | **45** | 2806 | 70.08 | 2 | 2 | 79061 | 3 | 2 | 4 | N | 140005 | Pyruvate carboxylase, mitochondrial OS=Rattus norvegicus OX=10116 GN=Pc PE=1 SV=1 |
| tr\|B2RYS8\| | **119** | 2857 | 69.95 | 8 | 8 | 134770 | 1 | 1 | 1 | N | 21959 | NADH dehydrogenase [ubiquinone] 1 beta subcomplex subunit 8, mitochondrial OS=Rattus norvegicus OX=10116 GN=Ndufb8 PE=1 SV=1 |
| P09605\|KC | **43** | 84 | 69.38 | 5 | 5 | 176570 | 2 | 2 | 4 | N | 47385 | Creatine kinase S-type, mitochondrial OS=Rattus norvegicus OX=10116 GN=Ckmt2 PE=1 SV=2 |
| tr\|Q7H115 | **36** | 2862 | 68.69 | 3 | 3 | 859140 | 2 | 2 | 6 | N | 29871 | Cytochrome c oxidase subunit 3 OS=Rattus norvegicus OX=10116 GN=Mt-co3 PE=3 SV=1 |
| tr\|I6V4L9\| | **36** | 2863 | 68.69 | 3 | 3 | 859140 | 2 | 2 | 6 | N | 29844 | Cytochrome c oxidase subunit 3 OS=Rattus norvegicus OX=10116 GN=COX3 PE=3 SV=1 |
| tr\|Q8M7G4 | **36** | 2864 | 68.69 | 3 | 3 | 859140 | 2 | 2 | 6 | N | 29861 | Cytochrome c oxidase subunit 3 OS=Rattus norvegicus OX=10116 PE=2 SV=1 |
| tr\|A0A096 | **36** | 2865 | 68.69 | 3 | 3 | 859140 | 2 | 2 | 6 | N | 29901 | Cytochrome c oxidase subunit 3 OS=Rattus norvegicus OX=10116 GN=COX3 PE=3 SV=1 |
| P05505\|CC | **36** | 2866 | 68.69 | 3 | 3 | 859140 | 2 | 2 | 6 | N | 29871 | Cytochrome c oxidase subunit 3 OS=Rattus norvegicus OX=10116 GN=Mtco3 PE=1 SV=5 |
| tr\|Q8SEZ2\| | **36** | 2867 | 68.69 | 3 | 3 | 859140 | 2 | 2 | 6 | N | 29870 | Cytochrome c oxidase subunit 3 (Fragment) OS=Rattus norvegicus OX=10116 GN=COIII PE=3 SV=1 |
| tr\|B1WBP7 | **51** | 25 | 68.23 | 18 | 18 | 853760 | 2 | 2 | 3 | Y | 12884 | ATP synthase subunit delta, mitochondrial OS=Rattus norvegicus OX=10116 GN=Atp5f1d PE=1 SV=1 |
| tr\|G3V7Y3 | **51** | 12 | 68.23 | 13 | 13 | 853760 | 2 | 2 | 3 | Y | 17563 | ATP synthase subunit delta, mitochondrial OS=Rattus norvegicus OX=10116 GN=Atp5f1d PE=1 SV=1 |
| P35434\|AT | **51** | 18 | 68.23 | 13 | 13 | 853760 | 2 | 2 | 3 | Y | 17595 | ATP synthase subunit delta, mitochondrial OS=Rattus norvegicus OX=10116 GN=Atp5f1d PE=1 SV=2 |
| Q02253\|M | **30** | 2801 | 67.09 | 12 | 12 | 798110 | 7 | 7 | 8 | Y | 57808 | Methylmalonate-semialdehyde dehydrogenase [acylating], mitochondrial OS=Rattus norvegicus OX=10116 GN=Aldh6a1 PE=1 SV=1 |
| tr\|G3V7J0\| | **30** | 2802 | 67.09 | 12 | 12 | 798110 | 7 | 7 | 8 | Y | 57748 | Aldehyde dehydrogenase family 6, subfamily A1, isoform CRA_b OS=Rattus norvegicus OX=10116 GN=Aldh6a1 PE=1 SV=1 |
| P85834\|EF | **96** | 2827 | 66.18 | 3 | 3 | 57920 | 1 | 1 | 1 | N | 49522 | Elongation factor Tu, mitochondrial OS=Rattus norvegicus OX=10116 GN=Tufm PE=1 SV=1 |
| P08461\|OD | **44** | 2804 | 50.17 | 4 | 4 | 262470 | 3 | 3 | 4 | N | 67166 | Dihydrolipoyllysine-residue acetyltransferase component of pyruvate dehydrogenase complex, mitochondrial OS=Rattus norvegicus OX=10116 GN=Dlat PE=1 SV=3 |
| P35171\|CX | **37** | 1062 | 49.4 | 14 | 14 | 283320 | 2 | 2 | 6 | N | 9353 | Cytochrome c oxidase subunit 7A2, mitochondrial OS=Rattus norvegicus OX=10116 GN=Cox7a2 PE=1 SV=1 |
| tr\|B2RYS0\| | **37** | 1063 | 49.4 | 14 | 14 | 283320 | 2 | 2 | 6 | N | 9353 | Cox7a2 protein OS=Rattus norvegicus OX=10116 GN=Cox7a2 PE=2 SV=1 |
| tr\|D3ZG43 | **50** | 2807 | 48.84 | 6 | 6 | 406480 | 2 | 2 | 3 | N | 30226 | NADH dehydrogenase (Ubiquinone) Fe-S protein 3 (Predicted), isoform CRA_c OS=Rattus norvegicus OX=10116 GN=Ndufs3 PE=1 SV=1 |
| P80432\|CC | **65** | 134 | 47.77 | 21 | 21 | 134270 | 1 | 1 | 2 | N | 7375 | Cytochrome c oxidase subunit 7C, mitochondrial OS=Rattus norvegicus OX=10116 GN=Cox7c PE=1 SV=2 |
| P63039\|CH | **70** | 142 | 47.17 | 2 | 2 | 412190 | 2 | 2 | 2 | N | 60956 | 60 kDa heat shock protein, mitochondrial OS=Rattus norvegicus OX=10116 GN=Hspd1 PE=1 SV=1 |
| tr\|A0A482 | **70** | 143 | 47.17 | 2 | 2 | 412190 | 2 | 2 | 2 | N | 60956 | Hsp60 OS=Rattus norvegicus OX=10116 GN=Hspd1 PE=2 SV=1 |

| | | | | | | | | | | | | |
|---|---|---|---|---|---|---|---|---|---|---|---|---|
| tr\|D4A7L4\| | **68** | 2872 | 46.96 | 9 | 9 | 201230 | 2 | 2 | 2 | N | 17634 | NADH dehydrogenase (Ubiquinone) 1 beta subcomplex, 11 (Predicted) OS=Rattus norvegicus OX=10116 GN=Ndufb11 PE=1 SV=1 |
| Q63704\|CF | **62** | 138 | 45.37 | 2 | 2 | 215760 | 1 | 1 | 2 | N | 88217 | Carnitine O-palmitoyltransferase 1, muscle isoform OS=Rattus norvegicus OX=10116 GN=Cpt1b PE=1 SV=1 |
| tr\|Q5PQZ9 | **59** | 2821 | 45.37 | 19 | 19 | 194600 | 2 | 2 | 2 | N | 14359 | NADH dehydrogenase [ubiquinone] 1 subunit C2 OS=Rattus norvegicus OX=10116 GN=Ndufc2 PE=1 SV=1 |
| tr\|B2RZD6 | **42** | 568 | 45.17 | 21 | 21 | 366830 | 3 | 3 | 4 | N | 9327 | NDUFA4, mitochondrial complex-associated OS=Rattus norvegicus OX=10116 GN=Ndufa4 PE=1 SV=1 |
| P02770\|AL | **97** | 2832 | 38.54 | 1 | 1 | 83100 | 1 | 1 | 1 | Y | 68731 | Serum albumin OS=Rattus norvegicus OX=10116 GN=Alb PE=1 SV=2 |
| tr\|A0A0G2 | **97** | 2833 | 38.54 | 1 | 1 | 83100 | 1 | 1 | 1 | Y | 68759 | Serum albumin OS=Rattus norvegicus OX=10116 GN=Alb PE=1 SV=1 |
| P19234\|ND | **120** | 2870 | 36.42 | 6 | 6 | 67052 | 1 | 1 | 1 | N | 27378 | NADH dehydrogenase [ubiquinone] flavoprotein 2, mitochondrial OS=Rattus norvegicus OX=10116 GN=Ndufv2 PE=1 SV=2 |
| tr\|A0A0G2 | **52** | 2830 | 35.54 | 4 | 4 | 225440 | 1 | 1 | 3 | N | 20750 | Mitochondrial ribosomal protein L11 OS=Rattus norvegicus OX=10116 GN=mrpl11 PE=1 SV=1 |
| Q5XIE3\|RM | **52** | 2831 | 35.54 | 4 | 4 | 225440 | 1 | 1 | 3 | N | 22420 | 39S ribosomal protein L11, mitochondrial OS=Rattus norvegicus OX=10116 GN=Mrpl11 PE=2 SV=1 |
| Q5HZA9\|T | **121** | 2871 | 35.32 | 5 | 5 | 83601 | 1 | 1 | 1 | N | 21657 | Transmembrane protein 126A OS=Rattus norvegicus OX=10116 GN=Tmem126a PE=2 SV=1 |
| P13086\|SU | **60** | 2822 | 35.23 | 7 | 7 | 82652 | 2 | 2 | 2 | N | 36148 | Succinate--CoA ligase [ADP/GDP-forming] subunit alpha, mitochondrial OS=Rattus norvegicus OX=10116 GN=Suclg1 PE=2 SV=2 |
| tr\|A0A0H2 | **60** | 2823 | 35.23 | 6 | 6 | 82652 | 2 | 2 | 2 | N | 37560 | Succinate--CoA ligase [ADP/GDP-forming] subunit alpha, mitochondrial OS=Rattus norvegicus OX=10116 GN=Suclg1 PE=1 SV=1 |
| tr\|B2RZ57\| | **74** | 2880 | 33.43 | 8 | 8 | 144400 | 2 | 2 | 2 | N | 20651 | Mitochondrial ribosomal protein L18 OS=Rattus norvegicus OX=10116 GN=Mrpl18 PE=1 SV=1 |
| tr\|D4A0T0 | **69** | 180 | 32.98 | 10 | 10 | 442030 | 1 | 1 | 2 | N | 20859 | NADH:ubiquinone oxidoreductase subunit B10 OS=Rattus norvegicus OX=10116 GN=Ndufb10 PE=1 SV=1 |
| P26284\|OD | **61** | 2826 | 32.88 | 5 | 5 | 193430 | 2 | 2 | 2 | N | 43227 | Pyruvate dehydrogenase E1 component subunit alpha, somatic form, mitochondrial OS=Rattus norvegicus OX=10116 GN=Pdha1 PE=1 SV=2 |
| Q7TQ16\|Q | **84** | 2828 | 29.98 | 10 | 10 | 173740 | 1 | 1 | 1 | N | 9849 | Cytochrome b-c1 complex subunit 8 OS=Rattus norvegicus OX=10116 GN=Uqcrq PE=3 SV=1 |
| tr\|A9UMV | **71** | 2873 | 29.5 | 11 | 11 | 121680 | 1 | 1 | 2 | N | 6539 | RCG29512 OS=Rattus norvegicus OX=10116 GN=Uqcr11 PE=1 SV=1 |
| P11530\|DM | **78** | 333 | 29.47 | 0 | 0 | 100010 | 2 | 2 | 2 | N | 425830 | Dystrophin OS=Rattus norvegicus OX=10116 GN=Dmd PE=1 SV=2 |
| Q5XIN6\|LE | **63** | 2855 | 27.95 | 2 | 2 | 215100 | 1 | 1 | 2 | N | 83060 | Mitochondrial proton/calcium exchanger protein OS=Rattus norvegicus OX=10116 GN=Letm1 PE=1 SV=1 |
| Q4V8F9\|HS | **72** | 121 | 27.91 | 2 | 2 | 149350 | 1 | 1 | 2 | N | 58344 | Hydroxysteroid dehydrogenase-like protein 2 OS=Rattus norvegicus OX=10116 GN=Hsdl2 PE=2 SV=1 |
| Q641Y2\|ND | **64** | 2856 | 27.68 | 4 | 4 | 216330 | 2 | 2 | 2 | N | 52562 | NADH dehydrogenase [ubiquinone] iron-sulfur protein 2, mitochondrial OS=Rattus norvegicus OX=10116 GN=Ndufs2 PE=1 SV=1 |
| tr\|M0R5K3 | **123** | 2876 | 44557 | 10 | 10 | 49381 | 1 | 1 | 1 | N | 9028 | Cytochrome c oxidase subunit 7A1 OS=Rattus norvegicus OX=10116 GN=LOC687508 PE=4 SV=1 |
| Q5M9I5\|Q | **122** | 2875 | 44282 | 12 | 12 | 162270 | 1 | 1 | 1 | N | 10424 | Cytochrome b-c1 complex subunit 6, mitochondrial OS=Rattus norvegicus OX=10116 GN=Uqcrh PE=3 SV=1 |
| tr\|B2RZ24\| | **99** | 2877 | 26.7 | 4 | 4 | 42281 | 1 | 1 | 1 | N | 47388 | Succinate-CoA ligase subunit beta (Fragment) OS=Rattus norvegicus OX=10116 GN=Sucla2 PE=2 SV=1 |
| tr\|F1LM47 | **99** | 2878 | 26.7 | 4 | 4 | 42281 | 1 | 1 | 1 | N | 50306 | Succinate--CoA ligase [ADP-forming] subunit beta, mitochondrial OS=Rattus norvegicus OX=10116 GN=Sucla2 PE=1 SV=1 |
| tr\|D3ZD09 | **73** | 2879 | 25.72 | 12 | 12 | 138060 | 1 | 1 | 2 | N | 10071 | Cytochrome c oxidase subunit OS=Rattus norvegicus OX=10116 GN=Cox6b1 PE=1 SV=1 |
| P07895\|SO | **124** | 2881 | 25.47 | 4 | 4 | 49403 | 1 | 1 | 1 | N | 24674 | Superoxide dismutase [Mn], mitochondrial OS=Rattus norvegicus OX=10116 GN=Sod2 PE=1 SV=2 |
| tr\|Q6P9Y4 | **82** | 1009 | 25.26 | 3 | 3 | 193010 | 1 | 1 | 1 | N | 32904 | ADP/ATP translocase 1 OS=Rattus norvegicus OX=10116 GN=Slc25a4 PE=1 SV=1 |
| Q05962\|AD | **82** | 1010 | 25.26 | 3 | 3 | 193010 | 1 | 1 | 1 | N | 32989 | ADP/ATP translocase 1 OS=Rattus norvegicus OX=10116 GN=Slc25a4 PE=1 SV=3 |
| Q09073\|AD | **82** | 1011 | 25.26 | 3 | 3 | 193010 | 1 | 1 | 1 | N | 32901 | ADP/ATP translocase 2 OS=Rattus norvegicus OX=10116 GN=Slc25a5 PE=1 SV=3 |
| tr\|D3Z900\| | **126** | 2885 | 24.мар | 3 | 3 | 0 | 1 | 1 | 1 | Y | 38239 | Mitochondrial amidoxime reducing component 2 OS=Rattus norvegicus OX=10116 GN=Marc2 PE=1 SV=2 |
| O88994\|M | **126** | 2886 | 24.мар | 3 | 3 | 0 | 1 | 1 | 1 | Y | 38249 | Mitochondrial amidoxime reducing component 2 OS=Rattus norvegicus OX=10116 GN=Marc2 PE=2 SV=1 |
| tr\|M0R6N2 | **126** | 2887 | 24.мар | 3 | 3 | 0 | 1 | 1 | 1 | Y | 38175 | MOSC domain-containing protein 2, mitochondrial-like OS=Rattus norvegicus OX=10116 GN=LOC100910481 PE=1 SV=1 |
