## Supplementary material for "Permeability transition pore-related changes in the proteome and channel activity of ATP synthase dimers and monomers": RHM

| Accession | Protein Group | Protein ID | -10lgP | Coverage (%) | Coverage (%) Sample 2 | IBAQ | #Peptides | #Unique | #Spec Sample 2 | PTM | Avg. Mass | Description |
|---|---|---|---|---|---|---|---|---|---|---|---|---|
| Q66HF1\|N | **1** | 3421 | 242.54 | 28 | 28 | 41174000 | 46 | 46 | 127 | Y | 79412 | NADH-ubiquinone oxidoreductase 75 kDa subunit, mitochondrial OS=Rattus norvegicus OX=10116 GN=Ndufs1 PE=1 SV=1 |
| P19234\|N | **2** | 2870 | 240.09 | 36 | 36 | 47076000 | 32 | 32 | 116 | Y | 27378 | NADH dehydrogenase [ubiquinone] flavoprotein 2, mitochondrial OS=Rattus norvegicus OX=10116 GN=Ndufv2 PE=1 SV=2 |
| Q5BK63\|N | **7** | 5406 | 215.02 | 31 | 31 | 13836000 | 22 | 21 | 70 | N | 42559 | NADH dehydrogenase [ubiquinone] 1 alpha subcomplex subunit 9, mitochondrial OS=Rattus norvegicus OX=10116 GN=Ndufa9 PE=1 SV=2 |
| P32551\|Q | **3** | 34 | 214.98 | 32 | 32 | 41186000 | 38 | 38 | 114 | Y | 48396 | Cytochrome b-c1 complex subunit 2, mitochondrial OS=Rattus norvegicus OX=10116 GN=Uqcrc2 PE=1 SV=2 |
| P10719\|AT | **10** | 1 | 214.82 | 14 | 14 | 13124000 | 17 | 17 | 44 | Y | 56354 | ATP synthase subunit beta, mitochondrial OS=Rattus norvegicus OX=10116 GN=Atp5f1b PE=1 SV=2 |
| tr\|G3V6D3 | **10** | 2 | 214.82 | 14 | 14 | 13124000 | 17 | 17 | 44 | Y | 56345 | ATP synthase subunit beta OS=Rattus norvegicus OX=10116 GN=Atp5f1b PE=1 SV=1 |
| Q68FY0\|Q | **4** | 118 | 203.19 | 29 | 29 | 31011000 | 36 | 36 | 95 | Y | 52849 | Cytochrome b-c1 complex subunit 1, mitochondrial OS=Rattus norvegicus OX=10116 GN=Uqcrc1 PE=1 SV=1 |
| Q641Y2\|N | **5** | 2856 | 199.64 | 22 | 22 | 23036000 | 27 | 27 | 81 | N | 52562 | NADH dehydrogenase [ubiquinone] iron-sulfur protein 2, mitochondrial OS=Rattus norvegicus OX=10116 GN=Ndufs2 PE=1 SV=1 |
| tr\|D3ZFQ8 | **15** | 133 | 195.71 | 23 | 23 | 16067000 | 13 | 13 | 34 | Y | 35435 | Cytochrome c-1 OS=Rattus norvegicus OX=10116 GN=Cyc1 PE=1 SV=3 |
| P02563\|M | **6** | 196 | 195.57 | 15 | 15 | 11436000 | 37 | 37 | 72 | Y | 223506 | Myosin-6 OS=Rattus norvegicus OX=10116 GN=Myh6 PE=1 SV=2 |
| P20788\|U | **18** | 132 | 186.44 | 20 | 20 | 9091700 | 11 | 11 | 32 | N | 29446 | Cytochrome b-c1 complex subunit Rieske, mitochondrial OS=Rattus norvegicus OX=10116 GN=Uqcrfs1 PE=1 SV=2 |
| tr\|B2RYS8\| | **23** | 2857 | 185.82 | 23 | 23 | 8790200 | 12 | 12 | 22 | Y | 21959 | NADH dehydrogenase [ubiquinone] 1 beta subcomplex subunit 8, mitochondrial OS=Rattus norvegicus OX=10116 GN=Ndufb8 PE=1 SV=1 |
| P11240\|C | **13** | 86 | 179.36 | 48 | 48 | 6966500 | 12 | 12 | 38 | N | 16130 | Cytochrome c oxidase subunit 5A, mitochondrial OS=Rattus norvegicus OX=10116 GN=Cox5a PE=1 SV=1 |
| tr\|D3ZF13\| | **14** | 3210 | 178.94 | 28 | 28 | 8608100 | 14 | 14 | 34 | Y | 17514 | Acyl carrier protein OS=Rattus norvegicus OX=10116 GN=Ndufab1 PE=1 SV=1 |
| tr\|Q5PQZ9 | **16** | 2821 | 175.83 | 36 | 36 | 14648000 | 10 | 10 | 33 | N | 14359 | NADH dehydrogenase [ubiquinone] 1 subunit C2 OS=Rattus norvegicus OX=10116 GN=Ndufc2 PE=1 SV=1 |
| tr\|D3ZG43 | **9** | 2807 | 162.53 | 30 | 30 | 17012000 | 21 | 21 | 48 | N | 30226 | NADH dehydrogenase (Ubiquinone) Fe-S protein 3 (Predicted), isoform CRA_c OS=Rattus norvegicus OX=10116 GN=Ndufs3 PE=1 SV=1 |
| Q80W89\| | **22** | 5440 | 161.18 | 21 | 21 | 6703000 | 8 | 8 | 23 | N | 14854 | NADH dehydrogenase [ubiquinone] 1 alpha subcomplex subunit 11 OS=Rattus norvegicus OX=10116 GN=Ndufa11 PE=2 SV=1 |
| tr\|B0BNE6 | **20** | 5411 | 159.58 | 17 | 17 | 4028900 | 9 | 9 | 25 | N | 23970 | NADH dehydrogenase (Ubiquinone) Fe-S protein 8 (Predicted), isoform CRA_a OS=Rattus norvegicus OX=10116 GN=Ndufs8 PE=1 SV=1 |
| Q561S0\|N | **12** | 5410 | 152.71 | 35 | 35 | 7850800 | 20 | 20 | 42 | Y | 40493 | NADH dehydrogenase [ubiquinone] 1 alpha subcomplex subunit 10, mitochondrial OS=Rattus norvegicus OX=10116 GN=Ndufa10 PE=1 SV=1 |
| tr\|A0A411 | **17** | 39 | 151.02 | 10 | 10 | 5777900 | 11 | 11 | 33 | N | 40144 | Cytochrome b (Fragment) OS=Rattus norvegicus OX=10116 GN=Cytb PE=3 SV=1 |
| tr\|A0A0U2 | **17** | 40 | 151.02 | 10 | 10 | 5777900 | 11 | 11 | 33 | N | 40981 | Cytochrome b (Fragment) OS=Rattus norvegicus OX=10116 PE=3 SV=1 |
| tr\|A0A0U2 | **17** | 41 | 151.02 | 10 | 10 | 5777900 | 11 | 11 | 33 | N | 41037 | Cytochrome b (Fragment) OS=Rattus norvegicus OX=10116 PE=3 SV=1 |
| tr\|A0A411 | **17** | 42 | 151.02 | 10 | 10 | 5777900 | 11 | 11 | 33 | N | 41017 | Cytochrome b (Fragment) OS=Rattus norvegicus OX=10116 GN=Cytb PE=3 SV=1 |
| tr\|F8QU31 | **17** | 43 | 151.02 | 9 | 9 | 5777900 | 11 | 11 | 33 | N | 42181 | Cytochrome b (Fragment) OS=Rattus norvegicus OX=10116 GN=cytb PE=3 SV=1 |
| tr\|A0A411 | **17** | 44 | 151.02 | 9 | 9 | 5777900 | 11 | 11 | 33 | N | 42238 | Cytochrome b (Fragment) OS=Rattus norvegicus OX=10116 GN=Cytb PE=3 SV=1 |
| tr\|L0L4L8\| | **17** | 45 | 151.02 | 9 | 9 | 5777900 | 11 | 11 | 33 | N | 42470 | Cytochrome b (Fragment) OS=Rattus norvegicus OX=10116 GN=cytb PE=3 SV=1 |
| tr\|A0A140 | **17** | 46 | 151.02 | 9 | 9 | 5777900 | 11 | 11 | 33 | N | 42574 | Cytochrome b (Fragment) OS=Rattus norvegicus OX=10116 PE=3 SV=1 |
| tr\|A0A140 | **17** | 47 | 151.02 | 9 | 9 | 5777900 | 11 | 11 | 33 | N | 42588 | Cytochrome b (Fragment) OS=Rattus norvegicus OX=10116 PE=3 SV=1 |
| tr\|A0A140 | **17** | 48 | 151.02 | 9 | 9 | 5777900 | 11 | 11 | 33 | N | 42527 | Cytochrome b (Fragment) OS=Rattus norvegicus OX=10116 PE=3 SV=1 |
| tr\|A0A385 | **17** | 49 | 151.02 | 9 | 9 | 5777900 | 11 | 11 | 33 | N | 42766 | Cytochrome b (Fragment) OS=Rattus norvegicus OX=10116 PE=3 SV=1 |
| tr\|A0A3Q8 | **17** | 50 | 151.02 | 9 | 9 | 5777900 | 11 | 11 | 33 | N | 42712 | Cytochrome b (Fragment) OS=Rattus norvegicus OX=10116 GN=Cytb PE=3 SV=1 |
| tr\|A0A3Q8 | **17** | 38 | 151.02 | 9 | 9 | 5777900 | 11 | 11 | 33 | N | 42622 | Cytochrome b (Fragment) OS=Rattus norvegicus OX=10116 GN=Cytb PE=3 SV=1 |
| tr\|A0A3S6 | **17** | 51 | 151.02 | 9 | 9 | 5777900 | 11 | 11 | 33 | N | 42702 | Cytochrome b (Fragment) OS=Rattus norvegicus OX=10116 GN=Cytb PE=3 SV=1 |
| tr\|A0A3S6 | **17** | 52 | 151.02 | 9 | 9 | 5777900 | 11 | 11 | 33 | N | 42698 | Cytochrome b (Fragment) OS=Rattus norvegicus OX=10116 GN=Cytb PE=3 SV=1 |
| tr\|A0A097 | **17** | 53 | 151.02 | 9 | 9 | 5777900 | 11 | 11 | 33 | N | 42865 | Cytochrome b OS=Rattus norvegicus OX=10116 GN=CYTB PE=3 SV=1 |
| tr\|A0A220 | **17** | 54 | 151.02 | 9 | 9 | 5777900 | 11 | 11 | 33 | N | 43018 | Cytochrome b OS=Rattus norvegicus OX=10116 PE=3 SV=1 |
| tr\|D6NSP2 | **17** | 55 | 151.02 | 9 | 9 | 5777900 | 11 | 11 | 33 | N | 42945 | Cytochrome b (Fragment) OS=Rattus norvegicus OX=10116 GN=cytb PE=3 SV=1 |
| tr\|D6NSS6 | **17** | 56 | 151.02 | 9 | 9 | 5777900 | 11 | 11 | 33 | N | 42993 | Cytochrome b (Fragment) OS=Rattus norvegicus OX=10116 GN=cytb PE=3 SV=1 |
| tr\|D6NSQ3 | **17** | 57 | 151.02 | 9 | 9 | 5777900 | 11 | 11 | 33 | N | 43012 | Cytochrome b (Fragment) OS=Rattus norvegicus OX=10116 GN=cytb PE=3 SV=1 |
| tr\|D6NSR3 | **17** | 58 | 151.02 | 9 | 9 | 5777900 | 11 | 11 | 33 | N | 42986 | Cytochrome b (Fragment) OS=Rattus norvegicus OX=10116 GN=cytb PE=3 SV=1 |
| tr\|D6NSP4 | **17** | 59 | 151.02 | 9 | 9 | 5777900 | 11 | 11 | 33 | N | 43016 | Cytochrome b (Fragment) OS=Rattus norvegicus OX=10116 GN=cytb PE=3 SV=1 |
| tr\|D6NSQ8 | **17** | 60 | 151.02 | 9 | 9 | 5777900 | 11 | 11 | 33 | N | 43016 | Cytochrome b (Fragment) OS=Rattus norvegicus OX=10116 GN=cytb PE=3 SV=1 |
| tr\|D6NSR8 | **17** | 61 | 151.02 | 9 | 9 | 5777900 | 11 | 11 | 33 | N | 42998 | Cytochrome b (Fragment) OS=Rattus norvegicus OX=10116 GN=cytb PE=3 SV=1 |
| tr\|F8QU37 | **17** | 62 | 151.02 | 9 | 9 | 5777900 | 11 | 11 | 33 | N | 42992 | Cytochrome b (Fragment) OS=Rattus norvegicus OX=10116 GN=cytb PE=3 SV=1 |
| tr\|F2Q6S6\| | **17** | 63 | 151.02 | 9 | 9 | 5777900 | 11 | 11 | 33 | N | 42938 | Cytochrome b (Fragment) OS=Rattus norvegicus OX=10116 GN=cytb PE=3 SV=1 |
| tr\|A0A0A1 | **17** | 64 | 151.02 | 9 | 9 | 5777900 | 11 | 11 | 33 | N | 42989 | Cytochrome b OS=Rattus norvegicus OX=10116 GN=CYTB PE=3 SV=1 |
| tr\|D6NSS7 | **17** | 65 | 151.02 | 9 | 9 | 5777900 | 11 | 11 | 33 | N | 42948 | Cytochrome b (Fragment) OS=Rattus norvegicus OX=10116 GN=cytb PE=3 SV=1 |
| tr\|Q8SEY9\| | **17** | 66 | 151.02 | 9 | 9 | 5777900 | 11 | 11 | 33 | N | 43015 | Cytochrome b OS=Rattus norvegicus OX=10116 GN=cytb PE=3 SV=1 |
| tr\|D6NSR7 | **17** | 67 | 151.02 | 9 | 9 | 5777900 | 11 | 11 | 33 | N | 42982 | Cytochrome b (Fragment) OS=Rattus norvegicus OX=10116 GN=cytb PE=3 SV=1 |
| tr\|A0A220 | **17** | 68 | 151.02 | 9 | 9 | 5777900 | 11 | 11 | 33 | N | 42968 | Cytochrome b OS=Rattus norvegicus OX=10116 PE=3 SV=1 |
| tr\|F2Q6S5\| | **17** | 69 | 151.02 | 9 | 9 | 5777900 | 11 | 11 | 33 | N | 42952 | Cytochrome b (Fragment) OS=Rattus norvegicus OX=10116 GN=cytb PE=3 SV=1 |
| tr\|D6NSQ6 | **17** | 70 | 151.02 | 9 | 9 | 5777900 | 11 | 11 | 33 | N | 42952 | Cytochrome b (Fragment) OS=Rattus norvegicus OX=10116 GN=cytb PE=3 SV=1 |
| tr\|D6NSQ0 | **17** | 71 | 151.02 | 9 | 9 | 5777900 | 11 | 11 | 33 | N | 42968 | Cytochrome b (Fragment) OS=Rattus norvegicus OX=10116 GN=cytb PE=3 SV=1 |
| tr\|Q5UAI7 | **17** | 72 | 151.02 | 9 | 9 | 5777900 | 11 | 11 | 33 | N | 42998 | Cytochrome b OS=Rattus norvegicus OX=10116 GN=CYTB PE=3 SV=1 |
| tr\|D6NSR2 | **17** | 73 | 151.02 | 9 | 9 | 5777900 | 11 | 11 | 33 | N | 43073 | Cytochrome b (Fragment) OS=Rattus norvegicus OX=10116 GN=cytb PE=3 SV=1 |
| tr\|A0A0S1 | **17** | 74 | 151.02 | 9 | 9 | 5777900 | 11 | 11 | 33 | N | 43002 | Cytochrome b OS=Rattus norvegicus OX=10116 GN=CYTB PE=3 SV=1 |
| tr\|D6NSR1 | **17** | 75 | 151.02 | 9 | 9 | 5777900 | 11 | 11 | 33 | N | 43002 | Cytochrome b (Fragment) OS=Rattus norvegicus OX=10116 GN=cytb PE=3 SV=1 |

| | | | | | | | | | | | |
|---|---|---|---|---|---|---|---|---|---|---|---|
| tr\|Q8HIC4\| | **17** | 76 | 151.02 | 9 | 9 | 5777900 | 11 | 11 | 33 | N | 43012 Cytochrome b OS=Rattus norvegicus OX=10116 GN=Mt-cyb PE=3 SV=1 |
| tr\|A0A220I | **17** | 77 | 151.02 | 9 | 9 | 5777900 | 11 | 11 | 33 | N | 42968 Cytochrome b OS=Rattus norvegicus OX=10116 PE=3 SV=1 |
| tr\|A0A0S1Z | **17** | 78 | 151.02 | 9 | 9 | 5777900 | 11 | 11 | 33 | N | 43016 Cytochrome b OS=Rattus norvegicus OX=10116 GN=CYTB PE=3 SV=1 |
| tr\|R9TKN1 | **17** | 119 | 151.02 | 9 | 9 | 5777900 | 11 | 11 | 33 | N | 43012 Cytochrome b OS=Rattus norvegicus OX=10116 GN=CYTB PE=3 SV=1 |
| P00159\|CY | **17** | 79 | 151.02 | 9 | 9 | 5777900 | 11 | 11 | 33 | N | 43012 Cytochrome b OS=Rattus norvegicus OX=10116 GN=Mt-Cyb PE=3 SV=3 |
| tr\|L0N311\| | **17** | 80 | 151.02 | 9 | 9 | 5777900 | 11 | 11 | 33 | N | 43045 Cytochrome b (Fragment) OS=Rattus norvegicus OX=10116 GN=cytb PE=3 SV=1 |
| tr\|A0A220I | **17** | 81 | 151.02 | 9 | 9 | 5777900 | 11 | 11 | 33 | N | 42970 Cytochrome b OS=Rattus norvegicus OX=10116 PE=3 SV=1 |
| tr\|H2KXA0 | **17** | 82 | 151.02 | 9 | 9 | 5777900 | 11 | 11 | 33 | N | 42978 Cytochrome b (Fragment) OS=Rattus norvegicus OX=10116 GN=cytb PE=3 SV=1 |
| tr\|A0A096X | **17** | 83 | 151.02 | 9 | 9 | 5777900 | 11 | 11 | 33 | N | 43042 Cytochrome b (Fragment) OS=Rattus norvegicus OX=10116 PE=3 SV=1 |
| tr\|D6NSP5 | **17** | 120 | 151.02 | 9 | 9 | 5777900 | 11 | 11 | 33 | N | 43012 Cytochrome b (Fragment) OS=Rattus norvegicus OX=10116 GN=cytb PE=3 SV=1 |
| P15999\|AT | **8** | 3 | 148.67 | 20 | 20 | 8088800 | 19 | 19 | 48 | N | 59754 ATP synthase subunit alpha, mitochondrial OS=Rattus norvegicus OX=10116 GN=Atp5f1a PE=1 SV=2 |
| tr\|F1LP05\| | **8** | 4 | 148.67 | 20 | 20 | 8088800 | 19 | 19 | 48 | N | 59813 ATP synthase subunit alpha OS=Rattus norvegicus OX=10116 GN=Atp5f1a PE=1 SV=1 |
| Q6AXV4\|SA | **32** | 2803 | 148.56 | 14 | 14 | 2832600 | 8 | 8 | 15 | N | 51960 Sorting and assembly machinery component 50 homolog OS=Rattus norvegicus OX=10116 GN=Samm50 PE=1 SV=1 |
| tr\|Q5XIH3\| | **11** | 5408 | 141.88 | 12 | 12 | 9537800 | 12 | 12 | 43 | N | 50731 NADH dehydrogenase [ubiquinone] flavoprotein 1, mitochondrial OS=Rattus norvegicus OX=10116 GN=Ndufv1 PE=1 SV=1 |
| P67779\|PH | **21** | 5407 | 141.46 | 28 | 28 | 3269800 | 12 | 12 | 24 | N | 29820 Prohibitin OS=Rattus norvegicus OX=10116 GN=Phb PE=1 SV=1 |
| tr\|A0A0G2 | **25** | 147 | 141.36 | 11 | 11 | 3368600 | 12 | 12 | 19 | N | 86230 MICOS complex subunit MIC60 OS=Rattus norvegicus OX=10116 GN=Immt PE=1 SV=1 |
| tr\|A0A140T | **25** | 145 | 141.36 | 14 | 14 | 3368600 | 12 | 12 | 19 | N | 67049 MICOS complex subunit MIC60 OS=Rattus norvegicus OX=10116 GN=Immt PE=1 SV=1 |
| Q3KR86\|M | **25** | 146 | 141.36 | 14 | 14 | 3368600 | 12 | 12 | 19 | N | 67177 MICOS complex subunit Mic60 (Fragment) OS=Rattus norvegicus OX=10116 GN=Immt PE=1 SV=1 |
| P00507\|AA | **30** | 19 | 139.71 | 11 | 11 | 2877000 | 8 | 8 | 16 | Y | 47314 Aspartate aminotransferase, mitochondrial OS=Rattus norvegicus OX=10116 GN=Got2 PE=1 SV=2 |
| tr\|D4A7L4\| | **28** | 2872 | 139.22 | 26 | 26 | 8714300 | 6 | 6 | 18 | Y | 17634 NADH dehydrogenase (Ubiquinone) 1 beta subcomplex, 11 (Predicted) OS=Rattus norvegicus OX=10116 GN=Ndufb11 PE=1 SV=1 |
| Q02253\|M | **33** | 2801 | 131.03 | 17 | 17 | 1652900 | 9 | 9 | 14 | Y | 57808 Methylmalonate-semialdehyde dehydrogenase [acylating], mitochondrial OS=Rattus norvegicus OX=10116 GN=Aldh6a1 PE=1 SV=1 |
| tr\|G3V7J0\| | **33** | 2802 | 131.03 | 17 | 17 | 1652900 | 9 | 9 | 14 | Y | 57748 Aldehyde dehydrogenase family 6, subfamily A1, isoform CRA_b OS=Rattus norvegicus OX=10116 GN=Aldh6a1 PE=1 SV=1 |
| tr\|F1LXA0\| | **40** | 5458 | 125.66 | 17 | 17 | 2298700 | 4 | 4 | 8 | N | 17178 NADH dehydrogenase [ubiquinone] 1 alpha subcomplex subunit 12 OS=Rattus norvegicus OX=10116 GN=Ndufa12 PE=1 SV=2 |
| tr\|A0A0G2 | **31** | 3028 | 123.31 | 33 | 33 | 4706400 | 7 | 7 | 15 | Y | 19965 NADH dehydrogenase [ubiquinone] 1 alpha subcomplex subunit 8 OS=Rattus norvegicus OX=10116 GN=Ndufa8 PE=1 SV=1 |
| tr\|D4A565 | **27** | 5412 | 122.44 | 30 | 30 | 3205900 | 10 | 10 | 18 | N | 21664 NADH dehydrogenase (Ubiquinone) 1 beta subcomplex, 5 (Predicted), isoform CRA_b OS=Rattus norvegicus OX=10116 GN=Ndufb5 PE=1 SV=1 |
| tr\|D4A0T0 | **24** | 180 | 120.28 | 26 | 26 | 10576000 | 9 | 9 | 20 | N | 20859 NADH:ubiquinone oxidoreductase subunit B10 OS=Rattus norvegicus OX=10116 GN=Ndufb10 PE=1 SV=1 |
| tr\|Q5UAJ6 | **19** | 24 | 116.69 | 32 | 32 | 6738900 | 14 | 14 | 29 | Y | 25942 Cytochrome c oxidase subunit 2 OS=Rattus norvegicus OX=10116 GN=COX2 PE=3 SV=1 |
| Q5M9I5\|Q | **26** | 2875 | 112.06 | 18 | 18 | 4097800 | 5 | 5 | 19 | N | 10424 Cytochrome b-c1 complex subunit 6, mitochondrial OS=Rattus norvegicus OX=10116 GN=Uqcrh PE=3 SV=1 |
| tr\|A0A0G2 | **29** | 5422 | 111.64 | 24 | 24 | 5076400 | 7 | 7 | 16 | Y | 33168 Prohibitin OS=Rattus norvegicus OX=10116 GN=Phb2 PE=1 SV=1 |
| Q5XIH7\|PH | **29** | 5423 | 111.64 | 24 | 24 | 5076400 | 7 | 7 | 16 | Y | 33312 Prohibitin-2 OS=Rattus norvegicus OX=10116 GN=Phb2 PE=1 SV=1 |
| tr\|D3ZS58\| | **34** | 5449 | 106.35 | 23 | 23 | 2032400 | 4 | 4 | 13 | N | 10845 NADH dehydrogenase [ubiquinone] 1 alpha subcomplex subunit 2 OS=Rattus norvegicus OX=10116 GN=Ndufa2 PE=1 SV=1 |
| tr\|D4A3V2 | **38** | 5424 | 100.01 | 22 | 22 | 2676700 | 4 | 4 | 9 | N | 15224 NADH dehydrogenase [ubiquinone] 1 alpha subcomplex subunit 6 OS=Rattus norvegicus OX=10116 GN=Ndufa6 PE=1 SV=1 |
| tr\|Q06QK5 | **36** | 5413 | 97.29 | 5 | 5 | 1684200 | 6 | 6 | 11 | Y | 68606 NADH-ubiquinone oxidoreductase chain 5 OS=Rattus norvegicus OX=10116 GN=ND5 PE=3 SV=1 |
| P11661\|NU | **36** | 5414 | 97.29 | 5 | 5 | 1684200 | 6 | 6 | 11 | Y | 68618 NADH-ubiquinone oxidoreductase chain 5 OS=Rattus norvegicus OX=10116 GN=Mtnd5 PE=3 SV=3 |
| tr\|Q06QG6 | **36** | 5415 | 97.29 | 5 | 5 | 1684200 | 6 | 6 | 11 | Y | 68588 NADH-ubiquinone oxidoreductase chain 5 OS=Rattus norvegicus OX=10116 GN=ND5 PE=3 SV=1 |
| tr\|Q06QA1 | **36** | 5416 | 97.29 | 5 | 5 | 1684200 | 6 | 6 | 11 | Y | 68574 NADH-ubiquinone oxidoreductase chain 5 OS=Rattus norvegicus OX=10116 GN=ND5 PE=3 SV=1 |
| tr\|Q8SEZ0\| | **36** | 5417 | 97.29 | 5 | 5 | 1684200 | 6 | 6 | 11 | Y | 68618 NADH-ubiquinone oxidoreductase chain 5 OS=Rattus norvegicus OX=10116 GN=Mt-nd5 PE=3 SV=1 |
| tr\|A0A096X | **36** | 5418 | 97.29 | 5 | 5 | 1684200 | 6 | 6 | 11 | Y | 68584 NADH-ubiquinone oxidoreductase chain 5 OS=Rattus norvegicus OX=10116 GN=ND5 PE=3 SV=1 |
| tr\|B5DEL8\| | **44** | 5442 | 96.29 | 23 | 23 | 2524100 | 4 | 4 | 7 | N | 12700 NADH dehydrogenase (Ubiquinone) Fe-S protein 5 OS=Rattus norvegicus OX=10116 GN=Ndufs5 PE=1 SV=1 |
| tr\|A0A0G2 | **44** | 5443 | 96.29 | 23 | 23 | 2524100 | 4 | 4 | 7 | N | 12730 Uncharacterized protein OS=Rattus norvegicus OX=10116 PE=4 SV=1 |
| P31399\|AT | **59** | 7 | 94.75 | 20 | 20 | 570370 | 3 | 3 | 4 | N | 18763 ATP synthase subunit d, mitochondrial OS=Rattus norvegicus OX=10116 GN=Atp5pd PE=1 SV=3 |
| Q63362\|NI | **35** | 5439 | 94 | 28 | 28 | 1728200 | 4 | 4 | 11 | N | 13412 NADH dehydrogenase [ubiquinone] 1 alpha subcomplex subunit 5 OS=Rattus norvegicus OX=10116 GN=Ndufa5 PE=1 SV=3 |
| P35435\|AT | **54** | 5 | 92.58 | 8 | 8 | 374190 | 3 | 3 | 5 | N | 30191 ATP synthase subunit gamma, mitochondrial OS=Rattus norvegicus OX=10116 GN=Atp5f1c PE=1 SV=2 |
| tr\|Q6QI09\| | **54** | 6 | 92.58 | 4 | 4 | 374190 | 3 | 3 | 5 | N | 67721 ATP synthase subunit gamma, mitochondrial OS=Rattus norvegicus OX=10116 GN=Taf3 PE=1 SV=1 |
| P10817\|CX | **41** | 151 | 91.57 | 17 | 17 | 1335400 | 5 | 5 | 8 | N | 10487 Cytochrome c oxidase subunit 6A2, mitochondrial (Fragment) OS=Rattus norvegicus OX=10116 GN=Cox6a2 PE=1 SV=3 |
| tr\|G3V8M4 | **41** | 152 | 91.57 | 16 | 16 | 1335400 | 5 | 5 | 8 | N | 10802 Cytochrome c oxidase subunit 6A, mitochondrial OS=Rattus norvegicus OX=10116 GN=Cox6a2 PE=3 SV=1 |
| tr\|D3ZZ21\| | **42** | 5448 | 90.48 | 34 | 34 | 1585200 | 6 | 6 | 7 | Y | 15638 NADH dehydrogenase (Ubiquinone) 1 beta subcomplex, 6 (Predicted) OS=Rattus norvegicus OX=10116 GN=Ndufb6 PE=1 SV=1 |
| Q60587\|EC | **43** | 20 | 89.86 | 9 | 9 | 676180 | 5 | 5 | 7 | N | 51414 Trifunctional enzyme subunit beta, mitochondrial OS=Rattus norvegicus OX=10116 GN=Hadhb PE=1 SV=1 |
| tr\|A0A0G2 | **43** | 21 | 89.86 | 9 | 9 | 676180 | 5 | 5 | 7 | N | 52568 Trifunctional enzyme subunit beta, mitochondrial OS=Rattus norvegicus OX=10116 GN=Hadhb PE=1 SV=1 |
| P10888\|CC | **37** | 290 | 82.08 | 15 | 15 | 1871200 | 5 | 5 | 11 | Y | 19515 Cytochrome c oxidase subunit 4 isoform 1, mitochondrial OS=Rattus norvegicus OX=10116 GN=Cox4i1 PE=1 SV=1 |
| tr\|F1LPG5\| | **69** | 5445 | 81.82 | 21 | 21 | 354200 | 2 | 2 | 3 | N | 15064 NADH:ubiquinone oxidoreductase subunit B4 OS=Rattus norvegicus OX=10116 GN=Ndufb4 PE=1 SV=1 |
| tr\|A9UMW | **52** | 5453 | 79.81 | 33 | 33 | 3890800 | 4 | 4 | 5 | Y | 9255 Ndufa3 protein (Fragment) OS=Rattus norvegicus OX=10116 GN=Ndufa3 PE=2 SV=1 |
| tr\|M0RB63 | **52** | 5454 | 79.81 | 32 | 32 | 3890800 | 4 | 4 | 5 | Y | 9372 RCG63041 OS=Rattus norvegicus OX=10116 GN=LOC684509 PE=4 SV=1 |
| tr\|A0A0G2 | **52** | 5455 | 79.81 | 30 | 30 | 3890800 | 4 | 4 | 5 | Y | 10170 NADH:ubiquinone oxidoreductase subunit A3 OS=Rattus norvegicus OX=10116 GN=Ndufa3 PE=1 SV=1 |
| tr\|D3ZE15\| | **45** | 5447 | 78.17 | 33 | 33 | 1235800 | 4 | 4 | 7 | Y | 16777 NADH:ubiquinone oxidoreductase subunit A13 OS=Rattus norvegicus OX=10116 GN=Ndufa13 PE=1 SV=1 |
| tr\|C8CHS6 | **61** | 15533 | 74.94 | 26 | 26 | 203820 | 2 | 2 | 4 | N | 7988 Mitochondrial superoxide dismutase 2 (Fragment) OS=Rattus norvegicus OX=10116 PE=2 SV=1 |
| P07895\|SO | **61** | 2881 | 74.94 | 9 | 9 | 203820 | 2 | 2 | 4 | N | 24674 Superoxide dismutase [Mn], mitochondrial OS=Rattus norvegicus OX=10116 GN=Sod2 PE=1 SV=2 |
| tr\|D3ZUX5 | **79** | 2815 | 74.57 | 9 | 9 | 1113500 | 2 | 2 | 2 | N | 26435 MICOS complex subunit OS=Rattus norvegicus OX=10116 GN=Chchd3 PE=1 SV=1 |
| P17764\|TH | **47** | 2787 | 71.56 | 8 | 8 | 1087200 | 4 | 4 | 6 | N | 44695 Acetyl-CoA acetyltransferase, mitochondrial OS=Rattus norvegicus OX=10116 GN=Acat1 PE=1 SV=1 |
| tr\|F1LN92\| | **39** | 7847 | 70.65 | 5 | 5 | 1056000 | 4 | 4 | 8 | N | 89352 AFG3-like matrix AAA peptidase subunit 2 OS=Rattus norvegicus OX=10116 GN=Afg3l2 PE=1 SV=1 |

| | | | | | | | | | | | |
|---|---|---|---|---|---|---|---|---|---|---|---|
| tr\|Q5RJN0 | **53** | 564 | 69.96 | 10 | 10 | 1507800 | 3 | 3 | 5 | N | 23945 NADH dehydrogenase (Ubiquinone) Fe-S protein 7 OS=Rattus norvegicus OX=10116 GN=Ndufs7 PE=1 SV=1 |
| Q64428\|EC | **46** | 9 | 68.1 | 4 | 4 | 745210 | 3 | 3 | 7 | N | 82665 Trifunctional enzyme subunit alpha, mitochondrial OS=Rattus norvegicus OX=10116 GN=Hadha PE=1 SV=2 |
| Q5XIF3\|ND | **57** | 5457 | 66.68 | 11 | 11 | 540880 | 2 | 2 | 4 | N | 19741 NADH dehydrogenase [ubiquinone] iron-sulfur protein 4, mitochondrial OS=Rattus norvegicus OX=10116 GN=Ndufs4 PE=1 SV=1 |
| P11951\|CX | **68** | 2816 | 66.38 | 36 | 36 | 432780 | 3 | 3 | 3 | Y | 8455 Cytochrome c oxidase subunit 6C-2 OS=Rattus norvegicus OX=10116 GN=Cox6c2 PE=1 SV=3 |
| Q9ER34\|AC | **76** | 88 | 63.98 | 4 | 4 | 436150 | 2 | 2 | 2 | N | 85433 Aconitate hydratase, mitochondrial OS=Rattus norvegicus OX=10116 GN=Aco2 PE=1 SV=2 |
| tr\|Q35733 | **62** | 5460 | 61.95 | 13 | 13 | 355130 | 3 | 3 | 4 | N | 8389 NADH-ubiquinone oxidoreductase chain 6 (Fragment) OS=Rattus norvegicus OX=10116 PE=3 SV=1 |
| tr\|Q06QD9 | **62** | 5461 | 61.95 | 6 | 6 | 355130 | 3 | 3 | 4 | N | 18943 NADH-ubiquinone oxidoreductase chain 6 OS=Rattus norvegicus OX=10116 GN=ND6 PE=3 SV=1 |
| tr\|Q7HKW | **62** | 5462 | 61.95 | 6 | 6 | 355130 | 3 | 3 | 4 | N | 18957 NADH-ubiquinone oxidoreductase chain 6 OS=Rattus norvegicus OX=10116 GN=NADH6 PE=3 SV=1 |
| P02770\|AL | **71** | 2832 | 58.72 | 2 | 2 | 447390 | 2 | 2 | 3 | Y | 68731 Serum albumin OS=Rattus norvegicus OX=10116 GN=Alb PE=1 SV=2 |
| tr\|A0A0G2 | **71** | 2833 | 58.72 | 2 | 2 | 447390 | 2 | 2 | 3 | Y | 68759 Serum albumin OS=Rattus norvegicus OX=10116 GN=Alb PE=1 SV=1 |
| P80432\|CC | **58** | 134 | 57.84 | 21 | 21 | 653930 | 2 | 2 | 4 | N | 7375 Cytochrome c oxidase subunit 7C, mitochondrial OS=Rattus norvegicus OX=10116 GN=Cox7c PE=1 SV=2 |
| Q01205\|OI | **73** | 13095 | 54.67 | 9 | 9 | 1205300 | 2 | 2 | 2 | Y | 48925 Dihydrolipoyllysine-residue succinyltransferase component of 2-oxoglutarate dehydrogenase complex, mitochondrial OS=Rattus norvegicus OX=10116 GN=Dlst PE=1 SV=2 |
| tr\|G3V6P2 | **73** | 13096 | 54.67 | 9 | 9 | 1205300 | 2 | 2 | 2 | Y | 48899 Dihydrolipoamide S-succinyltransferase (E2 component of 2-oxo-glutarate complex), isoform CRA_a OS=Rattus norvegicus OX=10116 GN=Dlst PE=1 SV=1 |
| tr\|A9UMV9 | **50** | 5450 | 53.72 | 20 | 20 | 3607100 | 3 | 3 | 6 | N | 12500 NADH:ubiquinone oxidoreductase subunit A7 OS=Rattus norvegicus OX=10116 GN=Ndufa7 PE=1 SV=1 |
| P19511\|AT | **49** | 10 | 51.52 | 9 | 9 | 1827600 | 2 | 2 | 6 | N | 28869 ATP synthase F(0) complex subunit B1, mitochondrial OS=Rattus norvegicus OX=10116 GN=Atp5pb PE=1 SV=1 |
| P12075\|CC | **48** | 139 | 51.26 | 11 | 11 | 4935500 | 3 | 3 | 6 | N | 13915 Cytochrome c oxidase subunit 5B, mitochondrial OS=Rattus norvegicus OX=10116 GN=Cox5b PE=1 SV=2 |
| tr\|D3ZCZ9\| | **70** | 5435 | 50.54 | 21 | 21 | 954580 | 2 | 2 | 3 | N | 13040 NADH dehydrogenase [ubiquinone] iron-sulfur protein 6, mitochondrial OS=Rattus norvegicus OX=10116 GN=LOC100912599 PE=1 SV=1 |
| P05508\|NU | **66** | 5425 | 46.79 | 4 | 4 | 481580 | 2 | 2 | 3 | N | 51783 NADH-ubiquinone oxidoreductase chain 4 OS=Rattus norvegicus OX=10116 GN=Mtnd4 PE=3 SV=3 |
| tr\|D2E6K0\| | **66** | 5426 | 46.79 | 4 | 4 | 481580 | 2 | 2 | 3 | N | 51833 NADH-ubiquinone oxidoreductase chain 4 OS=Rattus norvegicus OX=10116 GN=ND4 PE=3 SV=1 |
| tr\|A7XYB9\| | **66** | 5427 | 46.79 | 4 | 4 | 481580 | 2 | 2 | 3 | N | 51773 NADH-ubiquinone oxidoreductase chain 4 OS=Rattus norvegicus OX=10116 GN=Nd4 PE=3 SV=1 |
