## Supplementary material for "Permeability transition pore-related changes in the proteome and channel activity of ATP synthase dimers and monomers": RHM

| Accession | rotein Grou | Protein ID | -10lgP | overage (% | age (%) Sa | IBAQ | #Peptides | #Unique | pec Sample | PTM | Avg. Mass | Description |
|---|---|---|---|---|---|---|---|---|---|---|---|---|
| Q5BK63\|N | **7** | 5406 | 262.99 | 38 | 38 | 29784000 | 35 | 34 | 103 | Y | 42559 | NADH dehydrogenase [ubiquinone] 1 alpha subcomplex subunit 9, mitochondrial OS=Rattus norvegicus OX=10116 GN=Ndufa9 PE=1 SV=2 |
| Q68FY0\|Q | **1** | 118 | 262.68 | 38 | 38 | 77415000 | 61 | 61 | 202 | Y | 52849 | Cytochrome b-c1 complex subunit 1, mitochondrial OS=Rattus norvegicus OX=10116 GN=Uqcrc1 PE=1 SV=1 |
| Q66HF1\|N | **2** | 3421 | 253.83 | 38 | 38 | 83852000 | 67 | 66 | 174 | Y | 79412 | NADH-ubiquinone oxidoreductase 75 kDa subunit, mitochondrial OS=Rattus norvegicus OX=10116 GN=Ndufs1 PE=1 SV=1 |
| P19234\|ND | **4** | 2870 | 242.81 | 46 | 46 | 88531000 | 43 | 43 | 126 | Y | 27378 | NADH dehydrogenase [ubiquinone] flavoprotein 2, mitochondrial OS=Rattus norvegicus OX=10116 GN=Ndufv2 PE=1 SV=2 |
| P32551\|Q | **3** | 34 | 241.32 | 35 | 35 | 80073000 | 56 | 56 | 162 | Y | 48396 | Cytochrome b-c1 complex subunit 2, mitochondrial OS=Rattus norvegicus OX=10116 GN=Uqcrc2 PE=1 SV=2 |
| tr\|B2RYS8\| | **15** | 2857 | 238.52 | 37 | 37 | 25612000 | 18 | 18 | 49 | Y | 21959 | NADH dehydrogenase [ubiquinone] 1 beta subcomplex subunit 8, mitochondrial OS=Rattus norvegicus OX=10116 GN=Ndufb8 PE=1 SV=1 |
| tr\|D3ZFQ8 | **13** | 133 | 231.04 | 30 | 30 | 33694000 | 21 | 21 | 53 | Y | 35435 | Cytochrome c-1 OS=Rattus norvegicus OX=10116 GN=Cyc1 PE=1 SV=3 |
| P20788\|U | **12** | 132 | 227.85 | 35 | 35 | 29898000 | 20 | 20 | 59 | N | 29446 | Cytochrome b-c1 complex subunit Rieske, mitochondrial OS=Rattus norvegicus OX=10116 GN=Uqcrfs1 PE=1 SV=2 |
| Q641Y2\|N | **6** | 2856 | 212.21 | 31 | 31 | 53720000 | 37 | 36 | 110 | N | 52562 | NADH dehydrogenase [ubiquinone] iron-sulfur protein 2, mitochondrial OS=Rattus norvegicus OX=10116 GN=Ndufs2 PE=1 SV=1 |
| tr\|D3ZG43 | **5** | 2807 | 210.03 | 42 | 42 | 38871000 | 32 | 32 | 111 | N | 30226 | NADH dehydrogenase (Ubiquinone) Fe-S protein 3 (Predicted), isoform CRA_c OS=Rattus norvegicus OX=10116 GN=Ndufs3 PE=1 SV=1 |
| tr\|A0A097 | **8** | 53 | 202.18 | 18 | 18 | 3485500 | 22 | 1 | 77 | N | 42865 | Cytochrome b OS=Rattus norvegicus OX=10116 GN=CYTB PE=3 SV=1 |
| tr\|A0A411 | **8** | 39 | 202.18 | 19 | 19 | 3485500 | 22 | 1 | 77 | N | 40144 | Cytochrome b (Fragment) OS=Rattus norvegicus OX=10116 GN=Cytb PE=3 SV=1 |
| tr\|A0A0U2 | **8** | 40 | 202.18 | 19 | 19 | 3485500 | 22 | 1 | 77 | N | 40981 | Cytochrome b (Fragment) OS=Rattus norvegicus OX=10116 PE=3 SV=1 |
| tr\|A0A0U2 | **8** | 41 | 202.18 | 19 | 19 | 3485500 | 22 | 1 | 77 | N | 41037 | Cytochrome b (Fragment) OS=Rattus norvegicus OX=10116 PE=3 SV=1 |
| tr\|A0A411 | **8** | 42 | 202.18 | 19 | 19 | 3485500 | 22 | 1 | 77 | N | 41017 | Cytochrome b (Fragment) OS=Rattus norvegicus OX=10116 GN=Cytb PE=3 SV=1 |
| tr\|F8QU31 | **8** | 43 | 202.18 | 18 | 18 | 3485500 | 22 | 1 | 77 | N | 42181 | Cytochrome b (Fragment) OS=Rattus norvegicus OX=10116 GN=cytb PE=3 SV=1 |
| tr\|A0A411 | **8** | 44 | 202.18 | 18 | 18 | 3485500 | 22 | 1 | 77 | N | 42238 | Cytochrome b (Fragment) OS=Rattus norvegicus OX=10116 GN=Cytb PE=3 SV=1 |
| tr\|L0L4L8\| | **8** | 45 | 202.18 | 18 | 18 | 3485500 | 22 | 1 | 77 | N | 42470 | Cytochrome b (Fragment) OS=Rattus norvegicus OX=10116 GN=cytb PE=3 SV=1 |
| tr\|A0A140 | **8** | 46 | 202.18 | 18 | 18 | 3485500 | 22 | 1 | 77 | N | 42574 | Cytochrome b (Fragment) OS=Rattus norvegicus OX=10116 PE=3 SV=1 |
| tr\|A0A140 | **8** | 47 | 202.18 | 18 | 18 | 3485500 | 22 | 1 | 77 | N | 42588 | Cytochrome b (Fragment) OS=Rattus norvegicus OX=10116 PE=3 SV=1 |
| tr\|A0A140 | **8** | 48 | 202.18 | 18 | 18 | 3485500 | 22 | 1 | 77 | N | 42527 | Cytochrome b (Fragment) OS=Rattus norvegicus OX=10116 PE=3 SV=1 |
| tr\|A0A385 | **8** | 49 | 202.18 | 18 | 18 | 3485500 | 22 | 1 | 77 | N | 42766 | Cytochrome b (Fragment) OS=Rattus norvegicus OX=10116 PE=3 SV=1 |
| tr\|A0A3Q8 | **8** | 50 | 202.18 | 18 | 18 | 3485500 | 22 | 1 | 77 | N | 42712 | Cytochrome b (Fragment) OS=Rattus norvegicus OX=10116 GN=Cytb PE=3 SV=1 |
| tr\|A0A3Q8 | **8** | 38 | 202.18 | 18 | 18 | 3485500 | 22 | 1 | 77 | N | 42622 | Cytochrome b (Fragment) OS=Rattus norvegicus OX=10116 GN=Cytb PE=3 SV=1 |
| tr\|A0A3S6 | **8** | 52 | 202.18 | 18 | 18 | 3485500 | 22 | 1 | 77 | N | 42698 | Cytochrome b (Fragment) OS=Rattus norvegicus OX=10116 GN=Cytb PE=3 SV=1 |
| tr\|A0A220 | **8** | 54 | 202.18 | 18 | 18 | 3485500 | 22 | 1 | 77 | N | 43018 | Cytochrome b OS=Rattus norvegicus OX=10116 PE=3 SV=1 |
| tr\|D6NSP2 | **8** | 55 | 202.18 | 18 | 18 | 3485500 | 22 | 1 | 77 | N | 42945 | Cytochrome b (Fragment) OS=Rattus norvegicus OX=10116 GN=cytb PE=3 SV=1 |
| tr\|D6NSS6 | **8** | 56 | 202.18 | 18 | 18 | 3485500 | 22 | 1 | 77 | N | 42993 | Cytochrome b (Fragment) OS=Rattus norvegicus OX=10116 GN=cytb PE=3 SV=1 |
| tr\|D6NSQ3 | **8** | 57 | 202.18 | 18 | 18 | 3485500 | 22 | 1 | 77 | N | 43012 | Cytochrome b (Fragment) OS=Rattus norvegicus OX=10116 GN=cytb PE=3 SV=1 |
| tr\|D6NSR3 | **8** | 58 | 202.18 | 18 | 18 | 3485500 | 22 | 1 | 77 | N | 42986 | Cytochrome b (Fragment) OS=Rattus norvegicus OX=10116 GN=cytb PE=3 SV=1 |
| tr\|D6NSP4 | **8** | 59 | 202.18 | 18 | 18 | 3485500 | 22 | 1 | 77 | N | 43016 | Cytochrome b (Fragment) OS=Rattus norvegicus OX=10116 GN=cytb PE=3 SV=1 |
| tr\|D6NSR8 | **8** | 61 | 202.18 | 18 | 18 | 3485500 | 22 | 1 | 77 | N | 42998 | Cytochrome b (Fragment) OS=Rattus norvegicus OX=10116 GN=cytb PE=3 SV=1 |
| tr\|F2Q6S6 | **8** | 63 | 202.18 | 18 | 18 | 3485500 | 22 | 1 | 77 | N | 42938 | Cytochrome b (Fragment) OS=Rattus norvegicus OX=10116 GN=cytb PE=3 SV=1 |
| tr\|A0A0A1 | **8** | 64 | 202.18 | 18 | 18 | 3485500 | 22 | 1 | 77 | N | 42989 | Cytochrome b OS=Rattus norvegicus OX=10116 GN=CYTB PE=3 SV=1 |
| tr\|D6NSS7 | **8** | 65 | 202.18 | 18 | 18 | 3485500 | 22 | 1 | 77 | N | 42948 | Cytochrome b (Fragment) OS=Rattus norvegicus OX=10116 GN=cytb PE=3 SV=1 |
| tr\|Q8SEY9 | **8** | 66 | 202.18 | 18 | 18 | 3485500 | 22 | 1 | 77 | N | 43015 | Cytochrome b OS=Rattus norvegicus OX=10116 GN=cytb PE=3 SV=1 |
| tr\|D6NSR7 | **8** | 67 | 202.18 | 18 | 18 | 3485500 | 22 | 1 | 77 | N | 42982 | Cytochrome b (Fragment) OS=Rattus norvegicus OX=10116 GN=cytb PE=3 SV=1 |
| tr\|A0A220 | **8** | 68 | 202.18 | 18 | 18 | 3485500 | 22 | 1 | 77 | N | 42968 | Cytochrome b OS=Rattus norvegicus OX=10116 PE=3 SV=1 |
| tr\|F2Q6S5 | **8** | 69 | 202.18 | 18 | 18 | 3485500 | 22 | 1 | 77 | N | 42952 | Cytochrome b (Fragment) OS=Rattus norvegicus OX=10116 GN=cytb PE=3 SV=1 |
| tr\|D6NSQ0 | **8** | 71 | 202.18 | 18 | 18 | 3485500 | 22 | 1 | 77 | N | 42968 | Cytochrome b (Fragment) OS=Rattus norvegicus OX=10116 GN=cytb PE=3 SV=1 |
| tr\|Q5UAI7 | **8** | 72 | 202.18 | 18 | 18 | 3485500 | 22 | 1 | 77 | N | 42998 | Cytochrome b OS=Rattus norvegicus OX=10116 GN=CYTB PE=3 SV=1 |
| tr\|D6NSR2 | **8** | 73 | 202.18 | 18 | 18 | 3485500 | 22 | 1 | 77 | N | 43073 | Cytochrome b (Fragment) OS=Rattus norvegicus OX=10116 GN=cytb PE=3 SV=1 |
| tr\|A0A0S1 | **8** | 74 | 202.18 | 18 | 18 | 3485500 | 22 | 1 | 77 | N | 43002 | Cytochrome b OS=Rattus norvegicus OX=10116 GN=CYTB PE=3 SV=1 |
| tr\|D6NSR1 | **8** | 75 | 202.18 | 18 | 18 | 3485500 | 22 | 1 | 77 | N | 43002 | Cytochrome b (Fragment) OS=Rattus norvegicus OX=10116 GN=cytb PE=3 SV=1 |
| tr\|Q8HIC4 | **8** | 76 | 202.18 | 18 | 18 | 3485500 | 22 | 1 | 77 | N | 43012 | Cytochrome b OS=Rattus norvegicus OX=10116 GN=Mt-cyb PE=3 SV=1 |
| tr\|A0A220 | **8** | 77 | 202.18 | 18 | 18 | 3485500 | 22 | 1 | 77 | N | 42968 | Cytochrome b OS=Rattus norvegicus OX=10116 PE=3 SV=1 |
| tr\|A0A0S1 | **8** | 78 | 202.18 | 18 | 18 | 3485500 | 22 | 1 | 77 | N | 43016 | Cytochrome b OS=Rattus norvegicus OX=10116 GN=CYTB PE=3 SV=1 |
| tr\|R9TKN1 | **8** | 119 | 202.18 | 18 | 18 | 3485500 | 22 | 1 | 77 | N | 43012 | Cytochrome b OS=Rattus norvegicus OX=10116 GN=CYTB PE=3 SV=1 |
| P00159\|CY | **8** | 79 | 202.18 | 18 | 18 | 3485500 | 22 | 1 | 77 | N | 43012 | Cytochrome b OS=Rattus norvegicus OX=10116 GN=Mt-Cyb PE=3 SV=3 |
| tr\|L0N311 | **8** | 80 | 202.18 | 18 | 18 | 3485500 | 22 | 1 | 77 | N | 43045 | Cytochrome b (Fragment) OS=Rattus norvegicus OX=10116 GN=cytb PE=3 SV=1 |
| tr\|A0A220 | **8** | 81 | 202.18 | 18 | 18 | 3485500 | 22 | 1 | 77 | N | 42970 | Cytochrome b OS=Rattus norvegicus OX=10116 PE=3 SV=1 |
| tr\|H2KXA0 | **8** | 82 | 202.18 | 18 | 18 | 3485500 | 22 | 1 | 77 | N | 42978 | Cytochrome b (Fragment) OS=Rattus norvegicus OX=10116 GN=cytb PE=3 SV=1 |
| tr\|A0A096 | **8** | 83 | 202.18 | 18 | 18 | 3485500 | 22 | 1 | 77 | N | 43042 | Cytochrome b (Fragment) OS=Rattus norvegicus OX=10116 PE=3 SV=1 |
| tr\|D6NSP5 | **8** | 120 | 202.18 | 18 | 18 | 3485500 | 22 | 1 | 77 | N | 43012 | Cytochrome b (Fragment) OS=Rattus norvegicus OX=10116 GN=cytb PE=3 SV=1 |
| tr\|A0A385 | **9** | 212 | 194.82 | 18 | 18 | 196240 | 22 | 1 | 73 | N | 42751 | Cytochrome b (Fragment) OS=Rattus norvegicus OX=10116 PE=3 SV=1 |
| tr\|B0BNE6 | **17** | 5411 | 194.78 | 27 | 27 | 9077000 | 18 | 18 | 42 | N | 23970 | NADH dehydrogenase (Ubiquinone) Fe-S protein 8 (Predicted), isoform CRA_a OS=Rattus norvegicus OX=10116 GN=Ndufs8 PE=1 SV=1 |

| | | | | | | | | | | | | |
|---|---|---|---|---|---|---|---|---|---|---|---|---|
| tr\|D3ZF13\| | **14** | 3210 | 192.11 | 31 | 31 | 22300000 | 21 | 21 | 51 | Y | 17514 | Acyl carrier protein OS=Rattus norvegicus OX=10116 GN=Ndufab1 PE=1 SV=1 |
| tr\|D4A565 | **20** | 5412 | 190.07 | 37 | 37 | 11795000 | 16 | 16 | 35 | N | 21664 | NADH dehydrogenase (Ubiquinone) 1 beta subcomplex, 5 (Predicted), isoform CRA_b OS=Rattus norvegicus OX=10116 GN=Ndufb5 PE=1 SV=1 |
| tr\|Q5XIH3\| | **11** | 5408 | 186.57 | 22 | 22 | 21344000 | 20 | 20 | 65 | N | 50731 | NADH dehydrogenase [ubiquinone] flavoprotein 1, mitochondrial OS=Rattus norvegicus OX=10116 GN=Ndufv1 PE=1 SV=1 |
| tr\|Q5PQZ9 | **16** | 2821 | 185.99 | 39 | 39 | 30829000 | 13 | 13 | 42 | N | 14359 | NADH dehydrogenase [ubiquinone] 1 subunit C2 OS=Rattus norvegicus OX=10116 GN=Ndufc2 PE=1 SV=1 |
| tr\|A0A0G2 | **18** | 3028 | 183.81 | 37 | 37 | 18316000 | 18 | 18 | 37 | Y | 19965 | NADH dehydrogenase [ubiquinone] 1 alpha subcomplex subunit 8 OS=Rattus norvegicus OX=10116 GN=Ndufa8 PE=1 SV=1 |
| P10719\|AT | **23** | 1 | 181.93 | 10 | 10 | 10128000 | 13 | 13 | 26 | Y | 56354 | ATP synthase subunit beta, mitochondrial OS=Rattus norvegicus OX=10116 GN=Atp5f1b PE=1 SV=2 |
| tr\|G3V6D3 | **23** | 2 | 181.93 | 10 | 10 | 10128000 | 13 | 13 | 26 | Y | 56345 | ATP synthase subunit beta OS=Rattus norvegicus OX=10116 GN=Atp5f1b PE=1 SV=1 |
| Q561S0\|N | **10** | 5410 | 177.25 | 46 | 46 | 20602000 | 33 | 33 | 72 | Y | 40493 | NADH dehydrogenase [ubiquinone] 1 alpha subcomplex subunit 10, mitochondrial OS=Rattus norvegicus OX=10116 GN=Ndufa10 PE=1 SV=1 |
| tr\|D4A7L4 | **19** | 2872 | 168.95 | 29 | 29 | 18411000 | 11 | 11 | 37 | N | 17634 | NADH dehydrogenase (Ubiquinone) 1 beta subcomplex, 11 (Predicted) OS=Rattus norvegicus OX=10116 GN=Ndufb11 PE=1 SV=1 |
| Q80W89\|N | **25** | 5440 | 164.41 | 21 | 21 | 12554000 | 9 | 9 | 25 | N | 14854 | NADH dehydrogenase [ubiquinone] 1 alpha subcomplex subunit 11 OS=Rattus norvegicus OX=10116 GN=Ndufa11 PE=2 SV=1 |
| tr\|B5DEL8\| | **31** | 5442 | 157.61 | 23 | 23 | 8304200 | 7 | 7 | 17 | N | 12700 | NADH dehydrogenase (Ubiquinone) Fe-S protein 5 OS=Rattus norvegicus OX=10116 GN=Ndufs5 PE=1 SV=1 |
| tr\|A0A0G2 | **31** | 5443 | 157.61 | 23 | 23 | 8304200 | 7 | 7 | 17 | N | 12730 | Uncharacterized protein OS=Rattus norvegicus OX=10116 PE=4 SV=1 |
| tr\|D4A0T0 | **22** | 180 | 154.07 | 34 | 34 | 31942000 | 13 | 13 | 29 | N | 20859 | NADH:ubiquinone oxidoreductase subunit B10 OS=Rattus norvegicus OX=10116 GN=Ndufb10 PE=1 SV=1 |
| tr\|A9UMW | **26** | 5453 | 148.1 | 43 | 43 | 14590000 | 9 | 9 | 18 | Y | 9255 | Ndufa3 protein (Fragment) OS=Rattus norvegicus OX=10116 GN=Ndufa3 PE=2 SV=1 |
| tr\|M0RB63 | **26** | 5454 | 148.1 | 43 | 43 | 14590000 | 9 | 9 | 18 | Y | 9372 | RCG63041 OS=Rattus norvegicus OX=10116 GN=LOC684509 PE=4 SV=1 |
| tr\|A0A0G2 | **26** | 5455 | 148.1 | 40 | 40 | 14590000 | 9 | 9 | 18 | Y | 10170 | NADH:ubiquinone oxidoreductase subunit A3 OS=Rattus norvegicus OX=10116 GN=Ndufa3 PE=1 SV=1 |
| tr\|F1LXA0\| | **38** | 5458 | 146.06 | 17 | 17 | 4650700 | 5 | 5 | 11 | N | 17178 | NADH dehydrogenase [ubiquinone] 1 alpha subcomplex subunit 12 OS=Rattus norvegicus OX=10116 GN=Ndufa12 PE=1 SV=2 |
| tr\|Q06QK5 | **24** | 5413 | 139.43 | 11 | 11 | 12761000 | 13 | 13 | 25 | Y | 68606 | NADH-ubiquinone oxidoreductase chain 5 OS=Rattus norvegicus OX=10116 GN=ND5 PE=3 SV=1 |
| P11661\|NU | **24** | 5414 | 139.43 | 11 | 11 | 12761000 | 13 | 13 | 25 | Y | 68618 | NADH-ubiquinone oxidoreductase chain 5 OS=Rattus norvegicus OX=10116 GN=Mtnd5 PE=3 SV=3 |
| tr\|Q06QG6 | **24** | 5415 | 139.43 | 11 | 11 | 12761000 | 13 | 13 | 25 | Y | 68588 | NADH-ubiquinone oxidoreductase chain 5 OS=Rattus norvegicus OX=10116 GN=ND5 PE=3 SV=1 |
| tr\|Q06QA1 | **24** | 5416 | 139.43 | 11 | 11 | 12761000 | 13 | 13 | 25 | Y | 68574 | NADH-ubiquinone oxidoreductase chain 5 OS=Rattus norvegicus OX=10116 GN=ND5 PE=3 SV=1 |
| tr\|Q8SEZ0\| | **24** | 5417 | 139.43 | 11 | 11 | 12761000 | 13 | 13 | 25 | Y | 68618 | NADH-ubiquinone oxidoreductase chain 5 OS=Rattus norvegicus OX=10116 GN=Mt-nd5 PE=3 SV=1 |
| tr\|A0A096 | **24** | 5418 | 139.43 | 11 | 11 | 12761000 | 13 | 13 | 25 | Y | 68584 | NADH-ubiquinone oxidoreductase chain 5 OS=Rattus norvegicus OX=10116 GN=ND5 PE=3 SV=1 |
| P15999\|AT | **21** | 3 | 130.49 | 19 | 19 | 5484100 | 14 | 14 | 32 | N | 59754 | ATP synthase subunit alpha, mitochondrial OS=Rattus norvegicus OX=10116 GN=Atp5f1a PE=1 SV=2 |
| tr\|F1LP05\| | **21** | 4 | 130.49 | 19 | 19 | 5484100 | 14 | 14 | 32 | N | 59813 | ATP synthase subunit alpha OS=Rattus norvegicus OX=10116 GN=Atp5f1a PE=1 SV=1 |
| tr\|D3ZS58\| | **36** | 5449 | 125.37 | 31 | 31 | 3109900 | 5 | 5 | 11 | N | 10845 | NADH dehydrogenase [ubiquinone] 1 alpha subcomplex subunit 2 OS=Rattus norvegicus OX=10116 GN=Ndufa2 PE=1 SV=1 |
| P11240\|CO | **37** | 86 | 123.58 | 27 | 27 | 1870900 | 6 | 6 | 11 | N | 16130 | Cytochrome c oxidase subunit 5A, mitochondrial OS=Rattus norvegicus OX=10116 GN=Cox5a PE=1 SV=1 |
| Q5XIF3\|ND | **30** | 5457 | 122.21 | 12 | 12 | 1812700 | 7 | 7 | 17 | N | 19741 | NADH dehydrogenase [ubiquinone] iron-sulfur protein 4, mitochondrial OS=Rattus norvegicus OX=10116 GN=Ndufs4 PE=1 SV=1 |
| tr\|D3ZZ21\| | **29** | 5448 | 119.92 | 34 | 34 | 5471200 | 8 | 8 | 17 | Y | 15638 | NADH dehydrogenase (Ubiquinone) 1 beta subcomplex, 6 (Predicted) OS=Rattus norvegicus OX=10116 GN=Ndufb6 PE=1 SV=1 |
| tr\|D3ZE15\| | **33** | 5447 | 117.61 | 36 | 36 | 4475000 | 8 | 8 | 14 | Y | 16777 | NADH:ubiquinone oxidoreductase subunit A13 OS=Rattus norvegicus OX=10116 GN=Ndufa13 PE=1 SV=1 |
| tr\|D4A3V2 | **43** | 5424 | 116.44 | 22 | 22 | 1759200 | 5 | 5 | 8 | N | 15224 | NADH dehydrogenase [ubiquinone] 1 alpha subcomplex subunit 6 OS=Rattus norvegicus OX=10116 GN=Ndufa6 PE=1 SV=1 |
| tr\|Q5UAJ6 | **34** | 24 | 111.04 | 19 | 19 | 2728500 | 10 | 10 | 14 | Y | 25942 | Cytochrome c oxidase subunit 2 OS=Rattus norvegicus OX=10116 GN=COX2 PE=3 SV=1 |
| Q63362\|N | **41** | 5439 | 108.29 | 28 | 28 | 1982000 | 5 | 5 | 9 | N | 13412 | NADH dehydrogenase [ubiquinone] 1 alpha subcomplex subunit 5 OS=Rattus norvegicus OX=10116 GN=Ndufa5 PE=1 SV=3 |
| Q5M9I5\|Q | **27** | 2875 | 105.16 | 21 | 21 | 7211800 | 7 | 7 | 18 | N | 10424 | Cytochrome b-c1 complex subunit 6, mitochondrial OS=Rattus norvegicus OX=10116 GN=Uqcrh PE=3 SV=1 |
| tr\|A9UMV9 | **32** | 5450 | 104.46 | 21 | 21 | 11686000 | 4 | 4 | 16 | N | 12500 | NADH:ubiquinone oxidoreductase subunit A7 OS=Rattus norvegicus OX=10116 GN=Ndufa7 PE=1 SV=1 |
| P00507\|AA | **40** | 19 | 98.56 | 11 | 11 | 1665800 | 6 | 6 | 9 | Y | 47314 | Aspartate aminotransferase, mitochondrial OS=Rattus norvegicus OX=10116 GN=Got2 PE=1 SV=2 |
| tr\|Q35733 | **49** | 5460 | 96.51 | 14 | 14 | 1053700 | 4 | 4 | 5 | N | 8389 | NADH-ubiquinone oxidoreductase chain 6 (Fragment) OS=Rattus norvegicus OX=10116 PE=3 SV=1 |
| tr\|Q06QD9 | **49** | 5461 | 96.51 | 6 | 6 | 1053700 | 4 | 4 | 5 | N | 18943 | NADH-ubiquinone oxidoreductase chain 6 OS=Rattus norvegicus OX=10116 GN=ND6 PE=3 SV=1 |
| tr\|Q7HKW | **49** | 5462 | 96.51 | 6 | 6 | 1053700 | 4 | 4 | 5 | N | 18957 | NADH-ubiquinone oxidoreductase chain 6 OS=Rattus norvegicus OX=10116 GN=NADH6 PE=3 SV=1 |
| P02563\|MY | **45** | 196 | 96.23 | 2 | 2 | 1235500 | 5 | 5 | 7 | N | 223506 | Myosin-6 OS=Rattus norvegicus OX=10116 GN=Myh6 PE=1 SV=2 |
| tr\|B2RYW3 | **28** | 5441 | 93.62 | 35 | 35 | 10872000 | 6 | 6 | 17 | Y | 21892 | NADH dehydrogenase (Ubiquinone) 1 beta subcomplex, 9 OS=Rattus norvegicus OX=10116 GN=Ndufb9 PE=1 SV=1 |
| tr\|D3ZCZ9\| | **42** | 5435 | 88.86 | 29 | 29 | 1712900 | 4 | 4 | 9 | N | 13040 | NADH dehydrogenase [ubiquinone] iron-sulfur protein 6, mitochondrial OS=Rattus norvegicus OX=10116 GN=LOC100912599 PE=1 SV=1 |
| tr\|F1LPG5\| | **64** | 5445 | 84.44 | 29 | 29 | 975990 | 3 | 3 | 3 | N | 15064 | NADH:ubiquinone oxidoreductase subunit B4 OS=Rattus norvegicus OX=10116 GN=Ndufb4 PE=1 SV=1 |
| tr\|Q5RJN0 | **39** | 564 | 79.92 | 11 | 11 | 3815700 | 5 | 5 | 10 | N | 23945 | NADH dehydrogenase (Ubiquinone) Fe-S protein 7 OS=Rattus norvegicus OX=10116 GN=Ndufs7 PE=1 SV=1 |
| P67779\|PH | **55** | 5407 | 76.82 | 10 | 10 | 602730 | 3 | 3 | 4 | N | 29820 | Prohibitin OS=Rattus norvegicus OX=10116 GN=Phb PE=1 SV=1 |
| tr\|D4A4P3 | **44** | 5451 | 76.32 | 25 | 25 | 2427200 | 4 | 4 | 8 | Y | 11267 | NADH:ubiquinone oxidoreductase subunit B3 OS=Rattus norvegicus OX=10116 GN=Ndufb3 PE=1 SV=1 |
| P05508\|NU | **46** | 5425 | 74.52 | 7 | 7 | 1685000 | 5 | 5 | 7 | N | 51783 | NADH-ubiquinone oxidoreductase chain 4 OS=Rattus norvegicus OX=10116 GN=Mtnd4 PE=3 SV=3 |
| tr\|D2E6K0 | **46** | 5426 | 74.52 | 7 | 7 | 1685000 | 5 | 5 | 7 | N | 51833 | NADH-ubiquinone oxidoreductase chain 4 OS=Rattus norvegicus OX=10116 GN=ND4 PE=3 SV=1 |
| tr\|A7XYB9 | **46** | 5427 | 74.52 | 7 | 7 | 1685000 | 5 | 5 | 7 | N | 51773 | NADH-ubiquinone oxidoreductase chain 4 OS=Rattus norvegicus OX=10116 GN=Nd4 PE=3 SV=1 |
| tr\|Q06QE1 | **46** | 5428 | 74.52 | 7 | 7 | 1685000 | 5 | 5 | 7 | N | 51851 | NADH-ubiquinone oxidoreductase chain 4 OS=Rattus norvegicus OX=10116 GN=ND4 PE=3 SV=1 |
| tr\|Q06QA2 | **46** | 5429 | 74.52 | 7 | 7 | 1685000 | 5 | 5 | 7 | N | 51787 | NADH-ubiquinone oxidoreductase chain 4 OS=Rattus norvegicus OX=10116 GN=ND4 PE=3 SV=1 |
| tr\|Q7HKW | **46** | 5430 | 74.52 | 7 | 7 | 1685000 | 5 | 5 | 7 | N | 51801 | NADH-ubiquinone oxidoreductase chain 4 (Fragment) OS=Rattus norvegicus OX=10116 GN=NADH4 PE=3 SV=1 |
| tr\|Q06QG7 | **46** | 5431 | 74.52 | 7 | 7 | 1685000 | 5 | 5 | 7 | N | 51791 | NADH-ubiquinone oxidoreductase chain 4 OS=Rattus norvegicus OX=10116 GN=ND4 PE=3 SV=1 |
| tr\|Q06Q89 | **46** | 5432 | 74.52 | 7 | 7 | 1685000 | 5 | 5 | 7 | N | 51819 | NADH-ubiquinone oxidoreductase chain 4 OS=Rattus norvegicus OX=10116 GN=ND4 PE=3 SV=1 |
| tr\|Q8HIC6\| | **46** | 5434 | 74.52 | 7 | 7 | 1685000 | 5 | 5 | 7 | N | 51783 | NADH-ubiquinone oxidoreductase chain 4 OS=Rattus norvegicus OX=10116 GN=Mt-nd4 PE=3 SV=1 |
| tr\|Q35737 | **46** | 5433 | 74.52 | 7 | 7 | 1685000 | 5 | 5 | 7 | N | 51765 | NADH-ubiquinone oxidoreductase chain 4 OS=Rattus norvegicus OX=10116 PE=3 SV=1 |
| Q6AXV4\|SA | **75** | 2803 | 70.96 | 6 | 6 | 151070 | 2 | 2 | 2 | N | 51960 | Sorting and assembly machinery component 50 homolog OS=Rattus norvegicus OX=10116 GN=Samm50 PE=1 SV=1 |

| | | | | | | | | | | | | |
|---|---|---|---|---|---|---|---|---|---|---|---|---|
| tr\|Q8SEZ8\| | **35** | 671 | 61.93 | 6 | 6 | 5781300 | 5 | 5 | 12 | N | 36133 | NADH-ubiquinone oxidoreductase chain 1 (Fragment) OS=Rattus norvegicus OX=10116 GN=NADH1 PE=3 SV=1 |
| P03889\|NU | **35** | 672 | 61.93 | 6 | 6 | 5781300 | 5 | 5 | 12 | N | 36145 | NADH-ubiquinone oxidoreductase chain 1 OS=Rattus norvegicus OX=10116 GN=Mtnd1 PE=1 SV=3 |
| tr\|Q8HID1 | **35** | 673 | 61.93 | 6 | 6 | 5781300 | 5 | 5 | 12 | N | 36145 | NADH-ubiquinone oxidoreductase chain 1 OS=Rattus norvegicus OX=10116 GN=Mt-nd1 PE=3 SV=1 |
| tr\|D2E6L7\| | **35** | 5456 | 61.93 | 6 | 6 | 5781300 | 5 | 5 | 12 | N | 36062 | NADH-ubiquinone oxidoreductase chain 1 OS=Rattus norvegicus OX=10116 GN=ND1 PE=3 SV=1 |
| P35435\|AT | **77** | 5 | 57.75 | 8 | 8 | 218070 | 2 | 2 | 2 | N | 30191 | ATP synthase subunit gamma, mitochondrial OS=Rattus norvegicus OX=10116 GN=Atp5f1c PE=1 SV=2 |
| tr\|Q6QI09 | **77** | 6 | 57.75 | 4 | 4 | 218070 | 2 | 2 | 2 | N | 67721 | ATP synthase subunit gamma, mitochondrial OS=Rattus norvegicus OX=10116 GN=Taf3 PE=1 SV=1 |
| P19511\|AT | **53** | 10 | 57.14 | 9 | 9 | 812140 | 2 | 2 | 4 | N | 28869 | ATP synthase F(0) complex subunit B1, mitochondrial OS=Rattus norvegicus OX=10116 GN=Atp5pb PE=1 SV=1 |
| tr\|A0A140 | **65** | 145 | 56.86 | 4 | 4 | 524870 | 2 | 2 | 3 | N | 67049 | MICOS complex subunit MIC60 OS=Rattus norvegicus OX=10116 GN=Immt PE=1 SV=1 |
| Q3KR86\|M | **65** | 146 | 56.86 | 4 | 4 | 524870 | 2 | 2 | 3 | N | 67177 | MICOS complex subunit Mic60 (Fragment) OS=Rattus norvegicus OX=10116 GN=Immt PE=1 SV=1 |
| tr\|A0A0G2 | **65** | 147 | 56.86 | 3 | 3 | 524870 | 2 | 2 | 3 | N | 86230 | MICOS complex subunit MIC60 OS=Rattus norvegicus OX=10116 GN=Immt PE=1 SV=1 |
| Q60587\|EC | **66** | 20 | 56.1 | 5 | 5 | 240460 | 2 | 2 | 3 | N | 51414 | Trifunctional enzyme subunit beta, mitochondrial OS=Rattus norvegicus OX=10116 GN=Hadhb PE=1 SV=1 |
| tr\|A0A0G2 | **66** | 21 | 56.1 | 5 | 5 | 240460 | 2 | 2 | 3 | N | 52568 | Trifunctional enzyme subunit beta, mitochondrial OS=Rattus norvegicus OX=10116 GN=Hadhb PE=1 SV=1 |
| tr\|B2RYU0 | **48** | 5486 | 50.83 | 18 | 18 | 1243300 | 3 | 3 | 7 | N | 11842 | NADH dehydrogenase (Ubiquinone) 1 beta subcomplex, 2 (Predicted), isoform CRA_b OS=Rattus norvegicus OX=10116 GN=Ndufb2 PE=1 SV=1 |
| Q64428\|EC | **54** | 9 | 49.05 | 2 | 2 | 379450 | 2 | 2 | 4 | N | 82665 | Trifunctional enzyme subunit alpha, mitochondrial OS=Rattus norvegicus OX=10116 GN=Hadha PE=1 SV=2 |
| P31399\|AT | **78** | 7 | 48.22 | 12 | 12 | 293370 | 2 | 2 | 2 | N | 18763 | ATP synthase subunit d, mitochondrial OS=Rattus norvegicus OX=10116 GN=Atp5pd PE=1 SV=3 |
| P10888\|CC | **56** | 290 | 47.61 | 14 | 14 | 556290 | 3 | 3 | 4 | N | 19515 | Cytochrome c oxidase subunit 4 isoform 1, mitochondrial OS=Rattus norvegicus OX=10116 GN=Cox4i1 PE=1 SV=1 |
| tr\|A0A0G2 | **50** | 5422 | 47.03 | 13 | 13 | 1225000 | 3 | 3 | 4 | N | 33168 | Prohibitin OS=Rattus norvegicus OX=10116 GN=Phb2 PE=1 SV=1 |
| Q5XIH7\|PH | **50** | 5423 | 47.03 | 13 | 13 | 1225000 | 3 | 3 | 4 | N | 33312 | Prohibitin-2 OS=Rattus norvegicus OX=10116 GN=Phb2 PE=1 SV=1 |
| tr\|Q06Q97 | **51** | 1299 | 41.27 | 6 | 6 | 669860 | 3 | 3 | 4 | N | 38485 | NADH-ubiquinone oxidoreductase chain 2 OS=Rattus norvegicus OX=10116 GN=ND2 PE=3 SV=1 |
| tr\|A0A0A1 | **51** | 1300 | 41.27 | 6 | 6 | 669860 | 3 | 3 | 4 | N | 38542 | NADH-ubiquinone oxidoreductase chain 2 OS=Rattus norvegicus OX=10116 GN=ND2 PE=3 SV=1 |
| tr\|Q5UAJ8 | **51** | 1301 | 41.27 | 6 | 6 | 669860 | 3 | 3 | 4 | N | 38455 | NADH-ubiquinone oxidoreductase chain 2 OS=Rattus norvegicus OX=10116 GN=ND2 PE=3 SV=1 |
| tr\|Q8HID0 | **51** | 1303 | 41.27 | 6 | 6 | 669860 | 3 | 3 | 4 | N | 38653 | NADH-ubiquinone oxidoreductase chain 2 OS=Rattus norvegicus OX=10116 GN=Mt-nd2 PE=3 SV=1 |
| P11662\|NU | **51** | 1304 | 41.27 | 6 | 6 | 669860 | 3 | 3 | 4 | N | 38653 | NADH-ubiquinone oxidoreductase chain 2 OS=Rattus norvegicus OX=10116 GN=Mtnd2 PE=3 SV=3 |
| tr\|Q06QH5 | **51** | 1305 | 41.27 | 6 | 6 | 669860 | 3 | 3 | 4 | N | 38626 | NADH-ubiquinone oxidoreductase chain 2 OS=Rattus norvegicus OX=10116 GN=ND2 PE=3 SV=1 |
| tr\|Q8SEZ7\| | **51** | 1306 | 41.27 | 6 | 6 | 669860 | 3 | 3 | 4 | N | 38580 | NADH-ubiquinone oxidoreductase chain 2 (Fragment) OS=Rattus norvegicus OX=10116 GN=NADH2 PE=3 SV=1 |
| tr\|Q06QB0 | **51** | 5452 | 41.27 | 6 | 6 | 669860 | 3 | 3 | 4 | N | 38534 | NADH-ubiquinone oxidoreductase chain 2 OS=Rattus norvegicus OX=10116 GN=ND2 PE=3 SV=1 |
| tr\|D2E6P4 | **51** | 1307 | 41.27 | 6 | 6 | 669860 | 3 | 3 | 4 | N | 38623 | NADH-ubiquinone oxidoreductase chain 2 OS=Rattus norvegicus OX=10116 GN=ND2 PE=3 SV=1 |
| tr\|Q06QE9 | **51** | 1308 | 41.27 | 6 | 6 | 669860 | 3 | 3 | 4 | N | 38598 | NADH-ubiquinone oxidoreductase chain 2 OS=Rattus norvegicus OX=10116 GN=ND2 PE=3 SV=1 |
| P80432\|CC | **116** | 134 | 38.09 | 21 | 21 | 105400 | 1 | 1 | 1 | N | 7375 | Cytochrome c oxidase subunit 7C, mitochondrial OS=Rattus norvegicus OX=10116 GN=Cox7c PE=1 SV=2 |
| P02770\|AL | **117** | 2832 | 35.8 | 1 | 1 | 72142 | 1 | 1 | 1 | Y | 68731 | Serum albumin OS=Rattus norvegicus OX=10116 GN=Alb PE=1 SV=2 |
| tr\|A0A0G2 | **117** | 2833 | 35.8 | 1 | 1 | 72142 | 1 | 1 | 1 | Y | 68759 | Serum albumin OS=Rattus norvegicus OX=10116 GN=Alb PE=1 SV=1 |
| tr\|G3V644 | **67** | 1383 | 34.98 | 3 | 3 | 873790 | 2 | 2 | 3 | N | 49300 | NADH dehydrogenase (Ubiquinone) flavoprotein 3-like, isoform CRA_a OS=Rattus norvegicus OX=10116 GN=Ndufv3 PE=1 SV=1 |
| tr\|O54755 | **67** | 1384 | 34.98 | 3 | 3 | 873790 | 2 | 2 | 3 | N | 49312 | MIPP65 OS=Rattus norvegicus OX=10116 GN=Ndufv3 PE=2 SV=1 |
| Q7TQ16\|Q | **59** | 2828 | 33.56 | 20 | 20 | 1140400 | 2 | 2 | 3 | N | 9849 | Cytochrome b-c1 complex subunit 8 OS=Rattus norvegicus OX=10116 GN=Uqcrq PE=3 SV=1 |
| tr\|D3ZUX5 | **118** | 2815 | 33.3 | 4 | 4 | 122690 | 1 | 1 | 1 | N | 26435 | MICOS complex subunit OS=Rattus norvegicus OX=10116 GN=Chchd3 PE=1 SV=1 |
| P10817\|CX | **79** | 151 | 29.76 | 12 | 12 | 617270 | 1 | 1 | 2 | N | 10487 | Cytochrome c oxidase subunit 6A2, mitochondrial (Fragment) OS=Rattus norvegicus OX=10116 GN=Cox6a2 PE=1 SV=3 |
| tr\|G3V8M4 | **79** | 152 | 29.76 | 11 | 11 | 617270 | 1 | 1 | 2 | N | 10802 | Cytochrome c oxidase subunit 6A, mitochondrial OS=Rattus norvegicus OX=10116 GN=Cox6a2 PE=3 SV=1 |
| tr\|Q5UAJ5 | **119** | 32 | 27.72 | 13 | 13 | 478580 | 1 | 1 | 1 | N | 7642 | ATP synthase protein 8 OS=Rattus norvegicus OX=10116 GN=ATP8 PE=3 SV=1 |
| P11608\|AT | **119** | 35 | 27.72 | 13 | 13 | 478580 | 1 | 1 | 1 | N | 7630 | ATP synthase protein 8 OS=Rattus norvegicus OX=10116 GN=Mt-atp8 PE=1 SV=2 |
| tr\|Q8SEZ4\| | **119** | 33 | 27.72 | 13 | 13 | 478580 | 1 | 1 | 1 | N | 7632 | ATP synthase protein 8 OS=Rattus norvegicus OX=10116 GN=ATPase8 PE=3 SV=1 |
| tr\|Q8HIC8 | **119** | 36 | 27.72 | 13 | 13 | 478580 | 1 | 1 | 1 | N | 7630 | ATP synthase protein 8 OS=Rattus norvegicus OX=10116 GN=Mt-atp8 PE=3 SV=1 |
| P52631\|ST | **96** | 5489 | 27.16 | 1 | 1 | 1333400 | 1 | 1 | 1 | N | 88040 | Signal transducer and activator of transcription 3 OS=Rattus norvegicus OX=10116 GN=Stat3 PE=1 SV=1 |
| tr\|Q5XHZ7 | **80** | 12765 | 26.88 | 1 | 1 | 0 | 1 | 1 | 2 | N | 66660 | MMR_HSR1 domain containing protein RGD1359460 OS=Rattus norvegicus OX=10116 GN=Noa1 PE=2 SV=1 |
| tr\|A0A0G2 | **80** | 12766 | 26.88 | 1 | 1 | 0 | 1 | 1 | 2 | N | 66708 | Nitric oxide-associated 1 OS=Rattus norvegicus OX=10116 GN=Noa1 PE=4 SV=1 |
| tr\|A0A0G2 | **80** | 12767 | 26.88 | 1 | 1 | 0 | 1 | 1 | 2 | N | 77504 | MMR_HSR1 domain containing protein RGD1359460, isoform CRA_a OS=Rattus norvegicus OX=10116 GN=Noa1 PE=4 SV=1 |
| Q62600\|NO | **84** | 5463 | 26.58 | 1 | 1 | 466130 | 2 | 1 | 2 | N | 133290 | Nitric oxide synthase, endothelial OS=Rattus norvegicus OX=10116 GN=Nos3 PE=1 SV=4 |
| Q02253\|M | **82** | 2801 | 26.58 | 4 | 4 | 308360 | 2 | 2 | 2 | N | 57808 | Methylmalonate-semialdehyde dehydrogenase [acylating], mitochondrial OS=Rattus norvegicus OX=10116 GN=Aldh6a1 PE=1 SV=1 |
| tr\|G3V7J0\| | **82** | 2802 | 26.58 | 4 | 4 | 308360 | 2 | 2 | 2 | N | 57748 | Aldehyde dehydrogenase family 6, subfamily A1, isoform CRA_b OS=Rattus norvegicus OX=10116 GN=Aldh6a1 PE=1 SV=1 |
| tr\|A9UMV | **122** | 2873 | 44523 | 11 | 11 | 108350 | 1 | 1 | 1 | N | 6539 | RCG29512 OS=Rattus norvegicus OX=10116 GN=Uqcr11 PE=1 SV=1 |
| Q9ER34\|AC | **97** | 88 | 23 | 2 | 2 | 0 | 1 | 1 | 1 | N | 85433 | Aconitate hydratase, mitochondrial OS=Rattus norvegicus OX=10116 GN=Aco2 PE=1 SV=2 |
| Q62630\|EC | **123** | 12771 | 22.97 | 3 | 3 | 196260 | 1 | 1 | 1 | Y | 27242 | Egl nine homolog 3 OS=Rattus norvegicus OX=10116 GN=Egln3 PE=1 SV=2 |
| tr\|F1LN92\| | **120** | 7847 | 22.93 | 1 | 1 | 111170 | 1 | 1 | 1 | N | 89352 | AFG3-like matrix AAA peptidase subunit 2 OS=Rattus norvegicus OX=10116 GN=Afg3l2 PE=1 SV=1 |
| tr\|Q8SEZ1\| | **121** | 7779 | 22.82 | 6 | 6 | 94812 | 1 | 1 | 1 | N | 13087 | NADH-ubiquinone oxidoreductase chain 3 OS=Rattus norvegicus OX=10116 GN=Mt-nd3 PE=2 SV=1 |
| P05506\|NU | **121** | 7780 | 22.82 | 6 | 6 | 94812 | 1 | 1 | 1 | N | 13087 | NADH-ubiquinone oxidoreductase chain 3 OS=Rattus norvegicus OX=10116 GN=Mtnd3 PE=3 SV=3 |
| tr\|A0A0A1 | **121** | 7781 | 22.82 | 6 | 6 | 94812 | 1 | 1 | 1 | N | 13071 | NADH-ubiquinone oxidoreductase chain 3 OS=Rattus norvegicus OX=10116 GN=ND3 PE=3 SV=1 |
| tr\|Q06QG9 | **121** | 7782 | 22.82 | 6 | 6 | 94812 | 1 | 1 | 1 | N | 13075 | NADH-ubiquinone oxidoreductase chain 3 OS=Rattus norvegicus OX=10116 GN=ND3 PE=3 SV=1 |
| tr\|M0R6J0 | **124** | 12772 | 22.22 | 3 | 3 | 793730 | 1 | 1 | 1 | N | 38374 | Mitochondrial ribosomal protein L39 OS=Rattus norvegicus OX=10116 GN=Mrpl39 PE=1 SV=2 |

| | | | | | | | | | | | | |
|---|---|---|---|---|---|---|---|---|---|---|---|---|
| Q63560\|M | **127** | 3380 | 21.41 | 1 | 1 | 203950 | 1 | 1 | 1 | N | 100485 | Microtubule-associated protein 6 OS=Rattus norvegicus OX=10116 GN=Map6 PE=1 SV=1 |
| B2RYN7\|SF | **87** | 12768 | 44459.00 | 2 | 2 | 0 | 1 | 1 | 1 | Y | 63022 | Spastin OS=Rattus norvegicus OX=10116 GN=Spast PE=1 SV=1 |
