## Supplementary material for "Permeability transition pore-related changes in the proteome and channel activity of ATP synthase dimers and monomers": RHM

| Protein Gr | Protein ID | Accession | -10lgP | Coverage ( | Coverage ( | IBAQ | #Peptides | #Unique | #Spec Sam | PTM | Avg. Mass | Description |
|---|---|---|---|---|---|---|---|---|---|---|---|---|
| 8 | 5 | P10719\|AT | 311.76 | 22 | 22 | 8.10E+07 | 37 | 37 | 105 | Oxidation (M) | | |
| 8 | 6 | tr\|G3V6D3 | 311.76 | 22 | 22 | 8.10E+07 | 37 | 37 | 105 | Oxidation (M) | | |
| 7 | 59 | tr\|D3ZFQ8 | 303.3 | 35 | 35 | 1.61E+08 | 30 | 30 | 118 | Oxidation (M) | | |
| 5 | 58 | P20788\|U | 297.05 | 32 | 32 | 1.19E+08 | 36 | 35 | 132 | | 29446 | Cytochrome b-c1 complex subunit Rieske mitochondrial OS=Rattus norvegicus OX=10116 GN=Uqcrfs1 PE=1 SV=2 |
| 1 | 20 | Q66HF1\|N | 295.51 | 31 | 31 | 1.10E+08 | 79 | 78 | 206 | Carbamidomethylation | | |
| 15 | 212 | tr\|B2RYS8 | 293.12 | 53 | 53 | 5.18E+07 | 27 | 27 | 91 | Oxidation ( | 21959 | NADH dehydrogenase [ubiquinone] 1 beta subcomplex subunit 8 mitochondrial OS=Rattus norvegicus OX=10116 GN=Ndufb8 PE=1 SV=1 |
| 4 | 26 | Q641Y2\|N | 292.16 | 35 | 35 | 1.20E+08 | 47 | 46 | 151 | Carbamidomethylation | | |
| 2 | 37 | P19234\|NI | 284.02 | 46 | 46 | 2.78E+08 | 46 | 46 | 177 | | 27378 | NADH dehydrogenase [ubiquinone] flavoprotein 2 mitochondrial OS=Rattus norvegicus OX=10116 GN=Ndufv2 PE=1 SV=2 |
| 19 | 60 | tr\|D4A565 | 283.01 | 41 | 41 | 3.38E+07 | 28 | 27 | 73 | | 21664 | NADH dehydrogenase (Ubiquinone) 1 beta subcomplex 5 (Predicted) isoform CRA_b OS=Rattus norvegicus OX=10116 GN=Ndufb5 PE=1 SV=1 |
| 17 | 161 | tr\|A0A0G2 | 255.82 | 42 | 42 | 1.24E+08 | 31 | 31 | 86 | Carbamido | 19965 | NADH dehydrogenase [ubiquinone] 1 alpha subcomplex subunit 8 OS=Rattus norvegicus OX=10116 GN=Ndufa8 PE=1 SV=1 |
| 13 | 89 | tr\|L0L4L8\| | 251.9 | 19 | 19 | | 28 | 0 | 92 | | 42470 | Cytochrome b (Fragment) OS=Rattus norvegicus OX=10116 GN=cytb PE=3 SV=1 |
| 13 | 91 | tr\|A0A140 | 251.9 | 19 | 19 | | 28 | 0 | 92 | | 42574 | Cytochrome b (Fragment) OS=Rattus norvegicus OX=10116 PE=3 SV=1 |
| 13 | 92 | tr\|A0A140 | 251.9 | 19 | 19 | | 28 | 0 | 92 | | 42588 | Cytochrome b (Fragment) OS=Rattus norvegicus OX=10116 PE=3 SV=1 |
| 13 | 95 | tr\|A0A385 | 251.9 | 19 | 19 | | 28 | 0 | 92 | | 42766 | Cytochrome b (Fragment) OS=Rattus norvegicus OX=10116 PE=3 SV=1 |
| 13 | 96 | tr\|A0A3Q8 | 251.9 | 19 | 19 | | 28 | 0 | 92 | | 42712 | Cytochrome b (Fragment) OS=Rattus norvegicus OX=10116 GN=Cytb PE=3 SV=1 |
| 13 | 97 | tr\|A0A3Q8 | 251.9 | 19 | 19 | | 28 | 0 | 92 | | 42622 | Cytochrome b (Fragment) OS=Rattus norvegicus OX=10116 GN=Cytb PE=3 SV=1 |
| 13 | 98 | tr\|A0A3S6 | 251.9 | 19 | 19 | | 28 | 0 | 92 | | 42702 | Cytochrome b (Fragment) OS=Rattus norvegicus OX=10116 GN=Cytb PE=3 SV=1 |
| 13 | 100 | tr\|A0A3S6 | 251.9 | 19 | 19 | | 28 | 0 | 92 | | 42698 | Cytochrome b (Fragment) OS=Rattus norvegicus OX=10116 GN=Cytb PE=3 SV=1 |
| 13 | 102 | tr\|A0A220 | 251.9 | 19 | 19 | | 28 | 0 | 92 | | 43018 | Cytochrome b OS=Rattus norvegicus OX=10116 PE=3 SV=1 |
| 13 | 103 | tr\|D6NSP2 | 251.9 | 19 | 19 | | 28 | 0 | 92 | | 42945 | Cytochrome b (Fragment) OS=Rattus norvegicus OX=10116 GN=cytb PE=3 SV=1 |
| 13 | 104 | tr\|D6NSS6 | 251.9 | 19 | 19 | | 28 | 0 | 92 | | 42993 | Cytochrome b (Fragment) OS=Rattus norvegicus OX=10116 GN=cytb PE=3 SV=1 |
| 13 | 106 | tr\|D6NSQ3 | 251.9 | 19 | 19 | | 28 | 0 | 92 | | 43012 | Cytochrome b (Fragment) OS=Rattus norvegicus OX=10116 GN=cytb PE=3 SV=1 |
| 13 | 107 | tr\|D6NSR3 | 251.9 | 19 | 19 | | 28 | 0 | 92 | | 42986 | Cytochrome b (Fragment) OS=Rattus norvegicus OX=10116 GN=cytb PE=3 SV=1 |
| 13 | 109 | tr\|D6NSQ8 | 251.9 | 19 | 19 | | 28 | 0 | 92 | | 43016 | Cytochrome b (Fragment) OS=Rattus norvegicus OX=10116 GN=cytb PE=3 SV=1 |
| 13 | 110 | tr\|D6NSR8 | 251.9 | 19 | 19 | | 28 | 0 | 92 | | 42998 | Cytochrome b (Fragment) OS=Rattus norvegicus OX=10116 GN=cytb PE=3 SV=1 |
| 13 | 112 | tr\|F2Q6S6 | 251.9 | 19 | 19 | | 28 | 0 | 92 | | 42938 | Cytochrome b (Fragment) OS=Rattus norvegicus OX=10116 GN=cytb PE=3 SV=1 |
| 13 | 113 | tr\|A0A0A1 | 251.9 | 19 | 19 | | 28 | 0 | 92 | | 42989 | Cytochrome b OS=Rattus norvegicus OX=10116 GN=CYTB PE=3 SV=1 |
| 13 | 114 | tr\|D6NSS7 | 251.9 | 19 | 19 | | 28 | 0 | 92 | | 42948 | Cytochrome b (Fragment) OS=Rattus norvegicus OX=10116 GN=cytb PE=3 SV=1 |
| 13 | 115 | tr\|Q8SEY9 | 251.9 | 19 | 19 | | 28 | 0 | 92 | | 43015 | Cytochrome b OS=Rattus norvegicus OX=10116 GN=cytb PE=3 SV=1 |
| 13 | 116 | tr\|D6NSR7 | 251.9 | 19 | 19 | | 28 | 0 | 92 | | 42982 | Cytochrome b (Fragment) OS=Rattus norvegicus OX=10116 GN=cytb PE=3 SV=1 |
| 13 | 117 | tr\|A0A220 | 251.9 | 19 | 19 | | 28 | 0 | 92 | | 42968 | Cytochrome b OS=Rattus norvegicus OX=10116 PE=3 SV=1 |
| 13 | 119 | tr\|D6NSQ6 | 251.9 | 19 | 19 | | 28 | 0 | 92 | | 42952 | Cytochrome b (Fragment) OS=Rattus norvegicus OX=10116 GN=cytb PE=3 SV=1 |
| 13 | 120 | tr\|D6NSQ0 | 251.9 | 19 | 19 | | 28 | 0 | 92 | | 42968 | Cytochrome b (Fragment) OS=Rattus norvegicus OX=10116 GN=cytb PE=3 SV=1 |
| 13 | 121 | tr\|Q5UAI7 | 251.9 | 19 | 19 | | 28 | 0 | 92 | | 42998 | Cytochrome b OS=Rattus norvegicus OX=10116 GN=CYTB PE=3 SV=1 |
| 13 | 122 | tr\|D6NSR2 | 251.9 | 19 | 19 | | 28 | 0 | 92 | | 43073 | Cytochrome b (Fragment) OS=Rattus norvegicus OX=10116 GN=cytb PE=3 SV=1 |
| 13 | 123 | tr\|A0A0S1 | 251.9 | 19 | 19 | | 28 | 0 | 92 | | 43002 | Cytochrome b OS=Rattus norvegicus OX=10116 GN=CYTB PE=3 SV=1 |
| 13 | 125 | tr\|Q8HIC4 | 251.9 | 19 | 19 | | 28 | 0 | 92 | | 43012 | Cytochrome b OS=Rattus norvegicus OX=10116 GN=Mt-cyb PE=3 SV=1 |
| 13 | 126 | tr\|A0A220 | 251.9 | 19 | 19 | | 28 | 0 | 92 | | 42968 | Cytochrome b OS=Rattus norvegicus OX=10116 PE=3 SV=1 |
| 13 | 128 | tr\|A0A0S1 | 251.9 | 19 | 19 | | 28 | 0 | 92 | | 43016 | Cytochrome b OS=Rattus norvegicus OX=10116 GN=CYTB PE=3 SV=1 |
| 13 | 129 | tr\|R9TKN1 | 251.9 | 19 | 19 | | 28 | 0 | 92 | | 43012 | Cytochrome b OS=Rattus norvegicus OX=10116 GN=CYTB PE=3 SV=1 |
| 13 | 130 | P00159\|CY | 251.9 | 19 | 19 | | 28 | 0 | 92 | | 43012 | Cytochrome b OS=Rattus norvegicus OX=10116 GN=Mt-Cyb PE=3 SV=3 |
| 13 | 131 | tr\|L0N311 | 251.9 | 19 | 19 | | 28 | 0 | 92 | | 43045 | Cytochrome b (Fragment) OS=Rattus norvegicus OX=10116 GN=cytb PE=3 SV=1 |
| 13 | 132 | tr\|A0A220 | 251.9 | 19 | 19 | | 28 | 0 | 92 | | 42970 | Cytochrome b OS=Rattus norvegicus OX=10116 PE=3 SV=1 |
| 13 | 133 | tr\|H2KXA0 | 251.9 | 19 | 19 | | 28 | 0 | 92 | | 42978 | Cytochrome b (Fragment) OS=Rattus norvegicus OX=10116 GN=cytb PE=3 SV=1 |
| 13 | 134 | tr\|A0A096 | 251.9 | 19 | 19 | | 28 | 0 | 92 | | 43042 | Cytochrome b (Fragment) OS=Rattus norvegicus OX=10116 PE=3 SV=1 |
| 13 | 135 | tr\|D6NSP5 | 251.9 | 19 | 19 | | 28 | 0 | 92 | | 43012 | Cytochrome b (Fragment) OS=Rattus norvegicus OX=10116 GN=cytb PE=3 SV=1 |
| 13 | 86 | tr\|A0A411 | 251.9 | 20 | 20 | | 28 | 0 | 92 | | 41017 | Cytochrome b (Fragment) OS=Rattus norvegicus OX=10116 GN=Cytb PE=3 SV=1 |
| 13 | 88 | tr\|A0A411 | 251.9 | 20 | 20 | | 28 | 0 | 92 | | 42238 | Cytochrome b (Fragment) OS=Rattus norvegicus OX=10116 GN=Cytb PE=3 SV=1 |
| 13 | 108 | tr\|D6NSP4 | 251.9 | 19 | 19 | | 28 | 0 | 92 | | 43016 | Cytochrome b (Fragment) OS=Rattus norvegicus OX=10116 GN=cytb PE=3 SV=1 |
| 13 | 124 | tr\|D6NSR1 | 251.9 | 19 | 19 | | 28 | 0 | 92 | | 43002 | Cytochrome b (Fragment) OS=Rattus norvegicus OX=10116 GN=cytb PE=3 SV=1 |
| 14 | 101 | tr\|A0A097 | 251.81 | 19 | 19 | 2.92E+05 | 28 | 1 | 92 | | 42865 | Cytochrome b OS=Rattus norvegicus OX=10116 GN=CYTB PE=3 SV=1 |
| 6 | 25 | Q68FY0\|Q | 250 | 27 | 27 | 5.97E+07 | 44 | 43 | 129 | Carbamido | 52849 | Cytochrome b-c1 complex subunit 1 mitochondrial OS=Rattus norvegicus OX=10116 GN=Uqcrc1 PE=1 SV=1 |
| 16 | 99 | tr\|A0A385 | 248.84 | 20 | 20 | 1.11E+06 | 29 | 2 | 91 | | 42751 | Cytochrome b (Fragment) OS=Rattus norvegicus OX=10116 PE=3 SV=1 |
| 32 | 248 | tr\|A9UMW | 243.75 | 37 | 37 | 3.44E+07 | 12 | 12 | 39 | Oxidation (M) | | |
| 32 | 249 | tr\|M0RB63 | 243.75 | 37 | 37 | 3.44E+07 | 12 | 12 | 39 | Oxidation (M) | | |
| 32 | 250 | tr\|A0A0G2 | 243.75 | 34 | 34 | 3.44E+07 | 12 | 12 | 39 | Oxidation (M) | | |
| 10 | 77 | tr\|Q5PQZ9 | 228.73 | 48 | 48 | 6.71E+07 | 25 | 25 | 92 | | 14359 | NADH dehydrogenase [ubiquinone] 1 subunit C2 OS=Rattus norvegicus OX=10116 GN=Ndufc2 PE=1 SV=1 |
| 22 | 137 | tr\|Q06QK5 | 219.83 | 15 | 15 | 3.34E+07 | 25 | 25 | 67 | Oxidation ( | 68606 | NADH-ubiquinone oxidoreductase chain 5 OS=Rattus norvegicus OX=10116 GN=ND5 PE=3 SV=1 |
| 22 | 138 | P11661\|NU | 219.83 | 15 | 15 | 3.34E+07 | 25 | 25 | 67 | Oxidation ( | 68618 | NADH-ubiquinone oxidoreductase chain 5 OS=Rattus norvegicus OX=10116 GN=Mtnd5 PE=3 SV=3 |
| 22 | 139 | tr\|Q06QG6 | 219.83 | 15 | 15 | 3.34E+07 | 25 | 25 | 67 | Oxidation ( | 68588 | NADH-ubiquinone oxidoreductase chain 5 OS=Rattus norvegicus OX=10116 GN=ND5 PE=3 SV=1 |
| 22 | 140 | tr\|Q06QA1 | 219.83 | 15 | 15 | 3.34E+07 | 25 | 25 | 67 | Oxidation ( | 68574 | NADH-ubiquinone oxidoreductase chain 5 OS=Rattus norvegicus OX=10116 GN=ND5 PE=3 SV=1 |
| 22 | 141 | tr\|Q8SEZ0 | 219.83 | 15 | 15 | 3.34E+07 | 25 | 25 | 67 | Oxidation ( | 68618 | NADH-ubiquinone oxidoreductase chain 5 OS=Rattus norvegicus OX=10116 GN=Mt-nd5 PE=3 SV=1 |

| | | | | | | | | | | | | |
|---|---|---|---|---|---|---|---|---|---|---|---|---|
| 22 | 142 | tr\|A0A096 | 219.83 | 15 | 15 | 3.34E+07 | 25 | 25 | 67 | Oxidation ( | 68584 | NADH-ubiquinone oxidoreductase chain 5 OS=Rattus norvegicus OX=10116 GN=ND5 PE=3 SV=1 |
| 3 | 50 | P32551\|Q | 219.65 | 25 | 25 | 1.86E+08 | 40 | 40 | 170 | Oxidation ( | 48396 | Cytochrome b-c1 complex subunit 2 mitochondrial OS=Rattus norvegicus OX=10116 GN=Uqcrc2 PE=1 SV=2 |
| 12 | 49 | Q5BK63\|N | 215.14 | 34 | 34 | 2.80E+07 | 30 | 30 | 92 | | 42559 | NADH dehydrogenase [ubiquinone] 1 alpha subcomplex subunit 9 mitochondrial OS=Rattus norvegicus OX=10116 GN=Ndufa9 PE=1 SV=2 |
| 25 | 160 | tr\|D4A7L4 | 214.5 | 26 | 26 | 4.59E+07 | 14 | 14 | 54 | | 17634 | NADH dehydrogenase (Ubiquinone) 1 beta subcomplex 11 (Predicted) OS=Rattus norvegicus OX=10116 GN=Ndufb11 PE=1 SV=1 |
| 11 | 171 | Q561S0\|N | 214.48 | 35 | 35 | 2.19E+07 | 27 | 8 | 92 | Oxidation ( | 40493 | NADH dehydrogenase [ubiquinone] 1 alpha subcomplex subunit 10 mitochondrial OS=Rattus norvegicus OX=10116 GN=Ndufa10 PE=1 SV=1 |
| 27 | 252 | tr\|D3ZF13 | 212.98 | 26 | 26 | 3.15E+07 | 17 | 17 | 49 | Oxidation ( | 17514 | Acyl carrier protein OS=Rattus norvegicus OX=10116 GN=Ndufab1 PE=1 SV=1 |
| 24 | 211 | tr\|F1LXA0\| | 209.98 | 43 | 43 | 3.43E+07 | 14 | 13 | 54 | Oxidation ( | 17178 | NADH dehydrogenase [ubiquinone] 1 alpha subcomplex subunit 12 OS=Rattus norvegicus OX=10116 GN=Ndufa12 PE=1 SV=2 |
| 21 | 8 | P15999\|AT | 205.59 | 23 | 23 | 3.41E+07 | 25 | 25 | 67 | Oxidation ( | 59754 | ATP synthase subunit alpha mitochondrial OS=Rattus norvegicus OX=10116 GN=Atp5f1a PE=1 SV=2 |
| 21 | 9 | tr\|F1LP05\| | 205.59 | 23 | 23 | 3.41E+07 | 25 | 25 | 67 | Oxidation ( | 59813 | ATP synthase subunit alpha OS=Rattus norvegicus OX=10116 GN=Atp5f1a PE=1 SV=1 |
| 23 | 172 | tr\|D4A0T0 | 203.64 | 34 | 34 | 4.56E+07 | 18 | 18 | 65 | | 20859 | NADH:ubiquinone oxidoreductase subunit B10 OS=Rattus norvegicus OX=10116 GN=Ndufb10 PE=1 SV=1 |
| 18 | 313 | tr\|D3ZZ21 | 203.58 | 53 | 53 | 3.24E+07 | 19 | 19 | 75 | Oxidation ( | 15638 | NADH dehydrogenase (Ubiquinone) 1 beta subcomplex 6 (Predicted) OS=Rattus norvegicus OX=10116 GN=Ndufb6 PE=1 SV=1 |
| 31 | 198 | Q80W89\|N | 202.22 | 24 | 24 | 4.86E+07 | 12 | 12 | 41 | | 14854 | NADH dehydrogenase [ubiquinone] 1 alpha subcomplex subunit 11 OS=Rattus norvegicus OX=10116 GN=Ndufa11 PE=2 SV=1 |
| 9 | 74 | tr\|D3ZG43 | 195.58 | 42 | 42 | 6.68E+07 | 28 | 28 | 94 | | 30226 | NADH dehydrogenase (Ubiquinone) Fe-S protein 3 (Predicted) isoform CRA_c OS=Rattus norvegicus OX=10116 GN=Ndufs3 PE=1 SV=1 |
| 35 | 318 | Q5M9I5\|Q | 194.3 | 21 | 21 | 4.08E+07 | 11 | 11 | 38 | | 10424 | Cytochrome b-c1 complex subunit 6 mitochondrial OS=Rattus norvegicus OX=10116 GN=Uqcrh PE=3 SV=1 |
| 28 | 199 | tr\|B0BNE6 | 188.75 | 25 | 25 | 2.62E+07 | 16 | 16 | 49 | | 23970 | NADH dehydrogenase (Ubiquinone) Fe-S protein 8 (Predicted) isoform CRA_a OS=Rattus norvegicus OX=10116 GN=Ndufs8 PE=1 SV=1 |
| 26 | 274 | tr\|A0A1W | 186.16 | 31 | 31 | 1.38E+05 | 21 | 2 | 49 | Oxidation ( | 40544 | NADH dehydrogenase [ubiquinone] 1 alpha subcomplex subunit 10 mitochondrial OS=Rattus norvegicus OX=10116 GN=Ndufa10l1 PE=3 SV=1 |
| 34 | 219 | tr\|D3ZS58 | 182.98 | 24 | 24 | 2.27E+07 | 12 | 12 | 38 | | 10845 | NADH dehydrogenase [ubiquinone] 1 alpha subcomplex subunit 2 OS=Rattus norvegicus OX=10116 GN=Ndufa2 PE=1 SV=1 |
| 29 | 827 | tr\|D4A3V2 | 182.83 | 42 | 42 | 1.04E+07 | 20 | 20 | 43 | | 15224 | NADH dehydrogenase [ubiquinone] 1 alpha subcomplex subunit 6 OS=Rattus norvegicus OX=10116 GN=Ndufa6 PE=1 SV=1 |
| 40 | 16 | P02563\|M | 179.18 | 6 | 6 | 6.87E+06 | 16 | 16 | 29 | Oxidation ( | 223506 | Myosin-6 OS=Rattus norvegicus OX=10116 GN=Myh6 PE=1 SV=2 |
| 50 | 65 | P31399\|AT | 176.04 | 30 | 30 | 3.80E+06 | 8 | 8 | 17 | | 18763 | ATP synthase subunit d mitochondrial OS=Rattus norvegicus OX=10116 GN=Atp5pd PE=1 SV=3 |
| 33 | 56 | tr\|F1LPG5 | 170.71 | 31 | 31 | 1.95E+07 | 12 | 12 | 38 | | 15064 | NADH:ubiquinone oxidoreductase subunit B4 OS=Rattus norvegicus OX=10116 GN=Ndufb4 PE=1 SV=1 |
| 33 | 57 | tr\|F1M7T1 | 170.71 | 31 | 31 | 1.95E+07 | 12 | 12 | 38 | | 15160 | NADH dehydrogenase (ubiquinone) 1 beta subcomplex 4 OS=Rattus norvegicus OX=10116 GN=LOC100361934 PE=4 SV=2 |
| 20 | 80 | tr\|Q5XIH3 | 169.88 | 20 | 20 | 3.82E+07 | 22 | 22 | 70 | Oxidation ( | 50731 | NADH dehydrogenase [ubiquinone] flavoprotein 1 mitochondrial OS=Rattus norvegicus OX=10116 GN=Ndufv1 PE=1 SV=1 |
| 37 | 169 | tr\|A9UMV | 164.05 | 30 | 30 | 7.83E+07 | 9 | 9 | 35 | Oxidation ( | 12500 | NADH:ubiquinone oxidoreductase subunit A7 OS=Rattus norvegicus OX=10116 GN=Ndufa7 PE=1 SV=1 |
| 44 | 45 | P00507\|A | 160.95 | 16 | 16 | 4.41E+06 | 10 | 10 | 21 | Oxidation ( | 47314 | Aspartate aminotransferase mitochondrial OS=Rattus norvegicus OX=10116 GN=Got2 PE=1 SV=2 |
| 36 | 72 | tr\|D3ZE15 | 149.38 | 35 | 35 | 1.30E+07 | 10 | 10 | 35 | Oxidation ( | 16777 | NADH:ubiquinone oxidoreductase subunit A13 OS=Rattus norvegicus OX=10116 GN=Ndufa13 PE=1 SV=1 |
| 39 | 238 | Q5XIF3\|ND | 146.45 | 20 | 20 | 1.37E+07 | 8 | 8 | 30 | Oxidation ( | 19741 | NADH dehydrogenase [ubiquinone] iron-sulfur protein 4 mitochondrial OS=Rattus norvegicus OX=10116 GN=Ndufs4 PE=1 SV=1 |
| 38 | 73 | Q63362\|N | 141 | 48 | 48 | 6.18E+06 | 15 | 12 | 30 | Carbamido | 13412 | NADH dehydrogenase [ubiquinone] 1 alpha subcomplex subunit 5 OS=Rattus norvegicus OX=10116 GN=Ndufa5 PE=1 SV=3 |
| 41 | 3899 | tr\|B5DEL8 | 138.55 | 30 | 30 | 1.18E+07 | 9 | 9 | 28 | | 12700 | NADH dehydrogenase (Ubiquinone) Fe-S protein 5 OS=Rattus norvegicus OX=10116 GN=Ndufs5 PE=1 SV=1 |
| 41 | 3900 | tr\|A0A0G2 | 138.55 | 30 | 30 | 1.18E+07 | 9 | 9 | 28 | | 12730 | Uncharacterized protein OS=Rattus norvegicus OX=10116 PE=4 SV=1 |
| 59 | 251 | P80432\|C | 135.09 | 21 | 21 | 3.18E+06 | 4 | 4 | 10 | | 7375 | Cytochrome c oxidase subunit 7C mitochondrial OS=Rattus norvegicus OX=10116 GN=Cox7c PE=1 SV=2 |
| 30 | 3918 | tr\|D3ZLT1\| | 131.42 | 21 | 21 | 2.01E+07 | 6 | 6 | 42 | Oxidation ( | 16568 | NADH dehydrogenase (Ubiquinone) 1 beta subcomplex 7 (Predicted) OS=Rattus norvegicus OX=10116 GN=Ndufb7 PE=1 SV=1 |
| 64 | 14 | Q6AXV4\|S | 126.2 | 9 | 9 | 8.13E+05 | 4 | 4 | 7 | | 51960 | Sorting and assembly machinery component 50 homolog OS=Rattus norvegicus OX=10116 GN=Samm50 PE=1 SV=1 |
| 58 | 200 | P11240\|C | 118.7 | 27 | 27 | 1.95E+06 | 6 | 6 | 10 | | 16130 | Cytochrome c oxidase subunit 5A mitochondrial OS=Rattus norvegicus OX=10116 GN=Cox5a PE=1 SV=1 |
| 60 | 61 | Q63704\|C | 115.68 | 3 | 3 | 4.88E+05 | 3 | 3 | 9 | | 88217 | Carnitine O-palmitoyltransferase 1 muscle isoform OS=Rattus norvegicus OX=10116 GN=Cpt1b PE=1 SV=1 |
| 46 | 3911 | tr\|B2RYW | 112.45 | 29 | 29 | 5.52E+06 | 7 | 7 | 19 | Oxidation ( | 21892 | NADH dehydrogenase (Ubiquinone) 1 beta subcomplex 9 OS=Rattus norvegicus OX=10116 GN=Ndufb9 PE=1 SV=1 |
| 52 | 28 | tr\|Q5UAJ6 | 110.7 | 23 | 23 | 2.20E+06 | 7 | 7 | 14 | Oxidation ( | 25942 | Cytochrome c oxidase subunit 2 OS=Rattus norvegicus OX=10116 GN=COX2 PE=3 SV=1 |
| 52 | 33 | tr\|S5RZM8 | 110.7 | 23 | 23 | 2.20E+06 | 7 | 7 | 14 | Oxidation ( | 25958 | Cytochrome c oxidase subunit 2 OS=Rattus norvegicus OX=10116 GN=COX2 PE=3 SV=1 |
| 52 | 34 | P00406\|C | 110.7 | 23 | 23 | 2.20E+06 | 7 | 7 | 14 | Oxidation ( | 25928 | Cytochrome c oxidase subunit 2 OS=Rattus norvegicus OX=10116 GN=Mtco2 PE=1 SV=3 |
| 52 | 35 | tr\|Q8SEZ5 | 110.7 | 23 | 23 | 2.20E+06 | 7 | 7 | 14 | Oxidation ( | 25928 | Cytochrome c oxidase subunit 2 OS=Rattus norvegicus OX=10116 GN=Mt-co2 PE=1 SV=1 |
| 52 | 36 | tr\|A0A097 | 110.7 | 23 | 23 | 2.20E+06 | 7 | 7 | 14 | Oxidation ( | 25894 | Cytochrome c oxidase subunit 2 OS=Rattus norvegicus OX=10116 GN=COX2 PE=3 SV=1 |
| 45 | 3893 | tr\|B2RYS2 | 109.49 | 30 | 30 | 8.80E+06 | 7 | 7 | 20 | | 13559 | Cytochrome b-c1 complex subunit 7 OS=Rattus norvegicus OX=10116 GN=Uqcrb PE=1 SV=1 |
| 57 | 253 | P10817\|C | 108.57 | 23 | 23 | 4.97E+06 | 5 | 5 | 10 | | 10487 | Cytochrome c oxidase subunit 6A2 mitochondrial (Fragment) OS=Rattus norvegicus OX=10116 GN=Cox6a2 PE=1 SV=3 |
| 57 | 254 | tr\|G3V8M | 108.57 | 23 | 23 | 4.97E+06 | 5 | 5 | 10 | | 10802 | Cytochrome c oxidase subunit 6A mitochondrial OS=Rattus norvegicus OX=10116 GN=Cox6a2 PE=3 SV=1 |
| 53 | 168 | P19511\|AT | 108.29 | 11 | 11 | 8.23E+06 | 6 | 6 | 14 | | 28869 | ATP synthase F(0) complex subunit B1 mitochondrial OS=Rattus norvegicus OX=10116 GN=Atp5pb PE=1 SV=1 |
| 47 | 3906 | tr\|D3ZCZ9 | 107.61 | 28 | 28 | 5.52E+06 | 6 | 6 | 19 | | 13040 | NADH dehydrogenase [ubiquinone] iron-sulfur protein 6 mitochondrial OS=Rattus norvegicus OX=10116 GN=LOC100912599 PE=1 SV=1 |
| 49 | 3914 | tr\|B2RYU0 | 105.16 | 21 | 21 | 1.08E+07 | 5 | 5 | 18 | | 11842 | NADH dehydrogenase (Ubiquinone) 1 beta subcomplex 2 (Predicted) isoform CRA_b OS=Rattus norvegicus OX=10116 GN=Ndufb2 PE=1 SV=1 |
| 48 | 376 | tr\|Q5RJN0 | 100.49 | 18 | 18 | 9.10E+06 | 5 | 5 | 18 | Carbamidomethylation | | |
| 62 | 3930 | tr\|Q35733 | 97.8 | 14 | 14 | 2.73E+06 | 3 | 3 | 9 | | 8389 | NADH-ubiquinone oxidoreductase chain 6 (Fragment) OS=Rattus norvegicus OX=10116 PE=3 SV=1 |
| 62 | 3921 | tr\|Q06QD | 97.8 | 6 | 6 | 2.73E+06 | 3 | 3 | 9 | | 18943 | NADH-ubiquinone oxidoreductase chain 6 OS=Rattus norvegicus OX=10116 GN=ND6 PE=3 SV=1 |
| 62 | 3922 | tr\|Q7HKW | 97.8 | 6 | 6 | 2.73E+06 | 3 | 3 | 9 | | 18957 | NADH-ubiquinone oxidoreductase chain 6 OS=Rattus norvegicus OX=10116 GN=NADH6 PE=3 SV=1 |
| 51 | 385 | tr\|A0A0A1 | 97.25 | 13 | 13 | 2.79E+06 | 7 | 7 | 16 | Oxidation ( | 38542 | NADH-ubiquinone oxidoreductase chain 2 OS=Rattus norvegicus OX=10116 GN=ND2 PE=3 SV=1 |
| 51 | 384 | tr\|Q06Q97 | 97.25 | 13 | 13 | 2.79E+06 | 7 | 7 | 16 | Oxidation ( | 38485 | NADH-ubiquinone oxidoreductase chain 2 OS=Rattus norvegicus OX=10116 GN=ND2 PE=3 SV=1 |
| 51 | 386 | tr\|Q5UAJ8 | 97.25 | 13 | 13 | 2.79E+06 | 7 | 7 | 16 | Oxidation ( | 38455 | NADH-ubiquinone oxidoreductase chain 2 OS=Rattus norvegicus OX=10116 GN=ND2 PE=3 SV=1 |
| 51 | 388 | tr\|Q8HID0 | 97.25 | 13 | 13 | 2.79E+06 | 7 | 7 | 16 | Oxidation ( | 38653 | NADH-ubiquinone oxidoreductase chain 2 OS=Rattus norvegicus OX=10116 GN=Mt-nd2 PE=3 SV=1 |
| 51 | 389 | P11662\|N | 97.25 | 13 | 13 | 2.79E+06 | 7 | 7 | 16 | Oxidation ( | 38653 | NADH-ubiquinone oxidoreductase chain 2 OS=Rattus norvegicus OX=10116 GN=Mtnd2 PE=3 SV=3 |
| 51 | 390 | tr\|Q06QH | 97.25 | 13 | 13 | 2.79E+06 | 7 | 7 | 16 | Oxidation ( | 38626 | NADH-ubiquinone oxidoreductase chain 2 OS=Rattus norvegicus OX=10116 GN=ND2 PE=3 SV=1 |
| 51 | 391 | tr\|Q8SEZ7 | 97.25 | 13 | 13 | 2.79E+06 | 7 | 7 | 16 | Oxidation ( | 38580 | NADH-ubiquinone oxidoreductase chain 2 (Fragment) OS=Rattus norvegicus OX=10116 GN=NADH2 PE=3 SV=1 |
| 51 | 392 | tr\|Q06QB | 97.25 | 13 | 13 | 2.79E+06 | 7 | 7 | 16 | Oxidation ( | 38534 | NADH-ubiquinone oxidoreductase chain 2 OS=Rattus norvegicus OX=10116 GN=ND2 PE=3 SV=1 |
| 51 | 393 | tr\|D2E6P4 | 97.25 | 13 | 13 | 2.79E+06 | 7 | 7 | 16 | Oxidation ( | 38623 | NADH-ubiquinone oxidoreductase chain 2 OS=Rattus norvegicus OX=10116 GN=ND2 PE=3 SV=1 |
| 51 | 394 | tr\|Q06QE | 97.25 | 13 | 13 | 2.79E+06 | 7 | 7 | 16 | Oxidation ( | 38598 | NADH-ubiquinone oxidoreductase chain 2 OS=Rattus norvegicus OX=10116 GN=ND2 PE=3 SV=1 |
| 42 | 208 | Q7TQ16\|Q | 96.03 | 37 | 37 | 1.31E+07 | 9 | 9 | 27 | | 9849 | Cytochrome b-c1 complex subunit 8 OS=Rattus norvegicus OX=10116 GN=Uqcrq PE=3 SV=1 |
| 56 | 63 | P35435\|AT | 95.49 | 15 | 15 | 1.40E+06 | 6 | 6 | 11 | | 30191 | ATP synthase subunit gamma mitochondrial OS=Rattus norvegicus OX=10116 GN=Atp5f1c PE=1 SV=2 |

| | | | | | | | | | | | | |
|---|---|---|---|---|---|---|---|---|---|---|---|---|
| 56 | 64 | tr\|Q6QI09 | 95.49 | 7 | 7 | 1.40E+06 | 6 | 6 | 11 | | 67721 | ATP synthase subunit gamma mitochondrial OS=Rattus norvegicus OX=10116 GN=Taf3 PE=1 SV=1 |
| 54 | 363 | P05508\|NU | 95.13 | 11 | 11 | 2.91E+06 | 8 | 8 | 12 | Oxidation (M) | | |
| 54 | 364 | tr\|D2E6K0 | 95.13 | 11 | 11 | 2.91E+06 | 8 | 8 | 12 | Oxidation (M) | | |
| 54 | 365 | tr\|A7XYB9 | 95.13 | 11 | 11 | 2.91E+06 | 8 | 8 | 12 | Oxidation (M) | | |
| 54 | 366 | tr\|Q06QE1 | 95.13 | 11 | 11 | 2.91E+06 | 8 | 8 | 12 | Oxidation (M) | | |
| 54 | 367 | tr\|Q06QA2 | 95.13 | 11 | 11 | 2.91E+06 | 8 | 8 | 12 | Oxidation (M) | | |
| 54 | 368 | tr\|Q7HKW | 95.13 | 11 | 11 | 2.91E+06 | 8 | 8 | 12 | Oxidation (M) | | |
| 54 | 369 | tr\|Q06QG7 | 95.13 | 11 | 11 | 2.91E+06 | 8 | 8 | 12 | Oxidation (M) | | |
| 54 | 370 | tr\|Q06Q89 | 95.13 | 11 | 11 | 2.91E+06 | 8 | 8 | 12 | Oxidation (M) | | |
| 54 | 371 | tr\|Q35737 | 95.13 | 11 | 11 | 2.91E+06 | 8 | 8 | 12 | Oxidation (M) | | |
| 54 | 372 | tr\|Q8HIC6 | 95.13 | 11 | 11 | 2.91E+06 | 8 | 8 | 12 | Oxidation (M) | | |
| 43 | 3901 | tr\|Q8SEZ8 | 91.15 | 10 | 10 | 8.69E+06 | 9 | 9 | 22 | Oxidation ( | 36133 | NADH-ubiquinone oxidoreductase chain 1 (Fragment) OS=Rattus norvegicus OX=10116 GN=NADH1 PE=3 SV=1 |
| 43 | 3902 | P03889\|NU | 91.15 | 10 | 10 | 8.69E+06 | 9 | 9 | 22 | Oxidation ( | 36145 | NADH-ubiquinone oxidoreductase chain 1 OS=Rattus norvegicus OX=10116 GN=Mtnd1 PE=1 SV=3 |
| 43 | 3903 | tr\|Q8HID1 | 91.15 | 10 | 10 | 8.69E+06 | 9 | 9 | 22 | Oxidation ( | 36145 | NADH-ubiquinone oxidoreductase chain 1 OS=Rattus norvegicus OX=10116 GN=Mt-nd1 PE=3 SV=1 |
| 43 | 3904 | tr\|D2E6L7 | 91.15 | 10 | 10 | 8.69E+06 | 9 | 9 | 22 | Oxidation ( | 36062 | NADH-ubiquinone oxidoreductase chain 1 OS=Rattus norvegicus OX=10116 GN=ND1 PE=3 SV=1 |
| 76 | 17 | Q9ER34\|A | 87.7 | 3 | 3 | 2.62E+05 | 3 | 3 | 4 | | 85433 | Aconitate hydratase mitochondrial OS=Rattus norvegicus OX=10116 GN=Aco2 PE=1 SV=2 |
| 68 | 3924 | Q6PCU8\|N | 84.04 | 19 | 19 | 1.43E+06 | 3 | 3 | 5 | | 11942 | NADH dehydrogenase [ubiquinone] flavoprotein 3 mitochondrial OS=Rattus norvegicus OX=10116 GN=Ndufv3 PE=3 SV=1 |
| 55 | 467 | tr\|D4A4P3 | 77.45 | 30 | 30 | 4.73E+06 | 5 | 5 | 12 | Oxidation ( | 11267 | NADH:ubiquinone oxidoreductase subunit B3 OS=Rattus norvegicus OX=10116 GN=Ndufb3 PE=1 SV=1 |
| 72 | 76 | Q06647\|AT | 76.54 | 8 | 8 | 7.44E+05 | 2 | 2 | 4 | | 23398 | ATP synthase subunit O mitochondrial OS=Rattus norvegicus OX=10116 GN=Atp5po PE=1 SV=1 |
| 77 | 245 | tr\|C8CHS6 | 75.85 | 15 | 15 | 7.12E+05 | 1 | 1 | 4 | | 7988 | Mitochondrial superoxide dismutase 2 (Fragment) OS=Rattus norvegicus OX=10116 PE=2 SV=1 |
| 77 | 71 | P07895\|SC | 75.85 | 5 | 5 | 7.12E+05 | 1 | 1 | 4 | | 24674 | Superoxide dismutase [Mn] mitochondrial OS=Rattus norvegicus OX=10116 GN=Sod2 PE=1 SV=2 |
| 61 | 334 | tr\|A0A0G2 | 72.38 | 6 | 6 | 2.66E+06 | 4 | 4 | 9 | | 33168 | Prohibitin OS=Rattus norvegicus OX=10116 GN=Phb2 PE=1 SV=1 |
| 61 | 335 | Q5XIH7\|PH | 72.38 | 6 | 6 | 2.66E+06 | 4 | 4 | 9 | | 33312 | Prohibitin-2 OS=Rattus norvegicus OX=10116 GN=Phb2 PE=1 SV=1 |
| 65 | 21 | Q64428\|E | 71.7 | 4 | 4 | 6.00E+05 | 3 | 3 | 6 | | 82665 | Trifunctional enzyme subunit alpha mitochondrial OS=Rattus norvegicus OX=10116 GN=Hadha PE=1 SV=2 |
| 103 | 233 | tr\|Q5UAJ5 | 69.06 | 16 | 16 | 4.68E+05 | 2 | 2 | 2 | | 7642 | ATP synthase protein 8 OS=Rattus norvegicus OX=10116 GN=ATP8 PE=3 SV=1 |
| 103 | 234 | P11608\|AT | 69.06 | 16 | 16 | 4.68E+05 | 2 | 2 | 2 | | 7630 | ATP synthase protein 8 OS=Rattus norvegicus OX=10116 GN=Mt-atp8 PE=1 SV=2 |
| 103 | 235 | tr\|Q8SEZ4 | 69.06 | 16 | 16 | 4.68E+05 | 2 | 2 | 2 | | 7632 | ATP synthase protein 8 OS=Rattus norvegicus OX=10116 GN=ATPase8 PE=3 SV=1 |
| 103 | 236 | tr\|Q8HIC8 | 69.06 | 16 | 16 | 4.68E+05 | 2 | 2 | 2 | | 7630 | ATP synthase protein 8 OS=Rattus norvegicus OX=10116 GN=Mt-atp8 PE=3 SV=1 |
| 97 | 11 | tr\|A0A140 | 67.14 | 4 | 4 | 3.02E+05 | 2 | 2 | 2 | | 67049 | MICOS complex subunit MIC60 OS=Rattus norvegicus OX=10116 GN=Immt PE=1 SV=1 |
| 97 | 12 | Q3KR86\|M | 67.14 | 4 | 4 | 3.02E+05 | 2 | 2 | 2 | | 67177 | MICOS complex subunit Mic60 (Fragment) OS=Rattus norvegicus OX=10116 GN=Immt PE=1 SV=1 |
| 97 | 10 | tr\|A0A0G2 | 67.14 | 3 | 3 | 3.02E+05 | 2 | 2 | 2 | | 86230 | MICOS complex subunit MIC60 OS=Rattus norvegicus OX=10116 GN=Immt PE=1 SV=1 |
| 67 | 40 | tr\|A0A0G2 | 66.36 | 4 | 4 | 6.54E+05 | 3 | 3 | 5 | | 52568 | Trifunctional enzyme subunit beta mitochondrial OS=Rattus norvegicus OX=10116 GN=Hadhb PE=1 SV=1 |
| 67 | 29 | Q60587\|E | 66.36 | 4 | 4 | 6.54E+05 | 3 | 3 | 5 | | 51414 | Trifunctional enzyme subunit beta mitochondrial OS=Rattus norvegicus OX=10116 GN=Hadhb PE=1 SV=1 |
| 88 | 263 | P35171\|CX | 64.71 | 12 | 12 | 3.23E+05 | 2 | 2 | 3 | | 9353 | Cytochrome c oxidase subunit 7A2 mitochondrial OS=Rattus norvegicus OX=10116 GN=Cox7a2 PE=1 SV=1 |
| 88 | 264 | tr\|B2RYS0 | 64.71 | 12 | 12 | 3.23E+05 | 2 | 2 | 3 | | 9353 | Cox7a2 protein OS=Rattus norvegicus OX=10116 GN=Cox7a2 PE=2 SV=1 |
| 73 | 1021 | tr\|Q06QA9 | 58.61 | 3 | 3 | 3.29E+05 | 2 | 2 | 4 | | 56893 | Cytochrome c oxidase subunit 1 OS=Rattus norvegicus OX=10116 GN=CO1 PE=3 SV=1 |
| 73 | 1022 | tr\|Q06QK0 | 58.61 | 3 | 3 | 3.29E+05 | 2 | 2 | 4 | | 56880 | Cytochrome c oxidase subunit 1 OS=Rattus norvegicus OX=10116 GN=CO1 PE=3 SV=1 |
| 73 | 1023 | tr\|Q8SEZ6 | 58.61 | 3 | 3 | 3.29E+05 | 2 | 2 | 4 | | 56879 | Cytochrome c oxidase subunit 1 OS=Rattus norvegicus OX=10116 GN=COI PE=3 SV=1 |
| 73 | 1024 | tr\|A0A0A1 | 58.61 | 3 | 3 | 3.29E+05 | 2 | 2 | 4 | | 56937 | Cytochrome c oxidase subunit 1 OS=Rattus norvegicus OX=10116 GN=COX1 PE=3 SV=1 |
| 73 | 1025 | tr\|Q8HIC9 | 58.61 | 3 | 3 | 3.29E+05 | 2 | 2 | 4 | | 56845 | Cytochrome c oxidase subunit 1 OS=Rattus norvegicus OX=10116 GN=Mt-co1 PE=3 SV=1 |
| 73 | 1026 | P05503\|CO | 58.61 | 3 | 3 | 3.29E+05 | 2 | 2 | 4 | | 56845 | Cytochrome c oxidase subunit 1 OS=Rattus norvegicus OX=10116 GN=Mtco1 PE=2 SV=3 |
| 73 | 1027 | tr\|Q95938 | 58.61 | 3 | 3 | 3.29E+05 | 2 | 2 | 4 | | 56977 | Cytochrome c oxidase subunit 1 OS=Rattus norvegicus OX=10116 GN=Co I PE=3 SV=1 |
| 78 | 532 | P67779\|PH | 54.81 | 6 | 6 | 2.47E+05 | 2 | 2 | 3 | | 29820 | Prohibitin OS=Rattus norvegicus OX=10116 GN=Phb PE=1 SV=1 |
| 78 | 531 | tr\|D3ZFH6 | 54.81 | 6 | 6 | 2.47E+05 | 2 | 2 | 3 | | 29594 | Prohibitin OS=Rattus norvegicus OX=10116 GN=Phb-ps1 PE=3 SV=3 |
| 85 | 209 | tr\|B2RZD6 | 51.54 | 18 | 18 | 7.55E+05 | 2 | 2 | 3 | | 9327 | NDUFA4 mitochondrial complex-associated OS=Rattus norvegicus OX=10116 GN=Ndufa4 PE=1 SV=1 |
| 63 | 474 | tr\|F1LN92 | 50.9 | 5 | 5 | 1.54E+06 | 4 | 4 | 8 | | 89352 | AFG3-like matrix AAA peptidase subunit 2 OS=Rattus norvegicus OX=10116 GN=Afg3l2 PE=1 SV=1 |
| 153 | 4 | P08461\|OD | 50.08 | 2 | 2 | 2.47E+05 | 1 | 1 | 1 | | 67166 | Dihydrolipoyllysine-residue acetyltransferase component of pyruvate dehydrogenase complex mitochondrial OS=Rattus norvegicus OX=10116 GN=Dlat PE=1 SV=3 |
| 87 | 315 | tr\|G3V7I0\| | 46.62 | 5 | 5 | 3.54E+05 | 1 | 1 | 3 | | 28299 | Peroxiredoxin 3 OS=Rattus norvegicus OX=10116 GN=Prdx3 PE=1 SV=1 |
| 87 | 316 | Q9Z0V6\|PI | 46.62 | 5 | 5 | 3.54E+05 | 1 | 1 | 3 | | 28295 | Thioredoxin-dependent peroxide reductase mitochondrial OS=Rattus norvegicus OX=10116 GN=Prdx3 PE=1 SV=2 |
| 74 | 55 | tr\|A0A1W2 | 45.4 | 8 | 8 | 3.40E+05 | 3 | 3 | 4 | | 13534 | Solute carrier family 25 member 31 (Fragment) OS=Rattus norvegicus OX=10116 GN=Slc25a31 PE=3 SV=1 |
| 74 | 23 | tr\|Q6P9Y4 | 45.4 | 3 | 3 | 3.40E+05 | 3 | 3 | 4 | | 32904 | ADP/ATP translocase 1 OS=Rattus norvegicus OX=10116 GN=Slc25a4 PE=1 SV=1 |
| 74 | 24 | Q05962\|AI | 45.4 | 3 | 3 | 3.40E+05 | 3 | 3 | 4 | | 32989 | ADP/ATP translocase 1 OS=Rattus norvegicus OX=10116 GN=Slc25a4 PE=1 SV=3 |
| 74 | 27 | Q09073\|AI | 45.4 | 3 | 3 | 3.40E+05 | 3 | 3 | 4 | | 32901 | ADP/ATP translocase 2 OS=Rattus norvegicus OX=10116 GN=Slc25a5 PE=1 SV=3 |
| 74 | 43 | tr\|D3ZB81 | 45.4 | 3 | 3 | 3.40E+05 | 3 | 3 | 4 | | 35227 | Solute carrier family 25 member 31 OS=Rattus norvegicus OX=10116 GN=Slc25a31 PE=3 SV=3 |
| 102 | 215 | P33124\|AC | 41.6 | 2 | 2 | 0 | 2 | 1 | 2 | | 78180 | Long-chain-fatty-acid--CoA ligase 6 OS=Rattus norvegicus OX=10116 GN=Acsl6 PE=1 SV=1 |
| 102 | 47 | P18163\|AC | 41.6 | 2 | 2 | 0 | 2 | 1 | 2 | | 78179 | Long-chain-fatty-acid--CoA ligase 1 OS=Rattus norvegicus OX=10116 GN=Acsl1 PE=1 SV=1 |
| 99 | 8042 | tr\|Q4G067 | 40.68 | 3 | 3 | 1.77E+05 | 1 | 1 | 2 | | 37440 | Mitochondrial ribosomal protein L44 OS=Rattus norvegicus OX=10116 GN=Mrpl44 PE=1 SV=1 |
| 104 | 3915 | tr\|Q8SEZ1 | 40.25 | 6 | 6 | 3.54E+05 | 1 | 1 | 2 | | 13087 | NADH-ubiquinone oxidoreductase chain 3 OS=Rattus norvegicus OX=10116 GN=Mt-nd3 PE=2 SV=1 |
| 104 | 3916 | P05506\|NU | 40.25 | 6 | 6 | 3.54E+05 | 1 | 1 | 2 | | 13087 | NADH-ubiquinone oxidoreductase chain 3 OS=Rattus norvegicus OX=10116 GN=Mtnd3 PE=3 SV=3 |
| 104 | 3938 | tr\|A0A0A1 | 40.25 | 6 | 6 | 3.54E+05 | 1 | 1 | 2 | | 13071 | NADH-ubiquinone oxidoreductase chain 3 OS=Rattus norvegicus OX=10116 GN=ND3 PE=3 SV=1 |
| 104 | 3939 | tr\|Q06QG9 | 40.25 | 6 | 6 | 3.54E+05 | 1 | 1 | 2 | | 13075 | NADH-ubiquinone oxidoreductase chain 3 OS=Rattus norvegicus OX=10116 GN=ND3 PE=3 SV=1 |
| 82 | 44 | P17764\|TH | 39.12 | 3 | 3 | 5.67E+05 | 1 | 1 | 3 | | 44695 | Acetyl-CoA acetyltransferase mitochondrial OS=Rattus norvegicus OX=10116 GN=Acat1 PE=1 SV=1 |
| 154 | 2 | Q01205\|O | 38.58 | 4 | 4 | 2.52E+05 | 1 | 1 | 1 | | 48925 | Dihydrolipoyllysine-residue succinyltransferase component of 2-oxoglutarate dehydrogenase complex mitochondrial OS=Rattus norvegicus OX=10116 GN=Dlst PE=1 SV=2 |

| | | | | | | | | | | | | |
|---|---|---|---|---|---|---|---|---|---|---|---|---|
| 154 | 1 | tr\|G3V6P2 | 38.58 | 4 | 4 | 2.52E+05 | 1 | 1 | 1 | | 48899 | Dihydrolipoamide S-succinyltransferase (E2 component of 2-oxo-glutarate complex) isoform CRA_a OS=Rattus norvegicus OX=10116 GN=Dlst PE=1 SV=1 |
| 71 | 75 | Q6PDU7\|A | 38.18 | 16 | 16 | 4.27E+05 | 1 | 1 | 4 | | 11433 | ATP synthase subunit g mitochondrial OS=Rattus norvegicus OX=10116 GN=Atp5mg PE=1 SV=2 |
| 79 | 4322 | tr\|A0A0G2 | 34.08 | 1 | 1 | 1.71E+05 | 2 | 1 | 3 | | 121101 | Cation-transporting ATPase OS=Rattus norvegicus OX=10116 GN=Atp13a2 PE=3 SV=1 |
| 79 | 1051 | tr\|B5DEH6 | 34.08 | 1 | 1 | 1.71E+05 | 2 | 1 | 3 | | 124814 | Cation-transporting ATPase OS=Rattus norvegicus OX=10116 GN=Atp13a2 PE=2 SV=1 |
| 79 | 1052 | tr\|F1MAA4 | 34.08 | 1 | 1 | 1.71E+05 | 2 | 1 | 3 | | 124838 | Cation-transporting ATPase OS=Rattus norvegicus OX=10116 GN=Atp13a2 PE=3 SV=2 |
| 105 | 3932 | tr\|D3ZD09 | 33.98 | 12 | 12 | 2.15E+05 | 1 | 1 | 2 | | 10071 | Cytochrome c oxidase subunit OS=Rattus norvegicus OX=10116 GN=Cox6b1 PE=1 SV=1 |
| 80 | 3936 | tr\|A0A0G2 | 32.99 | 21 | 21 | 9.28E+05 | 2 | 2 | 3 | | 7099 | Ubiquinol-cytochrome c reductase complex III subunit X OS=Rattus norvegicus OX=10116 GN=Uqcr10 PE=1 SV=1 |
| 80 | 3937 | tr\|B2RYX1 | 32.99 | 20 | 20 | 9.28E+05 | 2 | 2 | 3 | | 7462 | LOC685322 protein OS=Rattus norvegicus OX=10116 GN=Uqcr10 PE=2 SV=1 |
| 69 | 3958 | tr\|D3ZD73 | 32.09 | 2 | 2 | | 2 | 0 | 5 | | 54245 | DEAD-box helicase 6 OS=Rattus norvegicus OX=10116 GN=Ddx6 PE=1 SV=1 |
| 66 | 3919 | tr\|A9UMV | 32.02 | 11 | 11 | 8.27E+05 | 1 | 1 | 6 | | 6539 | RCG29512 OS=Rattus norvegicus OX=10116 GN=Uqcr11 PE=1 SV=1 |
| 106 | 343 | Q924S5\|L | 31.79 | 1 | 1 | 5.36E+04 | 1 | 1 | 2 | | 105792 | Lon protease homolog mitochondrial OS=Rattus norvegicus OX=10116 GN=Lonp1 PE=2 SV=1 |
| 98 | 3951 | tr\|B1WBP | 31.03 | 13 | 13 | 1.03E+05 | 2 | 2 | 2 | | 12884 | ATP synthase subunit delta mitochondrial OS=Rattus norvegicus OX=10116 GN=Atp5f1d PE=1 SV=1 |
| 98 | 3953 | tr\|G3V7Y3 | 31.03 | 10 | 10 | 1.03E+05 | 2 | 2 | 2 | | 17563 | ATP synthase subunit delta mitochondrial OS=Rattus norvegicus OX=10116 GN=Atp5f1d PE=1 SV=1 |
| 120 | 524 | tr\|D4A054 | 30.11 | 0 | 0 | 4.17E+05 | 1 | 1 | 1 | | 341403 | RAN-binding protein 2 OS=Rattus norvegicus OX=10116 GN=Ranbp2 PE=1 SV=2 |
| 120 | 525 | tr\|M0R3M | 30.11 | 0 | 0 | 4.17E+05 | 1 | 1 | 1 | | 344396 | RAN-binding protein 2 OS=Rattus norvegicus OX=10116 GN=Ranbp2 PE=1 SV=1 |
| 90 | 151 | P12075\|C | 29.69 | 9 | 9 | 1.19E+06 | 2 | 2 | 2 | | 13915 | Cytochrome c oxidase subunit 5B mitochondrial OS=Rattus norvegicus OX=10116 GN=Cox5b PE=1 SV=2 |
| 155 | 3934 | Q66H47\|R | 28.92 | 6 | 6 | 0 | 1 | 1 | 1 | | 25001 | 39S ribosomal protein L24 mitochondrial OS=Rattus norvegicus OX=10116 GN=Mrpl24 PE=2 SV=1 |
| 93 | 377 | P97521\|M | 28.13 | 4 | 4 | 1.19E+06 | 1 | 1 | 2 | Formylatio | 33154 | Mitochondrial carnitine/acylcarnitine carrier protein OS=Rattus norvegicus OX=10116 GN=Slc25a20 PE=1 SV=1 |
| 93 | 241 | tr\|Q66HP8 | 28.13 | 4 | 4 | 1.19E+06 | 1 | 1 | 2 | Formylatio | 33071 | Mitochondrial carnitine/acylcarnitine carrier protein OS=Rattus norvegicus OX=10116 GN=Slc25a20 PE=1 SV=1 |
| 107 | 292 | P10888\|C | 27.42 | 4 | 4 | 1.84E+05 | 1 | 1 | 2 | | 19515 | Cytochrome c oxidase subunit 4 isoform 1 mitochondrial OS=Rattus norvegicus OX=10116 GN=Cox4i1 PE=1 SV=1 |
| 91 | 4670 | tr\|E9PU34 | 26.64 | 3 | 3 | 3.03E+06 | 2 | 1 | 2 | Formylatio | 51699 | RCG41110 OS=Rattus norvegicus OX=10116 GN=Rmnd1 PE=1 SV=2 |
| 141 | 66 | P56574\|ID | 26.19 | 2 | 2 | 1.15E+05 | 1 | 1 | 1 | | 50967 | Isocitrate dehydrogenase [NADP] mitochondrial OS=Rattus norvegicus OX=10116 GN=Idh2 PE=1 SV=2 |
| 138 | 266 | tr\|Q7H115 | 25.86 | 3 | 3 | 0 | 1 | 1 | 1 | | 29871 | Cytochrome c oxidase subunit 3 OS=Rattus norvegicus OX=10116 GN=Mt-co3 PE=3 SV=1 |
| 138 | 267 | tr\|I6V4L9\| | 25.86 | 3 | 3 | 0 | 1 | 1 | 1 | | 29844 | Cytochrome c oxidase subunit 3 OS=Rattus norvegicus OX=10116 GN=COX3 PE=3 SV=1 |
| 138 | 269 | tr\|A0A096 | 25.86 | 3 | 3 | 0 | 1 | 1 | 1 | | 29901 | Cytochrome c oxidase subunit 3 OS=Rattus norvegicus OX=10116 GN=COX3 PE=3 SV=1 |
| 138 | 270 | P05505\|C | 25.86 | 3 | 3 | 0 | 1 | 1 | 1 | | 29871 | Cytochrome c oxidase subunit 3 OS=Rattus norvegicus OX=10116 GN=Mtco3 PE=1 SV=5 |
| 138 | 271 | tr\|Q8SEZ2 | 25.86 | 3 | 3 | 0 | 1 | 1 | 1 | | 29870 | Cytochrome c oxidase subunit 3 (Fragment) OS=Rattus norvegicus OX=10116 GN=COIII PE=3 SV=1 |
| 138 | 268 | tr\|Q8M7G | 25.86 | 3 | 3 | 0 | 1 | 1 | 1 | | 29861 | Cytochrome c oxidase subunit 3 OS=Rattus norvegicus OX=10116 PE=2 SV=1 |
| 156 | 8046 | Q5M807\|F | 25.24 | 3 | 3 | 0 | 1 | 1 | 1 | | 19837 | E3 ubiquitin-protein ligase RNF5 OS=Rattus norvegicus OX=10116 GN=Rnf5 PE=2 SV=1 |
| 109 | 3985 | tr\|Q5XFW | 24.92 | 12 | 12 | 4.59E+05 | 2 | 2 | 2 | | 20541 | Mitochondrial ribosomal protein L13 OS=Rattus norvegicus OX=10116 GN=Mrpl13 PE=1 SV=1 |
| 83 | 4091 | P0DN35\|N | 24.66 | 18 | 18 | 7.58E+05 | 1 | 1 | 3 | | 6998 | NADH dehydrogenase [ubiquinone] 1 beta subcomplex subunit 1 OS=Rattus norvegicus OX=10116 GN=Ndufb1 PE=3 SV=1 |
| 139 | 8043 | Q8K1M7\|E | 23.5 | 1 | 1 | 0 | 1 | 1 | 1 | | 88497 | BMP/retinoic acid-inducible neural-specific protein 3 OS=Rattus norvegicus OX=10116 GN=Brinp3 PE=2 SV=1 |
| 110 | 398 | tr\|B2RYT5 | 23 | 8 | 8 | 2.75E+05 | 1 | 1 | 2 | | 12651 | Cox7a2l protein OS=Rattus norvegicus OX=10116 GN=Cox7a2l PE=2 SV=1 |
| 110 | 399 | tr\|D3ZYX8 | 23 | 8 | 8 | 2.75E+05 | 1 | 1 | 2 | | 13274 | Cytochrome c oxidase subunit 7A2-like OS=Rattus norvegicus OX=10116 GN=Cox7a2l PE=1 SV=1 |
| 157 | 3917 | tr\|D3ZJG4 | 22.08 | 1 | 1 | 7.28E+05 | 1 | 1 | 1 | Oxidation ( | 95972 | Phosphofurin acidic cluster sorting protein 2 OS=Rattus norvegicus OX=10116 GN=Pacs2 PE=1 SV=2 |
| 111 | 1034 | P05696\|KF | 20.89 | 1 | 1 | 7.59E+05 | 1 | 1 | 2 | | 76792 | Protein kinase C alpha type OS=Rattus norvegicus OX=10116 GN=Prkca PE=1 SV=3 |
