## Supplementary material for "Permeability transition pore-related changes in the proteome and channel activity of ATP synthase dimers and monomers": RHM

| Protein Gr | Protein ID | Accession | -10lgP | Coverage ( | Coverage ( | IBAQ | #Peptides | #Unique | #Spec Sam | PTM | Avg. Mass | Description |
|---|---|---|---|---|---|---|---|---|---|---|---|---|
| 1 | 5 | P10719\|AT | 339.53 | 29 | 29 | 1.65E+08 | 68 | 66 | 199 | Oxidation ( | 56354 | ATP synthase subunit beta mitochondrial OS=Rattus norvegicus OX=10116 GN=Atp5f1b PE=1 SV=2 |
| 1 | 6 | tr\|G3V6D3 | 339.53 | 29 | 29 | 1.65E+08 | 68 | 66 | 199 | Oxidation ( | 56345 | ATP synthase subunit beta OS=Rattus norvegicus OX=10116 GN=Atp5f1b PE=1 SV=1 |
| 7 | 58 | P20788\|U | 253.19 | 24 | 24 | 1.47E+07 | 26 | 26 | 57 | | 29446 | Cytochrome b-c1 complex subunit Rieske mitochondrial OS=Rattus norvegicus OX=10116 GN=Uqcrfs1 PE=1 SV=2 |
| 2 | 8 | P15999\|AT | 251.13 | 36 | 36 | 6.45E+07 | 63 | 62 | 152 | Oxidation ( | 59754 | ATP synthase subunit alpha mitochondrial OS=Rattus norvegicus OX=10116 GN=Atp5f1a PE=1 SV=2 |
| 2 | 9 | tr\|F1LP05\| | 251.13 | 36 | 36 | 6.45E+07 | 63 | 62 | 152 | Oxidation ( | 59813 | ATP synthase subunit alpha OS=Rattus norvegicus OX=10116 GN=Atp5f1a PE=1 SV=1 |
| 10 | 59 | tr\|D3ZFQ8 | 249.65 | 27 | 27 | 1.52E+07 | 23 | 23 | 52 | Oxidation (M) | | Cytochrome c-1 |
| 3 | 37 | P19234\|NI | 223.99 | 33 | 33 | 2.75E+07 | 24 | 24 | 65 | | 27378 | NADH dehydrogenase [ubiquinone] flavoprotein 2 mitochondrial OS=Rattus norvegicus OX=10116 GN=Ndufv2 PE=1 SV=2 |
| 18 | 64 | tr\|Q6QI09 | 219.17 | 10 | 10 | 5.32E+06 | 19 | 19 | 34 | Oxidation ( | 67721 | ATP synthase subunit gamma mitochondrial OS=Rattus norvegicus OX=10116 GN=Taf3 PE=1 SV=1 |
| 18 | 63 | P35435\|AT | 219.17 | 22 | 22 | 5.32E+06 | 19 | 19 | 34 | Oxidation ( | 30191 | ATP synthase subunit gamma mitochondrial OS=Rattus norvegicus OX=10116 GN=Atp5f1c PE=1 SV=2 |
| 21 | 212 | tr\|B2RYS8 | 217.7 | 27 | 27 | 6.18E+06 | 16 | 16 | 30 | Oxidation ( | 21959 | NADH dehydrogenase [ubiquinone] 1 beta subcomplex subunit 8 mitochondrial OS=Rattus norvegicus OX=10116 GN=Ndufb8 PE=1 SV=1 |
| 5 | 20 | Q66HF1\|N | 216.87 | 18 | 18 | 1.24E+07 | 31 | 31 | 59 | Carbamido | 79412 | NADH-ubiquinone oxidoreductase 75 kDa subunit mitochondrial OS=Rattus norvegicus OX=10116 GN=Ndufs1 PE=1 SV=1 |
| 24 | 200 | P11240\|C | 216.63 | 39 | 39 | 3.06E+06 | 15 | 15 | 27 | | 16130 | Cytochrome c oxidase subunit 5A mitochondrial OS=Rattus norvegicus OX=10116 GN=Cox5a PE=1 SV=1 |
| 4 | 45 | P00507\|A | 215.58 | 28 | 28 | 1.38E+07 | 34 | 34 | 64 | Oxidation ( | 47314 | Aspartate aminotransferase mitochondrial OS=Rattus norvegicus OX=10116 GN=Got2 PE=1 SV=2 |
| 17 | 65 | P31399\|AT | 213.43 | 38 | 38 | 1.31E+07 | 17 | 17 | 35 | Oxidation ( | 18763 | ATP synthase subunit d mitochondrial OS=Rattus norvegicus OX=10116 GN=Atp5pd PE=1 SV=3 |
| 15 | 101 | tr\|A0A097 | 210.95 | 16 | 16 | 1.12E+06 | 18 | 2 | 42 | | 42865 | Cytochrome b OS=Rattus norvegicus OX=10116 GN=CYTB PE=3 SV=1 |
| 6 | 405 | P14882\|P | 208.33 | 24 | 24 | 9.07E+05 | 32 | 2 | 58 | Carbamidomethylatior | | Propionyl-CoA carboxylase alpha chain, mitochondrial |
| 8 | 403 | tr\|A0A0G2 | 208.06 | 23 | 23 | 2.18E+05 | 32 | 2 | 57 | Carbamidomethylatior | | Propionyl-CoA carboxylase alpha chain, mitochondrial |
| 8 | 404 | tr\|A0A0G2 | 208.06 | 22 | 22 | 2.18E+05 | 32 | 2 | 57 | Carbamidomethylation | | |
| 16 | 99 | tr\|A0A385 | 203.25 | 14 | 14 | 1.33E+05 | 17 | 1 | 38 | | 42751 | Cytochrome b (Fragment) OS=Rattus norvegicus OX=10116 PE=3 SV=1 |
| 12 | 25 | Q68FY0\|Q | 201.21 | 20 | 20 | 9.14E+06 | 24 | 24 | 45 | Carbamido | 52849 | Cytochrome b-c1 complex subunit 1 mitochondrial OS=Rattus norvegicus OX=10116 GN=Uqcrc1 PE=1 SV=1 |
| 13 | 26 | Q641Y2\|N | 192.46 | 17 | 17 | 1.24E+07 | 21 | 21 | 45 | | 52562 | NADH dehydrogenase [ubiquinone] iron-sulfur protein 2 mitochondrial OS=Rattus norvegicus OX=10116 GN=Ndufs2 PE=1 SV=1 |
| 22 | 17 | Q9ER34\|A | 186.86 | 11 | 11 | 4.61E+06 | 18 | 18 | 28 | | 85433 | Aconitate hydratase mitochondrial OS=Rattus norvegicus OX=10116 GN=Aco2 PE=1 SV=2 |
| 25 | 161 | tr\|A0A0G2 | 176.99 | 34 | 34 | 1.13E+07 | 14 | 14 | 27 | Carbamido | 19965 | NADH dehydrogenase [ubiquinone] 1 alpha subcomplex subunit 8 OS=Rattus norvegicus OX=10116 GN=Ndufa8 PE=1 SV=1 |
| 44 | 248 | tr\|A9UMW | 175.25 | 42 | 42 | 5.27E+06 | 6 | 6 | 15 | Oxidation ( | 9255 | Ndufa3 protein (Fragment) OS=Rattus norvegicus OX=10116 GN=Ndufa3 PE=2 SV=1 |
| 44 | 250 | tr\|A0A0G2 | 175.25 | 38 | 38 | 5.27E+06 | 6 | 6 | 15 | Oxidation ( | 10170 | NADH:ubiquinone oxidoreductase subunit A3 OS=Rattus norvegicus OX=10116 GN=Ndufa3 PE=1 SV=1 |
| 44 | 249 | tr\|M0RB6 | 175.25 | 42 | 42 | 5.27E+06 | 6 | 6 | 15 | Oxidation ( | 9372 | RCG63041 OS=Rattus norvegicus OX=10116 GN=LOC684509 PE=4 SV=1 |
| 48 | 211 | tr\|F1LXA0\| | 167.34 | 21 | 21 | 2.39E+06 | 9 | 9 | 14 | | 17178 | NADH dehydrogenase [ubiquinone] 1 alpha subcomplex subunit 12 OS=Rattus norvegicus OX=10116 GN=Ndufa12 PE=1 SV=2 |
| 34 | 160 | tr\|D4A7L4 | 167.09 | 23 | 23 | 5.68E+06 | 9 | 9 | 19 | | 17634 | NADH dehydrogenase (Ubiquinone) 1 beta subcomplex 11 (Predicted) OS=Rattus norvegicus OX=10116 GN=Ndufb11 PE=1 SV=1 |
| 31 | 60 | tr\|D4A565 | 165.31 | 26 | 26 | 3.17E+06 | 12 | 12 | 20 | | 21664 | NADH dehydrogenase (Ubiquinone) 1 beta subcomplex 5 (Predicted) isoform CRA_b OS=Rattus norvegicus OX=10116 GN=Ndufb5 PE=1 SV=1 |
| 37 | 251 | P80432\|C | 163.96 | 21 | 21 | 5.05E+06 | 5 | 5 | 17 | | 7375 | Cytochrome c oxidase subunit 7C mitochondrial OS=Rattus norvegicus OX=10116 GN=Cox7c PE=1 SV=2 |
| 11 | 8294 | tr\|Q68FZ8 | 160.56 | 22 | 22 | 8.06E+06 | 23 | 23 | 46 | Carbamidomethylatior | | Propionyl coenzyme A carboxylase, beta polypeptide |
| 49 | 31 | P45953\|A | 160.07 | 9 | 9 | 1.75E+06 | 9 | 9 | 13 | | 70749 | Very long-chain specific acyl-CoA dehydrogenase mitochondrial OS=Rattus norvegicus OX=10116 GN=Acadvl PE=1 SV=1 |
| 49 | 32 | tr\|Q5M9H | 160.07 | 9 | 9 | 1.75E+06 | 9 | 9 | 13 | | 70821 | Acyl-Coenzyme A dehydrogenase very long chain OS=Rattus norvegicus OX=10116 GN=Acadvl PE=1 SV=1 |
| 19 | 29 | Q60587\|E | 159.32 | 15 | 15 | 4.52E+06 | 17 | 17 | 33 | | 51414 | Trifunctional enzyme subunit beta mitochondrial OS=Rattus norvegicus OX=10116 GN=Hadhb PE=1 SV=1 |
| 19 | 40 | tr\|A0A0G2 | 159.32 | 14 | 14 | 4.52E+06 | 17 | 17 | 33 | | 52568 | Trifunctional enzyme subunit beta mitochondrial OS=Rattus norvegicus OX=10116 GN=Hadhb PE=1 SV=1 |
| 35 | 198 | Q80W89\| | 156.27 | 18 | 18 | 4.62E+06 | 9 | 9 | 18 | | 14854 | NADH dehydrogenase [ubiquinone] 1 alpha subcomplex subunit 11 OS=Rattus norvegicus OX=10116 GN=Ndufa11 PE=2 SV=1 |
| 14 | 28 | tr\|Q5UAJ6 | 154.17 | 40 | 40 | 8.90E+06 | 22 | 22 | 43 | Oxidation ( | 25942 | Cytochrome c oxidase subunit 2 OS=Rattus norvegicus OX=10116 GN=COX2 PE=3 SV=1 |
| 14 | 33 | tr\|S5RZM8 | 154.17 | 40 | 40 | 8.90E+06 | 22 | 22 | 43 | Oxidation ( | 25958 | Cytochrome c oxidase subunit 2 OS=Rattus norvegicus OX=10116 GN=COX2 PE=3 SV=1 |
| 14 | 34 | P00406\|C | 154.17 | 40 | 40 | 8.90E+06 | 22 | 22 | 43 | Oxidation ( | 25928 | Cytochrome c oxidase subunit 2 OS=Rattus norvegicus OX=10116 GN=Mtco2 PE=1 SV=3 |
| 14 | 35 | tr\|Q8SEZ5 | 154.17 | 40 | 40 | 8.90E+06 | 22 | 22 | 43 | Oxidation ( | 25928 | Cytochrome c oxidase subunit 2 OS=Rattus norvegicus OX=10116 GN=Mt-co2 PE=1 SV=1 |
| 14 | 36 | tr\|A0A097 | 154.17 | 40 | 40 | 8.90E+06 | 22 | 22 | 43 | Oxidation ( | 25894 | Cytochrome c oxidase subunit 2 OS=Rattus norvegicus OX=10116 GN=COX2 PE=3 SV=1 |
| 47 | 14 | Q6AXV4\|S | 153.8 | 13 | 13 | 1.02E+06 | 11 | 11 | 14 | | 51960 | Sorting and assembly machinery component 50 homolog OS=Rattus norvegicus OX=10116 GN=Samm50 PE=1 SV=1 |
| 36 | 252 | tr\|D3ZF13 | 149.92 | 22 | 22 | 2.34E+06 | 10 | 10 | 17 | | 17514 | Acyl carrier protein OS=Rattus norvegicus OX=10116 GN=Ndufab1 PE=1 SV=1 |
| 23 | 74 | tr\|D3ZG43 | 149.55 | 29 | 29 | 6.25E+06 | 14 | 14 | 27 | | 30226 | NADH dehydrogenase (Ubiquinone) Fe-S protein 3 (Predicted) isoform CRA_c OS=Rattus norvegicus OX=10116 GN=Ndufs3 PE=1 SV=1 |
| 26 | 77 | tr\|Q5PQZ9 | 149.37 | 37 | 37 | 6.43E+06 | 10 | 10 | 24 | | 14359 | NADH dehydrogenase [ubiquinone] 1 subunit C2 OS=Rattus norvegicus OX=10116 GN=Ndufc2 PE=1 SV=1 |
| 9 | 50 | P32551\|Q | 148.38 | 13 | 13 | 2.86E+07 | 17 | 17 | 53 | Oxidation ( | 48396 | Cytochrome b-c1 complex subunit 2 mitochondrial OS=Rattus norvegicus OX=10116 GN=Uqcrc2 PE=1 SV=2 |
| 27 | 21 | Q64428\|E | 147.72 | 11 | 11 | 3.65E+06 | 14 | 14 | 24 | | 82665 | Trifunctional enzyme subunit alpha mitochondrial OS=Rattus norvegicus OX=10116 GN=Hadha PE=1 SV=2 |
| 39 | 10 | tr\|A0A0G2 | 141.73 | 10 | 10 | 1.53E+06 | 12 | 12 | 16 | | 86230 | MICOS complex subunit MIC60 OS=Rattus norvegicus OX=10116 GN=Immt PE=1 SV=1 |
| 30 | 253 | P10817\|C | 137.51 | 26 | 26 | 7.35E+06 | 8 | 6 | 21 | | 10487 | Cytochrome c oxidase subunit 6A2 mitochondrial (Fragment) OS=Rattus norvegicus OX=10116 GN=Cox6a2 PE=1 SV=3 |
| 30 | 254 | tr\|G3V8M | 137.51 | 25 | 25 | 7.35E+06 | 8 | 6 | 21 | | 10802 | Cytochrome c oxidase subunit 6A mitochondrial OS=Rattus norvegicus OX=10116 GN=Cox6a2 PE=3 SV=1 |
| 20 | 313 | tr\|D3ZZ21 | 133.85 | 40 | 40 | 5.50E+06 | 10 | 10 | 31 | Oxidation ( | 15638 | NADH dehydrogenase (Ubiquinone) 1 beta subcomplex 6 (Predicted) OS=Rattus norvegicus OX=10116 GN=Ndufb6 PE=1 SV=1 |
| 52 | 139 | tr\|Q06QG | 132.64 | 5 | 5 | 3.15E+06 | 9 | 9 | 12 | | 68588 | NADH-ubiquinone oxidoreductase chain 5 OS=Rattus norvegicus OX=10116 GN=ND5 PE=3 SV=1 |
| 52 | 137 | tr\|Q06QK | 132.64 | 5 | 5 | 3.15E+06 | 9 | 9 | 12 | | 68606 | NADH-ubiquinone oxidoreductase chain 5 OS=Rattus norvegicus OX=10116 GN=ND5 PE=3 SV=1 |
| 52 | 138 | P11661\|N | 132.64 | 5 | 5 | 3.15E+06 | 9 | 9 | 12 | | 68618 | NADH-ubiquinone oxidoreductase chain 5 OS=Rattus norvegicus OX=10116 GN=Mtnd5 PE=3 SV=3 |
| 52 | 140 | tr\|Q06QA1 | 132.64 | 5 | 5 | 3.15E+06 | 9 | 9 | 12 | | 68574 | NADH-ubiquinone oxidoreductase chain 5 OS=Rattus norvegicus OX=10116 GN=ND5 PE=3 SV=1 |
| 52 | 141 | tr\|Q8SEZ0 | 132.64 | 5 | 5 | 3.15E+06 | 9 | 9 | 12 | | 68618 | NADH-ubiquinone oxidoreductase chain 5 OS=Rattus norvegicus OX=10116 GN=Mt-nd5 PE=3 SV=1 |
| 52 | 142 | tr\|A0A096 | 132.64 | 5 | 5 | 3.15E+06 | 9 | 9 | 12 | | 68584 | NADH-ubiquinone oxidoreductase chain 5 OS=Rattus norvegicus OX=10116 GN=ND5 PE=3 SV=1 |
| 52 | 164 | tr\|A7XYD5 | 132.64 | 5 | 5 | 3.15E+06 | 9 | 9 | 12 | | 68856 | NADH-ubiquinone oxidoreductase chain 5 OS=Rattus norvegicus OX=10116 GN=Nd5 PE=3 SV=1 |
| 52 | 165 | tr\|A7XYC1 | 132.64 | 5 | 5 | 3.15E+06 | 9 | 9 | 12 | | 68841 | NADH-ubiquinone oxidoreductase chain 5 OS=Rattus norvegicus OX=10116 GN=Nd5 PE=3 SV=1 |
| 52 | 166 | tr\|A0A0A1 | 132.64 | 5 | 5 | 3.15E+06 | 9 | 9 | 12 | | 68970 | NADH-ubiquinone oxidoreductase chain 5 OS=Rattus norvegicus OX=10116 GN=ND5 PE=3 SV=1 |
| 32 | 46 | P09605\|K | 131.78 | 16 | 16 | 1.59E+06 | 10 | 10 | 19 | | 47385 | Creatine kinase S-type mitochondrial OS=Rattus norvegicus OX=10116 GN=Ckmt2 PE=1 SV=2 |
| 50 | 30 | tr\|Q5BJZ3 | 131.23 | 4 | 4 | 1.96E+06 | 6 | 6 | 13 | Oxidation ( | 113869 | Nicotinamide nucleotide transhydrogenase OS=Rattus norvegicus OX=10116 GN=Nnt PE=1 SV=1 |

| | | | | | | | | | | | | |
|---|---|---|---|---|---|---|---|---|---|---|---|---|
| 57 | 44 | P17764\|TH | 130.29 | 12 | 12 | 1.61E+06 | 6 | 6 | 10 | | 44695 | Acetyl-CoA acetyltransferase mitochondrial OS=Rattus norvegicus OX=10116 GN=Acat1 PE=1 SV=1 |
| 33 | 172 | tr\|D4A0T0 | 129.62 | 16 | 16 | 1.14E+07 | 6 | 6 | 19 | | 20859 | NADH:ubiquinone oxidoreductase subunit B10 OS=Rattus norvegicus OX=10116 GN=Ndufb10 PE=1 SV=1 |
| 43 | 76 | Q06647\|AT | 126.78 | 13 | 13 | 2.85E+06 | 8 | 8 | 15 | | 23398 | ATP synthase subunit O mitochondrial OS=Rattus norvegicus OX=10116 GN=Atp5po PE=1 SV=1 |
| 51 | 219 | tr\|D3ZS58 | 125.75 | 19 | 19 | 1.42E+06 | 6 | 6 | 13 | | 10845 | NADH dehydrogenase [ubiquinone] 1 alpha subcomplex subunit 2 OS=Rattus norvegicus OX=10116 GN=Ndufa2 PE=1 SV=1 |
| 64 | 61 | Q63704\|CI | 121.35 | 4 | 4 | 9.88E+05 | 4 | 4 | 8 | | 88217 | Carnitine O-palmitoyltransferase 1 muscle isoform OS=Rattus norvegicus OX=10116 GN=Cpt1b PE=1 SV=1 |
| 59 | 56 | tr\|F1LPG5 | 119.89 | 21 | 21 | 3.64E+06 | 6 | 6 | 10 | | 15064 | NADH:ubiquinone oxidoreductase subunit B4 OS=Rattus norvegicus OX=10116 GN=Ndufb4 PE=1 SV=1 |
| 59 | 57 | tr\|F1M7T1 | 119.89 | 21 | 21 | 3.64E+06 | 6 | 6 | 10 | | 15160 | NADH dehydrogenase (ubiquinone) 1 beta subcomplex 4 OS=Rattus norvegicus OX=10116 GN=LOC100361934 PE=4 SV=2 |
| 46 | 169 | tr\|A9UMV | 119.02 | 21 | 21 | 1.00E+07 | 4 | 4 | 15 | | 12500 | NADH:ubiquinone oxidoreductase subunit A7 OS=Rattus norvegicus OX=10116 GN=Ndufa7 PE=1 SV=1 |
| 28 | 168 | P19511\|AT | 118.26 | 9 | 9 | 1.37E+07 | 6 | 6 | 21 | | 28869 | ATP synthase F(0) complex subunit B1 mitochondrial OS=Rattus norvegicus OX=10116 GN=Atp5pb PE=1 SV=1 |
| 58 | 318 | Q5M9I5\|Q | 118.08 | 18 | 18 | 4.25E+06 | 5 | 5 | 10 | | 10424 | Cytochrome b-c1 complex subunit 6 mitochondrial OS=Rattus norvegicus OX=10116 GN=Uqcrh PE=3 SV=1 |
| 40 | 49 | Q5BK63\|N | 115.68 | 18 | 18 | 2.41E+06 | 10 | 10 | 16 | | 42559 | NADH dehydrogenase [ubiquinone] 1 alpha subcomplex subunit 9 mitochondrial OS=Rattus norvegicus OX=10116 GN=Ndufa9 PE=1 SV=2 |
| 61 | 199 | tr\|B0BNE6 | 115.53 | 9 | 9 | 1.74E+06 | 5 | 5 | 10 | | 23970 | NADH dehydrogenase (Ubiquinone) Fe-S protein 8 (Predicted) isoform CRA_a OS=Rattus norvegicus OX=10116 GN=Ndufs8 PE=1 SV=1 |
| 85 | 3941 | P10818\|CX | 114.46 | 21 | 21 | 2.05E+05 | 5 | 3 | 5 | Oxidation ( | 12301 | Cytochrome c oxidase subunit 6A1 mitochondrial OS=Rattus norvegicus OX=10116 GN=Cox6a1 PE=1 SV=2 |
| 29 | 80 | tr\|Q5XIH3 | 110.16 | 11 | 11 | 3.12E+06 | 11 | 11 | 21 | | 50731 | NADH dehydrogenase [ubiquinone] flavoprotein 1 mitochondrial OS=Rattus norvegicus OX=10116 GN=Ndufv1 PE=1 SV=1 |
| 69 | 4293 | P52873\|PY | 105.37 | 6 | 6 | 2.73E+05 | 7 | 5 | 8 | Formylatio | 129777 | Pyruvate carboxylase mitochondrial OS=Rattus norvegicus OX=10116 GN=Pc PE=1 SV=2 |
| 69 | 4294 | tr\|A0A0G2 | 105.37 | 5 | 5 | 2.73E+05 | 7 | 5 | 8 | Formylatio | 140005 | Pyruvate carboxylase mitochondrial OS=Rattus norvegicus OX=10116 GN=Pc PE=1 SV=1 |
| 42 | 263 | P35171\|CX | 105.29 | 12 | 12 | 1.49E+06 | 3 | 3 | 16 | | 9353 | Cytochrome c oxidase subunit 7A2 mitochondrial OS=Rattus norvegicus OX=10116 GN=Cox7a2 PE=1 SV=1 |
| 42 | 264 | tr\|B2RYS0 | 105.29 | 12 | 12 | 1.49E+06 | 3 | 3 | 16 | | 9353 | Cox7a2 protein OS=Rattus norvegicus OX=10116 GN=Cox7a2 PE=2 SV=1 |
| 53 | 1021 | tr\|Q06QA9 | 104.02 | 9 | 9 | 1.12E+06 | 8 | 8 | 12 | Carbamido | 56893 | Cytochrome c oxidase subunit 1 OS=Rattus norvegicus OX=10116 GN=CO1 PE=3 SV=1 |
| 53 | 1022 | tr\|Q06QK0 | 104.02 | 9 | 9 | 1.12E+06 | 8 | 8 | 12 | Carbamido | 56880 | Cytochrome c oxidase subunit 1 OS=Rattus norvegicus OX=10116 GN=CO1 PE=3 SV=1 |
| 53 | 1023 | tr\|Q8SEZ6 | 104.02 | 9 | 9 | 1.12E+06 | 8 | 8 | 12 | Carbamido | 56879 | Cytochrome c oxidase subunit 1 OS=Rattus norvegicus OX=10116 GN=COI PE=3 SV=1 |
| 53 | 1024 | tr\|A0A0A1 | 104.02 | 9 | 9 | 1.12E+06 | 8 | 8 | 12 | Carbamido | 56937 | Cytochrome c oxidase subunit 1 OS=Rattus norvegicus OX=10116 GN=COX1 PE=3 SV=1 |
| 53 | 1025 | tr\|Q8HIC9 | 104.02 | 9 | 9 | 1.12E+06 | 8 | 8 | 12 | Carbamido | 56845 | Cytochrome c oxidase subunit 1 OS=Rattus norvegicus OX=10116 GN=Mt-co1 PE=3 SV=1 |
| 53 | 1026 | P05503\|CC | 104.02 | 9 | 9 | 1.12E+06 | 8 | 8 | 12 | Carbamido | 56845 | Cytochrome c oxidase subunit 1 OS=Rattus norvegicus OX=10116 GN=Mtco1 PE=2 SV=3 |
| 53 | 1027 | tr\|Q95938 | 104.02 | 9 | 9 | 1.12E+06 | 8 | 8 | 12 | Carbamido | 56977 | Cytochrome c oxidase subunit 1 OS=Rattus norvegicus OX=10116 GN=Co I PE=3 SV=1 |
| 45 | 398 | tr\|B2RYT5 | 102.9 | 32 | 32 | 2.73E+06 | 9 | 9 | 15 | Oxidation ( | 12651 | Cox7a2l protein OS=Rattus norvegicus OX=10116 GN=Cox7a2l PE=2 SV=1 |
| 45 | 399 | tr\|D3ZYX8 | 102.9 | 30 | 30 | 2.73E+06 | 9 | 9 | 15 | Oxidation ( | 13274 | Cytochrome c oxidase subunit 7A2-like OS=Rattus norvegicus OX=10116 GN=Cox7a2l PE=1 SV=1 |
| 79 | 48 | Q4V8F9\|H | 100.07 | 8 | 8 | 8.07E+05 | 4 | 4 | 5 | | 58344 | Hydroxysteroid dehydrogenase-like protein 2 OS=Rattus norvegicus OX=10116 GN=Hsdl2 PE=2 SV=1 |
| 56 | 292 | P10888\|CC | 99.64 | 24 | 24 | 1.07E+06 | 9 | 9 | 11 | Oxidation ( | 19515 | Cytochrome c oxidase subunit 4 isoform 1 mitochondrial OS=Rattus norvegicus OX=10116 GN=Cox4i1 PE=1 SV=1 |
| 72 | 301 | Q5FVQ8\|N | 97.91 | 4 | 4 | 4.95E+05 | 5 | 4 | 7 | | 107590 | NLR family member X1 OS=Rattus norvegicus OX=10116 GN=Nlrx1 PE=2 SV=1 |
| 76 | 18 | P26284\|OI | 97.7 | 7 | 7 | 1.07E+06 | 4 | 4 | 6 | | 43227 | Pyruvate dehydrogenase E1 component subunit alpha somatic form mitochondrial OS=Rattus norvegicus OX=10116 GN=Pdha1 PE=1 SV=2 |
| 38 | 75 | Q6PDU7\|A | 94.42 | 22 | 22 | 2.67E+06 | 8 | 8 | 16 | | 11433 | ATP synthase subunit g mitochondrial OS=Rattus norvegicus OX=10116 GN=Atp5mg PE=1 SV=2 |
| 41 | 267 | tr\|I6V4L9\| | 91.98 | 11 | 11 | 1.12E+06 | 5 | 5 | 16 | Oxidation ( | 29844 | Cytochrome c oxidase subunit 3 OS=Rattus norvegicus OX=10116 GN=COX3 PE=3 SV=1 |
| 54 | 532 | P67779\|PH | 88.85 | 17 | 17 | 5.14E+05 | 8 | 8 | 12 | | 29820 | Prohibitin OS=Rattus norvegicus OX=10116 GN=Phb PE=1 SV=1 |
| 93 | 3906 | tr\|D3ZCZ9 | 87.47 | 16 | 16 | 5.41E+05 | 3 | 3 | 4 | | 13040 | NADH dehydrogenase [ubiquinone] iron-sulfur protein 6 mitochondrial OS=Rattus norvegicus OX=10116 GN=LOC100912599 PE=1 SV=1 |
| 71 | 233 | tr\|Q5UAJ5 | 86.48 | 19 | 19 | 4.07E+06 | 3 | 3 | 7 | Oxidation ( | 7642 | ATP synthase protein 8 OS=Rattus norvegicus OX=10116 GN=ATP8 PE=3 SV=1 |
| 71 | 234 | P11608\|AT | 86.48 | 19 | 19 | 4.07E+06 | 3 | 3 | 7 | Oxidation ( | 7630 | ATP synthase protein 8 OS=Rattus norvegicus OX=10116 GN=Mt-atp8 PE=1 SV=2 |
| 71 | 235 | tr\|Q8SEZ4 | 86.48 | 19 | 19 | 4.07E+06 | 3 | 3 | 7 | Oxidation ( | 7632 | ATP synthase protein 8 OS=Rattus norvegicus OX=10116 GN=ATPase8 PE=3 SV=1 |
| 71 | 236 | tr\|Q8HIC8 | 86.48 | 19 | 19 | 4.07E+06 | 3 | 3 | 7 | Oxidation ( | 7630 | ATP synthase protein 8 OS=Rattus norvegicus OX=10116 GN=Mt-atp8 PE=3 SV=1 |
| 75 | 152 | Q8VHF5\|C | 81.78 | 5 | 5 | 7.09E+05 | 5 | 5 | 6 | | 51867 | Citrate synthase mitochondrial OS=Rattus norvegicus OX=10116 GN=Cs PE=1 SV=1 |
| 75 | 153 | tr\|G3V936 | 81.78 | 5 | 5 | 7.09E+05 | 5 | 5 | 6 | | 51831 | Citrate synthase OS=Rattus norvegicus OX=10116 GN=Cs PE=1 SV=1 |
| 78 | 315 | tr\|G3V7I0\| | 81.76 | 12 | 12 | 1.10E+06 | 3 | 3 | 6 | | 28299 | Peroxiredoxin 3 OS=Rattus norvegicus OX=10116 GN=Prdx3 PE=1 SV=1 |
| 78 | 316 | Q9Z0V6\|PI | 81.76 | 12 | 12 | 1.10E+06 | 3 | 3 | 6 | | 28295 | Thioredoxin-dependent peroxide reductase mitochondrial OS=Rattus norvegicus OX=10116 GN=Prdx3 PE=1 SV=2 |
| 66 | 273 | P04636\|M | 81.68 | 9 | 9 | 5.04E+05 | 5 | 5 | 8 | | 35684 | Malate dehydrogenase mitochondrial OS=Rattus norvegicus OX=10116 GN=Mdh2 PE=1 SV=2 |
| 77 | 335 | Q5XIH7\|PH | 81.4 | 9 | 9 | 1.78E+06 | 4 | 4 | 6 | | 33312 | Prohibitin-2 OS=Rattus norvegicus OX=10116 GN=Phb2 PE=1 SV=1 |
| 63 | 72 | tr\|D3ZE15 | 80.33 | 32 | 32 | 6.98E+05 | 6 | 6 | 9 | Oxidation ( | 16777 | NADH:ubiquinone oxidoreductase subunit A13 OS=Rattus norvegicus OX=10116 GN=Ndufa13 PE=1 SV=1 |
| 73 | 827 | tr\|D4A3V2 | 79.44 | 8 | 8 | 8.10E+05 | 4 | 4 | 7 | | 15224 | NADH dehydrogenase [ubiquinone] 1 alpha subcomplex subunit 6 OS=Rattus norvegicus OX=10116 GN=Ndufa6 PE=1 SV=1 |
| 67 | 207 | P11951\|CX | 79.4 | 26 | 26 | 6.38E+05 | 5 | 5 | 8 | Oxidation ( | 8455 | Cytochrome c oxidase subunit 6C-2 OS=Rattus norvegicus OX=10116 GN=Cox6c2 PE=1 SV=3 |
| 87 | 296 | tr\|F1LP30\| | 75.63 | 3 | 3 | 7.84E+04 | 2 | 1 | 4 | | 79296 | Methylcrotonoyl-CoA carboxylase subunit alpha mitochondrial OS=Rattus norvegicus OX=10116 GN=Mccc1 PE=1 SV=1 |
| 87 | 297 | Q5I0C3\|M | 75.63 | 3 | 3 | 7.84E+04 | 2 | 1 | 4 | | 79330 | Methylcrotonoyl-CoA carboxylase subunit alpha mitochondrial OS=Rattus norvegicus OX=10116 GN=Mccc1 PE=1 SV=1 |
| 60 | 47 | P18163\|AC | 75.55 | 5 | 5 | 8.12E+05 | 5 | 4 | 10 | Carbamidomethylatio | | Long-chain-fatty-acid--CoA ligase 1 |
| 84 | 238 | Q5XIF3\|ND | 74.92 | 12 | 12 | 8.40E+05 | 3 | 3 | 5 | | 19741 | NADH dehydrogenase [ubiquinone] iron-sulfur protein 4 mitochondrial OS=Rattus norvegicus OX=10116 GN=Ndufs4 PE=1 SV=1 |
| 70 | 209 | tr\|B2RZD6 | 74.74 | 21 | 21 | 2.12E+06 | 4 | 4 | 7 | | 9327 | NDUFA4 mitochondrial complex-associated OS=Rattus norvegicus OX=10116 GN=Ndufa4 PE=1 SV=1 |
| 55 | 170 | tr\|Q6P6W | 72.91 | 16 | 16 | 1.24E+06 | 6 | 6 | 12 | | 31701 | NADH dehydrogenase [ubiquinone] 1 alpha subcomplex subunit 10 mitochondrial OS=Rattus norvegicus OX=10116 GN=Ndufa10 PE=2 SV=1 |
| 55 | 171 | Q561S0\|N | 72.91 | 13 | 13 | 1.24E+06 | 6 | 6 | 12 | | 40493 | NADH dehydrogenase [ubiquinone] 1 alpha subcomplex subunit 10 mitochondrial OS=Rattus norvegicus OX=10116 GN=Ndufa10 PE=1 SV=1 |
| 62 | 3901 | tr\|Q8SEZ8 | 71.94 | 6 | 6 | 1.86E+06 | 5 | 5 | 9 | | 36133 | NADH-ubiquinone oxidoreductase chain 1 (Fragment) OS=Rattus norvegicus OX=10116 GN=NADH1 PE=3 SV=1 |
| 62 | 3902 | P03889\|NU | 71.94 | 6 | 6 | 1.86E+06 | 5 | 5 | 9 | | 36145 | NADH-ubiquinone oxidoreductase chain 1 OS=Rattus norvegicus OX=10116 GN=Mtnd1 PE=1 SV=3 |
| 62 | 3903 | tr\|Q8HID1 | 71.94 | 6 | 6 | 1.86E+06 | 5 | 5 | 9 | | 36145 | NADH-ubiquinone oxidoreductase chain 1 OS=Rattus norvegicus OX=10116 GN=Mt-nd1 PE=3 SV=1 |
| 62 | 3904 | tr\|D2E6L7 | 71.94 | 6 | 6 | 1.86E+06 | 5 | 5 | 9 | | 36062 | NADH-ubiquinone oxidoreductase chain 1 OS=Rattus norvegicus OX=10116 GN=ND1 PE=3 SV=1 |
| 106 | 3930 | tr\|Q35733 | 67.62 | 13 | 13 | 1.77E+05 | 2 | 2 | 3 | | 8389 | NADH-ubiquinone oxidoreductase chain 6 (Fragment) OS=Rattus norvegicus OX=10116 PE=3 SV=1 |
| 106 | 3921 | tr\|Q06QD9 | 67.62 | 6 | 6 | 1.77E+05 | 2 | 2 | 3 | | 18943 | NADH-ubiquinone oxidoreductase chain 6 OS=Rattus norvegicus OX=10116 GN=ND6 PE=3 SV=1 |
| 106 | 3922 | tr\|Q7HKW | 67.62 | 6 | 6 | 1.77E+05 | 2 | 2 | 3 | | 18957 | NADH-ubiquinone oxidoreductase chain 6 OS=Rattus norvegicus OX=10116 GN=NADH6 PE=3 SV=1 |
| 99 | 4 | P08461\|OI | 63.83 | 3 | 3 | 4.62E+05 | 3 | 3 | 3 | | 67166 | Dihydrolipoyllysine-residue acetyltransferase component of pyruvate dehydrogenase complex mitochondrial OS=Rattus norvegicus OX=10116 GN=Dlat PE=1 SV=3 |

| | | | | | | | | | | | | |
|---|---|---|---|---|---|---|---|---|---|---|---|---|
| 161 | 245 | tr\|C8CHS6 | 63.48 | 15 | 15 | 1.18E+05 | 1 | 1 | 1 | | 7988 | Mitochondrial superoxide dismutase 2 (Fragment) OS=Rattus norvegicus OX=10116 PE=2 SV=1 |
| 161 | 71 | P07895\|SO | 63.48 | 5 | 5 | 1.18E+05 | 1 | 1 | 1 | | 24674 | Superoxide dismutase [Mn] mitochondrial OS=Rattus norvegicus OX=10116 GN=Sod2 PE=1 SV=2 |
| 68 | 3899 | tr\|B5DEL8 | 63.31 | 21 | 21 | 6.45E+05 | 4 | 4 | 8 | | 12700 | NADH dehydrogenase (Ubiquinone) Fe-S protein 5 OS=Rattus norvegicus OX=10116 GN=Ndufs5 PE=1 SV=1 |
| 68 | 3900 | tr\|A0A0G2 | 63.31 | 21 | 21 | 6.45E+05 | 4 | 4 | 8 | | 12730 | Uncharacterized protein OS=Rattus norvegicus OX=10116 PE=4 SV=1 |
| 96 | 8042 | tr\|Q4G067 | 62.48 | 4 | 4 | 1.61E+05 | 3 | 3 | 4 | | 37440 | Mitochondrial ribosomal protein L44 OS=Rattus norvegicus OX=10116 GN=Mrpl44 PE=1 SV=1 |
| 92 | 144 | P13086\|SU | 59.66 | 7 | 7 | 1.15E+05 | 3 | 3 | 4 | | 36148 | Succinate--CoA ligase [ADP/GDP-forming] subunit alpha mitochondrial OS=Rattus norvegicus OX=10116 GN=Suclg1 PE=2 SV=2 |
| 92 | 145 | tr\|A0A0H2 | 59.66 | 7 | 7 | 1.15E+05 | 3 | 3 | 4 | | 37560 | Succinate--CoA ligase [ADP/GDP-forming] subunit alpha mitochondrial OS=Rattus norvegicus OX=10116 GN=Suclg1 PE=1 SV=1 |
| 113 | 73 | Q63362\|N | 58.4 | 9 | 9 | 2.40E+05 | 2 | 2 | 2 | | 13412 | NADH dehydrogenase [ubiquinone] 1 alpha subcomplex subunit 5 OS=Rattus norvegicus OX=10116 GN=Ndufa5 PE=1 SV=3 |
| 83 | 326 | Q91XJ1\|BE | 54.38 | 3 | 3 | | 3 | 0 | 5 | | 51557 | Beclin-1 OS=Rattus norvegicus OX=10116 GN=Becn1 PE=1 SV=1 |
| 94 | 400 | Q704S8\|C | 53.12 | 3 | 3 | 1.52E+05 | 2 | 2 | 4 | | 70801 | Carnitine O-acetyltransferase OS=Rattus norvegicus OX=10116 GN=Crat PE=1 SV=1 |
| 94 | 401 | tr\|A0A0H2 | 53.12 | 3 | 3 | 1.52E+05 | 2 | 2 | 4 | | 71014 | Carnitine O-acetyltransferase OS=Rattus norvegicus OX=10116 GN=Crat PE=1 SV=1 |
| 86 | 3893 | tr\|B2RYS2 | 52.63 | 24 | 24 | 1.35E+06 | 4 | 4 | 4 | | 13559 | Cytochrome b-c1 complex subunit 7 OS=Rattus norvegicus OX=10116 GN=Uqcrb PE=1 SV=1 |
| 80 | 66 | P56574\|ID | 52.23 | 7 | 7 | 2.71E+05 | 4 | 4 | 5 | | 50967 | Isocitrate dehydrogenase [NADP] mitochondrial OS=Rattus norvegicus OX=10116 GN=Idh2 PE=1 SV=2 |
| 74 | 208 | Q7TQ16\|Q | 51.71 | 20 | 20 | 1.30E+06 | 4 | 4 | 6 | | 9849 | Cytochrome b-c1 complex subunit 8 OS=Rattus norvegicus OX=10116 GN=Uqcrq PE=3 SV=1 |
| 162 | 15874 | tr\|M0R5K3 | 49.53 | 11 | 11 | 9.12E+04 | 1 | 1 | 1 | | 9028 | Cytochrome c oxidase subunit 7A1 OS=Rattus norvegicus OX=10116 GN=LOC687508 PE=4 SV=1 |
| 91 | 4412 | Q02253\|M | 49.42 | 8 | 8 | 2.08E+05 | 4 | 4 | 4 | Oxidation ( | 57808 | Methylmalonate-semialdehyde dehydrogenase [acylating] mitochondrial OS=Rattus norvegicus OX=10116 GN=Aldh6a1 PE=1 SV=1 |
| 91 | 4413 | tr\|G3V7J0 | 49.42 | 8 | 8 | 2.08E+05 | 4 | 4 | 4 | Oxidation ( | 57748 | Aldehyde dehydrogenase family 6 subfamily A1 isoform CRA_b OS=Rattus norvegicus OX=10116 GN=Aldh6a1 PE=1 SV=1 |
| 81 | 467 | tr\|D4A4P3 | 48.34 | 22 | 22 | 7.98E+05 | 3 | 3 | 5 | Oxidation ( | 11267 | NADH:ubiquinone oxidoreductase subunit B3 OS=Rattus norvegicus OX=10116 GN=Ndufb3 PE=1 SV=1 |
| 88 | 319 | P29419\|AT | 46.66 | 21 | 21 | 3.70E+05 | 2 | 2 | 4 | | 8255 | ATP synthase subunit e mitochondrial OS=Rattus norvegicus OX=10116 GN=Atp5me PE=1 SV=3 |
| 98 | 159 | tr\|Q5U1W | 45.27 | 12 | 12 | 3.96E+05 | 2 | 2 | 3 | | 28238 | MICOS complex subunit OS=Rattus norvegicus OX=10116 GN=Apool PE=1 SV=1 |
| 104 | 78 | Q920L2\|SD | 41.59 | 3 | 3 | 1.26E+05 | 2 | 2 | 3 | | 71615 | Succinate dehydrogenase [ubiquinone] flavoprotein subunit mitochondrial OS=Rattus norvegicus OX=10116 GN=Sdha PE=1 SV=1 |
| 65 | 151 | P12075\|CO | 40.77 | 9 | 9 | 2.34E+06 | 3 | 3 | 8 | | 13915 | Cytochrome c oxidase subunit 5B mitochondrial OS=Rattus norvegicus OX=10116 GN=Cox5b PE=1 SV=2 |
| 110 | 23 | tr\|Q6P9Y4 | 40.63 | 5 | 5 | 1.66E+05 | 2 | 2 | 2 | | 32904 | ADP/ATP translocase 1 OS=Rattus norvegicus OX=10116 GN=Slc25a4 PE=1 SV=1 |
| 110 | 24 | Q05962\|AI | 40.63 | 5 | 5 | 1.66E+05 | 2 | 2 | 2 | | 32989 | ADP/ATP translocase 1 OS=Rattus norvegicus OX=10116 GN=Slc25a4 PE=1 SV=3 |
| 110 | 27 | Q09073\|AI | 40.63 | 5 | 5 | 1.66E+05 | 2 | 2 | 2 | | 32901 | ADP/ATP translocase 2 OS=Rattus norvegicus OX=10116 GN=Slc25a5 PE=1 SV=3 |
| 110 | 43 | tr\|D3ZB81 | 40.63 | 5 | 5 | 1.66E+05 | 2 | 2 | 2 | | 35227 | Solute carrier family 25 member 31 OS=Rattus norvegicus OX=10116 GN=Slc25a31 PE=3 SV=3 |
| 111 | 3914 | tr\|B2RYU0 | 39.33 | 21 | 21 | 7.13E+05 | 2 | 2 | 2 | | 11842 | NADH dehydrogenase (Ubiquinone) 1 beta subcomplex 2 (Predicted) isoform CRA_b OS=Rattus norvegicus OX=10116 GN=Ndufb2 PE=1 SV=1 |
| 95 | 3951 | tr\|B1WBP7 | 39.1 | 13 | 13 | 3.81E+05 | 2 | 2 | 4 | | 12884 | ATP synthase subunit delta mitochondrial OS=Rattus norvegicus OX=10116 GN=Atp5f1d PE=1 SV=1 |
| 95 | 3953 | tr\|G3V7Y3 | 39.1 | 10 | 10 | 3.81E+05 | 2 | 2 | 4 | | 17563 | ATP synthase subunit delta mitochondrial OS=Rattus norvegicus OX=10116 GN=Atp5f1d PE=1 SV=1 |
| 82 | 3923 | P80431\|CO | 39.07 | 24 | 24 | 3.27E+05 | 3 | 3 | 5 | | 8995 | Cytochrome c oxidase subunit 7B mitochondrial OS=Rattus norvegicus OX=10116 GN=Cox7b PE=1 SV=3 |
| 163 | 15875 | tr\|F1LWG4 | 38.24 | 4 | 4 | 2.74E+05 | 1 | 1 | 1 | | 37782 | NADH dehydrogenase (Ubiquinone) 1 alpha subcomplex assembly factor 1 (Predicted) isoform CRA_a OS=Rattus norvegicus OX=10116 GN=Ndufaf1 PE=1 SV=2 |
| 105 | 215 | P33124\|AC | 38.05 | 2 | 2 | 0 | 2 | 1 | 3 | Carbamido | 78180 | Long-chain-fatty-acid--CoA ligase 6 OS=Rattus norvegicus OX=10116 GN=Acsl6 PE=1 SV=1 |
| 117 | 343 | Q924S5\|LO | 36.69 | 1 | 1 | 6.41E+04 | 1 | 1 | 2 | | 105792 | Lon protease homolog mitochondrial OS=Rattus norvegicus OX=10116 GN=Lonp1 PE=2 SV=1 |
| 107 | 310 | Q75Q40\|T | 36.13 | 4 | 4 | 2.57E+05 | 1 | 1 | 3 | | 37919 | Mitochondrial import receptor subunit TOM40 homolog OS=Rattus norvegicus OX=10116 GN=Tomm40 PE=1 SV=1 |
| 107 | 311 | tr\|G3V8F5 | 36.13 | 4 | 4 | 2.57E+05 | 1 | 1 | 3 | | 37920 | Mitochondrial import receptor subunit TOM40 homolog OS=Rattus norvegicus OX=10116 GN=Tomm40 PE=1 SV=1 |
| 164 | 3934 | Q66H47\|R | 34.79 | 6 | 6 | 7.38E+04 | 1 | 1 | 1 | | 25001 | 39S ribosomal protein L24 mitochondrial OS=Rattus norvegicus OX=10116 GN=Mrpl24 PE=2 SV=1 |
| 165 | 23059 | Q5I0I4\|DM | 32.87 | 6 | 6 | 6.83E+04 | 1 | 1 | 1 | | 28508 | Distal membrane-arm assembly complex protein 2 OS=Rattus norvegicus OX=10116 GN=Dmac2 PE=2 SV=1 |
| 118 | 3985 | tr\|Q5XFW4 | 32.7 | 7 | 7 | 2.56E+04 | 1 | 1 | 2 | | 20541 | Mitochondrial ribosomal protein L13 OS=Rattus norvegicus OX=10116 GN=Mrpl13 PE=1 SV=1 |
| 141 | 8319 | Q9WVK7\|H | 32.24 | 5 | 5 | 3.54E+04 | 1 | 1 | 1 | Oxidation ( | 34448 | Hydroxyacyl-coenzyme A dehydrogenase mitochondrial OS=Rattus norvegicus OX=10116 GN=Hadh PE=2 SV=1 |
| 89 | 256 | tr\|Q8M7G | 31.21 | 8 | 8 | 1.72E+05 | 2 | 2 | 4 | Oxidation ( | 24240 | ATP synthase subunit a (Fragment) OS=Rattus norvegicus OX=10116 PE=2 SV=1 |
| 89 | 257 | tr\|Q549I0\| | 31.21 | 8 | 8 | 1.72E+05 | 2 | 2 | 4 | Oxidation ( | 25050 | ATP synthase subunit a OS=Rattus norvegicus OX=10116 GN=atp6 PE=4 SV=1 |
| 89 | 258 | P05504\|AT | 31.21 | 8 | 8 | 1.72E+05 | 2 | 2 | 4 | Oxidation ( | 25076 | ATP synthase subunit a OS=Rattus norvegicus OX=10116 GN=Mt-atp6 PE=1 SV=3 |
| 89 | 259 | tr\|Q8HIC7 | 31.21 | 8 | 8 | 1.72E+05 | 2 | 2 | 4 | Oxidation ( | 25076 | ATP synthase subunit a OS=Rattus norvegicus OX=10116 GN=Mt-atp6 PE=4 SV=1 |
| 89 | 260 | tr\|S5S1E9\| | 31.21 | 8 | 8 | 1.72E+05 | 2 | 2 | 4 | Oxidation ( | 25049 | ATP synthase subunit a OS=Rattus norvegicus OX=10116 GN=ATP6 PE=4 SV=1 |
| 89 | 262 | tr\|Q8SEZ3 | 31.21 | 8 | 8 | 1.72E+05 | 2 | 2 | 4 | Oxidation ( | 25030 | ATP synthase subunit a OS=Rattus norvegicus OX=10116 GN=ATPase6 PE=4 SV=1 |
| 119 | 1037 | Q08877\|D | 30.88 | 1 | 1 | 1.19E+05 | 1 | 1 | 2 | | 97914 | Dynamin-3 OS=Rattus norvegicus OX=10116 GN=Dnm3 PE=1 SV=2 |
| 101 | 376 | tr\|Q5RJN0 | 30.79 | 8 | 8 | 6.84E+05 | 2 | 2 | 3 | | 23945 | NADH dehydrogenase (Ubiquinone) Fe-S protein 7 OS=Rattus norvegicus OX=10116 GN=Ndufs7 PE=1 SV=1 |
| 149 | 79 | tr\|D3ZUX5 | 30.11 | 4 | 4 | 5.14E+04 | 1 | 1 | 1 | | 26435 | MICOS complex subunit OS=Rattus norvegicus OX=10116 GN=Chchd3 PE=1 SV=1 |
| 116 | 4487 | tr\|D4A4B1 | 29.17 | 6 | 6 | 9.13E+04 | 2 | 2 | 2 | | 23223 | Mitochondrial ribosomal protein L15 OS=Rattus norvegicus OX=10116 GN=Mrpl15 PE=1 SV=2 |
| 116 | 4488 | tr\|A0A0G2 | 29.17 | 4 | 4 | 9.13E+04 | 2 | 2 | 2 | | 33652 | Mitochondrial ribosomal protein L15 OS=Rattus norvegicus OX=10116 GN=Mrpl15 PE=1 SV=1 |
| 114 | 3919 | tr\|A9UMV | 29.05 | 11 | 11 | 1.15E+05 | 1 | 1 | 2 | | 6539 | RCG29512 OS=Rattus norvegicus OX=10116 GN=Uqcr11 PE=1 SV=1 |
| 166 | 3932 | tr\|D3ZD09 | 28.7 | 12 | 12 | 8.65E+04 | 1 | 1 | 1 | | 10071 | Cytochrome c oxidase subunit OS=Rattus norvegicus OX=10116 GN=Cox6b1 PE=1 SV=1 |
| 167 | 384 | tr\|Q06Q97 | 28.64 | 2 | 2 | 1.03E+05 | 1 | 1 | 1 | | 38485 | NADH-ubiquinone oxidoreductase chain 2 OS=Rattus norvegicus OX=10116 GN=ND2 PE=3 SV=1 |
| 167 | 385 | tr\|A0A0A1 | 28.64 | 2 | 2 | 1.03E+05 | 1 | 1 | 1 | | 38542 | NADH-ubiquinone oxidoreductase chain 2 OS=Rattus norvegicus OX=10116 GN=ND2 PE=3 SV=1 |
| 167 | 386 | tr\|Q5UAJ8 | 28.64 | 2 | 2 | 1.03E+05 | 1 | 1 | 1 | | 38455 | NADH-ubiquinone oxidoreductase chain 2 OS=Rattus norvegicus OX=10116 GN=ND2 PE=3 SV=1 |
| 167 | 388 | tr\|Q8HID0 | 28.64 | 2 | 2 | 1.03E+05 | 1 | 1 | 1 | | 38653 | NADH-ubiquinone oxidoreductase chain 2 OS=Rattus norvegicus OX=10116 GN=Mt-nd2 PE=3 SV=1 |
| 167 | 389 | P11662\|NU | 28.64 | 2 | 2 | 1.03E+05 | 1 | 1 | 1 | | 38653 | NADH-ubiquinone oxidoreductase chain 2 OS=Rattus norvegicus OX=10116 GN=Mtnd2 PE=3 SV=3 |
| 167 | 390 | tr\|Q06QH5 | 28.64 | 2 | 2 | 1.03E+05 | 1 | 1 | 1 | | 38626 | NADH-ubiquinone oxidoreductase chain 2 OS=Rattus norvegicus OX=10116 GN=ND2 PE=3 SV=1 |
| 167 | 391 | tr\|Q8SEZ7 | 28.64 | 2 | 2 | 1.03E+05 | 1 | 1 | 1 | | 38580 | NADH-ubiquinone oxidoreductase chain 2 (Fragment) OS=Rattus norvegicus OX=10116 GN=NADH2 PE=3 SV=1 |
| 167 | 392 | tr\|Q06QB0 | 28.64 | 2 | 2 | 1.03E+05 | 1 | 1 | 1 | | 38534 | NADH-ubiquinone oxidoreductase chain 2 OS=Rattus norvegicus OX=10116 GN=ND2 PE=3 SV=1 |
| 167 | 393 | tr\|D2E6P4 | 28.64 | 2 | 2 | 1.03E+05 | 1 | 1 | 1 | | 38623 | NADH-ubiquinone oxidoreductase chain 2 OS=Rattus norvegicus OX=10116 GN=ND2 PE=3 SV=1 |
| 167 | 394 | tr\|Q06QE9 | 28.64 | 2 | 2 | 1.03E+05 | 1 | 1 | 1 | | 38598 | NADH-ubiquinone oxidoreductase chain 2 OS=Rattus norvegicus OX=10116 GN=ND2 PE=3 SV=1 |
| 100 | 344 | tr\|D4A4K4 | 28.43 | 0 | 0 | 1.48E+06 | 2 | 2 | 3 | | 418626 | Vacuolar protein sorting 13 homolog C OS=Rattus norvegicus OX=10116 GN=Vps13c PE=1 SV=2 |

| | | | | | | | | | | | | |
|---|---|---|---|---|---|---|---|---|---|---|---|---|
| 148 | 12037 | P21571\|AT | 26.73 | 9 | 9 | 2.69E+04 | 1 | 1 | 1 | | 12494 | ATP synthase-coupling factor 6 mitochondrial OS=Rattus norvegicus OX=10116 GN=Atp5pf PE=1 SV=1 |
| 169 | 8178 | tr\|D4A9Z6 | 26.51 | 2 | 2 | 1.03E+05 | 1 | 1 | 1 | | 36207 | Mitochondrial ribosomal protein S35 OS=Rattus norvegicus OX=10116 GN=Mrps35 PE=1 SV=1 |
| 120 | 474 | tr\|F1LN92 | 25.3 | 2 | 2 | 1.16E+05 | 2 | 2 | 2 | | 89352 | AFG3-like matrix AAA peptidase subunit 2 OS=Rattus norvegicus OX=10116 GN=Afg3l2 PE=1 SV=1 |
| 108 | 3917 | tr\|D3ZJG4 | 25.09 | 1 | 1 | 2.61E+05 | 1 | 1 | 3 | Oxidation ( | 95972 | Phosphofurin acidic cluster sorting protein 2 OS=Rattus norvegicus OX=10116 GN=Pacs2 PE=1 SV=2 |
| 121 | 4158 | P46462\|TE | 24.78 | 1 | 1 | 1.52E+05 | 1 | 1 | 2 | | 89349 | Transitional endoplasmic reticulum ATPase OS=Rattus norvegicus OX=10116 GN=Vcp PE=1 SV=3 |
| 170 | 23060 | P80433\|CO | 24.13 | 14 | 14 | 2.14E+04 | 1 | 1 | 1 | | 7672 | Cytochrome c oxidase subunit 8A mitochondrial OS=Rattus norvegicus OX=10116 GN=Cox8a PE=1 SV=3 |
| 143 | 16 | P02563\|M | 24.12 | 1 | 1 | 8.00E+05 | 1 | 1 | 1 | Formylatio | 223506 | Myosin-6 OS=Rattus norvegicus OX=10116 GN=Myh6 PE=1 SV=2 |
| 171 | 12335 | tr\|D3ZFJ6\| | 23.94 | 4 | 4 | 3.36E+05 | 1 | 1 | 1 | | 60420 | Lactamase beta OS=Rattus norvegicus OX=10116 GN=Lactb PE=1 SV=1 |
| 115 | 363 | P05508\|NU | 23.45 | 2 | 2 | 7.97E+04 | 1 | 1 | 2 | | 51783 | NADH-ubiquinone oxidoreductase chain 4 OS=Rattus norvegicus OX=10116 GN=Mtnd4 PE=3 SV=3 |
| 115 | 364 | tr\|D2E6K0 | 23.45 | 2 | 2 | 7.97E+04 | 1 | 1 | 2 | | 51833 | NADH-ubiquinone oxidoreductase chain 4 OS=Rattus norvegicus OX=10116 GN=ND4 PE=3 SV=1 |
| 115 | 365 | tr\|A7XYB9 | 23.45 | 2 | 2 | 7.97E+04 | 1 | 1 | 2 | | 51773 | NADH-ubiquinone oxidoreductase chain 4 OS=Rattus norvegicus OX=10116 GN=Nd4 PE=3 SV=1 |
| 115 | 366 | tr\|Q06QE1 | 23.45 | 2 | 2 | 7.97E+04 | 1 | 1 | 2 | | 51851 | NADH-ubiquinone oxidoreductase chain 4 OS=Rattus norvegicus OX=10116 GN=ND4 PE=3 SV=1 |
| 115 | 367 | tr\|Q06QA2 | 23.45 | 2 | 2 | 7.97E+04 | 1 | 1 | 2 | | 51787 | NADH-ubiquinone oxidoreductase chain 4 OS=Rattus norvegicus OX=10116 GN=ND4 PE=3 SV=1 |
| 115 | 368 | tr\|Q7HKW | 23.45 | 2 | 2 | 7.97E+04 | 1 | 1 | 2 | | 51801 | NADH-ubiquinone oxidoreductase chain 4 (Fragment) OS=Rattus norvegicus OX=10116 GN=NADH4 PE=3 SV=1 |
| 115 | 369 | tr\|Q06QG7 | 23.45 | 2 | 2 | 7.97E+04 | 1 | 1 | 2 | | 51791 | NADH-ubiquinone oxidoreductase chain 4 OS=Rattus norvegicus OX=10116 GN=ND4 PE=3 SV=1 |
| 115 | 370 | tr\|Q06Q89 | 23.45 | 2 | 2 | 7.97E+04 | 1 | 1 | 2 | | 51819 | NADH-ubiquinone oxidoreductase chain 4 OS=Rattus norvegicus OX=10116 GN=ND4 PE=3 SV=1 |
| 115 | 371 | tr\|Q35737 | 23.45 | 2 | 2 | 7.97E+04 | 1 | 1 | 2 | | 51765 | NADH-ubiquinone oxidoreductase chain 4 OS=Rattus norvegicus OX=10116 PE=3 SV=1 |
| 115 | 372 | tr\|Q8HIC6 | 23.45 | 2 | 2 | 7.97E+04 | 1 | 1 | 2 | | 51783 | NADH-ubiquinone oxidoreductase chain 4 OS=Rattus norvegicus OX=10116 GN=Mt-nd4 PE=3 SV=1 |
| 122 | 19471 | Q75Q41\|T | 23.29 | 8 | 8 | 1.28E+05 | 1 | 1 | 2 | | 15491 | Mitochondrial import receptor subunit TOM22 homolog OS=Rattus norvegicus OX=10116 GN=Tomm22 PE=1 SV=1 |
| 109 | 1269 | Q5M934\|T | 22.59 | 2 | 2 | 1.65E+06 | 1 | 1 | 2 | | 36404 | Probable tRNA pseudouridine synthase 1 OS=Rattus norvegicus OX=10116 GN=Trub1 PE=2 SV=1 |
| 174 | 19460 | tr\|D3ZMR9 | 22.46 | 4 | 4 | 6.74E+04 | 1 | 1 | 1 | | 23390 | Mitochondrial ribosomal protein L21 OS=Rattus norvegicus OX=10116 GN=Mrpl21 PE=1 SV=1 |
| 142 | 13 | P49432\|OD | 21.9 | 3 | 3 | 9.26E+04 | 1 | 1 | 1 | | 38982 | Pyruvate dehydrogenase E1 component subunit beta mitochondrial OS=Rattus norvegicus OX=10116 GN=Pdhb PE=1 SV=2 |
| 142 | 15 | tr\|A0A0G2 | 21.9 | 2 | 2 | 9.26E+04 | 1 | 1 | 1 | | 46193 | Pyruvate dehydrogenase E1 component subunit beta OS=Rattus norvegicus OX=10116 GN=Pdhb PE=1 SV=1 |
| 127 | 524 | tr\|D4A054 | 21.54 | 0 | 0 | 2.73E+06 | 1 | 1 | 1 | | 341403 | RAN-binding protein 2 OS=Rattus norvegicus OX=10116 GN=Ranbp2 PE=1 SV=2 |
| 127 | 525 | tr\|M0R3M | 21.54 | 0 | 0 | 2.73E+06 | 1 | 1 | 1 | | 344396 | RAN-binding protein 2 OS=Rattus norvegicus OX=10116 GN=Ranbp2 PE=1 SV=1 |
| 176 | 12109 | Q6SKG1\|A | 21.3 | 1 | 1 | 5.59E+05 | 1 | 1 | 1 | Formylatio | 65713 | Acyl-coenzyme A synthetase ACSM3 mitochondrial OS=Rattus norvegicus OX=10116 GN=Acsm3 PE=2 SV=1 |
| 128 | 738 | tr\|A0A0G2 | 21.17 | 0 | 0 | | 1 | 0 | 1 | | 276393 | Acetyl-CoA carboxylase beta OS=Rattus norvegicus OX=10116 GN=Acacb PE=1 SV=1 |
| 128 | 739 | tr\|D3ZBE2 | 21.17 | 0 | 0 | | 1 | 0 | 1 | | 275967 | Acetyl-CoA carboxylase beta OS=Rattus norvegicus OX=10116 GN=Acacb PE=1 SV=3 |
| 128 | 740 | tr\|E9PSQ0 | 21.17 | 0 | 0 | | 1 | 0 | 1 | | 276255 | Acetyl-CoA carboxylase beta OS=Rattus norvegicus OX=10116 GN=Acacb PE=1 SV=2 |
| 128 | 426 | tr\|O70151 | 21.17 | 0 | 0 | | 1 | 0 | 1 | | 276097 | Acetyl-CoA carboxylase OS=Rattus norvegicus OX=10116 GN=Acacb PE=2 SV=1 |
| 128 | 737 | tr\|A0A0G2 | 21.17 | 0 | 0 | | 1 | 0 | 1 | | 275755 | Acetyl-CoA carboxylase beta OS=Rattus norvegicus OX=10116 GN=Acacb PE=1 SV=1 |
| 132 | 307 | P11497\|AC | 21.17 | 0 | 0 | | 1 | 0 | 1 | | 265191 | Acetyl-CoA carboxylase 1 OS=Rattus norvegicus OX=10116 GN=Acaca PE=1 SV=1 |
| 177 | 4381 | tr\|A0A0G2 | 20.92 | 1 | 1 | 8.42E+04 | 1 | 1 | 1 | | 63436 | Malic enzyme OS=Rattus norvegicus OX=10116 GN=Me3 PE=3 SV=1 |
| 177 | 1308 | P13697\|M | 20.92 | 1 | 1 | 8.42E+04 | 1 | 1 | 1 | | 64003 | NADP-dependent malic enzyme OS=Rattus norvegicus OX=10116 GN=Me1 PE=1 SV=2 |
| 177 | 4463 | tr\|D3ZJH9 | 20.92 | 1 | 1 | 8.42E+04 | 1 | 1 | 1 | | 65352 | Malic enzyme OS=Rattus norvegicus OX=10116 GN=Me2 PE=1 SV=1 |
| 177 | 4464 | tr\|A0A0G2 | 20.92 | 1 | 1 | 8.42E+04 | 1 | 1 | 1 | | 66310 | Malic enzyme OS=Rattus norvegicus OX=10116 GN=Me2 PE=1 SV=1 |
| 177 | 4382 | tr\|F1M5N4 | 20.92 | 1 | 1 | 8.42E+04 | 1 | 1 | 1 | | 67209 | Malic enzyme OS=Rattus norvegicus OX=10116 GN=Me3 PE=3 SV=2 |
| 90 | 410 | tr\|F1LNF0\| | 20.47 | 0 | 0 | 2.57E+05 | 1 | 1 | 4 | | 228912 | Myosin heavy chain 14 OS=Rattus norvegicus OX=10116 GN=Myh14 PE=1 SV=1 |
| 124 | 298 | tr\|Q5RKL4 | 20.39 | 1 | 1 | 0 | 1 | 1 | 2 | | 95977 | Dimethylglycine dehydrogenase OS=Rattus norvegicus OX=10116 GN=Dmgdh PE=1 SV=1 |
| 124 | 299 | Q63342\|M | 20.39 | 1 | 1 | 0 | 1 | 1 | 2 | | 96047 | Dimethylglycine dehydrogenase mitochondrial OS=Rattus norvegicus OX=10116 GN=Dmgdh PE=1 SV=1 |
| 124 | 303 | tr\|A0A0G2 | 20.39 | 1 | 1 | 0 | 1 | 1 | 2 | | 98864 | Dimethylglycine dehydrogenase mitochondrial OS=Rattus norvegicus OX=10116 GN=Dmgdh PE=1 SV=1 |
