## Supplementary material for "Permeability transition pore-related changes in the proteome and channel activity of ATP synthase dimers and monomers": RHM

| Protein Gr | Protein ID | Accession | -10lgP | Coverage ( | Coverage ( | IBAQ | #Peptides | #Unique | #Spec Sam | PTM | Avg. Mass | Description |
|---|---|---|---|---|---|---|---|---|---|---|---|---|
| 5 | 5 | P10719\|AT | 293.04 | 21 | 21 | 3.49E+07 | 44 | 44 | 98 | Oxidation ( | 56354 | ATP synthase subunit beta mitochondrial OS=Rattus norvegicus OX=10116 GN=Atp5f1b PE=1 SV=2 |
| 5 | 6 | tr\|G3V6D3 | 293.04 | 21 | 21 | 3.49E+07 | 44 | 44 | 98 | Oxidation ( | 56345 | ATP synthase subunit beta OS=Rattus norvegicus OX=10116 GN=Atp5f1b PE=1 SV=1 |
| 1 | 20 | Q66HF1\|N | 261.24 | 25 | 25 | 4.70E+07 | 57 | 56 | 134 | Carbamido | 79412 | NADH-ubiquinone oxidoreductase 75 kDa subunit mitochondrial OS=Rattus norvegicus OX=10116 GN=Ndufs1 PE=1 SV=1 |
| 6 | 58 | P20788\|U | 241.84 | 41 | 41 | 3.37E+07 | 31 | 31 | 81 | Oxidation ( | 29446 | Cytochrome b-c1 complex subunit Rieske mitochondrial OS=Rattus norvegicus OX=10116 GN=Uqcrfs1 PE=1 SV=2 |
| 2 | 37 | P19234\|N | 236.79 | 46 | 46 | 7.04E+07 | 36 | 36 | 109 | Formylatio | 27378 | NADH dehydrogenase [ubiquinone] flavoprotein 2 mitochondrial OS=Rattus norvegicus OX=10116 GN=Ndufv2 PE=1 SV=2 |
| 15 | 59 | tr\|D3ZFQ8 | 227.59 | 24 | 24 | 3.82E+07 | 19 | 19 | 55 | Oxidation ( | 35435 | Cytochrome c-1 OS=Rattus norvegicus OX=10116 GN=Cyc1 PE=1 SV=3 |
| 21 | 212 | tr\|B2RYS8 | 226.48 | 30 | 30 | 1.73E+07 | 18 | 18 | 41 | Oxidation ( | 21959 | NADH dehydrogenase [ubiquinone] 1 beta subcomplex subunit 8 mitochondrial OS=Rattus norvegicus OX=10116 GN=Ndufb8 PE=1 SV=1 |
| 9 | 89 | tr\|L0L4L8\| | 222.24 | 24 | 24 | 1.96E+06 | 28 | 1 | 66 | | 42470 | Cytochrome b (Fragment) OS=Rattus norvegicus OX=10116 GN=cytb PE=3 SV=1 |
| 9 | 91 | tr\|A0A140 | 222.24 | 24 | 24 | 1.96E+06 | 28 | 1 | 66 | | 42574 | Cytochrome b (Fragment) OS=Rattus norvegicus OX=10116 PE=3 SV=1 |
| 9 | 92 | tr\|A0A140 | 222.24 | 24 | 24 | 1.96E+06 | 28 | 1 | 66 | | 42588 | Cytochrome b (Fragment) OS=Rattus norvegicus OX=10116 PE=3 SV=1 |
| 9 | 95 | tr\|A0A385 | 222.24 | 24 | 24 | 1.96E+06 | 28 | 1 | 66 | | 42766 | Cytochrome b (Fragment) OS=Rattus norvegicus OX=10116 PE=3 SV=1 |
| 9 | 96 | tr\|A0A3Q8 | 222.24 | 24 | 24 | 1.96E+06 | 28 | 1 | 66 | | 42712 | Cytochrome b (Fragment) OS=Rattus norvegicus OX=10116 GN=Cytb PE=3 SV=1 |
| 9 | 97 | tr\|A0A3Q8 | 222.24 | 24 | 24 | 1.96E+06 | 28 | 1 | 66 | | 42622 | Cytochrome b (Fragment) OS=Rattus norvegicus OX=10116 GN=Cytb PE=3 SV=1 |
| 9 | 100 | tr\|A0A3S6 | 222.24 | 24 | 24 | 1.96E+06 | 28 | 1 | 66 | | 42698 | Cytochrome b (Fragment) OS=Rattus norvegicus OX=10116 GN=Cytb PE=3 SV=1 |
| 9 | 102 | tr\|A0A220 | 222.24 | 23 | 23 | 1.96E+06 | 28 | 1 | 66 | | 43018 | Cytochrome b OS=Rattus norvegicus OX=10116 PE=3 SV=1 |
| 9 | 103 | tr\|D6NSP2 | 222.24 | 23 | 23 | 1.96E+06 | 28 | 1 | 66 | | 42945 | Cytochrome b (Fragment) OS=Rattus norvegicus OX=10116 GN=cytb PE=3 SV=1 |
| 9 | 104 | tr\|D6NSS6 | 222.24 | 23 | 23 | 1.96E+06 | 28 | 1 | 66 | | 42993 | Cytochrome b (Fragment) OS=Rattus norvegicus OX=10116 GN=cytb PE=3 SV=1 |
| 9 | 106 | tr\|D6NSQ3 | 222.24 | 23 | 23 | 1.96E+06 | 28 | 1 | 66 | | 43012 | Cytochrome b (Fragment) OS=Rattus norvegicus OX=10116 GN=cytb PE=3 SV=1 |
| 9 | 107 | tr\|D6NSR3 | 222.24 | 23 | 23 | 1.96E+06 | 28 | 1 | 66 | | 42986 | Cytochrome b (Fragment) OS=Rattus norvegicus OX=10116 GN=cytb PE=3 SV=1 |
| 9 | 110 | tr\|D6NSR8 | 222.24 | 23 | 23 | 1.96E+06 | 28 | 1 | 66 | | 42998 | Cytochrome b (Fragment) OS=Rattus norvegicus OX=10116 GN=cytb PE=3 SV=1 |
| 9 | 113 | tr\|A0A0A1 | 222.24 | 23 | 23 | 1.96E+06 | 28 | 1 | 66 | | 42989 | Cytochrome b OS=Rattus norvegicus OX=10116 GN=CYTB PE=3 SV=1 |
| 9 | 114 | tr\|D6NSS7 | 222.24 | 23 | 23 | 1.96E+06 | 28 | 1 | 66 | | 42948 | Cytochrome b (Fragment) OS=Rattus norvegicus OX=10116 GN=cytb PE=3 SV=1 |
| 9 | 115 | tr\|Q8SEY9 | 222.24 | 23 | 23 | 1.96E+06 | 28 | 1 | 66 | | 43015 | Cytochrome b OS=Rattus norvegicus OX=10116 GN=cytb PE=3 SV=1 |
| 9 | 116 | tr\|D6NSR7 | 222.24 | 23 | 23 | 1.96E+06 | 28 | 1 | 66 | | 42982 | Cytochrome b (Fragment) OS=Rattus norvegicus OX=10116 GN=cytb PE=3 SV=1 |
| 9 | 117 | tr\|A0A220 | 222.24 | 23 | 23 | 1.96E+06 | 28 | 1 | 66 | | 42968 | Cytochrome b OS=Rattus norvegicus OX=10116 PE=3 SV=1 |
| 9 | 120 | tr\|D6NSQ | 222.24 | 23 | 23 | 1.96E+06 | 28 | 1 | 66 | | 42968 | Cytochrome b (Fragment) OS=Rattus norvegicus OX=10116 GN=cytb PE=3 SV=1 |
| 9 | 121 | tr\|Q5UAI7 | 222.24 | 23 | 23 | 1.96E+06 | 28 | 1 | 66 | | 42998 | Cytochrome b OS=Rattus norvegicus OX=10116 GN=CYTB PE=3 SV=1 |
| 9 | 122 | tr\|D6NSR2 | 222.24 | 23 | 23 | 1.96E+06 | 28 | 1 | 66 | | 43073 | Cytochrome b (Fragment) OS=Rattus norvegicus OX=10116 GN=cytb PE=3 SV=1 |
| 9 | 123 | tr\|A0A0S1 | 222.24 | 23 | 23 | 1.96E+06 | 28 | 1 | 66 | | 43002 | Cytochrome b OS=Rattus norvegicus OX=10116 GN=CYTB PE=3 SV=1 |
| 9 | 125 | tr\|Q8HIC4 | 222.24 | 23 | 23 | 1.96E+06 | 28 | 1 | 66 | | 43012 | Cytochrome b OS=Rattus norvegicus OX=10116 GN=Mt-cyb PE=3 SV=1 |
| 9 | 126 | tr\|A0A220 | 222.24 | 23 | 23 | 1.96E+06 | 28 | 1 | 66 | | 42968 | Cytochrome b OS=Rattus norvegicus OX=10116 PE=3 SV=1 |
| 9 | 128 | tr\|A0A0S1 | 222.24 | 23 | 23 | 1.96E+06 | 28 | 1 | 66 | | 43016 | Cytochrome b OS=Rattus norvegicus OX=10116 GN=CYTB PE=3 SV=1 |
| 9 | 129 | tr\|R9TKN1 | 222.24 | 23 | 23 | 1.96E+06 | 28 | 1 | 66 | | 43012 | Cytochrome b OS=Rattus norvegicus OX=10116 GN=CYTB PE=3 SV=1 |
| 9 | 130 | P00159\|CY | 222.24 | 23 | 23 | 1.96E+06 | 28 | 1 | 66 | | 43012 | Cytochrome b OS=Rattus norvegicus OX=10116 GN=Mt-Cyb PE=3 SV=3 |
| 9 | 131 | tr\|L0N311 | 222.24 | 23 | 23 | 1.96E+06 | 28 | 1 | 66 | | 43045 | Cytochrome b (Fragment) OS=Rattus norvegicus OX=10116 GN=cytb PE=3 SV=1 |
| 9 | 132 | tr\|A0A220 | 222.24 | 23 | 23 | 1.96E+06 | 28 | 1 | 66 | | 42970 | Cytochrome b OS=Rattus norvegicus OX=10116 PE=3 SV=1 |
| 9 | 133 | tr\|H2KXA0 | 222.24 | 23 | 23 | 1.96E+06 | 28 | 1 | 66 | | 42978 | Cytochrome b (Fragment) OS=Rattus norvegicus OX=10116 GN=cytb PE=3 SV=1 |
| 9 | 134 | tr\|A0A096 | 222.24 | 23 | 23 | 1.96E+06 | 28 | 1 | 66 | | 43042 | Cytochrome b (Fragment) OS=Rattus norvegicus OX=10116 PE=3 SV=1 |
| 9 | 135 | tr\|D6NSP5 | 222.24 | 23 | 23 | 1.96E+06 | 28 | 1 | 66 | | 43012 | Cytochrome b (Fragment) OS=Rattus norvegicus OX=10116 GN=cytb PE=3 SV=1 |
| 9 | 86 | tr\|A0A411 | 222.24 | 24 | 24 | 1.96E+06 | 28 | 1 | 66 | | 41017 | Cytochrome b (Fragment) OS=Rattus norvegicus OX=10116 GN=Cytb PE=3 SV=1 |
| 9 | 88 | tr\|A0A411 | 222.24 | 24 | 24 | 1.96E+06 | 28 | 1 | 66 | | 42238 | Cytochrome b (Fragment) OS=Rattus norvegicus OX=10116 GN=Cytb PE=3 SV=1 |
| 9 | 108 | tr\|D6NSP4 | 222.24 | 23 | 23 | 1.96E+06 | 28 | 1 | 66 | | 43016 | Cytochrome b (Fragment) OS=Rattus norvegicus OX=10116 GN=cytb PE=3 SV=1 |
| 9 | 124 | tr\|D6NSR1 | 222.24 | 23 | 23 | 1.96E+06 | 28 | 1 | 66 | | 43002 | Cytochrome b (Fragment) OS=Rattus norvegicus OX=10116 GN=cytb PE=3 SV=1 |
| 12 | 99 | tr\|A0A385 | 218.35 | 24 | 24 | 2.22E+05 | 28 | 1 | 63 | | 42751 | Cytochrome b (Fragment) OS=Rattus norvegicus OX=10116 PE=3 SV=1 |
| 4 | 26 | Q641Y2\|N | 217.68 | 29 | 29 | 4.21E+07 | 41 | 41 | 106 | Oxidation ( | 52562 | NADH dehydrogenase [ubiquinone] iron-sulfur protein 2 mitochondrial OS=Rattus norvegicus OX=10116 GN=Ndufs2 PE=1 SV=1 |
| 10 | 25 | Q68FY0\|Q | 216.29 | 23 | 23 | 2.10E+07 | 32 | 32 | 64 | Carbamido | 52849 | Cytochrome b-c1 complex subunit 1 mitochondrial OS=Rattus norvegicus OX=10116 GN=Uqcrc1 PE=1 SV=1 |
| 7 | 16 | P02563\|M | 214.15 | 15 | 15 | 1.13E+07 | 44 | 44 | 71 | Oxidation (M) | | |
| 23 | 60 | tr\|D4A565 | 207.02 | 37 | 37 | 1.05E+07 | 20 | 19 | 35 | | 21664 | NADH dehydrogenase (Ubiquinone) 1 beta subcomplex 5 (Predicted) isoform CRA_b OS=Rattus norvegicus OX=10116 GN=Ndufb5 PE=1 SV=1 |
| 3 | 50 | P32551\|Q | 206.62 | 24 | 24 | 6.90E+07 | 35 | 33 | 106 | Oxidation ( | 48396 | Cytochrome b-c1 complex subunit 2 mitochondrial OS=Rattus norvegicus OX=10116 GN=Uqcrc2 PE=1 SV=2 |
| 8 | 8 | P15999\|AT | 199.95 | 27 | 27 | 1.83E+07 | 29 | 29 | 71 | Oxidation ( | 59754 | ATP synthase subunit alpha mitochondrial OS=Rattus norvegicus OX=10116 GN=Atp5f1a PE=1 SV=2 |
| 8 | 9 | tr\|F1LP05\| | 199.95 | 27 | 27 | 1.83E+07 | 29 | 29 | 71 | Oxidation ( | 59813 | ATP synthase subunit alpha OS=Rattus norvegicus OX=10116 GN=Atp5f1a PE=1 SV=1 |
| 30 | 248 | tr\|A9UMW | 195.52 | 42 | 42 | 1.34E+07 | 11 | 11 | 29 | Oxidation ( | 9255 | Ndufa3 protein (Fragment) OS=Rattus norvegicus OX=10116 GN=Ndufa3 PE=2 SV=1 |
| 30 | 250 | tr\|A0A0G2 | 195.52 | 38 | 38 | 1.34E+07 | 11 | 11 | 29 | Oxidation ( | 10170 | NADH:ubiquinone oxidoreductase subunit A3 OS=Rattus norvegicus OX=10116 GN=Ndufa3 PE=1 SV=1 |
| 30 | 249 | tr\|M0RB63 | 195.52 | 42 | 42 | 1.34E+07 | 11 | 11 | 29 | Oxidation ( | 9372 | RCG63041 OS=Rattus norvegicus OX=10116 GN=LOC684509 PE=4 SV=1 |
| 26 | 14 | Q6AXV4\|S | 189.34 | 18 | 18 | 4.89E+06 | 17 | 17 | 31 | | 51960 | Sorting and assembly machinery component 50 homolog OS=Rattus norvegicus OX=10116 GN=Samm50 PE=1 SV=1 |
| 13 | 74 | tr\|D3ZG43 | 188.8 | 41 | 41 | 1.95E+07 | 27 | 27 | 58 | | 30226 | NADH dehydrogenase (Ubiquinone) Fe-S protein 3 (Predicted) isoform CRA_c OS=Rattus norvegicus OX=10116 GN=Ndufs3 PE=1 SV=1 |
| 16 | 136 | tr\|Q9G898 | 187.73 | 23 | 23 | 0 | 26 | 1 | 54 | | 42830 | Cytochrome b OS=Rattus norvegicus OX=10116 PE=2 SV=1 |
| 14 | 49 | Q5BK63\|N | 185.88 | 29 | 29 | 1.15E+07 | 25 | 25 | 56 | Oxidation ( | 42559 | NADH dehydrogenase [ubiquinone] 1 alpha subcomplex subunit 9 mitochondrial OS=Rattus norvegicus OX=10116 GN=Ndufa9 PE=1 SV=2 |
| 17 | 77 | tr\|Q5PQZ9 | 185.82 | 44 | 44 | 2.06E+07 | 16 | 16 | 48 | | 14359 | NADH dehydrogenase [ubiquinone] 1 subunit C2 OS=Rattus norvegicus OX=10116 GN=Ndufc2 PE=1 SV=1 |
| 38 | 200 | P11240\|C | 183.83 | 38 | 38 | 3.16E+06 | 11 | 11 | 20 | | 16130 | Cytochrome c oxidase subunit 5A mitochondrial OS=Rattus norvegicus OX=10116 GN=Cox5a PE=1 SV=1 |
| 22 | 137 | tr\|Q06QK5 | 182.41 | 11 | 11 | 1.24E+07 | 19 | 19 | 38 | Oxidation ( | 68606 | NADH-ubiquinone oxidoreductase chain 5 OS=Rattus norvegicus OX=10116 GN=ND5 PE=3 SV=1 |
| 22 | 138 | P11661\|N | 182.41 | 11 | 11 | 1.24E+07 | 19 | 19 | 38 | Oxidation ( | 68618 | NADH-ubiquinone oxidoreductase chain 5 OS=Rattus norvegicus OX=10116 GN=Mtnd5 PE=3 SV=3 |

| | | | | | | | | | | | | |
|---|---|---|---|---|---|---|---|---|---|---|---|---|
| 22 | 139 | tr\|Q06QG( | 182.41 | 11 | 11 | 1.24E+07 | 19 | 19 | 38 | Oxidation ( | 68588 | NADH-ubiquinone oxidoreductase chain 5 OS=Rattus norvegicus OX=10116 GN=ND5 PE=3 SV=1 |
| 22 | 140 | tr\|Q06QA1 | 182.41 | 11 | 11 | 1.24E+07 | 19 | 19 | 38 | Oxidation ( | 68574 | NADH-ubiquinone oxidoreductase chain 5 OS=Rattus norvegicus OX=10116 GN=ND5 PE=3 SV=1 |
| 22 | 141 | tr\|Q8SEZ0 | 182.41 | 11 | 11 | 1.24E+07 | 19 | 19 | 38 | Oxidation ( | 68618 | NADH-ubiquinone oxidoreductase chain 5 OS=Rattus norvegicus OX=10116 GN=Mt-nd5 PE=3 SV=1 |
| 22 | 142 | tr\|A0A096 | 182.41 | 11 | 11 | 1.24E+07 | 19 | 19 | 38 | Oxidation ( | 68584 | NADH-ubiquinone oxidoreductase chain 5 OS=Rattus norvegicus OX=10116 GN=ND5 PE=3 SV=1 |
| 28 | 161 | tr\|A0A0G2 | 179.77 | 42 | 42 | 1.56E+07 | 16 | 16 | 30 | Carbamido | 19965 | NADH dehydrogenase [ubiquinone] 1 alpha subcomplex subunit 8 OS=Rattus norvegicus OX=10116 GN=Ndufa8 PE=1 SV=1 |
| 11 | 171 | Q561S0\|N | 177.4 | 32 | 32 | 1.43E+07 | 25 | 23 | 64 | Oxidation (M) | | |
| 35 | 160 | tr\|D4A7L4 | 172.5 | 27 | 27 | 1.19E+07 | 10 | 10 | 22 | Oxidation ( | 17634 | NADH dehydrogenase (Ubiquinone) 1 beta subcomplex 11 (Predicted) OS=Rattus norvegicus OX=10116 GN=Ndufb11 PE=1 SV=1 |
| 29 | 45 | P00507\|A/ | 169.11 | 17 | 17 | 4.51E+06 | 16 | 16 | 29 | Oxidation ( | 47314 | Aspartate aminotransferase mitochondrial OS=Rattus norvegicus OX=10116 GN=Got2 PE=1 SV=2 |
| 24 | 199 | tr\|B0BNE6 | 168.87 | 25 | 25 | 8.65E+06 | 16 | 16 | 33 | Carbamido | 23970 | NADH dehydrogenase (Ubiquinone) Fe-S protein 8 (Predicted) isoform CRA_a OS=Rattus norvegicus OX=10116 GN=Ndufs8 PE=1 SV=1 |
| 32 | 252 | tr\|D3ZF13 | 168.64 | 23 | 23 | 7.49E+06 | 13 | 13 | 25 | Oxidation ( | 17514 | Acyl carrier protein OS=Rattus norvegicus OX=10116 GN=Ndufab1 PE=1 SV=1 |
| 39 | 211 | tr\|F1LXA0\| | 168.55 | 29 | 29 | 7.78E+06 | 10 | 10 | 20 | | 17178 | NADH dehydrogenase [ubiquinone] 1 alpha subcomplex subunit 12 OS=Rattus norvegicus OX=10116 GN=Ndufa12 PE=1 SV=2 |
| 37 | 198 | Q80W89\|N | 167.6 | 23 | 23 | 1.38E+07 | 9 | 9 | 21 | | 14854 | NADH dehydrogenase [ubiquinone] 1 alpha subcomplex subunit 11 OS=Rattus norvegicus OX=10116 GN=Ndufa11 PE=2 SV=1 |
| 20 | 33 | tr\|S5RZM8 | 164.42 | 40 | 40 | 1.02E+07 | 20 | 20 | 42 | Oxidation ( | 25958 | Cytochrome c oxidase subunit 2 OS=Rattus norvegicus OX=10116 GN=COX2 PE=3 SV=1 |
| 20 | 34 | P00406\|CC | 164.42 | 40 | 40 | 1.02E+07 | 20 | 20 | 42 | Oxidation ( | 25928 | Cytochrome c oxidase subunit 2 OS=Rattus norvegicus OX=10116 GN=Mtco2 PE=1 SV=3 |
| 20 | 28 | tr\|Q5UAJ6 | 164.42 | 40 | 40 | 1.02E+07 | 20 | 20 | 42 | Oxidation ( | 25942 | Cytochrome c oxidase subunit 2 OS=Rattus norvegicus OX=10116 GN=COX2 PE=3 SV=1 |
| 20 | 35 | tr\|Q8SEZ5 | 164.42 | 40 | 40 | 1.02E+07 | 20 | 20 | 42 | Oxidation ( | 25928 | Cytochrome c oxidase subunit 2 OS=Rattus norvegicus OX=10116 GN=Mt-co2 PE=1 SV=1 |
| 20 | 36 | tr\|A0A097 | 164.42 | 40 | 40 | 1.02E+07 | 20 | 20 | 42 | Oxidation ( | 25894 | Cytochrome c oxidase subunit 2 OS=Rattus norvegicus OX=10116 GN=COX2 PE=3 SV=1 |
| 18 | 313 | tr\|D3ZZ21 | 160.18 | 52 | 52 | 1.08E+07 | 16 | 16 | 44 | Oxidation ( | 15638 | NADH dehydrogenase (Ubiquinone) 1 beta subcomplex 6 (Predicted) OS=Rattus norvegicus OX=10116 GN=Ndufb6 PE=1 SV=1 |
| 34 | 827 | tr\|D4A3V2 | 157.67 | 44 | 44 | 4.88E+06 | 19 | 19 | 24 | | 15224 | NADH dehydrogenase [ubiquinone] 1 alpha subcomplex subunit 6 OS=Rattus norvegicus OX=10116 GN=Ndufa6 PE=1 SV=1 |
| 19 | 80 | tr\|Q5XIH3 | 152.05 | 19 | 19 | 1.26E+07 | 19 | 19 | 43 | Oxidation ( | 50731 | NADH dehydrogenase [ubiquinone] flavoprotein 1 mitochondrial OS=Rattus norvegicus OX=10116 GN=Ndufv1 PE=1 SV=1 |
| 53 | 65 | P31399\|AT | 151.97 | 29 | 29 | 2.29E+06 | 7 | 7 | 12 | | 18763 | ATP synthase subunit d mitochondrial OS=Rattus norvegicus OX=10116 GN=Atp5pd PE=1 SV=3 |
| 25 | 474 | tr\|F1LN92 | 148.92 | 12 | 12 | 6.05E+06 | 15 | 15 | 31 | Oxidation ( | 89352 | AFG3-like matrix AAA peptidase subunit 2 OS=Rattus norvegicus OX=10116 GN=Afg3l2 PE=1 SV=1 |
| 44 | 253 | P10817\|CX | 147.19 | 27 | 27 | 6.52E+06 | 8 | 8 | 17 | | 10487 | Cytochrome c oxidase subunit 6A2 mitochondrial (Fragment) OS=Rattus norvegicus OX=10116 GN=Cox6a2 PE=1 SV=3 |
| 44 | 254 | tr\|G3V8M4 | 147.19 | 26 | 26 | 6.52E+06 | 8 | 8 | 17 | | 10802 | Cytochrome c oxidase subunit 6A mitochondrial OS=Rattus norvegicus OX=10116 GN=Cox6a2 PE=3 SV=1 |
| 27 | 172 | tr\|D4A0T0 | 142.48 | 27 | 27 | 2.09E+07 | 10 | 10 | 31 | | 20859 | NADH:ubiquinone oxidoreductase subunit B10 OS=Rattus norvegicus OX=10116 GN=Ndufb10 PE=1 SV=1 |
| 50 | 2 | Q01205\|O | 141.91 | 17 | 17 | 4.05E+06 | 6 | 6 | 13 | Oxidation ( | 48925 | Dihydrolipoyllysine-residue succinyltransferase component of 2-oxoglutarate dehydrogenase complex mitochondrial OS=Rattus norvegicus OX=10116 GN=Dlst PE=1 SV=2 |
| 50 | 1 | tr\|G3V6P2 | 141.91 | 17 | 17 | 4.05E+06 | 6 | 6 | 13 | Oxidation ( | 48899 | Dihydrolipoamide S-succinyltransferase (E2 component of 2-oxo-glutarate complex) isoform CRA_a OS=Rattus norvegicus OX=10116 GN=Dlst PE=1 SV=1 |
| 54 | 251 | P80432\|CC | 141.51 | 21 | 21 | 3.44E+06 | 4 | 4 | 12 | | 7375 | Cytochrome c oxidase subunit 7C mitochondrial OS=Rattus norvegicus OX=10116 GN=Cox7c PE=1 SV=2 |
| 33 | 56 | tr\|F1LPG5 | 139.49 | 35 | 35 | 1.09E+07 | 9 | 9 | 24 | | 15064 | NADH:ubiquinone oxidoreductase subunit B4 OS=Rattus norvegicus OX=10116 GN=Ndufb4 PE=1 SV=1 |
| 33 | 57 | tr\|F1M7T1 | 139.49 | 35 | 35 | 1.09E+07 | 9 | 9 | 24 | | 15160 | NADH dehydrogenase (ubiquinone) 1 beta subcomplex 4 OS=Rattus norvegicus OX=10116 GN=LOC100361934 PE=4 SV=2 |
| 40 | 169 | tr\|A9UMV9 | 134.78 | 38 | 38 | 2.64E+07 | 7 | 7 | 20 | Oxidation ( | 12500 | NADH:ubiquinone oxidoreductase subunit A7 OS=Rattus norvegicus OX=10116 GN=Ndufa7 PE=1 SV=1 |
| 42 | 219 | tr\|D3ZS58 | 130.27 | 19 | 19 | 3.74E+06 | 7 | 7 | 18 | | 10845 | NADH dehydrogenase [ubiquinone] 1 alpha subcomplex subunit 2 OS=Rattus norvegicus OX=10116 GN=Ndufa2 PE=1 SV=1 |
| 36 | 29 | Q60587\|EC | 130.16 | 15 | 15 | 2.90E+06 | 14 | 14 | 21 | | 51414 | Trifunctional enzyme subunit beta mitochondrial OS=Rattus norvegicus OX=10116 GN=Hadhb PE=1 SV=1 |
| 31 | 72 | tr\|D3ZE15 | 126.31 | 38 | 38 | 4.35E+06 | 11 | 11 | 25 | Oxidation ( | 16777 | NADH:ubiquinone oxidoreductase subunit A13 OS=Rattus norvegicus OX=10116 GN=Ndufa13 PE=1 SV=1 |
| 49 | 318 | Q5M9I5\|Q | 122.61 | 21 | 21 | 4.60E+06 | 6 | 6 | 14 | | 10424 | Cytochrome b-c1 complex subunit 6 mitochondrial OS=Rattus norvegicus OX=10116 GN=Uqcrh PE=3 SV=1 |
| 43 | 10 | tr\|A0A0G2 | 120.33 | 12 | 12 | 1.86E+06 | 11 | 11 | 17 | | 86230 | MICOS complex subunit MIC60 OS=Rattus norvegicus OX=10116 GN=Immt PE=1 SV=1 |
| 46 | 532 | P67779\|PH | 119.94 | 17 | 17 | 1.96E+06 | 8 | 8 | 15 | | 29820 | Prohibitin OS=Rattus norvegicus OX=10116 GN=Phb PE=1 SV=1 |
| 48 | 3899 | tr\|B5DEL8 | 114.92 | 22 | 22 | 2.27E+06 | 6 | 6 | 15 | | 12700 | NADH dehydrogenase (Ubiquinone) Fe-S protein 5 OS=Rattus norvegicus OX=10116 GN=Ndufs5 PE=1 SV=1 |
| 48 | 3900 | tr\|A0A0G2 | 114.92 | 22 | 22 | 2.27E+06 | 6 | 6 | 15 | | 12730 | Uncharacterized protein OS=Rattus norvegicus OX=10116 PE=4 SV=1 |
| 68 | 17 | Q9ER34\|A | 109.74 | 3 | 3 | 1.09E+06 | 4 | 4 | 7 | | 85433 | Aconitate hydratase mitochondrial OS=Rattus norvegicus OX=10116 GN=Aco2 PE=1 SV=2 |
| 61 | 238 | Q5XIF3\|ND | 104.1 | 15 | 15 | 3.73E+06 | 4 | 4 | 9 | | 19741 | NADH dehydrogenase [ubiquinone] iron-sulfur protein 4 mitochondrial OS=Rattus norvegicus OX=10116 GN=Ndufs4 PE=1 SV=1 |
| 56 | 3906 | tr\|D3ZCZ9 | 103.87 | 21 | 21 | 1.37E+06 | 4 | 4 | 10 | | 13040 | NADH dehydrogenase [ubiquinone] iron-sulfur protein 6 mitochondrial OS=Rattus norvegicus OX=10116 GN=LOC100912599 PE=1 SV=1 |
| 66 | 61 | Q63704\|CI | 98.66 | 3 | 3 | 6.64E+05 | 3 | 3 | 7 | | 88217 | Carnitine O-palmitoyltransferase 1 muscle isoform OS=Rattus norvegicus OX=10116 GN=Cpt1b PE=1 SV=1 |
| 52 | 335 | Q5XIH7\|PH | 98.45 | 14 | 14 | 5.93E+06 | 9 | 9 | 12 | | 33312 | Prohibitin-2 OS=Rattus norvegicus OX=10116 GN=Phb2 PE=1 SV=1 |
| 55 | 3893 | tr\|B2RYS2 | 93.22 | 28 | 28 | 3.07E+06 | 8 | 7 | 11 | | 13559 | Cytochrome b-c1 complex subunit 7 OS=Rattus norvegicus OX=10116 GN=Uqcrb PE=1 SV=1 |
| 77 | 245 | tr\|C8CHS6 | 91.52 | 17 | 17 | 6.51E+05 | 2 | 2 | 5 | | 7988 | Mitochondrial superoxide dismutase 2 (Fragment) OS=Rattus norvegicus OX=10116 PE=2 SV=1 |
| 77 | 71 | P07895\|SC | 91.52 | 5 | 5 | 6.51E+05 | 2 | 2 | 5 | | 24674 | Superoxide dismutase [Mn] mitochondrial OS=Rattus norvegicus OX=10116 GN=Sod2 PE=1 SV=2 |
| 71 | 3921 | tr\|Q06QD9 | 90.75 | 6 | 6 | 9.06E+05 | 3 | 3 | 7 | | 18943 | NADH-ubiquinone oxidoreductase chain 6 OS=Rattus norvegicus OX=10116 GN=ND6 PE=3 SV=1 |
| 71 | 3922 | tr\|Q7HKW | 90.75 | 6 | 6 | 9.06E+05 | 3 | 3 | 7 | | 18957 | NADH-ubiquinone oxidoreductase chain 6 OS=Rattus norvegicus OX=10116 GN=NADH6 PE=3 SV=1 |
| 71 | 3930 | tr\|Q35733 | 90.75 | 14 | 14 | 9.06E+05 | 3 | 3 | 7 | | 8389 | NADH-ubiquinone oxidoreductase chain 6 (Fragment) OS=Rattus norvegicus OX=10116 PE=3 SV=1 |
| 57 | 168 | P19511\|AT | 90.5 | 9 | 9 | 3.08E+06 | 3 | 3 | 10 | | 28869 | ATP synthase F(0) complex subunit B1 mitochondrial OS=Rattus norvegicus OX=10116 GN=Atp5pb PE=1 SV=1 |
| 45 | 3901 | tr\|Q8SEZ8 | 89.63 | 7 | 7 | 5.21E+06 | 7 | 7 | 17 | | 36133 | NADH-ubiquinone oxidoreductase chain 1 (Fragment) OS=Rattus norvegicus OX=10116 GN=NADH1 PE=3 SV=1 |
| 45 | 3902 | P03889\|NU | 89.63 | 7 | 7 | 5.21E+06 | 7 | 7 | 17 | | 36145 | NADH-ubiquinone oxidoreductase chain 1 OS=Rattus norvegicus OX=10116 GN=Mtnd1 PE=1 SV=3 |
| 45 | 3903 | tr\|Q8HID1 | 89.63 | 7 | 7 | 5.21E+06 | 7 | 7 | 17 | | 36145 | NADH-ubiquinone oxidoreductase chain 1 OS=Rattus norvegicus OX=10116 GN=Mt-nd1 PE=3 SV=1 |
| 45 | 3904 | tr\|D2E6L7 | 89.63 | 7 | 7 | 5.21E+06 | 7 | 7 | 17 | | 36062 | NADH-ubiquinone oxidoreductase chain 1 OS=Rattus norvegicus OX=10116 GN=ND1 PE=3 SV=1 |
| 63 | 44 | P17764\|TH | 88.67 | 8 | 8 | 1.49E+06 | 4 | 4 | 8 | | 44695 | Acetyl-CoA acetyltransferase mitochondrial OS=Rattus norvegicus OX=10116 GN=Acat1 PE=1 SV=1 |
| 70 | 207 | P11951\|CX | 87.07 | 38 | 38 | 8.51E+05 | 5 | 5 | 7 | Oxidation ( | 8455 | Cytochrome c oxidase subunit 6C-2 OS=Rattus norvegicus OX=10116 GN=Cox6c2 PE=1 SV=3 |
| 65 | 263 | P35171\|CX | 86.66 | 12 | 12 | 8.34E+05 | 3 | 3 | 8 | | 9353 | Cytochrome c oxidase subunit 7A2 mitochondrial OS=Rattus norvegicus OX=10116 GN=Cox7a2 PE=1 SV=1 |
| 65 | 264 | tr\|B2RYS0 | 86.66 | 12 | 12 | 8.34E+05 | 3 | 3 | 8 | | 9353 | Cox7a2 protein OS=Rattus norvegicus OX=10116 GN=Cox7a2 PE=2 SV=1 |
| 75 | 4 | P08461\|OI | 84.98 | 5 | 5 | 7.00E+05 | 4 | 4 | 5 | | 67166 | Dihydrolipoyllysine-residue acetyltransferase component of pyruvate dehydrogenase complex mitochondrial OS=Rattus norvegicus OX=10116 GN=Dlat PE=1 SV=3 |
| 74 | 63 | P35435\|AT | 83.6 | 11 | 11 | 4.53E+05 | 4 | 4 | 6 | | 30191 | ATP synthase subunit gamma mitochondrial OS=Rattus norvegicus OX=10116 GN=Atp5f1c PE=1 SV=2 |
| 74 | 64 | tr\|Q6QI09 | 83.6 | 5 | 5 | 4.53E+05 | 4 | 4 | 6 | | 67721 | ATP synthase subunit gamma mitochondrial OS=Rattus norvegicus OX=10116 GN=Taf3 PE=1 SV=1 |
| 62 | 21 | Q64428\|EC | 82.88 | 6 | 6 | 9.06E+05 | 6 | 6 | 8 | | 82665 | Trifunctional enzyme subunit alpha mitochondrial OS=Rattus norvegicus OX=10116 GN=Hadha PE=1 SV=2 |

| | | | | | | | | | | | | |
|---|---|---|---|---|---|---|---|---|---|---|---|---|
| 58 | 384 | tr\|Q06Q97 | 81.46 | 8 | 8 | 9.70E+05 | 4 | 4 | 10 | | 38485 | NADH-ubiquinone oxidoreductase chain 2 OS=Rattus norvegicus OX=10116 GN=ND2 PE=3 SV=1 |
| 58 | 386 | tr\|Q5UAJ8 | 81.46 | 8 | 8 | 9.70E+05 | 4 | 4 | 10 | | 38455 | NADH-ubiquinone oxidoreductase chain 2 OS=Rattus norvegicus OX=10116 GN=ND2 PE=3 SV=1 |
| 58 | 388 | tr\|Q8HID0 | 81.46 | 8 | 8 | 9.70E+05 | 4 | 4 | 10 | | 38653 | NADH-ubiquinone oxidoreductase chain 2 OS=Rattus norvegicus OX=10116 GN=Mt-nd2 PE=3 SV=1 |
| 58 | 389 | P11662\|NU | 81.46 | 8 | 8 | 9.70E+05 | 4 | 4 | 10 | | 38653 | NADH-ubiquinone oxidoreductase chain 2 OS=Rattus norvegicus OX=10116 GN=Mtnd2 PE=3 SV=3 |
| 58 | 390 | tr\|Q06QH5 | 81.46 | 8 | 8 | 9.70E+05 | 4 | 4 | 10 | | 38626 | NADH-ubiquinone oxidoreductase chain 2 OS=Rattus norvegicus OX=10116 GN=ND2 PE=3 SV=1 |
| 58 | 391 | tr\|Q8SEZ7 | 81.46 | 8 | 8 | 9.70E+05 | 4 | 4 | 10 | | 38580 | NADH-ubiquinone oxidoreductase chain 2 (Fragment) OS=Rattus norvegicus OX=10116 GN=NADH2 PE=3 SV=1 |
| 58 | 392 | tr\|Q06QB0 | 81.46 | 8 | 8 | 9.70E+05 | 4 | 4 | 10 | | 38534 | NADH-ubiquinone oxidoreductase chain 2 OS=Rattus norvegicus OX=10116 GN=ND2 PE=3 SV=1 |
| 58 | 393 | tr\|D2E6P4 | 81.46 | 8 | 8 | 9.70E+05 | 4 | 4 | 10 | | 38623 | NADH-ubiquinone oxidoreductase chain 2 OS=Rattus norvegicus OX=10116 GN=ND2 PE=3 SV=1 |
| 58 | 394 | tr\|Q06QE9 | 81.46 | 8 | 8 | 9.70E+05 | 4 | 4 | 10 | | 38598 | NADH-ubiquinone oxidoreductase chain 2 OS=Rattus norvegicus OX=10116 GN=ND2 PE=3 SV=1 |
| 58 | 385 | tr\|A0A0A1 | 81.46 | 8 | 8 | 9.70E+05 | 4 | 4 | 10 | | 38542 | NADH-ubiquinone oxidoreductase chain 2 OS=Rattus norvegicus OX=10116 GN=ND2 PE=3 SV=1 |
| 59 | 363 | P05508\|NU | 80.63 | 12 | 12 | 1.18E+06 | 8 | 8 | 10 | Oxidation (M) | | |
| 59 | 364 | tr\|D2E6K0 | 80.63 | 12 | 12 | 1.18E+06 | 8 | 8 | 10 | Oxidation (M) | | |
| 59 | 365 | tr\|A7XYB9 | 80.63 | 12 | 12 | 1.18E+06 | 8 | 8 | 10 | Oxidation (M) | | |
| 59 | 366 | tr\|Q06QE1 | 80.63 | 12 | 12 | 1.18E+06 | 8 | 8 | 10 | Oxidation (M) | | |
| 59 | 367 | tr\|Q06QA2 | 80.63 | 12 | 12 | 1.18E+06 | 8 | 8 | 10 | Oxidation (M) | | |
| 59 | 368 | tr\|Q7HKW | 80.63 | 12 | 12 | 1.18E+06 | 8 | 8 | 10 | Oxidation (M) | | |
| 59 | 369 | tr\|Q06QG7 | 80.63 | 12 | 12 | 1.18E+06 | 8 | 8 | 10 | Oxidation (M) | | |
| 59 | 370 | tr\|Q06Q89 | 80.63 | 12 | 12 | 1.18E+06 | 8 | 8 | 10 | Oxidation (M) | | |
| 59 | 371 | tr\|Q35737 | 80.63 | 12 | 12 | 1.18E+06 | 8 | 8 | 10 | Oxidation (M) | | |
| 59 | 372 | tr\|Q8HIC6 | 80.63 | 12 | 12 | 1.18E+06 | 8 | 8 | 10 | Oxidation (M) | | |
| 51 | 376 | tr\|Q5RJN0 | 76.98 | 17 | 17 | 3.34E+06 | 4 | 4 | 13 | Oxidation ( | 23945 | NADH dehydrogenase (Ubiquinone) Fe-S protein 7 OS=Rattus norvegicus OX=10116 GN=Ndufs7 PE=1 SV=1 |
| 41 | 208 | Q7TQ16\|Q | 73.67 | 30 | 30 | 4.27E+06 | 6 | 6 | 19 | | 9849 | Cytochrome b-c1 complex subunit 8 OS=Rattus norvegicus OX=10116 GN=Uqcrq PE=3 SV=1 |
| 76 | 267 | tr\|I6V4L9\| | 73.39 | 7 | 7 | 1.67E+05 | 3 | 3 | 5 | | 29844 | Cytochrome c oxidase subunit 3 OS=Rattus norvegicus OX=10116 GN=COX3 PE=3 SV=1 |
| 76 | 266 | tr\|Q7H115 | 73.39 | 7 | 7 | 1.67E+05 | 3 | 3 | 5 | | 29871 | Cytochrome c oxidase subunit 3 OS=Rattus norvegicus OX=10116 GN=Mt-co3 PE=3 SV=1 |
| 76 | 268 | tr\|Q8M7G | 73.39 | 7 | 7 | 1.67E+05 | 3 | 3 | 5 | | 29861 | Cytochrome c oxidase subunit 3 OS=Rattus norvegicus OX=10116 PE=2 SV=1 |
| 76 | 269 | tr\|A0A096 | 73.39 | 7 | 7 | 1.67E+05 | 3 | 3 | 5 | | 29901 | Cytochrome c oxidase subunit 3 OS=Rattus norvegicus OX=10116 GN=COX3 PE=3 SV=1 |
| 76 | 270 | P05505\|CO | 73.39 | 7 | 7 | 1.67E+05 | 3 | 3 | 5 | | 29871 | Cytochrome c oxidase subunit 3 OS=Rattus norvegicus OX=10116 GN=Mtco3 PE=1 SV=5 |
| 76 | 271 | tr\|Q8SEZ2 | 73.39 | 7 | 7 | 1.67E+05 | 3 | 3 | 5 | | 29870 | Cytochrome c oxidase subunit 3 (Fragment) OS=Rattus norvegicus OX=10116 GN=COIII PE=3 SV=1 |
| 47 | 3918 | tr\|D3ZLT1\| | 70.05 | 16 | 16 | 4.23E+06 | 3 | 3 | 15 | Oxidation ( | 16568 | NADH dehydrogenase (Ubiquinone) 1 beta subcomplex 7 (Predicted) OS=Rattus norvegicus OX=10116 GN=Ndufb7 PE=1 SV=1 |
| 60 | 151 | P12075\|CO | 69.1 | 11 | 11 | 7.55E+06 | 6 | 6 | 10 | | 13915 | Cytochrome c oxidase subunit 5B mitochondrial OS=Rattus norvegicus OX=10116 GN=Cox5b PE=1 SV=2 |
| 67 | 1021 | tr\|Q06QA9 | 67.34 | 5 | 5 | 6.21E+05 | 4 | 3 | 7 | | 56893 | Cytochrome c oxidase subunit 1 OS=Rattus norvegicus OX=10116 GN=CO1 PE=3 SV=1 |
| 67 | 1022 | tr\|Q06QK0 | 67.34 | 5 | 5 | 6.21E+05 | 4 | 3 | 7 | | 56880 | Cytochrome c oxidase subunit 1 OS=Rattus norvegicus OX=10116 GN=CO1 PE=3 SV=1 |
| 67 | 1023 | tr\|Q8SEZ6 | 67.34 | 5 | 5 | 6.21E+05 | 4 | 3 | 7 | | 56879 | Cytochrome c oxidase subunit 1 OS=Rattus norvegicus OX=10116 GN=COI PE=3 SV=1 |
| 67 | 1024 | tr\|A0A0A1 | 67.34 | 5 | 5 | 6.21E+05 | 4 | 3 | 7 | | 56937 | Cytochrome c oxidase subunit 1 OS=Rattus norvegicus OX=10116 GN=COX1 PE=3 SV=1 |
| 67 | 1025 | tr\|Q8HIC9 | 67.34 | 5 | 5 | 6.21E+05 | 4 | 3 | 7 | | 56845 | Cytochrome c oxidase subunit 1 OS=Rattus norvegicus OX=10116 GN=Mt-co1 PE=3 SV=1 |
| 67 | 1026 | P05503\|CO | 67.34 | 5 | 5 | 6.21E+05 | 4 | 3 | 7 | | 56845 | Cytochrome c oxidase subunit 1 OS=Rattus norvegicus OX=10116 GN=Mtco1 PE=2 SV=3 |
| 67 | 1027 | tr\|Q95938 | 67.34 | 5 | 5 | 6.21E+05 | 4 | 3 | 7 | | 56977 | Cytochrome c oxidase subunit 1 OS=Rattus norvegicus OX=10116 GN=Co I PE=3 SV=1 |
| 69 | 292 | P10888\|CO | 64.61 | 18 | 18 | 8.05E+05 | 4 | 4 | 7 | Oxidation ( | 19515 | Cytochrome c oxidase subunit 4 isoform 1 mitochondrial OS=Rattus norvegicus OX=10116 GN=Cox4i1 PE=1 SV=1 |
| 81 | 233 | tr\|Q5UAJ5 | 62.86 | 16 | 16 | 7.14E+05 | 2 | 2 | 4 | | 7642 | ATP synthase protein 8 OS=Rattus norvegicus OX=10116 GN=ATP8 PE=3 SV=1 |
| 81 | 234 | P11608\|AT | 62.86 | 16 | 16 | 7.14E+05 | 2 | 2 | 4 | | 7630 | ATP synthase protein 8 OS=Rattus norvegicus OX=10116 GN=Mt-atp8 PE=1 SV=2 |
| 81 | 235 | tr\|Q8SEZ4 | 62.86 | 16 | 16 | 7.14E+05 | 2 | 2 | 4 | | 7632 | ATP synthase protein 8 OS=Rattus norvegicus OX=10116 GN=ATPase8 PE=3 SV=1 |
| 81 | 236 | tr\|Q8HIC8 | 62.86 | 16 | 16 | 7.14E+05 | 2 | 2 | 4 | | 7630 | ATP synthase protein 8 OS=Rattus norvegicus OX=10116 GN=Mt-atp8 PE=3 SV=1 |
| 80 | 3911 | tr\|B2RYW3 | 61.6 | 11 | 11 | 6.57E+05 | 2 | 2 | 4 | Oxidation ( | 21892 | NADH dehydrogenase (Ubiquinone) 1 beta subcomplex 9 OS=Rattus norvegicus OX=10116 GN=Ndufb9 PE=1 SV=1 |
| 64 | 398 | tr\|B2RYT5 | 61.2 | 30 | 30 | 1.45E+06 | 4 | 4 | 8 | Oxidation ( | 12651 | Cox7a2l protein OS=Rattus norvegicus OX=10116 GN=Cox7a2l PE=2 SV=1 |
| 64 | 399 | tr\|D3ZYX8 | 61.2 | 28 | 28 | 1.45E+06 | 4 | 4 | 8 | Oxidation ( | 13274 | Cytochrome c oxidase subunit 7A2-like OS=Rattus norvegicus OX=10116 GN=Cox7a2l PE=1 SV=1 |
| 78 | 3914 | tr\|B2RYU0 | 57.92 | 21 | 21 | 1.42E+06 | 3 | 3 | 4 | | 11842 | NADH dehydrogenase (Ubiquinone) 1 beta subcomplex 2 (Predicted) isoform CRA_b OS=Rattus norvegicus OX=10116 GN=Ndufb2 PE=1 SV=1 |
| 73 | 467 | tr\|D4A4P3 | 54.71 | 22 | 22 | 1.83E+06 | 3 | 3 | 6 | Oxidation ( | 11267 | NADH:ubiquinone oxidoreductase subunit B3 OS=Rattus norvegicus OX=10116 GN=Ndufb3 PE=1 SV=1 |
| 72 | 209 | tr\|B2RZD6 | 54.33 | 20 | 20 | 8.71E+05 | 3 | 3 | 7 | | 9327 | NDUFA4 mitochondrial complex-associated OS=Rattus norvegicus OX=10116 GN=Ndufa4 PE=1 SV=1 |
| 88 | 3909 | tr\|G3V644 | 52 | 4 | 4 | 3.93E+05 | 2 | 2 | 3 | | 49300 | NADH dehydrogenase (Ubiquinone) flavoprotein 3-like isoform CRA_a OS=Rattus norvegicus OX=10116 GN=Ndufv3 PE=1 SV=1 |
| 88 | 3910 | tr\|O54755 | 52 | 4 | 4 | 3.93E+05 | 2 | 2 | 3 | | 49312 | MIPP65 OS=Rattus norvegicus OX=10116 GN=Ndufv3 PE=2 SV=1 |
| 129 | 30 | tr\|Q5BJZ3\| | 49.53 | 1 | 1 | 9.86E+04 | 1 | 1 | 1 | | 113869 | Nicotinamide nucleotide transhydrogenase OS=Rattus norvegicus OX=10116 GN=Nnt PE=1 SV=1 |
| 87 | 159 | tr\|Q5U1W | 48.74 | 9 | 9 | 3.39E+05 | 3 | 3 | 3 | | 28238 | MICOS complex subunit OS=Rattus norvegicus OX=10116 GN=Apool PE=1 SV=1 |
| 89 | 18 | P26284\|OI | 48.39 | 5 | 5 | 2.97E+05 | 3 | 3 | 3 | | 43227 | Pyruvate dehydrogenase E1 component subunit alpha somatic form mitochondrial OS=Rattus norvegicus OX=10116 GN=Pdha1 PE=1 SV=2 |
| 147 | 76 | Q06647\|AT | 45.73 | 4 | 4 | 3.64E+05 | 1 | 1 | 1 | | 23398 | ATP synthase subunit O mitochondrial OS=Rattus norvegicus OX=10116 GN=Atp5po PE=1 SV=1 |
| 107 | 3936 | tr\|A0A0G2 | 42.62 | 13 | 13 | 1.54E+05 | 2 | 2 | 2 | | 7099 | Ubiquinol-cytochrome c reductase complex III subunit X OS=Rattus norvegicus OX=10116 GN=Uqcr10 PE=1 SV=1 |
| 107 | 3937 | tr\|B2RYX1 | 42.62 | 12 | 12 | 1.54E+05 | 2 | 2 | 2 | | 7462 | LOC685322 protein OS=Rattus norvegicus OX=10116 GN=Uqcr10 PE=2 SV=1 |
| 109 | 3915 | tr\|Q8SEZ1 | 42.05 | 20 | 20 | 1.04E+05 | 2 | 2 | 2 | | 13087 | NADH-ubiquinone oxidoreductase chain 3 OS=Rattus norvegicus OX=10116 GN=Mt-nd3 PE=2 SV=1 |
| 109 | 3916 | P05506\|NU | 42.05 | 20 | 20 | 1.04E+05 | 2 | 2 | 2 | | 13087 | NADH-ubiquinone oxidoreductase chain 3 OS=Rattus norvegicus OX=10116 GN=Mtnd3 PE=3 SV=3 |
| 137 | 31 | P45953\|AC | 40.59 | 2 | 2 | 1.37E+05 | 1 | 1 | 1 | | 70749 | Very long-chain specific acyl-CoA dehydrogenase mitochondrial OS=Rattus norvegicus OX=10116 GN=Acadvl PE=1 SV=1 |
| 137 | 32 | tr\|Q5M9H | 40.59 | 2 | 2 | 1.37E+05 | 1 | 1 | 1 | | 70821 | Acyl-Coenzyme A dehydrogenase very long chain OS=Rattus norvegicus OX=10116 GN=Acadvl PE=1 SV=1 |
| 79 | 43 | tr\|D3ZB81 | 39.45 | 6 | 6 | 2.99E+05 | 3 | 3 | 4 | | 35227 | Solute carrier family 25 member 31 OS=Rattus norvegicus OX=10116 GN=Slc25a31 PE=3 SV=3 |
| 79 | 23 | tr\|Q6P9Y4 | 39.45 | 6 | 6 | 2.99E+05 | 3 | 3 | 4 | | 32904 | ADP/ATP translocase 1 OS=Rattus norvegicus OX=10116 GN=Slc25a4 PE=1 SV=1 |
| 79 | 24 | Q05962\|AI | 39.45 | 6 | 6 | 2.99E+05 | 3 | 3 | 4 | | 32989 | ADP/ATP translocase 1 OS=Rattus norvegicus OX=10116 GN=Slc25a4 PE=1 SV=3 |

| | | | | | | | | | | | | |
|---|---|---|---|---|---|---|---|---|---|---|---|---|
| 79 | 27 | Q09073\|AI | 39.45 | 6 | 6 | 2.99E+05 | 3 | 3 | 4 | | 32901 | ADP/ATP translocase 2 OS=Rattus norvegicus OX=10116 GN=Slc25a5 PE=1 SV=3 |
| 84 | 73 | Q63362\|N | 39.17 | 14 | 14 | 5.50E+05 | 2 | 1 | 4 | | 13412 | NADH dehydrogenase [ubiquinone] 1 alpha subcomplex subunit 5 OS=Rattus norvegicus OX=10116 GN=Ndufa5 PE=1 SV=3 |
| 148 | 48 | Q4V8F9\|H | 37.54 | 2 | 2 | 3.93E+05 | 1 | 1 | 1 | | 58344 | Hydroxysteroid dehydrogenase-like protein 2 OS=Rattus norvegicus OX=10116 GN=Hsdl2 PE=2 SV=1 |
| 111 | 3932 | tr\|D3ZD09 | 36.05 | 12 | 12 | 1.57E+05 | 1 | 1 | 2 | | 10071 | Cytochrome c oxidase subunit OS=Rattus norvegicus OX=10116 GN=Cox6b1 PE=1 SV=1 |
| 110 | 321 | tr\|D4AE90 | 35.5 | 3 | 3 | 0 | 2 | 1 | 2 | Formylatio | 50113 | RCC1-like OS=Rattus norvegicus OX=10116 GN=Rcc1l PE=1 SV=1 |
| 95 | 410 | tr\|F1LNF0\| | 34.93 | 1 | 1 | 5.57E+05 | 2 | 2 | 2 | | 228912 | Myosin heavy chain 14 OS=Rattus norvegicus OX=10116 GN=Myh14 PE=1 SV=1 |
| 169 | 3934 | Q66H47\|R | 34.9 | 6 | 6 | 1.25E+05 | 1 | 1 | 1 | | 25001 | 39S ribosomal protein L24 mitochondrial OS=Rattus norvegicus OX=10116 GN=Mrpl24 PE=2 SV=1 |
| 112 | 143 | tr\|A0A0G2 | 34.42 | 2 | 2 | 5.13E+04 | 1 | 1 | 2 | | 47273 | Creatine kinase S-type mitochondrial OS=Rattus norvegicus OX=10116 GN=Ckmt2 PE=1 SV=1 |
| 112 | 46 | P09605\|KC | 34.42 | 2 | 2 | 5.13E+04 | 1 | 1 | 2 | | 47385 | Creatine kinase S-type mitochondrial OS=Rattus norvegicus OX=10116 GN=Ckmt2 PE=1 SV=2 |
| 101 | 75 | Q6PDU7\|A | 33.7 | 16 | 16 | 2.92E+05 | 1 | 1 | 2 | | 11433 | ATP synthase subunit g mitochondrial OS=Rattus norvegicus OX=10116 GN=Atp5mg PE=1 SV=2 |
| 105 | 13 | P49432\|OI | 30.79 | 5 | 5 | 1.14E+05 | 2 | 2 | 2 | | 38982 | Pyruvate dehydrogenase E1 component subunit beta mitochondrial OS=Rattus norvegicus OX=10116 GN=Pdhb PE=1 SV=2 |
| 105 | 15 | tr\|A0A0G2 | 30.79 | 4 | 4 | 1.14E+05 | 2 | 2 | 2 | | 46193 | Pyruvate dehydrogenase E1 component subunit beta OS=Rattus norvegicus OX=10116 GN=Pdhb PE=1 SV=1 |
| 138 | 3923 | P80431\|CC | 30.26 | 9 | 9 | 1.57E+05 | 1 | 1 | 1 | | 8995 | Cytochrome c oxidase subunit 7B mitochondrial OS=Rattus norvegicus OX=10116 GN=Cox7b PE=1 SV=3 |
| 106 | 3919 | tr\|A9UMV | 29.03 | 11 | 11 | 2.80E+05 | 1 | 1 | 2 | | 6539 | RCG29512 OS=Rattus norvegicus OX=10116 GN=Uqcr11 PE=1 SV=1 |
| 118 | 3958 | tr\|D3ZD73 | 28.78 | 2 | 2 | | 2 | 0 | 2 | | 54245 | DEAD-box helicase 6 OS=Rattus norvegicus OX=10116 GN=Ddx6 PE=1 SV=1 |
| 113 | 813 | Q68FU4\|S | 28.27 | 2 | 2 | | 2 | 0 | 2 | | 47549 | Succinate--hydroxymethylglutarate CoA-transferase OS=Rattus norvegicus OX=10116 GN=Sugct PE=2 SV=1 |
| 83 | 482 | tr\|D3ZRJ7\| | 28.04 | 1 | 1 | 1.07E+05 | 2 | 1 | 4 | | 225446 | Leucine-rich repeat kinase 1 OS=Rattus norvegicus OX=10116 GN=Lrrk1 PE=4 SV=2 |
| 170 | 79 | tr\|D3ZUX5 | 27.82 | 4 | 4 | 9.87E+04 | 1 | 1 | 1 | | 26435 | MICOS complex subunit OS=Rattus norvegicus OX=10116 GN=Chchd3 PE=1 SV=1 |
| 90 | 3917 | tr\|D3ZJG4 | 27.81 | 1 | 1 | 2.67E+05 | 1 | 1 | 3 | Oxidation ( | 95972 | Phosphofurin acidic cluster sorting protein 2 OS=Rattus norvegicus OX=10116 GN=Pacs2 PE=1 SV=2 |
| 92 | 256 | tr\|Q8M7G | 27.65 | 9 | 9 | 1.55E+06 | 2 | 2 | 3 | Oxidation ( | 24240 | ATP synthase subunit a (Fragment) OS=Rattus norvegicus OX=10116 PE=2 SV=1 |
| 92 | 258 | P05504\|AT | 27.65 | 9 | 9 | 1.55E+06 | 2 | 2 | 3 | Oxidation ( | 25076 | ATP synthase subunit a OS=Rattus norvegicus OX=10116 GN=Mt-atp6 PE=1 SV=3 |
| 92 | 259 | tr\|Q8HIC7 | 27.65 | 9 | 9 | 1.55E+06 | 2 | 2 | 3 | Oxidation ( | 25076 | ATP synthase subunit a OS=Rattus norvegicus OX=10116 GN=Mt-atp6 PE=4 SV=1 |
| 92 | 260 | tr\|S5S1E9\| | 27.65 | 9 | 9 | 1.55E+06 | 2 | 2 | 3 | Oxidation ( | 25049 | ATP synthase subunit a OS=Rattus norvegicus OX=10116 GN=ATP6 PE=4 SV=1 |
| 92 | 261 | tr\|Q06QE5 | 27.65 | 9 | 9 | 1.55E+06 | 2 | 2 | 3 | Oxidation ( | 25075 | ATP synthase subunit a OS=Rattus norvegicus OX=10116 GN=ATP6 PE=4 SV=1 |
| 92 | 262 | tr\|Q8SEZ3 | 27.65 | 9 | 9 | 1.55E+06 | 2 | 2 | 3 | Oxidation ( | 25030 | ATP synthase subunit a OS=Rattus norvegicus OX=10116 GN=ATPase6 PE=4 SV=1 |
| 86 | 224 | P11530\|DI | 27.34 | 0 | 0 | 4.24E+05 | 2 | 2 | 3 | | 425830 | Dystrophin OS=Rattus norvegicus OX=10116 GN=Dmd PE=1 SV=2 |
| 82 | 1034 | P05696\|KF | 25.58 | 1 | 1 | 8.13E+05 | 1 | 1 | 4 | | 76792 | Protein kinase C alpha type OS=Rattus norvegicus OX=10116 GN=Prkca PE=1 SV=3 |
| 98 | 497 | tr\|A0A0G2 | 25.13 | 0 | 0 | 0 | 2 | 1 | 2 | | 487857 | Vacuolar protein sorting 13 homolog D OS=Rattus norvegicus OX=10116 GN=Vps13d PE=1 SV=1 |
| 98 | 498 | tr\|D3ZKC6 | 25.13 | 0 | 0 | 0 | 2 | 1 | 2 | | 488962 | Vacuolar protein sorting 13 homolog D OS=Rattus norvegicus OX=10116 GN=Vps13d PE=1 SV=1 |
| 99 | 298 | tr\|Q5RKL4 | 23.9 | 1 | 1 | 0 | 1 | 1 | 2 | | 95977 | Dimethylglycine dehydrogenase OS=Rattus norvegicus OX=10116 GN=Dmgdh PE=1 SV=1 |
| 99 | 299 | Q63342\|M | 23.9 | 1 | 1 | 0 | 1 | 1 | 2 | | 96047 | Dimethylglycine dehydrogenase mitochondrial OS=Rattus norvegicus OX=10116 GN=Dmgdh PE=1 SV=1 |
| 99 | 303 | tr\|A0A0G2 | 23.9 | 1 | 1 | 0 | 1 | 1 | 2 | | 98864 | Dimethylglycine dehydrogenase mitochondrial OS=Rattus norvegicus OX=10116 GN=Dmgdh PE=1 SV=1 |
| 115 | 3943 | tr\|D4A6D9 | 23.7 | 3 | 3 | 3.24E+05 | 1 | 1 | 2 | | 43431 | HCLS1-binding protein 3 OS=Rattus norvegicus OX=10116 GN=Hs1bp3 PE=1 SV=1 |
| 173 | 332 | tr\|F1LNJ1\| | 21.14 | 0 | 0 | 1.30E+06 | 1 | 1 | 1 | | 284829 | Leucine-rich repeat kinase 2 OS=Rattus norvegicus OX=10116 GN=Lrrk2 PE=1 SV=3 |
| 116 | 524 | tr\|D4A054 | 21.14 | 0 | 0 | 5.15E+05 | 1 | 1 | 2 | | 341403 | RAN-binding protein 2 OS=Rattus norvegicus OX=10116 GN=Ranbp2 PE=1 SV=2 |
| 116 | 525 | tr\|M0R3M | 21.14 | 0 | 0 | 5.15E+05 | 1 | 1 | 2 | | 344396 | RAN-binding protein 2 OS=Rattus norvegicus OX=10116 GN=Ranbp2 PE=1 SV=1 |
| 174 | 3951 | tr\|B1WBP | 20.71 | 7 | 7 | 3.83E+04 | 1 | 1 | 1 | | 12884 | ATP synthase subunit delta mitochondrial OS=Rattus norvegicus OX=10116 GN=Atp5f1d PE=1 SV=1 |
| 174 | 3952 | P35434\|AT | 20.71 | 5 | 5 | 3.83E+04 | 1 | 1 | 1 | | 17595 | ATP synthase subunit delta mitochondrial OS=Rattus norvegicus OX=10116 GN=Atp5f1d PE=1 SV=2 |
| 174 | 3953 | tr\|G3V7Y3 | 20.71 | 5 | 5 | 3.83E+04 | 1 | 1 | 1 | | 17563 | ATP synthase subunit delta mitochondrial OS=Rattus norvegicus OX=10116 GN=Atp5f1d PE=1 SV=1 |
| 108 | 204 | Q499N5\|A | 20.6 | 1 | 1 | 1.44E+05 | 1 | 1 | 2 | Formylatio | 67887 | Medium-chain acyl-CoA ligase ACSF2 mitochondrial OS=Rattus norvegicus OX=10116 GN=Acsf2 PE=2 SV=1 |
| 93 | 3954 | P50554\|GA | 20.6 | 3 | 3 | 1.76E+06 | 1 | 1 | 3 | | 56456 | 4-aminobutyrate aminotransferase mitochondrial OS=Rattus norvegicus OX=10116 GN=Abat PE=1 SV=3 |
