## Supplementary material for "Permeability transition pore-related changes in the proteome and channel activity of ATP synthase dimers and monomers": RLM SM-BM

| Protein Gr | Protein ID | Accession | -10lgP | Coverage ( | Coverage ( | IBAQ | #Peptides | #Unique | #Spec Sam | PTM | Avg. Mass | Description |
|---|---|---|---|---|---|---|---|---|---|---|---|---|
| 1 | 1 | Q02253\|MMSA_RAT | 421.37 | 77 | 77 | 178060000.00 | 160 | 159 | 269 | Carbamidomethylatio | | Methylmalonate-semialdehyde dehydrogenase [acylating], mitochondrial |
| 1 | 2 | tr\|G3V7J0\|G3V7J0_RAT | 421.37 | 77 | 77 | 178060000.00 | 160 | 159 | 269 | Carbamidomethylatio | | Methylmalonate-semialdehyde dehydrogenase [acylating], mitochondrial |
| 2 | 3 | tr\|B2GV15\|B2GV15_RAT | 385.64 | 64 | 64 | 206960000.00 | 138 | 138 | 251 | Oxidation ( | 53274 | Dihydrolipoamide acetyltransferase component of pyruvate dehydrogenase complex OS=Rattus norvegicus OX=10116 GN=Dbt PE=1 SV=1 |
| 3 | 5 | tr\|G3V6P2\|G3V6P2_RAT | 338.24 | 62 | 62 | 106240000.00 | 87 | 87 | 152 | Oxidation ( | 48899 | Dihydrolipoamide S-succinyltransferase (E2 component of 2-oxo-glutarate complex) isoform CRA_a OS=Rattus norvegicus OX=10116 GN=Dlst PE=1 SV=1 |
| 4 | 10 | Q66HF1\|NDUS1_RAT | 323.83 | 51 | 51 | 55466000.00 | 88 | 88 | 123 | Oxidation ( | 79412 | NADH-ubiquinone oxidoreductase 75 kDa subunit mitochondrial OS=Rattus norvegicus OX=10116 GN=Ndufs1 PE=1 SV=1 |
| 11 | 9 | Q5BK63\|NDUA9_RAT | 302.61 | 49 | 49 | 24972000.00 | 47 | 47 | 73 | Oxidation ( | 42559 | NADH dehydrogenase [ubiquinone] 1 alpha subcomplex subunit 9 mitochondrial OS=Rattus norvegicus OX=10116 GN=Ndufa9 PE=1 SV=2 |
| 9 | 7 | Q4FZT0\|STML2_RAT | 301.36 | 63 | 63 | 36116000.00 | 58 | 58 | 81 | Oxidation ( | 38414 | Stomatin-like protein 2 mitochondrial OS=Rattus norvegicus OX=10116 GN=Stoml2 PE=1 SV=1 |
| 5 | 13 | P32551\|QCR2_RAT | 301.28 | 60 | 60 | 62909000.00 | 76 | 76 | 121 | Oxidation ( | 48396 | Cytochrome b-c1 complex subunit 2 mitochondrial OS=Rattus norvegicus OX=10116 GN=Uqcrc2 PE=1 SV=2 |
| 10 | 8 | P07756\|CPSM_RAT | 295.66 | 32 | 32 | 24577000.00 | 64 | 64 | 79 | Oxidation ( | 164579 | Carbamoyl-phosphate synthase [ammonia] mitochondrial OS=Rattus norvegicus OX=10116 GN=Cps1 PE=1 SV=1 |
| 8 | 30 | P19234\|NDUV2_RAT | 293.35 | 47 | 47 | 45081000.00 | 46 | 45 | 81 | Formylatio | 27378 | NADH dehydrogenase [ubiquinone] flavoprotein 2 mitochondrial OS=Rattus norvegicus OX=10116 GN=Ndufv2 PE=1 SV=2 |
| 7 | 11 | P67779\|PHB_RAT | 290.16 | 81 | 81 | 54220000.00 | 65 | 65 | 96 | | 29820 | Prohibitin OS=Rattus norvegicus OX=10116 GN=Phb PE=1 SV=1 |
| 17 | 22 | tr\|D3ZFQ8\|D3ZFQ8_RAT | 283.17 | 43 | 43 | 36962000.00 | 36 | 36 | 52 | Oxidation ( | 35435 | Cytochrome c-1 OS=Rattus norvegicus OX=10116 GN=Cyc1 PE=1 SV=3 |
| 6 | 12 | Q68FY0\|QCR1_RAT | 281.35 | 50 | 50 | 66322000.00 | 76 | 76 | 118 | Oxidation ( | 52849 | Cytochrome b-c1 complex subunit 1 mitochondrial OS=Rattus norvegicus OX=10116 GN=Uqcrc1 PE=1 SV=1 |
| 18 | 29 | P07895\|SODM_RAT | 270.29 | 61 | 61 | 33275000.00 | 30 | 30 | 48 | | 24674 | Superoxide dismutase [Mn] mitochondrial OS=Rattus norvegicus OX=10116 GN=Sod2 PE=1 SV=2 |
| 32 | 45 | tr\|B2RYS8\|B2RYS8_RAT | 259.39 | 46 | 46 | 10525000.00 | 21 | 21 | 25 | Oxidation ( | 21959 | NADH dehydrogenase [ubiquinone] 1 beta subcomplex subunit 8 mitochondrial OS=Rattus norvegicus OX=10116 GN=Ndufb8 PE=1 SV=1 |
| 20 | 17 | P10719\|ATPB_RAT | 258.97 | 42 | 42 | 20045000.00 | 35 | 35 | 46 | Oxidation ( | 56354 | ATP synthase subunit beta mitochondrial OS=Rattus norvegicus OX=10116 GN=Atp5f1b PE=1 SV=2 |
| 20 | 18 | tr\|G3V6D3\|G3V6D3_RAT | 258.97 | 42 | 42 | 20045000.00 | 35 | 35 | 46 | Oxidation ( | 56345 | ATP synthase subunit beta OS=Rattus norvegicus OX=10116 GN=Atp5f1b PE=1 SV=1 |
| 14 | 15 | P00507\|AATM_RAT | 258.07 | 54 | 54 | 27286000.00 | 44 | 44 | 58 | Oxidation ( | 47314 | Aspartate aminotransferase mitochondrial OS=Rattus norvegicus OX=10116 GN=Got2 PE=1 SV=2 |
| 21 | 16 | tr\|F1LN92\|F1LN92_RAT | 251.05 | 30 | 30 | 12495000.00 | 38 | 37 | 43 | Oxidation ( | 89352 | AFG3-like matrix AAA peptidase subunit 2 OS=Rattus norvegicus OX=10116 GN=Afg3l2 PE=1 SV=1 |
| 24 | 14 | P10860\|DHE3_RAT | 248.69 | 33 | 33 | 11732000.00 | 31 | 31 | 37 | Oxidation ( | 61416 | Glutamate dehydrogenase 1 mitochondrial OS=Rattus norvegicus OX=10116 GN=Glud1 PE=1 SV=2 |
| 19 | 20 | P15999\|ATPA_RAT | 242.36 | 38 | 38 | 14170000.00 | 37 | 37 | 47 | Oxidation ( | 59754 | ATP synthase subunit alpha mitochondrial OS=Rattus norvegicus OX=10116 GN=Atp5f1a PE=1 SV=2 |
| 19 | 21 | tr\|F1LP05\|F1LP05_RAT | 242.36 | 38 | 38 | 14170000.00 | 37 | 37 | 47 | Oxidation ( | 59813 | ATP synthase subunit alpha OS=Rattus norvegicus OX=10116 GN=Atp5f1a PE=1 SV=1 |
| 26 | 42 | tr\|D4A0T0\|D4A0T0_RAT | 238.01 | 54 | 54 | 12843000.00 | 18 | 18 | 32 | Oxidation ( | 20859 | NADH:ubiquinone oxidoreductase subunit B10 OS=Rattus norvegicus OX=10116 GN=Ndufb10 PE=1 SV=1 |
| 13 | 19 | Q5XIH7\|PHB2_RAT | 235.74 | 57 | 57 | 27551000.00 | 45 | 45 | 66 | Oxidation ( | 33312 | Prohibitin-2 OS=Rattus norvegicus OX=10116 GN=Phb2 PE=1 SV=1 |
| 27 | 164 | tr\|D3ZF13\|D3ZF13_RAT | 231.88 | 38 | 38 | 15809000.00 | 21 | 21 | 32 | Oxidation ( | 17514 | Acyl carrier protein OS=Rattus norvegicus OX=10116 GN=Ndufab1 PE=1 SV=1 |
| 16 | 26 | Q641Y2\|NDUS2_RAT | 224.67 | 42 | 42 | 23383000.00 | 35 | 35 | 53 | Oxidation ( | 52562 | NADH dehydrogenase [ubiquinone] iron-sulfur protein 2 mitochondrial OS=Rattus norvegicus OX=10116 GN=Ndufs2 PE=1 SV=1 |
| 22 | 23 | tr\|Q5XIH3\|Q5XIH3_RAT | 223.96 | 34 | 34 | 16564000.00 | 28 | 28 | 43 | Oxidation ( | 50731 | NADH dehydrogenase [ubiquinone] flavoprotein 1 mitochondrial OS=Rattus norvegicus OX=10116 GN=Ndufv1 PE=1 SV=1 |
| 15 | 24 | tr\|D3ZG43\|D3ZG43_RAT | 223.17 | 63 | 63 | 21013000.00 | 36 | 36 | 53 | | 30226 | NADH dehydrogenase (Ubiquinone) Fe-S protein 3 (Predicted) isoform CRA_c OS=Rattus norvegicus OX=10116 GN=Ndufs3 PE=1 SV=1 |
| 29 | 80 | P20788\|UCRI_RAT | 222.59 | 41 | 41 | 12944000.00 | 19 | 19 | 28 | Oxidation ( | 29446 | Cytochrome b-c1 complex subunit Rieske mitochondrial OS=Rattus norvegicus OX=10116 GN=Uqcrfs1 PE=1 SV=2 |
| 33 | 179 | P11240\|COX5A_RAT | 218.25 | 48 | 48 | 10992000.00 | 17 | 17 | 25 | | 16130 | Cytochrome c oxidase subunit 5A mitochondrial OS=Rattus norvegicus OX=10116 GN=Cox5a PE=1 SV=1 |
| 30 | 110 | tr\|A0A0G2JVL6\|A0A0G2JVL6_ | 216.77 | 38 | 38 | 11962000.00 | 18 | 18 | 28 | | 19965 | NADH dehydrogenase [ubiquinone] 1 alpha subcomplex subunit 8 OS=Rattus norvegicus OX=10116 GN=Ndufa8 PE=1 SV=1 |
| 25 | 44 | tr\|D6NSR7\|D6NSR7_RAT | 212.55 | 28 | 28 | 14231000.00 | 25 | 25 | 34 | Oxidation (M) | | Cytochrome b |
| 25 | 46 | tr\|L0L4L8\|L0L4L8_RAT | 212.55 | 28 | 28 | 14231000.00 | 25 | 25 | 34 | Oxidation (M) | | |
| 25 | 47 | tr\|A0A140GE10\|A0A140GE1 | 212.55 | 28 | 28 | 14231000.00 | 25 | 25 | 34 | Oxidation (M) | | |
| 25 | 48 | tr\|A0A140GE11\|A0A140GE1 | 212.55 | 28 | 28 | 14231000.00 | 25 | 25 | 34 | Oxidation (M) | | |
| 25 | 49 | tr\|A0A3Q8AGE8\|A0A3Q8AG | 212.55 | 28 | 28 | 14231000.00 | 25 | 25 | 34 | Oxidation (M) | | |
| 25 | 50 | tr\|A0A3Q8AC68\|A0A3Q8AC | 212.55 | 28 | 28 | 14231000.00 | 25 | 25 | 34 | Oxidation (M) | | |
| 25 | 51 | tr\|A0A3S6FM15\|A0A3S6FM1 | 212.55 | 28 | 28 | 14231000.00 | 25 | 25 | 34 | Oxidation (M) | | |
| 25 | 52 | tr\|A0A220D9Z6\|A0A220D9Z | 212.55 | 28 | 28 | 14231000.00 | 25 | 25 | 34 | Oxidation (M) | | |
| 25 | 53 | tr\|D6NSS6\|D6NSS6_RAT | 212.55 | 28 | 28 | 14231000.00 | 25 | 25 | 34 | Oxidation (M) | | |
| 25 | 54 | tr\|D6NSR3\|D6NSR3_RAT | 212.55 | 28 | 28 | 14231000.00 | 25 | 25 | 34 | Oxidation (M) | | |
| 25 | 55 | tr\|D6NSR8\|D6NSR8_RAT | 212.55 | 28 | 28 | 14231000.00 | 25 | 25 | 34 | Oxidation (M) | | |
| 25 | 56 | tr\|A0A0A1FZ42\|A0A0A1FZ42 | 212.55 | 28 | 28 | 14231000.00 | 25 | 25 | 34 | Oxidation (M) | | |
| 25 | 57 | tr\|D6NSS7\|D6NSS7_RAT | 212.55 | 28 | 28 | 14231000.00 | 25 | 25 | 34 | Oxidation (M) | | |
| 25 | 58 | tr\|Q8SEY9\|Q8SEY9_RAT | 212.55 | 28 | 28 | 14231000.00 | 25 | 25 | 34 | Oxidation (M) | | |
| 25 | 59 | tr\|A0A220DA44\|A0A220DA4 | 212.55 | 28 | 28 | 14231000.00 | 25 | 25 | 34 | Oxidation (M) | | |
| 25 | 60 | tr\|D6NSQ0\|D6NSQ0_RAT | 212.55 | 28 | 28 | 14231000.00 | 25 | 25 | 34 | Oxidation (M) | | |
| 25 | 61 | tr\|Q5UAI7\|Q5UAI7_RAT | 212.55 | 28 | 28 | 14231000.00 | 25 | 25 | 34 | Oxidation (M) | | |
| 25 | 62 | tr\|Q8HIC4\|Q8HIC4_RAT | 212.55 | 28 | 28 | 14231000.00 | 25 | 25 | 34 | Oxidation (M) | | |
| 25 | 63 | tr\|A0A220DA02\|A0A220DA0 | 212.55 | 28 | 28 | 14231000.00 | 25 | 25 | 34 | Oxidation (M) | | |
| 25 | 64 | tr\|A0A0S1Z1V4\|A0A0S1Z1V4 | 212.55 | 28 | 28 | 14231000.00 | 25 | 25 | 34 | Oxidation (M) | | |
| 25 | 65 | P00159\|CYB_RAT | 212.55 | 28 | 28 | 14231000.00 | 25 | 25 | 34 | Oxidation (M) | | |
| 25 | 66 | tr\|L0N311\|L0N311_RAT | 212.55 | 28 | 28 | 14231000.00 | 25 | 25 | 34 | Oxidation (M) | | |
| 25 | 67 | tr\|A0A220DA28\|A0A220DA2 | 212.55 | 28 | 28 | 14231000.00 | 25 | 25 | 34 | Oxidation (M) | | |
| 25 | 68 | tr\|H2KXA0\|H2KXA0_RAT | 212.55 | 28 | 28 | 14231000.00 | 25 | 25 | 34 | Oxidation (M) | | |
| 23 | 28 | Q561S0\|NDUAA_RAT | 207.81 | 38 | 38 | 11638000.00 | 22 | 22 | 38 | Oxidation ( | 40493 | NADH dehydrogenase [ubiquinone] 1 alpha subcomplex subunit 10 mitochondrial OS=Rattus norvegicus OX=10116 GN=Ndufa10 PE=1 SV=1 |
| 37 | 135 | tr\|B0BNE6\|B0BNE6_RAT | 205.64 | 27 | 27 | 5325100.00 | 14 | 14 | 20 | | 23970 | NADH dehydrogenase (Ubiquinone) Fe-S protein 8 (Predicted) isoform CRA_a OS=Rattus norvegicus OX=10116 GN=Ndufs8 PE=1 SV=1 |
| 31 | 36 | tr\|Q06QK5\|Q06QK5_RAT | 203.86 | 18 | 18 | 6938100.00 | 21 | 21 | 25 | Oxidation ( | 68606 | NADH-ubiquinone oxidoreductase chain 5 OS=Rattus norvegicus OX=10116 GN=ND5 PE=3 SV=1 |
| 31 | 37 | P11661\|NU5M_RAT | 203.86 | 18 | 18 | 6938100.00 | 21 | 21 | 25 | Oxidation ( | 68618 | NADH-ubiquinone oxidoreductase chain 5 OS=Rattus norvegicus OX=10116 GN=Mtnd5 PE=3 SV=3 |
| 31 | 38 | tr\|Q06QA1\|Q06QA1_RAT | 203.86 | 18 | 18 | 6938100.00 | 21 | 21 | 25 | Oxidation ( | 68574 | NADH-ubiquinone oxidoreductase chain 5 OS=Rattus norvegicus OX=10116 GN=ND5 PE=3 SV=1 |
| 31 | 39 | tr\|Q8SEZ0\|Q8SEZ0_RAT | 203.86 | 18 | 18 | 6938100.00 | 21 | 21 | 25 | Oxidation ( | 68618 | NADH-ubiquinone oxidoreductase chain 5 OS=Rattus norvegicus OX=10116 GN=Mt-nd5 PE=3 SV=1 |
| 31 | 40 | tr\|A0A096XKT9\|A0A096XKT9 | 203.86 | 18 | 18 | 6938100.00 | 21 | 21 | 25 | Oxidation ( | 68584 | NADH-ubiquinone oxidoreductase chain 5 OS=Rattus norvegicus OX=10116 GN=ND5 PE=3 SV=1 |
| 43 | 133 | tr\|F1LPG5\|F1LPG5_RAT | 200.05 | 51 | 51 | 8287400.00 | 14 | 14 | 17 | | 15064 | NADH:ubiquinone oxidoreductase subunit B4 OS=Rattus norvegicus OX=10116 GN=Ndufb4 PE=1 SV=1 |
| 28 | 41 | tr\|Q5UAJ6\|Q5UAJ6_RAT | 198.9 | 51 | 51 | 9188700.00 | 24 | 24 | 28 | Oxidation ( | 25942 | Cytochrome c oxidase subunit 2 OS=Rattus norvegicus OX=10116 GN=COX2 PE=3 SV=1 |
| 42 | 165 | tr\|D4A565\|D4A565_RAT | 196.9 | 39 | 39 | 7630700.00 | 13 | 13 | 17 | | 21664 | NADH dehydrogenase (Ubiquinone) 1 beta subcomplex 5 (Predicted) isoform CRA_b OS=Rattus norvegicus OX=10116 GN=Ndufb5 PE=1 SV=1 |
| 48 | 75 | P08461\|ODP2_RAT | 180.96 | 17 | 17 | 3516200.00 | 14 | 14 | 14 | | 67166 | Dihydrolipoyllysine-residue acetyltransferase component of pyruvate dehydrogenase complex mitochondrial OS=Rattus norvegicus OX=10116 GN=Dlat PE=1 SV=3 |
| 41 | 111 | Q5XIF3\|NDUS4_RAT | 180.95 | 38 | 38 | 5935600.00 | 15 | 15 | 18 | Oxidation ( | 19741 | NADH dehydrogenase [ubiquinone] iron-sulfur protein 4 mitochondrial OS=Rattus norvegicus OX=10116 GN=Ndufs4 PE=1 SV=1 |
| 36 | 100 | P17764\|THIL_RAT | 179.57 | 22 | 22 | 4587000.00 | 15 | 15 | 20 | | 44695 | Acetyl-CoA acetyltransferase mitochondrial OS=Rattus norvegicus OX=10116 GN=Acat1 PE=1 SV=1 |
| 34 | 34 | tr\|A0A0G2JTL5\|A0A0G2JTL5_ | 173.33 | 15 | 15 | 4533400.00 | 22 | 22 | 24 | Oxidation ( | 140005 | Pyruvate carboxylase mitochondrial OS=Rattus norvegicus OX=10116 GN=Pc PE=1 SV=1 |
| 45 | 183 | tr\|D4A7L4\|D4A7L4_RAT | 168.77 | 32 | 32 | 9803100.00 | 10 | 10 | 16 | | 17634 | NADH dehydrogenase (Ubiquinone) 1 beta subcomplex 11 (Predicted) OS=Rattus norvegicus OX=10116 GN=Ndufb11 PE=1 SV=1 |

| | | | | | | | | | | | | |
|---|---|---|---|---|---|---|---|---|---|---|---|---|
| 35 | 112 | tr\|D4A3V2\|D4A3V2_RAT | 166.64 | 50 | 50 | 5239000.00 | 17 | 17 | 21 | Oxidation ( | 15224 | NADH dehydrogenase [ubiquinone] 1 alpha subcomplex subunit 6 OS=Rattus norvegicus OX=10116 GN=Ndufa6 PE=1 SV=1 |
| 46 | 123 | Q5XI78\|ODO1_RAT | 164.66 | 13 | 13 | 2482200.00 | 10 | 10 | 16 | | 116296 | 2-oxoglutarate dehydrogenase mitochondrial OS=Rattus norvegicus OX=10116 GN=Ogdh PE=1 SV=1 |
| 40 | 33 | Q64428\|ECHA_RAT | 164.65 | 18 | 18 | 3419900.00 | 15 | 15 | 19 | | 82665 | Trifunctional enzyme subunit alpha mitochondrial OS=Rattus norvegicus OX=10116 GN=Hadha PE=1 SV=2 |
| 53 | 246 | tr\|Q5PQZ9\|Q5PQZ9_RAT | 164.55 | 36 | 36 | 11579000.00 | 9 | 9 | 13 | | 14359 | NADH dehydrogenase [ubiquinone] 1 subunit C2 OS=Rattus norvegicus OX=10116 GN=Ndufc2 PE=1 SV=1 |
| 38 | 184 | tr\|B2RYT5\|B2RYT5_RAT | 164.46 | 43 | 43 | 5949200.00 | 13 | 13 | 20 | | 12651 | Cox7a2l protein OS=Rattus norvegicus OX=10116 GN=Cox7a2l PE=2 SV=1 |
| 73 | 230 | tr\|B2RYS2\|B2RYS2_RAT | 156.38 | 34 | 34 | 2689100.00 | 6 | 6 | 6 | Oxidation ( | 13559 | Cytochrome b-c1 complex subunit 7 OS=Rattus norvegicus OX=10116 GN=Uqcrb PE=1 SV=1 |
| 52 | 242 | Q80W89\|NDUAB_RAT | 152.77 | 21 | 21 | 5940400.00 | 9 | 9 | 14 | | 14854 | NADH dehydrogenase [ubiquinone] 1 alpha subcomplex subunit 11 OS=Rattus norvegicus OX=10116 GN=Ndufa11 PE=2 SV=1 |
| 59 | 168 | tr\|F1LXA0\|F1LXA0_RAT | 150.86 | 43 | 43 | 3045100.00 | 10 | 10 | 11 | | 17178 | NADH dehydrogenase [ubiquinone] 1 alpha subcomplex subunit 12 OS=Rattus norvegicus OX=10116 GN=Ndufa12 PE=1 SV=2 |
| 56 | 182 | P10888\|COX41_RAT | 150.09 | 32 | 32 | 5775000.00 | 10 | 10 | 12 | Oxidation ( | 19515 | Cytochrome c oxidase subunit 4 isoform 1 mitochondrial OS=Rattus norvegicus OX=10116 GN=Cox4i1 PE=1 SV=1 |
| 49 | 76 | tr\|Q68G44\|Q68G44_RAT | 150.07 | 20 | 20 | 3789700.00 | 13 | 13 | 14 | | 56886 | 3-hydroxy-3-methylglutaryl coenzyme A synthase OS=Rattus norvegicus OX=10116 GN=Hmgcs2 PE=1 SV=1 |
| 49 | 77 | P22791\|HMCS2_RAT | 150.07 | 20 | 20 | 3789700.00 | 13 | 13 | 14 | | 56912 | Hydroxymethylglutaryl-CoA synthase mitochondrial OS=Rattus norvegicus OX=10116 GN=Hmgcs2 PE=1 SV=1 |
| 54 | 175 | tr\|D3ZCZ9\|D3ZCZ9_RAT | 146.42 | 51 | 51 | 4168000.00 | 9 | 9 | 13 | | 13040 | NADH dehydrogenase [ubiquinone] iron-sulfur protein 6 mitochondrial OS=Rattus norvegicus OX=10116 GN=LOC100912599 PE=1 SV=1 |
| 44 | 171 | tr\|Q8SEZ8\|Q8SEZ8_RAT | 144.78 | 21 | 21 | 3385700.00 | 11 | 11 | 16 | Oxidation ( | 36133 | NADH-ubiquinone oxidoreductase chain 1 (Fragment) OS=Rattus norvegicus OX=10116 GN=NADH1 PE=3 SV=1 |
| 44 | 172 | P03889\|NU1M_RAT | 144.78 | 21 | 21 | 3385700.00 | 11 | 11 | 16 | Oxidation ( | 36145 | NADH-ubiquinone oxidoreductase chain 1 OS=Rattus norvegicus OX=10116 GN=Mtnd1 PE=1 SV=3 |
| 44 | 173 | tr\|Q8HID1\|Q8HID1_RAT | 144.78 | 21 | 21 | 3385700.00 | 11 | 11 | 16 | Oxidation ( | 36145 | NADH-ubiquinone oxidoreductase chain 1 OS=Rattus norvegicus OX=10116 GN=Mt-nd1 PE=3 SV=1 |
| 44 | 174 | tr\|D2E6L7\|D2E6L7_RAT | 144.78 | 21 | 21 | 3385700.00 | 11 | 11 | 16 | Oxidation ( | 36062 | NADH-ubiquinone oxidoreductase chain 1 OS=Rattus norvegicus OX=10116 GN=ND1 PE=3 SV=1 |
| 51 | 148 | tr\|Q5RJN0\|Q5RJN0_RAT | 144.2 | 31 | 31 | 3573900.00 | 11 | 10 | 14 | Oxidation ( | 23945 | NADH dehydrogenase (Ubiquinone) Fe-S protein 7 OS=Rattus norvegicus OX=10116 GN=Ndufs7 PE=1 SV=1 |
| 70 | 226 | tr\|D3ZXF9\|D3ZXF9_RAT | 137.97 | 18 | 18 | 702860.00 | 6 | 6 | 7 | | 29441 | Mitochondrial ribosomal protein L12 OS=Rattus norvegicus OX=10116 GN=Mrpl12 PE=1 SV=1 |
| 58 | 119 | tr\|D3ZE15\|D3ZE15_RAT | 130.87 | 45 | 45 | 2909600.00 | 9 | 9 | 11 | Oxidation ( | 16777 | NADH:ubiquinone oxidoreductase subunit A13 OS=Rattus norvegicus OX=10116 GN=Ndufa13 PE=1 SV=1 |
| 39 | 138 | P05508\|NU4M_RAT | 129.68 | 22 | 22 | 3239700.00 | 16 | 16 | 20 | Oxidation (M) | | NADH-ubiquinone oxidoreductase chain 4 |
| 39 | 139 | tr\|D2E6K0\|D2E6K0_RAT | 129.68 | 22 | 22 | 3239700.00 | 16 | 16 | 20 | Oxidation (M) | | |
| 39 | 140 | tr\|Q06QE1\|Q06QE1_RAT | 129.68 | 22 | 22 | 3239700.00 | 16 | 16 | 20 | Oxidation (M) | | |
| 39 | 141 | tr\|Q06QA2\|Q06QA2_RAT | 129.68 | 22 | 22 | 3239700.00 | 16 | 16 | 20 | Oxidation (M) | | |
| 39 | 142 | tr\|Q7HKW3\|Q7HKW3_RAT | 129.68 | 22 | 22 | 3239700.00 | 16 | 16 | 20 | Oxidation (M) | | |
| 39 | 143 | tr\|Q06Q89\|Q06Q89_RAT | 129.68 | 22 | 22 | 3239700.00 | 16 | 16 | 20 | Oxidation (M) | | |
| 39 | 144 | tr\|Q35737\|Q35737_RAT | 129.68 | 22 | 22 | 3239700.00 | 16 | 16 | 20 | Oxidation (M) | | |
| 39 | 145 | tr\|Q8HIC6\|Q8HIC6_RAT | 129.68 | 22 | 22 | 3239700.00 | 16 | 16 | 20 | Oxidation (M) | | |
| 62 | 196 | Q63362\|NDUA5_RAT | 127.9 | 39 | 39 | 3530600.00 | 7 | 7 | 11 | | 13412 | NADH dehydrogenase [ubiquinone] 1 alpha subcomplex subunit 5 OS=Rattus norvegicus OX=10116 GN=Ndufa5 PE=1 SV=3 |
| 50 | 176 | P13086\|SUCA_RAT | 127.53 | 21 | 21 | 3197700.00 | 10 | 10 | 14 | | 36148 | Succinate--CoA ligase [ADP/GDP-forming] subunit alpha mitochondrial OS=Rattus norvegicus OX=10116 GN=Suclg1 PE=2 SV=2 |
| 50 | 177 | tr\|A0A0H2UHE1\|A0A0H2UHI | 127.53 | 20 | 20 | 3197700.00 | 10 | 10 | 14 | | 37560 | Succinate--CoA ligase [ADP/GDP-forming] subunit alpha mitochondrial OS=Rattus norvegicus OX=10116 GN=Suclg1 PE=1 SV=1 |
| 68 | 228 | P16970\|ABCD3_RAT | 127.43 | 8 | 8 | 2980300.00 | 7 | 7 | 7 | | 75316 | ATP-binding cassette sub-family D member 3 OS=Rattus norvegicus OX=10116 GN=Abcd3 PE=1 SV=3 |
| 79 | 269 | P35738\|ODBB_RAT | 127 | 9 | 9 | 861590.00 | 5 | 5 | 5 | | 42823 | 2-oxoisovalerate dehydrogenase subunit beta mitochondrial OS=Rattus norvegicus OX=10116 GN=Bckdhb PE=1 SV=3 |
| 79 | 270 | tr\|A0A0A0MXW1\|A0A0A0M) | 127 | 9 | 9 | 861590.00 | 5 | 5 | 5 | | 42960 | 2-oxoisovalerate dehydrogenase subunit beta mitochondrial OS=Rattus norvegicus OX=10116 GN=Bckdhb PE=1 SV=1 |
| 64 | 178 | tr\|D3ZZ21\|D3ZZ21_RAT | 126.29 | 51 | 51 | 3983400.00 | 7 | 6 | 10 | Oxidation ( | 15638 | NADH dehydrogenase (Ubiquinone) 1 beta subcomplex 6 (Predicted) OS=Rattus norvegicus OX=10116 GN=Ndufb6 PE=1 SV=1 |
| 66 | 218 | tr\|M0RAM5\|M0RAM5_RAT | 125.48 | 26 | 26 | 1776100.00 | 7 | 7 | 8 | | 22155 | Glutathione peroxidase OS=Rattus norvegicus OX=10116 GN=Gpx1 PE=1 SV=1 |
| 66 | 219 | P04041\|GPX1_RAT | 125.48 | 25 | 25 | 1776100.00 | 7 | 7 | 8 | | 22305 | Glutathione peroxidase 1 OS=Rattus norvegicus OX=10116 GN=Gpx1 PE=1 SV=4 |
| 47 | 169 | Q60587\|ECHB_RAT | 124.51 | 21 | 21 | 3338500.00 | 12 | 12 | 15 | | 51414 | Trifunctional enzyme subunit beta mitochondrial OS=Rattus norvegicus OX=10116 GN=Hadhb PE=1 SV=1 |
| 55 | 153 | tr\|Q06QA9\|Q06QA9_RAT | 119.32 | 14 | 14 | 2915300.00 | 10 | 10 | 12 | | 56893 | Cytochrome c oxidase subunit 1 OS=Rattus norvegicus OX=10116 GN=CO1 PE=3 SV=1 |
| 55 | 154 | tr\|Q06QK0\|Q06QK0_RAT | 119.32 | 14 | 14 | 2915300.00 | 10 | 10 | 12 | | 56880 | Cytochrome c oxidase subunit 1 OS=Rattus norvegicus OX=10116 GN=CO1 PE=3 SV=1 |
| 55 | 155 | tr\|Q8SEZ6\|Q8SEZ6_RAT | 119.32 | 14 | 14 | 2915300.00 | 10 | 10 | 12 | | 56879 | Cytochrome c oxidase subunit 1 OS=Rattus norvegicus OX=10116 GN=COI PE=3 SV=1 |
| 55 | 156 | tr\|A0A0A1FZ34\|A0A0A1FZ34 | 119.32 | 14 | 14 | 2915300.00 | 10 | 10 | 12 | | 56937 | Cytochrome c oxidase subunit 1 OS=Rattus norvegicus OX=10116 GN=COX1 PE=3 SV=1 |
| 55 | 157 | tr\|Q8HIC9\|Q8HIC9_RAT | 119.32 | 14 | 14 | 2915300.00 | 10 | 10 | 12 | | 56845 | Cytochrome c oxidase subunit 1 OS=Rattus norvegicus OX=10116 GN=Mt-co1 PE=3 SV=1 |
| 55 | 158 | P05503\|COX1_RAT | 119.32 | 14 | 14 | 2915300.00 | 10 | 10 | 12 | | 56845 | Cytochrome c oxidase subunit 1 OS=Rattus norvegicus OX=10116 GN=Mtco1 PE=2 SV=3 |
| 55 | 159 | tr\|Q95938\|Q95938_RAT | 119.32 | 14 | 14 | 2915300.00 | 10 | 10 | 12 | | 56977 | Cytochrome c oxidase subunit 1 OS=Rattus norvegicus OX=10116 GN=Co I PE=3 SV=1 |
| 61 | 192 | tr\|B5DEL8\|B5DEL8_RAT | 118.6 | 39 | 39 | 2182200.00 | 6 | 6 | 11 | | 12700 | NADH dehydrogenase (Ubiquinone) Fe-S protein 5 OS=Rattus norvegicus OX=10116 GN=Ndufs5 PE=1 SV=1 |
| 61 | 193 | tr\|A0A0G2JZJ9\|A0A0G2JZJ9_ | 118.6 | 39 | 39 | 2182200.00 | 6 | 6 | 11 | | 12730 | Uncharacterized protein OS=Rattus norvegicus OX=10116 PE=4 SV=1 |
| 67 | 194 | Q9R063\|PRDX5_RAT | 116.7 | 26 | 26 | 1220500.00 | 6 | 6 | 8 | Oxidation ( | 22179 | Peroxiredoxin-5 mitochondrial OS=Rattus norvegicus OX=10116 GN=Prdx5 PE=1 SV=1 |
| 67 | 195 | tr\|A0A0G2JSS8\|A0A0G2JSS8_ | 116.7 | 26 | 26 | 1220500.00 | 6 | 6 | 8 | Oxidation ( | 22207 | Peroxiredoxin OS=Rattus norvegicus OX=10116 GN=Prdx5 PE=1 SV=1 |
| 63 | 216 | Q7TQ16\|QCR8_RAT | 115.27 | 49 | 49 | 3614000.00 | 8 | 8 | 10 | | 9849 | Cytochrome b-c1 complex subunit 8 OS=Rattus norvegicus OX=10116 GN=Uqcrq PE=3 SV=1 |
| 57 | 220 | P12075\|COX5B_RAT | 114.67 | 32 | 32 | 5639800.00 | 8 | 8 | 11 | | 13915 | Cytochrome c oxidase subunit 5B mitochondrial OS=Rattus norvegicus OX=10116 GN=Cox5b PE=1 SV=2 |
| 86 | 261 | P24329\|THTR_RAT | 111.18 | 14 | 14 | 940480.00 | 4 | 4 | 4 | | 33407 | Thiosulfate sulfurtransferase OS=Rattus norvegicus OX=10116 GN=Tst PE=1 SV=3 |
| 76 | 267 | tr\|I6V4L9\|I6V4L9_RAT | 105.29 | 9 | 9 | 729190.00 | 4 | 4 | 5 | Oxidation ( | 29844 | Cytochrome c oxidase subunit 3 OS=Rattus norvegicus OX=10116 GN=COX3 PE=3 SV=1 |
| 80 | 272 | tr\|M0RB63\|M0RB63_RAT | 103.56 | 31 | 31 | 2033100.00 | 3 | 3 | 5 | Oxidation ( | 9372 | RCG63041 OS=Rattus norvegicus OX=10116 GN=LOC684509 PE=4 SV=1 |
| 80 | 281 | tr\|A9UMW2\|A9UMW2_RAT | 103.56 | 31 | 31 | 2033100.00 | 3 | 3 | 5 | Oxidation ( | 9255 | Ndufa3 protein (Fragment) OS=Rattus norvegicus OX=10116 GN=Ndufa3 PE=2 SV=1 |
| 80 | 282 | tr\|A0A0G2KAA3\|A0A0G2KAA | 103.56 | 29 | 29 | 2033100.00 | 3 | 3 | 5 | Oxidation ( | 10170 | NADH:ubiquinone oxidoreductase subunit A3 OS=Rattus norvegicus OX=10116 GN=Ndufa3 PE=1 SV=1 |
| 75 | 243 | tr\|D3ZS58\|D3ZS58_RAT | 102.75 | 34 | 34 | 6916200.00 | 5 | 5 | 6 | Oxidation ( | 10845 | NADH dehydrogenase [ubiquinone] 1 alpha subcomplex subunit 2 OS=Rattus norvegicus OX=10116 GN=Ndufa2 PE=1 SV=1 |
| 60 | 162 | tr\|Q5EBA4\|Q5EBA4_RAT | 101.2 | 22 | 22 | 2917400.00 | 9 | 9 | 11 | Oxidation ( | 33215 | Nipsnap1 protein (Fragment) OS=Rattus norvegicus OX=10116 GN=Nipsnap1 PE=2 SV=1 |
| 60 | 163 | tr\|G3V728\|G3V728_RAT | 101.2 | 22 | 22 | 2917400.00 | 9 | 9 | 11 | Oxidation ( | 33346 | 4-nitrophenylphosphatase domain and non-neuronal SNAP25-like protein homolog 1 (C. elegans) isoform CRA_b OS=Rattus norvegicus OX=10116 GN=Nipsnap1 PE=1 SV=1 |
| 69 | 231 | tr\|Q5M949\|Q5M949_RAT | 100.36 | 17 | 17 | 1926600.00 | 6 | 6 | 7 | | 28340 | Nipsnap homolog 3A (C. elegans) OS=Rattus norvegicus OX=10116 GN=Nipsnap3b PE=1 SV=1 |
| 85 | 232 | tr\|M0R6J0\|M0R6J0_RAT | 99.49 | 11 | 11 | 696830.00 | 3 | 3 | 4 | | 38374 | Mitochondrial ribosomal protein L39 OS=Rattus norvegicus OX=10116 GN=Mrpl39 PE=1 SV=2 |
| 81 | 279 | Q5M9I5\|QCR6_RAT | 98.98 | 21 | 21 | 2029400.00 | 4 | 4 | 5 | | 10424 | Cytochrome b-c1 complex subunit 6 mitochondrial OS=Rattus norvegicus OX=10116 GN=Uqcrh PE=3 SV=1 |
| 71 | 198 | P85834\|EFTU_RAT | 98.26 | 13 | 13 | 823640.00 | 5 | 5 | 6 | | 49522 | Elongation factor Tu mitochondrial OS=Rattus norvegicus OX=10116 GN=Tufm PE=1 SV=1 |
| 74 | 229 | P11960\|ODBA_RAT | 98.1 | 14 | 14 | 857650.00 | 5 | 5 | 6 | | 50165 | 2-oxoisovalerate dehydrogenase subunit alpha mitochondrial (Fragment) OS=Rattus norvegicus OX=10116 GN=Bckdha PE=1 SV=1 |
| 130 | 280 | P31399\|ATP5H_RAT | 95.44 | 15 | 15 | 386280.00 | 2 | 2 | 2 | | 18763 | ATP synthase subunit d mitochondrial OS=Rattus norvegicus OX=10116 GN=Atp5pd PE=1 SV=3 |
| 65 | 249 | tr\|Q06QE9\|Q06QE9_RAT | 94.01 | 14 | 14 | 1087900.00 | 5 | 5 | 8 | | 38598 | NADH-ubiquinone oxidoreductase chain 2 OS=Rattus norvegicus OX=10116 GN=ND2 PE=3 SV=1 |
| 65 | 250 | tr\|Q06Q97\|Q06Q97_RAT | 94.01 | 14 | 14 | 1087900.00 | 5 | 5 | 8 | | 38485 | NADH-ubiquinone oxidoreductase chain 2 OS=Rattus norvegicus OX=10116 GN=ND2 PE=3 SV=1 |
| 65 | 251 | tr\|A0A0A1G491\|A0A0A1G49 | 94.01 | 14 | 14 | 1087900.00 | 5 | 5 | 8 | | 38542 | NADH-ubiquinone oxidoreductase chain 2 OS=Rattus norvegicus OX=10116 GN=ND2 PE=3 SV=1 |
| 65 | 252 | tr\|Q5UAJ8\|Q5UAJ8_RAT | 94.01 | 14 | 14 | 1087900.00 | 5 | 5 | 8 | | 38455 | NADH-ubiquinone oxidoreductase chain 2 OS=Rattus norvegicus OX=10116 GN=ND2 PE=3 SV=1 |
| 65 | 253 | tr\|Q8HID0\|Q8HID0_RAT | 94.01 | 14 | 14 | 1087900.00 | 5 | 5 | 8 | | 38653 | NADH-ubiquinone oxidoreductase chain 2 OS=Rattus norvegicus OX=10116 GN=Mt-nd2 PE=3 SV=1 |
| 65 | 254 | P11662\|NU2M_RAT | 94.01 | 14 | 14 | 1087900.00 | 5 | 5 | 8 | | 38653 | NADH-ubiquinone oxidoreductase chain 2 OS=Rattus norvegicus OX=10116 GN=Mtnd2 PE=3 SV=3 |
| 65 | 255 | tr\|Q06QH5\|Q06QH5_RAT | 94.01 | 14 | 14 | 1087900.00 | 5 | 5 | 8 | | 38626 | NADH-ubiquinone oxidoreductase chain 2 OS=Rattus norvegicus OX=10116 GN=ND2 PE=3 SV=1 |

| | | | | | | | | | | | | |
|---|---|---|---|---|---|---|---|---|---|---|---|---|
| 65 | 256 | tr\|Q06QB0\|Q06QB0_RAT | 94.01 | 14 | 14 | 1087900.00 | 5 | 5 | 8 | | 38534 | NADH-ubiquinone oxidoreductase chain 2 OS=Rattus norvegicus OX=10116 GN=ND2 PE=3 SV=1 |
| 65 | 257 | tr\|D2E6P4\|D2E6P4_RAT | 94.01 | 14 | 14 | 1087900.00 | 5 | 5 | 8 | | 38623 | NADH-ubiquinone oxidoreductase chain 2 OS=Rattus norvegicus OX=10116 GN=ND2 PE=3 SV=1 |
| 65 | 258 | tr\|Q8SEZ7\|Q8SEZ7_RAT | 94.01 | 14 | 14 | 1087900.00 | 5 | 5 | 8 | | 38580 | NADH-ubiquinone oxidoreductase chain 2 (Fragment) OS=Rattus norvegicus OX=10116 GN=NADH2 PE=3 SV=1 |
| 78 | 263 | tr\|A0A0G2JSH5\|A0A0G2JSH5 | 84.93 | 4 | 4 | 1530300.00 | 4 | 4 | 5 | Formylatio | 68759 | Serum albumin OS=Rattus norvegicus OX=10116 GN=Alb PE=1 SV=1 |
| 78 | 264 | P02770\|ALBU_RAT | 84.93 | 4 | 4 | 1530300.00 | 4 | 4 | 5 | Formylatio | 68731 | Serum albumin OS=Rattus norvegicus OX=10116 GN=Alb PE=1 SV=2 |
| 87 | 233 | P29147\|BDH_RAT | 83.22 | 11 | 11 | 654630.00 | 4 | 4 | 4 | | 38202 | D-beta-hydroxybutyrate dehydrogenase mitochondrial OS=Rattus norvegicus OX=10116 GN=Bdh1 PE=1 SV=2 |
| 87 | 234 | tr\|A0A0G2JSH2\|A0A0G2JSH2 | 83.22 | 11 | 11 | 654630.00 | 4 | 4 | 4 | | 38333 | 3-hydroxybutyrate dehydrogenase type 1 isoform CRA_a OS=Rattus norvegicus OX=10116 GN=Bdh1 PE=1 SV=1 |
| 99 | 291 | tr\|Q5XFW4\|Q5XFW4_RAT | 77.86 | 15 | 15 | 440540.00 | 3 | 3 | 3 | | 20541 | Mitochondrial ribosomal protein L13 OS=Rattus norvegicus OX=10116 GN=Mrpl13 PE=1 SV=1 |
| 82 | 347 | P80433\|COX8A_RAT | 77.35 | 19 | 19 | 2079300.00 | 3 | 3 | 5 | | 7672 | Cytochrome c oxidase subunit 8A mitochondrial OS=Rattus norvegicus OX=10116 GN=Cox8a PE=1 SV=3 |
| 72 | 248 | tr\|B2RYW3\|B2RYW3_RAT | 76.03 | 23 | 23 | 1659100.00 | 5 | 5 | 6 | Oxidation ( | 21892 | NADH dehydrogenase (Ubiquinone) 1 beta subcomplex 9 OS=Rattus norvegicus OX=10116 GN=Ndufb9 PE=1 SV=1 |
| 97 | 268 | P19511\|AT5F1_RAT | 71.92 | 9 | 9 | 1783700.00 | 2 | 2 | 3 | | 28869 | ATP synthase F(0) complex subunit B1 mitochondrial OS=Rattus norvegicus OX=10116 GN=Atp5pb PE=1 SV=1 |
| 98 | 290 | tr\|Q4G067\|Q4G067_RAT | 69.7 | 8 | 8 | 545360.00 | 3 | 3 | 3 | | 37440 | Mitochondrial ribosomal protein L44 OS=Rattus norvegicus OX=10116 GN=Mrpl44 PE=1 SV=1 |
| 131 | 283 | P80431\|COX7B_RAT | 68.41 | 28 | 28 | 587520.00 | 2 | 2 | 2 | | 8995 | Cytochrome c oxidase subunit 7B mitochondrial OS=Rattus norvegicus OX=10116 GN=Cox7b PE=1 SV=3 |
| 100 | 284 | tr\|A0A0G2K8Q8\|A0A0G2K8C | 67.89 | 29 | 29 | 571820.00 | 3 | 3 | 3 | | 7099 | Ubiquinol-cytochrome c reductase complex III subunit X OS=Rattus norvegicus OX=10116 GN=Uqcr10 PE=1 SV=1 |
| 100 | 285 | tr\|B2RYX1\|B2RYX1_RAT | 67.89 | 28 | 28 | 571820.00 | 3 | 3 | 3 | | 7462 | LOC685322 protein OS=Rattus norvegicus OX=10116 GN=Uqcr10 PE=2 SV=1 |
| 101 | 286 | tr\|A9UMV9\|A9UMV9_RAT | 66.48 | 21 | 21 | 2811100.00 | 3 | 3 | 3 | | 12500 | NADH:ubiquinone oxidoreductase subunit A7 OS=Rattus norvegicus OX=10116 GN=Ndufa7 PE=1 SV=1 |
| 123 | 271 | Q7TT47\|SPG7_RAT | 65.42 | 3 | 3 | 222660.00 | 2 | 2 | 2 | | 86103 | Paraplegin OS=Rattus norvegicus OX=10116 GN=Spg7 PE=2 SV=2 |
| 171 | 300 | tr\|A0A0G2K9B4\|A0A0G2K9B4 | 62.86 | 6 | 6 | 0.00 | 1 | 1 | 1 | | 33652 | Mitochondrial ribosomal protein L15 OS=Rattus norvegicus OX=10116 GN=Mrpl15 PE=1 SV=1 |
| 171 | 328 | tr\|D4A4B1\|D4A4B1_RAT | 62.86 | 9 | 9 | 0.00 | 1 | 1 | 1 | | 23223 | Mitochondrial ribosomal protein L15 OS=Rattus norvegicus OX=10116 GN=Mrpl15 PE=1 SV=2 |
| 111 | 266 | P04762\|CATA_RAT | 62.17 | 4 | 4 | 146900.00 | 2 | 2 | 2 | | 59757 | Catalase OS=Rattus norvegicus OX=10116 GN=Cat PE=1 SV=3 |
| 77 | 245 | P18163\|ACSL1_RAT | 58.17 | 4 | 4 | 563690.00 | 4 | 3 | 5 | | 78179 | Long-chain-fatty-acid--CoA ligase 1 OS=Rattus norvegicus OX=10116 GN=Acsl1 PE=1 SV=1 |
| 205 | 322 | Q09073\|ADT2_RAT | 57.34 | 3 | 3 | 168630.00 | 1 | 1 | 1 | | 32901 | ADP/ATP translocase 2 OS=Rattus norvegicus OX=10116 GN=Slc25a5 PE=1 SV=3 |
| 96 | 244 | P07824\|ARGI1_RAT | 56.58 | 12 | 12 | 1022000.00 | 3 | 3 | 3 | | 34973 | Arginase-1 OS=Rattus norvegicus OX=10116 GN=Arg1 PE=1 SV=2 |
| 255 | 343 | tr\|D3ZX69\|D3ZX69_RAT | 56.07 | 5 | 5 | 350330.00 | 1 | 1 | 1 | | 29466 | 39S ribosomal protein L10 mitochondrial OS=Rattus norvegicus OX=10116 GN=Mrpl10 PE=1 SV=1 |
| 255 | 344 | P0C2C4\|RM10_RAT | 56.07 | 5 | 5 | 350330.00 | 1 | 1 | 1 | | 29940 | 39S ribosomal protein L10 mitochondrial OS=Rattus norvegicus OX=10116 GN=Mrpl10 PE=1 SV=1 |
| 256 | 345 | tr\|D3ZPE6\|D3ZPE6_RAT | 55.09 | 8 | 8 | 248030.00 | 1 | 1 | 1 | | 15047 | Mitochondrial ribosomal protein L51 OS=Rattus norvegicus OX=10116 GN=Mrpl51 PE=1 SV=1 |
| 125 | 325 | P11951\|CX6C2_RAT | 52.85 | 11 | 11 | 296770.00 | 2 | 2 | 2 | | 8455 | Cytochrome c oxidase subunit 6C-2 OS=Rattus norvegicus OX=10116 GN=Cox6c2 PE=1 SV=3 |
| 257 | 346 | Q63750\|RM23_RAT | 51.43 | 18 | 18 | 240890.00 | 1 | 1 | 1 | | 17050 | 39S ribosomal protein L23 mitochondrial OS=Rattus norvegicus OX=10116 GN=Mrpl23 PE=2 SV=1 |
| 132 | 289 | tr\|D3ZFJ6\|D3ZFJ6_RAT | 48.84 | 6 | 6 | 326800.00 | 2 | 2 | 2 | | 60420 | Lactamase beta OS=Rattus norvegicus OX=10116 GN=Lactb PE=1 SV=1 |
| 258 | 348 | P09139\|SPYA_RAT | 48.27 | 3 | 3 | 344780.00 | 1 | 1 | 1 | | 45834 | Serine--pyruvate aminotransferase mitochondrial OS=Rattus norvegicus OX=10116 GN=Agxt PE=1 SV=1 |
| 133 | 295 | Q8VID1\|DHRS4_RAT | 47.03 | 9 | 9 | 411400.00 | 2 | 2 | 2 | | 29822 | Dehydrogenase/reductase SDR family member 4 OS=Rattus norvegicus OX=10116 GN=Dhrs4 PE=2 SV=2 |
| 12 | 368 | tr\|A0A0G2JWK2\|A0A0G2JWK | 45.06 | 1 | 1 | 2837900.00 | 1 | 1 | 70 | | 53049 | Methyl-CpG-binding protein 2 OS=Rattus norvegicus OX=10116 GN=Mecp2 PE=1 SV=1 |
| 12 | 369 | Q00566\|MECP2_RAT | 45.06 | 1 | 1 | 2837900.00 | 1 | 1 | 70 | | 53048 | Methyl-CpG-binding protein 2 OS=Rattus norvegicus OX=10116 GN=Mecp2 PE=1 SV=1 |
| 91 | 273 | tr\|A0A0G2JVH4\|A0A0G2JVH4 | 44.45 | 3 | 3 | 2584700.00 | 2 | 2 | 3 | | 86230 | MICOS complex subunit MIC60 OS=Rattus norvegicus OX=10116 GN=Immt PE=1 SV=1 |
| 259 | 350 | tr\|Q35733\|Q35733_RAT | 44.41 | 13 | 13 | 0.00 | 1 | 1 | 1 | | 8389 | NADH-ubiquinone oxidoreductase chain 6 (Fragment) OS=Rattus norvegicus OX=10116 PE=3 SV=1 |
| 259 | 351 | tr\|Q06QD9\|Q06QD9_RAT | 44.41 | 6 | 6 | 0.00 | 1 | 1 | 1 | | 18943 | NADH-ubiquinone oxidoreductase chain 6 OS=Rattus norvegicus OX=10116 GN=ND6 PE=3 SV=1 |
| 259 | 352 | tr\|Q7HKW2\|Q7HKW2_RAT | 44.41 | 6 | 6 | 0.00 | 1 | 1 | 1 | | 18957 | NADH-ubiquinone oxidoreductase chain 6 OS=Rattus norvegicus OX=10116 GN=NADH6 PE=3 SV=1 |
| 134 | 312 | P0DN35\|NDUB1_RAT | 43.81 | 37 | 37 | 498660.00 | 2 | 2 | 2 | | 6998 | NADH dehydrogenase [ubiquinone] 1 beta subcomplex subunit 1 OS=Rattus norvegicus OX=10116 GN=Ndufb1 PE=3 SV=1 |
| 260 | 354 | tr\|Q3B8N9\|Q3B8N9_RAT | 42.77 | 3 | 3 | 130870.00 | 1 | 1 | 1 | | 32823 | Biphenyl hydrolase-like OS=Rattus norvegicus OX=10116 GN=Bphl PE=1 SV=1 |
| 261 | 355 | P80432\|COX7C_RAT | 42.07 | 16 | 16 | 194060.00 | 1 | 1 | 1 | | 7375 | Cytochrome c oxidase subunit 7C mitochondrial OS=Rattus norvegicus OX=10116 GN=Cox7c PE=1 SV=2 |
| 206 | 330 | tr\|F1M953\|F1M953_RAT | 41.52 | 2 | 2 | 107350.00 | 1 | 1 | 1 | | 73745 | Stress-70 protein mitochondrial OS=Rattus norvegicus OX=10116 GN=Hspa9 PE=1 SV=1 |
| 206 | 331 | P48721\|GRP75_RAT | 41.52 | 2 | 2 | 107350.00 | 1 | 1 | 1 | | 73858 | Stress-70 protein mitochondrial OS=Rattus norvegicus OX=10116 GN=Hspa9 PE=1 SV=3 |
| 172 | 307 | Q5XIA8\|GHITM_RAT | 41.14 | 3 | 3 | 103700.00 | 1 | 1 | 1 | | 37178 | Growth hormone-inducible transmembrane protein OS=Rattus norvegicus OX=10116 GN=Ghitm PE=2 SV=1 |
| 262 | 356 | P10818\|CX6A1_RAT | 40.12 | 10 | 10 | 228910.00 | 1 | 1 | 1 | Oxidation ( | 12301 | Cytochrome c oxidase subunit 6A1 mitochondrial OS=Rattus norvegicus OX=10116 GN=Cox6a1 PE=1 SV=2 |
| 114 | 326 | P05696\|KPCA_RAT | 39.15 | 1 | 1 | 555840.00 | 1 | 1 | 2 | | 76792 | Protein kinase C alpha type OS=Rattus norvegicus OX=10116 GN=Prkca PE=1 SV=3 |
| 136 | 357 | tr\|D3ZLT1\|D3ZLT1_RAT | 38.66 | 8 | 8 | 482450.00 | 1 | 1 | 2 | Oxidation ( | 16568 | NADH dehydrogenase (Ubiquinone) 1 beta subcomplex 7 (Predicted) OS=Rattus norvegicus OX=10116 GN=Ndufb7 PE=1 SV=1 |
| 135 | 324 | P33124\|ACSL6_RAT | 38.24 | 2 | 2 | | 2 | 0 | 2 | | 78180 | Long-chain-fatty-acid--CoA ligase 6 OS=Rattus norvegicus OX=10116 GN=Acsl6 PE=1 SV=1 |
| 263 | 358 | tr\|A0A0H2UHR5\|A0A0H2UHI | 38.03 | 3 | 3 | 44636.00 | 1 | 1 | 1 | | 46214 | Protein-serine/threonine kinase OS=Rattus norvegicus OX=10116 GN=Bckdk PE=1 SV=1 |
| 263 | 359 | Q00972\|BCKD_RAT | 38.03 | 3 | 3 | 44636.00 | 1 | 1 | 1 | | 46474 | [3-methyl-2-oxobutanoate dehydrogenase [lipoamide]] kinase mitochondrial OS=Rattus norvegicus OX=10116 GN=Bckdk PE=1 SV=2 |
| 264 | 362 | tr\|G3V9S6\|G3V9S6_RAT | 36.42 | 14 | 14 | 139720.00 | 1 | 1 | 1 | | 8148 | NADH dehydrogenase [ubiquinone] 1 alpha subcomplex subunit 1 OS=Rattus norvegicus OX=10116 GN=Ndufa1 PE=1 SV=2 |
| 102 | 406 | tr\|D3ZD23\|D3ZD23_RAT | 35.06 | 1 | 1 | 817600.00 | 2 | 2 | 3 | | 67300 | ATP-binding cassette subfamily E member 1 OS=Rattus norvegicus OX=10116 GN=Abce1 PE=1 SV=1 |
| 115 | 329 | Q06437\|ODPAT_RAT | 34.37 | 7 | 7 | 2018600.00 | 2 | 2 | 2 | Carbamidomethylatioı | | Pyruvate dehydrogenase E1 component subunit alpha, testis-specific form, mitochondrial |
| 117 | 293 | Q9WVK3\|PECR_RAT | 34.35 | 7 | 7 | 530840.00 | 2 | 2 | 2 | Oxidation ( | 32433 | Peroxisomal trans-2-enoyl-CoA reductase OS=Rattus norvegicus OX=10116 GN=Pecr PE=2 SV=1 |
| 117 | 294 | tr\|A0A0G2JVG4\|A0A0G2JVG4 | 34.35 | 7 | 7 | 530840.00 | 2 | 2 | 2 | Oxidation ( | 32737 | Peroxisomal trans-2-enoyl-CoA reductase OS=Rattus norvegicus OX=10116 GN=Pecr PE=1 SV=1 |
| 207 | 332 | Q9ER34\|ACON_RAT | 34.16 | 1 | 1 | 171590.00 | 1 | 1 | 1 | | 85433 | Aconitate hydratase mitochondrial OS=Rattus norvegicus OX=10116 GN=Aco2 PE=1 SV=2 |
| 106 | 380 | O70351\|HCD2_RAT | 33.38 | 8 | 8 | 30581.00 | 2 | 2 | 2 | | 27246 | 3-hydroxyacyl-CoA dehydrogenase type-2 OS=Rattus norvegicus OX=10116 GN=Hsd17b10 PE=1 SV=3 |
| 106 | 381 | tr\|B0BMW2\|B0BMW2_RAT | 33.38 | 8 | 8 | 30581.00 | 2 | 2 | 2 | | 27250 | 3-hydroxyacyl-CoA dehydrogenase type-2 OS=Rattus norvegicus OX=10116 GN=Hsd17b10 PE=1 SV=1 |
| 208 | 335 | P28494\|MA2A1_RAT | 32.84 | 1 | 1 | 69802.00 | 1 | 1 | 1 | | 131242 | Alpha-mannosidase 2 OS=Rattus norvegicus OX=10116 GN=Man2a1 PE=1 SV=2 |
| 89 | 317 | tr\|A0A0G2JYD4\|A0A0G2JYD4 | 31.49 | 1 | 1 | 651930.00 | 3 | 3 | 3 | Formylatio | 487857 | Vacuolar protein sorting 13 homolog D OS=Rattus norvegicus OX=10116 GN=Vps13d PE=1 SV=1 |
| 89 | 318 | tr\|D3ZKC6\|D3ZKC6_RAT | 31.49 | 1 | 1 | 651930.00 | 3 | 3 | 3 | Formylatio | 488962 | Vacuolar protein sorting 13 homolog D OS=Rattus norvegicus OX=10116 GN=Vps13d PE=1 SV=1 |
| 187 | 309 | P63039\|CH60_RAT | 30.71 | 2 | 2 | 145330.00 | 1 | 1 | 1 | | 60956 | 60 kDa heat shock protein mitochondrial OS=Rattus norvegicus OX=10116 GN=Hspd1 PE=1 SV=1 |
| 187 | 310 | tr\|A0A482IDN3\|A0A482IDN3 | 30.71 | 2 | 2 | 145330.00 | 1 | 1 | 1 | | 60956 | Hsp60 OS=Rattus norvegicus OX=10116 GN=Hspd1 PE=2 SV=1 |
| 266 | 373 | Q4KLP0\|DHTK1_RAT | 30.6 | 1 | 1 | 0.00 | 1 | 1 | 1 | | 102642 | Probable 2-oxoglutarate dehydrogenase E1 component DHKTD1 mitochondrial OS=Rattus norvegicus OX=10116 GN=Dhtkd1 PE=2 SV=1 |
| 170 | 297 | tr\|Q6QI09\|Q6QI09_RAT | 30.39 | 2 | 2 | 0.00 | 1 | 1 | 1 | | 67721 | ATP synthase subunit gamma mitochondrial OS=Rattus norvegicus OX=10116 GN=Taf3 PE=1 SV=1 |
| 170 | 301 | P35435\|ATPG_RAT | 30.39 | 4 | 4 | 0.00 | 1 | 1 | 1 | | 30191 | ATP synthase subunit gamma mitochondrial OS=Rattus norvegicus OX=10116 GN=Atp5f1c PE=1 SV=2 |
| 209 | 338 | tr\|A0A0G2JYU2\|A0A0G2JYU2 | 30.08 | 4 | 4 | 1354800.00 | 1 | 1 | 1 | | 20750 | Mitochondrial ribosomal protein L11 OS=Rattus norvegicus OX=10116 GN=mrpl11 PE=1 SV=1 |
| 209 | 339 | Q5XIE3\|RM11_RAT | 30.08 | 4 | 4 | 1354800.00 | 1 | 1 | 1 | | 22420 | 39S ribosomal protein L11 mitochondrial OS=Rattus norvegicus OX=10116 GN=Mrpl11 PE=2 SV=1 |
| 137 | 374 | P48037\|ANXA6_RAT | 29.99 | 1 | 1 | 441900.00 | 1 | 1 | 2 | | 75754 | Annexin A6 OS=Rattus norvegicus OX=10116 GN=Anxa6 PE=1 SV=2 |
| 137 | 375 | tr\|Q6IMZ3\|Q6IMZ3_RAT | 29.99 | 1 | 1 | 441900.00 | 1 | 1 | 2 | | 75756 | Annexin OS=Rattus norvegicus OX=10116 GN=Anxa6 PE=1 SV=1 |
| 94 | 341 | Q63563\|ABCC9_RAT | 28.83 | 1 | 1 | 1451100.00 | 2 | 2 | 3 | | 174117 | ATP-binding cassette sub-family C member 9 OS=Rattus norvegicus OX=10116 GN=Abcc9 PE=1 SV=1 |
| 267 | 377 | A0A0G2JZ79\|SIR1_RAT | 28.73 | 2 | 2 | 308790.00 | 1 | 1 | 1 | | 62059 | NAD-dependent protein deacetylase sirtuin-1 OS=Rattus norvegicus OX=10116 GN=Sirt1 PE=3 SV=2 |

| | | | | | | | | | | | | |
|---|---|---|---|---|---|---|---|---|---|---|---|---|
| 267 | 378 | tr\|A0A182DWI7\|A0A182DWI | 28.73 | 1 | 1 | 308790.00 | 1 | 1 | 1 | | 81778 | NAD-dependent protein deacetylase sirtuin-1 OS=Rattus norvegicus OX=10116 GN=Sirt1 PE=4 SV=1 |
| 268 | 382 | Q01062\|PDE2A_RAT | 28.44 | 2 | 2 | 8010800.00 | 1 | 1 | 1 | | 104664 | cGMP-dependent 3' 5'-cyclic phosphodiesterase OS=Rattus norvegicus OX=10116 GN=Pde2a PE=1 SV=2 |
| 268 | 383 | tr\|F8WFW5\|F8WFW5_RAT | 28.44 | 1 | 1 | 8010800.00 | 1 | 1 | 1 | | 105234 | Phosphodiesterase OS=Rattus norvegicus OX=10116 GN=Pde2a PE=1 SV=1 |
| 269 | 385 | tr\|B1WBP7\|B1WBP7_RAT | 27.42 | 7 | 7 | 95662.00 | 1 | 1 | 1 | | 12884 | ATP synthase subunit delta mitochondrial OS=Rattus norvegicus OX=10116 GN=Atp5f1d PE=1 SV=1 |
| 269 | 386 | P35434\|ATPD_RAT | 27.42 | 5 | 5 | 95662.00 | 1 | 1 | 1 | | 17595 | ATP synthase subunit delta mitochondrial OS=Rattus norvegicus OX=10116 GN=Atp5f1d PE=1 SV=2 |
| 269 | 387 | tr\|G3V7Y3\|G3V7Y3_RAT | 27.42 | 5 | 5 | 95662.00 | 1 | 1 | 1 | | 17563 | ATP synthase subunit delta mitochondrial OS=Rattus norvegicus OX=10116 GN=Atp5f1d PE=1 SV=1 |
| 270 | 388 | P83565\|RM40_RAT | 27 | 8 | 8 | 210140.00 | 1 | 1 | 1 | | 24397 | 39S ribosomal protein L40 mitochondrial OS=Rattus norvegicus OX=10116 GN=Mrpl40 PE=1 SV=2 |
| 107 | 441 | P11530\|DMD_RAT | 26.92 | 0 | 0 | 2142800.00 | 2 | 2 | 2 | | 425830 | Dystrophin OS=Rattus norvegicus OX=10116 GN=Dmd PE=1 SV=2 |
| 156 | 314 | tr\|D3ZHK4\|D3ZHK4_RAT | 25.68 | 0 | 0 | 0.00 | 1 | 1 | 1 | | 182226 | RB1-inducible coiled-coil 1 OS=Rattus norvegicus OX=10116 GN=Rb1cc1 PE=1 SV=1 |
| 271 | 396 | tr\|D3ZH23\|D3ZH23_RAT | 25.53 | 13 | 13 | 18559.00 | 1 | 1 | 1 | Oxidation ( | 15195 | Similar to mitochondrial ribosomal protein L41 OS=Rattus norvegicus OX=10116 GN=RGD1560917 PE=4 SV=1 |
| 271 | 397 | Q5BJX1\|RM41_RAT | 25.53 | 13 | 13 | 18559.00 | 1 | 1 | 1 | Oxidation ( | 15193 | 39S ribosomal protein L41 mitochondrial OS=Rattus norvegicus OX=10116 GN=Mrpl41 PE=1 SV=1 |
| 211 | 376 | tr\|D3ZDP2\|D3ZDP2_RAT | 25.29 | 4 | 4 | 130930.00 | 1 | 1 | 1 | | 23470 | Mitochondrial ribosomal protein L58 OS=Rattus norvegicus OX=10116 GN=Mrpl58 PE=1 SV=1 |
| 273 | 416 | tr\|Q5UAJ5\|Q5UAJ5_RAT | 24.18 | 13 | 13 | 267640.00 | 1 | 1 | 1 | | 7642 | ATP synthase protein 8 OS=Rattus norvegicus OX=10116 GN=ATP8 PE=3 SV=1 |
| 273 | 417 | P11608\|ATP8_RAT | 24.18 | 13 | 13 | 267640.00 | 1 | 1 | 1 | | 7630 | ATP synthase protein 8 OS=Rattus norvegicus OX=10116 GN=Mt-atp8 PE=1 SV=2 |
| 273 | 418 | tr\|Q8SEZ4\|Q8SEZ4_RAT | 24.18 | 13 | 13 | 267640.00 | 1 | 1 | 1 | | 7632 | ATP synthase protein 8 OS=Rattus norvegicus OX=10116 GN=ATPase8 PE=3 SV=1 |
| 273 | 419 | tr\|Q8HIC8\|Q8HIC8_RAT | 24.18 | 13 | 13 | 267640.00 | 1 | 1 | 1 | | 7630 | ATP synthase protein 8 OS=Rattus norvegicus OX=10116 GN=Mt-atp8 PE=3 SV=1 |
| 274 | 420 | tr\|F1LUC0\|F1LUC0_RAT | 24.08 | 2 | 2 | 79994.00 | 1 | 1 | 1 | Formylatio | 72937 | Similar to RIKEN cDNA 5730410E15 gene (Predicted) isoform CRA_a OS=Rattus norvegicus OX=10116 GN=Sybu PE=1 SV=2 |
| 277 | 431 | Q32PX9\|AFG1L_RAT | 23.34 | 2 | 2 | 789440.00 | 1 | 1 | 1 | Formylatio | 54503 | AFG1-like ATPase OS=Rattus norvegicus OX=10116 GN=Afg1l PE=2 SV=1 |
| 173 | 320 | tr\|D3ZYB6\|D3ZYB6_RAT | 23.23 | 1 | 1 | 149470.00 | 1 | 1 | 1 | | 135817 | DNA-directed RNA polymerase OS=Rattus norvegicus OX=10116 GN=Polrmt PE=3 SV=2 |
| 278 | 432 | P61980\|HNRPK_RAT | 22.99 | 2 | 2 | 48816.00 | 1 | 1 | 1 | | 50976 | Heterogeneous nuclear ribonucleoprotein K OS=Rattus norvegicus OX=10116 GN=Hnrnpk PE=1 SV=1 |
| 212 | 392 | Q6AYC2\|IRGM_RAT | 22.73 | 2 | 2 | 0.00 | 1 | 1 | 1 | | 46338 | Immunity-related GTPase family M protein OS=Rattus norvegicus OX=10116 GN=Irgm PE=2 SV=1 |
| 213 | 400 | tr\|D4A7Q5\|D4A7Q5_RAT | 22.58 | 1 | 1 | 890830.00 | 1 | 1 | 1 | | 59940 | DEAD (Asp-Glu-Ala-Asp) box polypeptide 28 (Predicted) OS=Rattus norvegicus OX=10116 GN=Ddx28 PE=4 SV=1 |
| 216 | 407 | tr\|D3ZXQ6\|D3ZXQ6_RAT | 22.58 | 3 | 3 | 761030.00 | 1 | 1 | 1 | | 46897 | LETM1 domain-containing protein LETM2 mitochondrial OS=Rattus norvegicus OX=10116 GN=Letm2 PE=4 SV=3 |
| 216 | 408 | Q5PQQ5\|LETM2_RAT | 22.58 | 3 | 3 | 761030.00 | 1 | 1 | 1 | | 52524 | LETM1 domain-containing protein LETM2 mitochondrial OS=Rattus norvegicus OX=10116 GN=Letm2 PE=2 SV=1 |
| 216 | 409 | tr\|F1LP77\|F1LP77_RAT | 22.58 | 3 | 3 | 761030.00 | 1 | 1 | 1 | | 52501 | LETM1 domain-containing protein LETM2 mitochondrial OS=Rattus norvegicus OX=10116 GN=Letm2 PE=4 SV=3 |
| 116 | 403 | Q01728\|NAC1_RAT | 22.12 | 1 | 1 | 4155100.00 | 1 | 1 | 2 | Formylatio | 108184 | Sodium/calcium exchanger 1 OS=Rattus norvegicus OX=10116 GN=Slc8a1 PE=1 SV=3 |
| 116 | 404 | P70549\|NAC3_RAT | 22.12 | 1 | 1 | 4155100.00 | 1 | 1 | 2 | Formylatio | 103163 | Sodium/calcium exchanger 3 OS=Rattus norvegicus OX=10116 GN=Slc8a3 PE=1 SV=1 |
| 148 | 536 | P15205\|MAP1B_RAT | 22.12 | 0 | 0 | | 1 | 0 | 1 | | 269640 | Microtubule-associated protein 1B OS=Rattus norvegicus OX=10116 GN=Map1b PE=1 SV=3 |
| 215 | 405 | tr\|D3ZVS2\|D3ZVS2_RAT | 22.08 | 2 | 2 | 405220.00 | 1 | 1 | 1 | | 50733 | L-2-hydroxyglutarate dehydrogenase OS=Rattus norvegicus OX=10116 GN=L2hgdh PE=1 SV=1 |
| 279 | 435 | tr\|A0A0G2K4M8\|A0A0G2K4I | 22.01 | 2 | 2 | 164510.00 | 1 | 1 | 1 | Formylatio | 47158 | Acyl-CoA thioesterase 3 OS=Rattus norvegicus OX=10116 GN=Acot3 PE=1 SV=1 |
| 181 | 456 | tr\|A0A0G2K5L6\|A0A0G2K5L | 21.77 | 0 | 0 | 103530.00 | 1 | 1 | 1 | | 275755 | Acetyl-CoA carboxylase beta OS=Rattus norvegicus OX=10116 GN=Acacb PE=1 SV=1 |
| 181 | 457 | tr\|A0A0G2K1F2\|A0A0G2K1F | 21.77 | 0 | 0 | 103530.00 | 1 | 1 | 1 | | 276393 | Acetyl-CoA carboxylase beta OS=Rattus norvegicus OX=10116 GN=Acacb PE=1 SV=1 |
| 181 | 458 | tr\|O70151\|O70151_RAT | 21.77 | 0 | 0 | 103530.00 | 1 | 1 | 1 | | 276097 | Acetyl-CoA carboxylase OS=Rattus norvegicus OX=10116 GN=Acacb PE=2 SV=1 |
| 181 | 459 | tr\|D3ZBE2\|D3ZBE2_RAT | 21.77 | 0 | 0 | 103530.00 | 1 | 1 | 1 | | 275967 | Acetyl-CoA carboxylase beta OS=Rattus norvegicus OX=10116 GN=Acacb PE=1 SV=3 |
| 181 | 460 | tr\|E9PSQ0\|E9PSQ0_RAT | 21.77 | 0 | 0 | 103530.00 | 1 | 1 | 1 | | 276255 | Acetyl-CoA carboxylase beta OS=Rattus norvegicus OX=10116 GN=Acacb PE=1 SV=2 |
| 181 | 545 | P11497\|ACACA_RAT | 21.77 | 0 | 0 | 103530.00 | 1 | 1 | 1 | | 265191 | Acetyl-CoA carboxylase 1 OS=Rattus norvegicus OX=10116 GN=Acaca PE=1 SV=1 |
| 145 | 498 | Q99NA5\|IDH3A_RAT | 20.6 | 4 | 4 | 0.00 | 1 | 1 | 1 | | 39614 | Isocitrate dehydrogenase [NAD] subunit alpha mitochondrial OS=Rattus norvegicus OX=10116 GN=Idh3a PE=1 SV=1 |
| 145 | 499 | tr\|F1LNF7\|F1LNF7_RAT | 20.6 | 4 | 4 | 0.00 | 1 | 1 | 1 | | 41177 | Isocitrate dehydrogenase [NAD] subunit mitochondrial OS=Rattus norvegicus OX=10116 GN=Idh3a PE=1 SV=2 |
| 198 | 502 | Q5BJT0\|ARGL1_RAT | 20.49 | 4 | 4 | 1221100.00 | 1 | 1 | 1 | Formylatio | 32887 | Arginine and glutamate-rich protein 1 OS=Rattus norvegicus OX=10116 GN=Arglu1 PE=1 SV=1 |
| 218 | 465 | P07633\|PCCB_RAT | 20.48 | 2 | 2 | 0.00 | 1 | 1 | 1 | Formylatio | 58626 | Propionyl-CoA carboxylase beta chain mitochondrial OS=Rattus norvegicus OX=10116 GN=Pccb PE=2 SV=1 |
| 218 | 466 | tr\|Q68FZ8\|Q68FZ8_RAT | 20.48 | 2 | 2 | 0.00 | 1 | 1 | 1 | Formylatio | 58678 | Propionyl coenzyme A carboxylase beta polypeptide OS=Rattus norvegicus OX=10116 GN=Pccb PE=1 SV=1 |
| 190 | 424 | P21575\|DYN1_RAT | 20.08 | 1 | 1 | 174940.00 | 1 | 1 | 1 | Formylatio | 97295 | Dynamin-1 OS=Rattus norvegicus OX=10116 GN=Dnm1 PE=1 SV=2 |
