## Supplementary material for "Permeability transition pore-related changes in the proteome and channel activity of ATP synthase dimers and monomers": RLM SM-BM

| Protein Gr | Protein ID | Accession | -10lgP | Coverage (' | Coverage (' | IBAQ | #Peptides | #Unique | #Spec Sam | PTM | Avg. Mass | Description |
|---|---|---|---|---|---|---|---|---|---|---|---|---|
| 1 | 1 | Q02253\|MMSA_RAT | 368.86 | 56 | 56 | 94026000 | 98 | 98 | 166 | Carbamidomethylatio | | Methylmalonate-semialdehyde dehydrogenase [acylating], mitochondrial |
| 1 | 2 | tr\|G3V7J0\|G3V7J0_RAT | 368.86 | 56 | 56 | 9.40E+07 | 98 | 98 | 166 | Carbamidomethylatio | | Aldehyde dehydrogenase family 6, subfamily A1, isoform CRA_b |
| 4 | 17 | P10719\|ATPB_RAT | 343.59 | 52 | 52 | 7.56E+07 | 70 | 70 | 103 | Oxidation ( | 56354 | ATP synthase subunit beta mitochondrial OS=Rattus norvegicus OX=10116 GN=Atp5f1b PE=1 SV=2 |
| 4 | 18 | tr\|G3V6D3\|G3V6D3_RAT | 343.59 | 52 | 52 | 7.56E+07 | 70 | 70 | 103 | Oxidation ( | 56345 | ATP synthase subunit beta OS=Rattus norvegicus OX=10116 GN=Atp5f1b PE=1 SV=1 |
| 7 | 9 | Q5BK63\|NDUA9_RAT | 336.82 | 50 | 50 | 3.99E+07 | 60 | 58 | 99 | Oxidation (M) | | NADH dehydrogenase [ubiquinone] 1 alpha subcomplex subunit 9, mitochondrial |
| 3 | 10 | Q66HF1\|NDUS1_RAT | 310.93 | 40 | 40 | 8.61E+07 | 88 | 87 | 139 | Oxidation ( | 79412 | NADH-ubiquinone oxidoreductase 75 kDa subunit mitochondrial OS=Rattus norvegicus OX=10116 GN=Ndufs1 PE=1 SV=1 |
| 2 | 13 | P32551\|QCR2_RAT | 308.41 | 43 | 43 | 9.60E+07 | 60 | 60 | 157 | Oxidation ( | 48396 | Cytochrome b-c1 complex subunit 2 mitochondrial OS=Rattus norvegicus OX=10116 GN=Uqcrc2 PE=1 SV=2 |
| 6 | 30 | P19234\|NDUV2_RAT | 293.78 | 48 | 48 | 8.33E+07 | 51 | 51 | 101 | Formylatio | 27378 | NADH dehydrogenase [ubiquinone] flavoprotein 2 mitochondrial OS=Rattus norvegicus OX=10116 GN=Ndufv2 PE=1 SV=2 |
| 5 | 20 | P15999\|ATPA_RAT | 291.03 | 46 | 46 | 6.28E+07 | 63 | 62 | 101 | Oxidation ( | 59754 | ATP synthase subunit alpha mitochondrial OS=Rattus norvegicus OX=10116 GN=Atp5f1a PE=1 SV=2 |
| 5 | 21 | tr\|F1LP05\|F1LP05_RAT | 291.03 | 46 | 46 | 6.28E+07 | 63 | 62 | 101 | Oxidation ( | 59813 | ATP synthase subunit alpha OS=Rattus norvegicus OX=10116 GN=Atp5f1a PE=1 SV=1 |
| 13 | 22 | tr\|D3ZFQ8\|D3ZFQ8_RAT | 290.52 | 44 | 44 | 6.14E+07 | 34 | 33 | 61 | Oxidation ( | 35435 | Cytochrome c-1 OS=Rattus norvegicus OX=10116 GN=Cyc1 PE=1 SV=3 |
| 20 | 45 | tr\|B2RYS8\|B2RYS8_RAT | 285.81 | 41 | 41 | 2.29E+07 | 26 | 26 | 40 | Oxidation ( | 21959 | NADH dehydrogenase [ubiquinone] 1 beta subcomplex subunit 8 mitochondrial OS=Rattus norvegicus OX=10116 GN=Ndufb8 PE=1 SV=1 |
| 12 | 8 | P07756\|CPSM_RAT | 283.66 | 25 | 25 | 3.18E+07 | 56 | 56 | 73 | Oxidation ( | 164579 | Carbamoyl-phosphate synthase [ammonia] mitochondrial OS=Rattus norvegicus OX=10116 GN=Cps1 PE=1 SV=1 |
| 8 | 12 | Q68FY0\|QCR1_RAT | 283.38 | 44 | 44 | 9.76E+07 | 60 | 59 | 97 | Carbamidomethylatio | | Cytochrome b-c1 complex subunit 1, mitochondrial |
| 10 | 26 | Q641Y2\|NDUS2_RAT | 280.71 | 53 | 53 | 5.11E+07 | 52 | 51 | 93 | Oxidation ( | 52562 | NADH dehydrogenase [ubiquinone] iron-sulfur protein 2 mitochondrial OS=Rattus norvegicus OX=10116 GN=Ndufs2 PE=1 SV=1 |
| 16 | 164 | tr\|D3ZF13\|D3ZF13_RAT | 254.65 | 36 | 36 | 2.88E+07 | 28 | 28 | 48 | Oxidation ( | 17514 | Acyl carrier protein OS=Rattus norvegicus OX=10116 GN=Ndufab1 PE=1 SV=1 |
| 19 | 80 | P20788\|UCRI_RAT | 254.3 | 42 | 42 | 2.98E+07 | 25 | 24 | 41 | Oxidation ( | 29446 | Cytochrome b-c1 complex subunit Rieske mitochondrial OS=Rattus norvegicus OX=10116 GN=Uqcrfs1 PE=1 SV=2 |
| 14 | 11 | P67779\|PHB_RAT | 254.11 | 60 | 60 | 2.72E+07 | 39 | 39 | 56 | | 29820 | Prohibitin OS=Rattus norvegicus OX=10116 GN=Phb PE=1 SV=1 |
| 11 | 24 | tr\|D3ZG43\|D3ZG43_RAT | 245.56 | 52 | 52 | 4.48E+07 | 44 | 43 | 77 | | 30226 | NADH dehydrogenase (Ubiquinone) Fe-S protein 3 (Predicted) isoform CRA_c OS=Rattus norvegicus OX=10116 GN=Ndufs3 PE=1 SV=1 |
| 36 | 111 | Q5XIF3\|NDUS4_RAT | 240.38 | 21 | 21 | 1.15E+07 | 16 | 16 | 20 | | 19741 | NADH dehydrogenase [ubiquinone] iron-sulfur protein 4 mitochondrial OS=Rattus norvegicus OX=10116 GN=Ndufs4 PE=1 SV=1 |
| 17 | 28 | Q561S0\|NDUAA_RAT | 238.77 | 38 | 38 | 2.45E+07 | 27 | 27 | 46 | Oxidation ( | 40493 | NADH dehydrogenase [ubiquinone] 1 alpha subcomplex subunit 10 mitochondrial OS=Rattus norvegicus OX=10116 GN=Ndufa10 PE=1 SV=1 |
| 18 | 46 | tr\|L0L4L8\|L0L4L8_RAT | 236.92 | 29 | 29 | 6.46E+06 | 27 | 3 | 44 | Oxidation (M) | | |
| 18 | 47 | tr\|A0A140GE10\|A0A140GE10_RAT | 236.92 | 29 | 29 | 6.46E+06 | 27 | 3 | 44 | Oxidation (M) | | |
| 18 | 48 | tr\|A0A140GE11\|A0A140GE11_RAT | 236.92 | 29 | 29 | 6.46E+06 | 27 | 3 | 44 | Oxidation (M) | | |
| 18 | 49 | tr\|A0A3Q8AGE8\|A0A3Q8AGE8_RA | 236.92 | 29 | 29 | 6.46E+06 | 27 | 3 | 44 | Oxidation (M) | | |
| 18 | 50 | tr\|A0A3Q8AC68\|A0A3Q8AC68_RA | 236.92 | 29 | 29 | 6.46E+06 | 27 | 3 | 44 | Oxidation (M) | | |
| 18 | 51 | tr\|A0A3S6FM15\|A0A3S6FM15_RA | 236.92 | 29 | 29 | 6.46E+06 | 27 | 3 | 44 | Oxidation (M) | | |
| 18 | 52 | tr\|A0A220D9Z6\|A0A220D9Z6_RAT | 236.92 | 28 | 28 | 6.46E+06 | 27 | 3 | 44 | Oxidation (M) | | |
| 18 | 99 | tr\|D6NSP2\|D6NSP2_RAT | 236.92 | 28 | 28 | 6.46E+06 | 27 | 3 | 44 | Oxidation (M) | | |
| 18 | 53 | tr\|D6NSS6\|D6NSS6_RAT | 236.92 | 28 | 28 | 6.46E+06 | 27 | 3 | 44 | Oxidation (M) | | |
| 18 | 54 | tr\|D6NSR3\|D6NSR3_RAT | 236.92 | 28 | 28 | 6.46E+06 | 27 | 3 | 44 | Oxidation (M) | | |
| 18 | 55 | tr\|D6NSR8\|D6NSR8_RAT | 236.92 | 28 | 28 | 6.46E+06 | 27 | 3 | 44 | Oxidation (M) | | |
| 18 | 56 | tr\|A0A0A1FZ42\|A0A0A1FZ42_RAT | 236.92 | 28 | 28 | 6.46E+06 | 27 | 3 | 44 | Oxidation (M) | | |
| 18 | 57 | tr\|D6NSS7\|D6NSS7_RAT | 236.92 | 28 | 28 | 6.46E+06 | 27 | 3 | 44 | Oxidation (M) | | |
| 18 | 58 | tr\|Q8SEY9\|Q8SEY9_RAT | 236.92 | 28 | 28 | 6.46E+06 | 27 | 3 | 44 | Oxidation (M) | | |
| 18 | 44 | tr\|D6NSR7\|D6NSR7_RAT | 236.92 | 28 | 28 | 6.46E+06 | 27 | 3 | 44 | Oxidation (M) | | |
| 18 | 59 | tr\|A0A220DA44\|A0A220DA44_RAT | 236.92 | 28 | 28 | 6.46E+06 | 27 | 3 | 44 | Oxidation (M) | | |
| 18 | 60 | tr\|D6NSQ0\|D6NSQ0_RAT | 236.92 | 28 | 28 | 6.46E+06 | 27 | 3 | 44 | Oxidation (M) | | |
| 18 | 61 | tr\|Q5UAI7\|Q5UAI7_RAT | 236.92 | 28 | 28 | 6.46E+06 | 27 | 3 | 44 | Oxidation (M) | | Cytochrome b |
| 18 | 62 | tr\|Q8HIC4\|Q8HIC4_RAT | 236.92 | 28 | 28 | 6.46E+06 | 27 | 3 | 44 | Oxidation (M) | | Cytochrome b |
| 18 | 63 | tr\|A0A220DA02\|A0A220DA02_RAT | 236.92 | 28 | 28 | 6.46E+06 | 27 | 3 | 44 | Oxidation (M) | | |
| 18 | 64 | tr\|A0A0S1Z1V4\|A0A0S1Z1V4_RAT | 236.92 | 28 | 28 | 6.46E+06 | 27 | 3 | 44 | Oxidation (M) | | |
| 18 | 65 | P00159\|CYB_RAT | 236.92 | 28 | 28 | 6.46E+06 | 27 | 3 | 44 | Oxidation (M) | | Cytochrome b |
| 18 | 66 | tr\|L0N311\|L0N311_RAT | 236.92 | 28 | 28 | 6.46E+06 | 27 | 3 | 44 | Oxidation (M) | | |
| 18 | 67 | tr\|A0A220DA28\|A0A220DA28_RAT | 236.92 | 28 | 28 | 6.46E+06 | 27 | 3 | 44 | Oxidation (M) | | |
| 18 | 68 | tr\|H2KXA0\|H2KXA0_RAT | 236.92 | 28 | 28 | 6.46E+06 | 27 | 3 | 44 | Oxidation (M) | | |
| 27 | 165 | tr\|D4A565\|D4A565_RAT | 236.7 | 43 | 43 | 1.84E+07 | 21 | 21 | 31 | | 21664 | NADH dehydrogenase (Ubiquinone) 1 beta subcomplex 5 (Predicted) isoform CRA_b OS=Rattus norvegicus OX=10116 GN=Ndufb5 PE=1 SV=1 |
| 31 | 135 | tr\|B0BNE6\|B0BNE6_RAT | 236.02 | 26 | 26 | 1.36E+07 | 20 | 18 | 27 | | 23970 | NADH dehydrogenase (Ubiquinone) Fe-S protein 8 (Predicted) isoform CRA_a OS=Rattus norvegicus OX=10116 GN=Ndufs8 PE=1 SV=1 |
| 21 | 19 | Q5XIH7\|PHB2_RAT | 234.49 | 46 | 46 | 1.63E+07 | 27 | 27 | 38 | Oxidation ( | 33312 | Prohibitin-2 OS=Rattus norvegicus OX=10116 GN=Phb2 PE=1 SV=1 |
| 29 | 14 | P10860\|DHE3_RAT | 233.57 | 27 | 27 | 8.90E+06 | 26 | 26 | 29 | Oxidation (M) | | Glutamate dehydrogenase 1, mitochondrial |
| 24 | 35 | P52873\|PYC_RAT | 229.8 | 15 | 15 | 9.05E+06 | 27 | 27 | 34 | Oxidation ( | 129777 | Pyruvate carboxylase mitochondrial OS=Rattus norvegicus OX=10116 GN=Pc PE=1 SV=2 |
| 24 | 34 | tr\|A0A0G2JTL5\|A0A0G2JTL5_RAT | 229.8 | 14 | 14 | 9.05E+06 | 27 | 27 | 34 | Oxidation ( | 140005 | Pyruvate carboxylase mitochondrial OS=Rattus norvegicus OX=10116 GN=Pc PE=1 SV=1 |
| 23 | 42 | tr\|D4A0T0\|D4A0T0_RAT | 229.5 | 45 | 45 | 2.41E+07 | 20 | 20 | 36 | | 20859 | NADH:ubiquinone oxidoreductase subunit B10 OS=Rattus norvegicus OX=10116 GN=Ndufb10 PE=1 SV=1 |
| 32 | 242 | Q80W89\|NDUAB_RAT | 226.94 | 28 | 28 | 1.23E+07 | 16 | 16 | 26 | | 14854 | NADH dehydrogenase [ubiquinone] 1 alpha subcomplex subunit 11 OS=Rattus norvegicus OX=10116 GN=Ndufa11 PE=2 SV=1 |
| 25 | 15 | P00507\|AATM_RAT | 226.22 | 30 | 30 | 1.45E+07 | 23 | 23 | 33 | Oxidation ( | 47314 | Aspartate aminotransferase mitochondrial OS=Rattus norvegicus OX=10116 GN=Got2 PE=1 SV=2 |
| 39 | 29 | P07895\|SODM_RAT | 217.75 | 36 | 36 | 1.27E+07 | 11 | 11 | 19 | | 24674 | Superoxide dismutase [Mn] mitochondrial OS=Rattus norvegicus OX=10116 GN=Sod2 PE=1 SV=2 |
| 34 | 246 | tr\|Q5PQZ9\|Q5PQZ9_RAT | 215.8 | 37 | 37 | 2.91E+07 | 13 | 13 | 25 | | 14359 | NADH dehydrogenase [ubiquinone] 1 subunit C2 OS=Rattus norvegicus OX=10116 GN=Ndufc2 PE=1 SV=1 |
| 35 | 183 | tr\|D4A7L4\|D4A7L4_RAT | 208.45 | 42 | 42 | 2.46E+07 | 14 | 14 | 23 | | 17634 | NADH dehydrogenase (Ubiquinone) 1 beta subcomplex 11 (Predicted) OS=Rattus norvegicus OX=10116 GN=Ndufb11 PE=1 SV=1 |
| 15 | 23 | tr\|Q5XIH3\|Q5XIH3_RAT | 206.98 | 27 | 27 | 4.29E+07 | 25 | 25 | 49 | Oxidation ( | 50731 | NADH dehydrogenase [ubiquinone] flavoprotein 1 mitochondrial OS=Rattus norvegicus OX=10116 GN=Ndufv1 PE=1 SV=1 |
| 22 | 74 | tr\|Q9G898\|Q9G898_RAT | 202.43 | 28 | 28 | 0 | 25 | 1 | 38 | Oxidation (M) | | Cytochrome b |
| 38 | 168 | tr\|F1LXA0\|F1LXA0_RAT | 201.7 | 37 | 37 | 1.77E+07 | 14 | 14 | 19 | Oxidation ( | 17178 | NADH dehydrogenase [ubiquinone] 1 alpha subcomplex subunit 12 OS=Rattus norvegicus OX=10116 GN=Ndufa12 PE=1 SV=2 |
| 37 | 112 | tr\|D4A3V2\|D4A3V2_RAT | 191.54 | 42 | 42 | 5.75E+06 | 17 | 17 | 19 | Oxidation ( | 15224 | NADH dehydrogenase [ubiquinone] 1 alpha subcomplex subunit 6 OS=Rattus norvegicus OX=10116 GN=Ndufa6 PE=1 SV=1 |
| 28 | 3 | tr\|B2GV15\|B2GV15_RAT | 189.62 | 25 | 25 | 1.55E+07 | 19 | 19 | 30 | Oxidation ( | 53274 | Dihydrolipoamide acetyltransferase component of pyruvate dehydrogenase complex OS=Rattus norvegicus OX=10116 GN=Dbt PE=1 SV=1 |
| 30 | 110 | tr\|A0A0G2JVL6\|A0A0G2JVL6_RAT | 187.14 | 37 | 37 | 1.52E+07 | 15 | 15 | 28 | | 19965 | NADH dehydrogenase [ubiquinone] 1 alpha subcomplex subunit 8 OS=Rattus norvegicus OX=10116 GN=Ndufa8 PE=1 SV=1 |
| 33 | 36 | tr\|Q06QK5\|Q06QK5_RAT | 180.32 | 17 | 17 | 1.20E+07 | 17 | 17 | 25 | Oxidation ( | 68606 | NADH-ubiquinone oxidoreductase chain 5 OS=Rattus norvegicus OX=10116 GN=ND5 PE=3 SV=1 |
| 33 | 37 | P11661\|NU5M_RAT | 180.32 | 17 | 17 | 1.20E+07 | 17 | 17 | 25 | Oxidation ( | 68618 | NADH-ubiquinone oxidoreductase chain 5 OS=Rattus norvegicus OX=10116 GN=Mtnd5 PE=3 SV=3 |
| 33 | 43 | tr\|Q06QG6\|Q06QG6_RAT | 180.32 | 17 | 17 | 1.20E+07 | 17 | 17 | 25 | Oxidation ( | 68588 | NADH-ubiquinone oxidoreductase chain 5 OS=Rattus norvegicus OX=10116 GN=ND5 PE=3 SV=1 |

<table>
<tr><td>33</td><td>38</td><td>tr|Q06QA1|Q06QA1_RAT</td><td>180.32</td><td>17</td><td>17</td><td>1.20E+07</td><td>17</td><td>17</td><td>25</td><td>Oxidation (</td><td>68574</td><td>NADH-ubiquinone oxidoreductase chain 5 OS=Rattus norvegicus OX=10116 GN=ND5 PE=3 SV=1</td></tr>
<tr><td>33</td><td>39</td><td>tr|Q8SEZ0|Q8SEZ0_RAT</td><td>180.32</td><td>17</td><td>17</td><td>1.20E+07</td><td>17</td><td>17</td><td>25</td><td>Oxidation (</td><td>68618</td><td>NADH-ubiquinone oxidoreductase chain 5 OS=Rattus norvegicus OX=10116 GN=Mt-nd5 PE=3 SV=1</td></tr>
<tr><td>33</td><td>40</td><td>tr|A0A096XKT9|A0A096XKT9_RAT</td><td>180.32</td><td>17</td><td>17</td><td>1.20E+07</td><td>17</td><td>17</td><td>25</td><td>Oxidation (</td><td>68584</td><td>NADH-ubiquinone oxidoreductase chain 5 OS=Rattus norvegicus OX=10116 GN=ND5 PE=3 SV=1</td></tr>
<tr><td>53</td><td>280</td><td>P31399|ATP5H_RAT</td><td>178.38</td><td>32</td><td>32</td><td>4.71E+06</td><td>8</td><td>8</td><td>11</td><td></td><td>18763</td><td>ATP synthase subunit d  mitochondrial OS=Rattus norvegicus OX=10116 GN=Atp5pd PE=1 SV=3</td></tr>
<tr><td>44</td><td>301</td><td>P35435|ATPG_RAT</td><td>175.9</td><td>24</td><td>24</td><td>5.03E+06</td><td>12</td><td>12</td><td>14</td><td></td><td>30191</td><td>ATP synthase subunit gamma  mitochondrial OS=Rattus norvegicus OX=10116 GN=Atp5f1c PE=1 SV=2</td></tr>
<tr><td>44</td><td>297</td><td>tr|Q6QI09|Q6QI09_RAT</td><td>175.9</td><td>11</td><td>11</td><td>5.03E+06</td><td>12</td><td>12</td><td>14</td><td></td><td>67721</td><td>ATP synthase subunit gamma  mitochondrial OS=Rattus norvegicus OX=10116 GN=Taf3 PE=1 SV=1</td></tr>
<tr><td>40</td><td>175</td><td>tr|D3ZCZ9|D3ZCZ9_RAT</td><td>171.89</td><td>48</td><td>48</td><td>7.12E+06</td><td>11</td><td>11</td><td>18</td><td></td><td>13040</td><td>NADH dehydrogenase [ubiquinone] iron-sulfur protein 6  mitochondrial OS=Rattus norvegicus OX=10116 GN=LOC100912599 PE=1 SV=1</td></tr>
<tr><td>55</td><td>281</td><td>tr|A9UMW2|A9UMW2_RAT</td><td>168.75</td><td>45</td><td>45</td><td>5.57E+06</td><td>8</td><td>8</td><td>11</td><td>Oxidation (</td><td>9255</td><td>Ndufa3 protein (Fragment) OS=Rattus norvegicus OX=10116 GN=Ndufa3 PE=2 SV=1</td></tr>
<tr><td>55</td><td>272</td><td>tr|M0RB63|M0RB63_RAT</td><td>168.75</td><td>44</td><td>44</td><td>5.57E+06</td><td>8</td><td>8</td><td>11</td><td>Oxidation (</td><td>9372</td><td>RCG63041 OS=Rattus norvegicus OX=10116 GN=LOC684509 PE=4 SV=1</td></tr>
<tr><td>55</td><td>282</td><td>tr|A0A0G2KAA3|A0A0G2KAA3_RA</td><td>168.75</td><td>41</td><td>41</td><td>5.57E+06</td><td>8</td><td>8</td><td>11</td><td>Oxidation (</td><td>10170</td><td>NADH:ubiquinone oxidoreductase subunit A3 OS=Rattus norvegicus OX=10116 GN=Ndufa3 PE=1 SV=1</td></tr>
<tr><td>41</td><td>119</td><td>tr|D3ZE15|D3ZE15_RAT</td><td>166.26</td><td>49</td><td>49</td><td>4.67E+06</td><td>14</td><td>14</td><td>17</td><td>Oxidation (</td><td>16777</td><td>NADH:ubiquinone oxidoreductase subunit A13 OS=Rattus norvegicus OX=10116 GN=Ndufa13 PE=1 SV=1</td></tr>
<tr><td>48</td><td>133</td><td>tr|F1LPG5|F1LPG5_RAT</td><td>164.49</td><td>35</td><td>35</td><td>7.63E+06</td><td>8</td><td>8</td><td>13</td><td></td><td>15064</td><td>NADH:ubiquinone oxidoreductase subunit B4 OS=Rattus norvegicus OX=10116 GN=Ndufb4 PE=1 SV=1</td></tr>
<tr><td>45</td><td>196</td><td>Q63362|NDUA5_RAT</td><td>158.49</td><td>42</td><td>42</td><td>6.46E+06</td><td>11</td><td>10</td><td>14</td><td></td><td>13412</td><td>NADH dehydrogenase [ubiquinone] 1 alpha subcomplex subunit 5 OS=Rattus norvegicus OX=10116 GN=Ndufa5 PE=1 SV=3</td></tr>
<tr><td>26</td><td>138</td><td>P05508|NU4M_RAT</td><td>158.36</td><td>17</td><td>17</td><td>7.30E+06</td><td>17</td><td>16</td><td>32</td><td>Oxidation (M)</td><td></td><td>NADH-ubiquinone oxidoreductase chain 4</td></tr>
<tr><td>26</td><td>139</td><td>tr|D2E6K0|D2E6K0_RAT</td><td>158.36</td><td>17</td><td>17</td><td>7.30E+06</td><td>17</td><td>16</td><td>32</td><td>Oxidation (M)</td><td></td><td></td></tr>
<tr><td>26</td><td>151</td><td>tr|A7XYB9|A7XYB9_RAT</td><td>158.36</td><td>17</td><td>17</td><td>7.30E+06</td><td>17</td><td>16</td><td>32</td><td>Oxidation (M)</td><td></td><td></td></tr>
<tr><td>26</td><td>140</td><td>tr|Q06QE1|Q06QE1_RAT</td><td>158.36</td><td>17</td><td>17</td><td>7.30E+06</td><td>17</td><td>16</td><td>32</td><td>Oxidation (M)</td><td></td><td></td></tr>
<tr><td>26</td><td>141</td><td>tr|Q06QA2|Q06QA2_RAT</td><td>158.36</td><td>17</td><td>17</td><td>7.30E+06</td><td>17</td><td>16</td><td>32</td><td>Oxidation (M)</td><td></td><td></td></tr>
<tr><td>26</td><td>142</td><td>tr|Q7HKW3|Q7HKW3_RAT</td><td>158.36</td><td>17</td><td>17</td><td>7.30E+06</td><td>17</td><td>16</td><td>32</td><td>Oxidation (M)</td><td></td><td></td></tr>
<tr><td>26</td><td>152</td><td>tr|Q06QG7|Q06QG7_RAT</td><td>158.36</td><td>17</td><td>17</td><td>7.30E+06</td><td>17</td><td>16</td><td>32</td><td>Oxidation (M)</td><td></td><td></td></tr>
<tr><td>26</td><td>143</td><td>tr|Q06Q89|Q06Q89_RAT</td><td>158.36</td><td>17</td><td>17</td><td>7.30E+06</td><td>17</td><td>16</td><td>32</td><td>Oxidation (M)</td><td></td><td></td></tr>
<tr><td>26</td><td>144</td><td>tr|Q35737|Q35737_RAT</td><td>158.36</td><td>17</td><td>17</td><td>7.30E+06</td><td>17</td><td>16</td><td>32</td><td>Oxidation (M)</td><td></td><td></td></tr>
<tr><td>26</td><td>145</td><td>tr|Q8HIC6|Q8HIC6_RAT</td><td>158.36</td><td>17</td><td>17</td><td>7.30E+06</td><td>17</td><td>16</td><td>32</td><td>Oxidation (M)</td><td></td><td></td></tr>
<tr><td>43</td><td>178</td><td>tr|D3ZZ21|D3ZZ21_RAT</td><td>155.36</td><td>49</td><td>49</td><td>8.28E+06</td><td>10</td><td>10</td><td>15</td><td>Oxidation (</td><td>15638</td><td>NADH dehydrogenase (Ubiquinone) 1 beta subcomplex  6 (Predicted) OS=Rattus norvegicus OX=10116 GN=Ndufb6 PE=1 SV=1</td></tr>
<tr><td>60</td><td>33</td><td>Q64428|ECHA_RAT</td><td>143.98</td><td>10</td><td>10</td><td>2.17E+06</td><td>8</td><td>8</td><td>8</td><td></td><td>82665</td><td>Trifunctional enzyme subunit alpha  mitochondrial OS=Rattus norvegicus OX=10116 GN=Hadha PE=1 SV=2</td></tr>
<tr><td>69</td><td>289</td><td>tr|D3ZFJ6|D3ZFJ6_RAT</td><td>139.35</td><td>9</td><td>9</td><td>1.67E+06</td><td>5</td><td>5</td><td>5</td><td></td><td>60420</td><td>Lactamase  beta OS=Rattus norvegicus OX=10116 GN=Lactb PE=1 SV=1</td></tr>
<tr><td>59</td><td>76</td><td>tr|Q68G44|Q68G44_RAT</td><td>138.29</td><td>13</td><td>13</td><td>2.75E+06</td><td>8</td><td>8</td><td>8</td><td></td><td>56886</td><td>3-hydroxy-3-methylglutaryl coenzyme A synthase OS=Rattus norvegicus OX=10116 GN=Hmgcs2 PE=1 SV=1</td></tr>
<tr><td>46</td><td>268</td><td>P19511|AT5F1_RAT</td><td>135.45</td><td>24</td><td>24</td><td>8.80E+06</td><td>10</td><td>10</td><td>13</td><td>Carbamidomethylatio</td><td></td><td>ATP synthase F(0) complex subunit B1, mitochondrial</td></tr>
<tr><td>51</td><td>192</td><td>tr|B5DEL8|B5DEL8_RAT</td><td>132.98</td><td>39</td><td>39</td><td>3.65E+06</td><td>8</td><td>8</td><td>11</td><td></td><td>12700</td><td>NADH dehydrogenase (Ubiquinone) Fe-S protein 5 OS=Rattus norvegicus OX=10116 GN=Ndufs5 PE=1 SV=1</td></tr>
<tr><td>51</td><td>193</td><td>tr|A0A0G2JZJ9|A0A0G2JZJ9_RAT</td><td>132.98</td><td>39</td><td>39</td><td>3.65E+06</td><td>8</td><td>8</td><td>11</td><td></td><td>12730</td><td>Uncharacterized protein OS=Rattus norvegicus OX=10116 PE=4 SV=1</td></tr>
<tr><td>57</td><td>176</td><td>P13086|SUCA_RAT</td><td>132.72</td><td>13</td><td>13</td><td>2.39E+06</td><td>7</td><td>7</td><td>9</td><td></td><td>36148</td><td>Succinate--CoA ligase [ADP/GDP-forming] subunit alpha  mitochondrial OS=Rattus norvegicus OX=10116 GN=Suclg1 PE=2 SV=2</td></tr>
<tr><td>57</td><td>177</td><td>tr|A0A0H2UHE1|A0A0H2UHE1_RA</td><td>132.72</td><td>13</td><td>13</td><td>2.39E+06</td><td>7</td><td>7</td><td>9</td><td></td><td>37560</td><td>Succinate--CoA ligase [ADP/GDP-forming] subunit alpha  mitochondrial OS=Rattus norvegicus OX=10116 GN=Suclg1 PE=1 SV=1</td></tr>
<tr><td>54</td><td>243</td><td>tr|D3ZS58|D3ZS58_RAT</td><td>127.97</td><td>33</td><td>33</td><td>4.11E+06</td><td>5</td><td>5</td><td>11</td><td>Oxidation (</td><td>10845</td><td>NADH dehydrogenase [ubiquinone] 1 alpha subcomplex subunit 2 OS=Rattus norvegicus OX=10116 GN=Ndufa2 PE=1 SV=1</td></tr>
<tr><td>50</td><td>148</td><td>tr|Q5RJN0|Q5RJN0_RAT</td><td>127.63</td><td>19</td><td>19</td><td>7.11E+06</td><td>8</td><td>8</td><td>12</td><td>Oxidation (</td><td>23945</td><td>NADH dehydrogenase (Ubiquinone) Fe-S protein 7 OS=Rattus norvegicus OX=10116 GN=Ndufs7 PE=1 SV=1</td></tr>
<tr><td>58</td><td>286</td><td>tr|A9UMV9|A9UMV9_RAT</td><td>126.35</td><td>21</td><td>21</td><td>9.45E+06</td><td>4</td><td>4</td><td>9</td><td></td><td>12500</td><td>NADH:ubiquinone oxidoreductase subunit A7 OS=Rattus norvegicus OX=10116 GN=Ndufa7 PE=1 SV=1</td></tr>
<tr><td>56</td><td>100</td><td>P17764|THIL_RAT</td><td>123.72</td><td>14</td><td>14</td><td>3.20E+06</td><td>8</td><td>8</td><td>9</td><td></td><td>44695</td><td>Acetyl-CoA acetyltransferase  mitochondrial OS=Rattus norvegicus OX=10116 GN=Acat1 PE=1 SV=1</td></tr>
<tr><td>42</td><td>171</td><td>tr|Q8SEZ8|Q8SEZ8_RAT</td><td>121.52</td><td>20</td><td>20</td><td>7.16E+06</td><td>11</td><td>11</td><td>15</td><td>Oxidation (</td><td>36133</td><td>NADH-ubiquinone oxidoreductase chain 1 (Fragment) OS=Rattus norvegicus OX=10116 GN=NADH1 PE=3 SV=1</td></tr>
<tr><td>42</td><td>172</td><td>P03889|NU1M_RAT</td><td>121.52</td><td>20</td><td>20</td><td>7.16E+06</td><td>11</td><td>11</td><td>15</td><td>Oxidation (</td><td>36145</td><td>NADH-ubiquinone oxidoreductase chain 1 OS=Rattus norvegicus OX=10116 GN=Mtnd1 PE=1 SV=3</td></tr>
<tr><td>42</td><td>173</td><td>tr|Q8HID1|Q8HID1_RAT</td><td>121.52</td><td>20</td><td>20</td><td>7.16E+06</td><td>11</td><td>11</td><td>15</td><td>Oxidation (</td><td>36145</td><td>NADH-ubiquinone oxidoreductase chain 1 OS=Rattus norvegicus OX=10116 GN=Mt-nd1 PE=3 SV=1</td></tr>
<tr><td>42</td><td>174</td><td>tr|D2E6L7|D2E6L7_RAT</td><td>121.52</td><td>20</td><td>20</td><td>7.16E+06</td><td>11</td><td>11</td><td>15</td><td>Oxidation (</td><td>36062</td><td>NADH-ubiquinone oxidoreductase chain 1 OS=Rattus norvegicus OX=10116 GN=ND1 PE=3 SV=1</td></tr>
<tr><td>66</td><td>231</td><td>tr|Q5M949|Q5M949_RAT</td><td>120.72</td><td>17</td><td>17</td><td>1.29E+06</td><td>6</td><td>6</td><td>7</td><td></td><td>28340</td><td>Nipsnap homolog 3A (C. elegans) OS=Rattus norvegicus OX=10116 GN=Nipsnap3b PE=1 SV=1</td></tr>
<tr><td>61</td><td>248</td><td>tr|B2RYW3|B2RYW3_RAT</td><td>110.97</td><td>23</td><td>23</td><td>5.44E+06</td><td>7</td><td>7</td><td>8</td><td>Oxidation (</td><td>21892</td><td>NADH dehydrogenase (Ubiquinone) 1 beta subcomplex  9 OS=Rattus norvegicus OX=10116 GN=Ndufb9 PE=1 SV=1</td></tr>
<tr><td>116</td><td>261</td><td>P24329|THTR_RAT</td><td>109.2</td><td>9</td><td>9</td><td>2.96E+05</td><td>2</td><td>2</td><td>2</td><td></td><td>33407</td><td>Thiosulfate sulfurtransferase OS=Rattus norvegicus OX=10116 GN=Tst PE=1 SV=3</td></tr>
<tr><td>47</td><td>251</td><td>tr|A0A0A1G491|A0A0A1G491_RA</td><td>107.04</td><td>21</td><td>21</td><td>2.05E+06</td><td>10</td><td>10</td><td>13</td><td></td><td>38542</td><td>NADH-ubiquinone oxidoreductase chain 2 OS=Rattus norvegicus OX=10116 GN=ND2 PE=3 SV=1</td></tr>
<tr><td>47</td><td>252</td><td>tr|Q5UAJ8|Q5UAJ8_RAT</td><td>107.04</td><td>21</td><td>21</td><td>2.05E+06</td><td>10</td><td>10</td><td>13</td><td></td><td>38455</td><td>NADH-ubiquinone oxidoreductase chain 2 OS=Rattus norvegicus OX=10116 GN=ND2 PE=3 SV=1</td></tr>
<tr><td>52</td><td>216</td><td>Q7TQ16|QCR8_RAT</td><td>106.31</td><td>43</td><td>43</td><td>4.23E+06</td><td>9</td><td>9</td><td>11</td><td></td><td>9849</td><td>Cytochrome b-c1 complex subunit 8 OS=Rattus norvegicus OX=10116 GN=Uqcrq PE=3 SV=1</td></tr>
<tr><td>62</td><td>291</td><td>tr|Q5XFW4|Q5XFW4_RAT</td><td>105.57</td><td>20</td><td>20</td><td>3.10E+06</td><td>5</td><td>5</td><td>8</td><td></td><td>20541</td><td>Mitochondrial ribosomal protein L13 OS=Rattus norvegicus OX=10116 GN=Mrpl13 PE=1 SV=1</td></tr>
<tr><td>64</td><td>169</td><td>Q60587|ECHB_RAT</td><td>103.51</td><td>14</td><td>14</td><td>1.93E+06</td><td>7</td><td>7</td><td>8</td><td></td><td>51414</td><td>Trifunctional enzyme subunit beta  mitochondrial OS=Rattus norvegicus OX=10116 GN=Hadhb PE=1 SV=1</td></tr>
<tr><td>72</td><td>350</td><td>tr|Q35733|Q35733_RAT</td><td>98.83</td><td>14</td><td>14</td><td>8.71E+05</td><td>4</td><td>4</td><td>5</td><td></td><td>8389</td><td>NADH-ubiquinone oxidoreductase chain 6 (Fragment) OS=Rattus norvegicus OX=10116 PE=3 SV=1</td></tr>
<tr><td>72</td><td>351</td><td>tr|Q06QD9|Q06QD9_RAT</td><td>98.83</td><td>6</td><td>6</td><td>8.71E+05</td><td>4</td><td>4</td><td>5</td><td></td><td>18943</td><td>NADH-ubiquinone oxidoreductase chain 6 OS=Rattus norvegicus OX=10116 GN=ND6 PE=3 SV=1</td></tr>
<tr><td>72</td><td>352</td><td>tr|Q7HKW2|Q7HKW2_RAT</td><td>98.83</td><td>6</td><td>6</td><td>8.71E+05</td><td>4</td><td>4</td><td>5</td><td></td><td>18957</td><td>NADH-ubiquinone oxidoreductase chain 6 OS=Rattus norvegicus OX=10116 GN=NADH6 PE=3 SV=1</td></tr>
<tr><td>82</td><td>179</td><td>P11240|COX5A_RAT</td><td>96.51</td><td>23</td><td>23</td><td>2.31E+06</td><td>3</td><td>3</td><td>4</td><td></td><td>16130</td><td>Cytochrome c oxidase subunit 5A  mitochondrial OS=Rattus norvegicus OX=10116 GN=Cox5a PE=1 SV=1</td></tr>
<tr><td>68</td><td>6</td><td>Q01205|ODO2_RAT</td><td>95.86</td><td>13</td><td>13</td><td>1.29E+06</td><td>5</td><td>5</td><td>5</td><td></td><td>48925</td><td>Dihydrolipoyllysine-residue succinyltransferase component of 2-oxoglutarate dehydrogenase complex  mitochondrial OS=Rattus norvegicus OX=10116 GN=Dlst PE=1 SV=2</td></tr>
<tr><td>68</td><td>5</td><td>tr|G3V6P2|G3V6P2_RAT</td><td>95.86</td><td>13</td><td>13</td><td>1.29E+06</td><td>5</td><td>5</td><td>5</td><td></td><td>48899</td><td>Dihydrolipoamide S-succinyltransferase (E2 component of 2-oxo-glutarate complex)  isoform CRA_a OS=Rattus norvegicus OX=10116 GN=Dlst PE=1 SV=1</td></tr>
<tr><td>63</td><td>266</td><td>P04762|CATA_RAT</td><td>94.02</td><td>6</td><td>6</td><td>9.55E+05</td><td>7</td><td>7</td><td>8</td><td></td><td>59757</td><td>Catalase OS=Rattus norvegicus OX=10116 GN=Cat PE=1 SV=3</td></tr>
<tr><td>109</td><td>273</td><td>tr|A0A0G2JVH4|A0A0G2JVH4_RAT</td><td>87.78</td><td>3</td><td>3</td><td>4.21E+05</td><td>2</td><td>2</td><td>2</td><td></td><td>86230</td><td>MICOS complex subunit MIC60 OS=Rattus norvegicus OX=10116 GN=Immt PE=1 SV=1</td></tr>
<tr><td>109</td><td>276</td><td>tr|A0A140TAG5|A0A140TAG5_RA</td><td>87.78</td><td>4</td><td>4</td><td>4.21E+05</td><td>2</td><td>2</td><td>2</td><td></td><td>67049</td><td>MICOS complex subunit MIC60 OS=Rattus norvegicus OX=10116 GN=Immt PE=1 SV=1</td></tr>
<tr><td>109</td><td>277</td><td>Q3KR86|MIC60_RAT</td><td>87.78</td><td>4</td><td>4</td><td>4.21E+05</td><td>2</td><td>2</td><td>2</td><td></td><td>67177</td><td>MICOS complex subunit Mic60 (Fragment) OS=Rattus norvegicus OX=10116 GN=Immt PE=1 SV=1</td></tr>
<tr><td>87</td><td>198</td><td>P85834|EFTU_RAT</td><td>83.9</td><td>5</td><td>5</td><td>3.66E+05</td><td>2</td><td>2</td><td>3</td><td></td><td>49522</td><td>Elongation factor Tu  mitochondrial OS=Rattus norvegicus OX=10116 GN=Tufm PE=1 SV=1</td></tr>
<tr><td>89</td><td>228</td><td>P16970|ABCD3_RAT</td><td>79.65</td><td>4</td><td>4</td><td>4.12E+06</td><td>3</td><td>3</td><td>3</td><td></td><td>75316</td><td>ATP-binding cassette sub-family D member 3 OS=Rattus norvegicus OX=10116 GN=Abcd3 PE=1 SV=3</td></tr>
<tr><td>70</td><td>244</td><td>P07824|ARGI1_RAT</td><td>79.62</td><td>13</td><td>13</td><td>1.17E+06</td><td>4</td><td>4</td><td>5</td><td></td><td>34973</td><td>Arginase-1 OS=Rattus norvegicus OX=10116 GN=Arg1 PE=1 SV=2</td></tr>
<tr><td>65</td><td>230</td><td>tr|B2RYS2|B2RYS2_RAT</td><td>78.1</td><td>28</td><td>28</td><td>5.63E+06</td><td>6</td><td>6</td><td>8</td><td></td><td>13559</td><td>Cytochrome b-c1 complex subunit 7 OS=Rattus norvegicus OX=10116 GN=Uqcrb PE=1 SV=1</td></tr>
<tr><td>74</td><td>16</td><td>tr|F1LN92|F1LN92_RAT</td><td>73.65</td><td>5</td><td>5</td><td>2.11E+06</td><td>4</td><td>4</td><td>4</td><td></td><td>89352</td><td>AFG3-like matrix AAA peptidase subunit 2 OS=Rattus norvegicus OX=10116 GN=Afg3l2 PE=1 SV=1</td></tr>
<tr><td>80</td><td>7</td><td>Q4FZT0|STML2_RAT</td><td>73.61</td><td>7</td><td>7</td><td>4.36E+05</td><td>3</td><td>3</td><td>4</td><td></td><td>38414</td><td>Stomatin-like protein 2  mitochondrial OS=Rattus norvegicus OX=10116 GN=Stoml2 PE=1 SV=1</td></tr>
<tr><td>98</td><td>194</td><td>Q9R063|PRDX5_RAT</td><td>73.05</td><td>13</td><td>13</td><td>8.15E+05</td><td>3</td><td>3</td><td>3</td><td></td><td>22179</td><td>Peroxiredoxin-5  mitochondrial OS=Rattus norvegicus OX=10116 GN=Prdx5 PE=1 SV=1</td></tr>
<tr><td>98</td><td>195</td><td>tr|A0A0G2JSS8|A0A0G2JSS8_RAT</td><td>73.05</td><td>13</td><td>13</td><td>8.15E+05</td><td>3</td><td>3</td><td>3</td><td></td><td>22207</td><td>Peroxiredoxin OS=Rattus norvegicus OX=10116 GN=Prdx5 PE=1 SV=1</td></tr>
<tr><td>49</td><td>1309</td><td>tr|B2GUW4|B2GUW4_RAT</td><td>71.58</td><td>2</td><td>2</td><td>1.57E+07</td><td>6</td><td>6</td><td>12</td><td></td><td>74060</td><td>Exdl2 protein OS=Rattus norvegicus OX=10116 GN=Exd2 PE=2 SV=1</td></tr>
<tr><td>194</td><td>814</td><td>P45953|ACADV_RAT</td><td>69.12</td><td>3</td><td>3</td><td>1.58E+05</td><td>1</td><td>1</td><td>1</td><td></td><td>70749</td><td>Very long-chain specific acyl-CoA dehydrogenase  mitochondrial OS=Rattus norvegicus OX=10116 GN=Acadvl PE=1 SV=1</td></tr>
</table>

| | | | | | | | | | | | | |
|---|---|---|---|---|---|---|---|---|---|---|---|---|
| 194 | 815 | tr\|Q5M9H2\|Q5M9H2_RAT | 69.12 | 3 | 3 | 1.58E+05 | 1 | 1 | 1 | | 70821 | Acyl-Coenzyme A dehydrogenase very long chain OS=Rattus norvegicus OX=10116 GN=Acadvl PE=1 SV=1 |
| 99 | 5949 | tr\|B2RYU0\|B2RYU0_RAT | 68.6 | 9 | 9 | 1.33E+06 | 3 | 3 | 3 | | 11842 | NADH dehydrogenase (Ubiquinone) 1 beta subcomplex 2 (Predicted) isoform CRA_b OS=Rattus norvegicus OX=10116 GN=Ndufb2 PE=1 SV=1 |
| 145 | 232 | tr\|M0R6J0\|M0R6J0_RAT | 67.31 | 6 | 6 | 4.06E+05 | 2 | 2 | 2 | | 38374 | Mitochondrial ribosomal protein L39 OS=Rattus norvegicus OX=10116 GN=Mrpl39 PE=1 SV=2 |
| 112 | 5937 | Q6PDU7\|ATP5L_RAT | 65.67 | 17 | 17 | 7.48E+05 | 2 | 2 | 2 | | 11433 | ATP synthase subunit g mitochondrial OS=Rattus norvegicus OX=10116 GN=Atp5mg PE=1 SV=2 |
| 78 | 385 | tr\|B1WBP7\|B1WBP7_RAT | 61.44 | 24 | 24 | 2.45E+06 | 3 | 3 | 4 | | 12884 | ATP synthase subunit delta mitochondrial OS=Rattus norvegicus OX=10116 GN=Atp5f1d PE=1 SV=1 |
| 78 | 387 | tr\|G3V7Y3\|G3V7Y3_RAT | 61.44 | 17 | 17 | 2.45E+06 | 3 | 3 | 4 | | 17563 | ATP synthase subunit delta mitochondrial OS=Rattus norvegicus OX=10116 GN=Atp5f1d PE=1 SV=1 |
| 146 | 153 | tr\|Q06QA9\|Q06QA9_RAT | 59.76 | 4 | 4 | 3.89E+05 | 2 | 2 | 2 | | 56893 | Cytochrome c oxidase subunit 1 OS=Rattus norvegicus OX=10116 GN=CO1 PE=3 SV=1 |
| 146 | 154 | tr\|Q06QK0\|Q06QK0_RAT | 59.76 | 4 | 4 | 3.89E+05 | 2 | 2 | 2 | | 56880 | Cytochrome c oxidase subunit 1 OS=Rattus norvegicus OX=10116 GN=CO1 PE=3 SV=1 |
| 146 | 155 | tr\|Q8SEZ6\|Q8SEZ6_RAT | 59.76 | 4 | 4 | 3.89E+05 | 2 | 2 | 2 | | 56879 | Cytochrome c oxidase subunit 1 OS=Rattus norvegicus OX=10116 GN=COI PE=3 SV=1 |
| 146 | 156 | tr\|A0A0A1FZ34\|A0A0A1FZ34_RAT | 59.76 | 4 | 4 | 3.89E+05 | 2 | 2 | 2 | | 56937 | Cytochrome c oxidase subunit 1 OS=Rattus norvegicus OX=10116 GN=COX1 PE=3 SV=1 |
| 146 | 157 | tr\|Q8HIC9\|Q8HIC9_RAT | 59.76 | 4 | 4 | 3.89E+05 | 2 | 2 | 2 | | 56845 | Cytochrome c oxidase subunit 1 OS=Rattus norvegicus OX=10116 GN=Mt-co1 PE=3 SV=1 |
| 146 | 158 | P05503\|COX1_RAT | 59.76 | 4 | 4 | 3.89E+05 | 2 | 2 | 2 | | 56845 | Cytochrome c oxidase subunit 1 OS=Rattus norvegicus OX=10116 GN=Mtco1 PE=2 SV=3 |
| 146 | 159 | tr\|Q95938\|Q95938_RAT | 59.76 | 4 | 4 | 3.89E+05 | 2 | 2 | 2 | | 56977 | Cytochrome c oxidase subunit 1 OS=Rattus norvegicus OX=10116 GN=Co I PE=3 SV=1 |
| 132 | 343 | tr\|D3ZX69\|D3ZX69_RAT | 55.71 | 5 | 5 | 4.84E+05 | 1 | 1 | 2 | | 29466 | 39S ribosomal protein L10 mitochondrial OS=Rattus norvegicus OX=10116 GN=Mrpl10 PE=1 SV=1 |
| 132 | 344 | P0C2C4\|RM10_RAT | 55.71 | 5 | 5 | 4.84E+05 | 1 | 1 | 2 | | 29940 | 39S ribosomal protein L10 mitochondrial OS=Rattus norvegicus OX=10116 GN=Mrpl10 PE=1 SV=1 |
| 149 | 5951 | tr\|B2RZ57\|B2RZ57_RAT | 52.7 | 8 | 8 | 2.67E+05 | 2 | 2 | 2 | | 20651 | Mitochondrial ribosomal protein L18 OS=Rattus norvegicus OX=10116 GN=Mrpl18 PE=1 SV=1 |
| 264 | 1398 | Q5PQN9\|RM38_RAT | 52.03 | 3 | 3 | 0 | 1 | 1 | 1 | | 44838 | 39S ribosomal protein L38 mitochondrial OS=Rattus norvegicus OX=10116 GN=Mrpl38 PE=2 SV=2 |
| 147 | 290 | tr\|Q4G067\|Q4G067_RAT | 50.43 | 8 | 8 | 1.80E+05 | 2 | 2 | 2 | | 37440 | Mitochondrial ribosomal protein L44 OS=Rattus norvegicus OX=10116 GN=Mrpl44 PE=1 SV=1 |
| 195 | 5938 | Q06647\|ATPO_RAT | 50.22 | 4 | 4 | 1.62E+05 | 1 | 1 | 1 | | 23398 | ATP synthase subunit O mitochondrial OS=Rattus norvegicus OX=10116 GN=Atp5po PE=1 SV=1 |
| 9 | 368 | tr\|A0A0G2JWK2\|A0A0G2JWK2_RA | 49 | 1 | 1 | 2.27E+06 | 1 | 1 | 97 | | 53049 | Methyl-CpG-binding protein 2 OS=Rattus norvegicus OX=10116 GN=Mecp2 PE=1 SV=1 |
| 9 | 369 | Q00566\|MECP2_RAT | 49 | 1 | 1 | 2.27E+06 | 1 | 1 | 97 | | 53048 | Methyl-CpG-binding protein 2 OS=Rattus norvegicus OX=10116 GN=Mecp2 PE=1 SV=1 |
| 83 | 483 | tr\|D4A4P3\|D4A4P3_RAT | 45.99 | 15 | 15 | 1.96E+06 | 2 | 2 | 4 | | 11267 | NADH:ubiquinone oxidoreductase subunit B3 OS=Rattus norvegicus OX=10116 GN=Ndufb3 PE=1 SV=1 |
| 117 | 245 | P18163\|ACSL1_RAT | 45.77 | 3 | 3 | 2.83E+05 | 2 | 1 | 2 | | 78179 | Long-chain-fatty-acid--CoA ligase 1 OS=Rattus norvegicus OX=10116 GN=Acsl1 PE=1 SV=1 |
| 131 | 416 | tr\|Q5UAJ5\|Q5UAJ5_RAT | 45.76 | 13 | 13 | 1.40E+06 | 1 | 1 | 2 | | 7642 | ATP synthase protein 8 OS=Rattus norvegicus OX=10116 GN=ATP8 PE=3 SV=1 |
| 131 | 418 | tr\|Q8SEZ4\|Q8SEZ4_RAT | 45.76 | 13 | 13 | 1.40E+06 | 1 | 1 | 2 | | 7632 | ATP synthase protein 8 OS=Rattus norvegicus OX=10116 GN=ATPase8 PE=3 SV=1 |
| 131 | 417 | P11608\|ATP8_RAT | 45.76 | 13 | 13 | 1.40E+06 | 1 | 1 | 2 | | 7630 | ATP synthase protein 8 OS=Rattus norvegicus OX=10116 GN=Mt-atp8 PE=1 SV=2 |
| 131 | 419 | tr\|Q8HIC8\|Q8HIC8_RAT | 45.76 | 13 | 13 | 1.40E+06 | 1 | 1 | 2 | | 7630 | ATP synthase protein 8 OS=Rattus norvegicus OX=10116 GN=Mt-atp8 PE=3 SV=1 |
| 67 | 816 | tr\|Q5U2T0\|Q5U2T0_RAT | 45.3 | 5 | 5 | 1.56E+05 | 3 | 2 | 5 | | 44494 | Death associated protein 3 OS=Rattus norvegicus OX=10116 GN=Dap3 PE=2 SV=1 |
| 67 | 817 | tr\|F7EZZ0\|F7EZZ0_RAT | 45.3 | 5 | 5 | 1.56E+05 | 3 | 2 | 5 | | 45112 | Death-associated protein 3 OS=Rattus norvegicus OX=10116 GN=Dap3 PE=1 SV=1 |
| 217 | 284 | tr\|A0A0G2K8Q8\|A0A0G2K8Q8_RA | 44.38 | 13 | 13 | 1.93E+05 | 1 | 1 | 1 | | 7099 | Ubiquinol-cytochrome c reductase complex III subunit X OS=Rattus norvegicus OX=10116 GN=Uqcr10 PE=1 SV=1 |
| 217 | 285 | tr\|B2RYX1\|B2RYX1_RAT | 44.38 | 12 | 12 | 1.93E+05 | 1 | 1 | 1 | | 7462 | LOC685322 protein OS=Rattus norvegicus OX=10116 GN=Uqcr10 PE=2 SV=1 |
| 129 | 81 | tr\|S5RZM8\|S5RZM8_RAT | 44.18 | 9 | 9 | 4.46E+05 | 2 | 2 | 2 | Oxidation ( | 25958 | Cytochrome c oxidase subunit 2 OS=Rattus norvegicus OX=10116 GN=COX2 PE=3 SV=1 |
| 129 | 82 | P00406\|COX2_RAT | 44.18 | 9 | 9 | 4.46E+05 | 2 | 2 | 2 | Oxidation ( | 25928 | Cytochrome c oxidase subunit 2 OS=Rattus norvegicus OX=10116 GN=Mtco2 PE=1 SV=3 |
| 129 | 41 | tr\|Q5UAJ6\|Q5UAJ6_RAT | 44.18 | 9 | 9 | 4.46E+05 | 2 | 2 | 2 | Oxidation ( | 25942 | Cytochrome c oxidase subunit 2 OS=Rattus norvegicus OX=10116 GN=COX2 PE=3 SV=1 |
| 129 | 83 | tr\|Q8SEZ5\|Q8SEZ5_RAT | 44.18 | 9 | 9 | 4.46E+05 | 2 | 2 | 2 | Oxidation ( | 25928 | Cytochrome c oxidase subunit 2 OS=Rattus norvegicus OX=10116 GN=Mt-co2 PE=1 SV=1 |
| 129 | 84 | tr\|A0A097PE04\|A0A097PE04_RAT | 44.18 | 9 | 9 | 4.46E+05 | 2 | 2 | 2 | Oxidation ( | 25894 | Cytochrome c oxidase subunit 2 OS=Rattus norvegicus OX=10116 GN=COX2 PE=3 SV=1 |
| 266 | 354 | tr\|Q3B8N9\|Q3B8N9_RAT | 43.61 | 3 | 3 | 1.67E+05 | 1 | 1 | 1 | | 32823 | Biphenyl hydrolase-like OS=Rattus norvegicus OX=10116 GN=Bphl PE=1 SV=1 |
| 265 | 300 | tr\|A0A0G2K9B4\|A0A0G2K9B4_RAT | 42.99 | 4 | 4 | 2.81E+05 | 1 | 1 | 1 | | 33652 | Mitochondrial ribosomal protein L15 OS=Rattus norvegicus OX=10116 GN=Mrpl15 PE=1 SV=1 |
| 215 | 5939 | P56571\|ES1_RAT | 40.42 | 4 | 4 | 1.71E+05 | 1 | 1 | 1 | | 28173 | ES1 protein homolog mitochondrial OS=Rattus norvegicus OX=10116 PE=1 SV=2 |
| 107 | 324 | P33124\|ACSL6_RAT | 39.77 | 2 | 2 | | 2 | 0 | 2 | | 78180 | Long-chain-fatty-acid--CoA ligase 6 OS=Rattus norvegicus OX=10116 GN=Acsl6 PE=1 SV=1 |
| 118 | 326 | P05696\|KPCA_RAT | 39.51 | 1 | 1 | 4.88E+05 | 1 | 1 | 2 | | 76792 | Protein kinase C alpha type OS=Rattus norvegicus OX=10116 GN=Prkca PE=1 SV=3 |
| 73 | 5959 | D3ZS74\|OMA1_RAT | 38.19 | 1 | 1 | 1.35E+06 | 2 | 1 | 5 | | 57145 | Metalloendopeptidase OMA1 mitochondrial OS=Rattus norvegicus OX=10116 GN=Oma1 PE=3 SV=1 |
| 268 | 233 | P29147\|BDH_RAT | 37.75 | 3 | 3 | 1.53E+05 | 1 | 1 | 1 | | 38202 | D-beta-hydroxybutyrate dehydrogenase mitochondrial OS=Rattus norvegicus OX=10116 GN=Bdh1 PE=1 SV=2 |
| 268 | 234 | tr\|A0A0G2JSH2\|A0A0G2JSH2_RAT | 37.75 | 3 | 3 | 1.53E+05 | 1 | 1 | 1 | | 38333 | 3-hydroxybutyrate dehydrogenase type 1 isoform CRA_a OS=Rattus norvegicus OX=10116 GN=Bdh1 PE=1 SV=1 |
| 267 | 357 | tr\|D3ZLT1\|D3ZLT1_RAT | 37.73 | 16 | 16 | 2.65E+05 | 1 | 1 | 1 | Oxidation ( | 16568 | NADH dehydrogenase (Ubiquinone) 1 beta subcomplex 7 (Predicted) OS=Rattus norvegicus OX=10116 GN=Ndufb7 PE=1 SV=1 |
| 269 | 346 | Q63750\|RM23_RAT | 36.9 | 17 | 17 | 1.47E+05 | 1 | 1 | 1 | | 17050 | 39S ribosomal protein L23 mitochondrial OS=Rattus norvegicus OX=10116 GN=Mrpl23 PE=2 SV=1 |
| 148 | 495 | tr\|D4ACN9\|D4ACN9_RAT | 36.42 | 4 | 4 | 5.61E+05 | 2 | 1 | 2 | | 34267 | Solute carrier family 25 member 36 OS=Rattus norvegicus OX=10116 GN=Slc25a36 PE=3 SV=1 |
| 218 | 226 | tr\|D3ZXF9\|D3ZXF9_RAT | 36.28 | 4 | 4 | 0 | 1 | 1 | 1 | | 29441 | Mitochondrial ribosomal protein L12 OS=Rattus norvegicus OX=10116 GN=Mrpl12 PE=1 SV=1 |
| 151 | 406 | tr\|D3ZD23\|D3ZD23_RAT | 34.51 | 1 | 1 | 9.87E+05 | 2 | 2 | 2 | | 67300 | ATP-binding cassette subfamily E member 1 OS=Rattus norvegicus OX=10116 GN=Abce1 PE=1 SV=1 |
| 270 | 312 | P0DN35\|NDUB1_RAT | 34.45 | 19 | 19 | 2.20E+05 | 1 | 1 | 1 | | 6998 | NADH dehydrogenase [ubiquinone] 1 beta subcomplex subunit 1 OS=Rattus norvegicus OX=10116 GN=Ndufb1 PE=3 SV=1 |
| 128 | 341 | Q63563\|ABCC9_RAT | 34.26 | 1 | 1 | 1.25E+06 | 2 | 1 | 2 | | 174117 | ATP-binding cassette sub-family C member 9 OS=Rattus norvegicus OX=10116 GN=Abcc9 PE=1 SV=1 |
| 75 | 382 | Q01062\|PDE2A_RAT | 32.62 | 2 | 2 | 2.22E+06 | 2 | 1 | 4 | | 104664 | cGMP-dependent 3' 5'-cyclic phosphodiesterase OS=Rattus norvegicus OX=10116 GN=Pde2a PE=1 SV=2 |
| 75 | 383 | tr\|F8WFW5\|F8WFW5_RAT | 32.62 | 2 | 2 | 2.22E+06 | 2 | 1 | 4 | | 105234 | Phosphodiesterase OS=Rattus norvegicus OX=10116 GN=Pde2a PE=1 SV=1 |
| 216 | 293 | Q9WVK3\|PECR_RAT | 32.33 | 3 | 3 | 3.77E+05 | 1 | 1 | 1 | Oxidation ( | 32433 | Peroxisomal trans-2-enoyl-CoA reductase OS=Rattus norvegicus OX=10116 GN=Pecr PE=2 SV=1 |
| 216 | 294 | tr\|A0A0G2JVG4\|A0A0G2JVG4_RAT | 32.33 | 3 | 3 | 3.77E+05 | 1 | 1 | 1 | Oxidation ( | 32737 | Peroxisomal trans-2-enoyl-CoA reductase OS=Rattus norvegicus OX=10116 GN=Pecr PE=1 SV=1 |
| 93 | 5952 | tr\|Q99PW9\|Q99PW9_RAT | 31.72 | 2 | 2 | 1.06E+06 | 3 | 2 | 3 | | 120572 | Huntingtin interacting protein 1 related (Fragment) OS=Rattus norvegicus OX=10116 GN=Hip1r PE=2 SV=1 |
| 141 | 5978 | tr\|A0A0G2K2T2\|A0A0G2K2T2_RAT | 31.29 | 1 | 1 | 1.86E+05 | 2 | 1 | 2 | | 67482 | Mitoguardin 1 OS=Rattus norvegicus OX=10116 GN=Miga1 PE=4 SV=1 |
| 141 | 6015 | tr\|D4A5P3\|D4A5P3_RAT | 31.29 | 1 | 1 | 1.86E+05 | 2 | 1 | 2 | | 59813 | Mitoguardin 1 OS=Rattus norvegicus OX=10116 GN=Miga1 PE=4 SV=1 |
| 136 | 696 | Q62600\|NOS3_RAT | 30.17 | 1 | 1 | 2.09E+06 | 2 | 1 | 2 | Formylatio | 133290 | Nitric oxide synthase endothelial OS=Rattus norvegicus OX=10116 GN=Nos3 PE=1 SV=4 |
| 271 | 279 | Q5M9I5\|QCR6_RAT | 30.июл | 12 | 12 | 3.91E+05 | 1 | 1 | 1 | | 10424 | Cytochrome b-c1 complex subunit 6 mitochondrial OS=Rattus norvegicus OX=10116 GN=Uqcrh PE=3 SV=1 |
| 101 | 5976 | P04694\|ATTY_RAT | 29.27 | 5 | 5 | 2.19E+05 | 2 | 1 | 3 | Carbamido | 50635 | Tyrosine aminotransferase OS=Rattus norvegicus OX=10116 GN=Tat PE=1 SV=1 |
| 91 | 752 | tr\|D4A5Q9\|D4A5Q9_RAT | 29.25 | 2 | 2 | 2.25E+05 | 2 | 1 | 3 | | 98548 | Glycine decarboxylase OS=Rattus norvegicus OX=10116 GN=Gldc PE=1 SV=1 |
| 272 | 347 | P80433\|COX8A_RAT | 29.16 | 14 | 14 | 1.84E+05 | 1 | 1 | 1 | | 7672 | Cytochrome c oxidase subunit 8A mitochondrial OS=Rattus norvegicus OX=10116 GN=Cox8a PE=1 SV=3 |
| 85 | 441 | P11530\|DMD_RAT | 28.69 | 0 | 0 | 1.02E+06 | 2 | 1 | 3 | | 425830 | Dystrophin OS=Rattus norvegicus OX=10116 GN=Dmd PE=1 SV=2 |
| 100 | 5972 | tr\|A0A0G2KAB9\|A0A0G2KAB9_RAT | 28.53 | 4 | 4 | 0 | 2 | 1 | 3 | | 28715 | Autophagy protein 5 OS=Rattus norvegicus OX=10116 GN=Atg5 PE=1 SV=1 |
| 100 | 5977 | Q3MQ06\|ATG5_RAT | 28.53 | 4 | 4 | 0 | 2 | 1 | 3 | | 32398 | Autophagy protein 5 OS=Rattus norvegicus OX=10116 GN=Atg5 PE=2 SV=1 |
| 103 | 6010 | P60924\|DELE1_RAT | 28.41 | 3 | 3 | 6.78E+05 | 2 | 1 | 3 | | 55250 | DAP3-binding cell death enhancer 1 OS=Rattus norvegicus OX=10116 GN=Dele1 PE=2 SV=1 |
| 273 | 338 | tr\|A0A0G2JYU2\|A0A0G2JYU2_RAT | 28.24 | 4 | 4 | 1.05E+06 | 1 | 1 | 1 | | 20750 | Mitochondrial ribosomal protein L11 OS=Rattus norvegicus OX=10116 GN=mrpl11 PE=1 SV=1 |

<table>
<tr><td>273</td><td>339</td><td>Q5XIE3|RM11_RAT</td><td>28.24</td><td>4</td><td>4</td><td>1.05E+06</td><td>1</td><td>1</td><td>1</td><td></td><td>22420</td><td>39S ribosomal protein L11  mitochondrial OS=Rattus norvegicus OX=10116 GN=Mrpl11 PE=2 SV=1</td></tr>
<tr><td>120</td><td>5940</td><td>tr|Q8M7G5|Q8M7G5_RAT</td><td>27.79</td><td>5</td><td>5</td><td>3.57E+05</td><td>1</td><td>1</td><td>2</td><td>Oxidation (</td><td>24240</td><td>ATP synthase subunit a (Fragment) OS=Rattus norvegicus OX=10116 PE=2 SV=1</td></tr>
<tr><td>120</td><td>5941</td><td>tr|Q549I0|Q549I0_RAT</td><td>27.79</td><td>5</td><td>5</td><td>3.57E+05</td><td>1</td><td>1</td><td>2</td><td>Oxidation (</td><td>25050</td><td>ATP synthase subunit a OS=Rattus norvegicus OX=10116 GN=atp6 PE=4 SV=1</td></tr>
<tr><td>120</td><td>5942</td><td>P05504|ATP6_RAT</td><td>27.79</td><td>5</td><td>5</td><td>3.57E+05</td><td>1</td><td>1</td><td>2</td><td>Oxidation (</td><td>25076</td><td>ATP synthase subunit a OS=Rattus norvegicus OX=10116 GN=Mt-atp6 PE=1 SV=3</td></tr>
<tr><td>120</td><td>5943</td><td>tr|Q8HIC7|Q8HIC7_RAT</td><td>27.79</td><td>5</td><td>5</td><td>3.57E+05</td><td>1</td><td>1</td><td>2</td><td>Oxidation (</td><td>25076</td><td>ATP synthase subunit a OS=Rattus norvegicus OX=10116 GN=Mt-atp6 PE=4 SV=1</td></tr>
<tr><td>120</td><td>5944</td><td>tr|S5S1E9|S5S1E9_RAT</td><td>27.79</td><td>5</td><td>5</td><td>3.57E+05</td><td>1</td><td>1</td><td>2</td><td>Oxidation (</td><td>25049</td><td>ATP synthase subunit a OS=Rattus norvegicus OX=10116 GN=ATP6 PE=4 SV=1</td></tr>
<tr><td>120</td><td>5945</td><td>tr|Q06QE5|Q06QE5_RAT</td><td>27.79</td><td>5</td><td>5</td><td>3.57E+05</td><td>1</td><td>1</td><td>2</td><td>Oxidation (</td><td>25075</td><td>ATP synthase subunit a OS=Rattus norvegicus OX=10116 GN=ATP6 PE=4 SV=1</td></tr>
<tr><td>120</td><td>5946</td><td>tr|Q8SEZ3|Q8SEZ3_RAT</td><td>27.79</td><td>5</td><td>5</td><td>3.57E+05</td><td>1</td><td>1</td><td>2</td><td>Oxidation (</td><td>25030</td><td>ATP synthase subunit a OS=Rattus norvegicus OX=10116 GN=ATPase6 PE=4 SV=1</td></tr>
<tr><td>274</td><td>162</td><td>tr|Q5EBA4|Q5EBA4_RAT</td><td>27.41</td><td>2</td><td>2</td><td>0</td><td>1</td><td>1</td><td>1</td><td></td><td>33215</td><td>Nipsnap1 protein (Fragment) OS=Rattus norvegicus OX=10116 GN=Nipsnap1 PE=2 SV=1</td></tr>
<tr><td>274</td><td>163</td><td>tr|G3V728|G3V728_RAT</td><td>27.41</td><td>2</td><td>2</td><td>0</td><td>1</td><td>1</td><td>1</td><td></td><td>33346</td><td>4-nitrophenylphosphatase domain and non-neuronal SNAP25-like protein homolog 1 (C. elegans)  isoform CRA_b OS=Rattus norvegicus OX=10116 GN=Nipsnap1 PE=1 SV=1</td></tr>
<tr><td>135</td><td>525</td><td>tr|A0A096MJ87|A0A096MJ87_RAT</td><td>27.35</td><td>9</td><td>9</td><td>4.15E+06</td><td>2</td><td>2</td><td>2</td><td>Formylatio</td><td>21509</td><td>Septin 4 (Fragment) OS=Rattus norvegicus OX=10116 GN=Sept4 PE=1 SV=6</td></tr>
<tr><td>275</td><td>5960</td><td>tr|D3ZJH9|D3ZJH9_RAT</td><td>26.99</td><td>1</td><td>1</td><td>3.98E+06</td><td>1</td><td>1</td><td>1</td><td>Formylatio</td><td>65352</td><td>Malic enzyme OS=Rattus norvegicus OX=10116 GN=Me2 PE=1 SV=1</td></tr>
<tr><td>275</td><td>5961</td><td>tr|A0A0G2K502|A0A0G2K502_RAT</td><td>26.99</td><td>1</td><td>1</td><td>3.98E+06</td><td>1</td><td>1</td><td>1</td><td>Formylatio</td><td>66310</td><td>Malic enzyme OS=Rattus norvegicus OX=10116 GN=Me2 PE=1 SV=1</td></tr>
<tr><td>176</td><td>5933</td><td>O35800|HIF1A_RAT</td><td>26.18</td><td>1</td><td>1</td><td>0</td><td>1</td><td>1</td><td>1</td><td></td><td>92319</td><td>Hypoxia-inducible factor 1-alpha OS=Rattus norvegicus OX=10116 GN=Hif1a PE=2 SV=1</td></tr>
<tr><td>176</td><td>5934</td><td>tr|D4A8P8|D4A8P8_RAT</td><td>26.18</td><td>1</td><td>1</td><td>0</td><td>1</td><td>1</td><td>1</td><td></td><td>92336</td><td>Hypoxia-inducible factor 1-alpha OS=Rattus norvegicus OX=10116 GN=Hif1a PE=4 SV=2</td></tr>
<tr><td>197</td><td>5947</td><td>Q924S5|LONM_RAT</td><td>25.авг</td><td>1</td><td>1</td><td>9.31E+04</td><td>1</td><td>1</td><td>1</td><td></td><td>105792</td><td>Lon protease homolog  mitochondrial OS=Rattus norvegicus OX=10116 GN=Lonp1 PE=2 SV=1</td></tr>
<tr><td>168</td><td>317</td><td>tr|A0A0G2JYD4|A0A0G2JYD4_RAT</td><td>25.май</td><td>0</td><td>0</td><td>0</td><td>1</td><td>1</td><td>1</td><td></td><td>487857</td><td>Vacuolar protein sorting 13 homolog D OS=Rattus norvegicus OX=10116 GN=Vps13d PE=1 SV=1</td></tr>
<tr><td>168</td><td>318</td><td>tr|D3ZKC6|D3ZKC6_RAT</td><td>25.май</td><td>0</td><td>0</td><td>0</td><td>1</td><td>1</td><td>1</td><td></td><td>488962</td><td>Vacuolar protein sorting 13 homolog D OS=Rattus norvegicus OX=10116 GN=Vps13d PE=1 SV=1</td></tr>
<tr><td>155</td><td>455</td><td>P35559|IDE_RAT</td><td>24.14</td><td>2</td><td>2</td><td>1.88E+05</td><td>2</td><td>2</td><td>2</td><td>Carbamidomethylatio</td><td></td><td>Insulin-degrading enzyme</td></tr>
<tr><td>144</td><td>864</td><td>Q05343|RXRA_RAT</td><td>24.ноя</td><td>3</td><td>3</td><td>1.42E+05</td><td>2</td><td>1</td><td>2</td><td></td><td>51266</td><td>Retinoic acid receptor RXR-alpha OS=Rattus norvegicus OX=10116 GN=Rxra PE=1 SV=1</td></tr>
<tr><td>108</td><td>1371</td><td>tr|M0RAQ6|M0RAQ6_RAT</td><td>23.57</td><td>2</td><td>2</td><td>3.13E+05</td><td>2</td><td>2</td><td>2</td><td></td><td>99557</td><td>Hexokinase-1 OS=Rattus norvegicus OX=10116 GN=Hk1 PE=1 SV=2</td></tr>
<tr><td>108</td><td>1372</td><td>P05708|HXK1_RAT</td><td>23.57</td><td>2</td><td>2</td><td>3.13E+05</td><td>2</td><td>2</td><td>2</td><td></td><td>102408</td><td>Hexokinase-1 OS=Rattus norvegicus OX=10116 GN=Hk1 PE=1 SV=4</td></tr>
<tr><td>123</td><td>1292</td><td>tr|G3V6I4|G3V6I4_RAT</td><td>23.29</td><td>5</td><td>5</td><td>8.34E+04</td><td>1</td><td>1</td><td>2</td><td></td><td>37791</td><td>Mitochondrial amidoxime reducing component 1 OS=Rattus norvegicus OX=10116 GN=Marc1 PE=1 SV=1</td></tr>
<tr><td>138</td><td>374</td><td>P48037|ANXA6_RAT</td><td>23.ноя</td><td>1</td><td>1</td><td>1.15E+06</td><td>1</td><td>1</td><td>2</td><td></td><td>75754</td><td>Annexin A6 OS=Rattus norvegicus OX=10116 GN=Anxa6 PE=1 SV=2</td></tr>
<tr><td>138</td><td>375</td><td>tr|Q6IMZ3|Q6IMZ3_RAT</td><td>23.ноя</td><td>1</td><td>1</td><td>1.15E+06</td><td>1</td><td>1</td><td>2</td><td></td><td>75756</td><td>Annexin OS=Rattus norvegicus OX=10116 GN=Anxa6 PE=1 SV=1</td></tr>
<tr><td>221</td><td>793</td><td>tr|A0A0G2KAN7|A0A0G2KAN7_RA</td><td>22.92</td><td>1</td><td>1</td><td>4.70E+05</td><td>1</td><td>1</td><td>1</td><td></td><td>65838</td><td>Glutaminase kidney isoform  mitochondrial OS=Rattus norvegicus OX=10116 GN=Gls PE=1 SV=1</td></tr>
<tr><td>221</td><td>1207</td><td>tr|A0A0G2K1T0|A0A0G2K1T0_RAT</td><td>22.92</td><td>1</td><td>1</td><td>4.70E+05</td><td>1</td><td>1</td><td>1</td><td></td><td>73866</td><td>Glutaminase kidney isoform  mitochondrial OS=Rattus norvegicus OX=10116 GN=Gls PE=1 SV=1</td></tr>
<tr><td>221</td><td>1208</td><td>P13264|GLSK_RAT</td><td>22.92</td><td>1</td><td>1</td><td>4.70E+05</td><td>1</td><td>1</td><td>1</td><td></td><td>74024</td><td>Glutaminase kidney isoform  mitochondrial OS=Rattus norvegicus OX=10116 GN=Gls PE=1 SV=2</td></tr>
<tr><td>220</td><td>5964</td><td>tr|B0BNI6|B0BNI6_RAT</td><td>22.91</td><td>4</td><td>4</td><td>3.93E+05</td><td>1</td><td>1</td><td>1</td><td></td><td>32492</td><td>Similar to solute carrier family 25  member 35  isoform CRA_a OS=Rattus norvegicus OX=10116 GN=Slc25a35 PE=1 SV=1</td></tr>
<tr><td>169</td><td>456</td><td>tr|A0A0G2K5L6|A0A0G2K5L6_RAT</td><td>22.75</td><td>0</td><td>0</td><td>0</td><td>1</td><td>1</td><td>1</td><td></td><td>275755</td><td>Acetyl-CoA carboxylase beta OS=Rattus norvegicus OX=10116 GN=Acacb PE=1 SV=1</td></tr>
<tr><td>169</td><td>457</td><td>tr|A0A0G2K1F2|A0A0G2K1F2_RAT</td><td>22.75</td><td>0</td><td>0</td><td>0</td><td>1</td><td>1</td><td>1</td><td></td><td>276393</td><td>Acetyl-CoA carboxylase beta OS=Rattus norvegicus OX=10116 GN=Acacb PE=1 SV=1</td></tr>
<tr><td>169</td><td>459</td><td>tr|D3ZBE2|D3ZBE2_RAT</td><td>22.75</td><td>0</td><td>0</td><td>0</td><td>1</td><td>1</td><td>1</td><td></td><td>275967</td><td>Acetyl-CoA carboxylase beta OS=Rattus norvegicus OX=10116 GN=Acacb PE=1 SV=3</td></tr>
<tr><td>169</td><td>460</td><td>tr|E9PSQ0|E9PSQ0_RAT</td><td>22.75</td><td>0</td><td>0</td><td>0</td><td>1</td><td>1</td><td>1</td><td></td><td>276255</td><td>Acetyl-CoA carboxylase beta OS=Rattus norvegicus OX=10116 GN=Acacb PE=1 SV=2</td></tr>
<tr><td>169</td><td>458</td><td>tr|O70151|O70151_RAT</td><td>22.75</td><td>0</td><td>0</td><td>0</td><td>1</td><td>1</td><td>1</td><td></td><td>276097</td><td>Acetyl-CoA carboxylase OS=Rattus norvegicus OX=10116 GN=Acacb PE=2 SV=1</td></tr>
<tr><td>169</td><td>545</td><td>P11497|ACACA_RAT</td><td>22.75</td><td>0</td><td>0</td><td>0</td><td>1</td><td>1</td><td>1</td><td></td><td>265191</td><td>Acetyl-CoA carboxylase 1 OS=Rattus norvegicus OX=10116 GN=Acaca PE=1 SV=1</td></tr>
<tr><td>224</td><td>753</td><td>P08011|MGST1_RAT</td><td>22.33</td><td>8</td><td>8</td><td>2.49E+04</td><td>1</td><td>1</td><td>1</td><td></td><td>17472</td><td>Microsomal glutathione S-transferase 1 OS=Rattus norvegicus OX=10116 GN=Mgst1 PE=1 SV=3</td></tr>
<tr><td>224</td><td>754</td><td>tr|B6DYQ4|B6DYQ4_RAT</td><td>22.33</td><td>8</td><td>8</td><td>2.49E+04</td><td>1</td><td>1</td><td>1</td><td></td><td>17472</td><td>Microsomal glutathione S-transferase OS=Rattus norvegicus OX=10116 GN=Mgst1 PE=2 SV=1</td></tr>
<tr><td>94</td><td>5968</td><td>Q9Z2P5|RIPK3_RAT</td><td>22.31</td><td>2</td><td>2</td><td>3.54E+06</td><td>1</td><td>1</td><td>3</td><td></td><td>52218</td><td>Receptor-interacting serine/threonine-protein kinase 3 OS=Rattus norvegicus OX=10116 GN=Ripk3 PE=1 SV=3</td></tr>
<tr><td>222</td><td>774</td><td>tr|D3ZB81|D3ZB81_RAT</td><td>22.сен</td><td>3</td><td>3</td><td>0</td><td>1</td><td>1</td><td>1</td><td></td><td>35227</td><td>Solute carrier family 25 member 31 OS=Rattus norvegicus OX=10116 GN=Slc25a31 PE=3 SV=3</td></tr>
<tr><td>222</td><td>1195</td><td>tr|A0A1W2Q6F5|A0A1W2Q6F5_R</td><td>22.сен</td><td>5</td><td>5</td><td>0</td><td>1</td><td>1</td><td>1</td><td></td><td>20418</td><td>Solute carrier family 25 member 31 (Fragment) OS=Rattus norvegicus OX=10116 GN=Slc25a31 PE=3 SV=1</td></tr>
<tr><td>202</td><td>1217</td><td>tr|D4AB75|D4AB75_RAT</td><td>21.89</td><td>1</td><td>1</td><td>2.03E+05</td><td>1</td><td>1</td><td>1</td><td></td><td>113160</td><td>Zinc finger protein 217 OS=Rattus norvegicus OX=10116 GN=Zfp217 PE=1 SV=1</td></tr>
<tr><td>279</td><td>557</td><td>Q9WVK7|HCDH_RAT</td><td>21.57</td><td>3</td><td>3</td><td>1.58E+05</td><td>1</td><td>1</td><td>1</td><td></td><td>34448</td><td>Hydroxyacyl-coenzyme A dehydrogenase  mitochondrial OS=Rattus norvegicus OX=10116 GN=Hadh PE=2 SV=1</td></tr>
<tr><td>77</td><td>987</td><td>Q9JIR0|RIMB1_RAT</td><td>21.5</td><td>0</td><td>0</td><td>2.47E+06</td><td>1</td><td>1</td><td>4</td><td></td><td>200202</td><td>Peripheral-type benzodiazepine receptor-associated protein 1 OS=Rattus norvegicus OX=10116 GN=Tspoap1 PE=1 SV=2</td></tr>
<tr><td>77</td><td>688</td><td>tr|F1LPI6|F1LPI6_RAT</td><td>21.5</td><td>0</td><td>0</td><td>2.47E+06</td><td>1</td><td>1</td><td>4</td><td></td><td>200075</td><td>Peripheral-type benzodiazepine receptor-associated protein 1 OS=Rattus norvegicus OX=10116 GN=Tspoap1 PE=4 SV=1</td></tr>
<tr><td>200</td><td>622</td><td>Q62812|MYH9_RAT</td><td>21.25</td><td>1</td><td>1</td><td>8.25E+04</td><td>1</td><td>1</td><td>1</td><td></td><td>226336</td><td>Myosin-9 OS=Rattus norvegicus OX=10116 GN=Myh9 PE=1 SV=3</td></tr>
<tr><td>183</td><td>541</td><td>tr|F1M9H4|F1M9H4_RAT</td><td>20.49</td><td>0</td><td>0</td><td>5.63E+06</td><td>1</td><td>1</td><td>1</td><td></td><td>225602</td><td>HIVEP zinc finger 1 OS=Rattus norvegicus OX=10116 GN=Hivep1 PE=1 SV=2</td></tr>
<tr><td>227</td><td>770</td><td>tr|Q5XHZ7|Q5XHZ7_RAT</td><td>20.45</td><td>1</td><td>1</td><td>0</td><td>1</td><td>1</td><td>1</td><td></td><td>66660</td><td>MMR_HSR1 domain containing protein RGD1359460 OS=Rattus norvegicus OX=10116 GN=Noa1 PE=2 SV=1</td></tr>
<tr><td>227</td><td>771</td><td>tr|A0A0G2JUI4|A0A0G2JUI4_RAT</td><td>20.45</td><td>1</td><td>1</td><td>0</td><td>1</td><td>1</td><td>1</td><td></td><td>66708</td><td>Nitric oxide-associated 1 OS=Rattus norvegicus OX=10116 GN=Noa1 PE=4 SV=1</td></tr>
<tr><td>227</td><td>789</td><td>tr|A0A0G2K602|A0A0G2K602_RAT</td><td>20.45</td><td>1</td><td>1</td><td>0</td><td>1</td><td>1</td><td>1</td><td></td><td>77504</td><td>MMR_HSR1 domain containing protein RGD1359460  isoform CRA_a OS=Rattus norvegicus OX=10116 GN=Noa1 PE=4 SV=1</td></tr>
</table>
