## Supplementary material for "Permeability transition pore-related changes in the proteome and channel activity of ATP synthase dimers and monomers": RLM SM-BM

| Protein Gr | Protein ID | Accession | -10lgP | Coverage ( | Coverage ( | IBAQ | #Peptides | #Unique | #Spec Sam | PTM | Avg. Mass | Description |
|---|---|---|---|---|---|---|---|---|---|---|---|---|
| 1 | 35 | P52873\|PYC_RAT | 390.62 | 56 | 56 | 2.97E+08 | 232 | 227 | 461 | Oxidation (M) | | |
| 1 | 34 | tr\|A0A0G2JTL5\|A0A0G2JTL5_ | 390.62 | 51 | 51 | 2.97E+08 | 232 | 227 | 461 | Oxidation (M) | | |
| 4 | 17 | P10719\|ATPB_RAT | 351.65 | 57 | 57 | 2.67E+08 | 126 | 124 | 249 | Oxidation (M) | | |
| 4 | 18 | tr\|G3V6D3\|G3V6D3_RAT | 351.65 | 57 | 57 | 2.67E+08 | 126 | 124 | 249 | Oxidation (M) | | |
| 3 | 8 | P07756\|CPSM_RAT | 350.04 | 48 | 48 | 1.57E+08 | 190 | 186 | 307 | Carbamidomethylatio | | Carbamoyl-phosphate synthase [ammonia], mitochondrial |
| 2 | 20 | P15999\|ATPA_RAT | 329.29 | 55 | 55 | 2.71E+08 | 141 | 140 | 366 | Oxidation ( | 59754 | ATP synthase subunit alpha mitochondrial OS=Rattus norvegicus OX=10116 GN=Atp5f1a PE=1 SV=2 |
| 2 | 21 | tr\|F1LP05\|F1LP05_RAT | 329.29 | 55 | 55 | 2.71E+08 | 141 | 140 | 366 | Oxidation ( | 59813 | ATP synthase subunit alpha OS=Rattus norvegicus OX=10116 GN=Atp5f1a PE=1 SV=1 |
| 5 | 1 | Q02253\|MMSA_RAT | 293.76 | 50 | 50 | 4.37E+07 | 70 | 69 | 120 | Carbamidomethylatio | | Methylmalonate-semialdehyde dehydrogenase [acylating], mitochondrial |
| 5 | 2 | tr\|G3V7J0\|G3V7J0_RAT | 293.76 | 50 | 50 | 4.37E+07 | 70 | 69 | 120 | Carbamidomethylatio | | Aldehyde dehydrogenase family 6, subfamily A1, isoform CRA_b |
| 10 | 289 | tr\|D3ZFJ6\|D3ZFJ6_RAT | 273.45 | 34 | 34 | 3.07E+07 | 43 | 43 | 66 | Carbamido | 60420 | Lactamase beta OS=Rattus norvegicus OX=10116 GN=Lactb PE=1 SV=1 |
| 17 | 280 | P31399\|ATP5H_RAT | 251.7 | 55 | 55 | 2.27E+07 | 28 | 28 | 44 | Carbamidomethylatio | | ATP synthase subunit d, mitochondrial |
| 7 | 15 | P00507\|AATM_RAT | 246.46 | 44 | 44 | 4.83E+07 | 44 | 42 | 78 | Carbamidomethylatio | | Aspartate aminotransferase, mitochondrial |
| 13 | 1940 | A2VCW9\|AASS_RAT | 245.83 | 29 | 29 | 1.46E+07 | 39 | 39 | 51 | Oxidation ( | 102908 | Alpha-aminoadipic semialdehyde synthase mitochondrial OS=Rattus norvegicus OX=10116 GN=Aass PE=2 SV=1 |
| 13 | 1939 | tr\|D4ACE9\|D4ACE9_RAT | 245.83 | 29 | 29 | 1.46E+07 | 39 | 39 | 51 | Oxidation ( | 103115 | Alpha-aminoadipic semialdehyde synthase mitochondrial OS=Rattus norvegicus OX=10116 GN=Aass PE=1 SV=3 |
| 11 | 301 | P35435\|ATPG_RAT | 245.24 | 42 | 42 | 3.26E+07 | 32 | 32 | 63 | Oxidation ( | 30191 | ATP synthase subunit gamma mitochondrial OS=Rattus norvegicus OX=10116 GN=Atp5f1c PE=1 SV=2 |
| 11 | 297 | tr\|Q6QI09\|Q6QI09_RAT | 245.24 | 19 | 19 | 3.26E+07 | 32 | 32 | 63 | Oxidation ( | 67721 | ATP synthase subunit gamma mitochondrial OS=Rattus norvegicus OX=10116 GN=Taf3 PE=1 SV=1 |
| 22 | 22 | tr\|D3ZFQ8\|D3ZFQ8_RAT | 238.43 | 38 | 38 | 1.80E+07 | 24 | 24 | 37 | Oxidation ( | 35435 | Cytochrome c-1 OS=Rattus norvegicus OX=10116 GN=Cyc1 PE=1 SV=3 |
| 15 | 266 | P04762\|CATA_RAT | 237.41 | 31 | 31 | 1.44E+07 | 30 | 30 | 48 | Oxidation ( | 59757 | Catalase OS=Rattus norvegicus OX=10116 GN=Cat PE=1 SV=3 |
| 9 | 13 | P32551\|QCR2_RAT | 235.06 | 30 | 30 | 3.01E+07 | 36 | 36 | 67 | Oxidation ( | 48396 | Cytochrome b-c1 complex subunit 2 mitochondrial OS=Rattus norvegicus OX=10116 GN=Uqcrc2 PE=1 SV=2 |
| 8 | 33 | Q64428\|ECHA_RAT | 233.95 | 29 | 29 | 1.98E+07 | 47 | 46 | 76 | Carbamidomethylatio | | Trifunctional enzyme subunit alpha, mitochondrial |
| 26 | 14 | P10860\|DHE3_RAT | 224.97 | 26 | 26 | 7.03E+06 | 27 | 27 | 29 | Oxidation ( | 61416 | Glutamate dehydrogenase 1 mitochondrial OS=Rattus norvegicus OX=10116 GN=Glud1 PE=1 SV=2 |
| 16 | 76 | tr\|Q68G44\|Q68G44_RAT | 219.55 | 32 | 32 | 1.46E+07 | 33 | 33 | 45 | Formylatio | 56886 | 3-hydroxy-3-methylglutaryl coenzyme A synthase OS=Rattus norvegicus OX=10116 GN=Hmgcs2 PE=1 SV=1 |
| 12 | 169 | Q60587\|ECHB_RAT | 215.35 | 28 | 28 | 1.58E+07 | 32 | 31 | 55 | Oxidation ( | 51414 | Trifunctional enzyme subunit beta mitochondrial OS=Rattus norvegicus OX=10116 GN=Hadhb PE=1 SV=1 |
| 6 | 12 | Q68FY0\|QCR1_RAT | 214.74 | 36 | 36 | 4.06E+07 | 44 | 44 | 79 | Carbamidomethylatio | | Cytochrome b-c1 complex subunit 1, mitochondrial |
| 18 | 46 | tr\|L0L4L8\|L0L4L8_RAT | 210.67 | 30 | 30 | 3.04E+06 | 30 | 3 | 43 | Oxidation (M) | | |
| 18 | 47 | tr\|A0A140GE10\|A0A140GE1 | 210.67 | 29 | 29 | 3.04E+06 | 30 | 3 | 43 | Oxidation (M) | | |
| 18 | 48 | tr\|A0A140GE11\|A0A140GE1 | 210.67 | 29 | 29 | 3.04E+06 | 30 | 3 | 43 | Oxidation (M) | | |
| 18 | 49 | tr\|A0A3Q8AGE8\|A0A3Q8AG | 210.67 | 29 | 29 | 3.04E+06 | 30 | 3 | 43 | Oxidation (M) | | |
| 18 | 50 | tr\|A0A3Q8AC68\|A0A3Q8AC | 210.67 | 29 | 29 | 3.04E+06 | 30 | 3 | 43 | Oxidation (M) | | |
| 18 | 51 | tr\|A0A3S6FM15\|A0A3S6FM1 | 210.67 | 29 | 29 | 3.04E+06 | 30 | 3 | 43 | Oxidation (M) | | |
| 18 | 55 | tr\|D6NSR8\|D6NSR8_RAT | 210.67 | 29 | 29 | 3.04E+06 | 30 | 3 | 43 | Oxidation (M) | | |
| 18 | 56 | tr\|A0A0A1FZ42\|A0A0A1FZ42 | 210.67 | 29 | 29 | 3.04E+06 | 30 | 3 | 43 | Oxidation (M) | | |
| 18 | 57 | tr\|D6NSS7\|D6NSS7_RAT | 210.67 | 29 | 29 | 3.04E+06 | 30 | 3 | 43 | Oxidation (M) | | |
| 18 | 58 | tr\|Q8SEY9\|Q8SEY9_RAT | 210.67 | 29 | 29 | 3.04E+06 | 30 | 3 | 43 | Oxidation (M) | | |
| 18 | 44 | tr\|D6NSR7\|D6NSR7_RAT | 210.67 | 29 | 29 | 3.04E+06 | 30 | 3 | 43 | Oxidation (M) | | |
| 18 | 59 | tr\|A0A220DA44\|A0A220DA4 | 210.67 | 29 | 29 | 3.04E+06 | 30 | 3 | 43 | Oxidation (M) | | |
| 18 | 60 | tr\|D6NSQ0\|D6NSQ0_RAT | 210.67 | 29 | 29 | 3.04E+06 | 30 | 3 | 43 | Oxidation (M) | | |
| 18 | 61 | tr\|Q5UAI7\|Q5UAI7_RAT | 210.67 | 29 | 29 | 3.04E+06 | 30 | 3 | 43 | Oxidation (M) | | |
| 18 | 62 | tr\|Q8HIC4\|Q8HIC4_RAT | 210.67 | 29 | 29 | 3.04E+06 | 30 | 3 | 43 | Oxidation (M) | | |
| 18 | 63 | tr\|A0A220DA02\|A0A220DA0 | 210.67 | 29 | 29 | 3.04E+06 | 30 | 3 | 43 | Oxidation (M) | | |
| 18 | 64 | tr\|A0A0S1Z1V4\|A0A0S1Z1V4 | 210.67 | 29 | 29 | 3.04E+06 | 30 | 3 | 43 | Oxidation (M) | | |
| 18 | 65 | P00159\|CYB_RAT | 210.67 | 29 | 29 | 3.04E+06 | 30 | 3 | 43 | Oxidation (M) | | |
| 18 | 67 | tr\|A0A220DA28\|A0A220DA2 | 210.67 | 29 | 29 | 3.04E+06 | 30 | 3 | 43 | Oxidation (M) | | |
| 18 | 68 | tr\|H2KXA0\|H2KXA0_RAT | 210.67 | 29 | 29 | 3.04E+06 | 30 | 3 | 43 | Oxidation (M) | | |
| 18 | 99 | tr\|D6NSP2\|D6NSP2_RAT | 210.67 | 29 | 29 | 3.04E+06 | 30 | 3 | 43 | Oxidation (M) | | |
| 20 | 117 | tr\|A0A385HC93\|A0A385HC9 | 204.51 | 25 | 25 | 1.75E+05 | 28 | 1 | 38 | | 42751 | Cytochrome b (Fragment) OS=Rattus norvegicus OX=10116 PE=3 SV=1 |
| 14 | 268 | P19511\|AT5F1_RAT | 196.15 | 28 | 28 | 2.53E+07 | 24 | 24 | 50 | | 28869 | ATP synthase F(0) complex subunit B1 mitochondrial OS=Rattus norvegicus OX=10116 GN=Atp5pb PE=1 SV=1 |
| 35 | 179 | P11240\|COX5A_RAT | 191.9 | 43 | 43 | 8.24E+06 | 12 | 12 | 18 | | 16130 | Cytochrome c oxidase subunit 5A mitochondrial OS=Rattus norvegicus OX=10116 GN=Cox5a PE=1 SV=1 |
| 23 | 228 | P16970\|ABCD3_RAT | 188.18 | 25 | 25 | 1.30E+07 | 28 | 28 | 36 | | 75316 | ATP-binding cassette sub-family D member 3 OS=Rattus norvegicus OX=10116 GN=Abcd3 PE=1 SV=3 |
| 21 | 309 | P63039\|CH60_RAT | 185.06 | 25 | 25 | 1.12E+07 | 26 | 26 | 37 | Oxidation ( | 60956 | 60 kDa heat shock protein mitochondrial OS=Rattus norvegicus OX=10116 GN=Hspd1 PE=1 SV=1 |
| 21 | 310 | tr\|A0A482IDN3\|A0A482IDN3 | 185.06 | 25 | 25 | 1.12E+07 | 26 | 26 | 37 | Oxidation ( | 60956 | Hsp60 OS=Rattus norvegicus OX=10116 GN=Hspd1 PE=2 SV=1 |
| 34 | 80 | P20788\|UCRI_RAT | 181.6 | 32 | 32 | 8.94E+06 | 12 | 12 | 19 | Oxidation ( | 29446 | Cytochrome b-c1 complex subunit Rieske mitochondrial OS=Rattus norvegicus OX=10116 GN=Uqcrfs1 PE=1 SV=2 |
| 19 | 1119 | P14882\|PCCA_RAT | 180.39 | 22 | 22 | 7.31E+06 | 28 | 25 | 39 | Oxidation ( | 81623 | Propionyl-CoA carboxylase alpha chain mitochondrial OS=Rattus norvegicus OX=10116 GN=Pcca PE=1 SV=3 |
| 25 | 466 | tr\|Q68FZ8\|Q68FZ8_RAT | 173.51 | 26 | 26 | 7.08E+06 | 21 | 21 | 29 | Carbamidomethylatio | | Propionyl coenzyme A carboxylase, beta polypeptide |
| 28 | 41 | tr\|Q5UAJ6\|Q5UAJ6_RAT | 168.26 | 46 | 46 | 6.20E+06 | 20 | 20 | 23 | Oxidation ( | 25942 | Cytochrome c oxidase subunit 2 OS=Rattus norvegicus OX=10116 GN=COX2 PE=3 SV=1 |
| 24 | 245 | P18163\|ACSL1_RAT | 168.26 | 15 | 15 | 4.51E+06 | 18 | 15 | 30 | | 78179 | Long-chain-fatty-acid--CoA ligase 1 OS=Rattus norvegicus OX=10116 GN=Acsl1 PE=1 SV=1 |
| 27 | 5938 | Q06647\|ATPO_RAT | 161.06 | 43 | 43 | 8.28E+06 | 20 | 20 | 29 | Oxidation ( | 23398 | ATP synthase subunit O mitochondrial OS=Rattus norvegicus OX=10116 GN=Atp5po PE=1 SV=1 |
| 43 | 261 | P24329\|THTR_RAT | 159.16 | 14 | 14 | 2.23E+06 | 7 | 7 | 10 | | 33407 | Thiosulfate sulfurtransferase OS=Rattus norvegicus OX=10116 GN=Tst PE=1 SV=3 |
| 32 | 5937 | Q6PDU7\|ATP5L_RAT | 151.3 | 46 | 46 | 1.18E+07 | 11 | 11 | 20 | Oxidation ( | 11433 | ATP synthase subunit g mitochondrial OS=Rattus norvegicus OX=10116 GN=Atp5mg PE=1 SV=2 |
| 33 | 100 | P17764\|THIL_RAT | 150.99 | 21 | 21 | 5.56E+06 | 14 | 14 | 19 | | 44695 | Acetyl-CoA acetyltransferase mitochondrial OS=Rattus norvegicus OX=10116 GN=Acat1 PE=1 SV=1 |
| 49 | 814 | P45953\|ACADV_RAT | 149.84 | 5 | 5 | 9.27E+05 | 6 | 6 | 7 | | 70749 | Very long-chain specific acyl-CoA dehydrogenase mitochondrial OS=Rattus norvegicus OX=10116 GN=Acadvl PE=1 SV=1 |
| 49 | 815 | tr\|Q5M9H2\|Q5M9H2_RAT | 149.84 | 5 | 5 | 9.27E+05 | 6 | 6 | 7 | | 70821 | Acyl-Coenzyme A dehydrogenase very long chain OS=Rattus norvegicus OX=10116 GN=Acadvl PE=1 SV=1 |
| 38 | 6503 | B3DMA2\|ACD11_RAT | 149.45 | 10 | 10 | 2.53E+06 | 12 | 11 | 14 | Oxidation ( | 87371 | Acyl-CoA dehydrogenase family member 11 OS=Rattus norvegicus OX=10116 GN=Acad11 PE=1 SV=1 |
| 31 | 367 | P0C2X9\|AL4A1_RAT | 148.38 | 24 | 24 | 6.33E+06 | 18 | 17 | 20 | | 61869 | Delta-1-pyrroline-5-carboxylate dehydrogenase mitochondrial OS=Rattus norvegicus OX=10116 GN=Aldh4a1 PE=1 SV=1 |
| 37 | 184 | tr\|B2RYT5\|B2RYT5_RAT | 148.32 | 34 | 34 | 3.69E+06 | 12 | 11 | 15 | | 12651 | Cox7a2l protein OS=Rattus norvegicus OX=10116 GN=Cox7a2l PE=2 SV=1 |

| | | | | | | | | | | | | |
|---|---|---|---|---|---|---|---|---|---|---|---|---|
| 30 | 153 | tr\|Q06QA9\|Q06QA9_RAT | 147.02 | 20 | 20 | 5.28E+06 | 15 | 15 | 20 | Oxidation ( | 56893 | Cytochrome c oxidase subunit 1 OS=Rattus norvegicus OX=10116 GN=CO1 PE=3 SV=1 |
| 30 | 154 | tr\|Q06QK0\|Q06QK0_RAT | 147.02 | 20 | 20 | 5.28E+06 | 15 | 15 | 20 | Oxidation ( | 56880 | Cytochrome c oxidase subunit 1 OS=Rattus norvegicus OX=10116 GN=CO1 PE=3 SV=1 |
| 30 | 155 | tr\|Q8SEZ6\|Q8SEZ6_RAT | 147.02 | 20 | 20 | 5.28E+06 | 15 | 15 | 20 | Oxidation ( | 56879 | Cytochrome c oxidase subunit 1 OS=Rattus norvegicus OX=10116 GN=COI PE=3 SV=1 |
| 30 | 156 | tr\|A0A0A1FZ34\|A0A0A1FZ34 | 147.02 | 20 | 20 | 5.28E+06 | 15 | 15 | 20 | Oxidation ( | 56937 | Cytochrome c oxidase subunit 1 OS=Rattus norvegicus OX=10116 GN=COX1 PE=3 SV=1 |
| 30 | 157 | tr\|Q8HIC9\|Q8HIC9_RAT | 147.02 | 20 | 20 | 5.28E+06 | 15 | 15 | 20 | Oxidation ( | 56845 | Cytochrome c oxidase subunit 1 OS=Rattus norvegicus OX=10116 GN=Mt-co1 PE=3 SV=1 |
| 30 | 158 | P05503\|COX1_RAT | 147.02 | 20 | 20 | 5.28E+06 | 15 | 15 | 20 | Oxidation ( | 56845 | Cytochrome c oxidase subunit 1 OS=Rattus norvegicus OX=10116 GN=Mtco1 PE=2 SV=3 |
| 30 | 159 | tr\|Q95938\|Q95938_RAT | 147.02 | 20 | 20 | 5.28E+06 | 15 | 15 | 20 | Oxidation ( | 56977 | Cytochrome c oxidase subunit 1 OS=Rattus norvegicus OX=10116 GN=Co I PE=3 SV=1 |
| 29 | 387 | tr\|G3V7Y3\|G3V7Y3_RAT | 146.37 | 30 | 30 | 7.37E+06 | 12 | 12 | 21 | Oxidation ( | 17563 | ATP synthase subunit delta mitochondrial OS=Rattus norvegicus OX=10116 GN=Atp5f1d PE=1 SV=1 |
| 39 | 218 | tr\|M0RAM5\|M0RAM5_RAT | 144.66 | 25 | 25 | 3.10E+06 | 11 | 11 | 14 | | 22155 | Glutathione peroxidase OS=Rattus norvegicus OX=10116 GN=Gpx1 PE=1 SV=1 |
| 39 | 219 | P04041\|GPX1_RAT | 144.66 | 25 | 25 | 3.10E+06 | 11 | 11 | 14 | | 22305 | Glutathione peroxidase 1 OS=Rattus norvegicus OX=10116 GN=Gpx1 PE=1 SV=4 |
| 36 | 176 | P13086\|SUCA_RAT | 144.21 | 15 | 15 | 5.36E+06 | 10 | 10 | 15 | | 36148 | Succinate--CoA ligase [ADP/GDP-forming] subunit alpha mitochondrial OS=Rattus norvegicus OX=10116 GN=Suclg1 PE=2 SV=2 |
| 36 | 177 | tr\|A0A0H2UHE1\|A0A0H2UH | 144.21 | 14 | 14 | 5.36E+06 | 10 | 10 | 15 | | 37560 | Succinate--CoA ligase [ADP/GDP-forming] subunit alpha mitochondrial OS=Rattus norvegicus OX=10116 GN=Suclg1 PE=1 SV=1 |
| 48 | 332 | Q9ER34\|ACON_RAT | 131.32 | 10 | 10 | 7.56E+05 | 7 | 7 | 7 | | 85433 | Aconitate hydratase mitochondrial OS=Rattus norvegicus OX=10116 GN=Aco2 PE=1 SV=2 |
| 42 | 244 | P07824\|ARGI1_RAT | 128.38 | 18 | 18 | 1.17E+06 | 8 | 7 | 10 | | 34973 | Arginase-1 OS=Rattus norvegicus OX=10116 GN=Arg1 PE=1 SV=2 |
| 41 | 380 | O70351\|HCD2_RAT | 104.05 | 27 | 27 | 1.95E+06 | 8 | 8 | 10 | | 27246 | 3-hydroxyacyl-CoA dehydrogenase type-2 OS=Rattus norvegicus OX=10116 GN=Hsd17b10 PE=1 SV=3 |
| 41 | 381 | tr\|B0BMW2\|B0BMW2_RAT | 104.05 | 27 | 27 | 1.95E+06 | 8 | 8 | 10 | | 27250 | 3-hydroxyacyl-CoA dehydrogenase type-2 OS=Rattus norvegicus OX=10116 GN=Hsd17b10 PE=1 SV=1 |
| 45 | 1292 | tr\|G3V6I4\|G3V6I4_RAT | 102.71 | 8 | 8 | 8.29E+05 | 5 | 5 | 9 | | 37791 | Mitochondrial amidoxime reducing component 1 OS=Rattus norvegicus OX=10116 GN=Marc1 PE=1 SV=1 |
| 55 | 511 | P32198\|CPT1A_RAT | 99.58 | 4 | 4 | 1.10E+06 | 4 | 4 | 5 | | 88126 | Carnitine O-palmitoyltransferase 1 liver isoform OS=Rattus norvegicus OX=10116 GN=Cpt1a PE=1 SV=2 |
| 50 | 416 | tr\|Q5UAJ5\|Q5UAJ5_RAT | 98.81 | 30 | 30 | 5.32E+06 | 4 | 4 | 7 | Oxidation ( | 7642 | ATP synthase protein 8 OS=Rattus norvegicus OX=10116 GN=ATP8 PE=3 SV=1 |
| 50 | 418 | tr\|Q8SEZ4\|Q8SEZ4_RAT | 98.81 | 30 | 30 | 5.32E+06 | 4 | 4 | 7 | Oxidation ( | 7632 | ATP synthase protein 8 OS=Rattus norvegicus OX=10116 GN=ATPase8 PE=3 SV=1 |
| 53 | 267 | tr\|I6V4L9\|I6V4L9_RAT | 98.81 | 13 | 13 | 5.92E+05 | 6 | 6 | 6 | Oxidation ( | 29844 | Cytochrome c oxidase subunit 3 OS=Rattus norvegicus OX=10116 GN=COX3 PE=3 SV=1 |
| 54 | 273 | tr\|A0A0G2JVH4\|A0A0G2JVH | 97.52 | 6 | 6 | 7.38E+05 | 5 | 5 | 5 | | 86230 | MICOS complex subunit MIC60 OS=Rattus norvegicus OX=10116 GN=Immt PE=1 SV=1 |
| 54 | 276 | tr\|A0A140TAG5\|A0A140TAG | 97.52 | 8 | 8 | 7.38E+05 | 5 | 5 | 5 | | 67049 | MICOS complex subunit MIC60 OS=Rattus norvegicus OX=10116 GN=Immt PE=1 SV=1 |
| 54 | 277 | Q3KR86\|MIC60_RAT | 97.52 | 8 | 8 | 7.38E+05 | 5 | 5 | 5 | | 67177 | MICOS complex subunit Mic60 (Fragment) OS=Rattus norvegicus OX=10116 GN=Immt PE=1 SV=1 |
| 65 | 29 | P07895\|SODM_RAT | 94.27 | 14 | 14 | 4.21E+05 | 4 | 4 | 4 | | 24674 | Superoxide dismutase [Mn] mitochondrial OS=Rattus norvegicus OX=10116 GN=Sod2 PE=1 SV=2 |
| 40 | 5940 | tr\|Q8M7G5\|Q8M7G5_RAT | 89.22 | 23 | 23 | 1.39E+06 | 7 | 7 | 13 | Oxidation ( | 24240 | ATP synthase subunit a (Fragment) OS=Rattus norvegicus OX=10116 PE=2 SV=1 |
| 40 | 5942 | P05504\|ATP6_RAT | 89.22 | 23 | 23 | 1.39E+06 | 7 | 7 | 13 | Oxidation ( | 25076 | ATP synthase subunit a OS=Rattus norvegicus OX=10116 GN=Mt-atp6 PE=1 SV=3 |
| 40 | 5943 | tr\|Q8HIC7\|Q8HIC7_RAT | 89.22 | 23 | 23 | 1.39E+06 | 7 | 7 | 13 | Oxidation ( | 25076 | ATP synthase subunit a OS=Rattus norvegicus OX=10116 GN=Mt-atp6 PE=4 SV=1 |
| 40 | 5944 | tr\|S5S1E9\|S5S1E9_RAT | 89.22 | 23 | 23 | 1.39E+06 | 7 | 7 | 13 | Oxidation ( | 25049 | ATP synthase subunit a OS=Rattus norvegicus OX=10116 GN=ATP6 PE=4 SV=1 |
| 40 | 5945 | tr\|Q06QE5\|Q06QE5_RAT | 89.22 | 23 | 23 | 1.39E+06 | 7 | 7 | 13 | Oxidation ( | 25075 | ATP synthase subunit a OS=Rattus norvegicus OX=10116 GN=ATP6 PE=4 SV=1 |
| 51 | 5947 | Q924S5\|LONM_RAT | 89.01 | 4 | 4 | 6.38E+05 | 5 | 5 | 6 | | 105792 | Lon protease homolog mitochondrial OS=Rattus norvegicus OX=10116 GN=Lonp1 PE=2 SV=1 |
| 89 | 1173 | tr\|A0A0G2JWI5\|A0A0G2JWI | 87.9 | 3 | 3 | 3.43E+05 | 2 | 2 | 2 | | 39453 | Glycine N-acyltransferase OS=Rattus norvegicus OX=10116 GN=Glyat PE=1 SV=1 |
| 89 | 1171 | Q5PQT3\|GLYAT_RAT | 87.9 | 4 | 4 | 3.43E+05 | 2 | 2 | 2 | | 33899 | Glycine N-acyltransferase OS=Rattus norvegicus OX=10116 GN=Glyat PE=2 SV=1 |
| 89 | 1172 | tr\|A4PB92\|A4PB92_RAT | 87.9 | 4 | 4 | 3.43E+05 | 2 | 2 | 2 | | 33899 | Glycine N-acyltransferase OS=Rattus norvegicus OX=10116 GN=Glyat PE=2 SV=1 |
| 89 | 1170 | tr\|B1H250\|B1H250_RAT | 87.9 | 4 | 4 | 3.43E+05 | 2 | 2 | 2 | | 33986 | Glycine-N-acyltransferase-like 1 OS=Rattus norvegicus OX=10116 GN=Glyatl1 PE=1 SV=1 |
| 100 | 6074 | B1WC61\|ACAD9_RAT | 87.38 | 4 | 4 | 4.67E+05 | 2 | 2 | 2 | | 68843 | Complex I assembly factor ACAD9 mitochondrial OS=Rattus norvegicus OX=10116 GN=Acad9 PE=1 SV=1 |
| 56 | 231 | tr\|Q5M949\|Q5M949_RAT | 86.85 | 13 | 13 | 4.80E+05 | 5 | 5 | 5 | | 28340 | Nipsnap homolog 3A (C. elegans) OS=Rattus norvegicus OX=10116 GN=Nipsnap3b PE=1 SV=1 |
| 63 | 233 | P29147\|BDH_RAT | 86.69 | 8 | 8 | 4.94E+05 | 3 | 3 | 4 | | 38202 | D-beta-hydroxybutyrate dehydrogenase mitochondrial OS=Rattus norvegicus OX=10116 GN=Bdh1 PE=1 SV=2 |
| 63 | 234 | tr\|A0A0G2JSH2\|A0A0G2JSH | 86.69 | 8 | 8 | 4.94E+05 | 3 | 3 | 4 | | 38333 | 3-hydroxybutyrate dehydrogenase type 1 isoform CRA_a OS=Rattus norvegicus OX=10116 GN=Bdh1 PE=1 SV=1 |
| 88 | 1610 | A0A0G2K047\|ACSS3_RAT | 84.61 | 4 | 4 | 2.56E+05 | 2 | 2 | 2 | | 74675 | Acyl-CoA synthetase short-chain family member 3 mitochondrial OS=Rattus norvegicus OX=10116 GN=Acss3 PE=1 SV=1 |
| 58 | 182 | P10888\|COX41_RAT | 84.37 | 20 | 20 | 1.77E+06 | 4 | 4 | 5 | Oxidation ( | 19515 | Cytochrome c oxidase subunit 4 isoform 1 mitochondrial OS=Rattus norvegicus OX=10116 GN=Cox4i1 PE=1 SV=1 |
| 67 | 198 | P85834\|EFTU_RAT | 78.37 | 8 | 8 | 2.55E+05 | 3 | 3 | 4 | | 49522 | Elongation factor Tu mitochondrial OS=Rattus norvegicus OX=10116 GN=Tufm PE=1 SV=1 |
| 71 | 348 | P09139\|SPYA_RAT | 78.25 | 8 | 8 | 2.79E+05 | 3 | 3 | 3 | Formylatio | 45834 | Serine--pyruvate aminotransferase mitochondrial OS=Rattus norvegicus OX=10116 GN=Agxt PE=1 SV=1 |
| 61 | 264 | P02770\|ALBU_RAT | 72.78 | 3 | 3 | 1.07E+06 | 3 | 3 | 4 | Formylatio | 68731 | Serum albumin OS=Rattus norvegicus OX=10116 GN=Alb PE=1 SV=2 |
| 61 | 263 | tr\|A0A0G2JSH5\|A0A0G2JSH | 72.78 | 3 | 3 | 1.07E+06 | 3 | 3 | 4 | Formylatio | 68759 | Serum albumin OS=Rattus norvegicus OX=10116 GN=Alb PE=1 SV=1 |
| 68 | 279 | Q5M9I5\|QCR6_RAT | 72.43 | 18 | 18 | 2.93E+06 | 2 | 2 | 4 | | 10424 | Cytochrome b-c1 complex subunit 6 mitochondrial OS=Rattus norvegicus OX=10116 GN=Uqcrh PE=3 SV=1 |
| 44 | 324 | P33124\|ACSL6_RAT | 67.62 | 3 | 3 | 1.38E+05 | 4 | 1 | 9 | Formylatio | 78180 | Long-chain-fatty-acid--CoA ligase 6 OS=Rattus norvegicus OX=10116 GN=Acsl6 PE=1 SV=1 |
| 78 | 2021 | tr\|G3V7I0\|G3V7I0_RAT | 66.8 | 14 | 14 | 1.04E+06 | 3 | 3 | 3 | | 28299 | Peroxiredoxin 3 OS=Rattus norvegicus OX=10116 GN=Prdx3 PE=1 SV=1 |
| 78 | 2022 | Q9Z0V6\|PRDX3_RAT | 66.8 | 14 | 14 | 1.04E+06 | 3 | 3 | 3 | | 28295 | Thioredoxin-dependent peroxide reductase mitochondrial OS=Rattus norvegicus OX=10116 GN=Prdx3 PE=1 SV=2 |
| 102 | 183 | tr\|D4A7L4\|D4A7L4_RAT | 63.82 | 9 | 9 | 4.12E+05 | 1 | 1 | 2 | | 17634 | NADH dehydrogenase (Ubiquinone) 1 beta subcomplex 11 (Predicted) OS=Rattus norvegicus OX=10116 GN=Ndufb11 PE=1 SV=1 |
| 79 | 295 | Q8VID1\|DHRS4_RAT | 63.57 | 13 | 13 | 3.51E+05 | 3 | 3 | 3 | | 29822 | Dehydrogenase/reductase SDR family member 4 OS=Rattus norvegicus OX=10116 GN=Dhrs4 PE=2 SV=2 |
| 47 | 427 | tr\|G3V9J8\|G3V9J8_RAT | 62.73 | 5 | 5 | 7.17E+06 | 3 | 3 | 7 | Formylatio | 87521 | Glycerol-3-phosphate acyltransferase 1 mitochondrial OS=Rattus norvegicus OX=10116 GN=Gpam PE=1 SV=2 |
| 47 | 413 | tr\|A0A0G2K2U7\|A0A0G2K2U | 62.73 | 4 | 4 | 7.17E+06 | 3 | 3 | 7 | Formylatio | 93728 | Glycerol-3-phosphate acyltransferase 1 mitochondrial OS=Rattus norvegicus OX=10116 GN=Gpam PE=1 SV=1 |
| 47 | 414 | P97564\|GPAT1_RAT | 62.73 | 4 | 4 | 7.17E+06 | 3 | 3 | 7 | Formylatio | 93715 | Glycerol-3-phosphate acyltransferase 1 mitochondrial OS=Rattus norvegicus OX=10116 GN=Gpam PE=1 SV=3 |
| 92 | 36 | tr\|Q06QK5\|Q06QK5_RAT | 61.6 | 2 | 2 | 2.10E+05 | 2 | 2 | 2 | | 68606 | NADH-ubiquinone oxidoreductase chain 5 OS=Rattus norvegicus OX=10116 GN=ND5 PE=3 SV=1 |
| 92 | 37 | P11661\|NU5M_RAT | 61.6 | 2 | 2 | 2.10E+05 | 2 | 2 | 2 | | 68618 | NADH-ubiquinone oxidoreductase chain 5 OS=Rattus norvegicus OX=10116 GN=Mtnd5 PE=3 SV=3 |
| 92 | 43 | tr\|Q06QG6\|Q06QG6_RAT | 61.6 | 2 | 2 | 2.10E+05 | 2 | 2 | 2 | | 68588 | NADH-ubiquinone oxidoreductase chain 5 OS=Rattus norvegicus OX=10116 GN=ND5 PE=3 SV=1 |
| 92 | 38 | tr\|Q06QA1\|Q06QA1_RAT | 61.6 | 2 | 2 | 2.10E+05 | 2 | 2 | 2 | | 68574 | NADH-ubiquinone oxidoreductase chain 5 OS=Rattus norvegicus OX=10116 GN=ND5 PE=3 SV=1 |
| 92 | 131 | tr\|A7XYD5\|A7XYD5_RAT | 61.6 | 2 | 2 | 2.10E+05 | 2 | 2 | 2 | | 68856 | NADH-ubiquinone oxidoreductase chain 5 OS=Rattus norvegicus OX=10116 GN=Nd5 PE=3 SV=1 |
| 92 | 39 | tr\|Q8SEZ0\|Q8SEZ0_RAT | 61.6 | 2 | 2 | 2.10E+05 | 2 | 2 | 2 | | 68618 | NADH-ubiquinone oxidoreductase chain 5 OS=Rattus norvegicus OX=10116 GN=Mt-nd5 PE=3 SV=1 |
| 92 | 40 | tr\|A0A096XKT9\|A0A096XKT | 61.6 | 2 | 2 | 2.10E+05 | 2 | 2 | 2 | | 68584 | NADH-ubiquinone oxidoreductase chain 5 OS=Rattus norvegicus OX=10116 GN=ND5 PE=3 SV=1 |
| 92 | 132 | tr\|A7XYC1\|A7XYC1_RAT | 61.6 | 2 | 2 | 2.10E+05 | 2 | 2 | 2 | | 68841 | NADH-ubiquinone oxidoreductase chain 5 OS=Rattus norvegicus OX=10116 GN=Nd5 PE=3 SV=1 |
| 92 | 134 | tr\|A0A0A1FZN8\|A0A0A1FZN | 61.6 | 2 | 2 | 2.10E+05 | 2 | 2 | 2 | | 68970 | NADH-ubiquinone oxidoreductase chain 5 OS=Rattus norvegicus OX=10116 GN=ND5 PE=3 SV=1 |
| 64 | 293 | Q9WVK3\|PECR_RAT | 60.55 | 9 | 9 | 7.81E+05 | 3 | 3 | 4 | Oxidation ( | 32433 | Peroxisomal trans-2-enoyl-CoA reductase OS=Rattus norvegicus OX=10116 GN=Pecr PE=2 SV=1 |
| 64 | 294 | tr\|A0A0G2JVG4\|A0A0G2JVG | 60.55 | 9 | 9 | 7.81E+05 | 3 | 3 | 4 | Oxidation ( | 32737 | Peroxisomal trans-2-enoyl-CoA reductase OS=Rattus norvegicus OX=10116 GN=Pecr PE=1 SV=1 |
| 46 | 220 | P12075\|COX5B_RAT | 60.48 | 22 | 22 | 6.51E+06 | 4 | 4 | 8 | | 13915 | Cytochrome c oxidase subunit 5B mitochondrial OS=Rattus norvegicus OX=10116 GN=Cox5b PE=1 SV=2 |

<table>
<tr><td>59</td><td>216</td><td>Q7TQ16|QCR8_RAT</td><td>59.61</td><td>29</td><td>29</td><td>8.57E+05</td><td>3</td><td>3</td><td>5</td><td></td><td>9849</td><td>Cytochrome b-c1 complex subunit 8 OS=Rattus norvegicus OX=10116 GN=Uqcrq PE=3 SV=1</td></tr>
<tr><td>66</td><td>12010</td><td>P04636|MDHM_RAT</td><td>55.36</td><td>5</td><td>5</td><td>3.17E+05</td><td>3</td><td>3</td><td>4</td><td></td><td>35684</td><td>Malate dehydrogenase  mitochondrial OS=Rattus norvegicus OX=10116 GN=Mdh2 PE=1 SV=2</td></tr>
<tr><td>72</td><td>399</td><td>Q91XJ1|BECN1_RAT</td><td>54.76</td><td>3</td><td>3</td><td></td><td>3</td><td>0</td><td>3</td><td></td><td>51557</td><td>Beclin-1 OS=Rattus norvegicus OX=10116 GN=Becn1 PE=1 SV=1</td></tr>
<tr><td>91</td><td>26</td><td>Q641Y2|NDUS2_RAT</td><td>53.33</td><td>4</td><td>4</td><td>3.31E+05</td><td>2</td><td>2</td><td>2</td><td></td><td>52562</td><td>NADH dehydrogenase [ubiquinone] iron-sulfur protein 2  mitochondrial OS=Rattus norvegicus OX=10116 GN=Ndufs2 PE=1 SV=1</td></tr>
<tr><td>191</td><td>110</td><td>tr|A0A0G2JVL6|A0A0G2JVL6</td><td>50.08</td><td>7</td><td>7</td><td>1.40E+05</td><td>1</td><td>1</td><td>1</td><td></td><td>19965</td><td>NADH dehydrogenase [ubiquinone] 1 alpha subcomplex subunit 8 OS=Rattus norvegicus OX=10116 GN=Ndufa8 PE=1 SV=1</td></tr>
<tr><td>76</td><td>705</td><td>Q64565|AGT2_RAT</td><td>49.15</td><td>3</td><td>3</td><td>5.03E+04</td><td>2</td><td>2</td><td>3</td><td></td><td>57201</td><td>Alanine--glyoxylate aminotransferase 2  mitochondrial OS=Rattus norvegicus OX=10116 GN=Agxt2 PE=1 SV=2</td></tr>
<tr><td>213</td><td>246</td><td>tr|Q5PQZ9|Q5PQZ9_RAT</td><td>48.34</td><td>11</td><td>11</td><td>5.06E+04</td><td>1</td><td>1</td><td>1</td><td></td><td>14359</td><td>NADH dehydrogenase [ubiquinone] 1 subunit C2 OS=Rattus norvegicus OX=10116 GN=Ndufc2 PE=1 SV=1</td></tr>
<tr><td>111</td><td>11922</td><td>Q9JJW3|ATPMD_RAT</td><td>47.66</td><td>21</td><td>21</td><td>6.95E+05</td><td>1</td><td>1</td><td>2</td><td></td><td>6408</td><td>ATP synthase membrane subunit DAPIT  mitochondrial OS=Rattus norvegicus OX=10116 GN=Atp5md PE=1 SV=1</td></tr>
<tr><td>80</td><td>162</td><td>tr|Q5EBA4|Q5EBA4_RAT</td><td>47.18</td><td>7</td><td>7</td><td>5.43E+05</td><td>2</td><td>2</td><td>3</td><td></td><td>33215</td><td>Nipsnap1 protein (Fragment) OS=Rattus norvegicus OX=10116 GN=Nipsnap1 PE=2 SV=1</td></tr>
<tr><td>80</td><td>163</td><td>tr|G3V728|G3V728_RAT</td><td>47.18</td><td>7</td><td>7</td><td>5.43E+05</td><td>2</td><td>2</td><td>3</td><td></td><td>33346</td><td>4-nitrophenylphosphatase domain and non-neuronal SNAP25-like protein homolog 1 (C. elegans)  isoform CRA_b OS=Rattus norvegicus OX=10116 GN=Nipsnap1 PE=1 SV=1</td></tr>
<tr><td>90</td><td>1422</td><td>tr|F1LPV8|F1LPV8_RAT</td><td>45.45</td><td>4</td><td>4</td><td>0</td><td>2</td><td>2</td><td>2</td><td></td><td>46639</td><td>Succinate--CoA ligase [GDP-forming] subunit beta  mitochondrial OS=Rattus norvegicus OX=10116 GN=Suclg2 PE=1 SV=2</td></tr>
<tr><td>90</td><td>1423</td><td>tr|B1H270|B1H270_RAT</td><td>45.45</td><td>4</td><td>4</td><td>0</td><td>2</td><td>2</td><td>2</td><td></td><td>46988</td><td>Succinate--CoA ligase [GDP-forming] subunit beta  mitochondrial OS=Rattus norvegicus OX=10116 GN=Suclg2 PE=2 SV=1</td></tr>
<tr><td>214</td><td>11</td><td>P67779|PHB_RAT</td><td>44.94</td><td>6</td><td>6</td><td>1.35E+05</td><td>1</td><td>1</td><td>1</td><td></td><td>29820</td><td>Prohibitin OS=Rattus norvegicus OX=10116 GN=Phb PE=1 SV=1</td></tr>
<tr><td>215</td><td>354</td><td>tr|Q3B8N9|Q3B8N9_RAT</td><td>44.05</td><td>3</td><td>3</td><td>3.10E+05</td><td>1</td><td>1</td><td>1</td><td></td><td>32823</td><td>Biphenyl hydrolase-like OS=Rattus norvegicus OX=10116 GN=Bphl PE=1 SV=1</td></tr>
<tr><td>52</td><td>888</td><td>O70600|RSAD2_RAT</td><td>42.63</td><td>6</td><td>6</td><td>1.45E+06</td><td>2</td><td>2</td><td>6</td><td>Formylatio</td><td>41255</td><td>Radical S-adenosyl methionine domain-containing protein 2 OS=Rattus norvegicus OX=10116 GN=Rsad2 PE=1 SV=1</td></tr>
<tr><td>52</td><td>889</td><td>tr|A0A0H2UHF4|A0A0H2UHI</td><td>42.63</td><td>6</td><td>6</td><td>1.45E+06</td><td>2</td><td>2</td><td>6</td><td>Formylatio</td><td>41444</td><td>RCG62278 OS=Rattus norvegicus OX=10116 GN=Rsad2 PE=4 SV=1</td></tr>
<tr><td>187</td><td>17822</td><td>P05182|CP2E1_RAT</td><td>40.67</td><td>3</td><td>3</td><td>0</td><td>1</td><td>1</td><td>1</td><td></td><td>56627</td><td>Cytochrome P450 2E1 OS=Rattus norvegicus OX=10116 GN=Cyp2e1 PE=1 SV=4</td></tr>
<tr><td>144</td><td>6630</td><td>tr|F1LZW6|F1LZW6_RAT</td><td>39.94</td><td>2</td><td>2</td><td>4.71E+04</td><td>1</td><td>1</td><td>1</td><td></td><td>54099</td><td>Solute carrier family 25 member 13 OS=Rattus norvegicus OX=10116 GN=Slc25a13 PE=1 SV=2</td></tr>
<tr><td>144</td><td>11913</td><td>tr|F1LX07|F1LX07_RAT</td><td>39.94</td><td>2</td><td>2</td><td>4.71E+04</td><td>1</td><td>1</td><td>1</td><td></td><td>71917</td><td>Solute carrier family 25 member 12 OS=Rattus norvegicus OX=10116 GN=Slc25a12 PE=1 SV=3</td></tr>
<tr><td>144</td><td>11912</td><td>tr|A0A0G2K2J7|A0A0G2K2J7</td><td>39.94</td><td>3</td><td>3</td><td>4.71E+04</td><td>1</td><td>1</td><td>1</td><td></td><td>41504</td><td>Solute carrier family 25 member 12 OS=Rattus norvegicus OX=10116 GN=Slc25a12 PE=1 SV=1</td></tr>
<tr><td>57</td><td>230</td><td>tr|B2RYS2|B2RYS2_RAT</td><td>39.92</td><td>26</td><td>26</td><td>6.00E+05</td><td>4</td><td>3</td><td>5</td><td></td><td>13559</td><td>Cytochrome b-c1 complex subunit 7 OS=Rattus norvegicus OX=10116 GN=Uqcrb PE=1 SV=1</td></tr>
<tr><td>110</td><td>854</td><td>tr|D3Z900|D3Z900_RAT</td><td>38.67</td><td>7</td><td>7</td><td>7.90E+04</td><td>2</td><td>2</td><td>2</td><td>Oxidation (</td><td>38239</td><td>Mitochondrial amidoxime reducing component 2 OS=Rattus norvegicus OX=10116 GN=Marc2 PE=1 SV=2</td></tr>
<tr><td>110</td><td>855</td><td>O88994|MARC2_RAT</td><td>38.67</td><td>7</td><td>7</td><td>7.90E+04</td><td>2</td><td>2</td><td>2</td><td>Oxidation (</td><td>38249</td><td>Mitochondrial amidoxime reducing component 2 OS=Rattus norvegicus OX=10116 GN=Marc2 PE=2 SV=1</td></tr>
<tr><td>93</td><td>326</td><td>P05696|KPCA_RAT</td><td>38.24</td><td>1</td><td>1</td><td>2.89E+05</td><td>1</td><td>1</td><td>2</td><td></td><td>76792</td><td>Protein kinase C alpha type OS=Rattus norvegicus OX=10116 GN=Prkca PE=1 SV=3</td></tr>
<tr><td>188</td><td>165</td><td>tr|D4A565|D4A565_RAT</td><td>36.59</td><td>7</td><td>7</td><td>0</td><td>1</td><td>1</td><td>1</td><td></td><td>21664</td><td>NADH dehydrogenase (Ubiquinone) 1 beta subcomplex  5 (Predicted)  isoform CRA_b OS=Rattus norvegicus OX=10116 GN=Ndufb5 PE=1 SV=1</td></tr>
<tr><td>186</td><td>283</td><td>P80431|COX7B_RAT</td><td>36.17</td><td>15</td><td>15</td><td>1.22E+05</td><td>1</td><td>1</td><td>1</td><td></td><td>8995</td><td>Cytochrome c oxidase subunit 7B  mitochondrial OS=Rattus norvegicus OX=10116 GN=Cox7b PE=1 SV=3</td></tr>
<tr><td>152</td><td>42</td><td>tr|D4A0T0|D4A0T0_RAT</td><td>35.81</td><td>10</td><td>10</td><td>1.42E+05</td><td>1</td><td>1</td><td>1</td><td></td><td>20859</td><td>NADH:ubiquinone oxidoreductase subunit B10 OS=Rattus norvegicus OX=10116 GN=Ndufb10 PE=1 SV=1</td></tr>
<tr><td>153</td><td>624</td><td>tr|Q2TVU3|Q2TVU3_RAT</td><td>35.46</td><td>3</td><td>3</td><td>9.74E+04</td><td>1</td><td>1</td><td>1</td><td></td><td>49412</td><td>TID1 OS=Rattus norvegicus OX=10116 GN=Dnaja3 PE=2 SV=1</td></tr>
<tr><td>153</td><td>625</td><td>tr|A0A0G2K4Y1|A0A0G2K4Y</td><td>35.46</td><td>3</td><td>3</td><td>9.74E+04</td><td>1</td><td>1</td><td>1</td><td></td><td>49416</td><td>DnaJ heat shock protein family (Hsp40) member A3 OS=Rattus norvegicus OX=10116 GN=Dnaja3 PE=1 SV=1</td></tr>
<tr><td>153</td><td>626</td><td>tr|Q2UZS7|Q2UZS7_RAT</td><td>35.46</td><td>2</td><td>2</td><td>9.74E+04</td><td>1</td><td>1</td><td>1</td><td></td><td>52399</td><td>Tid-1 long isoform OS=Rattus norvegicus OX=10116 GN=Dnaja3 PE=2 SV=1</td></tr>
<tr><td>153</td><td>627</td><td>tr|A0A0G2K5E4|A0A0G2K5E</td><td>35.46</td><td>2</td><td>2</td><td>9.74E+04</td><td>1</td><td>1</td><td>1</td><td></td><td>52403</td><td>DnaJ heat shock protein family (Hsp40) member A3 OS=Rattus norvegicus OX=10116 GN=Dnaja3 PE=1 SV=1</td></tr>
<tr><td>153</td><td>623</td><td>tr|G3V6I5|G3V6I5_RAT</td><td>35.46</td><td>3</td><td>3</td><td>9.74E+04</td><td>1</td><td>1</td><td>1</td><td></td><td>46777</td><td>DnaJ heat shock protein family (Hsp40) member A3 OS=Rattus norvegicus OX=10116 GN=Dnaja3 PE=1 SV=2</td></tr>
<tr><td>69</td><td>753</td><td>P08011|MGST1_RAT</td><td>35.1</td><td>8</td><td>8</td><td>2.91E+05</td><td>2</td><td>2</td><td>4</td><td></td><td>17472</td><td>Microsomal glutathione S-transferase 1 OS=Rattus norvegicus OX=10116 GN=Mgst1 PE=1 SV=3</td></tr>
<tr><td>69</td><td>754</td><td>tr|B6DYQ4|B6DYQ4_RAT</td><td>35.1</td><td>8</td><td>8</td><td>2.91E+05</td><td>2</td><td>2</td><td>4</td><td></td><td>17472</td><td>Microsomal glutathione S-transferase OS=Rattus norvegicus OX=10116 GN=Mgst1 PE=2 SV=1</td></tr>
<tr><td>83</td><td>11914</td><td>P29419|ATP5I_RAT</td><td>35.06</td><td>13</td><td>13</td><td>4.51E+05</td><td>1</td><td>1</td><td>3</td><td></td><td>8255</td><td>ATP synthase subunit e  mitochondrial OS=Rattus norvegicus OX=10116 GN=Atp5me PE=1 SV=3</td></tr>
<tr><td>216</td><td>12296</td><td>tr|G3V734|G3V734_RAT</td><td>34.07</td><td>3</td><td>3</td><td>0</td><td>1</td><td>1</td><td>1</td><td></td><td>36133</td><td>2 4-dienoyl CoA reductase 1  mitochondrial  isoform CRA_a OS=Rattus norvegicus OX=10116 GN=Decr1 PE=1 SV=1</td></tr>
<tr><td>216</td><td>12297</td><td>Q64591|DECR_RAT</td><td>34.07</td><td>3</td><td>3</td><td>0</td><td>1</td><td>1</td><td>1</td><td></td><td>36133</td><td>2 4-dienoyl-CoA reductase  mitochondrial OS=Rattus norvegicus OX=10116 GN=Decr1 PE=1 SV=2</td></tr>
<tr><td>166</td><td>373</td><td>Q4KLP0|DHTK1_RAT</td><td>31.97</td><td>1</td><td>1</td><td>7.76E+05</td><td>1</td><td>1</td><td>1</td><td></td><td>102642</td><td>Probable 2-oxoglutarate dehydrogenase E1 component DHKTD1  mitochondrial OS=Rattus norvegicus OX=10116 GN=Dhtkd1 PE=2 SV=1</td></tr>
<tr><td>217</td><td>1796</td><td>P22071|3BHS1_RAT</td><td>30.95</td><td>3</td><td>3</td><td>4.97E+04</td><td>1</td><td>1</td><td>1</td><td></td><td>42037</td><td>3 beta-hydroxysteroid dehydrogenase/Delta 5-->4-isomerase type 1 OS=Rattus norvegicus OX=10116 GN=Hsd3b1 PE=1 SV=3</td></tr>
<tr><td>217</td><td>1797</td><td>P22072|3BHS2_RAT</td><td>30.95</td><td>3</td><td>3</td><td>4.97E+04</td><td>1</td><td>1</td><td>1</td><td></td><td>42277</td><td>3 beta-hydroxysteroid dehydrogenase/Delta 5-->4-isomerase type 2 OS=Rattus norvegicus OX=10116 GN=Hsd3b PE=2 SV=3</td></tr>
<tr><td>217</td><td>28053</td><td>P27364|3BHS5_RAT</td><td>30.95</td><td>3</td><td>3</td><td>4.97E+04</td><td>1</td><td>1</td><td>1</td><td></td><td>42206</td><td>NADPH-dependent 3-keto-steroid reductase Hsd3b5 OS=Rattus norvegicus OX=10116 GN=Hsd3b5 PE=1 SV=3</td></tr>
<tr><td>189</td><td>368</td><td>tr|A0A0G2JWK2|A0A0G2JWI</td><td>29.45</td><td>1</td><td>1</td><td>0</td><td>1</td><td>1</td><td>1</td><td></td><td>53049</td><td>Methyl-CpG-binding protein 2 OS=Rattus norvegicus OX=10116 GN=Mecp2 PE=1 SV=1</td></tr>
<tr><td>189</td><td>369</td><td>Q00566|MECP2_RAT</td><td>29.45</td><td>1</td><td>1</td><td>0</td><td>1</td><td>1</td><td>1</td><td></td><td>53048</td><td>Methyl-CpG-binding protein 2 OS=Rattus norvegicus OX=10116 GN=Mecp2 PE=1 SV=1</td></tr>
<tr><td>74</td><td>595</td><td>D4A929|WDR81_RAT</td><td>29.42</td><td>1</td><td>1</td><td>1.09E+06</td><td>2</td><td>2</td><td>3</td><td>Carbamidomethylatio</td><td></td><td>WD repeat-containing protein 81</td></tr>
<tr><td>219</td><td>11916</td><td>Q5I0C5|FMT_RAT</td><td>29.18</td><td>2</td><td>2</td><td>4.28E+05</td><td>1</td><td>1</td><td>1</td><td>Formylatio</td><td>42997</td><td>Methionyl-tRNA formyltransferase  mitochondrial OS=Rattus norvegicus OX=10116 GN=Mtfmt PE=2 SV=1</td></tr>
<tr><td>109</td><td>406</td><td>tr|D3ZD23|D3ZD23_RAT</td><td>29.ноя</td><td>1</td><td>1</td><td>9.21E+05</td><td>2</td><td>2</td><td>2</td><td></td><td>67300</td><td>ATP-binding cassette subfamily E member 1 OS=Rattus norvegicus OX=10116 GN=Abce1 PE=1 SV=1</td></tr>
<tr><td>113</td><td>23328</td><td>tr|Q5BK11|Q5BK11_RAT</td><td>29.мар</td><td>5</td><td>5</td><td>9.17E+04</td><td>2</td><td>1</td><td>2</td><td></td><td>34506</td><td>Signal transducing adaptor family member 1 OS=Rattus norvegicus OX=10116 GN=Stap1 PE=2 SV=1</td></tr>
<tr><td>218</td><td>23266</td><td>P0CG51|UBB_RAT</td><td>29.янв</td><td>4</td><td>4</td><td>9.47E+04</td><td>1</td><td>1</td><td>1</td><td></td><td>34369</td><td>Polyubiquitin-B OS=Rattus norvegicus OX=10116 GN=Ubb PE=1 SV=1</td></tr>
<tr><td>220</td><td>24</td><td>tr|D3ZG43|D3ZG43_RAT</td><td>28.86</td><td>3</td><td>3</td><td>3.82E+05</td><td>1</td><td>1</td><td>1</td><td></td><td>30226</td><td>NADH dehydrogenase (Ubiquinone) Fe-S protein 3 (Predicted)  isoform CRA_c OS=Rattus norvegicus OX=10116 GN=Ndufs3 PE=1 SV=1</td></tr>
<tr><td>103</td><td>338</td><td>tr|A0A0G2JYU2|A0A0G2JYU</td><td>28.57</td><td>4</td><td>4</td><td>8.45E+05</td><td>1</td><td>1</td><td>2</td><td></td><td>20750</td><td>Mitochondrial ribosomal protein L11 OS=Rattus norvegicus OX=10116 GN=mrpl11 PE=1 SV=1</td></tr>
<tr><td>103</td><td>339</td><td>Q5XIE3|RM11_RAT</td><td>28.57</td><td>4</td><td>4</td><td>8.45E+05</td><td>1</td><td>1</td><td>2</td><td></td><td>22420</td><td>39S ribosomal protein L11  mitochondrial OS=Rattus norvegicus OX=10116 GN=Mrpl11 PE=2 SV=1</td></tr>
<tr><td>116</td><td>763</td><td>O35303|DNM1L_RAT</td><td>28.37</td><td>2</td><td>2</td><td>9.48E+04</td><td>2</td><td>1</td><td>2</td><td></td><td>83908</td><td>Dynamin-1-like protein OS=Rattus norvegicus OX=10116 GN=Dnm1l PE=1 SV=1</td></tr>
<tr><td>77</td><td>347</td><td>P80433|COX8A_RAT</td><td>26.49</td><td>14</td><td>14</td><td>4.03E+05</td><td>1</td><td>1</td><td>3</td><td></td><td>7672</td><td>Cytochrome c oxidase subunit 8A  mitochondrial OS=Rattus norvegicus OX=10116 GN=Cox8a PE=1 SV=3</td></tr>
<tr><td>222</td><td>325</td><td>P11951|CX6C2_RAT</td><td>26.16</td><td>11</td><td>11</td><td>7.41E+04</td><td>1</td><td>1</td><td>1</td><td></td><td>8455</td><td>Cytochrome c oxidase subunit 6C-2 OS=Rattus norvegicus OX=10116 GN=Cox6c2 PE=1 SV=3</td></tr>
<tr><td>60</td><td>317</td><td>tr|A0A0G2JYD4|A0A0G2JYD4</td><td>25.97</td><td>0</td><td>0</td><td>1.10E+06</td><td>2</td><td>2</td><td>4</td><td>Formylatio</td><td>487857</td><td>Vacuolar protein sorting 13 homolog D OS=Rattus norvegicus OX=10116 GN=Vps13d PE=1 SV=1</td></tr>
<tr><td>60</td><td>318</td><td>tr|D3ZKC6|D3ZKC6_RAT</td><td>25.97</td><td>0</td><td>0</td><td>1.10E+06</td><td>2</td><td>2</td><td>4</td><td>Formylatio</td><td>488962</td><td>Vacuolar protein sorting 13 homolog D OS=Rattus norvegicus OX=10116 GN=Vps13d PE=1 SV=1</td></tr>
<tr><td>190</td><td>250</td><td>tr|Q06Q97|Q06Q97_RAT</td><td>25.15</td><td>4</td><td>4</td><td>1.56E+04</td><td>1</td><td>1</td><td>1</td><td></td><td>38485</td><td>NADH-ubiquinone oxidoreductase chain 2 OS=Rattus norvegicus OX=10116 GN=ND2 PE=3 SV=1</td></tr>
<tr><td>190</td><td>251</td><td>tr|A0A0A1G491|A0A0A1G49</td><td>25.15</td><td>4</td><td>4</td><td>1.56E+04</td><td>1</td><td>1</td><td>1</td><td></td><td>38542</td><td>NADH-ubiquinone oxidoreductase chain 2 OS=Rattus norvegicus OX=10116 GN=ND2 PE=3 SV=1</td></tr>
<tr><td>190</td><td>252</td><td>tr|Q5UAJ8|Q5UAJ8_RAT</td><td>25.15</td><td>4</td><td>4</td><td>1.56E+04</td><td>1</td><td>1</td><td>1</td><td></td><td>38455</td><td>NADH-ubiquinone oxidoreductase chain 2 OS=Rattus norvegicus OX=10116 GN=ND2 PE=3 SV=1</td></tr>
<tr><td>190</td><td>265</td><td>tr|S5RF74|S5RF74_RAT</td><td>25.15</td><td>4</td><td>4</td><td>1.56E+04</td><td>1</td><td>1</td><td>1</td><td></td><td>38641</td><td>NADH-ubiquinone oxidoreductase chain 2 OS=Rattus norvegicus OX=10116 GN=ND2 PE=3 SV=1</td></tr>
<tr><td>190</td><td>253</td><td>tr|Q8HID0|Q8HID0_RAT</td><td>25.15</td><td>4</td><td>4</td><td>1.56E+04</td><td>1</td><td>1</td><td>1</td><td></td><td>38653</td><td>NADH-ubiquinone oxidoreductase chain 2 OS=Rattus norvegicus OX=10116 GN=Mt-nd2 PE=3 SV=1</td></tr>
<tr><td>190</td><td>254</td><td>P11662|NU2M_RAT</td><td>25.15</td><td>4</td><td>4</td><td>1.56E+04</td><td>1</td><td>1</td><td>1</td><td></td><td>38653</td><td>NADH-ubiquinone oxidoreductase chain 2 OS=Rattus norvegicus OX=10116 GN=Mtnd2 PE=3 SV=3</td></tr>
<tr><td>190</td><td>255</td><td>tr|Q06QH5|Q06QH5_RAT</td><td>25.15</td><td>4</td><td>4</td><td>1.56E+04</td><td>1</td><td>1</td><td>1</td><td></td><td>38626</td><td>NADH-ubiquinone oxidoreductase chain 2 OS=Rattus norvegicus OX=10116 GN=ND2 PE=3 SV=1</td></tr>
<tr><td>190</td><td>256</td><td>tr|Q06QB0|Q06QB0_RAT</td><td>25.15</td><td>4</td><td>4</td><td>1.56E+04</td><td>1</td><td>1</td><td>1</td><td></td><td>38534</td><td>NADH-ubiquinone oxidoreductase chain 2 OS=Rattus norvegicus OX=10116 GN=ND2 PE=3 SV=1</td></tr>
<tr><td>190</td><td>257</td><td>tr|D2E6P4|D2E6P4_RAT</td><td>25.15</td><td>4</td><td>4</td><td>1.56E+04</td><td>1</td><td>1</td><td>1</td><td></td><td>38623</td><td>NADH-ubiquinone oxidoreductase chain 2 OS=Rattus norvegicus OX=10116 GN=ND2 PE=3 SV=1</td></tr>
<tr><td>190</td><td>249</td><td>tr|Q06QE9|Q06QE9_RAT</td><td>25.15</td><td>4</td><td>4</td><td>1.56E+04</td><td>1</td><td>1</td><td>1</td><td></td><td>38598</td><td>NADH-ubiquinone oxidoreductase chain 2 OS=Rattus norvegicus OX=10116 GN=ND2 PE=3 SV=1</td></tr>
</table>

| | | | | | | | | | | | | |
|---|---|---|---|---|---|---|---|---|---|---|---|---|
| 190 | 258 | tr\|Q8SEZ7\|Q8SEZ7_RAT | 25.15 | 4 | 4 | 1.56E+04 | 1 | 1 | 1 | | 38580 | NADH-ubiquinone oxidoreductase chain 2 (Fragment) OS=Rattus norvegicus OX=10116 GN=NADH2 PE=3 SV=1 |
| 223 | 164 | tr\|D3ZF13\|D3ZF13_RAT | 24.89 | 9 | 9 | 3.11E+04 | 1 | 1 | 1 | | 17514 | Acyl carrier protein OS=Rattus norvegicus OX=10116 GN=Ndufab1 PE=1 SV=1 |
| 97 | 1838 | O35132\|CP27B_RAT | 24.16 | 2 | 2 | | 2 | 0 | 2 | | 55369 | 25-hydroxyvitamin D-1 alpha hydroxylase mitochondrial OS=Rattus norvegicus OX=10116 GN=Cyp27b1 PE=2 SV=2 |
| 97 | 1839 | tr\|M0R3V1\|M0R3V1_RAT | 24.16 | 2 | 2 | | 2 | 0 | 2 | | 55368 | 25-hydroxyvitamin D-1 alpha hydroxylase mitochondrial OS=Rattus norvegicus OX=10116 GN=Cyp27b1 PE=3 SV=1 |
| 94 | 1055 | tr\|D3ZTP0\|D3ZTP0_RAT | 23.43 | 1 | 1 | 1.14E+05 | 1 | 1 | 2 | | 101806 | 10-formyltetrahydrofolate dehydrogenase OS=Rattus norvegicus OX=10116 GN=Aldh1l2 PE=1 SV=3 |
| 225 | 11911 | P21571\|ATP5J_RAT | 23.22 | 9 | 9 | 1.49E+05 | 1 | 1 | 1 | | 12494 | ATP synthase-coupling factor 6 mitochondrial OS=Rattus norvegicus OX=10116 GN=Atp5pf PE=1 SV=1 |
| 165 | 11963 | tr\|Q5BJZ3\|Q5BJZ3_RAT | 23.18 | 1 | 1 | 0 | 1 | 1 | 1 | | 113869 | Nicotinamide nucleotide transhydrogenase OS=Rattus norvegicus OX=10116 GN=Nnt PE=1 SV=1 |
| 112 | 382 | Q01062\|PDE2A_RAT | 23.13 | 2 | 2 | 7.94E+05 | 1 | 1 | 2 | | 104664 | cGMP-dependent 3' 5'-cyclic phosphodiesterase OS=Rattus norvegicus OX=10116 GN=Pde2a PE=1 SV=2 |
| 112 | 383 | tr\|F8WFW5\|F8WFW5_RAT | 23.13 | 1 | 1 | 7.94E+05 | 1 | 1 | 2 | | 105234 | Phosphodiesterase OS=Rattus norvegicus OX=10116 GN=Pde2a PE=1 SV=1 |
| 227 | 6383 | tr\|B2RZD6\|B2RZD6_RAT | 22.89 | 10 | 10 | 1.27E+05 | 1 | 1 | 1 | | 9327 | NDUFA4 mitochondrial complex-associated OS=Rattus norvegicus OX=10116 GN=Ndufa4 PE=1 SV=1 |
| 122 | 456 | tr\|A0A0G2K5L6\|A0A0G2K5L6 | 22.26 | 0 | 0 | 6.99E+05 | 1 | 1 | 1 | | 275755 | Acetyl-CoA carboxylase beta OS=Rattus norvegicus OX=10116 GN=Acacb PE=1 SV=1 |
| 122 | 457 | tr\|A0A0G2K1F2\|A0A0G2K1F2 | 22.26 | 0 | 0 | 6.99E+05 | 1 | 1 | 1 | | 276393 | Acetyl-CoA carboxylase beta OS=Rattus norvegicus OX=10116 GN=Acacb PE=1 SV=1 |
| 122 | 459 | tr\|D3ZBE2\|D3ZBE2_RAT | 22.26 | 0 | 0 | 6.99E+05 | 1 | 1 | 1 | | 275967 | Acetyl-CoA carboxylase beta OS=Rattus norvegicus OX=10116 GN=Acacb PE=1 SV=3 |
| 122 | 460 | tr\|E9PSQ0\|E9PSQ0_RAT | 22.26 | 0 | 0 | 6.99E+05 | 1 | 1 | 1 | | 276255 | Acetyl-CoA carboxylase beta OS=Rattus norvegicus OX=10116 GN=Acacb PE=1 SV=2 |
| 122 | 458 | tr\|O70151\|O70151_RAT | 22.26 | 0 | 0 | 6.99E+05 | 1 | 1 | 1 | | 276097 | Acetyl-CoA carboxylase OS=Rattus norvegicus OX=10116 GN=Acacb PE=2 SV=1 |
| 122 | 545 | P11497\|ACACA_RAT | 22.26 | 0 | 0 | 6.99E+05 | 1 | 1 | 1 | | 265191 | Acetyl-CoA carboxylase 1 OS=Rattus norvegicus OX=10116 GN=Acaca PE=1 SV=1 |
| 229 | 374 | P48037\|ANXA6_RAT | 22.2 | 1 | 1 | 4.10E+05 | 1 | 1 | 1 | | 75754 | Annexin A6 OS=Rattus norvegicus OX=10116 GN=Anxa6 PE=1 SV=2 |
| 229 | 375 | tr\|Q6IMZ3\|Q6IMZ3_RAT | 22.2 | 1 | 1 | 4.10E+05 | 1 | 1 | 1 | | 75756 | Annexin OS=Rattus norvegicus OX=10116 GN=Anxa6 PE=1 SV=1 |
| 230 | 1810 | tr\|D3ZD73\|D3ZD73_RAT | 22.мар | 2 | 2 | 1.52E+05 | 1 | 1 | 1 | | 54245 | DEAD-box helicase 6 OS=Rattus norvegicus OX=10116 GN=Ddx6 PE=1 SV=1 |
| 193 | 135 | tr\|B0BNE6\|B0BNE6_RAT | 21.98 | 6 | 6 | 0 | 1 | 1 | 1 | | 23970 | NADH dehydrogenase (Ubiquinone) Fe-S protein 8 (Predicted) isoform CRA_a OS=Rattus norvegicus OX=10116 GN=Ndufs8 PE=1 SV=1 |
| 164 | 17950 | Q5I0L3\|SYYM_RAT | 21.94 | 2 | 2 | 5.50E+05 | 1 | 1 | 1 | Oxidation (M) | | Tyrosine--tRNA ligase, mitochondrial |
| 231 | 25 | tr\|A0A0G2KB63\|A0A0G2KB6 | 21.84 | 4 | 4 | 1.17E+05 | 1 | 1 | 1 | | 33168 | Prohibitin OS=Rattus norvegicus OX=10116 GN=Phb2 PE=1 SV=1 |
| 231 | 19 | Q5XIH7\|PHB2_RAT | 21.84 | 4 | 4 | 1.17E+05 | 1 | 1 | 1 | | 33312 | Prohibitin-2 OS=Rattus norvegicus OX=10116 GN=Phb2 PE=1 SV=1 |
| 107 | 1261 | tr\|Q5RKL4\|Q5RKL4_RAT | 21.83 | 1 | 1 | 0 | 1 | 1 | 2 | | 95977 | Dimethylglycine dehydrogenase OS=Rattus norvegicus OX=10116 GN=Dmgdh PE=1 SV=1 |
| 107 | 1262 | Q63342\|M2GD_RAT | 21.83 | 1 | 1 | 0 | 1 | 1 | 2 | | 96047 | Dimethylglycine dehydrogenase mitochondrial OS=Rattus norvegicus OX=10116 GN=Dmgdh PE=1 SV=1 |
| 107 | 1268 | tr\|A0A0G2K9Y2\|A0A0G2K9Y2 | 21.83 | 1 | 1 | 0 | 1 | 1 | 2 | | 98864 | Dimethylglycine dehydrogenase mitochondrial OS=Rattus norvegicus OX=10116 GN=Dmgdh PE=1 SV=1 |
| 155 | 6035 | Q68FU7\|COQ6_RAT | 21.68 | 3 | 3 | 0 | 1 | 1 | 1 | | 51496 | Ubiquinone biosynthesis monooxygenase COQ6 mitochondrial OS=Rattus norvegicus OX=10116 GN=Coq6 PE=2 SV=1 |
| 131 | 816 | tr\|Q5U2T0\|Q5U2T0_RAT | 21.68 | 3 | 3 | 0 | 1 | 1 | 1 | | 44494 | Death associated protein 3 OS=Rattus norvegicus OX=10116 GN=Dap3 PE=2 SV=1 |
| 131 | 817 | tr\|F7EZZ0\|F7EZZ0_RAT | 21.68 | 3 | 3 | 0 | 1 | 1 | 1 | | 45112 | Death-associated protein 3 OS=Rattus norvegicus OX=10116 GN=Dap3 PE=1 SV=1 |
| 131 | 1036 | tr\|A0A0G2K264\|A0A0G2K264 | 21.68 | 2 | 2 | 0 | 1 | 1 | 1 | | 46666 | Death-associated protein 3 OS=Rattus norvegicus OX=10116 GN=Dap3 PE=1 SV=1 |
| 232 | 11926 | D3ZAF6\|ATPK_RAT | 21.21 | 10 | 10 | 0 | 1 | 1 | 1 | | 10452 | ATP synthase subunit f mitochondrial OS=Rattus norvegicus OX=10116 GN=Atp5mf PE=1 SV=1 |
| 128 | 441 | P11530\|DMD_RAT | 21.янв | 0 | 0 | 2.97E+05 | 1 | 1 | 1 | | 425830 | Dystrophin OS=Rattus norvegicus OX=10116 GN=Dmd PE=1 SV=2 |
| 128 | 592 | tr\|B0BN02\|B0BN02_RAT | 21.янв | 1 | 1 | 2.97E+05 | 1 | 1 | 1 | | 48444 | Metaxin OS=Rattus norvegicus OX=10116 GN=Mtx1 PE=1 SV=1 |
| 233 | 11970 | P24473\|GSTK1_RAT | 20.94 | 4 | 4 | 1.05E+05 | 1 | 1 | 1 | | 25493 | Glutathione S-transferase kappa 1 OS=Rattus norvegicus OX=10116 GN=Gstk1 PE=1 SV=3 |
| 233 | 11971 | tr\|B6DYQ0\|B6DYQ0_RAT | 20.94 | 4 | 4 | 1.05E+05 | 1 | 1 | 1 | | 25493 | Glutathione S-transferase kappa OS=Rattus norvegicus OX=10116 GN=Gstk1 PE=2 SV=1 |
