## Supplementary material for "Permeability transition pore-related changes in the proteome and channel activity of ATP synthase dimers and monomers": RLM SM-BM

| Protein Gro | Protein ID | Accession | -10lgP | Coverage ( | Coverage ( | IBAQ | #Peptides | #Unique | #Spec Sam | PTM | Avg. Mass | Description |
|---|---|---|---|---|---|---|---|---|---|---|---|---|
| 2 | 1 | P10860\|DI | 381.48 | 57 | 57 | 3.81E+08 | 160 | 158 | 291 | Carbamidomethylatior | | Glutamate dehydrogenase 1, mitochondrial |
| 1 | 30 | P52873\|PY | 358.29 | 46 | 46 | 3.42E+08 | 177 | 172 | 328 | Oxidation ( | 129777 | Pyruvate carboxylase mitochondrial OS=Rattus norvegicus OX=10116 GN=Pc PE=1 SV=2 |
| 1 | 31 | tr\|A0A0G2 | 358.29 | 42 | 42 | 3.42E+08 | 177 | 172 | 328 | Oxidation ( | 140005 | Pyruvate carboxylase mitochondrial OS=Rattus norvegicus OX=10116 GN=Pc PE=1 SV=1 |
| 4 | 8 | P10719\|AT | 356.81 | 45 | 45 | 4.28E+08 | 118 | 115 | 223 | Oxidation ( | 56354 | ATP synthase subunit beta mitochondrial OS=Rattus norvegicus OX=10116 GN=Atp5f1b PE=1 SV=2 |
| 4 | 9 | tr\|G3V6D3 | 356.81 | 45 | 45 | 4.28E+08 | 118 | 115 | 223 | Oxidation ( | 56345 | ATP synthase subunit beta OS=Rattus norvegicus OX=10116 GN=Atp5f1b PE=1 SV=1 |
| 3 | 10 | P15999\|AT | 323.73 | 49 | 49 | 3.64E+08 | 120 | 118 | 259 | Carbamidomethylatior | | ATP synthase subunit alpha, mitochondrial |
| 3 | 11 | tr\|F1LP05\| | 323.73 | 49 | 49 | 3.64E+08 | 120 | 118 | 259 | Carbamidomethylation | | |
| 12 | 171 | tr\|D3ZFJ6\| | 307.73 | 29 | 29 | 5.02E+07 | 40 | 40 | 64 | Carbamido | 60420 | Lactamase beta OS=Rattus norvegicus OX=10116 GN=Lactb PE=1 SV=1 |
| 9 | 742 | A2VCW9\|A | 303.36 | 33 | 33 | 3.16E+07 | 57 | 57 | 80 | Carbamidomethylatior | | Alpha-aminoadipic semialdehyde synthase, mitochondrial |
| 5 | 4 | P07756\|CF | 295.28 | 34 | 34 | 1.01E+08 | 106 | 105 | 153 | Carbamidomethylatior | | Carbamoyl-phosphate synthase [ammonia], mitochondrial |
| 7 | 26 | P00507\|A/ | 266.63 | 42 | 42 | 1.05E+08 | 59 | 55 | 101 | Oxidation ( | 47314 | Aspartate aminotransferase mitochondrial OS=Rattus norvegicus OX=10116 GN=Got2 PE=1 SV=2 |
| 19 | 119 | P31399\|AT | 257.35 | 51 | 51 | 3.94E+07 | 25 | 24 | 46 | Carbamidomethylatior | | ATP synthase subunit d, mitochondrial |
| 10 | 63 | tr\|Q6QI09 | 253.72 | 16 | 16 | 5.10E+07 | 31 | 31 | 72 | Oxidation ( | 67721 | ATP synthase subunit gamma mitochondrial OS=Rattus norvegicus OX=10116 GN=Taf3 PE=1 SV=1 |
| 11 | 24 | Q64428\|E( | 243.93 | 27 | 27 | 3.62E+07 | 42 | 41 | 66 | Carbamidomethylatior | | Trifunctional enzyme subunit alpha, mitochondrial |
| 22 | 38 | tr\|D3ZFQ8 | 240.17 | 33 | 33 | 4.19E+07 | 28 | 28 | 41 | Oxidation ( | 35435 | Cytochrome c-1 OS=Rattus norvegicus OX=10116 GN=Cyc1 PE=1 SV=3 |
| 13 | 5806 | P14882\|PC | 239.1 | 29 | 29 | 2.21E+06 | 45 | 4 | 63 | Carbamidomethylatior | | Propionyl-CoA carboxylase alpha chain, mitochondrial |
| 16 | 27 | Q60587\|E( | 238.56 | 29 | 29 | 2.71E+07 | 36 | 35 | 55 | Oxidation ( | 51414 | Trifunctional enzyme subunit beta mitochondrial OS=Rattus norvegicus OX=10116 GN=Hadhb PE=1 SV=1 |
| 15 | 45 | P04762\|CA | 234.4 | 33 | 33 | 3.05E+07 | 36 | 36 | 58 | Oxidation ( | 59757 | Catalase OS=Rattus norvegicus OX=10116 GN=Cat PE=1 SV=3 |
| 26 | 42 | P20788\|U( | 233.43 | 36 | 36 | 2.95E+07 | 21 | 21 | 34 | Oxidation ( | 29446 | Cytochrome b-c1 complex subunit Rieske mitochondrial OS=Rattus norvegicus OX=10116 GN=Uqcrfs1 PE=1 SV=2 |
| 8 | 16 | Q68FY0\|Q | 230.64 | 34 | 34 | 7.64E+07 | 44 | 43 | 87 | Carbamido | 52849 | Cytochrome b-c1 complex subunit 1 mitochondrial OS=Rattus norvegicus OX=10116 GN=Uqcrc1 PE=1 SV=1 |
| 14 | 5804 | tr\|A0A0G2 | 229.44 | 28 | 28 | 1.96E+05 | 42 | 1 | 60 | Carbamidomethylatior | | Propionyl-CoA carboxylase alpha chain, mitochondrial |
| 14 | 5805 | tr\|A0A0G2 | 229.44 | 28 | 28 | 1.96E+05 | 42 | 1 | 60 | Carbamidomethylation | | |
| 6 | 21 | P32551\|Q( | 225.37 | 26 | 26 | 6.80E+07 | 35 | 35 | 111 | | 48396 | Cytochrome b-c1 complex subunit 2 mitochondrial OS=Rattus norvegicus OX=10116 GN=Uqcrc2 PE=1 SV=2 |
| 24 | 218 | P63039\|CH | 222.92 | 27 | 27 | 2.01E+07 | 26 | 26 | 36 | Oxidation ( | 60956 | 60 kDa heat shock protein mitochondrial OS=Rattus norvegicus OX=10116 GN=Hspd1 PE=1 SV=1 |
| 24 | 219 | tr\|A0A482 | 222.92 | 27 | 27 | 2.01E+07 | 26 | 26 | 36 | Oxidation ( | 60956 | Hsp60 OS=Rattus norvegicus OX=10116 GN=Hspd1 PE=2 SV=1 |
| 18 | 50 | Q02253\|M | 221.29 | 27 | 27 | 1.54E+07 | 30 | 29 | 47 | Carbamidomethylatior | | Methylmalonate-semialdehyde dehydrogenase [acylating], mitochondrial |
| 18 | 51 | tr\|G3V7J0 | 221.29 | 27 | 27 | 1.54E+07 | 30 | 29 | 47 | Carbamidomethylation | | |
| 23 | 124 | tr\|A0A411 | 220.22 | 30 | 30 | 6.19E+06 | 27 | 4 | 39 | Oxidation ( | 42238 | Cytochrome b (Fragment) OS=Rattus norvegicus OX=10116 GN=Cytb PE=3 SV=1 |
| 23 | 67 | tr\|L0L4L8\| | 220.22 | 30 | 30 | 6.19E+06 | 27 | 4 | 39 | Oxidation ( | 42470 | Cytochrome b (Fragment) OS=Rattus norvegicus OX=10116 GN=cytb PE=3 SV=1 |
| 23 | 68 | tr\|A0A140 | 220.22 | 30 | 30 | 6.19E+06 | 27 | 4 | 39 | Oxidation ( | 42574 | Cytochrome b (Fragment) OS=Rattus norvegicus OX=10116 PE=3 SV=1 |
| 23 | 69 | tr\|A0A140 | 220.22 | 30 | 30 | 6.19E+06 | 27 | 4 | 39 | Oxidation ( | 42588 | Cytochrome b (Fragment) OS=Rattus norvegicus OX=10116 PE=3 SV=1 |
| 23 | 125 | tr\|A0A385 | 220.22 | 30 | 30 | 6.19E+06 | 27 | 4 | 39 | Oxidation ( | 42766 | Cytochrome b (Fragment) OS=Rattus norvegicus OX=10116 PE=3 SV=1 |
| 23 | 71 | tr\|A0A3Q8 | 220.22 | 30 | 30 | 6.19E+06 | 27 | 4 | 39 | Oxidation ( | 42712 | Cytochrome b (Fragment) OS=Rattus norvegicus OX=10116 GN=Cytb PE=3 SV=1 |
| 23 | 72 | tr\|A0A3Q8 | 220.22 | 30 | 30 | 6.19E+06 | 27 | 4 | 39 | Oxidation ( | 42622 | Cytochrome b (Fragment) OS=Rattus norvegicus OX=10116 GN=Cytb PE=3 SV=1 |
| 23 | 74 | tr\|A0A3S6 | 220.22 | 30 | 30 | 6.19E+06 | 27 | 4 | 39 | Oxidation ( | 42698 | Cytochrome b (Fragment) OS=Rattus norvegicus OX=10116 GN=Cytb PE=3 SV=1 |
| 23 | 76 | tr\|A0A220 | 220.22 | 30 | 30 | 6.19E+06 | 27 | 4 | 39 | Oxidation ( | 43018 | Cytochrome b OS=Rattus norvegicus OX=10116 PE=3 SV=1 |
| 23 | 78 | tr\|D6NSS6 | 220.22 | 30 | 30 | 6.19E+06 | 27 | 4 | 39 | Oxidation ( | 42993 | Cytochrome b (Fragment) OS=Rattus norvegicus OX=10116 GN=cytb PE=3 SV=1 |
| 23 | 80 | tr\|D6NSR3 | 220.22 | 30 | 30 | 6.19E+06 | 27 | 4 | 39 | Oxidation ( | 42986 | Cytochrome b (Fragment) OS=Rattus norvegicus OX=10116 GN=cytb PE=3 SV=1 |
| 23 | 127 | tr\|D6NSP4 | 220.22 | 30 | 30 | 6.19E+06 | 27 | 4 | 39 | Oxidation ( | 43016 | Cytochrome b (Fragment) OS=Rattus norvegicus OX=10116 GN=cytb PE=3 SV=1 |
| 23 | 82 | tr\|D6NSR8 | 220.22 | 30 | 30 | 6.19E+06 | 27 | 4 | 39 | Oxidation ( | 42998 | Cytochrome b (Fragment) OS=Rattus norvegicus OX=10116 GN=cytb PE=3 SV=1 |
| 23 | 84 | tr\|A0A0A1 | 220.22 | 30 | 30 | 6.19E+06 | 27 | 4 | 39 | Oxidation ( | 42989 | Cytochrome b OS=Rattus norvegicus OX=10116 GN=CYTB PE=3 SV=1 |
| 23 | 85 | tr\|D6NSS7 | 220.22 | 30 | 30 | 6.19E+06 | 27 | 4 | 39 | Oxidation ( | 42948 | Cytochrome b (Fragment) OS=Rattus norvegicus OX=10116 GN=cytb PE=3 SV=1 |
| 23 | 86 | tr\|Q8SEY9 | 220.22 | 30 | 30 | 6.19E+06 | 27 | 4 | 39 | Oxidation ( | 43015 | Cytochrome b OS=Rattus norvegicus OX=10116 GN=cytb PE=3 SV=1 |
| 23 | 87 | tr\|D6NSR7 | 220.22 | 30 | 30 | 6.19E+06 | 27 | 4 | 39 | Oxidation ( | 42982 | Cytochrome b (Fragment) OS=Rattus norvegicus OX=10116 GN=cytb PE=3 SV=1 |
| 23 | 88 | tr\|A0A220 | 220.22 | 30 | 30 | 6.19E+06 | 27 | 4 | 39 | Oxidation ( | 42968 | Cytochrome b OS=Rattus norvegicus OX=10116 PE=3 SV=1 |
| 23 | 116 | tr\|D6NSQ( | 220.22 | 30 | 30 | 6.19E+06 | 27 | 4 | 39 | Oxidation ( | 42968 | Cytochrome b (Fragment) OS=Rattus norvegicus OX=10116 GN=cytb PE=3 SV=1 |
| 23 | 91 | tr\|Q5UAI7 | 220.22 | 30 | 30 | 6.19E+06 | 27 | 4 | 39 | Oxidation ( | 42998 | Cytochrome b OS=Rattus norvegicus OX=10116 GN=CYTB PE=3 SV=1 |
| 23 | 128 | tr\|D6NSR2 | 220.22 | 30 | 30 | 6.19E+06 | 27 | 4 | 39 | Oxidation ( | 43073 | Cytochrome b (Fragment) OS=Rattus norvegicus OX=10116 GN=cytb PE=3 SV=1 |
| 23 | 129 | tr\|D6NSR1 | 220.22 | 30 | 30 | 6.19E+06 | 27 | 4 | 39 | Oxidation ( | 43002 | Cytochrome b (Fragment) OS=Rattus norvegicus OX=10116 GN=cytb PE=3 SV=1 |
| 23 | 93 | tr\|Q8HIC4 | 220.22 | 30 | 30 | 6.19E+06 | 27 | 4 | 39 | Oxidation ( | 43012 | Cytochrome b OS=Rattus norvegicus OX=10116 GN=Mt-cyb PE=3 SV=1 |
| 23 | 94 | tr\|A0A220 | 220.22 | 30 | 30 | 6.19E+06 | 27 | 4 | 39 | Oxidation ( | 42968 | Cytochrome b OS=Rattus norvegicus OX=10116 PE=3 SV=1 |
| 23 | 95 | tr\|A0A0S1 | 220.22 | 30 | 30 | 6.19E+06 | 27 | 4 | 39 | Oxidation ( | 43016 | Cytochrome b OS=Rattus norvegicus OX=10116 GN=CYTB PE=3 SV=1 |
| 23 | 97 | P00159\|CY | 220.22 | 30 | 30 | 6.19E+06 | 27 | 4 | 39 | Oxidation ( | 43012 | Cytochrome b OS=Rattus norvegicus OX=10116 GN=Mt-Cyb PE=3 SV=3 |
| 23 | 98 | tr\|L0N311 | 220.22 | 30 | 30 | 6.19E+06 | 27 | 4 | 39 | Oxidation ( | 43045 | Cytochrome b (Fragment) OS=Rattus norvegicus OX=10116 GN=cytb PE=3 SV=1 |
| 23 | 99 | tr\|A0A220 | 220.22 | 30 | 30 | 6.19E+06 | 27 | 4 | 39 | Oxidation ( | 42970 | Cytochrome b OS=Rattus norvegicus OX=10116 PE=3 SV=1 |
| 23 | 100 | tr\|H2KXA0 | 220.22 | 30 | 30 | 6.19E+06 | 27 | 4 | 39 | Oxidation ( | 42978 | Cytochrome b (Fragment) OS=Rattus norvegicus OX=10116 GN=cytb PE=3 SV=1 |
| 27 | 126 | tr\|A0A385 | 214.87 | 26 | 26 | 3.47E+05 | 24 | 1 | 34 | | 42751 | Cytochrome b (Fragment) OS=Rattus norvegicus OX=10116 PE=3 SV=1 |
| 34 | 44 | P11240\|C( | 214.03 | 39 | 39 | 1.27E+07 | 13 | 13 | 20 | | 16130 | Cytochrome c oxidase subunit 5A mitochondrial OS=Rattus norvegicus OX=10116 GN=Cox5a PE=1 SV=1 |
| 20 | 22 | tr\|Q68G44 | 202.94 | 28 | 28 | 1.89E+07 | 33 | 33 | 45 | | 56886 | 3-hydroxy-3-methylglutaryl coenzyme A synthase OS=Rattus norvegicus OX=10116 GN=Hmgcs2 PE=1 SV=1 |
| 40 | 136 | P24329\|TH | 202.33 | 16 | 16 | 7.06E+06 | 11 | 11 | 15 | | 33407 | Thiosulfate sulfurtransferase OS=Rattus norvegicus OX=10116 GN=Tst PE=1 SV=3 |
| 17 | 246 | P19511\|AT | 200.27 | 23 | 23 | 4.99E+07 | 24 | 24 | 51 | | 28869 | ATP synthase F(0) complex subunit B1 mitochondrial OS=Rattus norvegicus OX=10116 GN=Atp5pb PE=1 SV=1 |
| 38 | 201 | P45953\|A( | 181.03 | 12 | 12 | 3.35E+06 | 12 | 11 | 15 | | 70749 | Very long-chain specific acyl-CoA dehydrogenase mitochondrial OS=Rattus norvegicus OX=10116 GN=Acadvl PE=1 SV=1 |

| | | | | | | | | | | | | |
|---|---|---|---|---|---|---|---|---|---|---|---|---|
| 38 | 202 | tr\|Q5M9H | 181.03 | 12 | 12 | 3.35E+06 | 12 | 11 | 15 | | 70821 | Acyl-Coenzyme A dehydrogenase very long chain OS=Rattus norvegicus OX=10116 GN=Acadvl PE=1 SV=1 |
| 33 | 3 | P85834\|EF | 178.71 | 18 | 18 | 7.83E+06 | 15 | 15 | 23 | | 49522 | Elongation factor Tu mitochondrial OS=Rattus norvegicus OX=10116 GN=Tufm PE=1 SV=1 |
| 25 | 54 | P18163\|AC | 168.05 | 17 | 17 | 1.73E+07 | 22 | 22 | 35 | Oxidation ( | 78179 | Long-chain-fatty-acid--CoA ligase 1 OS=Rattus norvegicus OX=10116 GN=Acsl1 PE=1 SV=1 |
| 35 | 35 | P17764\|TH | 165.28 | 18 | 18 | 9.88E+06 | 12 | 12 | 19 | | 44695 | Acetyl-CoA acetyltransferase mitochondrial OS=Rattus norvegicus OX=10116 GN=Acat1 PE=1 SV=1 |
| 21 | 6258 | tr\|Q68FZ8 | 162.08 | 24 | 24 | 1.81E+07 | 28 | 28 | 42 | Carbamidomethylatior | | Propionyl coenzyme A carboxylase, beta polypeptide |
| 32 | 1646 | Q6PDU7\|A | 157.92 | 38 | 38 | 1.61E+07 | 12 | 11 | 23 | Oxidation ( | 11433 | ATP synthase subunit g mitochondrial OS=Rattus norvegicus OX=10116 GN=Atp5mg PE=1 SV=2 |
| 61 | 277 | tr\|B2RZ24 | 157.69 | 5 | 5 | 8.39E+05 | 4 | 4 | 5 | | 47388 | Succinate-CoA ligase subunit beta (Fragment) OS=Rattus norvegicus OX=10116 GN=Sucla2 PE=2 SV=1 |
| 61 | 278 | tr\|F1LM47 | 157.69 | 5 | 5 | 8.39E+05 | 4 | 4 | 5 | | 50306 | Succinate--CoA ligase [ADP-forming] subunit beta mitochondrial OS=Rattus norvegicus OX=10116 GN=Sucla2 PE=1 SV=1 |
| 31 | 46 | P13086\|SU | 152.5 | 22 | 22 | 7.82E+06 | 15 | 15 | 23 | | 36148 | Succinate--CoA ligase [ADP/GDP-forming] subunit alpha mitochondrial OS=Rattus norvegicus OX=10116 GN=Suclg1 PE=2 SV=2 |
| 31 | 47 | tr\|A0A0H2 | 152.5 | 21 | 21 | 7.82E+06 | 15 | 15 | 23 | | 37560 | Succinate--CoA ligase [ADP/GDP-forming] subunit alpha mitochondrial OS=Rattus norvegicus OX=10116 GN=Suclg1 PE=1 SV=1 |
| 43 | 52 | tr\|M0RAM | 150.63 | 25 | 25 | 3.41E+06 | 10 | 10 | 14 | | 22155 | Glutathione peroxidase OS=Rattus norvegicus OX=10116 GN=Gpx1 PE=1 SV=1 |
| 43 | 53 | P04041\|GI | 150.63 | 25 | 25 | 3.41E+06 | 10 | 10 | 14 | | 22305 | Glutathione peroxidase 1 OS=Rattus norvegicus OX=10116 GN=Gpx1 PE=1 SV=4 |
| 41 | 115 | tr\|A0A0G2 | 150.14 | 11 | 11 | 3.15E+06 | 11 | 11 | 14 | | 86230 | MICOS complex subunit MIC60 OS=Rattus norvegicus OX=10116 GN=Immt PE=1 SV=1 |
| 39 | 11785 | B3DMA2\|A | 149.24 | 10 | 10 | 4.58E+06 | 13 | 11 | 15 | Oxidation ( | 87371 | Acyl-CoA dehydrogenase family member 11 OS=Rattus norvegicus OX=10116 GN=Acad11 PE=1 SV=1 |
| 28 | 5802 | Q06647\|AT | 147.56 | 31 | 31 | 1.20E+07 | 18 | 17 | 29 | Carbamidomethylatior | | ATP synthase subunit O, mitochondrial |
| 30 | 352 | P0C2X9\|AL | 144.1 | 19 | 19 | 8.63E+06 | 16 | 15 | 26 | | 61869 | Delta-1-pyrroline-5-carboxylate dehydrogenase mitochondrial OS=Rattus norvegicus OX=10116 GN=Aldh4a1 PE=1 SV=1 |
| 29 | 111 | tr\|S5RZM8 | 142.74 | 40 | 40 | 1.45E+07 | 18 | 18 | 28 | Oxidation ( | 25958 | Cytochrome c oxidase subunit 2 OS=Rattus norvegicus OX=10116 GN=COX2 PE=3 SV=1 |
| 29 | 112 | P00406\|CC | 142.74 | 40 | 40 | 1.45E+07 | 18 | 18 | 28 | Oxidation ( | 25928 | Cytochrome c oxidase subunit 2 OS=Rattus norvegicus OX=10116 GN=Mtco2 PE=1 SV=3 |
| 29 | 104 | tr\|Q5UAJ6 | 142.74 | 40 | 40 | 1.45E+07 | 18 | 18 | 28 | Oxidation ( | 25942 | Cytochrome c oxidase subunit 2 OS=Rattus norvegicus OX=10116 GN=COX2 PE=3 SV=1 |
| 29 | 113 | tr\|Q8SEZ5 | 142.74 | 40 | 40 | 1.45E+07 | 18 | 18 | 28 | Oxidation ( | 25928 | Cytochrome c oxidase subunit 2 OS=Rattus norvegicus OX=10116 GN=Mt-co2 PE=1 SV=1 |
| 29 | 114 | tr\|A0A097 | 142.74 | 40 | 40 | 1.45E+07 | 18 | 18 | 28 | Oxidation ( | 25894 | Cytochrome c oxidase subunit 2 OS=Rattus norvegicus OX=10116 GN=COX2 PE=3 SV=1 |
| 37 | 350 | tr\|G3V7Y3 | 135.6 | 23 | 23 | 1.15E+07 | 12 | 12 | 17 | Oxidation ( | 17563 | ATP synthase subunit delta mitochondrial OS=Rattus norvegicus OX=10116 GN=Atp5f1d PE=1 SV=1 |
| 37 | 314 | tr\|B1WBP7 | 135.6 | 31 | 31 | 1.15E+07 | 12 | 12 | 17 | Oxidation ( | 12884 | ATP synthase subunit delta mitochondrial OS=Rattus norvegicus OX=10116 GN=Atp5f1d PE=1 SV=1 |
| 49 | 377 | P80431\|CC | 130.02 | 30 | 30 | 3.74E+06 | 5 | 5 | 8 | | 8995 | Cytochrome c oxidase subunit 7B mitochondrial OS=Rattus norvegicus OX=10116 GN=Cox7b PE=1 SV=3 |
| 36 | 307 | P16970\|AE | 125.83 | 17 | 17 | 9.13E+06 | 16 | 16 | 17 | | 75316 | ATP-binding cassette sub-family D member 3 OS=Rattus norvegicus OX=10116 GN=Abcd3 PE=1 SV=3 |
| 42 | 238 | tr\|B2RYT5 | 124.03 | 34 | 34 | 7.20E+06 | 10 | 9 | 14 | | 12651 | Cox7a2l protein OS=Rattus norvegicus OX=10116 GN=Cox7a2l PE=2 SV=1 |
| 54 | 1891 | P04636\|M | 115.81 | 12 | 12 | 1.22E+06 | 6 | 6 | 6 | | 35684 | Malate dehydrogenase mitochondrial OS=Rattus norvegicus OX=10116 GN=Mdh2 PE=1 SV=2 |
| 47 | 61 | P29147\|BD | 113.62 | 8 | 8 | 2.16E+06 | 6 | 5 | 9 | | 38202 | D-beta-hydroxybutyrate dehydrogenase mitochondrial OS=Rattus norvegicus OX=10116 GN=Bdh1 PE=1 SV=2 |
| 47 | 62 | tr\|A0A0G2 | 113.62 | 8 | 8 | 2.16E+06 | 6 | 5 | 9 | | 38333 | 3-hydroxybutyrate dehydrogenase type 1 isoform CRA_a OS=Rattus norvegicus OX=10116 GN=Bdh1 PE=1 SV=1 |
| 57 | 338 | Q5M9I5\|Q | 109.84 | 18 | 18 | 4.66E+06 | 4 | 4 | 6 | | 10424 | Cytochrome b-c1 complex subunit 6 mitochondrial OS=Rattus norvegicus OX=10116 GN=Uqcrh PE=3 SV=1 |
| 74 | 5862 | tr\|G3V7I0\| | 109.69 | 12 | 12 | 2.08E+06 | 4 | 4 | 4 | | 28299 | Peroxiredoxin 3 OS=Rattus norvegicus OX=10116 GN=Prdx3 PE=1 SV=1 |
| 74 | 5863 | Q9Z0V6\|PI | 109.69 | 12 | 12 | 2.08E+06 | 4 | 4 | 4 | | 28295 | Thioredoxin-dependent peroxide reductase mitochondrial OS=Rattus norvegicus OX=10116 GN=Prdx3 PE=1 SV=2 |
| 48 | 241 | tr\|Q06QA9 | 109.06 | 11 | 11 | 1.40E+06 | 8 | 8 | 8 | | 56893 | Cytochrome c oxidase subunit 1 OS=Rattus norvegicus OX=10116 GN=CO1 PE=3 SV=1 |
| 48 | 229 | tr\|Q06QK0 | 109.06 | 11 | 11 | 1.40E+06 | 8 | 8 | 8 | | 56880 | Cytochrome c oxidase subunit 1 OS=Rattus norvegicus OX=10116 GN=CO1 PE=3 SV=1 |
| 48 | 230 | tr\|Q8SEZ6 | 109.06 | 11 | 11 | 1.40E+06 | 8 | 8 | 8 | | 56879 | Cytochrome c oxidase subunit 1 OS=Rattus norvegicus OX=10116 GN=COI PE=3 SV=1 |
| 48 | 231 | tr\|A0A0A1 | 109.06 | 11 | 11 | 1.40E+06 | 8 | 8 | 8 | | 56937 | Cytochrome c oxidase subunit 1 OS=Rattus norvegicus OX=10116 GN=COX1 PE=3 SV=1 |
| 48 | 232 | tr\|Q8HIC9 | 109.06 | 11 | 11 | 1.40E+06 | 8 | 8 | 8 | | 56845 | Cytochrome c oxidase subunit 1 OS=Rattus norvegicus OX=10116 GN=Mt-co1 PE=3 SV=1 |
| 48 | 233 | P05503\|CC | 109.06 | 11 | 11 | 1.40E+06 | 8 | 8 | 8 | | 56845 | Cytochrome c oxidase subunit 1 OS=Rattus norvegicus OX=10116 GN=Mtco1 PE=2 SV=3 |
| 48 | 234 | tr\|Q95938 | 109.06 | 11 | 11 | 1.40E+06 | 8 | 8 | 8 | | 56977 | Cytochrome c oxidase subunit 1 OS=Rattus norvegicus OX=10116 GN=Co I PE=3 SV=1 |
| 77 | 137 | tr\|D4A7L4 | 107.65 | 9 | 9 | 6.32E+05 | 3 | 3 | 4 | | 17634 | NADH dehydrogenase (Ubiquinone) 1 beta subcomplex 11 (Predicted) OS=Rattus norvegicus OX=10116 GN=Ndufb11 PE=1 SV=1 |
| 66 | 1617 | Q5RJR8\|LF | 105.06 | 7 | 7 | 5.29E+05 | 4 | 4 | 4 | | 34869 | Leucine-rich repeat-containing protein 59 OS=Rattus norvegicus OX=10116 GN=Lrrc59 PE=1 SV=1 |
| 53 | 1095 | tr\|Q5UAJ5 | 100.41 | 30 | 30 | 7.49E+06 | 4 | 4 | 7 | Oxidation ( | 7642 | ATP synthase protein 8 OS=Rattus norvegicus OX=10116 GN=ATP8 PE=3 SV=1 |
| 53 | 1097 | tr\|Q8SEZ4 | 100.41 | 30 | 30 | 7.49E+06 | 4 | 4 | 7 | Oxidation ( | 7632 | ATP synthase protein 8 OS=Rattus norvegicus OX=10116 GN=ATPase8 PE=3 SV=1 |
| 64 | 240 | A0A0G2K0 | 98.95 | 5 | 5 | 5.54E+05 | 4 | 4 | 4 |  | 74675 | Acyl-CoA synthetase short-chain family member 3 mitochondrial OS=Rattus norvegicus OX=10116 GN=Acss3 PE=1 SV=1 |
| 59 | 19 | Q641Y2\|N | 97.98 | 10 | 10 | 9.23E+06 | 5 | 5 | 5 | | 52562 | NADH dehydrogenase [ubiquinone] iron-sulfur protein 2 mitochondrial OS=Rattus norvegicus OX=10116 GN=Ndufs2 PE=1 SV=1 |
| 51 | 1322 | P32198\|CF | 97.71 | 6 | 6 | 1.61E+06 | 5 | 5 | 7 | | 88126 | Carnitine O-palmitoyltransferase 1 liver isoform OS=Rattus norvegicus OX=10116 GN=Cpt1a PE=1 SV=2 |
| 52 | 247 | P10888\|CC | 95.59 | 17 | 17 | 3.91E+06 | 6 | 6 | 7 | | 19515 | Cytochrome c oxidase subunit 4 isoform 1 mitochondrial OS=Rattus norvegicus OX=10116 GN=Cox4i1 PE=1 SV=1 |
| 67 | 59 | P17178\|CF | 95.03 | 10 | 10 | 8.82E+04 | 4 | 4 | 4 | | 60733 | Sterol 26-hydroxylase mitochondrial OS=Rattus norvegicus OX=10116 GN=Cyp27a1 PE=1 SV=1 |
| 67 | 60 | tr\|A0A0H2 | 95.03 | 10 | 10 | 8.82E+04 | 4 | 4 | 4 | | 62492 | RCG24013 isoform CRA_a OS=Rattus norvegicus OX=10116 GN=Cyp27a1 PE=1 SV=1 |
| 69 | 725 | tr\|F1LP30\| | 89.44 | 3 | 3 | 1.06E+05 | 3 | 1 | 4 | | 79296 | Methylcrotonoyl-CoA carboxylase subunit alpha mitochondrial OS=Rattus norvegicus OX=10116 GN=Mccc1 PE=1 SV=1 |
| 69 | 726 | Q5I0C3\|M | 89.44 | 3 | 3 | 1.06E+05 | 3 | 1 | 4 | | 79330 | Methylcrotonoyl-CoA carboxylase subunit alpha mitochondrial OS=Rattus norvegicus OX=10116 GN=Mccc1 PE=1 SV=1 |
| 44 | 322 | P12075\|CC | 89.05 | 22 | 22 | 1.26E+07 | 7 | 7 | 11 | | 13915 | Cytochrome c oxidase subunit 5B mitochondrial OS=Rattus norvegicus OX=10116 GN=Cox5b PE=1 SV=2 |
| 55 | 207 | tr\|G3V6I4\| | 87.31 | 5 | 5 | 8.68E+05 | 3 | 3 | 6 | | 37791 | Mitochondrial amidoxime reducing component 1 OS=Rattus norvegicus OX=10116 GN=Marc1 PE=1 SV=1 |
| 58 | 14 | tr\|F1LN92 | 84.72 | 5 | 5 | 5.08E+06 | 4 | 4 | 5 | | 89352 | AFG3-like matrix AAA peptidase subunit 2 OS=Rattus norvegicus OX=10116 GN=Afg3l2 PE=1 SV=1 |
| 60 | 190 | P07824\|AF | 84.56 | 13 | 13 | 1.24E+06 | 4 | 3 | 5 | | 34973 | Arginase-1 OS=Rattus norvegicus OX=10116 GN=Arg1 PE=1 SV=2 |
| 46 | 134 | tr\|B2RYS2 | 84.22 | 37 | 37 | 5.04E+06 | 7 | 6 | 9 | Oxidation ( | 13559 | Cytochrome b-c1 complex subunit 7 OS=Rattus norvegicus OX=10116 GN=Uqcrb PE=1 SV=1 |
| 45 | 250 | P08011\|M | 84.15 | 23 | 23 | 2.06E+06 | 8 | 8 | 9 | | 17472 | Microsomal glutathione S-transferase 1 OS=Rattus norvegicus OX=10116 GN=Mgst1 PE=1 SV=3 |
| 45 | 251 | tr\|B6DYQ4 | 84.15 | 23 | 23 | 2.06E+06 | 8 | 8 | 9 | | 17472 | Microsomal glutathione S-transferase OS=Rattus norvegicus OX=10116 GN=Mgst1 PE=2 SV=1 |
| 62 | 244 | P09139\|SF | 83.27 | 7 | 7 | 8.30E+05 | 4 | 4 | 5 | | 45834 | Serine--pyruvate aminotransferase mitochondrial OS=Rattus norvegicus OX=10116 GN=Agxt PE=1 SV=1 |
| 88 | 15 | P67779\|PH | 79.81 | 10 | 10 | 5.27E+05 | 3 | 3 | 3 | | 29820 | Prohibitin OS=Rattus norvegicus OX=10116 GN=Phb PE=1 SV=1 |
| 70 | 139 | Q6AXV4\|S. | 78.98 | 10 | 10 | 1.47E+06 | 4 | 4 | 4 | | 51960 | Sorting and assembly machinery component 50 homolog OS=Rattus norvegicus OX=10116 GN=Samm50 PE=1 SV=1 |
| 91 | 298 | P11951\|CX | 77.94 | 24 | 24 | 8.29E+05 | 3 | 3 | 3 | | 8455 | Cytochrome c oxidase subunit 6C-2 OS=Rattus norvegicus OX=10116 GN=Cox6c2 PE=1 SV=3 |
| 128 | 1959 | tr\|A0A0G2 | 73.21 | 6 | 6 | 4.89E+05 | 2 | 2 | 2 | | 36057 | Mitochondrial calcium uniporter OS=Rattus norvegicus OX=10116 GN=Mcu PE=1 SV=1 |

| | | | | | | | | | | | | |
|---|---|---|---|---|---|---|---|---|---|---|---|---|
| 128 | 1960 | tr\|M0RDI5 | 73.21 | 6 | 6 | 4.89E+05 | 2 | 2 | 2 | | 39758 | Mitochondrial calcium uniporter OS=Rattus norvegicus OX=10116 GN=Mcu PE=1 SV=1 |
| 71 | 255 | tr\|I6V4L9\| | 71.43 | 7 | 7 | 2.54E+05 | 3 | 3 | 4 | | 29844 | Cytochrome c oxidase subunit 3 OS=Rattus norvegicus OX=10116 GN=COX3 PE=3 SV=1 |
| 71 | 388 | tr\|Q7H115 | 71.43 | 7 | 7 | 2.54E+05 | 3 | 3 | 4 | | 29871 | Cytochrome c oxidase subunit 3 OS=Rattus norvegicus OX=10116 GN=Mt-co3 PE=3 SV=1 |
| 71 | 389 | tr\|Q8M7G | 71.43 | 7 | 7 | 2.54E+05 | 3 | 3 | 4 | | 29861 | Cytochrome c oxidase subunit 3 OS=Rattus norvegicus OX=10116 PE=2 SV=1 |
| 71 | 390 | tr\|A0A096 | 71.43 | 7 | 7 | 2.54E+05 | 3 | 3 | 4 | | 29901 | Cytochrome c oxidase subunit 3 OS=Rattus norvegicus OX=10116 GN=COX3 PE=3 SV=1 |
| 71 | 391 | P05505\|C | 71.43 | 7 | 7 | 2.54E+05 | 3 | 3 | 4 | | 29871 | Cytochrome c oxidase subunit 3 OS=Rattus norvegicus OX=10116 GN=Mtco3 PE=1 SV=5 |
| 71 | 392 | tr\|Q8SEZ2 | 71.43 | 7 | 7 | 2.54E+05 | 3 | 3 | 4 | | 29870 | Cytochrome c oxidase subunit 3 (Fragment) OS=Rattus norvegicus OX=10116 GN=COIII PE=3 SV=1 |
| 75 | 140 | tr\|F1LN88 | 71.07 | 6 | 6 | 4.18E+05 | 3 | 2 | 4 | | 56517 | Aldehyde dehydrogenase mitochondrial OS=Rattus norvegicus OX=10116 GN=Aldh2 PE=1 SV=2 |
| 75 | 141 | P11884\|AL | 71.07 | 6 | 6 | 4.18E+05 | 3 | 2 | 4 | | 56488 | Aldehyde dehydrogenase mitochondrial OS=Rattus norvegicus OX=10116 GN=Aldh2 PE=1 SV=1 |
| 68 | 358 | P06761\|BI | 70.39 | 5 | 5 | 9.37E+05 | 4 | 4 | 4 | Formylatio | 72347 | Endoplasmic reticulum chaperone BiP OS=Rattus norvegicus OX=10116 GN=Hspa5 PE=1 SV=1 |
| 50 | 188 | O70351\|H | 69.72 | 17 | 17 | 1.34E+06 | 5 | 5 | 7 | | 27246 | 3-hydroxyacyl-CoA dehydrogenase type-2 OS=Rattus norvegicus OX=10116 GN=Hsd17b10 PE=1 SV=3 |
| 50 | 189 | tr\|B0BMW | 69.72 | 17 | 17 | 1.34E+06 | 5 | 5 | 7 | | 27250 | 3-hydroxyacyl-CoA dehydrogenase type-2 OS=Rattus norvegicus OX=10116 GN=Hsd17b10 PE=1 SV=1 |
| 95 | 5817 | P21571\|AT | 68.32 | 19 | 19 | 1.04E+06 | 2 | 2 | 3 | | 12494 | ATP synthase-coupling factor 6 mitochondrial OS=Rattus norvegicus OX=10116 GN=Atp5pf PE=1 SV=1 |
| 85 | 28 | tr\|D3ZG43 | 67.43 | 9 | 9 | 1.08E+06 | 3 | 3 | 3 | | 30226 | NADH dehydrogenase (Ubiquinone) Fe-S protein 3 (Predicted) isoform CRA_c OS=Rattus norvegicus OX=10116 GN=Ndufs3 PE=1 SV=1 |
| 56 | 286 | Q7TQ16\|Q | 65.99 | 35 | 35 | 1.27E+06 | 4 | 4 | 6 | | 9849 | Cytochrome b-c1 complex subunit 8 OS=Rattus norvegicus OX=10116 GN=Uqcrq PE=3 SV=1 |
| 106 | 342 | tr\|Q2TVU3 | 65.97 | 3 | 3 | 2.76E+05 | 2 | 2 | 2 | | 49412 | TID1 OS=Rattus norvegicus OX=10116 GN=Dnaja3 PE=2 SV=1 |
| 106 | 343 | tr\|A0A0G2 | 65.97 | 3 | 3 | 2.76E+05 | 2 | 2 | 2 | | 49416 | DnaJ heat shock protein family (Hsp40) member A3 OS=Rattus norvegicus OX=10116 GN=Dnaja3 PE=1 SV=1 |
| 106 | 344 | tr\|Q2UZS7 | 65.97 | 3 | 3 | 2.76E+05 | 2 | 2 | 2 | | 52399 | Tid-1 long isoform OS=Rattus norvegicus OX=10116 GN=Dnaja3 PE=2 SV=1 |
| 106 | 345 | tr\|A0A0G2 | 65.97 | 3 | 3 | 2.76E+05 | 2 | 2 | 2 | | 52403 | DnaJ heat shock protein family (Hsp40) member A3 OS=Rattus norvegicus OX=10116 GN=Dnaja3 PE=1 SV=1 |
| 106 | 341 | tr\|G3V6I5\| | 65.97 | 3 | 3 | 2.76E+05 | 2 | 2 | 2 | | 46777 | DnaJ heat shock protein family (Hsp40) member A3 OS=Rattus norvegicus OX=10116 GN=Dnaja3 PE=1 SV=2 |
| 72 | 283 | Q9WVK3\| | 65.08 | 9 | 9 | 1.92E+06 | 4 | 4 | 4 | Oxidation ( | 32433 | Peroxisomal trans-2-enoyl-CoA reductase OS=Rattus norvegicus OX=10116 GN=Pecr PE=2 SV=1 |
| 72 | 284 | tr\|A0A0G2 | 65.08 | 9 | 9 | 1.92E+06 | 4 | 4 | 4 | Oxidation ( | 32737 | Peroxisomal trans-2-enoyl-CoA reductase OS=Rattus norvegicus OX=10116 GN=Pecr PE=1 SV=1 |
| 81 | 48 | tr\|D3ZF13 | 65.04 | 16 | 16 | 2.15E+05 | 3 | 3 | 3 | | 17514 | Acyl carrier protein OS=Rattus norvegicus OX=10116 GN=Ndufab1 PE=1 SV=1 |
| 126 | 5788 | tr\|Q3B8N9 | 64.97 | 6 | 6 | 5.33E+05 | 2 | 2 | 2 | | 32823 | Biphenyl hydrolase-like OS=Rattus norvegicus OX=10116 GN=Bphl PE=1 SV=1 |
| 127 | 58 | tr\|D4A565 | 61.95 | 13 | 13 | 6.54E+05 | 2 | 2 | 2 | | 21664 | NADH dehydrogenase (Ubiquinone) 1 beta subcomplex 5 (Predicted) isoform CRA_b OS=Rattus norvegicus OX=10116 GN=Ndufb5 PE=1 SV=1 |
| 87 | 143 | tr\|Q5EBA4 | 59.59 | 7 | 7 | 1.09E+06 | 3 | 3 | 3 | | 33215 | Nipsnap1 protein (Fragment) OS=Rattus norvegicus OX=10116 GN=Nipsnap1 PE=2 SV=1 |
| 87 | 144 | tr\|G3V728 | 59.59 | 7 | 7 | 1.09E+06 | 3 | 3 | 3 | | 33346 | 4-nitrophenylphosphatase domain and non-neuronal SNAP25-like protein homolog 1 (C. elegans) isoform CRA_b OS=Rattus norvegicus OX=10116 GN=Nipsnap1 PE=1 SV=1 |
| 129 | 11921 | Q9Z2L0\|VD | 59.29 | 6 | 6 | 4.95E+05 | 2 | 2 | 2 | | 30756 | Voltage-dependent anion-selective channel protein 1 OS=Rattus norvegicus OX=10116 GN=Vdac1 PE=1 SV=4 |
| 96 | 300 | Q5PQT3\|G | 58.51 | 8 | 8 | 1.10E+05 | 3 | 3 | 3 | | 33899 | Glycine N-acyltransferase OS=Rattus norvegicus OX=10116 GN=Glyat PE=2 SV=1 |
| 96 | 301 | tr\|A4PB92 | 58.51 | 8 | 8 | 1.10E+05 | 3 | 3 | 3 | | 33899 | Glycine N-acyltransferase OS=Rattus norvegicus OX=10116 GN=Glyat PE=2 SV=1 |
| 96 | 302 | tr\|A0A0G2 | 58.51 | 7 | 7 | 1.10E+05 | 3 | 3 | 3 | | 39453 | Glycine N-acyltransferase OS=Rattus norvegicus OX=10116 GN=Glyat PE=1 SV=1 |
| 90 | 446 | Q91XJ1\|BE | 58.34 | 3 | 3 | | 3 | 0 | 3 | | 51557 | Beclin-1 OS=Rattus norvegicus OX=10116 GN=Becn1 PE=1 SV=1 |
| 108 | 66 | tr\|D4A0T0 | 58.03 | 16 | 16 | 1.07E+06 | 2 | 2 | 2 | | 20859 | NADH:ubiquinone oxidoreductase subunit B10 OS=Rattus norvegicus OX=10116 GN=Ndufb10 PE=1 SV=1 |
| 224 | 6486 | tr\|D3ZBM3 | 57.69 | 3 | 3 | 2.83E+05 | 1 | 1 | 1 | | 47552 | Ferrochelatase OS=Rattus norvegicus OX=10116 GN=Fech PE=1 SV=1 |
| 115 | 426 | tr\|A0A0G2 | 56.31 | 2 | 2 | 1.98E+05 | 2 | 1 | 2 | | 107334 | NLR family member X1 OS=Rattus norvegicus OX=10116 GN=Nlrx1 PE=1 SV=1 |
| 115 | 427 | Q5FVQ8\|N | 56.31 | 2 | 2 | 1.98E+05 | 2 | 1 | 2 | | 107590 | NLR family member X1 OS=Rattus norvegicus OX=10116 GN=Nlrx1 PE=2 SV=1 |
| 63 | 11740 | P29419\|AT | 54.82 | 27 | 27 | 2.02E+06 | 2 | 2 | 5 | | 8255 | ATP synthase subunit e mitochondrial OS=Rattus norvegicus OX=10116 GN=Atp5me PE=1 SV=3 |
| 116 | 5782 | Q09073\|A | 53.71 | 5 | 5 | 2.20E+06 | 2 | 2 | 2 | | 32901 | ADP/ATP translocase 2 OS=Rattus norvegicus OX=10116 GN=Slc25a5 PE=1 SV=3 |
| 132 | 253 | Q9ER34\|A | 53.25 | 4 | 4 | 5.07E+05 | 2 | 2 | 2 | | 85433 | Aconitate hydratase mitochondrial OS=Rattus norvegicus OX=10116 GN=Aco2 PE=1 SV=2 |
| 269 | 309 | P22071\|3B | 52.58 | 3 | 3 | 4.39E+05 | 1 | 1 | 1 | | 42037 | 3 beta-hydroxysteroid dehydrogenase/Delta 5-->4-isomerase type 1 OS=Rattus norvegicus OX=10116 GN=Hsd3b1 PE=1 SV=3 |
| 269 | 310 | P22072\|3B | 52.58 | 3 | 3 | 4.39E+05 | 1 | 1 | 1 | | 42277 | 3 beta-hydroxysteroid dehydrogenase/Delta 5-->4-isomerase type 2 OS=Rattus norvegicus OX=10116 GN=Hsd3b PE=2 SV=3 |
| 269 | 311 | P27364\|3B | 52.58 | 3 | 3 | 4.39E+05 | 1 | 1 | 1 | | 42206 | NADPH-dependent 3-keto-steroid reductase Hsd3b5 OS=Rattus norvegicus OX=10116 GN=Hsd3b5 PE=1 SV=3 |
| 135 | 321 | tr\|Q5BJZ3\| | 52.15 | 2 | 2 | 2.36E+05 | 2 | 2 | 2 | | 113869 | Nicotinamide nucleotide transhydrogenase OS=Rattus norvegicus OX=10116 GN=Nnt PE=1 SV=1 |
| 133 | 49 | tr\|B2RYS8 | 51.68 | 13 | 13 | 6.36E+05 | 2 | 2 | 2 | | 21959 | NADH dehydrogenase [ubiquinone] 1 beta subcomplex subunit 8 mitochondrial OS=Rattus norvegicus OX=10116 GN=Ndufb8 PE=2 SV=1 |
| 134 | 191 | P02770\|AL | 51.19 | 3 | 3 | 4.76E+05 | 2 | 2 | 2 | Formylatio | 68731 | Serum albumin OS=Rattus norvegicus OX=10116 GN=Alb PE=1 SV=2 |
| 134 | 192 | tr\|A0A0G2 | 51.19 | 3 | 3 | 4.76E+05 | 2 | 2 | 2 | Formylatio | 68759 | Serum albumin OS=Rattus norvegicus OX=10116 GN=Alb PE=1 SV=1 |
| 219 | 34 | tr\|A0A0G2 | 50.91 | 7 | 7 | 2.95E+05 | 1 | 1 | 1 | | 19965 | NADH dehydrogenase [ubiquinone] 1 alpha subcomplex subunit 8 OS=Rattus norvegicus OX=10116 GN=Ndufa8 PE=1 SV=1 |
| 131 | 210 | tr\|Q5M94 | 50.51 | 6 | 6 | 3.95E+05 | 2 | 2 | 2 | | 28340 | Nipsnap homolog 3A (C. elegans) OS=Rattus norvegicus OX=10116 GN=Nipsnap3b PE=1 SV=1 |
| 65 | 1178 | tr\|G3V9J8 | 49.34 | 5 | 5 | 2.41E+06 | 3 | 3 | 4 | Formylatio | 87521 | Glycerol-3-phosphate acyltransferase 1 mitochondrial OS=Rattus norvegicus OX=10116 GN=Gpam PE=1 SV=2 |
| 65 | 1179 | tr\|A0A0G2 | 49.34 | 4 | 4 | 2.41E+06 | 3 | 3 | 4 | Formylatio | 93728 | Glycerol-3-phosphate acyltransferase 1 mitochondrial OS=Rattus norvegicus OX=10116 GN=Gpam PE=1 SV=1 |
| 65 | 1180 | P97564\|GP | 49.34 | 4 | 4 | 2.41E+06 | 3 | 3 | 4 | Formylatio | 93715 | Glycerol-3-phosphate acyltransferase 1 mitochondrial OS=Rattus norvegicus OX=10116 GN=Gpam PE=1 SV=3 |
| 98 | 353 | Q9WUS0\| | 47.58 | 9 | 9 | 0 | 2 | 2 | 3 | Formylatio | 25203 | Adenylate kinase 4 mitochondrial OS=Rattus norvegicus OX=10116 GN=Ak4 PE=2 SV=1 |
| 139 | 208 | P13803\|ET | 46.94 | 3 | 3 | 4.93E+05 | 1 | 1 | 2 | | 34951 | Electron transfer flavoprotein subunit alpha mitochondrial OS=Rattus norvegicus OX=10116 GN=Etfa PE=1 SV=4 |
| 73 | 297 | Q924S5\|LO | 46.64 | 2 | 2 | 8.40E+05 | 2 | 2 | 4 | | 105792 | Lon protease homolog mitochondrial OS=Rattus norvegicus OX=10116 GN=Lonp1 PE=2 SV=1 |
| 80 | 7 | Q5BK63\|N | 46.48 | 5 | 5 | 1.30E+05 | 3 | 3 | 3 | | 42559 | NADH dehydrogenase [ubiquinone] 1 alpha subcomplex subunit 9 mitochondrial OS=Rattus norvegicus OX=10116 GN=Ndufa9 PE=1 SV=2 |
| 130 | 281 | tr\|F1LPV8\| | 45.62 | 5 | 5 | 5.20E+05 | 2 | 2 | 2 | | 46639 | Succinate--CoA ligase [GDP-forming] subunit beta mitochondrial OS=Rattus norvegicus OX=10116 GN=Suclg2 PE=1 SV=2 |
| 130 | 282 | tr\|B1H270 | 45.62 | 5 | 5 | 5.20E+05 | 2 | 2 | 2 | | 46988 | Succinate--CoA ligase [GDP-forming] subunit beta mitochondrial OS=Rattus norvegicus OX=10116 GN=Suclg2 PE=2 SV=1 |
| 140 | 11743 | P35171\|CX | 45.36 | 11 | 11 | 1.09E+05 | 1 | 1 | 2 | | 9353 | Cytochrome c oxidase subunit 7A2 mitochondrial OS=Rattus norvegicus OX=10116 GN=Cox7a2 PE=1 SV=1 |
| 140 | 11744 | tr\|B2RYS0 | 45.36 | 11 | 11 | 1.09E+05 | 1 | 1 | 2 | | 9353 | Cox7a2 protein OS=Rattus norvegicus OX=10116 GN=Cox7a2 PE=2 SV=1 |
| 76 | 361 | P80433\|CO | 45.32 | 19 | 19 | 1.02E+06 | 2 | 2 | 4 | | 7672 | Cytochrome c oxidase subunit 8A mitochondrial OS=Rattus norvegicus OX=10116 GN=Cox8a PE=1 SV=3 |
| 270 | 371 | tr\|B5DFA3 | 44.08 | 2 | 2 | 2.82E+05 | 1 | 1 | 1 | | 65357 | Bucs1 protein OS=Rattus norvegicus OX=10116 GN=Acsm1 PE=2 SV=1 |
| 270 | 372 | tr\|F1M1W | 44.08 | 2 | 2 | 2.82E+05 | 1 | 1 | 1 | | 65347 | Acyl-coenzyme A synthetase ACSM2 mitochondrial OS=Rattus norvegicus OX=10116 GN=Acsm1 PE=1 SV=3 |
| 270 | 373 | tr\|A0A0G2 | 44.08 | 2 | 2 | 2.82E+05 | 1 | 1 | 1 | | 65319 | Acyl-coenzyme A synthetase ACSM2 mitochondrial OS=Rattus norvegicus OX=10116 GN=Acsm1 PE=1 SV=1 |

| | | | | | | | | | | | |
|---|---|---|---|---|---|---|---|---|---|---|---|
| 270 | 374 | tr\|A0A0G2 | 44.08 | 2 | 2 | 2.82E+05 | 1 | 1 | 1 | | 70943 Acyl-coenzyme A synthetase ACSM2 mitochondrial OS=Rattus norvegicus OX=10116 GN=Acsm1 PE=1 SV=1 |
| 86 | 1701 | Q6UPE0\|C | 42.06 | 5 | 5 | 3.30E+05 | 3 | 3 | 3 | | 66389 Choline dehydrogenase mitochondrial OS=Rattus norvegicus OX=10116 GN=Chdh PE=1 SV=1 |
| 271 | 5916 | Q58FK9\|KA | 41.57 | 3 | 3 | 3.97E+04 | 1 | 1 | 1 | | 51044 Kynurenine--oxoglutarate transaminase 3 OS=Rattus norvegicus OX=10116 GN=Kyat3 PE=2 SV=1 |
| 168 | 306 | Q2V057\|H | 41.1 | 2 | 2 | 2.31E+05 | 1 | 1 | 1 | | 51002 Hydroxyproline dehydrogenase OS=Rattus norvegicus OX=10116 GN=Prodh2 PE=2 SV=1 |
| 272 | 17 | tr\|A0A0G2 | 40.91 | 3 | 3 | 2.19E+05 | 1 | 1 | 1 | | 33168 Prohibitin OS=Rattus norvegicus OX=10116 GN=Phb2 PE=1 SV=1 |
| 272 | 12 | Q5XIH7\|PH | 40.91 | 3 | 3 | 2.19E+05 | 1 | 1 | 1 | | 33312 Prohibitin-2 OS=Rattus norvegicus OX=10116 GN=Phb2 PE=1 SV=1 |
| 89 | 369 | tr\|G3V734 | 40.26 | 4 | 4 | 9.82E+05 | 2 | 2 | 3 | | 36133 2 4-dienoyl CoA reductase 1 mitochondrial isoform CRA_a OS=Rattus norvegicus OX=10116 GN=Decr1 PE=1 SV=1 |
| 89 | 370 | Q64591\|D | 40.26 | 4 | 4 | 9.82E+05 | 2 | 2 | 3 | | 36133 2 4-dienoyl-CoA reductase mitochondrial OS=Rattus norvegicus OX=10116 GN=Decr1 PE=1 SV=2 |
| 220 | 1716 | P56522\|AD | 39.88 | 3 | 3 | 1.14E+05 | 1 | 1 | 1 | | 54363 NADPH:adrenodoxin oxidoreductase mitochondrial OS=Rattus norvegicus OX=10116 GN=Fdxr PE=1 SV=1 |
| 273 | 337 | P80432\|CC | 38.79 | 16 | 16 | 5.84E+05 | 1 | 1 | 1 | | 7375 Cytochrome c oxidase subunit 7C mitochondrial OS=Rattus norvegicus OX=10116 GN=Cox7c PE=1 SV=2 |
| 97 | 6375 | tr\|D3ZTP0 | 38.72 | 3 | 3 | 3.95E+05 | 2 | 2 | 3 | | 101806 10-formyltetrahydrofolate dehydrogenase OS=Rattus norvegicus OX=10116 GN=Aldh1l2 PE=1 SV=3 |
| 136 | 333 | tr\|F1LZW6 | 38.71 | 3 | 3 | 0 | 2 | 2 | 2 | | 54099 Solute carrier family 25 member 13 OS=Rattus norvegicus OX=10116 GN=Slc25a13 PE=1 SV=2 |
| 199 | 1268 | Q2TA68\|O | 38.16 | 1 | 1 | 0 | 1 | 1 | 1 | | 111307 Dynamin-like 120 kDa protein mitochondrial OS=Rattus norvegicus OX=10116 GN=Opa1 PE=1 SV=1 |
| 152 | 1896 | tr\|D3ZMG( | 37.74 | 1 | 1 | | 2 | 0 | 2 | | 112437 Serine/threonine-protein kinase OS=Rattus norvegicus OX=10116 GN=Ulk1 PE=1 SV=2 |
| 82 | 356 | O70600\|RS | 36.21 | 3 | 3 | 1.31E+06 | 1 | 1 | 3 | Formylatio | 41255 Radical S-adenosyl methionine domain-containing protein 2 OS=Rattus norvegicus OX=10116 GN=Rsad2 PE=1 SV=1 |
| 82 | 357 | tr\|A0A0H2 | 36.21 | 3 | 3 | 1.31E+06 | 1 | 1 | 3 | Formylatio | 41444 RCG62278 OS=Rattus norvegicus OX=10116 GN=Rsad2 PE=4 SV=1 |
| 275 | 349 | Q5I0C5\|FM | 35.73 | 2 | 2 | 0 | 1 | 1 | 1 | Formylatio | 42997 Methionyl-tRNA formyltransferase mitochondrial OS=Rattus norvegicus OX=10116 GN=Mtfmt PE=2 SV=1 |
| 110 | 6517 | P56574\|ID | 34.89 | 4 | 4 | 3.51E+05 | 2 | 2 | 2 | | 50967 Isocitrate dehydrogenase [NADP] mitochondrial OS=Rattus norvegicus OX=10116 GN=Idh2 PE=1 SV=2 |
| 274 | 193 | P08461\|OD | 34.74 | 2 | 2 | 2.03E+05 | 1 | 1 | 1 | | 67166 Dihydrolipoyllysine-residue acetyltransferase component of pyruvate dehydrogenase complex mitochondrial OS=Rattus norvegicus OX=10116 GN=Dlat PE=1 SV=3 |
| 283 | 590 | tr\|F1MAR6 | 34.69 | 1 | 1 | 0 | 1 | 1 | 1 | | 68095 Proline dehydrogenase OS=Rattus norvegicus OX=10116 GN=Prodh1 PE=1 SV=2 |
| 114 | 6174 | tr\|D3Z900 | 34.14 | 7 | 7 | 8.22E+04 | 2 | 1 | 2 | Oxidation ( | 38239 Mitochondrial amidoxime reducing component 2 OS=Rattus norvegicus OX=10116 GN=Marc2 PE=1 SV=2 |
| 114 | 6175 | O88994\|M | 34.14 | 7 | 7 | 8.22E+04 | 2 | 1 | 2 | Oxidation ( | 38249 Mitochondrial amidoxime reducing component 2 OS=Rattus norvegicus OX=10116 GN=Marc2 PE=2 SV=1 |
| 119 | 6119 | A0A0G2JZ7 | 34.04 | 3 | 3 | 1.44E+06 | 2 | 2 | 2 | | 62059 NAD-dependent protein deacetylase sirtuin-1 OS=Rattus norvegicus OX=10116 GN=Sirt1 PE=3 SV=2 |
| 119 | 6120 | tr\|A0A182 | 34.04 | 2 | 2 | 1.44E+06 | 2 | 2 | 2 | | 81778 NAD-dependent protein deacetylase sirtuin-1 OS=Rattus norvegicus OX=10116 GN=Sirt1 PE=4 SV=1 |
| 100 | 531 | tr\|D3ZHK4 | 33.55 | 1 | 1 | | 2 | 0 | 3 | | 182226 RB1-inducible coiled-coil 1 OS=Rattus norvegicus OX=10116 GN=Rb1cc1 PE=1 SV=1 |
| 120 | 656 | tr\|D3ZD23 | 32.62 | 1 | 1 | 5.46E+05 | 2 | 2 | 2 | | 67300 ATP-binding cassette subfamily E member 1 OS=Rattus norvegicus OX=10116 GN=Abce1 PE=1 SV=1 |
| 276 | 28531 | Q5I0I4\|DM | 31.92 | 6 | 6 | 1.73E+05 | 1 | 1 | 1 | | 28508 Distal membrane-arm assembly complex protein 2 OS=Rattus norvegicus OX=10116 GN=Dmac2 PE=2 SV=1 |
| 118 | 745 | F8WLE0\|K | 31.81 | 2 | 2 | 8.41E+05 | 2 | 1 | 2 | | 115542 Kinesin-like protein KIF28P OS=Rattus norvegicus OX=10116 GN=Kif28p PE=2 SV=1 |
| 99 | 514 | tr\|Q5RKL4 | 31.23 | 1 | 1 | 0 | 1 | 1 | 3 | | 95977 Dimethylglycine dehydrogenase OS=Rattus norvegicus OX=10116 GN=Dmgdh PE=1 SV=1 |
| 99 | 515 | Q63342\|M | 31.23 | 1 | 1 | 0 | 1 | 1 | 3 | | 96047 Dimethylglycine dehydrogenase mitochondrial OS=Rattus norvegicus OX=10116 GN=Dmgdh PE=1 SV=1 |
| 99 | 516 | tr\|A0A0G2 | 31.23 | 1 | 1 | 0 | 1 | 1 | 3 | | 98864 Dimethylglycine dehydrogenase mitochondrial OS=Rattus norvegicus OX=10116 GN=Dmgdh PE=1 SV=1 |
| 141 | 11767 | tr\|Q64592 | 31.04 | 3 | 3 | | 2 | 0 | 2 | | 32894 Delta3 delta2-enoyl-CoA isomerase OS=Rattus norvegicus OX=10116 PE=2 SV=1 |
| 124 | 1677 | Q6QI44\|NA | 30.89 | 1 | 1 | | 2 | 0 | 2 | | 111258 N-alpha-acetyltransferase 25 NatB auxiliary subunit OS=Rattus norvegicus OX=10116 GN=Naa25 PE=2 SV=1 |
| 79 | 688 | tr\|Q8M7G | 30.76 | 5 | 5 | 4.41E+05 | 1 | 1 | 3 | Oxidation ( | 24240 ATP synthase subunit a (Fragment) OS=Rattus norvegicus OX=10116 PE=2 SV=1 |
| 79 | 689 | tr\|Q549I0\| | 30.76 | 5 | 5 | 4.41E+05 | 1 | 1 | 3 | Oxidation ( | 25050 ATP synthase subunit a OS=Rattus norvegicus OX=10116 GN=atp6 PE=4 SV=1 |
| 79 | 690 | P05504\|AT | 30.76 | 5 | 5 | 4.41E+05 | 1 | 1 | 3 | Oxidation ( | 25076 ATP synthase subunit a OS=Rattus norvegicus OX=10116 GN=Mt-atp6 PE=1 SV=3 |
| 79 | 691 | tr\|Q8HIC7 | 30.76 | 5 | 5 | 4.41E+05 | 1 | 1 | 3 | Oxidation ( | 25076 ATP synthase subunit a OS=Rattus norvegicus OX=10116 GN=Mt-atp6 PE=4 SV=1 |
| 79 | 692 | tr\|S5S1E9\| | 30.76 | 5 | 5 | 4.41E+05 | 1 | 1 | 3 | Oxidation ( | 25049 ATP synthase subunit a OS=Rattus norvegicus OX=10116 GN=ATP6 PE=4 SV=1 |
| 79 | 693 | tr\|Q06QE5 | 30.76 | 5 | 5 | 4.41E+05 | 1 | 1 | 3 | Oxidation ( | 25075 ATP synthase subunit a OS=Rattus norvegicus OX=10116 GN=ATP6 PE=4 SV=1 |
| 79 | 459 | tr\|Q8SEZ3 | 30.76 | 5 | 5 | 4.41E+05 | 1 | 1 | 3 | Oxidation ( | 25030 ATP synthase subunit a OS=Rattus norvegicus OX=10116 GN=ATPase6 PE=4 SV=1 |
| 221 | 11891 | Q64565\|A( | 30.03 | 2 | 2 | 1.03E+05 | 1 | 1 | 1 | | 57201 Alanine--glyoxylate aminotransferase 2 mitochondrial OS=Rattus norvegicus OX=10116 GN=Agxt2 PE=1 SV=2 |
| 277 | 1552 | O88813\|A( | 29.94 | 2 | 2 | 0 | 1 | 1 | 1 | | 76405 Long-chain-fatty-acid--CoA ligase 5 OS=Rattus norvegicus OX=10116 GN=Acsl5 PE=1 SV=1 |
| 103 | 555 | tr\|D4A1D3 | 29.88 | 0 | 0 | 2.01E+06 | 2 | 1 | 2 | | 521487 Sacsin molecular chaperone OS=Rattus norvegicus OX=10116 GN=Sacs PE=1 SV=2 |
| 146 | 482 | tr\|D3ZHF8 | 29.69 | 2 | 2 | | 2 | 0 | 2 | | 73207 Translation factor GUF1 mitochondrial OS=Rattus norvegicus OX=10116 GN=Guf1 PE=1 SV=1 |
| 142 | 23226 | Q6QBQ4\|F | 29.34 | 3 | 3 | | 2 | 0 | 2 | | 31646 Phospholipid scramblase 3 OS=Rattus norvegicus OX=10116 GN=Plscr3 PE=2 SV=1 |
| 83 | 432 | tr\|F1LSP2\| | 28.77 | 1 | 1 | 8.90E+04 | 2 | 1 | 3 | | 119190 Acyl-CoA dehydrogenase family member 10 OS=Rattus norvegicus OX=10116 GN=Acad10 PE=1 SV=2 |
| 278 | 12149 | tr\|A0A0G2 | 28.14 | 7 | 7 | 5.20E+05 | 1 | 1 | 1 | | 19554 NADH:ubiquinone oxidoreductase complex assembly factor 2 OS=Rattus norvegicus OX=10116 GN=Ndufaf2 PE=1 SV=1 |
| 279 | 204 | tr\|Q5PQZ9 | 28 | 9 | 9 | 7.31E+05 | 1 | 1 | 1 | | 14359 NADH dehydrogenase [ubiquinone] 1 subunit C2 OS=Rattus norvegicus OX=10116 GN=Ndufc2 PE=1 SV=1 |
| 92 | 1348 | tr\|D4AB75 | 27.92 | 2 | 2 | 5.86E+05 | 2 | 2 | 3 | Formylatio | 113160 Zinc finger protein 217 OS=Rattus norvegicus OX=10116 GN=Zfp217 PE=1 SV=1 |
| 222 | 5796 | B1WC61\|A | 27.74 | 2 | 2 | 1.52E+05 | 1 | 1 | 1 | | 68843 Complex I assembly factor ACAD9 mitochondrial OS=Rattus norvegicus OX=10116 GN=Acad9 PE=1 SV=1 |
| 226 | 355 | tr\|D4AE90 | 26.79 | 2 | 2 | 4.59E+05 | 1 | 1 | 1 | Formylatio | 50113 RCC1-like OS=Rattus norvegicus OX=10116 GN=Rcc1l PE=1 SV=1 |
| 280 | 28532 | tr\|A0A0G2 | 26.34 | 16 | 16 | 8.70E+04 | 1 | 1 | 1 | | 9388 10 kDa heat shock protein mitochondrial OS=Rattus norvegicus OX=10116 GN=Hspe1 PE=1 SV=1 |
| 280 | 28533 | P26772\|CH | 26.34 | 14 | 14 | 8.70E+04 | 1 | 1 | 1 | | 10902 10 kDa heat shock protein mitochondrial OS=Rattus norvegicus OX=10116 GN=Hspe1 PE=1 SV=3 |
| 94 | 393 | Q63563\|AI | 26.24 | 1 | 1 | 1.90E+06 | 2 | 2 | 3 | | 174117 ATP-binding cassette sub-family C member 9 OS=Rattus norvegicus OX=10116 GN=Abcc9 PE=1 SV=1 |
| 84 | 1248 | Q80ZG6\|B | 25.76 | 6 | 6 | 1.54E+07 | 2 | 2 | 3 | | 20718 Bcl-2-binding component 3 OS=Rattus norvegicus OX=10116 GN=Bbc3 PE=1 SV=1 |
| 282 | 513 | Q8K1M7\|E | 25.45 | 1 | 1 | 0 | 1 | 1 | 1 | | 88497 BMP/retinoic acid-inducible neural-specific protein 3 OS=Rattus norvegicus OX=10116 GN=Brinp3 PE=2 SV=1 |
| 281 | 290 | tr\|A0A0G2 | 25.44 | 4 | 4 | 7.73E+05 | 1 | 1 | 1 | | 20750 Mitochondrial ribosomal protein L11 OS=Rattus norvegicus OX=10116 GN=mrpl11 PE=1 SV=1 |
| 281 | 291 | Q5XIE3\|RM | 25.44 | 4 | 4 | 7.73E+05 | 1 | 1 | 1 | | 22420 39S ribosomal protein L11 mitochondrial OS=Rattus norvegicus OX=10116 GN=Mrpl11 PE=2 SV=1 |
| 145 | 1243 | D4A693\|AZ | 24.4 | 3 | 3 | 2.53E+05 | 2 | 2 | 2 | Formylatio | 49420 Antizyme inhibitor 2 OS=Rattus norvegicus OX=10116 GN=Azin2 PE=1 SV=2 |
| 227 | 1945 | P26284\|OD | 22.73 | 3 | 3 | 0 | 1 | 1 | 1 | | 43227 Pyruvate dehydrogenase E1 component subunit alpha somatic form mitochondrial OS=Rattus norvegicus OX=10116 GN=Pdha1 PE=1 SV=2 |
| 143 | 1374 | tr\|A0A0G2 | 22.06 | 3 | 3 | 1.27E+05 | 1 | 1 | 2 | | 53760 von Willebrand factor A domain-containing 8 OS=Rattus norvegicus OX=10116 GN=Vwa8 PE=1 SV=1 |
| 235 | 11768 | Q68FU7\|C( | 21.82 | 2 | 2 | 2.19E+04 | 1 | 1 | 1 | | 51496 Ubiquinone biosynthesis monooxygenase COQ6 mitochondrial OS=Rattus norvegicus OX=10116 GN=Coq6 PE=2 SV=1 |
| 232 | 1013 | tr\|D3ZJH9 | 21.59 | 1 | 1 | 1.62E+06 | 1 | 1 | 1 | Formylatio | 65352 Malic enzyme OS=Rattus norvegicus OX=10116 GN=Me2 PE=1 SV=1 |

| | | | | | | | | | | | | |
|---|---|---|---|---|---|---|---|---|---|---|---|---|
| 232 | 1014 | tr\|A0A0G2 | 21.59 | 1 | 1 | 1.62E+06 | 1 | 1 | 1 | Formylatio | 66310 | Malic enzyme OS=Rattus norvegicus OX=10116 GN=Me2 PE=1 SV=1 |
| 229 | 1100 | Q05961\|O | 21.5 | 3 | 3 | 0 | 1 | 1 | 1 | | 41666 | 2'-5'-oligoadenylate synthase 1A OS=Rattus norvegicus OX=10116 GN=Oas1a PE=2 SV=1 |
| 285 | 398 | P48037\|AN | 21.31 | 1 | 1 | 7.57E+05 | 1 | 1 | 1 | | 75754 | Annexin A6 OS=Rattus norvegicus OX=10116 GN=Anxa6 PE=1 SV=2 |
| 285 | 399 | tr\|Q6IMZ3 | 21.31 | 1 | 1 | 7.57E+05 | 1 | 1 | 1 | | 75756 | Annexin OS=Rattus norvegicus OX=10116 GN=Anxa6 PE=1 SV=1 |
| 198 | 5848 | tr\|Q5FVL8 | 21.22 | 1 | 1 | 2.30E+05 | 1 | 1 | 1 | Formylatio | 77394 | ATP-binding cassette subfamily B member 10 OS=Rattus norvegicus OX=10116 GN=Abcb10 PE=1 SV=1 |
| 225 | 351 | Q8VID1\|DI | 21.21 | 3 | 3 | 5.17E+04 | 1 | 1 | 1 | | 29822 | Dehydrogenase/reductase SDR family member 4 OS=Rattus norvegicus OX=10116 GN=Dhrs4 PE=2 SV=2 |
| 144 | 428 | P24473\|GS | 21.05 | 4 | 4 | 1.77E+05 | 1 | 1 | 2 | | 25493 | Glutathione S-transferase kappa 1 OS=Rattus norvegicus OX=10116 GN=Gstk1 PE=1 SV=3 |
| 144 | 429 | tr\|B6DYQ0 | 21.05 | 4 | 4 | 1.77E+05 | 1 | 1 | 2 | | 25493 | Glutathione S-transferase kappa OS=Rattus norvegicus OX=10116 GN=Gstk1 PE=2 SV=1 |
| 122 | 1214 | Q5RKH7\|S | 21.04 | 3 | 3 | 6.37E+05 | 1 | 1 | 2 | Carbamido | 41092 | Solute carrier family 35 member F6 OS=Rattus norvegicus OX=10116 GN=Slc35f6 PE=2 SV=1 |
| 228 | 324 | P11497\|AC | 20.95 | 0 | 0 | 0 | 1 | 1 | 1 | | 265191 | Acetyl-CoA carboxylase 1 OS=Rattus norvegicus OX=10116 GN=Acaca PE=1 SV=1 |
| 231 | 197 | tr\|M0R6J0 | 20.87 | 3 | 3 | 0 | 1 | 1 | 1 | | 38374 | Mitochondrial ribosomal protein L39 OS=Rattus norvegicus OX=10116 GN=Mrpl39 PE=1 SV=2 |
| 286 | 55 | tr\|B0BNE6 | 20.81 | 6 | 6 | 1.83E+05 | 1 | 1 | 1 | | 23970 | NADH dehydrogenase (Ubiquinone) Fe-S protein 8 (Predicted) isoform CRA_a OS=Rattus norvegicus OX=10116 GN=Ndufs8 PE=1 SV=1 |
| 233 | 6062 | P15304\|LII | 20.8 | 1 | 1 | 2.62E+05 | 1 | 1 | 1 | | 116812 | Hormone-sensitive lipase OS=Rattus norvegicus OX=10116 GN=Lipe PE=1 SV=3 |
| 181 | 491 | P46462\|TE | 20.65 | 1 | 1 | 1.96E+04 | 1 | 1 | 1 | Oxidation (M) | | Transitional endoplasmic reticulum ATPase |
