## Supplementary material for "Permeability transition pore-related changes in the proteome and channel activity of ATP synthase dimers and monomers": RLM SM-BM

| Protein Gro | Protein ID | Accession | -10lgP | Coverage ( | Coverage ( | IBAQ | #Peptides | #Unique | #Spec Sam | PTM | Avg. Mass | Description |
|---|---|---|---|---|---|---|---|---|---|---|---|---|
| 1 | 1 | P10860\|DH | 395.27 | 67 | 67 | 5.62E+08 | 242 | 238 | 526 | Carbamidomethylation | | Glutamate dehydrogenase 1, mitochondrial |
| 3 | 2 | tr\|B2GV15 | 278 | 49 | 49 | 6.29E+07 | 69 | 68 | 123 | Oxidation ( | 53274 | Dihydrolipoamide acetyltransferase component of pyruvate dehydrogenase complex OS=Rattus norvegicus OX=10116 GN=Dbt PE=1 SV=1 |
| 2 | 4 | P07756\|CP | 272.24 | 31 | 31 | 5.32E+07 | 94 | 94 | 127 | Oxidation ( | 164579 | Carbamoyl-phosphate synthase [ammonia] mitochondrial OS=Rattus norvegicus OX=10116 GN=Cps1 PE=1 SV=1 |
| 7 | 5 | tr\|G3V6P2 | 252.3 | 43 | 43 | 2.86E+07 | 43 | 43 | 63 | Oxidation (M) | | Dihydrolipoyllysine-residue succinyltransferase component of 2-oxoglutarate dehydrogenase complex, mitochondrial |
| 7 | 6 | Q01205\|O | 252.3 | 43 | 43 | 2.86E+07 | 43 | 43 | 63 | Oxidation (M) | | |
| 11 | 8 | P10719\|AT | 248.73 | 28 | 28 | 2.50E+07 | 31 | 31 | 56 | Oxidation ( | 56354 | ATP synthase subunit beta mitochondrial OS=Rattus norvegicus OX=10116 GN=Atp5f1b PE=1 SV=2 |
| 11 | 9 | tr\|G3V6D3 | 248.73 | 28 | 28 | 2.50E+07 | 31 | 31 | 56 | Oxidation ( | 56345 | ATP synthase subunit beta OS=Rattus norvegicus OX=10116 GN=Atp5f1b PE=1 SV=1 |
| 6 | 3 | P85834\|EF | 241.69 | 41 | 41 | 2.38E+07 | 44 | 44 | 65 | Carbamidomethylation | | Elongation factor Tu, mitochondrial |
| 10 | 13 | Q66HF1\|N | 223.38 | 24 | 24 | 1.94E+07 | 39 | 39 | 57 | Carbamido | 79412 | NADH-ubiquinone oxidoreductase 75 kDa subunit mitochondrial OS=Rattus norvegicus OX=10116 GN=Ndufs1 PE=1 SV=1 |
| 8 | 20 | P19234\|NI | 223.05 | 39 | 39 | 2.69E+07 | 30 | 30 | 62 | Formylatio | 27378 | NADH dehydrogenase [ubiquinone] flavoprotein 2 mitochondrial OS=Rattus norvegicus OX=10116 GN=Ndufv2 PE=1 SV=2 |
| 9 | 16 | Q68FY0\|Q | 222.01 | 35 | 35 | 2.54E+07 | 39 | 39 | 61 | Carbamido | 52849 | Cytochrome b-c1 complex subunit 1 mitochondrial OS=Rattus norvegicus OX=10116 GN=Uqcrc1 PE=1 SV=1 |
| 14 | 15 | P67779\|PH | 221.53 | 49 | 49 | 2.27E+07 | 31 | 31 | 49 | | 29820 | Prohibitin OS=Rattus norvegicus OX=10116 GN=Phb PE=1 SV=1 |
| 20 | 38 | tr\|D3ZFQ8 | 219.54 | 26 | 26 | 1.40E+07 | 19 | 19 | 29 | Oxidation ( | 35435 | Cytochrome c-1 OS=Rattus norvegicus OX=10116 GN=Cyc1 PE=1 SV=3 |
| 5 | 21 | P32551\|Q | 214.75 | 30 | 30 | 2.69E+07 | 36 | 36 | 67 | | 48396 | Cytochrome b-c1 complex subunit 2 mitochondrial OS=Rattus norvegicus OX=10116 GN=Uqcrc2 PE=1 SV=2 |
| 17 | 26 | P00507\|AA | 214.72 | 26 | 26 | 1.69E+07 | 31 | 31 | 40 | Oxidation ( | 47314 | Aspartate aminotransferase mitochondrial OS=Rattus norvegicus OX=10116 GN=Got2 PE=1 SV=2 |
| 19 | 32 | P07895\|SC | 213.38 | 41 | 41 | 1.13E+07 | 18 | 18 | 32 | | 24674 | Superoxide dismutase [Mn] mitochondrial OS=Rattus norvegicus OX=10116 GN=Sod2 PE=1 SV=2 |
| 16 | 14 | tr\|F1LN92 | 212.72 | 26 | 26 | 9.01E+06 | 34 | 23 | 43 | Oxidation ( | 89352 | AFG3-like matrix AAA peptidase subunit 2 OS=Rattus norvegicus OX=10116 GN=Afg3l2 PE=1 SV=1 |
| 4 | 10 | P15999\|AT | 211.86 | 29 | 29 | 2.71E+07 | 38 | 37 | 94 | Oxidation ( | 59754 | ATP synthase subunit alpha mitochondrial OS=Rattus norvegicus OX=10116 GN=Atp5f1a PE=1 SV=2 |
| 4 | 11 | tr\|F1LP05\| | 211.86 | 29 | 29 | 2.71E+07 | 38 | 37 | 94 | Oxidation ( | 59813 | ATP synthase subunit alpha OS=Rattus norvegicus OX=10116 GN=Atp5f1a PE=1 SV=1 |
| 12 | 7 | Q5BK63\|N | 209.89 | 36 | 36 | 9.30E+06 | 32 | 31 | 54 | Oxidation ( | 42559 | NADH dehydrogenase [ubiquinone] 1 alpha subcomplex subunit 9 mitochondrial OS=Rattus norvegicus OX=10116 GN=Ndufa9 PE=1 SV=2 |
| 15 | 12 | Q5XIH7\|PH | 205.46 | 39 | 39 | 1.60E+07 | 32 | 32 | 49 | Oxidation ( | 33312 | Prohibitin-2 OS=Rattus norvegicus OX=10116 GN=Phb2 PE=1 SV=1 |
| 13 | 18 | Q4FZT0\|ST | 200.69 | 37 | 37 | 1.62E+07 | 38 | 38 | 51 | Oxidation ( | 38414 | Stomatin-like protein 2 mitochondrial OS=Rattus norvegicus OX=10116 GN=Stoml2 PE=1 SV=1 |
| 18 | 19 | Q641Y2\|N | 192.95 | 29 | 29 | 1.19E+07 | 27 | 27 | 36 | Oxidation ( | 52562 | NADH dehydrogenase [ubiquinone] iron-sulfur protein 2 mitochondrial OS=Rattus norvegicus OX=10116 GN=Ndufs2 PE=1 SV=1 |
| 33 | 49 | tr\|B2RYS8 | 190.59 | 26 | 26 | 4.30E+06 | 10 | 10 | 15 | Oxidation ( | 21959 | NADH dehydrogenase [ubiquinone] 1 beta subcomplex subunit 8 mitochondrial OS=Rattus norvegicus OX=10116 GN=Ndufb8 PE=1 SV=1 |
| 24 | 48 | tr\|D3ZF13 | 185.4 | 30 | 30 | 6.65E+06 | 16 | 15 | 25 | Oxidation ( | 17514 | Acyl carrier protein OS=Rattus norvegicus OX=10116 GN=Ndufab1 PE=1 SV=1 |
| 26 | 42 | P20788\|UC | 177.16 | 24 | 24 | 9.41E+06 | 12 | 12 | 22 | Oxidation ( | 29446 | Cytochrome b-c1 complex subunit Rieske mitochondrial OS=Rattus norvegicus OX=10116 GN=Uqcrfs1 PE=1 SV=2 |
| 21 | 22 | tr\|Q68G44 | 174.89 | 26 | 26 | 6.34E+06 | 24 | 24 | 28 | | 56886 | 3-hydroxy-3-methylglutaryl coenzyme A synthase OS=Rattus norvegicus OX=10116 GN=Hmgcs2 PE=1 SV=1 |
| 21 | 23 | P22791\|HI | 174.89 | 26 | 26 | 6.34E+06 | 24 | 24 | 28 | | 56912 | Hydroxymethylglutaryl-CoA synthase mitochondrial OS=Rattus norvegicus OX=10116 GN=Hmgcs2 PE=1 SV=1 |
| 28 | 34 | tr\|A0A0G2 | 174.86 | 38 | 38 | 3.85E+06 | 14 | 14 | 21 | Carbamido | 19965 | NADH dehydrogenase [ubiquinone] 1 alpha subcomplex subunit 8 OS=Rattus norvegicus OX=10116 GN=Ndufa8 PE=1 SV=1 |
| 27 | 24 | Q64428\|EC | 173.49 | 16 | 16 | 6.06E+06 | 17 | 17 | 21 | Oxidation ( | 82665 | Trifunctional enzyme subunit alpha mitochondrial OS=Rattus norvegicus OX=10116 GN=Hadha PE=1 SV=2 |
| 37 | 44 | P11240\|CC | 170.12 | 40 | 40 | 4.84E+06 | 10 | 10 | 13 | | 16130 | Cytochrome c oxidase subunit 5A mitochondrial OS=Rattus norvegicus OX=10116 GN=Cox5a PE=1 SV=1 |
| 43 | 136 | P24329\|TH | 167.7 | 16 | 16 | 2.42E+06 | 8 | 8 | 10 | | 33407 | Thiosulfate sulfurtransferase OS=Rattus norvegicus OX=10116 GN=Tst PE=1 SV=3 |
| 23 | 30 | P52873\|PY | 157.16 | 14 | 14 | 3.91E+06 | 20 | 20 | 25 | | 129777 | Pyruvate carboxylase mitochondrial OS=Rattus norvegicus OX=10116 GN=Pc PE=1 SV=2 |
| 23 | 31 | tr\|A0A0G2 | 157.16 | 13 | 13 | 3.91E+06 | 20 | 20 | 25 | | 140005 | Pyruvate carboxylase OS=Rattus norvegicus OX=10116 GN=Pc PE=1 SV=1 |
| 22 | 28 | tr\|D3ZG43 | 153.34 | 37 | 37 | 7.92E+06 | 21 | 21 | 28 | | 30226 | NADH dehydrogenase (Ubiquinone) Fe-S protein 3 (Predicted) isoform CRA_c OS=Rattus norvegicus OX=10116 GN=Ndufs3 PE=1 SV=1 |
| 46 | 58 | tr\|D4A565 | 152.99 | 20 | 20 | 2.36E+06 | 8 | 8 | 10 | | 21664 | NADH dehydrogenase (Ubiquinone) 1 beta subcomplex 5 (Predicted) isoform CRA_b OS=Rattus norvegicus OX=10116 GN=Ndufb5 PE=1 SV=1 |
| 25 | 27 | Q60587\|EC | 148.52 | 24 | 24 | 4.00E+06 | 16 | 16 | 22 | | 51414 | Trifunctional enzyme subunit beta mitochondrial OS=Rattus norvegicus OX=10116 GN=Hadhb PE=1 SV=1 |
| 31 | 66 | tr\|D4A0T0 | 142.49 | 33 | 33 | 5.38E+06 | 11 | 11 | 17 | | 20859 | NADH:ubiquinone oxidoreductase subunit B10 OS=Rattus norvegicus OX=10116 GN=Ndufb10 PE=1 SV=1 |
| 41 | 138 | Q5XIF3\|ND | 139.6 | 17 | 17 | 3.37E+06 | 7 | 7 | 10 | | 19741 | NADH dehydrogenase [ubiquinone] iron-sulfur protein 4 mitochondrial OS=Rattus norvegicus OX=10116 GN=Ndufs4 PE=1 SV=1 |
| 55 | 119 | P31399\|AT | 139.27 | 30 | 30 | 1.37E+06 | 8 | 8 | 8 | | 18763 | ATP synthase subunit d mitochondrial OS=Rattus norvegicus OX=10116 GN=Atp5pd PE=1 SV=3 |
| 50 | 63 | tr\|Q6QI09 | 137.3 | 9 | 9 | 1.50E+06 | 9 | 9 | 9 | | 67721 | ATP synthase subunit gamma mitochondrial OS=Rattus norvegicus OX=10116 GN=Taf3 PE=1 SV=1 |
| 50 | 64 | P35435\|AT | 137.3 | 19 | 19 | 1.50E+06 | 9 | 9 | 9 | | 30191 | ATP synthase subunit gamma mitochondrial OS=Rattus norvegicus OX=10116 GN=Atp5f1c PE=1 SV=2 |
| 32 | 46 | P13086\|SU | 135.95 | 13 | 13 | 3.40E+06 | 9 | 9 | 15 | | 36148 | Succinate--CoA ligase [ADP/GDP-forming] subunit alpha mitochondrial OS=Rattus norvegicus OX=10116 GN=Suclg1 PE=2 SV=2 |
| 32 | 47 | tr\|A0A0H2 | 135.95 | 13 | 13 | 3.40E+06 | 9 | 9 | 15 | | 37560 | Succinate--CoA ligase [ADP/GDP-forming] subunit alpha mitochondrial OS=Rattus norvegicus OX=10116 GN=Suclg1 PE=1 SV=1 |
| 45 | 137 | tr\|D4A7L4 | 135.39 | 23 | 23 | 7.15E+06 | 7 | 7 | 10 | | 17634 | NADH dehydrogenase (Ubiquinone) 1 beta subcomplex 11 (Predicted) OS=Rattus norvegicus OX=10116 GN=Ndufb11 PE=1 SV=1 |
| 48 | 181 | Q80W89\|N | 135.19 | 18 | 18 | 2.86E+06 | 7 | 7 | 10 | | 14854 | NADH dehydrogenase [ubiquinone] 1 alpha subcomplex subunit 11 OS=Rattus norvegicus OX=10116 GN=Ndufa11 PE=2 SV=1 |
| 39 | 204 | tr\|Q5PQZ9 | 134.72 | 21 | 21 | 4.23E+06 | 9 | 9 | 12 | | 14359 | NADH dehydrogenase [ubiquinone] 1 subunit C2 OS=Rattus norvegicus OX=10116 GN=Ndufc2 PE=1 SV=1 |
| 29 | 43 | tr\|Q5XIH3 | 134.64 | 11 | 11 | 7.99E+06 | 10 | 10 | 18 | | 50731 | NADH dehydrogenase [ubiquinone] flavoprotein 1 mitochondrial OS=Rattus norvegicus OX=10116 GN=Ndufv1 PE=1 SV=1 |
| 30 | 35 | P17764\|TH | 133.75 | 19 | 19 | 5.63E+06 | 10 | 10 | 17 | | 44695 | Acetyl-CoA acetyltransferase mitochondrial OS=Rattus norvegicus OX=10116 GN=Acat1 PE=1 SV=1 |
| 36 | 67 | tr\|L0L4L8\| | 131.95 | 11 | 11 | 3.98E+06 | 10 | 10 | 13 | | 42470 | Cytochrome b (Fragment) OS=Rattus norvegicus OX=10116 GN=cytb PE=3 SV=1 |
| 36 | 68 | tr\|A0A140 | 131.95 | 11 | 11 | 3.98E+06 | 10 | 10 | 13 | | 42574 | Cytochrome b (Fragment) OS=Rattus norvegicus OX=10116 PE=3 SV=1 |
| 36 | 69 | tr\|A0A140 | 131.95 | 11 | 11 | 3.98E+06 | 10 | 10 | 13 | | 42588 | Cytochrome b (Fragment) OS=Rattus norvegicus OX=10116 PE=3 SV=1 |
| 36 | 70 | tr\|A0A140 | 131.95 | 11 | 11 | 3.98E+06 | 10 | 10 | 13 | | 42527 | Cytochrome b (Fragment) OS=Rattus norvegicus OX=10116 PE=3 SV=1 |
| 36 | 71 | tr\|A0A3Q8 | 131.95 | 11 | 11 | 3.98E+06 | 10 | 10 | 13 | | 42712 | Cytochrome b (Fragment) OS=Rattus norvegicus OX=10116 GN=Cytb PE=3 SV=1 |
| 36 | 72 | tr\|A0A3Q8 | 131.95 | 11 | 11 | 3.98E+06 | 10 | 10 | 13 | | 42622 | Cytochrome b (Fragment) OS=Rattus norvegicus OX=10116 GN=Cytb PE=3 SV=1 |
| 36 | 73 | tr\|A0A3S6 | 131.95 | 11 | 11 | 3.98E+06 | 10 | 10 | 13 | | 42702 | Cytochrome b (Fragment) OS=Rattus norvegicus OX=10116 GN=Cytb PE=3 SV=1 |
| 36 | 74 | tr\|A0A3S6 | 131.95 | 11 | 11 | 3.98E+06 | 10 | 10 | 13 | | 42698 | Cytochrome b (Fragment) OS=Rattus norvegicus OX=10116 GN=Cytb PE=3 SV=1 |
| 36 | 75 | tr\|A0A097 | 131.95 | 11 | 11 | 3.98E+06 | 10 | 10 | 13 | | 42865 | Cytochrome b OS=Rattus norvegicus OX=10116 GN=CYTB PE=3 SV=1 |
| 36 | 76 | tr\|A0A220 | 131.95 | 11 | 11 | 3.98E+06 | 10 | 10 | 13 | | 43018 | Cytochrome b OS=Rattus norvegicus OX=10116 PE=3 SV=1 |
| 36 | 77 | tr\|D6NSP2 | 131.95 | 11 | 11 | 3.98E+06 | 10 | 10 | 13 | | 42945 | Cytochrome b (Fragment) OS=Rattus norvegicus OX=10116 GN=cytb PE=3 SV=1 |
| 36 | 78 | tr\|D6NSS6 | 131.95 | 11 | 11 | 3.98E+06 | 10 | 10 | 13 | | 42993 | Cytochrome b (Fragment) OS=Rattus norvegicus OX=10116 GN=cytb PE=3 SV=1 |
| 36 | 79 | tr\|D6NSQ3 | 131.95 | 11 | 11 | 3.98E+06 | 10 | 10 | 13 | | 43012 | Cytochrome b (Fragment) OS=Rattus norvegicus OX=10116 GN=cytb PE=3 SV=1 |

| | | | | | | | | | | | | |
|---|---|---|---|---|---|---|---|---|---|---|---|---|
| 36 | 80 | tr\|D6NSR3 | 131.95 | 11 | 11 | 3.98E+06 | 10 | 10 | 13 | | 42986 | Cytochrome b (Fragment) OS=Rattus norvegicus OX=10116 GN=cytb PE=3 SV=1 |
| 36 | 81 | tr\|D6NSQ8 | 131.95 | 11 | 11 | 3.98E+06 | 10 | 10 | 13 | | 43016 | Cytochrome b (Fragment) OS=Rattus norvegicus OX=10116 GN=cytb PE=3 SV=1 |
| 36 | 82 | tr\|D6NSR8 | 131.95 | 11 | 11 | 3.98E+06 | 10 | 10 | 13 | | 42998 | Cytochrome b (Fragment) OS=Rattus norvegicus OX=10116 GN=cytb PE=3 SV=1 |
| 36 | 83 | tr\|F2Q6S6 | 131.95 | 11 | 11 | 3.98E+06 | 10 | 10 | 13 | | 42938 | Cytochrome b (Fragment) OS=Rattus norvegicus OX=10116 GN=cytb PE=3 SV=1 |
| 36 | 84 | tr\|A0A0A1 | 131.95 | 11 | 11 | 3.98E+06 | 10 | 10 | 13 | | 42989 | Cytochrome b OS=Rattus norvegicus OX=10116 GN=CYTB PE=3 SV=1 |
| 36 | 85 | tr\|D6NSS7 | 131.95 | 11 | 11 | 3.98E+06 | 10 | 10 | 13 | | 42948 | Cytochrome b (Fragment) OS=Rattus norvegicus OX=10116 GN=cytb PE=3 SV=1 |
| 36 | 86 | tr\|Q8SEY9 | 131.95 | 11 | 11 | 3.98E+06 | 10 | 10 | 13 | | 43015 | Cytochrome b OS=Rattus norvegicus OX=10116 GN=cytb PE=3 SV=1 |
| 36 | 87 | tr\|D6NSR7 | 131.95 | 11 | 11 | 3.98E+06 | 10 | 10 | 13 | | 42982 | Cytochrome b (Fragment) OS=Rattus norvegicus OX=10116 GN=cytb PE=3 SV=1 |
| 36 | 88 | tr\|A0A220 | 131.95 | 11 | 11 | 3.98E+06 | 10 | 10 | 13 | | 42968 | Cytochrome b OS=Rattus norvegicus OX=10116 PE=3 SV=1 |
| 36 | 89 | tr\|F2Q6S5 | 131.95 | 11 | 11 | 3.98E+06 | 10 | 10 | 13 | | 42952 | Cytochrome b (Fragment) OS=Rattus norvegicus OX=10116 GN=cytb PE=3 SV=1 |
| 36 | 90 | tr\|D6NSQ6 | 131.95 | 11 | 11 | 3.98E+06 | 10 | 10 | 13 | | 42952 | Cytochrome b (Fragment) OS=Rattus norvegicus OX=10116 GN=cytb PE=3 SV=1 |
| 36 | 91 | tr\|Q5UAI7 | 131.95 | 11 | 11 | 3.98E+06 | 10 | 10 | 13 | | 42998 | Cytochrome b OS=Rattus norvegicus OX=10116 GN=CYTB PE=3 SV=1 |
| 36 | 92 | tr\|A0A0S1 | 131.95 | 11 | 11 | 3.98E+06 | 10 | 10 | 13 | | 43002 | Cytochrome b OS=Rattus norvegicus OX=10116 GN=CYTB PE=3 SV=1 |
| 36 | 93 | tr\|Q8HIC4 | 131.95 | 11 | 11 | 3.98E+06 | 10 | 10 | 13 | | 43012 | Cytochrome b OS=Rattus norvegicus OX=10116 GN=Mt-cyb PE=3 SV=1 |
| 36 | 94 | tr\|A0A220 | 131.95 | 11 | 11 | 3.98E+06 | 10 | 10 | 13 | | 42968 | Cytochrome b OS=Rattus norvegicus OX=10116 PE=3 SV=1 |
| 36 | 95 | tr\|A0A0S1 | 131.95 | 11 | 11 | 3.98E+06 | 10 | 10 | 13 | | 43016 | Cytochrome b OS=Rattus norvegicus OX=10116 GN=CYTB PE=3 SV=1 |
| 36 | 96 | tr\|R9TKN1 | 131.95 | 11 | 11 | 3.98E+06 | 10 | 10 | 13 | | 43012 | Cytochrome b OS=Rattus norvegicus OX=10116 GN=CYTB PE=3 SV=1 |
| 36 | 97 | P00159\|CY | 131.95 | 11 | 11 | 3.98E+06 | 10 | 10 | 13 | | 43012 | Cytochrome b OS=Rattus norvegicus OX=10116 GN=Mt-Cyb PE=3 SV=3 |
| 36 | 98 | tr\|L0N311 | 131.95 | 11 | 11 | 3.98E+06 | 10 | 10 | 13 | | 43045 | Cytochrome b (Fragment) OS=Rattus norvegicus OX=10116 GN=cytb PE=3 SV=1 |
| 36 | 99 | tr\|A0A220 | 131.95 | 11 | 11 | 3.98E+06 | 10 | 10 | 13 | | 42970 | Cytochrome b OS=Rattus norvegicus OX=10116 PE=3 SV=1 |
| 36 | 100 | tr\|H2KXA0 | 131.95 | 11 | 11 | 3.98E+06 | 10 | 10 | 13 | | 42978 | Cytochrome b (Fragment) OS=Rattus norvegicus OX=10116 GN=cytb PE=3 SV=1 |
| 36 | 101 | tr\|A0A096 | 131.95 | 11 | 11 | 3.98E+06 | 10 | 10 | 13 | | 43042 | Cytochrome b (Fragment) OS=Rattus norvegicus OX=10116 PE=3 SV=1 |
| 36 | 102 | tr\|D6NSP5 | 131.95 | 11 | 11 | 3.98E+06 | 10 | 10 | 13 | | 43012 | Cytochrome b (Fragment) OS=Rattus norvegicus OX=10116 GN=cytb PE=3 SV=1 |
| 36 | 103 | tr\|F8QU37 | 131.95 | 11 | 11 | 3.98E+06 | 10 | 10 | 13 | | 42992 | Cytochrome b (Fragment) OS=Rattus norvegicus OX=10116 GN=cytb PE=3 SV=1 |
| 36 | 116 | tr\|D6NSQ0 | 131.95 | 11 | 11 | 3.98E+06 | 10 | 10 | 13 | | 42968 | Cytochrome b (Fragment) OS=Rattus norvegicus OX=10116 GN=cytb PE=3 SV=1 |
| 36 | 120 | tr\|A0A411 | 131.95 | 11 | 11 | 3.98E+06 | 10 | 10 | 13 | | 40144 | Cytochrome b (Fragment) OS=Rattus norvegicus OX=10116 GN=Cytb PE=3 SV=1 |
| 36 | 121 | tr\|A0A0U2 | 131.95 | 11 | 11 | 3.98E+06 | 10 | 10 | 13 | | 41037 | Cytochrome b (Fragment) OS=Rattus norvegicus OX=10116 PE=3 SV=1 |
| 36 | 122 | tr\|A0A411 | 131.95 | 11 | 11 | 3.98E+06 | 10 | 10 | 13 | | 41017 | Cytochrome b (Fragment) OS=Rattus norvegicus OX=10116 GN=Cytb PE=3 SV=1 |
| 36 | 123 | tr\|F8QU31 | 131.95 | 11 | 11 | 3.98E+06 | 10 | 10 | 13 | | 42181 | Cytochrome b (Fragment) OS=Rattus norvegicus OX=10116 GN=cytb PE=3 SV=1 |
| 36 | 124 | tr\|A0A411 | 131.95 | 11 | 11 | 3.98E+06 | 10 | 10 | 13 | | 42238 | Cytochrome b (Fragment) OS=Rattus norvegicus OX=10116 GN=Cytb PE=3 SV=1 |
| 36 | 125 | tr\|A0A385 | 131.95 | 11 | 11 | 3.98E+06 | 10 | 10 | 13 | | 42766 | Cytochrome b (Fragment) OS=Rattus norvegicus OX=10116 PE=3 SV=1 |
| 36 | 127 | tr\|D6NSP4 | 131.95 | 11 | 11 | 3.98E+06 | 10 | 10 | 13 | | 43016 | Cytochrome b (Fragment) OS=Rattus norvegicus OX=10116 GN=cytb PE=3 SV=1 |
| 36 | 128 | tr\|D6NSR2 | 131.95 | 11 | 11 | 3.98E+06 | 10 | 10 | 13 | | 43073 | Cytochrome b (Fragment) OS=Rattus norvegicus OX=10116 GN=cytb PE=3 SV=1 |
| 36 | 129 | tr\|D6NSR1 | 131.95 | 11 | 11 | 3.98E+06 | 10 | 10 | 13 | | 43002 | Cytochrome b (Fragment) OS=Rattus norvegicus OX=10116 GN=cytb PE=3 SV=1 |
| 36 | 132 | tr\|A0A0U2 | 131.95 | 11 | 11 | 3.98E+06 | 10 | 10 | 13 | | 40981 | Cytochrome b (Fragment) OS=Rattus norvegicus OX=10116 PE=3 SV=1 |
| 34 | 36 | tr\|B5DEY1 | 131.88 | 11 | 11 | 4.43E+04 | 12 | 1 | 14 | | 84183 | Afg3l1 protein (Fragment) OS=Rattus norvegicus OX=10116 GN=Afg3l1 PE=2 SV=1 |
| 34 | 37 | tr\|D3ZZ20 | 131.88 | 10 | 10 | 4.43E+04 | 12 | 1 | 14 | | 87203 | AFG3(ATPase family gene 3)-like 1 (S. cerevisiae) OS=Rattus norvegicus OX=10116 GN=Afg3l1 PE=1 SV=2 |
| 42 | 41 | tr\|D4A3V2 | 127.28 | 42 | 42 | 1.38E+06 | 8 | 8 | 10 | Oxidation ( | 15224 | NADH dehydrogenase [ubiquinone] 1 alpha subcomplex subunit 6 OS=Rattus norvegicus OX=10116 GN=Ndufa6 PE=1 SV=1 |
| 69 | 277 | tr\|B2RZ24 | 125.76 | 6 | 6 | 3.24E+05 | 5 | 5 | 5 | | 47388 | Succinate-CoA ligase subunit beta (Fragment) OS=Rattus norvegicus OX=10116 GN=Sucla2 PE=2 SV=1 |
| 69 | 278 | tr\|F1LM47 | 125.76 | 6 | 6 | 3.24E+05 | 5 | 5 | 5 | | 50306 | Succinate--CoA ligase [ADP-forming] subunit beta mitochondrial OS=Rattus norvegicus OX=10116 GN=Sucla2 PE=1 SV=1 |
| 38 | 55 | tr\|B0BNE6 | 125.04 | 16 | 16 | 1.86E+06 | 10 | 9 | 12 | | 23970 | NADH dehydrogenase (Ubiquinone) Fe-S protein 8 (Predicted) isoform CRA_a OS=Rattus norvegicus OX=10116 GN=Ndufs8 PE=1 SV=1 |
| 65 | 142 | tr\|D3ZCZ9 | 123.05 | 21 | 21 | 1.92E+06 | 5 | 5 | 6 | | 13040 | NADH dehydrogenase [ubiquinone] iron-sulfur protein 6 mitochondrial OS=Rattus norvegicus OX=10116 GN=LOC100912599 PE=1 SV=1 |
| 53 | 52 | tr\|M0RAM | 119.1 | 24 | 24 | 2.10E+06 | 8 | 8 | 8 | | 22155 | Glutathione peroxidase OS=Rattus norvegicus OX=10116 GN=Gpx1 PE=1 SV=1 |
| 53 | 53 | P04041\|GP | 119.1 | 24 | 24 | 2.10E+06 | 8 | 8 | 8 | | 22305 | Glutathione peroxidase 1 OS=Rattus norvegicus OX=10116 GN=Gpx1 PE=1 SV=4 |
| 44 | 45 | P04762\|CA | 117.9 | 9 | 9 | 1300500 | 9 | 9 | 10 | | 59757 | Catalase OS=Rattus norvegicus OX=10116 GN=Cat PE=1 SV=3 |
| 66 | 145 | tr\|F1LPG5 | 116.18 | 24 | 24 | 1.72E+06 | 5 | 5 | 6 | | 15064 | NADH:ubiquinone oxidoreductase subunit B4 OS=Rattus norvegicus OX=10116 GN=Ndufb4 PE=1 SV=1 |
| 58 | 61 | P29147\|BD | 112.91 | 13 | 13 | 7.99E+05 | 6 | 5 | 7 | | 38202 | D-beta-hydroxybutyrate dehydrogenase mitochondrial OS=Rattus norvegicus OX=10116 GN=Bdh1 PE=1 SV=2 |
| 58 | 62 | tr\|A0A0G2 | 112.91 | 12 | 12 | 7.99E+05 | 6 | 5 | 7 | | 38333 | 3-hydroxybutyrate dehydrogenase type 1 isoform CRA_a OS=Rattus norvegicus OX=10116 GN=Bdh1 PE=1 SV=1 |
| 63 | 109 | tr\|A0A140 | 112.35 | 8 | 8 | 1.03E+06 | 6 | 6 | 6 | | 67049 | MICOS complex subunit MIC60 OS=Rattus norvegicus OX=10116 GN=Immt PE=1 SV=1 |
| 63 | 110 | Q3KR86\|M | 112.35 | 8 | 8 | 1.03E+06 | 6 | 6 | 6 | | 67177 | MICOS complex subunit Mic60 (Fragment) OS=Rattus norvegicus OX=10116 GN=Immt PE=1 SV=1 |
| 63 | 115 | tr\|A0A0G2 | 112.35 | 6 | 6 | 1.03E+06 | 6 | 6 | 6 | | 86230 | MICOS complex subunit MIC60 OS=Rattus norvegicus OX=10116 GN=Immt PE=1 SV=1 |
| 56 | 182 | tr\|Q06QG6 | 107.66 | 5 | 5 | 1.23E+06 | 5 | 5 | 7 | | 68588 | NADH-ubiquinone oxidoreductase chain 5 OS=Rattus norvegicus OX=10116 GN=ND5 PE=3 SV=1 |
| 56 | 183 | tr\|Q06QK5 | 107.66 | 5 | 5 | 1.23E+06 | 5 | 5 | 7 | | 68606 | NADH-ubiquinone oxidoreductase chain 5 OS=Rattus norvegicus OX=10116 GN=ND5 PE=3 SV=1 |
| 56 | 184 | P11661\|NU | 107.66 | 5 | 5 | 1.23E+06 | 5 | 5 | 7 | | 68618 | NADH-ubiquinone oxidoreductase chain 5 OS=Rattus norvegicus OX=10116 GN=Mtnd5 PE=3 SV=3 |
| 56 | 185 | tr\|Q06QA1 | 107.66 | 5 | 5 | 1.23E+06 | 5 | 5 | 7 | | 68574 | NADH-ubiquinone oxidoreductase chain 5 OS=Rattus norvegicus OX=10116 GN=ND5 PE=3 SV=1 |
| 56 | 186 | tr\|Q8SEZ0 | 107.66 | 5 | 5 | 1.23E+06 | 5 | 5 | 7 | | 68618 | NADH-ubiquinone oxidoreductase chain 5 OS=Rattus norvegicus OX=10116 GN=Mt-nd5 PE=3 SV=1 |
| 56 | 187 | tr\|A0A096 | 107.66 | 5 | 5 | 1.23E+06 | 5 | 5 | 7 | | 68584 | NADH-ubiquinone oxidoreductase chain 5 OS=Rattus norvegicus OX=10116 GN=ND5 PE=3 SV=1 |
| 60 | 133 | Q63362\|N | 104.74 | 26 | 26 | 1.52E+06 | 5 | 5 | 7 | | 13412 | NADH dehydrogenase [ubiquinone] 1 alpha subcomplex subunit 5 OS=Rattus norvegicus OX=10116 GN=Ndufa5 PE=1 SV=3 |
| 57 | 59 | P17178\|CP | 100.49 | 13 | 13 | 6.57E+05 | 6 | 6 | 7 | Oxidation ( | 60733 | Sterol 26-hydroxylase mitochondrial OS=Rattus norvegicus OX=10116 GN=Cyp27a1 PE=1 SV=1 |
| 57 | 60 | tr\|A0A0H2 | 100.49 | 12 | 12 | 6.57E+05 | 6 | 6 | 7 | Oxidation ( | 62492 | RCG24013 isoform CRA_a OS=Rattus norvegicus OX=10116 GN=Cyp27a1 PE=1 SV=1 |
| 67 | 338 | Q5M9I5\|Q | 100.41 | 18 | 18 | 2.77E+06 | 4 | 4 | 6 | | 10424 | Cytochrome b-c1 complex subunit 6 mitochondrial OS=Rattus norvegicus OX=10116 GN=Uqcrh PE=3 SV=1 |
| 35 | 56 | tr\|Q6P6W | 99.17 | 19 | 19 | 4.70E+06 | 7 | 7 | 13 | | 31701 | NADH dehydrogenase [ubiquinone] 1 alpha subcomplex subunit 10 mitochondrial OS=Rattus norvegicus OX=10116 GN=Ndufa10 PE=2 SV=1 |
| 35 | 57 | Q561S0\|N | 99.17 | 14 | 14 | 4.70E+06 | 7 | 7 | 13 | | 40493 | NADH dehydrogenase [ubiquinone] 1 alpha subcomplex subunit 10 mitochondrial OS=Rattus norvegicus OX=10116 GN=Ndufa10 PE=1 SV=1 |

| | | | | | | | | | | | | |
|---|---|---|---|---|---|---|---|---|---|---|---|---|
| 72 | 139 | Q6AXV4\|S | 98.69 | 10 | 10 | 7.47E+05 | 4 | 4 | 5 | | 51960 | Sorting and assembly machinery component 50 homolog OS=Rattus norvegicus OX=10116 GN=Samm50 PE=1 SV=1 |
| 40 | 50 | Q02253\|M | 97.94 | 13 | 13 | 1061600 | 10 | 9 | 11 | Carbamidomethylatior | | Methylmalonate-semialdehyde dehydrogenase [acylating], mitochondrial |
| 40 | 51 | tr\|G3V7J0 | 97.94 | 13 | 13 | 1.06E+06 | 10 | 9 | 11 | Carbamidomethylatior | | Methylmalonate-semialdehyde dehydrogenase [acylating], mitochondrial |
| 88 | 257 | tr\|A9UMW | 95.87 | 28 | 28 | 7.65E+05 | 3 | 2 | 4 | Oxidation ( | 9255 | Ndufa3 protein (Fragment) OS=Rattus norvegicus OX=10116 GN=Ndufa3 PE=2 SV=1 |
| 88 | 258 | tr\|M0RB63 | 95.87 | 27 | 27 | 7.65E+05 | 3 | 2 | 4 | Oxidation ( | 9372 | RCG63041 OS=Rattus norvegicus OX=10116 GN=LOC684509 PE=4 SV=1 |
| 88 | 259 | tr\|A0A0G2 | 95.87 | 25 | 25 | 7.65E+05 | 3 | 2 | 4 | Oxidation ( | 10170 | NADH:ubiquinone oxidoreductase subunit A3 OS=Rattus norvegicus OX=10116 GN=Ndufa3 PE=1 SV=1 |
| 49 | 104 | tr\|Q5UAJ6 | 95 | 18 | 18 | 2.46E+06 | 8 | 8 | 9 | Oxidation ( | 25942 | Cytochrome c oxidase subunit 2 OS=Rattus norvegicus OX=10116 GN=COX2 PE=3 SV=1 |
| 49 | 111 | tr\|S5RZM8 | 95 | 18 | 18 | 2.46E+06 | 8 | 8 | 9 | Oxidation ( | 25958 | Cytochrome c oxidase subunit 2 OS=Rattus norvegicus OX=10116 GN=COX2 PE=3 SV=1 |
| 49 | 112 | P00406\|CC | 95 | 18 | 18 | 2.46E+06 | 8 | 8 | 9 | Oxidation ( | 25928 | Cytochrome c oxidase subunit 2 OS=Rattus norvegicus OX=10116 GN=Mtco2 PE=1 SV=3 |
| 49 | 113 | tr\|Q8SEZ5 | 95 | 18 | 18 | 2.46E+06 | 8 | 8 | 9 | Oxidation ( | 25928 | Cytochrome c oxidase subunit 2 OS=Rattus norvegicus OX=10116 GN=Mt-co2 PE=1 SV=1 |
| 49 | 114 | tr\|A0A097 | 95 | 18 | 18 | 2.46E+06 | 8 | 8 | 9 | Oxidation ( | 25894 | Cytochrome c oxidase subunit 2 OS=Rattus norvegicus OX=10116 GN=COX2 PE=3 SV=1 |
| 90 | 200 | Q5XI78\|OD | 89.24 | 3 | 3 | 3.53E+05 | 3 | 2 | 3 | | 116296 | 2-oxoglutarate dehydrogenase mitochondrial OS=Rattus norvegicus OX=10116 GN=Ogdh PE=1 SV=1 |
| 59 | 65 | tr\|D3ZE15 | 88.79 | 26 | 26 | 5.78E+05 | 6 | 6 | 7 | Oxidation ( | 16777 | NADH:ubiquinone oxidoreductase subunit A13 OS=Rattus norvegicus OX=10116 GN=Ndufa13 PE=1 SV=1 |
| 52 | 54 | P18163\|AC | 86.21 | 7 | 7 | 1.69E+06 | 6 | 6 | 8 | | 78179 | Long-chain-fatty-acid--CoA ligase 1 OS=Rattus norvegicus OX=10116 GN=Acsl1 PE=1 SV=1 |
| 47 | 134 | tr\|B2RYS2 | 83.94 | 39 | 39 | 2.62E+06 | 6 | 6 | 10 | Oxidation ( | 13559 | Cytochrome b-c1 complex subunit 7 OS=Rattus norvegicus OX=10116 GN=Uqcrb PE=1 SV=1 |
| 62 | 135 | tr\|D3ZZ21 | 80.06 | 34 | 34 | 1.70E+06 | 4 | 4 | 6 | | 15638 | NADH dehydrogenase (Ubiquinone) 1 beta subcomplex 6 (Predicted) OS=Rattus norvegicus OX=10116 GN=Ndufb6 PE=1 SV=1 |
| 95 | 210 | tr\|Q5M949 | 79.1 | 6 | 6 | 6.57E+05 | 3 | 3 | 3 | | 28340 | Nipsnap homolog 3A (C. elegans) OS=Rattus norvegicus OX=10116 GN=Nipsnap3b PE=1 SV=1 |
| 111 | 201 | P45953\|AC | 78.74 | 4 | 4 | 2.16E+04 | 2 | 2 | 2 | | 70749 | Very long-chain specific acyl-CoA dehydrogenase mitochondrial OS=Rattus norvegicus OX=10116 GN=Acadvl PE=1 SV=1 |
| 111 | 202 | tr\|Q5M9H | 78.74 | 4 | 4 | 2.16E+04 | 2 | 2 | 2 | | 70821 | Acyl-Coenzyme A dehydrogenase very long chain OS=Rattus norvegicus OX=10116 GN=Acadvl PE=1 SV=1 |
| 68 | 193 | P08461\|OD | 75.51 | 8 | 8 | 8.71E+05 | 4 | 4 | 5 | | 67166 | Dihydrolipoyllysine-residue acetyltransferase component of pyruvate dehydrogenase complex mitochondrial OS=Rattus norvegicus OX=10116 GN=Dlat PE=1 SV=3 |
| 79 | 140 | tr\|F1LN88 | 74.94 | 8 | 8 | 3.70E+05 | 4 | 3 | 4 | | 56517 | Aldehyde dehydrogenase mitochondrial OS=Rattus norvegicus OX=10116 GN=Aldh2 PE=1 SV=2 |
| 79 | 141 | P11884\|AL | 74.94 | 8 | 8 | 3.70E+05 | 4 | 3 | 4 | | 56488 | Aldehyde dehydrogenase mitochondrial OS=Rattus norvegicus OX=10116 GN=Aldh2 PE=1 SV=1 |
| 54 | 147 | P05508\|NU | 74.05 | 12 | 12 | 5.88E+05 | 6 | 5 | 8 | Oxidation (M) | | NADH-ubiquinone oxidoreductase chain 4 |
| 54 | 148 | tr\|D2E6K0 | 74.05 | 12 | 12 | 5.88E+05 | 6 | 5 | 8 | Oxidation (M) | | |
| 54 | 149 | tr\|A7XYB9 | 74.05 | 12 | 12 | 5.88E+05 | 6 | 5 | 8 | Oxidation (M) | | |
| 54 | 150 | tr\|Q06QE1 | 74.05 | 12 | 12 | 5.88E+05 | 6 | 5 | 8 | Oxidation (M) | | |
| 54 | 151 | tr\|Q06QA2 | 74.05 | 12 | 12 | 5.88E+05 | 6 | 5 | 8 | Oxidation (M) | | |
| 54 | 152 | tr\|Q7HKW | 74.05 | 12 | 12 | 5.88E+05 | 6 | 5 | 8 | Oxidation (M) | | |
| 54 | 153 | tr\|Q06QG7 | 74.05 | 12 | 12 | 5.88E+05 | 6 | 5 | 8 | Oxidation (M) | | |
| 54 | 154 | tr\|Q06Q89 | 74.05 | 12 | 12 | 5.88E+05 | 6 | 5 | 8 | Oxidation (M) | | |
| 54 | 155 | tr\|Q35737 | 74.05 | 12 | 12 | 5.88E+05 | 6 | 5 | 8 | Oxidation (M) | | |
| 54 | 156 | tr\|Q8HIC6 | 74.05 | 12 | 12 | 5.88E+05 | 6 | 5 | 8 | Oxidation (M) | | |
| 83 | 190 | P07824\|AR | 73.82 | 12 | 12 | 4.10E+06 | 3 | 3 | 4 | | 34973 | Arginase-1 OS=Rattus norvegicus OX=10116 GN=Arg1 PE=1 SV=2 |
| 85 | 247 | P10888\|CO | 73.24 | 15 | 15 | 1.05E+06 | 4 | 4 | 4 | | 19515 | Cytochrome c oxidase subunit 4 isoform 1 mitochondrial OS=Rattus norvegicus OX=10116 GN=Cox4i1 PE=1 SV=1 |
| 64 | 191 | P02770\|AL | 72.68 | 4 | 4 | 1.60E+06 | 5 | 5 | 6 | Formylatio | 68731 | Serum albumin OS=Rattus norvegicus OX=10116 GN=Alb PE=1 SV=2 |
| 64 | 192 | tr\|A0A0G2 | 72.68 | 4 | 4 | 1.60E+06 | 5 | 5 | 6 | Formylatio | 68759 | Serum albumin OS=Rattus norvegicus OX=10116 GN=Alb PE=1 SV=1 |
| 73 | 245 | tr\|A9UMV9 | 69.21 | 19 | 19 | 3.13E+06 | 2 | 2 | 5 | | 12500 | NADH:ubiquinone oxidoreductase subunit A7 OS=Rattus norvegicus OX=10116 GN=Ndufa7 PE=1 SV=1 |
| 101 | 224 | tr\|D3ZXF9 | 68.89 | 8 | 8 | 4.63E+05 | 3 | 3 | 3 | | 29441 | Mitochondrial ribosomal protein L12 OS=Rattus norvegicus OX=10116 GN=Mrpl12 PE=1 SV=1 |
| 86 | 249 | tr\|F1LXA0\| | 68.47 | 14 | 14 | 1.30E+06 | 2 | 2 | 4 | | 17178 | NADH dehydrogenase [ubiquinone] 1 alpha subcomplex subunit 12 OS=Rattus norvegicus OX=10116 GN=Ndufa12 PE=1 SV=2 |
| 51 | 246 | P19511\|AT | 68 | 9 | 9 | 2.38E+06 | 3 | 3 | 8 | | 28869 | ATP synthase F(0) complex subunit B1 mitochondrial OS=Rattus norvegicus OX=10116 GN=Atp5pb PE=1 SV=1 |
| 70 | 143 | tr\|Q5EBA4 | 65.79 | 13 | 13 | 7.54E+05 | 4 | 4 | 5 | | 33215 | Nipsnap1 protein (Fragment) OS=Rattus norvegicus OX=10116 GN=Nipsnap1 PE=2 SV=1 |
| 70 | 144 | tr\|G3V728 | 65.79 | 13 | 13 | 7.54E+05 | 4 | 4 | 5 | | 33346 | 4-nitrophenylphosphatase domain and non-neuronal SNAP25-like protein homolog 1 (C. elegans) isoform CRA_b OS=Rattus norvegicus OX=10116 GN=Nipsnap1 PE=1 SV=1 |
| 141 | 340 | tr\|B2RZ57 | 63.58 | 8 | 8 | 2.18E+05 | 2 | 2 | 2 | | 20651 | Mitochondrial ribosomal protein L18 OS=Rattus norvegicus OX=10116 GN=Mrpl18 PE=1 SV=1 |
| 117 | 265 | Q63750\|RI | 61.94 | 22 | 22 | 4.23E+05 | 2 | 2 | 2 | | 17050 | 39S ribosomal protein L23 mitochondrial OS=Rattus norvegicus OX=10116 GN=Mrpl23 PE=2 SV=1 |
| 106 | 346 | tr\|D3ZS58 | 61.42 | 11 | 11 | 6.60E+05 | 3 | 3 | 3 | | 10845 | NADH dehydrogenase [ubiquinone] 1 alpha subcomplex subunit 2 OS=Rattus norvegicus OX=10116 GN=Ndufa2 PE=1 SV=1 |
| 100 | 223 | P0C2C4\|RM | 60.76 | 8 | 8 | 3.58E+05 | 2 | 2 | 3 | | 29940 | 39S ribosomal protein L10 mitochondrial OS=Rattus norvegicus OX=10116 GN=Mrpl10 PE=1 SV=1 |
| 100 | 287 | tr\|D3ZX69 | 60.76 | 8 | 8 | 3.58E+05 | 2 | 2 | 3 | | 29466 | 39S ribosomal protein L10 mitochondrial OS=Rattus norvegicus OX=10116 GN=Mrpl10 PE=1 SV=1 |
| 71 | 168 | Q7TT47\|SP | 59.97 | 6 | 6 | 2.58E+05 | 4 | 4 | 5 | | 86103 | Paraplegin OS=Rattus norvegicus OX=10116 GN=Spg7 PE=2 SV=2 |
| 82 | 188 | O70351\|HC | 59.9 | 10 | 10 | 4.04E+05 | 3 | 3 | 4 | | 27246 | 3-hydroxyacyl-CoA dehydrogenase type-2 OS=Rattus norvegicus OX=10116 GN=Hsd17b10 PE=1 SV=3 |
| 82 | 189 | tr\|B0BMW | 59.9 | 10 | 10 | 4.04E+05 | 3 | 3 | 4 | | 27250 | 3-hydroxyacyl-CoA dehydrogenase type-2 OS=Rattus norvegicus OX=10116 GN=Hsd17b10 PE=1 SV=1 |
| 84 | 195 | P49432\|OD | 59.81 | 8 | 8 | 3.54E+05 | 3 | 3 | 4 | | 38982 | Pyruvate dehydrogenase E1 component subunit beta mitochondrial OS=Rattus norvegicus OX=10116 GN=Pdhb PE=1 SV=2 |
| 84 | 196 | tr\|A0A0G2 | 59.81 | 7 | 7 | 3.54E+05 | 3 | 3 | 4 | | 46193 | Pyruvate dehydrogenase E1 component subunit beta OS=Rattus norvegicus OX=10116 GN=Pdhb PE=1 SV=1 |
| 75 | 289 | tr\|Q5XFW4 | 59.8 | 14 | 14 | 3.67E+05 | 3 | 3 | 5 | | 20541 | Mitochondrial ribosomal protein L13 OS=Rattus norvegicus OX=10116 GN=Mrpl13 PE=1 SV=1 |
| 98 | 198 | tr\|B5DEL8 | 59.13 | 21 | 21 | 6.65E+05 | 3 | 3 | 3 | | 12700 | NADH dehydrogenase (Ubiquinone) Fe-S protein 5 OS=Rattus norvegicus OX=10116 GN=Ndufs5 PE=1 SV=1 |
| 98 | 199 | tr\|A0A0G2 | 59.13 | 21 | 21 | 6.65E+05 | 3 | 3 | 3 | | 12730 | Uncharacterized protein OS=Rattus norvegicus OX=10116 PE=4 SV=1 |
| 104 | 197 | tr\|M0R6J0 | 59.1 | 9 | 9 | 2.38E+05 | 3 | 3 | 3 | | 38374 | Mitochondrial ribosomal protein L39 OS=Rattus norvegicus OX=10116 GN=Mrpl39 PE=1 SV=2 |
| 114 | 208 | P13803\|ET | 58.36 | 7 | 7 | 3.81E+05 | 2 | 2 | 2 | | 34951 | Electron transfer flavoprotein subunit alpha mitochondrial OS=Rattus norvegicus OX=10116 GN=Etfa PE=1 SV=4 |
| 89 | 286 | Q7TQ16\|Q | 58.35 | 22 | 22 | 6.15E+05 | 4 | 4 | 4 | | 9849 | Cytochrome b-c1 complex subunit 8 OS=Rattus norvegicus OX=10116 GN=Uqcrq PE=3 SV=1 |
| 87 | 250 | P08011\|M | 58.31 | 15 | 15 | 4.68E+05 | 4 | 4 | 4 | | 17472 | Microsomal glutathione S-transferase 1 OS=Rattus norvegicus OX=10116 GN=Mgst1 PE=1 SV=3 |
| 87 | 251 | tr\|B6DYQ4 | 58.31 | 15 | 15 | 4.68E+05 | 4 | 4 | 4 | | 17472 | Microsomal glutathione S-transferase OS=Rattus norvegicus OX=10116 GN=Mgst1 PE=2 SV=1 |
| 99 | 222 | tr\|Q4G067 | 57.98 | 8 | 8 | 3.25E+05 | 3 | 3 | 3 | | 37440 | Mitochondrial ribosomal protein L44 OS=Rattus norvegicus OX=10116 GN=Mrpl44 PE=1 SV=1 |
| 80 | 146 | tr\|B0BMT9 | 57.6 | 6 | 6 | 7.85E+05 | 3 | 3 | 4 | | 50202 | Sqrdl protein OS=Rattus norvegicus OX=10116 GN=Sqor PE=1 SV=1 |
| 61 | 171 | tr\|D3ZFJ6\| | 57.43 | 12 | 12 | 7.62E+05 | 5 | 5 | 7 | | 60420 | Lactamase beta OS=Rattus norvegicus OX=10116 GN=Lactb PE=1 SV=1 |

| | | | | | | | | | | | |
|---|---|---|---|---|---|---|---|---|---|---|---|
| 259 | 337 | P80432\|CC | 57.02 | 21 | 21 | 2.26E+05 | 1 | 1 | 1 | | 7375 Cytochrome c oxidase subunit 7C mitochondrial OS=Rattus norvegicus OX=10116 GN=Cox7c PE=1 SV=2 |
| 138 | 264 | tr\|D3ZTW8 | 56.73 | 14 | 14 | 7.69E+05 | 2 | 2 | 2 | | 15823 Mitochondrial ribosomal protein L27 OS=Rattus norvegicus OX=10116 GN=Mrpl27 PE=1 SV=2 |
| 97 | 194 | tr\|Q5RJN0 | 56.57 | 11 | 11 | 6.70E+05 | 3 | 3 | 3 | Oxidation ( | 23945 NADH dehydrogenase (Ubiquinone) Fe-S protein 7 OS=Rattus norvegicus OX=10116 GN=Ndufs7 PE=1 SV=1 |
| 187 | 248 | tr\|D3ZUX5 | 55.22 | 5 | 5 | 2.41E+05 | 1 | 1 | 1 | | 26435 MICOS complex subunit OS=Rattus norvegicus OX=10116 GN=Chchd3 PE=1 SV=1 |
| 126 | 211 | P70584\|AC | 54.8 | 5 | 5 | 5.71E+05 | 2 | 2 | 2 | | 47824 Short/branched chain specific acyl-CoA dehydrogenase mitochondrial OS=Rattus norvegicus OX=10116 GN=Acadsb PE=1 SV=1 |
| 126 | 212 | tr\|A0A0A0 | 54.8 | 4 | 4 | 5.71E+05 | 2 | 2 | 2 | | 52043 Short/branched chain-specific acyl-CoA dehydrogenase mitochondrial OS=Rattus norvegicus OX=10116 GN=Acadsb PE=1 SV=2 |
| 77 | 216 | Q9R063\|PI | 53.78 | 10 | 10 | 1.06E+06 | 3 | 3 | 4 | | 22179 Peroxiredoxin-5 mitochondrial OS=Rattus norvegicus OX=10116 GN=Prdx5 PE=1 SV=1 |
| 77 | 217 | tr\|A0A0G2 | 53.78 | 10 | 10 | 1.06E+06 | 3 | 3 | 4 | | 22207 Peroxiredoxin OS=Rattus norvegicus OX=10116 GN=Prdx5 PE=1 SV=1 |
| 78 | 238 | tr\|B2RYT5 | 52.19 | 11 | 11 | 1.62E+06 | 3 | 3 | 4 | | 12651 Cox7a2l protein OS=Rattus norvegicus OX=10116 GN=Cox7a2l PE=2 SV=1 |
| 78 | 239 | tr\|D3ZYX8 | 52.19 | 10 | 10 | 1.62E+06 | 3 | 3 | 4 | | 13274 Cytochrome c oxidase subunit 7A2-like OS=Rattus norvegicus OX=10116 GN=Cox7a2l PE=1 SV=1 |
| 93 | 220 | P11960\|OI | 50.57 | 6 | 6 | 1.45E+05 | 3 | 2 | 3 | | 50165 2-oxoisovalerate dehydrogenase subunit alpha mitochondrial (Fragment) OS=Rattus norvegicus OX=10116 GN=Bckdha PE=1 SV=1 |
| 127 | 218 | P63039\|CH | 49.87 | 3 | 3 | 6.35E+05 | 2 | 2 | 2 | | 60956 60 kDa heat shock protein mitochondrial OS=Rattus norvegicus OX=10116 GN=Hspd1 PE=1 SV=1 |
| 127 | 219 | tr\|A0A482 | 49.87 | 3 | 3 | 6.35E+05 | 2 | 2 | 2 | | 60956 Hsp60 OS=Rattus norvegicus OX=10116 GN=Hspd1 PE=2 SV=1 |
| 74 | 283 | Q9WVK3\|I | 48.47 | 9 | 9 | 8.52E+05 | 4 | 4 | 5 | Oxidation ( | 32433 Peroxisomal trans-2-enoyl-CoA reductase OS=Rattus norvegicus OX=10116 GN=Pecr PE=2 SV=1 |
| 74 | 284 | tr\|A0A0G2 | 48.47 | 9 | 9 | 8.52E+05 | 4 | 4 | 5 | Oxidation ( | 32737 Peroxisomal trans-2-enoyl-CoA reductase OS=Rattus norvegicus OX=10116 GN=Pecr PE=1 SV=1 |
| 171 | 240 | A0A0G2K0 | 45.64 | 1 | 1 | 1.59E+05 | 1 | 1 | 1 | | 74675 Acyl-CoA synthetase short-chain family member 3 mitochondrial OS=Rattus norvegicus OX=10116 GN=Acss3 PE=1 SV=1 |
| 176 | 254 | P0CG51\|U | 45.49 | 4 | 4 | 0 | 1 | 1 | 1 | | 34369 Polyubiquitin-B OS=Rattus norvegicus OX=10116 GN=Ubb PE=1 SV=1 |
| 160 | 229 | tr\|Q06QK0 | 45.31 | 2 | 2 | 2.24E+05 | 1 | 1 | 1 | | 56880 Cytochrome c oxidase subunit 1 OS=Rattus norvegicus OX=10116 GN=CO1 PE=3 SV=1 |
| 160 | 230 | tr\|Q8SEZ6 | 45.31 | 2 | 2 | 2.24E+05 | 1 | 1 | 1 | | 56879 Cytochrome c oxidase subunit 1 OS=Rattus norvegicus OX=10116 GN=COI PE=3 SV=1 |
| 160 | 231 | tr\|A0A0A1 | 45.31 | 2 | 2 | 2.24E+05 | 1 | 1 | 1 | | 56937 Cytochrome c oxidase subunit 1 OS=Rattus norvegicus OX=10116 GN=COX1 PE=3 SV=1 |
| 160 | 232 | tr\|Q8HIC9 | 45.31 | 2 | 2 | 2.24E+05 | 1 | 1 | 1 | | 56845 Cytochrome c oxidase subunit 1 OS=Rattus norvegicus OX=10116 GN=Mt-co1 PE=3 SV=1 |
| 160 | 233 | P05503\|CC | 45.31 | 2 | 2 | 2.24E+05 | 1 | 1 | 1 | | 56845 Cytochrome c oxidase subunit 1 OS=Rattus norvegicus OX=10116 GN=Mtco1 PE=2 SV=3 |
| 160 | 234 | tr\|Q95938 | 45.31 | 2 | 2 | 2.24E+05 | 1 | 1 | 1 | | 56977 Cytochrome c oxidase subunit 1 OS=Rattus norvegicus OX=10116 GN=Co I PE=3 SV=1 |
| 160 | 241 | tr\|Q06QA9 | 45.31 | 2 | 2 | 2.24E+05 | 1 | 1 | 1 | | 56893 Cytochrome c oxidase subunit 1 OS=Rattus norvegicus OX=10116 GN=CO1 PE=3 SV=1 |
| 125 | 207 | tr\|G3V6I4\| | 44.65 | 5 | 5 | 4.00E+05 | 2 | 2 | 2 | | 37791 Mitochondrial amidoxime reducing component 1 OS=Rattus norvegicus OX=10116 GN=Marc1 PE=1 SV=1 |
| 260 | 341 | tr\|G3V6I5\| | 43.54 | 4 | 4 | 2.04E+05 | 1 | 1 | 1 | | 46777 DnaJ heat shock protein family (Hsp40) member A3 OS=Rattus norvegicus OX=10116 GN=Dnaja3 PE=1 SV=2 |
| 260 | 342 | tr\|Q2TVU3 | 43.54 | 4 | 4 | 2.04E+05 | 1 | 1 | 1 | | 49412 TID1 OS=Rattus norvegicus OX=10116 GN=Dnaja3 PE=2 SV=1 |
| 260 | 343 | tr\|A0A0G2 | 43.54 | 4 | 4 | 2.04E+05 | 1 | 1 | 1 | | 49416 DnaJ heat shock protein family (Hsp40) member A3 OS=Rattus norvegicus OX=10116 GN=Dnaja3 PE=1 SV=1 |
| 260 | 344 | tr\|Q2UZS7 | 43.54 | 3 | 3 | 2.04E+05 | 1 | 1 | 1 | | 52399 Tid-1 long isoform OS=Rattus norvegicus OX=10116 GN=Dnaja3 PE=2 SV=1 |
| 260 | 345 | tr\|A0A0G2 | 43.54 | 3 | 3 | 2.04E+05 | 1 | 1 | 1 | | 52403 DnaJ heat shock protein family (Hsp40) member A3 OS=Rattus norvegicus OX=10116 GN=Dnaja3 PE=1 SV=1 |
| 220 | 300 | Q5PQT3\|G | 42.89 | 4 | 4 | 1.05E+05 | 1 | 1 | 1 | | 33899 Glycine N-acyltransferase OS=Rattus norvegicus OX=10116 GN=Glyat PE=2 SV=1 |
| 220 | 301 | tr\|A4PB92 | 42.89 | 4 | 4 | 1.05E+05 | 1 | 1 | 1 | | 33899 Glycine N-acyltransferase OS=Rattus norvegicus OX=10116 GN=Glyat PE=2 SV=1 |
| 220 | 302 | tr\|A0A0G2 | 42.89 | 3 | 3 | 1.05E+05 | 1 | 1 | 1 | | 39453 Glycine N-acyltransferase OS=Rattus norvegicus OX=10116 GN=Glyat PE=1 SV=1 |
| 220 | 303 | tr\|B1H250 | 42.89 | 4 | 4 | 1.05E+05 | 1 | 1 | 1 | | 33986 Glycine-N-acyltransferase-like 1 OS=Rattus norvegicus OX=10116 GN=Glyatl1 PE=1 SV=1 |
| 136 | 255 | tr\|I6V4L9\| | 42.82 | 6 | 6 | 9.35E+04 | 2 | 2 | 2 | | 29844 Cytochrome c oxidase subunit 3 OS=Rattus norvegicus OX=10116 GN=COX3 PE=3 SV=1 |
| 137 | 260 | P35738\|OI | 42.82 | 5 | 5 | 9.06E+04 | 2 | 2 | 2 | | 42823 2-oxoisovalerate dehydrogenase subunit beta mitochondrial OS=Rattus norvegicus OX=10116 GN=Bckdhb PE=1 SV=3 |
| 137 | 261 | tr\|A0A0A0 | 42.82 | 5 | 5 | 9.06E+04 | 2 | 2 | 2 | | 42960 2-oxoisovalerate dehydrogenase subunit beta mitochondrial OS=Rattus norvegicus OX=10116 GN=Bckdhb PE=1 SV=1 |
| 140 | 297 | Q924S5\|LC | 41.96 | 2 | 2 | 1.38E+05 | 2 | 2 | 2 | | 105792 Lon protease homolog mitochondrial OS=Rattus norvegicus OX=10116 GN=Lonp1 PE=2 SV=1 |
| 129 | 298 | P11951\|CX | 41.25 | 17 | 17 | 2.59E+05 | 2 | 2 | 2 | | 8455 Cytochrome c oxidase subunit 6C-2 OS=Rattus norvegicus OX=10116 GN=Cox6c2 PE=1 SV=3 |
| 261 | 347 | tr\|D4A3E8 | 40.96 | 3 | 3 | 2.35E+05 | 1 | 1 | 1 | | 47648 Mitochondrial ribosomal protein S27 OS=Rattus norvegicus OX=10116 GN=Mrps27 PE=1 SV=1 |
| 222 | 309 | P22071\|3E | 40.96 | 3 | 3 | 5.49E+04 | 1 | 1 | 1 | | 42037 3 beta-hydroxysteroid dehydrogenase/Delta 5-->4-isomerase type 1 OS=Rattus norvegicus OX=10116 GN=Hsd3b1 PE=1 SV=3 |
| 222 | 310 | P22072\|3E | 40.96 | 3 | 3 | 5.49E+04 | 1 | 1 | 1 | | 42277 3 beta-hydroxysteroid dehydrogenase/Delta 5-->4-isomerase type 2 OS=Rattus norvegicus OX=10116 GN=Hsd3b PE=2 SV=3 |
| 222 | 311 | P27364\|3E | 40.96 | 3 | 3 | 5.49E+04 | 1 | 1 | 1 | | 42206 NADPH-dependent 3-keto-steroid reductase Hsd3b5 OS=Rattus norvegicus OX=10116 GN=Hsd3b5 PE=1 SV=3 |
| 105 | 267 | tr\|Q06Q97 | 39.36 | 6 | 6 | 1.56E+05 | 2 | 2 | 3 | | 38485 NADH-ubiquinone oxidoreductase chain 2 OS=Rattus norvegicus OX=10116 GN=ND2 PE=3 SV=1 |
| 105 | 268 | tr\|A0A0A1 | 39.36 | 6 | 6 | 1.56E+05 | 2 | 2 | 3 | | 38542 NADH-ubiquinone oxidoreductase chain 2 OS=Rattus norvegicus OX=10116 GN=ND2 PE=3 SV=1 |
| 105 | 269 | tr\|Q5UAJ8 | 39.36 | 6 | 6 | 1.56E+05 | 2 | 2 | 3 | | 38455 NADH-ubiquinone oxidoreductase chain 2 OS=Rattus norvegicus OX=10116 GN=ND2 PE=3 SV=1 |
| 105 | 270 | tr\|Q8HID0 | 39.36 | 6 | 6 | 1.56E+05 | 2 | 2 | 3 | | 38653 NADH-ubiquinone oxidoreductase chain 2 OS=Rattus norvegicus OX=10116 GN=Mt-nd2 PE=3 SV=1 |
| 105 | 271 | P11662\|NU | 39.36 | 6 | 6 | 1.56E+05 | 2 | 2 | 3 | | 38653 NADH-ubiquinone oxidoreductase chain 2 OS=Rattus norvegicus OX=10116 GN=Mtnd2 PE=3 SV=3 |
| 105 | 272 | tr\|Q06QH5 | 39.36 | 6 | 6 | 1.56E+05 | 2 | 2 | 3 | | 38626 NADH-ubiquinone oxidoreductase chain 2 OS=Rattus norvegicus OX=10116 GN=ND2 PE=3 SV=1 |
| 105 | 273 | tr\|Q8SEZ7 | 39.36 | 6 | 6 | 1.56E+05 | 2 | 2 | 3 | | 38580 NADH-ubiquinone oxidoreductase chain 2 (Fragment) OS=Rattus norvegicus OX=10116 GN=NADH2 PE=3 SV=1 |
| 105 | 274 | tr\|Q06QB0 | 39.36 | 6 | 6 | 1.56E+05 | 2 | 2 | 3 | | 38534 NADH-ubiquinone oxidoreductase chain 2 OS=Rattus norvegicus OX=10116 GN=ND2 PE=3 SV=1 |
| 105 | 275 | tr\|D2E6P4 | 39.36 | 6 | 6 | 1.56E+05 | 2 | 2 | 3 | | 38623 NADH-ubiquinone oxidoreductase chain 2 OS=Rattus norvegicus OX=10116 GN=ND2 PE=3 SV=1 |
| 105 | 276 | tr\|Q06QE9 | 39.36 | 6 | 6 | 1.56E+05 | 2 | 2 | 3 | | 38598 NADH-ubiquinone oxidoreductase chain 2 OS=Rattus norvegicus OX=10116 GN=ND2 PE=3 SV=1 |
| 92 | 225 | tr\|Q8SEZ8 | 38.73 | 7 | 7 | 5.66E+05 | 3 | 3 | 3 | Oxidation ( | 36133 NADH-ubiquinone oxidoreductase chain 1 (Fragment) OS=Rattus norvegicus OX=10116 GN=NADH1 PE=3 SV=1 |
| 92 | 226 | P03889\|NU | 38.73 | 7 | 7 | 5.66E+05 | 3 | 3 | 3 | Oxidation ( | 36145 NADH-ubiquinone oxidoreductase chain 1 OS=Rattus norvegicus OX=10116 GN=Mtnd1 PE=1 SV=3 |
| 92 | 227 | tr\|Q8HID1 | 38.73 | 7 | 7 | 5.66E+05 | 3 | 3 | 3 | Oxidation ( | 36145 NADH-ubiquinone oxidoreductase chain 1 OS=Rattus norvegicus OX=10116 GN=Mt-nd1 PE=3 SV=1 |
| 92 | 228 | tr\|D2E6L7 | 38.73 | 7 | 7 | 5.66E+05 | 3 | 3 | 3 | Oxidation ( | 36062 NADH-ubiquinone oxidoreductase chain 1 OS=Rattus norvegicus OX=10116 GN=ND1 PE=3 SV=1 |
| 94 | 322 | P12075\|CC | 37.07 | 22 | 22 | 2.18E+06 | 3 | 3 | 3 | | 13915 Cytochrome c oxidase subunit 5B mitochondrial OS=Rattus norvegicus OX=10116 GN=Cox5b PE=1 SV=2 |
| 110 | 236 | tr\|Q5U2T0 | 36.95 | 3 | 3 | 1.81E+04 | 1 | 1 | 2 | | 44494 Death associated protein 3 OS=Rattus norvegicus OX=10116 GN=Dap3 PE=2 SV=1 |
| 110 | 237 | tr\|F7EZZ0\| | 36.95 | 3 | 3 | 1.81E+04 | 1 | 1 | 2 | | 45112 Death-associated protein 3 OS=Rattus norvegicus OX=10116 GN=Dap3 PE=1 SV=1 |
| 110 | 242 | tr\|A0A0G2 | 36.95 | 2 | 2 | 1.81E+04 | 1 | 1 | 2 | | 46666 Death-associated protein 3 OS=Rattus norvegicus OX=10116 GN=Dap3 PE=1 SV=1 |
| 139 | 290 | tr\|A0A0G2 | 36.62 | 7 | 7 | 1.29E+06 | 2 | 2 | 2 | | 20750 Mitochondrial ribosomal protein L11 OS=Rattus norvegicus OX=10116 GN=mrpl11 PE=1 SV=1 |
| 139 | 291 | Q5XIE3\|RM | 36.62 | 7 | 7 | 1.29E+06 | 2 | 2 | 2 | | 22420 39S ribosomal protein L11 mitochondrial OS=Rattus norvegicus OX=10116 GN=Mrpl11 PE=2 SV=1 |

| | | | | | | | | | | | | |
|---|---|---|---|---|---|---|---|---|---|---|---|---|
| 224 | 314 | tr\|B1WBP | 35.98 | 8 | 8 | 3.54E+05 | 1 | 1 | 1 | | 12884 | ATP synthase subunit delta mitochondrial OS=Rattus norvegicus OX=10116 GN=Atp5f1d PE=1 SV=1 |
| 224 | 350 | tr\|G3V7Y3 | 35.98 | 6 | 6 | 3.54E+05 | 1 | 1 | 1 | | 17563 | ATP synthase subunit delta mitochondrial OS=Rattus norvegicus OX=10116 GN=Atp5f1d PE=1 SV=1 |
| 262 | 351 | Q8VID1\|DI | 35.13 | 3 | 3 | 9.71E+04 | 1 | 1 | 1 | | 29822 | Dehydrogenase/reductase SDR family member 4 OS=Rattus norvegicus OX=10116 GN=Dhrs4 PE=2 SV=2 |
| 107 | 418 | Q5XIA8\|GI | 34.52 | 3 | 3 | 0 | 2 | 1 | 3 | | 37178 | Growth hormone-inducible transmembrane protein OS=Rattus norvegicus OX=10116 GN=Ghitm PE=2 SV=1 |
| 128 | 244 | P09139\|SP | 33.48 | 6 | 6 | 3.20E+04 | 2 | 2 | 2 | | 45834 | Serine--pyruvate aminotransferase mitochondrial OS=Rattus norvegicus OX=10116 GN=Agxt PE=1 SV=1 |
| 223 | 312 | tr\|D3ZGM | 33.07 | 1 | 1 | 1.39E+05 | 1 | 1 | 1 | | 77891 | Pentatricopeptide repeat domain 3 OS=Rattus norvegicus OX=10116 GN=Ptcd3 PE=1 SV=1 |
| 223 | 313 | tr\|A0A0G2 | 33.07 | 1 | 1 | 1.39E+05 | 1 | 1 | 1 | | 84557 | Pentatricopeptide repeat domain 3 OS=Rattus norvegicus OX=10116 GN=Ptcd3 PE=1 SV=1 |
| 221 | 306 | Q2V057\|H | 32.72 | 2 | 2 | 0 | 1 | 1 | 1 | | 51002 | Hydroxyproline dehydrogenase OS=Rattus norvegicus OX=10116 GN=Prodh2 PE=2 SV=1 |
| 263 | 353 | Q9WUS0\|I | 32.43 | 4 | 4 | 0 | 1 | 1 | 1 | | 25203 | Adenylate kinase 4 mitochondrial OS=Rattus norvegicus OX=10116 GN=Ak4 PE=2 SV=1 |
| 96 | 383 | P42346\|M | 32.43 | 1 | 1 | 2.57E+05 | 3 | 2 | 3 | Formylatio | 288792 | Serine/threonine-protein kinase mTOR OS=Rattus norvegicus OX=10116 GN=Mtor PE=1 SV=1 |
| 103 | 656 | tr\|D3ZD23 | 32.2 | 1 | 1 | 5.82E+05 | 3 | 3 | 3 | | 67300 | ATP-binding cassette subfamily E member 1 OS=Rattus norvegicus OX=10116 GN=Abce1 PE=1 SV=1 |
| 81 | 482 | tr\|D3ZHF8 | 32.11 | 1 | 1 | 7.05E+05 | 2 | 1 | 4 | Formylatio | 73207 | Translation factor GUF1 mitochondrial OS=Rattus norvegicus OX=10116 GN=Guf1 PE=1 SV=1 |
| 188 | 253 | Q9ER34\|A | 31.98 | 3 | 3 | 2.25E+05 | 1 | 1 | 1 | | 85433 | Aconitate hydratase mitochondrial OS=Rattus norvegicus OX=10116 GN=Aco2 PE=1 SV=2 |
| 170 | 235 | tr\|B2RYW | 31.26 | 7 | 7 | 3.09E+05 | 1 | 1 | 1 | Oxidation ( | 21892 | NADH dehydrogenase (Ubiquinone) 1 beta subcomplex 9 OS=Rattus norvegicus OX=10116 GN=Ndufb9 PE=1 SV=1 |
| 264 | 358 | P06761\|BI | 31.19 | 1 | 1 | 8.78E+04 | 1 | 1 | 1 | | 72347 | Endoplasmic reticulum chaperone BiP OS=Rattus norvegicus OX=10116 GN=Hspa5 PE=1 SV=1 |
| 230 | 356 | O70600\|R | 30.96 | 3 | 3 | 4.07E+05 | 1 | 1 | 1 | Formylatio | 41255 | Radical S-adenosyl methionine domain-containing protein 2 OS=Rattus norvegicus OX=10116 GN=Rsad2 PE=1 SV=1 |
| 230 | 357 | tr\|A0A0H2 | 30.96 | 3 | 3 | 4.07E+05 | 1 | 1 | 1 | Formylatio | 41444 | RCG62278 OS=Rattus norvegicus OX=10116 GN=Rsad2 PE=4 SV=1 |
| 142 | 361 | P80433\|C | 30.83 | 14 | 14 | 2.94E+05 | 1 | 1 | 2 | | 7672 | Cytochrome c oxidase subunit 8A mitochondrial OS=Rattus norvegicus OX=10116 GN=Cox8a PE=1 SV=3 |
| 144 | 396 | P97521\|M | 30.38 | 4 | 4 | | 2 | 0 | 2 | | 33154 | Mitochondrial carnitine/acylcarnitine carrier protein OS=Rattus norvegicus OX=10116 GN=Slc25a20 PE=1 SV=1 |
| 144 | 397 | tr\|Q66HP8 | 30.38 | 4 | 4 | | 2 | 0 | 2 | | 33071 | Mitochondrial carnitine/acylcarnitine carrier protein OS=Rattus norvegicus OX=10116 GN=Slc25a20 PE=1 SV=1 |
| 146 | 432 | tr\|F1LSP2\| | 30.17 | 1 | 1 | 7.50E+04 | 2 | 1 | 2 | | 119190 | Acyl-CoA dehydrogenase family member 10 OS=Rattus norvegicus OX=10116 GN=Acad10 PE=1 SV=2 |
| 227 | 321 | tr\|Q5BJZ3 | 29.9 | 1 | 1 | 3.78E+05 | 1 | 1 | 1 | | 113869 | Nicotinamide nucleotide transhydrogenase OS=Rattus norvegicus OX=10116 GN=Nnt PE=1 SV=1 |
| 265 | 367 | tr\|D3ZXF8 | 29.34 | 5 | 5 | 2.09E+05 | 1 | 1 | 1 | | 17575 | Mitochondrial ribosomal protein L43 OS=Rattus norvegicus OX=10116 GN=Mrpl43 PE=1 SV=3 |
| 190 | 281 | tr\|F1LPV8\| | 28.4 | 2 | 2 | 1.26E+05 | 1 | 1 | 1 | | 46639 | Succinate--CoA ligase [GDP-forming] subunit beta mitochondrial OS=Rattus norvegicus OX=10116 GN=Suclg2 PE=1 SV=2 |
| 190 | 282 | tr\|B1H270 | 28.4 | 2 | 2 | 1.26E+05 | 1 | 1 | 1 | | 46988 | Succinate--CoA ligase [GDP-forming] subunit beta mitochondrial OS=Rattus norvegicus OX=10116 GN=Suclg2 PE=2 SV=1 |
| 266 | 368 | tr\|A0A0G2 | 28.27 | 4 | 4 | 3.54E+05 | 1 | 1 | 1 | | 33652 | Mitochondrial ribosomal protein L15 OS=Rattus norvegicus OX=10116 GN=Mrpl15 PE=1 SV=1 |
| 131 | 333 | tr\|F1LZW6 | 28.18 | 3 | 3 | 9.72E+04 | 2 | 1 | 2 | | 54099 | Solute carrier family 25 member 13 OS=Rattus norvegicus OX=10116 GN=Slc25a13 PE=1 SV=2 |
| 131 | 448 | tr\|A0A0G2 | 28.18 | 3 | 3 | 9.72E+04 | 2 | 1 | 2 | | 41504 | Solute carrier family 25 member 12 OS=Rattus norvegicus OX=10116 GN=Slc25a12 PE=1 SV=1 |
| 131 | 456 | tr\|F1LX07\| | 28.18 | 2 | 2 | 9.72E+04 | 2 | 1 | 2 | | 71917 | Solute carrier family 25 member 12 OS=Rattus norvegicus OX=10116 GN=Slc25a12 PE=1 SV=3 |
| 172 | 324 | P11497\|A | 27.62 | 0 | 0 | 1.47E+05 | 1 | 1 | 1 | | 265191 | Acetyl-CoA carboxylase 1 OS=Rattus norvegicus OX=10116 GN=Acaca PE=1 SV=1 |
| 172 | 327 | tr\|A0A0G2 | 27.62 | 0 | 0 | 1.47E+05 | 1 | 1 | 1 | | 276393 | Acetyl-CoA carboxylase beta OS=Rattus norvegicus OX=10116 GN=Acacb PE=1 SV=1 |
| 172 | 328 | tr\|D3ZBE2 | 27.62 | 0 | 0 | 1.47E+05 | 1 | 1 | 1 | | 275967 | Acetyl-CoA carboxylase beta OS=Rattus norvegicus OX=10116 GN=Acacb PE=1 SV=3 |
| 172 | 329 | tr\|E9PSQ0 | 27.62 | 0 | 0 | 1.47E+05 | 1 | 1 | 1 | | 276255 | Acetyl-CoA carboxylase beta OS=Rattus norvegicus OX=10116 GN=Acacb PE=1 SV=2 |
| 172 | 401 | tr\|A0A0G2 | 27.62 | 0 | 0 | 1.47E+05 | 1 | 1 | 1 | | 275755 | Acetyl-CoA carboxylase beta OS=Rattus norvegicus OX=10116 GN=Acacb PE=1 SV=1 |
| 172 | 402 | tr\|O70151 | 27.62 | 0 | 0 | 1.47E+05 | 1 | 1 | 1 | | 276097 | Acetyl-CoA carboxylase OS=Rattus norvegicus OX=10116 GN=Acacb PE=2 SV=1 |
| 225 | 315 | P04797\|G | 27.6 | 3 | 3 | 1.66E+05 | 1 | 1 | 1 | | 35828 | Glyceraldehyde-3-phosphate dehydrogenase OS=Rattus norvegicus OX=10116 GN=Gapdh PE=1 SV=3 |
| 130 | 307 | P16970\|A | 27.15 | 3 | 3 | 3.45E+05 | 2 | 2 | 2 | | 75316 | ATP-binding cassette sub-family D member 3 OS=Rattus norvegicus OX=10116 GN=Abcd3 PE=1 SV=3 |
| 267 | 369 | tr\|G3V734 | 27.11 | 3 | 3 | 2.24E+05 | 1 | 1 | 1 | | 36133 | 2 4-dienoyl CoA reductase 1 mitochondrial isoform CRA_a OS=Rattus norvegicus OX=10116 GN=Decr1 PE=1 SV=1 |
| 267 | 370 | Q64591\|D | 27.11 | 3 | 3 | 2.24E+05 | 1 | 1 | 1 | | 36133 | 2 4-dienoyl-CoA reductase mitochondrial OS=Rattus norvegicus OX=10116 GN=Decr1 PE=1 SV=2 |
| 268 | 371 | tr\|B5DFA3 | 26.86 | 2 | 2 | 9.55E+04 | 1 | 1 | 1 | | 65357 | Bucs1 protein OS=Rattus norvegicus OX=10116 GN=Acsm1 PE=2 SV=1 |
| 268 | 372 | tr\|F1M1W | 26.86 | 2 | 2 | 9.55E+04 | 1 | 1 | 1 | | 65347 | Acyl-coenzyme A synthetase ACSM2 mitochondrial OS=Rattus norvegicus OX=10116 GN=Acsm1 PE=1 SV=3 |
| 268 | 373 | tr\|A0A0G2 | 26.86 | 2 | 2 | 9.55E+04 | 1 | 1 | 1 | | 65319 | Acyl-coenzyme A synthetase ACSM2 mitochondrial OS=Rattus norvegicus OX=10116 GN=Acsm1 PE=1 SV=1 |
| 268 | 374 | tr\|A0A0G2 | 26.86 | 2 | 2 | 9.55E+04 | 1 | 1 | 1 | | 70943 | Acyl-coenzyme A synthetase ACSM2 mitochondrial OS=Rattus norvegicus OX=10116 GN=Acsm1 PE=1 SV=1 |
| 231 | 375 | tr\|A0A0G2 | 26.76 | 4 | 4 | 0 | 1 | 1 | 1 | | 45197 | Glycine C-acetyltransferase OS=Rattus norvegicus OX=10116 GN=Gcat PE=1 SV=1 |
| 147 | 480 | Q4V8B0\|O | 26.65 | 2 | 2 | 7.22E+04 | 2 | 2 | 2 | Formylatio | 92810 | Oxidation resistance protein 1 OS=Rattus norvegicus OX=10116 GN=Oxr1 PE=1 SV=3 |
| 143 | 377 | P80431\|C | 26.23 | 14 | 14 | 7.18E+05 | 1 | 1 | 2 | | 8995 | Cytochrome c oxidase subunit 7B mitochondrial OS=Rattus norvegicus OX=10116 GN=Cox7b PE=1 SV=3 |
| 143 | 378 | tr\|A0A0G2 | 26.23 | 14 | 14 | 7.18E+05 | 1 | 1 | 2 | | 9070 | Cytochrome c oxidase subunit 7B2 OS=Rattus norvegicus OX=10116 GN=Cox7b2 PE=4 SV=1 |
| 269 | 379 | tr\|D3ZDP2 | 26.2 | 4 | 4 | 1.93E+05 | 1 | 1 | 1 | | 23470 | Mitochondrial ribosomal protein L58 OS=Rattus norvegicus OX=10116 GN=Mrpl58 PE=1 SV=1 |
| 108 | 380 | P11530\|D | 25.47 | 0 | 0 | 1.27E+06 | 2 | 2 | 2 | | 425830 | Dystrophin OS=Rattus norvegicus OX=10116 GN=Dmd PE=1 SV=2 |
| 270 | 394 | P83565\|R | 25.39 | 8 | 8 | 1.32E+05 | 1 | 1 | 1 | | 24397 | 39S ribosomal protein L40 mitochondrial OS=Rattus norvegicus OX=10116 GN=Mrpl40 PE=1 SV=2 |
| 148 | 517 | tr\|F1M4Y5 | 25.19 | 1 | 1 | 2.69E+05 | 2 | 2 | 2 | Formylatio | 108218 | Storkhead box 1 OS=Rattus norvegicus OX=10116 GN=Stox1 PE=4 SV=2 |
| 145 | 398 | P48037\|A | 25.13 | 1 | 1 | 6.63E+05 | 1 | 1 | 2 | | 75754 | Annexin A6 OS=Rattus norvegicus OX=10116 GN=Anxa6 PE=1 SV=2 |
| 145 | 399 | tr\|Q6IMZ3 | 25.13 | 1 | 1 | 6.63E+05 | 1 | 1 | 2 | | 75756 | Annexin OS=Rattus norvegicus OX=10116 GN=Anxa6 PE=1 SV=1 |
| 132 | 349 | Q5I0C5\|F | 25.01 | 2 | 2 | 5.08E+05 | 1 | 1 | 2 | Formylatio | 42997 | Methionyl-tRNA formyltransferase mitochondrial OS=Rattus norvegicus OX=10116 GN=Mtfmt PE=2 SV=1 |
| 274 | 413 | tr\|M0RA2 | 23.92 | 7 | 7 | 2.76E+05 | 1 | 1 | 1 | | 10982 | Uncharacterized protein OS=Rattus norvegicus OX=10116 PE=4 SV=2 |
| 274 | 414 | tr\|D4A4P3 | 23.92 | 7 | 7 | 2.76E+05 | 1 | 1 | 1 | | 11267 | NADH:ubiquinone oxidoreductase subunit B3 OS=Rattus norvegicus OX=10116 GN=Ndufb3 PE=1 SV=1 |
| 229 | 355 | tr\|D4AE90 | 23.78 | 2 | 2 | 0 | 1 | 1 | 1 | Formylatio | 50113 | RCC1-like OS=Rattus norvegicus OX=10116 GN=Rcc1l PE=1 SV=1 |
| 273 | 412 | tr\|B2GV62 | 23.74 | 7 | 7 | 1.83E+05 | 1 | 1 | 1 | | 17526 | Mitochondrial ribosomal protein L20 OS=Rattus norvegicus OX=10116 GN=Mrpl20 PE=1 SV=1 |
| 115 | 294 | Q01728\|N | 23.58 | 1 | 1 | 5.43E+06 | 1 | 1 | 2 | Formylatio | 108184 | Sodium/calcium exchanger 1 OS=Rattus norvegicus OX=10116 GN=Slc8a1 PE=1 SV=3 |
| 115 | 359 | P70549\|N | 23.58 | 1 | 1 | 5.43E+06 | 1 | 1 | 2 | Formylatio | 103163 | Sodium/calcium exchanger 3 OS=Rattus norvegicus OX=10116 GN=Slc8a3 PE=1 SV=1 |
| 133 | 419 | Q01062\|P | 23.4 | 2 | 2 | 5.12E+05 | 1 | 1 | 2 | | 104664 | cGMP-dependent 3' 5'-cyclic phosphodiesterase OS=Rattus norvegicus OX=10116 GN=Pde2a PE=1 SV=2 |
| 133 | 420 | tr\|F8WFW | 23.4 | 1 | 1 | 5.12E+05 | 1 | 1 | 2 | | 105234 | Phosphodiesterase OS=Rattus norvegicus OX=10116 GN=Pde2a PE=1 SV=1 |
| 275 | 428 | P24473\|G | 23.11 | 4 | 4 | 6.97E+04 | 1 | 1 | 1 | | 25493 | Glutathione S-transferase kappa 1 OS=Rattus norvegicus OX=10116 GN=Gstk1 PE=1 SV=3 |

| | | | | | | | | | | | |
|---|---|---|---|---|---|---|---|---|---|---|---|
| 275 | 429 | tr\|B6DYQ0 | 23.11 | 4 | 4 | 6.97E+04 | 1 | 1 | 1 | 25493 | Glutathione S-transferase kappa OS=Rattus norvegicus OX=10116 GN=Gstk1 PE=2 SV=1 |
| 276 | 431 | D3ZHR2\|A | 22.69 | 1 | 1 | 2.13E+05 | 1 | 1 | 1 | 81938 | ATP-binding cassette sub-family D member 1 OS=Rattus norvegicus OX=10116 GN=Abcd1 PE=1 SV=1 |
| 278 | 443 | tr\|D4A5X7 | 22.3 | 3 | 3 | 8.54E+04 | 1 | 1 | 1 | 41217 | Ganglioside-induced differentiation-associated-protein 1 OS=Rattus norvegicus OX=10116 GN=Gdap1 PE=3 SV=1 |
| 279 | 447 | tr\|D3ZMR9 | 22 | 4 | 4 | 2.35E+05 | 1 | 1 | 1 | 23390 | Mitochondrial ribosomal protein L21 OS=Rattus norvegicus OX=10116 GN=Mrpl21 PE=1 SV=1 |
| 232 | 406 | P28494\|M | 21.8 | 1 | 1 | 3.07E+04 | 1 | 1 | 1 | 131242 | Alpha-mannosidase 2 OS=Rattus norvegicus OX=10116 GN=Man2a1 PE=1 SV=2 |
| 180 | 407 | tr\|G3V996 | 21.06 | 2 | 2 | 8.92E+04 | 1 | 1 | 1 | 41967 | LETM1 domain-containing 1 OS=Rattus norvegicus OX=10116 GN=Letmd1 PE=1 SV=1 |
| 180 | 408 | tr\|B0K026 | 21.06 | 2 | 2 | 8.92E+04 | 1 | 1 | 1 | 43121 | LETM1 domain-containing 1 OS=Rattus norvegicus OX=10116 GN=Letmd1 PE=1 SV=1 |
| 282 | 520 | tr\|A0A0G2 | 20.67 | 1 | 1 | 1.21E+05 | 1 | 1 | 1 | 74235 | Selenoprotein O OS=Rattus norvegicus OX=10116 GN=Selenoo PE=1 SV=1 |
| 282 | 521 | tr\|B2GVA1 | 20.67 | 1 | 1 | 1.21E+05 | 1 | 1 | 1 | 74385 | Selenoprotein O OS=Rattus norvegicus OX=10116 GN=Selenoo PE=2 SV=1 |
