## Supplementary material for "Permeability transition pore-related changes in the proteome and channel activity of ATP synthase dimers and monomers": RLM SM-BM

| Protein Gro | Protein ID | Accession | -10lgP | Coverage ( | Coverage ( | IBAQ | #Peptides | #Unique | #Spec Sam | PTM | Avg. Mass | Description |
|---|---|---|---|---|---|---|---|---|---|---|---|---|
| 1 | 1 | P10860\|DH | 360.33 | 52 | 52 | 3.65E+08 | 151 | 150 | 327 | Carbamidomethylatior | | Glutamate dehydrogenase 1, mitochondrial |
| 11 | 38 | tr\|D3ZFQ8 | 260.23 | 22 | 22 | 3.05E+07 | 27 | 27 | 49 | Oxidation ( | 35435 | Cytochrome c-1 OS=Rattus norvegicus OX=10116 GN=Cyc1 PE=1 SV=3 |
| 17 | 49 | tr\|B2RYS8 | 255.9 | 39 | 39 | 2.57E+07 | 27 | 27 | 44 | Oxidation ( | 21959 | NADH dehydrogenase [ubiquinone] 1 beta subcomplex subunit 8 mitochondrial OS=Rattus norvegicus OX=10116 GN=Ndufb8 PE=1 SV=1 |
| 5 | 16 | Q68FY0\|Q | 250.22 | 28 | 28 | 6.47E+07 | 46 | 45 | 83 | Carbamido | 52849 | Cytochrome b-c1 complex subunit 1 mitochondrial OS=Rattus norvegicus OX=10116 GN=Uqcrc1 PE=1 SV=1 |
| 10 | 8 | P10719\|AT | 248.87 | 19 | 19 | 2.87E+07 | 30 | 29 | 50 | Oxidation ( | 56354 | ATP synthase subunit beta mitochondrial OS=Rattus norvegicus OX=10116 GN=Atp5f1b PE=1 SV=2 |
| 10 | 9 | tr\|G3V6D3 | 248.87 | 19 | 19 | 2.87E+07 | 30 | 29 | 50 | Oxidation ( | 56345 | ATP synthase subunit beta OS=Rattus norvegicus OX=10116 GN=Atp5f1b PE=1 SV=1 |
| 18 | 42 | P20788\|U | 247.71 | 29 | 29 | 2.95E+07 | 25 | 25 | 43 | | 29446 | Cytochrome b-c1 complex subunit Rieske mitochondrial OS=Rattus norvegicus OX=10116 GN=Uqcrfs1 PE=1 SV=2 |
| 2 | 20 | P19234\|NI | 247.46 | 37 | 37 | 6.66E+07 | 36 | 36 | 95 | | 27378 | NADH dehydrogenase [ubiquinone] flavoprotein 2 mitochondrial OS=Rattus norvegicus OX=10116 GN=Ndufv2 PE=1 SV=2 |
| 4 | 13 | Q66HF1\|N | 240.07 | 26 | 26 | 4.85E+07 | 55 | 55 | 85 | Oxidation ( | 79412 | NADH-ubiquinone oxidoreductase 75 kDa subunit mitochondrial OS=Rattus norvegicus OX=10116 GN=Ndufs1 PE=1 SV=1 |
| 7 | 19 | Q641Y2\|N | 231.78 | 31 | 31 | 3.76E+07 | 42 | 41 | 72 | Oxidation ( | 52562 | NADH dehydrogenase [ubiquinone] iron-sulfur protein 2 mitochondrial OS=Rattus norvegicus OX=10116 GN=Ndufs2 PE=1 SV=1 |
| 14 | 3 | P85834\|EF | 216.92 | 19 | 19 | 1.75E+07 | 23 | 21 | 46 | | 49522 | Elongation factor Tu mitochondrial OS=Rattus norvegicus OX=10116 GN=Tufm PE=1 SV=1 |
| 22 | 58 | tr\|D4A565 | 216.37 | 38 | 38 | 1.91E+07 | 20 | 20 | 33 | | 21664 | NADH dehydrogenase (Ubiquinone) 1 beta subcomplex 5 (Predicted) isoform CRA_b OS=Rattus norvegicus OX=10116 GN=Ndufb5 PE=1 SV=1 |
| 9 | 67 | tr\|L0L4L8\| | 215.68 | 23 | 23 | 3.59E+06 | 27 | 1 | 51 | | 42470 | Cytochrome b (Fragment) OS=Rattus norvegicus OX=10116 GN=cytb PE=3 SV=1 |
| 9 | 68 | tr\|A0A140 | 215.68 | 23 | 23 | 3.59E+06 | 27 | 1 | 51 | | 42574 | Cytochrome b (Fragment) OS=Rattus norvegicus OX=10116 PE=3 SV=1 |
| 9 | 69 | tr\|A0A140 | 215.68 | 23 | 23 | 3.59E+06 | 27 | 1 | 51 | | 42588 | Cytochrome b (Fragment) OS=Rattus norvegicus OX=10116 PE=3 SV=1 |
| 9 | 125 | tr\|A0A385 | 215.68 | 23 | 23 | 3.59E+06 | 27 | 1 | 51 | | 42766 | Cytochrome b (Fragment) OS=Rattus norvegicus OX=10116 PE=3 SV=1 |
| 9 | 71 | tr\|A0A3Q8 | 215.68 | 23 | 23 | 3.59E+06 | 27 | 1 | 51 | | 42712 | Cytochrome b (Fragment) OS=Rattus norvegicus OX=10116 GN=Cytb PE=3 SV=1 |
| 9 | 72 | tr\|A0A3Q8 | 215.68 | 23 | 23 | 3.59E+06 | 27 | 1 | 51 | | 42622 | Cytochrome b (Fragment) OS=Rattus norvegicus OX=10116 GN=Cytb PE=3 SV=1 |
| 9 | 74 | tr\|A0A3S6 | 215.68 | 23 | 23 | 3.59E+06 | 27 | 1 | 51 | | 42698 | Cytochrome b (Fragment) OS=Rattus norvegicus OX=10116 GN=Cytb PE=3 SV=1 |
| 9 | 76 | tr\|A0A220 | 215.68 | 23 | 23 | 3.59E+06 | 27 | 1 | 51 | | 43018 | Cytochrome b OS=Rattus norvegicus OX=10116 PE=3 SV=1 |
| 9 | 77 | tr\|D6NSP2 | 215.68 | 23 | 23 | 3.59E+06 | 27 | 1 | 51 | | 42945 | Cytochrome b (Fragment) OS=Rattus norvegicus OX=10116 GN=cytb PE=3 SV=1 |
| 9 | 78 | tr\|D6NSS6 | 215.68 | 23 | 23 | 3.59E+06 | 27 | 1 | 51 | | 42993 | Cytochrome b (Fragment) OS=Rattus norvegicus OX=10116 GN=cytb PE=3 SV=1 |
| 9 | 79 | tr\|D6NSQ3 | 215.68 | 23 | 23 | 3.59E+06 | 27 | 1 | 51 | | 43012 | Cytochrome b (Fragment) OS=Rattus norvegicus OX=10116 GN=cytb PE=3 SV=1 |
| 9 | 80 | tr\|D6NSR3 | 215.68 | 23 | 23 | 3.59E+06 | 27 | 1 | 51 | | 42986 | Cytochrome b (Fragment) OS=Rattus norvegicus OX=10116 GN=cytb PE=3 SV=1 |
| 9 | 82 | tr\|D6NSR8 | 215.68 | 23 | 23 | 3.59E+06 | 27 | 1 | 51 | | 42998 | Cytochrome b (Fragment) OS=Rattus norvegicus OX=10116 GN=cytb PE=3 SV=1 |
| 9 | 84 | tr\|A0A0A1 | 215.68 | 23 | 23 | 3.59E+06 | 27 | 1 | 51 | | 42989 | Cytochrome b OS=Rattus norvegicus OX=10116 GN=CYTB PE=3 SV=1 |
| 9 | 85 | tr\|D6NSS7 | 215.68 | 23 | 23 | 3.59E+06 | 27 | 1 | 51 | | 42948 | Cytochrome b (Fragment) OS=Rattus norvegicus OX=10116 GN=cytb PE=3 SV=1 |
| 9 | 86 | tr\|Q8SEY9 | 215.68 | 23 | 23 | 3.59E+06 | 27 | 1 | 51 | | 43015 | Cytochrome b OS=Rattus norvegicus OX=10116 GN=cytb PE=3 SV=1 |
| 9 | 87 | tr\|D6NSR7 | 215.68 | 23 | 23 | 3.59E+06 | 27 | 1 | 51 | | 42982 | Cytochrome b (Fragment) OS=Rattus norvegicus OX=10116 GN=cytb PE=3 SV=1 |
| 9 | 88 | tr\|A0A220 | 215.68 | 23 | 23 | 3.59E+06 | 27 | 1 | 51 | | 42968 | Cytochrome b OS=Rattus norvegicus OX=10116 PE=3 SV=1 |
| 9 | 116 | tr\|D6NSQ0 | 215.68 | 23 | 23 | 3.59E+06 | 27 | 1 | 51 | | 42968 | Cytochrome b (Fragment) OS=Rattus norvegicus OX=10116 GN=cytb PE=3 SV=1 |
| 9 | 91 | tr\|Q5UAI7 | 215.68 | 23 | 23 | 3.59E+06 | 27 | 1 | 51 | | 42998 | Cytochrome b OS=Rattus norvegicus OX=10116 GN=CYTB PE=3 SV=1 |
| 9 | 128 | tr\|D6NSR2 | 215.68 | 23 | 23 | 3.59E+06 | 27 | 1 | 51 | | 43073 | Cytochrome b (Fragment) OS=Rattus norvegicus OX=10116 GN=cytb PE=3 SV=1 |
| 9 | 92 | tr\|A0A0S1 | 215.68 | 23 | 23 | 3.59E+06 | 27 | 1 | 51 | | 43002 | Cytochrome b OS=Rattus norvegicus OX=10116 GN=CYTB PE=3 SV=1 |
| 9 | 93 | tr\|Q8HIC4 | 215.68 | 23 | 23 | 3.59E+06 | 27 | 1 | 51 | | 43012 | Cytochrome b OS=Rattus norvegicus OX=10116 GN=Mt-cyb PE=3 SV=1 |
| 9 | 94 | tr\|A0A220 | 215.68 | 23 | 23 | 3.59E+06 | 27 | 1 | 51 | | 42968 | Cytochrome b OS=Rattus norvegicus OX=10116 PE=3 SV=1 |
| 9 | 95 | tr\|A0A0S1 | 215.68 | 23 | 23 | 3.59E+06 | 27 | 1 | 51 | | 43016 | Cytochrome b OS=Rattus norvegicus OX=10116 GN=CYTB PE=3 SV=1 |
| 9 | 96 | tr\|R9TKN1 | 215.68 | 23 | 23 | 3.59E+06 | 27 | 1 | 51 | | 43012 | Cytochrome b OS=Rattus norvegicus OX=10116 GN=CYTB PE=3 SV=1 |
| 9 | 97 | P00159\|CY | 215.68 | 23 | 23 | 3.59E+06 | 27 | 1 | 51 | | 43012 | Cytochrome b OS=Rattus norvegicus OX=10116 GN=Mt-Cyb PE=3 SV=3 |
| 9 | 98 | tr\|L0N311 | 215.68 | 23 | 23 | 3.59E+06 | 27 | 1 | 51 | | 43045 | Cytochrome b (Fragment) OS=Rattus norvegicus OX=10116 GN=cytb PE=3 SV=1 |
| 9 | 99 | tr\|A0A220 | 215.68 | 23 | 23 | 3.59E+06 | 27 | 1 | 51 | | 42970 | Cytochrome b OS=Rattus norvegicus OX=10116 PE=3 SV=1 |
| 9 | 100 | tr\|H2KXA0 | 215.68 | 23 | 23 | 3.59E+06 | 27 | 1 | 51 | | 42978 | Cytochrome b (Fragment) OS=Rattus norvegicus OX=10116 GN=cytb PE=3 SV=1 |
| 9 | 101 | tr\|A0A096 | 215.68 | 23 | 23 | 3.59E+06 | 27 | 1 | 51 | | 43042 | Cytochrome b (Fragment) OS=Rattus norvegicus OX=10116 PE=3 SV=1 |
| 9 | 102 | tr\|D6NSP5 | 215.68 | 23 | 23 | 3.59E+06 | 27 | 1 | 51 | | 43012 | Cytochrome b (Fragment) OS=Rattus norvegicus OX=10116 GN=cytb PE=3 SV=1 |
| 9 | 124 | tr\|A0A411 | 215.68 | 23 | 23 | 3.59E+06 | 27 | 1 | 51 | | 42238 | Cytochrome b (Fragment) OS=Rattus norvegicus OX=10116 GN=Cytb PE=3 SV=1 |
| 9 | 127 | tr\|D6NSP4 | 215.68 | 23 | 23 | 3.59E+06 | 27 | 1 | 51 | | 43016 | Cytochrome b (Fragment) OS=Rattus norvegicus OX=10116 GN=cytb PE=3 SV=1 |
| 9 | 129 | tr\|D6NSR1 | 215.68 | 23 | 23 | 3.59E+06 | 27 | 1 | 51 | | 43002 | Cytochrome b (Fragment) OS=Rattus norvegicus OX=10116 GN=cytb PE=3 SV=1 |
| 12 | 126 | tr\|A0A385 | 206.71 | 23 | 23 | 3.27E+05 | 27 | 1 | 48 | | 42751 | Cytochrome b (Fragment) OS=Rattus norvegicus OX=10116 PE=3 SV=1 |
| 6 | 4 | P07756\|CF | 205.76 | 17 | 17 | 2.17E+07 | 48 | 48 | 75 | Carbamidomethylatior | | Carbamoyl-phosphate synthase [ammonia], mitochondrial |
| 8 | 7 | Q5BK63\|N | 201.23 | 32 | 32 | 1.73E+07 | 32 | 31 | 57 | | 42559 | NADH dehydrogenase [ubiquinone] 1 alpha subcomplex subunit 9 mitochondrial OS=Rattus norvegicus OX=10116 GN=Ndufa9 PE=1 SV=2 |
| 20 | 204 | tr\|Q5PQZ9 | 196.91 | 45 | 45 | 2.36E+07 | 18 | 18 | 38 | | 14359 | NADH dehydrogenase [ubiquinone] 1 subunit C2 OS=Rattus norvegicus OX=10116 GN=Ndufc2 PE=1 SV=1 |
| 21 | 30 | P52873\|PY | 194.27 | 15 | 15 | 5.77E+06 | 28 | 27 | 36 | | 129777 | Pyruvate carboxylase mitochondrial OS=Rattus norvegicus OX=10116 GN=Pc PE=1 SV=2 |
| 21 | 31 | tr\|A0A0G2 | 194.27 | 13 | 13 | 5.77E+06 | 28 | 27 | 36 | | 140005 | Pyruvate carboxylase mitochondrial OS=Rattus norvegicus OX=10116 GN=Pc PE=1 SV=1 |
| 19 | 34 | tr\|A0A0G2 | 191.93 | 28 | 28 | 2.00E+07 | 17 | 17 | 40 | | 19965 | NADH dehydrogenase [ubiquinone] 1 alpha subcomplex subunit 8 OS=Rattus norvegicus OX=10116 GN=Ndufa8 PE=1 SV=1 |
| 26 | 26 | P00507\|AA | 191.31 | 21 | 21 | 1.22E+07 | 19 | 19 | 29 | Oxidation ( | 47314 | Aspartate aminotransferase mitochondrial OS=Rattus norvegicus OX=10116 GN=Got2 PE=1 SV=2 |
| 16 | 57 | Q561S0\|N | 190.22 | 30 | 30 | 1.56E+07 | 26 | 25 | 44 | Oxidation ( | 40493 | NADH dehydrogenase [ubiquinone] 1 alpha subcomplex subunit 10 mitochondrial OS=Rattus norvegicus OX=10116 GN=Ndufa10 PE=1 SV=1 |
| 13 | 10 | P15999\|AT | 185.56 | 24 | 24 | 2.12E+07 | 32 | 30 | 47 | Oxidation ( | 59754 | ATP synthase subunit alpha mitochondrial OS=Rattus norvegicus OX=10116 GN=Atp5f1a PE=1 SV=2 |
| 13 | 11 | tr\|F1LP05\| | 185.56 | 24 | 24 | 2.12E+07 | 32 | 30 | 47 | Oxidation ( | 59813 | ATP synthase subunit alpha OS=Rattus norvegicus OX=10116 GN=Atp5f1a PE=1 SV=1 |
| 38 | 136 | P24329\|TH | 182.97 | 20 | 20 | 2.35E+06 | 9 | 9 | 14 | | 33407 | Thiosulfate sulfurtransferase OS=Rattus norvegicus OX=10116 GN=Tst PE=1 SV=3 |
| 3 | 21 | P32551\|Q | 178.29 | 24 | 24 | 4.58E+07 | 32 | 31 | 92 | Carbamidomethylatior | | Cytochrome b-c1 complex subunit 2, mitochondrial |
| 15 | 28 | tr\|D3ZG43 | 176.38 | 34 | 34 | 2.98E+07 | 25 | 25 | 45 | | 30226 | NADH dehydrogenase (Ubiquinone) Fe-S protein 3 (Predicted) isoform CRA_c OS=Rattus norvegicus OX=10116 GN=Ndufs3 PE=1 SV=1 |
| 24 | 48 | tr\|D3ZF13 | 174.41 | 28 | 28 | 1.98E+07 | 17 | 17 | 30 | | 17514 | Acyl carrier protein OS=Rattus norvegicus OX=10116 GN=Ndufab1 PE=1 SV=1 |

| | | | | | | | | | | | |
|---|---|---|---|---|---|---|---|---|---|---|---|
| 28 | 182 | tr\|Q06QG | 173.92 | 12 | 12 | 1.13E+07 | 17 | 17 | 24 | Oxidation ( | 68588 NADH-ubiquinone oxidoreductase chain 5 OS=Rattus norvegicus OX=10116 GN=ND5 PE=3 SV=1 |
| 28 | 183 | tr\|Q06QK5 | 173.92 | 12 | 12 | 1.13E+07 | 17 | 17 | 24 | Oxidation ( | 68606 NADH-ubiquinone oxidoreductase chain 5 OS=Rattus norvegicus OX=10116 GN=ND5 PE=3 SV=1 |
| 28 | 184 | P11661\|NU | 173.92 | 12 | 12 | 1.13E+07 | 17 | 17 | 24 | Oxidation ( | 68618 NADH-ubiquinone oxidoreductase chain 5 OS=Rattus norvegicus OX=10116 GN=Mtnd5 PE=3 SV=3 |
| 28 | 185 | tr\|Q06QA1 | 173.92 | 12 | 12 | 1.13E+07 | 17 | 17 | 24 | Oxidation ( | 68574 NADH-ubiquinone oxidoreductase chain 5 OS=Rattus norvegicus OX=10116 GN=ND5 PE=3 SV=1 |
| 28 | 186 | tr\|Q8SEZ0 | 173.92 | 12 | 12 | 1.13E+07 | 17 | 17 | 24 | Oxidation ( | 68618 NADH-ubiquinone oxidoreductase chain 5 OS=Rattus norvegicus OX=10116 GN=Mt-nd5 PE=3 SV=1 |
| 28 | 187 | tr\|A0A096 | 173.92 | 12 | 12 | 1.13E+07 | 17 | 17 | 24 | Oxidation ( | 68584 NADH-ubiquinone oxidoreductase chain 5 OS=Rattus norvegicus OX=10116 GN=ND5 PE=3 SV=1 |
| 46 | 249 | tr\|F1LXA0\| | 173.16 | 28 | 28 | 7.58E+06 | 8 | 8 | 12 | Oxidation ( | 17178 NADH dehydrogenase [ubiquinone] 1 alpha subcomplex subunit 12 OS=Rattus norvegicus OX=10116 GN=Ndufa12 PE=1 SV=2 |
| 23 | 15 | P67779\|PH | 171.76 | 26 | 26 | 9.57E+06 | 20 | 19 | 31 | | 29820 Prohibitin OS=Rattus norvegicus OX=10116 GN=Phb PE=1 SV=1 |
| 29 | 41 | tr\|D4A3V2 | 169.52 | 34 | 34 | 8.57E+06 | 16 | 16 | 24 | | 15224 NADH dehydrogenase [ubiquinone] 1 alpha subcomplex subunit 6 OS=Rattus norvegicus OX=10116 GN=Ndufa6 PE=1 SV=1 |
| 35 | 137 | tr\|D4A7L4 | 164.06 | 23 | 23 | 1.38E+07 | 9 | 9 | 15 | | 17634 NADH dehydrogenase (Ubiquinone) 1 beta subcomplex 11 (Predicted) OS=Rattus norvegicus OX=10116 GN=Ndufb11 PE=1 SV=1 |
| 32 | 181 | Q80W89\| | 163.87 | 18 | 18 | 9.78E+06 | 9 | 9 | 17 | | 14854 NADH dehydrogenase [ubiquinone] 1 alpha subcomplex subunit 11 OS=Rattus norvegicus OX=10116 GN=Ndufa11 PE=2 SV=1 |
| 30 | 66 | tr\|D4A0T0 | 159.46 | 38 | 38 | 2.36E+07 | 13 | 13 | 23 | | 20859 NADH:ubiquinone oxidoreductase subunit B10 OS=Rattus norvegicus OX=10116 GN=Ndufb10 PE=1 SV=1 |
| 27 | 55 | tr\|B0BNE6 | 151.31 | 26 | 26 | 7.92E+06 | 19 | 18 | 26 | | 23970 NADH dehydrogenase (Ubiquinone) Fe-S protein 8 (Predicted) isoform CRA_a OS=Rattus norvegicus OX=10116 GN=Ndufs8 PE=1 SV=1 |
| 25 | 43 | tr\|Q5XIH3 | 144.68 | 19 | 19 | 2.40E+07 | 15 | 15 | 30 | Oxidation ( | 50731 NADH dehydrogenase [ubiquinone] flavoprotein 1 mitochondrial OS=Rattus norvegicus OX=10116 GN=Ndufv1 PE=1 SV=1 |
| 34 | 245 | tr\|A9UMV | 144.28 | 30 | 30 | 1.51E+07 | 8 | 8 | 16 | Oxidation ( | 12500 NADH:ubiquinone oxidoreductase subunit A7 OS=Rattus norvegicus OX=10116 GN=Ndufa7 PE=1 SV=1 |
| 49 | 142 | tr\|D3ZCZ9 | 140.5 | 31 | 31 | 5.01E+06 | 6 | 6 | 11 | | 13040 NADH dehydrogenase [ubiquinone] iron-sulfur protein 6 mitochondrial OS=Rattus norvegicus OX=10116 GN=LOC100912599 PE=1 SV=1 |
| 48 | 138 | Q5XIF3\|ND | 138.6 | 15 | 15 | 5.65E+06 | 7 | 7 | 11 | | 19741 NADH dehydrogenase [ubiquinone] iron-sulfur protein 4 mitochondrial OS=Rattus norvegicus OX=10116 GN=Ndufs4 PE=1 SV=1 |
| 44 | 35 | P17764\|TH | 136.01 | 10 | 10 | 3.92E+06 | 9 | 9 | 12 | | 44695 Acetyl-CoA acetyltransferase mitochondrial OS=Rattus norvegicus OX=10116 GN=Acat1 PE=1 SV=1 |
| 40 | 65 | tr\|D3ZE15 | 134.46 | 35 | 35 | 3.46E+06 | 9 | 8 | 13 | Oxidation ( | 16777 NADH:ubiquinone oxidoreductase subunit A13 OS=Rattus norvegicus OX=10116 GN=Ndufa13 PE=1 SV=1 |
| 56 | 63 | tr\|Q6QI09 | 130.8 | 7 | 7 | 2.86E+06 | 8 | 8 | 9 | | 67721 ATP synthase subunit gamma mitochondrial OS=Rattus norvegicus OX=10116 GN=Taf3 PE=1 SV=1 |
| 51 | 145 | tr\|F1LPG5 | 129.82 | 21 | 21 | 5.74E+06 | 7 | 7 | 11 | | 15064 NADH:ubiquinone oxidoreductase subunit B4 OS=Rattus norvegicus OX=10116 GN=Ndufb4 PE=1 SV=1 |
| 51 | 157 | tr\|F1M7T1 | 129.82 | 21 | 21 | 5.74E+06 | 7 | 7 | 11 | | 15160 NADH dehydrogenase (ubiquinone) 1 beta subcomplex 4 OS=Rattus norvegicus OX=10116 GN=LOC100361934 PE=4 SV=2 |
| 47 | 338 | Q5M9I5\|Q | 129.8 | 21 | 21 | 1.24E+07 | 5 | 5 | 12 | | 10424 Cytochrome b-c1 complex subunit 6 mitochondrial OS=Rattus norvegicus OX=10116 GN=Uqcrh PE=3 SV=1 |
| 37 | 27 | Q60587\|E | 128.86 | 12 | 12 | 2.51E+06 | 11 | 11 | 14 | | 51414 Trifunctional enzyme subunit beta mitochondrial OS=Rattus norvegicus OX=10116 GN=Hadhb PE=1 SV=1 |
| 37 | 29 | tr\|A0A0G2 | 128.86 | 12 | 12 | 2.51E+06 | 11 | 11 | 14 | | 52568 Trifunctional enzyme subunit beta mitochondrial OS=Rattus norvegicus OX=10116 GN=Hadhb PE=1 SV=1 |
| 60 | 258 | tr\|M0RB6 | 127.66 | 48 | 48 | 4.36E+06 | 5 | 5 | 8 | Oxidation ( | 9372 RCG63041 OS=Rattus norvegicus OX=10116 GN=LOC684509 PE=4 SV=1 |
| 60 | 257 | tr\|A9UMW | 127.66 | 48 | 48 | 4.36E+06 | 5 | 5 | 8 | Oxidation ( | 9255 Ndufa3 protein (Fragment) OS=Rattus norvegicus OX=10116 GN=Ndufa3 PE=2 SV=1 |
| 60 | 259 | tr\|A0A0G2 | 127.66 | 44 | 44 | 4.36E+06 | 5 | 5 | 8 | Oxidation ( | 10170 NADH:ubiquinone oxidoreductase subunit A3 OS=Rattus norvegicus OX=10116 GN=Ndufa3 PE=1 SV=1 |
| 36 | 24 | Q64428\|E | 126.44 | 9 | 9 | 4.43E+06 | 9 | 9 | 14 | Carbamido | 82665 Trifunctional enzyme subunit alpha mitochondrial OS=Rattus norvegicus OX=10116 GN=Hadha PE=1 SV=2 |
| 33 | 135 | tr\|D3ZZ21 | 125.1 | 37 | 37 | 5.81E+06 | 10 | 9 | 16 | Oxidation ( | 15638 NADH dehydrogenase (Ubiquinone) 1 beta subcomplex 6 (Predicted) OS=Rattus norvegicus OX=10116 GN=Ndufb6 PE=1 SV=1 |
| 39 | 17 | tr\|A0A0G2 | 119.52 | 15 | 15 | 7.55E+06 | 10 | 9 | 14 | | 33168 Prohibitin OS=Rattus norvegicus OX=10116 GN=Phb2 PE=1 SV=1 |
| 39 | 12 | Q5XIH7\|PH | 119.52 | 15 | 15 | 7.55E+06 | 10 | 9 | 14 | | 33312 Prohibitin-2 OS=Rattus norvegicus OX=10116 GN=Phb2 PE=1 SV=1 |
| 58 | 194 | tr\|Q5RJN0 | 116.63 | 19 | 19 | 1.39E+06 | 8 | 8 | 8 | Oxidation ( | 23945 NADH dehydrogenase (Ubiquinone) Fe-S protein 7 OS=Rattus norvegicus OX=10116 GN=Ndufs7 PE=1 SV=1 |
| 45 | 45 | P04762\|CA | 112.9 | 10 | 10 | 1.94E+06 | 9 | 9 | 12 | | 59757 Catalase OS=Rattus norvegicus OX=10116 GN=Cat PE=1 SV=3 |
| 65 | 32 | P07895\|SO | 112.56 | 12 | 12 | 2.03E+06 | 4 | 4 | 7 | | 24674 Superoxide dismutase [Mn] mitochondrial OS=Rattus norvegicus OX=10116 GN=Sod2 PE=1 SV=2 |
| 62 | 198 | tr\|B5DEL8 | 112.53 | 21 | 21 | 2.94E+06 | 5 | 5 | 8 | Carbamido | 12700 NADH dehydrogenase (Ubiquinone) Fe-S protein 5 OS=Rattus norvegicus OX=10116 GN=Ndufs5 PE=1 SV=1 |
| 55 | 346 | tr\|D3ZS58 | 112.17 | 11 | 11 | 5.29E+06 | 4 | 4 | 10 | | 10845 NADH dehydrogenase [ubiquinone] 1 alpha subcomplex subunit 2 OS=Rattus norvegicus OX=10116 GN=Ndufa2 PE=1 SV=1 |
| 42 | 23 | P22791\|HI | 111.6 | 12 | 12 | 1.67E+06 | 11 | 11 | 13 | Carbamidomethylatior | Hydroxymethylglutaryl-CoA synthase, mitochondrial |
| 42 | 22 | tr\|Q68G44 | 111.6 | 12 | 12 | 1.67E+06 | 11 | 11 | 13 | Carbamidomethylatior | Hydroxymethylglutaryl-CoA synthase, mitochondrial |
| 61 | 119 | P31399\|AT | 110.27 | 21 | 21 | 2.18E+06 | 5 | 5 | 8 | | 18763 ATP synthase subunit d mitochondrial OS=Rattus norvegicus OX=10116 GN=Atp5pd PE=1 SV=3 |
| 41 | 171 | tr\|D3ZFJ6\| | 109.77 | 16 | 16 | 2.59E+06 | 9 | 9 | 13 | | 60420 Lactamase beta OS=Rattus norvegicus OX=10116 GN=Lactb PE=1 SV=1 |
| 64 | 5783 | tr\|Q35733 | 104.7 | 14 | 14 | 1.05E+06 | 7 | 7 | 8 | | 8389 NADH-ubiquinone oxidoreductase chain 6 (Fragment) OS=Rattus norvegicus OX=10116 PE=3 SV=1 |
| 64 | 5784 | tr\|Q06QD | 104.7 | 6 | 6 | 1.05E+06 | 7 | 7 | 8 | | 18943 NADH-ubiquinone oxidoreductase chain 6 OS=Rattus norvegicus OX=10116 GN=ND6 PE=3 SV=1 |
| 64 | 5785 | tr\|Q7HKW | 104.7 | 6 | 6 | 1.05E+06 | 7 | 7 | 8 | | 18957 NADH-ubiquinone oxidoreductase chain 6 OS=Rattus norvegicus OX=10116 GN=NADH6 PE=3 SV=1 |
| 52 | 2 | tr\|B2GV15 | 104.24 | 11 | 11 | 2.43E+06 | 6 | 6 | 10 | | 53274 Dihydrolipoamide acetyltransferase component of pyruvate dehydrogenase complex OS=Rattus norvegicus OX=10116 GN=Dbt PE=1 SV=1 |
| 31 | 225 | tr\|Q8SEZ8 | 102.51 | 17 | 17 | 4.35E+06 | 12 | 12 | 17 | Oxidation ( | 36133 NADH-ubiquinone oxidoreductase chain 1 (Fragment) OS=Rattus norvegicus OX=10116 GN=NADH1 PE=3 SV=1 |
| 31 | 226 | P03889\|NU | 102.51 | 17 | 17 | 4.35E+06 | 12 | 12 | 17 | Oxidation ( | 36145 NADH-ubiquinone oxidoreductase chain 1 OS=Rattus norvegicus OX=10116 GN=Mtnd1 PE=1 SV=3 |
| 31 | 227 | tr\|Q8HID1 | 102.51 | 17 | 17 | 4.35E+06 | 12 | 12 | 17 | Oxidation ( | 36145 NADH-ubiquinone oxidoreductase chain 1 OS=Rattus norvegicus OX=10116 GN=Mt-nd1 PE=3 SV=1 |
| 31 | 228 | tr\|D2E6L7 | 102.51 | 17 | 17 | 4.35E+06 | 12 | 12 | 17 | Oxidation ( | 36062 NADH-ubiquinone oxidoreductase chain 1 OS=Rattus norvegicus OX=10116 GN=ND1 PE=3 SV=1 |
| 59 | 46 | P13086\|SU | 99.25 | 8 | 8 | 5.92E+05 | 5 | 5 | 8 | | 36148 Succinate--CoA ligase [ADP/GDP-forming] subunit alpha mitochondrial OS=Rattus norvegicus OX=10116 GN=Suclg1 PE=2 SV=2 |
| 59 | 47 | tr\|A0A0H2 | 99.25 | 7 | 7 | 5.92E+05 | 5 | 5 | 8 | | 37560 Succinate--CoA ligase [ADP/GDP-forming] subunit alpha mitochondrial OS=Rattus norvegicus OX=10116 GN=Suclg1 PE=1 SV=1 |
| 84 | 6 | Q01205\|O | 95.82 | 8 | 8 | 5.58E+05 | 3 | 3 | 3 | | 48925 Dihydrolipoyllysine-residue succinyltransferase component of 2-oxoglutarate dehydrogenase complex mitochondrial OS=Rattus norvegicus OX=10116 GN=Dlst PE=1 SV=2 |
| 84 | 5 | tr\|G3V6P2 | 95.82 | 8 | 8 | 5.58E+05 | 3 | 3 | 3 | | 48899 Dihydrolipoamide S-succinyltransferase (E2 component of 2-oxo-glutarate complex) isoform CRA_a OS=Rattus norvegicus OX=10116 GN=Dlst PE=1 SV=1 |
| 74 | 115 | tr\|A0A0G2 | 90.57 | 3 | 3 | 6.85E+05 | 4 | 4 | 4 | | 86230 MICOS complex subunit MIC60 OS=Rattus norvegicus OX=10116 GN=Immt PE=1 SV=1 |
| 74 | 109 | tr\|A0A140 | 90.57 | 4 | 4 | 6.85E+05 | 4 | 4 | 4 | | 67049 MICOS complex subunit MIC60 OS=Rattus norvegicus OX=10116 GN=Immt PE=1 SV=1 |
| 74 | 110 | Q3KR86\|M | 90.57 | 4 | 4 | 6.85E+05 | 4 | 4 | 4 | | 67177 MICOS complex subunit Mic60 (Fragment) OS=Rattus norvegicus OX=10116 GN=Immt PE=1 SV=1 |
| 50 | 133 | Q63362\|N | 88.02 | 30 | 30 | 1.23E+06 | 8 | 8 | 11 | Carbamido | 13412 NADH dehydrogenase [ubiquinone] 1 alpha subcomplex subunit 5 OS=Rattus norvegicus OX=10116 GN=Ndufa5 PE=1 SV=3 |
| 75 | 5779 | tr\|D3ZYT2 | 85.53 | 3 | 3 | 5.29E+05 | 2 | 2 | 4 | | 48136 Mitochondrial ribosomal protein S5 OS=Rattus norvegicus OX=10116 GN=Mrps5 PE=1 SV=1 |
| 111 | 277 | tr\|B2RZ24 | 84.12 | 4 | 4 | 3.40E+05 | 2 | 2 | 2 | | 47388 Succinate-CoA ligase subunit beta (Fragment) OS=Rattus norvegicus OX=10116 GN=Sucla2 PE=2 SV=1 |
| 111 | 278 | tr\|F1LM47 | 84.12 | 3 | 3 | 3.40E+05 | 2 | 2 | 2 | | 50306 Succinate--CoA ligase [ADP-forming] subunit beta mitochondrial OS=Rattus norvegicus OX=10116 GN=Sucla2 PE=1 SV=1 |
| 72 | 385 | Q5U2U0\|C | 81.44 | 5 | 5 | 2.44E+05 | 3 | 3 | 4 | | 69208 ATP-dependent Clp protease ATP-binding subunit clpX-like mitochondrial OS=Rattus norvegicus OX=10116 GN=Clpx PE=2 SV=1 |
| 63 | 50 | Q02253\|M | 81.16 | 9 | 9 | 6.89E+05 | 7 | 7 | 8 | Oxidation ( | 57808 Methylmalonate-semialdehyde dehydrogenase [acylating] mitochondrial OS=Rattus norvegicus OX=10116 GN=Aldh6a1 PE=1 SV=1 |
| 63 | 51 | tr\|G3V7J0 | 81.16 | 9 | 9 | 6.89E+05 | 7 | 7 | 8 | Oxidation ( | 57748 Aldehyde dehydrogenase family 6 subfamily A1 isoform CRA_b OS=Rattus norvegicus OX=10116 GN=Aldh6a1 PE=1 SV=1 |

| | | | | | | | | | | | | |
|---|---|---|---|---|---|---|---|---|---|---|---|---|
| 43 | 147 | P05508\|NU | 78.18 | 8 | 8 | 2.13E+06 | 7 | 7 | 12 | Oxidation (M) | | NADH-ubiquinone oxidoreductase chain 4 |
| 43 | 148 | tr\|D2E6K0 | 78.18 | 8 | 8 | 2.13E+06 | 7 | 7 | 12 | Oxidation (M) | | |
| 43 | 149 | tr\|A7XYB9 | 78.18 | 8 | 8 | 2.13E+06 | 7 | 7 | 12 | Oxidation (M) | | |
| 43 | 150 | tr\|Q06QE1 | 78.18 | 8 | 8 | 2.13E+06 | 7 | 7 | 12 | Oxidation (M) | | |
| 43 | 151 | tr\|Q06QA2 | 78.18 | 8 | 8 | 2.13E+06 | 7 | 7 | 12 | Oxidation (M) | | |
| 43 | 152 | tr\|Q7HKW | 78.18 | 8 | 8 | 2.13E+06 | 7 | 7 | 12 | Oxidation (M) | | |
| 43 | 153 | tr\|Q06QG7 | 78.18 | 8 | 8 | 2.13E+06 | 7 | 7 | 12 | Oxidation (M) | | |
| 43 | 154 | tr\|Q06Q89 | 78.18 | 8 | 8 | 2.13E+06 | 7 | 7 | 12 | Oxidation (M) | | |
| 43 | 155 | tr\|Q35737 | 78.18 | 8 | 8 | 2.13E+06 | 7 | 7 | 12 | Oxidation (M) | | |
| 43 | 156 | tr\|Q8HIC6 | 78.18 | 8 | 8 | 2.13E+06 | 7 | 7 | 12 | Oxidation (M) | | |
| 78 | 52 | tr\|M0RAM | 77.53 | 18 | 18 | 5.82E+05 | 4 | 4 | 4 | | 22155 | Glutathione peroxidase OS=Rattus norvegicus OX=10116 GN=Gpx1 PE=1 SV=1 |
| 78 | 53 | P04041\|GP | 77.53 | 18 | 18 | 5.82E+05 | 4 | 4 | 4 | | 22305 | Glutathione peroxidase 1 OS=Rattus norvegicus OX=10116 GN=Gpx1 PE=1 SV=4 |
| 69 | 54 | P18163\|AC | 77.48 | 4 | 4 | 9.19E+05 | 3 | 2 | 6 | | 78179 | Long-chain-fatty-acid--CoA ligase 1 OS=Rattus norvegicus OX=10116 GN=Acsl1 PE=1 SV=1 |
| 67 | 207 | tr\|G3V6I4\| | 75.45 | 10 | 10 | 2.72E+05 | 4 | 4 | 6 | | 37791 | Mitochondrial amidoxime reducing component 1 OS=Rattus norvegicus OX=10116 GN=Marc1 PE=1 SV=1 |
| 66 | 267 | tr\|Q06Q97 | 73.13 | 8 | 8 | 6.15E+05 | 5 | 5 | 7 | | 38485 | NADH-ubiquinone oxidoreductase chain 2 OS=Rattus norvegicus OX=10116 GN=ND2 PE=3 SV=1 |
| 66 | 268 | tr\|A0A0A1 | 73.13 | 8 | 8 | 6.15E+05 | 5 | 5 | 7 | | 38542 | NADH-ubiquinone oxidoreductase chain 2 OS=Rattus norvegicus OX=10116 GN=ND2 PE=3 SV=1 |
| 66 | 269 | tr\|Q5UAJ8 | 73.13 | 8 | 8 | 6.15E+05 | 5 | 5 | 7 | | 38455 | NADH-ubiquinone oxidoreductase chain 2 OS=Rattus norvegicus OX=10116 GN=ND2 PE=3 SV=1 |
| 66 | 270 | tr\|Q8HID0 | 73.13 | 8 | 8 | 6.15E+05 | 5 | 5 | 7 | | 38653 | NADH-ubiquinone oxidoreductase chain 2 OS=Rattus norvegicus OX=10116 GN=Mt-nd2 PE=3 SV=1 |
| 66 | 271 | P11662\|NU | 73.13 | 8 | 8 | 6.15E+05 | 5 | 5 | 7 | | 38653 | NADH-ubiquinone oxidoreductase chain 2 OS=Rattus norvegicus OX=10116 GN=Mtnd2 PE=3 SV=3 |
| 66 | 272 | tr\|Q06QH5 | 73.13 | 8 | 8 | 6.15E+05 | 5 | 5 | 7 | | 38626 | NADH-ubiquinone oxidoreductase chain 2 OS=Rattus norvegicus OX=10116 GN=ND2 PE=3 SV=1 |
| 66 | 274 | tr\|Q06QB0 | 73.13 | 8 | 8 | 6.15E+05 | 5 | 5 | 7 | | 38534 | NADH-ubiquinone oxidoreductase chain 2 OS=Rattus norvegicus OX=10116 GN=ND2 PE=3 SV=1 |
| 66 | 275 | tr\|D2E6P4 | 73.13 | 8 | 8 | 6.15E+05 | 5 | 5 | 7 | | 38623 | NADH-ubiquinone oxidoreductase chain 2 OS=Rattus norvegicus OX=10116 GN=ND2 PE=3 SV=1 |
| 66 | 276 | tr\|Q06QE9 | 73.13 | 8 | 8 | 6.15E+05 | 5 | 5 | 7 | | 38598 | NADH-ubiquinone oxidoreductase chain 2 OS=Rattus norvegicus OX=10116 GN=ND2 PE=3 SV=1 |
| 66 | 273 | tr\|Q8SEZ7 | 73.13 | 8 | 8 | 6.15E+05 | 5 | 5 | 7 | | 38580 | NADH-ubiquinone oxidoreductase chain 2 (Fragment) OS=Rattus norvegicus OX=10116 GN=NADH2 PE=3 SV=1 |
| 71 | 246 | P19511\|AT | 72.84 | 9 | 9 | 4.66E+06 | 3 | 3 | 5 | | 28869 | ATP synthase F(0) complex subunit B1 mitochondrial OS=Rattus norvegicus OX=10116 GN=Atp5pb PE=1 SV=1 |
| 73 | 190 | P07824\|AR | 72.36 | 11 | 11 | 1.59E+06 | 3 | 3 | 4 | | 34973 | Arginase-1 OS=Rattus norvegicus OX=10116 GN=Arg1 PE=1 SV=2 |
| 53 | 235 | tr\|B2RYW3 | 70.71 | 19 | 19 | 3.44E+06 | 4 | 4 | 10 | | 21892 | NADH dehydrogenase (Ubiquinone) 1 beta subcomplex 9 OS=Rattus norvegicus OX=10116 GN=Ndufb9 PE=1 SV=1 |
| 54 | 134 | tr\|B2RYS2 | 70.53 | 24 | 24 | 5.92E+06 | 6 | 5 | 10 | | 13559 | Cytochrome b-c1 complex subunit 7 OS=Rattus norvegicus OX=10116 GN=Uqcrb PE=1 SV=1 |
| 81 | 222 | tr\|Q4G067 | 69.99 | 4 | 4 | 9.15E+05 | 2 | 2 | 4 | | 37440 | Mitochondrial ribosomal protein L44 OS=Rattus norvegicus OX=10116 GN=Mrpl44 PE=1 SV=1 |
| 57 | 286 | Q7TQ16\|Q | 65.36 | 18 | 18 | 2.54E+06 | 5 | 5 | 9 | | 9849 | Cytochrome b-c1 complex subunit 8 OS=Rattus norvegicus OX=10116 GN=Uqcrq PE=3 SV=1 |
| 98 | 218 | P63039\|CH | 64.67 | 6 | 6 | 1.37E+06 | 3 | 3 | 3 | Oxidation ( | 60956 | 60 kDa heat shock protein mitochondrial OS=Rattus norvegicus OX=10116 GN=Hspd1 PE=1 SV=1 |
| 98 | 219 | tr\|A0A482 | 64.67 | 6 | 6 | 1.37E+06 | 3 | 3 | 3 | Oxidation ( | 60956 | Hsp60 OS=Rattus norvegicus OX=10116 GN=Hspd1 PE=2 SV=1 |
| 80 | 5780 | tr\|D4A4B1 | 63.07 | 8 | 8 | 1.84E+05 | 4 | 4 | 4 | | 23223 | Mitochondrial ribosomal protein L15 OS=Rattus norvegicus OX=10116 GN=Mrpl15 PE=1 SV=2 |
| 80 | 368 | tr\|A0A0G2 | 63.07 | 5 | 5 | 1.84E+05 | 4 | 4 | 4 | | 33652 | Mitochondrial ribosomal protein L15 OS=Rattus norvegicus OX=10116 GN=Mrpl15 PE=1 SV=1 |
| 76 | 111 | tr\|S5RZM8 | 59.89 | 7 | 7 | 7.06E+05 | 3 | 3 | 4 | | 25958 | Cytochrome c oxidase subunit 2 OS=Rattus norvegicus OX=10116 GN=COX2 PE=3 SV=1 |
| 76 | 112 | P00406\|CO | 59.89 | 7 | 7 | 7.06E+05 | 3 | 3 | 4 | | 25928 | Cytochrome c oxidase subunit 2 OS=Rattus norvegicus OX=10116 GN=Mtco2 PE=1 SV=3 |
| 76 | 104 | tr\|Q5UAJ6 | 59.89 | 7 | 7 | 7.06E+05 | 3 | 3 | 4 | | 25942 | Cytochrome c oxidase subunit 2 OS=Rattus norvegicus OX=10116 GN=COX2 PE=3 SV=1 |
| 76 | 113 | tr\|Q8SEZ5 | 59.89 | 7 | 7 | 7.06E+05 | 3 | 3 | 4 | | 25928 | Cytochrome c oxidase subunit 2 OS=Rattus norvegicus OX=10116 GN=Mt-co2 PE=1 SV=1 |
| 76 | 114 | tr\|A0A097 | 59.89 | 7 | 7 | 7.06E+05 | 3 | 3 | 4 | | 25894 | Cytochrome c oxidase subunit 2 OS=Rattus norvegicus OX=10116 GN=COX2 PE=3 SV=1 |
| 68 | 14 | tr\|F1LN92 | 59.26 | 5 | 5 | 4.87E+06 | 6 | 6 | 6 | Oxidation ( | 89352 | AFG3-like matrix AAA peptidase subunit 2 OS=Rattus norvegicus OX=10116 GN=Afg3l2 PE=1 SV=1 |
| 70 | 414 | tr\|D4A4P3 | 53.7 | 22 | 22 | 1.97E+06 | 3 | 3 | 6 | Oxidation ( | 11267 | NADH:ubiquinone oxidoreductase subunit B3 OS=Rattus norvegicus OX=10116 GN=Ndufb3 PE=1 SV=1 |
| 122 | 253 | Q9ER34\|AC | 53.06 | 2 | 2 | 2.98E+05 | 2 | 2 | 2 | | 85433 | Aconitate hydratase mitochondrial OS=Rattus norvegicus OX=10116 GN=Aco2 PE=1 SV=2 |
| 261 | 287 | tr\|D3ZX69 | 47.58 | 5 | 5 | 3.38E+05 | 1 | 1 | 1 | | 29466 | 39S ribosomal protein L10 mitochondrial OS=Rattus norvegicus OX=10116 GN=Mrpl10 PE=1 SV=1 |
| 261 | 223 | P0C2C4\|RM | 47.58 | 5 | 5 | 3.38E+05 | 1 | 1 | 1 | | 29940 | 39S ribosomal protein L10 mitochondrial OS=Rattus norvegicus OX=10116 GN=Mrpl10 PE=1 SV=1 |
| 133 | 146 | tr\|B0BMT9 | 44.76 | 4 | 4 | 2.46E+05 | 2 | 2 | 2 | | 50202 | Sqrdl protein OS=Rattus norvegicus OX=10116 GN=Sqor PE=1 SV=1 |
| 93 | 265 | Q63750\|RI | 44.68 | 19 | 19 | 3.64E+05 | 3 | 3 | 3 | | 17050 | 39S ribosomal protein L23 mitochondrial OS=Rattus norvegicus OX=10116 GN=Mrpl23 PE=2 SV=1 |
| 99 | 424 | Q9Z0W7\|C | 44.57 | 5 | 5 | 8.15E+04 | 3 | 1 | 3 | | 28633 | Chloride intracellular channel protein 4 OS=Rattus norvegicus OX=10116 GN=Clic4 PE=1 SV=3 |
| 99 | 425 | tr\|G3V8C4 | 44.57 | 5 | 5 | 8.15E+04 | 3 | 1 | 3 | | 28669 | Chloride intracellular channel protein OS=Rattus norvegicus OX=10116 GN=Clic4 PE=1 SV=1 |
| 79 | 59 | P17178\|CP | 43.75 | 5 | 5 | 5.96E+05 | 3 | 2 | 4 | | 60733 | Sterol 26-hydroxylase mitochondrial OS=Rattus norvegicus OX=10116 GN=Cyp27a1 PE=1 SV=1 |
| 79 | 60 | tr\|A0A0H2 | 43.75 | 5 | 5 | 5.96E+05 | 3 | 2 | 4 | | 62492 | RCG24013 isoform CRA_a OS=Rattus norvegicus OX=10116 GN=Cyp27a1 PE=1 SV=1 |
| 131 | 210 | tr\|Q5M949 | 41.6 | 6 | 6 | 5.63E+05 | 2 | 2 | 2 | | 28340 | Nipsnap homolog 3A (C. elegans) OS=Rattus norvegicus OX=10116 GN=Nipsnap3b PE=1 SV=1 |
| 112 | 341 | tr\|G3V6I5\| | 40.45 | 3 | 3 | 1.38E+05 | 1 | 1 | 2 | | 46777 | DnaJ heat shock protein family (Hsp40) member A3 OS=Rattus norvegicus OX=10116 GN=Dnaja3 PE=1 SV=2 |
| 112 | 342 | tr\|Q2TVU3 | 40.45 | 3 | 3 | 1.38E+05 | 1 | 1 | 2 | | 49412 | TID1 OS=Rattus norvegicus OX=10116 GN=Dnaja3 PE=2 SV=1 |
| 112 | 343 | tr\|A0A0G2 | 40.45 | 3 | 3 | 1.38E+05 | 1 | 1 | 2 | | 49416 | DnaJ heat shock protein family (Hsp40) member A3 OS=Rattus norvegicus OX=10116 GN=Dnaja3 PE=1 SV=1 |
| 112 | 344 | tr\|Q2UZS7 | 40.45 | 2 | 2 | 1.38E+05 | 1 | 1 | 2 | | 52399 | Tid-1 long isoform OS=Rattus norvegicus OX=10116 GN=Dnaja3 PE=2 SV=1 |
| 112 | 345 | tr\|A0A0G2 | 40.45 | 2 | 2 | 1.38E+05 | 1 | 1 | 2 | | 52403 | DnaJ heat shock protein family (Hsp40) member A3 OS=Rattus norvegicus OX=10116 GN=Dnaja3 PE=1 SV=1 |
| 132 | 216 | Q9R063\|PI | 40.42 | 9 | 9 | 2.17E+05 | 2 | 2 | 2 | | 22179 | Peroxiredoxin-5 mitochondrial OS=Rattus norvegicus OX=10116 GN=Prdx5 PE=1 SV=1 |
| 132 | 217 | tr\|A0A0G2 | 40.42 | 9 | 9 | 2.17E+05 | 2 | 2 | 2 | | 22207 | Peroxiredoxin OS=Rattus norvegicus OX=10116 GN=Prdx5 PE=1 SV=1 |
| 134 | 379 | tr\|D3ZDP2 | 40.31 | 4 | 4 | 4.28E+05 | 2 | 2 | 2 | | 23470 | Mitochondrial ribosomal protein L58 OS=Rattus norvegicus OX=10116 GN=Mrpl58 PE=1 SV=1 |
| 136 | 322 | P12075\|CO | 39.16 | 9 | 9 | 5.96E+05 | 2 | 2 | 2 | | 13915 | Cytochrome c oxidase subunit 5B mitochondrial OS=Rattus norvegicus OX=10116 GN=Cox5b PE=1 SV=2 |
| 262 | 224 | tr\|D3ZXF9 | 38.96 | 4 | 4 | 8.79E+04 | 1 | 1 | 1 | | 29441 | Mitochondrial ribosomal protein L12 OS=Rattus norvegicus OX=10116 GN=Mrpl12 PE=1 SV=1 |
| 120 | 143 | tr\|Q5EBA4 | 38.62 | 5 | 5 | 5.36E+05 | 2 | 2 | 2 | | 33215 | Nipsnap1 protein (Fragment) OS=Rattus norvegicus OX=10116 GN=Nipsnap1 PE=2 SV=1 |
| 120 | 144 | tr\|G3V728 | 38.62 | 5 | 5 | 5.36E+05 | 2 | 2 | 2 | | 33346 | 4-nitrophenylphosphatase domain and non-neuronal SNAP25-like protein homolog 1 (C. elegans) isoform CRA_b OS=Rattus norvegicus OX=10116 GN=Nipsnap1 PE=1 SV=1 |

| | | | | | | | | | | | | |
|---|---|---|---|---|---|---|---|---|---|---|---|---|
| 108 | 324 | P11497\|AC | 38.48 | 1 | 1 | 1.08E+05 | 2 | 1 | 2 | | 265191 | Acetyl-CoA carboxylase 1 OS=Rattus norvegicus OX=10116 GN=Acaca PE=1 SV=1 |
| 89 | 725 | tr\|F1LP30\| | 38.27 | 3 | 3 | 2.14E+04 | 2 | 1 | 3 | | 79296 | Methylcrotonoyl-CoA carboxylase subunit alpha mitochondrial OS=Rattus norvegicus OX=10116 GN=Mccc1 PE=1 SV=1 |
| 89 | 726 | Q5I0C3\|M | 38.27 | 3 | 3 | 2.14E+04 | 2 | 1 | 3 | | 79330 | Methylcrotonoyl-CoA carboxylase subunit alpha mitochondrial OS=Rattus norvegicus OX=10116 GN=Mccc1 PE=1 SV=1 |
| 119 | 195 | P49432\|OI | 37.65 | 5 | 5 | 9.28E+04 | 2 | 2 | 2 | | 38982 | Pyruvate dehydrogenase E1 component subunit beta mitochondrial OS=Rattus norvegicus OX=10116 GN=Pdhb PE=1 SV=2 |
| 119 | 196 | tr\|A0A0G2 | 37.65 | 4 | 4 | 9.28E+04 | 2 | 2 | 2 | | 46193 | Pyruvate dehydrogenase E1 component subunit beta OS=Rattus norvegicus OX=10116 GN=Pdhb PE=1 SV=1 |
| 171 | 306 | Q2V057\|H | 36.8 | 2 | 2 | 7.19E+04 | 1 | 1 | 1 | | 51002 | Hydroxyproline dehydrogenase OS=Rattus norvegicus OX=10116 GN=Prodh2 PE=2 SV=1 |
| 100 | 366 | P33124\|AC | 36.34 | 3 | 3 | 1.70E+05 | 2 | 1 | 3 | Formylatio | 78180 | Long-chain-fatty-acid--CoA ligase 6 OS=Rattus norvegicus OX=10116 GN=Acsl6 PE=1 SV=1 |
| 92 | 375 | tr\|A0A0G2 | 36.28 | 4 | 4 | 8.64E+05 | 2 | 2 | 3 | | 45197 | Glycine C-acetyltransferase OS=Rattus norvegicus OX=10116 GN=Gcat PE=1 SV=1 |
| 123 | 289 | tr\|Q5XFW4 | 35.43 | 7 | 7 | 2.75E+05 | 1 | 1 | 2 | | 20541 | Mitochondrial ribosomal protein L13 OS=Rattus norvegicus OX=10116 GN=Mrpl13 PE=1 SV=1 |
| 225 | 5798 | tr\|Q8SEZ1 | 35.15 | 6 | 6 | 0 | 1 | 1 | 1 | | 13087 | NADH-ubiquinone oxidoreductase chain 3 OS=Rattus norvegicus OX=10116 GN=Mt-nd3 PE=2 SV=1 |
| 225 | 5799 | P05506\|NU | 35.15 | 6 | 6 | 0 | 1 | 1 | 1 | | 13087 | NADH-ubiquinone oxidoreductase chain 3 OS=Rattus norvegicus OX=10116 GN=Mtnd3 PE=3 SV=3 |
| 225 | 5800 | tr\|A0A0A1 | 35.15 | 6 | 6 | 0 | 1 | 1 | 1 | | 13071 | NADH-ubiquinone oxidoreductase chain 3 OS=Rattus norvegicus OX=10116 GN=ND3 PE=3 SV=1 |
| 225 | 5801 | tr\|Q06QG9 | 35.15 | 6 | 6 | 0 | 1 | 1 | 1 | | 13075 | NADH-ubiquinone oxidoreductase chain 3 OS=Rattus norvegicus OX=10116 GN=ND3 PE=3 SV=1 |
| 95 | 656 | tr\|D3ZD23 | 34.78 | 1 | 1 | 4.81E+05 | 2 | 2 | 3 | | 67300 | ATP-binding cassette subfamily E member 1 OS=Rattus norvegicus OX=10116 GN=Abce1 PE=1 SV=1 |
| 263 | 5797 | tr\|D4A830 | 34.64 | 3 | 3 | 1.31E+05 | 1 | 1 | 1 | | 35047 | Inorganic pyrophosphatase 2 OS=Rattus norvegicus OX=10116 GN=Ppa2 PE=1 SV=2 |
| 101 | 446 | Q91XJ1\|BE | 34.45 | 2 | 2 | | 2 | 0 | 3 | | 51557 | Beclin-1 OS=Rattus norvegicus OX=10116 GN=Becn1 PE=1 SV=1 |
| 219 | 5790 | Q66H47\|R | 34.33 | 6 | 6 | 9.40E+04 | 1 | 1 | 1 | | 25001 | 39S ribosomal protein L24 mitochondrial OS=Rattus norvegicus OX=10116 GN=Mrpl24 PE=2 SV=1 |
| 265 | 367 | tr\|D3ZXF8 | 34.13 | 5 | 5 | 1.66E+05 | 1 | 1 | 1 | | 17575 | Mitochondrial ribosomal protein L43 OS=Rattus norvegicus OX=10116 GN=Mrpl43 PE=1 SV=3 |
| 266 | 211 | P70584\|AC | 33.43 | 2 | 2 | 1.48E+05 | 1 | 1 | 1 | | 47824 | Short/branched chain specific acyl-CoA dehydrogenase mitochondrial OS=Rattus norvegicus OX=10116 GN=Acadsb PE=1 SV=1 |
| 266 | 212 | tr\|A0A0A0 | 33.43 | 2 | 2 | 1.48E+05 | 1 | 1 | 1 | | 52043 | Short/branched chain-specific acyl-CoA dehydrogenase mitochondrial OS=Rattus norvegicus OX=10116 GN=Acadsb PE=1 SV=2 |
| 217 | 5788 | tr\|Q3B8N9 | 33.32 | 3 | 3 | 2.26E+05 | 1 | 1 | 1 | | 32823 | Biphenyl hydrolase-like OS=Rattus norvegicus OX=10116 GN=Bphl PE=1 SV=1 |
| 264 | 264 | tr\|D3ZTW8 | 33.17 | 8 | 8 | 0 | 1 | 1 | 1 | | 15823 | Mitochondrial ribosomal protein L27 OS=Rattus norvegicus OX=10116 GN=Mrpl27 PE=1 SV=2 |
| 124 | 349 | Q5I0C5\|FN | 32.97 | 2 | 2 | 7.67E+05 | 1 | 1 | 2 | Formylatio | 42997 | Methionyl-tRNA formyltransferase mitochondrial OS=Rattus norvegicus OX=10116 GN=Mtfmt PE=2 SV=1 |
| 267 | 5802 | Q06647\|AT | 32.88 | 4 | 4 | 7.76E+04 | 1 | 1 | 1 | | 23398 | ATP synthase subunit O mitochondrial OS=Rattus norvegicus OX=10116 GN=Atp5po PE=1 SV=1 |
| 190 | 208 | P13803\|ET | 32.42 | 3 | 3 | 4.17E+05 | 1 | 1 | 1 | | 34951 | Electron transfer flavoprotein subunit alpha mitochondrial OS=Rattus norvegicus OX=10116 GN=Etfa PE=1 SV=4 |
| 269 | 312 | tr\|D3ZGM1 | 31.96 | 1 | 1 | 1.00E+05 | 1 | 1 | 1 | | 77891 | Pentatricopeptide repeat domain 3 OS=Rattus norvegicus OX=10116 GN=Ptcd3 PE=1 SV=1 |
| 269 | 313 | tr\|A0A0G2 | 31.96 | 1 | 1 | 1.00E+05 | 1 | 1 | 1 | | 84557 | Pentatricopeptide repeat domain 3 OS=Rattus norvegicus OX=10116 GN=Ptcd3 PE=1 SV=1 |
| 270 | 448 | tr\|A0A0G2 | 31.81 | 2 | 2 | 9.61E+04 | 1 | 1 | 1 | | 41504 | Solute carrier family 25 member 12 OS=Rattus norvegicus OX=10116 GN=Slc25a12 PE=1 SV=1 |
| 270 | 333 | tr\|F1LZW6 | 31.81 | 2 | 2 | 9.61E+04 | 1 | 1 | 1 | | 54099 | Solute carrier family 25 member 13 OS=Rattus norvegicus OX=10116 GN=Slc25a13 PE=1 SV=2 |
| 270 | 456 | tr\|F1LX07\| | 31.81 | 1 | 1 | 9.61E+04 | 1 | 1 | 1 | | 71917 | Solute carrier family 25 member 12 OS=Rattus norvegicus OX=10116 GN=Slc25a12 PE=1 SV=3 |
| 191 | 5781 | Q5XIC0\|EC | 30.23 | 2 | 2 | 0 | 1 | 1 | 1 | | 43021 | Enoyl-CoA delta isomerase 2 mitochondrial OS=Rattus norvegicus OX=10116 GN=Eci2 PE=1 SV=1 |
| 191 | 5793 | tr\|Q5M884 | 30.23 | 3 | 3 | 0 | 1 | 1 | 1 | | 33686 | Enoyl-CoA delta isomerase 2 mitochondrial OS=Rattus norvegicus OX=10116 GN=Eci3 PE=1 SV=1 |
| 191 | 5794 | tr\|A0A0G2 | 30.23 | 2 | 2 | 0 | 1 | 1 | 1 | | 42325 | Enoyl-CoA delta isomerase 2 mitochondrial OS=Rattus norvegicus OX=10116 GN=Eci3 PE=1 SV=1 |
| 125 | 1095 | tr\|Q5UAJ5 | 29.81 | 13 | 13 | 7.13E+05 | 1 | 1 | 2 | | 7642 | ATP synthase protein 8 OS=Rattus norvegicus OX=10116 GN=ATP8 PE=3 SV=1 |
| 125 | 1096 | P11608\|AT | 29.81 | 13 | 13 | 7.13E+05 | 1 | 1 | 2 | | 7630 | ATP synthase protein 8 OS=Rattus norvegicus OX=10116 GN=Mt-atp8 PE=1 SV=2 |
| 125 | 1097 | tr\|Q8SEZ4 | 29.81 | 13 | 13 | 7.13E+05 | 1 | 1 | 2 | | 7632 | ATP synthase protein 8 OS=Rattus norvegicus OX=10116 GN=ATPase8 PE=3 SV=1 |
| 125 | 1098 | tr\|Q8HIC8 | 29.81 | 13 | 13 | 7.13E+05 | 1 | 1 | 2 | | 7630 | ATP synthase protein 8 OS=Rattus norvegicus OX=10116 GN=Mt-atp8 PE=3 SV=1 |
| 109 | 61 | P29147\|BD | 29.73 | 3 | 3 | 7.40E+04 | 2 | 2 | 2 | | 38202 | D-beta-hydroxybutyrate dehydrogenase mitochondrial OS=Rattus norvegicus OX=10116 GN=Bdh1 PE=1 SV=2 |
| 109 | 62 | tr\|A0A0G2 | 29.73 | 3 | 3 | 7.40E+04 | 2 | 2 | 2 | | 38333 | 3-hydroxybutyrate dehydrogenase type 1 isoform CRA_a OS=Rattus norvegicus OX=10116 GN=Bdh1 PE=1 SV=1 |
| 271 | 5804 | tr\|A0A0G2 | 29.69 | 1 | 1 | 0 | 1 | 1 | 1 | | 79812 | Propionyl-CoA carboxylase alpha chain mitochondrial OS=Rattus norvegicus OX=10116 GN=Pcca PE=1 SV=1 |
| 271 | 5805 | tr\|A0A0G2 | 29.69 | 1 | 1 | 0 | 1 | 1 | 1 | | 81413 | Propionyl-CoA carboxylase alpha chain mitochondrial OS=Rattus norvegicus OX=10116 GN=Pcca PE=1 SV=1 |
| 271 | 5806 | P14882\|PC | 29.69 | 1 | 1 | 0 | 1 | 1 | 1 | | 81623 | Propionyl-CoA carboxylase alpha chain mitochondrial OS=Rattus norvegicus OX=10116 GN=Pcca PE=1 SV=3 |
| 104 | 595 | Q9Z286\|AI | 29.49 | 1 | 1 | 0 | 2 | 1 | 2 | | 185852 | Adenylate cyclase type 10 OS=Rattus norvegicus OX=10116 GN=Adcy10 PE=1 SV=1 |
| 90 | 5782 | Q09073\|AI | 29.26 | 11 | 11 | 1.05E+06 | 3 | 3 | 3 | Formylatio | 32901 | ADP/ATP translocase 2 OS=Rattus norvegicus OX=10116 GN=Slc25a5 PE=1 SV=3 |
| 220 | 5791 | tr\|F1LSY7\| | 29.22 | 1 | 1 | 1.62E+05 | 1 | 1 | 1 | | 109031 | Endoplasmic reticulum to nucleus-signaling 1 OS=Rattus norvegicus OX=10116 GN=Ern1 PE=4 SV=2 |
| 220 | 5792 | tr\|A0A0G2 | 29.22 | 1 | 1 | 1.62E+05 | 1 | 1 | 1 | | 110150 | Endoplasmic reticulum to nucleus-signaling 1 OS=Rattus norvegicus OX=10116 GN=Ern1 PE=2 SV=1 |
| 223 | 369 | tr\|G3V734 | 28.95 | 3 | 3 | 1.60E+05 | 1 | 1 | 1 | | 36133 | 2 4-dienoyl CoA reductase 1 mitochondrial isoform CRA_a OS=Rattus norvegicus OX=10116 GN=Decr1 PE=1 SV=1 |
| 223 | 370 | Q64591\|D | 28.95 | 3 | 3 | 1.60E+05 | 1 | 1 | 1 | | 36133 | 2 4-dienoyl-CoA reductase mitochondrial OS=Rattus norvegicus OX=10116 GN=Decr1 PE=1 SV=2 |
| 218 | 140 | tr\|F1LN88 | 28.91 | 2 | 2 | 3.16E+05 | 1 | 1 | 1 | | 56517 | Aldehyde dehydrogenase mitochondrial OS=Rattus norvegicus OX=10116 GN=Aldh2 PE=1 SV=2 |
| 218 | 141 | P11884\|AL | 28.91 | 2 | 2 | 3.16E+05 | 1 | 1 | 1 | | 56488 | Aldehyde dehydrogenase mitochondrial OS=Rattus norvegicus OX=10116 GN=Aldh2 PE=1 SV=1 |
| 222 | 5796 | B1WC61\|A | 28.73 | 2 | 2 | 8.10E+04 | 1 | 1 | 1 | | 68843 | Complex I assembly factor ACAD9 mitochondrial OS=Rattus norvegicus OX=10116 GN=Acad9 PE=1 SV=1 |
| 196 | 907 | P70705\|AT | 28.38 | 1 | 1 | 0 | 1 | 1 | 1 | | 162092 | Copper-transporting ATPase 1 OS=Rattus norvegicus OX=10116 GN=Atp7a PE=1 SV=1 |
| 172 | 1044 | tr\|D4ACE9 | 28.05 | 2 | 2 | 7.45E+04 | 1 | 1 | 1 | | 103115 | Alpha-aminoadipic semialdehyde synthase mitochondrial OS=Rattus norvegicus OX=10116 GN=Aass PE=1 SV=3 |
| 172 | 742 | A2VCW9\|A | 28.05 | 2 | 2 | 7.45E+04 | 1 | 1 | 1 | | 102908 | Alpha-aminoadipic semialdehyde synthase mitochondrial OS=Rattus norvegicus OX=10116 GN=Aass PE=2 SV=1 |
| 115 | 283 | Q9WVK3\|F | 27.03 | 4 | 4 | 2.97E+05 | 1 | 1 | 2 | | 32433 | Peroxisomal trans-2-enoyl-CoA reductase OS=Rattus norvegicus OX=10116 GN=Pecr PE=2 SV=1 |
| 115 | 284 | tr\|A0A0G2 | 27.03 | 4 | 4 | 2.97E+05 | 1 | 1 | 2 | | 32737 | Peroxisomal trans-2-enoyl-CoA reductase OS=Rattus norvegicus OX=10116 GN=Pecr PE=1 SV=1 |
| 221 | 5795 | tr\|A9UMV | 26.45 | 11 | 11 | 9.68E+04 | 1 | 1 | 1 | | 6539 | RCG29512 OS=Rattus norvegicus OX=10116 GN=Uqcr11 PE=1 SV=1 |
| 106 | 1081 | tr\|Q5XHZ7 | 26.31 | 2 | 2 | 1.31E+05 | 2 | 1 | 2 | | 66660 | MMR_HSR1 domain containing protein RGD1359460 OS=Rattus norvegicus OX=10116 GN=Noa1 PE=2 SV=1 |
| 106 | 1082 | tr\|A0A0G2 | 26.31 | 2 | 2 | 1.31E+05 | 2 | 1 | 2 | | 66708 | Nitric oxide-associated 1 OS=Rattus norvegicus OX=10116 GN=Noa1 PE=4 SV=1 |
| 106 | 1089 | tr\|A0A0G2 | 26.31 | 2 | 2 | 1.31E+05 | 2 | 1 | 2 | | 77504 | MMR_HSR1 domain containing protein RGD1359460 isoform CRA_a OS=Rattus norvegicus OX=10116 GN=Noa1 PE=4 SV=1 |
| 142 | 5823 | Q68FU4\|SU | 26.28 | 2 | 2 | | 2 | 0 | 2 | | 47549 | Succinate--hydroxymethylglutarate CoA-transferase OS=Rattus norvegicus OX=10116 GN=Sugct PE=2 SV=1 |
| 137 | 1384 | P97615\|TH | 26.16 | 7 | 7 | 9.74E+04 | 2 | 1 | 2 | | 18232 | Thioredoxin mitochondrial OS=Rattus norvegicus OX=10116 GN=Txn2 PE=2 SV=1 |
| 114 | 5786 | tr\|D4A5P3 | 25.95 | 3 | 3 | 0 | 2 | 1 | 2 | | 59813 | Mitoguardin 1 OS=Rattus norvegicus OX=10116 GN=Miga1 PE=4 SV=1 |

| | | | | | | | | | | | | |
|---|---|---|---|---|---|---|---|---|---|---|---|---|
| 114 | 5787 | tr\|A0A0G2 | 25.95 | 2 | 2 | 0 | 2 | 1 | 2 | | 67482 | Mitoguardin 1 OS=Rattus norvegicus OX=10116 GN=Miga1 PE=4 SV=1 |
| 83 | 555 | tr\|D4A1D3 | 25.04 | 0 | 0 | 4.11E+05 | 2 | 1 | 3 | | 521487 | Sacsin molecular chaperone OS=Rattus norvegicus OX=10116 GN=Sacs PE=1 SV=2 |
| 144 | 1485 | tr\|D3ZA68 | 24.92 | 2 | 2 | 0 | 2 | 1 | 2 | | 76890 | Progressive external ophthalmoplegia 1 homolog (Human) (Predicted) OS=Rattus norvegicus OX=10116 GN=Twnk PE=4 SV=1 |
| 116 | 5810 | tr\|A0A0H2 | 24.61 | 4 | 4 | 0 | 2 | 1 | 2 | | 25865 | 39S ribosomal protein L22 mitochondrial OS=Rattus norvegicus OX=10116 GN=Mrpl22 PE=1 SV=1 |
| 116 | 5841 | P0C2C0\|RI | 24.61 | 5 | 5 | 0 | 2 | 1 | 2 | | 24039 | 39S ribosomal protein L22 mitochondrial OS=Rattus norvegicus OX=10116 GN=Mrpl22 PE=1 SV=1 |
| 91 | 236 | tr\|Q5U2T0 | 23.94 | 3 | 3 | 7.66E+03 | 2 | 2 | 3 | | 44494 | Death associated protein 3 OS=Rattus norvegicus OX=10116 GN=Dap3 PE=2 SV=1 |
| 91 | 237 | tr\|F7EZZ0\| | 23.94 | 3 | 3 | 7.66E+03 | 2 | 2 | 3 | | 45112 | Death-associated protein 3 OS=Rattus norvegicus OX=10116 GN=Dap3 PE=1 SV=1 |
| 91 | 242 | tr\|A0A0G2 | 23.94 | 2 | 2 | 7.66E+03 | 2 | 2 | 3 | | 46666 | Death-associated protein 3 OS=Rattus norvegicus OX=10116 GN=Dap3 PE=1 SV=1 |
| 224 | 139 | Q6AXV4\|S | 23.79 | 2 | 2 | 1.41E+05 | 1 | 1 | 1 | | 51960 | Sorting and assembly machinery component 50 homolog OS=Rattus norvegicus OX=10116 GN=Samm50 PE=1 SV=1 |
| 128 | 1105 | tr\|B5DFI9\| | 23.69 | 3 | 3 | 3.18E+05 | 2 | 1 | 2 | | 47944 | Protein-serine/threonine kinase OS=Rattus norvegicus OX=10116 GN=Pdk3 PE=2 SV=1 |
| 273 | 5817 | P21571\|AT | 23.6 | 9 | 9 | 7.20E+04 | 1 | 1 | 1 | | 12494 | ATP synthase-coupling factor 6 mitochondrial OS=Rattus norvegicus OX=10116 GN=Atp5pf PE=1 SV=1 |
| 96 | 5819 | tr\|B2RYU0 | 23.08 | 12 | 12 | 1.40E+06 | 1 | 1 | 3 | | 11842 | NADH dehydrogenase (Ubiquinone) 1 beta subcomplex 2 (Predicted) isoform CRA_b OS=Rattus norvegicus OX=10116 GN=Ndufb2 PE=1 SV=1 |
| 226 | 1690 | tr\|B1WBY5 | 22.53 | 1 | 1 | 0 | 1 | 1 | 1 | | 63205 | DnaJ (Hsp40) homolog subfamily C member 11 OS=Rattus norvegicus OX=10116 GN=Dnajc11 PE=1 SV=1 |
| 82 | 599 | Q00918\|LT | 22.51 | 1 | 1 | 2.25E+06 | 1 | 1 | 4 | Carbamido | 186598 | Latent-transforming growth factor beta-binding protein 1 OS=Rattus norvegicus OX=10116 GN=Ltbp1 PE=1 SV=1 |
| 274 | 398 | P48037\|AN | 22.4 | 1 | 1 | 3.14E+05 | 1 | 1 | 1 | | 75754 | Annexin A6 OS=Rattus norvegicus OX=10116 GN=Anxa6 PE=1 SV=2 |
| 274 | 399 | tr\|Q6IMZ3 | 22.4 | 1 | 1 | 3.14E+05 | 1 | 1 | 1 | | 75756 | Annexin OS=Rattus norvegicus OX=10116 GN=Anxa6 PE=1 SV=1 |
| 194 | 188 | O70351\|H | 22.09 | 3 | 3 | 1.61E+05 | 1 | 1 | 1 | | 27246 | 3-hydroxyacyl-CoA dehydrogenase type-2 OS=Rattus norvegicus OX=10116 GN=Hsd17b10 PE=1 SV=3 |
| 194 | 189 | tr\|B0BMW | 22.09 | 3 | 3 | 1.61E+05 | 1 | 1 | 1 | | 27250 | 3-hydroxyacyl-CoA dehydrogenase type-2 OS=Rattus norvegicus OX=10116 GN=Hsd17b10 PE=1 SV=1 |
| 174 | 393 | Q63563\|A | 21.99 | 1 | 1 | 1.31E+06 | 1 | 1 | 1 | | 174117 | ATP-binding cassette sub-family C member 9 OS=Rattus norvegicus OX=10116 GN=Abcc9 PE=1 SV=1 |
| 129 | 890 | tr\|D3ZAE6 | 21.73 | 1 | 1 | 4.06E+05 | 1 | 1 | 2 | | 72321 | RCG49849 OS=Rattus norvegicus OX=10116 GN=Vasn PE=1 SV=1 |
| 129 | 6092 | Q2Q0I9\|FN | 21.73 | 0 | 0 | 4.06E+05 | 1 | 1 | 2 | | 194048 | Fibronectin type III domain-containing protein 1 OS=Rattus norvegicus OX=10116 GN=Fndc1 PE=1 SV=2 |
| 97 | 497 | tr\|Q9JID3\| | 21.51 | 2 | 2 | 3.11E+05 | 1 | 1 | 3 | Oxidation ( | 43884 | Protein-serine/threonine kinase OS=Rattus norvegicus OX=10116 PE=2 SV=1 |
| 97 | 729 | tr\|F1LMM | 21.51 | 2 | 2 | 3.11E+05 | 1 | 1 | 3 | Oxidation ( | 46240 | Protein-serine/threonine kinase OS=Rattus norvegicus OX=10116 GN=Pdk2 PE=1 SV=3 |
| 97 | 498 | Q64536\|P | 21.51 | 2 | 2 | 3.11E+05 | 1 | 1 | 3 | Oxidation ( | 46106 | [Pyruvate dehydrogenase (acetyl-transferring)] kinase isozyme 2 mitochondrial OS=Rattus norvegicus OX=10116 GN=Pdk2 PE=1 SV=1 |
| 127 | 5826 | tr\|A0A0G2 | 21.46 | 2 | 2 | 4.74E+04 | 1 | 1 | 2 | | 28715 | Autophagy protein 5 OS=Rattus norvegicus OX=10116 GN=Atg5 PE=1 SV=1 |
| 127 | 5827 | Q3MQ06\| | 21.46 | 2 | 2 | 4.74E+04 | 1 | 1 | 2 | | 32398 | Autophagy protein 5 OS=Rattus norvegicus OX=10116 GN=Atg5 PE=2 SV=1 |
| 143 | 584 | tr\|D3ZYM7 | 21.24 | 10 | 10 | 1.64E+06 | 1 | 1 | 2 | | 10816 | LYR motif-containing 4 OS=Rattus norvegicus OX=10116 GN=Lyrm4 PE=1 SV=1 |
| 156 | 489 | tr\|D3ZL86\| | 21.19 | 0 | 0 | | 1 | 0 | 1 | | 241219 | HEAT repeat-containing protein 1 OS=Rattus norvegicus OX=10116 GN=Heatr1 PE=1 SV=2 |
| 192 | 612 | tr\|A0A0G2 | 21.09 | 1 | 1 | 3.76E+05 | 1 | 1 | 1 | | 62571 | 5-aminolevulinate synthase OS=Rattus norvegicus OX=10116 GN=Alas1 PE=1 SV=1 |
| 192 | 613 | tr\|A0A0G2 | 21.09 | 1 | 1 | 3.76E+05 | 1 | 1 | 1 | | 71039 | 5-aminolevulinate synthase OS=Rattus norvegicus OX=10116 GN=Alas1 PE=1 SV=1 |
| 192 | 614 | P13195\|HE | 21.09 | 1 | 1 | 3.76E+05 | 1 | 1 | 1 | | 71021 | 5-aminolevulinate synthase nonspecific mitochondrial OS=Rattus norvegicus OX=10116 GN=Alas1 PE=2 SV=2 |
| 276 | 361 | P80433\|CC | 20.66 | 14 | 14 | 1.01E+05 | 1 | 1 | 1 | | 7672 | Cytochrome c oxidase subunit 8A mitochondrial OS=Rattus norvegicus OX=10116 GN=Cox8a PE=1 SV=3 |
| 230 | 5820 | tr\|D4A8P8 | 20.55 | 1 | 1 | 3.22E+05 | 1 | 1 | 1 | | 92336 | Hypoxia-inducible factor 1-alpha OS=Rattus norvegicus OX=10116 GN=Hif1a PE=4 SV=2 |
| 230 | 5821 | O35800\|H | 20.55 | 1 | 1 | 3.22E+05 | 1 | 1 | 1 | | 92319 | Hypoxia-inducible factor 1-alpha OS=Rattus norvegicus OX=10116 GN=Hif1a PE=2 SV=1 |
| 182 | 1280 | tr\|Q5U2M | 20.55 | 1 | 1 | 0 | 1 | 1 | 1 | | 105443 | DNA ligase OS=Rattus norvegicus OX=10116 GN=Lig3 PE=1 SV=1 |
| 182 | 1281 | tr\|A0A096 | 20.55 | 1 | 1 | 0 | 1 | 1 | 1 | | 112017 | DNA ligase OS=Rattus norvegicus OX=10116 GN=Lig3 PE=1 SV=1 |
| 162 | 1672 | tr\|F1LSE5\| | 20.41 | 1 | 1 | 0 | 1 | 1 | 1 | | 109803 | Neighbor of Brca1 gene 1 isoform CRA_a OS=Rattus norvegicus OX=10116 GN=Nbr1 PE=1 SV=2 |
| 278 | 197 | tr\|M0R6J0 | 20.31 | 3 | 3 | 8.19E+04 | 1 | 1 | 1 | | 38374 | Mitochondrial ribosomal protein L39 OS=Rattus norvegicus OX=10116 GN=Mrpl39 PE=1 SV=2 |
| 280 | 5834 | tr\|D4A131 | 20.03 | 2 | 2 | 7.96E+04 | 1 | 1 | 1 | | 33082 | Mitochondrial ribosomal protein L4 OS=Rattus norvegicus OX=10116 GN=Mrpl4 PE=1 SV=1 |
