## Supplementary material for "Permeability transition pore-related changes in the proteome and channel activity of ATP synthase dimers and monomers": RLM KCl-BM

| Protein Group | Protein ID | Accession | -10lgP | Coverage (%) | Coverage (%) Sample 6 | IBAQ | #Peptides | #Unique | #Spec Sample 6 | PTM | Avg. Mass | Description |
|---|---|---|---|---|---|---|---|---|---|---|---|---|
| **5** | 14 | P15999\|ATPA_RAT | 374.67 | 55 | 55 | 96658000 | 37 | 37 | 61 | Y | 59754 | ATP synthase subunit alpha, mitochondrial OS=Rattus norvegicus OX=10116 GN=Atp5f1a PE=1 SV=2 |
| **3** | 13 | P07756\|CPSM_RAT | 352.11 | 32 | 32 | 35426000 | 38 | 38 | 71 | Y | 164579 | Carbamoyl-phosphate synthase [ammonia], mitochondrial OS=Rattus norvegicus OX=10116 GN=Cps1 PE=1 SV=1 |
| **2** | 7 | P10719\|ATPB_RAT | 336.91 | 75 | 75 | 98213000 | 33 | 33 | 82 | Y | 56354 | ATP synthase subunit beta, mitochondrial OS=Rattus norvegicus OX=10116 GN=Atp5f1b PE=1 SV=2 |
| **2** | 8 | tr\|G3V6D3\|G3V6D3_RAT | 336.91 | 75 | 75 | 98213000 | 33 | 33 | 82 | Y | 56345 | ATP synthase subunit beta OS=Rattus norvegicus OX=10116 GN=Atp5f1b PE=1 SV=1 |
| **1** | 53 | P52873\|PYC_RAT | 334.87 | 56 | 56 | 102670000 | 61 | 61 | 105 | Y | 129777 | Pyruvate carboxylase, mitochondrial OS=Rattus norvegicus OX=10116 GN=Pc PE=1 SV=2 |
| **1** | 54 | tr\|A0A0G2JTL5\|A0A0G2JTL5_RAT | 334.87 | 51 | 51 | 102670000 | 61 | 61 | 105 | Y | 140005 | Pyruvate carboxylase, mitochondrial OS=Rattus norvegicus OX=10116 GN=Pc PE=1 SV=1 |
| **6** | 12 | Q68FY0\|QCR1_RAT | 312.6 | 59 | 59 | 34117000 | 23 | 23 | 52 | Y | 52849 | Cytochrome b-c1 complex subunit 1, mitochondrial OS=Rattus norvegicus OX=10116 GN=Uqcrc1 PE=1 SV=1 |
| **4** | 21 | Q64428\|ECHA_RAT | 312.36 | 44 | 44 | 37818000 | 34 | 34 | 64 | Y | 82665 | Trifunctional enzyme subunit alpha, mitochondrial OS=Rattus norvegicus OX=10116 GN=Hadha PE=1 SV=2 |
| **7** | 9 | P00507\|AATM_RAT | 299.08 | 65 | 65 | 81734000 | 28 | 28 | 44 | Y | 47314 | Aspartate aminotransferase, mitochondrial OS=Rattus norvegicus OX=10116 GN=Got2 PE=1 SV=2 |
| **9** | 1 | P32551\|QCR2_RAT | 286.87 | 69 | 69 | 49716000 | 25 | 25 | 35 | Y | 48396 | Cytochrome b-c1 complex subunit 2, mitochondrial OS=Rattus norvegicus OX=10116 GN=Uqcrc2 PE=1 SV=2 |
| **10** | 6796 | P04762\|CATA_RAT | 273.78 | 45 | 45 | 31692000 | 21 | 21 | 34 | Y | 59757 | Catalase OS=Rattus norvegicus OX=10116 GN=Cat PE=1 SV=3 |
| **8** | 2 | Q02253\|MMSA_RAT | 267.47 | 50 | 50 | 33668000 | 23 | 23 | 38 | Y | 57808 | Methylmalonate-semialdehyde dehydrogenase [acylating], mitochondrial OS=Rattus norvegicus OX=10116 GN=Aldh6a1 PE=1 SV=1 |
| **8** | 3 | tr\|G3V7J0\|G3V7J0_RAT | 267.47 | 50 | 50 | 33668000 | 23 | 23 | 38 | Y | 57748 | Aldehyde dehydrogenase family 6, subfamily A1, isoform CRA_b OS=Rattus norvegicus OX=10116 GN=Aldh6a1 PE=1 SV=1 |
| **11** | 229 | A2VCW9\|AASS_RAT | 236.85 | 22 | 22 | 13489000 | 19 | 19 | 28 | Y | 102908 | Alpha-aminoadipic semialdehyde synthase, mitochondrial OS=Rattus norvegicus OX=10116 GN=Aass PE=2 SV=1 |
| **24** | 4338 | P63039\|CH60_RAT | 227.54 | 18 | 18 | 2694500 | 8 | 8 | 11 | N | 60956 | 60 kDa heat shock protein, mitochondrial OS=Rattus norvegicus OX=10116 GN=Hspd1 PE=1 SV=1 |
| **24** | 4339 | tr\|A0A482IDN3\|A0A482IDN3_RAT | 227.54 | 18 | 18 | 2694500 | 8 | 8 | 11 | N | 60956 | Hsp60 OS=Rattus norvegicus OX=10116 GN=Hspd1 PE=2 SV=1 |
| **15** | 4373 | P0C2X9\|AL4A1_RAT | 216.75 | 28 | 28 | 7200300 | 10 | 10 | 18 | Y | 61869 | Delta-1-pyrroline-5-carboxylate dehydrogenase, mitochondrial OS=Rattus norvegicus OX=10116 GN=Aldh4a1 PE=1 SV=1 |
| **22** | 85 | tr\|A0A0G2JVH4\|A0A0G2JVH4_RAT | 209.76 | 12 | 12 | 3129900 | 9 | 9 | 11 | N | 86230 | MICOS complex subunit MIC60 OS=Rattus norvegicus OX=10116 GN=Immt PE=1 SV=1 |
| **17** | 6795 | tr\|Q68FZ8\|Q68FZ8_RAT | 207.14 | 28 | 28 | 9730100 | 12 | 12 | 18 | N | 58678 | Propionyl coenzyme A carboxylase, beta polypeptide OS=Rattus norvegicus OX=10116 GN=Pccb PE=1 SV=1 |
| **21** | 35 | P13086\|SUCA_RAT | 205.52 | 31 | 31 | 4547800 | 8 | 8 | 11 | Y | 36148 | Succinate--CoA ligase [ADP/GDP-forming] subunit alpha, mitochondrial OS=Rattus norvegicus OX=10116 GN=Suclg1 PE=2 SV=2 |
| **21** | 36 | tr\|A0A0H2UHE1\|A0A0H2UHE1_RAT | 205.52 | 30 | 30 | 4547800 | 8 | 8 | 11 | Y | 37560 | Succinate--CoA ligase [ADP/GDP-forming] subunit alpha, mitochondrial OS=Rattus norvegicus OX=10116 GN=Suclg1 PE=1 SV=1 |
| **13** | 19 | P22791\|HMCS2_RAT | 203.92 | 36 | 36 | 12073000 | 16 | 16 | 23 | Y | 56912 | Hydroxymethylglutaryl-CoA synthase, mitochondrial OS=Rattus norvegicus OX=10116 GN=Hmgcs2 PE=1 SV=1 |
| **13** | 20 | tr\|Q68G44\|Q68G44_RAT | 203.92 | 36 | 36 | 12073000 | 16 | 16 | 23 | Y | 56886 | 3-hydroxy-3-methylglutaryl coenzyme A synthase OS=Rattus norvegicus OX=10116 GN=Hmgcs2 PE=1 SV=1 |
| **20** | 39 | P17764\|THIL_RAT | 203.46 | 26 | 26 | 4574300 | 8 | 8 | 12 | N | 44695 | Acetyl-CoA acetyltransferase, mitochondrial OS=Rattus norvegicus OX=10116 GN=Acat1 PE=1 SV=1 |
| **19** | 2302 | P16970\|ABCD3_RAT | 201.57 | 19 | 19 | 2930900 | 9 | 9 | 13 | Y | 75316 | ATP-binding cassette sub-family D member 3 OS=Rattus norvegicus OX=10116 GN=Abcd3 PE=1 SV=3 |
| **12** | 40 | Q60587\|ECHB_RAT | 200.5 | 38 | 38 | 17524000 | 16 | 16 | 27 | Y | 51414 | Trifunctional enzyme subunit beta, mitochondrial OS=Rattus norvegicus OX=10116 GN=Hadhb PE=1 SV=1 |
| **16** | 116 | P35435\|ATPG_RAT | 199.46 | 48 | 48 | 12988000 | 12 | 12 | 18 | Y | 30191 | ATP synthase subunit gamma, mitochondrial OS=Rattus norvegicus OX=10116 GN=Atp5f1c PE=1 SV=2 |

| | | | | | | | | | | | | |
|---|---|---|---|---|---|---|---|---|---|---|---|---|
| **18** | 28 | tr\|D3ZFQ8\|D3ZFQ8_RAT | 191.16 | 38 | 38 | 13900000 | 10 | 10 | 14 | Y | 35435 | Cytochrome c-1 OS=Rattus norvegicus OX=10116 GN=Cyc1 PE=1 SV=3 |
| **23** | 41 | P19511\|AT5F1_RAT | 168.44 | 30 | 30 | 20815000 | 9 | 9 | 11 | N | 28869 | ATP synthase F(0) complex subunit B1, mitochondrial OS=Rattus norvegicus OX=10116 GN=Atp5pb PE=1 SV=1 |
| **29** | 47 | Q9WVK3\|PECR_RAT | 166.94 | 22 | 22 | 1395600 | 4 | 4 | 7 | N | 32433 | Peroxisomal trans-2-enoyl-CoA reductase OS=Rattus norvegicus OX=10116 GN=Pecr PE=2 SV=1 |
| **29** | 48 | tr\|A0A0G2JVG4\|A0A0G2JVG4_RAT | 166.94 | 22 | 22 | 1395600 | 4 | 4 | 7 | N | 32737 | Peroxisomal trans-2-enoyl-CoA reductase OS=Rattus norvegicus OX=10116 GN=Pecr PE=1 SV=1 |
| **14** | 9289 | P24329\|THTR_RAT | 165.11 | 33 | 33 | 9651800 | 8 | 8 | 21 | Y | 33407 | Thiosulfate sulfurtransferase OS=Rattus norvegicus OX=10116 GN=Tst PE=1 SV=3 |
| **26** | 4602 | B3DMA2\|ACD11_RAT | 162.27 | 9 | 9 | 3166500 | 7 | 7 | 9 | N | 87371 | Acyl-CoA dehydrogenase family member 11 OS=Rattus norvegicus OX=10116 GN=Acad11 PE=1 SV=1 |
| **28** | 2163 | Q06647\|ATPO_RAT | 142.69 | 29 | 29 | 5665400 | 6 | 6 | 8 | N | 23398 | ATP synthase subunit O, mitochondrial OS=Rattus norvegicus OX=10116 GN=Atp5po PE=1 SV=1 |
| **30** | 37 | P20788\|UCRI_RAT | 138.25 | 28 | 28 | 3454900 | 6 | 6 | 7 | N | 29446 | Cytochrome b-c1 complex subunit Rieske, mitochondrial OS=Rattus norvegicus OX=10116 GN=Uqcrfs1 PE=1 SV=2 |
| **41** | 109 | P11240\|COX5A_RAT | 137.4 | 30 | 30 | 999100 | 3 | 3 | 4 | N | 16130 | Cytochrome c oxidase subunit 5A, mitochondrial OS=Rattus norvegicus OX=10116 GN=Cox5a PE=1 SV=1 |
| **32** | 93 | tr\|A0A0G2K8Q8\|A0A0G2K8Q8_RAT | 128.24 | 52 | 52 | 2463400 | 3 | 3 | 7 | N | 7099 | Ubiquinol-cytochrome c reductase, complex III subunit X OS=Rattus norvegicus OX=10116 GN=Uqcr10 PE=1 SV=1 |
| **32** | 94 | tr\|B2RYX1\|B2RYX1_RAT | 128.24 | 50 | 50 | 2463400 | 3 | 3 | 7 | N | 7462 | LOC685322 protein OS=Rattus norvegicus OX=10116 GN=Uqcr10 PE=2 SV=1 |
| **40** | 289 | P07872\|ACOX1_RAT | 123.16 | 8 | 8 | 242590 | 4 | 4 | 4 | Y | 74679 | Peroxisomal acyl-coenzyme A oxidase 1 OS=Rattus norvegicus OX=10116 GN=Acox1 PE=1 SV=1 |
| **31** | 51 | P10888\|COX41_RAT | 120.98 | 29 | 29 | 3057200 | 5 | 5 | 7 | N | 19515 | Cytochrome c oxidase subunit 4 isoform 1, mitochondrial OS=Rattus norvegicus OX=10116 GN=Cox4i1 PE=1 SV=1 |
| **38** | 4261 | tr\|G3V7Y3\|G3V7Y3_RAT | 117.25 | 27 | 27 | 3782200 | 4 | 4 | 4 | N | 17563 | ATP synthase subunit delta, mitochondrial OS=Rattus norvegicus OX=10116 GN=Atp5f1d PE=1 SV=1 |
| **38** | 4262 | P35434\|ATPD_RAT | 117.25 | 27 | 27 | 3782200 | 4 | 4 | 4 | N | 17595 | ATP synthase subunit delta, mitochondrial OS=Rattus norvegicus OX=10116 GN=Atp5f1d PE=1 SV=2 |
| **47** | 775 | tr\|A0A0G2K401\|A0A0G2K401_RAT | 108.57 | 4 | 4 | 1445200 | 2 | 2 | 2 | N | 79812 | Propionyl-CoA carboxylase alpha chain, mitochondrial OS=Rattus norvegicus OX=10116 GN=Pcca PE=1 SV=1 |
| **47** | 777 | tr\|A0A0G2K5P9\|A0A0G2K5P9_RAT | 108.57 | 4 | 4 | 1445200 | 2 | 2 | 2 | N | 81413 | Propionyl-CoA carboxylase alpha chain, mitochondrial OS=Rattus norvegicus OX=10116 GN=Pcca PE=1 SV=1 |
| **47** | 776 | P14882\|PCCA_RAT | 108.57 | 4 | 4 | 1445200 | 2 | 2 | 2 | N | 81623 | Propionyl-CoA carboxylase alpha chain, mitochondrial OS=Rattus norvegicus OX=10116 GN=Pcca PE=1 SV=3 |
| **27** | 30 | Q7TQ16\|QCR8_RAT | 107.64 | 34 | 34 | 2211200 | 4 | 4 | 9 | N | 9849 | Cytochrome b-c1 complex subunit 8 OS=Rattus norvegicus OX=10116 GN=Uqcrq PE=3 SV=1 |
| **34** | 9290 | P04636\|MDHM_RAT | 107.58 | 14 | 14 | 1372600 | 4 | 4 | 5 | N | 35684 | Malate dehydrogenase, mitochondrial OS=Rattus norvegicus OX=10116 GN=Mdh2 PE=1 SV=2 |
| **33** | 4260 | P31399\|ATP5H_RAT | 106.2 | 30 | 30 | 1583500 | 4 | 4 | 5 | Y | 18763 | ATP synthase subunit d, mitochondrial OS=Rattus norvegicus OX=10116 GN=Atp5pd PE=1 SV=3 |
| **35** | 101 | tr\|B2RYT5\|B2RYT5_RAT | 105.26 | 35 | 35 | 5930900 | 4 | 4 | 5 | N | 12651 | Cox7a2l protein OS=Rattus norvegicus OX=10116 GN=Cox7a2l PE=2 SV=1 |
| **43** | 603 | A0A0G2K047\|ACSS3_RAT | 102.24 | 4 | 4 | 362900 | 3 | 3 | 3 | N | 74675 | Acyl-CoA synthetase short-chain family member 3, mitochondrial OS=Rattus norvegicus OX=10116 GN=Acss3 PE=1 SV=1 |
| **37** | 98 | P29147\|BDH_RAT | 100.97 | 12 | 12 | 1706300 | 3 | 3 | 5 | Y | 38202 | D-beta-hydroxybutyrate dehydrogenase, mitochondrial OS=Rattus norvegicus OX=10116 GN=Bdh1 PE=1 SV=2 |
| **37** | 99 | tr\|A0A0G2JSH2\|A0A0G2JSH2_RAT | 100.97 | 12 | 12 | 1706300 | 3 | 3 | 5 | Y | 38333 | 3-hydroxybutyrate dehydrogenase, type 1, isoform CRA_a OS=Rattus norvegicus OX=10116 GN=Bdh1 PE=1 SV=1 |
| **42** | 106 | tr\|G3V6I4\|G3V6I4_RAT | 100.4 | 9 | 9 | 1088300 | 1 | 1 | 4 | N | 37791 | Mitochondrial amidoxime reducing component 1 OS=Rattus norvegicus OX=10116 GN=Marc1 PE=1 SV=1 |
| **25** | 278 | tr\|D3ZFJ6\|D3ZFJ6_RAT | 97.52 | 17 | 17 | 978600 | 5 | 5 | 9 | Y | 60420 | Lactamase, beta OS=Rattus norvegicus OX=10116 GN=Lactb PE=1 SV=1 |
| **54** | 11610 | tr\|Q7H115\|Q7H115_RAT | 84.76 | 5 | 5 | 497400 | 1 | 1 | 2 | N | 29871 | Cytochrome c oxidase subunit 3 OS=Rattus norvegicus OX=10116 GN=Mt-co3 PE=3 SV=1 |

| | | | | | | | | | | | | |
|---|---|---|---|---|---|---|---|---|---|---|---|---|
| **54** | 11611 | tr\|I6V4L9\|I6V4L9_RAT | 84.76 | 5 | 5 | 497400 | 1 | 1 | 2 | N | 29844 | Cytochrome c oxidase subunit 3 OS=Rattus norvegicus OX=10116 GN=COX3 PE=3 SV=1 |
| **54** | 11612 | tr\|Q8M7G4\|Q8M7G4_RAT | 84.76 | 5 | 5 | 497400 | 1 | 1 | 2 | N | 29861 | Cytochrome c oxidase subunit 3 OS=Rattus norvegicus OX=10116 PE=2 SV=1 |
| **54** | 11613 | tr\|A0A096XKT2\|A0A096XKT2_RAT | 84.76 | 5 | 5 | 497400 | 1 | 1 | 2 | N | 29901 | Cytochrome c oxidase subunit 3 OS=Rattus norvegicus OX=10116 GN=COX3 PE=3 SV=1 |
| **54** | 11614 | P05505\|COX3_RAT | 84.76 | 5 | 5 | 497400 | 1 | 1 | 2 | N | 29871 | Cytochrome c oxidase subunit 3 OS=Rattus norvegicus OX=10116 GN=Mtco3 PE=1 SV=5 |
| **54** | 11615 | tr\|Q8SEZ2\|Q8SEZ2_RAT | 84.76 | 5 | 5 | 497400 | 1 | 1 | 2 | N | 29870 | Cytochrome c oxidase subunit 3 (Fragment) OS=Rattus norvegicus OX=10116 GN=COIII PE=3 SV=1 |
| **39** | 2210 | Q8VID1\|DHRS4_RAT | 78.03 | 8 | 8 | 254190 | 2 | 2 | 4 | N | 29822 | Dehydrogenase/reductase SDR family member 4 OS=Rattus norvegicus OX=10116 GN=Dhrs4 PE=2 SV=2 |
| **36** | 2174 | O70351\|HCD2_RAT | 74.56 | 23 | 23 | 997660 | 3 | 3 | 5 | Y | 27246 | 3-hydroxyacyl-CoA dehydrogenase type-2 OS=Rattus norvegicus OX=10116 GN=Hsd17b10 PE=1 SV=3 |
| **36** | 2175 | tr\|B0BMW2\|B0BMW2_RAT | 74.56 | 23 | 23 | 997660 | 3 | 3 | 5 | Y | 27250 | 3-hydroxyacyl-CoA dehydrogenase type-2 OS=Rattus norvegicus OX=10116 GN=Hsd17b10 PE=1 SV=1 |
| **91** | 96 | P08461\|ODP2_RAT | 73.22 | 2 | 2 | 112270 | 1 | 1 | 1 | N | 67166 | Dihydrolipoyllysine-residue acetyltransferase component of pyruvate dehydrogenase complex, mitochondrial OS=Rattus norvegicus OX=10116 GN=Dlat PE=1 SV=3 |
| **92** | 4472 | P45953\|ACADV_RAT | 70.38 | 3 | 3 | 43860 | 1 | 1 | 1 | N | 70749 | Very long-chain specific acyl-CoA dehydrogenase, mitochondrial OS=Rattus norvegicus OX=10116 GN=Acadvl PE=1 SV=1 |
| **92** | 4473 | tr\|Q5M9H2\|Q5M9H2_RAT | 70.38 | 3 | 3 | 43860 | 1 | 1 | 1 | N | 70821 | Acyl-Coenzyme A dehydrogenase, very long chain OS=Rattus norvegicus OX=10116 GN=Acadvl PE=1 SV=1 |
| **93** | 25 | tr\|D3ZG43\|D3ZG43_RAT | 65.27 | 4 | 4 | 193260 | 1 | 1 | 1 | N | 30226 | NADH dehydrogenase (Ubiquinone) Fe-S protein 3 (Predicted), isoform CRA_c OS=Rattus norvegicus OX=10116 GN=Ndufs3 PE=1 SV=1 |
| **45** | 4264 | tr\|Q5UAJ5\|Q5UAJ5_RAT | 61.25 | 27 | 27 | 414950 | 2 | 2 | 3 | N | 7642 | ATP synthase protein 8 OS=Rattus norvegicus OX=10116 GN=ATP8 PE=3 SV=1 |
| **46** | 542 | P10860\|DHE3_RAT | 57.08 | 3 | 3 | 450270 | 1 | 1 | 2 | N | 61416 | Glutamate dehydrogenase 1, mitochondrial OS=Rattus norvegicus OX=10116 GN=Glud1 PE=1 SV=2 |
| **52** | 9291 | tr\|D3ZUX5\|D3ZUX5_RAT | 54.76 | 7 | 7 | 160850 | 2 | 2 | 2 | N | 26435 | MICOS complex subunit OS=Rattus norvegicus OX=10116 GN=Chchd3 PE=1 SV=1 |
| **94** | 11616 | P10818\|CX6A1_RAT | 52.78 | 11 | 11 | 115170 | 1 | 1 | 1 | N | 12301 | Cytochrome c oxidase subunit 6A1, mitochondrial OS=Rattus norvegicus OX=10116 GN=Cox6a1 PE=1 SV=2 |
| **72** | 760 | tr\|F1M953\|F1M953_RAT | 50.51 | 3 | 3 | 29534 | 1 | 1 | 1 | N | 73745 | Stress-70 protein, mitochondrial OS=Rattus norvegicus OX=10116 GN=Hspa9 PE=1 SV=1 |
| **72** | 761 | P48721\|GRP75_RAT | 50.51 | 3 | 3 | 29534 | 1 | 1 | 1 | N | 73858 | Stress-70 protein, mitochondrial OS=Rattus norvegicus OX=10116 GN=Hspa9 PE=1 SV=3 |
| **95** | 11617 | P80432\|COX7C_RAT | 49.48 | 14 | 14 | 106690 | 1 | 1 | 1 | N | 7375 | Cytochrome c oxidase subunit 7C, mitochondrial OS=Rattus norvegicus OX=10116 GN=Cox7c PE=1 SV=2 |
| **53** | 34 | tr\|Q5RJN0\|Q5RJN0_RAT | 46 | 12 | 12 | 62023 | 2 | 2 | 2 | N | 23945 | NADH dehydrogenase (Ubiquinone) Fe-S protein 7 OS=Rattus norvegicus OX=10116 GN=Ndufs7 PE=1 SV=1 |
| **44** | 56 | tr\|S5RZM8\|S5RZM8_RAT | 43.34 | 7 | 7 | 2752800 | 2 | 2 | 3 | Y | 25958 | Cytochrome c oxidase subunit 2 OS=Rattus norvegicus OX=10116 GN=COX2 PE=3 SV=1 |
| **44** | 57 | P00406\|COX2_RAT | 43.34 | 7 | 7 | 2752800 | 2 | 2 | 3 | Y | 25928 | Cytochrome c oxidase subunit 2 OS=Rattus norvegicus OX=10116 GN=Mtco2 PE=1 SV=3 |
| **44** | 58 | tr\|Q37652\|Q37652_RAT | 43.34 | 7 | 7 | 2752800 | 2 | 2 | 3 | Y | 26007 | Cytochrome c oxidase subunit 2 OS=Rattus norvegicus OX=10116 GN=COXII PE=3 SV=1 |
| **44** | 59 | tr\|Q5UAJ6\|Q5UAJ6_RAT | 43.34 | 7 | 7 | 2752800 | 2 | 2 | 3 | Y | 25942 | Cytochrome c oxidase subunit 2 OS=Rattus norvegicus OX=10116 GN=COX2 PE=3 SV=1 |
| **44** | 60 | tr\|Q8SEZ5\|Q8SEZ5_RAT | 43.34 | 7 | 7 | 2752800 | 2 | 2 | 3 | Y | 25928 | Cytochrome c oxidase subunit 2 OS=Rattus norvegicus OX=10116 GN=Mt-co2 PE=1 SV=1 |
| **44** | 61 | tr\|A0A097PE04\|A0A097PE04_RAT | 43.34 | 7 | 7 | 2752800 | 2 | 2 | 3 | Y | 25894 | Cytochrome c oxidase subunit 2 OS=Rattus norvegicus OX=10116 GN=COX2 PE=3 SV=1 |
| **96** | 11618 | P11951\|CX6C2_RAT | 38.97 | 12 | 12 | 0 | 1 | 1 | 1 | N | 8455 | Cytochrome c oxidase subunit 6C-2 OS=Rattus norvegicus OX=10116 GN=Cox6c2 PE=1 SV=3 |
| **99** | 26 | tr\|F1LN92\|F1LN92_RAT | 37.16 | 1 | 1 | 0 | 1 | 1 | 1 | N | 89352 | AFG3-like matrix AAA peptidase subunit 2 OS=Rattus norvegicus OX=10116 GN=Afg3l2 PE=1 SV=1 |

| | | | | | | | | | | | | |
|---|---|---|---|---|---|---|---|---|---|---|---|---|
| **97** | 4263 | Q6PDU7\|ATP5L_RAT | 36.33 | 12 | 12 | 0 | 1 | 1 | 1 | N | 11433 | ATP synthase subunit g, mitochondrial OS=Rattus norvegicus OX=10116 GN=Atp5mg PE=1 SV=2 |
| **98** | 4266 | D3ZAF6\|ATPK_RAT | 36.01 | 14 | 14 | 648680 | 1 | 1 | 1 | N | 10452 | ATP synthase subunit f, mitochondrial OS=Rattus norvegicus OX=10116 GN=Atp5mf PE=1 SV=1 |
| **73** | 7185 | Q924S5\|LONM_RAT | 34.44 | 4 | 4 | 429260 | 1 | 1 | 1 | N | 105792 | Lon protease homolog, mitochondrial OS=Rattus norvegicus OX=10116 GN=Lonp1 PE=2 SV=1 |
| **74** | 4857 | P32198\|CPT1A_RAT | 32.15 | 1 | 1 | 536720 | 1 | 1 | 1 | N | 88126 | Carnitine O-palmitoyltransferase 1, liver isoform OS=Rattus norvegicus OX=10116 GN=Cpt1a PE=1 SV=2 |
| **48** | 401 | tr\|B1WBY5\|B1WBY5_RAT | 31.43 | 2 | 2 | 1557500 | 2 | 2 | 2 | N | 63205 | DnaJ (Hsp40) homolog, subfamily C, member 11 OS=Rattus norvegicus OX=10116 GN=Dnajc11 PE=1 SV=1 |
| **100** | 11619 | P80431\|COX7B_RAT | 30.72 | 9 | 9 | 328570 | 1 | 1 | 1 | N | 8995 | Cytochrome c oxidase subunit 7B, mitochondrial OS=Rattus norvegicus OX=10116 GN=Cox7b PE=1 SV=3 |
| **55** | 157 | tr\|F1LTG5\|F1LTG5_RAT | 28.25 | 14 | 14 | 2537700 | 1 | 1 | 2 | N | 13829 | Uncharacterized protein OS=Rattus norvegicus OX=10116 PE=4 SV=2 |
| **55** | 158 | P12075\|COX5B_RAT | 28.25 | 14 | 14 | 2537700 | 1 | 1 | 2 | N | 13915 | Cytochrome c oxidase subunit 5B, mitochondrial OS=Rattus norvegicus OX=10116 GN=Cox5b PE=1 SV=2 |
| **126** | 140 | tr\|F1LPI6\|F1LPI6_RAT | 25.48 | 0 | 0 | 390360 | 1 | 1 | 1 | N | 200075 | Peripheral-type benzodiazepine receptor-associated protein 1 OS=Rattus norvegicus OX=10116 GN=Tspoap1 PE=4 SV=1 |
| **126** | 141 | Q9JIR0\|RIMB1_RAT | 25.48 | 0 | 0 | 390360 | 1 | 1 | 1 | N | 200202 | Peripheral-type benzodiazepine receptor-associated protein 1 OS=Rattus norvegicus OX=10116 GN=Tspoap1 PE=1 SV=2 |
| **75** | 11621 | tr\|Q6AY49\|Q6AY49_RAT | 25.23 | 2 | 2 | 0 | 1 | 1 | 1 | N | 39538 | Tumor suppressor candidate 3 OS=Rattus norvegicus OX=10116 GN=Tusc3 PE=1 SV=1 |
| **104** | 181 | P61980\|HNRPK_RAT | 43855 | 2 | 2 | 463410 | 1 | 1 | 1 | N | 50976 | Heterogeneous nuclear ribonucleoprotein K OS=Rattus norvegicus OX=10116 GN=Hnrnpk PE=1 SV=1 |
| **108** | 2187 | Q06645\|AT5G1_RAT | 23.15 | 5 | 5 | 782860 | 1 | 1 | 1 | N | 14244 | ATP synthase F(0) complex subunit C1, mitochondrial OS=Rattus norvegicus OX=10116 GN=Atp5mc1 PE=1 SV=1 |
| **108** | 2188 | Q06646\|AT5G2_RAT | 23.15 | 5 | 5 | 782860 | 1 | 1 | 1 | N | 14918 | ATP synthase F(0) complex subunit C2, mitochondrial OS=Rattus norvegicus OX=10116 GN=Atp5mc2 PE=1 SV=1 |
| **108** | 2189 | tr\|Q499S2\|Q499S2_RAT | 23.15 | 5 | 5 | 782860 | 1 | 1 | 1 | N | 14745 | ATP synthase F(0) complex subunit C3, mitochondrial OS=Rattus norvegicus OX=10116 GN=Atp5mc3 PE=2 SV=1 |
| **108** | 2190 | Q71S46\|AT5G3_RAT | 23.15 | 5 | 5 | 782860 | 1 | 1 | 1 | N | 14693 | ATP synthase F(0) complex subunit C3, mitochondrial OS=Rattus norvegicus OX=10116 GN=Atp5mc3 PE=1 SV=1 |
| **108** | 2191 | tr\|A0A0G2JTN8\|A0A0G2JTN8_RAT | 23.15 | 5 | 5 | 782860 | 1 | 1 | 1 | N | 15465 | ATP synthase F(0) complex subunit C2, mitochondrial OS=Rattus norvegicus OX=10116 GN=Atp5mc2 PE=3 SV=1 |
| **109** | 622 | P18163\|ACSL1_RAT | 22.19 | 2 | 2 | 1324400 | 1 | 1 | 1 | N | 78179 | Long-chain-fatty-acid--CoA ligase 1 OS=Rattus norvegicus OX=10116 GN=Acsl1 PE=1 SV=1 |
| **60** | 369 | P11530\|DMD_RAT | 44034 | 0 | 0 | 459950 | 1 | 1 | 1 | N | 425830 | Dystrophin OS=Rattus norvegicus OX=10116 GN=Dmd PE=1 SV=2 |
| **111** | 7191 | Q6UPE0\|CHDH_RAT | 21.75 | 1 | 1 | 24171 | 1 | 1 | 1 | N | 66389 | Choline dehydrogenase, mitochondrial OS=Rattus norvegicus OX=10116 GN=Chdh PE=1 SV=1 |
| **112** | 9303 | Q63538\|MK12_RAT | 20.78 | 4 | 4 | 4843900 | 1 | 1 | 1 | Y | 41985 | Mitogen-activated protein kinase 12 OS=Rattus norvegicus OX=10116 GN=Mapk12 PE=1 SV=1 |
| **114** | 654 | tr\|A0A0G2K1W1\|A0A0G2K1W1_RAT | 20.45 | 1 | 1 | 1356800 | 1 | 1 | 1 | Y | 123811 | RAB11 family-interacting protein 5 OS=Rattus norvegicus OX=10116 GN=Rab11fip5 PE=1 SV=1 |
| **49** | 712 | D3ZLY0\|ACKMT_RAT | 20.21 | 6 | 6 | 68486 | 1 | 1 | 2 | N | 24053 | ATP synthase subunit C lysine N-methyltransferase OS=Rattus norvegicus OX=10116 GN=Atpsckmt PE=3 SV=1 |
| **77** | 2241 | tr\|Q99PW9\|Q99PW9_RAT | 44124 | 1 | 1 | 643740 | 1 | 1 | 1 | Y | 120572 | Huntingtin interacting protein 1 related (Fragment) OS=Rattus norvegicus OX=10116 GN=Hip1r PE=2 SV=1 |
| **77** | 2238 | tr\|B5DFK5\|B5DFK5_RAT | 44124 | 1 | 1 | 643740 | 1 | 1 | 1 | Y | 119424 | Hip1r protein OS=Rattus norvegicus OX=10116 GN=Hip1r PE=2 SV=1 |
| **77** | 2240 | tr\|F1LML7\|F1LML7_RAT | 44124 | 1 | 1 | 643740 | 1 | 1 | 1 | Y | 119553 | Huntingtin-interacting protein 1-related OS=Rattus norvegicus OX=10116 GN=Hip1r PE=1 SV=1 |
| **90** | 284 | Q62812\|MYH9_RAT | 44002 | 0 | 0 | 0 | 1 | 1 | 1 | N | 226336 | Myosin-9 OS=Rattus norvegicus OX=10116 GN=Myh9 PE=1 SV=3 |
| **total 110 proteins** | | | | | | | | | | | | |
