## Supplementary material for "Permeability transition pore-related changes in the proteome and channel activity of ATP synthase dimers and monomers": RLM KCl-BM

| Protein Group | Protein ID | Accession | -10lgP | Coverage (%) | Coverage (%) Sample 2 | IBAQ | #Peptides | #Unique | #Spec Sample 2 | PTM | Avg. Mass | Description |
|---|---|---|---|---|---|---|---|---|---|---|---|---|
| **2** | 12 | Q68FY0\|QCR1_RAT | 348.07 | 64 | 64 | 52406000 | 33 | 33 | 66 | Y | 52849 | Cytochrome b-c1 complex subunit 1, mitochondrial OS=Rattus norvegicus OX=10116 GN=Uqcrc1 PE=1 SV=1 |
| **3** | 1 | P32551\|QCR2_RAT | 341.51 | 80 | 80 | 110240000 | 35 | 35 | 64 | Y | 48396 | Cytochrome b-c1 complex subunit 2, mitochondrial OS=Rattus norvegicus OX=10116 GN=Uqcrc2 PE=1 SV=2 |
| **5** | 6 | Q66HF1\|NDUS1_RAT | 327.69 | 61 | 61 | 55992000 | 37 | 37 | 58 | Y | 79412 | NADH-ubiquinone oxidoreductase 75 kDa subunit, mitochondrial OS=Rattus norvegicus OX=10116 GN=Ndufs1 PE=1 SV=1 |
| **6** | 14 | P15999\|ATPA_RAT | 320.78 | 46 | 46 | 27436000 | 28 | 28 | 51 | Y | 59754 | ATP synthase subunit alpha, mitochondrial OS=Rattus norvegicus OX=10116 GN=Atp5f1a PE=1 SV=2 |
| **10** | 9 | P00507\|AATM_RAT | 304.68 | 49 | 49 | 43236000 | 22 | 22 | 37 | Y | 47314 | Aspartate aminotransferase, mitochondrial OS=Rattus norvegicus OX=10116 GN=Got2 PE=1 SV=2 |
| **1** | 7 | P10719\|ATPB_RAT | 303.01 | 79 | 79 | 36456000 | 28 | 28 | 68 | Y | 56354 | ATP synthase subunit beta, mitochondrial OS=Rattus norvegicus OX=10116 GN=Atp5f1b PE=1 SV=2 |
| **1** | 8 | tr\|G3V6D3\|G3V6D3_RAT | 303.01 | 79 | 79 | 36456000 | 28 | 28 | 68 | Y | 56345 | ATP synthase subunit beta OS=Rattus norvegicus OX=10116 GN=Atp5f1b PE=1 SV=1 |
| **4** | 2 | Q02253\|MMSA_RAT | 300.11 | 59 | 59 | 68020000 | 30 | 30 | 59 | Y | 57808 | Methylmalonate-semialdehyde dehydrogenase [acylating], mitochondrial OS=Rattus norvegicus OX=10116 GN=Aldh6a1 PE=1 SV=1 |
| **4** | 3 | tr\|G3V7J0\|G3V7J0_RAT | 300.11 | 59 | 59 | 68020000 | 30 | 30 | 59 | Y | 57748 | Aldehyde dehydrogenase family 6, subfamily A1, isoform CRA_b OS=Rattus norvegicus OX=10116 GN=Aldh6a1 PE=1 SV=1 |
| **11** | 33 | Q561S0\|NDUAA_RAT | 285.35 | 60 | 60 | 11323000 | 20 | 20 | 35 | Y | 40493 | NADH dehydrogenase [ubiquinone] 1 alpha subcomplex subunit 10, mitochondrial OS=Rattus norvegicus OX=10116 GN=Ndufa10 PE=1 SV=1 |
| **13** | 27 | Q641Y2\|NDUS2_RAT | 283.74 | 39 | 39 | 22143000 | 16 | 16 | 29 | Y | 52562 | NADH dehydrogenase [ubiquinone] iron-sulfur protein 2, mitochondrial OS=Rattus norvegicus OX=10116 GN=Ndufs2 PE=1 SV=1 |
| **7** | 15 | Q5BK63\|NDUA9_RAT | 279.21 | 71 | 71 | 28429000 | 29 | 29 | 47 | Y | 42559 | NADH dehydrogenase [ubiquinone] 1 alpha subcomplex subunit 9, mitochondrial OS=Rattus norvegicus OX=10116 GN=Ndufa9 PE=1 SV=2 |
| **9** | 4 | P67779\|PHB_RAT | 275.87 | 86 | 86 | 30873000 | 23 | 23 | 37 | N | 29820 | Prohibitin OS=Rattus norvegicus OX=10116 GN=Phb PE=1 SV=1 |
| **8** | 23 | tr\|Q5XIH3\|Q5XIH3_RAT | 261.44 | 44 | 44 | 21231000 | 20 | 20 | 38 | Y | 50731 | NADH dehydrogenase [ubiquinone] flavoprotein 1, mitochondrial OS=Rattus norvegicus OX=10116 GN=Ndufv1 PE=1 SV=1 |
| **12** | 25 | tr\|D3ZG43\|D3ZG43_RAT | 257.54 | 56 | 56 | 40764000 | 20 | 20 | 31 | N | 30226 | NADH dehydrogenase (Ubiquinone) Fe-S protein 3 (Predicted), isoform CRA_c OS=Rattus norvegicus OX=10116 GN=Ndufs3 PE=1 SV=1 |
| **18** | 29 | tr\|D4A0T0\|D4A0T0_RAT | 238.03 | 55 | 55 | 8099900 | 9 | 9 | 20 | Y | 20859 | NADH:ubiquinone oxidoreductase subunit B10 OS=Rattus norvegicus OX=10116 GN=Ndufb10 PE=1 SV=1 |
| **14** | 11 | Q5XIH7\|PHB2_RAT | 236.25 | 61 | 61 | 26326000 | 20 | 20 | 27 | Y | 33312 | Prohibitin-2 OS=Rattus norvegicus OX=10116 GN=Phb2 PE=1 SV=1 |
| **23** | 42 | tr\|D4A565\|D4A565_RAT | 231.84 | 33 | 33 | 9080800 | 8 | 8 | 14 | N | 21664 | NADH dehydrogenase (Ubiquinone) 1 beta subcomplex, 5 (Predicted), isoform CRA_b OS=Rattus norvegicus OX=10116 GN=Ndufb5 PE=1 SV=1 |
| **15** | 13 | P07756\|CPSM_RAT | 229.44 | 11 | 11 | 4242100 | 11 | 11 | 27 | N | 164579 | Carbamoyl-phosphate synthase [ammonia], mitochondrial OS=Rattus norvegicus OX=10116 GN=Cps1 PE=1 SV=1 |

| | | | | | | | | | | | | |
|---|---|---|---|---|---|---|---|---|---|---|---|---|
| **22** | 21 | Q64428\|ECHA_RAT | 225.73 | 18 | 18 | 3929700 | 10 | 10 | 14 | N | 82665 | Trifunctional enzyme subunit alpha, mitochondrial OS=Rattus norvegicus OX=10116 GN=Hadha PE=1 SV=2 |
| **19** | 28 | tr\|D3ZFQ8\|D3ZFQ8_RAT | 223.25 | 52 | 52 | 27068000 | 13 | 13 | 19 | Y | 35435 | Cytochrome c-1 OS=Rattus norvegicus OX=10116 GN=Cyc1 PE=1 SV=3 |
| **33** | 95 | Q63362\|NDUA5_RAT | 207.47 | 46 | 46 | 467140 | 6 | 6 | 8 | N | 13412 | NADH dehydrogenase [ubiquinone] 1 alpha subcomplex subunit 5 OS=Rattus norvegicus OX=10116 GN=Ndufa5 PE=1 SV=3 |
| **17** | 49 | tr\|B2RYW3\|B2RYW3_RAT | 203 | 53 | 53 | 11627000 | 12 | 12 | 22 | Y | 21892 | NADH dehydrogenase (Ubiquinone) 1 beta subcomplex, 9 OS=Rattus norvegicus OX=10116 GN=Ndufb9 PE=1 SV=1 |
| **20** | 34 | tr\|Q5RJN0\|Q5RJN0_RAT | 193.53 | 31 | 31 | 9952800 | 11 | 11 | 16 | Y | 23945 | NADH dehydrogenase (Ubiquinone) Fe-S protein 7 OS=Rattus norvegicus OX=10116 GN=Ndufs7 PE=1 SV=1 |
| **26** | 31 | tr\|B2RYS8\|B2RYS8_RAT | 187.58 | 40 | 40 | 13460000 | 6 | 6 | 12 | Y | 21959 | NADH dehydrogenase [ubiquinone] 1 beta subcomplex subunit 8, mitochondrial OS=Rattus norvegicus OX=10116 GN=Ndufb8 PE=1 SV=1 |
| **24** | 37 | P20788\|UCRI_RAT | 186.52 | 36 | 36 | 12306000 | 11 | 11 | 13 | N | 29446 | Cytochrome b-c1 complex subunit Rieske, mitochondrial OS=Rattus norvegicus OX=10116 GN=Uqcrfs1 PE=1 SV=2 |
| **16** | 22 | P19234\|NDUV2_RAT | 181.43 | 43 | 43 | 25341000 | 13 | 13 | 23 | Y | 27378 | NADH dehydrogenase [ubiquinone] flavoprotein 2, mitochondrial OS=Rattus norvegicus OX=10116 GN=Ndufv2 PE=1 SV=2 |
| **21** | 19 | P22791\|HMCS2_RAT | 181.15 | 21 | 21 | 6116700 | 9 | 9 | 15 | Y | 56912 | Hydroxymethylglutaryl-CoA synthase, mitochondrial OS=Rattus norvegicus OX=10116 GN=Hmgcs2 PE=1 SV=1 |
| **21** | 20 | tr\|Q68G44\|Q68G44_RAT | 181.15 | 21 | 21 | 6116700 | 9 | 9 | 15 | Y | 56886 | 3-hydroxy-3-methylglutaryl coenzyme A synthase OS=Rattus norvegicus OX=10116 GN=Hmgcs2 PE=1 SV=1 |
| **25** | 52 | tr\|B0BNE6\|B0BNE6_RAT | 167.4 | 40 | 40 | 6960400 | 10 | 10 | 12 | Y | 23970 | NADH dehydrogenase (Ubiquinone) Fe-S protein 8 (Predicted), isoform CRA_a OS=Rattus norvegicus OX=10116 GN=Ndufs8 PE=1 SV=1 |
| **31** | 113 | tr\|D3ZCZ9\|D3ZCZ9_RAT | 161.8 | 46 | 46 | 2062400 | 4 | 4 | 9 | Y | 13040 | NADH dehydrogenase [ubiquinone] iron-sulfur protein 6, mitochondrial OS=Rattus norvegicus OX=10116 GN=LOC100912599 PE=1 SV=1 |
| **27** | 39 | P17764\|THIL_RAT | 151.22 | 16 | 16 | 1063400 | 4 | 4 | 11 | N | 44695 | Acetyl-CoA acetyltransferase, mitochondrial OS=Rattus norvegicus OX=10116 GN=Acat1 PE=1 SV=1 |
| **32** | 30 | Q7TQ16\|QCR8_RAT | 147.19 | 50 | 50 | 4341900 | 6 | 6 | 9 | N | 9849 | Cytochrome b-c1 complex subunit 8 OS=Rattus norvegicus OX=10116 GN=Uqcrq PE=3 SV=1 |
| **29** | 46 | tr\|F1LXA0\|F1LXA0_RAT | 145.12 | 42 | 42 | 10398000 | 7 | 7 | 10 | N | 17178 | NADH dehydrogenase [ubiquinone] 1 alpha subcomplex subunit 12 OS=Rattus norvegicus OX=10116 GN=Ndufa12 PE=1 SV=2 |
| **39** | 35 | P13086\|SUCA_RAT | 143.48 | 12 | 12 | 1500300 | 4 | 4 | 6 | N | 36148 | Succinate--CoA ligase [ADP/GDP-forming] subunit alpha, mitochondrial OS=Rattus norvegicus OX=10116 GN=Suclg1 PE=2 SV=2 |
| **39** | 36 | tr\|A0A0H2UHE1\|A0A0H2UHE1_RA | 143.48 | 12 | 12 | 1500300 | 4 | 4 | 6 | N | 37560 | Succinate--CoA ligase [ADP/GDP-forming] subunit alpha, mitochondrial OS=Rattus norvegicus OX=10116 GN=Suclg1 PE=1 SV=1 |
| **38** | 116 | P35435\|ATPG_RAT | 138.08 | 28 | 28 | 1479200 | 5 | 5 | 6 | N | 30191 | ATP synthase subunit gamma, mitochondrial OS=Rattus norvegicus OX=10116 GN=Atp5f1c PE=1 SV=2 |
| **47** | 77 | tr\|Q06QK5\|Q06QK5_RAT | 127.78 | 8 | 8 | 4045400 | 4 | 4 | 4 | Y | 68606 | NADH-ubiquinone oxidoreductase chain 5 OS=Rattus norvegicus OX=10116 GN=ND5 PE=3 SV=1 |
| **47** | 78 | tr\|Q8SEZ0\|Q8SEZ0_RAT | 127.78 | 8 | 8 | 4045400 | 4 | 4 | 4 | Y | 68618 | NADH-ubiquinone oxidoreductase chain 5 OS=Rattus norvegicus OX=10116 GN=Mt-nd5 PE=3 SV=1 |

| | | | | | | | | | | | | |
|---|---|---|---|---|---|---|---|---|---|---|---|---|
| **47** | 79 | tr\|A0A096XKT9\|A0A096XKT9_RAT | 127.78 | 8 | 8 | 4045400 | 4 | 4 | 4 | Y | 68584 | NADH-ubiquinone oxidoreductase chain 5 OS=Rattus norvegicus OX=10116 GN=ND5 PE=3 SV=1 |
| **47** | 81 | P11661\|NU5M_RAT | 127.78 | 8 | 8 | 4045400 | 4 | 4 | 4 | Y | 68618 | NADH-ubiquinone oxidoreductase chain 5 OS=Rattus norvegicus OX=10116 GN=Mtnd5 PE=3 SV=3 |
| **47** | 82 | tr\|Q06QG6\|Q06QG6_RAT | 127.78 | 8 | 8 | 4045400 | 4 | 4 | 4 | Y | 68588 | NADH-ubiquinone oxidoreductase chain 5 OS=Rattus norvegicus OX=10116 GN=ND5 PE=3 SV=1 |
| **47** | 83 | tr\|Q06QA1\|Q06QA1_RAT | 127.78 | 8 | 8 | 4045400 | 4 | 4 | 4 | Y | 68574 | NADH-ubiquinone oxidoreductase chain 5 OS=Rattus norvegicus OX=10116 GN=ND5 PE=3 SV=1 |
| **42** | 86 | tr\|F1LPG5\|F1LPG5_RAT | 127.78 | 34 | 34 | 2771400 | 3 | 3 | 5 | N | 15064 | NADH:ubiquinone oxidoreductase subunit B4 OS=Rattus norvegicus OX=10116 GN=Ndufb4 PE=1 SV=1 |
| **28** | 2162 | tr\|A9UMV9\|A9UMV9_RAT | 127.36 | 54 | 54 | 2745200 | 5 | 5 | 10 | Y | 12500 | NADH:ubiquinone oxidoreductase subunit A7 OS=Rattus norvegicus OX=10116 GN=Ndufa7 PE=1 SV=1 |
| **36** | 97 | tr\|Q5PQZ9\|Q5PQZ9_RAT | 124.87 | 27 | 27 | 4741200 | 5 | 5 | 7 | N | 14359 | NADH dehydrogenase [ubiquinone] 1 subunit C2 OS=Rattus norvegicus OX=10116 GN=Ndufc2 PE=1 SV=1 |
| **57** | 47 | Q9WVK3\|PECR_RAT | 119.59 | 15 | 15 | 73229 | 3 | 3 | 3 | N | 32433 | Peroxisomal trans-2-enoyl-CoA reductase OS=Rattus norvegicus OX=10116 GN=Pecr PE=2 SV=1 |
| **57** | 48 | tr\|A0A0G2JVG4\|A0A0G2JVG4_RAT | 119.59 | 15 | 15 | 73229 | 3 | 3 | 3 | N | 32737 | Peroxisomal trans-2-enoyl-CoA reductase OS=Rattus norvegicus OX=10116 GN=Pecr PE=1 SV=1 |
| **53** | 93 | tr\|A0A0G2K8Q8\|A0A0G2K8Q8_RA | 119.32 | 52 | 52 | 2061800 | 3 | 3 | 3 | N | 7099 | Ubiquinol-cytochrome c reductase, complex III subunit X OS=Rattus norvegicus OX=10116 GN=Uqcr10 PE=1 SV=1 |
| **53** | 94 | tr\|B2RYX1\|B2RYX1_RAT | 119.32 | 50 | 50 | 2061800 | 3 | 3 | 3 | N | 7462 | LOC685322 protein OS=Rattus norvegicus OX=10116 GN=Uqcr10 PE=2 SV=1 |
| **41** | 693 | tr\|A0A0G2JVL6\|A0A0G2JVL6_RAT | 118.01 | 33 | 33 | 2506900 | 4 | 4 | 5 | N | 19965 | NADH dehydrogenase [ubiquinone] 1 alpha subcomplex subunit 8 OS=Rattus norvegicus OX=10116 GN=Ndufa8 PE=1 SV=1 |
| **35** | 150 | tr\|A0A0G2JZJ9\|A0A0G2JZJ9_RAT | 115.61 | 33 | 33 | 3999000 | 3 | 3 | 7 | N | 12730 | Uncharacterized protein OS=Rattus norvegicus OX=10116 PE=4 SV=1 |
| **35** | 149 | tr\|B5DEL8\|B5DEL8_RAT | 115.61 | 33 | 33 | 3999000 | 3 | 3 | 7 | N | 12700 | NADH dehydrogenase (Ubiquinone) Fe-S protein 5 OS=Rattus norvegicus OX=10116 GN=Ndufs5 PE=1 SV=1 |
| **34** | 2161 | tr\|D3ZE15\|D3ZE15_RAT | 114.42 | 41 | 41 | 2302400 | 6 | 6 | 8 | Y | 16777 | NADH:ubiquinone oxidoreductase subunit A13 OS=Rattus norvegicus OX=10116 GN=Ndufa13 PE=1 SV=1 |
| **69** | 98 | P29147\|BDH_RAT | 112.67 | 9 | 9 | 482530 | 2 | 2 | 2 | N | 38202 | D-beta-hydroxybutyrate dehydrogenase, mitochondrial OS=Rattus norvegicus OX=10116 GN=Bdh1 PE=1 SV=2 |
| **69** | 99 | tr\|A0A0G2JSH2\|A0A0G2JSH2_RAT | 112.67 | 9 | 9 | 482530 | 2 | 2 | 2 | N | 38333 | 3-hydroxybutyrate dehydrogenase, type 1, isoform CRA_a OS=Rattus norvegicus OX=10116 GN=Bdh1 PE=1 SV=1 |
| **68** | 2166 | Q5XIF3\|NDUS4_RAT | 110.72 | 21 | 21 | 185470 | 2 | 2 | 2 | N | 19741 | NADH dehydrogenase [ubiquinone] iron-sulfur protein 4, mitochondrial OS=Rattus norvegicus OX=10116 GN=Ndufs4 PE=1 SV=1 |
| **46** | 38 | tr\|D4A3V2\|D4A3V2_RAT | 104.02 | 33 | 33 | 3079800 | 4 | 4 | 4 | N | 15224 | NADH dehydrogenase [ubiquinone] 1 alpha subcomplex subunit 6 OS=Rattus norvegicus OX=10116 GN=Ndufa6 PE=1 SV=1 |
| **30** | 55 | tr\|D3ZZ21\|D3ZZ21_RAT | 95.6 | 32 | 32 | 5977100 | 6 | 6 | 9 | Y | 15638 | NADH dehydrogenase (Ubiquinone) 1 beta subcomplex, 6 (Predicted) OS=Rattus norvegicus OX=10116 GN=Ndufb6 PE=1 SV=1 |

| | | | | | | | | | | | | |
|---|---|---|---|---|---|---|---|---|---|---|---|---|
| **54** | 53 | P52873\|PYC_RAT | 93.57 | 5 | 5 | 1052800 | 3 | 3 | 3 | N | 129777 | Pyruvate carboxylase, mitochondrial OS=Rattus norvegicus OX=10116 GN=Pc PE=1 SV=2 |
| **54** | 54 | tr\|A0A0G2JTL5\|A0A0G2JTL5_RAT | 93.57 | 4 | 4 | 1052800 | 3 | 3 | 3 | N | 140005 | Pyruvate carboxylase, mitochondrial OS=Rattus norvegicus OX=10116 GN=Pc PE=1 SV=1 |
| **56** | 2163 | Q06647\|ATPO_RAT | 92.71 | 16 | 16 | 1485400 | 3 | 3 | 3 | N | 23398 | ATP synthase subunit O, mitochondrial OS=Rattus norvegicus OX=10116 GN=Atp5po PE=1 SV=1 |
| **50** | 5 | tr\|B2GV15\|B2GV15_RAT | 90.35 | 6 | 6 | 813930 | 2 | 2 | 4 | N | 53274 | Dihydrolipoamide acetyltransferase component of pyruvate dehydrogenase complex OS=Rattus norvegicus OX=10116 GN=Dbt PE=1 SV=1 |
| **52** | 85 | tr\|A0A0G2JVH4\|A0A0G2JVH4_RAT | 88.79 | 4 | 4 | 537270 | 3 | 3 | 3 | N | 86230 | MICOS complex subunit MIC60 OS=Rattus norvegicus OX=10116 GN=Immt PE=1 SV=1 |
| **40** | 63 | tr\|D4A7L4\|D4A7L4_RAT | 88.05 | 32 | 32 | 5648800 | 4 | 4 | 6 | Y | 17634 | NADH dehydrogenase (Ubiquinone) 1 beta subcomplex, 11 (Predicted) OS=Rattus norvegicus OX=10116 GN=Ndufb11 PE=1 SV=1 |
| **58** | 100 | tr\|D3ZS58\|D3ZS58_RAT | 83.41 | 29 | 29 | 1991500 | 3 | 3 | 3 | N | 10845 | NADH dehydrogenase [ubiquinone] 1 alpha subcomplex subunit 2 OS=Rattus norvegicus OX=10116 GN=Ndufa2 PE=1 SV=1 |
| **49** | 41 | P19511\|AT5F1_RAT | 82.59 | 10 | 10 | 1872200 | 2 | 2 | 4 | N | 28869 | ATP synthase F(0) complex subunit B1, mitochondrial OS=Rattus norvegicus OX=10116 GN=Atp5pb PE=1 SV=1 |
| **59** | 2167 | Q5M9I5\|QCR6_RAT | 81.53 | 29 | 29 | 0 | 2 | 2 | 3 | Y | 10424 | Cytochrome b-c1 complex subunit 6, mitochondrial OS=Rattus norvegicus OX=10116 GN=Uqcrh PE=3 SV=1 |
| **48** | 40 | Q60587\|ECHB_RAT | 78.47 | 5 | 5 | 322560 | 2 | 2 | 4 | N | 51414 | Trifunctional enzyme subunit beta, mitochondrial OS=Rattus norvegicus OX=10116 GN=Hadhb PE=1 SV=1 |
| **48** | 50 | tr\|A0A0G2K330\|A0A0G2K330_RAT | 78.47 | 5 | 5 | 322560 | 2 | 2 | 4 | N | 52568 | Trifunctional enzyme subunit beta, mitochondrial OS=Rattus norvegicus OX=10116 GN=Hadhb PE=1 SV=1 |
| **43** | 2164 | tr\|D3ZF13\|D3ZF13_RAT | 74.64 | 15 | 15 | 2424900 | 2 | 2 | 5 | N | 17514 | Acyl carrier protein OS=Rattus norvegicus OX=10116 GN=Ndufab1 PE=1 SV=1 |
| **51** | 103 | Q80W89\|NDUAB_RAT | 70.34 | 11 | 11 | 2360700 | 1 | 1 | 4 | N | 14854 | NADH dehydrogenase [ubiquinone] 1 alpha subcomplex subunit 11 OS=Rattus norvegicus OX=10116 GN=Ndufa11 PE=2 SV=1 |
| **55** | 2165 | tr\|D3ZLT1\|D3ZLT1_RAT | 69.67 | 31 | 31 | 605430 | 2 | 2 | 3 | Y | 16568 | NADH dehydrogenase (Ubiquinone) 1 beta subcomplex, 7 (Predicted) OS=Rattus norvegicus OX=10116 GN=Ndufb7 PE=1 SV=1 |
| **37** | 43 | tr\|A0A0G2JSH5\|A0A0G2JSH5_RAT | 66.1 | 6 | 6 | 460430 | 3 | 1 | 6 | Y | 68759 | Serum albumin OS=Rattus norvegicus OX=10116 GN=Alb PE=1 SV=1 |
| **70** | 2170 | tr\|Q8SEZ1\|Q8SEZ1_RAT | 61.52 | 13 | 13 | 213650 | 1 | 1 | 2 | Y | 13087 | NADH-ubiquinone oxidoreductase chain 3 OS=Rattus norvegicus OX=10116 GN=Mt-nd3 PE=2 SV=1 |
| **70** | 2171 | P05506\|NU3M_RAT | 61.52 | 13 | 13 | 213650 | 1 | 1 | 2 | Y | 13087 | NADH-ubiquinone oxidoreductase chain 3 OS=Rattus norvegicus OX=10116 GN=Mtnd3 PE=3 SV=3 |
| **70** | 2172 | tr\|A0A0A1FZ38\|A0A0A1FZ38_RAT | 61.52 | 13 | 13 | 213650 | 1 | 1 | 2 | Y | 13071 | NADH-ubiquinone oxidoreductase chain 3 OS=Rattus norvegicus OX=10116 GN=ND3 PE=3 SV=1 |
| **70** | 2173 | tr\|Q06QG9\|Q06QG9_RAT | 61.52 | 13 | 13 | 213650 | 1 | 1 | 2 | Y | 13075 | NADH-ubiquinone oxidoreductase chain 3 OS=Rattus norvegicus OX=10116 GN=ND3 PE=3 SV=1 |
| **64** | 66 | P05508\|NU4M_RAT | 55.32 | 6 | 6 | 348350 | 2 | 2 | 2 | N | 51783 | NADH-ubiquinone oxidoreductase chain 4 OS=Rattus norvegicus OX=10116 GN=Mtnd4 PE=3 SV=3 |

| | | | | | | | | | | | | |
|---|---|---|---|---|---|---|---|---|---|---|---|---|
| **64** | 67 | tr\|Q06QA2\|Q06QA2_RAT | 55.32 | 6 | 6 | 348350 | 2 | 2 | 2 | N | 51787 | NADH-ubiquinone oxidoreductase chain 4 OS=Rattus norvegicus OX=10116 GN=ND4 PE=3 SV=1 |
| **64** | 68 | tr\|Q7HKW3\|Q7HKW3_RAT | 55.32 | 6 | 6 | 348350 | 2 | 2 | 2 | N | 51801 | NADH-ubiquinone oxidoreductase chain 4 (Fragment) OS=Rattus norvegicus OX=10116 GN=NADH4 PE=3 SV=1 |
| **64** | 70 | tr\|D2E6K0\|D2E6K0_RAT | 55.32 | 6 | 6 | 348350 | 2 | 2 | 2 | N | 51833 | NADH-ubiquinone oxidoreductase chain 4 OS=Rattus norvegicus OX=10116 GN=ND4 PE=3 SV=1 |
| **64** | 71 | tr\|Q06Q89\|Q06Q89_RAT | 55.32 | 6 | 6 | 348350 | 2 | 2 | 2 | N | 51819 | NADH-ubiquinone oxidoreductase chain 4 OS=Rattus norvegicus OX=10116 GN=ND4 PE=3 SV=1 |
| **64** | 72 | tr\|Q35737\|Q35737_RAT | 55.32 | 6 | 6 | 348350 | 2 | 2 | 2 | N | 51765 | NADH-ubiquinone oxidoreductase chain 4 OS=Rattus norvegicus OX=10116 PE=3 SV=1 |
| **64** | 74 | tr\|Q8HIC6\|Q8HIC6_RAT | 55.32 | 6 | 6 | 348350 | 2 | 2 | 2 | N | 51783 | NADH-ubiquinone oxidoreductase chain 4 OS=Rattus norvegicus OX=10116 GN=Mt-nd4 PE=3 SV=1 |
| **64** | 75 | tr\|Q06QE1\|Q06QE1_RAT | 55.32 | 6 | 6 | 348350 | 2 | 2 | 2 | N | 51851 | NADH-ubiquinone oxidoreductase chain 4 OS=Rattus norvegicus OX=10116 GN=ND4 PE=3 SV=1 |
| **64** | 69 | tr\|Q06QG7\|Q06QG7_RAT | 55.32 | 6 | 6 | 348350 | 2 | 2 | 2 | N | 51791 | NADH-ubiquinone oxidoreductase chain 4 OS=Rattus norvegicus OX=10116 GN=ND4 PE=3 SV=1 |
| **64** | 73 | tr\|A7XYB9\|A7XYB9_RAT | 55.32 | 6 | 6 | 348350 | 2 | 2 | 2 | N | 51773 | NADH-ubiquinone oxidoreductase chain 4 OS=Rattus norvegicus OX=10116 GN=Nd4 PE=3 SV=1 |
| **45** | 44 | P02770\|ALBU_RAT | 54.91 | 5 | 5 | 1197800 | 3 | 1 | 4 | Y | 68731 | Serum albumin OS=Rattus norvegicus OX=10116 GN=Alb PE=1 SV=2 |
| **79** | 89 | tr\|Q5UAJ8\|Q5UAJ8_RAT | 51.36 | 3 | 3 | 821350 | 1 | 1 | 1 | Y | 38455 | NADH-ubiquinone oxidoreductase chain 2 OS=Rattus norvegicus OX=10116 GN=ND2 PE=3 SV=1 |
| **79** | 90 | tr\|Q06Q97\|Q06Q97_RAT | 51.36 | 3 | 3 | 821350 | 1 | 1 | 1 | Y | 38485 | NADH-ubiquinone oxidoreductase chain 2 OS=Rattus norvegicus OX=10116 GN=ND2 PE=3 SV=1 |
| **79** | 91 | tr\|A0A0A1G491\|A0A0A1G491_RAT | 51.36 | 3 | 3 | 821350 | 1 | 1 | 1 | Y | 38542 | NADH-ubiquinone oxidoreductase chain 2 OS=Rattus norvegicus OX=10116 GN=ND2 PE=3 SV=1 |
| **60** | 106 | tr\|G3V6I4\|G3V6I4_RAT | 49.43 | 9 | 9 | 604220 | 1 | 1 | 3 | N | 37791 | Mitochondrial amidoxime reducing component 1 OS=Rattus norvegicus OX=10116 GN=Marc1 PE=1 SV=1 |
| **94** | 26 | tr\|F1LN92\|F1LN92_RAT | 48.83 | 3 | 3 | 0 | 1 | 1 | 1 | N | 89352 | AFG3-like matrix AAA peptidase subunit 2 OS=Rattus norvegicus OX=10116 GN=Afg3l2 PE=1 SV=1 |
| **95** | 2174 | O70351\|HCD2_RAT | 43.64 | 8 | 8 | 66864 | 1 | 1 | 1 | N | 27246 | 3-hydroxyacyl-CoA dehydrogenase type-2 OS=Rattus norvegicus OX=10116 GN=Hsd17b10 PE=1 SV=3 |
| **95** | 2175 | tr\|B0BMW2\|B0BMW2_RAT | 43.64 | 8 | 8 | 66864 | 1 | 1 | 1 | N | 27250 | 3-hydroxyacyl-CoA dehydrogenase type-2 OS=Rattus norvegicus OX=10116 GN=Hsd17b10 PE=1 SV=1 |
| **80** | 64 | tr\|D3ZJY1\|D3ZJY1_RAT | 34.9 | 12 | 12 | 23666 | 1 | 1 | 1 | N | 30237 | Mitochondrial ribosomal protein L28 OS=Rattus norvegicus OX=10116 GN=Mrpl28 PE=1 SV=1 |
| **96** | 2176 | tr\|B0BN02\|B0BN02_RAT | 33.6 | 4 | 4 | 0 | 1 | 1 | 1 | N | 48444 | Metaxin OS=Rattus norvegicus OX=10116 GN=Mtx1 PE=1 SV=1 |
| **97** | 2177 | tr\|D4A4P3\|D4A4P3_RAT | 33.25 | 8 | 8 | 599210 | 1 | 1 | 1 | N | 11267 | NADH:ubiquinone oxidoreductase subunit B3 OS=Rattus norvegicus OX=10116 GN=Ndufb3 PE=1 SV=1 |

| | | | | | | | | | | | | |
|---|---|---|---|---|---|---|---|---|---|---|---|---|
| **63** | 167 | tr\|D4A054\|D4A054_RAT | 43921 | 1 | 1 | 225610 | 2 | 2 | 2 | N | 341403 | RAN-binding protein 2 OS=Rattus norvegicus OX=10116 GN=Ranbp2 PE=1 SV=2 |
| **63** | 168 | tr\|M0R3M4\|M0R3M4_RAT | 43921 | 1 | 1 | 225610 | 2 | 2 | 2 | N | 344396 | RAN-binding protein 2 OS=Rattus norvegicus OX=10116 GN=Ranbp2 PE=1 SV=1 |
| **98** | 2179 | tr\|G3V8T9\|G3V8T9_RAT | 28.84 | 9 | 9 | 0 | 1 | 1 | 1 | Y | 21445 | Apoptosis regulator BAX OS=Rattus norvegicus OX=10116 GN=Bax PE=1 SV=1 |
| **81** | 2180 | tr\|Q5BJZ3\|Q5BJZ3_RAT | 28.22 | 1 | 1 | 352640 | 1 | 1 | 1 | Y | 113869 | Nicotinamide nucleotide transhydrogenase OS=Rattus norvegicus OX=10116 GN=Nnt PE=1 SV=1 |
| **72** | 2181 | tr\|B2RYS2\|B2RYS2_RAT | 26.7 | 9 | 9 | 759290 | 1 | 1 | 2 | N | 13559 | Cytochrome b-c1 complex subunit 7 OS=Rattus norvegicus OX=10116 GN=Uqcrb PE=1 SV=1 |
| **99** | 65 | Q06437\|ODPAT_RAT | 26.68 | 3 | 3 | 712170 | 1 | 1 | 1 | N | 43393 | Pyruvate dehydrogenase E1 component subunit alpha, testis-specific form, mitochondrial OS=Rattus norvegicus OX=10116 GN=Pdha2 PE=1 SV=1 |
| **113** | 140 | tr\|F1LPI6\|F1LPI6_RAT | 44161 | 0 | 0 | 327450 | 1 | 1 | 1 | N | 200075 | Peripheral-type benzodiazepine receptor-associated protein 1 OS=Rattus norvegicus OX=10116 GN=Tspoap1 PE=4 SV=1 |
| **113** | 141 | Q9JIR0\|RIMB1_RAT | 44161 | 0 | 0 | 327450 | 1 | 1 | 1 | N | 200202 | Peripheral-type benzodiazepine receptor-associated protein 1 OS=Rattus norvegicus OX=10116 GN=Tspoap1 PE=1 SV=2 |
| **101** | 2184 | tr\|B2RZ31\|B2RZ31_RAT | 24.54 | 4 | 4 | 111200 | 1 | 1 | 1 | Y | 28497 | NAD-dependent protein deacetylase OS=Rattus norvegicus OX=10116 GN=Sirt3 PE=2 SV=1 |
| **101** | 2185 | tr\|C6ZII9\|C6ZII9_RAT | 24.54 | 3 | 3 | 111200 | 1 | 1 | 1 | Y | 35000 | NAD-dependent protein deacetylase OS=Rattus norvegicus OX=10116 GN=Sirt3 PE=1 SV=1 |
| **73** | 246 | Q63704\|CPT1B_RAT | 24.5 | 1 | 1 | 0 | 1 | 1 | 2 | N | 88217 | Carnitine O-palmitoyltransferase 1, muscle isoform OS=Rattus norvegicus OX=10116 GN=Cpt1b PE=1 SV=1 |
| **73** | 288 | F1LN46\|CPT1C_RAT | 24.5 | 1 | 1 | 0 | 1 | 1 | 2 | N | 90170 | Carnitine O-palmitoyltransferase 1, brain isoform OS=Rattus norvegicus OX=10116 GN=Cpt1c PE=1 SV=1 |
| **73** | 330 | tr\|M0RAQ6\|M0RAQ6_RAT | 24.5 | 1 | 1 | 0 | 1 | 1 | 2 | N | 99557 | Hexokinase-1 OS=Rattus norvegicus OX=10116 GN=Hk1 PE=1 SV=2 |
| **73** | 333 | P05708\|HXK1_RAT | 24.5 | 1 | 1 | 0 | 1 | 1 | 2 | N | 102408 | Hexokinase-1 OS=Rattus norvegicus OX=10116 GN=Hk1 PE=1 SV=4 |
| **102** | 111 | Q5XJW2\|G45IP_RAT | 24.25 | 4 | 4 | 0 | 1 | 1 | 1 | N | 26467 | Growth arrest and DNA damage-inducible proteins-interacting protein 1 OS=Rattus norvegicus OX=10116 GN=Gadd45gip1 PE=2 SV=2 |
| **103** | 157 | tr\|F1LTG5\|F1LTG5_RAT | 24.17 | 14 | 14 | 152290 | 1 | 1 | 1 | N | 13829 | Uncharacterized protein OS=Rattus norvegicus OX=10116 PE=4 SV=2 |
| **103** | 158 | P12075\|COX5B_RAT | 24.17 | 14 | 14 | 152290 | 1 | 1 | 1 | N | 13915 | Cytochrome c oxidase subunit 5B, mitochondrial OS=Rattus norvegicus OX=10116 GN=Cox5b PE=1 SV=2 |
| **104** | 2186 | P07895\|SODM_RAT | 44098 | 6 | 6 | 0 | 1 | 1 | 1 | N | 24674 | Superoxide dismutase [Mn], mitochondrial OS=Rattus norvegicus OX=10116 GN=Sod2 PE=1 SV=2 |
| **106** | 2187 | Q06645\|AT5G1_RAT | 43975 | 5 | 5 | 0 | 1 | 1 | 1 | N | 14244 | ATP synthase F(0) complex subunit C1, mitochondrial OS=Rattus norvegicus OX=10116 GN=Atp5mc1 PE=1 SV=1 |
| **106** | 2188 | Q06646\|AT5G2_RAT | 43975 | 5 | 5 | 0 | 1 | 1 | 1 | N | 14918 | ATP synthase F(0) complex subunit C2, mitochondrial OS=Rattus norvegicus OX=10116 GN=Atp5mc2 PE=1 SV=1 |

| | | | | | | | | | | | | |
|---|---|---|---|---|---|---|---|---|---|---|---|---|
| **106** | 2189 | tr\|Q499S2\|Q499S2_RAT | 43975 | 5 | 5 | 0 | 1 | 1 | 1 | N | 14745 | ATP synthase F(0) complex subunit C3, mitochondrial OS=Rattus norvegicus OX=10116 GN=Atp5mc3 PE=2 SV=1 |
| **106** | 2190 | Q71S46\|AT5G3_RAT | 43975 | 5 | 5 | 0 | 1 | 1 | 1 | N | 14693 | ATP synthase F(0) complex subunit C3, mitochondrial OS=Rattus norvegicus OX=10116 GN=Atp5mc3 PE=1 SV=1 |
| **106** | 2191 | tr\|A0A0G2JTN8\|A0A0G2JTN8_RAT | 43975 | 5 | 5 | 0 | 1 | 1 | 1 | N | 15465 | ATP synthase F(0) complex subunit C2, mitochondrial OS=Rattus norvegicus OX=10116 GN=Atp5mc2 PE=3 SV=1 |
| **105** | 108 | O70595\|ABCB6_RAT | 23.93 | 1 | 1 | 302320 | 1 | 1 | 1 | Y | 93305 | ATP-binding cassette sub-family B member 6, mitochondrial OS=Rattus norvegicus OX=10116 GN=Abcb6 PE=2 SV=1 |
| **109** | 2194 | tr\|Q5U2T0\|Q5U2T0_RAT | 22.28 | 5 | 5 | 36644 | 1 | 1 | 1 | N | 44494 | Death associated protein 3 OS=Rattus norvegicus OX=10116 GN=Dap3 PE=2 SV=1 |
| **109** | 2195 | tr\|F7EZZ0\|F7EZZ0_RAT | 22.28 | 5 | 5 | 36644 | 1 | 1 | 1 | N | 45112 | Death-associated protein 3 OS=Rattus norvegicus OX=10116 GN=Dap3 PE=1 SV=1 |
| **109** | 256 | tr\|A0A0G2K264\|A0A0G2K264_RAT | 22.28 | 5 | 5 | 36644 | 1 | 1 | 1 | N | 46666 | Death-associated protein 3 OS=Rattus norvegicus OX=10116 GN=Dap3 PE=1 SV=1 |
| **87** | 2193 | Q5U310\|ARMX1_RAT | 21.96 | 2 | 2 | 171140 | 1 | 1 | 1 | N | 51049 | Armadillo repeat-containing X-linked protein 1 OS=Rattus norvegicus OX=10116 GN=Armcx1 PE=2 SV=1 |
| **110** | 2196 | tr\|A0A0U1RRQ1\|A0A0U1RRQ1_RA | 21.41 | 9 | 9 | 368300 | 1 | 1 | 1 | N | 7692 | NADH dehydrogenase [ubiquinone] 1 alpha subcomplex subunit 1 OS=Rattus norvegicus OX=10116 GN=Ndufa1 PE=1 SV=1 |
| **110** | 2197 | tr\|G3V9S6\|G3V9S6_RAT | 21.41 | 9 | 9 | 368300 | 1 | 1 | 1 | N | 8148 | NADH dehydrogenase [ubiquinone] 1 alpha subcomplex subunit 1 OS=Rattus norvegicus OX=10116 GN=Ndufa1 PE=1 SV=2 |
| **110** | 2198 | tr\|B4F7B9\|B4F7B9_RAT | 21.41 | 7 | 7 | 368300 | 1 | 1 | 1 | N | 9778 | Uncharacterized protein (Fragment) OS=Rattus norvegicus OX=10116 GN=Ndufa1 PE=2 SV=1 |
| **88** | 191 | Q8R5G2\|MUTYH_RAT | 21.22 | 2 | 2 | 845700 | 1 | 1 | 1 | N | 57919 | Adenine DNA glycosylase OS=Rattus norvegicus OX=10116 GN=Mutyh PE=2 SV=1 |
| **112** | 2199 | Q32Q86\|CRY1_RAT | 20.96 | 1 | 1 | 2160700 | 1 | 1 | 1 | N | 66231 | Cryptochrome-1 OS=Rattus norvegicus OX=10116 GN=Cry1 PE=1 SV=1 |
| **115** | 2201 | Q9Z2L9\|NDRG4_RAT | 20.46 | 3 | 3 | 0 | 1 | 1 | 1 | N | 38487 | Protein NDRG4 OS=Rattus norvegicus OX=10116 GN=Ndrg4 PE=1 SV=1 |
| **116** | 382 | tr\|A0A0G2JYY2\|A0A0G2JYY2_RAT | 20.27 | 2 | 2 | 1501900 | 1 | 1 | 1 | N | 62571 | 5-aminolevulinate synthase OS=Rattus norvegicus OX=10116 GN=Alas1 PE=1 SV=1 |
| **116** | 383 | tr\|A0A0G2K962\|A0A0G2K962_RAT | 20.27 | 2 | 2 | 1501900 | 1 | 1 | 1 | N | 71039 | 5-aminolevulinate synthase OS=Rattus norvegicus OX=10116 GN=Alas1 PE=1 SV=1 |
| **116** | 384 | P13195\|HEM1_RAT | 20.27 | 2 | 2 | 1501900 | 1 | 1 | 1 | N | 71021 | 5-aminolevulinate synthase, nonspecific, mitochondrial OS=Rattus norvegicus OX=10116 GN=Alas1 PE=2 SV=2 |
| **total 136 proteins** | | | | | | | | | | | | |
