## Supplementary material for "Permeability transition pore-related changes in the proteome and channel activity of ATP synthase dimers and monomers": RLM KCl-BM

| rotein Grou | Protein ID | Accession | -10lgP | overage (% | age (%) Sa | IBAQ | #Peptides | #Unique | pec Sampl | PTM | Avg. Mass | Description |
|---|---|---|---|---|---|---|---|---|---|---|---|---|
| **2** | 13 | P15999\|AT | 360.38 | 52 | 52 | 99869000 | 38 | 38 | 73 | Y | 59754 | ATP synthase subunit alpha, mitochondrial OS=Rattus norvegicus OX=10116 GN=Atp5f1a PE=1 SV=2 |
| **1** | 6 | P52873\|PY | 341.35 | 48 | 48 | 79987000 | 60 | 59 | 74 | Y | 129777 | Pyruvate carboxylase, mitochondrial OS=Rattus norvegicus OX=10116 GN=Pc PE=1 SV=2 |
| **1** | 7 | tr\|A0A0G2 | 341.35 | 44 | 44 | 79987000 | 60 | 59 | 74 | Y | 140005 | Pyruvate carboxylase, mitochondrial OS=Rattus norvegicus OX=10116 GN=Pc PE=1 SV=1 |
| **3** | 3 | P07756\|CP | 338.51 | 35 | 35 | 54889000 | 45 | 45 | 54 | Y | 164579 | Carbamoyl-phosphate synthase [ammonia], mitochondrial OS=Rattus norvegicus OX=10116 GN=Cps1 PE=1 SV=1 |
| **4** | 15 | Q68FY0\|Q | 312.74 | 51 | 51 | 64196000 | 27 | 27 | 53 | Y | 52849 | Cytochrome b-c1 complex subunit 1, mitochondrial OS=Rattus norvegicus OX=10116 GN=Uqcrc1 PE=1 SV=1 |
| **7** | 5 | Q64428\|E | 311.16 | 36 | 36 | 26912000 | 28 | 28 | 35 | Y | 82665 | Trifunctional enzyme subunit alpha, mitochondrial OS=Rattus norvegicus OX=10116 GN=Hadha PE=1 SV=2 |
| **5** | 9 | tr\|G3V6D3 | 292.82 | 64 | 64 | 95063000 | 30 | 30 | 47 | Y | 56345 | ATP synthase subunit beta OS=Rattus norvegicus OX=10116 GN=Atp5f1b PE=1 SV=1 |
| **5** | 8 | P10719\|AT | 292.82 | 64 | 64 | 95063000 | 30 | 30 | 47 | Y | 56354 | ATP synthase subunit beta, mitochondrial OS=Rattus norvegicus OX=10116 GN=Atp5f1b PE=1 SV=2 |
| **6** | 1 | P32551\|Q | 291.19 | 63 | 63 | 72356000 | 30 | 30 | 36 | Y | 48396 | Cytochrome b-c1 complex subunit 2, mitochondrial OS=Rattus norvegicus OX=10116 GN=Uqcrc2 PE=1 SV=2 |
| **9** | 103 | P04762\|CA | 271.11 | 43 | 43 | 24425000 | 22 | 21 | 27 | Y | 59757 | Catalase OS=Rattus norvegicus OX=10116 GN=Cat PE=1 SV=3 |
| **8** | 10 | P22791\|H | 263.3 | 47 | 47 | 34999000 | 26 | 26 | 29 | Y | 56912 | Hydroxymethylglutaryl-CoA synthase, mitochondrial OS=Rattus norvegicus OX=10116 GN=Hmgcs2 PE=1 SV=1 |
| **8** | 11 | tr\|Q68G44 | 263.3 | 47 | 47 | 34999000 | 26 | 26 | 29 | Y | 56886 | 3-hydroxy-3-methylglutaryl coenzyme A synthase OS=Rattus norvegicus OX=10116 GN=Hmgcs2 PE=1 SV=1 |
| **10** | 31 | Q60587\|E | 229.74 | 32 | 32 | 12872000 | 14 | 13 | 14 | Y | 51414 | Trifunctional enzyme subunit beta, mitochondrial OS=Rattus norvegicus OX=10116 GN=Hadhb PE=1 SV=1 |
| **18** | 2 | P10860\|D | 224.24 | 19 | 19 | 8337700 | 8 | 8 | 10 | N | 61416 | Glutamate dehydrogenase 1, mitochondrial OS=Rattus norvegicus OX=10116 GN=Glud1 PE=1 SV=2 |
| **12** | 45 | P20788\|U | 215.16 | 41 | 41 | 15370000 | 12 | 12 | 14 | N | 29446 | Cytochrome b-c1 complex subunit Rieske, mitochondrial OS=Rattus norvegicus OX=10116 GN=Uqcrfs1 PE=1 SV=2 |
| **15** | 61 | P63039\|C | 193.4 | 21 | 21 | 5668300 | 12 | 11 | 12 | N | 60956 | 60 kDa heat shock protein, mitochondrial OS=Rattus norvegicus OX=10116 GN=Hspd1 PE=1 SV=1 |
| **15** | 62 | tr\|A0A482 | 193.4 | 21 | 21 | 5668300 | 12 | 11 | 12 | N | 60956 | Hsp60 OS=Rattus norvegicus OX=10116 GN=Hspd1 PE=2 SV=1 |
| **13** | 28 | P18163\|A | 187.16 | 13 | 13 | 7044600 | 11 | 7 | 13 | Y | 78179 | Long-chain-fatty-acid--CoA ligase 1 OS=Rattus norvegicus OX=10116 GN=Acsl1 PE=1 SV=1 |
| **11** | 42 | tr\|D3ZFQ8 | 183.71 | 39 | 39 | 16026000 | 10 | 10 | 14 | Y | 35435 | Cytochrome c-1 OS=Rattus norvegicus OX=10116 GN=Cyc1 PE=1 SV=3 |
| **14** | 55 | P24329\|TH | 183.23 | 32 | 32 | 8548700 | 9 | 8 | 13 | N | 33407 | Thiosulfate sulfurtransferase OS=Rattus norvegicus OX=10116 GN=Tst PE=1 SV=3 |
| **35** | 64 | Q02253\|M | 180.77 | 13 | 13 | 3605700 | 5 | 5 | 5 | N | 57808 | Methylmalonate-semialdehyde dehydrogenase [acylating], mitochondrial OS=Rattus norvegicus OX=10116 GN=Aldh6a1 PE=1 SV=1 |
| **35** | 65 | tr\|G3V7J0\| | 180.77 | 13 | 13 | 3605700 | 5 | 5 | 5 | N | 57748 | Aldehyde dehydrogenase family 6, subfamily A1, isoform CRA_b OS=Rattus norvegicus OX=10116 GN=Aldh6a1 PE=1 SV=1 |
| **20** | 1579 | tr\|D4ACE9 | 173.93 | 10 | 10 | 4838200 | 9 | 8 | 9 | Y | 103115 | Alpha-aminoadipic semialdehyde synthase, mitochondrial OS=Rattus norvegicus OX=10116 GN=Aass PE=1 SV=3 |
| **20** | 1580 | A2VCW9\|A | 173.93 | 10 | 10 | 4838200 | 9 | 8 | 9 | Y | 102908 | Alpha-aminoadipic semialdehyde synthase, mitochondrial OS=Rattus norvegicus OX=10116 GN=Aass PE=2 SV=1 |
| **16** | 114 | P19511\|AT | 170.49 | 30 | 30 | 11113000 | 8 | 8 | 11 | N | 28869 | ATP synthase F(0) complex subunit B1, mitochondrial OS=Rattus norvegicus OX=10116 GN=Atp5pb PE=1 SV=1 |
| **45** | 46 | P17764\|TH | 170.37 | 13 | 13 | 1632300 | 4 | 4 | 4 | N | 44695 | Acetyl-CoA acetyltransferase, mitochondrial OS=Rattus norvegicus OX=10116 GN=Acat1 PE=1 SV=1 |
| **21** | 97 | P35435\|AT | 169.68 | 29 | 29 | 6072900 | 9 | 9 | 9 | Y | 30191 | ATP synthase subunit gamma, mitochondrial OS=Rattus norvegicus OX=10116 GN=Atp5f1c PE=1 SV=2 |
| **21** | 95 | tr\|Q6QI09\| | 169.68 | 13 | 13 | 6072900 | 9 | 9 | 9 | Y | 67721 | ATP synthase subunit gamma, mitochondrial OS=Rattus norvegicus OX=10116 GN=Taf3 PE=1 SV=1 |
| **30** | 44 | P16970\|AB | 168.52 | 14 | 14 | 3178700 | 6 | 6 | 6 | N | 75316 | ATP-binding cassette sub-family D member 3 OS=Rattus norvegicus OX=10116 GN=Abcd3 PE=1 SV=3 |
| **29** | 204 | P31399\|AT | 168.3 | 41 | 41 | 3042300 | 6 | 6 | 6 | N | 18763 | ATP synthase subunit d, mitochondrial OS=Rattus norvegicus OX=10116 GN=Atp5pd PE=1 SV=3 |
| **17** | 4 | P00507\|AA | 167.1 | 20 | 20 | 8105600 | 9 | 9 | 11 | Y | 47314 | Aspartate aminotransferase, mitochondrial OS=Rattus norvegicus OX=10116 GN=Got2 PE=1 SV=2 |
| **22** | 188 | Q06647\|A | 166.12 | 34 | 34 | 7368900 | 8 | 8 | 9 | Y | 23398 | ATP synthase subunit O, mitochondrial OS=Rattus norvegicus OX=10116 GN=Atp5po PE=1 SV=1 |
| **25** | 47 | P0C2X9\|AL | 162.06 | 13 | 13 | 3877700 | 6 | 6 | 6 | N | 61869 | Delta-1-pyrroline-5-carboxylate dehydrogenase, mitochondrial OS=Rattus norvegicus OX=10116 GN=Aldh4a1 PE=1 SV=1 |
| **23** | 32 | P13086\|SU | 150.81 | 16 | 16 | 4132500 | 6 | 6 | 7 | N | 36148 | Succinate--CoA ligase [ADP/GDP-forming] subunit alpha, mitochondrial OS=Rattus norvegicus OX=10116 GN=Suclg1 PE=2 SV=2 |
| **23** | 33 | tr\|A0A0H2 | 150.81 | 16 | 16 | 4132500 | 6 | 6 | 7 | N | 37560 | Succinate--CoA ligase [ADP/GDP-forming] subunit alpha, mitochondrial OS=Rattus norvegicus OX=10116 GN=Suclg1 PE=1 SV=1 |
| **27** | 91 | P45953\|A | 147.05 | 10 | 10 | 4548300 | 5 | 5 | 6 | N | 70749 | Very long-chain specific acyl-CoA dehydrogenase, mitochondrial OS=Rattus norvegicus OX=10116 GN=Acadvl PE=1 SV=1 |
| **27** | 92 | tr\|Q5M9H | 147.05 | 10 | 10 | 4548300 | 5 | 5 | 6 | N | 70821 | Acyl-Coenzyme A dehydrogenase, very long chain OS=Rattus norvegicus OX=10116 GN=Acadvl PE=1 SV=1 |
| **19** | 50 | P10888\|CC | 143.7 | 35 | 35 | 7154100 | 8 | 8 | 10 | N | 19515 | Cytochrome c oxidase subunit 4 isoform 1, mitochondrial OS=Rattus norvegicus OX=10116 GN=Cox4i1 PE=1 SV=1 |
| **33** | 133 | P12075\|CC | 133.5 | 37 | 37 | 4379500 | 5 | 5 | 6 | Y | 13915 | Cytochrome c oxidase subunit 5B, mitochondrial OS=Rattus norvegicus OX=10116 GN=Cox5b PE=1 SV=2 |
| **43** | 76 | O70351\|H | 131.08 | 22 | 22 | 1861400 | 3 | 3 | 4 | N | 27246 | 3-hydroxyacyl-CoA dehydrogenase type-2 OS=Rattus norvegicus OX=10116 GN=Hsd17b10 PE=1 SV=3 |
| **43** | 77 | tr\|B0BMW | 131.08 | 22 | 22 | 1861400 | 3 | 3 | 4 | N | 27250 | 3-hydroxyacyl-CoA dehydrogenase type-2 OS=Rattus norvegicus OX=10116 GN=Hsd17b10 PE=1 SV=1 |
| **26** | 12418 | tr\|A0A0G2 | 126.42 | 8 | 8 | 426670 | 6 | 1 | 6 | N | 79812 | Propionyl-CoA carboxylase alpha chain, mitochondrial OS=Rattus norvegicus OX=10116 GN=Pcca PE=1 SV=1 |
| **26** | 12419 | tr\|A0A0G2 | 126.42 | 7 | 7 | 426670 | 6 | 1 | 6 | N | 81413 | Propionyl-CoA carboxylase alpha chain, mitochondrial OS=Rattus norvegicus OX=10116 GN=Pcca PE=1 SV=1 |
| **28** | 908 | P14882\|PC | 124.78 | 8 | 8 | 1582100 | 6 | 1 | 6 | Y | 81623 | Propionyl-CoA carboxylase alpha chain, mitochondrial OS=Rattus norvegicus OX=10116 GN=Pcca PE=1 SV=3 |
| **46** | 12420 | P07633\|PC | 121.69 | 7 | 7 | 2410500 | 3 | 3 | 3 | N | 58626 | Propionyl-CoA carboxylase beta chain, mitochondrial OS=Rattus norvegicus OX=10116 GN=Pccb PE=2 SV=1 |
| **46** | 12417 | tr\|Q68FZ8\| | 121.69 | 7 | 7 | 2410500 | 3 | 3 | 3 | N | 58678 | Propionyl coenzyme A carboxylase, beta polypeptide OS=Rattus norvegicus OX=10116 GN=Pccb PE=1 SV=1 |
| **37** | 70 | tr\|B2RYT5\| | 117.43 | 45 | 45 | 8005400 | 5 | 5 | 5 | N | 12651 | Cox7a2l protein OS=Rattus norvegicus OX=10116 GN=Cox7a2l PE=2 SV=1 |
| **42** | 158 | P11240\|CC | 100.74 | 25 | 25 | 1510100 | 4 | 4 | 4 | N | 16130 | Cytochrome c oxidase subunit 5A, mitochondrial OS=Rattus norvegicus OX=10116 GN=Cox5a PE=1 SV=1 |
| **32** | 101 | O88813\|A | 100.56 | 10 | 10 | 1463700 | 6 | 3 | 6 | Y | 76405 | Long-chain-fatty-acid--CoA ligase 5 OS=Rattus norvegicus OX=10116 GN=Acsl5 PE=1 SV=1 |
| **32** | 102 | tr\|F1LPB3\| | 100.56 | 10 | 10 | 1463700 | 6 | 3 | 6 | Y | 76559 | Long-chain-fatty-acid--CoA ligase 5 OS=Rattus norvegicus OX=10116 GN=Acsl5 PE=1 SV=3 |
| **40** | 63 | Q09073\|A | 96.09 | 13 | 13 | 2830300 | 4 | 4 | 4 | N | 32901 | ADP/ATP translocase 2 OS=Rattus norvegicus OX=10116 GN=Slc25a5 PE=1 SV=3 |
| **36** | 109 | tr\|Q37652 | 93.98 | 15 | 15 | 5559100 | 4 | 4 | 5 | Y | 26007 | Cytochrome c oxidase subunit 2 OS=Rattus norvegicus OX=10116 GN=COXII PE=3 SV=1 |
| **36** | 80 | tr\|S5RZM8 | 93.98 | 15 | 15 | 5559100 | 4 | 4 | 5 | Y | 25958 | Cytochrome c oxidase subunit 2 OS=Rattus norvegicus OX=10116 GN=COX2 PE=3 SV=1 |
| **36** | 81 | P00406\|CC | 93.98 | 15 | 15 | 5559100 | 4 | 4 | 5 | Y | 25928 | Cytochrome c oxidase subunit 2 OS=Rattus norvegicus OX=10116 GN=Mtco2 PE=1 SV=3 |
| **36** | 68 | tr\|Q5UAJ6 | 93.98 | 15 | 15 | 5559100 | 4 | 4 | 5 | Y | 25942 | Cytochrome c oxidase subunit 2 OS=Rattus norvegicus OX=10116 GN=COX2 PE=3 SV=1 |
| **36** | 82 | tr\|Q8SEZ5\| | 93.98 | 15 | 15 | 5559100 | 4 | 4 | 5 | Y | 25928 | Cytochrome c oxidase subunit 2 OS=Rattus norvegicus OX=10116 GN=Mt-co2 PE=1 SV=1 |
| **36** | 83 | tr\|A0A097 | 93.98 | 15 | 15 | 5559100 | 4 | 4 | 5 | Y | 25894 | Cytochrome c oxidase subunit 2 OS=Rattus norvegicus OX=10116 GN=COX2 PE=3 SV=1 |
| **76** | 205 | Q5M9I5\|Q | 90.61 | 21 | 21 | 2255800 | 1 | 1 | 2 | N | 10424 | Cytochrome b-c1 complex subunit 6, mitochondrial OS=Rattus norvegicus OX=10116 GN=Uqcrh PE=3 SV=1 |
| **64** | 51 | tr\|Q5EBA4 | 90.35 | 8 | 8 | 835520 | 2 | 2 | 2 | N | 33215 | Nipsnap1 protein (Fragment) OS=Rattus norvegicus OX=10116 GN=Nipsnap1 PE=2 SV=1 |

<table>
<tr><td>64</td><td>52</td><td>tr|G3V728</td><td>90.35</td><td>8</td><td>8</td><td>835520</td><td>2</td><td>2</td><td>2</td><td>N</td><td>33346</td><td>4-nitrophenylphosphatase domain and non-neuronal SNAP25-like protein homolog 1 (C. elegans), isoform CRA_b OS=Rattus norvegicus OX=10116 GN=Nipsnap1 PE=1 SV=1</td></tr>
<tr><td>75</td><td>248</td><td>tr|G3V6I4|</td><td>86.14</td><td>9</td><td>9</td><td>511570</td><td>1</td><td>1</td><td>2</td><td>N</td><td>37791</td><td>Mitochondrial amidoxime reducing component 1 OS=Rattus norvegicus OX=10116 GN=Marc1 PE=1 SV=1</td></tr>
<tr><td>153</td><td>12439</td><td>Q5XIT9|M</td><td>81.51</td><td>4</td><td>4</td><td>139210</td><td>1</td><td>1</td><td>1</td><td>N</td><td>61517</td><td>Methylcrotonoyl-CoA carboxylase beta chain, mitochondrial OS=Rattus norvegicus OX=10116 GN=Mccc2 PE=2 SV=1</td></tr>
<tr><td>44</td><td>39</td><td>P29147|BD</td><td>80.43</td><td>8</td><td>8</td><td>2888000</td><td>4</td><td>4</td><td>4</td><td>N</td><td>38202</td><td>D-beta-hydroxybutyrate dehydrogenase, mitochondrial OS=Rattus norvegicus OX=10116 GN=Bdh1 PE=1 SV=2</td></tr>
<tr><td>44</td><td>40</td><td>tr|A0A0G2</td><td>80.43</td><td>8</td><td>8</td><td>2888000</td><td>4</td><td>4</td><td>4</td><td>N</td><td>38333</td><td>3-hydroxybutyrate dehydrogenase, type 1, isoform CRA_a OS=Rattus norvegicus OX=10116 GN=Bdh1 PE=1 SV=1</td></tr>
<tr><td>47</td><td>167</td><td>P35434|AT</td><td>78.79</td><td>17</td><td>17</td><td>4553400</td><td>3</td><td>3</td><td>3</td><td>N</td><td>17595</td><td>ATP synthase subunit delta, mitochondrial OS=Rattus norvegicus OX=10116 GN=Atp5f1d PE=1 SV=2</td></tr>
<tr><td>47</td><td>168</td><td>tr|G3V7Y3</td><td>78.79</td><td>17</td><td>17</td><td>4553400</td><td>3</td><td>3</td><td>3</td><td>N</td><td>17563</td><td>ATP synthase subunit delta, mitochondrial OS=Rattus norvegicus OX=10116 GN=Atp5f1d PE=1 SV=1</td></tr>
<tr><td>73</td><td>48</td><td>Q9WVK3|P</td><td>75.64</td><td>6</td><td>6</td><td>515410</td><td>2</td><td>2</td><td>2</td><td>N</td><td>32433</td><td>Peroxisomal trans-2-enoyl-CoA reductase OS=Rattus norvegicus OX=10116 GN=Pecr PE=2 SV=1</td></tr>
<tr><td>73</td><td>49</td><td>tr|A0A0G2</td><td>75.64</td><td>6</td><td>6</td><td>515410</td><td>2</td><td>2</td><td>2</td><td>N</td><td>32737</td><td>Peroxisomal trans-2-enoyl-CoA reductase OS=Rattus norvegicus OX=10116 GN=Pecr PE=1 SV=1</td></tr>
<tr><td>58</td><td>53</td><td>B3DMA2|A</td><td>74.9</td><td>2</td><td>2</td><td>644530</td><td>2</td><td>2</td><td>2</td><td>N</td><td>87371</td><td>Acyl-CoA dehydrogenase family member 11 OS=Rattus norvegicus OX=10116 GN=Acad11 PE=1 SV=1</td></tr>
<tr><td>48</td><td>96</td><td>tr|D3ZFJ6|</td><td>71.18</td><td>10</td><td>10</td><td>649370</td><td>3</td><td>3</td><td>3</td><td>N</td><td>60420</td><td>Lactamase, beta OS=Rattus norvegicus OX=10116 GN=Lactb PE=1 SV=1</td></tr>
<tr><td>52</td><td>29</td><td>tr|A0A140</td><td>70.98</td><td>6</td><td>6</td><td>500310</td><td>3</td><td>3</td><td>3</td><td>N</td><td>67049</td><td>MICOS complex subunit MIC60 OS=Rattus norvegicus OX=10116 GN=Immt PE=1 SV=1</td></tr>
<tr><td>52</td><td>30</td><td>Q3KR86|M</td><td>70.98</td><td>6</td><td>6</td><td>500310</td><td>3</td><td>3</td><td>3</td><td>N</td><td>67177</td><td>MICOS complex subunit Mic60 (Fragment) OS=Rattus norvegicus OX=10116 GN=Immt PE=1 SV=1</td></tr>
<tr><td>52</td><td>22</td><td>tr|A0A0G2</td><td>70.98</td><td>5</td><td>5</td><td>500310</td><td>3</td><td>3</td><td>3</td><td>N</td><td>86230</td><td>MICOS complex subunit MIC60 OS=Rattus norvegicus OX=10116 GN=Immt PE=1 SV=1</td></tr>
<tr><td>103</td><td>1310</td><td>P07872|AC</td><td>69.84</td><td>3</td><td>3</td><td>285760</td><td>1</td><td>1</td><td>1</td><td>N</td><td>74679</td><td>Peroxisomal acyl-coenzyme A oxidase 1 OS=Rattus norvegicus OX=10116 GN=Acox1 PE=1 SV=1</td></tr>
<tr><td>213</td><td>8227</td><td>Q6UPE0|C</td><td>69.29</td><td>4</td><td>4</td><td>213210</td><td>1</td><td>1</td><td>1</td><td>N</td><td>66389</td><td>Choline dehydrogenase, mitochondrial OS=Rattus norvegicus OX=10116 GN=Chdh PE=1 SV=1</td></tr>
<tr><td>74</td><td>36</td><td>tr|D3ZG43</td><td>64.2</td><td>6</td><td>6</td><td>692750</td><td>2</td><td>2</td><td>2</td><td>N</td><td>30226</td><td>NADH dehydrogenase (Ubiquinone) Fe-S protein 3 (Predicted), isoform CRA_c OS=Rattus norvegicus OX=10116 GN=Ndufs3 PE=1 SV=1</td></tr>
<tr><td>123</td><td>456</td><td>A0A0G2K0</td><td>63.46</td><td>2</td><td>2</td><td>234970</td><td>1</td><td>1</td><td>1</td><td>N</td><td>74675</td><td>Acyl-CoA synthetase short-chain family member 3, mitochondrial OS=Rattus norvegicus OX=10116 GN=Acss3 PE=1 SV=1</td></tr>
<tr><td>53</td><td>94</td><td>Q7TQ16|Q</td><td>60.59</td><td>30</td><td>30</td><td>1922000</td><td>3</td><td>3</td><td>3</td><td>N</td><td>9849</td><td>Cytochrome b-c1 complex subunit 8 OS=Rattus norvegicus OX=10116 GN=Uqcrq PE=3 SV=1</td></tr>
<tr><td>80</td><td>189</td><td>tr|B2RYS2|</td><td>60.08</td><td>14</td><td>14</td><td>1110900</td><td>2</td><td>2</td><td>2</td><td>N</td><td>13559</td><td>Cytochrome b-c1 complex subunit 7 OS=Rattus norvegicus OX=10116 GN=Uqcrb PE=1 SV=1</td></tr>
<tr><td>63</td><td>56</td><td>P04636|M</td><td>58.33</td><td>5</td><td>5</td><td>869950</td><td>2</td><td>2</td><td>2</td><td>N</td><td>35684</td><td>Malate dehydrogenase, mitochondrial OS=Rattus norvegicus OX=10116 GN=Mdh2 PE=1 SV=2</td></tr>
<tr><td>79</td><td>154</td><td>tr|A0A0G2</td><td>58</td><td>24</td><td>24</td><td>3819700</td><td>2</td><td>2</td><td>2</td><td>N</td><td>7099</td><td>Ubiquinol-cytochrome c reductase, complex III subunit X OS=Rattus norvegicus OX=10116 GN=Uqcr10 PE=1 SV=1</td></tr>
<tr><td>79</td><td>172</td><td>tr|B2RYX1|</td><td>58</td><td>23</td><td>23</td><td>3819700</td><td>2</td><td>2</td><td>2</td><td>N</td><td>7462</td><td>LOC685322 protein OS=Rattus norvegicus OX=10116 GN=Uqcr10 PE=2 SV=1</td></tr>
<tr><td>214</td><td>254</td><td>tr|Q7H115</td><td>53.03</td><td>5</td><td>5</td><td>179840</td><td>1</td><td>1</td><td>1</td><td>N</td><td>29871</td><td>Cytochrome c oxidase subunit 3 OS=Rattus norvegicus OX=10116 GN=Mt-co3 PE=3 SV=1</td></tr>
<tr><td>214</td><td>255</td><td>tr|I6V4L9|</td><td>53.03</td><td>5</td><td>5</td><td>179840</td><td>1</td><td>1</td><td>1</td><td>N</td><td>29844</td><td>Cytochrome c oxidase subunit 3 OS=Rattus norvegicus OX=10116 GN=COX3 PE=3 SV=1</td></tr>
<tr><td>214</td><td>256</td><td>tr|Q8M7G</td><td>53.03</td><td>5</td><td>5</td><td>179840</td><td>1</td><td>1</td><td>1</td><td>N</td><td>29861</td><td>Cytochrome c oxidase subunit 3 OS=Rattus norvegicus OX=10116 PE=2 SV=1</td></tr>
<tr><td>214</td><td>257</td><td>tr|A0A096</td><td>53.03</td><td>5</td><td>5</td><td>179840</td><td>1</td><td>1</td><td>1</td><td>N</td><td>29901</td><td>Cytochrome c oxidase subunit 3 OS=Rattus norvegicus OX=10116 GN=COX3 PE=3 SV=1</td></tr>
<tr><td>214</td><td>258</td><td>P05505|CC</td><td>53.03</td><td>5</td><td>5</td><td>179840</td><td>1</td><td>1</td><td>1</td><td>N</td><td>29871</td><td>Cytochrome c oxidase subunit 3 OS=Rattus norvegicus OX=10116 GN=Mtco3 PE=1 SV=5</td></tr>
<tr><td>214</td><td>259</td><td>tr|Q8SEZ2|</td><td>53.03</td><td>5</td><td>5</td><td>179840</td><td>1</td><td>1</td><td>1</td><td>N</td><td>29870</td><td>Cytochrome c oxidase subunit 3 (Fragment) OS=Rattus norvegicus OX=10116 GN=COIII PE=3 SV=1</td></tr>
<tr><td>215</td><td>274</td><td>P11951|CX</td><td>52.32</td><td>12</td><td>12</td><td>266410</td><td>1</td><td>1</td><td>1</td><td>N</td><td>8455</td><td>Cytochrome c oxidase subunit 6C-2 OS=Rattus norvegicus OX=10116 GN=Cox6c2 PE=1 SV=3</td></tr>
<tr><td>216</td><td>4486</td><td>P29419|AT</td><td>48.86</td><td>15</td><td>15</td><td>0</td><td>1</td><td>1</td><td>1</td><td>N</td><td>8255</td><td>ATP synthase subunit e, mitochondrial OS=Rattus norvegicus OX=10116 GN=Atp5me PE=1 SV=3</td></tr>
<tr><td>31</td><td>1344</td><td>D3ZG52|D</td><td>47.18</td><td>3</td><td>3</td><td>3065600</td><td>5</td><td>4</td><td>6</td><td>Y</td><td>119588</td><td>DNA replication ATP-dependent helicase/nuclease DNA2 OS=Rattus norvegicus OX=10116 GN=Dna2 PE=3 SV=1</td></tr>
<tr><td>31</td><td>1345</td><td>tr|A0A0H2</td><td>47.18</td><td>2</td><td>2</td><td>3065600</td><td>5</td><td>4</td><td>6</td><td>Y</td><td>136526</td><td>Graves disease carrier protein OS=Rattus norvegicus OX=10116 GN=Slc25a16 PE=1 SV=1</td></tr>
<tr><td>41</td><td>408</td><td>P33124|AC</td><td>45.99</td><td>4</td><td>4</td><td>4272700</td><td>3</td><td>2</td><td>4</td><td>N</td><td>78180</td><td>Long-chain-fatty-acid--CoA ligase 6 OS=Rattus norvegicus OX=10116 GN=Acsl6 PE=1 SV=1</td></tr>
<tr><td>122</td><td>1035</td><td>Q9Z2L0|VD</td><td>45.7</td><td>4</td><td>4</td><td>0</td><td>1</td><td>1</td><td>1</td><td>N</td><td>30756</td><td>Voltage-dependent anion-selective channel protein 1 OS=Rattus norvegicus OX=10116 GN=Vdac1 PE=1 SV=4</td></tr>
<tr><td>121</td><td>134</td><td>tr|A0A0G2</td><td>45.68</td><td>1</td><td>1</td><td>237960</td><td>1</td><td>1</td><td>1</td><td>N</td><td>68759</td><td>Serum albumin OS=Rattus norvegicus OX=10116 GN=Alb PE=1 SV=1</td></tr>
<tr><td>38</td><td>459</td><td>tr|F1LPI6|</td><td>45.17</td><td>2</td><td>2</td><td>1302900</td><td>4</td><td>3</td><td>4</td><td>N</td><td>200075</td><td>Peripheral-type benzodiazepine receptor-associated protein 1 OS=Rattus norvegicus OX=10116 GN=Tspoap1 PE=4 SV=1</td></tr>
<tr><td>217</td><td>296</td><td>P19527|NF</td><td>40.5</td><td>3</td><td>3</td><td>289740</td><td>1</td><td>1</td><td>1</td><td>Y</td><td>61335</td><td>Neurofilament light polypeptide OS=Rattus norvegicus OX=10116 GN=Nefl PE=1 SV=3</td></tr>
<tr><td>218</td><td>4487</td><td>tr|Q5UAJ5</td><td>36.98</td><td>12</td><td>12</td><td>802170</td><td>1</td><td>1</td><td>1</td><td>N</td><td>7642</td><td>ATP synthase protein 8 OS=Rattus norvegicus OX=10116 GN=ATP8 PE=3 SV=1</td></tr>
<tr><td>218</td><td>4491</td><td>P11608|AT</td><td>36.98</td><td>12</td><td>12</td><td>802170</td><td>1</td><td>1</td><td>1</td><td>N</td><td>7630</td><td>ATP synthase protein 8 OS=Rattus norvegicus OX=10116 GN=Mt-atp8 PE=1 SV=2</td></tr>
<tr><td>218</td><td>4492</td><td>tr|Q8SEZ4|</td><td>36.98</td><td>12</td><td>12</td><td>802170</td><td>1</td><td>1</td><td>1</td><td>N</td><td>7632</td><td>ATP synthase protein 8 OS=Rattus norvegicus OX=10116 GN=ATPase8 PE=3 SV=1</td></tr>
<tr><td>218</td><td>4493</td><td>tr|Q8HIC8|</td><td>36.98</td><td>12</td><td>12</td><td>802170</td><td>1</td><td>1</td><td>1</td><td>N</td><td>7630</td><td>ATP synthase protein 8 OS=Rattus norvegicus OX=10116 GN=Mt-atp8 PE=3 SV=1</td></tr>
<tr><td>222</td><td>586</td><td>tr|C7EMI4</td><td>34.09</td><td>3</td><td>3</td><td>0</td><td>1</td><td>1</td><td>1</td><td>N</td><td>20146</td><td>Cytochrome b (Fragment) OS=Rattus norvegicus OX=10116 PE=4 SV=1</td></tr>
<tr><td>222</td><td>587</td><td>tr|U3M4V1</td><td>34.09</td><td>3</td><td>3</td><td>0</td><td>1</td><td>1</td><td>1</td><td>N</td><td>20795</td><td>Cytochrome b (Fragment) OS=Rattus norvegicus OX=10116 GN=cyt b PE=4 SV=1</td></tr>
<tr><td>222</td><td>588</td><td>tr|A4UHZ3</td><td>34.09</td><td>3</td><td>3</td><td>0</td><td>1</td><td>1</td><td>1</td><td>N</td><td>25311</td><td>Cytochrome b (Fragment) OS=Rattus norvegicus OX=10116 GN=cytb PE=4 SV=1</td></tr>
<tr><td>222</td><td>589</td><td>tr|A4UHZ2</td><td>34.09</td><td>3</td><td>3</td><td>0</td><td>1</td><td>1</td><td>1</td><td>N</td><td>25312</td><td>Cytochrome b (Fragment) OS=Rattus norvegicus OX=10116 GN=cytb PE=4 SV=1</td></tr>
<tr><td>222</td><td>590</td><td>tr|B3Y999|</td><td>34.09</td><td>2</td><td>2</td><td>0</td><td>1</td><td>1</td><td>1</td><td>N</td><td>28378</td><td>Cytochrome b (Fragment) OS=Rattus norvegicus OX=10116 GN=cytb PE=4 SV=1</td></tr>
<tr><td>222</td><td>594</td><td>tr|M9TEK6</td><td>34.09</td><td>2</td><td>2</td><td>0</td><td>1</td><td>1</td><td>1</td><td>N</td><td>35982</td><td>Cytochrome b (Fragment) OS=Rattus norvegicus OX=10116 GN=cytb PE=3 SV=1</td></tr>
<tr><td>222</td><td>596</td><td>tr|E0A0C7|</td><td>34.09</td><td>2</td><td>2</td><td>0</td><td>1</td><td>1</td><td>1</td><td>N</td><td>41009</td><td>Cytochrome b (Fragment) OS=Rattus norvegicus OX=10116 GN=cytb PE=3 SV=1</td></tr>
<tr><td>222</td><td>600</td><td>tr|A0A411</td><td>34.09</td><td>2</td><td>2</td><td>0</td><td>1</td><td>1</td><td>1</td><td>N</td><td>41017</td><td>Cytochrome b (Fragment) OS=Rattus norvegicus OX=10116 GN=Cytb PE=3 SV=1</td></tr>
<tr><td>222</td><td>601</td><td>tr|E0A016|</td><td>34.09</td><td>2</td><td>2</td><td>0</td><td>1</td><td>1</td><td>1</td><td>N</td><td>41634</td><td>Cytochrome b (Fragment) OS=Rattus norvegicus OX=10116 GN=cytb PE=3 SV=1</td></tr>
<tr><td>222</td><td>603</td><td>tr|A0A411</td><td>34.09</td><td>2</td><td>2</td><td>0</td><td>1</td><td>1</td><td>1</td><td>N</td><td>42238</td><td>Cytochrome b (Fragment) OS=Rattus norvegicus OX=10116 GN=Cytb PE=3 SV=1</td></tr>
<tr><td>222</td><td>604</td><td>tr|E0A075|</td><td>34.09</td><td>2</td><td>2</td><td>0</td><td>1</td><td>1</td><td>1</td><td>N</td><td>42410</td><td>Cytochrome b (Fragment) OS=Rattus norvegicus OX=10116 GN=cytb PE=3 SV=1</td></tr>
<tr><td>222</td><td>605</td><td>tr|L0L4L8|</td><td>34.09</td><td>2</td><td>2</td><td>0</td><td>1</td><td>1</td><td>1</td><td>N</td><td>42470</td><td>Cytochrome b (Fragment) OS=Rattus norvegicus OX=10116 GN=cytb PE=3 SV=1</td></tr>
<tr><td>222</td><td>606</td><td>tr|A0A140</td><td>34.09</td><td>2</td><td>2</td><td>0</td><td>1</td><td>1</td><td>1</td><td>N</td><td>42569</td><td>Cytochrome b (Fragment) OS=Rattus norvegicus OX=10116 PE=3 SV=1</td></tr>
<tr><td>222</td><td>607</td><td>tr|A0A140</td><td>34.09</td><td>2</td><td>2</td><td>0</td><td>1</td><td>1</td><td>1</td><td>N</td><td>42574</td><td>Cytochrome b (Fragment) OS=Rattus norvegicus OX=10116 PE=3 SV=1</td></tr>
<tr><td>222</td><td>608</td><td>tr|A0A140</td><td>34.09</td><td>2</td><td>2</td><td>0</td><td>1</td><td>1</td><td>1</td><td>N</td><td>42588</td><td>Cytochrome b (Fragment) OS=Rattus norvegicus OX=10116 PE=3 SV=1</td></tr>
<tr><td>222</td><td>610</td><td>tr|A0A385</td><td>34.09</td><td>2</td><td>2</td><td>0</td><td>1</td><td>1</td><td>1</td><td>N</td><td>42780</td><td>Cytochrome b (Fragment) OS=Rattus norvegicus OX=10116 PE=3 SV=1</td></tr>
<tr><td>222</td><td>611</td><td>tr|A0A385</td><td>34.09</td><td>2</td><td>2</td><td>0</td><td>1</td><td>1</td><td>1</td><td>N</td><td>42766</td><td>Cytochrome b (Fragment) OS=Rattus norvegicus OX=10116 PE=3 SV=1</td></tr>
<tr><td>222</td><td>612</td><td>tr|A0A3Q8</td><td>34.09</td><td>2</td><td>2</td><td>0</td><td>1</td><td>1</td><td>1</td><td>N</td><td>42712</td><td>Cytochrome b (Fragment) OS=Rattus norvegicus OX=10116 GN=Cytb PE=3 SV=1</td></tr>
</table>

| | | | | | | | | | | | | |
|---|---|---|---|---|---|---|---|---|---|---|---|---|
| **222** | 613 | tr\|A0A3Q8 | 34.09 | 2 | 2 | 0 | 1 | 1 | 1 | N | 42622 | Cytochrome b (Fragment) OS=Rattus norvegicus OX=10116 GN=Cytb PE=3 SV=1 |
| **222** | 614 | tr\|A0A3S6 | 34.09 | 2 | 2 | 0 | 1 | 1 | 1 | N | 42702 | Cytochrome b (Fragment) OS=Rattus norvegicus OX=10116 GN=Cytb PE=3 SV=1 |
| **222** | 615 | tr\|A0A385 | 34.09 | 2 | 2 | 0 | 1 | 1 | 1 | N | 42824 | Cytochrome b (Fragment) OS=Rattus norvegicus OX=10116 PE=3 SV=1 |
| **222** | 616 | tr\|A0A3Q8 | 34.09 | 2 | 2 | 0 | 1 | 1 | 1 | N | 42723 | Cytochrome b (Fragment) OS=Rattus norvegicus OX=10116 GN=Cytb PE=3 SV=1 |
| **222** | 617 | tr\|A0A3S6 | 34.09 | 2 | 2 | 0 | 1 | 1 | 1 | N | 42726 | Cytochrome b (Fragment) OS=Rattus norvegicus OX=10116 GN=Cytb PE=3 SV=1 |
| **222** | 618 | tr\|A0A385 | 34.09 | 2 | 2 | 0 | 1 | 1 | 1 | N | 42751 | Cytochrome b (Fragment) OS=Rattus norvegicus OX=10116 PE=3 SV=1 |
| **222** | 619 | tr\|A0A3S6 | 34.09 | 2 | 2 | 0 | 1 | 1 | 1 | N | 42698 | Cytochrome b (Fragment) OS=Rattus norvegicus OX=10116 GN=Cytb PE=3 SV=1 |
| **222** | 621 | tr\|A0A097 | 34.09 | 2 | 2 | 0 | 1 | 1 | 1 | N | 42865 | Cytochrome b OS=Rattus norvegicus OX=10116 GN=CYTB PE=3 SV=1 |
| **222** | 622 | tr\|A0A220 | 34.09 | 2 | 2 | 0 | 1 | 1 | 1 | N | 43018 | Cytochrome b OS=Rattus norvegicus OX=10116 PE=3 SV=1 |
| **222** | 623 | tr\|D6NSP2 | 34.09 | 2 | 2 | 0 | 1 | 1 | 1 | N | 42945 | Cytochrome b (Fragment) OS=Rattus norvegicus OX=10116 GN=cytb PE=3 SV=1 |
| **222** | 624 | tr\|D6NSS6 | 34.09 | 2 | 2 | 0 | 1 | 1 | 1 | N | 42993 | Cytochrome b (Fragment) OS=Rattus norvegicus OX=10116 GN=cytb PE=3 SV=1 |
| **222** | 625 | tr\|Q06Q99 | 34.09 | 2 | 2 | 0 | 1 | 1 | 1 | N | 42997 | Cytochrome b OS=Rattus norvegicus OX=10116 GN=CYB PE=3 SV=1 |
| **222** | 626 | tr\|L0N490\| | 34.09 | 2 | 2 | 0 | 1 | 1 | 1 | N | 43056 | Cytochrome b (Fragment) OS=Rattus norvegicus OX=10116 GN=cytb PE=3 SV=1 |
| **222** | 627 | tr\|D6NSQ3 | 34.09 | 2 | 2 | 0 | 1 | 1 | 1 | N | 43012 | Cytochrome b (Fragment) OS=Rattus norvegicus OX=10116 GN=cytb PE=3 SV=1 |
| **222** | 628 | tr\|D6NSR3 | 34.09 | 2 | 2 | 0 | 1 | 1 | 1 | N | 42986 | Cytochrome b (Fragment) OS=Rattus norvegicus OX=10116 GN=cytb PE=3 SV=1 |
| **222** | 629 | tr\|D6NSP4 | 34.09 | 2 | 2 | 0 | 1 | 1 | 1 | N | 43016 | Cytochrome b (Fragment) OS=Rattus norvegicus OX=10116 GN=cytb PE=3 SV=1 |
| **222** | 630 | tr\|D6NSQ8 | 34.09 | 2 | 2 | 0 | 1 | 1 | 1 | N | 43016 | Cytochrome b (Fragment) OS=Rattus norvegicus OX=10116 GN=cytb PE=3 SV=1 |
| **222** | 631 | tr\|D6NSR8 | 34.09 | 2 | 2 | 0 | 1 | 1 | 1 | N | 42998 | Cytochrome b (Fragment) OS=Rattus norvegicus OX=10116 GN=cytb PE=3 SV=1 |
| **222** | 632 | tr\|F8QU37 | 34.09 | 2 | 2 | 0 | 1 | 1 | 1 | N | 42992 | Cytochrome b (Fragment) OS=Rattus norvegicus OX=10116 GN=cytb PE=3 SV=1 |
| **222** | 633 | tr\|F2Q6S6\| | 34.09 | 2 | 2 | 0 | 1 | 1 | 1 | N | 42938 | Cytochrome b (Fragment) OS=Rattus norvegicus OX=10116 GN=cytb PE=3 SV=1 |
| **222** | 634 | tr\|A0A0A1 | 34.09 | 2 | 2 | 0 | 1 | 1 | 1 | N | 42989 | Cytochrome b OS=Rattus norvegicus OX=10116 GN=CYTB PE=3 SV=1 |
| **222** | 635 | tr\|D6NSS7 | 34.09 | 2 | 2 | 0 | 1 | 1 | 1 | N | 42948 | Cytochrome b (Fragment) OS=Rattus norvegicus OX=10116 GN=cytb PE=3 SV=1 |
| **222** | 636 | tr\|Q8SEY9\| | 34.09 | 2 | 2 | 0 | 1 | 1 | 1 | N | 43015 | Cytochrome b OS=Rattus norvegicus OX=10116 GN=cytb PE=3 SV=1 |
| **222** | 637 | tr\|D6NSR7 | 34.09 | 2 | 2 | 0 | 1 | 1 | 1 | N | 42982 | Cytochrome b (Fragment) OS=Rattus norvegicus OX=10116 GN=cytb PE=3 SV=1 |
| **222** | 638 | tr\|A0A220 | 34.09 | 2 | 2 | 0 | 1 | 1 | 1 | N | 42968 | Cytochrome b OS=Rattus norvegicus OX=10116 PE=3 SV=1 |
| **222** | 640 | tr\|D6NSP0 | 34.09 | 2 | 2 | 0 | 1 | 1 | 1 | N | 43023 | Cytochrome b (Fragment) OS=Rattus norvegicus OX=10116 GN=cytb PE=3 SV=1 |
| **222** | 641 | tr\|D6NSQ6 | 34.09 | 2 | 2 | 0 | 1 | 1 | 1 | N | 42952 | Cytochrome b (Fragment) OS=Rattus norvegicus OX=10116 GN=cytb PE=3 SV=1 |
| **222** | 642 | tr\|L0N495\| | 34.09 | 2 | 2 | 0 | 1 | 1 | 1 | N | 42992 | Cytochrome b (Fragment) OS=Rattus norvegicus OX=10116 GN=cytb PE=3 SV=1 |
| **222** | 643 | tr\|D6NSQ0 | 34.09 | 2 | 2 | 0 | 1 | 1 | 1 | N | 42968 | Cytochrome b (Fragment) OS=Rattus norvegicus OX=10116 GN=cytb PE=3 SV=1 |
| **222** | 644 | tr\|Q5UAI7 | 34.09 | 2 | 2 | 0 | 1 | 1 | 1 | N | 42998 | Cytochrome b OS=Rattus norvegicus OX=10116 GN=CYTB PE=3 SV=1 |
| **222** | 645 | tr\|D6NSR2 | 34.09 | 2 | 2 | 0 | 1 | 1 | 1 | N | 43073 | Cytochrome b (Fragment) OS=Rattus norvegicus OX=10116 GN=cytb PE=3 SV=1 |
| **222** | 646 | tr\|A0A0S1 | 34.09 | 2 | 2 | 0 | 1 | 1 | 1 | N | 43002 | Cytochrome b OS=Rattus norvegicus OX=10116 GN=CYTB PE=3 SV=1 |
| **222** | 647 | tr\|B0M1Q | 34.09 | 2 | 2 | 0 | 1 | 1 | 1 | N | 43026 | Cytochrome b (Fragment) OS=Rattus norvegicus OX=10116 GN=cytb PE=3 SV=1 |
| **222** | 648 | tr\|D6NSR9 | 34.09 | 2 | 2 | 0 | 1 | 1 | 1 | N | 42970 | Cytochrome b (Fragment) OS=Rattus norvegicus OX=10116 GN=cytb PE=3 SV=1 |
| **222** | 649 | tr\|D6NSR1 | 34.09 | 2 | 2 | 0 | 1 | 1 | 1 | N | 43002 | Cytochrome b (Fragment) OS=Rattus norvegicus OX=10116 GN=cytb PE=3 SV=1 |
| **222** | 650 | tr\|Q8HIC4\| | 34.09 | 2 | 2 | 0 | 1 | 1 | 1 | N | 43012 | Cytochrome b OS=Rattus norvegicus OX=10116 GN=Mt-cyb PE=3 SV=1 |
| **222** | 651 | tr\|A0A220 | 34.09 | 2 | 2 | 0 | 1 | 1 | 1 | N | 42968 | Cytochrome b OS=Rattus norvegicus OX=10116 PE=3 SV=1 |
| **222** | 652 | tr\|S5RKC8\| | 34.09 | 2 | 2 | 0 | 1 | 1 | 1 | N | 42997 | Cytochrome b OS=Rattus norvegicus OX=10116 GN=CYTB PE=3 SV=1 |
| **222** | 653 | tr\|A0A0S1 | 34.09 | 2 | 2 | 0 | 1 | 1 | 1 | N | 43016 | Cytochrome b OS=Rattus norvegicus OX=10116 GN=CYTB PE=3 SV=1 |
| **222** | 654 | tr\|R9TKN1 | 34.09 | 2 | 2 | 0 | 1 | 1 | 1 | N | 43012 | Cytochrome b OS=Rattus norvegicus OX=10116 GN=CYTB PE=3 SV=1 |
| **222** | 655 | P00159\|CY | 34.09 | 2 | 2 | 0 | 1 | 1 | 1 | N | 43012 | Cytochrome b OS=Rattus norvegicus OX=10116 GN=Mt-Cyb PE=3 SV=3 |
| **222** | 656 | tr\|D6NSP8 | 34.09 | 2 | 2 | 0 | 1 | 1 | 1 | N | 43100 | Cytochrome b (Fragment) OS=Rattus norvegicus OX=10116 GN=cytb PE=3 SV=1 |
| **222** | 657 | tr\|L0N311\| | 34.09 | 2 | 2 | 0 | 1 | 1 | 1 | N | 43045 | Cytochrome b (Fragment) OS=Rattus norvegicus OX=10116 GN=cytb PE=3 SV=1 |
| **222** | 658 | tr\|A0A220 | 34.09 | 2 | 2 | 0 | 1 | 1 | 1 | N | 43010 | Cytochrome b OS=Rattus norvegicus OX=10116 PE=3 SV=1 |
| **222** | 659 | tr\|A0A220 | 34.09 | 2 | 2 | 0 | 1 | 1 | 1 | N | 43028 | Cytochrome b OS=Rattus norvegicus OX=10116 GN=cyt-b PE=3 SV=1 |
| **222** | 660 | tr\|A0A220 | 34.09 | 2 | 2 | 0 | 1 | 1 | 1 | N | 42970 | Cytochrome b OS=Rattus norvegicus OX=10116 PE=3 SV=1 |
| **222** | 661 | tr\|H6RXT5 | 34.09 | 2 | 2 | 0 | 1 | 1 | 1 | N | 43044 | Cytochrome b OS=Rattus norvegicus OX=10116 GN=cytb PE=3 SV=1 |
| **222** | 662 | tr\|H2KXA0 | 34.09 | 2 | 2 | 0 | 1 | 1 | 1 | N | 42978 | Cytochrome b (Fragment) OS=Rattus norvegicus OX=10116 GN=cytb PE=3 SV=1 |
| **222** | 663 | tr\|L0N1V8\| | 34.09 | 2 | 2 | 0 | 1 | 1 | 1 | N | 43058 | Cytochrome b (Fragment) OS=Rattus norvegicus OX=10116 GN=cytb PE=3 SV=1 |
| **222** | 664 | tr\|A0A096 | 34.09 | 2 | 2 | 0 | 1 | 1 | 1 | N | 43042 | Cytochrome b (Fragment) OS=Rattus norvegicus OX=10116 PE=3 SV=1 |
| **222** | 665 | tr\|D6NSP7 | 34.09 | 2 | 2 | 0 | 1 | 1 | 1 | N | 43046 | Cytochrome b (Fragment) OS=Rattus norvegicus OX=10116 GN=cytb PE=3 SV=1 |
| **222** | 666 | tr\|A0A220 | 34.09 | 2 | 2 | 0 | 1 | 1 | 1 | N | 43024 | Cytochrome b OS=Rattus norvegicus OX=10116 PE=3 SV=1 |
| **222** | 667 | tr\|D6NSP5 | 34.09 | 2 | 2 | 0 | 1 | 1 | 1 | N | 43012 | Cytochrome b (Fragment) OS=Rattus norvegicus OX=10116 GN=cytb PE=3 SV=1 |
| **222** | 558 | tr\|D6NSP6 | 34.09 | 2 | 2 | 0 | 1 | 1 | 1 | N | 43050 | Cytochrome b (Fragment) OS=Rattus norvegicus OX=10116 GN=cytb PE=3 SV=1 |
| **222** | 552 | tr\|Q9G898 | 34.09 | 2 | 2 | 0 | 1 | 1 | 1 | N | 42830 | Cytochrome b OS=Rattus norvegicus OX=10116 PE=2 SV=1 |
| **152** | 1110 | Q06437\|O | 33.4 | 3 | 3 | 330540 | 1 | 1 | 1 | N | 43393 | Pyruvate dehydrogenase E1 component subunit alpha, testis-specific form, mitochondrial OS=Rattus norvegicus OX=10116 GN=Pdha2 PE=1 SV=1 |
| **221** | 12926 | tr\|D4A9V7 | 33.37 | 1 | 1 | 213730 | 1 | 1 | 1 | N | 65753 | Methionine--tRNA ligase, mitochondrial-like OS=Rattus norvegicus OX=10116 GN=LOC100911305 PE=1 SV=1 |
| **219** | 282 | P19804\|N | 33.26 | 8 | 8 | 234580 | 1 | 1 | 1 | N | 17283 | Nucleoside diphosphate kinase B OS=Rattus norvegicus OX=10116 GN=Nme2 PE=1 SV=1 |
| **219** | 283 | Q05982\|N | 33.26 | 8 | 8 | 234580 | 1 | 1 | 1 | N | 17193 | Nucleoside diphosphate kinase A OS=Rattus norvegicus OX=10116 GN=Nme1 PE=1 SV=1 |
| **78** | 4527 | Q6RI88\|S4 | 33.15 | 1 | 1 | 21633000 | 2 | 1 | 2 | N | 123944 | Electrogenic sodium bicarbonate cotransporter 4 OS=Rattus norvegicus OX=10116 GN=Slc4a5 PE=1 SV=1 |

| | | | | | | | | | | | | |
|---|---|---|---|---|---|---|---|---|---|---|---|---|
| **81** | 478 | Q64578\|AT | 32.86 | 1 | 1 | 3992200 | 1 | 1 | 2 | Y | 109409 | Sarcoplasmic/endoplasmic reticulum calcium ATPase 1 OS=Rattus norvegicus OX=10116 GN=Atp2a1 PE=1 SV=1 |
| **83** | 410 | tr\|F1LRY0\| | 32.41 | 7 | 7 | 587880 | 2 | 1 | 2 | Y | 26077 | Max dimerization protein 1 OS=Rattus norvegicus OX=10116 GN=Mxd1 PE=4 SV=1 |
| **67** | 121 | P08461\|OD | 32.13 | 3 | 3 | 333050 | 2 | 1 | 2 | Y | 67166 | Dihydrolipoyllysine-residue acetyltransferase component of pyruvate dehydrogenase complex, mitochondrial OS=Rattus norvegicus OX=10116 GN=Dlat PE=1 SV=3 |
| **55** | 834 | tr\|A0A0G2 | 31.22 | 0 | 0 | 3130200 | 2 | 2 | 2 | N | 487857 | Vacuolar protein sorting 13 homolog D OS=Rattus norvegicus OX=10116 GN=Vps13d PE=1 SV=1 |
| **55** | 835 | tr\|D3ZKC6 | 31.22 | 0 | 0 | 3130200 | 2 | 2 | 2 | N | 488962 | Vacuolar protein sorting 13 homolog D OS=Rattus norvegicus OX=10116 GN=Vps13d PE=1 SV=1 |
| **71** | 450 | tr\|D4A4K4 | 31.11 | 0 | 0 | 1163800 | 2 | 2 | 2 | N | 418626 | Vacuolar protein sorting 13 homolog C OS=Rattus norvegicus OX=10116 GN=Vps13c PE=1 SV=2 |
| **220** | 5083 | Q6AY30\|SC | 30.98 | 7 | 7 | 7817.8 | 1 | 1 | 1 | N | 47088 | Saccharopine dehydrogenase-like oxidoreductase OS=Rattus norvegicus OX=10116 GN=Sccpdh PE=1 SV=1 |
| **157** | 8233 | tr\|F1LN88\| | 30.96 | 2 | 2 | 504540 | 1 | 1 | 1 | N | 56517 | Aldehyde dehydrogenase, mitochondrial OS=Rattus norvegicus OX=10116 GN=Aldh2 PE=1 SV=2 |
| **157** | 8234 | P11884\|AL | 30.96 | 2 | 2 | 504540 | 1 | 1 | 1 | N | 56488 | Aldehyde dehydrogenase, mitochondrial OS=Rattus norvegicus OX=10116 GN=Aldh2 PE=1 SV=1 |
| **84** | 12529 | Q3B8Q1\|D | 30.81 | 2 | 2 | 0 | 2 | 2 | 2 | N | 85966 | Nucleolar RNA helicase 2 OS=Rattus norvegicus OX=10116 GN=Ddx21 PE=2 SV=1 |
| **223** | 23454 | P18886\|CP | 30.22 | 1 | 1 | 0 | 1 | 1 | 1 | N | 74110 | Carnitine O-palmitoyltransferase 2, mitochondrial OS=Rattus norvegicus OX=10116 GN=Cpt2 PE=1 SV=1 |
| **223** | 23455 | tr\|G3V7N5 | 30.22 | 1 | 1 | 0 | 1 | 1 | 1 | N | 74078 | Carnitine O-palmitoyltransferase 2, mitochondrial OS=Rattus norvegicus OX=10116 GN=Cpt2 PE=1 SV=1 |
| **54** | 1365 | tr\|D3ZXY2\| | 29.91 | 3 | 3 | 2458000 | 3 | 3 | 3 | Y | 128251 | PDZ domain containing 8 (Predicted) OS=Rattus norvegicus OX=10116 GN=Pdzd8 PE=1 SV=1 |
| **224** | 23457 | Q5XIA8\|GH | 28.77 | 4 | 4 | 0 | 1 | 1 | 1 | N | 37178 | Growth hormone-inducible transmembrane protein OS=Rattus norvegicus OX=10116 GN=Ghitm PE=2 SV=1 |
| **111** | 555 | P15205\|MA | 28.окт | 0 | 0 | 115790 | 1 | 1 | 1 | N | 269640 | Microtubule-associated protein 1B OS=Rattus norvegicus OX=10116 GN=Map1b PE=1 SV=3 |
| **86** | 1281 | tr\|F1M9H4 | 28.сен | 1 | 1 | 972630 | 2 | 2 | 2 | N | 225602 | HIVEP zinc finger 1 OS=Rattus norvegicus OX=10116 GN=Hivep1 PE=1 SV=2 |
| **159** | 4619 | P35559\|ID | 27.7 | 1 | 1 | 0 | 1 | 1 | 1 | N | 117710 | Insulin-degrading enzyme OS=Rattus norvegicus OX=10116 GN=Ide PE=1 SV=1 |
| **66** | 741 | tr\|A0A0G2 | 26.19 | 2 | 2 | 1006500 | 2 | 2 | 2 | Y | 205333 | Golgi brefeldin A-resistant guanine nucleotide exchange factor 1 OS=Rattus norvegicus OX=10116 GN=Gbf1 PE=1 SV=1 |
| **66** | 742 | tr\|F1M8X9 | 26.19 | 2 | 2 | 1006500 | 2 | 2 | 2 | Y | 206522 | Golgi brefeldin A-resistant guanine nucleotide exchange factor 1 OS=Rattus norvegicus OX=10116 GN=Gbf1 PE=1 SV=2 |
| **77** | 5209 | tr\|Q5U2M | 26.17 | 1 | 1 | 0 | 2 | 1 | 2 | N | 105443 | DNA ligase OS=Rattus norvegicus OX=10116 GN=Lig3 PE=1 SV=1 |
| **77** | 1338 | tr\|A0A096 | 26.17 | 1 | 1 | 0 | 2 | 1 | 2 | N | 112017 | DNA ligase OS=Rattus norvegicus OX=10116 GN=Lig3 PE=1 SV=1 |
| **61** | 1506 | tr\|D4A1D3 | 25.15 | 0 | 0 | 672090 | 2 | 2 | 2 | Y | 521487 | Sacsin molecular chaperone OS=Rattus norvegicus OX=10116 GN=Sacs PE=1 SV=2 |
| **163** | 141 | Q924S5\|LC | 24.69 | 1 | 1 | 41430 | 1 | 1 | 1 | N | 105792 | Lon protease homolog, mitochondrial OS=Rattus norvegicus OX=10116 GN=Lonp1 PE=2 SV=1 |
| **69** | 769 | Q64680\|CP | 24.5 | 3 | 3 | 3700700 | 2 | 2 | 2 | N | 56684 | Cytochrome P450 2D4 OS=Rattus norvegicus OX=10116 GN=Cyp2d4 PE=2 SV=1 |
| **231** | 23460 | tr\|Q3B8P7 | 24.29 | 3 | 3 | 2003500 | 1 | 1 | 1 | Y | 27522 | RCG58555, isoform CRA_a OS=Rattus norvegicus OX=10116 GN=LOC683897 PE=2 SV=1 |
| **231** | 23461 | tr\|M0R7R2 | 24.29 | 3 | 3 | 2003500 | 1 | 1 | 1 | Y | 29265 | Similar to Protein C6orf203 OS=Rattus norvegicus OX=10116 GN=LOC683897 PE=1 SV=1 |
| **100** | 372 | tr\|F1LNJ1\| | 24.26 | 1 | 1 | 262740 | 1 | 1 | 1 | Y | 284829 | Leucine-rich repeat kinase 2 OS=Rattus norvegicus OX=10116 GN=Lrrk2 PE=1 SV=3 |
| **229** | 12586 | tr\|A0A0G2 | 23.91 | 10 | 10 | 0 | 1 | 1 | 1 | Y | 9388 | 10 kDa heat shock protein, mitochondrial OS=Rattus norvegicus OX=10116 GN=Hspe1 PE=1 SV=1 |
| **229** | 12587 | tr\|P97601\| | 23.91 | 9 | 9 | 0 | 1 | 1 | 1 | Y | 10876 | Chaperonin 10 OS=Rattus norvegicus OX=10116 GN=Hspe1 PE=3 SV=1 |
| **229** | 12588 | P26772\|CH | 23.91 | 9 | 9 | 0 | 1 | 1 | 1 | Y | 10902 | 10 kDa heat shock protein, mitochondrial OS=Rattus norvegicus OX=10116 GN=Hspe1 PE=1 SV=3 |
| **124** | 162 | tr\|D3Z900\| | 23.47 | 3 | 3 | 0 | 1 | 1 | 1 | N | 38239 | Mitochondrial amidoxime reducing component 2 OS=Rattus norvegicus OX=10116 GN=Marc2 PE=1 SV=2 |
| **124** | 163 | O88994\|M | 23.47 | 3 | 3 | 0 | 1 | 1 | 1 | N | 38249 | Mitochondrial amidoxime reducing component 2 OS=Rattus norvegicus OX=10116 GN=Marc2 PE=2 SV=1 |
| **124** | 164 | tr\|M0R6N2 | 23.47 | 3 | 3 | 0 | 1 | 1 | 1 | N | 38175 | MOSC domain-containing protein 2, mitochondrial-like OS=Rattus norvegicus OX=10116 GN=LOC100910481 PE=1 SV=1 |
| **234** | 4650 | Q5I0L3\|SY | 23.46 | 1 | 1 | 304950 | 1 | 1 | 1 | N | 52628 | Tyrosine--tRNA ligase, mitochondrial OS=Rattus norvegicus OX=10116 GN=Yars2 PE=2 SV=1 |
| **232** | 23462 | Q5XIG0\|NU | 23.35 | 3 | 3 | 554110 | 1 | 1 | 1 | Y | 38562 | ADP-ribose pyrophosphatase, mitochondrial OS=Rattus norvegicus OX=10116 GN=Nudt9 PE=2 SV=1 |
| **134** | 1188 | P04694\|AT | 23.19 | 2 | 2 | 614700 | 1 | 1 | 1 | Y | 50635 | Tyrosine aminotransferase OS=Rattus norvegicus OX=10116 GN=Tat PE=1 SV=1 |
| **164** | 233 | Q5U2U0\|C | 23.16 | 1 | 1 | 384440 | 1 | 1 | 1 | N | 69208 | ATP-dependent Clp protease ATP-binding subunit clpX-like, mitochondrial OS=Rattus norvegicus OX=10116 GN=Clpx PE=2 SV=1 |
| **237** | 348 | tr\|B2GUZ6 | 23.июн | 2 | 2 | 231090 | 1 | 1 | 1 | Y | 43606 | Reticulon 4-interacting protein 1 OS=Rattus norvegicus OX=10116 GN=Rtn4ip1 PE=1 SV=1 |
| **141** | 264 | B1WC61\|A | 22.86 | 2 | 2 | 29812 | 1 | 1 | 1 | Y | 68843 | Complex I assembly factor ACAD9, mitochondrial OS=Rattus norvegicus OX=10116 GN=Acad9 PE=1 SV=1 |
| **236** | 23466 | Q5M7W1\| | 22.46 | 2 | 2 | 0 | 1 | 1 | 1 | N | 44018 | Thioredoxin-interacting protein OS=Rattus norvegicus OX=10116 GN=Txnip PE=2 SV=1 |
| **105** | 394 | tr\|Q6TXF6 | 22.25 | 1 | 1 | 176760 | 1 | 1 | 1 | N | 95435 | LRRGT00043 OS=Rattus norvegicus OX=10116 GN=Decr2 PE=1 SV=1 |
| **165** | 421 | P11497\|AC | 21.67 | 1 | 1 | 0 | 1 | 1 | 1 | Y | 265191 | Acetyl-CoA carboxylase 1 OS=Rattus norvegicus OX=10116 GN=Acaca PE=1 SV=1 |
| **241** | 749 | tr\|D3ZJX5\| | 21.39 | 2 | 2 | 0 | 1 | 1 | 1 | N | 39855 | Mitochondrial import inner membrane translocase subunit TIM50 OS=Rattus norvegicus OX=10116 GN=Timm50 PE=1 SV=2 |
| **243** | 4992 | P80431\|CC | 20.94 | 9 | 9 | 247390 | 1 | 1 | 1 | N | 8995 | Cytochrome c oxidase subunit 7B, mitochondrial OS=Rattus norvegicus OX=10116 GN=Cox7b PE=1 SV=3 |
| **242** | 4519 | tr\|D3ZA45 | 20.91 | 2 | 2 | 559110 | 1 | 1 | 1 | N | 52491 | Autophagy-related protein 13 OS=Rattus norvegicus OX=10116 GN=Atg13 PE=1 SV=2 |
| **244** | 1077 | Q9QYV8\|D | 20.72 | 1 | 1 | 145750 | 1 | 1 | 1 | N | 136855 | DNA polymerase subunit gamma-1 OS=Rattus norvegicus OX=10116 GN=Polg PE=2 SV=1 |
| **128** | 1019 | Q2M2R8\|P | 20.55 | 1 | 1 | 187350 | 1 | 1 | 1 | N | 71022 | Peroxisomal targeting signal 1 receptor OS=Rattus norvegicus OX=10116 GN=Pex5 PE=1 SV=2 |
| **245** | 498 | tr\|Q5U2X8 | 20.45 | 2 | 2 | 1233700 | 1 | 1 | 1 | N | 50444 | Acyl-CoA thioesterase 9 OS=Rattus norvegicus OX=10116 GN=Acot9 PE=1 SV=1 |
| **172** | 12880 | P16310\|GH | 20.14 | 2 | 2 | 679350 | 1 | 1 | 1 | N | 71237 | Growth hormone receptor OS=Rattus norvegicus OX=10116 GN=Ghr PE=1 SV=1 |
| **101** | 12752 | tr\|D3ZAE6 | 20.ноя | 1 | 1 | 671140 | 1 | 1 | 1 | Y | 72321 | RCG49849 OS=Rattus norvegicus OX=10116 GN=Vasn PE=1 SV=1 |
