## Supplementary material for "Permeability transition pore-related changes in the proteome and channel activity of ATP synthase dimers and monomers": RLM KCl-BM

| Protein Group | Protein ID | Accession | -10lgP | Coverage (%) | Coverage (%) Sample 1 | IBAQ | #Peptides | #Unique | #Spec Sample 1 | PTM | Avg. Mass | Description |
|---|---|---|---|---|---|---|---|---|---|---|---|---|
| **1** | 3 | P07756\|CP | 429.18 | 40 | 40 | 85768000 | 54 | 54 | 66 | Y | 164579 | Carbamoyl-phosphate synthase [ammonia], mitochondrial OS=Rattus norvegicus OX=10116 GN=Cps1 PE=1 SV=1 |
| **8** | 5 | Q64428\|EC | 407.29 | 46 | 46 | 64588000 | 31 | 30 | 41 | Y | 82665 | Trifunctional enzyme subunit alpha, mitochondrial OS=Rattus norvegicus OX=10116 GN=Hadha PE=1 SV=2 |
| **5** | 1 | P32551\|QC | 386.85 | 79 | 79 | 147150000 | 39 | 39 | 52 | Y | 48396 | Cytochrome b-c1 complex subunit 2, mitochondrial OS=Rattus norvegicus OX=10116 GN=Uqcrc2 PE=1 SV=2 |
| **3** | 2 | P10860\|DH | 382.62 | 66 | 66 | 199000000 | 47 | 47 | 59 | Y | 61416 | Glutamate dehydrogenase 1, mitochondrial OS=Rattus norvegicus OX=10116 GN=Glud1 PE=1 SV=2 |
| **6** | 15 | Q68FY0\|QC | 345.05 | 58 | 58 | 72040000 | 33 | 33 | 51 | Y | 52849 | Cytochrome b-c1 complex subunit 1, mitochondrial OS=Rattus norvegicus OX=10116 GN=Uqcrc1 PE=1 SV=1 |
| **7** | 4 | P00507\|AA | 343.62 | 71 | 71 | 144720000 | 31 | 31 | 45 | Y | 47314 | Aspartate aminotransferase, mitochondrial OS=Rattus norvegicus OX=10116 GN=Got2 PE=1 SV=2 |
| **9** | 10 | P22791\|HN | 322.6 | 50 | 50 | 101990000 | 35 | 34 | 41 | Y | 56912 | Hydroxymethylglutaryl-CoA synthase, mitochondrial OS=Rattus norvegicus OX=10116 GN=Hmgcs2 PE=1 SV=1 |
| **9** | 11 | tr\|Q68G44 | 322.6 | 50 | 50 | 101990000 | 35 | 34 | 41 | Y | 56886 | 3-hydroxy-3-methylglutaryl coenzyme A synthase OS=Rattus norvegicus OX=10116 GN=Hmgcs2 PE=1 SV=1 |
| **13** | 8 | P10719\|AT | 321.39 | 69 | 69 | 43450000 | 25 | 25 | 30 | Y | 56354 | ATP synthase subunit beta, mitochondrial OS=Rattus norvegicus OX=10116 GN=Atp5f1b PE=1 SV=2 |
| **13** | 9 | tr\|G3V6D3 | 321.39 | 69 | 69 | 43450000 | 25 | 25 | 30 | Y | 56345 | ATP synthase subunit beta OS=Rattus norvegicus OX=10116 GN=Atp5f1b PE=1 SV=1 |
| **11** | 13 | P15999\|AT | 320.03 | 45 | 45 | 36563000 | 26 | 26 | 33 | Y | 59754 | ATP synthase subunit alpha, mitochondrial OS=Rattus norvegicus OX=10116 GN=Atp5f1a PE=1 SV=2 |
| **2** | 16 | P67779\|PH | 310.02 | 89 | 89 | 147040000 | 32 | 31 | 62 | Y | 29820 | Prohibitin OS=Rattus norvegicus OX=10116 GN=Phb PE=1 SV=1 |
| **12** | 6 | P52873\|PY | 302.63 | 30 | 30 | 28604000 | 26 | 26 | 32 | Y | 129777 | Pyruvate carboxylase, mitochondrial OS=Rattus norvegicus OX=10116 GN=Pc PE=1 SV=2 |
| **12** | 7 | tr\|A0A0G2 | 302.63 | 28 | 28 | 28604000 | 26 | 26 | 32 | Y | 140005 | Pyruvate carboxylase, mitochondrial OS=Rattus norvegicus OX=10116 GN=Pc PE=1 SV=1 |
| **20** | 24 | Q641Y2\|NI | 297.4 | 37 | 37 | 28281000 | 16 | 16 | 20 | Y | 52562 | NADH dehydrogenase [ubiquinone] iron-sulfur protein 2, mitochondrial OS=Rattus norvegicus OX=10116 GN=Ndufs2 PE=1 SV=1 |
| **25** | 23 | Q561S0\|NI | 293.13 | 46 | 46 | 15831000 | 17 | 17 | 18 | Y | 40493 | NADH dehydrogenase [ubiquinone] 1 alpha subcomplex subunit 10, mitochondrial OS=Rattus norvegicus OX=10116 GN=Ndufa10 PE=1 SV=1 |
| **14** | 21 | tr\|Q5XIH3\| | 274.37 | 40 | 40 | 22204000 | 20 | 20 | 26 | Y | 50731 | NADH dehydrogenase [ubiquinone] flavoprotein 1, mitochondrial OS=Rattus norvegicus OX=10116 GN=Ndufv1 PE=1 SV=1 |
| **10** | 12 | Q66HF1\|NI | 272.99 | 49 | 49 | 78123000 | 33 | 33 | 40 | Y | 79412 | NADH-ubiquinone oxidoreductase 75 kDa subunit, mitochondrial OS=Rattus norvegicus OX=10116 GN=Ndufs1 PE=1 SV=1 |
| **17** | 14 | Q5BK63\|NI | 268.38 | 55 | 55 | 29079000 | 21 | 21 | 21 | Y | 42559 | NADH dehydrogenase [ubiquinone] 1 alpha subcomplex subunit 9, mitochondrial OS=Rattus norvegicus OX=10116 GN=Ndufa9 PE=1 SV=2 |
| **18** | 20 | tr\|F1LN92\| | 265.95 | 32 | 32 | 16500000 | 21 | 20 | 21 | Y | 89352 | AFG3-like matrix AAA peptidase subunit 2 OS=Rattus norvegicus OX=10116 GN=Afg3l2 PE=1 SV=1 |
| **4** | 18 | Q5XIH7\|PH | 265.54 | 82 | 82 | 126220000 | 30 | 30 | 53 | Y | 33312 | Prohibitin-2 OS=Rattus norvegicus OX=10116 GN=Phb2 PE=1 SV=1 |
| **19** | 22 | tr\|A0A0G2 | 264 | 25 | 25 | 17778000 | 18 | 18 | 20 | Y | 86230 | MICOS complex subunit MIC60 OS=Rattus norvegicus OX=10116 GN=Immt PE=1 SV=1 |
| **29** | 28 | P18163\|AC | 263.52 | 20 | 20 | 17423000 | 14 | 14 | 14 | Y | 78179 | Long-chain-fatty-acid--CoA ligase 1 OS=Rattus norvegicus OX=10116 GN=Acsl1 PE=1 SV=1 |
| **24** | 36 | tr\|D3ZG43 | 261.82 | 51 | 51 | 45523000 | 13 | 13 | 18 | N | 30226 | NADH dehydrogenase (Ubiquinone) Fe-S protein 3 (Predicted), isoform CRA_c OS=Rattus norvegicus OX=10116 GN=Ndufs3 PE=1 SV=1 |
| **16** | 45 | P20788\|UC | 252.67 | 41 | 41 | 31501000 | 15 | 15 | 22 | N | 29446 | Cytochrome b-c1 complex subunit Rieske, mitochondrial OS=Rattus norvegicus OX=10116 GN=Uqcrfs1 PE=1 SV=2 |
| **15** | 31 | Q60587\|EC | 251.23 | 35 | 35 | 22242000 | 17 | 17 | 25 | Y | 51414 | Trifunctional enzyme subunit beta, mitochondrial OS=Rattus norvegicus OX=10116 GN=Hadhb PE=1 SV=1 |
| **26** | 27 | P85834\|EF | 250.81 | 33 | 33 | 12366000 | 14 | 14 | 15 | Y | 49522 | Elongation factor Tu, mitochondrial OS=Rattus norvegicus OX=10116 GN=Tufm PE=1 SV=1 |
| **28** | 32 | P13086\|SU | 247.11 | 41 | 41 | 17347000 | 13 | 13 | 14 | Y | 36148 | Succinate--CoA ligase [ADP/GDP-forming] subunit alpha, mitochondrial OS=Rattus norvegicus OX=10116 GN=Suclg1 PE=2 SV=2 |
| **28** | 33 | tr\|A0A0H2 | 247.11 | 39 | 39 | 17347000 | 13 | 13 | 14 | Y | 37560 | Succinate--CoA ligase [ADP/GDP-forming] subunit alpha, mitochondrial OS=Rattus norvegicus OX=10116 GN=Suclg1 PE=1 SV=1 |
| **23** | 37 | Q4FZT0\|ST | 237.69 | 39 | 39 | 29327000 | 16 | 16 | 18 | Y | 38414 | Stomatin-like protein 2, mitochondrial OS=Rattus norvegicus OX=10116 GN=Stoml2 PE=1 SV=1 |
| **38** | 93 | tr\|D4A565 | 229.37 | 31 | 31 | 10615000 | 8 | 8 | 10 | N | 21664 | NADH dehydrogenase (Ubiquinone) 1 beta subcomplex, 5 (Predicted), isoform CRA_b OS=Rattus norvegicus OX=10116 GN=Ndufb5 PE=1 SV=1 |
| **21** | 25 | tr\|B2GV15 | 225.55 | 49 | 49 | 20838000 | 17 | 17 | 20 | N | 53274 | Dihydrolipoamide acetyltransferase component of pyruvate dehydrogenase complex OS=Rattus norvegicus OX=10116 GN=Dbt PE=1 SV=1 |
| **27** | 42 | tr\|D3ZFQ8 | 224.57 | 38 | 38 | 32944000 | 11 | 11 | 15 | Y | 35435 | Cytochrome c-1 OS=Rattus norvegicus OX=10116 GN=Cyc1 PE=1 SV=3 |
| **30** | 48 | Q9WVK3\|P | 222.16 | 32 | 32 | 18283000 | 7 | 7 | 11 | Y | 32433 | Peroxisomal trans-2-enoyl-CoA reductase OS=Rattus norvegicus OX=10116 GN=Pecr PE=2 SV=1 |
| **30** | 49 | tr\|A0A0G2 | 222.16 | 31 | 31 | 18283000 | 7 | 7 | 11 | Y | 32737 | Peroxisomal trans-2-enoyl-CoA reductase OS=Rattus norvegicus OX=10116 GN=Pecr PE=1 SV=1 |
| **31** | 47 | P0C2X9\|AL | 217.83 | 17 | 17 | 6482400 | 9 | 9 | 11 | Y | 61869 | Delta-1-pyrroline-5-carboxylate dehydrogenase, mitochondrial OS=Rattus norvegicus OX=10116 GN=Aldh4a1 PE=1 SV=1 |
| **32** | 46 | P17764\|TH | 215.94 | 25 | 25 | 7123200 | 8 | 8 | 10 | Y | 44695 | Acetyl-CoA acetyltransferase, mitochondrial OS=Rattus norvegicus OX=10116 GN=Acat1 PE=1 SV=1 |
| **33** | 44 | P16970\|AB | 212.7 | 16 | 16 | 5556100 | 10 | 10 | 10 | Y | 75316 | ATP-binding cassette sub-family D member 3 OS=Rattus norvegicus OX=10116 GN=Abcd3 PE=1 SV=3 |
| **41** | 54 | tr\|D4A0T0 | 208.11 | 43 | 43 | 10083000 | 7 | 7 | 8 | Y | 20859 | NADH:ubiquinone oxidoreductase subunit B10 OS=Rattus norvegicus OX=10116 GN=Ndufb10 PE=1 SV=1 |
| **84** | 170 | Q63362\|NI | 203.96 | 41 | 41 | 2868600 | 3 | 3 | 4 | N | 13412 | NADH dehydrogenase [ubiquinone] 1 alpha subcomplex subunit 5 OS=Rattus norvegicus OX=10116 GN=Ndufa5 PE=1 SV=3 |
| **36** | 51 | tr\|Q5EBA4 | 202.91 | 27 | 27 | 10255000 | 8 | 8 | 10 | Y | 33215 | Nipsnap1 protein (Fragment) OS=Rattus norvegicus OX=10116 GN=Nipsnap1 PE=2 SV=1 |
| **36** | 52 | tr\|G3V728 | 202.91 | 27 | 27 | 10255000 | 8 | 8 | 10 | Y | 33346 | 4-nitrophenylphosphatase domain and non-neuronal SNAP25-like protein homolog 1 (C. elegans), isoform CRA_b OS=Rattus norvegicus OX=10116 GN=Nipsnap1 PE=1 SV=1 |
| **64** | 64 | Q02253\|M | 195.86 | 13 | 13 | 4306200 | 5 | 5 | 5 | Y | 57808 | Methylmalonate-semialdehyde dehydrogenase [acylating], mitochondrial OS=Rattus norvegicus OX=10116 GN=Aldh6a1 PE=1 SV=1 |
| **64** | 65 | tr\|G3V7J0\| | 195.86 | 13 | 13 | 4306200 | 5 | 5 | 5 | Y | 57748 | Aldehyde dehydrogenase family 6, subfamily A1, isoform CRA_b OS=Rattus norvegicus OX=10116 GN=Aldh6a1 PE=1 SV=1 |
| **22** | 39 | P29147\|BD | 195.15 | 29 | 29 | 18391000 | 16 | 15 | 19 | Y | 38202 | D-beta-hydroxybutyrate dehydrogenase, mitochondrial OS=Rattus norvegicus OX=10116 GN=Bdh1 PE=1 SV=2 |
| **22** | 40 | tr\|A0A0G2 | 195.15 | 29 | 29 | 18391000 | 16 | 15 | 19 | Y | 38333 | 3-hydroxybutyrate dehydrogenase, type 1, isoform CRA_a OS=Rattus norvegicus OX=10116 GN=Bdh1 PE=1 SV=1 |
| **35** | 38 | tr\|B0BMT9 | 194.51 | 24 | 24 | 7208900 | 9 | 9 | 10 | N | 50202 | Sqrdl protein OS=Rattus norvegicus OX=10116 GN=Sqor PE=1 SV=1 |
| **37** | 60 | tr\|B0BNE6 | 194.08 | 29 | 29 | 9149700 | 8 | 8 | 10 | Y | 23970 | NADH dehydrogenase (Ubiquinone) Fe-S protein 8 (Predicted), isoform CRA_a OS=Rattus norvegicus OX=10116 GN=Ndufs8 PE=1 SV=1 |
| **40** | 55 | P24329\|TH | 191.86 | 29 | 29 | 8038400 | 8 | 8 | 9 | N | 33407 | Thiosulfate sulfurtransferase OS=Rattus norvegicus OX=10116 GN=Tst PE=1 SV=3 |
| **60** | 61 | P63039\|CH | 187.49 | 13 | 13 | 2185500 | 5 | 5 | 5 | N | 60956 | 60 kDa heat shock protein, mitochondrial OS=Rattus norvegicus OX=10116 GN=Hspd1 PE=1 SV=1 |
| **60** | 62 | tr\|A0A482I | 187.49 | 13 | 13 | 2185500 | 5 | 5 | 5 | N | 60956 | Hsp60 OS=Rattus norvegicus OX=10116 GN=Hspd1 PE=2 SV=1 |
| **73** | 91 | P45953\|AC | 182.91 | 10 | 10 | 1476700 | 3 | 3 | 4 | N | 70749 | Very long-chain specific acyl-CoA dehydrogenase, mitochondrial OS=Rattus norvegicus OX=10116 GN=Acadvl PE=1 SV=1 |
| **73** | 92 | tr\|Q5M9H2 | 182.91 | 10 | 10 | 1476700 | 3 | 3 | 4 | N | 70821 | Acyl-Coenzyme A dehydrogenase, very long chain OS=Rattus norvegicus OX=10116 GN=Acadvl PE=1 SV=1 |
| **54** | 69 | Q5XIF3\|ND | 182.02 | 34 | 34 | 2413200 | 6 | 6 | 6 | N | 19741 | NADH dehydrogenase [ubiquinone] iron-sulfur protein 4, mitochondrial OS=Rattus norvegicus OX=10116 GN=Ndufs4 PE=1 SV=1 |
| **52** | 73 | tr\|B2RYS8\| | 180.1 | 28 | 28 | 8363800 | 4 | 4 | 6 | Y | 21959 | NADH dehydrogenase [ubiquinone] 1 beta subcomplex subunit 8, mitochondrial OS=Rattus norvegicus OX=10116 GN=Ndufb8 PE=1 SV=1 |
| **48** | 59 | tr\|D3ZJY1\| | 178.84 | 35 | 35 | 3394800 | 6 | 6 | 7 | N | 30237 | Mitochondrial ribosomal protein L28 OS=Rattus norvegicus OX=10116 GN=Mrpl28 PE=1 SV=1 |
| **53** | 53 | B3DMA2\|A | 178.16 | 12 | 12 | 2023800 | 6 | 6 | 6 | N | 87371 | Acyl-CoA dehydrogenase family member 11 OS=Rattus norvegicus OX=10116 GN=Acad11 PE=1 SV=1 |

| | | | | | | | | | | | | |
|---|---|---|---|---|---|---|---|---|---|---|---|---|
| **39** | 67 | tr\|Q5RJN0 | 170.89 | 26 | 26 | 13246000 | 8 | 8 | 9 | N | 23945 | NADH dehydrogenase (Ubiquinone) Fe-S protein 7 OS=Rattus norvegicus OX=10116 GN=Ndufs7 PE=1 SV=1 |
| **57** | 76 | O70351\|H( | 168.95 | 32 | 32 | 9910400 | 5 | 5 | 6 | Y | 27246 | 3-hydroxyacyl-CoA dehydrogenase type-2 OS=Rattus norvegicus OX=10116 GN=Hsd17b10 PE=1 SV=3 |
| **57** | 77 | tr\|B0BMW | 168.95 | 32 | 32 | 9910400 | 5 | 5 | 6 | Y | 27250 | 3-hydroxyacyl-CoA dehydrogenase type-2 OS=Rattus norvegicus OX=10116 GN=Hsd17b10 PE=1 SV=1 |
| **67** | 106 | tr\|D3ZXF9\| | 168.55 | 18 | 18 | 2117100 | 4 | 4 | 5 | N | 29441 | Mitochondrial ribosomal protein L12 OS=Rattus norvegicus OX=10116 GN=Mrpl12 PE=1 SV=1 |
| **90** | 145 | P62909\|RS | 166 | 17 | 17 | 4154300 | 3 | 3 | 3 | Y | 26674 | 40S ribosomal protein S3 OS=Rattus norvegicus OX=10116 GN=Rps3 PE=1 SV=1 |
| **55** | 57 | tr\|D4A131 | 154.79 | 35 | 35 | 3910000 | 6 | 6 | 6 | N | 33082 | Mitochondrial ribosomal protein L4 OS=Rattus norvegicus OX=10116 GN=Mrpl4 PE=1 SV=1 |
| **81** | 84 | tr\|Q2TVU3 | 153.85 | 19 | 19 | 997960 | 4 | 4 | 4 | Y | 49412 | TID1 OS=Rattus norvegicus OX=10116 GN=Dnaja3 PE=2 SV=1 |
| **81** | 85 | tr\|A0A0G2 | 153.85 | 19 | 19 | 997960 | 4 | 4 | 4 | Y | 49416 | DnaJ heat shock protein family (Hsp40) member A3 OS=Rattus norvegicus OX=10116 GN=Dnaja3 PE=1 SV=1 |
| **81** | 86 | tr\|Q2UZS7 | 153.85 | 18 | 18 | 997960 | 4 | 4 | 4 | Y | 52399 | Tid-1 long isoform OS=Rattus norvegicus OX=10116 GN=Dnaja3 PE=2 SV=1 |
| **81** | 87 | tr\|A0A0G2 | 153.85 | 18 | 18 | 997960 | 4 | 4 | 4 | Y | 52403 | DnaJ heat shock protein family (Hsp40) member A3 OS=Rattus norvegicus OX=10116 GN=Dnaja3 PE=1 SV=1 |
| **42** | 72 | tr\|F1LXA0\| | 153.63 | 38 | 38 | 7140800 | 6 | 6 | 8 | N | 17178 | NADH dehydrogenase [ubiquinone] 1 alpha subcomplex subunit 12 OS=Rattus norvegicus OX=10116 GN=Ndufa12 PE=1 SV=2 |
| **59** | 113 | tr\|B2RYW3 | 148.13 | 25 | 25 | 6365300 | 6 | 6 | 6 | N | 21892 | NADH dehydrogenase (Ubiquinone) 1 beta subcomplex, 9 OS=Rattus norvegicus OX=10116 GN=Ndufb9 PE=1 SV=1 |
| **47** | 63 | Q09073\|AI | 147.82 | 24 | 24 | 4899900 | 7 | 7 | 7 | Y | 32901 | ADP/ATP translocase 2 OS=Rattus norvegicus OX=10116 GN=Slc25a5 PE=1 SV=3 |
| **91** | 144 | tr\|A0A0G2 | 147.66 | 33 | 33 | 2387400 | 3 | 3 | 3 | N | 19965 | NADH dehydrogenase [ubiquinone] 1 alpha subcomplex subunit 8 OS=Rattus norvegicus OX=10116 GN=Ndufa8 PE=1 SV=1 |
| **51** | 68 | tr\|Q5UAJ6 | 147.14 | 25 | 25 | 9661900 | 5 | 5 | 6 | Y | 25942 | Cytochrome c oxidase subunit 2 OS=Rattus norvegicus OX=10116 GN=COX2 PE=3 SV=1 |
| **34** | 58 | P19234\|NI | 143.9 | 31 | 31 | 30551000 | 9 | 9 | 10 | N | 27378 | NADH dehydrogenase [ubiquinone] flavoprotein 2, mitochondrial OS=Rattus norvegicus OX=10116 GN=Ndufv2 PE=1 SV=2 |
| **43** | 50 | P10888\|CC | 135.95 | 38 | 38 | 8531200 | 7 | 7 | 8 | N | 19515 | Cytochrome c oxidase subunit 4 isoform 1, mitochondrial OS=Rattus norvegicus OX=10116 GN=Cox4i1 PE=1 SV=1 |
| **58** | 110 | tr\|D3ZCZ9\| | 135.49 | 34 | 34 | 3687500 | 4 | 4 | 6 | Y | 13040 | NADH dehydrogenase [ubiquinone] iron-sulfur protein 6, mitochondrial OS=Rattus norvegicus OX=10116 GN=LOC100912599 PE=1 SV=1 |
| **70** | 88 | tr\|D4A3V2 | 134.14 | 33 | 33 | 4426400 | 4 | 4 | 5 | N | 15224 | NADH dehydrogenase [ubiquinone] 1 alpha subcomplex subunit 6 OS=Rattus norvegicus OX=10116 GN=Ndufa6 PE=1 SV=1 |
| **65** | 66 | Q8VID1\|DI | 133.76 | 18 | 18 | 2671700 | 5 | 5 | 5 | Y | 29822 | Dehydrogenase/reductase SDR family member 4 OS=Rattus norvegicus OX=10116 GN=Dhrs4 PE=2 SV=2 |
| **49** | 56 | P04636\|MI | 133.59 | 18 | 18 | 2311900 | 6 | 6 | 6 | Y | 35684 | Malate dehydrogenase, mitochondrial OS=Rattus norvegicus OX=10116 GN=Mdh2 PE=1 SV=2 |
| **82** | 90 | tr\|D3ZMR9 | 125.22 | 24 | 24 | 1815400 | 4 | 4 | 4 | N | 23390 | Mitochondrial ribosomal protein L21 OS=Rattus norvegicus OX=10116 GN=Mrpl21 PE=1 SV=1 |
| **94** | 156 | P70584\|AC | 124.78 | 7 | 7 | 2063800 | 3 | 3 | 3 | Y | 47824 | Short/branched chain specific acyl-CoA dehydrogenase, mitochondrial OS=Rattus norvegicus OX=10116 GN=Acadsb PE=1 SV=1 |
| **94** | 157 | tr\|A0A0A0 | 124.78 | 7 | 7 | 2063800 | 3 | 3 | 3 | Y | 52043 | Short/branched chain-specific acyl-CoA dehydrogenase, mitochondrial OS=Rattus norvegicus OX=10116 GN=Acadsb PE=1 SV=2 |
| **78** | 96 | tr\|D3ZFJ6\| | 122.91 | 12 | 12 | 2279000 | 3 | 3 | 4 | N | 60420 | Lactamase, beta OS=Rattus norvegicus OX=10116 GN=Lactb PE=1 SV=1 |
| **93** | 152 | tr\|D3ZX69\| | 122.75 | 15 | 15 | 856680 | 2 | 2 | 3 | Y | 29466 | 39S ribosomal protein L10, mitochondrial OS=Rattus norvegicus OX=10116 GN=Mrpl10 PE=1 SV=1 |
| **93** | 153 | P0C2C4\|RN | 122.75 | 15 | 15 | 856680 | 2 | 2 | 3 | Y | 29940 | 39S ribosomal protein L10, mitochondrial OS=Rattus norvegicus OX=10116 GN=Mrpl10 PE=1 SV=1 |
| **170** | 181 | Q6AXV4\|SA | 121.41 | 4 | 4 | 788140 | 1 | 1 | 1 | N | 51960 | Sorting and assembly machinery component 50 homolog OS=Rattus norvegicus OX=10116 GN=Samm50 PE=1 SV=1 |
| **83** | 105 | tr\|D3ZF13\| | 121.33 | 21 | 21 | 5391600 | 4 | 4 | 4 | Y | 17514 | Acyl carrier protein OS=Rattus norvegicus OX=10116 GN=Ndufab1 PE=1 SV=1 |
| **66** | 100 | tr\|D3ZE15\| | 120.53 | 25 | 25 | 6146700 | 5 | 5 | 5 | Y | 16777 | NADH:ubiquinone oxidoreductase subunit A13 OS=Rattus norvegicus OX=10116 GN=Ndufa13 PE=1 SV=1 |
| **46** | 95 | tr\|Q6QI09\| | 119.69 | 9 | 9 | 1089200 | 3 | 3 | 7 | Y | 67721 | ATP synthase subunit gamma, mitochondrial OS=Rattus norvegicus OX=10116 GN=Taf3 PE=1 SV=1 |
| **46** | 97 | P35435\|AT | 119.69 | 20 | 20 | 1089200 | 3 | 3 | 7 | Y | 30191 | ATP synthase subunit gamma, mitochondrial OS=Rattus norvegicus OX=10116 GN=Atp5f1c PE=1 SV=2 |
| **45** | 75 | tr\|B1WBY5 | 118.4 | 12 | 12 | 9331300 | 6 | 6 | 7 | N | 63205 | DnaJ (Hsp40) homolog, subfamily C, member 11 OS=Rattus norvegicus OX=10116 GN=Dnajc11 PE=1 SV=1 |
| **56** | 70 | tr\|B2RYT5\| | 118.39 | 44 | 44 | 5012600 | 5 | 5 | 6 | N | 12651 | Cox7a2l protein OS=Rattus norvegicus OX=10116 GN=Cox7a2l PE=2 SV=1 |
| **114** | 154 | tr\|A0A0G2 | 116.94 | 39 | 39 | 2333600 | 2 | 2 | 2 | N | 7099 | Ubiquinol-cytochrome c reductase, complex III subunit X OS=Rattus norvegicus OX=10116 GN=Uqcr10 PE=1 SV=1 |
| **114** | 172 | tr\|B2RYX1\| | 116.94 | 38 | 38 | 2333600 | 2 | 2 | 2 | N | 7462 | LOC685322 protein OS=Rattus norvegicus OX=10116 GN=Uqcr10 PE=2 SV=1 |
| **50** | 112 | tr\|Q5PQZ9 | 115.26 | 27 | 27 | 3205900 | 5 | 5 | 6 | N | 14359 | NADH dehydrogenase [ubiquinone] 1 subunit C2 OS=Rattus norvegicus OX=10116 GN=Ndufc2 PE=1 SV=1 |
| **80** | 71 | tr\|F1LPG5\| | 114.95 | 47 | 47 | 4271700 | 4 | 4 | 4 | N | 15064 | NADH:ubiquinone oxidoreductase subunit B4 OS=Rattus norvegicus OX=10116 GN=Ndufb4 PE=1 SV=1 |
| **62** | 78 | P17178\|CP | 113.64 | 14 | 14 | 2531000 | 5 | 5 | 5 | N | 60733 | Sterol 26-hydroxylase, mitochondrial OS=Rattus norvegicus OX=10116 GN=Cyp27a1 PE=1 SV=1 |
| **62** | 79 | tr\|A0A0H2 | 113.64 | 13 | 13 | 2531000 | 5 | 5 | 5 | N | 62492 | RCG24013, isoform CRA_a OS=Rattus norvegicus OX=10116 GN=Cyp27a1 PE=1 SV=1 |
| **44** | 104 | tr\|A9UMV9 | 113.41 | 33 | 33 | 4859600 | 4 | 4 | 8 | N | 12500 | NADH:ubiquinone oxidoreductase subunit A7 OS=Rattus norvegicus OX=10116 GN=Ndufa7 PE=1 SV=1 |
| **74** | 74 | tr\|D3ZS58\| | 112.49 | 40 | 40 | 2699900 | 4 | 4 | 4 | N | 10845 | NADH dehydrogenase [ubiquinone] 1 alpha subcomplex subunit 2 OS=Rattus norvegicus OX=10116 GN=Ndufa2 PE=1 SV=1 |
| **119** | 205 | Q5M9I5\|Q | 111.86 | 21 | 21 | 2307300 | 1 | 1 | 2 | N | 10424 | Cytochrome b-c1 complex subunit 6, mitochondrial OS=Rattus norvegicus OX=10116 GN=Uqcrh PE=3 SV=1 |
| **69** | 133 | P12075\|CC | 111.85 | 35 | 35 | 7311700 | 4 | 4 | 5 | Y | 13915 | Cytochrome c oxidase subunit 5B, mitochondrial OS=Rattus norvegicus OX=10116 GN=Cox5b PE=1 SV=2 |
| **76** | 121 | P08461\|OI | 111.55 | 7 | 7 | 494620 | 4 | 3 | 4 | N | 67166 | Dihydrolipoyllysine-residue acetyltransferase component of pyruvate dehydrogenase complex, mitochondrial OS=Rattus norvegicus OX=10116 GN=Dlat PE=1 SV=3 |
| **92** | 148 | F1LU71\|AU | 111.09 | 12 | 12 | 639520 | 3 | 3 | 3 | Y | 33341 | Methylglutaconyl-CoA hydratase, mitochondrial OS=Rattus norvegicus OX=10116 GN=Auh PE=1 SV=2 |
| **63** | 122 | P05508\|NU | 109.96 | 10 | 10 | 1275800 | 4 | 4 | 5 | Y | 51783 | NADH-ubiquinone oxidoreductase chain 4 OS=Rattus norvegicus OX=10116 GN=Mtnd4 PE=3 SV=3 |
| **63** | 123 | tr\|A7XYB9\| | 109.96 | 10 | 10 | 1275800 | 4 | 4 | 5 | Y | 51773 | NADH-ubiquinone oxidoreductase chain 4 OS=Rattus norvegicus OX=10116 GN=Nd4 PE=3 SV=1 |
| **63** | 124 | tr\|Q06QA2 | 109.96 | 10 | 10 | 1275800 | 4 | 4 | 5 | Y | 51787 | NADH-ubiquinone oxidoreductase chain 4 OS=Rattus norvegicus OX=10116 GN=ND4 PE=3 SV=1 |
| **63** | 125 | tr\|Q7HKW3 | 109.96 | 10 | 10 | 1275800 | 4 | 4 | 5 | Y | 51801 | NADH-ubiquinone oxidoreductase chain 4 (Fragment) OS=Rattus norvegicus OX=10116 GN=NADH4 PE=3 SV=1 |
| **63** | 126 | tr\|Q06QG7 | 109.96 | 10 | 10 | 1275800 | 4 | 4 | 5 | Y | 51791 | NADH-ubiquinone oxidoreductase chain 4 OS=Rattus norvegicus OX=10116 GN=ND4 PE=3 SV=1 |
| **63** | 127 | tr\|Q06Q89 | 109.96 | 10 | 10 | 1275800 | 4 | 4 | 5 | Y | 51819 | NADH-ubiquinone oxidoreductase chain 4 OS=Rattus norvegicus OX=10116 GN=ND4 PE=3 SV=1 |
| **63** | 128 | tr\|Q35737 | 109.96 | 10 | 10 | 1275800 | 4 | 4 | 5 | Y | 51765 | NADH-ubiquinone oxidoreductase chain 4 OS=Rattus norvegicus OX=10116 PE=3 SV=1 |
| **63** | 129 | tr\|Q8HIC6\| | 109.96 | 10 | 10 | 1275800 | 4 | 4 | 5 | Y | 51783 | NADH-ubiquinone oxidoreductase chain 4 OS=Rattus norvegicus OX=10116 GN=Mt-nd4 PE=3 SV=1 |
| **63** | 130 | tr\|D2E6K0\| | 109.96 | 10 | 10 | 1275800 | 4 | 4 | 5 | Y | 51833 | NADH-ubiquinone oxidoreductase chain 4 OS=Rattus norvegicus OX=10116 GN=ND4 PE=3 SV=1 |
| **63** | 131 | tr\|Q06QE1 | 109.96 | 10 | 10 | 1275800 | 4 | 4 | 5 | Y | 51851 | NADH-ubiquinone oxidoreductase chain 4 OS=Rattus norvegicus OX=10116 GN=ND4 PE=3 SV=1 |
| **71** | 94 | Q7TQ16\|Q | 108.19 | 46 | 46 | 4040100 | 5 | 5 | 5 | N | 9849 | Cytochrome b-c1 complex subunit 8 OS=Rattus norvegicus OX=10116 GN=Uqcrq PE=3 SV=1 |
| **75** | 119 | P32198\|CP | 103.08 | 5 | 5 | 1299400 | 3 | 3 | 4 | N | 88126 | Carnitine O-palmitoyltransferase 1, liver isoform OS=Rattus norvegicus OX=10116 GN=Cpt1a PE=1 SV=2 |
| **99** | 120 | tr\|A0A0G2 | 102.04 | 8 | 8 | 728990 | 3 | 1 | 3 | N | 46666 | Death-associated protein 3 OS=Rattus norvegicus OX=10116 GN=Dap3 PE=1 SV=1 |
| **68** | 115 | tr\|D3ZIL6\| | 101.55 | 18 | 18 | 2891300 | 4 | 3 | 5 | N | 31779 | Enoyl CoA hydratase domain-containing 2 OS=Rattus norvegicus OX=10116 GN=Echdc2 PE=1 SV=1 |

| | | | | | | | | | | | | |
|---|---|---|---|---|---|---|---|---|---|---|---|---|
| **101** | 138 | tr\|Q5U2T0 | 100.69 | 7 | 7 | 7114400 | 3 | 1 | 3 | N | 44494 | Death associated protein 3 OS=Rattus norvegicus OX=10116 GN=Dap3 PE=2 SV=1 |
| **101** | 139 | tr\|F7EZZ0\| | 100.69 | 7 | 7 | 7114400 | 3 | 1 | 3 | N | 45112 | Death-associated protein 3 OS=Rattus norvegicus OX=10116 GN=Dap3 PE=1 SV=1 |
| **102** | 140 | tr\|M0R6J0 | 100.52 | 9 | 9 | 2367400 | 3 | 3 | 3 | N | 38374 | Mitochondrial ribosomal protein L39 OS=Rattus norvegicus OX=10116 GN=Mrpl39 PE=1 SV=2 |
| **197** | 206 | P13803\|ET | 98.35 | 8 | 8 | 641310 | 1 | 1 | 1 | N | 34951 | Electron transfer flavoprotein subunit alpha, mitochondrial OS=Rattus norvegicus OX=10116 GN=Etfa PE=1 SV=4 |
| **134** | 247 | tr\|B2GV62 | 96.32 | 14 | 14 | 123010 | 1 | 1 | 2 | N | 17526 | Mitochondrial ribosomal protein L20 OS=Rattus norvegicus OX=10116 GN=Mrpl20 PE=1 SV=1 |
| **171** | 196 | tr\|B0BN52 | 94.67 | 5 | 5 | 384060 | 1 | 1 | 1 | N | 33528 | Mitochondrial carrier 2 OS=Rattus norvegicus OX=10116 GN=Mtch2 PE=2 SV=1 |
| **171** | 197 | tr\|A0A0G2 | 94.67 | 5 | 5 | 384060 | 1 | 1 | 1 | N | 33758 | Mitochondrial carrier 2 OS=Rattus norvegicus OX=10116 GN=Mtch2 PE=1 SV=1 |
| **171** | 212 | tr\|A0A0G2 | 94.67 | 7 | 7 | 384060 | 1 | 1 | 1 | N | 26999 | Mitochondrial carrier 2 OS=Rattus norvegicus OX=10116 GN=Mtch2 PE=1 SV=1 |
| **86** | 111 | Q5RJR8\|LR | 93.61 | 10 | 10 | 1102700 | 3 | 3 | 3 | N | 34869 | Leucine-rich repeat-containing protein 59 OS=Rattus norvegicus OX=10116 GN=Lrrc59 PE=1 SV=1 |
| **123** | 171 | tr\|Q5M94 | 93.59 | 13 | 13 | 714950 | 2 | 2 | 2 | Y | 28340 | Nipsnap homolog 3A (C. elegans) OS=Rattus norvegicus OX=10116 GN=Nipsnap3b PE=1 SV=1 |
| **250** | 249 | tr\|D3ZY44\| | 91.57 | 5 | 5 | 149740 | 1 | 1 | 1 | N | 32176 | Mitochondrial ribosomal protein S2 OS=Rattus norvegicus OX=10116 GN=Mrps2 PE=1 SV=1 |
| **249** | 248 | tr\|G3V6I4\| | 91.37 | 9 | 9 | 2932800 | 1 | 1 | 1 | N | 37791 | Mitochondrial amidoxime reducing component 1 OS=Rattus norvegicus OX=10116 GN=Marc1 PE=1 SV=1 |
| **111** | 159 | tr\|Q4G067 | 90.92 | 8 | 8 | 1687500 | 2 | 2 | 2 | N | 37440 | Mitochondrial ribosomal protein L44 OS=Rattus norvegicus OX=10116 GN=Mrpl44 PE=1 SV=1 |
| **135** | 250 | Q80W89\|N | 90.19 | 11 | 11 | 1599300 | 1 | 1 | 2 | N | 14854 | NADH dehydrogenase [ubiquinone] 1 alpha subcomplex subunit 11 OS=Rattus norvegicus OX=10116 GN=Ndufa11 PE=2 SV=1 |
| **172** | 219 | tr\|Q8HN20 | 88.81 | 26 | 26 | 498640 | 1 | 1 | 1 | Y | 8534 | NADH-ubiquinone oxidoreductase chain 1 (Fragment) OS=Rattus norvegicus OX=10116 PE=2 SV=1 |
| **172** | 220 | tr\|Q8SEZ8\| | 88.81 | 6 | 6 | 498640 | 1 | 1 | 1 | Y | 36133 | NADH-ubiquinone oxidoreductase chain 1 (Fragment) OS=Rattus norvegicus OX=10116 GN=NADH1 PE=3 SV=1 |
| **172** | 221 | P03889\|NU | 88.81 | 6 | 6 | 498640 | 1 | 1 | 1 | Y | 36145 | NADH-ubiquinone oxidoreductase chain 1 OS=Rattus norvegicus OX=10116 GN=Mtnd1 PE=1 SV=3 |
| **172** | 222 | tr\|Q8HID1 | 88.81 | 6 | 6 | 498640 | 1 | 1 | 1 | Y | 36145 | NADH-ubiquinone oxidoreductase chain 1 OS=Rattus norvegicus OX=10116 GN=Mt-nd1 PE=3 SV=1 |
| **172** | 223 | tr\|D2E6L7\| | 88.81 | 6 | 6 | 498640 | 1 | 1 | 1 | Y | 36062 | NADH-ubiquinone oxidoreductase chain 1 OS=Rattus norvegicus OX=10116 GN=ND1 PE=3 SV=1 |
| **87** | 99 | tr\|G3V6P2 | 85.05 | 8 | 8 | 1347300 | 3 | 3 | 3 | N | 48899 | Dihydrolipoamide S-succinyltransferase (E2 component of 2-oxo-glutarate complex), isoform CRA_a OS=Rattus norvegicus OX=10116 GN=Dlst PE=1 SV=1 |
| **104** | 182 | tr\|Q06Q97 | 84 | 10 | 10 | 1461900 | 2 | 2 | 3 | Y | 38485 | NADH-ubiquinone oxidoreductase chain 2 OS=Rattus norvegicus OX=10116 GN=ND2 PE=3 SV=1 |
| **104** | 183 | tr\|A0A0A1 | 84 | 10 | 10 | 1461900 | 2 | 2 | 3 | Y | 38542 | NADH-ubiquinone oxidoreductase chain 2 OS=Rattus norvegicus OX=10116 GN=ND2 PE=3 SV=1 |
| **104** | 184 | tr\|Q5UAJ8 | 84 | 10 | 10 | 1461900 | 2 | 2 | 3 | Y | 38455 | NADH-ubiquinone oxidoreductase chain 2 OS=Rattus norvegicus OX=10116 GN=ND2 PE=3 SV=1 |
| **77** | 134 | tr\|A0A0G2 | 83.59 | 6 | 6 | 3129800 | 4 | 4 | 4 | Y | 68759 | Serum albumin OS=Rattus norvegicus OX=10116 GN=Alb PE=1 SV=1 |
| **155** | 193 | Q5XI78\|OD | 82.53 | 3 | 3 | 205310 | 1 | 1 | 1 | N | 116296 | 2-oxoglutarate dehydrogenase, mitochondrial OS=Rattus norvegicus OX=10116 GN=Ogdh PE=1 SV=1 |
| **89** | 137 | Q9WVK7\|H | 82.41 | 13 | 13 | 2132000 | 3 | 3 | 3 | N | 34448 | Hydroxyacyl-coenzyme A dehydrogenase, mitochondrial OS=Rattus norvegicus OX=10116 GN=Hadh PE=2 SV=1 |
| **109** | 141 | Q924S5\|LC | 79.52 | 5 | 5 | 1406100 | 2 | 2 | 2 | N | 105792 | Lon protease homolog, mitochondrial OS=Rattus norvegicus OX=10116 GN=Lonp1 PE=2 SV=1 |
| **251** | 251 | tr\|A0A0G2 | 79.02 | 5 | 5 | 601910 | 1 | 1 | 1 | N | 33652 | Mitochondrial ribosomal protein L15 OS=Rattus norvegicus OX=10116 GN=Mrpl15 PE=1 SV=1 |
| **96** | 169 | tr\|A0A0G2 | 78.9 | 22 | 22 | 2363100 | 2 | 2 | 3 | N | 12730 | Uncharacterized protein OS=Rattus norvegicus OX=10116 PE=4 SV=1 |
| **96** | 175 | tr\|B5DEL8\| | 78.9 | 22 | 22 | 2363100 | 2 | 2 | 3 | N | 12700 | NADH dehydrogenase (Ubiquinone) Fe-S protein 5 OS=Rattus norvegicus OX=10116 GN=Ndufs5 PE=1 SV=1 |
| **95** | 166 | tr\|D4A4A9 | 78.66 | 11 | 11 | 446310 | 2 | 2 | 3 | N | 33633 | Mitochondrial ribosomal protein L19 OS=Rattus norvegicus OX=10116 GN=Mrpl19 PE=1 SV=1 |
| **97** | 107 | P0C2C0\|RN | 77.28 | 18 | 18 | 1458500 | 3 | 3 | 3 | N | 24039 | 39S ribosomal protein L22, mitochondrial OS=Rattus norvegicus OX=10116 GN=Mrpl22 PE=1 SV=1 |
| **97** | 108 | tr\|A0A0H2 | 77.28 | 17 | 17 | 1458500 | 3 | 3 | 3 | N | 25865 | 39S ribosomal protein L22, mitochondrial OS=Rattus norvegicus OX=10116 GN=Mrpl22 PE=1 SV=1 |
| **204** | 227 | tr\|F1LZW6 | 76.89 | 5 | 5 | 132240 | 1 | 1 | 1 | N | 54099 | Solute carrier family 25 member 13 OS=Rattus norvegicus OX=10116 GN=Slc25a13 PE=1 SV=2 |
| **88** | 114 | P19511\|AT | 74.76 | 13 | 13 | 2820800 | 3 | 3 | 3 | N | 28869 | ATP synthase F(0) complex subunit B1, mitochondrial OS=Rattus norvegicus OX=10116 GN=Atp5pb PE=1 SV=1 |
| **117** | 192 | tr\|D4A7L4\| | 74.47 | 21 | 21 | 6584700 | 2 | 2 | 2 | Y | 17634 | NADH dehydrogenase (Ubiquinone) 1 beta subcomplex, 11 (Predicted) OS=Rattus norvegicus OX=10116 GN=Ndufb11 PE=1 SV=1 |
| **98** | 117 | tr\|G3V734 | 74.19 | 10 | 10 | 2722200 | 3 | 3 | 3 | N | 36133 | 2,4-dienoyl CoA reductase 1, mitochondrial, isoform CRA_a OS=Rattus norvegicus OX=10116 GN=Decr1 PE=1 SV=1 |
| **98** | 118 | Q64591\|DE | 74.19 | 10 | 10 | 2722200 | 3 | 3 | 3 | N | 36133 | 2,4-dienoyl-CoA reductase, mitochondrial OS=Rattus norvegicus OX=10116 GN=Decr1 PE=1 SV=2 |
| **255** | 260 | tr\|D3ZUX5 | 73.61 | 5 | 5 | 921530 | 1 | 1 | 1 | N | 26435 | MICOS complex subunit OS=Rattus norvegicus OX=10116 GN=Chchd3 PE=1 SV=1 |
| **200** | 214 | Q5I0K8\|RT | 73.36 | 7 | 7 | 436930 | 1 | 1 | 1 | N | 28197 | 28S ribosomal protein S7, mitochondrial OS=Rattus norvegicus OX=10116 GN=Mrps7 PE=2 SV=2 |
| **130** | 188 | Q06647\|AT | 72.56 | 12 | 12 | 1102900 | 2 | 2 | 2 | N | 23398 | ATP synthase subunit O, mitochondrial OS=Rattus norvegicus OX=10116 GN=Atp5po PE=1 SV=1 |
| **79** | 103 | P04762\|CA | 71.62 | 9 | 9 | 676110 | 4 | 4 | 4 | N | 59757 | Catalase OS=Rattus norvegicus OX=10116 GN=Cat PE=1 SV=3 |
| **196** | 204 | P31399\|AT | 71.56 | 8 | 8 | 351130 | 1 | 1 | 1 | N | 18763 | ATP synthase subunit d, mitochondrial OS=Rattus norvegicus OX=10116 GN=Atp5pd PE=1 SV=3 |
| **131** | 189 | tr\|B2RYS2\| | 71.48 | 14 | 14 | 2586100 | 2 | 2 | 2 | N | 13559 | Cytochrome b-c1 complex subunit 7 OS=Rattus norvegicus OX=10116 GN=Uqcrb PE=1 SV=1 |
| **205** | 229 | tr\|A0A0G2 | 70.95 | 3 | 3 | 265100 | 1 | 1 | 1 | N | 46686 | 39S ribosomal protein L37, mitochondrial OS=Rattus norvegicus OX=10116 GN=Mrpl37 PE=1 SV=1 |
| **205** | 230 | Q6AXT0\|RI | 70.95 | 3 | 3 | 265100 | 1 | 1 | 1 | N | 48366 | 39S ribosomal protein L37, mitochondrial OS=Rattus norvegicus OX=10116 GN=Mrpl37 PE=2 SV=1 |
| **124** | 173 | Q63750\|RN | 69.92 | 36 | 36 | 0 | 2 | 2 | 2 | N | 17050 | 39S ribosomal protein L23, mitochondrial OS=Rattus norvegicus OX=10116 GN=Mrpl23 PE=2 SV=1 |
| **100** | 132 | tr\|D4A3E8 | 68.54 | 7 | 7 | 167400 | 3 | 3 | 3 | N | 47648 | Mitochondrial ribosomal protein S27 OS=Rattus norvegicus OX=10116 GN=Mrps27 PE=1 SV=1 |
| **129** | 187 | tr\|D3ZDP2 | 68.21 | 11 | 11 | 441290 | 2 | 2 | 2 | N | 23470 | Mitochondrial ribosomal protein L58 OS=Rattus norvegicus OX=10116 GN=Mrpl58 PE=1 SV=1 |
| **125** | 174 | tr\|D4A104 | 67.61 | 8 | 8 | 1016800 | 2 | 2 | 2 | N | 35440 | Mitochondrial ribosomal protein L45 OS=Rattus norvegicus OX=10116 GN=Mrpl45 PE=1 SV=1 |
| **253** | 253 | tr\|D3ZTW8 | 67.38 | 10 | 10 | 391700 | 1 | 1 | 1 | N | 15823 | Mitochondrial ribosomal protein L27 OS=Rattus norvegicus OX=10116 GN=Mrpl27 PE=1 SV=2 |
| **252** | 252 | tr\|D3ZLT1\| | 67.37 | 12 | 12 | 967070 | 1 | 1 | 1 | Y | 16568 | NADH dehydrogenase (Ubiquinone) 1 beta subcomplex, 7 (Predicted) OS=Rattus norvegicus OX=10116 GN=Ndufb7 PE=1 SV=1 |
| **254** | 254 | tr\|Q7H115 | 66.73 | 5 | 5 | 650340 | 1 | 1 | 1 | N | 29871 | Cytochrome c oxidase subunit 3 OS=Rattus norvegicus OX=10116 GN=Mt-co3 PE=3 SV=1 |
| **254** | 255 | tr\|I6V4L9\| | 66.73 | 5 | 5 | 650340 | 1 | 1 | 1 | N | 29844 | Cytochrome c oxidase subunit 3 OS=Rattus norvegicus OX=10116 GN=COX3 PE=3 SV=1 |
| **254** | 256 | tr\|Q8M7G | 66.73 | 5 | 5 | 650340 | 1 | 1 | 1 | N | 29861 | Cytochrome c oxidase subunit 3 OS=Rattus norvegicus OX=10116 PE=2 SV=1 |
| **254** | 257 | tr\|A0A096 | 66.73 | 5 | 5 | 650340 | 1 | 1 | 1 | N | 29901 | Cytochrome c oxidase subunit 3 OS=Rattus norvegicus OX=10116 GN=COX3 PE=3 SV=1 |
| **254** | 258 | P05505\|CC | 66.73 | 5 | 5 | 650340 | 1 | 1 | 1 | N | 29871 | Cytochrome c oxidase subunit 3 OS=Rattus norvegicus OX=10116 GN=Mtco3 PE=1 SV=5 |
| **254** | 259 | tr\|Q8SEZ2\| | 66.73 | 5 | 5 | 650340 | 1 | 1 | 1 | N | 29870 | Cytochrome c oxidase subunit 3 (Fragment) OS=Rattus norvegicus OX=10116 GN=COIII PE=3 SV=1 |
| **207** | 244 | Q66H15\|RI | 66.57 | 5 | 5 | 155180 | 1 | 1 | 1 | N | 52312 | Regulator of microtubule dynamics protein 3 OS=Rattus norvegicus OX=10116 GN=Rmdn3 PE=1 SV=1 |
| **126** | 176 | P08011\|M | 63.47 | 17 | 17 | 1226700 | 2 | 2 | 2 | N | 17472 | Microsomal glutathione S-transferase 1 OS=Rattus norvegicus OX=10116 GN=Mgst1 PE=1 SV=3 |

| | | | | | | | | | | | | |
|---|---|---|---|---|---|---|---|---|---|---|---|---|
| **126** | 177 | tr\|B6DYQ4 | 63.47 | 17 | 17 | 1226700 | 2 | 2 | 2 | N | 17472 | Microsomal glutathione S-transferase OS=Rattus norvegicus OX=10116 GN=Mgst1 PE=2 SV=1 |
| **116** | 160 | Q8VHF5\|CI | 63.15 | 5 | 5 | 1411300 | 2 | 2 | 2 | N | 51867 | Citrate synthase, mitochondrial OS=Rattus norvegicus OX=10116 GN=Cs PE=1 SV=1 |
| **116** | 161 | tr\|G3V936 | 63.15 | 5 | 5 | 1411300 | 2 | 2 | 2 | N | 51831 | Citrate synthase OS=Rattus norvegicus OX=10116 GN=Cs PE=1 SV=1 |
| **256** | 261 | P18665\|RM | 62.84 | 6 | 6 | 872600 | 1 | 1 | 1 | N | 38253 | 39S ribosomal protein L3, mitochondrial OS=Rattus norvegicus OX=10116 GN=Mrpl3 PE=2 SV=1 |
| **256** | 262 | tr\|G3V7P3 | 62.84 | 6 | 6 | 872600 | 1 | 1 | 1 | N | 38492 | 39S ribosomal protein L3, mitochondrial OS=Rattus norvegicus OX=10116 GN=Mrpl3 PE=1 SV=1 |
| **118** | 198 | tr\|A0A0G2 | 62.41 | 3 | 3 | 750110 | 2 | 2 | 2 | N | 67320 | Glycerol-3-phosphate dehydrogenase, mitochondrial OS=Rattus norvegicus OX=10116 GN=Gpd2 PE=1 SV=1 |
| **118** | 199 | tr\|A0A0G2 | 62.41 | 3 | 3 | 750110 | 2 | 2 | 2 | N | 69062 | Glycerol-3-phosphate dehydrogenase, mitochondrial OS=Rattus norvegicus OX=10116 GN=Gpd2 PE=1 SV=1 |
| **118** | 200 | P35571\|GF | 62.41 | 3 | 3 | 750110 | 2 | 2 | 2 | N | 80973 | Glycerol-3-phosphate dehydrogenase, mitochondrial OS=Rattus norvegicus OX=10116 GN=Gpd2 PE=1 SV=1 |
| **257** | 263 | tr\|B0BN02 | 61.12 | 4 | 4 | 1602800 | 1 | 1 | 1 | N | 48444 | Metaxin OS=Rattus norvegicus OX=10116 GN=Mtx1 PE=1 SV=1 |
| **127** | 180 | Q5PQN9\|R | 59.74 | 4 | 4 | 433170 | 2 | 2 | 2 | N | 44838 | 39S ribosomal protein L38, mitochondrial OS=Rattus norvegicus OX=10116 GN=Mrpl38 PE=2 SV=2 |
| **115** | 158 | P11240\|CC | 59.57 | 10 | 10 | 1140500 | 2 | 2 | 2 | N | 16130 | Cytochrome c oxidase subunit 5A, mitochondrial OS=Rattus norvegicus OX=10116 GN=Cox5a PE=1 SV=1 |
| **258** | 264 | B1WC61\|A | 57.09 | 4 | 4 | 247330 | 1 | 1 | 1 | N | 68843 | Complex I assembly factor ACAD9, mitochondrial OS=Rattus norvegicus OX=10116 GN=Acad9 PE=1 SV=1 |
| **259** | 265 | tr\|Q8SEZ1\| | 56.6 | 13 | 13 | 487810 | 1 | 1 | 1 | Y | 13087 | NADH-ubiquinone oxidoreductase chain 3 OS=Rattus norvegicus OX=10116 GN=Mt-nd3 PE=2 SV=1 |
| **259** | 266 | P05506\|NU | 56.6 | 13 | 13 | 487810 | 1 | 1 | 1 | Y | 13087 | NADH-ubiquinone oxidoreductase chain 3 OS=Rattus norvegicus OX=10116 GN=Mtnd3 PE=3 SV=3 |
| **259** | 267 | tr\|A0A0A1 | 56.6 | 13 | 13 | 487810 | 1 | 1 | 1 | Y | 13071 | NADH-ubiquinone oxidoreductase chain 3 OS=Rattus norvegicus OX=10116 GN=ND3 PE=3 SV=1 |
| **259** | 268 | tr\|Q06QG9 | 56.6 | 13 | 13 | 487810 | 1 | 1 | 1 | Y | 13075 | NADH-ubiquinone oxidoreductase chain 3 OS=Rattus norvegicus OX=10116 GN=ND3 PE=3 SV=1 |
| **260** | 269 | tr\|A0A0G2 | 55.93 | 4 | 4 | 401380 | 1 | 1 | 1 | N | 41754 | 3-ketoacyl-CoA thiolase, mitochondrial OS=Rattus norvegicus OX=10116 GN=Acaa2 PE=1 SV=1 |
| **260** | 270 | P13437\|TH | 55.93 | 4 | 4 | 401380 | 1 | 1 | 1 | N | 41871 | 3-ketoacyl-CoA thiolase, mitochondrial OS=Rattus norvegicus OX=10116 GN=Acaa2 PE=1 SV=1 |
| **260** | 271 | tr\|G3V9U2 | 55.93 | 4 | 4 | 401380 | 1 | 1 | 1 | N | 41885 | 3-ketoacyl-CoA thiolase, mitochondrial OS=Rattus norvegicus OX=10116 GN=Acaa2 PE=1 SV=3 |
| **128** | 185 | tr\|D3ZZ21\| | 55.56 | 12 | 12 | 1374600 | 2 | 2 | 2 | N | 15638 | NADH dehydrogenase (Ubiquinone) 1 beta subcomplex, 6 (Predicted) OS=Rattus norvegicus OX=10116 GN=Ndufb6 PE=1 SV=1 |
| **206** | 243 | tr\|G3V644 | 53.41 | 6 | 6 | 685530 | 1 | 1 | 1 | N | 49300 | NADH dehydrogenase (Ubiquinone) flavoprotein 3-like, isoform CRA_a OS=Rattus norvegicus OX=10116 GN=Ndufv3 PE=1 SV=1 |
| **206** | 272 | tr\|O54755 | 53.41 | 6 | 6 | 685530 | 1 | 1 | 1 | N | 49312 | MIPP65 OS=Rattus norvegicus OX=10116 GN=Ndufv3 PE=2 SV=1 |
| **261** | 273 | Q7TT47\|SP | 52.61 | 1 | 1 | 0 | 1 | 1 | 1 | N | 86103 | Paraplegin OS=Rattus norvegicus OX=10116 GN=Spg7 PE=2 SV=2 |
| **136** | 274 | P11951\|CX | 51.17 | 12 | 12 | 299720 | 1 | 1 | 2 | N | 8455 | Cytochrome c oxidase subunit 6C-2 OS=Rattus norvegicus OX=10116 GN=Cox6c2 PE=1 SV=3 |
| **169** | 179 | tr\|Q5XFW4 | 51 | 6 | 6 | 457390 | 1 | 1 | 1 | N | 20541 | Mitochondrial ribosomal protein L13 OS=Rattus norvegicus OX=10116 GN=Mrpl13 PE=1 SV=1 |
| **110** | 142 | tr\|B5DEY1\| | 50.73 | 3 | 3 | 396480 | 2 | 1 | 2 | N | 84183 | Afg3l1 protein (Fragment) OS=Rattus norvegicus OX=10116 GN=Afg3l1 PE=2 SV=1 |
| **110** | 143 | tr\|D3ZZ20\| | 50.73 | 3 | 3 | 396480 | 2 | 1 | 2 | N | 87203 | AFG3(ATPase family gene 3)-like 1 (S. cerevisiae) OS=Rattus norvegicus OX=10116 GN=Afg3l1 PE=1 SV=2 |
| **132** | 191 | tr\|B2RZD6 | 46.83 | 22 | 22 | 265170 | 2 | 2 | 2 | N | 9327 | NDUFA4, mitochondrial complex-associated OS=Rattus norvegicus OX=10116 GN=Ndufa4 PE=1 SV=1 |
| **262** | 275 | B0BN56\|RT | 44.82 | 7 | 7 | 531390 | 1 | 1 | 1 | N | 43962 | 28S ribosomal protein S31, mitochondrial OS=Rattus norvegicus OX=10116 GN=Mrps31 PE=2 SV=1 |
| **198** | 209 | P10818\|CX | 44.23 | 11 | 11 | 322910 | 1 | 1 | 1 | N | 12301 | Cytochrome c oxidase subunit 6A1, mitochondrial OS=Rattus norvegicus OX=10116 GN=Cox6a1 PE=1 SV=2 |
| **263** | 277 | P0CG51\|UI | 42.64 | 3 | 3 | 413390 | 1 | 1 | 1 | N | 34369 | Polyubiquitin-B OS=Rattus norvegicus OX=10116 GN=Ubb PE=1 SV=1 |
| **199** | 213 | tr\|A0A0G2 | 41.64 | 8 | 8 | 0 | 1 | 1 | 1 | N | 34662 | RCG47744, isoform CRA_c OS=Rattus norvegicus OX=10116 GN=LOC100911440 PE=1 SV=1 |
| **168** | 167 | P35434\|AT | 41.33 | 5 | 5 | 379080 | 1 | 1 | 1 | N | 17595 | ATP synthase subunit delta, mitochondrial OS=Rattus norvegicus OX=10116 GN=Atp5f1d PE=1 SV=2 |
| **168** | 168 | tr\|G3V7Y3 | 41.33 | 5 | 5 | 379080 | 1 | 1 | 1 | N | 17563 | ATP synthase subunit delta, mitochondrial OS=Rattus norvegicus OX=10116 GN=Atp5f1d PE=1 SV=1 |
| **264** | 278 | tr\|B5DFA3 | 41.15 | 2 | 2 | 195130 | 1 | 1 | 1 | N | 65357 | Bucs1 protein OS=Rattus norvegicus OX=10116 GN=Acsm1 PE=2 SV=1 |
| **264** | 279 | tr\|F1M1W1 | 41.15 | 2 | 2 | 195130 | 1 | 1 | 1 | N | 65347 | Acyl-coenzyme A synthetase ACSM2, mitochondrial OS=Rattus norvegicus OX=10116 GN=Acsm1 PE=1 SV=3 |
| **264** | 280 | tr\|A0A0G2 | 41.15 | 2 | 2 | 195130 | 1 | 1 | 1 | N | 65319 | Acyl-coenzyme A synthetase ACSM2, mitochondrial OS=Rattus norvegicus OX=10116 GN=Acsm1 PE=1 SV=1 |
| **264** | 281 | tr\|A0A0G2 | 41.15 | 2 | 2 | 195130 | 1 | 1 | 1 | N | 70943 | Acyl-coenzyme A synthetase ACSM2, mitochondrial OS=Rattus norvegicus OX=10116 GN=Acsm1 PE=1 SV=1 |
| **160** | 234 | tr\|Q06QK5 | 40.91 | 2 | 2 | 5054400 | 1 | 1 | 1 | N | 68606 | NADH-ubiquinone oxidoreductase chain 5 OS=Rattus norvegicus OX=10116 GN=ND5 PE=3 SV=1 |
| **160** | 235 | P11661\|NU | 40.91 | 2 | 2 | 5054400 | 1 | 1 | 1 | N | 68618 | NADH-ubiquinone oxidoreductase chain 5 OS=Rattus norvegicus OX=10116 GN=Mtnd5 PE=3 SV=3 |
| **160** | 236 | tr\|Q06QA1 | 40.91 | 2 | 2 | 5054400 | 1 | 1 | 1 | N | 68574 | NADH-ubiquinone oxidoreductase chain 5 OS=Rattus norvegicus OX=10116 GN=ND5 PE=3 SV=1 |
| **160** | 237 | tr\|Q8SEZ0\| | 40.91 | 2 | 2 | 5054400 | 1 | 1 | 1 | N | 68618 | NADH-ubiquinone oxidoreductase chain 5 OS=Rattus norvegicus OX=10116 GN=Mt-nd5 PE=3 SV=1 |
| **160** | 238 | tr\|A0A096X | 40.91 | 2 | 2 | 5054400 | 1 | 1 | 1 | N | 68584 | NADH-ubiquinone oxidoreductase chain 5 OS=Rattus norvegicus OX=10116 GN=ND5 PE=3 SV=1 |
| **160** | 239 | tr\|Q06QG6 | 40.91 | 2 | 2 | 5054400 | 1 | 1 | 1 | N | 68588 | NADH-ubiquinone oxidoreductase chain 5 OS=Rattus norvegicus OX=10116 GN=ND5 PE=3 SV=1 |
| **160** | 240 | tr\|A7XYD5 | 40.91 | 2 | 2 | 5054400 | 1 | 1 | 1 | N | 68856 | NADH-ubiquinone oxidoreductase chain 5 OS=Rattus norvegicus OX=10116 GN=Nd5 PE=3 SV=1 |
| **160** | 241 | tr\|A7XYC1\| | 40.91 | 2 | 2 | 5054400 | 1 | 1 | 1 | N | 68841 | NADH-ubiquinone oxidoreductase chain 5 OS=Rattus norvegicus OX=10116 GN=Nd5 PE=3 SV=1 |
| **160** | 242 | tr\|A0A0A1 | 40.91 | 2 | 2 | 5054400 | 1 | 1 | 1 | N | 68970 | NADH-ubiquinone oxidoreductase chain 5 OS=Rattus norvegicus OX=10116 GN=ND5 PE=3 SV=1 |
| **265** | 282 | P19804\|ND | 40.82 | 8 | 8 | 265680 | 1 | 1 | 1 | N | 17283 | Nucleoside diphosphate kinase B OS=Rattus norvegicus OX=10116 GN=Nme2 PE=1 SV=1 |
| **265** | 283 | Q05982\|NI | 40.82 | 8 | 8 | 265680 | 1 | 1 | 1 | N | 17193 | Nucleoside diphosphate kinase A OS=Rattus norvegicus OX=10116 GN=Nme1 PE=1 SV=1 |
| **203** | 226 | tr\|D3ZYT2\| | 39.85 | 3 | 3 | 602050 | 1 | 1 | 1 | N | 48136 | Mitochondrial ribosomal protein S5 OS=Rattus norvegicus OX=10116 GN=Mrps5 PE=1 SV=1 |
| **105** | 246 | O54902\|NI | 39.68 | 3 | 3 | 1023400 | 2 | 1 | 3 | N | 62277 | Natural resistance-associated macrophage protein 2 OS=Rattus norvegicus OX=10116 GN=Slc11a2 PE=1 SV=1 |
| **201** | 215 | tr\|D3ZXF8\| | 39.63 | 7 | 7 | 369480 | 1 | 1 | 1 | N | 17575 | Mitochondrial ribosomal protein L43 OS=Rattus norvegicus OX=10116 GN=Mrpl43 PE=1 SV=3 |
| **112** | 210 | tr\|D3ZGM1 | 39.28 | 3 | 3 | 535970 | 2 | 2 | 2 | N | 77891 | Pentatricopeptide repeat domain 3 OS=Rattus norvegicus OX=10116 GN=Ptcd3 PE=1 SV=1 |
| **112** | 211 | tr\|A0A0G2 | 39.28 | 3 | 3 | 535970 | 2 | 2 | 2 | N | 84557 | Pentatricopeptide repeat domain 3 OS=Rattus norvegicus OX=10116 GN=Ptcd3 PE=1 SV=1 |
| **106** | 162 | tr\|D3Z900\| | 39.12 | 5 | 5 | 981390 | 2 | 2 | 2 | N | 38239 | Mitochondrial amidoxime reducing component 2 OS=Rattus norvegicus OX=10116 GN=Marc2 PE=1 SV=2 |
| **106** | 163 | O88994\|M | 39.12 | 5 | 5 | 981390 | 2 | 2 | 2 | N | 38249 | Mitochondrial amidoxime reducing component 2 OS=Rattus norvegicus OX=10116 GN=Marc2 PE=2 SV=1 |
| **106** | 164 | tr\|M0R6N2 | 39.12 | 5 | 5 | 981390 | 2 | 2 | 2 | N | 38175 | MOSC domain-containing protein 2, mitochondrial-like OS=Rattus norvegicus OX=10116 GN=LOC100910481 PE=1 SV=1 |
| **266** | 284 | tr\|A0A0G2 | 38.87 | 34 | 34 | 0 | 1 | 1 | 1 | N | 10511 | Mitochondrial ribosomal protein S18B (Fragment) OS=Rattus norvegicus OX=10116 GN=Mrps18b PE=1 SV=1 |
| **266** | 285 | tr\|A0A0G2 | 38.87 | 21 | 21 | 0 | 1 | 1 | 1 | N | 17268 | Mitochondrial ribosomal protein S18B (Fragment) OS=Rattus norvegicus OX=10116 GN=Mrps18b PE=1 SV=4 |
| **266** | 286 | tr\|Q6MG10 | 38.87 | 12 | 12 | 0 | 1 | 1 | 1 | N | 28682 | Mitochondrial ribosomal protein S18B OS=Rattus norvegicus OX=10116 GN=Mrps18b PE=2 SV=1 |

| | | | | | | | | | | | | |
|---|---|---|---|---|---|---|---|---|---|---|---|---|
| **266** | 287 | tr\|F6Q5K7 | 38.87 | 12 | 12 | 0 | 1 | 1 | 1 | N | 29159 | Mitochondrial ribosomal protein S18B OS=Rattus norvegicus OX=10116 GN=Mrps18b PE=1 SV=1 |
| **268** | 289 | P80433\|CC | 38.82 | 10 | 10 | 263390 | 1 | 1 | 1 | N | 7672 | Cytochrome c oxidase subunit 8A, mitochondrial OS=Rattus norvegicus OX=10116 GN=Cox8a PE=1 SV=3 |
| **267** | 288 | tr\|B2RZ57\| | 38.01 | 5 | 5 | 353210 | 1 | 1 | 1 | N | 20651 | Mitochondrial ribosomal protein L18 OS=Rattus norvegicus OX=10116 GN=Mrpl18 PE=1 SV=1 |
| **271** | 293 | Q641X9\|R | 37.8 | 3 | 3 | 279950 | 1 | 1 | 1 | N | 30076 | 39S ribosomal protein L9, mitochondrial OS=Rattus norvegicus OX=10116 GN=Mrpl9 PE=2 SV=1 |
| **269** | 290 | tr\|M0RAM | 37.11 | 6 | 6 | 457190 | 1 | 1 | 1 | N | 22155 | Glutathione peroxidase OS=Rattus norvegicus OX=10116 GN=Gpx1 PE=1 SV=1 |
| **269** | 291 | P04041\|G | 37.11 | 6 | 6 | 457190 | 1 | 1 | 1 | N | 22305 | Glutathione peroxidase 1 OS=Rattus norvegicus OX=10116 GN=Gpx1 PE=1 SV=4 |
| **270** | 292 | Q66H47\|R | 36.78 | 4 | 4 | 366240 | 1 | 1 | 1 | N | 25001 | 39S ribosomal protein L24, mitochondrial OS=Rattus norvegicus OX=10116 GN=Mrpl24 PE=2 SV=1 |
| **108** | 231 | P70705\|AT | 35.7 | 2 | 2 | 1513600 | 2 | 2 | 2 | Y | 162092 | Copper-transporting ATPase 1 OS=Rattus norvegicus OX=10116 GN=Atp7a PE=1 SV=1 |
| **202** | 224 | Q6AYC2\|IR | 35.1 | 2 | 2 | 526220 | 1 | 1 | 1 | N | 46338 | Immunity-related GTPase family M protein OS=Rattus norvegicus OX=10116 GN=Irgm PE=2 SV=1 |
| **121** | 365 | tr\|G3V8G1 | 34.64 | 1 | 1 | 2557800 | 2 | 1 | 2 | Y | 137113 | ATP/GTP binding protein 1 (Predicted), isoform CRA_a OS=Rattus norvegicus OX=10116 GN=Agtpbp1 PE=4 SV=1 |
| **272** | 296 | P19527\|N | 34.55 | 3 | 3 | 311400 | 1 | 1 | 1 | Y | 61335 | Neurofilament light polypeptide OS=Rattus norvegicus OX=10116 GN=Nefl PE=1 SV=3 |
| **273** | 297 | tr\|A9UMW | 33.76 | 13 | 13 | 552160 | 1 | 1 | 1 | N | 9255 | Ndufa3 protein (Fragment) OS=Rattus norvegicus OX=10116 GN=Ndufa3 PE=2 SV=1 |
| **273** | 298 | tr\|M0RB63 | 33.76 | 13 | 13 | 552160 | 1 | 1 | 1 | N | 9372 | RCG63041 OS=Rattus norvegicus OX=10116 GN=LOC684509 PE=4 SV=1 |
| **273** | 299 | tr\|A0A0G2 | 33.76 | 12 | 12 | 552160 | 1 | 1 | 1 | N | 10170 | NADH:ubiquinone oxidoreductase subunit A3 OS=Rattus norvegicus OX=10116 GN=Ndufa3 PE=1 SV=1 |
| **275** | 302 | tr\|D4A4P3 | 32.64 | 8 | 8 | 641070 | 1 | 1 | 1 | N | 11267 | NADH:ubiquinone oxidoreductase subunit B3 OS=Rattus norvegicus OX=10116 GN=Ndufb3 PE=1 SV=1 |
| **274** | 301 | tr\|B2RZ24\| | 32.64 | 2 | 2 | 763930 | 1 | 1 | 1 | N | 47388 | Succinate-CoA ligase subunit beta (Fragment) OS=Rattus norvegicus OX=10116 GN=Sucla2 PE=2 SV=1 |
| **274** | 303 | tr\|F1LM47 | 32.64 | 2 | 2 | 763930 | 1 | 1 | 1 | N | 50306 | Succinate--CoA ligase [ADP-forming] subunit beta, mitochondrial OS=Rattus norvegicus OX=10116 GN=Sucla2 PE=1 SV=1 |
| **142** | 468 | Q63704\|C | 30.76 | 1 | 1 | 0 | 1 | 1 | 2 | N | 88217 | Carnitine O-palmitoyltransferase 1, muscle isoform OS=Rattus norvegicus OX=10116 GN=Cpt1b PE=1 SV=1 |
| **142** | 489 | F1LN46\|CP | 30.76 | 1 | 1 | 0 | 1 | 1 | 2 | N | 90170 | Carnitine O-palmitoyltransferase 1, brain isoform OS=Rattus norvegicus OX=10116 GN=Cpt1c PE=1 SV=1 |
| **142** | 533 | tr\|M0RAQ | 30.76 | 1 | 1 | 0 | 1 | 1 | 2 | N | 99557 | Hexokinase-1 OS=Rattus norvegicus OX=10116 GN=Hk1 PE=1 SV=2 |
| **142** | 549 | P05708\|H | 30.76 | 1 | 1 | 0 | 1 | 1 | 2 | N | 102408 | Hexokinase-1 OS=Rattus norvegicus OX=10116 GN=Hk1 PE=1 SV=4 |
| **159** | 233 | Q5U2U0\|C | 30.15 | 2 | 2 | 0 | 1 | 1 | 1 | N | 69208 | ATP-dependent Clp protease ATP-binding subunit clpX-like, mitochondrial OS=Rattus norvegicus OX=10116 GN=Clpx PE=2 SV=1 |
| **211** | 310 | tr\|Q5BJZ3\| | 29.97 | 1 | 1 | 715240 | 1 | 1 | 1 | Y | 113869 | Nicotinamide nucleotide transhydrogenase OS=Rattus norvegicus OX=10116 GN=Nnt PE=1 SV=1 |
| **140** | 366 | P32089\|TX | 29.18 | 4 | 4 | 307380 | 2 | 1 | 2 | N | 33835 | Tricarboxylate transport protein, mitochondrial OS=Rattus norvegicus OX=10116 GN=Slc25a1 PE=1 SV=1 |
| **277** | 320 | tr\|D3ZYL4\| | 26.81 | 7 | 7 | 0 | 1 | 1 | 1 | N | 18319 | Mitochondrial ribosomal protein L50 OS=Rattus norvegicus OX=10116 GN=Mrpl50 PE=1 SV=1 |
| **212** | 311 | tr\|D3ZK81\| | 26.75 | 4 | 4 | 0 | 1 | 1 | 1 | Y | 34694 | Mitochondrial rRNA methyltransferase 1 OS=Rattus norvegicus OX=10116 GN=Mrm1 PE=4 SV=1 |
| **278** | 328 | Q5M9G9\| | 25.59 | 2 | 2 | 287680 | 1 | 1 | 1 | Y | 71181 | FAST kinase domain-containing protein 4 OS=Rattus norvegicus OX=10116 GN=Tbrg4 PE=2 SV=1 |
| **279** | 330 | tr\|D3ZXQ6 | 25.26 | 3 | 3 | 0 | 1 | 1 | 1 | N | 46897 | LETM1 domain-containing protein LETM2, mitochondrial OS=Rattus norvegicus OX=10116 GN=Letm2 PE=4 SV=3 |
| **279** | 331 | Q5PQQ5\|L | 25.26 | 3 | 3 | 0 | 1 | 1 | 1 | N | 52524 | LETM1 domain-containing protein LETM2, mitochondrial OS=Rattus norvegicus OX=10116 GN=Letm2 PE=2 SV=1 |
| **279** | 332 | tr\|F1LP77\| | 25.26 | 3 | 3 | 0 | 1 | 1 | 1 | N | 52501 | LETM1 domain-containing protein LETM2, mitochondrial OS=Rattus norvegicus OX=10116 GN=Letm2 PE=4 SV=3 |
| **143** | 416 | tr\|F1M953 | 24.52 | 4 | 4 | 2410700 | 2 | 2 | 2 | Y | 73745 | Stress-70 protein, mitochondrial OS=Rattus norvegicus OX=10116 GN=Hspa9 PE=1 SV=1 |
| **143** | 417 | P48721\|G | 24.52 | 4 | 4 | 2410700 | 2 | 2 | 2 | Y | 73858 | Stress-70 protein, mitochondrial OS=Rattus norvegicus OX=10116 GN=Hspa9 PE=1 SV=3 |
| **162** | 321 | tr\|F1LNF0\| | 24.48 | 0 | 0 | 851370 | 1 | 1 | 1 | Y | 228912 | Myosin heavy chain 14 OS=Rattus norvegicus OX=10116 GN=Myh14 PE=1 SV=1 |
| **284** | 339 | tr\|G3V783 | 24.май | 1 | 1 | 505150 | 1 | 1 | 1 | N | 60778 | Receptor-interacting serine/threonine-protein kinase 2 OS=Rattus norvegicus OX=10116 GN=Ripk2 PE=1 SV=1 |
| **286** | 346 | tr\|A0A0G2 | 24.фев | 4 | 4 | 442780 | 1 | 1 | 1 | N | 18468 | Uncharacterized protein OS=Rattus norvegicus OX=10116 PE=3 SV=1 |
| **280** | 334 | P0DMW0\| | 23.91 | 2 | 2 | 301130 | 1 | 1 | 1 | Y | 70185 | Heat shock 70 kDa protein 1A OS=Rattus norvegicus OX=10116 GN=Hspa1a PE=2 SV=1 |
| **280** | 335 | P0DMW1\| | 23.91 | 2 | 2 | 301130 | 1 | 1 | 1 | Y | 70185 | Heat shock 70 kDa protein 1B OS=Rattus norvegicus OX=10116 GN=Hspa1b PE=2 SV=1 |
| **281** | 336 | Q5FVP8\|D | 23.89 | 3 | 3 | 486090 | 1 | 1 | 1 | Y | 43795 | Diacylglycerol O-acyltransferase 2 OS=Rattus norvegicus OX=10116 GN=Dgat2 PE=2 SV=1 |
| **210** | 308 | P03957\|M | 23.83 | 3 | 3 | 0 | 1 | 1 | 1 | Y | 53428 | Stromelysin-1 OS=Rattus norvegicus OX=10116 GN=Mmp3 PE=1 SV=1 |
| **120** | 349 | tr\|A0A0G2 | 23.53 | 1 | 1 | 443580 | 1 | 1 | 2 | N | 112677 | Isoleucyl-tRNA synthetase 2, mitochondrial OS=Rattus norvegicus OX=10116 GN=Iars2 PE=1 SV=1 |
| **287** | 363 | tr\|D4A5E7 | 22.94 | 2 | 2 | 1124800 | 1 | 1 | 1 | Y | 80721 | Inhibitor of kappaB kinase epsilon (Predicted) OS=Rattus norvegicus OX=10116 GN=Ikbke PE=1 SV=1 |
| **179** | 358 | P51111\|H | 22.35 | 0 | 0 | 386830 | 1 | 1 | 1 | Y | 343761 | Huntingtin OS=Rattus norvegicus OX=10116 GN=Htt PE=1 SV=1 |
| **179** | 359 | tr\|G3V9P7 | 22.35 | 0 | 0 | 386830 | 1 | 1 | 1 | Y | 344745 | Huntingtin OS=Rattus norvegicus OX=10116 GN=Htt PE=1 SV=1 |
| **180** | 373 | Q641Y8\|D | 22.18 | 1 | 1 | 316030 | 1 | 1 | 1 | N | 82497 | ATP-dependent RNA helicase DDX1 OS=Rattus norvegicus OX=10116 GN=Ddx1 PE=1 SV=1 |
| **290** | 372 | tr\|F1LNJ1\| | 22.фев | 1 | 1 | 526780 | 1 | 1 | 1 | Y | 284829 | Leucine-rich repeat kinase 2 OS=Rattus norvegicus OX=10116 GN=Lrrk2 PE=1 SV=3 |
| **306** | 412 | P48037\|A | 21.77 | 1 | 1 | 193960 | 1 | 1 | 1 | N | 75754 | Annexin A6 OS=Rattus norvegicus OX=10116 GN=Anxa6 PE=1 SV=2 |
| **306** | 413 | tr\|Q6IMZ3 | 21.77 | 1 | 1 | 193960 | 1 | 1 | 1 | N | 75756 | Annexin OS=Rattus norvegicus OX=10116 GN=Anxa6 PE=1 SV=1 |
| **122** | 377 | tr\|G3V827 | 21.66 | 2 | 2 | 1387700 | 1 | 1 | 2 | N | 51581 | Cysteine conjugate-beta lyase 1, isoform CRA_a OS=Rattus norvegicus OX=10116 GN=Kyat1 PE=1 SV=1 |
| **122** | 378 | Q08415\|K | 21.66 | 2 | 2 | 1387700 | 1 | 1 | 2 | N | 51653 | Kynurenine--oxoglutarate transaminase 1, mitochondrial OS=Rattus norvegicus OX=10116 GN=Kyat1 PE=1 SV=1 |
| **304** | 405 | P29476\|N | 21.59 | 0 | 0 | 406760 | 1 | 1 | 1 | Y | 160559 | Nitric oxide synthase, brain OS=Rattus norvegicus OX=10116 GN=Nos1 PE=1 SV=1 |
| **304** | 406 | tr\|D3ZEW7 | 21.59 | 0 | 0 | 406760 | 1 | 1 | 1 | Y | 164329 | Nitric oxide synthase, brain OS=Rattus norvegicus OX=10116 GN=Nos1 PE=1 SV=2 |
| **292** | 380 | P63088\|PP | 21.34 | 5 | 5 | 463420 | 1 | 1 | 1 | Y | 36984 | Serine/threonine-protein phosphatase PP1-gamma catalytic subunit OS=Rattus norvegicus OX=10116 GN=Ppp1cc PE=1 SV=1 |
| **292** | 381 | tr\|Q6AYZ4 | 21.34 | 4 | 4 | 463420 | 1 | 1 | 1 | Y | 38504 | Serine/threonine-protein phosphatase OS=Rattus norvegicus OX=10116 GN=Ppp1cc PE=2 SV=1 |
| **295** | 384 | tr\|D4A7Q3 | 21.23 | 8 | 8 | 6737900 | 1 | 1 | 1 | N | 11813 | RCG43539, isoform CRA_a OS=Rattus norvegicus OX=10116 GN=Sdhaf4 PE=1 SV=2 |
| **215** | 348 | tr\|B2GUZ6 | 21.21 | 2 | 2 | 810750 | 1 | 1 | 1 | Y | 43606 | Reticulon 4-interacting protein 1 OS=Rattus norvegicus OX=10116 GN=Rtn4ip1 PE=1 SV=1 |
| **297** | 389 | tr\|F1LVW9 | 20.97 | 2 | 2 | 1946800 | 1 | 1 | 1 | Y | 64260 | 5-aminolevulinate synthase OS=Rattus norvegicus OX=10116 GN=Alas2 PE=3 SV=3 |
| **297** | 390 | Q63147\|H | 20.97 | 2 | 2 | 1946800 | 1 | 1 | 1 | Y | 64842 | 5-aminolevulinate synthase, erythroid-specific, mitochondrial OS=Rattus norvegicus OX=10116 GN=Alas2 PE=1 SV=1 |
| **297** | 391 | tr\|Q642C1 | 20.97 | 2 | 2 | 1946800 | 1 | 1 | 1 | Y | 64870 | 5-aminolevulinate synthase OS=Rattus norvegicus OX=10116 GN=Alas2 PE=2 SV=1 |
| **178** | 354 | tr\|B5DFK5 | 20.52 | 1 | 1 | 4349500 | 1 | 1 | 1 | Y | 119424 | Hip1r protein OS=Rattus norvegicus OX=10116 GN=Hip1r PE=2 SV=1 |
| **178** | 355 | tr\|F1LML7 | 20.52 | 1 | 1 | 4349500 | 1 | 1 | 1 | Y | 119553 | Huntingtin-interacting protein 1-related OS=Rattus norvegicus OX=10116 GN=Hip1r PE=1 SV=1 |

| | | | | | | | | | | | | |
|---|---|---|---|---|---|---|---|---|---|---|---|---|
| **178** | 356 | tr\|Q99PW9 | 20.52 | 1 | 1 | 4349500 | 1 | 1 | 1 | Y | 120572 | Huntingtin interacting protein 1 related (Fragment) OS=Rattus norvegicus OX=10116 GN=Hip1r PE=2 SV=1 |
| **303** | 403 | tr\|D4A676 | 20.39 | 4 | 4 | 462420 | 1 | 1 | 1 | N | 24898 | GrpE protein homolog OS=Rattus norvegicus OX=10116 GN=Grpel2 PE=3 SV=1 |
| **305** | 409 | P56574\|IDI | 20.29 | 2 | 2 | 989050 | 1 | 1 | 1 | Y | 50967 | Isocitrate dehydrogenase [NADP], mitochondrial OS=Rattus norvegicus OX=10116 GN=Idh2 PE=1 SV=2 |
| **total 300 proteins** | | | | | | | | | | | | |
