## Supplementary material for "Permeability transition pore-related changes in the proteome and channel activity of ATP synthase dimers and monomers": RLM KCl-BM

| rotein Grou | Protein ID | Accession | -10lgP | overage (% | age (%) Sa | IBAQ | #Peptides | #Unique | Spec Sample | PTM | Avg. Mass | Description |
|---|---|---|---|---|---|---|---|---|---|---|---|---|
| **2** | 13 | P15999\|ATPA_RA | 460.42 | 70 | 70 | 1.42E+09 | 75 | 75 | 245 | Y | 59754 | ATP synthase subunit alpha, mitochondrial OS=Rattus norvegicus OX=10116 GN=Atp5f1a PE=1 SV=2 |
| **1** | 8 | P10719\|ATPB_RA | 449.64 | 81 | 81 | 1.69E+09 | 64 | 64 | 308 | Y | 56354 | ATP synthase subunit beta, mitochondrial OS=Rattus norvegicus OX=10116 GN=Atp5f1b PE=1 SV=2 |
| **1** | 9 | tr\|G3V6D3\|G3V6 | 449.64 | 81 | 81 | 1.69E+09 | 64 | 64 | 308 | Y | 56345 | ATP synthase subunit beta OS=Rattus norvegicus OX=10116 GN=Atp5f1b PE=1 SV=1 |
| **6** | 3 | P07756\|CPSM_RA | 355.54 | 32 | 32 | 1.07E+08 | 37 | 37 | 47 | Y | 164579 | Carbamoyl-phosphate synthase [ammonia], mitochondrial OS=Rattus norvegicus OX=10116 GN=Cps1 PE=1 SV=1 |
| **8** | 10 | P22791\|HMCS2_ | 330.14 | 44 | 44 | 8.71E+07 | 27 | 27 | 32 | Y | 56912 | Hydroxymethylglutaryl-CoA synthase, mitochondrial OS=Rattus norvegicus OX=10116 GN=Hmgcs2 PE=1 SV=1 |
| **8** | 11 | tr\|Q68G44\|Q68G | 330.14 | 44 | 44 | 8.71E+07 | 27 | 27 | 32 | Y | 56886 | 3-hydroxy-3-methylglutaryl coenzyme A synthase OS=Rattus norvegicus OX=10116 GN=Hmgcs2 PE=1 SV=1 |
| **16** | 2 | P10860\|DHE3_RA | 319.64 | 36 | 36 | 6.13E+07 | 19 | 19 | 24 | Y | 61416 | Glutamate dehydrogenase 1, mitochondrial OS=Rattus norvegicus OX=10116 GN=Glud1 PE=1 SV=2 |
| **10** | 5 | Q64428\|ECHA_RA | 315.18 | 36 | 36 | 3.64E+07 | 26 | 25 | 30 | Y | 82665 | Trifunctional enzyme subunit alpha, mitochondrial OS=Rattus norvegicus OX=10116 GN=Hadha PE=1 SV=2 |
| **5** | 16 | P67779\|PHB_RAT | 293.28 | 85 | 85 | 2.99E+08 | 30 | 29 | 57 | Y | 29820 | Prohibitin OS=Rattus norvegicus OX=10116 GN=Phb PE=1 SV=1 |
| **7** | 97 | P35435\|ATPG_RA | 283.88 | 65 | 65 | 1.64E+07 | 19 | 1 | 37 | Y | 30191 | ATP synthase subunit gamma, mitochondrial OS=Rattus norvegicus OX=10116 GN=Atp5f1c PE=1 SV=2 |
| **11** | 95 | tr\|Q6QI09\|Q6QI0 | 277.73 | 30 | 30 | 1.45E+07 | 20 | 2 | 29 | Y | 67721 | ATP synthase subunit gamma, mitochondrial OS=Rattus norvegicus OX=10116 GN=Taf3 PE=1 SV=1 |
| **3** | 18 | Q5XIH7\|PHB2_RA | 275.42 | 75 | 75 | 4.40E+06 | 30 | 1 | 62 | Y | 33312 | Prohibitin-2 OS=Rattus norvegicus OX=10116 GN=Phb2 PE=1 SV=1 |
| **9** | 114 | P19511\|AT5F1_R | 275.26 | 55 | 55 | 2.51E+08 | 19 | 19 | 31 | Y | 28869 | ATP synthase F(0) complex subunit B1, mitochondrial OS=Rattus norvegicus OX=10116 GN=Atp5pb PE=1 SV=1 |
| **4** | 19 | tr\|A0A0G2KB63\| | 274.47 | 76 | 76 | 2.53E+06 | 30 | 1 | 61 | Y | 33168 | Prohibitin OS=Rattus norvegicus OX=10116 GN=Phb2 PE=1 SV=1 |
| **13** | 204 | P31399\|ATP5H_R | 266.61 | 80 | 80 | 1.16E+08 | 16 | 16 | 26 | Y | 18763 | ATP synthase subunit d, mitochondrial OS=Rattus norvegicus OX=10116 GN=Atp5pd PE=1 SV=3 |
| **17** | 168 | tr\|G3V7Y3\|G3V7 | 258.55 | 61 | 61 | 7.66E+07 | 11 | 11 | 15 | Y | 17563 | ATP synthase subunit delta, mitochondrial OS=Rattus norvegicus OX=10116 GN=Atp5f1d PE=1 SV=1 |
| **14** | 6 | P52873\|PYC_RAT | 258 | 20 | 20 | 3.40E+07 | 20 | 20 | 25 | Y | 129777 | Pyruvate carboxylase, mitochondrial OS=Rattus norvegicus OX=10116 GN=Pc PE=1 SV=2 |
| **14** | 7 | tr\|A0A0G2JTL5\|A | 258 | 19 | 19 | 3.40E+07 | 20 | 20 | 25 | Y | 140005 | Pyruvate carboxylase, mitochondrial OS=Rattus norvegicus OX=10116 GN=Pc PE=1 SV=1 |
| **12** | 188 | Q06647\|ATPO_RA | 245.52 | 65 | 65 | 2.18E+08 | 22 | 22 | 28 | Y | 23398 | ATP synthase subunit O, mitochondrial OS=Rattus norvegicus OX=10116 GN=Atp5po PE=1 SV=1 |
| **19** | 12 | Q66HF1\|NDUS1_ | 242.11 | 22 | 22 | 3.25E+07 | 13 | 13 | 14 | N | 79412 | NADH-ubiquinone oxidoreductase 75 kDa subunit, mitochondrial OS=Rattus norvegicus OX=10116 GN=Ndufs1 PE=1 SV=1 |
| **15** | 4 | P00507\|AATM_R | 238.5 | 39 | 39 | 8.32E+07 | 18 | 18 | 25 | Y | 47314 | Aspartate aminotransferase, mitochondrial OS=Rattus norvegicus OX=10116 GN=Got2 PE=1 SV=2 |
| **20** | 28 | P18163\|ACSL1_R | 213.5 | 15 | 15 | 1.02E+07 | 11 | 8 | 12 | N | 78179 | Long-chain-fatty-acid--CoA ligase 1 OS=Rattus norvegicus OX=10116 GN=Acsl1 PE=1 SV=1 |
| **23** | 48 | Q9WVK3\|PECR_R | 204.95 | 37 | 37 | 1.19E+07 | 8 | 8 | 10 | Y | 32433 | Peroxisomal trans-2-enoyl-CoA reductase OS=Rattus norvegicus OX=10116 GN=Pecr PE=2 SV=1 |
| **23** | 49 | tr\|A0A0G2JVG4\| | 204.95 | 36 | 36 | 1.19E+07 | 8 | 8 | 10 | Y | 32737 | Peroxisomal trans-2-enoyl-CoA reductase OS=Rattus norvegicus OX=10116 GN=Pecr PE=1 SV=1 |
| **34** | 54 | tr\|D4A0T0\|D4A0 | 200.87 | 28 | 28 | 6.54E+06 | 5 | 5 | 5 | Y | 20859 | NADH:ubiquinone oxidoreductase subunit B10 OS=Rattus norvegicus OX=10116 GN=Ndufb10 PE=1 SV=1 |
| **22** | 14 | Q5BK63\|NDUA9_ | 196.06 | 27 | 27 | 8.45E+06 | 9 | 9 | 10 | Y | 42559 | NADH dehydrogenase [ubiquinone] 1 alpha subcomplex subunit 9, mitochondrial OS=Rattus norvegicus OX=10116 GN=Ndufa9 PE=1 SV=2 |
| **24** | 32 | P13086\|SUCA_RA | 193.77 | 17 | 17 | 8.04E+06 | 5 | 5 | 8 | Y | 36148 | Succinate--CoA ligase [ADP/GDP-forming] subunit alpha, mitochondrial OS=Rattus norvegicus OX=10116 GN=Suclg1 PE=2 SV=2 |
| **24** | 33 | tr\|A0A0H2UHE1\| | 193.77 | 16 | 16 | 8.04E+06 | 5 | 5 | 8 | Y | 37560 | Succinate--CoA ligase [ADP/GDP-forming] subunit alpha, mitochondrial OS=Rattus norvegicus OX=10116 GN=Suclg1 PE=1 SV=1 |
| **40** | 36 | tr\|D3ZG43\|D3ZG | 185.13 | 21 | 21 | 1.41E+07 | 4 | 4 | 4 | N | 30226 | NADH dehydrogenase (Ubiquinone) Fe-S protein 3 (Predicted), isoform CRA_c OS=Rattus norvegicus OX=10116 GN=Ndufs3 PE=1 SV=1 |
| **21** | 31 | Q60587\|ECHB_RA | 176.17 | 22 | 22 | 1.35E+07 | 9 | 9 | 11 | Y | 51414 | Trifunctional enzyme subunit beta, mitochondrial OS=Rattus norvegicus OX=10116 GN=Hadhb PE=1 SV=1 |
| **37** | 47 | P0C2X9\|AL4A1_R | 171.46 | 9 | 9 | 3.73E+06 | 4 | 4 | 4 | N | 61869 | Delta-1-pyrroline-5-carboxylate dehydrogenase, mitochondrial OS=Rattus norvegicus OX=10116 GN=Aldh4a1 PE=1 SV=1 |
| **27** | 55 | P24329\|THTR_RA | 169.31 | 23 | 23 | 7.44E+06 | 5 | 5 | 6 | N | 33407 | Thiosulfate sulfurtransferase OS=Rattus norvegicus OX=10116 GN=Tst PE=1 SV=3 |
| **44** | 93 | tr\|D4A565\|D4A5 | 167.21 | 28 | 28 | 4.24E+06 | 4 | 4 | 4 | N | 21664 | NADH dehydrogenase (Ubiquinone) 1 beta subcomplex, 5 (Predicted), isoform CRA_b OS=Rattus norvegicus OX=10116 GN=Ndufb5 PE=1 SV=1 |
| **41** | 46 | P17764\|THIL_RAT | 162.52 | 12 | 12 | 2.19E+06 | 4 | 4 | 4 | N | 44695 | Acetyl-CoA acetyltransferase, mitochondrial OS=Rattus norvegicus OX=10116 GN=Acat1 PE=1 SV=1 |
| **32** | 39 | P29147\|BDH_RAT | 155.31 | 15 | 15 | 6.00E+06 | 5 | 5 | 5 | N | 38202 | D-beta-hydroxybutyrate dehydrogenase, mitochondrial OS=Rattus norvegicus OX=10116 GN=Bdh1 PE=1 SV=2 |
| **32** | 40 | tr\|A0A0G2JSH2\|A | 155.31 | 15 | 15 | 6.00E+06 | 5 | 5 | 5 | N | 38333 | 3-hydroxybutyrate dehydrogenase, type 1, isoform CRA_a OS=Rattus norvegicus OX=10116 GN=Bdh1 PE=1 SV=1 |
| **55** | 64 | Q02253\|MMSA_R | 153.04 | 10 | 10 | 3458800 | 3 | 3 | 3 | N | 57808 | Methylmalonate-semialdehyde dehydrogenase [acylating], mitochondrial OS=Rattus norvegicus OX=10116 GN=Aldh6a1 PE=1 SV=1 |
| **55** | 65 | tr\|G3V7J0\|G3V7J | 153.04 | 10 | 10 | 3.46E+06 | 3 | 3 | 3 | N | 57748 | Aldehyde dehydrogenase family 6, subfamily A1, isoform CRA_b OS=Rattus norvegicus OX=10116 GN=Aldh6a1 PE=1 SV=1 |
| **26** | 21 | tr\|Q5XIH3\|Q5XIH | 151.14 | 18 | 18 | 4.73E+06 | 7 | 7 | 7 | N | 50731 | NADH dehydrogenase [ubiquinone] flavoprotein 1, mitochondrial OS=Rattus norvegicus OX=10116 GN=Ndufv1 PE=1 SV=1 |
| **30** | 27 | P85834\|EFTU_RA | 145 | 13 | 13 | 4.04E+06 | 5 | 5 | 5 | N | 49522 | Elongation factor Tu, mitochondrial OS=Rattus norvegicus OX=10116 GN=Tufm PE=1 SV=1 |
| **29** | 61 | P63039\|CH60_RA | 139.8 | 7 | 7 | 1.90E+07 | 4 | 4 | 5 | N | 60956 | 60 kDa heat shock protein, mitochondrial OS=Rattus norvegicus OX=10116 GN=Hspd1 PE=1 SV=1 |
| **29** | 62 | tr\|A0A482IDN3\|A | 139.8 | 7 | 7 | 1.90E+07 | 4 | 4 | 5 | N | 60956 | Hsp60 OS=Rattus norvegicus OX=10116 GN=Hspd1 PE=2 SV=1 |
| **90** | 4499 | Q9JJW3\|ATPMD_ | 139.63 | 28 | 28 | 1.03E+07 | 2 | 2 | 2 | N | 6408 | ATP synthase membrane subunit DAPIT, mitochondrial OS=Rattus norvegicus OX=10116 GN=Atp5md PE=1 SV=1 |
| **35** | 4490 | Q6PDU7\|ATP5L_R | 133.19 | 34 | 34 | 1.10E+07 | 2 | 2 | 5 | Y | 11433 | ATP synthase subunit g, mitochondrial OS=Rattus norvegicus OX=10116 GN=Atp5mg PE=1 SV=2 |
| **58** | 35 | tr\|Q6P6W6\|Q6P6 | 128.01 | 20 | 20 | 1.17E+06 | 3 | 3 | 3 | N | 31701 | NADH dehydrogenase [ubiquinone] 1 alpha subcomplex subunit 10, mitochondrial OS=Rattus norvegicus OX=10116 GN=Ndufa10 PE=2 SV=1 |
| **58** | 23 | Q561S0\|NDUAA_ | 128.01 | 15 | 15 | 1.17E+06 | 3 | 3 | 3 | N | 40493 | NADH dehydrogenase [ubiquinone] 1 alpha subcomplex subunit 10, mitochondrial OS=Rattus norvegicus OX=10116 GN=Ndufa10 PE=1 SV=1 |
| **58** | 26 | tr\|A0A1W2Q6F8\| | 128.01 | 15 | 15 | 1.17E+06 | 3 | 3 | 3 | N | 40544 | NADH dehydrogenase [ubiquinone] 1 alpha subcomplex subunit 10, mitochondrial OS=Rattus norvegicus OX=10116 GN=Ndufa10l1 PE=3 SV=1 |
| **33** | 58 | P19234\|NDUV2_ | 126.7 | 24 | 24 | 1.48E+07 | 4 | 4 | 5 | N | 27378 | NADH dehydrogenase [ubiquinone] flavoprotein 2, mitochondrial OS=Rattus norvegicus OX=10116 GN=Ndufv2 PE=1 SV=2 |
| **42** | 50 | P10888\|COX41_R | 126.18 | 24 | 24 | 4.13E+06 | 4 | 4 | 4 | N | 19515 | Cytochrome c oxidase subunit 4 isoform 1, mitochondrial OS=Rattus norvegicus OX=10116 GN=Cox4i1 PE=1 SV=1 |
| **61** | 4494 | D3Z9R8\|ATP68_R | 123.36 | 32 | 32 | 4.40E+06 | 3 | 3 | 3 | Y | 6914 | ATP synthase subunit ATP5MPL, mitochondrial OS=Rattus norvegicus OX=10116 GN=Atp5mpl PE=1 SV=1 |
| **49** | 44 | P16970\|ABCD3_R | 122.68 | 5 | 5 | 1.97E+06 | 2 | 2 | 3 | N | 75316 | ATP-binding cassette sub-family D member 3 OS=Rattus norvegicus OX=10116 GN=Abcd3 PE=1 SV=3 |
| **48** | 76 | O70351\|HCD2_RA | 121.91 | 16 | 16 | 2.75E+06 | 3 | 3 | 3 | N | 27246 | 3-hydroxyacyl-CoA dehydrogenase type-2 OS=Rattus norvegicus OX=10116 GN=Hsd17b10 PE=1 SV=3 |
| **48** | 77 | tr\|B0BMW2\|B0B | 121.91 | 16 | 16 | 2.75E+06 | 3 | 3 | 3 | N | 27250 | 3-hydroxyacyl-CoA dehydrogenase type-2 OS=Rattus norvegicus OX=10116 GN=Hsd17b10 PE=1 SV=1 |
| **56** | 105 | tr\|D3ZF13\|D3ZF1 | 118.09 | 15 | 15 | 3.26E+06 | 3 | 3 | 3 | N | 17514 | Acyl carrier protein OS=Rattus norvegicus OX=10116 GN=Ndufab1 PE=1 SV=1 |
| **71** | 24 | Q641Y2\|NDUS2_ | 108.84 | 5 | 5 | 4.16E+06 | 2 | 2 | 2 | N | 52562 | NADH dehydrogenase [ubiquinone] iron-sulfur protein 2, mitochondrial OS=Rattus norvegicus OX=10116 GN=Ndufs2 PE=1 SV=1 |
| **85** | 73 | tr\|B2RYS8\|B2RYS | 106.22 | 15 | 15 | 1.99E+06 | 2 | 2 | 2 | N | 21959 | NADH dehydrogenase [ubiquinone] 1 beta subcomplex subunit 8, mitochondrial OS=Rattus norvegicus OX=10116 GN=Ndufb8 PE=1 SV=1 |
| **45** | 4487 | tr\|Q5UAJ5\|Q5UA | 105.48 | 46 | 46 | 2.69E+07 | 3 | 3 | 4 | N | 7642 | ATP synthase protein 8 OS=Rattus norvegicus OX=10116 GN=ATP8 PE=3 SV=1 |
| **83** | 1 | P32551\|QCR2_RA | 104.52 | 6 | 6 | 0 | 2 | 2 | 2 | N | 48396 | Cytochrome b-c1 complex subunit 2, mitochondrial OS=Rattus norvegicus OX=10116 GN=Uqcrc2 PE=1 SV=2 |
| **43** | 4486 | P29419\|ATP5I_RA | 101.05 | 52 | 52 | 1.54E+07 | 4 | 4 | 4 | N | 8255 | ATP synthase subunit e, mitochondrial OS=Rattus norvegicus OX=10116 GN=Atp5me PE=1 SV=3 |
| **63** | 4496 | P21571\|ATP5J_RA | 98.83 | 28 | 28 | 8.31E+05 | 2 | 2 | 3 | N | 12494 | ATP synthase-coupling factor 6, mitochondrial OS=Rattus norvegicus OX=10116 GN=Atp5pf PE=1 SV=1 |
| **106** | 248 | tr\|G3V6I4\|G3V6I | 98.24 | 9 | 9 | 1.56E+06 | 1 | 1 | 1 | N | 37791 | Mitochondrial amidoxime reducing component 1 OS=Rattus norvegicus OX=10116 GN=Marc1 PE=1 SV=1 |
| **196** | 170 | Q63362\|NDUA5_ | 97.71 | 22 | 22 | 0 | 1 | 1 | 1 | N | 13412 | NADH dehydrogenase [ubiquinone] 1 alpha subcomplex subunit 5 OS=Rattus norvegicus OX=10116 GN=Ndufa5 PE=1 SV=3 |

| | | | | | | | | | | | | |
|---|---|---|---|---|---|---|---|---|---|---|---|---|
| **64** | 112 | tr\|Q5PQZ9\|Q5PQ | 97.41 | 21 | 21 | 6.39E+05 | 3 | 3 | 3 | N | 14359 | NADH dehydrogenase [ubiquinone] 1 subunit C2 OS=Rattus norvegicus OX=10116 GN=Ndufc2 PE=1 SV=1 |
| **25** | 4488 | D3ZAF6\|ATPK_RA | 95.42 | 40 | 40 | 2.51E+07 | 7 | 7 | 8 | Y | 10452 | ATP synthase subunit f, mitochondrial OS=Rattus norvegicus OX=10116 GN=Atp5mf PE=1 SV=1 |
| **38** | 102 | tr\|F1LPB3\|F1LPB3 | 94.17 | 4 | 4 | 2.89E+05 | 3 | 1 | 4 | N | 76559 | Long-chain-fatty-acid--CoA ligase 5 OS=Rattus norvegicus OX=10116 GN=Acsl5 PE=1 SV=3 |
| **38** | 101 | O88813\|ACSL5_R | 94.17 | 4 | 4 | 2.89E+05 | 3 | 1 | 4 | N | 76405 | Long-chain-fatty-acid--CoA ligase 5 OS=Rattus norvegicus OX=10116 GN=Acsl5 PE=1 SV=1 |
| **39** | 53 | B3DMA2\|ACD11_ | 89.68 | 4 | 4 | 3.25E+06 | 4 | 4 | 4 | N | 87371 | Acyl-CoA dehydrogenase family member 11 OS=Rattus norvegicus OX=10116 GN=Acad11 PE=1 SV=1 |
| **77** | 51 | tr\|Q5EBA4\|Q5EB | 88.7 | 8 | 8 | 2.66E+06 | 2 | 2 | 2 | N | 33215 | Nipsnap1 protein (Fragment) OS=Rattus norvegicus OX=10116 GN=Nipsnap1 PE=2 SV=1 |
| **77** | 52 | tr\|G3V728\|G3V72 | 88.7 | 8 | 8 | 2.66E+06 | 2 | 2 | 2 | N | 33346 | 4-nitrophenylphosphatase domain and non-neuronal SNAP25-like protein homolog 1 (C. elegans), isoform CRA_b OS=Rattus norvegicus OX=10116 GN=Nipsnap1 PE=1 SV=1 |
| **84** | 71 | tr\|F1LPG5\|F1LPG5 | 82.67 | 29 | 29 | 1.85E+06 | 2 | 2 | 2 | N | 15064 | NADH:ubiquinone oxidoreductase subunit B4 OS=Rattus norvegicus OX=10116 GN=Ndufb4 PE=1 SV=1 |
| **36** | 29 | tr\|A0A140TAG5\|A | 81.55 | 6 | 6 | 5.97E+06 | 3 | 3 | 4 | N | 67049 | MICOS complex subunit MIC60 OS=Rattus norvegicus OX=10116 GN=Immt PE=1 SV=1 |
| **36** | 22 | tr\|A0A0G2JVH4\|A | 81.55 | 5 | 5 | 5.97E+06 | 3 | 3 | 4 | N | 86230 | MICOS complex subunit MIC60 OS=Rattus norvegicus OX=10116 GN=Immt PE=1 SV=1 |
| **36** | 30 | Q3KR86\|MIC60_R | 81.55 | 6 | 6 | 5.97E+06 | 3 | 3 | 4 | N | 67177 | MICOS complex subunit Mic60 (Fragment) OS=Rattus norvegicus OX=10116 GN=Immt PE=1 SV=1 |
| **88** | 106 | tr\|D3ZXF9\|D3ZXF | 80.57 | 9 | 9 | 9.52E+05 | 2 | 2 | 2 | Y | 29441 | Mitochondrial ribosomal protein L12 OS=Rattus norvegicus OX=10116 GN=Mrpl12 PE=1 SV=1 |
| **59** | 4489 | P29418\|ATP5E_R | 79.32 | 41 | 41 | 5.36E+06 | 3 | 3 | 3 | N | 5767 | ATP synthase subunit epsilon, mitochondrial OS=Rattus norvegicus OX=10116 GN=Atp5f1e PE=1 SV=2 |
| **62** | 67 | tr\|Q5RJN0\|Q5RJN | 78.15 | 11 | 11 | 2.35E+06 | 2 | 2 | 3 | N | 23945 | NADH dehydrogenase (Ubiquinone) Fe-S protein 7 OS=Rattus norvegicus OX=10116 GN=Ndufs7 PE=1 SV=1 |
| **60** | 103 | P04762\|CATA_RA | 75.08 | 8 | 8 | 1.31E+06 | 3 | 3 | 3 | N | 59757 | Catalase OS=Rattus norvegicus OX=10116 GN=Cat PE=1 SV=3 |
| **51** | 63 | Q09073\|ADT2_RA | 69.12 | 10 | 10 | 2.65E+06 | 3 | 3 | 3 | N | 32901 | ADP/ATP translocase 2 OS=Rattus norvegicus OX=10116 GN=Slc25a5 PE=1 SV=3 |
| **153** | 66 | Q8VID1\|DHRS4_R | 67.48 | 5 | 5 | 3.20E+05 | 1 | 1 | 1 | N | 29822 | Dehydrogenase/reductase SDR family member 4 OS=Rattus norvegicus OX=10116 GN=Dhrs4 PE=2 SV=2 |
| **197** | 158 | P11240\|COX5A_R | 62.99 | 17 | 17 | 0 | 1 | 1 | 1 | N | 16130 | Cytochrome c oxidase subunit 5A, mitochondrial OS=Rattus norvegicus OX=10116 GN=Cox5a PE=1 SV=1 |
| **152** | 104 | tr\|A9UMV9\|A9UN | 61.39 | 11 | 11 | 4.08E+05 | 1 | 1 | 1 | N | 12500 | NADH:ubiquinone oxidoreductase subunit A7 OS=Rattus norvegicus OX=10116 GN=Ndufa7 PE=1 SV=1 |
| **151** | 110 | tr\|D3ZCZ9\|D3ZCZ | 58.66 | 9 | 9 | 8.65E+05 | 1 | 1 | 1 | N | 13040 | NADH dehydrogenase [ubiquinone] iron-sulfur protein 6, mitochondrial OS=Rattus norvegicus OX=10116 GN=LOC100912599 PE=1 SV=1 |
| **151** | 178 | P52504\|NDUS6_R | 58.66 | 9 | 9 | 8.65E+05 | 1 | 1 | 1 | N | 12783 | NADH dehydrogenase [ubiquinone] iron-sulfur protein 6, mitochondrial OS=Rattus norvegicus OX=10116 GN=Ndufs6 PE=3 SV=1 |
| **91** | 100 | tr\|D3ZE15\|D3ZE1 | 54.88 | 7 | 7 | 1.86E+06 | 2 | 2 | 2 | N | 16777 | NADH:ubiquinone oxidoreductase subunit A13 OS=Rattus norvegicus OX=10116 GN=Ndufa13 PE=1 SV=1 |
| **124** | 111 | Q5RJR8\|LRC59_R | 54.45 | 4 | 4 | 7.23E+05 | 1 | 1 | 1 | N | 34869 | Leucine-rich repeat-containing protein 59 OS=Rattus norvegicus OX=10116 GN=Lrrc59 PE=1 SV=1 |
| **150** | 160 | Q8VHF5\|CISY_RA | 50.03 | 2 | 2 | 0 | 1 | 1 | 1 | N | 51867 | Citrate synthase, mitochondrial OS=Rattus norvegicus OX=10116 GN=Cs PE=1 SV=1 |
| **150** | 161 | tr\|G3V936\|G3V93 | 50.03 | 2 | 2 | 0 | 1 | 1 | 1 | N | 51831 | Citrate synthase OS=Rattus norvegicus OX=10116 GN=Cs PE=1 SV=1 |
| **198** | 234 | tr\|Q06QK5\|Q06Q | 46.65 | 2 | 2 | 2.81E+06 | 1 | 1 | 1 | N | 68606 | NADH-ubiquinone oxidoreductase chain 5 OS=Rattus norvegicus OX=10116 GN=ND5 PE=3 SV=1 |
| **198** | 235 | P11661\|NU5M_R | 46.65 | 2 | 2 | 2.81E+06 | 1 | 1 | 1 | N | 68618 | NADH-ubiquinone oxidoreductase chain 5 OS=Rattus norvegicus OX=10116 GN=Mtnd5 PE=3 SV=3 |
| **198** | 239 | tr\|Q06QG6\|Q06Q | 46.65 | 2 | 2 | 2.81E+06 | 1 | 1 | 1 | N | 68588 | NADH-ubiquinone oxidoreductase chain 5 OS=Rattus norvegicus OX=10116 GN=ND5 PE=3 SV=1 |
| **198** | 236 | tr\|Q06QA1\|Q06Q | 46.65 | 2 | 2 | 2.81E+06 | 1 | 1 | 1 | N | 68574 | NADH-ubiquinone oxidoreductase chain 5 OS=Rattus norvegicus OX=10116 GN=ND5 PE=3 SV=1 |
| **198** | 240 | tr\|A7XYD5\|A7XYD | 46.65 | 2 | 2 | 2.81E+06 | 1 | 1 | 1 | N | 68856 | NADH-ubiquinone oxidoreductase chain 5 OS=Rattus norvegicus OX=10116 GN=Nd5 PE=3 SV=1 |
| **198** | 237 | tr\|Q8SEZ0\|Q8SEZ | 46.65 | 2 | 2 | 2.81E+06 | 1 | 1 | 1 | N | 68618 | NADH-ubiquinone oxidoreductase chain 5 OS=Rattus norvegicus OX=10116 GN=Mt-nd5 PE=3 SV=1 |
| **198** | 238 | tr\|A0A096XKT9\|A | 46.65 | 2 | 2 | 2.81E+06 | 1 | 1 | 1 | N | 68584 | NADH-ubiquinone oxidoreductase chain 5 OS=Rattus norvegicus OX=10116 GN=ND5 PE=3 SV=1 |
| **198** | 241 | tr\|A7XYC1\|A7XYC | 46.65 | 2 | 2 | 2.81E+06 | 1 | 1 | 1 | N | 68841 | NADH-ubiquinone oxidoreductase chain 5 OS=Rattus norvegicus OX=10116 GN=Nd5 PE=3 SV=1 |
| **198** | 242 | tr\|A0A0A1FZN8\|A | 46.65 | 2 | 2 | 2.81E+06 | 1 | 1 | 1 | N | 68970 | NADH-ubiquinone oxidoreductase chain 5 OS=Rattus norvegicus OX=10116 GN=ND5 PE=3 SV=1 |
| **89** | 113 | tr\|B2RYW3\|B2RY | 44.46 | 11 | 11 | 1.70E+06 | 2 | 2 | 2 | N | 21892 | NADH dehydrogenase (Ubiquinone) 1 beta subcomplex, 9 OS=Rattus norvegicus OX=10116 GN=Ndufb9 PE=1 SV=1 |
| **86** | 80 | tr\|S5RZM8\|S5RZN | 44.27 | 7 | 7 | 2.40E+06 | 2 | 2 | 2 | Y | 25958 | Cytochrome c oxidase subunit 2 OS=Rattus norvegicus OX=10116 GN=COX2 PE=3 SV=1 |
| **86** | 81 | P00406\|COX2_RA | 44.27 | 7 | 7 | 2.40E+06 | 2 | 2 | 2 | Y | 25928 | Cytochrome c oxidase subunit 2 OS=Rattus norvegicus OX=10116 GN=Mtco2 PE=1 SV=3 |
| **86** | 68 | tr\|Q5UAJ6\|Q5UA | 44.27 | 7 | 7 | 2.40E+06 | 2 | 2 | 2 | Y | 25942 | Cytochrome c oxidase subunit 2 OS=Rattus norvegicus OX=10116 GN=COX2 PE=3 SV=1 |
| **86** | 109 | tr\|Q37652\|Q3765 | 44.27 | 7 | 7 | 2.40E+06 | 2 | 2 | 2 | Y | 26007 | Cytochrome c oxidase subunit 2 OS=Rattus norvegicus OX=10116 GN=COXII PE=3 SV=1 |
| **86** | 82 | tr\|Q8SEZ5\|Q8SEZ | 44.27 | 7 | 7 | 2.40E+06 | 2 | 2 | 2 | Y | 25928 | Cytochrome c oxidase subunit 2 OS=Rattus norvegicus OX=10116 GN=Mt-co2 PE=1 SV=1 |
| **86** | 83 | tr\|A0A097PE04\|A | 44.27 | 7 | 7 | 2.40E+06 | 2 | 2 | 2 | Y | 25894 | Cytochrome c oxidase subunit 2 OS=Rattus norvegicus OX=10116 GN=COX2 PE=3 SV=1 |
| **87** | 60 | tr\|B0BNE6\|B0BNE | 43.56 | 8 | 8 | 1.30E+05 | 2 | 2 | 2 | Y | 23970 | NADH dehydrogenase (Ubiquinone) Fe-S protein 8 (Predicted), isoform CRA_a OS=Rattus norvegicus OX=10116 GN=Ndufs8 PE=1 SV=1 |
| **154** | 15 | Q68FY0\|QCR1_RA | 43.38 | 5 | 5 | 0 | 1 | 1 | 1 | N | 52849 | Cytochrome b-c1 complex subunit 1, mitochondrial OS=Rattus norvegicus OX=10116 GN=Uqcrc1 PE=1 SV=1 |
| **50** | 1061 | tr\|D3ZKG9\|D3ZKG | 41.98 | 2 | 2 | 6.71E+06 | 3 | 2 | 3 | Y | 145348 | Clustered mitochondria protein homolog OS=Rattus norvegicus OX=10116 GN=Cluh PE=1 SV=3 |
| **50** | 1062 | tr\|A0A0U1RRV5\|A | 41.98 | 2 | 2 | 6.71E+06 | 3 | 2 | 3 | Y | 151165 | Clustered mitochondria protein homolog OS=Rattus norvegicus OX=10116 GN=Cluh PE=1 SV=1 |
| **199** | 88 | tr\|D4A3V2\|D4A3V | 41.62 | 8 | 8 | 6.21E+05 | 1 | 1 | 1 | N | 15224 | NADH dehydrogenase [ubiquinone] 1 alpha subcomplex subunit 6 OS=Rattus norvegicus OX=10116 GN=Ndufa6 PE=1 SV=1 |
| **149** | 1110 | Q06437\|ODPAT_F | 39.87 | 3 | 3 | 1.10E+06 | 1 | 1 | 1 | N | 43393 | Pyruvate dehydrogenase E1 component subunit alpha, testis-specific form, mitochondrial OS=Rattus norvegicus OX=10116 GN=Pdha2 PE=1 SV=1 |
| **200** | 228 | tr\|Q7TP12\|Q7TP1 | 39.81 | 10 | 10 | 1.25E+06 | 1 | 1 | 1 | N | 14969 | Cc2-17 OS=Rattus norvegicus OX=10116 GN=Sqor PE=1 SV=1 |
| **200** | 38 | tr\|B0BMT9\|B0BM | 39.81 | 3 | 3 | 1.25E+06 | 1 | 1 | 1 | N | 50202 | Sqrdl protein OS=Rattus norvegicus OX=10116 GN=Sqor PE=1 SV=1 |
| **125** | 96 | tr\|D3ZFJ6\|D3ZFJ6 | 37.93 | 7 | 7 | 8.89E+05 | 1 | 1 | 1 | N | 60420 | Lactamase, beta OS=Rattus norvegicus OX=10116 GN=Lactb PE=1 SV=1 |
| **201** | 276 | tr\|F1LTG5\|F1LTG5 | 37.15 | 14 | 14 | 1.81E+06 | 1 | 1 | 1 | N | 13829 | Uncharacterized protein OS=Rattus norvegicus OX=10116 PE=4 SV=2 |
| **201** | 133 | P12075\|COX5B_R | 37.15 | 14 | 14 | 1.81E+06 | 1 | 1 | 1 | N | 13915 | Cytochrome c oxidase subunit 5B, mitochondrial OS=Rattus norvegicus OX=10116 GN=Cox5b PE=1 SV=2 |
| **31** | 539 | F1M775\|DIAP1_R | 36.97 | 2 | 2 | 2.17E+07 | 3 | 3 | 5 | Y | 140418 | Protein diaphanous homolog 1 OS=Rattus norvegicus OX=10116 GN=Diaph1 PE=1 SV=3 |
| **95** | 365 | tr\|G3V8G1\|G3V8 | 33.48 | 1 | 1 | 1.28E+07 | 2 | 1 | 2 | Y | 137113 | ATP/GTP binding protein 1 (Predicted), isoform CRA_a OS=Rattus norvegicus OX=10116 GN=Agtpbp1 PE=4 SV=1 |
| **93** | 484 | Q80W57\|ABCG2_ | 33.48 | 3 | 3 | 3.65E+06 | 2 | 1 | 2 | N | 72961 | Broad substrate specificity ATP-binding cassette transporter ABCG2 OS=Rattus norvegicus OX=10116 GN=Abcg2 PE=1 SV=1 |
| **202** | 373 | Q641Y8\|DDX1_RA | 31.84 | 2 | 2 | 1.12E+06 | 1 | 1 | 1 | Y | 82497 | ATP-dependent RNA helicase DDX1 OS=Rattus norvegicus OX=10116 GN=Ddx1 PE=1 SV=1 |
| **74** | 482 | tr\|D3ZXA6\|D3ZXA | 29.61 | 3 | 3 | 2.05E+06 | 2 | 2 | 2 | N | 98819 | Pyruvate dehydrogenase phosphatase regulatory subunit OS=Rattus norvegicus OX=10116 GN=Pdpr PE=1 SV=1 |
| **69** | 321 | tr\|F1LNF0\|F1LNF | 29.17 | 1 | 1 | 4.37E+06 | 2 | 2 | 2 | N | 228912 | Myosin heavy chain 14 OS=Rattus norvegicus OX=10116 GN=Myh14 PE=1 SV=1 |
| **47** | 421 | P11497\|ACACA_R | 28.67 | 1 | 1 | 1.61E+06 | 3 | 2 | 3 | Y | 265191 | Acetyl-CoA carboxylase 1 OS=Rattus norvegicus OX=10116 GN=Acaca PE=1 SV=1 |
| **203** | 941 | tr\|D4A6J4\|D4A6J | 28.сен | 1 | 1 | 1.02E+06 | 1 | 1 | 1 | N | 111741 | DNA topoisomerase OS=Rattus norvegicus OX=10116 GN=Top3a PE=3 SV=3 |
| **75** | 515 | tr\|A0A0G2K5L6\|A | 27.72 | 1 | 1 | 0 | 2 | 1 | 2 | Y | 275755 | Acetyl-CoA carboxylase beta OS=Rattus norvegicus OX=10116 GN=Acacb PE=1 SV=1 |
| **75** | 516 | tr\|A0A0G2K1F2\|A | 27.72 | 1 | 1 | 0 | 2 | 1 | 2 | Y | 276393 | Acetyl-CoA carboxylase beta OS=Rattus norvegicus OX=10116 GN=Acacb PE=1 SV=1 |
| **75** | 518 | tr\|D3ZBE2\|D3ZBE | 27.72 | 1 | 1 | 0 | 2 | 1 | 2 | Y | 275967 | Acetyl-CoA carboxylase beta OS=Rattus norvegicus OX=10116 GN=Acacb PE=1 SV=3 |

| | | | | | | | | | | | | |
|---|---|---|---|---|---|---|---|---|---|---|---|---|
| **75** | 519 | tr\|E9PSQ0\|E9PSQ | 27.72 | 1 | 1 | 0 | 2 | 1 | 2 | Y | 276255 | Acetyl-CoA carboxylase beta OS=Rattus norvegicus OX=10116 GN=Acacb PE=1 SV=2 |
| **75** | 517 | tr\|O70151\|O7015 | 27.72 | 1 | 1 | 0 | 2 | 1 | 2 | Y | 276097 | Acetyl-CoA carboxylase OS=Rattus norvegicus OX=10116 GN=Acacb PE=2 SV=1 |
| **112** | 1624 | tr\|D3ZUJ5\|D3ZUJ | 25.99 | 8 | 8 | 2.02E+06 | 1 | 1 | 1 | Y | 23973 | Deoxythymidylate kinase OS=Rattus norvegicus OX=10116 GN=Dtymk PE=1 SV=1 |
| **204** | 119 | P32198\|CPT1A_R | 25.76 | 2 | 2 | 2.92E+05 | 1 | 1 | 1 | N | 88126 | Carnitine O-palmitoyltransferase 1, liver isoform OS=Rattus norvegicus OX=10116 GN=Cpt1a PE=1 SV=2 |
| **163** | 4506 | Q5BJQ0\|COQ8A_ | 25.69 | 1 | 1 | 0 | 1 | 1 | 1 | N | 72226 | Atypical kinase COQ8A, mitochondrial OS=Rattus norvegicus OX=10116 GN=Coq8a PE=2 SV=1 |
| **159** | 784 | tr\|A0A0G2JX15\|A | 25.окт | 1 | 1 | 7.16E+05 | 1 | 1 | 1 | N | 121101 | Cation-transporting ATPase OS=Rattus norvegicus OX=10116 GN=Atp13a2 PE=3 SV=1 |
| **92** | 4509 | Q06645\|AT5G1_R | 24.35 | 5 | 5 | 1.13E+07 | 1 | 1 | 2 | N | 14244 | ATP synthase F(0) complex subunit C1, mitochondrial OS=Rattus norvegicus OX=10116 GN=Atp5mc1 PE=1 SV=1 |
| **92** | 4510 | tr\|Q499S2\|Q499S | 24.35 | 5 | 5 | 1.13E+07 | 1 | 1 | 2 | N | 14745 | ATP synthase F(0) complex subunit C3, mitochondrial OS=Rattus norvegicus OX=10116 GN=Atp5mc3 PE=2 SV=1 |
| **92** | 4511 | Q06646\|AT5G2_R | 24.35 | 5 | 5 | 1.13E+07 | 1 | 1 | 2 | N | 14918 | ATP synthase F(0) complex subunit C2, mitochondrial OS=Rattus norvegicus OX=10116 GN=Atp5mc2 PE=1 SV=1 |
| **92** | 4512 | Q71S46\|AT5G3_R | 24.35 | 5 | 5 | 1.13E+07 | 1 | 1 | 2 | N | 14693 | ATP synthase F(0) complex subunit C3, mitochondrial OS=Rattus norvegicus OX=10116 GN=Atp5mc3 PE=1 SV=1 |
| **92** | 4513 | tr\|A0A0G2JTN8\|A | 24.35 | 5 | 5 | 1.13E+07 | 1 | 1 | 2 | N | 15465 | ATP synthase F(0) complex subunit C2, mitochondrial OS=Rattus norvegicus OX=10116 GN=Atp5mc2 PE=3 SV=1 |
| **81** | 701 | tr\|F1LMJ8\|F1LMJ | 24.29 | 5 | 5 | 6.00E+06 | 2 | 2 | 2 | N | 49215 | Calcium uptake protein 2, mitochondrial OS=Rattus norvegicus OX=10116 GN=Micu2 PE=1 SV=1 |
| **160** | 1534 | tr\|D3ZSM9\|D3ZS | 24.2 | 1 | 1 | 9.34E+05 | 1 | 1 | 1 | Y | 88476 | BRCA1-associated ATM activator 1 OS=Rattus norvegicus OX=10116 GN=Brat1 PE=1 SV=3 |
| **120** | 549 | P05708\|HXK1_RA | 24.14 | 1 | 1 | 5.40E+05 | 1 | 1 | 1 | N | 102408 | Hexokinase-1 OS=Rattus norvegicus OX=10116 GN=Hk1 PE=1 SV=4 |
| **120** | 533 | tr\|M0RAQ6\|M0R | 24.14 | 1 | 1 | 5.40E+05 | 1 | 1 | 1 | N | 99557 | Hexokinase-1 OS=Rattus norvegicus OX=10116 GN=Hk1 PE=1 SV=2 |
| **120** | 468 | Q63704\|CPT1B_R | 24.14 | 1 | 1 | 5.40E+05 | 1 | 1 | 1 | N | 88217 | Carnitine O-palmitoyltransferase 1, muscle isoform OS=Rattus norvegicus OX=10116 GN=Cpt1b PE=1 SV=1 |
| **120** | 489 | F1LN46\|CPT1C_R | 24.14 | 1 | 1 | 5.40E+05 | 1 | 1 | 1 | N | 90170 | Carnitine O-palmitoyltransferase 1, brain isoform OS=Rattus norvegicus OX=10116 GN=Cpt1c PE=1 SV=1 |
| **208** | 137 | Q9WVK7\|HCDH_ | 23.96 | 3 | 3 | 0 | 1 | 1 | 1 | N | 34448 | Hydroxyacyl-coenzyme A dehydrogenase, mitochondrial OS=Rattus norvegicus OX=10116 GN=Hadh PE=2 SV=1 |
| **94** | 4538 | tr\|B1WBM1\|B1W | 23.77 | 3 | 3 | 9.50E+05 | 2 | 1 | 2 | Y | 58354 | FAST kinase domain-containing protein 3, mitochondrial OS=Rattus norvegicus OX=10116 GN=Fastkd3 PE=2 SV=1 |
| **94** | 4541 | Q68FN9\|FAKD3_ | 23.77 | 2 | 2 | 9.50E+05 | 2 | 1 | 2 | Y | 75248 | FAST kinase domain-containing protein 3, mitochondrial OS=Rattus norvegicus OX=10116 GN=Fastkd3 PE=2 SV=2 |
| **209** | 4519 | tr\|D3ZA45\|D3ZA4 | 23.63 | 2 | 2 | 8.43E+06 | 1 | 1 | 1 | N | 52491 | Autophagy-related protein 13 OS=Rattus norvegicus OX=10116 GN=Atg13 PE=1 SV=2 |
| **107** | 162 | tr\|D3Z900\|D3Z90 | 23.39 | 3 | 3 | 0 | 1 | 1 | 1 | N | 38239 | Mitochondrial amidoxime reducing component 2 OS=Rattus norvegicus OX=10116 GN=Marc2 PE=1 SV=2 |
| **107** | 163 | O88994\|MARC2_ | 23.39 | 3 | 3 | 0 | 1 | 1 | 1 | N | 38249 | Mitochondrial amidoxime reducing component 2 OS=Rattus norvegicus OX=10116 GN=Marc2 PE=2 SV=1 |
| **107** | 164 | tr\|M0R6N2\|M0R | 23.39 | 3 | 3 | 0 | 1 | 1 | 1 | N | 38175 | MOSC domain-containing protein 2, mitochondrial-like OS=Rattus norvegicus OX=10116 GN=LOC100910481 PE=1 SV=1 |
| **127** | 4501 | tr\|F1LUC0\|F1LUC | 23.мар | 1 | 1 | 0 | 1 | 1 | 1 | Y | 72937 | Similar to RIKEN cDNA 5730410E15 gene (Predicted), isoform CRA_a OS=Rattus norvegicus OX=10116 GN=Sybu PE=1 SV=2 |
| **28** | 4514 | tr\|F1LSY7\|F1LSY7 | 22.67 | 1 | 1 | 7.68E+07 | 1 | 1 | 5 | Y | 109031 | Endoplasmic reticulum to nucleus-signaling 1 OS=Rattus norvegicus OX=10116 GN=Ern1 PE=4 SV=2 |
| **28** | 4515 | tr\|A0A0G2K2H4\| | 22.67 | 1 | 1 | 7.68E+07 | 1 | 1 | 5 | Y | 110150 | Endoplasmic reticulum to nucleus-signaling 1 OS=Rattus norvegicus OX=10116 GN=Ern1 PE=2 SV=1 |
| **168** | 1019 | Q2M2R8\|PEX5_R | 22.32 | 1 | 1 | 1.91E+06 | 1 | 1 | 1 | N | 71022 | Peroxisomal targeting signal 1 receptor OS=Rattus norvegicus OX=10116 GN=Pex5 PE=1 SV=2 |
| **170** | 1370 | Q6IMY1\|MTUS1_ | 21.96 | 2 | 2 | 0 | 1 | 1 | 1 | N | 50745 | Microtubule-associated tumor suppressor 1 homolog OS=Rattus norvegicus OX=10116 GN=Mtus1 PE=1 SV=1 |
| **170** | 1371 | tr\|B2GVA4\|B2GV | 21.96 | 2 | 2 | 0 | 1 | 1 | 1 | N | 50728 | Mitochondrial tumor suppressor 1 OS=Rattus norvegicus OX=10116 GN=Mtus1 PE=2 SV=1 |
| **169** | 4525 | tr\|R9PXR4\|R9PXR | 21.95 | 1 | 1 | 9.34E+06 | 1 | 1 | 1 | N | 61939 | Mitochondrial import receptor subunit TOM70 OS=Rattus norvegicus OX=10116 GN=Tomm70 PE=1 SV=1 |
| **169** | 4526 | Q75Q39\|TOM70_ | 21.95 | 1 | 1 | 9.34E+06 | 1 | 1 | 1 | N | 67445 | Mitochondrial import receptor subunit TOM70 OS=Rattus norvegicus OX=10116 GN=Tomm70 PE=1 SV=1 |
| **100** | 396 | P11530\|DMD_RA | 21.17 | 0 | 0 | 1.78E+06 | 1 | 1 | 1 | N | 425830 | Dystrophin OS=Rattus norvegicus OX=10116 GN=Dmd PE=1 SV=2 |
| **98** | 834 | tr\|A0A0G2JYD4\|A | 20.72 | 0 | 0 | 4.63E+06 | 1 | 1 | 1 | N | 487857 | Vacuolar protein sorting 13 homolog D OS=Rattus norvegicus OX=10116 GN=Vps13d PE=1 SV=1 |
| **98** | 835 | tr\|D3ZKC6\|D3ZKC | 20.72 | 0 | 0 | 4.63E+06 | 1 | 1 | 1 | N | 488962 | Vacuolar protein sorting 13 homolog D OS=Rattus norvegicus OX=10116 GN=Vps13d PE=1 SV=1 |
| **213** | 4555 | Q5EBA0\|ERAL1_R | 20.48 | 2 | 2 | 0 | 1 | 1 | 1 | N | 48379 | GTPase Era, mitochondrial OS=Rattus norvegicus OX=10116 GN=Eral1 PE=2 SV=2 |
| **214** | 4556 | Q8VIJ5\|OGA_RAT | 20.14 | 1 | 1 | 1.93E+06 | 1 | 1 | 1 | N | 102918 | Protein O-GlcNAcase OS=Rattus norvegicus OX=10116 GN=Oga PE=1 SV=1 |
