## Supplementary material for "Permeability transition pore-related changes in the proteome and channel activity of ATP synthase dimers and monomers": RLM KCl-BM

Protein Accession
Protein Descriptio
Protein Sample

| Protein Group | Protein ID | Accession | -10lgP | Coverage (%) | Coverage (%) Sample 1 | IBAQ | #Peptides | #Unique | #Spec Sample 1 | PTM | Avg. Mass | Description |
|---|---|---|---|---|---|---|---|---|---|---|---|---|
| **Protein Group** | **Protein ID** | **Accession** | **-10lgP** | **Coverage (%)** | **Coverage (%) Sample 1** | **Area Sample 1** | **#Peptides** | **#Unique** | **#Spec Sample 1** | **PTM** | **Avg. Mass** | **Description** |
| **1** | 2 | x | 300.15 | 61 | 61 | 53189000 | 35 | 35 | 68 | Y | 57808 | Methylmalonate-semialdehyde dehydrogenase [acylating], mitochondrial OS=Rattus norvegicus OX=10116 GN=Aldh6a1 PE=1 SV=1 |
| **1** | 3 | tr\|G3V7J0\|G3V7J0_RAT | 300.15 | 61 | 61 | 53189000 | 35 | 35 | 68 | Y | 57748 | Aldehyde dehydrogenase family 6, subfamily A1, isoform CRA_b OS=Rattus norvegicus OX=10116 GN=Aldh6a1 PE=1 SV=1 |
| **5** | 1 | P32551\|QCR2_RAT | 280.47 | 53 | 53 | 14655000 | 20 | 20 | 32 | Y | 48396 | Cytochrome b-c1 complex subunit 2, mitochondrial OS=Rattus norvegicus OX=10116 GN=Uqcrc2 PE=1 SV=2 |
| **2** | 5 | tr\|B2GV15\|B2GV15_RAT | 248.12 | 55 | 55 | 19824000 | 26 | 26 | 46 | Y | 53274 | Dihydrolipoamide acetyltransferase component of pyruvate dehydrogenase complex OS=Rattus norvegicus OX=10116 GN=Dbt PE=1 SV=1 |
| **4** | 4 | P67779\|PHB_RAT | 247.01 | 86 | 86 | 25218000 | 22 | 22 | 33 | N | 29820 | Prohibitin OS=Rattus norvegicus OX=10116 GN=Phb PE=1 SV=1 |
| **10** | 14 | P15999\|ATPA_RAT | 246 | 29 | 29 | 4011600 | 12 | 12 | 21 | Y | 59754 | ATP synthase subunit alpha, mitochondrial OS=Rattus norvegicus OX=10116 GN=Atp5f1a PE=1 SV=2 |
| **3** | 9 | P00507\|AATM_RAT | 244.24 | 56 | 56 | 21538000 | 23 | 23 | 36 | Y | 47314 | Aspartate aminotransferase, mitochondrial OS=Rattus norvegicus OX=10116 GN=Got2 PE=1 SV=2 |
| **8** | 12 | Q68FY0\|QCR1_RAT | 230.44 | 44 | 44 | 7338000 | 17 | 17 | 27 | Y | 52849 | Cytochrome b-c1 complex subunit 1, mitochondrial OS=Rattus norvegicus OX=10116 GN=Uqcrc1 PE=1 SV=1 |
| **9** | 6 | Q66HF1\|NDUS1_RAT | 230.37 | 35 | 35 | 8916300 | 17 | 17 | 27 | Y | 79412 | NADH-ubiquinone oxidoreductase 75 kDa subunit, mitochondrial OS=Rattus norvegicus OX=10116 GN=Ndufs1 PE=1 SV=1 |
| **7** | 7 | P10719\|ATPB_RAT | 224.82 | 59 | 59 | 6808300 | 16 | 16 | 29 | Y | 56354 | ATP synthase subunit beta, mitochondrial OS=Rattus norvegicus OX=10116 GN=Atp5f1b PE=1 SV=2 |
| **7** | 8 | tr\|G3V6D3\|G3V6D3_RAT | 224.82 | 59 | 59 | 6808300 | 16 | 16 | 29 | Y | 56345 | ATP synthase subunit beta OS=Rattus norvegicus OX=10116 GN=Atp5f1b PE=1 SV=1 |
| **6** | 10 | tr\|A0A0G2KB63\|A0A0G2KB63_R | 220.05 | 63 | 63 | 15965000 | 20 | 20 | 31 | Y | 33168 | Prohibitin OS=Rattus norvegicus OX=10116 GN=Phb2 PE=1 SV=1 |
| **6** | 11 | Q5XIH7\|PHB2_RAT | 220.05 | 63 | 63 | 15965000 | 20 | 20 | 31 | Y | 33312 | Prohibitin-2 OS=Rattus norvegicus OX=10116 GN=Phb2 PE=1 SV=1 |
| **16** | 18 | Q4FZT0\|STML2_RAT | 213.81 | 39 | 39 | 5135600 | 10 | 10 | 14 | Y | 38414 | Stomatin-like protein 2, mitochondrial OS=Rattus norvegicus OX=10116 GN=Stoml2 PE=1 SV=1 |
| **14** | 21 | Q64428\|ECHA_RAT | 213 | 18 | 18 | 1148100 | 9 | 9 | 14 | N | 82665 | Trifunctional enzyme subunit alpha, mitochondrial OS=Rattus norvegicus OX=10116 GN=Hadha PE=1 SV=2 |
| **12** | 15 | Q5BK63\|NDUA9_RAT | 206.52 | 34 | 34 | 3039700 | 11 | 11 | 17 | Y | 42559 | NADH dehydrogenase [ubiquinone] 1 alpha subcomplex subunit 9, mitochondrial OS=Rattus norvegicus OX=10116 GN=Ndufa9 PE=1 SV=2 |
| **19** | 25 | tr\|D3ZG43\|D3ZG43_RAT | 190.92 | 33 | 33 | 5884400 | 8 | 8 | 11 | N | 30226 | NADH dehydrogenase (Ubiquinone) Fe-S protein 3 (Predicted), isoform CRA_c OS=Rattus norvegicus OX=10116 GN=Ndufs3 PE=1 SV=1 |
| **11** | 13 | P07756\|CPSM_RAT | 187.66 | 14 | 14 | 2159100 | 12 | 12 | 19 | Y | 164579 | Carbamoyl-phosphate synthase [ammonia], mitochondrial OS=Rattus norvegicus OX=10116 GN=Cps1 PE=1 SV=1 |
| **20** | 29 | tr\|D4A0T0\|D4A0T0_RAT | 173.64 | 36 | 36 | 1052200 | 5 | 5 | 9 | N | 20859 | NADH:ubiquinone oxidoreductase subunit B10 OS=Rattus norvegicus OX=10116 GN=Ndufb10 PE=1 SV=1 |
| **18** | 27 | Q641Y2\|NDUS2_RAT | 166.55 | 19 | 19 | 4118600 | 7 | 7 | 12 | N | 52562 | NADH dehydrogenase [ubiquinone] iron-sulfur protein 2, mitochondrial OS=Rattus norvegicus OX=10116 GN=Ndufs2 PE=1 SV=1 |
| **22** | 26 | tr\|F1LN92\|F1LN92_RAT | 158.98 | 12 | 12 | 1078300 | 6 | 6 | 8 | N | 89352 | AFG3-like matrix AAA peptidase subunit 2 OS=Rattus norvegicus OX=10116 GN=Afg3l2 PE=1 SV=1 |
| **13** | 22 | P19234\|NDUV2_RAT | 158.31 | 33 | 33 | 5004800 | 8 | 8 | 15 | N | 27378 | NADH dehydrogenase [ubiquinone] flavoprotein 2, mitochondrial OS=Rattus norvegicus OX=10116 GN=Ndufv2 PE=1 SV=2 |
| **15** | 23 | tr\|Q5XIH3\|Q5XIH3_RAT | 151.87 | 19 | 19 | 2008100 | 7 | 7 | 14 | N | 50731 | NADH dehydrogenase [ubiquinone] flavoprotein 1, mitochondrial OS=Rattus norvegicus OX=10116 GN=Ndufv1 PE=1 SV=1 |
| **24** | 42 | tr\|D4A565\|D4A565_RAT | 151.66 | 24 | 24 | 969090 | 5 | 5 | 8 | N | 21664 | NADH dehydrogenase (Ubiquinone) 1 beta subcomplex, 5 (Predicted), isoform CRA_b OS=Rattus norvegicus OX=10116 GN=Ndufb5 PE=1 SV=1 |
| **23** | 31 | tr\|B2RYS8\|B2RYS8_RAT | 147.4 | 24 | 24 | 1923800 | 3 | 3 | 8 | N | 21959 | NADH dehydrogenase [ubiquinone] 1 beta subcomplex subunit 8, mitochondrial OS=Rattus norvegicus OX=10116 GN=Ndufb8 PE=1 SV=1 |
| **21** | 28 | tr\|D3ZFQ8\|D3ZFQ8_RAT | 146.08 | 22 | 22 | 5324200 | 6 | 6 | 8 | Y | 35435 | Cytochrome c-1 OS=Rattus norvegicus OX=10116 GN=Cyc1 PE=1 SV=3 |
| **17** | 19 | P22791\|HMCS2_RAT | 140.46 | 22 | 22 | 3592200 | 9 | 9 | 13 | Y | 56912 | Hydroxymethylglutaryl-CoA synthase, mitochondrial OS=Rattus norvegicus OX=10116 GN=Hmgcs2 PE=1 SV=1 |
| **17** | 20 | tr\|Q68G44\|Q68G44_RAT | 140.46 | 22 | 22 | 3592200 | 9 | 9 | 13 | Y | 56886 | 3-hydroxy-3-methylglutaryl coenzyme A synthase OS=Rattus norvegicus OX=10116 GN=Hmgcs2 PE=1 SV=1 |
| **30** | 32 | tr\|Q6P6W6\|Q6P6W6_RAT | 126.85 | 26 | 26 | 755760 | 3 | 3 | 5 | N | 31701 | NADH dehydrogenase [ubiquinone] 1 alpha subcomplex subunit 10, mitochondrial OS=Rattus norvegicus OX=10116 GN=Ndufa10 PE=2 SV=1 |
| **30** | 33 | Q561S0\|NDUAA_RAT | 126.85 | 20 | 20 | 755760 | 3 | 3 | 5 | N | 40493 | NADH dehydrogenase [ubiquinone] 1 alpha subcomplex subunit 10, mitochondrial OS=Rattus norvegicus OX=10116 GN=Ndufa10 PE=1 SV=1 |
| **32** | 35 | P13086\|SUCA_RAT | 124.18 | 11 | 11 | 766110 | 3 | 3 | 4 | N | 36148 | Succinate--CoA ligase [ADP/GDP-forming] subunit alpha, mitochondrial OS=Rattus norvegicus OX=10116 GN=Suclg1 PE=2 SV=2 |
| **32** | 36 | tr\|A0A0H2UHE1\|A0A0H2UHE1_I | 124.18 | 10 | 10 | 766110 | 3 | 3 | 4 | N | 37560 | Succinate--CoA ligase [ADP/GDP-forming] subunit alpha, mitochondrial OS=Rattus norvegicus OX=10116 GN=Suclg1 PE=1 SV=1 |
| **25** | 30 | Q7TQ16\|QCR8_RAT | 117.67 | 49 | 49 | 861460 | 5 | 5 | 7 | N | 9849 | Cytochrome b-c1 complex subunit 8 OS=Rattus norvegicus OX=10116 GN=Uqcrq PE=3 SV=1 |
| **31** | 39 | P17764\|THIL_RAT | 110.34 | 12 | 12 | 652570 | 3 | 3 | 5 | N | 44695 | Acetyl-CoA acetyltransferase, mitochondrial OS=Rattus norvegicus OX=10116 GN=Acat1 PE=1 SV=1 |
| **28** | 40 | Q60587\|ECHB_RAT | 99.26 | 8 | 8 | 323320 | 3 | 3 | 5 | N | 51414 | Trifunctional enzyme subunit beta, mitochondrial OS=Rattus norvegicus OX=10116 GN=Hadhb PE=1 SV=1 |
| **42** | 47 | Q9WVK3\|PECR_RAT | 88.59 | 10 | 10 | 498870 | 2 | 2 | 3 | N | 32433 | Peroxisomal trans-2-enoyl-CoA reductase OS=Rattus norvegicus OX=10116 GN=Pecr PE=2 SV=1 |
| **42** | 48 | tr\|A0A0G2JVG4\|A0A0G2JVG4_R | 88.59 | 10 | 10 | 498870 | 2 | 2 | 3 | N | 32737 | Peroxisomal trans-2-enoyl-CoA reductase OS=Rattus norvegicus OX=10116 GN=Pecr PE=1 SV=1 |
| **41** | 46 | tr\|F1LXA0\|F1LXA0_RAT | 84.32 | 17 | 17 | 638530 | 2 | 2 | 3 | N | 17178 | NADH dehydrogenase [ubiquinone] 1 alpha subcomplex subunit 12 OS=Rattus norvegicus OX=10116 GN=Ndufa12 PE=1 SV=2 |
| **27** | 37 | P20788\|UCRI_RAT | 83.1 | 16 | 16 | 1041900 | 3 | 3 | 6 | N | 29446 | Cytochrome b-c1 complex subunit Rieske, mitochondrial OS=Rattus norvegicus OX=10116 GN=Uqcrfs1 PE=1 SV=2 |
| **97** | 92 | tr\|Q4G067\|Q4G067_RAT | 80.06 | 6 | 6 | 206200 | 1 | 1 | 1 | N | 37440 | Mitochondrial ribosomal protein L44 OS=Rattus norvegicus OX=10116 GN=Mrpl44 PE=1 SV=1 |
| **26** | 34 | tr\|Q5RJN0\|Q5RJN0_RAT | 79.79 | 27 | 27 | 731870 | 4 | 4 | 6 | Y | 23945 | NADH dehydrogenase (Ubiquinone) Fe-S protein 7 OS=Rattus norvegicus OX=10116 GN=Ndufs7 PE=1 SV=1 |
| **98** | 93 | tr\|A0A0G2K8Q8\|A0A0G2K8Q8_F | 74.65 | 27 | 27 | 359800 | 1 | 1 | 1 | N | 7099 | Ubiquinol-cytochrome c reductase, complex III subunit X OS=Rattus norvegicus OX=10116 GN=Uqcr10 PE=1 SV=1 |

| | | | | | | | | | | | | |
|---|---|---|---|---|---|---|---|---|---|---|---|---|
| **98** | 94 | tr\|B2RYX1\|B2RYX1_RAT | 74.65 | 27 | 27 | 359800 | 1 | 1 | 1 | N | 7462 | LOC685322 protein OS=Rattus norvegicus OX=10116 GN=Uqcr10 PE=2 SV=1 |
| **99** | 95 | Q63362\|NDUA5_RAT | 72.33 | 22 | 22 | 0 | 1 | 1 | 1 | N | 13412 | NADH dehydrogenase [ubiquinone] 1 alpha subcomplex subunit 5 OS=Rattus norvegicus OX=10116 GN=Ndufa5 PE=1 SV=3 |
| **38** | 43 | tr\|A0A0G2JSH5\|A0A0G2JSH5_RA | 70.23 | 6 | 6 | 386050 | 3 | 1 | 3 | Y | 68759 | Serum albumin OS=Rattus norvegicus OX=10116 GN=Alb PE=1 SV=1 |
| **47** | 86 | tr\|F1LPG5\|F1LPG5_RAT | 69.49 | 19 | 19 | 1195600 | 1 | 1 | 2 | N | 15064 | NADH:ubiquinone oxidoreductase subunit B4 OS=Rattus norvegicus OX=10116 GN=Ndufb4 PE=1 SV=1 |
| **33** | 41 | P19511\|AT5F1_RAT | 69.1 | 26 | 26 | 957100 | 3 | 3 | 4 | N | 28869 | ATP synthase F(0) complex subunit B1, mitochondrial OS=Rattus norvegicus OX=10116 GN=Atp5pb PE=1 SV=1 |
| **43** | 51 | P10888\|COX41_RAT | 69 | 14 | 14 | 904460 | 2 | 2 | 3 | N | 19515 | Cytochrome c oxidase subunit 4 isoform 1, mitochondrial OS=Rattus norvegicus OX=10116 GN=Cox4i1 PE=1 SV=1 |
| **100** | 96 | P08461\|ODP2_RAT | 68.17 | 2 | 2 | 185680 | 1 | 1 | 1 | N | 67166 | Dihydrolipoyllysine-residue acetyltransferase component of pyruvate dehydrogenase complex, mitochondrial OS=Rattus norvegicus OX=10116 GN=Dlat PE=1 SV=3 |
| **101** | 97 | tr\|Q5PQZ9\|Q5PQZ9_RAT | 67.4 | 11 | 11 | 567510 | 1 | 1 | 1 | N | 14359 | NADH dehydrogenase [ubiquinone] 1 subunit C2 OS=Rattus norvegicus OX=10116 GN=Ndufc2 PE=1 SV=1 |
| **57** | 98 | P29147\|BDH_RAT | 64.79 | 6 | 6 | 671700 | 1 | 1 | 2 | N | 38202 | D-beta-hydroxybutyrate dehydrogenase, mitochondrial OS=Rattus norvegicus OX=10116 GN=Bdh1 PE=1 SV=2 |
| **57** | 99 | tr\|A0A0G2JSH2\|A0A0G2JSH2_RA | 64.79 | 6 | 6 | 671700 | 1 | 1 | 2 | N | 38333 | 3-hydroxybutyrate dehydrogenase, type 1, isoform CRA_a OS=Rattus norvegicus OX=10116 GN=Bdh1 PE=1 SV=1 |
| **29** | 49 | tr\|B2RYW3\|B2RYW3_RAT | 64.36 | 16 | 16 | 490610 | 2 | 2 | 5 | N | 21892 | NADH dehydrogenase (Ubiquinone) 1 beta subcomplex, 9 OS=Rattus norvegicus OX=10116 GN=Ndufb9 PE=1 SV=1 |
| **58** | 100 | tr\|D3ZS58\|D3ZS58_RAT | 63.23 | 14 | 14 | 65872 | 1 | 1 | 2 | N | 10845 | NADH dehydrogenase [ubiquinone] 1 alpha subcomplex subunit 2 OS=Rattus norvegicus OX=10116 GN=Ndufa2 PE=1 SV=1 |
| **34** | 38 | tr\|D4A3V2\|D4A3V2_RAT | 62.49 | 21 | 21 | 410840 | 3 | 3 | 4 | N | 15224 | NADH dehydrogenase [ubiquinone] 1 alpha subcomplex subunit 6 OS=Rattus norvegicus OX=10116 GN=Ndufa6 PE=1 SV=1 |
| **39** | 44 | P02770\|ALBU_RAT | 56.42 | 5 | 5 | 533210 | 3 | 1 | 3 | Y | 68731 | Serum albumin OS=Rattus norvegicus OX=10116 GN=Alb PE=1 SV=2 |
| **46** | 76 | tr\|A7XYD5\|A7XYD5_RAT | 55.8 | 2 | 2 | 293410 | 1 | 1 | 2 | N | 68856 | NADH-ubiquinone oxidoreductase chain 5 OS=Rattus norvegicus OX=10116 GN=Nd5 PE=3 SV=1 |
| **46** | 77 | tr\|Q06QK5\|Q06QK5_RAT | 55.8 | 2 | 2 | 293410 | 1 | 1 | 2 | N | 68606 | NADH-ubiquinone oxidoreductase chain 5 OS=Rattus norvegicus OX=10116 GN=ND5 PE=3 SV=1 |
| **46** | 78 | tr\|Q8SEZ0\|Q8SEZ0_RAT | 55.8 | 2 | 2 | 293410 | 1 | 1 | 2 | N | 68618 | NADH-ubiquinone oxidoreductase chain 5 OS=Rattus norvegicus OX=10116 GN=Mt-nd5 PE=3 SV=1 |
| **46** | 79 | tr\|A0A096XKT9\|A0A096XKT9_RA | 55.8 | 2 | 2 | 293410 | 1 | 1 | 2 | N | 68584 | NADH-ubiquinone oxidoreductase chain 5 OS=Rattus norvegicus OX=10116 GN=ND5 PE=3 SV=1 |
| **46** | 80 | tr\|A7XYC1\|A7XYC1_RAT | 55.8 | 2 | 2 | 293410 | 1 | 1 | 2 | N | 68841 | NADH-ubiquinone oxidoreductase chain 5 OS=Rattus norvegicus OX=10116 GN=Nd5 PE=3 SV=1 |
| **46** | 81 | P11661\|NU5M_RAT | 55.8 | 2 | 2 | 293410 | 1 | 1 | 2 | N | 68618 | NADH-ubiquinone oxidoreductase chain 5 OS=Rattus norvegicus OX=10116 GN=Mtnd5 PE=3 SV=3 |
| **46** | 82 | tr\|Q06QG6\|Q06QG6_RAT | 55.8 | 2 | 2 | 293410 | 1 | 1 | 2 | N | 68588 | NADH-ubiquinone oxidoreductase chain 5 OS=Rattus norvegicus OX=10116 GN=ND5 PE=3 SV=1 |
| **46** | 83 | tr\|Q06QA1\|Q06QA1_RAT | 55.8 | 2 | 2 | 293410 | 1 | 1 | 2 | N | 68574 | NADH-ubiquinone oxidoreductase chain 5 OS=Rattus norvegicus OX=10116 GN=ND5 PE=3 SV=1 |
| **46** | 84 | tr\|A0A0A1FZN8\|A0A0A1FZN8_R | 55.8 | 2 | 2 | 293410 | 1 | 1 | 2 | N | 68970 | NADH-ubiquinone oxidoreductase chain 5 OS=Rattus norvegicus OX=10116 GN=ND5 PE=3 SV=1 |
| **102** | 101 | tr\|B2RYT5\|B2RYT5_RAT | 55.47 | 11 | 11 | 648880 | 1 | 1 | 1 | N | 12651 | Cox7a2l protein OS=Rattus norvegicus OX=10116 GN=Cox7a2l PE=2 SV=1 |
| **102** | 102 | tr\|D3ZYX8\|D3ZYX8_RAT | 55.47 | 10 | 10 | 648880 | 1 | 1 | 1 | N | 13274 | Cytochrome c oxidase subunit 7A2-like OS=Rattus norvegicus OX=10116 GN=Cox7a2l PE=1 SV=1 |
| **40** | 52 | tr\|B0BNE6\|B0BNE6_RAT | 51.74 | 9 | 9 | 369880 | 2 | 2 | 3 | N | 23970 | NADH dehydrogenase (Ubiquinone) Fe-S protein 8 (Predicted), isoform CRA_a OS=Rattus norvegicus OX=10116 GN=Ndufs8 PE=1 SV=1 |
| **59** | 103 | Q80W89\|NDUAB_RAT | 47.9 | 11 | 11 | 134820 | 1 | 1 | 2 | N | 14854 | NADH dehydrogenase [ubiquinone] 1 alpha subcomplex subunit 11 OS=Rattus norvegicus OX=10116 GN=Ndufa11 PE=2 SV=1 |
| **50** | 55 | tr\|D3ZZ21\|D3ZZ21_RAT | 45.71 | 26 | 26 | 777740 | 2 | 2 | 2 | Y | 15638 | NADH dehydrogenase (Ubiquinone) 1 beta subcomplex, 6 (Predicted) OS=Rattus norvegicus OX=10116 GN=Ndufb6 PE=1 SV=1 |
| **75** | 89 | tr\|Q5UAJ8\|Q5UAJ8_RAT | 45.55 | 3 | 3 | 0 | 1 | 1 | 1 | Y | 38455 | NADH-ubiquinone oxidoreductase chain 2 OS=Rattus norvegicus OX=10116 GN=ND2 PE=3 SV=1 |
| **75** | 90 | tr\|Q06Q97\|Q06Q97_RAT | 45.55 | 3 | 3 | 0 | 1 | 1 | 1 | Y | 38485 | NADH-ubiquinone oxidoreductase chain 2 OS=Rattus norvegicus OX=10116 GN=ND2 PE=3 SV=1 |
| **75** | 91 | tr\|A0A0A1G491\|A0A0A1G491_R | 45.55 | 3 | 3 | 0 | 1 | 1 | 1 | Y | 38542 | NADH-ubiquinone oxidoreductase chain 2 OS=Rattus norvegicus OX=10116 GN=ND2 PE=3 SV=1 |
| **49** | 53 | P52873\|PYC_RAT | 45.14 | 3 | 3 | 0 | 2 | 2 | 2 | N | 129777 | Pyruvate carboxylase, mitochondrial OS=Rattus norvegicus OX=10116 GN=Pc PE=1 SV=2 |
| **49** | 54 | tr\|A0A0G2JTL5\|A0A0G2JTL5_RAT | 45.14 | 3 | 3 | 0 | 2 | 2 | 2 | N | 140005 | Pyruvate carboxylase, mitochondrial OS=Rattus norvegicus OX=10116 GN=Pc PE=1 SV=1 |
| **55** | 63 | tr\|D4A7L4\|D4A7L4_RAT | 41.16 | 21 | 21 | 490190 | 2 | 2 | 2 | Y | 17634 | NADH dehydrogenase (Ubiquinone) 1 beta subcomplex, 11 (Predicted) OS=Rattus norvegicus OX=10116 GN=Ndufb11 PE=1 SV=1 |
| **73** | 85 | tr\|A0A0G2JVH4\|A0A0G2JVH4_R | 39.7 | 1 | 1 | 32045 | 1 | 1 | 1 | N | 86230 | MICOS complex subunit MIC60 OS=Rattus norvegicus OX=10116 GN=Immt PE=1 SV=1 |
| **73** | 104 | tr\|A0A140TAG5\|A0A140TAG5_R | 39.7 | 2 | 2 | 32045 | 1 | 1 | 1 | N | 67049 | MICOS complex subunit MIC60 OS=Rattus norvegicus OX=10116 GN=Immt PE=1 SV=1 |
| **73** | 105 | Q3KR86\|MIC60_RAT | 39.7 | 2 | 2 | 32045 | 1 | 1 | 1 | N | 67177 | MICOS complex subunit Mic60 (Fragment) OS=Rattus norvegicus OX=10116 GN=Immt PE=1 SV=1 |
| **103** | 106 | tr\|G3V6I4\|G3V6I4_RAT | 39.2 | 9 | 9 | 81929 | 1 | 1 | 1 | N | 37791 | Mitochondrial amidoxime reducing component 1 OS=Rattus norvegicus OX=10116 GN=Marc1 PE=1 SV=1 |
| **44** | 56 | tr\|S5RZM8\|S5RZM8_RAT | 38 | 7 | 7 | 515070 | 2 | 2 | 3 | Y | 25958 | Cytochrome c oxidase subunit 2 OS=Rattus norvegicus OX=10116 GN=COX2 PE=3 SV=1 |
| **44** | 57 | P00406\|COX2_RAT | 38 | 7 | 7 | 515070 | 2 | 2 | 3 | Y | 25928 | Cytochrome c oxidase subunit 2 OS=Rattus norvegicus OX=10116 GN=Mtco2 PE=1 SV=3 |
| **44** | 58 | tr\|Q37652\|Q37652_RAT | 38 | 7 | 7 | 515070 | 2 | 2 | 3 | Y | 26007 | Cytochrome c oxidase subunit 2 OS=Rattus norvegicus OX=10116 GN=COXII PE=3 SV=1 |
| **44** | 59 | tr\|Q5UAJ6\|Q5UAJ6_RAT | 38 | 7 | 7 | 515070 | 2 | 2 | 3 | Y | 25942 | Cytochrome c oxidase subunit 2 OS=Rattus norvegicus OX=10116 GN=COX2 PE=3 SV=1 |
| **44** | 60 | tr\|Q8SEZ5\|Q8SEZ5_RAT | 38 | 7 | 7 | 515070 | 2 | 2 | 3 | Y | 25928 | Cytochrome c oxidase subunit 2 OS=Rattus norvegicus OX=10116 GN=Mt-co2 PE=1 SV=1 |
| **44** | 61 | tr\|A0A097PE04\|A0A097PE04_RA | 38 | 7 | 7 | 515070 | 2 | 2 | 3 | Y | 25894 | Cytochrome c oxidase subunit 2 OS=Rattus norvegicus OX=10116 GN=COX2 PE=3 SV=1 |
| **104** | 107 | tr\|G3V6P2\|G3V6P2_RAT | 37.62 | 3 | 3 | 77137 | 1 | 1 | 1 | N | 48899 | Dihydrolipoamide S-succinyltransferase (E2 component of 2-oxo-glutarate complex), isoform CRA_a OS=Rattus norvegicus OX=10116 GN=Dlst PE=1 SV=1 |
| **72** | 65 | Q06437\|ODPAT_RAT | 35.87 | 3 | 3 | 988510 | 1 | 1 | 1 | N | 43393 | Pyruvate dehydrogenase E1 component subunit alpha, testis-specific form, mitochondrial OS=Rattus norvegicus OX=10116 GN=Pdha2 PE=1 SV=1 |
| **56** | 66 | P05508\|NU4M_RAT | 35.57 | 6 | 6 | 139630 | 2 | 2 | 2 | N | 51783 | NADH-ubiquinone oxidoreductase chain 4 OS=Rattus norvegicus OX=10116 GN=Mtnd4 PE=3 SV=3 |

| | | | | | | | | | | | | |
|---|---|---|---|---|---|---|---|---|---|---|---|---|
| **56** | 67 | tr\|Q06QA2\|Q06QA2_RAT | 35.57 | 6 | 6 | 139630 | 2 | 2 | 2 | N | 51787 | NADH-ubiquinone oxidoreductase chain 4 OS=Rattus norvegicus OX=10116 GN=ND4 PE=3 SV=1 |
| **56** | 68 | tr\|Q7HKW3\|Q7HKW3_RAT | 35.57 | 6 | 6 | 139630 | 2 | 2 | 2 | N | 51801 | NADH-ubiquinone oxidoreductase chain 4 (Fragment) OS=Rattus norvegicus OX=10116 GN=NADH4 PE=3 SV=1 |
| **56** | 69 | tr\|Q06QG7\|Q06QG7_RAT | 35.57 | 6 | 6 | 139630 | 2 | 2 | 2 | N | 51791 | NADH-ubiquinone oxidoreductase chain 4 OS=Rattus norvegicus OX=10116 GN=ND4 PE=3 SV=1 |
| **56** | 70 | tr\|D2E6K0\|D2E6K0_RAT | 35.57 | 6 | 6 | 139630 | 2 | 2 | 2 | N | 51833 | NADH-ubiquinone oxidoreductase chain 4 OS=Rattus norvegicus OX=10116 GN=ND4 PE=3 SV=1 |
| **56** | 71 | tr\|Q06Q89\|Q06Q89_RAT | 35.57 | 6 | 6 | 139630 | 2 | 2 | 2 | N | 51819 | NADH-ubiquinone oxidoreductase chain 4 OS=Rattus norvegicus OX=10116 GN=ND4 PE=3 SV=1 |
| **56** | 72 | tr\|Q35737\|Q35737_RAT | 35.57 | 6 | 6 | 139630 | 2 | 2 | 2 | N | 51765 | NADH-ubiquinone oxidoreductase chain 4 OS=Rattus norvegicus OX=10116 PE=3 SV=1 |
| **56** | 73 | tr\|A7XYB9\|A7XYB9_RAT | 35.57 | 6 | 6 | 139630 | 2 | 2 | 2 | N | 51773 | NADH-ubiquinone oxidoreductase chain 4 OS=Rattus norvegicus OX=10116 GN=Nd4 PE=3 SV=1 |
| **56** | 74 | tr\|Q8HIC6\|Q8HIC6_RAT | 35.57 | 6 | 6 | 139630 | 2 | 2 | 2 | N | 51783 | NADH-ubiquinone oxidoreductase chain 4 OS=Rattus norvegicus OX=10116 GN=Mt-nd4 PE=3 SV=1 |
| **56** | 75 | tr\|Q06QE1\|Q06QE1_RAT | 35.57 | 6 | 6 | 139630 | 2 | 2 | 2 | N | 51851 | NADH-ubiquinone oxidoreductase chain 4 OS=Rattus norvegicus OX=10116 GN=ND4 PE=3 SV=1 |
| **60** | 109 | P11240\|COX5A_RAT | 35.01 | 21 | 21 | 0 | 1 | 1 | 2 | N | 16130 | Cytochrome c oxidase subunit 5A, mitochondrial OS=Rattus norvegicus OX=10116 GN=Cox5a PE=1 SV=1 |
| **61** | 110 | Q9WVK7\|HCDH_RAT | 34.57 | 4 | 4 | 44969 | 1 | 1 | 2 | N | 34448 | Hydroxyacyl-coenzyme A dehydrogenase, mitochondrial OS=Rattus norvegicus OX=10116 GN=Hadh PE=2 SV=1 |
| **105** | 111 | Q5XJW2\|G45IP_RAT | 34.39 | 4 | 4 | 89396 | 1 | 1 | 1 | N | 26467 | Growth arrest and DNA damage-inducible proteins-interacting protein 1 OS=Rattus norvegicus OX=10116 GN=Gadd45gip1 PE=2 SV=2 |
| **76** | 108 | O70595\|ABCB6_RAT | 33.13 | 1 | 1 | 300700 | 1 | 1 | 1 | Y | 93305 | ATP-binding cassette sub-family B member 6, mitochondrial OS=Rattus norvegicus OX=10116 GN=Abcb6 PE=2 SV=1 |
| **106** | 113 | tr\|D3ZCZ9\|D3ZCZ9_RAT | 32.42 | 21 | 21 | 0 | 1 | 1 | 1 | N | 13040 | NADH dehydrogenase [ubiquinone] iron-sulfur protein 6, mitochondrial OS=Rattus norvegicus OX=10116 GN=LOC100912599 PE=1 SV=1 |
| **67** | 64 | tr\|D3ZJY1\|D3ZJY1_RAT | 31.55 | 9 | 9 | 10862 | 1 | 1 | 1 | N | 30237 | Mitochondrial ribosomal protein L28 OS=Rattus norvegicus OX=10116 GN=Mrpl28 PE=1 SV=1 |
| **45** | 116 | P35435\|ATPG_RAT | 29.96 | 12 | 12 | 276970 | 1 | 1 | 3 | N | 30191 | ATP synthase subunit gamma, mitochondrial OS=Rattus norvegicus OX=10116 GN=Atp5f1c PE=1 SV=2 |
| **45** | 117 | tr\|Q6QI09\|Q6QI09_RAT | 29.96 | 5 | 5 | 276970 | 1 | 1 | 3 | N | 67721 | ATP synthase subunit gamma, mitochondrial OS=Rattus norvegicus OX=10116 GN=Taf3 PE=1 SV=1 |
| **108** | 121 | tr\|F1M9H4\|F1M9H4_RAT | 27.16 | 0 | 0 | 309460 | 1 | 1 | 1 | N | 225602 | HIVEP zinc finger 1 OS=Rattus norvegicus OX=10116 GN=Hivep1 PE=1 SV=2 |
| **74** | 88 | Q64578\|AT2A1_RAT | 25.35 | 1 | 1 | 174900 | 1 | 1 | 1 | Y | 109409 | Sarcoplasmic/endoplasmic reticulum calcium ATPase 1 OS=Rattus norvegicus OX=10116 GN=Atp2a1 PE=1 SV=1 |
| **116** | 140 | tr\|F1LPI6\|F1LPI6_RAT | 25.24 | 0 | 0 | 0 | 1 | 1 | 1 | N | 200075 | Peripheral-type benzodiazepine receptor-associated protein 1 OS=Rattus norvegicus OX=10116 GN=Tspoap1 PE=4 SV=1 |
| **116** | 141 | Q9JIR0\|RIMB1_RAT | 25.24 | 0 | 0 | 0 | 1 | 1 | 1 | N | 200202 | Peripheral-type benzodiazepine receptor-associated protein 1 OS=Rattus norvegicus OX=10116 GN=Tspoap1 PE=1 SV=2 |
| **110** | 128 | Q6AXW1\|GLRX2_RAT | 24.59 | 8 | 8 | 261060 | 1 | 1 | 1 | Y | 17275 | Glutaredoxin-2, mitochondrial OS=Rattus norvegicus OX=10116 GN=Glrx2 PE=2 SV=2 |
| **86** | 169 | tr\|Q68FQ5\|Q68FQ5_RAT | 22.95 | 1 | 1 | 0 | 1 | 1 | 1 | N | 81255 | Mitochondrial translational initiation factor 2 OS=Rattus norvegicus OX=10116 GN=Mtif2 PE=2 SV=1 |
| **86** | 170 | tr\|F7FDM3\|F7FDM3_RAT | 22.95 | 1 | 1 | 0 | 1 | 1 | 1 | N | 81470 | Mitochondrial translational initiation factor 2 OS=Rattus norvegicus OX=10116 GN=Mtif2 PE=1 SV=2 |
| **66** | 167 | tr\|D4A054\|D4A054_RAT | 21.33 | 0 | 0 | 150670 | 1 | 1 | 1 | N | 341403 | RAN-binding protein 2 OS=Rattus norvegicus OX=10116 GN=Ranbp2 PE=1 SV=2 |
| **66** | 168 | tr\|M0R3M4\|M0R3M4_RAT | 21.33 | 0 | 0 | 150670 | 1 | 1 | 1 | N | 344396 | RAN-binding protein 2 OS=Rattus norvegicus OX=10116 GN=Ranbp2 PE=1 SV=1 |
| **80** | 136 | tr\|Q5U2X8\|Q5U2X8_RAT | 20.96 | 2 | 2 | 788320 | 1 | 1 | 1 | N | 50444 | Acyl-CoA thioesterase 9 OS=Rattus norvegicus OX=10116 GN=Acot9 PE=1 SV=1 |
| **total 116 proteins** | | | | | | | | | | | | |
