## Supplementary material for "Permeability transition pore-related changes in the proteome and channel activity of ATP synthase dimers and monomers": SI Kruglov

**This PDF file includes:**

Figures S1 to S7
Tables S1 to S2
Legends for Datasets S1 to S10

**Other supplementary materials for this manuscript include the following:**

Datasets S1 to S10

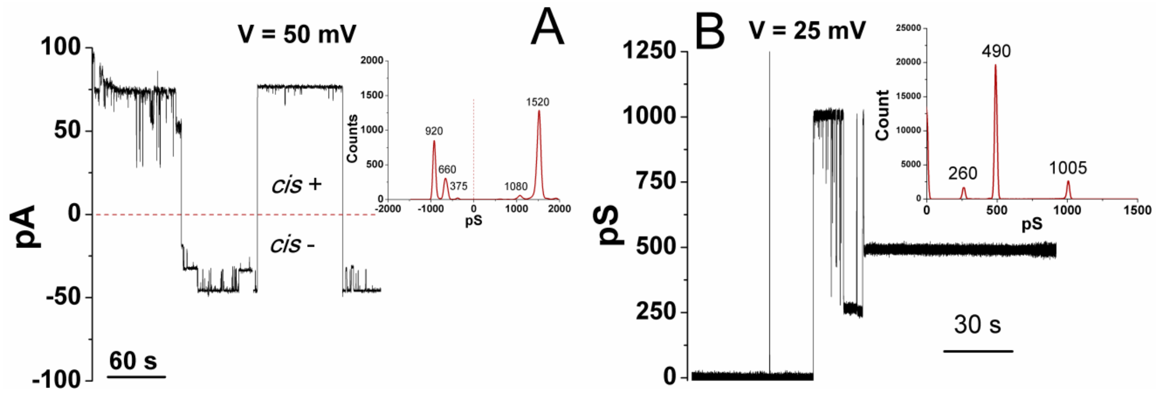


**Fig. S1.** Representative channel-forming activity of mitochondrial F-ATP synthase dimer from control samples in the media supplemented with 200 (A) and 150 mM KCl (B). Currents were recorded at 50 mV *cis* +/- (A) and at 25 mV *cis* +. Protein eluates (90 µl/ml) and $CaCl_2$ (300 µM) were added from the *trans* side of the membrane. Inserts are the corresponding amplitude histograms of conductance.

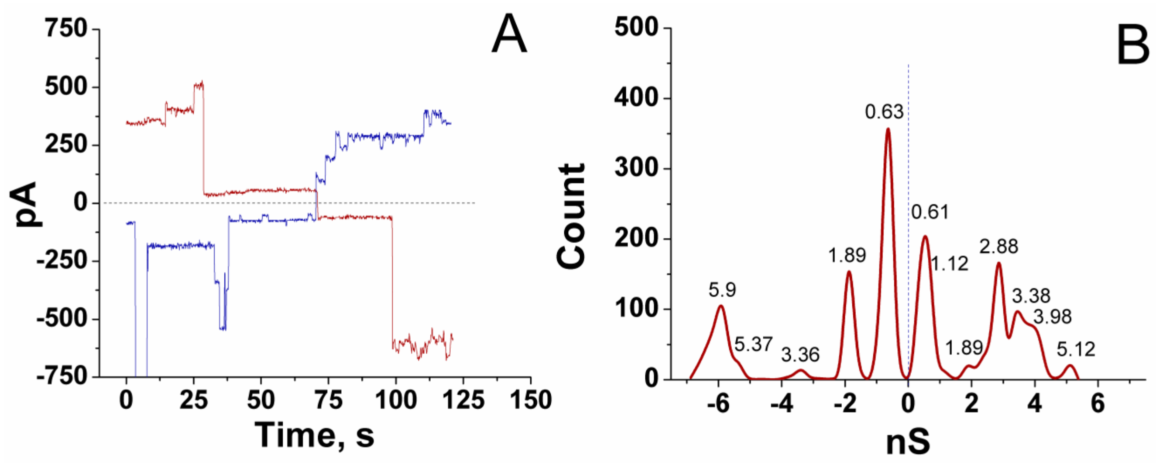


**Fig. S2.** Multiple channel activity of mitochondrial F-ATP synthase dimer from control samples (A) and the corresponding amplitude histogram of conductance (B). Currents were recorded at 100 mV *cis* +/-. In panel B, negative conductance designates the *cis* negative voltage. Protein concentrations in eluates were three times higher than in standard experiments. Protein eluates (90 µl/ml) and $CaCl_2$ (300 µM) were added from the *trans* side of the membrane.

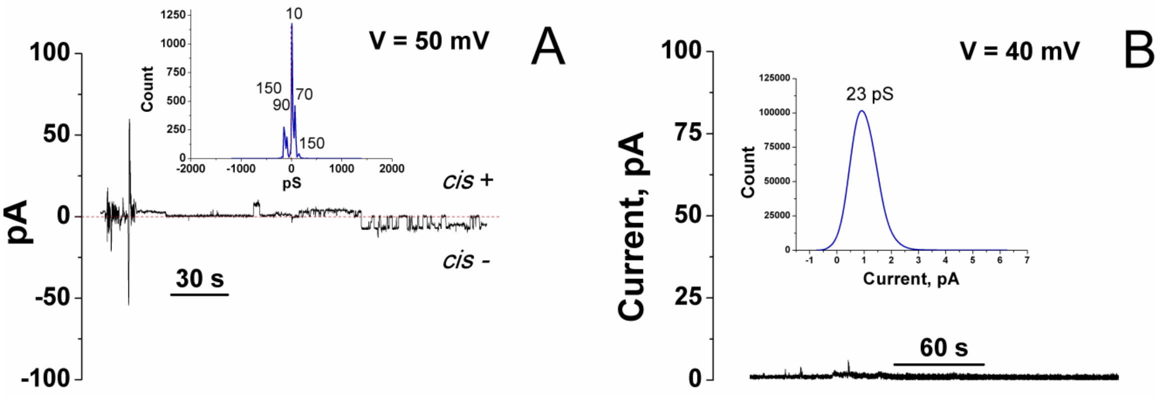


**Fig. S3.** F-ATP synthase dimer from PTP samples with (A) and without channel activity (B). Currents were recorded at 50 mV *cis* +/- (A) and at 40 mV *cis* + (B). Protein eluates (90 µl/ml) and $CaCl_2$ (300 µM) were added from the *trans* side of the membrane. Inserts are the corresponding amplitude histograms of conductance.

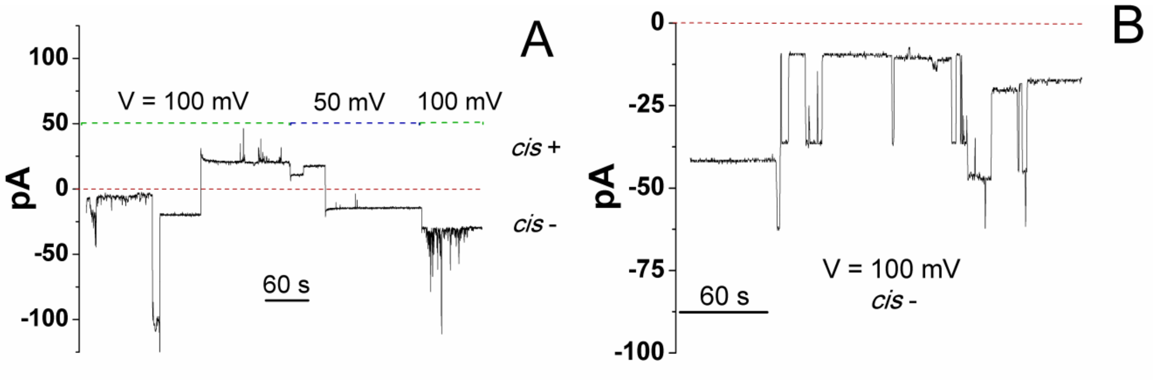


**Fig. S4.** Representative channel activity of F-ATP synthase monomer from control samples. Currents were recorded at 100/50 mV *cis* +/- (A) and at 100 mV *cis* - (B). Red dashed line shows the closed state of the channel. Green and blue dashed lines indicate the time when 100 and 50 mV voltage was applied, respectively. Protein eluates (30 µl/ml) and $CaCl_2$ (300 µM) were added from the *trans* side of the membrane.

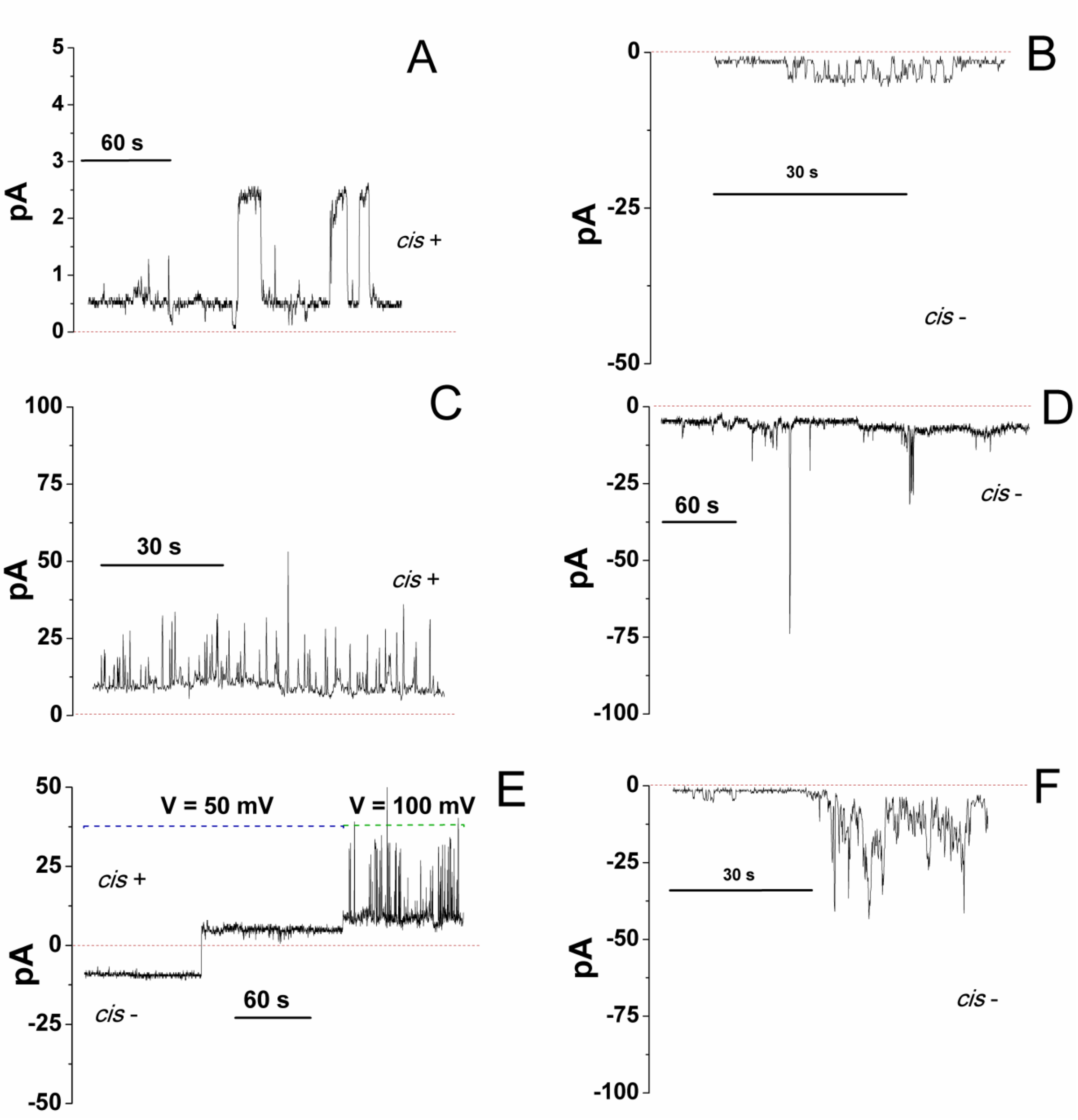


**Fig. S5.** Variations in the channel activity of F-ATP synthase monomer from PTP samples. Currents were recorded at 100 (panels A-D and F), and 50/100 mV *cis* +/- (E). Protein eluates (30 μl/ml) and $CaCl_2$ (300 μM) were added from the *trans* side of the membrane. Red dashed lines indicate the closed state of the channel. Green and blue dashed lines indicate the time when 100 and 50 mV voltage was applied, respectively.

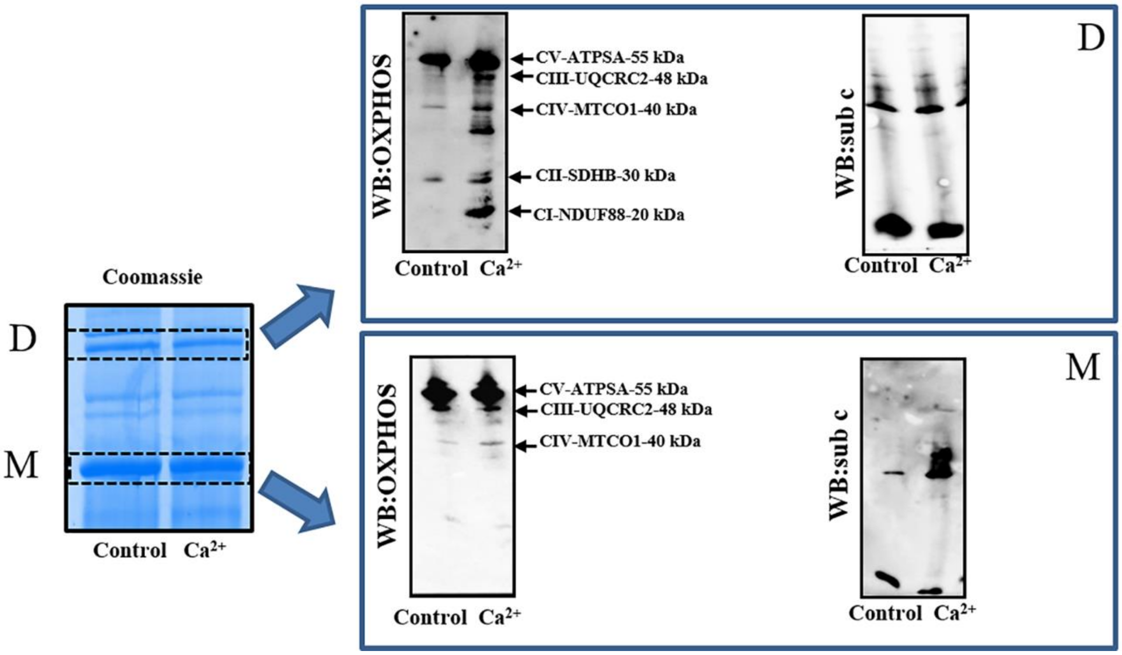


**Fig. S6.** Contamination of F-ATP synthase monomer and dimer bands from control and PTP samples by OXPHOS proteins. F-ATP synthase monomer (M) and dimer bands (D) from RLM control and PTP samples ($Ca^{2+}$) incubated in SM-BM were subjected to SDS-PAGE and immunoblotting for OXPHOS subunits Ndufb8 (CI), Sdhb (CII), Uqcrc2(CIII), Mt-co1(CIV), Atp5a (CV) (WB: OXPHOS), and ATP5MC1 (CV) (WB: sub c). All the figures are representative of at least three independent experiments.

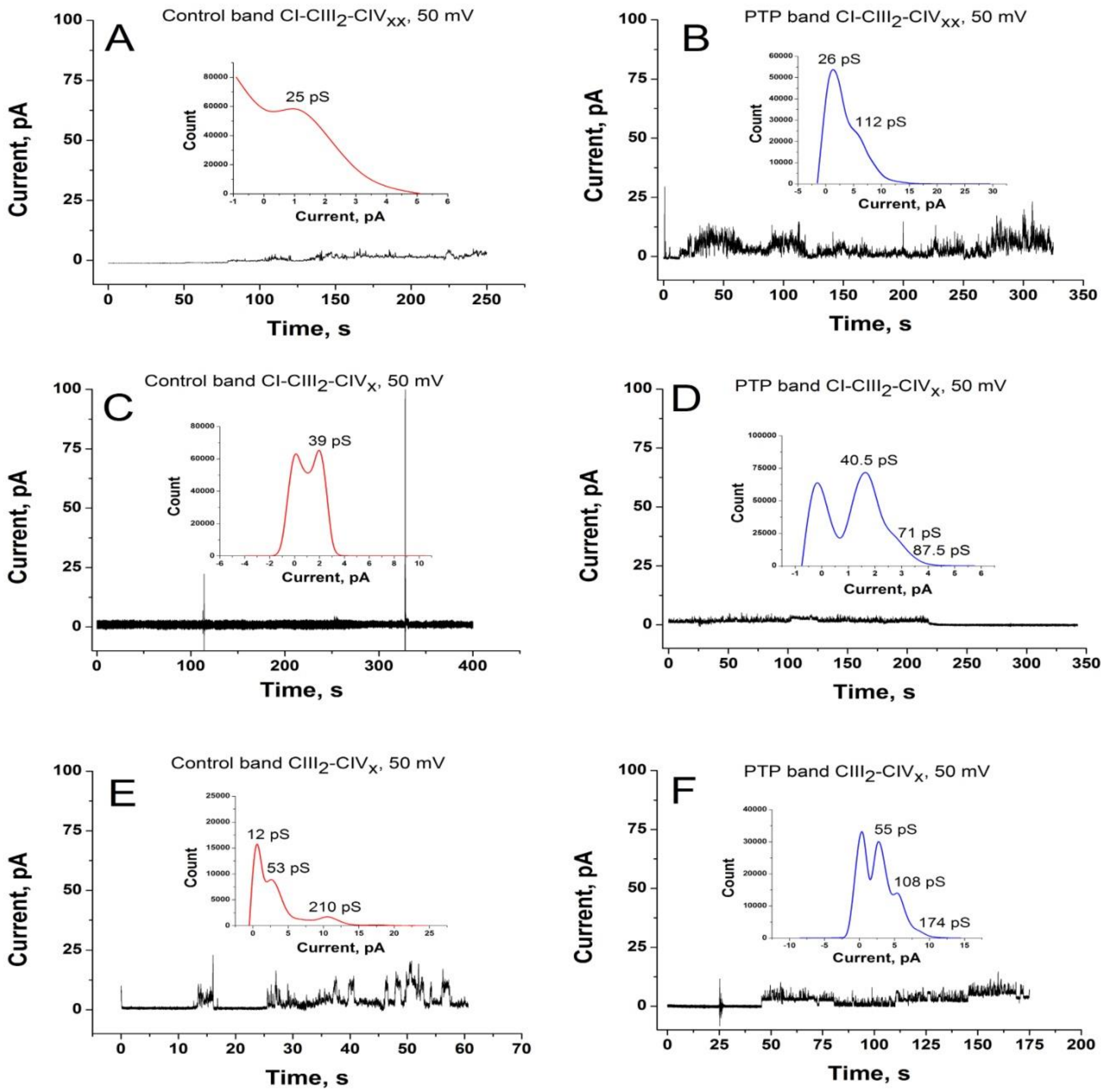


**Fig. S7.** Representative channel-forming activity of mitochondrial supercomplexes from control and mPTP samples eluted from the bands enriched with high (A and B) and low (C and D) molecular weight supercomplexes CI-CIII$_2$-CIV$_x$ and with CIII$_2$-CIV$_x$ supercomplex (E and F). The elution buffer for CI-CIII$_2$-CIV$_x$ supercomplexes was the same as for F-ATP synthase dimers and monomers except ATP was replaced by 5 mM NADH. Currents were recorded at +50 mV (*cis*). Protein eluates (90 µl/ml) and $CaCl_2$ (300 µM) were added from the *trans* side of the membrane. Inserts are the corresponding amplitude histograms of conductance.

**Table S1.** Mitochondrial ion channels and exchangers associated with (super)complexes. Samples were obtained from the RHM and RLM incubated in SM-BM and RLM incubated in KCl-BM in the absence (Control samples) and presence of $Ca^{2+}$ (PTP samples). Data from the corresponding raw MS datasets were extracted by PEAKS Studio 7.5/XPro. Designations: asterisk shows the presence of a protein at the trace quantity; H and L indexes at the CI-$CIII_2$-$CIV_x$ supercomplex indicate high and low molecular weight of the latter. Clic1, chloride intracellular channel protein 1; Clic4, chloride intracellular channel protein 4 (intracellular chloride ion channel protein p64H1); Letm1, mitochondrial proton/calcium exchanger protein (leucine zipper-EF-hand-containing transmembrane protein 1); Letmd1, LETM1 domain-containing 1; Letm2, LETM1 domain-containing protein LETM2, mitochondrial; Kcnj8, ATP-sensitive inward rectifier potassium channel 8 (inward rectifier K(+) channel Kir6.1); Slc8a2, sodium/calcium exchanger 2; Slc8a3, sodium/calcium exchanger 3; Slc8b1, mitochondrial sodium/calcium exchanger protein (solute carrier family 24 member 6); Vdac 1-3, voltage-dependent anion-selective channel protein 1-3.

| **(Super)complex** | **RHM** | | **RLM SM-BM** | | **RLM KCl-BM** | |
|---|---|---|---|---|---|---|
| | Control | PTP | Control | PTP | Control | PTP |
| CI-$CIII_2$-$CIV_x$ (H) | | | Letm2<br>Slc8b1 | Letmd1<br>Slc8b1 | | Letm2 |
| CI-$CIII_2$-$CIV_x$ (L) | | | | Clic4 | | |
| $CV_2$ | | | Slc8b1* | Slc8b1* | | |
| CI | | Vdac 1 | Slc8a3 | | | Clic1<br>Slc8b1 |
| $CIII_2$-$CIV_x$ | Letm1* | | | Vdac 1 | | Vdac 1 |
| CV | | Vdac 3<br>Kcnj8* | Vdac 3 | Slc8a2*<br>Slc8b1* | | Letm1<br>Vdac 1<br>Vdac 2<br>Vdac 3* |

**Table S2.** Changes in the level of prohibitin, prohibitin 2, and methylmalonate-semialdehyde dehydrogenase in the bands of mitochondrial (super)complexes after the PTP opening. Samples were obtained from the RHM and RLM incubated in SM-BM and RLM incubated in KCl-BM in the absence (Control samples) and presence of $Ca^{2+}$ (PTP samples). Data from the corresponding raw MS datasets were extracted by PEAKS Studio 7.5/XPro. In pairs where a protein was absent (ND) either in a control or a PTP sample, the rIBAQ value is given for the present protein. H and L indexes at the CI-$CIII_2$-$CIV_x$ supercomplex indicate high and low molecular weight of the latter.

| | **rIBAQ PTP/rIBAQ Control** | | | | | | | | | | | | | | | | | |
|---|---|---|---|---|---|---|---|---|---|---|---|---|---|---|---|---|---|---|
| | **RHM SM-BM** | | | | | | **RLM SM-BM** | | | | | | **RLM KCl-BM** | | | | | |
| | **CI-$CIII_2$-$CIV_x$ (H)** | **CI-$CIII_2$-$CIV_x$ (L)** | **$CV_2$** | **CI** | **$CIII_2$-$CIV_x$** | **CV** | **CI-$CIII_2$-$CIV_x$ (H)** | **CI-$CIII_2$-$CIV_x$ (L)** | **$CV_2$** | **CI** | **$CIII_2$-$CIV_x$** | **CV** | **CI-$CIII_2$-$CIV_x$ (H)** | **CI-$CIII_2$-$CIV_x$ (L)** | **$CV_2$** | **CI** | **$CIII_2$-$CIV_x$** | **CV** |
| **Prohibitin 2** | 0.650 | 0.834 | 0.184 | 1.019 | + / (0.29) | ND | 1.298 | 0.755 | 0.512 | 0.584 | ND | ND | 0.016 | 0.747 | 0.013 | 0.183 | ND | ND |
| **Prohibitin** | 0.334 | 0.158 | 0.507 | 0.464 | + / (0.085) | ND | 0.936 | 0.571 | 0.609 | 1.063 | 2.712 | ND | 0.742 | 0.880 | 0.834 | 0.159 | ND | ND |
| **Methylmalonate-semialdehyde dehydrogenase** | - / (0.36) | ND | - / (0.12) | ND | 0.115 | ND | 0.013 | 0.012 | 0.028 | 0.11 | 0.244 | 0.401 | 0.010 | 0.018 | 0.029 | 0.165 | 0.112 | 0.348 |

**Dataset S1 (separate file).** Mass-spectrometry data of protein content of F-ATP synthase monomer and dimer bands from control samples of RHM. Raw data were processed using Thermo Xcalibur Qual Browser software.

**Dataset S2 (separate file).** Mass-spectrometry data of protein content of F-ATP synthase monomer and dimer bands from PTP samples of RHM. Raw data were processed using Thermo Xcalibur Qual Browser software.

**Dataset S3 (separate file).** Mass-spectrometry data of protein content of F-ATP synthase monomer and dimer bands from Control samples of RLM incubated in KCl-BM. Raw data were processed using Thermo Xcalibur Qual Browser software.

**Dataset S4 (separate file).** Mass-spectrometry data of protein content of F-ATP synthase monomer and dimer bands from PTP samples of RLM incubated in KCl-BM. Raw data were processed using Thermo Xcalibur Qual Browser software.

**Dataset S5 (separate file).** Mass-spectrometry data of protein content of F-ATP synthase monomer and dimer bands from Control samples of RLM incubated in SM-BM. Raw data were processed using Thermo Xcalibur Qual Browser software.

**Dataset S6 (separate file).** Mass-spectrometry data of protein content of F-ATP synthase monomer and dimer bands from PTP samples of RLM incubated in SM-BM. Raw data were processed using Thermo Xcalibur Qual Browser software.

**Dataset S7 (separate file).** Comparison of protein composition of F-ATP synthase dimers and monomers from control and PTP samples. Samples were obtained from the RHM and RLM incubated in SM-BM and RLM incubated in KCl-BM in the absence (Control samples) and presence of $Ca^{2+}$ (PTP samples). The table summarizes the data from Datasets S1-S6 extracted by PEAKS Studio 7.5/XPro. Sheet titles indicate F-ATP synthase form (Dimer or Monomer), mitochondrial type (RHM or RLM), sample type (Control or PTP), incubation medium (SM-BM or KCl-BM), and whether the list contains information about all true proteins (All) or only those whose PTP/control rIBAQ ratio is greater than two (Big Difference).

**Dataset S8 (separate file).** Protein composition of high- (H) and low-molecular weight (L) CI-$CIII_2$-$CIV_x$ supercomplexes and $CIII_2$-$CIV_x$ supercomplex from control and PTP samples isolated from RHM. Data from the corresponding raw MS datasets were extracted by PEAKS Studio 7.5/XPro. Sheet titles indicate the supercomplex and sample type.

**Dataset S9 (separate file).** Protein composition of high- (H) and low-molecular weight (L) CI-$CIII_2$-$CIV_x$ supercomplexes and $CIII_2$-$CIV_x$ supercomplex from control and PTP samples isolated from RLM incubated in SM-BM. Data from the corresponding raw MS datasets were extracted by PEAKS Studio 7.5/XPro. Sheet titles indicate the supercomplex and sample type.

**Dataset S10 (separate file).** Protein composition of high- (H) and low-molecular weight (L) CI-$CIII_2$-$CIV_x$ supercomplexes and $CIII_2$-$CIV_x$ supercomplex from control and PTP samples isolated from RLM incubated in KCl-BM. Data from the corresponding raw MS datasets were extracted by PEAKS Studio 7.5/XPro. Sheet titles indicate the supercomplex and sample type.
